# OpenSplice: the impact of half a million mutations on the alternative splicing of 600 human exons

**DOI:** 10.64898/2026.05.22.727141

**Authors:** Gioia Quarantani, Joseph Clarke, Mike Thompson, Fei Sang, Juan Valcárcel, Ben Lehner

## Abstract

Alternative splicing of mRNA precursors is an important step in gene regulation and a major mechanism by which genetic variants cause human disease. However, changes in splicing have only been quantified for a tiny fraction of possible variants in the human genome, limiting our ability to interpret clinical variants, evaluate and develop machine learning models, and understand the splicing regulatory code. Here, to address this data gap, we present OpenSplice, a well-calibrated experimental dataset that quantifies the impact of >590,000 variants on the splicing of >600 human alternatively spliced exons. OpenSplice increases the number of exons with site saturation mutagenesis data ∼28-fold, quantifying the impact of all possible exonic and proximal intronic single nucleotide (nt) substitutions, as well as all 1, 3, 6 and 21nt deletions. Hundreds of thousands of variants affect splicing and we use the data to evaluate machine learning models, to interpret clinical variants, and to map splicing regulatory architectures. Exons and introns exhibit a broad spectrum of regulatory landscapes, including configurations dominated by enhancers or silencers, checkerboard-like patterns with interleaved enhancers and silencers, and sparse architectures with minimal regulatory content. Silencers are particularly important for determining inclusion levels. OpenSplice provides a complete atlas of variants that impact splicing, a rich testing and training dataset for machine learning models, and a comprehensive map of regulatory elements for mechanistic studies.

## Introduction

The splicing of precursor messenger RNAs (pre-mRNAs) is a key step in gene regulation, removing introns and joining exons to produce mature transcripts. Splicing is catalyzed by the spliceosome, a large ribonucleoprotein complex that recognizes conserved splice site motifs at exon–intron boundaries. At the 3′ end of introns, splice site recognition also depends on the polypyrimidine tract (PPT) and the branch point (BP), where an intronic adenosine initiates formation of the lariat intermediate. Beyond these core motifs, however, numerous additional cis-regulatory elements can influence splicing, including enhancer and silencer elements distributed across exons and introns^1^.

Disruption of splicing is a major mechanism by which variants cause human genetic diseases^2,3^. Variants in splice sites are relatively straightforward to classify as splice altering, with more than 40,000 splice site-disrupting pathogenic variants listed in the ClinVar database^4^. However, beyond splice sites, the identification of splice-altering variants within exons and introns is challenging^5^. Machine learning methods have been developed to predict variant effects on splicing^6^. However, under current clinical interpretation guidelines, computational predictions can provide only supporting evidence and are not sufficient to classify a variant as pathogenic^7,8^. Consequently, most variants outside canonical splice sites are considered variants of uncertain significance with respect to their impact on splicing and pathogenicity.

Large-scale experimental mutagenesis provides one approach to address this shortcoming. Saturation mutagenesis experiments, where every nt in an exon or intron is mutated, have revealed that many variants within exons and introns can alter splicing, particularly for alternatively spliced exons with intermediate level of inclusion^9–11^. Similarly, large-scale testing has suggested that up to 10% of pathogenic protein sequence-altering missense variants may also alter splicing^5^ and that ∼3% of common variants alter exon inclusion^12^. However, these previous studies have either tested many variants in only one or a few exons or a very small number of variants in and around many different exons, leaving the broader landscape of splice-disrupting variants largely unexplored for most genes.

Here, to address this data gap, we present OpenSplice, a large-scale experimental resource that systematically quantifies the impact of variants on splicing. Using DNA synthesis and massively parallel testing, we quantified the effects of 598,953 variants on the splicing of 608 diverse human exons. The data reveal a remarkably diverse architecture of splicing regulatory elements across exons and introns, highlight the importance of silencers in determining inclusion levels, and provide a powerful atlas for clinical variant interpretation, the evaluation and training of machine learning models, and the study of mechanisms of splice site selection.

## Results

### Massively parallel site-saturation mutagenesis of human exons and proximal introns

Previous site-saturation mutagenesis of human exons has quantified the impact of variants in one^10,11,13–17^, three^18^, or, at most, 15^19^ alternatively spliced exons. Across these studies, between 4,000^16^ and 9,670^19^ variants were tested, together comprising 44,836 mutations across 22 exons. To accelerate data production and obtain more generalizable principles of regulation, we established a platform that quantifies the splicing of tens of thousands of variants in hundreds of exons in each experiment. We used DNA synthesis to construct pooled variant libraries covering 70nt of upstream intron, the target exon, and 25nt of downstream intron in a minigene construct derived from the FAS gene with variant libraries replacing alternatively spliced exon 6 and its flanking intronic sequences^20,21^. Random barcodes inserted after the last exon enable the splicing products of each genotype to be quantified, even when the target exon is skipped (**Fig. 1a**).

**Figure 1.**
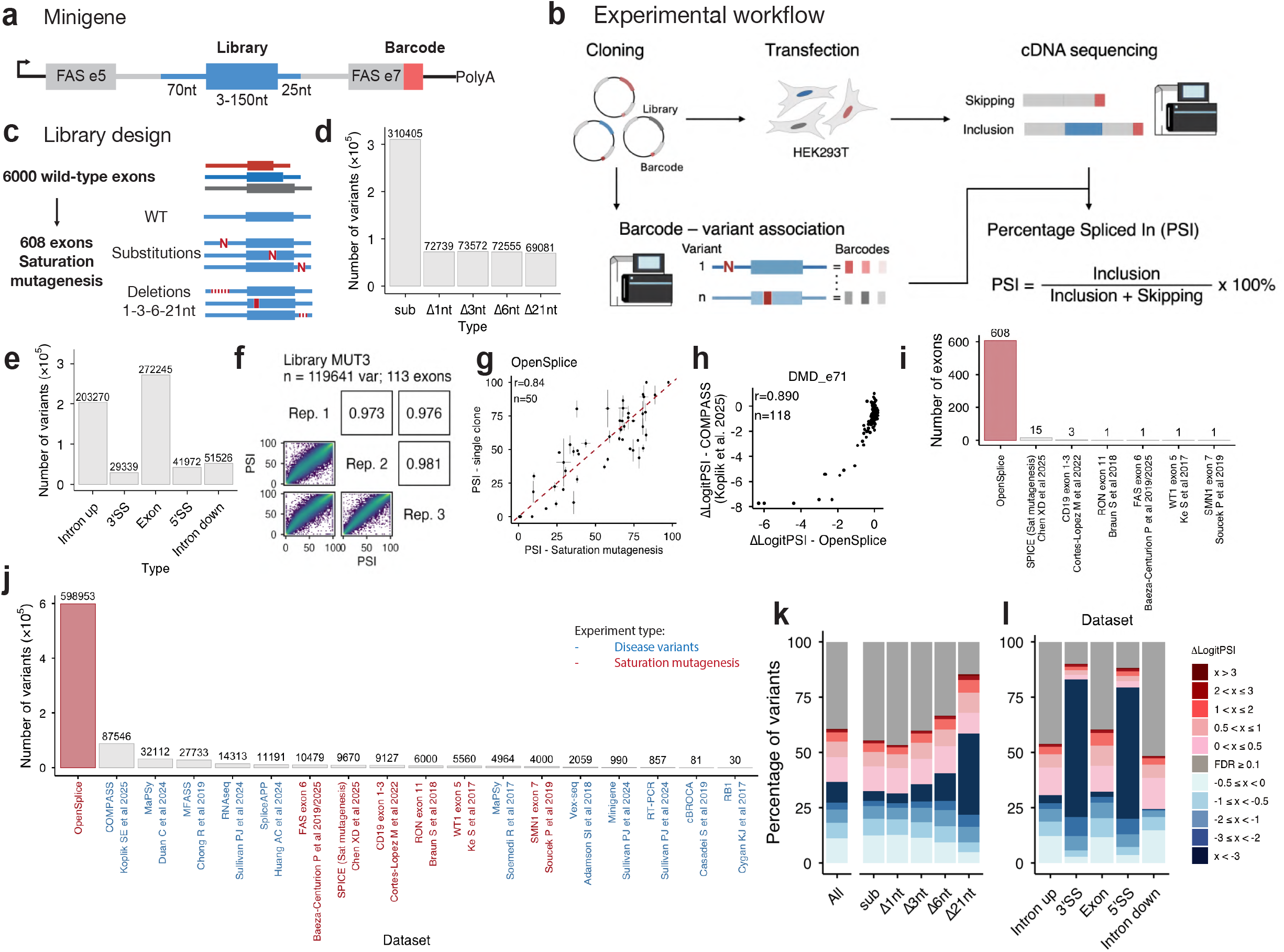
OpenSplice: massively parallel site-saturation mutagenesis of 608 human exons. **(a)** Schematic of the minigene reporter system. **(b)** Experimental workflow. **(c)** Library design for the 608 exons. **(d)** Number of variants per mutation type in the final OpenSplice dataset: substitutions (sub, n = 310,405), and deletions of 1 nt (Δ1nt, n = 72,739), 3nt (Δ3nt, n = 73,572), 6nt (Δ6nt, n = 72,555), and 21 nt (Δ21nt, n = 69,081). **(e)** Distribution of variants across genomic regions: upstream intron (n = 203,270), 3′ splice site (n = 29,339), exon (n = 272,245), 5′ splice site (n = 41,972), and downstream intron (n = 51,526). **(f)** Reproducibility of PSI measurements across three biological replicates for one mutagenesis library (MUT3, n = 117,507 variants in 113 exons). Pairwise Pearson r values are shown. **(g)** Correlation between PSI values measured by saturation mutagenesis in OpenSplice and by individual single-clone experiments (r = 0.84, n = 50). Variants tested span 25 exons from 23 genes and include wild-type constructs and variants representing substitutions (*BRAF* exon 9 G6T; *BRCA1* exon 17 A43T; *CCDC7* exon 12 G82C; *COL18A1* exon 19 G5T; *COL7A1* exon 75 A3C, A91C; *CREBBP* exon 11 C65T; *FOXP1* exon 18 G103A; *LRRFIP2* exon 15 G127C; *PCIF1* exon 9 A121T; *ROBO1* exon 20 A62G, A6T; *VEGFA* exon 7 G78T; *WAS* exon 5 G137T; *ZNF345* exon 3; *CAPRIN2* exon 6 T33G) and deletions of 1 to 21 nt (*BRAF* exon 9 del6-26, del44; *CCDC178* exon 21 del197-217; *FSTL5* exon 11 del95; *HFM1* exon 3 del132-134; *LRRFIP2* exon 15 del124-144, del185; *MYO6* exon 29B del26-28; *PAX6* exon 6 del51-53, del63; *PCIF1* exon 9 del137-157; *SPDL1* exon 2A del39-41; *SUZ12* exon 4 del141-143; *WAS* exon 5 del26-31; *ZNF345* exon 3 del167-169). **(h)** Correlation between ΔLogitPSI values from OpenSplice and an independent dataset^23^ for DMD exon 71 (r = 0.890, n = 118). Note that baseline exon inclusion differs between experiments. **(i)** Comparison of the number of exons with site-saturation mutagenesis data in OpenSplice (n = 608) versus 6 prior studies^11,14–18^. Numbers above bars indicate exon counts per study. **(j)** Comparison of the total number of variants quantified in OpenSplice (n = 598,953) versus prior disease-variant (blue) and saturation mutagenesis (red) datasets. **(k)** Percentage of variants per mutation class causing significant changes in exon inclusion (FDR = 0.1, Benjamini–Hochberg), stratified by direction and magnitude of ΔLogitPSI. Longer deletions affect splicing at progressively higher rates: substitutions (54.5%), Δ1nt (53.4%), Δ3nt (59.9%), Δ6nt (66.7%), Δ21nt (85.5%). **(l)** Percentage of variants per genomic region causing significant changes in exon inclusion (FDR = 0.1), stratified by direction and magnitude of ΔLogitPSI.

In a pilot selection we designed three libraries containing 23,231 site-saturation variants for 22 alternatively spliced human exons (P1-3, **Supplementary Table 1**) and we re-generated a previously characterized library^15^ with variant-associated barcodes (**Supplementary Table 2**). We transfected HEK293T cells in triplicate, extracted RNA 48h post-transfection, and reverse-transcribed RNA into cDNA that was sequenced. In parallel, we directly sequenced the plasmids to obtain a dictionary of variant:barcode pairs. By combining the two sequencing methods, we calculated the percentage spliced in (PSI) for each variant (**Fig. 1b**). PSI values quantified by barcode-sequencing in pooled exon libraries were highly reproducible (**Extended Data Fig. 1a-d**) and very well correlated with inclusion quantified by direct sequencing (Pearson r = 0.92, n = 5,838; **Extended Data Fig. 1e**) and individual variant testing (r = 0.89, n = 40; **Extended Data Fig. 1f**).

We next constructed three libraries containing the wild-type (WT) intron-exon-intron sequences of 6,000 human exons (WT1-3; **Supplementary Table 3**) with lengths between 3 and 150 nts and enriched for alternatively spliced exons: 3,364 have mean PSI across tissues between 10% and 90% in VastDB^22^, 674 mean PSI < 10%, and 1962 mean PSI > 90%. The last group was also enriched for exons with clinical variants^4^. PSI values in the minigene construct were very well correlated between replicates (median r=0.897, **Extended Data Fig. 1g-i**), with a median PSI = 64% and interquartile range (IQR) = 79%. We used these WT PSI measurements to select exons for site-saturation mutagenesis.

### OpenSplice: the impact of >590,000 variants on the splicing of 608 exons

We used pooled DNA synthesis to construct six additional site-saturation mutagenesis libraries for 586 of the 6000 tested WT exons, prioritising short exons with clinical variants and intermediate WT PSI values (**Extended Data Fig. 1j-k, Supplementary Table 1**), with exons pooled according to length. For each exonic and proximal intronic region the libraries contained all single nt substitutions and all possible deletions of length one, three, six and 21 nts (**Fig. 1c**). An intron-exon-intron construct of length 193 nt therefore requires measuring 7 x 193 = 1351 variants.

After quality control, the final OpenSplice dataset quantifies PSI values for 598,953 variants in and around 608 exons (92.7% of designed genotypes, **Supplementary Table 4**), including 310,405 substitutions, 72,739 1nt deletions, 73,572 3nt deletions, 72,555 6nt deletions and 69,081 21nt deletions (**Fig. 1d**). 272,245 variants are in the exons, 203,270 in the upstream introns, 51,526 in the downstream introns, 29,339 in the 3’ splice site (defined here as the last 4nt of the upstream intron and first nt of the exon), and 41,972 in the 5’ splice site (defined here as the last 3nt of the exon and first 6nt of the downstream intron) (**Fig. 1e**).

PSI values were very well correlated between replicate experiments (median r=0.976, **Fig. 1f, Extended Data Fig. 1l-p**). They also agree very well with testing of individual variants (r=0.84, n=50, **Supplementary Table 5, Fig. 1g**) and with an independent dataset of variants across 13 exons^23^, with one representative exon shown in **Fig. 1h** (*DMD* exon 71: r=0.89, n=118) and the remaining 12 shown in **Extended Data Fig. 1q**. Changes in PSI (ΔPSI values) in OpenSplice also classify ‘splice-altering’ from ‘neutral’ variants in SpliceVarDB^24^ extremely well, with Receiver Operating Characteristic - Area Under the Curve, ROC-AUC = 0.914 (n=190 splice-altering variants, n=129 neutral variants, **Extended Data Fig. 1r-s**).

The OpenSplice dataset increases the number of exons and proximal intronic regions with site-saturation mutagenesis data 28-fold, from 22 to 630 (**Fig. 1i**). It also increases the total number of variants quantified in site-saturation studies 13-fold, from 44,836 to 598,953 (**Fig. 1j**).

### Hundreds of thousands of variants affect splicing

In total, 362,199 of the 598,953 tested variants (60.5%) altered exon inclusion (False Discovery Rate, FDR=0.1, Benjamini–Hochberg procedure (**Fig. 1k, Extended Data Fig. 2a**), with 45.3% of variants (270,951) causing a change in inclusion of greater than 5 PSI units, 32.9% (196,931) a change of >10 PSI units, and 20.5% (122,749) a change of >20 PSI units (FDR=0.1). Variants disrupting splice sites were most likely to affect splicing (3’ splice site: 26,459/29,339, 90.2%; 5’ splice site: 36,982/41,972, 88.2%), followed by variants within exons (164,602/272,245, 60.4%), upstream introns (109,261/203,270, 53.8%), and downstream introns (24,895/51,526, 48.4%) (**Fig. 1l**). Variants in exons affected splicing more often than variants in upstream introns (60.4% vs 53.8%, p < 2.2 × 10^−16^, Fisher’s exact test, FET) and downstream introns (60.4% vs 48.4%, p < 2.2 × 10^−16^, FET).

In the upstream intron, distal intronic variants (1-26 nt) altered inclusion less frequently than exonic variants (37,029/85,071, 43.5%; p < 2.2 × 10^−16^, FET), with only 23.9% producing PSI changes greater than 5 units. Variants within the region where branch point sequences are typically located (27-51 nt; -44 to -19 nt relative to the 3′ splice site) had intermediate effects (46,210/79,986, 57.8%; 41% with PSI change >5 units) and were still less likely to alter splicing than exonic variants (p < 2.2 × 10^−16^, FET). In contrast, variants in the PPT region (52-66 nt) were more likely to affect inclusion than exonic variants (26,589/38,927, 68.3%; 55.4% with PSI change > 5 units; p < 2.2 × 10^−16^, FET, **Extended Data Fig 2b**).

Substitutions (171,987/310,405, 55.4%) and single-nucleotide deletions (38,801/72,739, 53.3%) affected inclusion at similar rates, whereas longer deletions were progressively more impactful: 3-nt deletions (44,062/73,572, 59.9%), 6-nt deletions (48,298/72,555, 66.6%), and 21-nt deletions (59,051/69,081, 85.5%) (p < 2.2 × 10^−16^, χ^2^ test; **Fig. 1k**).

Mutations were overall more likely to decrease exon inclusion than to increase it (36.8% vs 23.8%, FDR = 0.1, **Fig. 1k, Extended Data Fig 2c**). This bias toward decreased inclusion was observed for variants in exons (32.1% vs 28.3%) and upstream introns (30.8% vs 23.1%), after removing variants disrupting canonical splice sites (**Fig. 1l, Extended Data Fig 2c**). In contrast, variants in downstream introns were similarly likely to increase or decrease inclusion (24.0% vs 24.3%, **Fig. 1l, Extended Data Fig. 2c**). In addition, while most mutation types tended to decrease inclusion, exceptions were observed: 1-nt deletions in upstream introns were similarly likely to increase or decrease inclusion (23.9% vs 23.8%), and 21-nt deletions within exons more frequently increased inclusion (37.9% vs 48.8%, **Extended Data Fig. 2c**).

### Complete mutational landscapes

We provide the individual mutational maps for all 608 exons in **Supplementary Fig. 1**. These maps are also available through an interactive web interface, ExonExplorer (https://results.hgi.sanger.ac.uk/OpenSplice/, **Extended Data Fig. 3**). The data for 585 exons with most complete coverage are shown as a single heatmap in **Fig. 2a** ordered by WT PSI. Mutational maps for five example exons, spanning a range of inclusion levels, are shown in **Fig. 2b-f**, including an exon with very low inclusion, *SNRNP70* exon 8, where mutations identify an intronic and an exonic splicing silencer, and *SMN2* exon 7 where mutations identify a well-characterized overlapping SRSF2-binding enhancer, an hnRNP A1-binding silencer^25,26^, and a known Tra2β-dependent exonic splicing enhancer^26^.

**Figure 2.**
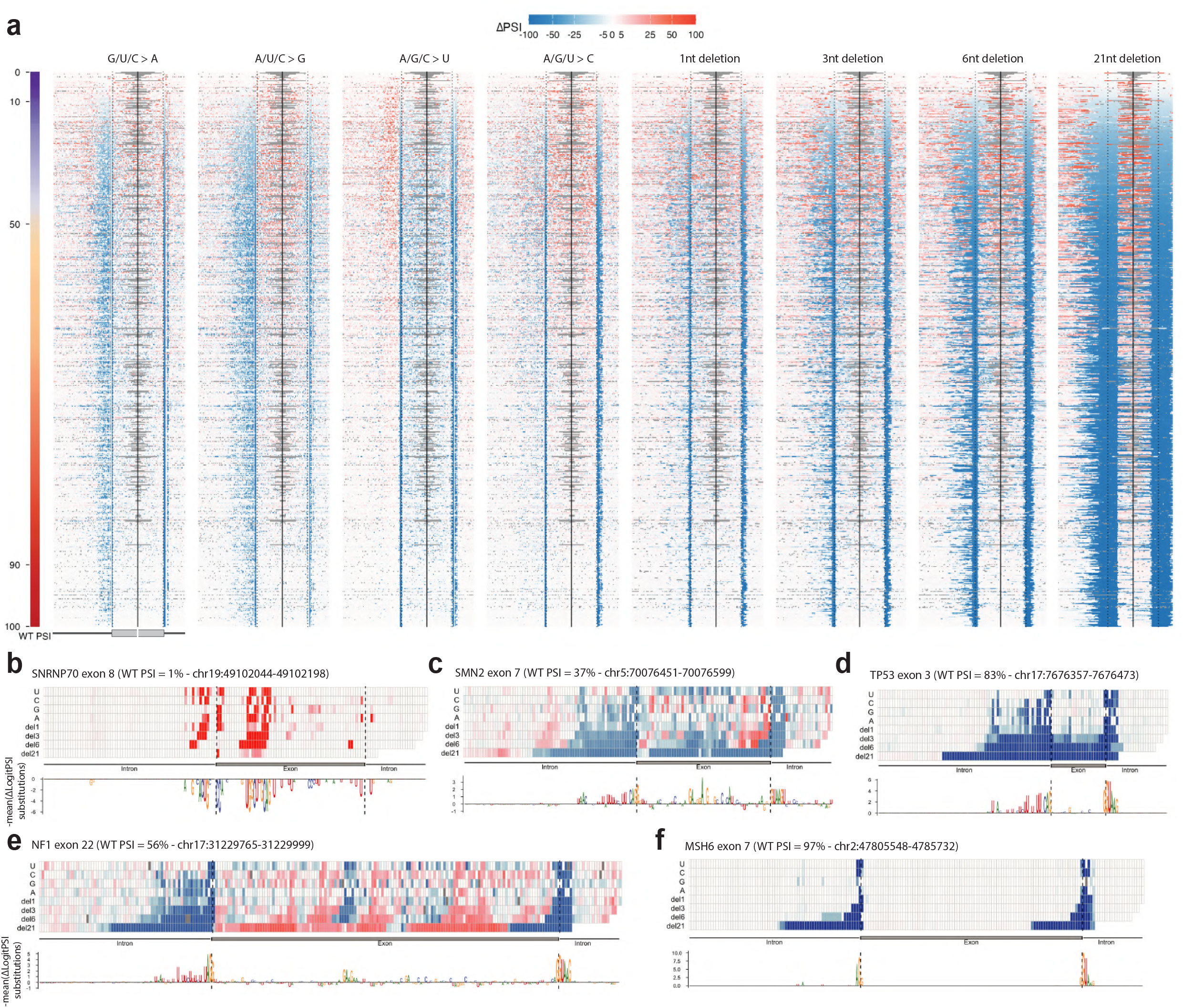
Complete mutational landscapes of human exons. **(a)** Heatmap of ΔPSI values for all 585 most completely characterized exons (>50% designed variants recovered), ordered by wild-type (WT) PSI (left color bar). Columns represent individual nucleotide positions split by mutation type (substitutions to A, G, U, C and deletions of 1, 3, 6, and 21 nt); rows represent exons. Exons are aligned at the splice sites; only the first and last 50 nt of each exon are shown. Deletions are plotted by the middle position. The color scale ranges from blue (exon skipping) to red (increased inclusion). Dashed black lines indicate splice site boundaries. **(b-f)** Mutational maps for five exons spanning a range of wild-type inclusion levels. For each exon, the upper panel shows per-position negative absolute mean ΔLogitPSI of substitution values (above the x-axis = favor inclusion; below x-axis = favor skipping), colored by nucleotide identity (A, green; G, yellow; U, blue; C, red). The lower heatmap shows effects of all substitution and deletion classes (rows: U, C, G, A substitutions and Δ1, Δ3, Δ6, Δ21 nt deletions). Deletion effects are plotted at their starting position. Dashed black lines indicate exon boundaries. (**b**) *SNRNP70* exon 8: WT PSI = 1% - chr19: 49102044-49102198. (**c**) *SMN2* exon 7: WT PSI = 37% - chr5:70076451-70076599. (**d**) *NF1* exon 22: WT PSI = 54% - chr17:31229765-31229999. (**e**) *TP53* exon 3: WT PSI = 80% - chr17:7676357-7676473. (**f**) *MSH6* exon 7: WT PSI = 97% - chr2: 47805548-47805732

### Exons with intermediate inclusion are more sensitive to mutation

The proportion of variants changing inclusion by >10 PSI units differs extensively across exons ranging from 0% to 77.1% (median=31.5%, IQR=36.5%, **Extended Data Fig. 2d**). Consistent with previous work^9,14^, alternatively spliced exons with intermediate inclusion were more sensitive to mutation than those with very low or very high inclusion. This scaling of mutational sensitivity is observed for all mutation types (**Fig. 3a**) and for both exonic and intronic variants (**Extended Data Fig. 2e**). For example, whereas 9.8% and 5.8% of substitutions alter PSI in exons with <10% inclusion (n=35 exons) and >90% (n=88 exons) WT inclusion, respectively, a median of 31% do so in exons with between 10% and 90% WT inclusion (n=478 exons). To account for the scaling of ΔPSI values with WT exon inclusion, we quantify them as ΔLogitPSI in most subsequent analyses. However, even in exons with similar intermediate WT PSI (40–60%), the fraction of variants changing inclusion by >10 PSI units is highly variable, ranging from 13.6% to 72.4% (median = 57.3%, IQR = 13.6%) (**Extended Data Fig 2d**). Baseline PSI alone does not, therefore, fully explain mutational sensitivity.

**Figure 3.**
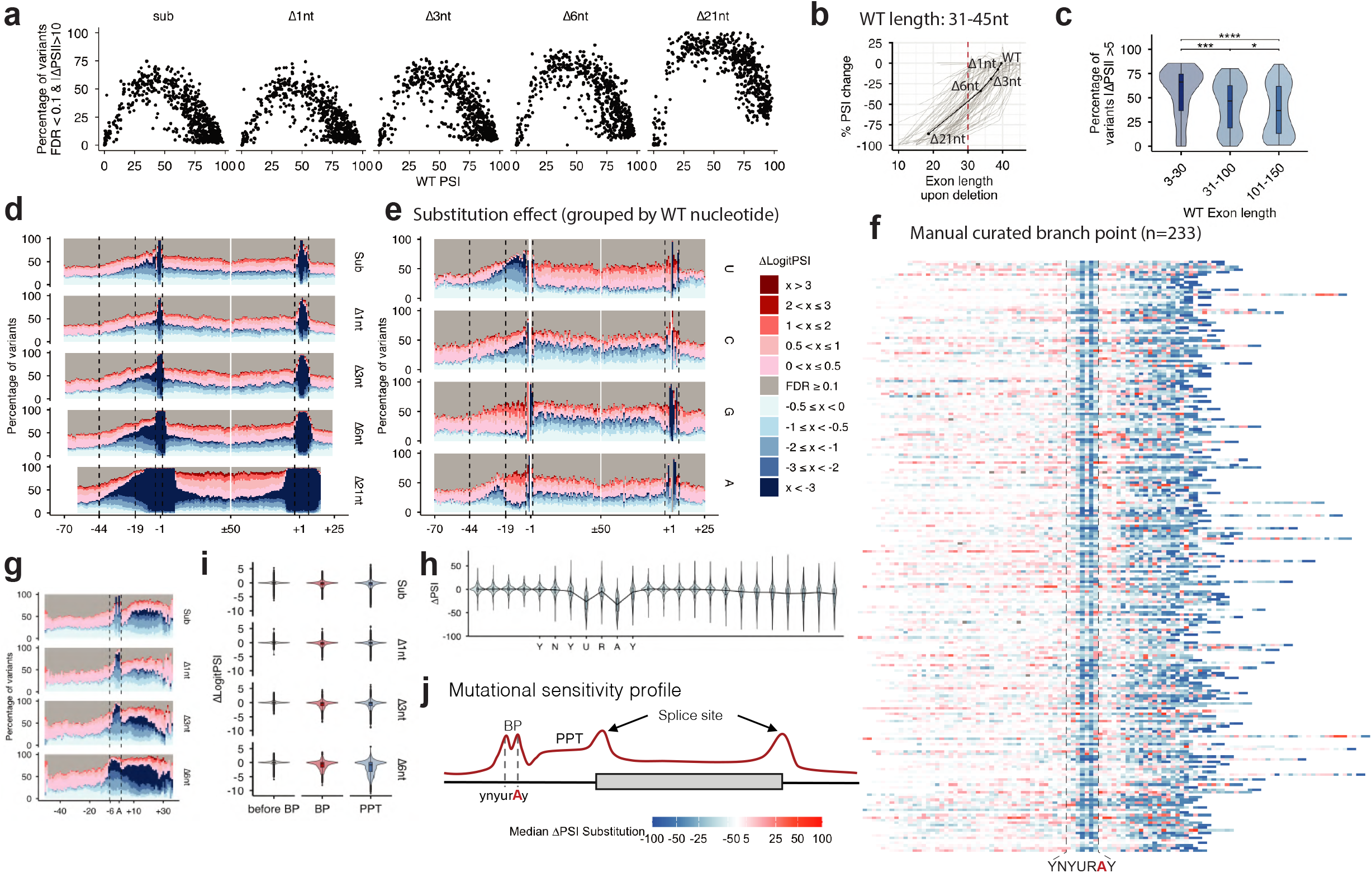
Determinants of mutational sensitivity. **(a)** Relationship between WT PSI and the percentage of non-neutral variants (FDR = 0.1) per exon, shown for each mutation class (substitutions, Δ1nt, Δ3nt, Δ6nt, Δ21nt). **(b)** Effect of exon shortening on exon inclusion, shown for exons with WT length 31-45 nt (remaining length bins in Extended Data Fig. 2g). Each line represents a single exon; points correspond to the resulting exon length after exonic deletions (excluding deletions that disrupt splice sites). The y-axis shows the mean ΔPSI at each resulting exon length, normalized to the WT PSI. **(c)** Percentage of variants with |ΔPSI| > 5 (FDR = 0.1) per exon for microexons (≤30 nt, n = 42) compared to longer exon length bins (31-100 nt and 101-150 nt). **(d)** Mutational sensitivity profiles across the pre-mRNA. For each of the five mutation classes (substitutions, Δ1, Δ3, Δ6, Δ21 nt), the percentage of variants significantly increasing (red) or decreasing (blue) exon inclusion is plotted as a function of position. Exons are aligned at their splice sites and split at their midpoint, showing the first and last 50 nt together with 70 nt upstream and 25 nt downstream. Three functional upstream intron regions are demarcated: PPT region (positions −1 to −19), BP region (−19 to −44), and distal intronic region (−44 to −70). **(e)** Substitution effect profiles across the pre-mRNA, grouped by wild-type nucleotide identity (rows: U, C, G, A). For each WT nucleotide, the percentage of substitutions significantly increasing (red) or decreasing (blue) exon inclusion is plotted per position as in (e). **(f)** Heatmap of median ΔPSI for all substitutions in the upstream intron of 233 exons with a manually curated single dominant branch point, aligned at the putative BP adenosine (position 6 of the YNYURAY consensus motif). Each row is one intron, colored by median ΔPSI per position (blue: decreased inclusion; red: increased inclusion). **(g)** Mutational sensitivity profiles in the upstream intron aligned at the BP adenosine, for all mutation classes. **(h)** Violin plots showing ΔLogitPSI distributions for substitutions and short deletions (1-,3-,6-nt) in the region before the BP, within the BP motif, and within the PPT. **(i)** Violin plots of ΔLogitPSI for substitutions at positions before the BP, within the BP motif, and within the PPT. **(j)** Summary schematic of mutational sensitivity across the pre-mRNA, illustrating peaks at the BP (ynyurAy), PPT, and splice sites, and the low sensitivity of the distal upstream intronic region.

### Exon length and splice site strength

Using exonic deletions that do not disrupt splice sites we examined the relationship between exon length and splicing. Plotting the mean PSI for all variants with the same length normalized to the WT PSI (**Extended Data Fig 2f**) shows that inclusion progressively decreases in exons shorter than 60 nt = 0.55, p < 2.2 × 10^−16^) (**Fig. 3B, Extended Data Fig 2g**). This is consistent with previous proposals of an optimal length for exon definition^27^. In contrast, only a weak negative relationship was observed for longer exons (R = −0.17, p = 6.8 × 10^-14^, **Extended Data Fig 2g**). When exon length is below ∼30 nt, inclusion is almost completely lost for most exons (median PSI =1% below 30nt vs 59% above 30nt; Wilcoxon p = 2.3 × 10^−16^, **Fig. 3b, Extended Data Fig 2h**). This threshold corresponds to the definition of microexons (≤30 nt), which require specialized regulatory mechanisms^28^. Nevertheless, a small subset of variants <30nt retained substantial inclusion, with n = 546 variants across 27 exons having PSI > 10% (**Extended Data Fig 2i**). Most of these derived from WT exons with high baseline inclusion (median WT PSI = 75.5% vs 59.6%; Wilcoxon p = 7.07 × 10^−4^, **Extended Data Fig. 2j**) and the mechanisms that retain their inclusion will be interesting to investigate in future studies.

OpenSplice included 42 wild-type exons of length ≤30 nt. These microexons were more sensitive to sequence variation than longer exons, with a higher median percentage of variants per exon producing |ΔPSI| ≥ 5 than exons with length 31-100nt or 101-150nt (66.3% vs 46.6% and 36.8%, respectively; Wilcoxon p = 4.3 × 10^−4^ for 3-30nt vs 31-100nt, **Fig. 3c**). This increased sensitivity was observed for both substitutions and deletions (median percentage of variants per exon with |ΔPSI| ≥ 5, 3-30nt vs 31-100nt: substitutions, 63.8% vs 41.6%, adjusted p = 3 × 10^−4^; Δ6nt deletions, 75.9% vs 41.6%, adjusted p = 1 × 10^−4^; **Extended Data Fig. 2k**). Microexons were also more sensitive to mutations in the flanking introns (median percentage of variants per exon with |ΔPSI| ≥ 5 upstream intron, 3-30nt vs 31-100nt: 66.1% vs 38.2%, adjusted p = 9.9 × 10^−6^; **Extended Data Fig. 2l**).

Mutations in splices sites were well predicted by the change in MaxEntScan^29^ score (3’ splice site: r=0.72, n=38874; 5’ splice site: r=0.75, n=15199; **Extended Data Fig. 4a-d**). However, MaxEntScan^29^ scores showed only weak positive correlations with WT PSI in both the OpenSplice minigene dataset (3′ splice site: r = 0.16, p = 8.37 × 10^−5^; 5′ splice site: r = 0.19, p = 4.92 × 10^−6^; mean: r = 0.24, p = 2.02 × 10^−9^; **Extended Data Fig. 4e-g**) and endogenous exons from VastDB^22^ (3′ splice site: r = 0.12; 5′ splice site: r = 0.11; mean: r = 0.16; p < 2.2 × 10^−16^; **Extended Data Fig. 4h-j**), explaining only ∼4% and ∼2.6% of variance in inclusion levels, respectively. Splice-site strength was also only weakly associated with overall mutational sensitivity, measured as median |ΔLogitPSI|, with stronger splice sites showing slightly lower sensitivity (3′ splice site: r = −0.22, p = 3.56 × 10^−8^; 5′ splice site: r = −0.17, p = 2.63 × 10^−5^; mean: r = −0.28, p = 3.45 × 10^−12^; **Extended Data Fig. 4k-m**), explaining ∼5% of the variance. Thus, while changes in splice-site strength predict the local effects of splice-site mutations, splice-site strength itself explains only a small fraction of WT inclusion levels and overall exon sensitivity to mutation.

### Mutational sensitivity profiles

Considering all exons, sensitivity was quite uniform along exons (excluding the first and last 3 nt) and in the downstream exon, but was reduced in the upstream intron further from the 3’ splice site (**Fig. 3d,j**). Within exons, substituting or deleting a U increases inclusion more often than decreasing it (U>A/C/G: 36.4% vs 19.6%; ΔU: 33% vs 19.5%), whereas substituting or deleting a C more often reduce inclusion (C>A/G/U 18.9% vs 39.7%, ΔC: 17.9% vs 37.5%). Mutations affecting G also more often decrease inclusion (G>A: 20.1% vs 40.4%; G>U: 16.3% vs 47.1%; ΔG: 21.7% vs 37.2%), except for G>C substitutions, which more often increase inclusion (30.2% vs 26.8%). For A, the effect depends on the specific mutation: ΔA and A>U more often decreases inclusion (ΔA: 23.6% vs 27.8%; A>U: 12.9% vs 41.7%), whereas A>C and A>G more often increases inclusion (A>C: 34% vs 20.8%; A>G: 30.8% vs 26.7%) (**Fig. 3e, Extended Data Fig. 5a-b**). Together with the effects of three and six nucleotides deletions (**Extended Data Fig. 5c-e**), this is consistent with the disruption of purine-rich exonic splicing enhancers^30^ and U-rich exonic silencer elements such as PTB-binding site^15,31^. The decreased inclusion associated with C mutations is consistent with cytosine-containing motifs acting as exonic splicing enhancers, potentially through A/C-rich ACE elements bound by YB-1^32^.

### Branchpoints and upstream introns

The first catalytic step of splicing occurs when the 2’-OH of the branchpoint (BP) A attacks the upstream 5’ splice site to form a lariat structure^33^. Approximately 30% of human introns use multiple BPs^34–36^. In OpenSplice, 233 of 585 upstream introns (39.8%) have a mutational signature consistent with usage of a single major BP (**Supplementary Table 6**), defined by deleterious effects of A>G,U,C substitution at the putative branch site A and potential base pairing to U2 snRNA of the flanking nucleotides. We re-aligned these introns at their BPs. Within the YNYURAY BP motif mutations at position 4 (U) and 6 (A) are most detrimental (**Fig. 3f-h**), consistent with evolutionary conservation^34–36^. Mutations at positions 1, 2, 3, 5, and 7 follow a pattern consistent with U2 snRNA pairing (**Extended data Fig. 5f**): variants that improve U2 complementarity increase inclusion, whereas variants that impair pairing decrease inclusion (**Extended Data Fig. 5g**). Interestingly, variants upstream of the BP are much less likely to have large effects on splicing, with only 1% of variants upstream of the BP having |ΔLogitPSI| > 0.5, compared to 44% in the BP motif and 47% in the downstream polypyrimidine tract. The region upstream of the BP is thus particularly insensitive to mutation and likely devoid of strong splicing regulatory elements (**Fig. 3f,i**), as has been previously observed in some examples^37^.

### Evaluating splicing variant effect predictors

We next used OpenSplice to evaluate the performance of computational methods for predicting splicing, evaluating four state-of-the-art neural network models: SpliceAI^38^, Pangolin^39^, SpliceTransformer^40^, and AlphaGenome^41^. All four models provided good predictions (**Fig. 4a, Extended Data Fig 6a-b**). The best performing model was SpliceAI (median per-exon Spearman R=0.73), followed by Pangolin (R=0.71). Interestingly the exons poorly predicted by one model tended to also be poorly predicted by all models (R=0.55-0.84 for exons with median inter-replicate experimental Spearman rho >0.8,. **Extended Data Fig. 6c-f**). Predictive performance was highest for all models in the 5’ splice sites (R=0.59-0.77 when all mutations pooled across all constructs) and 3’ splice sites (R=0.52-0.73), and lowest in the downstream intron (R=0.24-0.40). Performance in the upstream intron (R = 0.41-0.58) and exon (R=0.41-0.60) was intermediate (**Fig. 4b**). Longer deletions - which have larger effects - were better predicted than substitutions (R=0.74-0.85 vs. R=0.49-0.64, respectively).

**Figure 4.**
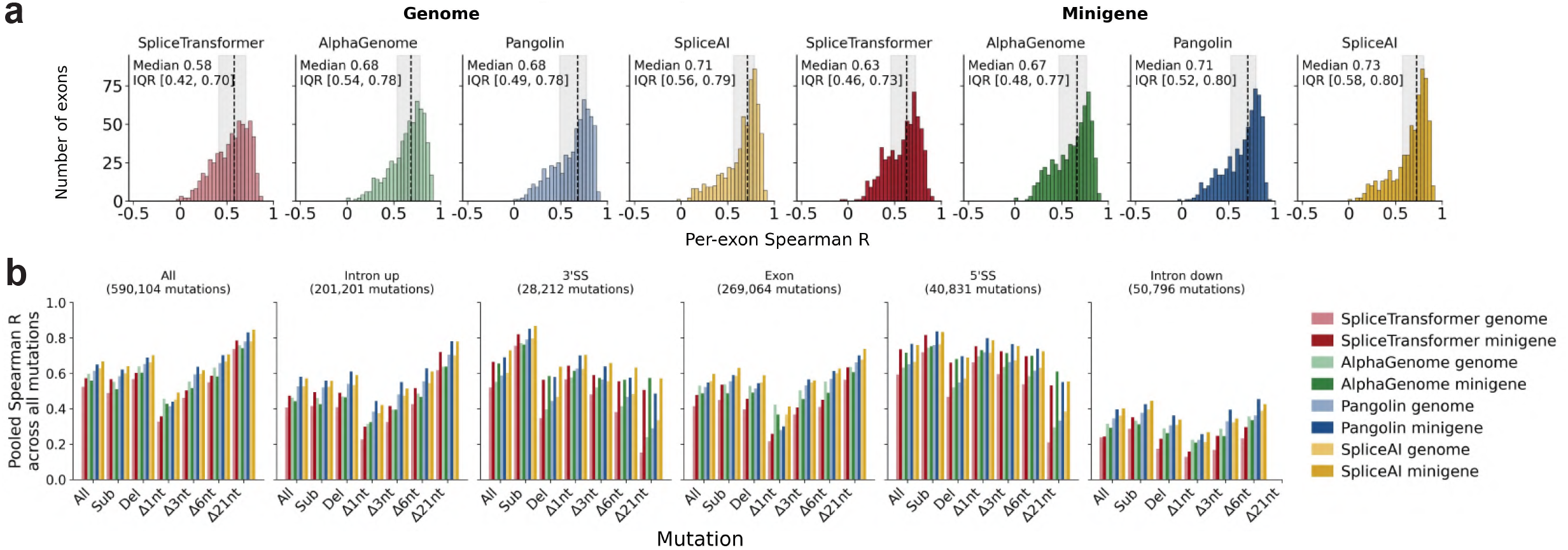
Evaluation of deep learning splicing variant effect predictors. **(a)** Distributions of per-exon Spearman R between predicted and experimentally measured ΔPSI for four models (SpliceTransformer, AlphaGenome, Pangolin, and SpliceAI) run in genome mode (top row) or minigene mode (bottom row), across n = 599 exons. Median R and IQR are indicated. **(b)** Pooled Spearman R across all mutations, stratified by genomic region (columns: all, upstream intron, 3′SS, exon, 5′SS, downstream intron) and mutation type (x-axis), for all eight model-context combinations.

Splicing models therefore provide very good prediction of variant effects but their performance varies across exons, regions, and mutation types, suggesting where additional data and re-training may be most beneficial.

### SpliceMaps: splicing regulatory landscapes

We used the substitution and short (1-, 3-, and 6-nt) deletion data to map the location of putative splicing cis-regulatory elements. We refer to these summary maps as SpliceMaps (**Fig. 5a**). To generate SpliceMaps, we used the most negative and most positive change in splicing for mutations at each position. We defined stretches of four or more nts with minimum ΔLogitPSI < −1 as putative enhancers (E) and four or more nts with maximum ΔLogitPSI > 1 as putative silencers (S). Since these two states are not mutually exclusive, nucleotides satisfying both criteria are labeled as overlapping (O). The remaining sequence is classified as neutral (N) (**Fig. 5a**).

**Figure 5.**
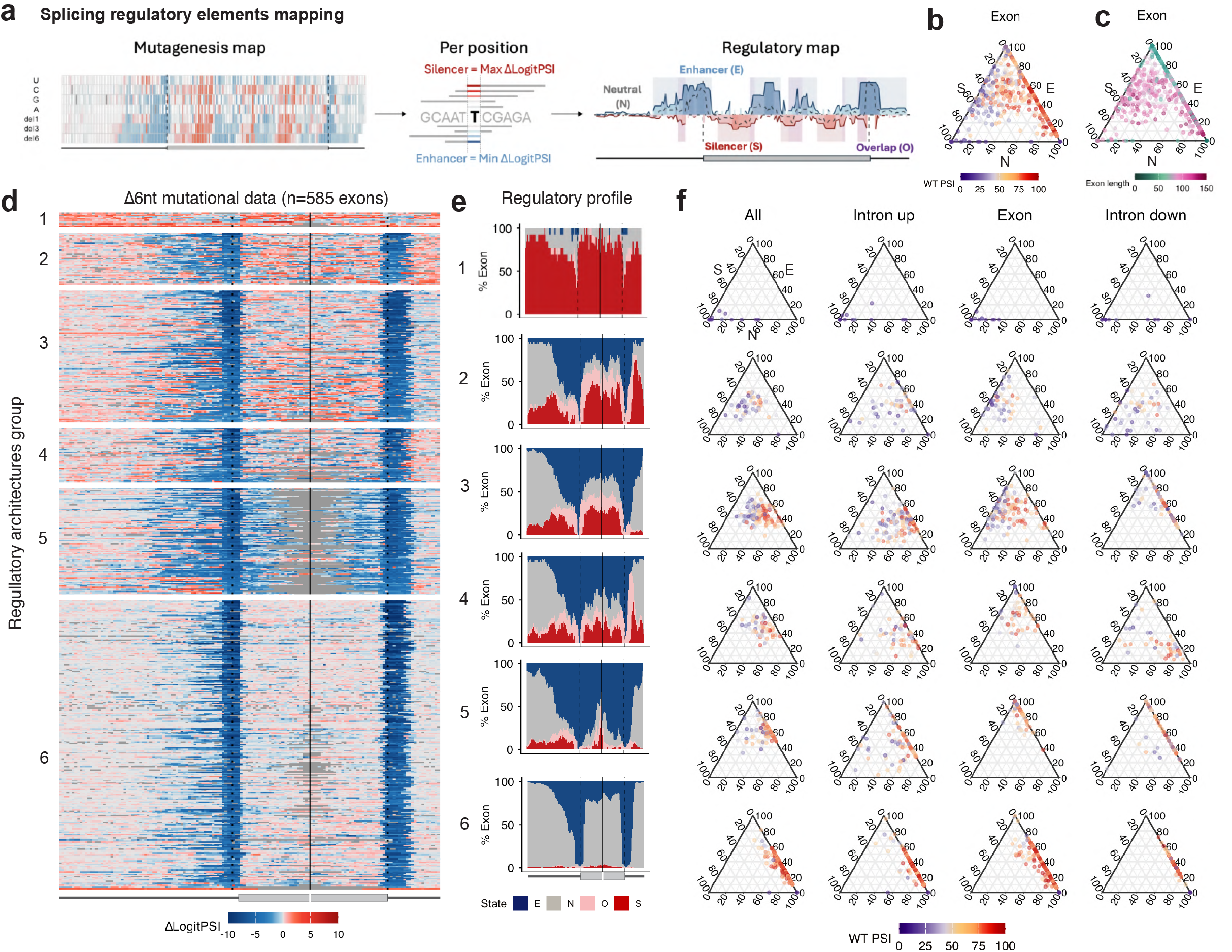
SpliceMaps reveal diverse regulatory architectures. **(a)** Schematic of the SpliceMap generation pipeline using *EHBP1* exon 17 as example. For each position, the minimum (most negative = Enhancer) and maximum (most positive = Silencer) ΔLogitPSI across all mutations are extracted. Stretches of ≥ 4nt with minimum ΔLogitPSI < −1 are classified as Enhancer (E); stretches with maximum ΔLogitPSI > 1 as Silencer (S); regions meeting both criteria as Overlap (O); the remainder as Neutral (N). **(b)** Ternary plot showing the proportion of Enhancer (E), Silencer (S), and Neutral (N) nucleotides per exon (exon body only), colored by WT PSI. **(c)** Same ternary plot as (b), colored by exon length. **(d)** Heatmap of 6nt deletion data for all 585 exons (rows), ordered by regulatory architecture group (1-6). Columns represent positions across the upstream intron, exon, and downstream intron, aligned at the splice sites. The color scale ranges from blue (increased skipping) to red (increased inclusion). **(e)** Average regulatory profiles (% nucleotides in each state: E = blue, N = grey, O = pink, S = red) per group (1-6), shown across the upstream intron, exon, and downstream intron. Vertical lines indicate splice site positions. **(f)** Ternary plots showing the E/S/N composition per exon for each group (rows 1-6), separately for all regions combined, upstream intron, exon body, and downstream intron (columns). Points are colored by WT PSI.

Using this approach identified 4,793 putative enhancers with median length = 6 nt, 1,473 putative silencers with median length = 6 nt, 3,100 neutral regions with median length = 9 nt, and 1,852 overlap regions with median length = 3 nt (**Extended Data Fig. 7a-b, Supplementary Table 7**). In total, 37,290 nts are classified as E (34.5%), 11,188 as S (10.3%), 6,726 as O (6.2%), and 52,881 as N (48.9%) (**Extended Data Fig. 7b**). The larger number of E than S nts is also seen for exons with intermediate inclusion (WT PSI = 20-80, n exons = 327, E= 38.4% and S = 11.1%, **Extended Data Fig. 7b**).

The regulatory state composition differs across transcript regions: exons have a heterogeneous architecture, with 33% enhancer, 13% silencer, 8% overlapping, and 44% neutral nucleotides (**Extended Data Fig. 7b**). Upstream and downstream introns are dominated by neutral sequence (59% and 63% N state, respectively), with silencers and enhancers similarly abundant in the two intron locations (S: 9% upstream and 8.5% downstream; E: 28% and 25%, respectively) (**Extended Data Fig. 7b**). In the upstream intron, the majority of the neutral nucleotides are located in the distal region (85.8% N in nt 1–26), and this percentage decreases when moving toward the 3′ splice site (BP region, nt 27–51: 39.4% N state; PPT region, nt 52–66: 19.4% N state) (**Extended Data Fig. 7c**).

### Regulatory-state composition and transitions

The number of nucleotides classified as S and O decreases with increasing WT PSI across all regions (exon: S, R = −0.49; O, R = −0.52; upstream intron: S, R = −0.57; O, R = −0.51; downstream intron: S, R = −0.45; O, R = −0.38; all p < 2.2 × 10^−16^) (**Extended Data Fig. 7d**). In contrast, the number of N increases with WT PSI in exons (R = 0.6), as well as in the upstream (R = 0.59) and downstream introns (R = 0.48) (**Extended Data Fig. 7d**). E content shows only a weak negative association with exon inclusion within the exon (exon: R = −0.14) but no significant relationship when considering the introns (upstream: R = -0.09, downstream: R =0.01) (**Extended Data Fig. 7d**). Exon inclusion is therefore more strongly associated with depletion of silencer elements and increased neutral content than with enrichment of enhancer elementst, suggesting that silencer and neutral content are major determinants of whether an exon is alternatively or highly included^15^ (**Fig. 5B; Extended Data Fig. 7d,e**).

We next considered the arrangement of regulatory states. The total number of state transitions decreases with increasing WT PSI (R = −0.5, p < 2.2 × 10^−16^), indicating that exons with low and intermediate inclusion have more segmented regulatory maps than highly included exons (**Extended Data Fig. 7f**). This decrease is strongly associated with transitions involving silencer states: the number of transitions to or from S states is highly correlated with the total number of transitions (R = 0.88, p < 2.2 × 10^−16^ **Extended Data Fig. 7g**), and both S/E and S/N transitions decrease with increasing WT PSI (R = −0.59 and R = −0.49 p < 2.2 × 10^−16^, respectively, **Extended Data Fig. 7h**), whereas E/N transitions moderately increase (R = 0.36, p < 2.2 × 10^−16^, **Extended Data Fig. 7h**), in agreement with computational predictions^15^. The number of transitions per 100 nt shows no correlation with exon length (R = −0.05, p > 0.01, **Extended Data Fig. 7i**). Alternatively included exons have complex regulatory maps with interspersed silencers and enhancers. In contrast, highly included exons have fewer transitions and are dominated by E/N organization.

### Exons differ in the use of exonic, upstream and downstream regulatory elements

Clustering the exons by the proportion of nucleotides in the E/O/S/N state in the upstream intron, exon, and downstream intron, identified six groups of exons (**Fig. 5d, Extended Data Fig. 8a-d, Supplementary Table 8**). Group 1 contains exons with very low inclusion (n=13 exons, median WT PSI = 1%) with silencer-dominated regulatory maps (median S = 144 nt, 81.1%) (**Fig. 5e-f, Extended Data Fig. 9a,c**). These exons have a median of 6.6 transitions per 100 nt (**Extended Data Fig. 9b**). The majority of transitions are between silencer and neutral states.

Groups 2-4 comprise alternatively spliced exons with similar inclusion levels, splice site strength distributions, overall regulatory-state composition, and total transition densities (median WT PSI: 35.5%, 55.5%, and 63.2%; median transitions per 100 nt: 12.7, 11.8, and 10.6 for Groups 2-4, respectively) (**Extended Data Fig. 9a-c**). The three groups differ in how the states are distributed across regions (**Fig. 5e-f, Extended Data Fig. 9c**). Within exons, Group 2 has similar contributions of enhancer and silencer sequence (median E = 30.9%, S = 35.6%), whereas Groups 3 and 4 are progressively more enhancer-enriched (Group 3: E = 39.6%, S = 25.3%; Group 4: E = 54.1%, S = 17.7%) (**Extended Data Fig. 9C**). In the downstream intron, Group 2 is enriched in silencer sequence (S = 36%), Group 3 is dominated by enhancer and neutral sequence (E = 52%, N = 40%), whereas Group 4 has a more balanced contribution of enhancer and silencer states (E = 16%, N = 14%) together with a substantial overlap fraction (∼20%) (**Extended Data Fig. 9C**).

Groups 5 and 6 contain highly included exons (median WT PSI: 64.2% and 83.2%, respectively) but differ in exon length, regulatory composition, and transition density (**Extended Data Fig. 9a-c**). Group 5 consists of short exons (median length = 42nt) with enhancer-rich sequences (E = 56%; N = 35.1%), whereas Group 6 contains longer exons (median length = 109 nt) with predominantly neutral sequence (E = 26.9%; N = 72.8%, **Extended Data Fig. 9c**). Consistent with this difference in composition, Group 5 shows a higher transition density than Group 6 (median 6.3 vs 4 transitions per 100 nt, **Extended Data Fig. 9b**). Regional differences in regulatory composition further distinguish these groups (**Fig. 5e-f**). In the upstream intron, Group 5 has a higher fraction of enhancer sequence (median ∼40%) compared to Group 6 (∼26%, **Extended Data Fig. 9c**), with enhancer elements enriched near the proximal 3′ splice site region in Group 5 (BP region 27-51nt: 63.4%, PPT region 52-66nt: 74.2%). This suggests that exon inclusion in the longer Group 6 exons is determined more by core splice site signals (**Extended Data Fig. 8d**).

Together, these results indicate that exons with similarly high inclusion levels can be supported by distinct regulatory architectures, with shorter exons relying on higher enhancer density despite a comparable number of enhancer nucleotides, and longer exons associated with sparser regulatory landscapes (**Fig. 5c, Extended Data Fig. 7j-l**).

### Splicing regulatory architectures are diverse

Despite similar percentages of nucleotides in each regulatory state, the exons within each group differ extensively in the number, length, and spatial distribution of individual regulatory elements. For example, *SCN5A* exon 6 and *CCDC112* exon 8 are both in group 3 (**Fig. 6A - 3**) and have similar WT PSI (50% and 45%, respectively) and similar overall state composition (*SCN5A* exon 6: E = 72nt, 38%; N = 51nt, 27.2%; O = 13nt, 6.9%; S = 51 nt, 27.3%; *CCDC112* exon 8: E = 75nt, 39.3%; N = 69nt, 36.1%; O = 15nt, 7.9%; S = 32 nt, 16.8%). However, they differ in the spatial organization of these states. In the upstream intron, *SCN5A* exon 6 contains multiple regulatory elements with distal silencers at 13–24 nt and 27–33 nt and a more proximal silencer at 43-54 nt, separated by neutral segments. The proximal 3′ splice site region is predominantly in the enhancer state (55–58 nt and 62–70 nt), corresponding to the branch point and polypyrimidine tract. In contrast, in *CCDC112* exon 8, upstream intronic regulatory elements are concentrated in the proximal region near the 3′ splice site (40–66 nt), whereas the distal region is largely neutral (1–39 nt). Within the exon, *SCN5A* exon 6 has higher enhancer density than *CCDC112* exon 8 (67.3% vs 48.9%). This difference is not due to the number of elements (4 vs 5) or median length (8 in both), but instead reflects a longer central region in *SCN5A* (35nt). Silencer organization also differs: *SCN5A* exon 6 contains three short silencers (median = 4nt), whereas *CCDC112* exon 8 contains three silencers with a median length of 11 nt.

**Figure 6.**
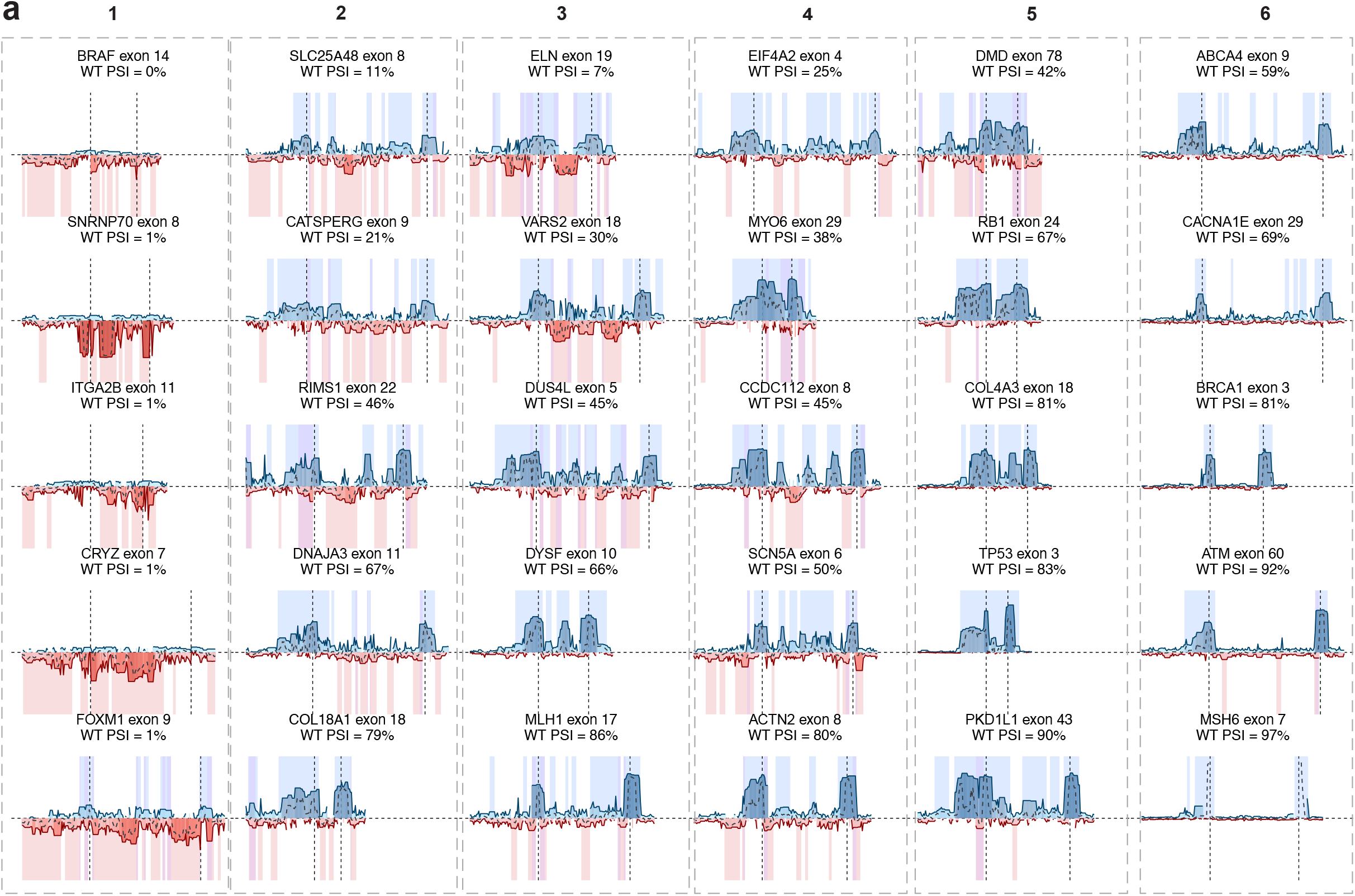
Example SpliceMaps. **(a)** SpliceMaps for five representative exons from each of the six regulatory architecture groups (columns 1-6, rows represent individual exons per cluster). Each SpliceMap displays the per-position regulatory state (Enhancer E, Silencer S, Overlap O, Neutral N) across the upstream intron, exon, and downstream intron, derived from combined substitution and short-deletion (1-, 3-, 6-nt) mutagenesis data. Exon boundaries are marked with dashed lines. Exons shown are Group 1: *GPAA1* exon 4 (0%), *BRAF* exon 14 (1%), *SNRNP70* exon 8 (1%), *INTS3* exon 27 (5%), *COL4A5* exon 12 (4%); Group 2: *ELN* exon 19 (7%), *PAX* exon 6 (10%), *RBM15* exon 2 (35%), *KDM6A* exon 16 (55%), *NCAM1* exon 9 (71%); Group 3: *SLC25A48* exon 8 (11%), *DMD* exon 78 (42%), *CCDC112* exon 8 (45%), *SCN5A* exon 6 (50%), *MLH1* exon 17 (86%); Group 4: *TMEM161B* exon 2 (14%), *VARS2* exon 18 (30%), *BBS1* exon 5 (45%), *RYR1* exon 8 (68%), *EYA4* exon 19 (82%); Group 5: *ABCA4* exon 9 (59%), *HPS1* exon 11 (72%), *POLE* exon 11 (71%), *COL1A1* exon 8 (77%), *SLC26A4* exon 15 (94%); Group 6: *RET* exon 19 (60%), *CACNA1E* exon 29 (69%), *BRCA1* exon 3 (81%), *ATM* exon 60 (92%), *MSH6* exon 7 (97%).

Similar diversity is also observed among highly included exons. For example, *MSH6* exon 7 (group 6; WT PSI = 97%; exon length = 90 nt; **Fig. 6a - 6**) has a sparse architecture dominated by neutral sequence (78.4% N, 21.6% E), with regulatory elements restricted to enhancer located in the nearby the 3’ splice site (47-50nt) and at the splice junctions (a 21-nt enhancer in the 3’ splice site region and a 15-nt enhancer spanning the end of the exon - 5′ splice site - beginning downstream intron). In contrast, *PKD1L1* exon 43 (group 5; WT PSI = 90%; exon length = 86nt; **Fig. 6a - 5**) has an enhancer-rich architecture (66.1% E, 25.6% N), with multiple enhancer segments (n = 7) distributed across both the upstream intron and the exon, interspersed with neutral segments.

### Clinical variants

The identification of splice-disrupting variants outside canonical splice sites is a major challenge in clinical variant interpretation^42,43^. Current American College of Medical Genetics (ACMG) guidelines only allow computational predictions to be used as supporting evidence for variant interpretation, necessitating the experimental testing of putative splice-altering variants located outside of canonical splice sites^7^. In total, OpenSplice includes 11,623 ClinVar variants, comprising 1,246 splice-site, 5,841 intronic and 4,536 synonymous variants. Pathogenic and likely pathogenic variants are strongly enriched for large decreases in exon inclusion (ΔLogitPSI < -1) in OpenSplice, including 98.3% of splice-site variants, 83% of intronic variants and 96% of synonymous variants. By contrast, only 4.7% of benign intronic variants and 5.7% of benign synonymous variants showed similarly strong effects. Variants of uncertain significance or conflicting interpretation showed intermediate behaviour, with ΔLogitPSI < -1 for 94.8% of splice-site, 41% of intronic and 20.3% of synonymous variants (**Fig. 7a; Extended Data Fig. 10a; Supplementary Table 9**).

**Figure 7.**
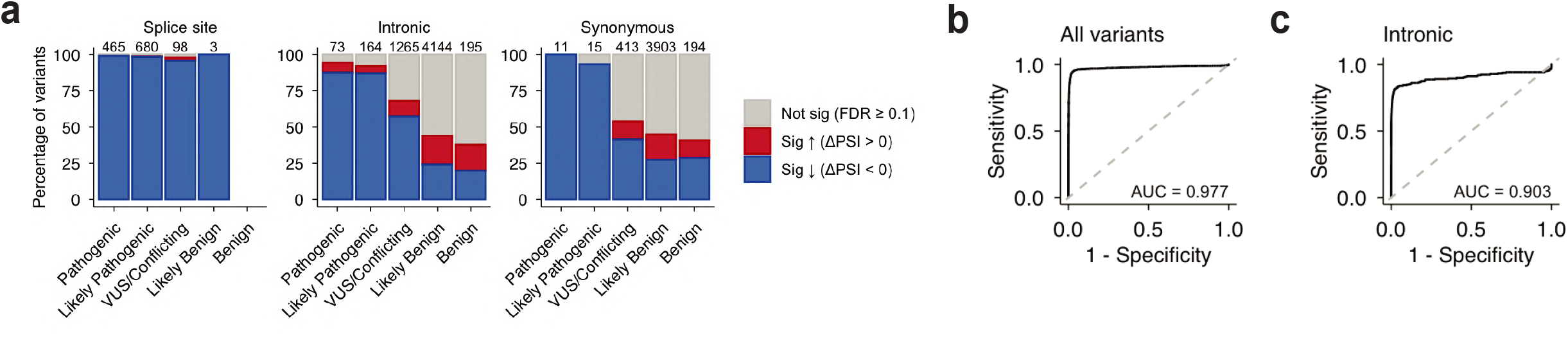
OpenSplice accurately classifies clinical variants. **(a)** Stacked bar plots showing the distribution of splicing effects for ClinVar variants stratified by clinical classification (Pathogenic, Likely Pathogenic, VUS/Conflicting, Likely Benign, Benign) and variant category: splice-site variants (left), intronic variants (middle), and synonymous variants (right). Bars are colored by significance and direction of effect: not significant (FDR = 0.1, grey), significant increase in inclusion (ΔPSI > 0, red), and significant decrease in inclusion (ΔPSI < 0, blue). Numbers above bars indicate variant counts per category. **(b)** ROC curve for classification of (Likely) pathogenic versus (Likely) benign ClinVar variants using OpenSplice ΔLogitPSI values across all variant types combined (AUC = 0.977). **(c)** ROC curve for classification of (Likely) pathogenic versus (Likely) benign ClinVar variants restricted to intronic variants outside canonical splice site dinucleotides (AUC = 0.903).

Across variants with clinical classifications, OpenSplice accurately discriminates pathogenic and likely pathogenic (n = 1,408) from benign and likely benign (n = 8,439) variants (ROC–AUC = 0.977; **Fig. 7b**). Performance remains high when restricting to intronic variants outside canonical splice sites (ROC–AUC = 0.903; n = 237 (likely) pathogenic and n = 4,339 (likely) benign; **Fig. 7c**). This illustrates the power of OpenSplice experimental data to extend clinically actionable interpretation beyond canonical splice sites.

## Discussion

We have presented here OpenSplice, a large and well-calibrated experimental resource that quantifies the impact of more than 590,000 variants on the splicing of 608 human exons. By expanding saturation mutagenesis data by more than an order of magnitude, OpenSplice provides high-resolution functional maps of splicing regulatory elements and a resource for the interpretation of clinical variants.

Hundreds of thousands of tested variants altered exon inclusion, highlighting the remarkable sensitivity of splicing to sequence perturbation^5,12^. Importantly, this sensitivity extends beyond canonical splice sites and encompasses both exonic and intronic regions, reinforcing the view that splicing is not determined solely by core splice signals but instead reflects the combined effects of numerous auxiliary regulatory elements distributed across a pre-mRNA^1,30^.

Mutational effects were well predicted by state-of-the-art machine learning models, demonstrating the power of these models to learn complex regulatory landscapes. There is, however, still substantial room for improvement, with large synthetic experimental datasets an important strategy to extend training data beyond natural sequences^44–46^. Clinical variant interpretation requires experimental evidence^7^ and OpenSplice provides a large atlas of splice-altering variants for this purpose, accurately classifying known pathogenic variants. Expanding OpenSplice to cover the entire human transcriptome, or to a size sufficient to train highly accurate predictive models, is an important goal for future research. An additional challenge for future work will be the expansion of site-saturation mutagenesis into deep intronic regions.

Beyond variant interpretation, OpenSplice also provides the largest systematic analysis of splicing regulatory architectures. Saturation mutagenesis resolved splice sites, polypyrimidine tracts and branch points, with mutational patterns indicating ∼40% of exons use a single major branch point. Reducing exon length reduced inclusion, particularly below 30nt, consistent with geometric constraints on spliceosome assembly^27^. Short exons were more sensitive to both exonic and intronic mutations, demonstrating that they require denser regulatory information to ensure efficient recognition.

One striking insight from these mutational maps is the sheer diversity of splicing regulatory architectures. Exons differ substantially in how regulatory information is encoded. While some exons - particularly alternatively spliced exons - contain dense regulatory landscapes with multiple enhancer and silencer elements, others have sparse architectures in which most positions are functionally neutral and mutational sensitivity is largely restricted to splice sites^15^. Unexpectedly, splice-site strength explained only part of exon inclusion and did not predict the overall sensitivity of exons to mutation, with many highly included exons also dependent on additional regulatory elements. Rather, OpenSplice demonstrates that silencers are particularly important for determining exon inclusion levels.

An important future direction will be to extend OpenSplice to additional cell types. Cell type-specific variant effects on splicing are rare^47,48^ and not well predicted by computational models^49^. An additional important challenge for future work will be to mechanistically connect splicing cis regulatory architectures to the trans factors that bind them. Combining systematic trans factor perturbation^50^ with saturation mutagenesis libraries may be one method to achieve this^11^.

## Extended Data Figure legends

**Extended Data Figure 1.**
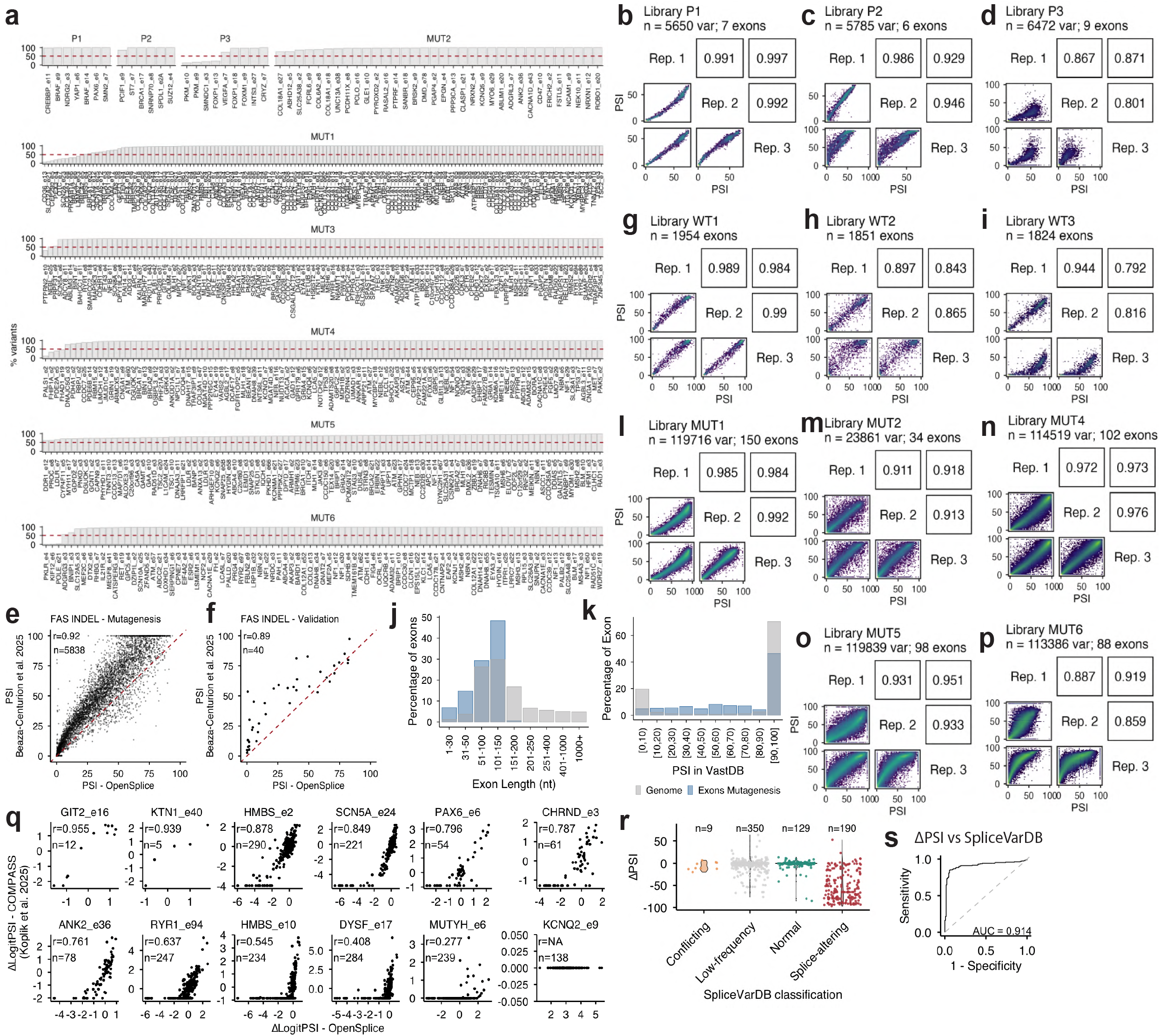
Reproducibility and validation of the OpenSplice experimental platform. **(a)** Percentage of designed variants recovered per exon across the three pilot libraries (P1, P2, P3) and six mutagenesis libraries (MUT1-6). The dashed red line indicates the 50% recovery threshold. **(b–d)** Pairwise scatter plots of PSI values across three biological replicates for pilot libraries P1 (n = 5,628 variants in 7 exons; **b**), P2 (n = 5,634 variants in 6 exons; **c**), and P3 (n = 6,472 variants in 9 exons; **d**). Pearson r values are shown for each replicate pair. **(e)** Correlation between PSI values measured by OpenSplice and by the FAS indel mutagenesis dataset^15^ (r = 0.92, n = 5,838). **(f)** Correlation between PSI values measured by OpenSplice saturation mutagenesis and individual single-clone validation experiments for variants from the FAS indel library^15^ (r = 0.89, n = 40). **(g–i)** Pairwise replicate scatter plots for the three wild-type exon screening libraries: WT1 (n = 1,954; **g**), WT2 (n = 1,851; **h**), and WT3 (n = 1,824; **i**). Pearson r values are indicated for each replicate pair. **(j)** Distribution of exon lengths for all human genome exons (grey) and the 608 exons selected for mutagenesis (blue). **(k)** Distribution of PSI values in the VastDB database for all annotated events (grey) and the 608 exons selected for mutagenesis (blue). **(l–p)** Pairwise replicate scatter plots for mutagenesis libraries MUT1 (n = 109,533; **l**), MUT2 (n = 23,038; **m**), MUT4 (n = 114,519; **n**), MUT5 (n = 103,269; **o**), and MUT6 (n = 113,386; **p**). Pearson r values are shown for each replicate pair. **(q)** Pairwise scatter plots of ΔLogitPSI values from OpenSplice versus the COMPASS dataset (Koplik et al. 2025) for 12 individual exons with overlapping variant measurements. Pearson r and n values are shown for each exon. **(r)** Distribution of ΔPSI values from OpenSplice for variants classified in SpliceVarDB as conflicting (n = 9), low frequency (n = 350), normal/neutral (n = 129), and splice-altering (n = 190). **(s)** ROC curve for classification of splice-altering versus neutral variants in SpliceVarDB using OpenSplice ΔPSI values (AUC = 0.914).

**Extended Data Figure 2.**
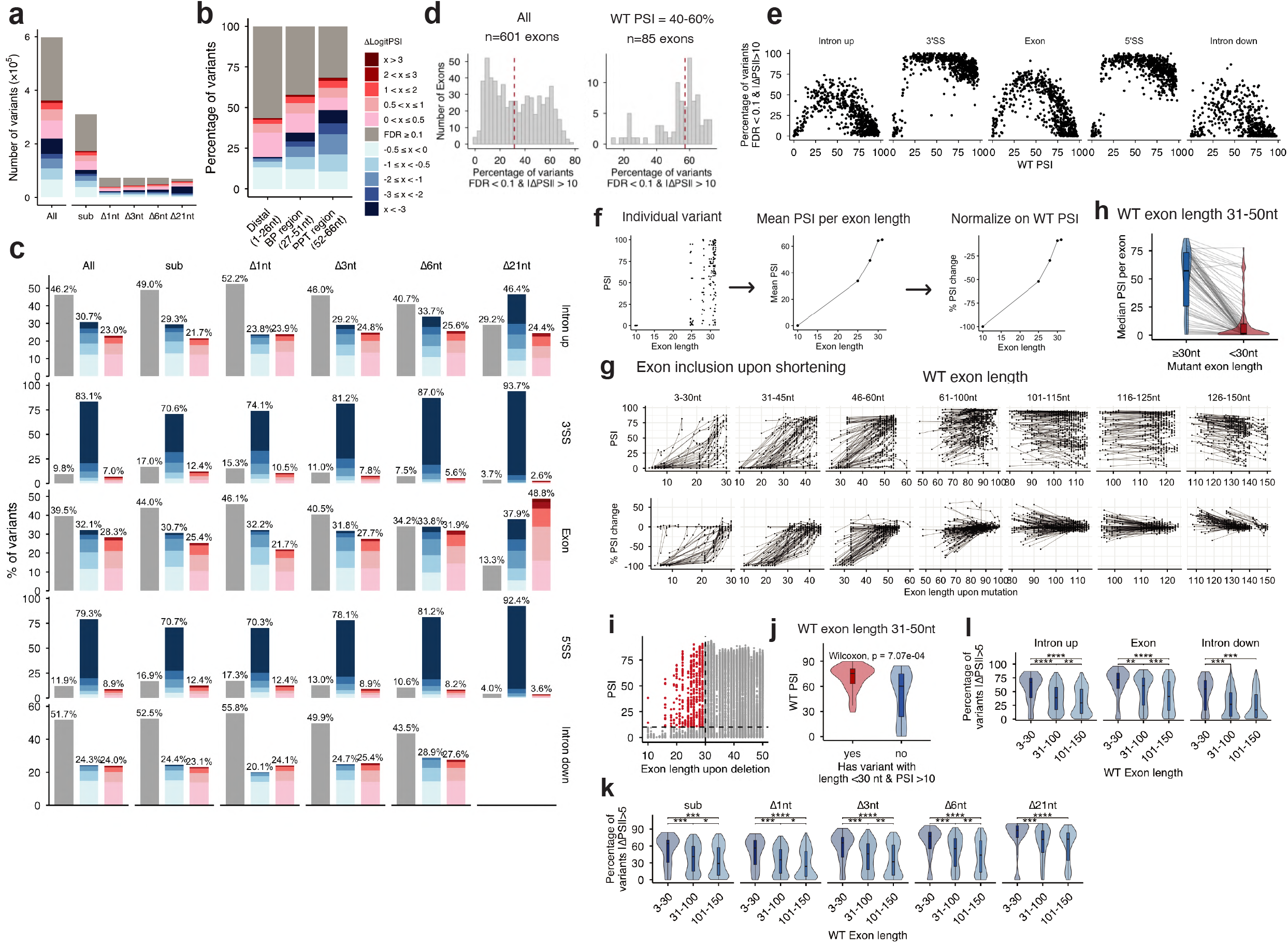
Distribution of variant effects and exon-length-dependent splicing sensitivity. **(a)** Total number of variants per mutation class (substitutions, 1-,3-6-,21-nt deletions) stratified by ΔLogitPSI magnitude and direction. Grey indicates variants not reaching significance (FDR = 0.1); blue shades indicate decreased inclusion; red shades indicate increased inclusion. **(b)** Stacked bar plots showing the percentage of variants causing significant increases or decreases in exon inclusion (FDR = 0.1), stratified by upstream intron region (distal intron, BP region, PPT region) and mutation class. **(c)** Stacked bar plots showing the percentage of variants causing significant increases or decreases in exon inclusion (FDR = 0.1), stratified by genomic region (rows: upstream intron, 3’SS, exon, 5’SS, downstream intron) and mutation class (columns: all, substitutions, 1-,3-6-,21-nt deletions). Percentages for each ΔLogitPSI bin are indicated. **(d)** Distribution of the percentage of non-neutral variants per exon across all exons (left, n=601) and exons with WT PSI = 40-60% (right; n=85 dashed red line = median). **(e)** Relationship between median |ΔPSI| per exon and WT PSI, shown separately for each genomic region (upstream intron, 3’SS, exon, 5’SS, downstream intron). **(f)** Schematic illustrating the three-step normalization used to quantify the effect of exon shortening (left: individual variant PSI values; middle: mean PSI per resulting exon length; right: % PSI change normalized to WT PSI). **(g)** Exon inclusion upon shortening across all WT exon length bins (3-30 nt, 31-45 nt, 46-60 nt, 61-100 nt, 101-115 nt, 116-125 nt, 126-150 nt). Top row: raw PSI values per resulting exon length; bottom row: % PSI change normalized to WT PSI. Each line represents one exon. **(h)** Violin plot comparing median PSI per exon when exon length upon deletion is =30 nt or <30 nt in exons with WT length = 31-50 nt. Each grey line is an exon. **(i)** Scatter plot of WT PSI versus resulting exon length for variants retaining PSI > 10 at lengths below 30 nt (red dashed box), highlighting rare exceptions to the microexon inclusion threshold. **(j)** Violin plot showing the WT PSI distribution of exons that contain at least one variant maintaining PSI > 10 at lengths below 30 nt (Wilcoxon test, p = 7.07 x 10-4). **(k)** Violin plots comparing median |ΔPSI| per exon across WT exon length bins (3-30, 31-100, 101-150 nt) for intronic variants (upstream intron, exon, downstream intron). **(l)** Violin plots comparing median |ΔPSI| per exon across WT exon length bins (3-30, 31-100, 101-150 nt) for each mutation class (substitutions, 1-,3-6-,21-nt deletions).

**Extended Data Figure 3.**
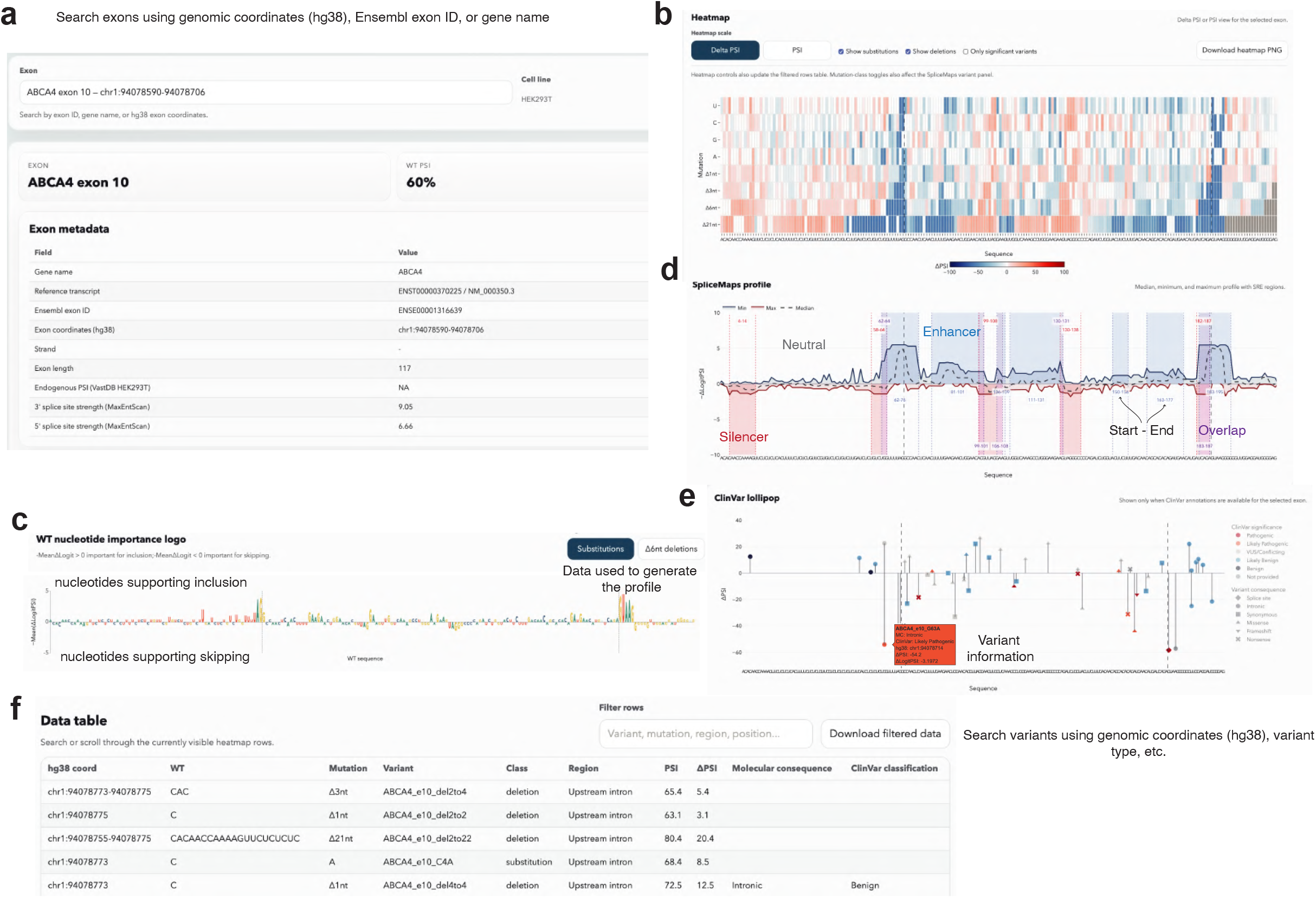
ExonExplorer interactive web interface. **(a)** Exon metadata panel for a selected exon, displaying gene name, reference transcript, Ensembl exon ID, genomic coordinates (hg38), exon length, endogenous PSI from VastDB, and 3’/5’ splice site strengths (MaxEntScan). Exons can be searched by genomic coordinates (hg38), Ensembl exon ID, or gene name. **(b)** Interactive mutational heatmap showing ΔPSI or PSI values for all variants in the selected exon, with options to display substitutions only, deletions only, or only significant variants. The heatmap can be downloaded as PNG. **(c)** Wild-type nucleotide importance logo derived from the mean ΔLogitPSI of substitution data. Positions above the x-axis favor exon inclusion; positions below favour exon skipping. Colored by nucleotide identity (A, green; G, yellow; U, blue; C, red). Users can toggle between substitutions and 6nt deletion data. **(d)** SpliceMap profile displaying the median, minimum, and maximum ΔLogitPSI across all mutations at each position. Regulatory states (Enhancer = blue, Silencer = red, Overlap = purple) are shown as color-coded blocks below the profile, with start-end coordinates for each element. **(e)** ClinVar lollipop plot showing ΔPSI values for ClinVar-annotated variants at the selected exon, colored by clinical significance. Displayed only when ClinVar annotations are available. **(f)** Searchable and downloadable data table listing all variants for the selected exon with their genomic coordinates (hg38), wild-type nucleotide, mutation, variant identifier, class, region, PSI, ΔPSI, molecular consequence, and ClinVar classification. Rows can be filtered by variant, mutation type, region, or position.

**Extended Data Figure 4.**
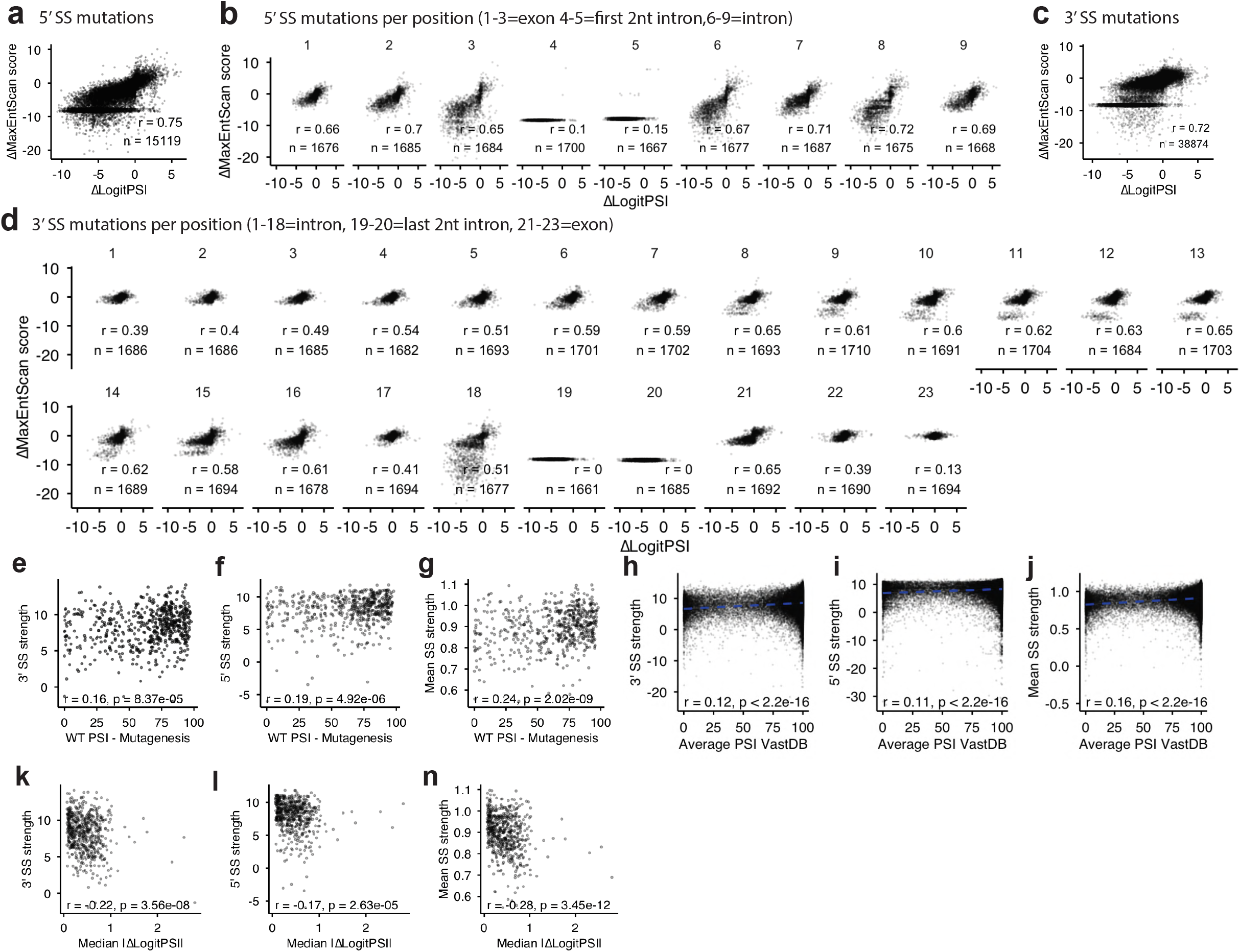
Splice site strength and its relationship to exon inclusion and mutational sensitivity. **(a)** Correlation between ΔMaxEntScan score and ΔLogitPSI for all 5’ splice site (SS) mutations (r = 0.75, n = 15,119). Mutation-induced changes in 5’SS strength are strongly predictive of changes in exon inclusion. **(b)** Correlation between ΔMaxEntScan score and ΔLogitPSI for 5’SS mutations at each individual position (positions 1-3 = exon; 4-5 = first 2 nt of intron; 6-9 = intron). Pearson r and n values are shown per position. **(c)** Correlation between ΔMaxEntScan score and ΔLogitPSI for all 3’ splice site (SS) mutations (r = 0.72, n = 38,874). **(d)** Correlation between ΔMaxEntScan score and ΔLogitPSI for 3’ splice site mutations at each individual position (positions 1-18 = intron; 19-20 = last 2 nt of intron; 21-23 = exon). Pearson r and n values are shown per position. **(e-g)** Scatter plots of 3’SS strength **(e)**, 5’SS strength **(f)**, and mean SS strength **(g)** versus WT PSI in the OpenSplice mutagenesis dataset. Weak but significant correlations are observed (r = 0.16 - p = 8.37 × 10^-5^, r = 0.19 - p = 4.92 × 10^-6^, r = 0.24 - p = 2.02 × 10^-9^). **(h-j)** Scatter plots of 3’SS strength **(h)**, 5’SS strength **(i)**, and mean SS strength **(j)** versus average PSI from VastDB for endogenous exons. Weak but significant correlations are observed (r = 0.12, 0.11, 0.16, respectively; p < 2.2 × 10^-16^). **(k-m)** Scatter plots of 3’SS strength **(k)**, mean SS strength **(l)**, and 5’SS strength **(m)** versus median |ΔLogitPSI| per exon (overall mutational sensitivity). Weak but significant correlations are (r = -0.22 - p = 3.56 × 10^-8^, r = -0.17 - p = 2.63 × 10^-5^, r = -0.28 - p = 3.45 × 10^-12^).

**Extended Data Figure 5.**
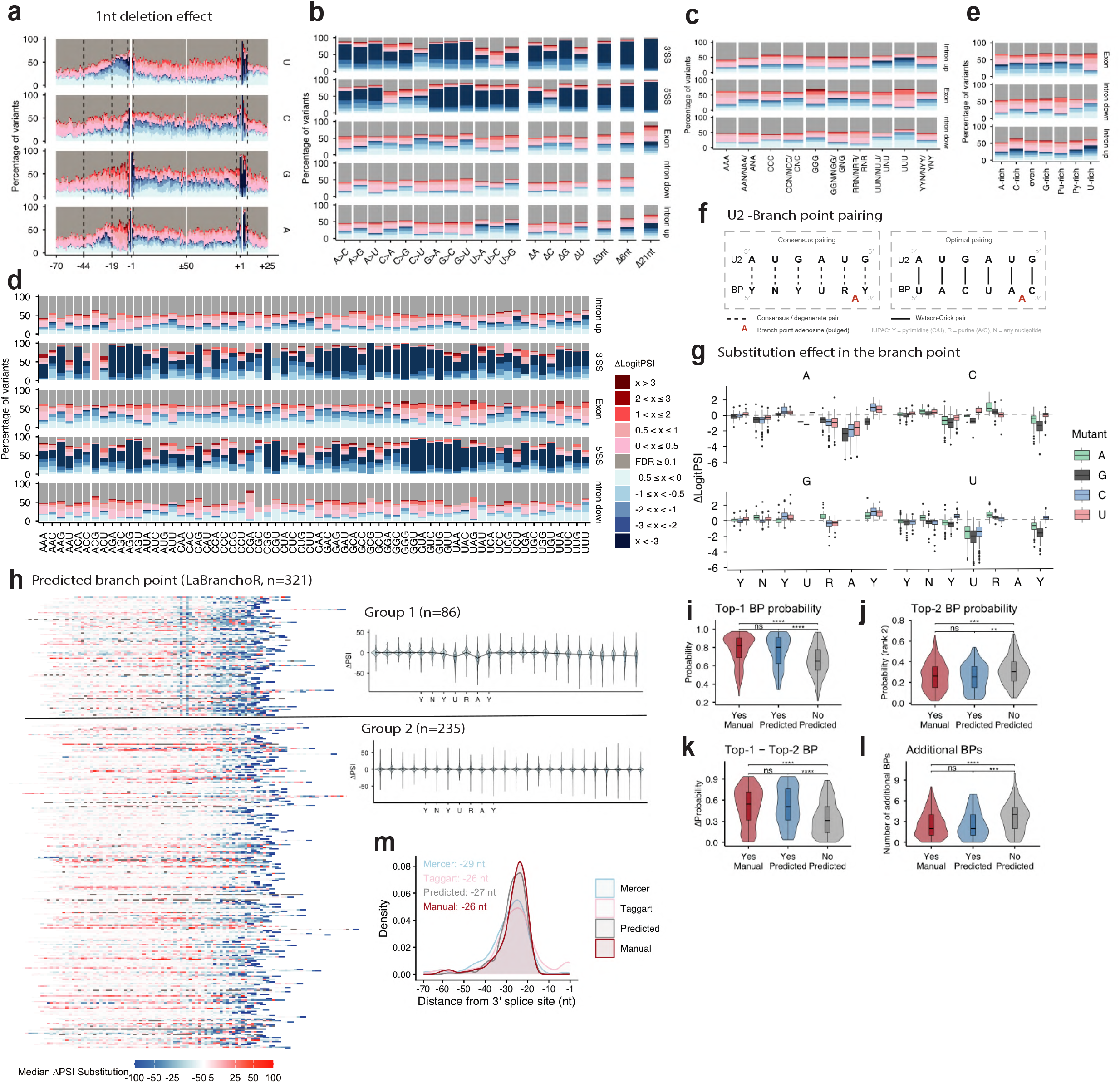
Nucleotide-specific and context-specific effects of mutations on splicing. **(a)** Mutational sensitivity profiles for 1nt deletions across the pre-mRNA (upstream intron: -70 to -1; exon: first and last 50 nt; downstream intron: +1 to +25), stratified by WT nucleotide identity (rows: U, C, G, A). The percentage of variants significantly increasing (red) or decreasing (blue) exon inclusion is plotted per position. **(b)** Stacked bar plots showing the percentage of substitutions and deletions causing significant changes in exon inclusion (FDR = 0.1), stratified by substitution type (rows: each specific nucleotide change) and genomic region (upstream intron, 3’SS, exon, 5’SS, downstream intron). **(c)** Effect of 3nt deletions on exon inclusion grouped by trinucleotide sequence composition (where N=A/C/G/U, R=A/G, Y=U/C), shown separately for upstream intron, exon, and downstream intron. Within exons, deletion of U-rich trinucleotides more frequently increased inclusion (UUU: 58.9% increase vs 10.1% decrease), whereas deletion of G-rich (GGN/NGG/GNG: 22.3% vs 40.3%), A-rich (AAN/NAA/ANA: 25.6% vs 35.2%), and C-rich (CCN/NCC/CNC: 19.4% vs 40.2%) trinucleotides more often decreased inclusion. **(d)** Effect of 3nt deletions on exon inclusion per trinucleotide class and genomic region (rows: upstream intron, 3’SS, exon, 5’SS, downstream intron; columns: trinucleotide classes). In the downstream intron, deletion of UUU predominantly decreased exon inclusion (50.4% decrease vs 9% increase), whereas deletion of purine-rich motifs (RRN/NRR/RNR, R=A/G) more often increased inclusion (39.9% vs 13.1%), consistent with U-rich intronic splicing enhancers and repressive purine-rich elements. **(e)** Effect of 6nt deletions on exon inclusion grouped by nucleotide composition (A-rich, C-rich, G-rich, even, Py-rich, U-rich), shown separately for upstream intron, exon, and downstream intron. Within exons, 6-nt deletions removing three or more U residues more often increased inclusion (50.1% vs 19% decrease), while deletions enriched in A and C content more often decreased inclusion. **(f)** Schematic illustrating U2 snRNA-BP base-pairing. Left: consensus pairing between U2 snRNA and the YNYURAY BP motif, with the BP adenosine (position 6) bulged out. Right: optimal pairing configuration showing the strongest predicted U2 snRNA complementarity. IUPAC notation: Y = pyrimidine (C/U), R = purine (A/G), N = any nucleotide. **(g)** ΔLogitPSI distributions for all substitutions at each position of the YNYURAY BP motif (positions 1-7), stratified by WT nucleotide (panels: A, C, G, U) and mutant nucleotide identity (colors: A, green; G, dark; C, blue; U, red). **(h)** heatmap of median ΔPSI substitution effects across the upstream introns of all 321 exons with LaBranchoR-predicted branch points, aligned at the predicted BP adenosine. Group 1 (n = 86, clear single-BP signature) and Group 2 (n = 235, no clear dominant BP) are shown separately. **(i–k)** Violin plots comparing top-ranked LaBranchoR BP probability **(i)**, second-ranked BP probability **(j)**, and Δ-probability (top minus second **(k)**) between manually annotated (yes), group 1 predicted (yes predicted), and group 2 (no predicted) introns. **(l)** Violin plots showing the number of additional BP-like 7-mers (within two mismatches of YNYURAY, excluding invariant positions) for manually annotated, group 1, and group 2 introns. **(m)** BP position distributions from Mercer^34^, Taggart^35^, predicted (LaBranchoR^36^), and manual datasets.

**Extended Data Figure 6.**
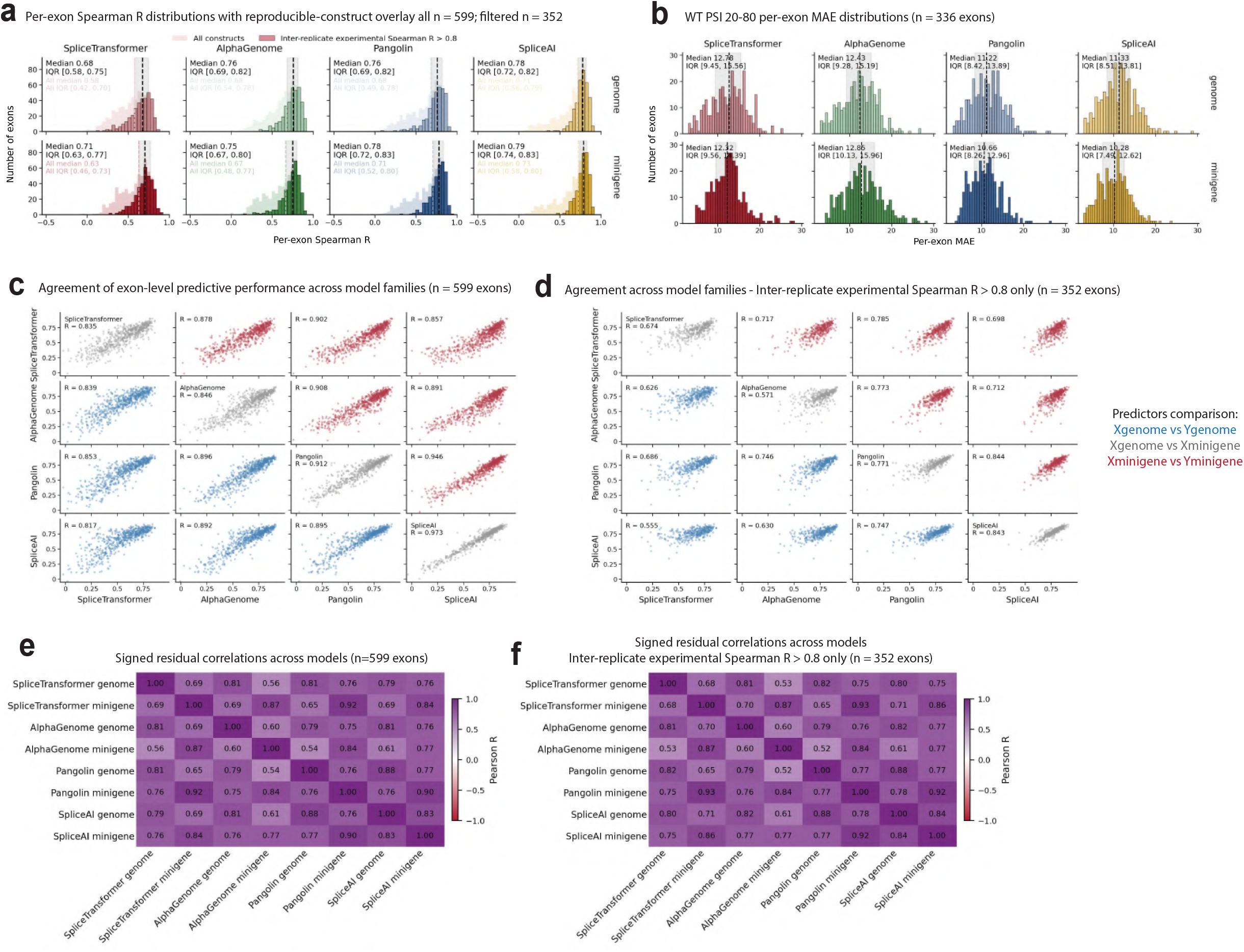
Model performance on alternatively spliced exons and correlated prediction errors. **(a)** Distributions of per-exon Spearman R between predicted and experimentally measured ΔPSI for four models (SpliceTransformer, AlphaGenome, Pangolin, and SpliceAI) run in genome mode (top row) or minigene mode (bottom row), across exons with experimental inter-replicate Spearman R>0.8 (n = 352 exons). **(b)** Distributions of per-exon mean absolute error (MAE, ΔPSI scale) for four splicing variant effect predictors (SpliceTransformer,AlphaGenome, Pangolin, SpliceAI) run in genome mode (top row) and minigene mode (bottom row), restricted to exons with WT PSI 20-80% (n = 336). Median MAE and IQR are indicated for each model. **(c)** Pairwise scatter plots of median per-exon Spearman R comparing all four models across two sequence contexts (n = 599 exons). Points are colored by comparison type: same-model genome versus minigene mode (grey), cross-model genome versus genome (blue), and cross-model minigene versus minigene (red). **(d)** same as **c** but considering only exons with experimental inter-replicate Spearman R>0.8 (n = 352 exons). **(e)** Heatmap of pairwise Pearson correlations between signed prediction residuals (predicted minus observed ΔLogitPSI) across all eight model-context combinations (genome and minigene modes for each of the four models). **(f)** same as (d) but considering only exons with experimental inter-replicate Spearman R>0.8 (n = 352 exons).

**Extended Data Figure 7.**
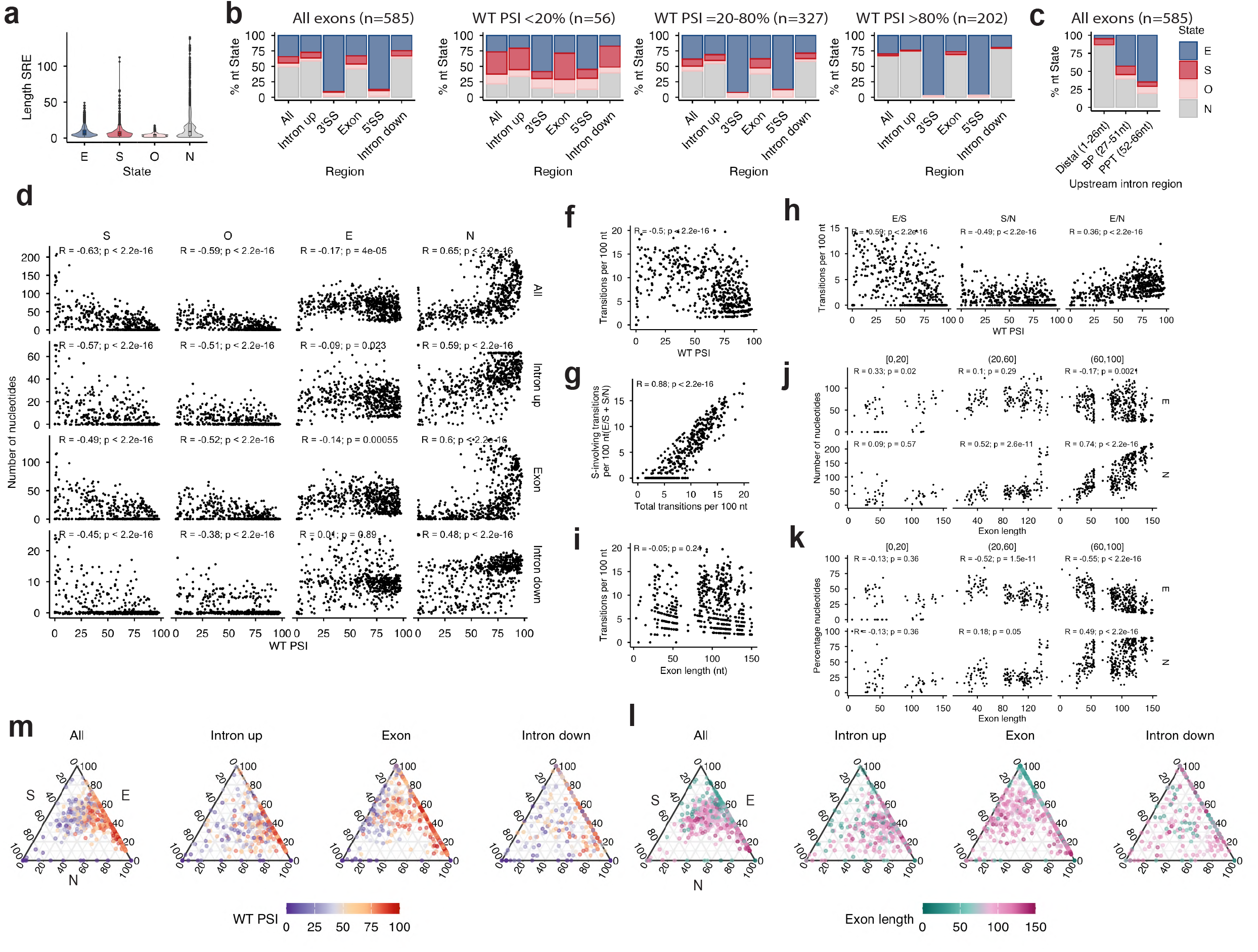
Regulatory state composition and transition architecture of SpliceMaps. **(a)** Violin plots showing the length distributions of splicing regulatory element segments classified as Enhancer (E), Silencer (S), Overlap (O), or Neutral (N) across all 585 exons. **(b)** Stacked bar plots showing the percentage of nucleotides in each regulatory state (E, S, O, N) per genomic region (all, upstream intron, 3’SS, exon, 5’SS, downstream intron), shown for all exons combined (n = 585) and stratified by WT PSI bins: <20% (n = 56), 20-80% (n = 327), and >80% (n = 202). **(c)** Regulatory state composition within the upstream intron subdivided into three functional sub-regions: distal (positions 1-26 nt from the 3’SS), BP region (27-51 nt), and PPT region (52-66 nt), for all 585 exons. **(d)** Scatter plots of the number of nucleotides in each regulatory state (S, O, E, N) versus WT PSI, shown separately for all regions combined and for each region (upstream intron, exon, downstream intron). Spearman R and p-values are indicated. **(e)** Ternary plots of E/S/N regulatory state composition per exon for all regions combined and for each individual region (upstream intron, exon, downstream intron), colored by WT PSI. **(f)** Scatter plot of the total number of regulatory state transitions per 100 nt versus WT PSI (R = -0.52, p < 2.2 × 10^-16^). **(g)** Scatter plot of the number of silencer-involving (S) transitions per 100 nt versus the total number of transitions per 100 nt (R = 0.88, p < 2.2 × 10^-16^) **(h)** Scatter plots of the number of E/S, S/N, and E/N transitions per 100 nt versus WT PSI. **(i)** Scatter plot of the number of transitions per 100 nt versus exon length (R = -0.09, p = 0.039). **(j)** Scatter plots of regulatory state composition (number of nucleotides) versus exon length, stratified by WT PSI bin ([0,20], (20,60], (60,100]), for E and N. **(k)** Same scatter plots as j but using percentage of nucleotides in E and N. **(l)** Same ternary plots as d, colored by exon length.

**Extended Data Figure 8.**
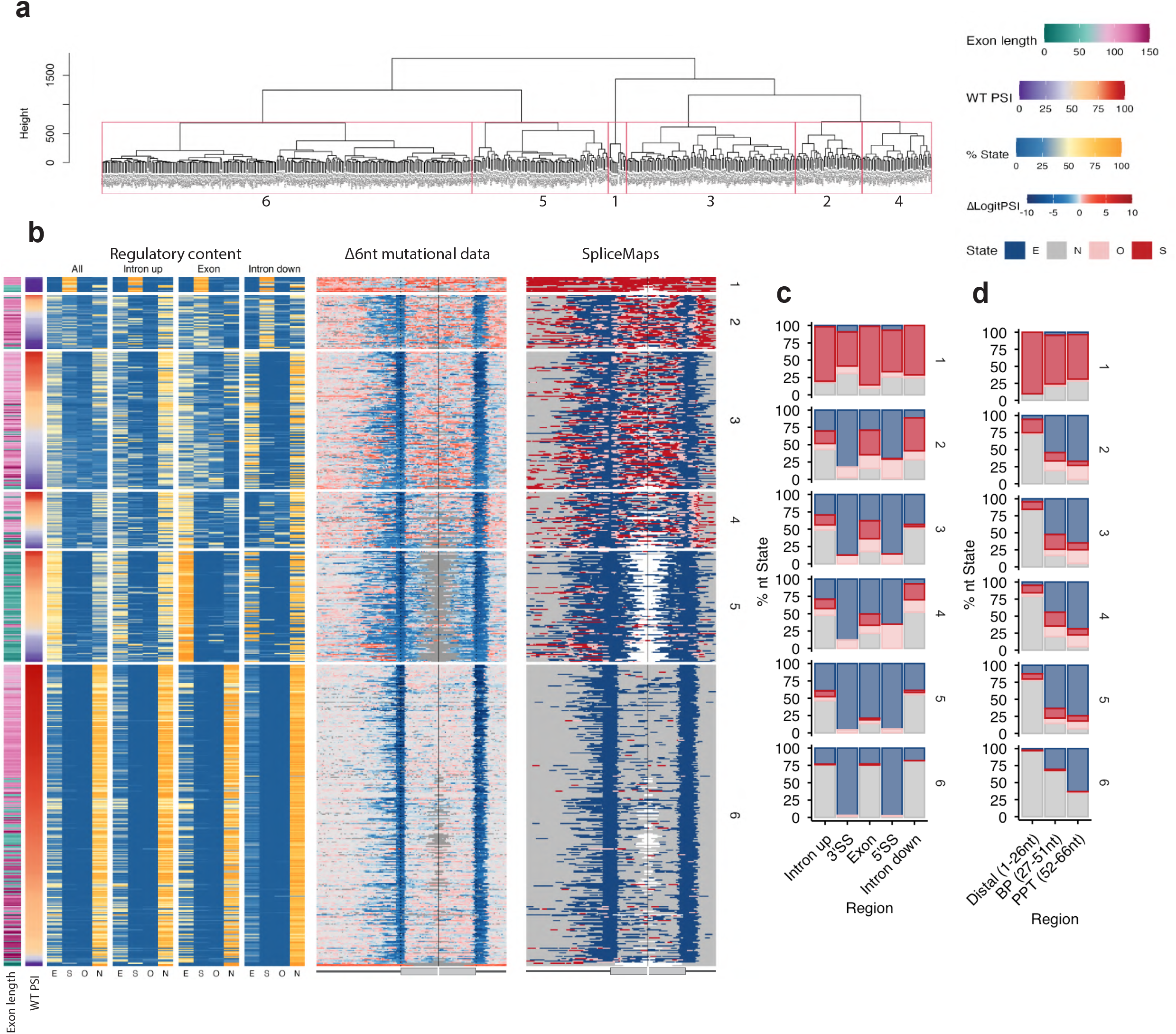
Per-cluster SpliceMap heatmaps and regional regulatory state composition. **(a)** Dendrogram from hierarchical clustering of 585 exons by regulatory state composition (E, S, O fractions across upstream intron, exon, and downstream intron). Six clusters are defined (red boxes). **(b)** Left: per-exon regulatory state composition (E, S, O, N) shown as color-coded bars for all regions combined and separately for upstream intron, exon, and downstream intron, with rows ordered by cluster (1-6). Centre: 6nt deletion mutational data for the same exons showing across upstream intron, exon, and downstream intron positions (blue: decreased inclusion; red: increased inclusion). Right: SpliceMap per exon. **(c)** Stacked bar plots showing the percentage of nucleotides in each regulatory state (E, S, O, N) per genomic region (all, upstream intron, 3’SS, exon, 5’SS, downstream intron) for each cluster (1-6). **(d)** same as **(c)** but focusing on upstream intron subdivided into the distal region (1-26 nt), BP region (27-51 nt), and PPT region (52-66 nt) for each cluster (1-6).

**Extended Data Figure 9.**
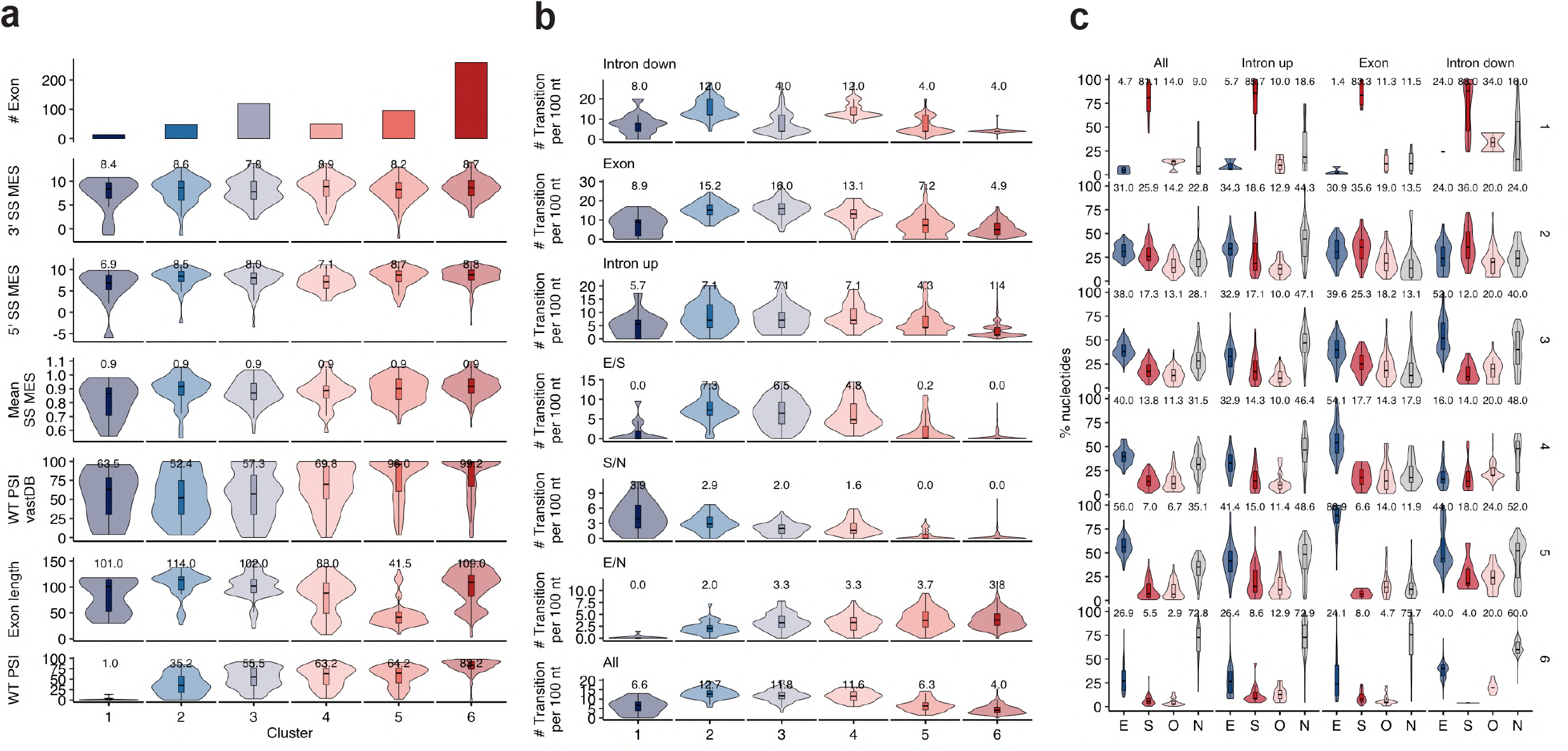
Characterization of regulatory architecture clusters. **(a)** Violin plots comparing key properties across the six regulatory architecture clusters: number of exons per cluster (bar chart), 3’SS MaxEntScan score, 5’SS MaxEntScan score, mean SS MaxEntScan score, WT PSI from VastDB, exon length, and WT PSI in the minigene system. Median values are indicated above each violin. **(b)** Violin plots of the number of regulatory state transitions per 100 nt for each cluster, shown separately for downstream intron, exon, upstream intron, and for each transition type (E/S, S/N, E/N, all). Median values are indicated. **(c)** Violin plots of the percentage of nucleotides in each regulatory state (E, S, O, N) per cluster, shown separately for all regions combined, upstream intron, exon, and downstream intron. Median percentages are annotated.

**Extended Data Figure 10.**
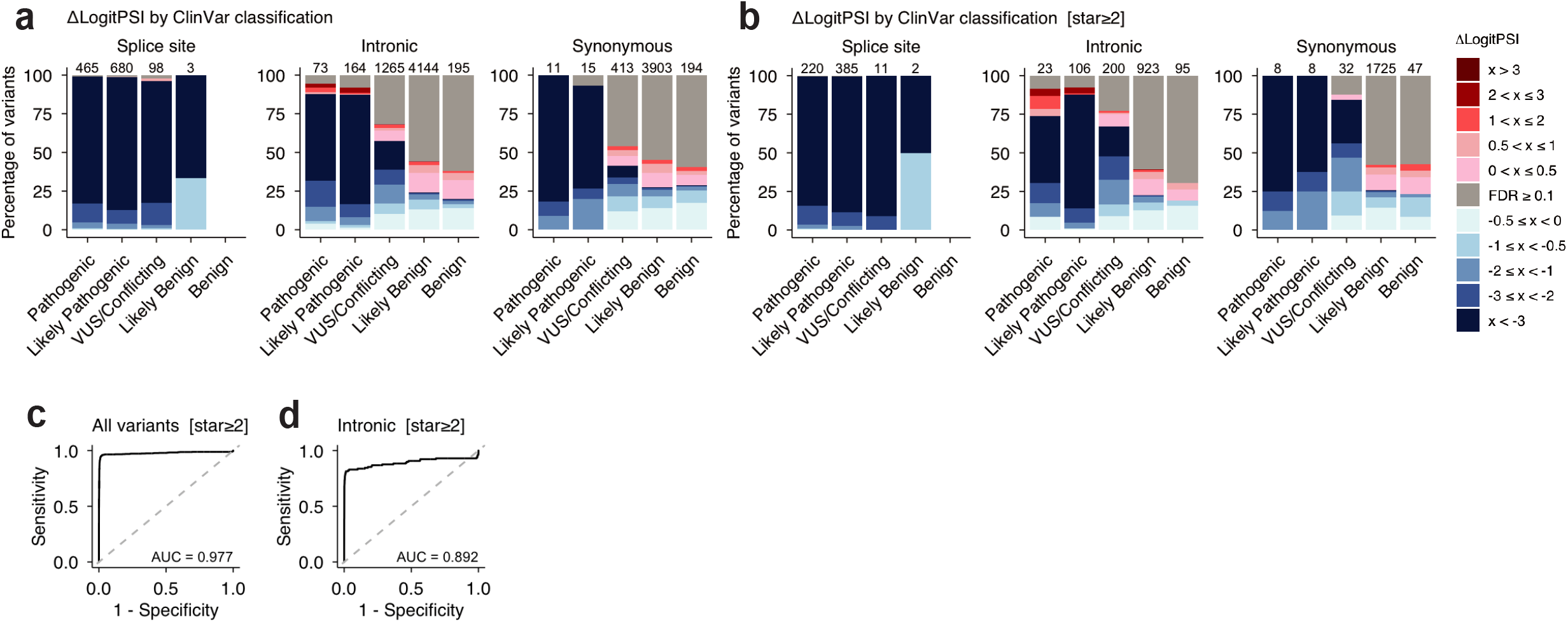
ClinVar variant classification with OpenSplice. **(a)** Stacked bar plots showing the distribution of ΔLogitPSI effects for all ClinVar variants, stratified by clinical classification (Pathogenic, Likely Pathogenic, VUS/Conflicting, Likely Benign, Benign) and variant category: splice-site, intronic and synonymous. Variant counts are shown above each bar. Colour scale indicates ΔLogitPSI magnitude and direction. **(b)** Same as **(a)**, restricted to higher-confidence ClinVar submissions with review status ≥ 2 stars. The separation between pathogenic and benign variant distributions is maintained. **(c)** ROC curve for classification of (Likely) pathogenic versus (Likely) benign ClinVar variants using OpenSplice ΔLogitPSI, restricted to ≥ 2 stars submissions, all variant types combined (AUC = 0.977). **(d)** ROC curve as in **(c)**, restricted to intronic variants outside canonical splice sites in ≥ 2 stars ClinVar submissions (AUC = 0.892).

## Supplementary Figure legends

**Supplementary Figure 1. Complete mutational maps for all 608 exons**.

Complete mutational maps for all 608 exons in the OpenSplice dataset, ordered by WT PSI. Each panel corresponds to one exon and displays two sub-panels. Upper panel: mutational heatmap showing the effect of every tested variant on exon inclusion. Rows correspond to mutation types (substitutions to U, C, G, and A; 1nt deletions, Δ1nt; 3nt deletions, Δ3nt; 6nt deletions, Δ6nt; and 21nt deletions, Δ21nt). Columns correspond to nucleotide positions across the construct: 70 nt of upstream intron, the exon, and 25 nt of downstream intron. Deletions are plotted at their starting position. The colour scale represents ΔPSI relative to the wild-type sequence: red indicates increased exon inclusion and blue indicates decreased exon inclusion; dark grey indicates missing data. Vertical dashed lines mark exon-intron boundaries (3′ and 5′ splice sites). The 3′ and 5′ splice site strengths (ss3 and ss5, respectively), calculated with MaxEntScan are annotated above the corresponding boundary. The wild-type nucleotide sequence of the mutagenised construct is shown below the heatmap, with the exon region indicated by a grey bar. Lower panel: wild-type nucleotide importance logo derived from substitution data only. For each position, the value plotted is −mean(ΔLogitPSI) calculated across all three substitutions at that position. Positive values (above the x-axis) indicate that the wild-type nucleotide promotes exon inclusion, whereas negative values (below the x-axis) indicate that it promotes exon skipping. Nucleotides are coloured by identity (A, green; G, yellow; U, blue; C, red). The x-axis corresponds to the wild-type sequence position along the construct; the y-axis shows −mean(ΔLogitPSI) of substitutions. Exon boundaries are indicated by vertical dashed lines as in the upper panel. The panel label indicates the exon name, wild-type PSI in the minigene system, and genomic coordinates (hg38).

**Supplementary Figure 2. Regulatory profiles for all 585 exons with complete coverage**.

Each panel corresponds to one exon and displays two sub-panels; exons are ordered by exon WT PSI. The panel title indicates the exon name and wild-type PSI in the minigene system. The x-axis shows the wild-type nucleotide sequence of the mutagenised construct (70 nt upstream intron, target exon, 25 nt downstream intron); vertical dashed lines mark exon-intron boundaries. Upper panel: the solid blue line shows, for each position, the minimum ΔLogitPSI across all mutations at that position (i.e. the most inclusion-decreasing effect); this profile is used to identify putative enhancer elements (regions where the minimum ΔLogitPSI < −1 for ≥4 consecutive nucleotides). The solid red line shows the maximum ΔLogitPSI per position (i.e. the most inclusion-increasing effect), used to identify putative silencer elements (regions where the maximum ΔLogitPSI > 1 for ≥4 consecutive nucleotides). The black dashed line shows the median ΔLogitPSI across all mutations at each position, providing a summary of the overall directional effect of mutating that nucleotide. The y-axis shows ΔLogitPSI. Lower panel: individual variant data underlying the profile shown in the upper panel. Each point represents a single variant, plotted at its position along the construct and coloured by direction of effect: blue points indicate variants that decrease exon inclusion (ΔLogitPSI < 0) and red points indicate variants that increase exon inclusion (ΔLogitPSI > 0). Shape distinguishes mutation type: filled squares = substitutions to A; filled circles = substitutions to G; filled triangles = substitutions to C; cross = substitutions to U; open diamonds = 1nt deletions; solid lines = 3nt deletions; dashed lines = 6nt deletions

## Acknowledgements

This work was funded by Open Targets (OTAR3087), Wellcome (220540/Z/20/A), the European Research Council (ERC, Advanced Grant 883742, Synergy Grant 101071936), the Spanish Ministry of Science and Innovation (PID2023-146691NB-I00 and LCF/PR/HR21/52410004, EMBL Partnership, Severo Ochoa Centre of Excellence), AGAUR (2021 SGR 01226), and the CERCA Programme/Generalitat de Catalunya. GQ was supported by a FPI fellowship PRE2022-102744, financed by MCIN/AEI/10.13039/501100011033 and FSE +. J.C. was funded by a BBSRC DTP (Bio-technology and Biological Sciences Research Council, Biosciences Doctoral Training Programme, Cambridge, UK). We thank the Wellcome Sanger Institute DNA pipelines for sequencing and all members of the Lehner Lab for discussion and suggestions.

## Data availability

All DNA and cDNA sequencing data have been deposited in the European Nucleotide Archive (ENA) at EMBL-EBI under accession number **PRJEB111846**. PSI values are available in **Supplementary Table 4**, processed predictions scores are available in **Supplementary Table 12**. Figshare contains read counts at the barcode level and additional input files used to run splicing predictor inference, including exon/variant level sequences and their flanking genomic regions and unprocessed model prediction scores: 10.6084/m9.figshare.32337414.

LaBranchoR scores: http://bejerano.stanford.edu/labranchor/; ClinVar data: https://www.ncbi.nlm.nih.gov/clinvar/docs/downloads/;PSI values (VastDB): https://vastdb.crg.eu/wiki/Downloads; SpliceVarDB variant classification: https://splicevardb.org.

ExonExplorer web interface to explore the data: https://results.hgi.sanger.ac.uk/OpenSplice/

## Code availability

All scripts used in this study are available at: https://github.com/lehner-lab/OpenSplice

## Author contributions

G.Q. performed all experiments and analyses except the evaluation of splicing predictors which was performed by J.C.. M.T. and F.S. developed the data processing pipeline. B.L. and J.V. supervised the project. G.Q. and B.L. designed analyses and wrote the manuscript, with input from all authors.

## Methods

### OpenSplice reporter construct

The experiments have been carried out using a minigene derived from the *FAS* human gene^20,21^. The original minigene was composed of (all the coordinates are in the hg38):

- FAS exon 5 (62nt - chr10:89,010,539-89,010,600)
- FAS intron 5 (152nt - chr10:89,010,601-89,010,752)
- FAS exon 6 (63nt - chr10:89,010,753-89,010,815),
- a short version of FAS intron 6 comprising the first 52nt (chr10:89,010,816-89,010,867) and the last 80nt (chr10:89,011,919-89,011,998)
- FAS exon 7 (47nt - hg38 chr10:89,011,999-89,012,045).

We replaced the last 68nt of the intron 5, the entire exon 6 and the first 25nt of intron 6 with a SpeI restriction site to allow library cloning. We also added an AsiSI restriction site right after the exon 7 to allow the barcode cloning. Restriction sites were added using the Q5^®^ Site-Directed Mutagenesis Kit Quick (NEB - #E0554) using the manufacturing protocol. Mutagenic primers are listed in **Supplementary Table 10**: SpeI_Fw and SpeI_Rv were used to insert SpeI site and had an annealing temperature of 56°C; AsisI_Fw and AsisI_Rv were used to insert AsisI site and had an annealing temperature of 56°C. The extension time of the PCR was 3min.

Nonsense-mediated decay is unlikely to contribute substantially to the observed mutational effects because the minigene construct lacks a natural AUG start codon. Moreover, any AUGs created upon pre-mRNA splicing of the variant libraries are unlikely to be in a strong Kozak context or to be followed by long open reading frames.

### Library design

All the libraries analyzed in this work were synthesized by Twist Bioscience. For each exon, the entire exon sequence along with 70 nucleotides (nt) of the upstream and 25 nt of the downstream intronic region were included in the design. For the mutagenesis libraries, saturation mutagenesis was performed across the target regions to generate the following variant types: all possible 1-nt substitutions and deletions of 1, 3, 6, and 21nt applied in a sliding window. To allow cloning via Gibson assembly in the FAS minigene, constant sequences complementary to the plasmid were added to the beginning (5’ AAAAACCAATCACTCTTGATTACTA 3’) and end (5’ CAGATTGAAATAACTTGGGAAGTAG 3’) of each library construct. The exons were categorized into three synthesis pools based on their size to optimize oligonucleotide synthesis and reduce amplification bias: exons shorter than 56 nt, exons 56–100 nt and exons longer than 100 nt.

### Pilot experiments

To test the feasibility of analysing several exons in one experiment, we selected 22 exons known to undergo alternative splicing and for which the WT PSI in a FAS minigene was previously measured (**Supplementary Table 1**).

The exons were categorized into 3 libraries based on their length: < 56 nt (n=7 - P1), 56–100 nt (n=6 - P2) and > 100 nt (n=9 - P3). In P3, for the 4 exons longer than 150nt the upstream intronic regions were truncated to ensure compatibility with synthesis length constraints imposed by the synthesis platform.

In addition, we reanalysed the FAS indel library^15^ to evaluate the effect of barcoding on the PSI values. For this library we keep the minigene used in the previous study (it has a longer intron 6 = 1182nt) and we add the barcodes at the end of exons 7.

### 6000 WT exons screen (to select 1k to mutate)

To select the 586 human exons targeted for mutagenesis, 6000 WT exons were tested in a minigene system to assess their PSI. The exons were divided in 3 libraries of 2000 each: exons < 56 nt (WT1), exons 56–105 nt (WT2), and exons 106–150 nt (WT3). We excluded all the exons longer than 150nt since in the pilot experiments very few variants were retrieved among the 4 exons longer than 150nt and to ensure that the upstream intronic region analyzed was constant across the whole dataset.

The information of all human exons was retrieved using the R package of BiomaRt ^51,52^ using the following criteria:

- filters = “biotype”, “transcript_gencode_basic”,
- values = “protein_coding”,
- attributes=“ensembl_gene_id”,”external_gene_name”,”transcript_mane_select”,”ens embl_transcript_id”,”ensembl_exon_id”,”chromosome_name”,”strand”, “exon_chrom_start”, “exon_chrom_end”,
- Filter the column “chromosome_name” for exons belonging only to the nuclear DNA (autosome + sexual chromosome).

Exon and intron sequences were obtained using the R packages BSgenome and BSgenome.Hsapiens.UCSC.hg38. We then added exon inclusion values from VastDB ^22^ and the number of Pathogenic, Benign and VUS variants from ClinVar 20230722 version ^4^. All the codes to create the table with all the human exons can be found in the github repository.

Once all the information was retrieved, we filtered the exons to be maximum 150nt due to synthesis constraints. We included all the exons with an average PSI across tissues between 25% and 75% (extremes included), yielding 599 exons < 56 nt, 1,235 exons 56-100 nt, and 882 exons 101-150 nt. We then prioritized the remaining exons based on the number of pathogenic ClinVar variants, to include exons belonging to clinical relevant genes. At last we picked exons with a PSI lower than 25% and higher than 75% but with a high PSI range across tissues. The list of the 6000 tested exons and the PSI values in the minigene can be found in the **Supplementary Table 3**.

### Site saturation mutagenesis libraries 1 to 6

Starting from the screening results of 6000 WT exons (**Supplementary. Table 3**), we selected a total of 586 exons. Design of the big mutagenesis libraries was done in two rounds.

In the first round we selected 150 exons from the WT1 library (< 56 nt) and synthesized them in a single oligo pool (MUT1). First, exons were ranked by the number of pathogenic mutations annotated in ClinVar, and the top three exons per gene were selected. Then we considered the PSI obtained from the WT screen and included: 10 exons with an average PSI [0,1), 20 exons with an average PSI [1,20), 90 exons with an average PSI [20,80] and 30 exons with an average PSI (80,100]. This first library has then been tested in our experimental setup to finalize the experimental conditions and improve the design of the other libraries.

In the second round, 436 exons were selected. Based on the result of the MUT1 library we excluded exons shorter than 27nt. We prioritized exons that are alternatively spliced in our system. We also took into account the PSI in the endogenous context to include exons that are alternatively spliced in the genome. For the exons with an inclusion level higher than 0% and we made sure to include those with clinically annotated variants.

In this second round we designed a total of 5 libraries (MUT2-6) and we also added variants that were tested in the pilot experiments and were covering the entire dynamic range. The list of the mutagenized exons can be found in **Supplementary Table 1** and the full list of synthetised variants in **Supplementary Table 11**.

### Variant libraries cloning

#### Oligonucleotide pool amplification

ssDNA oligonucleotides pools from Twist were resuspended in 10 mM Tris buffer, pH 8.0 to a concentration of 20 ng/µL. 5 ng of template were amplified with Q5 High-fidelity polymerase (NEB - #M0491) in a total reaction volume of 50 ul in quadruplicate, using the following primers: FAS_i5_GC_F and FAS_i6_GC_R (sequences in **Supplementary Table 10**). The PCR program was: 98 °C for 1 min; then 12 cycles of 98 °C for 10s, 55 °C for 20 s, 72 °C for 20 s; then 72 °C for 2 min, followed by 10 °C thereafter. To remove excess primers and dNTPs, PCR were incubated with 2 µL ExoSap at 37 °C for 1h, followed by an inactivation step at 80 °C for 20min. PCR reactions were combined and cleaned-up with the Quiaquick PCR purification kit, eluted with 30µL elution buffer and dsDNA measured with a NanoDrop spectrophotometer.

#### Plasmid backbone linearization

The minigene were linearized using 2µL of SpeI-HF (NEB - #R3133) and in the mix was added also 1µL of phosphatase (Quick-CIP NEB - #M0525) to prevent re-ligation during bacterial transformation. The mix was incubated at 37 °C for 1h, followed by an inactivation step at 80 °C for 20min. The linearized plasmid was loaded on a 1% agarose gel and purified using QIAquick gel cleanup kit (QIAGEN) and measured with a NanoDrop spectrophotometer.

#### Library cloning with Gibson assembly

The amplified library was recombined with the linearized minigene. We used a vector:insert ratio of 1:8. We used 150 ng of vector backbone for pilot (P1-3) and WT libraries (WT1-3), and 300ng for mutagenesis libraries (MUT1-6) split in 2 reactions. We added 10µL/reaction of Gibson master mix developed at the CRG Protein Technologies Unit, which contains a mix of T5 exonuclease (T5E4111K 1000U from Epicentre Biotech-Ecogen), Phusion polymerase (F530s 100U from VITRO and a Taq DNA ligase (Protein Technologies Unit CRG, homemade). The mix was incubated at 50 °C for 3 h for DNA assembly. 2h dialysis was performed, followed by concentration step to reach a final volume of 5µL, and then 1.5µL were transformed into 35µL of of 10-beta high-efficiency electrocompetent Escherichia coli cells (NEB - #C3020K), followed by 1 h recovery with 2 ml of super optimal broth with catabolite repression (SOC) medium. For MUT1-6 libraries the transformation was done in duplicates.

After recovery, 1µL was plated onto an LB agar plate with ampicillin and the rest was inoculated into 100 ml of LB liquid with ampicillin. The following morning, colony count was performed and if the number of estimated colonies was at least 20x the number of variants, the plasmid was isolated using the Qiagen Plasmid Plus Midi kit.

To check for cloning efficiency, 10µL of plasmids were sent to Plasmidsaurus for whole plasmids sequencing and we moved to the next steps only when at least 90% of the reads contained the library.

### Barcode cloning

After library cloning, the plasmid was linearized with AsisI (NEB - #R0630) following the protocol described above. The barcode was ordered from IDT as ssDNA. It had 26 degenerate nucleotides separated with constant AT dinucleotides, and it was flanked by sequences complementary to the backbone plasmids (sequence in **Supplementary Table 10**).

The barcode was diluted in NEB buffer r2.1 to a final concentration of 0.2µM, and cloned as ssDNA into the plasmids using the HiFi DNA Assembly (NEB - #E2621), the cloning for MUT1-6 libraries were done in duplicates. 5µL of ssDNA barcode (0.2µM) were mixed with 0.005pmol of vectors (∼12.6ng for a vector of 4.1kb), 10µL of NEBuilder HiFi DNA Assembly Master Mix and water (final volume 20µL). The mix was incubated at 50 °C for 1h. 2h dialysis was performed and the dialyzed mix was concentrated up to 1.5µL which was transformed into 30µL of of 10-beta high-efficiency electrocompetent Escherichia coli cells (NEB - #C3020K), followed by 1 h recovery with 2 ml of super optimal broth with catabolite repression (SOC) medium. After recovery, 1µL and 0.1µL were plated onto an LB agar plates with ampicillin, and the rest was split into 10 flask of 10 ml of LB liquid with ampicillin. The following morning, colony count was performed and flasks were combined to have a final number of transformants equal to 20x the number of variants in the library; this was done to ensure that the mean number of barcodes associated with a specific variant was 20. The plasmid was isolated using the Qiagen Plasmid Plus Midi kit and 10µL were sent to Plasmidsaurus for whole plasmids sequencing to ensure that at least 90% of the reads contained the library and the barcodes.

### Individual variants cloning

To validate splicing measurements from the saturation mutagenesis libraries, 70 variants were selected to span the full dynamic range of PSI values, with priority given to variants shared across multiple libraries. Each variant was cloned into the FAS minigene backbone via Gibson assembly. To enable cloning, constant sequences complementary to the plasmid were appended to the 5’ (5’-AAAAACCAATCACTCTTGATTACTA-3’) and 3’ (5’-CAGATTGAAATAACTTGGGAAGTAG-3’) ends of each construct, and oligopools were ordered from IDT.

Single-stranded DNA cloning was performed using the NEBuilder HiFi DNA Assembly Kit (NEB, #E2621). The minigine vector was linearized as described above. The oligopool was diluted in NEB Buffer r2.1 to 0.2 µM, and 5 µL of ssDNA were mixed with 0.005 pmol of linearized vector (∼12.6 ng for a 4.1 kb plasmid) and 10 µL of HiFi Master Mix (final volume 20 µL, prepared in duplicate). The reaction was incubated at 50°C for 1 hour. Following 2 hours of dialysis, the product was concentrated to 1.5 µL and electroporated into 30 µL of NEB 10-beta high-efficiency electrocompetent E. coli (#C3020K). Cells were recovered for 1 hour in 2 mL SOC medium, and 1 µL and 0.1 µL of the recovery culture were plated on LB-ampicillin agar. Individual colonies were picked the following morning, grown overnight in LB, and plasmid DNA was extracted by miniprep (Qiagen) and Sanger sequenced to confirm insert identity. Of the 70 designed variants, 50 were successfully recovered (**Supplementary Table 5**).

### Sequencing to associate variants with barcodes

#### Illumina sequencing

To associate barcodes with variants in pilot (P1-3), wild-type (WT1-3) and big mutagenesis (MUT1-6) libraries, 2.5ng of plasmids were amplified with Q5 High-fidelity polymerase (NEB - #M0491) in a total reaction volume of 50 ul in quadruplicate. Primers contained partial Illumina adapters, variable degenerate bases to promote complexity on the flowcell, and sequence complementary to the minigene (seq_FAS_i5_[3-5N]_Fw and seq_PT2_[3-5N]_Rv, **Supplementary Table 10**). The PCR program was 98 °C for 30 s; then 15 cycles of 98 °C for 10s, 62 °C for 20 s, 72 °C for 40 s; then 72 °C for 2min followed by 10°C thereafter. The products from this reaction were cleaned up 1.2× of SPRI beads and eluted in 30μl water. Two microliters of the eluate were then amplified in a second PCR that appended the rest of the Illumina sequencing adapter as well as index sequences (PCR2_i[5/7]). The PCR program was 98 °C for 30 s; then 10 cycles of 98 °C for 15 s, 62 °C for 30 s, 72 °C for 30 s; then 72 °C for 2min, followed by 10 °C thereafter. The products from this reaction were cleaned up 1.2× of SPRI beads and eluted in 30μl water. Libraries were sequenced on an Illumina NovaSeq 6000 with 2×250 paired end reads.

Barcodes were retained if (i) have been associated with design variants, (ii) supported by ≥5 reads.

#### Long read sequencing

Since the FAS indel library from ^15^ was not suitable for illumina sequencing due to the length (1182bp) of intron 6, we performed the barcode-variant association using Pacific Biosciences (PacBio) sequencing.

To isolate the barcoded minigene from the rest of the plasmid, 10 μg of plasmid was digested with 2.5µL of SbfI-HF (NEB - #R3642) and 2.5µL of CspCI (NEB - #R0645). The mix was incubated at 37 °C for 1h, followed by an inactivation step at 80 °C for 20min. The digested plasmid was loaded on a 1% agarose gel and the fragment containing the barcoded minigene purified using QIAquick gel cleanup kit (QIAGEN) and measured with a NanoDrop spectrophotometer.

SMRTbell libraries were prepared using the SMRTbell Express Template Prep Kit 2.0 (PacBio) at the sequencing unit of Wellcome Sanger Institute. The library was sequenced with two SMRT Cells on the Sequel IIe instrument (PacBio). The PacBio sequencing data were analyzed with alignparse ^53^. Reads were quality filtered by removing any read with an estimated error rate above 1× 10^−4^ for the minigine sequence, or above 1×10^−3^ for the barcode. Then, consensus sequences were called using the alignparse.consensus.simple_mutconsensus method with default parameter settings. Barcodes were retained if (i) have been associated with design variants, (ii) supported by ≥2 reads.

### HEK293T transfection

HEK293T (purchased from ATCC #CRL-3216) were cultured in Dulbecco’s Modified Eagle’s Medium (DMEM) supplemented with 10% fetal bovine serum (FBS). For each experiment, cells were seeded on day 1, transfected on day 2, the medium was replaced on day 3, and RNA was extracted on day 4.

The number of cells seeded and the culture vessel size were scaled according to the complexity of each library to ensure sufficient variant coverage. WT libraries (WT1–3, 2k variants) were seeded at 6 × 10^5^ cells per well in 6-well plates; pilot libraries P1 and P2 (∼6k variants) were seeded at 1.5 × 10^6^ cells in T25 flasks; pilot library P3 was seeded at 4.5 × 10^6^ cells in T75 flasks; MUT2 (∼24k variants) was seeded at 15 × 10^6^ cells in T175 flasks; and MUT1 and MUT3–6 (∼120k variants) were seeded at 20 × 10^6^ cells in T225 flasks. Single-clone validation experiments were performed by seeding 10^5^ cells/well in 96-well plates. All transfections were performed in biological triplicate.

Transfections were performed using Lipofectamine 3000 (Thermo Fisher Scientific), with all quantities scaled proportionally to the culture vessel surface area relative to one well of a 6-well plate. Briefly, for a 6-well plate format, 500 ng of plasmid DNA was combined with 5 µL of P3000 reagent in 15 µL of Opti-MEM (Solution A). Separately, 5 µL of Lipofectamine 3000 was diluted in 15 µL of Opti-MEM (Solution B). Solutions A and B were combined, mixed gently, and incubated for 10 minutes at room temperature. The resulting mixture (∼40 µL per well) was added dropwise to the cells.

### RNA extraction and reverse transcription

For variant libraries, mRNA was isolated using Dynabeads Oligo (dT)_25_ (Thermo Fisher). Cells were pelleted, washed in PBS, and lysed in Lysis/Binding Buffer. For samples exceeding 5 × 10^5^ cells (libraries P3, MUT1-6), lysates were sheared by repeated passage through a 21-gauge needle to reduce viscosity. Pre-washed beads were incubated with the lysate for 10 minutes at room temperature to allow poly(A) mRNA capture, followed by sequential washes with Washing Buffers A and B. mRNA was eluted in 80 µL of 10 mM Tris-HCl (pH 7.5) at 75°C for 2 minutes and transferred to a new tube on ice.

For individual variants, total RNA was extracted using the Quick-RNA 96 Kit (Zymo Research, R1052). Cells were lysed directly in the culture plate, and lysates were mixed 1:1 with ethanol before binding to the Silicon A plate. On-column DNase I treatment was performed to remove genomic DNA contamination, followed by wash steps with RNA Prep and Wash Buffers. RNA was eluted in 25 µL of DNase/RNase-free water.

cDNA was synthesized from extracted RNA using a minigene-specific primer (PT2, **Supplementary Table 10**) and SuperScript III reverse transcriptase. For each reaction, 11 µL of RNA was combined with 1 µL dNTPs (10 mM), 0.2 µL PT2 primer (10 µM), and water to a volume of 13 µL. Samples were denatured at 65°C for 5 minutes, then snap-cooled to 4°C. A master mix containing 5x First-Strand Buffer, DTT (100 mM), RNaseIN, and SuperScript III was then added to a final volume of 20 µL. The reverse transcription program proceeded as follows: 25°C for 12.5 minutes, 50°C for 1 hour, and a final extension at 72°C for 15 minutes. To ensure sufficient cDNA yield, reactions were performed in duplicate for WT and P libraries, and in quadruplicate for MUT libraries.

### Individual variants PSI calculation

From the cDNA of each individual variant, the minigene splicing product was amplified by PCR using primers PT1 and PT2 (**Supplementary Table 10**), which are complementary to the plasmid backbone. PCR products were resolved by electrophoresis on 6% polyacrylamide gels in 1× TBE buffer and stained with SYBR Safe (Thermo Fisher Scientific, S33102). Bands corresponding to exon inclusion and skipping isoforms were quantified using ImageJ v1.47 (NIH, USA), and PSI values were calculated as the inclusion band intensity divided by the total intensity of all bands. PSI measurements for all validated variants are reported in **Supplementary Table 5**.

### cDNA sequencing

Prior to Illumina library preparation, cDNA concentration was quantified by qPCR for each replicate of the variant libraries using primers Seq_O_F and PT2 (**Supplementary Table 10**). A calibration curve was generated from serial dilutions (1:5, 8 points) of an equimolar mix of spike-in gBlocks (IDT) spanning a range of amplicon sizes (**Supplementary Table 10**). Absolute quantification was used to determine the number of cDNA molecules per µL in each sample, ensuring that at least 300-fold coverage relative to the number of barcodes in each library was loaded into the Illumina library preparation.

Illumina libraries were prepared in two rounds of PCR using Q5 High-Fidelity DNA Polymerase (NEB, #M0491). In PCR1, primers containing partial Illumina adapters, 3–5 degenerate bases to promote flow cell complexity, and sequences complementary to the barcode flanks were used to amplify the barcode region (seq_PT1_[3-5N]_Fw and seq_PT2_[3-5N]_Rv, **Supplementary Table 10**). The thermocycling program was: 98°C for 30 seconds; 12 cycles of 98°C for 10 seconds, 62°C for 20 seconds, 72°C for 1 minute; final extension at 72°C for 2 minutes. Products were cleaned up with 1.2× SPRI beads and eluted in 30 µL water. In PCR2, 2.5 µL of the eluate was amplified in duplicate to append the remaining Illumina sequencing adapters and index sequences (PCR2_i[5/7]). The thermocycling program was: 98°C for 30 seconds; 10 cycles of 98°C for 15 seconds, 64°C for 30 seconds, 72°C for 1 minute; final extension at 72°C for 2 minutes. Products were cleaned up by double size selection (0.8x - 1.2× SPRI beads) and eluted in 30 µL water. Libraries were quantified using the TapeStation (D5000 ScreenTape) and sequenced on an Illumina NovaSeq 6000 with 2×250 paired-end reads.

### PSI calculation

#### Read counting with DiMSum

Splicing-isoform read counts were obtained by processing cDNA sequencing data through DiMSum^54^, run from stage 1 to stage 4. DiMSum was designed for deep mutational scanning of protein fitness and requires a paired input/output experimental design; since our experiment lacks a true input library, we adapted the study design by assigning two biological replicates as the “input” condition and the third as the “output”. For sequencing libraries spanning multiple FASTQ files (e.g. _run1, _run2), individual runs were encoded as technical replicates within DiMSum. A dummy input file, the smallest available FASTQ file among the libraries, was supplied as the second input, and an arbitrary sequence present in the data was designated as the wild-type reference, which is required by DiMSum for stage 5 but is not used in stages 1–4. The stage 4 output is a count table in which each row corresponds to a unique observed sequence and the columns report the read count per replicate.

#### PSI per barcode

The DiMSum output was filtered to retain only rows whose barcode was present in the barcode–variant association file, discarding reads that could not be mapped to a known synthesised variant. Isoform identity was then assigned by exact match of the trimmed sequence, following DiMSum stage 2 adapter trimming, to one of two expected structures: the skipping isoform, comprising FAS exon 7 followed by the barcode, or the inclusion isoform, comprising the variable middle exon, FAS exon 7, and the barcode. For exonic deletion variants, the middle exon in the inclusion isoform corresponds to the residual exon sequence remaining after the deletion. Per-barcode inclusion (N_inc_) and skipping (N_skip_) read counts were then extracted for each biological replicate, and a raw PSI estimate was computed per barcode as N_inc_ / (N_inc_ + N_skip_).

#### Count aggregation

Each synthesized variant was represented by multiple barcodes in the library. Per-barcode inclusion and skipping read counts were first filtered to retain only barcodes with a minimum of 5 sequencing reads. Counts were then summed across barcodes to obtain per-variant (*v*), per-replicate (*r*) inclusion (*N*_*inc*_) and skipping (*N*_*skip*_) counts for each of three biological replicates.

PSI is defined as the proportion of transcripts that include the variant and is constrained to the interval [0, 1]. Sequencing reads are assumed to arise from a binomial sampling process. For a variant (*v*) in a replicate (*r*) with total read depth *N*_*v,r*_, we have:

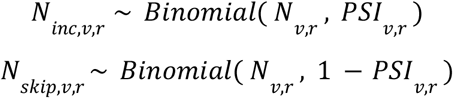

#### PSI estimation

Splicing measurements derived from RNA sequencing are affected by technical uncertainty due to finite read depth, low-coverage variants, and variability between replicate libraries. To obtain a robust estimate of the percent spliced in (PSI) for each variant, we applied a hierarchical error model on the logit scale, both sampling noise and replicate-specific variability within a unified error framework.

To avoid the boundary issue when N is 0, a small pseudocount ε is added to both inclusion and skipping reads. ε is set to 0.5, corresponding to the symmetric Jeffreys prior Beta (0.5, 0.5) used as a symmetric pseudo-count prior for a binomial proportion, which stabilizes estimates for low-coverage variants

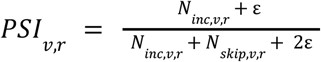

PSI is bounded between 0 and 1. Its variance depends on the mean and becomes asymmetric near the boundaries. To stabilize the variance and better approximate Gaussian errors, estimates were transformed to the logit scale

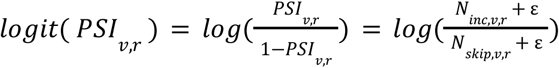

#### Sampling variance of logit PSI

Giving

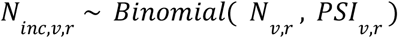

The variance of the number of inclusion reads is given by the binomial variance.

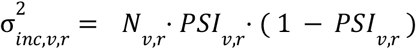

The observed PSI is

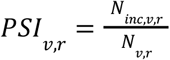

and its variance is

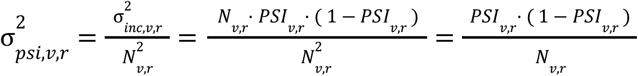

Then the variance of logit PSI is

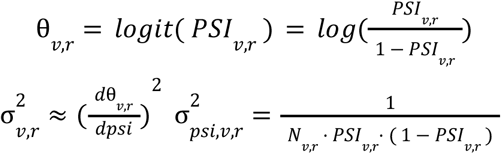

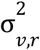 represents sampling uncertainty due to finite read depth. It captures multiplicative (depth-dependent) uncertainty due to finite read counts.

#### Replicate variance

Replicate-specific additive variances 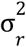 are assumed to be shared across variants and are estimated by pooling information across all variants. To ensure identifiability, the error model is fit jointly across all replicate subsets (e.g., for three replicates, S ∈ {{1,2,3},{1,2},{1,3},{2,3}}), and parameters are learned by maximizing the joint likelihood.

#### Error model

For a variant (v) in a replicate (r), the observed logit-PSI is modeled as

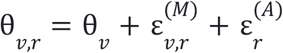

where:

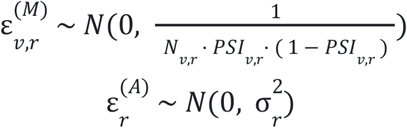

θ_*v*_ is the true underlying logit-PSI of variant (v)

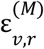 is a multiplicative error term arising from sampling noise

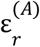 is an additive replicate-specific error term

The total variance of θ_*v,r*_ is therefore

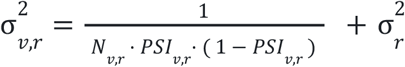

#### Estimation

To ensure identifiability and robust estimation of 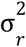, parameters are estimated jointly across all replicate subsets. For each subset S, the negative log-likelihood is

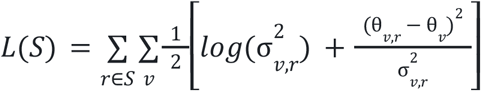

where:

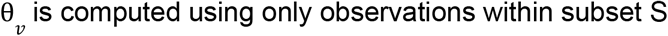

The total objective function is defined as the sum over all subsets:

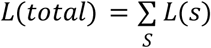

Then following the DiMSum framework ^54^, a joint negative log-likelihood was minimised over all replicate subsets simultaneously using the BFGS algorithm

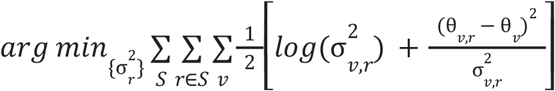

Then under the Gaussian error model

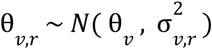

For a given set of variances, the maximum likelihood estimate of θ_*v*_ is given by the inverse-variance weighted mean.

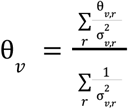

#### Empirical Bayes shrinkage

To reduce noise in variants with low coverage, we applied empirical Bayes shrinkage toward the global mean, estimated from the full variant population.

Assuming a normal prior θ _*v*_ ∼ *N* (μ, τ^2^) where μ and τ^2^ are the mean and variance of logit-PSI across all variants, the shrinkage factor for each variant is:

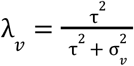

The posterior (shrunk) estimate and variance are given by:

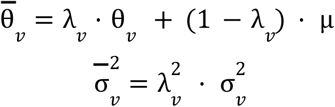

The final PSI estimate and 95% confidence interval were obtained by back-transforming through the logistic function. Only variants with at least one replicate meeting a minimum coverage of 10 reads were retained for downstream analysis.

#### Cross-library normalisation

Variants were assayed across multiple sequencing libraries, each potentially subject to a systematic offset reflecting differences in transfection efficiency or library composition. To remove this batch effect while preserving relative variant effects, we computed an inverse-variance weighted median of logit-PSI within each library and re-centred all estimates to a common global mean:

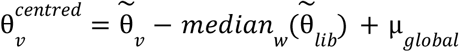

where the weighted median uses per-variant weights proportional to the inverse variance, and the global mean is the inverse-variance weighted mean across all libraries.

#### Calibration to orthogonal validation

To anchor PSI estimates to an absolute, interpretable scale, we performed linear regression of logit-transformed PSI values measured by single-clone RT-PCR gel quantification against the centred logit-PSI estimates:

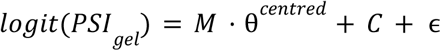

The regression coefficients were then applied to all variants to produce calibrated estimates, with variance propagated accordingly:

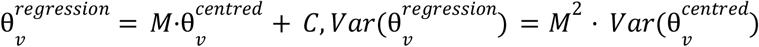

The final PSI value for each variant was obtained by back-transforming the calibrated logit estimate through the logistic function.

#### Variant effect testing

For each exon, the effect of each variant was quantified as the difference in PSI and in logit-PSI relative to the wild-type sequence measured in the same library. Statistical significance was assessed on the logit scale using a z-test, where the test statistic was the logit-scale difference divided by the combined standard error of the mutant and wild-type estimates:

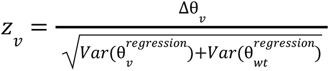

Two-sided p-values were derived from the standard normal distribution and corrected for multiple testing within each exon using the Benjamini–Hochberg procedure. A variant was called a significant splicing-altering when the adjusted p-value < 0.1

### Scoring splice site strength

To score splice sites (SS) strength we used MaxEntScan^29^. Since 3’ SS and 5’ SS scoring model are independent and not comparable (2 different scales, so they cannot cannot be directly combine), we normalized the scores of the splice site (WT and mutant) in our experiments over the score of all coding exon in the human genome:

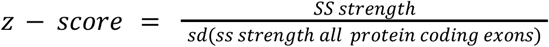

After normalization, we combined the scores:

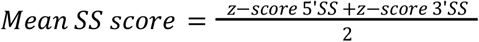

### Branch point (BP) mapping

#### Manual curation

We manually annotated branch points (BPs) using the substitution part of the mutational maps. For each exon, we scanned the upstream intron to identify candidate BP adenines defined by a wild-type A for which all three alternative substitutions (C/G/U) were strongly deleterious for exon inclusion. We then examined the surrounding positions relative to this A in light of the canonical BP consensus (YNYURAY with Y=U/C, R=A/G). In particular, we required evidence of a deleterious U at position A–2 and characteristic context-dependent effects at the three neighboring positions: (i) at A–3 (Y), mutations to U/C tended to increase PSI, whereas mutations to A/G decreased PSI; (ii) at A–1 (R), mutations to A/G increased PSI and mutations to U/C decreased PSI; and (iii) at A+1 (Y), mutations to U/C increased PSI and mutations to A/G decreased PSI. Positions were called as BPs when the wild-type sequence and/or the pattern of mutational effects increased PSI when making the site more similar to the consensus and decreased PSI when moving away from it. List of all manually curated BP in **Supplementary Table 6**.

Manually annotated BPs were validated against two experimental BP datasets^34,35^, matching 82% and 73% of shared exons within ±2 nt, respectively, and with similar position distributions (**Extended data Fig. 5m)**

#### LaBranchoR

To extend BP annotation beyond the 233 manually curated exons, we applied LaBranchoR^36^, a deep-learning model trained to predict BP positions from intronic sequence. For each intron, LaBranchoR returns a probability score at each position within a 70-nt window upstream of the 3′ splice site. The top-ranked prediction per intron was taken as the primary BP (**Supplementary Table 6**). We observed 2 groups: group 1 displaying the mutational signatures observed in the manual curated BP, while group 2 has no mutational signature (**Extended data Fig. 5h-l**).

### Splicing Predictors

#### Model selection

We selected end-to-end deep learning splicing predictors to benchmark that i) take direct encoding of the sequence as input ii) predict splice site probabilities as output at single nucleotide resolution and are iii) amenable to in-silico mutagenesis (ISM) for SNV’s and deletions. Rather than exhaustively benchmark all 12 models^6^, we focus on a subset of widely used models that differ in architecture, training and optimisation strategies.

#### Data generation

SpliceAI, Pangolin, SpliceTransformer were each run using only using their custom sequence scoring option where raw output scores can be extracted. They were evaluated twice, in their native genomic sequence context and in the experimental FAS minigene reporter construct sequence. While the minigene context more accurately reflects the experimental constructs we used to measure splicing outcomes, the reduced length and synthetic minigene context may impair some models performance. Alphagenome was evaluated in native genomic context via the API and in minigene context via the custom sequence mode.

#### Model-specific settings

We used SpliceAI with 10kb of sequence context. It predicts only splice acceptor/donor probabilities. Pangolin predicts splice acceptor and donor probabilities and splice site usage across four tissues. We used splice site probabilities only and for each variant used the tissue with the largest absolute change in acceptor or donor score. For SpliceAI and Pangolin we batched input sequences for computational efficiency. Splice transformer predicts splice acceptor/donor/neither probabilities and provides boolean tissue specific classifications. We used the splice site probabilities only. Alphagenome was evaluated in genomic mode using official API functions predict_interval (reference) and predict_variant (variant) using a 16,384 bp context and 1,048,576 bp context, with the 16,384 bp context yielding the best performance and used in subsequent analyses. Alphagenome predictions were generated on a fixed genomic window centered on the target exon midpoint to ensure consistent WT acceptor and donor predictions. AlphaGenome was run in minigene mode using the research offline model, evaluating each construct with the predict_sequence function on custom sequences padded with Ns to 16,384 bp. Alpha genome predicts splice sites, splice site usage and splice junctions, we use splice site probabilities only to maintain consistency across models. All the predictions are available in **Supplementary Table 12**.

#### Data processing and model evaluation

Canonical splice sites were defined as the exon start (acceptor) and exon end (donor) positions in a strand aware manner. For each variant, splice site scores at these positions were extracted for reference (wild-type) and alternate (mutant) sequences. For deletions, zeroes were inserted in the output arrays at the location of the deletion. Variant effects were calculated as the difference between mutant and reference scores at the canonical sites. The mean of the resultant Δ canonical acceptor and Δ canonical donor scores were compared with experimentally measured ΔPSI values.

Agreement between model predictions and experimental data was assessed using per-construct Spearman R correlations and MAE for both minigene and genome modes. Restricting evaluation to WT exons with intermediate 20-80 PSI values (those most sensitive to mutation, n=336), quantifying performance using mean absolute error (MAE) also yielded the same ranking (**Extended Data Fig. 6b**, Splice AI minigene MAE per exon 10.28, IQR 7.49-12.62). Median pairwise Spearman R correlations across mutations between the experimental replicate pairs were calculated for each exon independently and model performance was re-evaluated on exons for which median of pairwise inter-replicate experimental Spearman R correlations were >0.8 (n=352 exons), this yielded the same ranking (SpliceAI minigene per exon median R 0.79, IQR 0.74-0.83 **Extended Data Fig. 6a)**. Model performance was also evaluated using Spearman R across all mutations (not per construct), stratified by mutation type and transcript region. Model performance was evaluated on 599 exons and on 590,104 unique variants with available experimental measurements and non-null predictions from all models. Residual agreement across models was assessed by calculating signed and absolute errors (experimental ΔPSI minus predicted ΔPSI) and calculating their Spearman R (**Extended Data Fig. 6c,d**).

### Mapping regulatory states

To map enhancer and silencer, we considered all mutation types except 21nt deletions. For exons shorter than 30nt we excluded exonic nt deletions, because we assumed that the effect of mutation can be driven by the excessive exon shortening instead of the presence of a real cis-regulatory element. We then proceed to map the regulatory elements evaluating each nt independently for enhancer and silencer. To map enhancers we considered per each position the median negative ΔLogitPSI values across all the mutations encompassing that position, and we called an enhancer when 4 or more consecutive nt have a ΔLogitPSI < -1 and ΔPSI < -1. For silencer mapping we applied the same process but the threshold was ΔLogitPSI > 1 and ΔPSI > 1. Since enhancer and silencer were mapped separately, we observed that some regions were called in both states and so we added a third state called overlap to identify those sequences. The parts of the sequence in which we cannot find an enhancer/silencer were called neutral. A list of the mapped regulatory elements is available in **Supplementary Table 7**.

#### Hierarchical clustering of regulatory maps

For each exon, the fraction of nucleotide positions annotated as enhancer (E), silencer (S), or overlap (O) was calculated separately for the upstream intron, exon body, and downstream intron, such that E + S + O + N = 1 within each region. The S/E/O fractions were used as input features for hierarchical clustering, with pairwise distances computed using Manhattan distance and clusters assembled using Ward’s D2 linkage criterion (hclust, method = “ward.D2”). The resulting dendrogram was cut at k = 6 to define six exon clusters (**Supplementary Table 8**).

### Clinical variants

ClinVar variant data were downloaded on February 26, 2026 (release clinvar_20260226.vcf.gz). The VCF file was parsed in Python to extract, for each variant, the chromosome, position, allele identifier (ALLELEID), gene name (GENEINFO), genomic HGVS notation (CLNHGVS), variant type (CLNVC), molecular consequence (MC), clinical significance (CLNSIG), and review status (CLNREVSTAT). When multiple molecular consequence terms were present, the SO ontology prefix was removed and terms were concatenated with a pipe separator. The resulting table was filtered to retain only single nucleotide variants and deletions overlapping positions covered by the saturation mutagenesis library (**Supplementary Table 4**).

Clinical significance (CLNSIG) was collapsed into five categories: “*Benign*”, “*Likely Benign*” (including Benign/Likely_benign and Likely_benign), “*Likely Pathogenic*” (including Pathogenic/Likely_pathogenic, Likely_pathogenic), “*Pathogenic*”, and “*VUS/Conflicting*” (covering Uncertain_significance and Conflicting_classifications_of_pathogenicity). Variants lacking a classification (not_provided, no_classification_for_the_single_variant, or missing values) were labelled “*Not provided*”.

Molecular consequence (MC) was simplified into eight categories: Missense, Synonymous, Frameshift, Nonsense, Splice site (splice_acceptor_variant or splice_donor_variant), Intronic, Non-coding (UTR and genic up/downstream transcripts), and Inframe deletion. MC annotations were subsequently re-evaluated against the experimentally determined position and mutation type of each variant in the library to correct cases where the ClinVar annotation was based on a different transcript model.

Review status (CLNREVSTAT) was converted to an ordinal confidence score: 0: no assertion criteria or classification provided; 1: criteria provided by a single submitter or with conflicting classifications; 2: criteria provided by multiple submitters without conflicts; 3: reviewed by an expert panel. Analyses were conducted on the full set of variants, as well as on the subset supported by multiple submitters or expert panel review (review status score > 1) (**Extended Data Fig. 10b-d**).

## Supplementary Tables

**Supplementary Table 1: List of the 608 mutagenized exons with metadata (and wild-type sequences**. Metadata provided: Ensembl gene, transcript and exon IDs, MANE-select RefSeq accession, genomic coordinates (hg38), VastDB event ID and PSI values (mean across tissues and HEK293T-specific).

**Supplementary Table 2: FAS indel library PSI values**. PSI measurements for ∼6,000 variants from the previously published FAS indel mutagenesis library^15^, re-quantified with the OpenSplice barcode-sequencing pipeline.

**Supplementary Table 3**: **6**,**000 wild-type exon screen PSI values and metadata**. Columns include genomic coordinates (hg38), library identifier, Ensembl and VastDB annotation, sequences, ClinVar variant counts, and PSI measurements from the minigene system (logit-PSI estimate, and calibrated PSI)

**Supplementary Table 4: OpenSplice dataset**. Contains variants metadata and PSI values for all the measured variants (598,953 variants across 608 exons)

**Supplementary Table 5: PSI validation of 50 variants**

**Supplementary Table 6: Branch point annotations**. Branch point annotations for 554 introns: 233 manually curated and 321 LaBranchoR-predicted.

**Supplementary Table 7: SpliceMap regulatory elements list**

**Supplementary Table 8: Regulatory architecture groups assignments.**

**Supplementary Table 9: ClinVar variants summary**.

**Supplementary Table 10: Oligonucleotide and primer sequences**.

**Supplementary Table 11: Synthesized oligonucleotide sequences for all variant libraries**.

**Supplementary Table 12**: **OpenSplice dataset with splicing variant effect predictions**. Contains variants metadata, PSI values and model predictions at canonical splice sites for 590,104 variants across 599 exons, for which all models have predictions.

