## Supplementary Figure 1 for "OpenSplice: the impact of half a million mutations on the alternative splicing of 600 human exons"

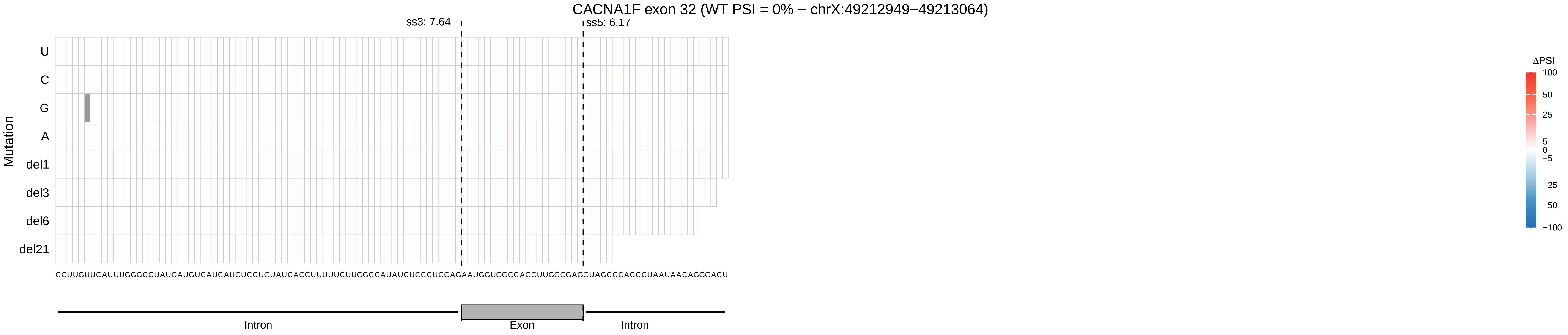

CACNA1F exon 32 (WT PSI = 0% − chrX:49212949−49213064)
ss3: 7.64
ss5: 6.17
Mutation
U
C
G
A
del1
del3
del6
del21
CCUUGUUCAUUUGGGCCUAUGAUGUCAUCAUCUCCUGUAUCACCUUUUUCUUGGCCAUAUCUCCCUCCAGAAUGGUGGCCACCUUGGCGAGGUAGCCCACCCUAAUAACAGGGACU
Intron
Exon
Intron
ΔPSI
100
50
25
5
0
−5
−25
−50
−100

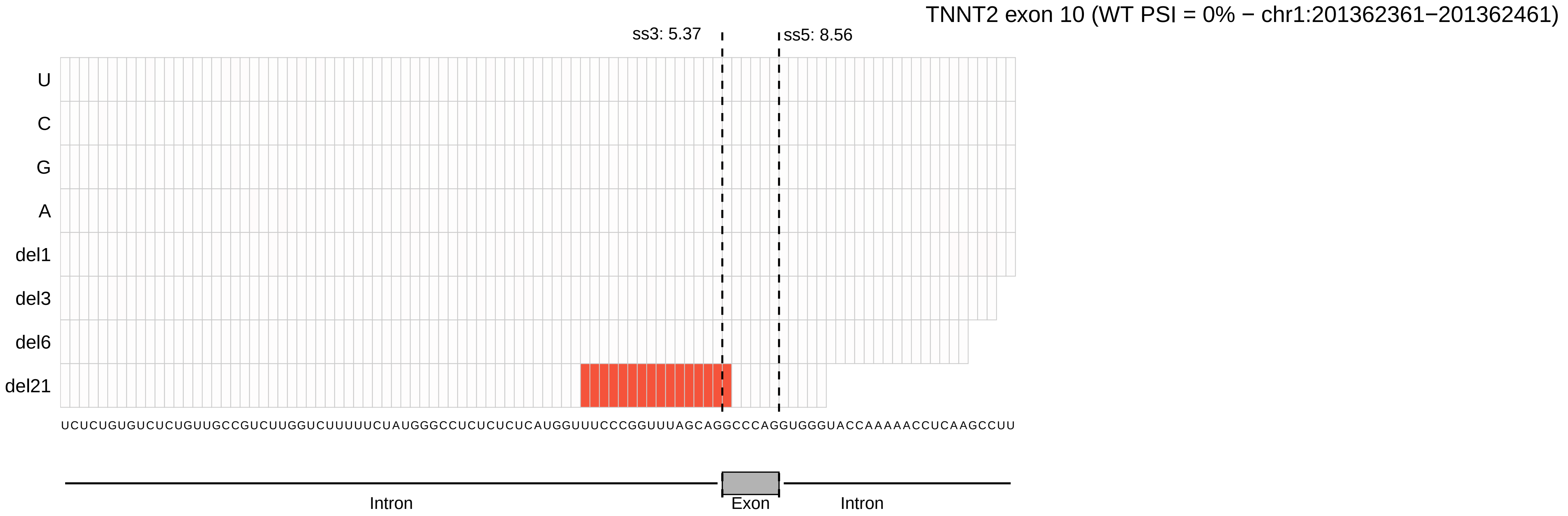

TNNT2 exon 10 (WT PSI = 0% − chr1:201362361−201362461)
ss3: 5.37
ss5: 8.56
U
C
G
A
del1
del3
del6
del21
UCUCUGUGUCUCUGUUGCCGUCUUGGUCUUUUUCUAUGGGCCUCUCUCUCAUGGUUUCCCGGUUUAGCAGGCCCAGGUGGGUACCAAAAACCUCAAGCCUU
Intron
Exon
Intron

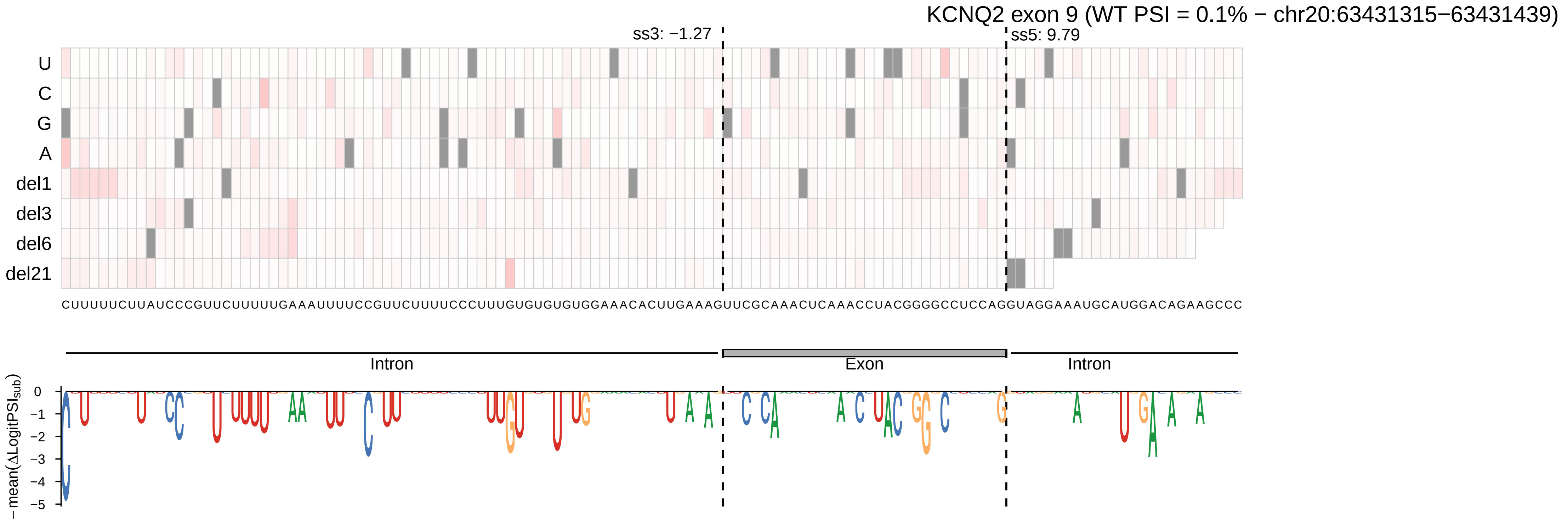

KCNQ2 exon 9 (WT PSI = 0.1% − chr20:63431315−63431439)
ss3: −1.27
ss5: 9.79
U
C
G
A
del1
del3
del6
del21
CUUUUUCUUAUCCCGUUCUUUUUGAAAUUUUCCGUUCUUUUCCCUUUGUGUGUGUGGAAACACUUGAAAGUUCGCAAACUCAAACCUACGGGGCCUCCAGGUAGGAAAUGCAUGGACAGAAGCCC
Intron
Exon
Intron
− mean(ΔLogitPSI$_{sub}$)
0
−1
−2
−3
−4
−5

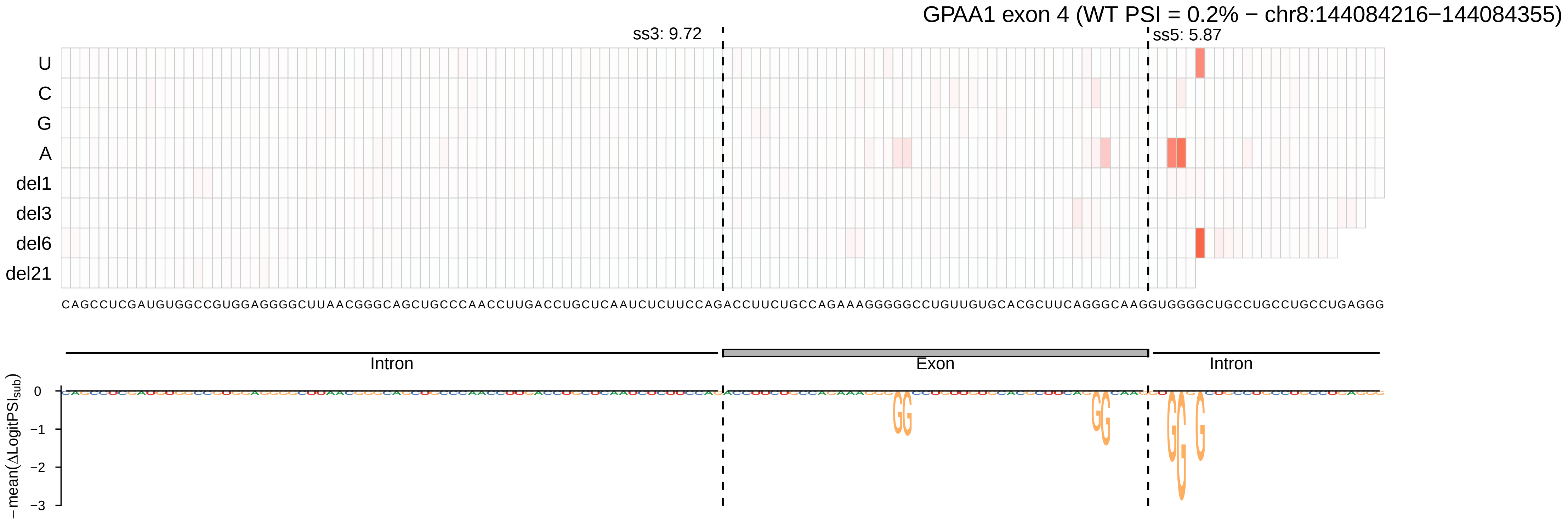
GPAA1 exon 4 (WT PSI = 0.2% − chr8:144084216−144084355)
ss3: 9.72
ss5: 5.87
U
C
G
A
del1
del3
del6
del21
Intron
Exon
Intron
− mean(ΔLogitPSI_sub)
0
−1
−2
−3

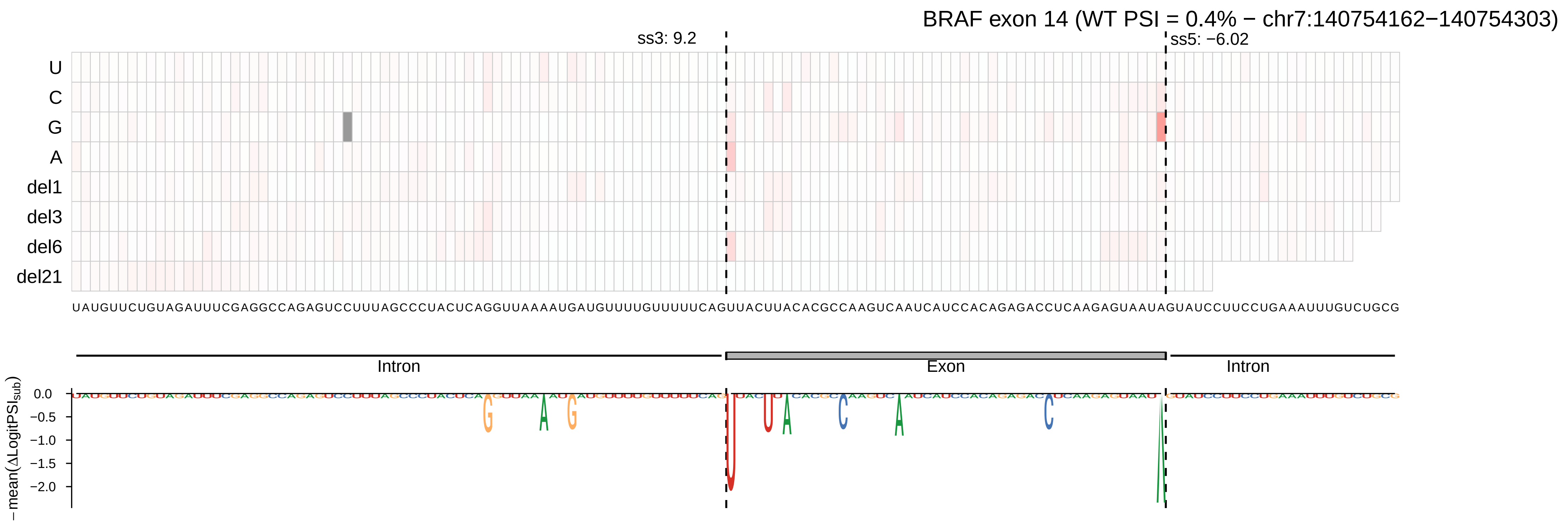
BRAF exon 14 (WT PSI = 0.4% − chr7:140754162−140754303)
ss3: 9.2
ss5: −6.02
U
C
G
A
del1
del3
del6
del21
UAUGUUCUGUAGAUUUCGAGGCCAGAGUCCUUUAGCCCUACUCAGGUUAAAAUGAUGUUUUGUUUUUCAGUUACUUACACGCCAAGUCAAUCAUCCACAGAGACCUCAAGAGUAAUAGUAUCCUUCCUGAAAUUUGUCUGCG
Intron
Exon
Intron
0.0
−0.5
−1.0
−1.5
−2.0
− mean(ΔLogitPSI_sub)

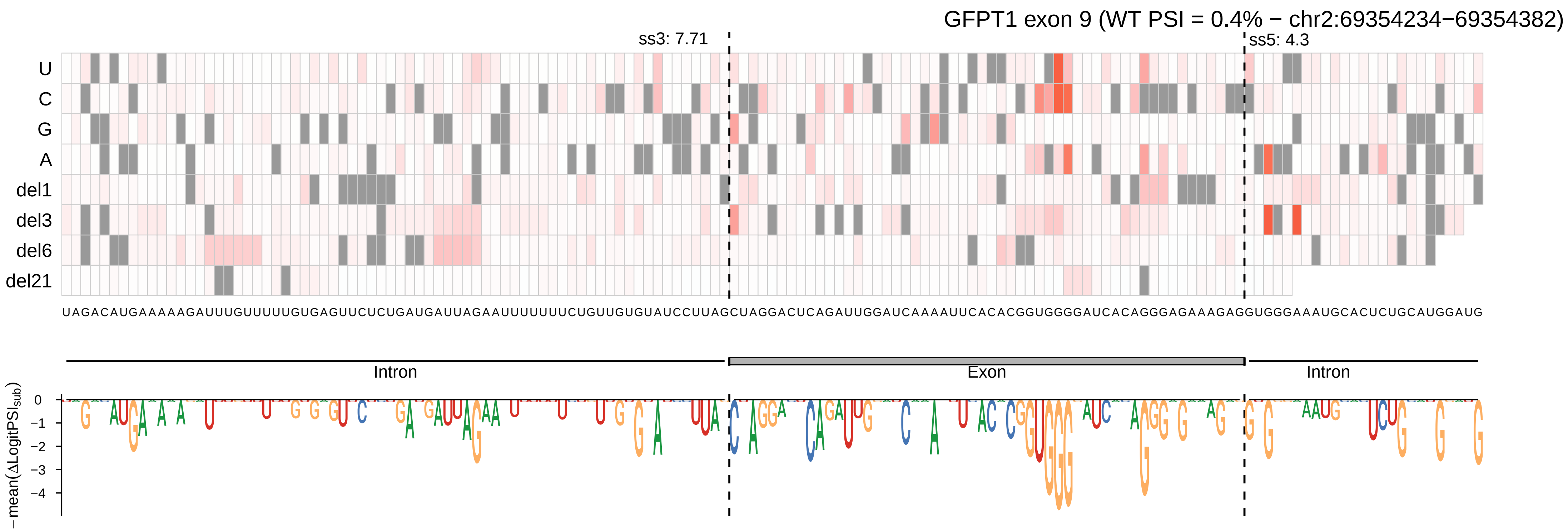

GFPT1 exon 9 (WT PSI = 0.4% − chr2:69354234−69354382)
ss3: 7.71
ss5: 4.3
U
C
G
A
del1
del3
del6
del21
UAGACAUGAAAAAGAUUUGUUUUUGUGAGUUCUCUGAUGAUUAGAAUUUUUUUCUGUUGUGUAUCCUUAGCUAGGACUCAGAUUGGAUCAAAAUUCACACGGUGGGAUCACAGGGAGAAAGAGGUGGGAAAUGCACUCUGCAUGGAUG
Intron
Exon
Intron
−mean(ΔLogitPSI_sub)
0
−1
−2
−3
−4

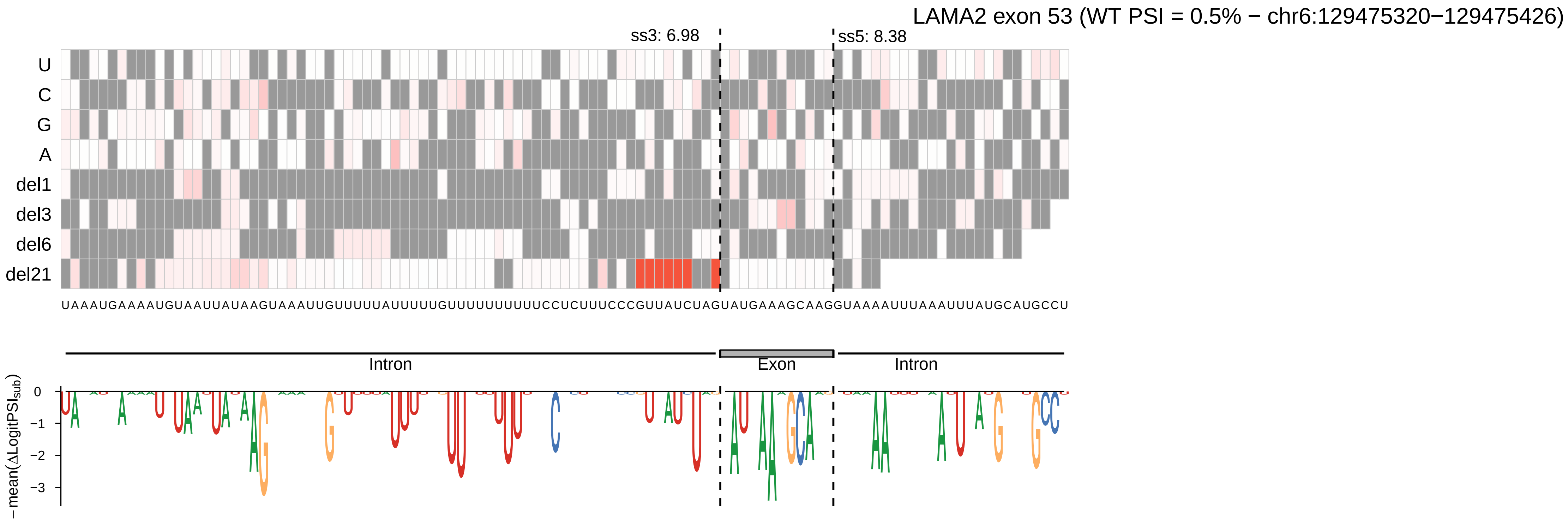
LAMA2 exon 53 (WT PSI = 0.5% − chr6:129475320−129475426)
ss3: 6.98
ss5: 8.38
U
C
G
A
del1
del3
del6
del21
UAAAUGAAAAUGUAAUUAUAAGUAAAUUGUUUUUAUUUUUGUUUUUUUUUUCCUCUUUCCCGUUAUCUAGUAUGAAAGCAAGGUAAAAUUUAAAUUUAUGCAUGCCU
Intron
Exon
Intron
−mean(ΔLogitPSI_sub)
0
−1
−2
−3

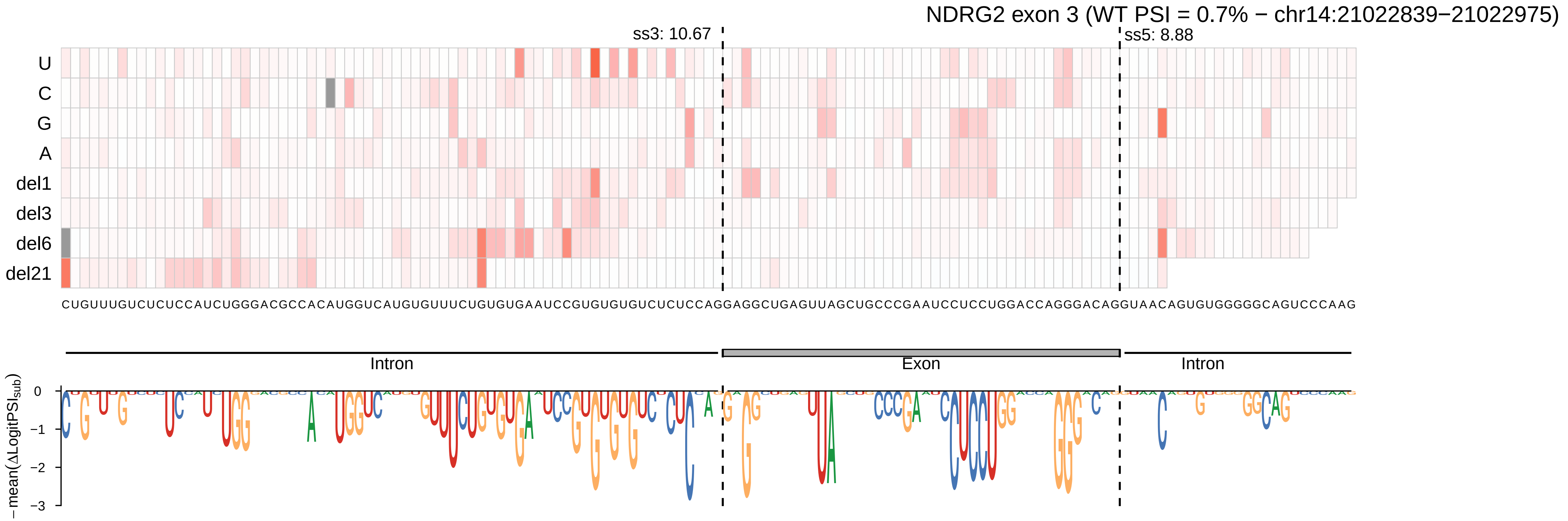

NDRG2 exon 3 (WT PSI = 0.7% − chr14:21022839−21022975)
ss3: 10.67
ss5: 8.88
U
C
G
A
del1
del3
del6
del21
CUGUUUGUCUCUCCAUCUGGGACGCCACAUGGUCAUGUGUUUCUGUGUGAAUCCGUGUGUGUCUCUCCAGGAGGCUGAGUUAGCUGCCCGAAUCCUCCUGGACCAGGGACAGGUAACAGUGUGGGGGCAGUCCCAAG
Intron
Exon
Intron
− mean(ΔLogitPSI_sub)
0
−1
−2
−3

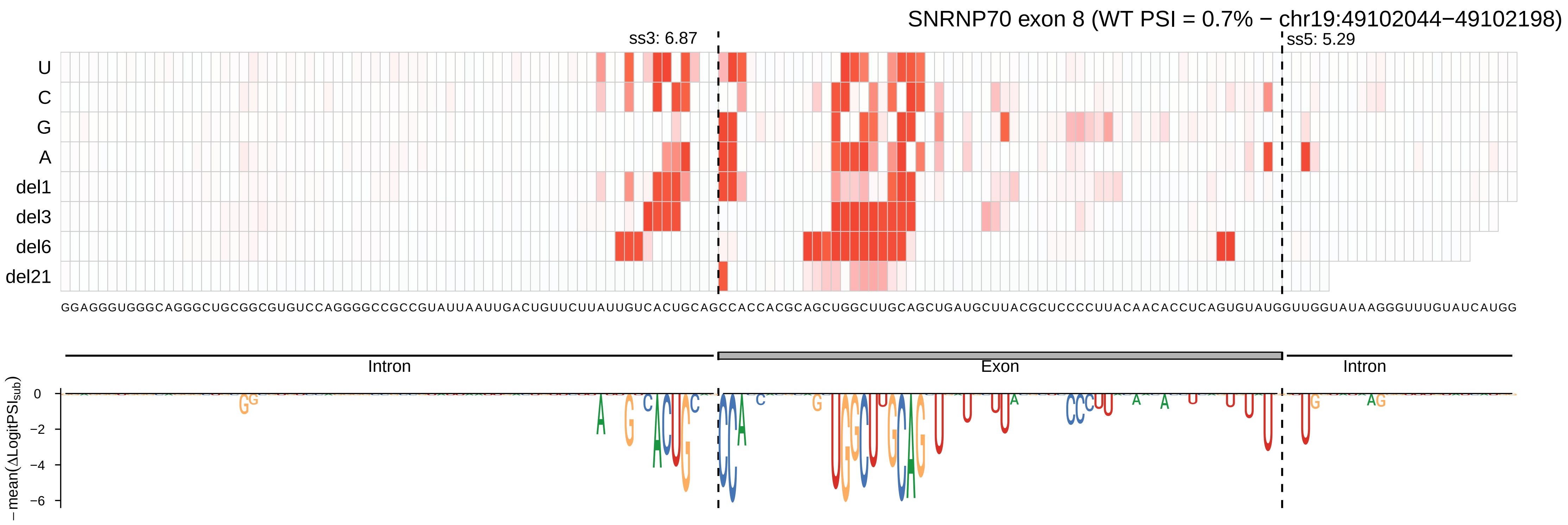
SNRNP70 exon 8 (WT PSI = 0.7% − chr19:49102044−49102198)
ss3: 6.87
ss5: 5.29
U
C
G
A
del1
del3
del6
del21
GGAGGGUGGGCAGGGCUGCGGCGUGUCCAGGGGCCGCCGUAUUAAUUGACUGUUCUUAUUGUCACUGCAGCCACCACGCAGCUGGCUUGCAGCUGAUGCUUACGCUCCCCUUACAACACCUCAGUGUAUGGUUGGUAUAAGGGUUUGUAUCAUGG
Intron
Exon
Intron
−mean(ΔLogitPSI_sub)
0
−2
−4
−6

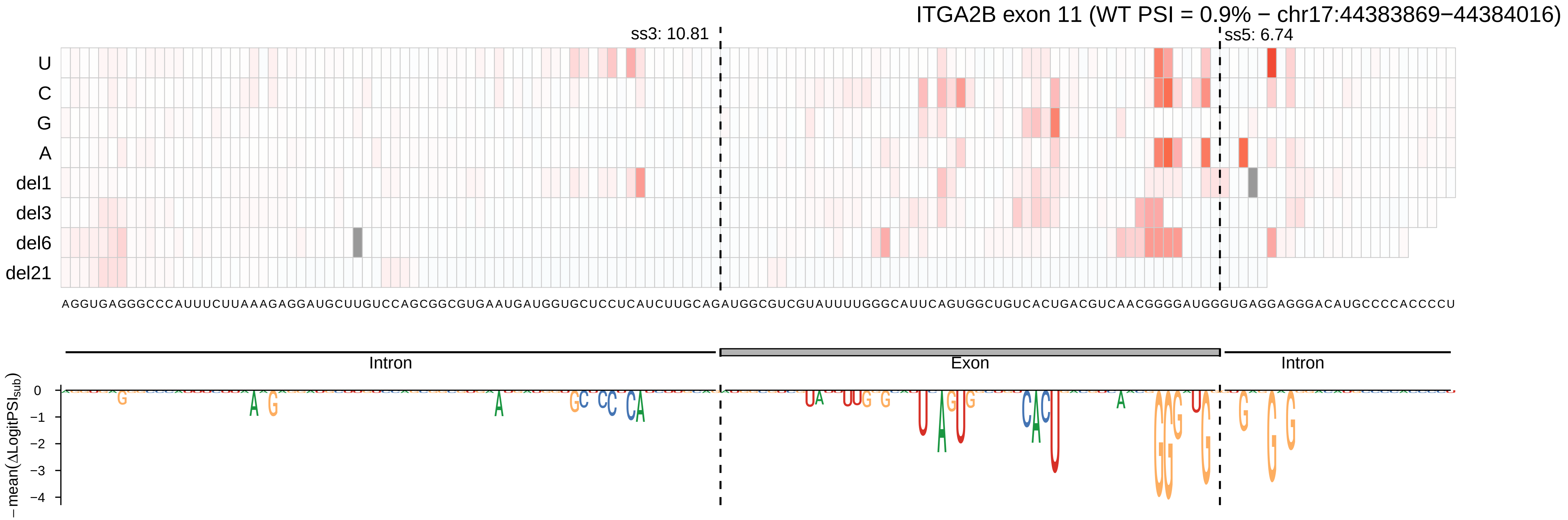

ITGA2B exon 11 (WT PSI = 0.9% − chr17:44383869−44384016)
ss3: 10.81
ss5: 6.74
U
C
G
A
del1
del3
del6
del21
AGGUGAGGGCCCAUUUCUUAAAGAGGAUGCUUGUCCAGCGGCGUGAAUGAUGGUGCUCCUCAUCUUGCAGAUGGCGUCGUAUUUUGGGCAUUCAGUGGCUGUCACUGACGUCAACGGGGAUGGGUGAGGAGGGACAUGCCCCACCCCU
Intron
Exon
Intron
−mean(ΔLogitPSI$_{sub}$)
0
−1
−2
−3
−4

CDH23 exon 12 (WT PSI = 1% − chr10:71643791−71643891)

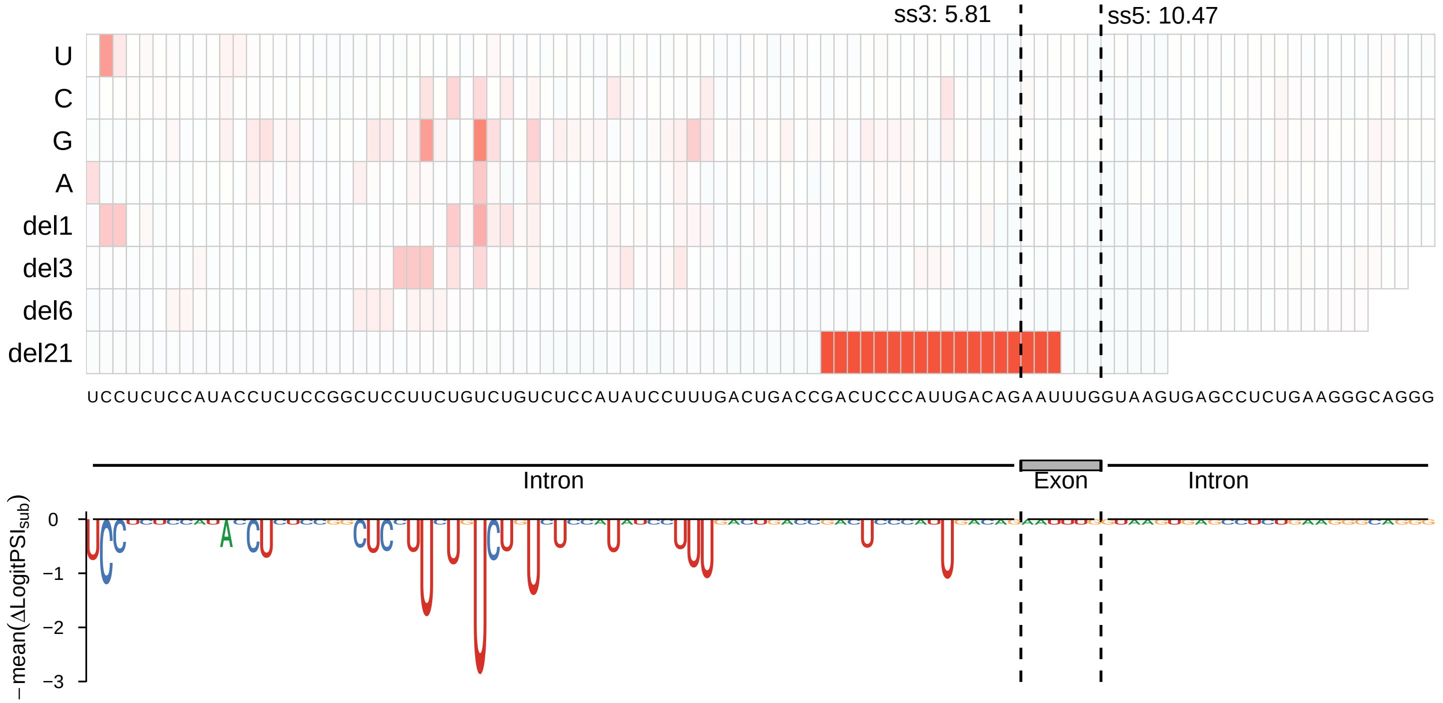

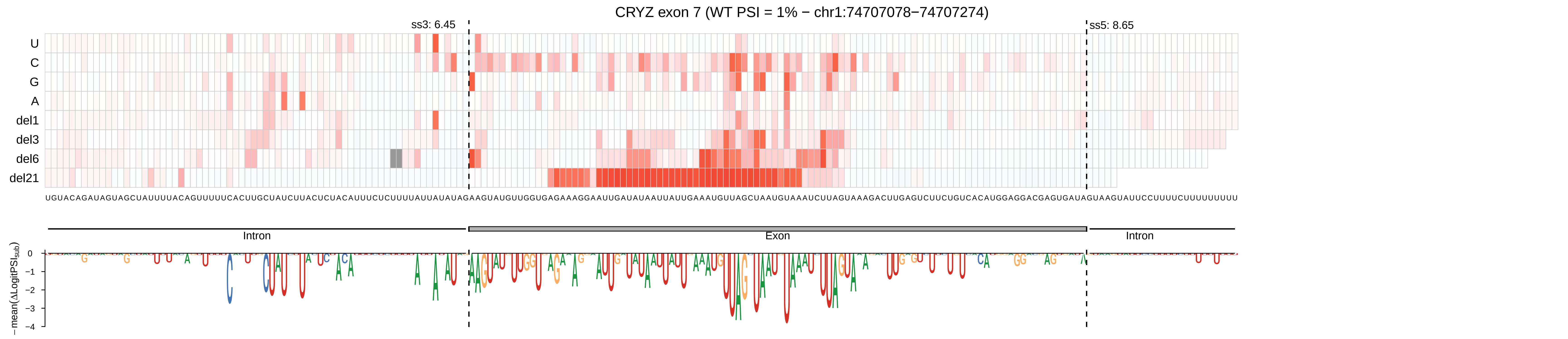

CRYZ exon 7 (WT PSI = 1% − chr1:74707078−74707274)
ss3: 6.45
ss5: 8.65
U
C
G
A
del1
del3
del6
del21
UGUACAGAUAGUAGCUAUUUUACAGUUUUUCACUUGCUAUCUUACUCUACAUUUCUCUUUUAUUAUAUAGAAGUAUGUUGGUGAGAAAGGAAUUGAUAUAAUUAUUGAAAUGUUAGCUAAUGUAAAUCUUAGUAAAGACUUGAGUCUUCUGUCACAUGGAGGACGAGUGAUAGUAAGUAUUCCUUUUCUUUUUUUUU
Intron
Exon
Intron
− mean(ΔLogitPSI_sub)
0
−1
−2
−3
−4

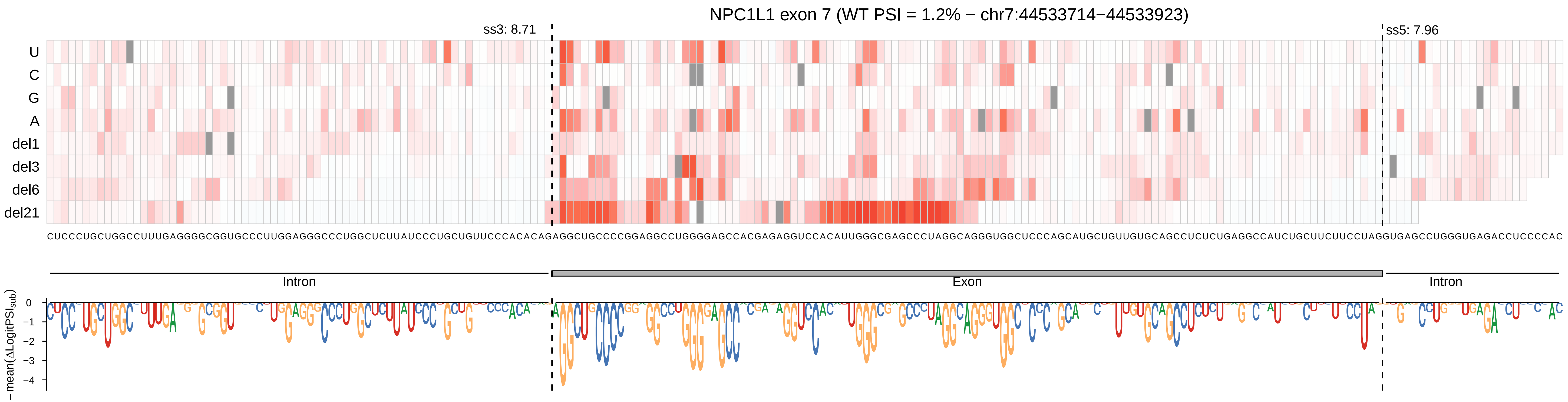
NPC1L1 exon 7 (WT PSI = 1.2% − chr7:44533714−44533923)
ss3: 8.71
ss5: 7.96
U
C
G
A
del1
del3
del6
del21
Intron
Exon
Intron
−mean(ΔLogitPSI_sub)
0
−1
−2
−3
−4

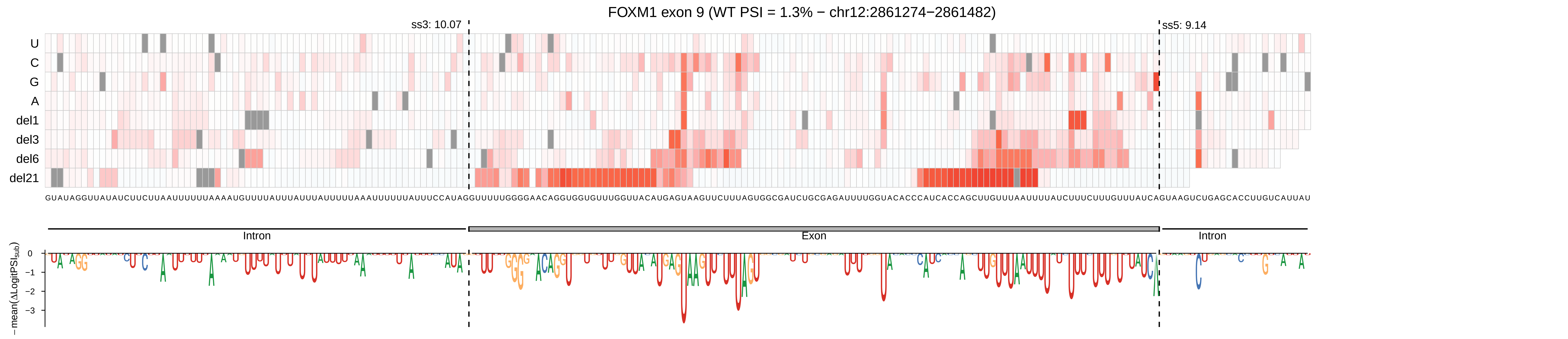

FOXM1 exon 9 (WT PSI = 1.3% − chr12:2861274−2861482)
ss3: 10.07
ss5: 9.14
U
C
G
A
del1
del3
del6
del21
Intron
Exon
Intron
−mean(ΔLogitPSI_sub)
0
−1
−2
−3

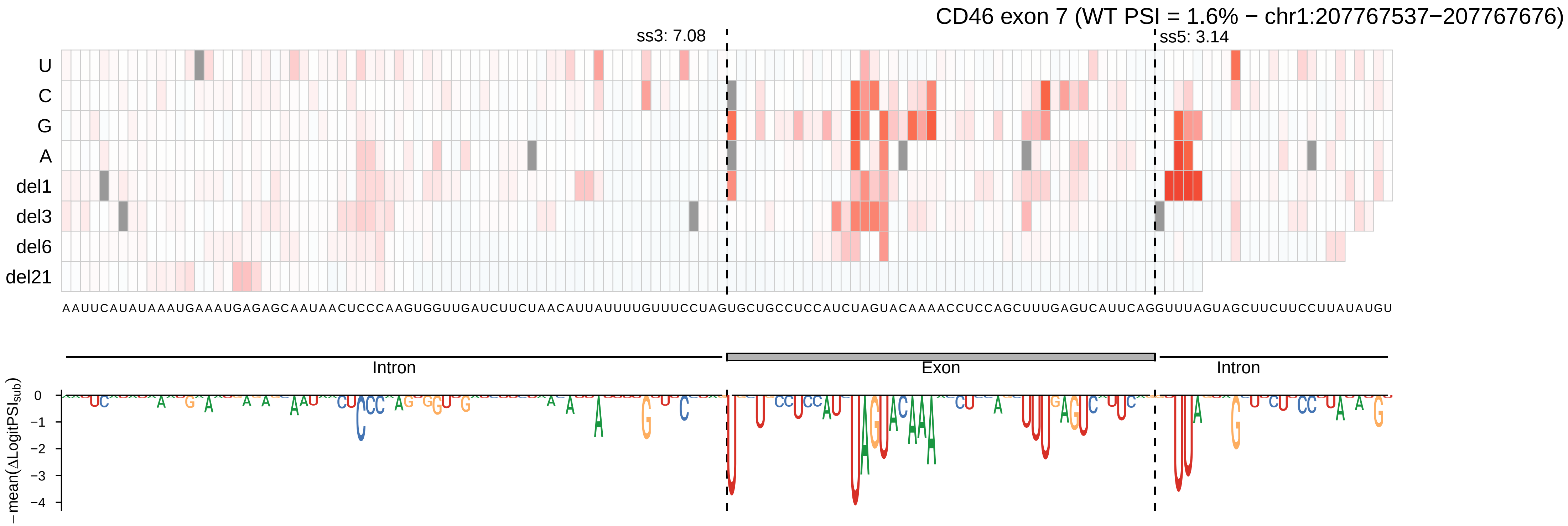

CD46 exon 7 (WT PSI = 1.6% − chr1:207767537−207767676)
ss3: 7.08
ss5: 3.14
U
C
G
A
del1
del3
del6
del21
AAUUCAUAUAAAUGAAAUGAGAGCAAUAACUCCCAAGUGGUUGAUCUUCUAACAUUAUUUUGUUUCCUAGUGCUGCCUCCAUCUAGUACAAAACCUCCAGCUUUGAGUCAUUCAGGUUUAGUAGCUUCUUCCUUAUAUGU
Intron
Exon
Intron
−mean(ΔLogitPSI_sub)
0
−1
−2
−3
−4

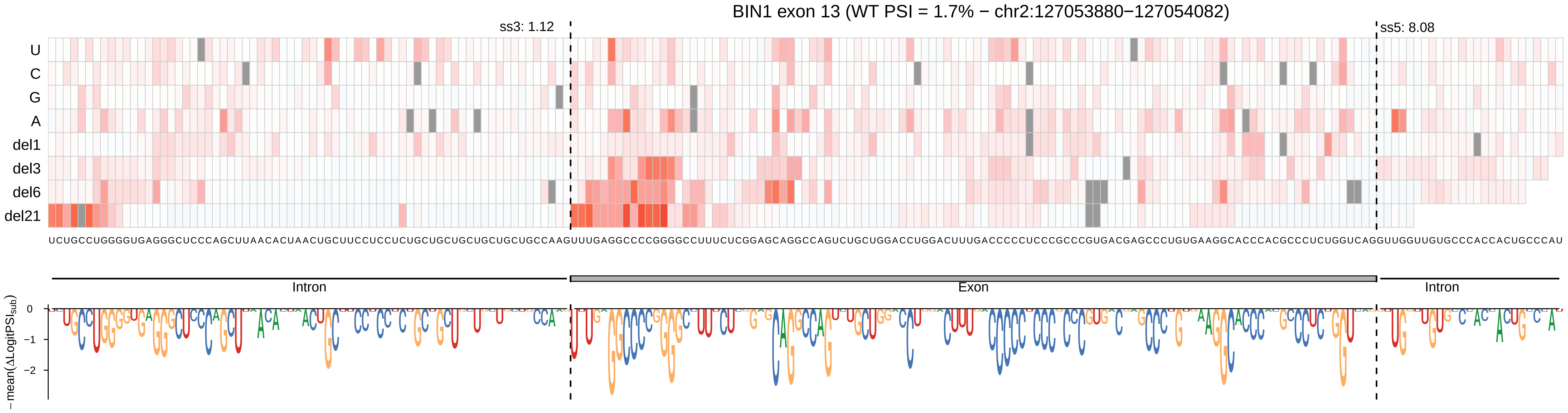

BIN1 exon 13 (WT PSI = 1.7% − chr2:127053880−127054082)
ss3: 1.12
ss5: 8.08
U
C
G
A
del1
del3
del6
del21
Intron
Exon
Intron
0
−1
−2
− mean(ΔLogitPSI_sub)

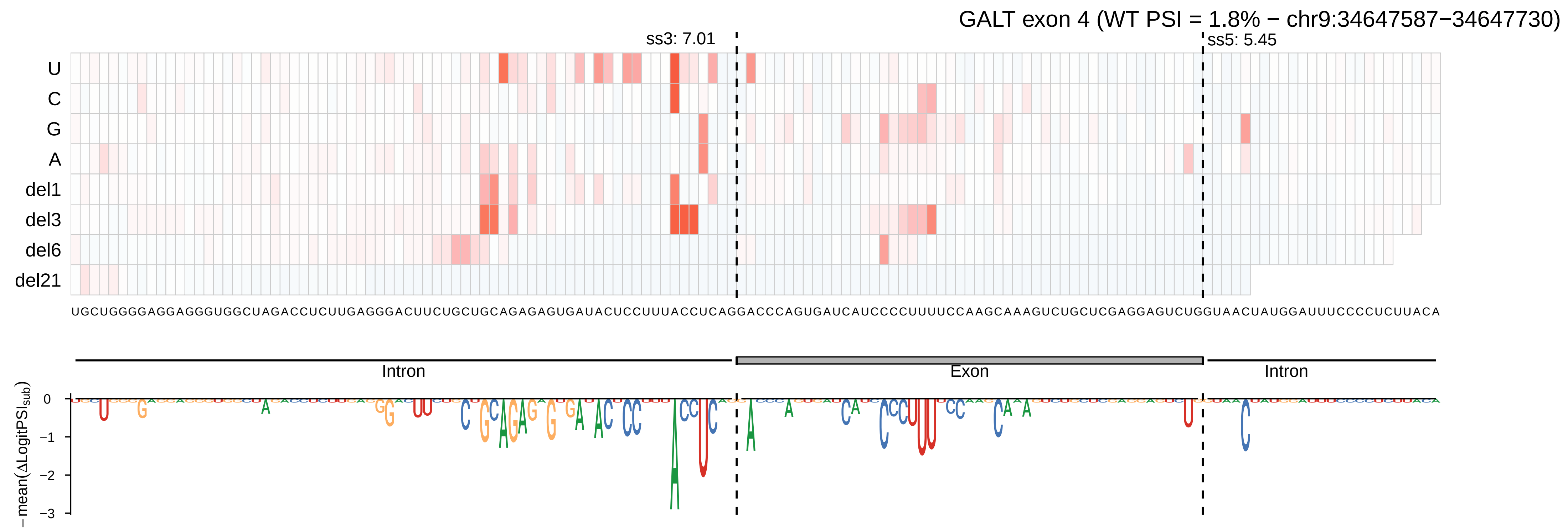
GALT exon 4 (WT PSI = 1.8% − chr9:34647587−34647730)
ss3: 7.01
ss5: 5.45
U
C
G
A
del1
del3
del6
del21
UGCUGGGGAGGAGGGUGGCUAGACCUCUUGAGGGACUUCUGCUGCAGAGAGUGAUACUCCUUUACCUCAGGACCCAGUGAUCAUCCCCUUUUCCAAGCAAAGUCUGCUCGAGGAGUCUGGUAACUAUGGAUUUCCCCUCUUACA
Intron
Exon
Intron
− mean(ΔLogitPSI_sub)
0
−1
−2
−3

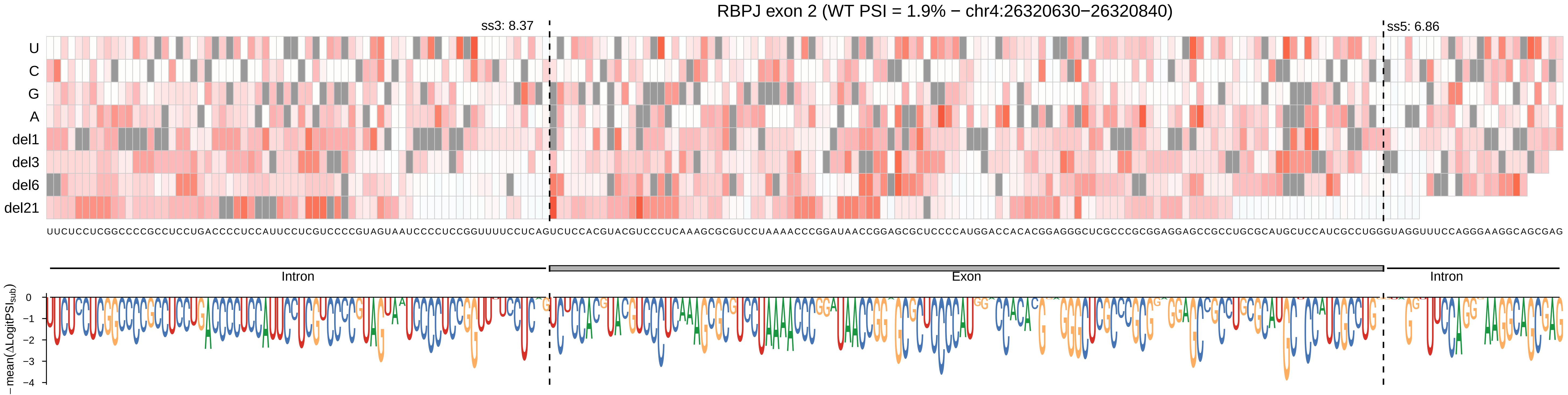
RBPJ exon 2 (WT PSI = 1.9% − chr4:26320630−26320840)
ss3: 8.37
ss5: 6.86
U
C
G
A
del1
del3
del6
del21
UUCUCCUCGGCCCCGCCUCCUGACCCCUCCAUUCCUCGUCCCCGUAGUAAUCCCCUCCGGUUUUCCUCAGUCUCCACGUACGUCCCUCAAAGCGCGUCCUAAAACCCGGAUAACCGGAGCGCUCCCCAUGGACCACACGGAGGGCUCGCCCGCGGAGGAGCCGCCUGCGCAUGCUCCAUCGCCUGGGUAGGUUUCCAGGGAAGGCAGCGAG
Intron
Exon
Intron
−mean(ΔLogitPSI_sub)
0
−1
−2
−3
−4

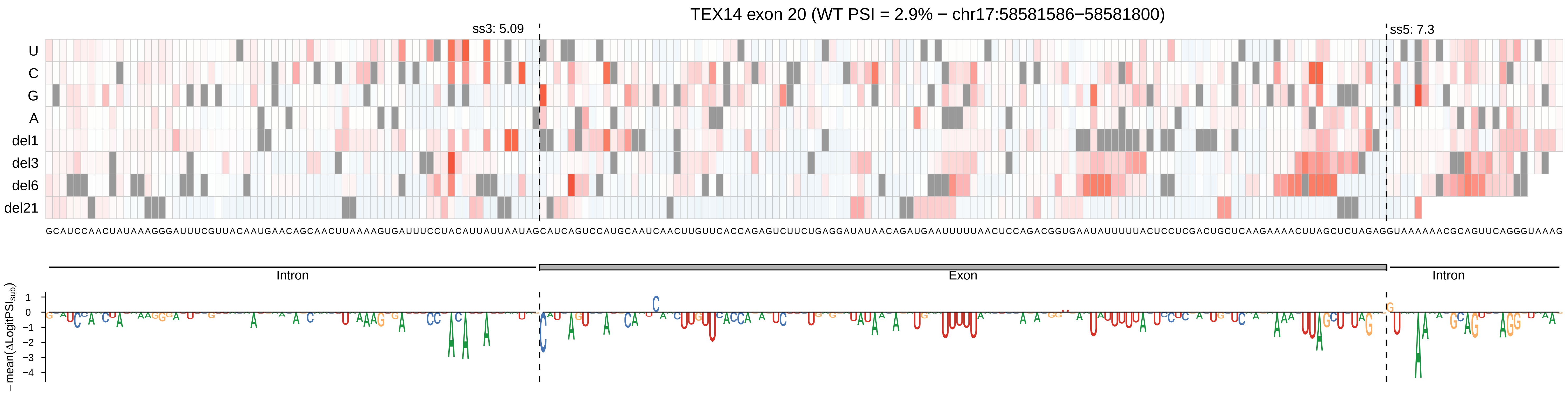
TEX14 exon 20 (WT PSI = 2.9% − chr17:58581586−58581800)
ss3: 5.09
ss5: 7.3
U
C
G
A
del1
del3
del6
del21
GCAUCCAACUAUAAAGGGAUUUCGUUACAAUGAACAGCAACUUAAAAGUGAUUUCCUACAUUAUUAAUAGCAUCAGUCCAUGCAAUCAACUUGUUCACCAGAGUCUUCUGAGGAUAUAACAGAUGAAUUUUUAACUCCAGACGGUGAAUAUUUUUACUCCUCGACUGCUCAAGAAAACUUAGCUCUAGAGGUAAAAAACGCAGUUCAGGGUAAAG
Intron
Exon
Intron
−mean(ΔLogitPSI_sub)
1
0
−1
−2
−3
−4

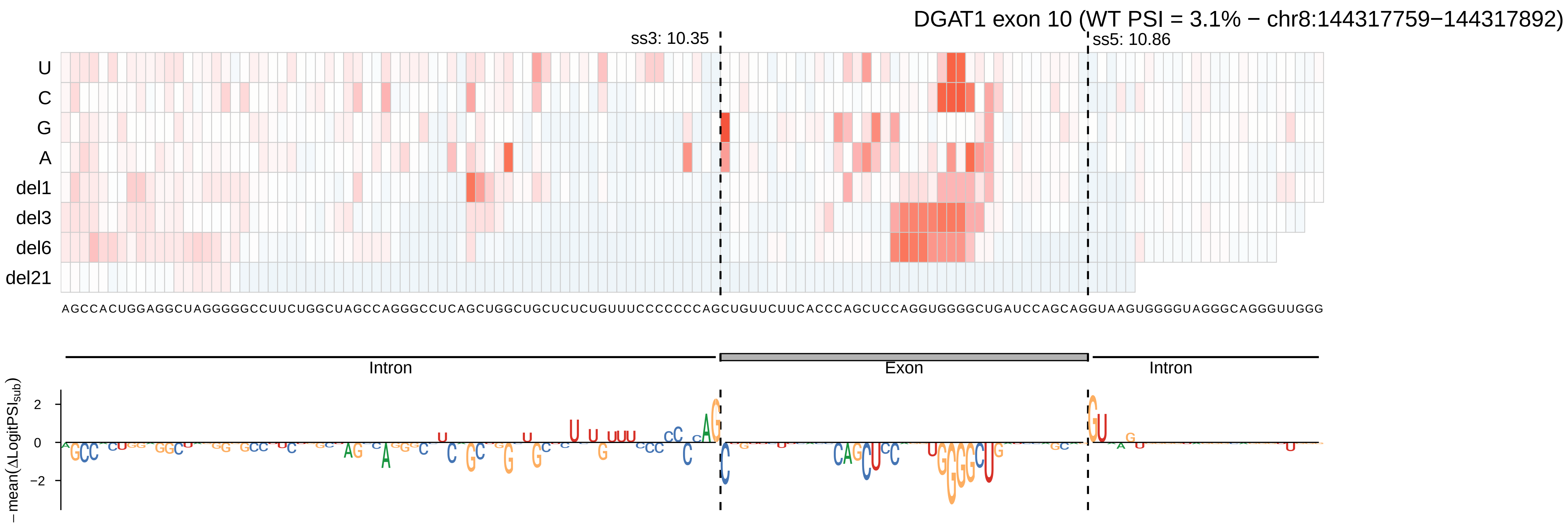
DGAT1 exon 10 (WT PSI = 3.1% − chr8:144317759−144317892)
ss3: 10.35
ss5: 10.86
U
C
G
A
del1
del3
del6
del21
AGCCACUGGAGGCUAGGGGGCCUUCUGGCUAGCCAGGGCCUCAGCUGGCUGCUCUCUGUUUCCCCCCCAGCUGUUCUUCACCCAGCUCCAGGUGGGGCUGAUCCAGCAGGUAAGUGGGGUAGGGCAGGGUUGGG
Intron
Exon
Intron
2
0
−2
− mean(ΔLogitPSI_sub)

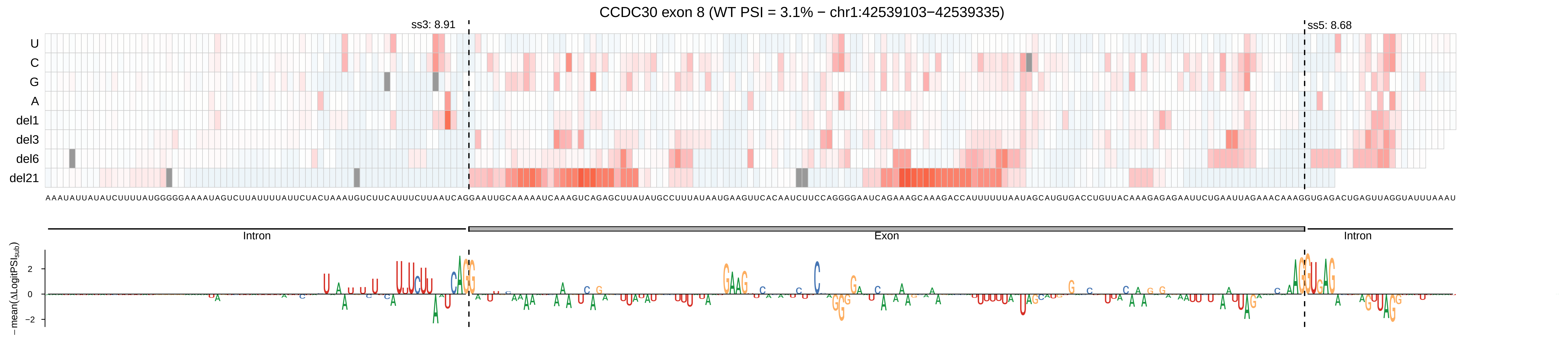

CCDC30 exon 8 (WT PSI = 3.1% − chr1:42539103−42539335)
ss3: 8.91
ss5: 8.68
U
C
G
A
del1
del3
del6
del21
AAAUAUUAUAUCUUUUAUGGGGGAAAAUAGUCUUAUUUUAUUCUACUAAAUGUCUUCAUUUCUUAAUCAGGAAUUGCAAAAAUCAAAGUCAGAGCUUAUAUGCCUUUUAUAAUGAAGUUCACAAUCUUCCAGGGGAAUCAGAAAGCAAAGACCAUUUUUUAAUAGCAUGUGACCUGUUACAAAGAGAGAAUUCUGAAUUAGAAACAAAGGUGAGACUGAGUUAGGUAUUUAAAU
Intron
Exon
Intron
−mean(ΔLogitPSI_sub)
2
0
−2

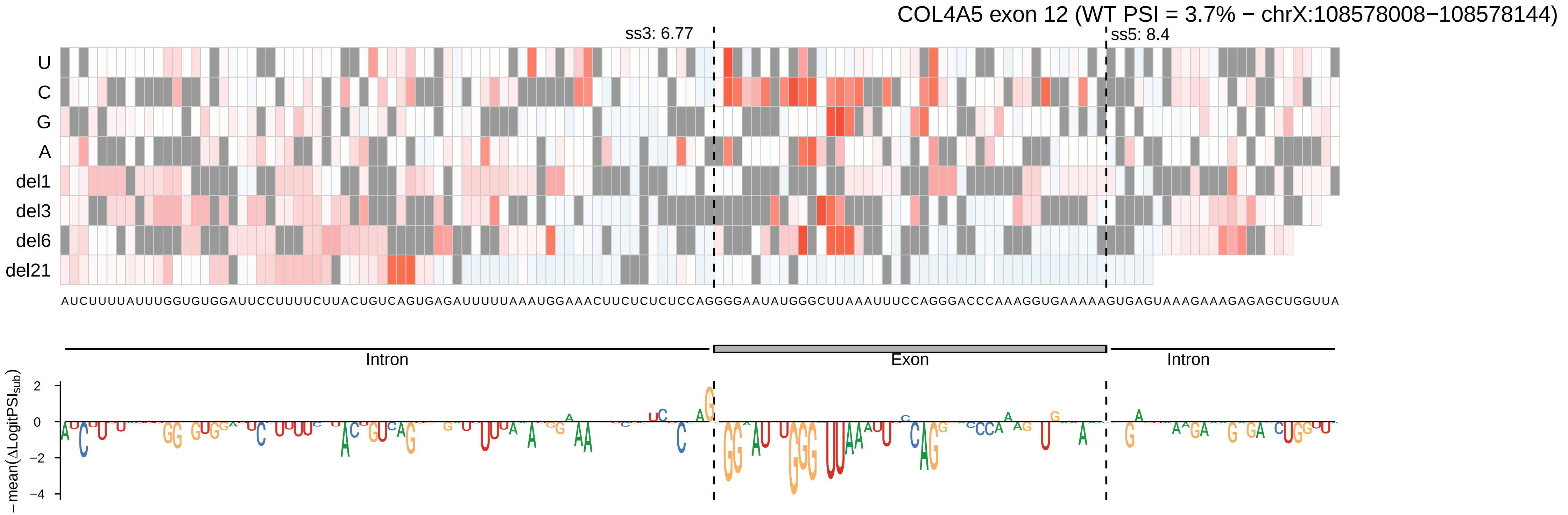
COL4A5 exon 12 (WT PSI = 3.7% − chrX:108578008−108578144)
ss3: 6.77
ss5: 8.4
U
C
G
A
del1
del3
del6
del21
AUCUUUUAUUUGGUGUGGAUUCCUUUUCUUACUGUCAGUGAGAUUUUUAAAUGGAAACUUCUCUCUCCAGGGGAAUAUGGGCUUAAAUUUCCAGGGACCCAAAGGUGAAAAAGUGAGUAAAGAAAGAGAGCUGGUUA
Intron
Exon
Intron
2
0
−2
−4
− mean(ΔLogitPSI_sub)

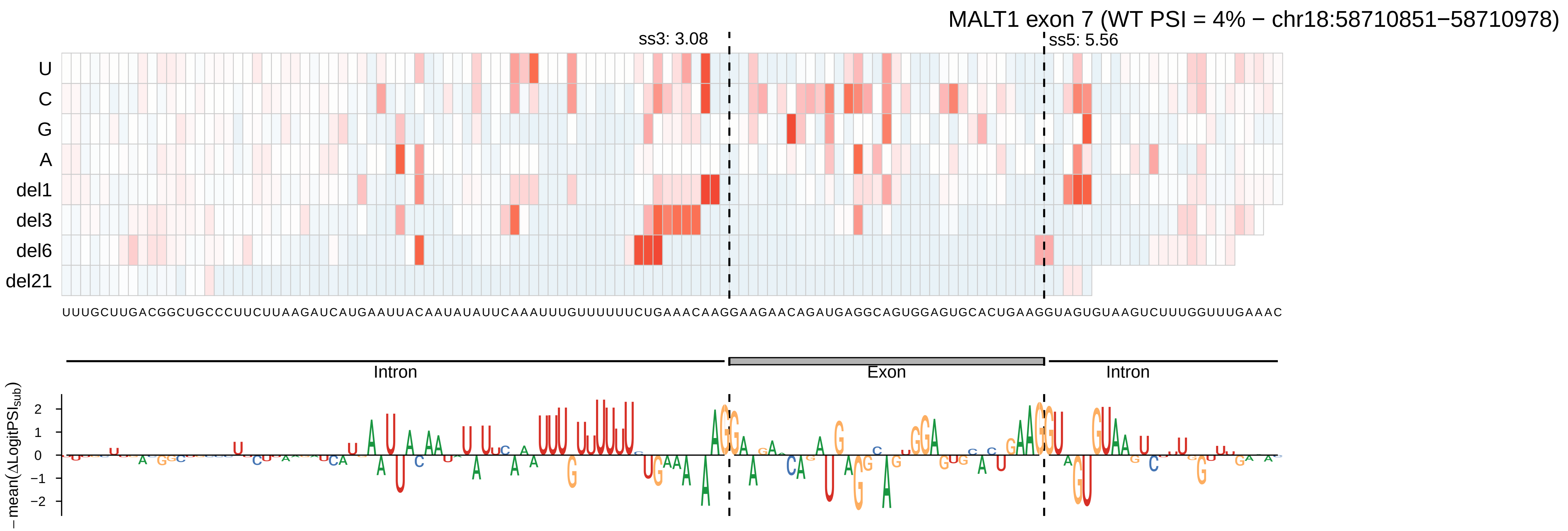
MALT1 exon 7 (WT PSI = 4% − chr18:58710851−58710978)
ss3: 3.08
ss5: 5.56
U
C
G
A
del1
del3
del6
del21
UUUGCUUGACGGCUGCCCUUCUUAAGAUCAUGAAUUACAAUAUAUUCAAAUUUGUUUUUUCUGAAACAAGGAAGAACAGAUGAGGCAGUGGAGUGCACUGAAGGUAGUGUAAGUCUUUGGUUUGAAAC
Intron
Exon
Intron
− mean(ΔLogitPSI_sub)
2
1
0
−1
−2

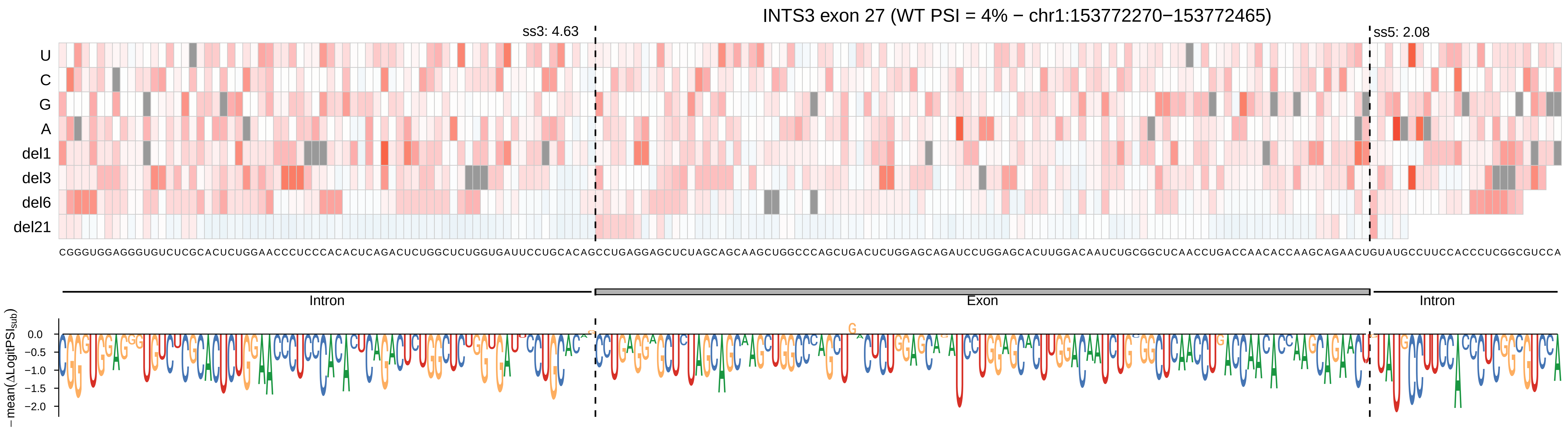
INTS3 exon 27 (WT PSI = 4% − chr1:153772270−153772465)
ss3: 4.63
ss5: 2.08
U
C
G
A
del1
del3
del6
del21
CGGGUGGAGGGUGUCUCGCACUCUGGAACCCUCCCACACUCAGACUCUGGCUCUGGUGAUUCCUGCACAGCCUGAGGAGCUCUAGCAGCAAGCUGGCCCAGCUGACUCUGGAGCAGAUCCUGGAGCACUUGGACAAUCUGCGGCUCAACCUGACCAACACCAAGCAGAACUGUAUGCCUUCCACCCUCGGCGUCCA
Intron
Exon
Intron
0.0
−0.5
−1.0
−1.5
−2.0
− mean(ΔLogitPSI_sub)

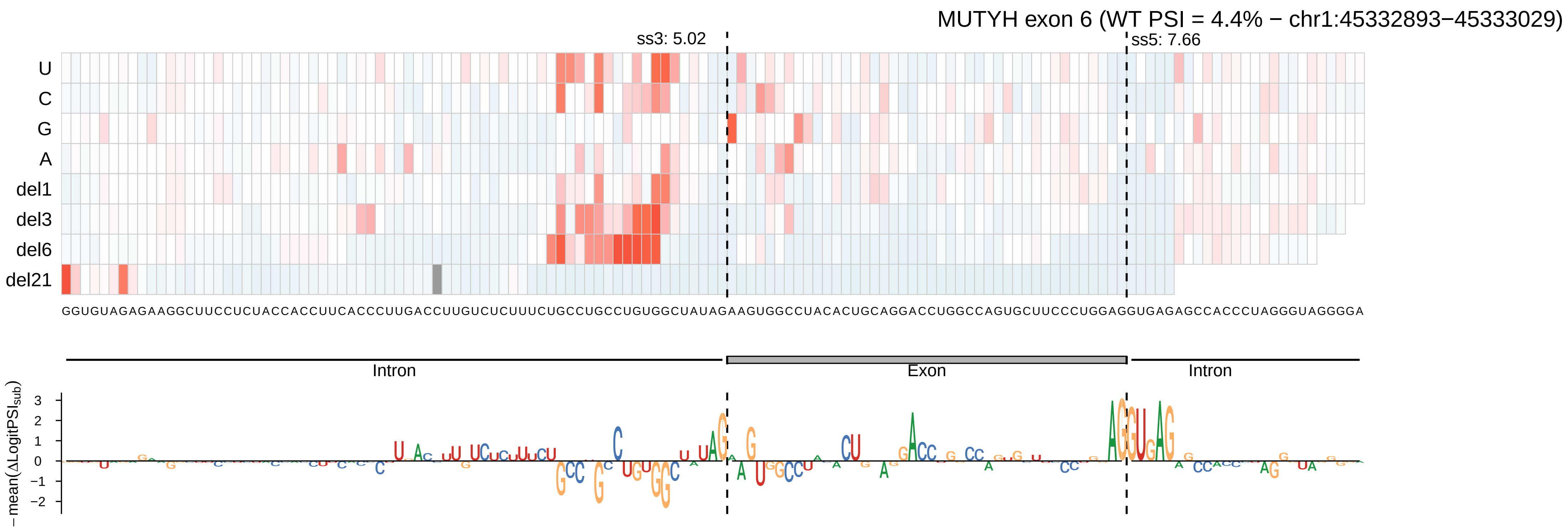

MUTYH exon 6 (WT PSI = 4.4% − chr1:45332893−45333029)
ss3: 5.02
ss5: 7.66
U
C
G
A
del1
del3
del6
del21
GGUGUAGAGAAGGCUUCCUCUACCACCUUCACCCUUGACCUUGUCUCUUUCUGCCUGCCUGUGGCUAUAGAAGUGGCCUACACUGCAGGACCUGGCCAGUGCUUCCCUGGAGGUGAGAGCCACCCUAGGGUAGGGGA
Intron
Exon
Intron
− mean(ΔLogitPSI_sub)

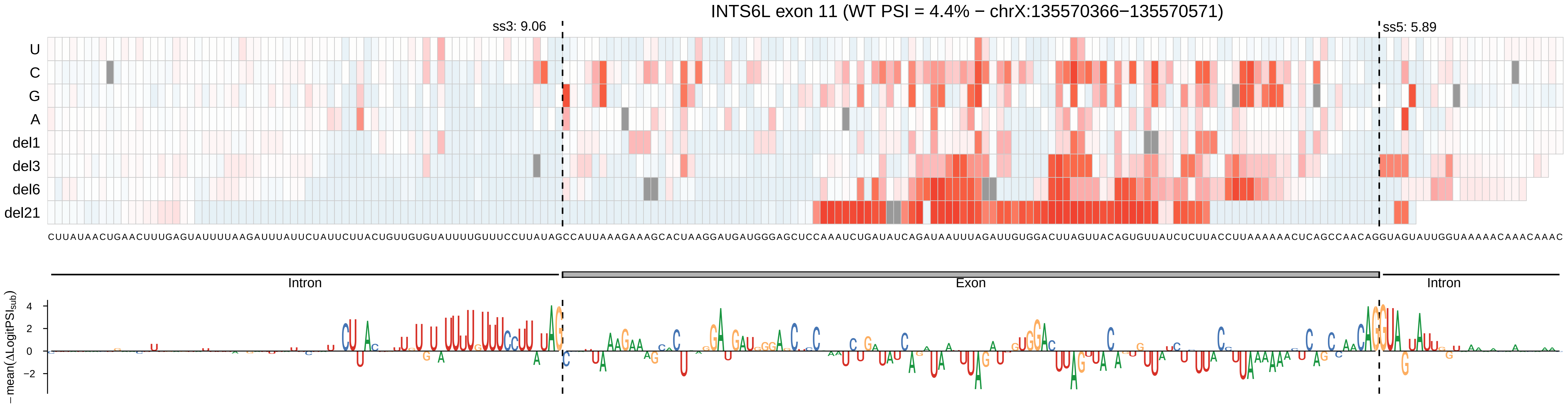
INTS6L exon 11 (WT PSI = 4.4% − chrX:135570366−135570571)
ss3: 9.06
ss5: 5.89
U
C
G
A
del1
del3
del6
del21
CUUAUAACUGAACUUUGAGUAUUUUAAGAUUUAUUCUAUUCUUACUGUUGUGUAUUUUGUUUCCUUAUAGCCAUUAAAGAAAGCACUAAGGAUGAUGGGAGCUCCAAAUCUGAUAUCAGAUAAUUUAGAUUGUGGACUUAGUUACAGUGUUAUCUCUUACCUUAAAAAACUCAGCCAACAGGUAGUAUUGGUAAAAACAAACAAAC
Intron
Exon
Intron
− mean(ΔLogitPSI_sub)
4
2
0
−2

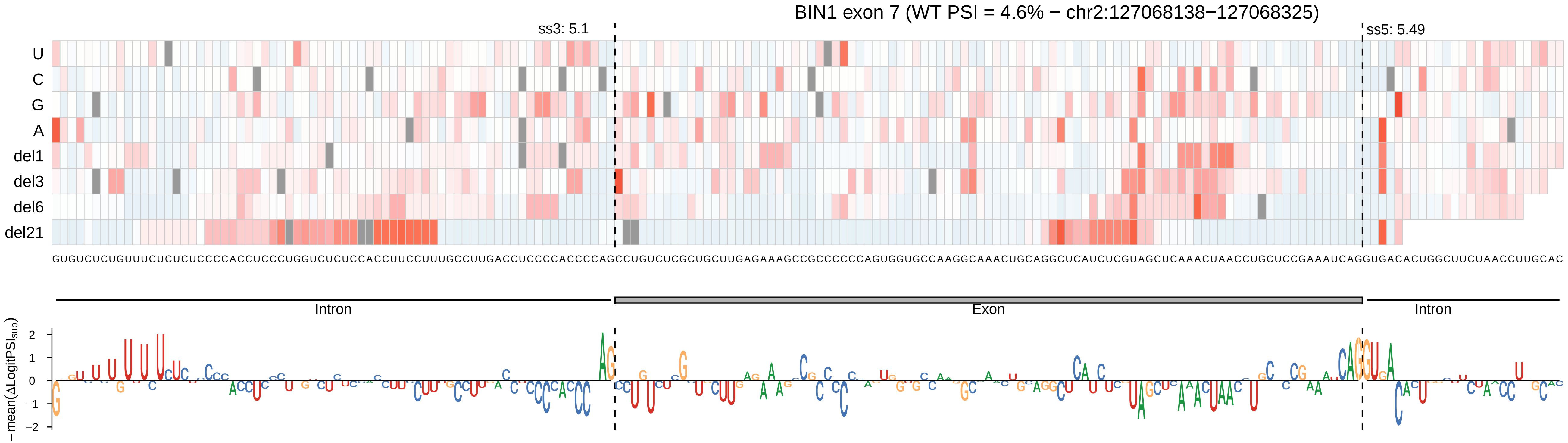

BIN1 exon 7 (WT PSI = 4.6% − chr2:127068138−127068325)
ss3: 5.1
ss5: 5.49
U
C
G
A
del1
del3
del6
del21
Intron
Exon
Intron
−mean(ΔLogitPSI_sub)
2
1
0
−1
−2

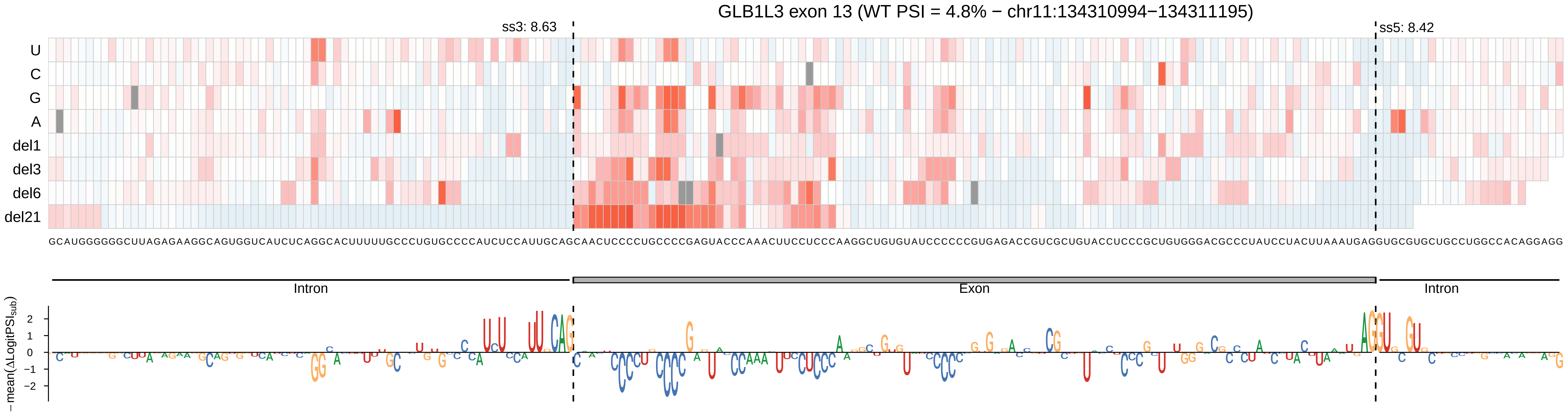

GLB1L3 exon 13 (WT PSI = 4.8% − chr11:134310994−134311195)
ss3: 8.63
ss5: 8.42
U
C
G
A
del1
del3
del6
del21
GCAUGGGGGGCUUAGAGAAGGCAGUGGUCAUCUCAGGCACUUUUUGCCCUGUGCCCCAUCUCCAUUGCAGCAACUCCCCUGCCCCGAGUACCCAAACUUCCUCCCAAGGCUGUGUAUCCCCCCGUGAGACCGUCGCUGUACCUCCCGCUGUGGGACGCCCUAUCCUACUUAAAUGAGGUGCGUGCUGCCUGGCCACAGGAGG
Intron
Exon
Intron
− mean(ΔLogitPSI_sub)
2
1
0
−1
−2

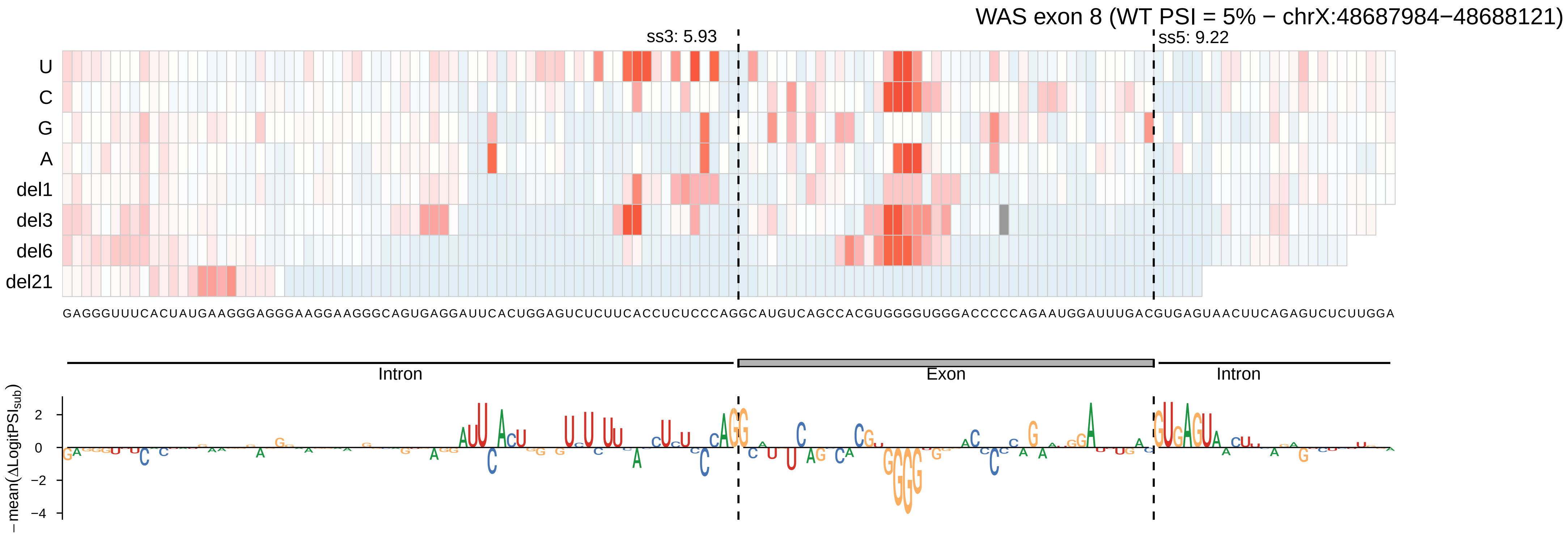
WAS exon 8 (WT PSI = 5% − chrX:48687984−48688121)
ss3: 5.93
ss5: 9.22
U
C
G
A
del1
del3
del6
del21
GAGGGUUUCACUAUGAAGGGAGGGAAGGAAGGGCAGUGAGGAUUCACUGGAGUCUCUUCACCUCUCCCAGGCAUGUCAGCCACGUGGGGUGGGACCCCCAGAAUGGAUUUGACGUGAGUAACUUCAGAGUCUCUUGGA
Intron
Exon
Intron
− mean(ΔLogitPSI_sub)
2
0
−2
−4

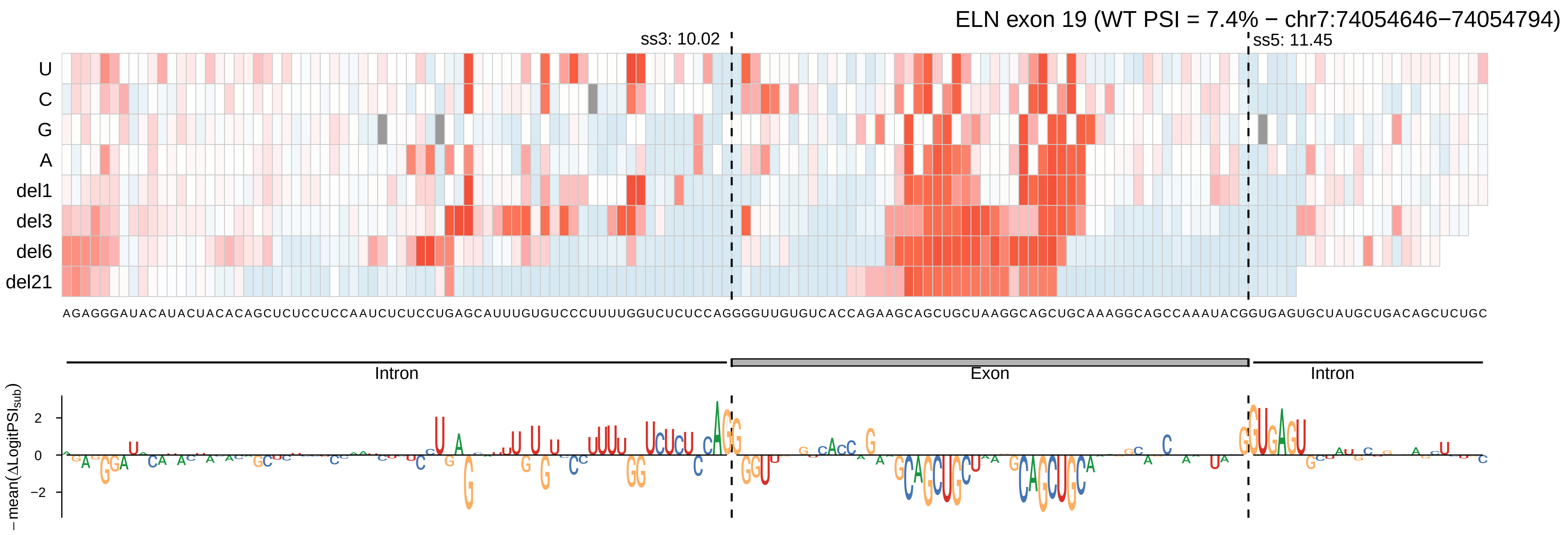
ELN exon 19 (WT PSI = 7.4% − chr7:74054646−74054794)
ss3: 10.02
ss5: 11.45
U
C
G
A
del1
del3
del6
del21
AGAGGGAUACAUACUACACAGCUCUCCUCCAAUCUCUCCUGAGCAUUUGUGUCCCUUUUGGUCUCUCCAGGGGUUGUGUCACCAGAAGCAGCUGCUAAGGCAGCUGCAAAGGCAGCCAAAUACGGUGAGUGCUAUGCUGACAGCUCUGC
Intron
Exon
Intron
− mean(ΔLogitPSI_sub)
2
0
−2

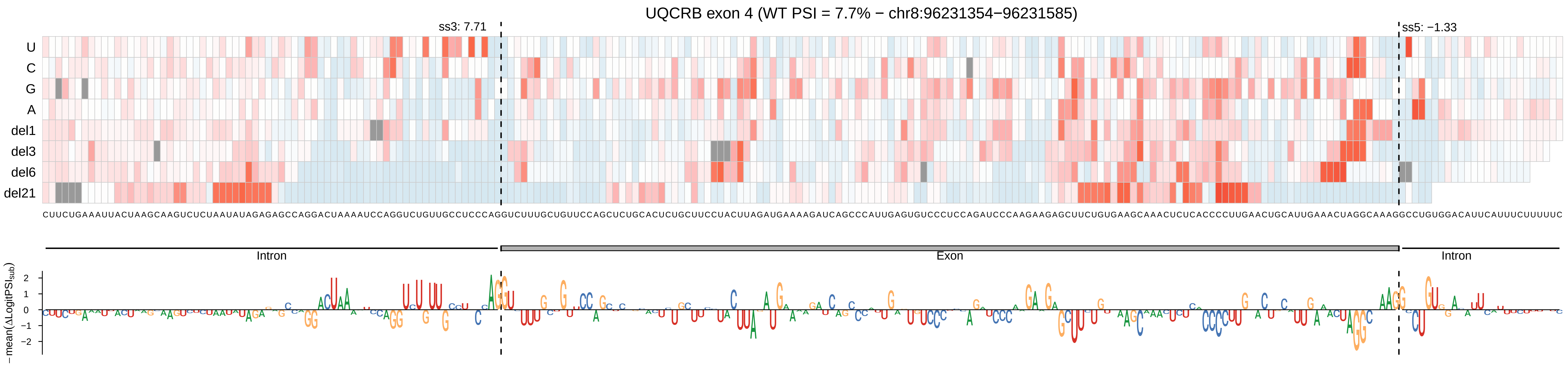
UQCRB exon 4 (WT PSI = 7.7% − chr8:96231354−96231585)
ss3: 7.71
ss5: −1.33
U
C
G
A
del1
del3
del6
del21
Intron
Exon
Intron
−mean(ΔLogitPSI_sub)
2
1
0
−1
−2

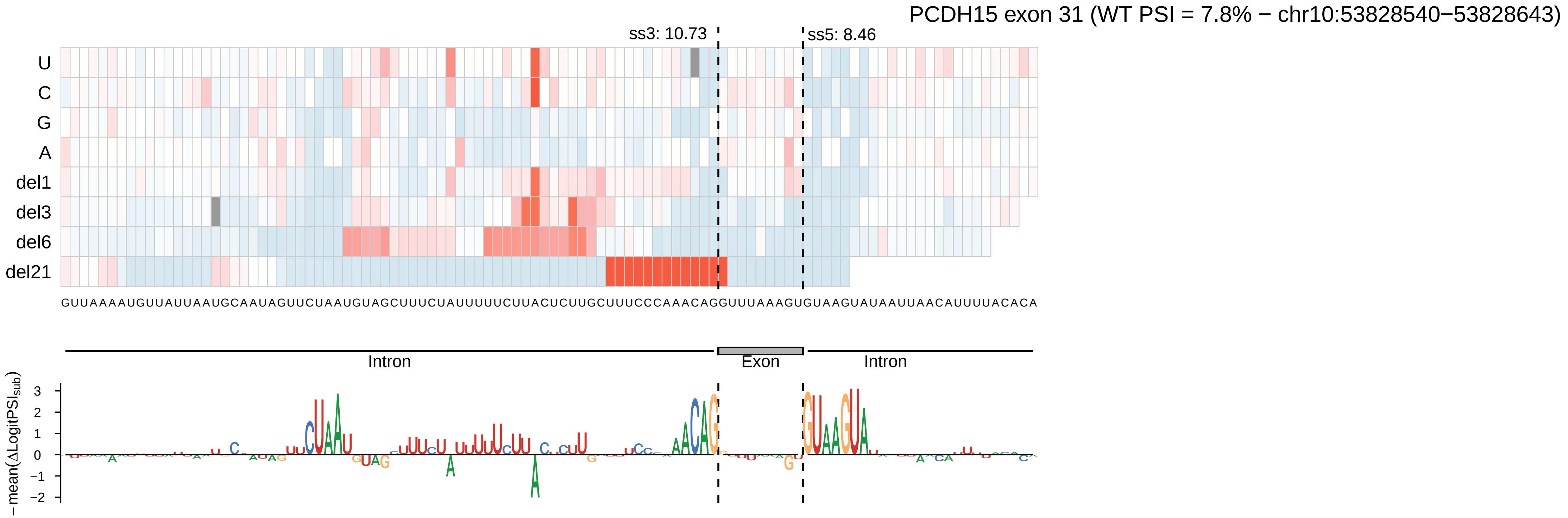
PCDH15 exon 31 (WT PSI = 7.8% − chr10:53828540−53828643)
ss3: 10.73
ss5: 8.46
U
C
G
A
del1
del3
del6
del21
GUUAAAAUGUUAUUAAUGCAAUAGUUCUAAUGUAGCUUUCUAUUUUUCUUACUCUUGCUUUCCCAAACAGGUUUAAAGUGUAAGUAUAAUUAACAUUUUACACA
Intron
Exon
Intron
− mean(ΔLogitPSI_sub)

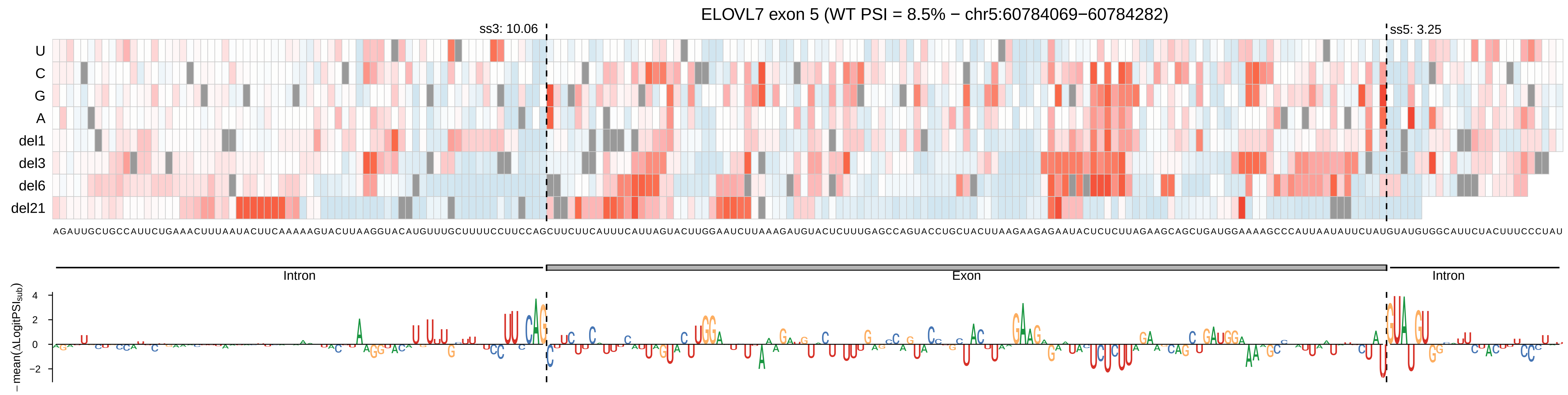
ELOVL7 exon 5 (WT PSI = 8.5% − chr5:60784069−60784282)
ss3: 10.06
ss5: 3.25
U
C
G
A
del1
del3
del6
del21
AGAUUGCUGCCAUUCUGAAACUUUAAUACUUCAAAAAGUACUUAAGGUACAUGUUUGCUUUUCCUUCCAGCUUCUUCAUUUCAUUAGUACUUGGAAUCUUAAAGAUGUACUCUUUGAGCCAGUACCUGCUACUUAAGAAGAGAAUACUCUCUUAGAAGCAGCUGAUGGAAAAGCCCAUUAAUAUUCUAUGUAUGUGGCAUUCUACUUUCCCUAU
Intron
Exon
Intron
−mean(ΔLogitPSI_sub)
4
2
0
−2

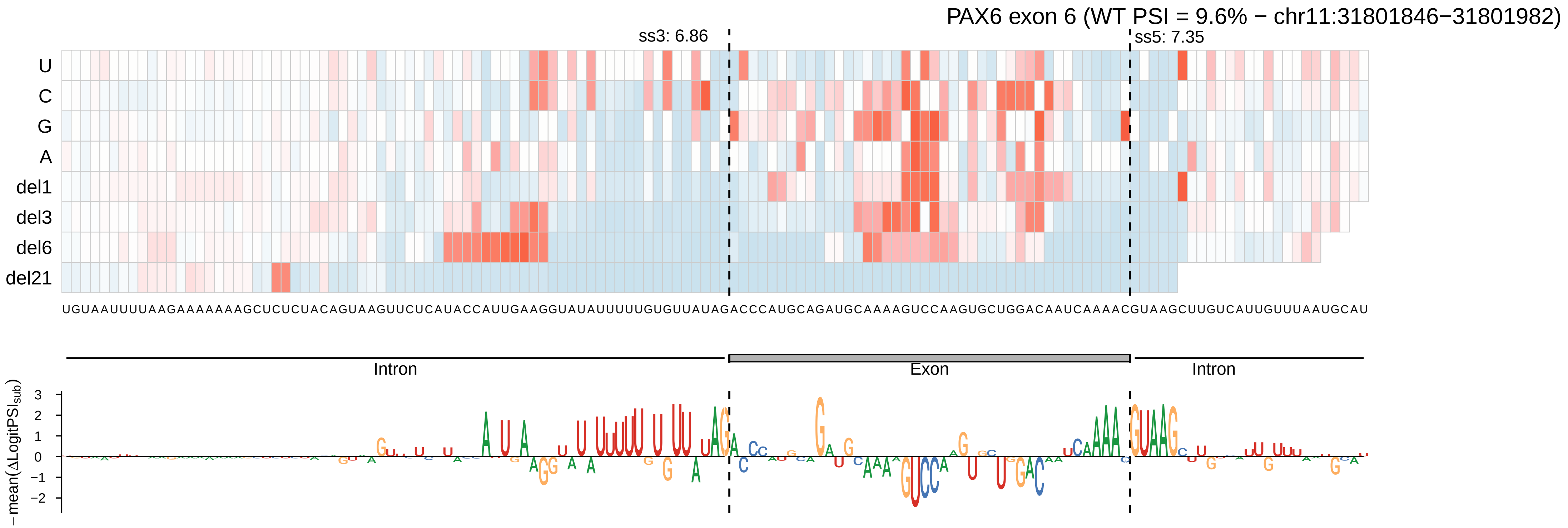

PAX6 exon 6 (WT PSI = 9.6% − chr11:31801846−31801982)
ss3: 6.86
ss5: 7.35
U
C
G
A
del1
del3
del6
del21
Intron
Exon
Intron
−mean(ΔLogitPSI_sub)
3
2
1
0
−1
−2

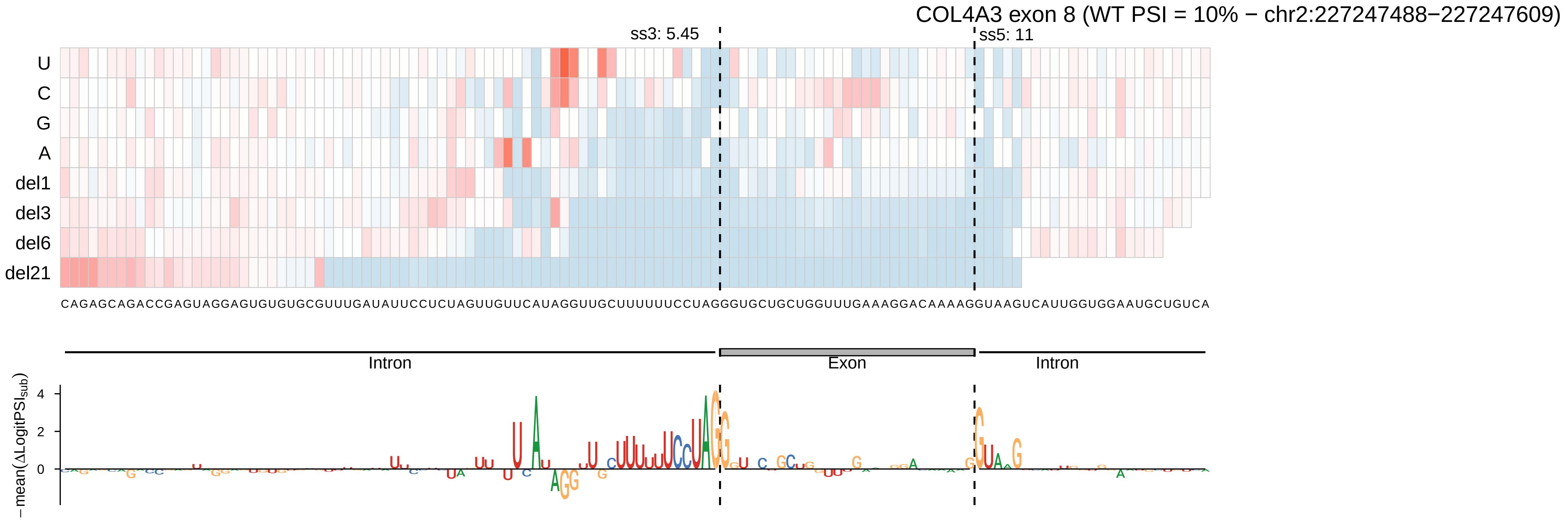

COL4A3 exon 8 (WT PSI = 10% − chr2:227247488−227247609)
ss3: 5.45
ss5: 11
U
C
G
A
del1
del3
del6
del21
CAGAGCAGACCGAGUAGGAGUGUGUGCGUUUGAUAUUCCUCUAGUUGUUCAUAGGUUGCUUUUUUCCUAGGGUGCUGCUGGUUUGAAAGGACAAAAGGUAAGUCAUUGGUGGAAUGCUGUCA
Intron
Exon
Intron
−mean(ΔLogitPSI_sub)
4
2
0

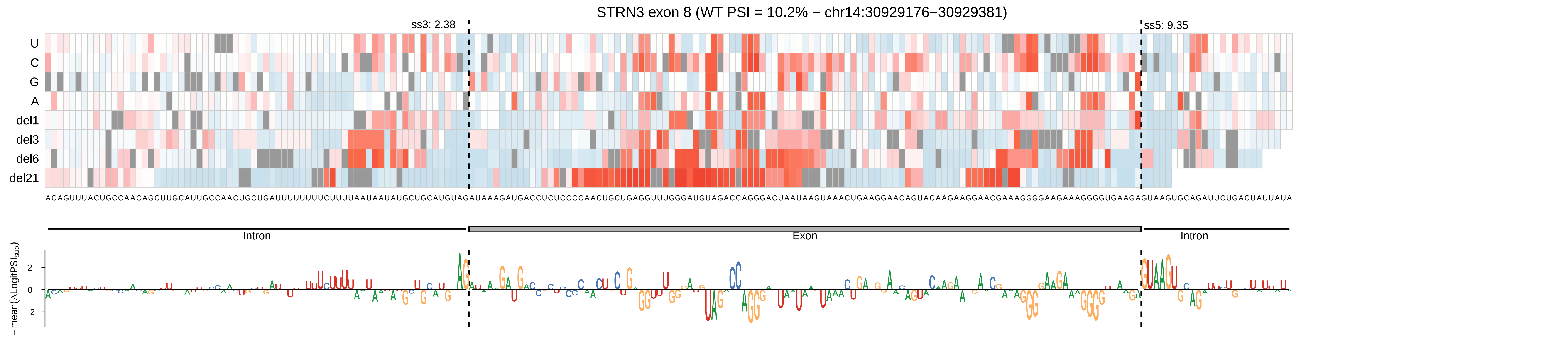
STRN3 exon 8 (WT PSI = 10.2% − chr14:30929176−30929381)
ss3: 2.38
ss5: 9.35
U
C
G
A
del1
del3
del6
del21
ACAGUUUACUGCCAACAGCUUGCAUUGCCAACUGCUGAUUUUUUUUCUUUUAAUAAUAUGCUGCAUGUAGAUAAAGAUGACCUCUCCCCAACUGCUGAGGUUUGGGAUGUAGACCAGGGACUAAUAAGUAAACUGAAGGAACAGUACAAGAAGGAACGAAAGGGGAAGAAAGGGGUGAAGAGUAAGUGCAGAUUCUGACUAUUAUA
Intron
Exon
Intron
−mean(ΔLogitPSI_sub)
2
0
−2

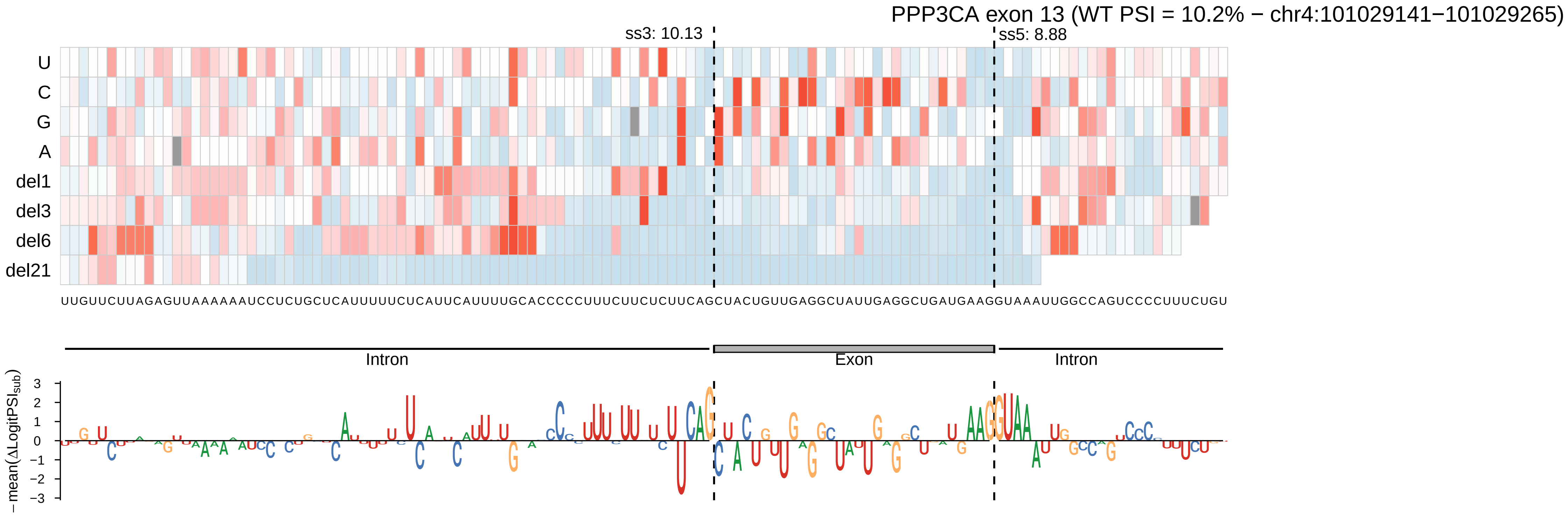
PPP3CA exon 13 (WT PSI = 10.2% − chr4:101029141−101029265)
ss3: 10.13
ss5: 8.88
U
C
G
A
del1
del3
del6
del21
UUGUUCUUAGAGUUAAAAAAUCCUCUGCUCAUUUUUCUCAUUCAUUUUGCACCCCCUUUCUUCUCUUCAGCUACUGUUGAGGCUAUUGAGGCUGAUGAAGGUAAAUUGGCCAGUCCCCUUUCUGU
Intron
Exon
Intron
− mean(ΔLogitPSI_sub)
3
2
1
0
−1
−2
−3

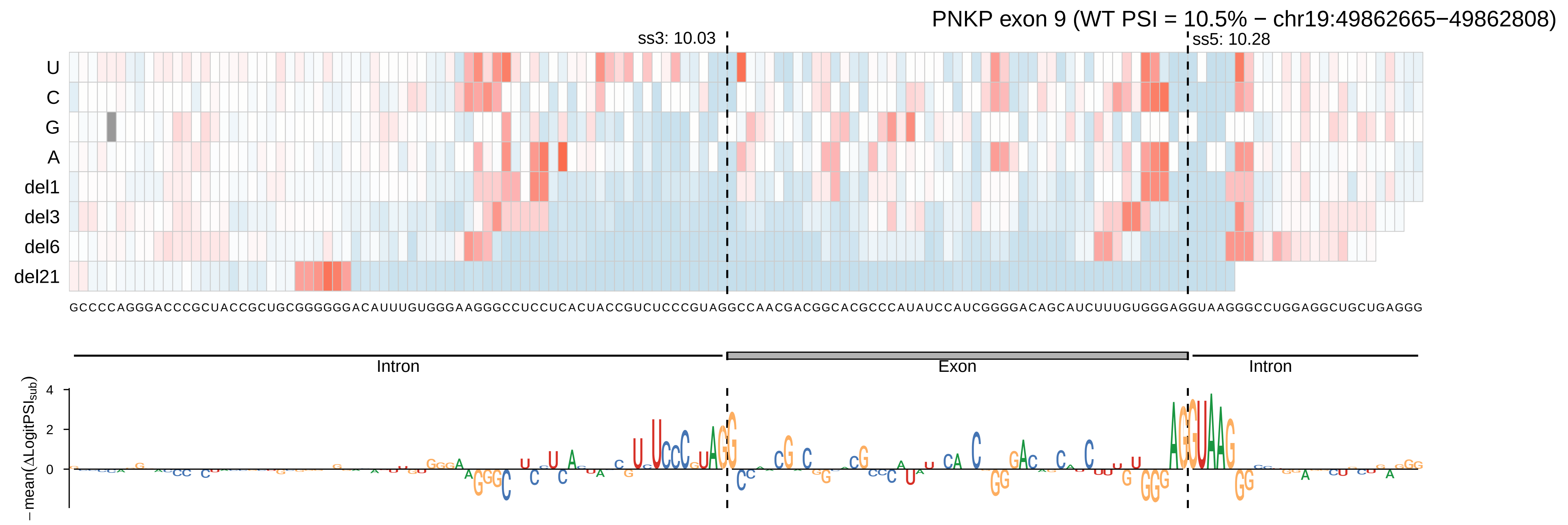

PNKP exon 9 (WT PSI = 10.5% − chr19:49862665−49862808)
ss3: 10.03
ss5: 10.28
U
C
G
A
del1
del3
del6
del21
GCCCCAGGGACCCGCUACCGCUGCGGGGGGACAUUUGUGGGAAGGGCCUCCUCACUACCGUCUCCCGUAGGCCAACGACGGCACGCCCAUAUCCAUCGGGGACAGCAUCUUUGUGGGAGGUAAGGGCCUGGAGGCUGCUGAGGG
Intron
Exon
Intron
−mean(ΔLogitPSI_sub)
4
2
0

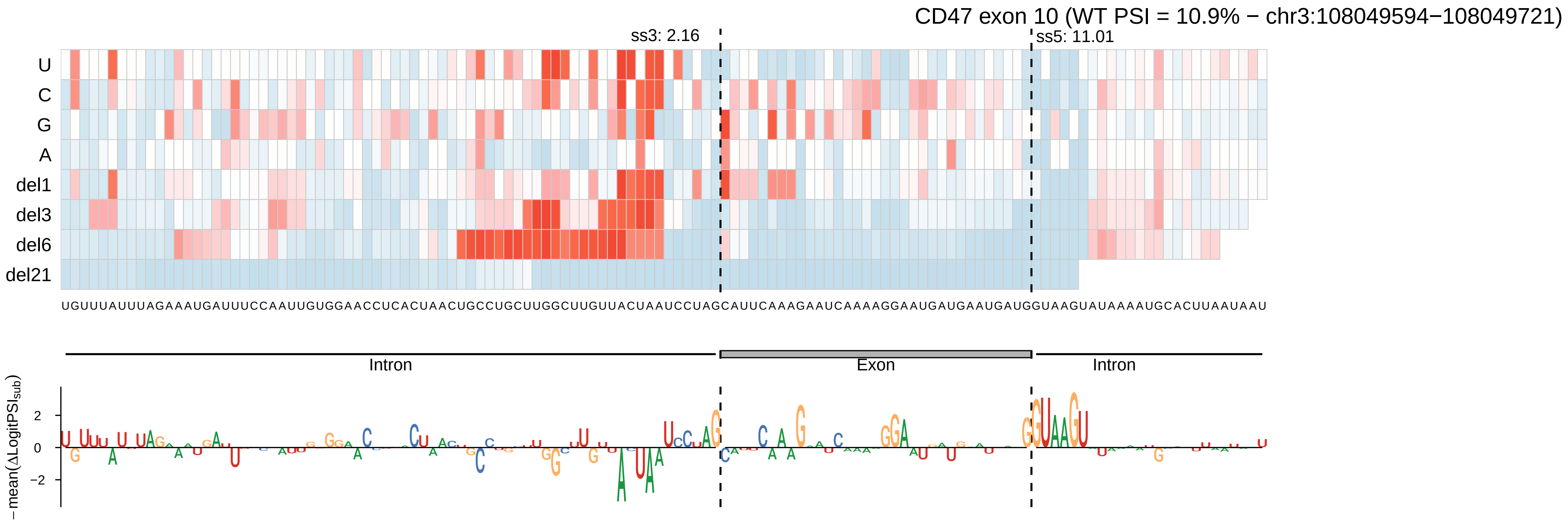

CD47 exon 10 (WT PSI = 10.9% − chr3:108049594−108049721)
ss3: 2.16
ss5: 11.01
U
C
G
A
del1
del3
del6
del21
UGUUUAUUUAGAAAUGAUUUCCAAUUGUGGAACCUCACUAACUGCCUGCUUGGCUUGUUACUAAUCCUAGCAUUCAAAGAAUCAAAAGGAAUGAUGAAUGAUGGUAAGUAUAAAAUGCACUUAAUAAU
Intron
Exon
Intron
− mean(ΔLogitPSI_sub)
2
0
−2

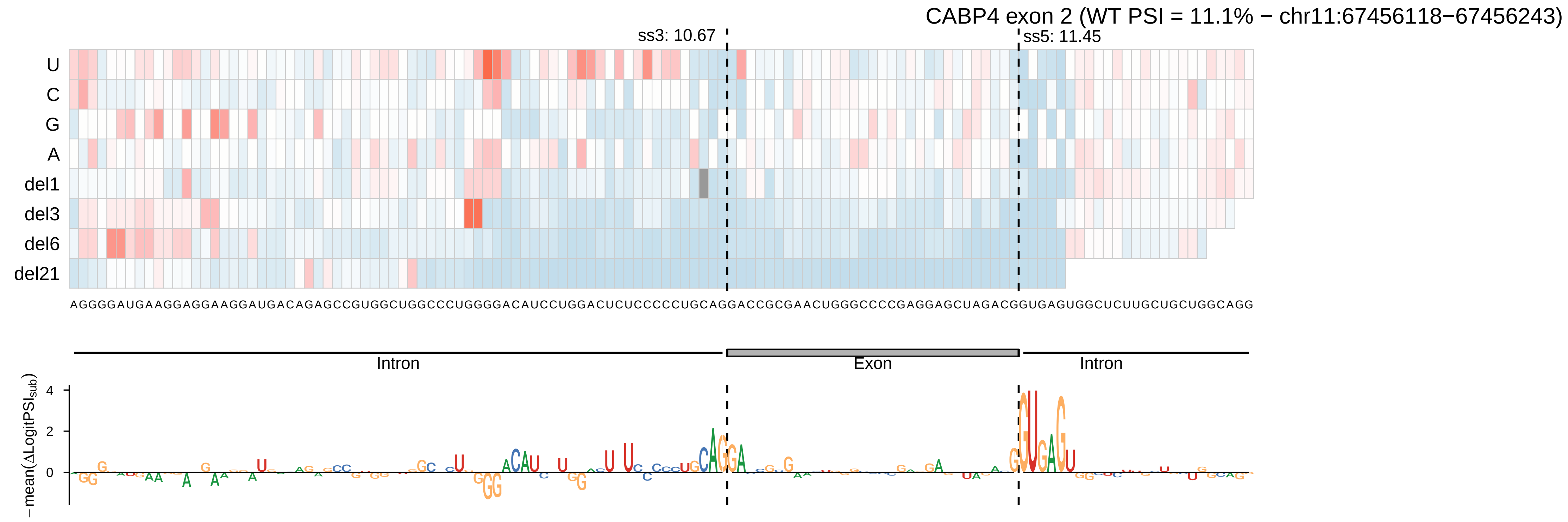

CABP4 exon 2 (WT PSI = 11.1% − chr11:67456118−67456243)
ss3: 10.67
ss5: 11.45
U
C
G
A
del1
del3
del6
del21
AGGGGAUGAAGGAGGAAGGAUGACAGAGCCGUGGCUGGCCCUGGGGACAUCCUGGACUCUCCCCCUGCAGGACCGCGAACUGGGCCCCGAGGAGCUAGACGGUGAGUGGCUCUUGCUGCUGGCAGG
Intron
Exon
Intron
−mean(ΔLogitPSI_sub)
4
2
0

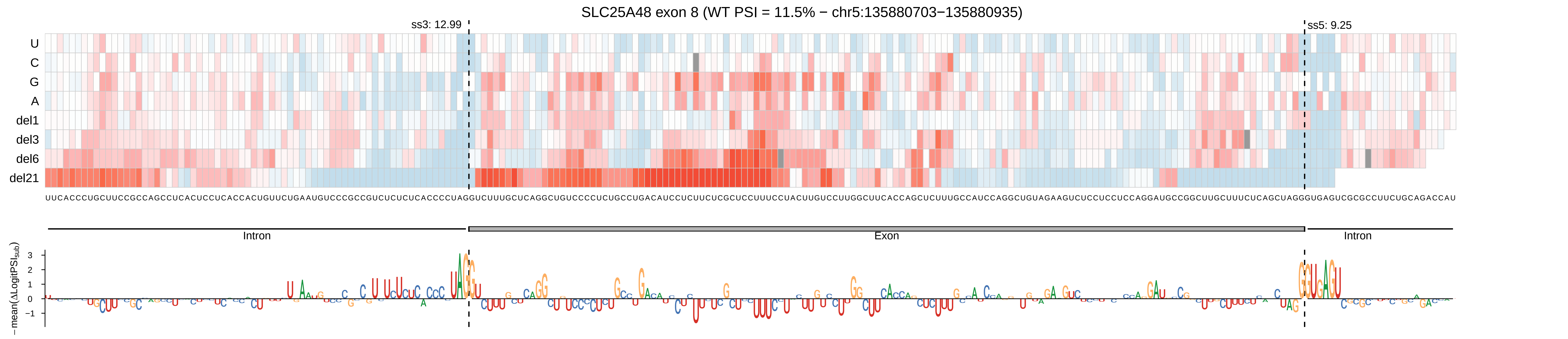

SLC25A48 exon 8 (WT PSI = 11.5% − chr5:135880703−135880935)
ss3: 12.99
ss5: 9.25
U
C
G
A
del1
del3
del6
del21
Intron
Exon
Intron
−mean(ΔLogitPSI_sub)

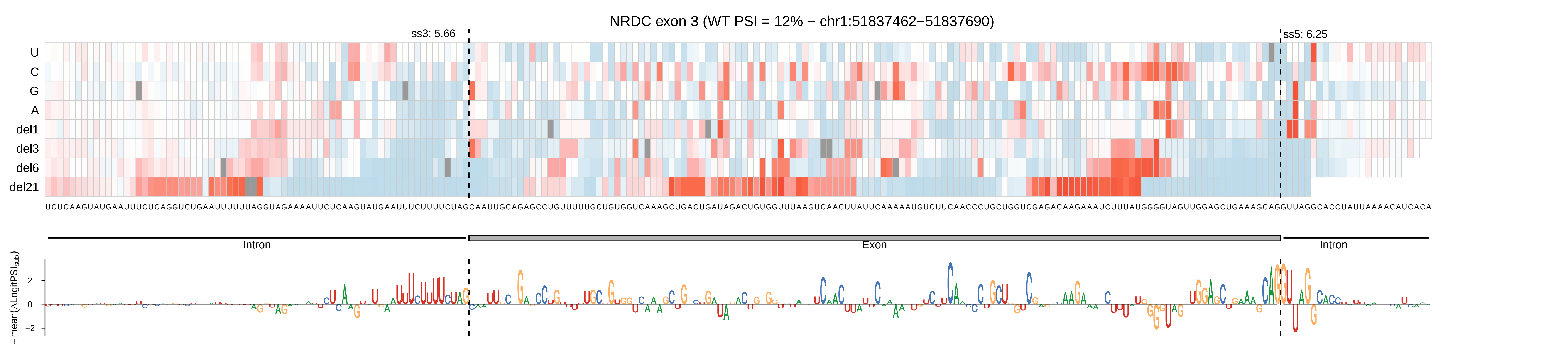

NRDC exon 3 (WT PSI = 12% − chr1:51837462−51837690)
ss3: 5.66
ss5: 6.25
U
C
G
A
del1
del3
del6
del21
Intron
Exon
Intron
−mean(ΔLogitPSIsub)
2
0
−2

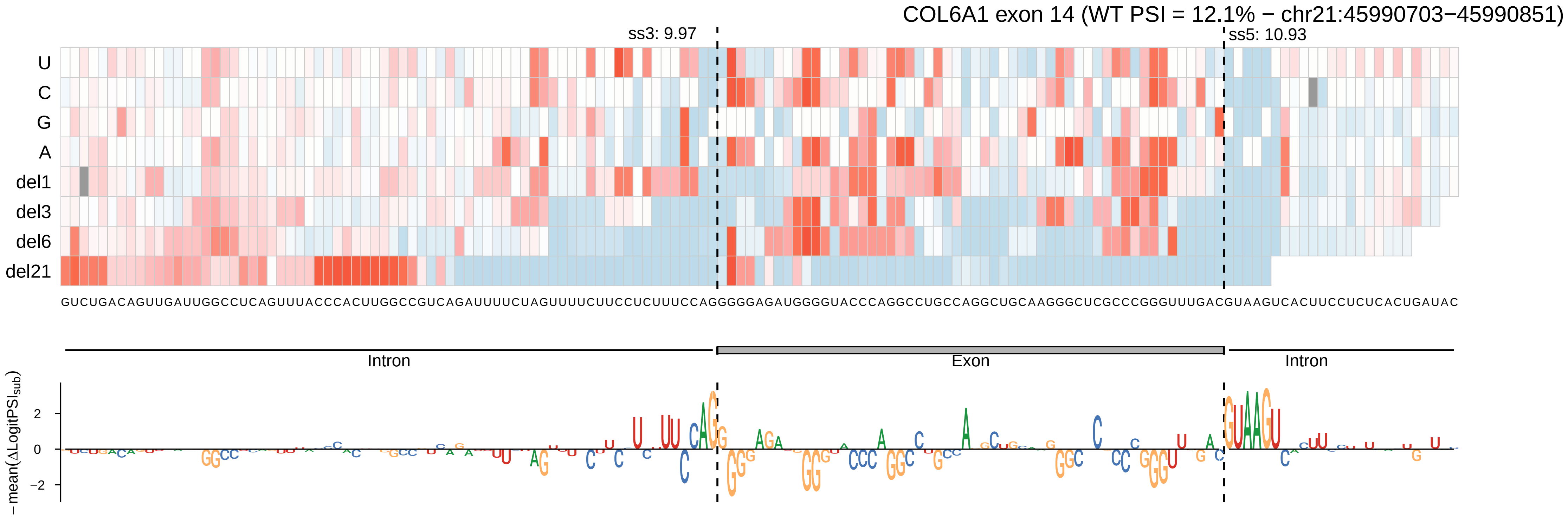

COL6A1 exon 14 (WT PSI = 12.1% − chr21:45990703−45990851)
ss3: 9.97
ss5: 10.93
U
C
G
A
del1
del3
del6
del21
GUCUGACAGUUGAUUGGCCUCAGUUUACCCACUUGGCCGUCAGAUUUUCUAGUUUUCUUCCUCUUUCCAGGGGGAGAUGGGGUACCCAGGCCUGCCAGGCUGCAAGGGCUCGCCCGGGUUUGACGUAAGUCACUUCCUCUCACUGAUAC
Intron
Exon
Intron
−mean(ΔLogitPSI_sub)
2
0
−2

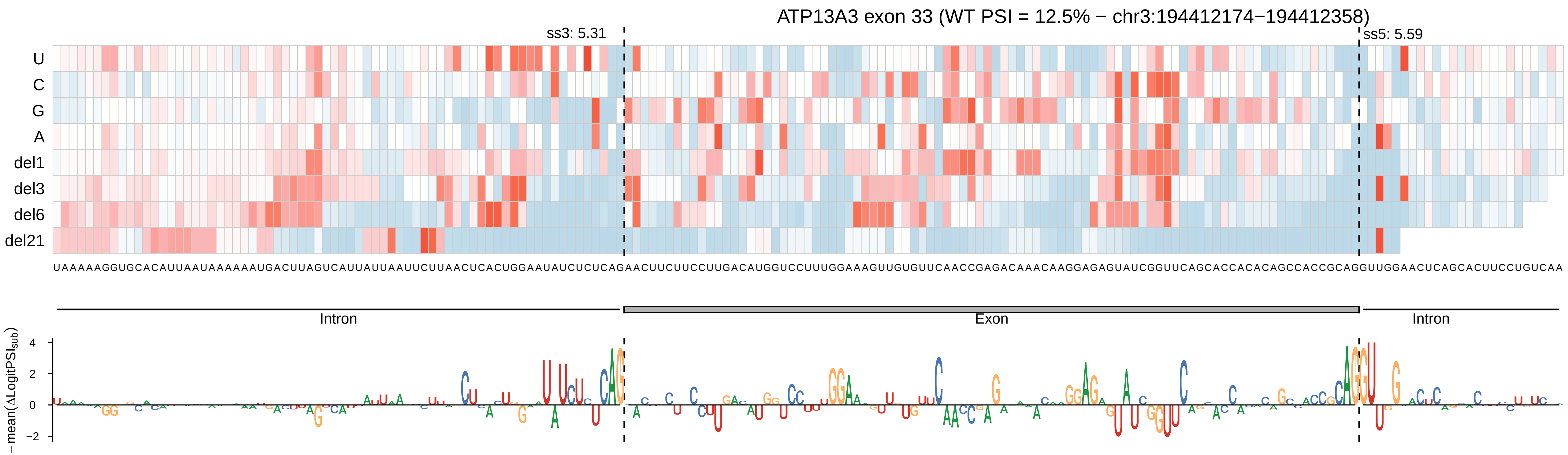

ATP13A3 exon 33 (WT PSI = 12.5% − chr3:194412174−194412358)
ss3: 5.31
ss5: 5.59
U
C
G
A
del1
del3
del6
del21
UAAAAAGGUGCACAUUAAUAAAAAAUGACUUAGUCAUUAUUAAUUCUUAACUCACUGGAAUAUCUCUCAGAACUUCUUCCUUGACAUGGUCCUUUGGAAAGUUGUGUUCAACCGAGACAAACAAGGAGAGUAUCGGUUCAGCACCACACAGCCACCGCAGGUUGGAACUCAGCACUUCCUGUCAA
Intron
Exon
Intron
−mean(ΔLogitPSI_sub)
4
2
0
−2

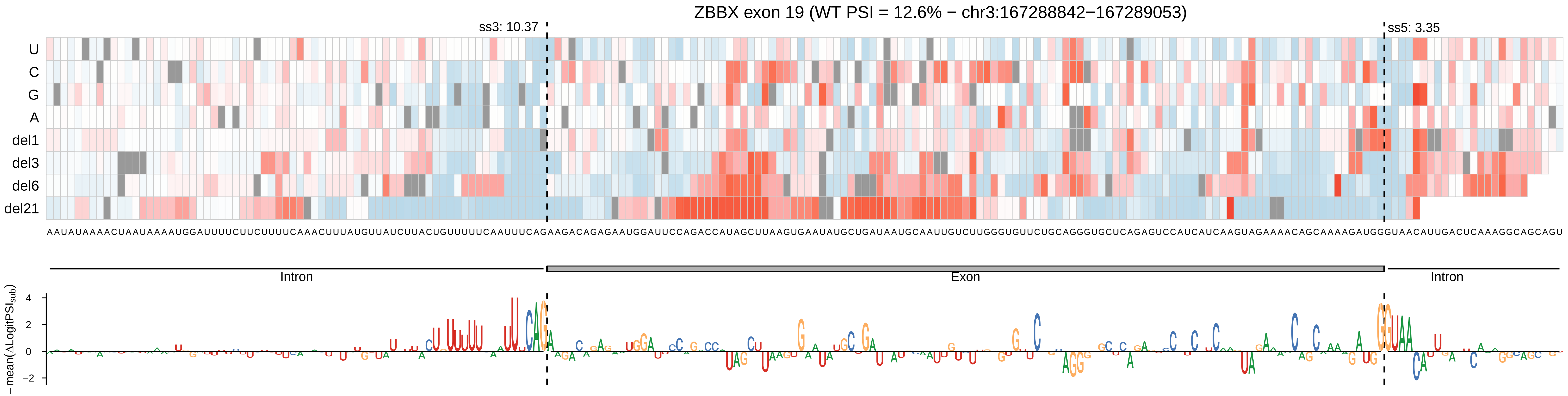

ZBBX exon 19 (WT PSI = 12.6% − chr3:167288842−167289053)
ss3: 10.37
ss5: 3.35
U
C
G
A
del1
del3
del6
del21
AAUAUAAAACUAAUAAAAUGGAUUUUCUUCUUUUCAAACUUUAUGUUAUCUUACUGUUUUUCAAUUUCAGAAGACAGAGAAUGGAUUCCAGACCAUAGCUUAAGUGAAUAUGCUGAUAAUGCAAUUGUCUUGGGUGUUCUGCAGGGUGCUCAGAGUCCAUCAUCAAGUAGAAAACAGCAAAAGAUGGGUAACAUUGACUCAAAGGCAGCAGU
Intron
Exon
Intron
−mean(ΔLogitPSI_sub)
4
2
0
−2

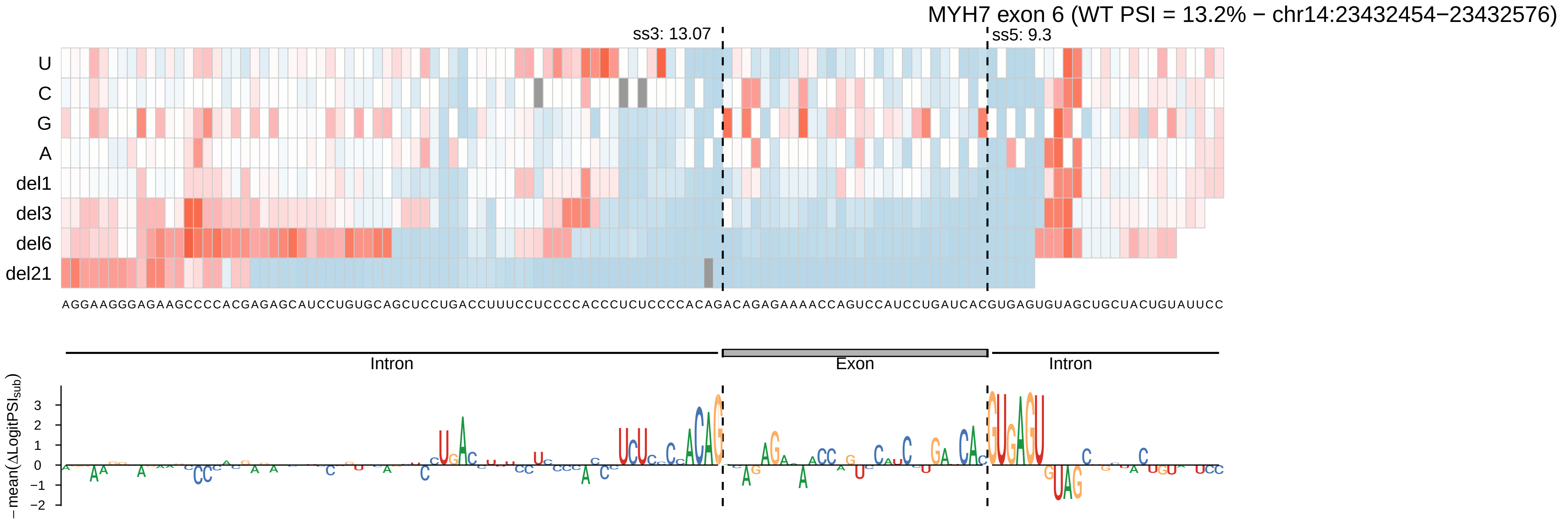

MYH7 exon 6 (WT PSI = 13.2% − chr14:23432454−23432576)
ss3: 13.07
ss5: 9.3
U
C
G
A
del1
del3
del6
del21
AGGAAGGGAGAAGCCCCACGAGAGCAUCCUGUGCAGCUCCUGACCUUUCCUCCCCACCCUCUCCCCACAGACAGAGAAAACCAGUCCAUCCUGAUCACGUGAGUGUAGCUGCUACUGUAUUCC
Intron
Exon
Intron
− mean(ΔLogitPSI_sub)
3
2
1
0
−1
−2

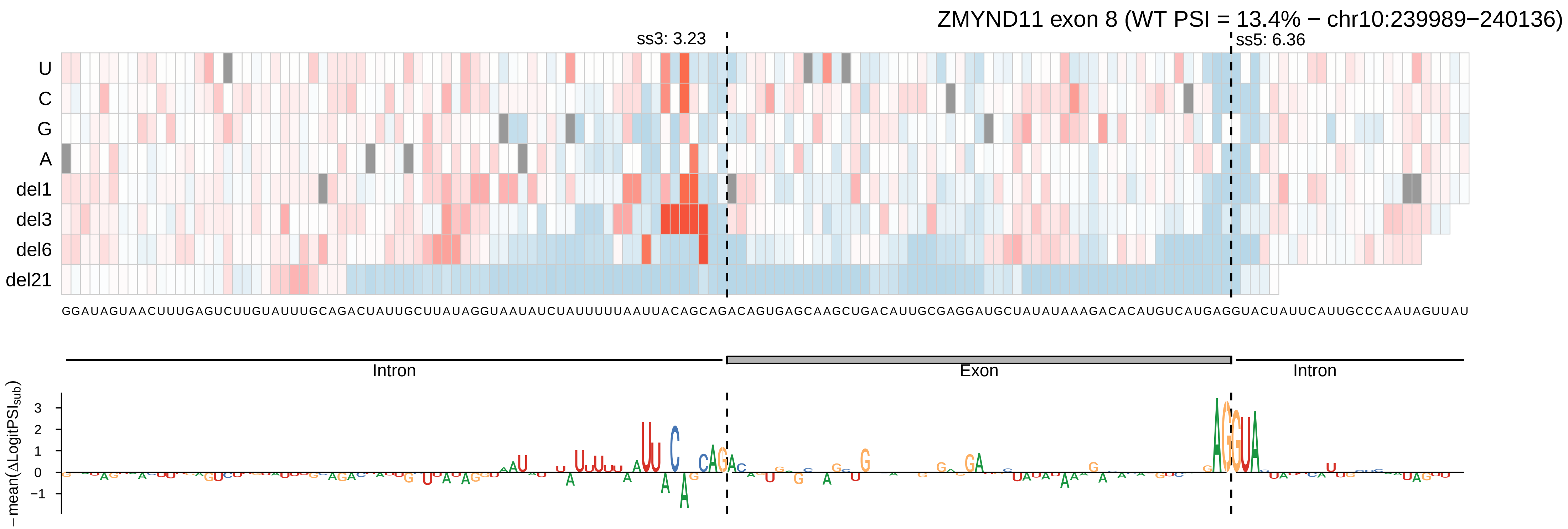
ZMYND11 exon 8 (WT PSI = 13.4% − chr10:239989−240136)
ss3: 3.23
ss5: 6.36
U
C
G
A
del1
del3
del6
del21
GGAUAGUAACUUUGAGUCUUGUAUUUGCAGACUAUUGCUUAUAGGUAAUAUCUAUUUUUAAUUACAGCAGACAGUGAGCAAGCUGACAUUGCGAGGAUGCUAUAUAAAGACACAUGUCAUGAGGUACUAUUCAUUGCCCAAUAGUUAU
Intron
Exon
Intron
−mean(ΔLogitPSI_sub)
3
2
1
0
−1

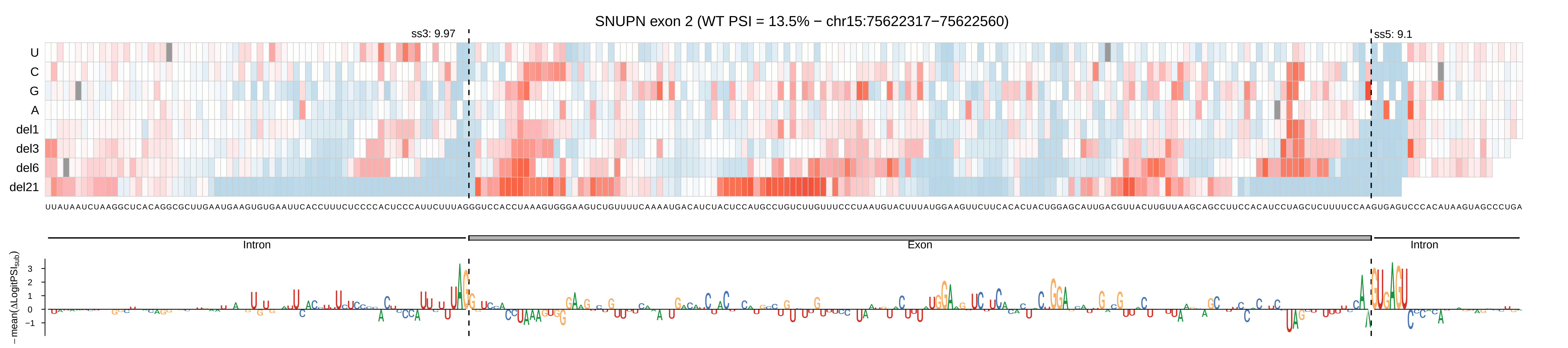
SNUPN exon 2 (WT PSI = 13.5% − chr15:75622317−75622560)
ss3: 9.97
ss5: 9.1
U
C
G
A
del1
del3
del6
del21
Intron
Exon
Intron
−mean(ΔLogitPSI_sub)
3
2
1
0
−1

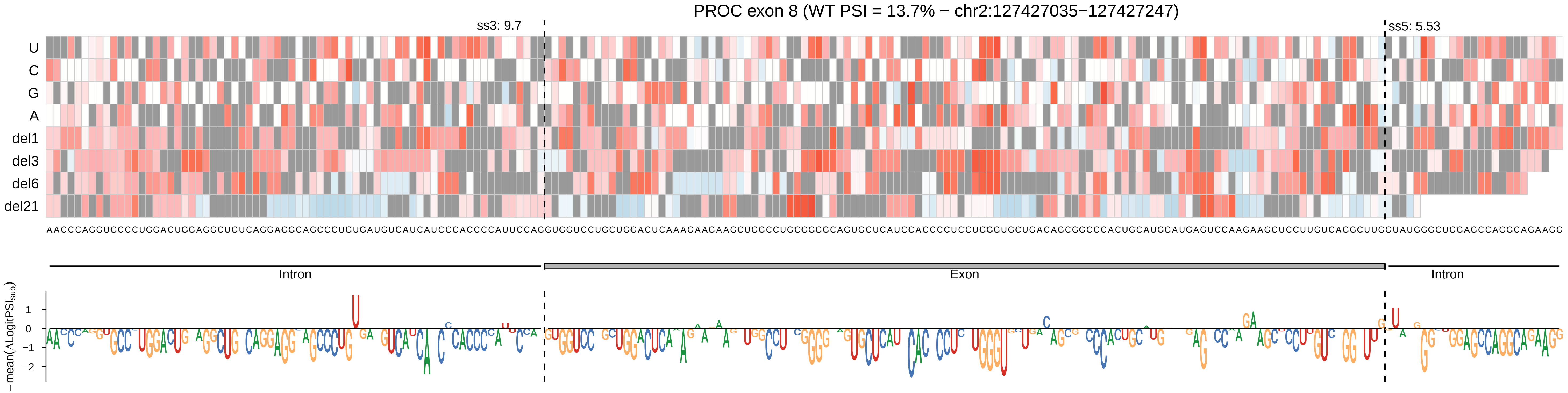

PROC exon 8 (WT PSI = 13.7% − chr2:127427035−127427247)
ss3: 9.7
ss5: 5.53
U
C
G
A
del1
del3
del6
del21
AACCCAGGUGCCCUGGACUGGAGGCUGUCAGGAGGCAGCCCUGUGAUGUCAUCAUCCCACCCCAUUCCAGGUGGUCCUGCUGGACUCAAAGAAGAAGCUGGCCUGCGGGGCAGUGCUCAUCCACCCCUCCUGGGUGCUGACAGCGGCCCACUGCAUGGAUGAGUCCAAGAAGCUCCUUGUCAGGCUUGGUAUGGGCUGGAGCCAGGCAGAAGG
Intron
Exon
Intron
−mean(ΔLogitPSI_sub)
1
0
−1
−2

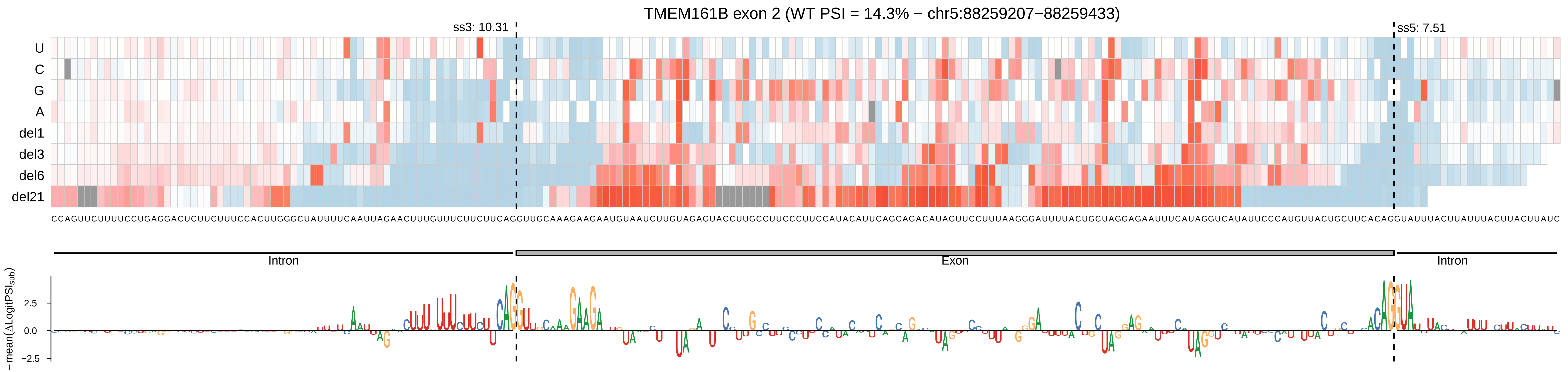
TMEM161B exon 2 (WT PSI = 14.3% − chr5:88259207−88259433)
ss3: 10.31
ss5: 7.51
U
C
G
A
del1
del3
del6
del21
CCAGUUCUUUUCCUGAGGACUCUUCUUUCCACUUGGGCUAUUUUCAAUUAGAACUUUGUUUCUUCUUCAGGUUGCAAAGAAGAAUGUAAUCUUGUAGAGUACCUUGCCUUCCCUUCCAUACAUUCAGCAGACAUAGUUCCUUUAAGGGAUUUUACUGCUAGGAGAAUUUCAUAGGUCAUAUUCCCAUGUUACUGCUUCACAGGUAUUUACUUAUUUACUUACUUAUC
Intron
Exon
Intron
−mean(ΔLogitPSI_sub)
2.5
0.0
−2.5

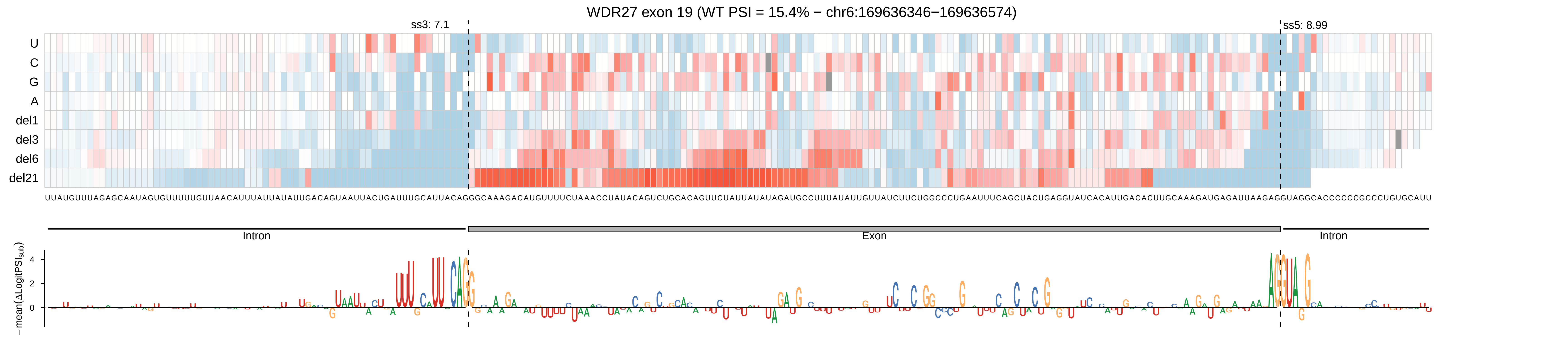
WDR27 exon 19 (WT PSI = 15.4% − chr6:169636346−169636574)
ss3: 7.1
ss5: 8.99
U
C
G
A
del1
del3
del6
del21
UUAUGUUUAGAGCAAUAGUGUUUUUGUUAACAUUUAUUAUAUUGACAGUAAUUACUGAUUUGCAUUACAGGGCAAAGACAUGUUUUCUAAACCUAUACAGUCUGCACAGUUCUAUUAUAUAGAUGCCUUUAUAUUGUUAUCUUCUGGCCCUGAAUUUCAGCUACUGAGGUAUCACAUUGACACUUGCAAAGAUGAGAUUAAGAGGUAGGCACCCCCCGCCCUGUGCAUU
Intron
Exon
Intron
−mean(ΔLogitPSI_sub)
4
2
0

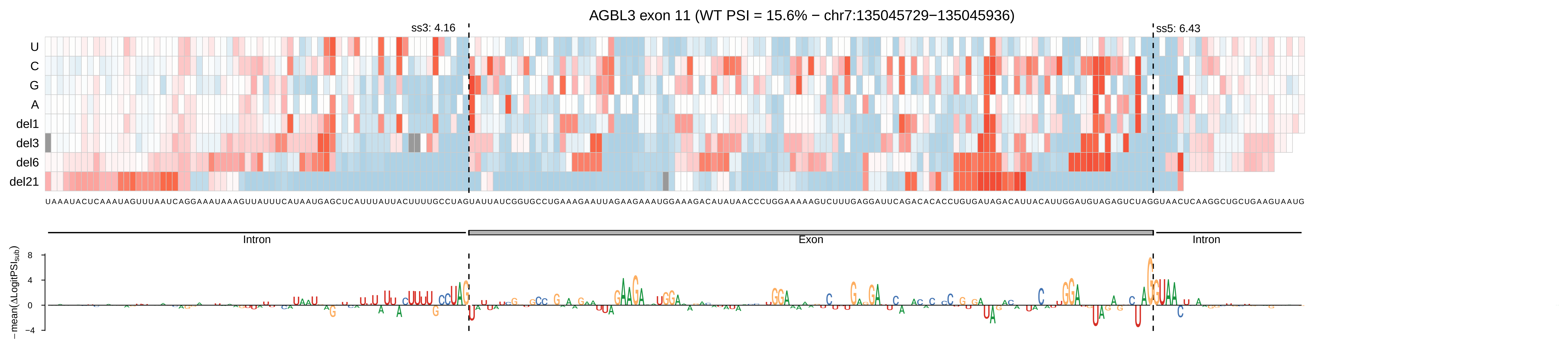

AGBL3 exon 11 (WT PSI = 15.6% − chr7:135045729−135045936)
ss3: 4.16
ss5: 6.43
U
C
G
A
del1
del3
del6
del21
UAAAUACUCAAAUAGUUUAAUCAGGAAAUAAAGUUAUUUCAUAAUGAGCUCAUUUAUUACUUUUGCCUAGUAUUAUCGGUGCCUGAAAGAAUUAGAAGAAAUGGAAAGACAUAUAACCCUGGAAAAAGUCUUUGAGGAUUCAGACACACCUGUGAUAGACAUUACAUUGGAUGUAGAGUCUAGGUAACUCAAGGCUGCUGAAGUAAUG
Intron
Exon
Intron
− mean(ΔLogitPSI_sub)
8
4
0
−4

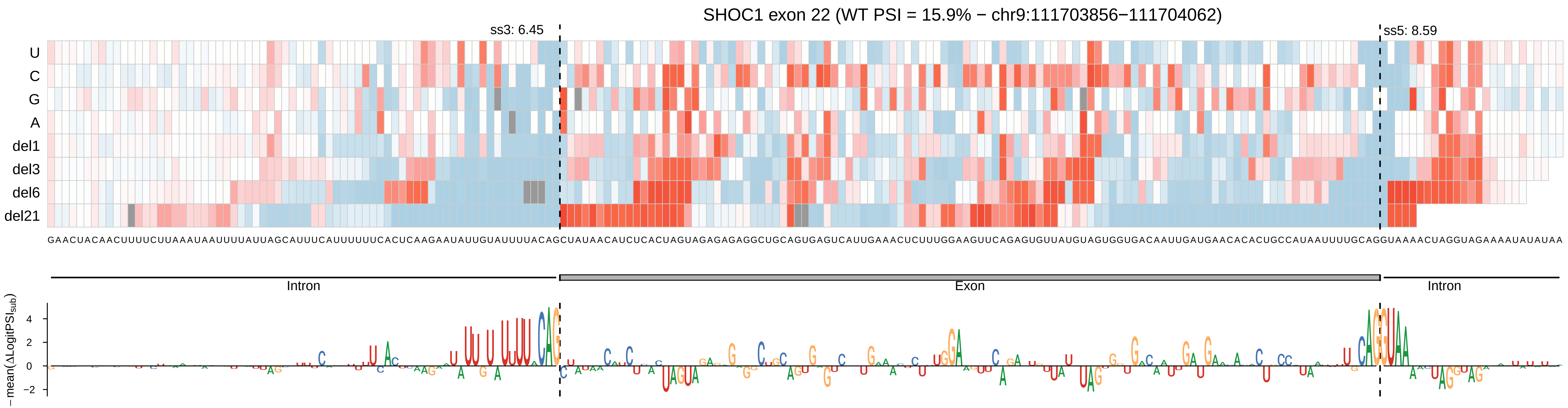

SHOC1 exon 22 (WT PSI = 15.9% − chr9:111703856−111704062)
ss3: 6.45
ss5: 8.59
U
C
G
A
del1
del3
del6
del21
GAACUACAACUUUUCUUAAAUAAUUUUAUUAGCAUUUCAUUUUUUCACUCAAGAAUAUUGUAUUUUACAGCUAUAACAUCUCACUAGUAGAGAGAGGCUGCAGUGAGUCAUUGAAACUCUUUGGAAGUUCAGAGUGUUAUGUAGUGGUGACAAUUGAUGAACACACUGCCAUAAUUUUGCAGGUAAAACUAGGUAGAAAAUAUAUAA
Intron
Exon
Intron
−mean(ΔLogitPSI_sub)
4
2
0
−2

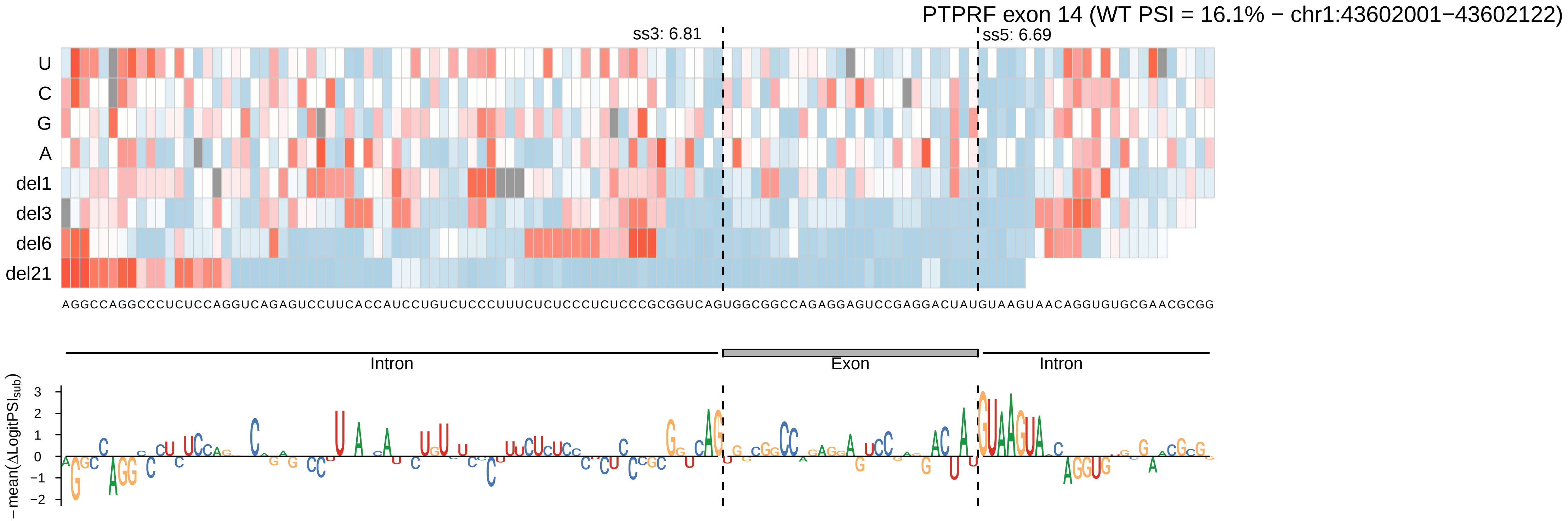

PTPRF exon 14 (WT PSI = 16.1% − chr1:43602001−43602122)
ss3: 6.81
ss5: 6.69
U
C
G
A
del1
del3
del6
del21
AGGCCAGGCCCUCUCCAGGUCAGAGUCCUUCACCAUCCUGUCUCCCUUUCUCUCCCUCUCCCGCGGUCAGUGGCGGCCAGAGGAGUCCGAGGACUAUGUAAGUAACAGGUGUGCGAACGCGG
Intron
Exon
Intron
− mean(ΔLogitPSI_sub)
3
2
1
0
−1
−2

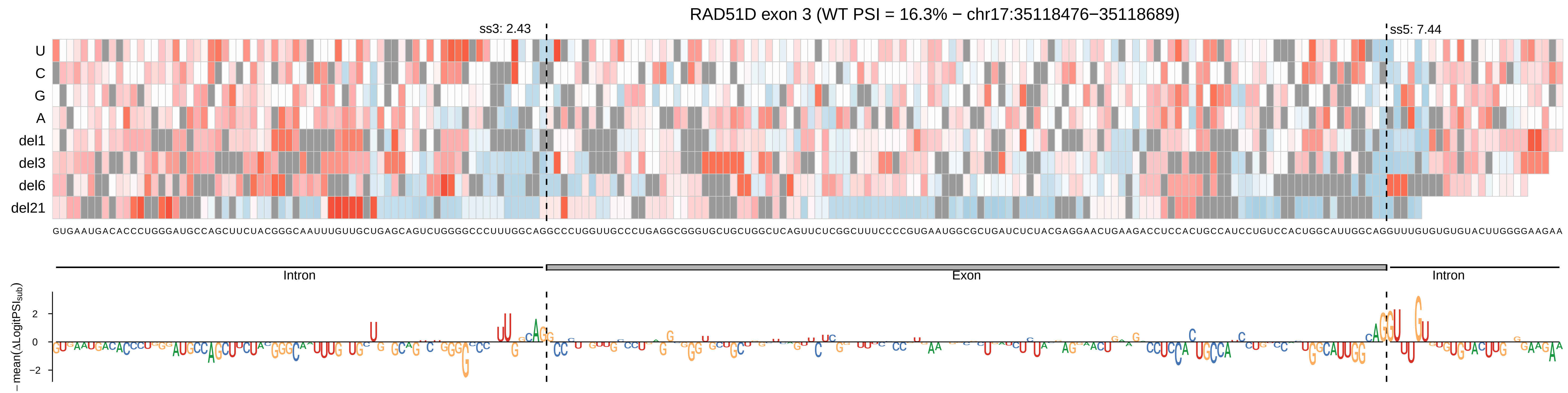

RAD51D exon 3 (WT PSI = 16.3% − chr17:35118476−35118689)
ss3: 2.43
ss5: 7.44
U
C
G
A
del1
del3
del6
del21
Intron
Exon
Intron
−mean(ΔLogitPSI_sub)

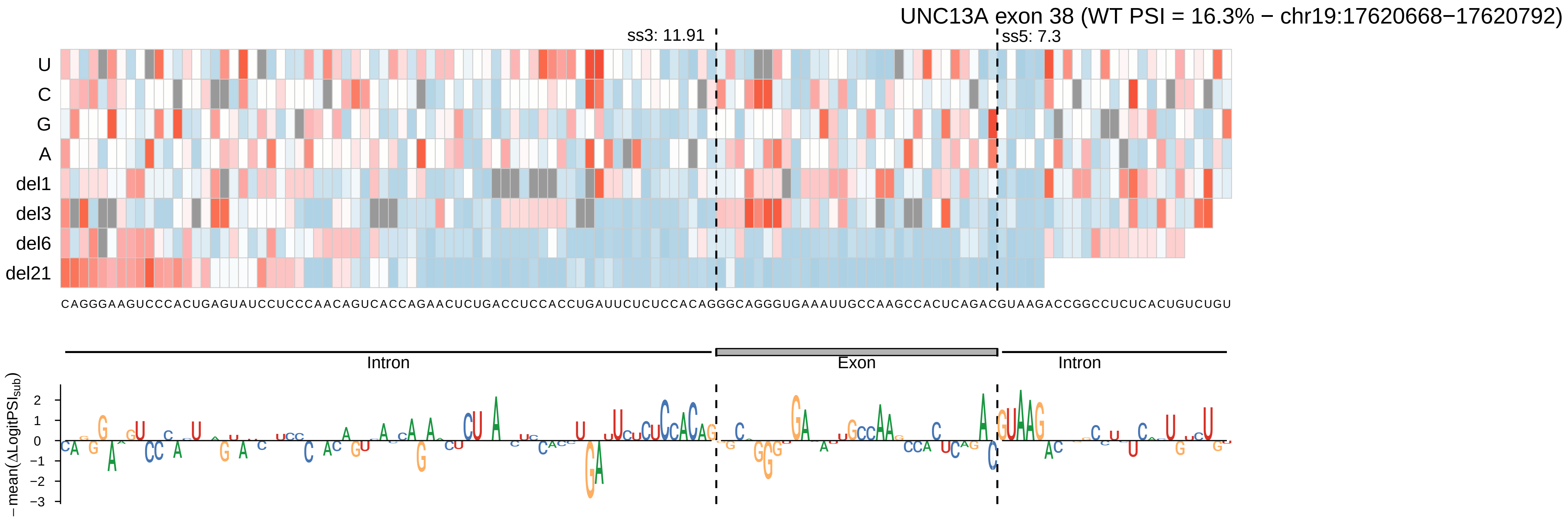

UNC13A exon 38 (WT PSI = 16.3% − chr19:17620668−17620792)
ss3: 11.91
ss5: 7.3
U
C
G
A
del1
del3
del6
del21
CAGGGAAGUCCCACUGAGUAUCCUCCCAACAGUCACCAGAACUCUGACCUCCACCUGAUUCUCUCCACAGGGCAGGGUGAAAUUGCCAAGCCACUCAGACGUAAGACCGGCCUCUCACUGUCUGU
Intron
Exon
Intron
− mean(ΔLogitPSI_sub)
2
1
0
−1
−2
−3

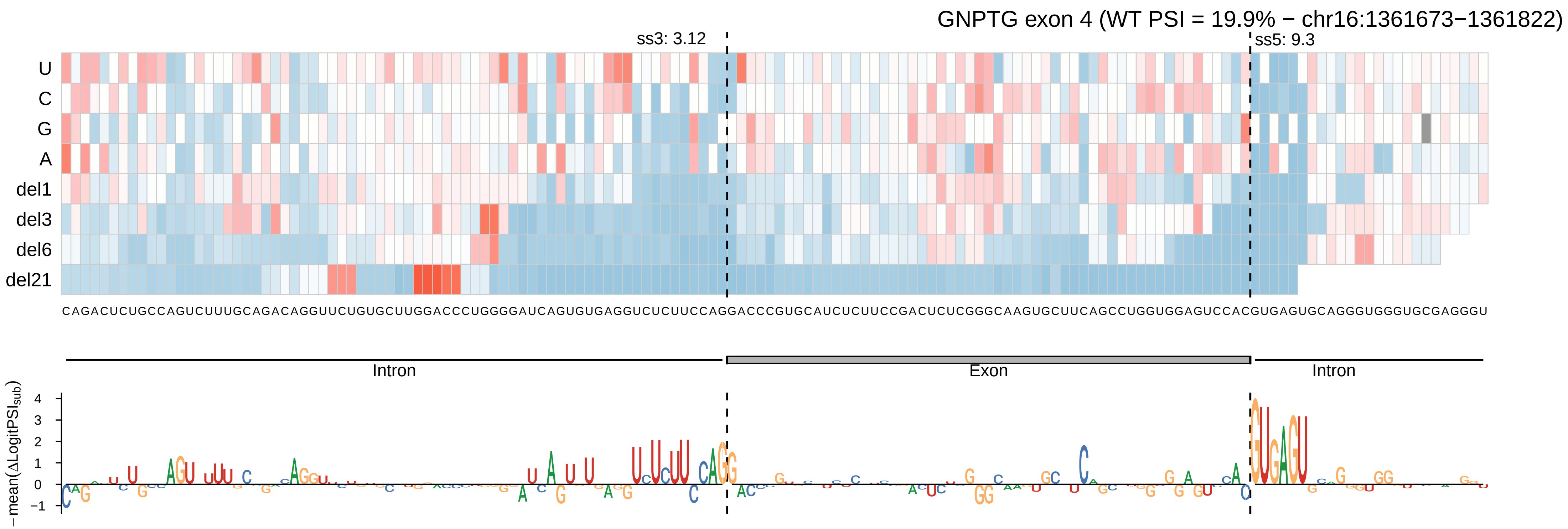
GNPTG exon 4 (WT PSI = 19.9% − chr16:1361673−1361822)
ss3: 3.12
ss5: 9.3
U
C
G
A
del1
del3
del6
del21
CAGACUCUGCCAGUCUUUGCAGACAGGUUCUGUGCUUGGACCCUGGGGAUCAGUGUGAGGUCUCUUCCAGGACCCGUGCAUCUCUUCCGACUCUCGGGCAAGUGCUUCAGCCUGGUGGAGUCCACGUGAGUGCAGGGUGGGUGCGAGGGU
Intron
Exon
Intron
− mean(ΔLogitPSIsub)
4
3
2
1
0
−1

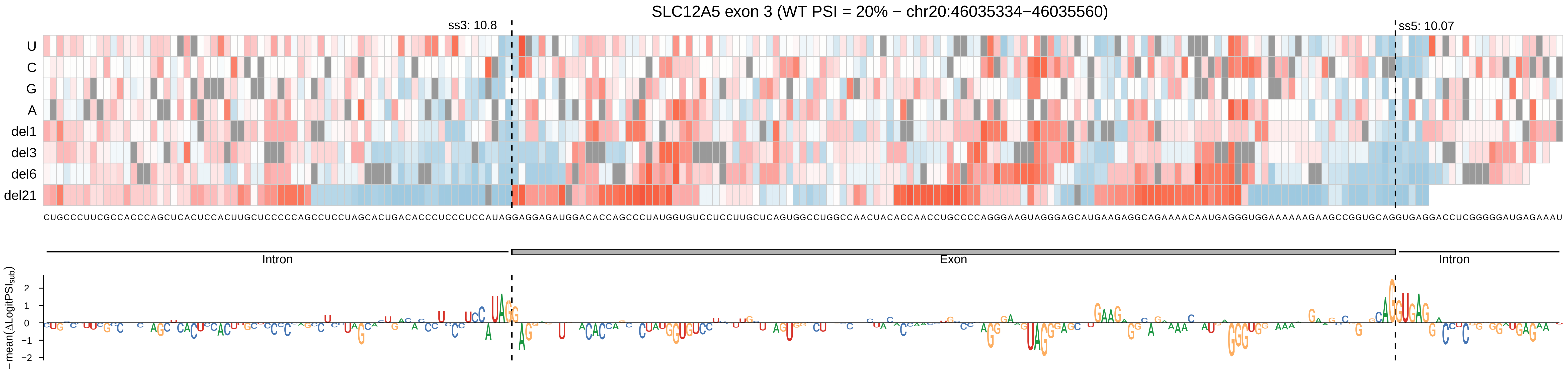
SLC12A5 exon 3 (WT PSI = 20% − chr20:46035334−46035560)
ss3: 10.8
ss5: 10.07
U
C
G
A
del1
del3
del6
del21
CUGCCCUUCGCCACCCAGCUCACUCCACUUGCUCCCCCAGCCUCCUAGCACUGACACCCUCCCUCCAUAGGAGGAGAUGGACACCAGCCCUAUGGUGUCCUCCUUGCUCAGUGGCCUGGCCAACUACACCAACCUGCCCCAGGGAAGUAGGGAGCAUGAAGAGGCAGAAAACAAUGAGGGUGGAAAAAAGAAGCCGGUGCAGGUGAGGACCUCGGGGAUGAGAAAU
Intron
Exon
Intron
−mean(ΔLogitPSI_sub)
2
1
0
−1
−2

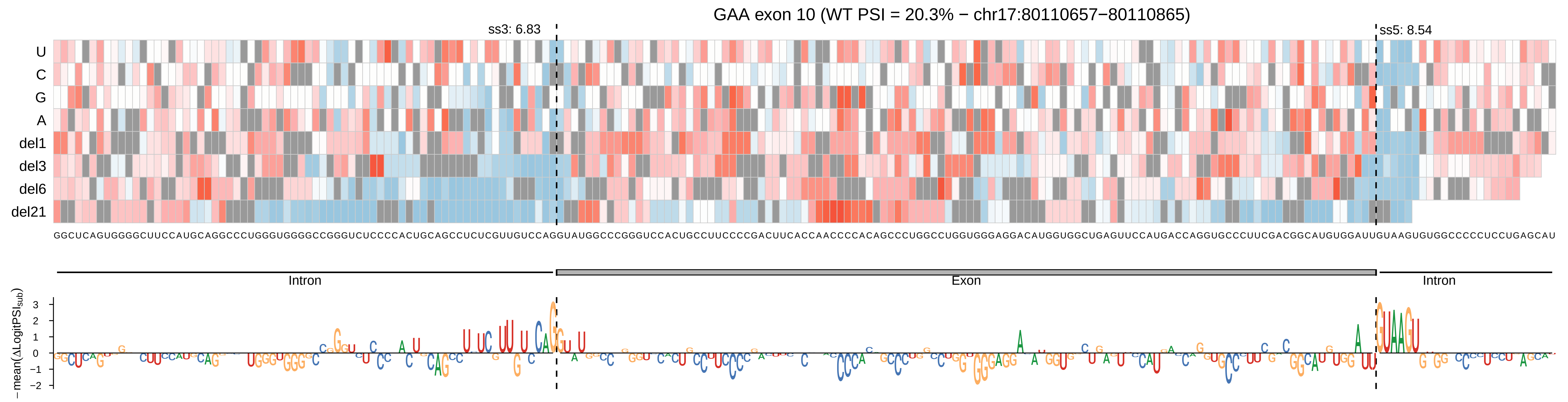
GAA exon 10 (WT PSI = 20.3% − chr17:80110657−80110865)
ss3: 6.83
ss5: 8.54
U
C
G
A
del1
del3
del6
del21
GGCUCAGUGGGGCUUCCAUGCAGGCCCUGGGUGGGGCCGGGUCUCCCCACUGCAGCCUCUCGUUGUCCAGGUAUGGCCCGGGUCCACUGCCUUCCCCGACUUCACCAACCCCACAGCCCUGGCCUGGUGGGAGGACAUGGUGGCUGAGUUCCAUGACCAGGUGCCCUUCGACGGCAUGUGGAUUGUAAGUGUGGCCCCCUCCUGAGCAU
Intron
Exon
Intron
− mean(ΔLogitPSI$_{sub}$)
3
2
1
0
−1
−2

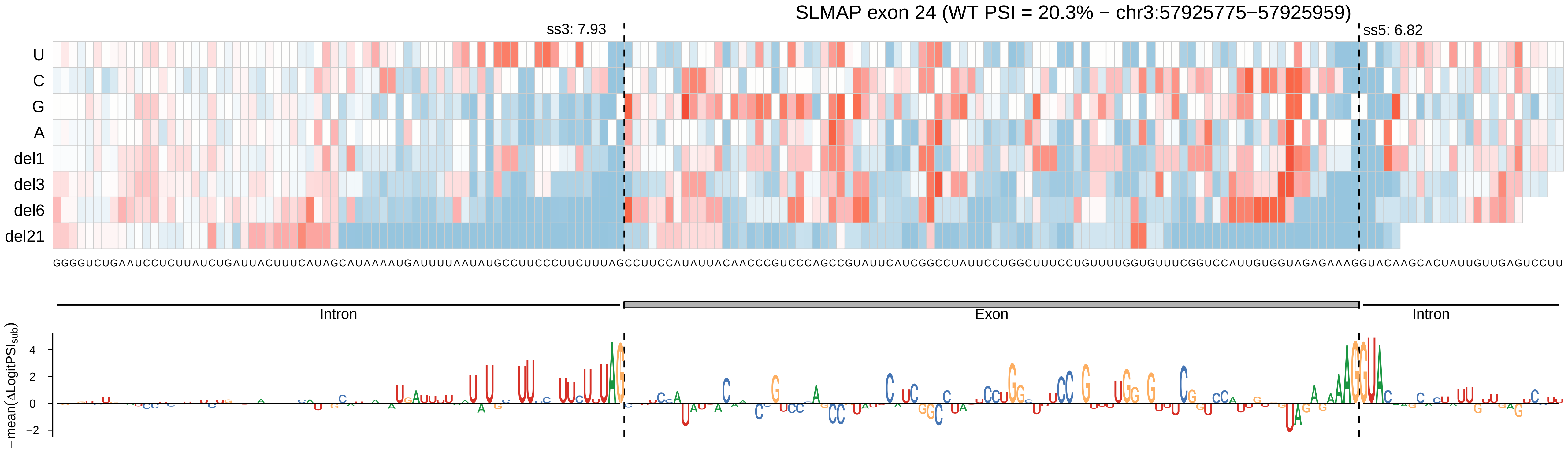
SLMAP exon 24 (WT PSI = 20.3% − chr3:57925775−57925959)
ss3: 7.93
ss5: 6.82
U
C
G
A
del1
del3
del6
del21
GGGGUCUGAAUCCUCUUAUCUGAUUACUUUCAUAGCAUAAAAUGAUUUUAAUAUGCCUUCCCUUCUUUAGCCUUCCAUAUUACAACCCGUCCCAGCCGUAUUCAUCGGCCUAUUCCUGGCUUUCCUGUUUUGGUGUUUCGGUCCAUUGUGGUAGAGAAAGGUACAAGCACUAUUGUUGAGUCCUU
Intron
Exon
Intron
−mean(ΔLogitPSI_sub)
4
2
0
−2

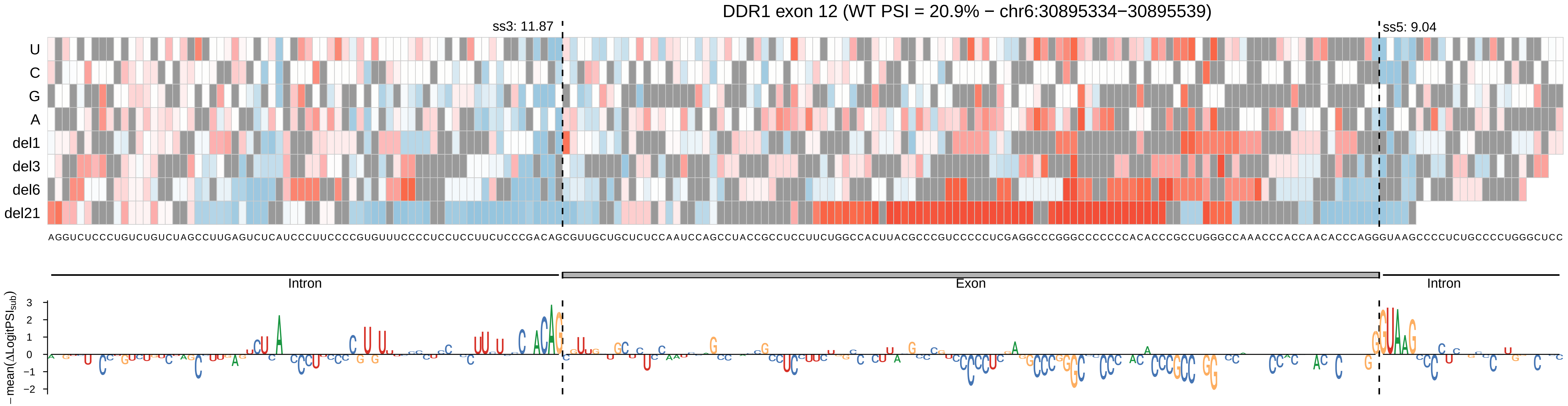
DDR1 exon 12 (WT PSI = 20.9% − chr6:30895334−30895539)
ss3: 11.87
ss5: 9.04
U
C
G
A
del1
del3
del6
del21
Intron
Exon
Intron
−mean(ΔLogitPSI_sub)
3
2
1
0
−1
−2

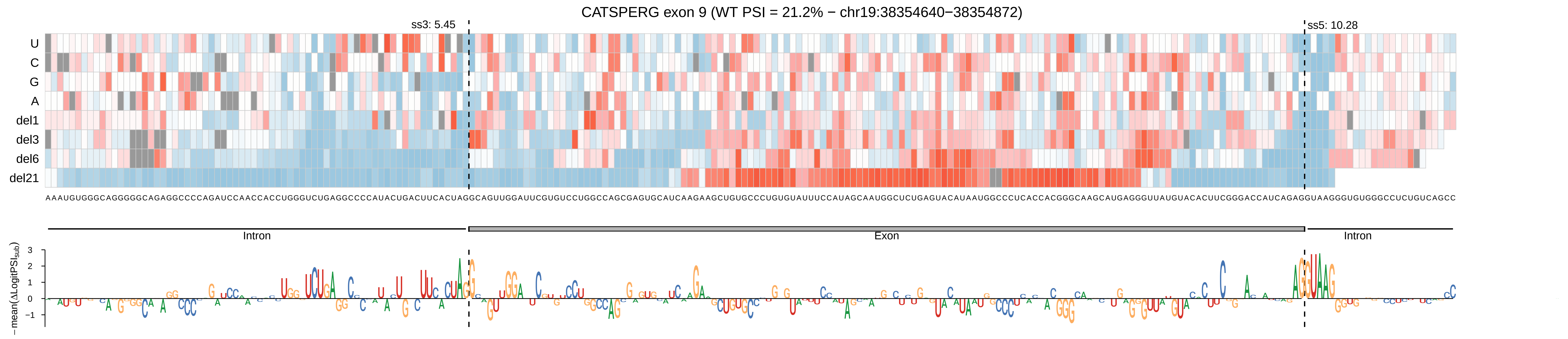

CATSPERG exon 9 (WT PSI = 21.2% − chr19:38354640−38354872)
ss3: 5.45
ss5: 10.28
U
C
G
A
del1
del3
del6
del21
Intron
Exon
Intron
−mean(ΔLogitPSI_sub)
3
2
1
0
−1

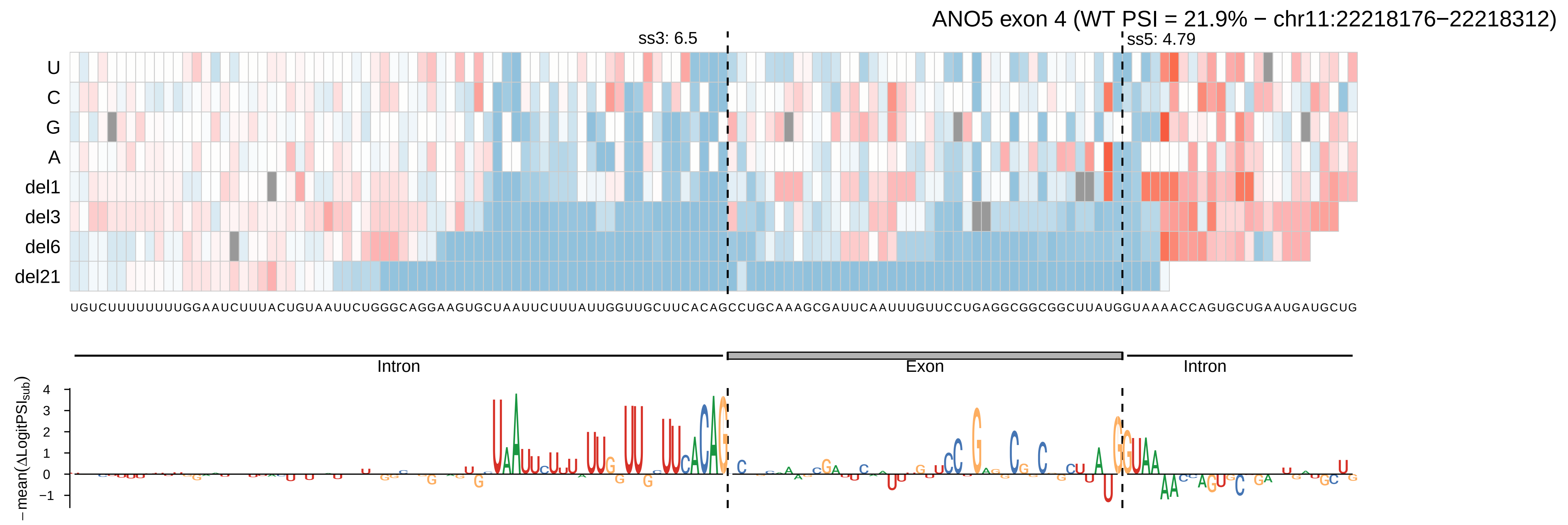
ANO5 exon 4 (WT PSI = 21.9% − chr11:22218176−22218312)
ss3: 6.5
ss5: 4.79
U
C
G
A
del1
del3
del6
del21
UGUCUUUUUUUUGGAAUCUUUACUGUAAUUCUGGGCAGGAAGUGCUAAUUCUUUAUUGGUUGCUUCACAGCCUGCAAAGCGAUUCAAUUUGUUCCUGAGGCGGCGGCUUAUGGUAAAACCAGUGCUGAAUGAUGCUG
Intron
Exon
Intron
−mean(ΔLogitPSI_sub)
4
3
2
1
0
−1

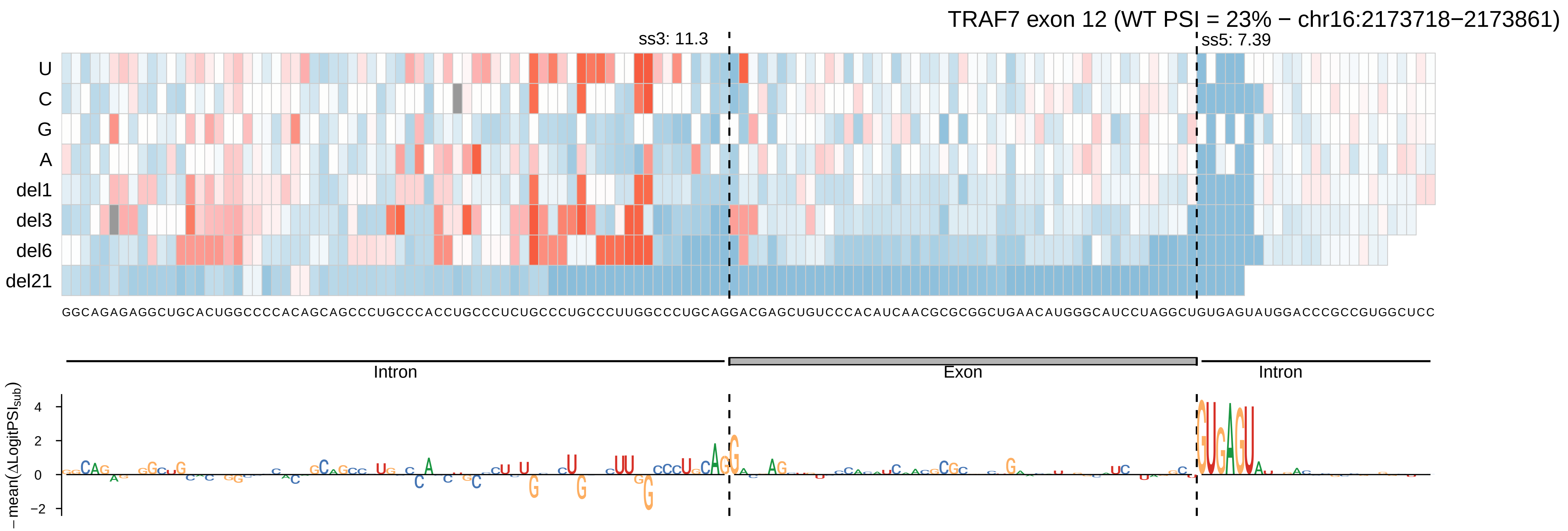
TRAF7 exon 12 (WT PSI = 23% − chr16:2173718−2173861)
ss3: 11.3
ss5: 7.39
U
C
G
A
del1
del3
del6
del21
GGCAGAGAGGCUGCACUGGCCCCACAGCAGCCCUGCCCACCUGCCCUCUGCCCUGCCCUUGGCCCUGCAGGACGAGCUGUCCCACAUCAACGCGCGGCUGAACAUGGGCAUCCUAGGCUGUGAGUAUGGACCCGCCGUGGCUCC
Intron
Exon
Intron
−mean(ΔLogitPSI_sub)
4
2
0
−2

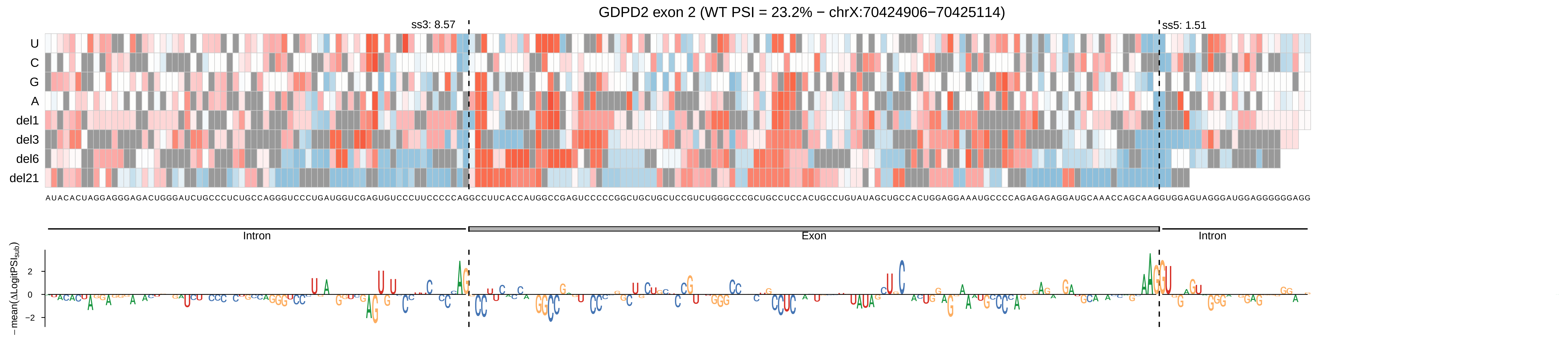

GDPD2 exon 2 (WT PSI = 23.2% − chrX:70424906−70425114)
ss3: 8.57
ss5: 1.51
U
C
G
A
del1
del3
del6
del21
AUACACUAGGAGGGAGACUGGGAUCUGCCCUCUGCCAGGGUCCCUGAUGGUCGAGUGUCCCUUCCCCCAGGCCUUCACCAUGGCCGAGUCCCCCGGCUGCUGCUCCGUCUGGGCCCGCUGCCUCCACUGCCUGUAUAGCUGCCACUGGAGGAAAUGCCCCAGAGAGAGGAUGCAAACCAGCAAGGUGGAGUAGGGAUGGAGGGGGGAGG
Intron
Exon
Intron
−mean(ΔLogitPSI_sub)
2
0
−2

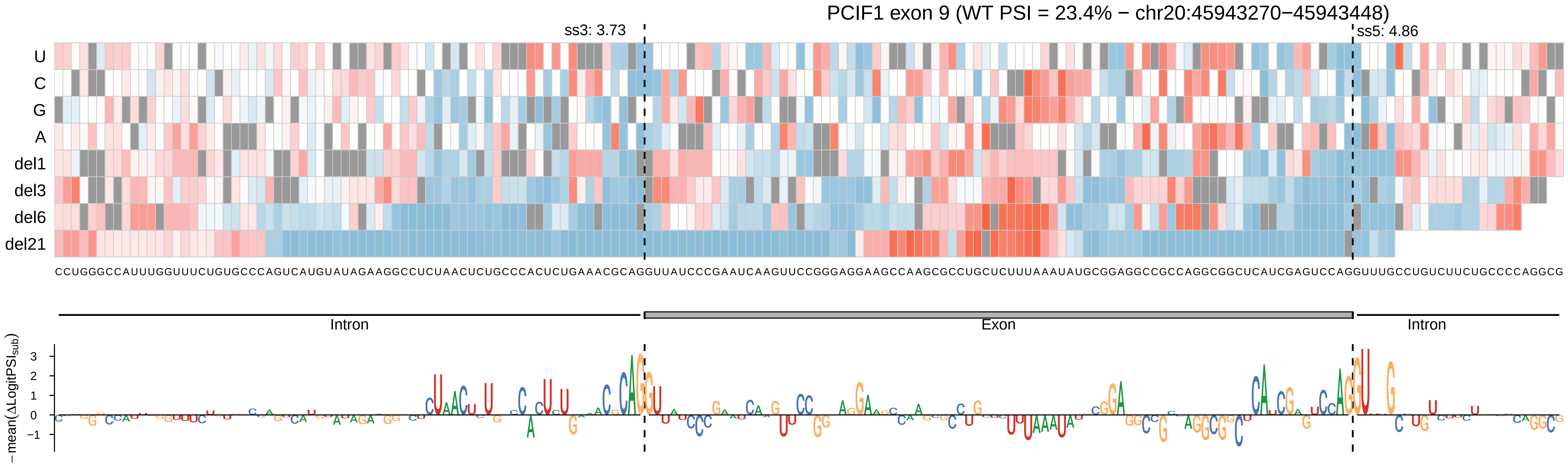

PCIF1 exon 9 (WT PSI = 23.4% − chr20:45943270−45943448)
ss3: 3.73
ss5: 4.86
U
C
G
A
del1
del3
del6
del21
Intron
Exon
Intron
−mean(ΔLogitPSI_sub)
3
2
1
0
−1

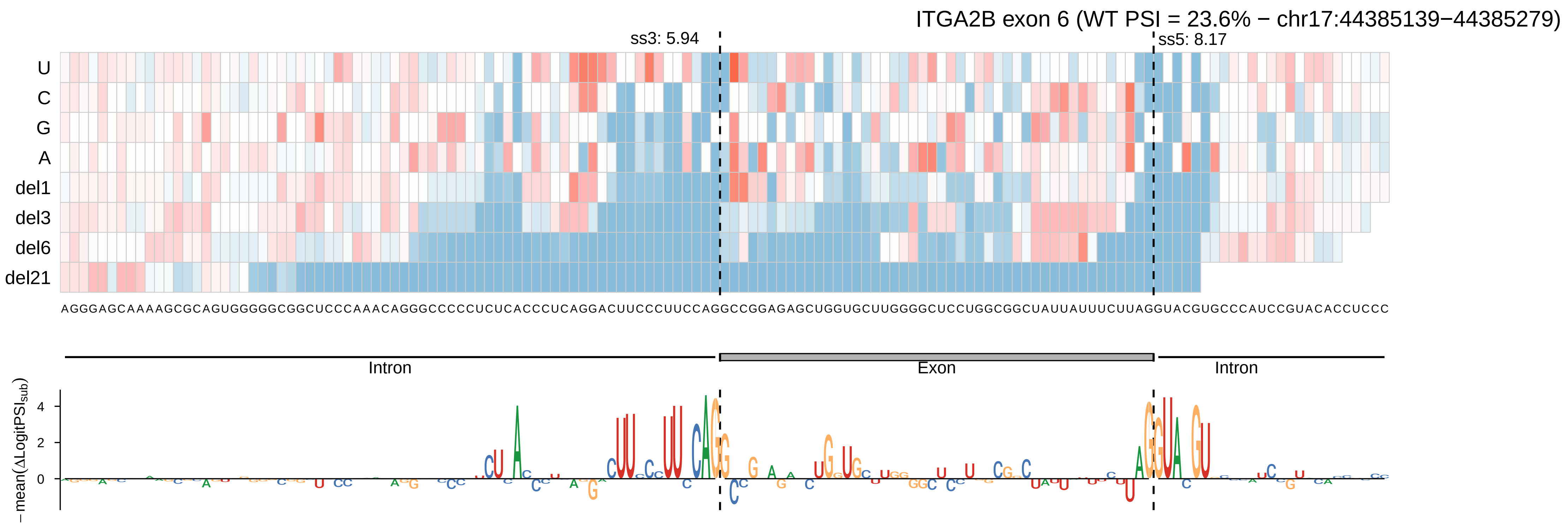
ITGA2B exon 6 (WT PSI = 23.6% − chr17:44385139−44385279)
ss3: 5.94
ss5: 8.17
U
C
G
A
del1
del3
del6
del21
AGGGAGCAAAAGCGCAGUGGGGGCGGCUCCCAAACAGGGCCCCCUCUCACCCUCAGGACUUCCCUUCCAGGCCGGAGAGCUGGUGCUUGGGGCUCCUGGCGGCUAUUAUUUCUUAGGUACGUGCCCAUCCGUACACCUCCC
Intron
Exon
Intron
−mean(ΔLogitPSI_sub)
4
2
0

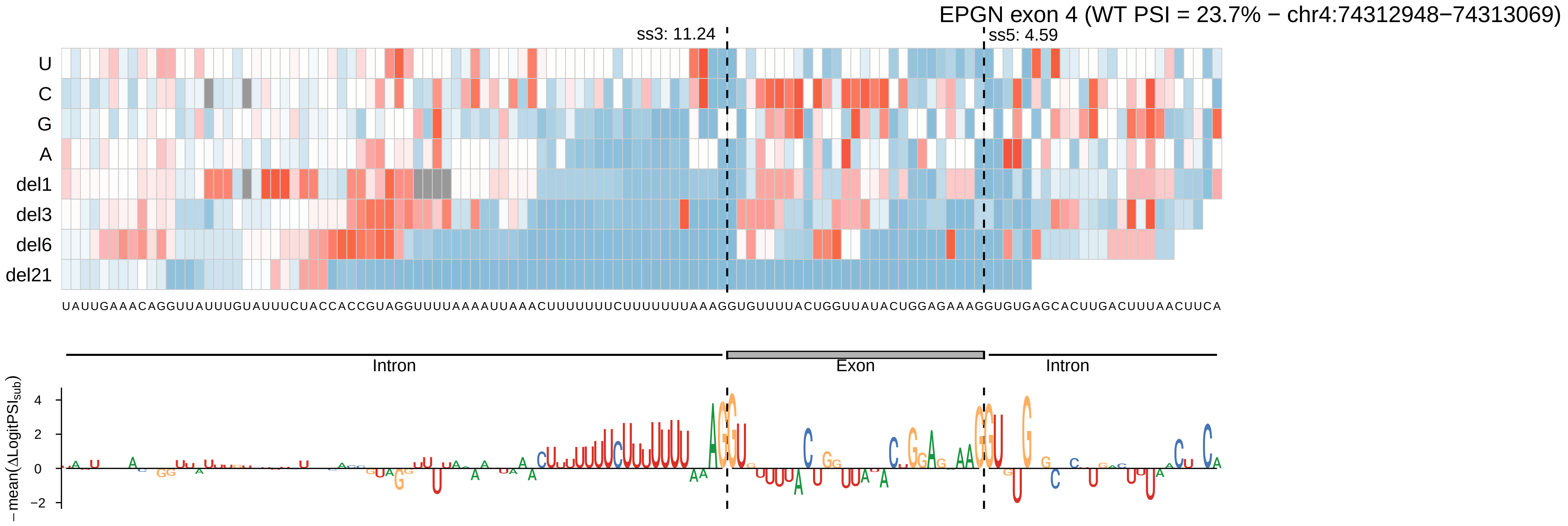

EPGN exon 4 (WT PSI = 23.7% − chr4:74312948−74313069)
ss3: 11.24
ss5: 4.59
U
C
G
A
del1
del3
del6
del21
UAUUGAAACAGGUUAUUUGUAUUUCUACCACCGUAGGUUUUAAAAUUAAACUUUUUUUCUUUUUUUAAAGGUGUUUUACUGGUUAUACUGGAGAAAGGUGUGAGCACUUGACUUUAACUUCA
Intron
Exon
Intron
− mean(ΔLogitPSI_sub)
4
2
0
−2

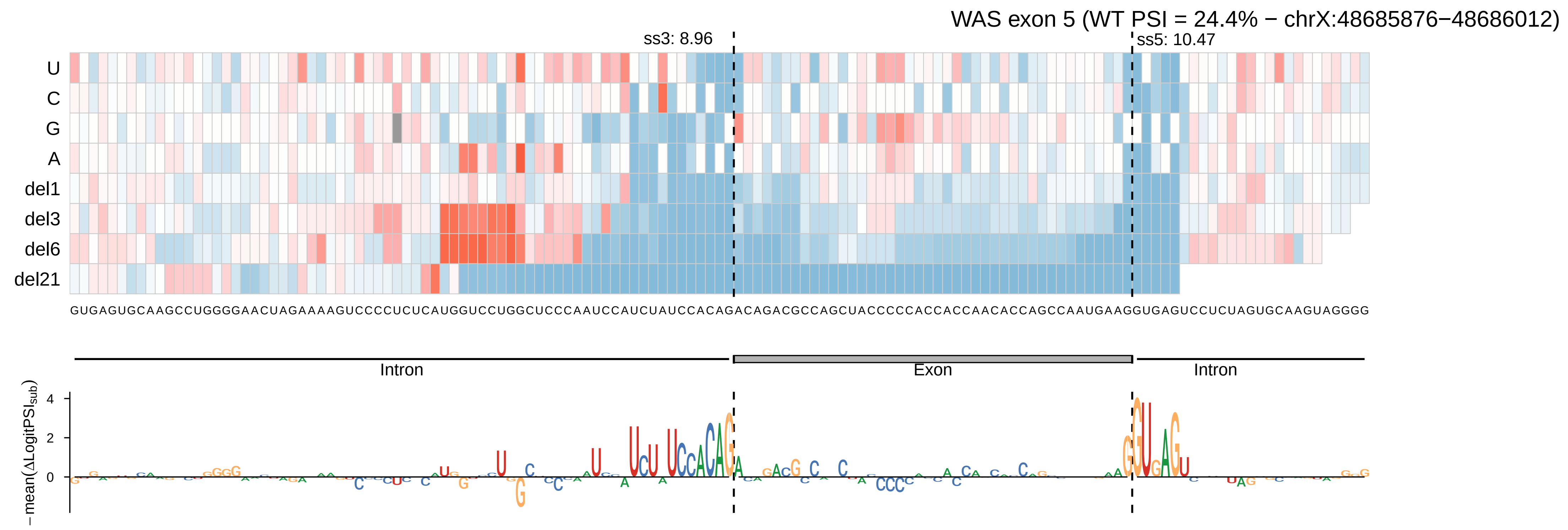
WAS exon 5 (WT PSI = 24.4% − chrX:48685876−48686012)
ss3: 8.96
ss5: 10.47
U
C
G
A
del1
del3
del6
del21
Intron
Exon
Intron
−mean(ΔLogitPSI_sub)
4
2
0

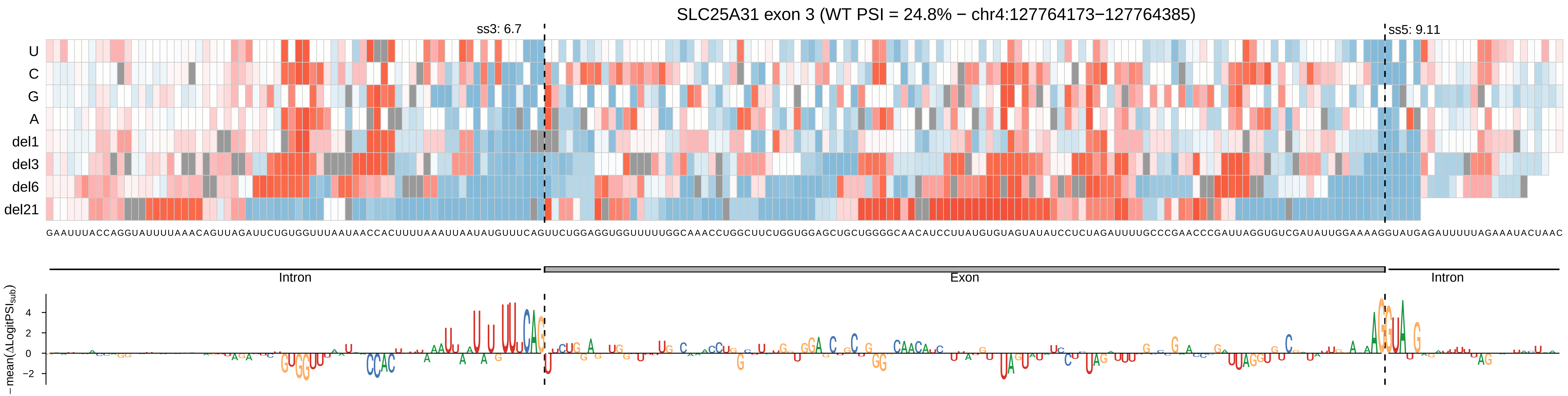

SLC25A31 exon 3 (WT PSI = 24.8% − chr4:127764173−127764385)
ss3: 6.7
ss5: 9.11
U
C
G
A
del1
del3
del6
del21
Intron
Exon
Intron
−mean(ΔLogitPSI$_{sub}$)
4
2
0
−2

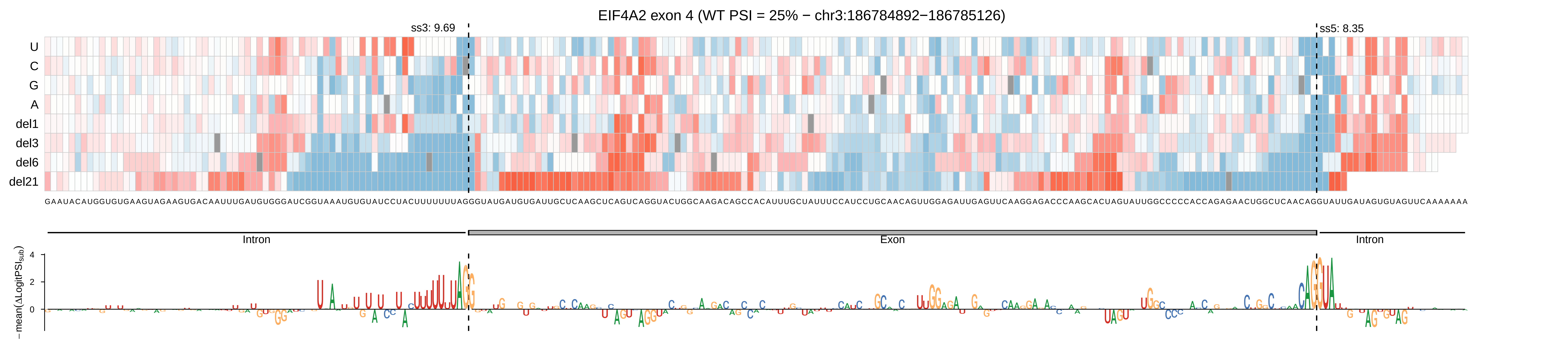

EIF4A2 exon 4 (WT PSI = 25% − chr3:186784892−186785126)
ss3: 9.69
ss5: 8.35
U
C
G
A
del1
del3
del6
del21
Intron
Exon
Intron
−mean(ΔLogitPSI_sub)
4
2
0

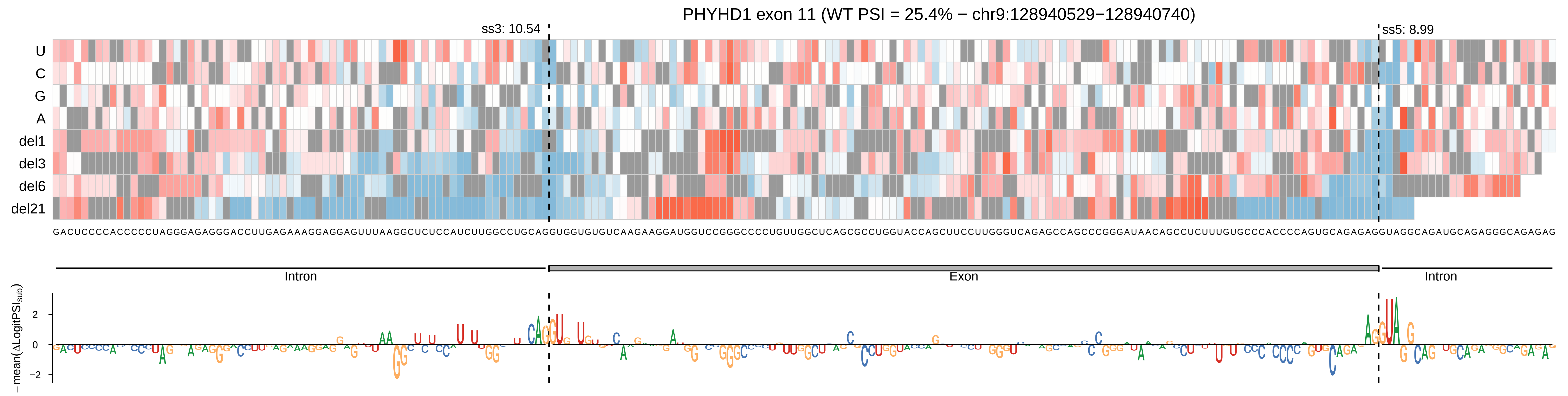
PHYHD1 exon 11 (WT PSI = 25.4% − chr9:128940529−128940740)
ss3: 10.54
ss5: 8.99
U
C
G
A
del1
del3
del6
del21
GACUCCCCACCCCCUAGGGAGAGGGACCUUGAGAAAGGAGGAGUUUAAGGCUCUCCAUCUUGGCCUGCAGGUGGUGUGUCAAGAAGGAUGGUCCGGGCCCCUGUUGGCUCAGCGCCUGGUACCAGCUUCCUUGGGUCAGAGCCAGCCCGGGAUAACAGCCUCUUUGUGCCCACCCCAGUGCAGAGAGGUAGGCAGAUGCAGAGGGCAGAGAG
Intron
Exon
Intron
−mean(ΔLogitPSI_sub)
2
0
−2

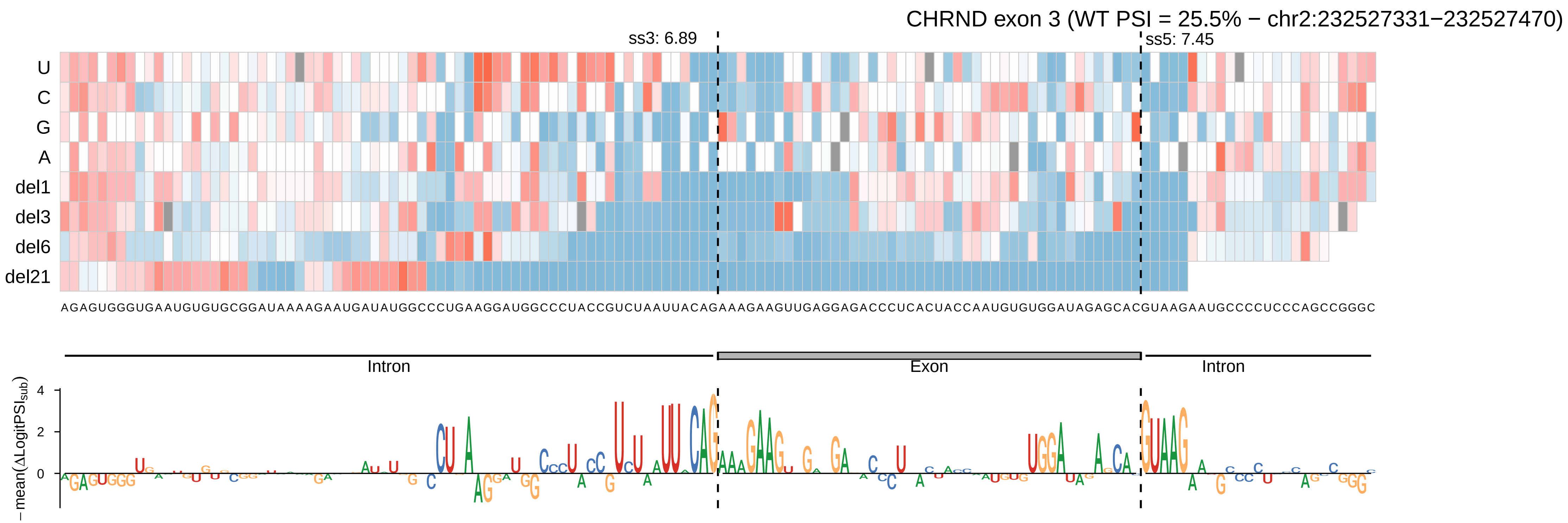
CHRND exon 3 (WT PSI = 25.5% − chr2:232527331−232527470)
ss3: 6.89
ss5: 7.45
U
C
G
A
del1
del3
del6
del21
AGAGUGGGUGAAUGUGUGCGGAUAAAAGAAUGAUAUGGCCCUGAAGGAUGGCCCUACCGUCUAAUUACAGAAAGAAGUUGAGGAGACCCUCACUACCAAUGUGUGGAUAGAGCACGUAAGAAUGCCCCUCCCAGCCGGGC
Intron
Exon
Intron
−mean(ΔLogitPSI_sub)
4
2
0

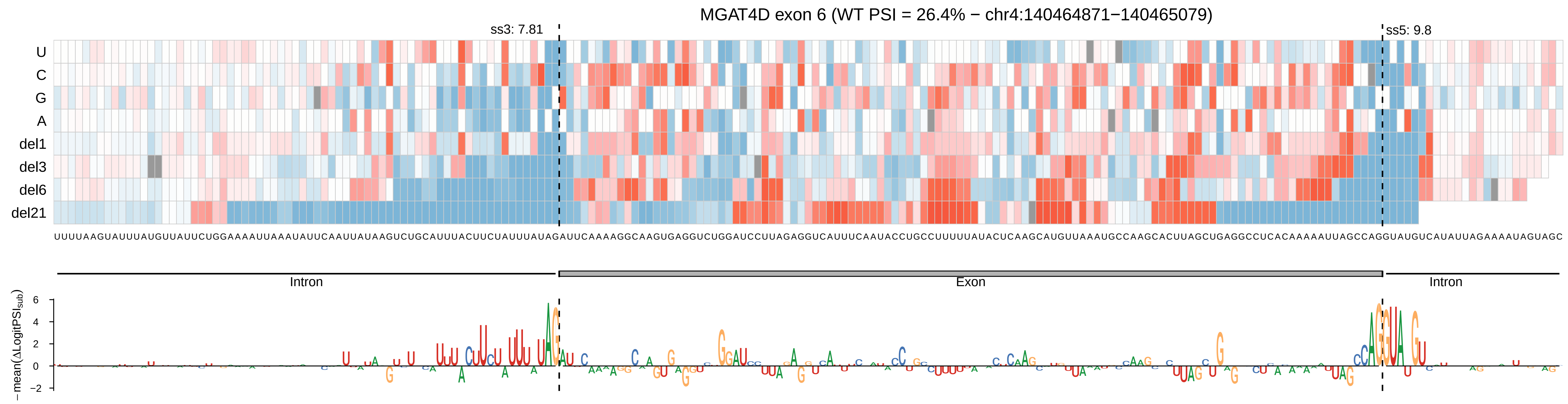
MGAT4D exon 6 (WT PSI = 26.4% − chr4:140464871−140465079)
ss3: 7.81
ss5: 9.8
U
C
G
A
del1
del3
del6
del21
UUUUAAGUAUUUAUGUUAUUCUGGAAAAUUAAAUAUUCAAUUAUAAGUCUGCAUUUACUUCUAUUUAUAGAUUCAAAAGGCAAGUGAGGUCUGGAUCCUUAGAGGUCAUUUCAAUACCUGCCUUUUUAUACUCAAGCAUGUUAAAUGCCAAGCACUUAGCUGAGGCCUCACAAAAAUUAGCCAGGUAUGUCAUAUUAGAAAAUAGUAGC
Intron
Exon
Intron
−mean(ΔLogitPSI_sub)
6
4
2
0
−2

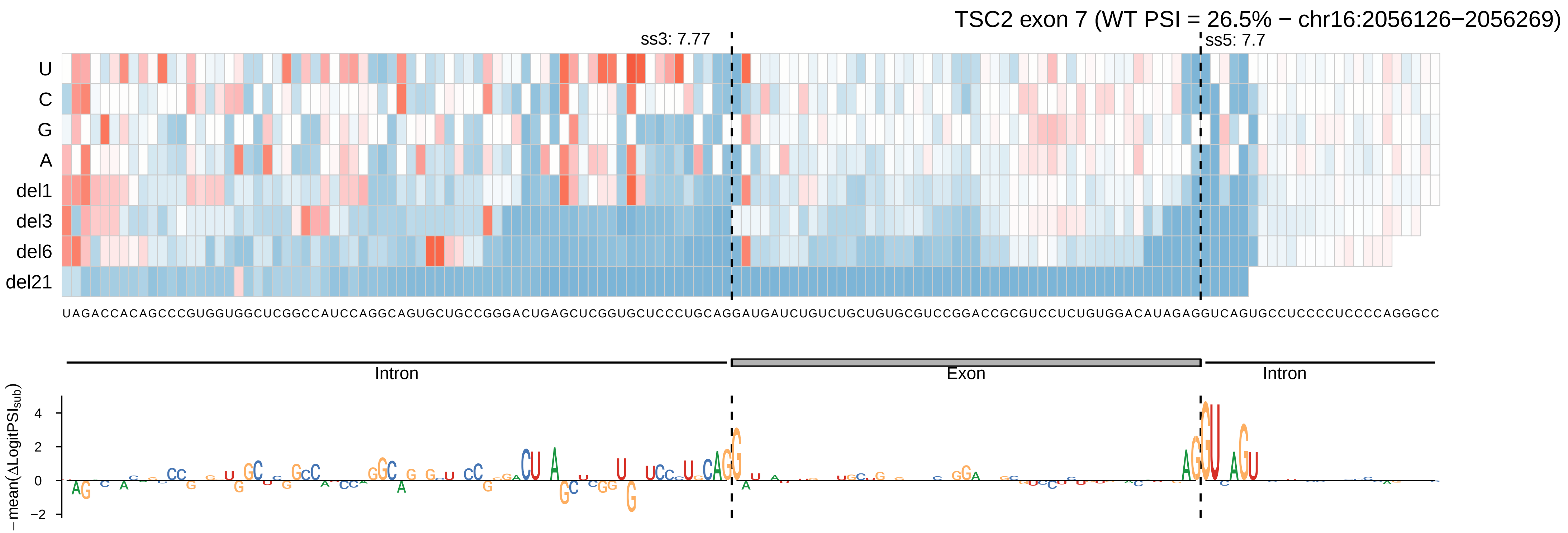

TSC2 exon 7 (WT PSI = 26.5% − chr16:2056126−2056269)
ss3: 7.77
ss5: 7.7
U
C
G
A
del1
del3
del6
del21
UAGACCACAGCCCGUGGUGGCUCGGCCAUCCAGGCAGUGCUGCCGGGACUGAGCUCGGUGCUCCCUGCAGGAUGAUCUGUCUGCUGUGCGUCCGGACCGCGUCCUCUGUGGACAUAGAGGUCAGUGCCUCCCCUCCCCAGGGCC
Intron
Exon
Intron
−mean(ΔLogitPSI_sub)
4
2
0
−2

PTS exon 3 (WT PSI = 26.6% − chr11:112230138−112230255)

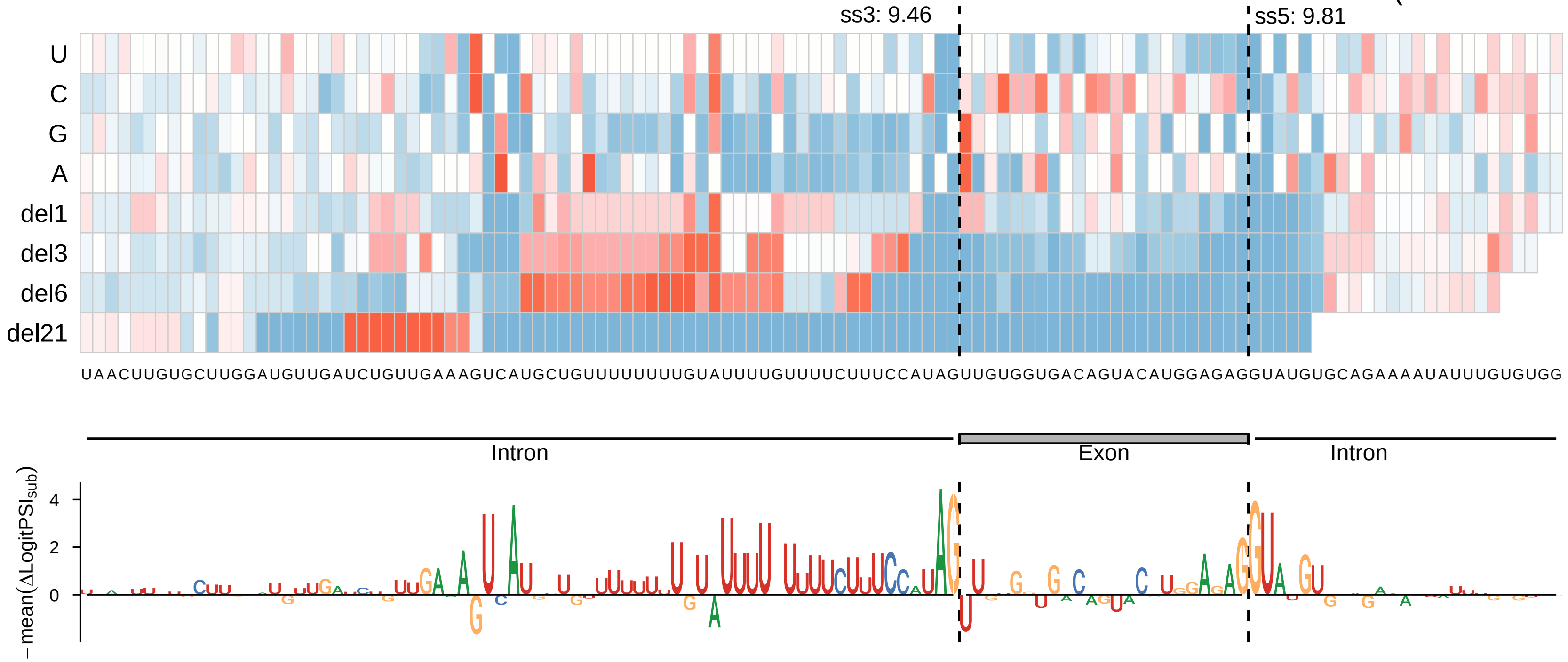

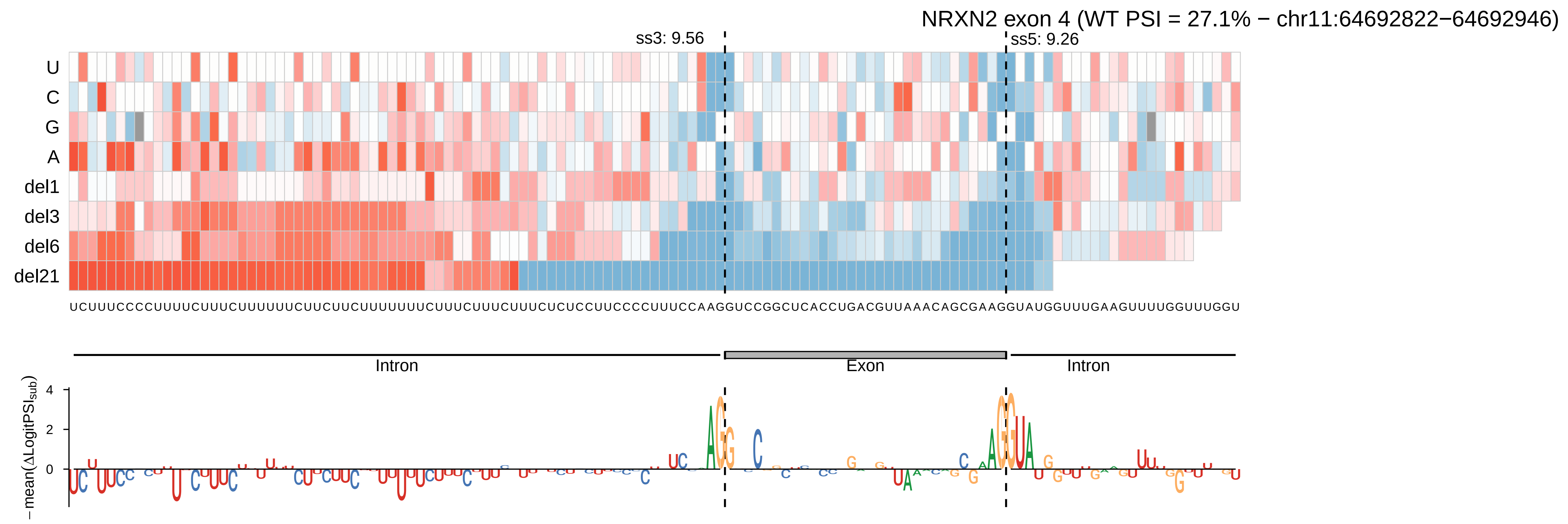

NRXN2 exon 4 (WT PSI = 27.1% − chr11:64692822−64692946)
ss3: 9.56
ss5: 9.26
U
C
G
A
del1
del3
del6
del21
UCUUUCCCCUUUUCUUUCUUUUUUCUUCUUCUUUUUUUCUUUCUUUCUUUCUCUCCUUCCCCUUUCCAAGGUCCGGCUCACCUGACGUUAAACAGCGAAGGUAUGGUUUGAAGUUUUGGUUUGGU
Intron
Exon
Intron
4
2
0
− mean(ΔLogitPSI_sub)

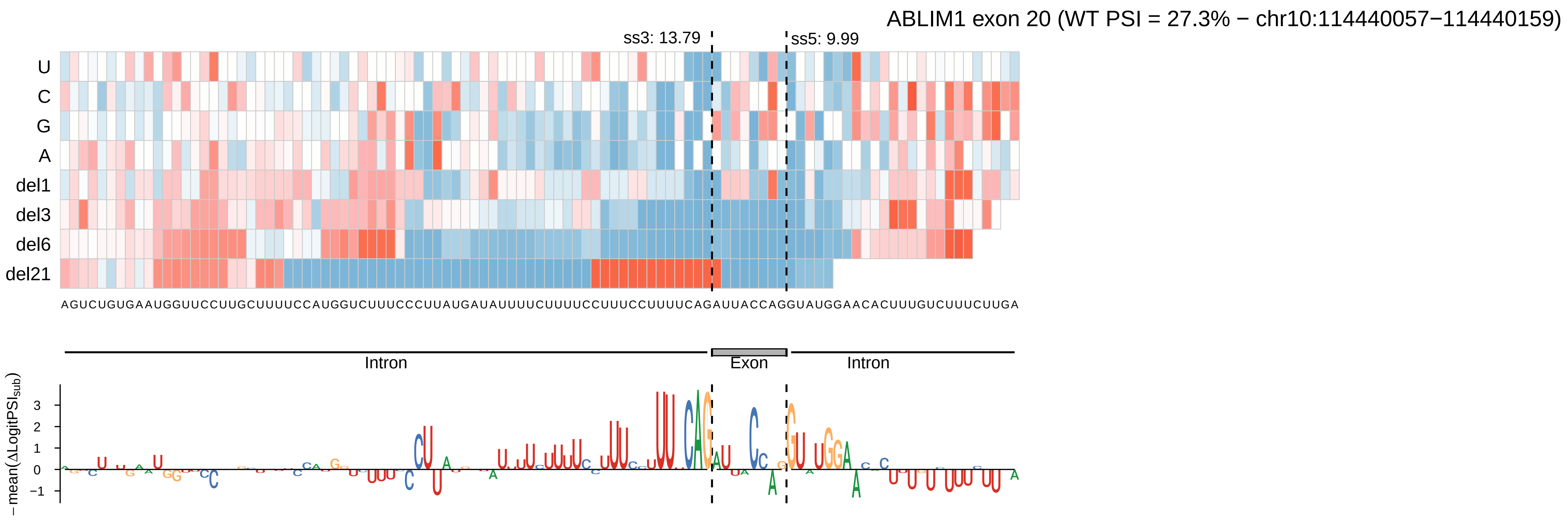
ABLIM1 exon 20 (WT PSI = 27.3% − chr10:114440057−114440159)
ss3: 13.79
ss5: 9.99
U
C
G
A
del1
del3
del6
del21
AGUCUGUGAAUGGUUCCUUGCUUUUCCAUGGUCUUUCCCUUAUGAUAUUUUCUUUUCCUUUCCUUUUCAGAUUACCAGGUAUGGAACACUUUGUCUUUCUUGA
Intron
Exon
Intron
−mean(ΔLogitPSI_sub)
3
2
1
0
−1

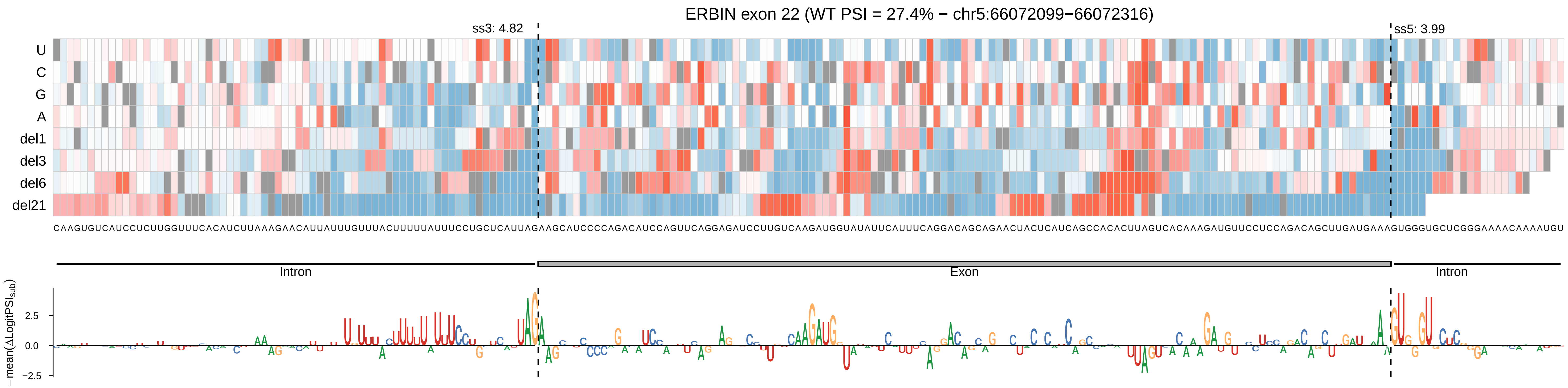

ERBIN exon 22 (WT PSI = 27.4% − chr5:66072099−66072316)
ss3: 4.82
ss5: 3.99
U
C
G
A
del1
del3
del6
del21
CAAGUGUCAUCCUCUUGGUUUCACAUCUUAAAGAACAUUAUUUGUUUACUUUUUAUUUCCUGCUCAUUAGAAGCAUCCCCAGACAUCCAGUUCAGGAGAUCCUUGUCAAGAUGGUAUAUUCAUUUCAGGACAGCAGAACUACUCAUCAGCCACACUUAGUCACAAAGAUGUUCCUCCAGACAGCUUGAUGAAAGUGGGUGCUCGGGAAAACAAAAUGU
Intron
Exon
Intron
−mean(ΔLogitPSI_sub)
2.5
0.0
−2.5

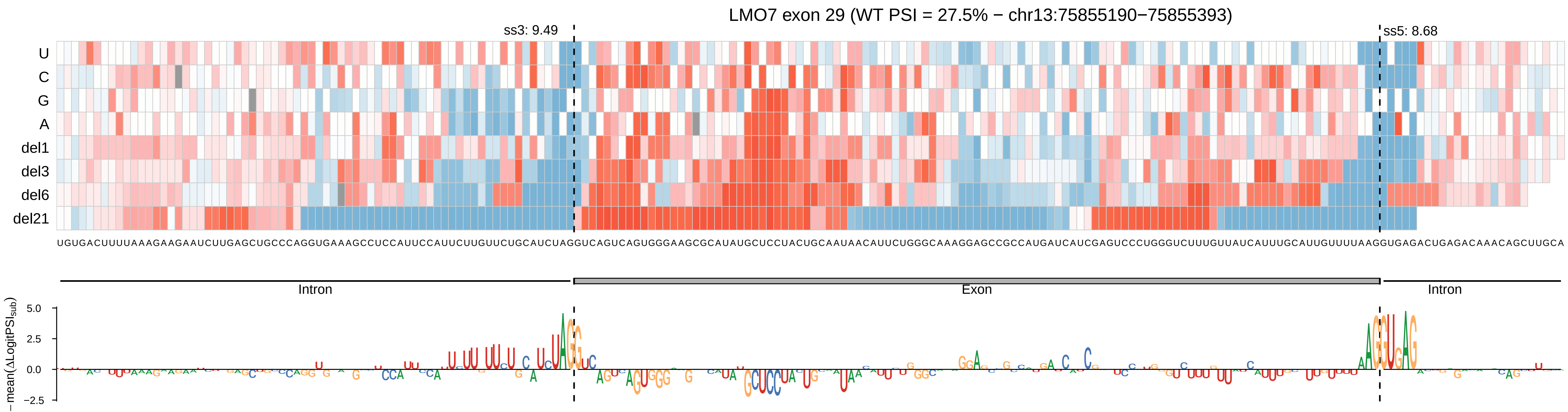

LMO7 exon 29 (WT PSI = 27.5% − chr13:75855190−75855393)
ss3: 9.49
ss5: 8.68
U
C
G
A
del1
del3
del6
del21
UGUGACUUUUAAAGAAGAAUCUUGAGCUGCCCAGGUGAAAGCCUCCAUUCCAUUCUUGUUCUGCAUCUAGGUCAGUCAGUGGGAAGCGCAUAUGCUCCUACUGCAAUAACAUUCUGGGCAAAGGAGCCGCCAUGAUCAUCGAGUCCCUGGGUCUUUGUUAUCAUUUGCAUUGUUUUAAGGUGAGACUGAGACAAACAGCUUGCA
Intron
Exon
Intron
− mean(ΔLogitPSI_Sub)
5.0
2.5
0.0
−2.5

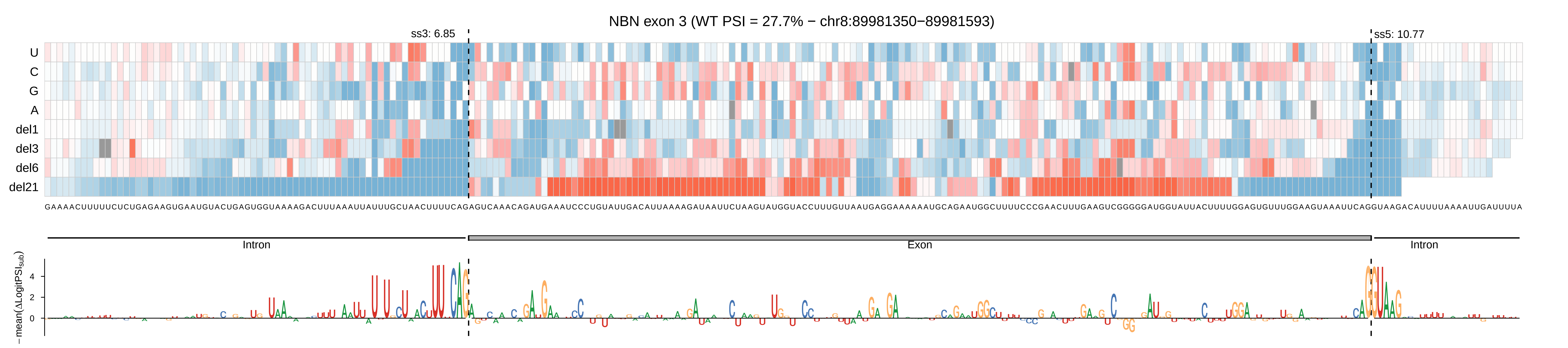

NBN exon 3 (WT PSI = 27.7% − chr8:89981350−89981593)
ss3: 6.85
ss5: 10.77
U
C
G
A
del1
del3
del6
del21
Intron
Exon
Intron
−mean(ΔLogitPSI_sub)
4
2
0

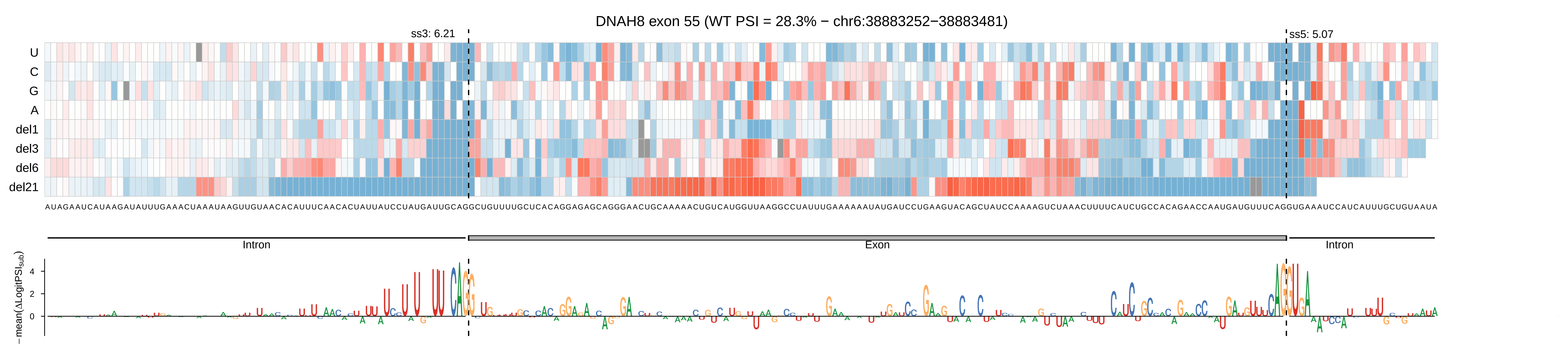
DNAH8 exon 55 (WT PSI = 28.3% − chr6:38883252−38883481)
ss3: 6.21
ss5: 5.07
U
C
G
A
del1
del3
del6
del21
Intron
Exon
Intron
−mean(ΔLogitPSI_sub)
4
2
0

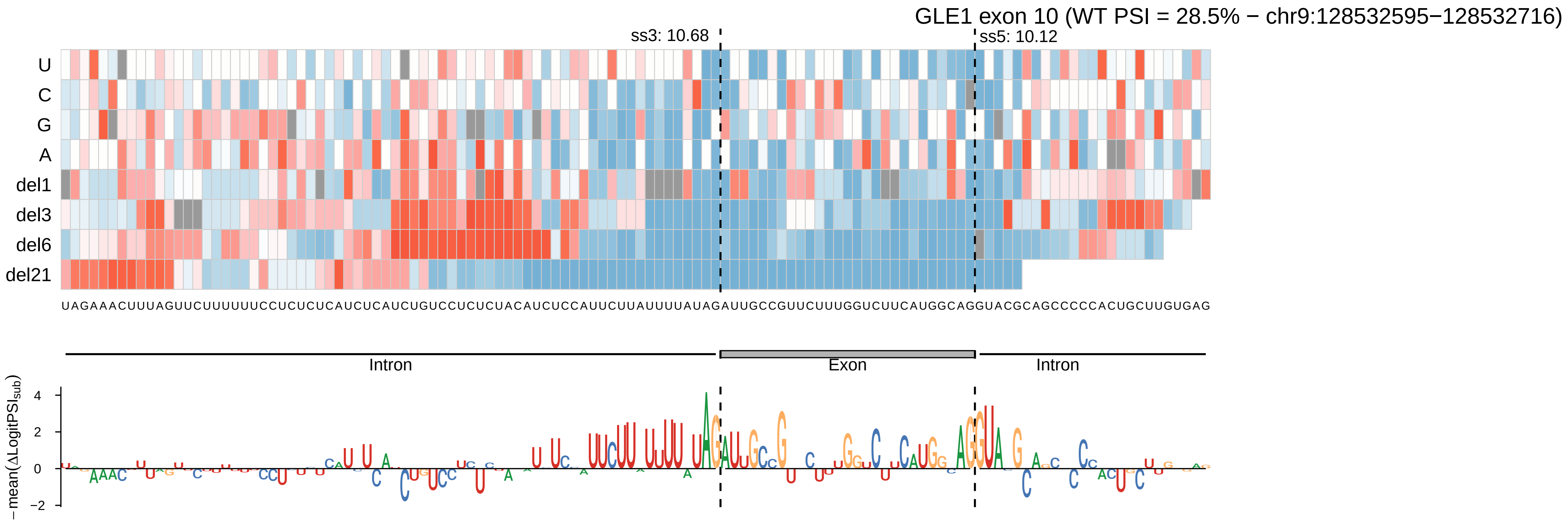
GLE1 exon 10 (WT PSI = 28.5% − chr9:128532595−128532716)
ss3: 10.68
ss5: 10.12
U
C
G
A
del1
del3
del6
del21
UAGAAACUUUAGUUCUUUUUUCCUCUCUCAUCUCAUCUGUCCUCUCUACAUCUCCAUUCUUAUUUUAUAGAUUGCCGUUCUUUGGUCUUCAUGGCAGGUACGCAGCCCCCACUGCUUGUGAG
Intron
Exon
Intron
− mean(ΔLogitPSI_sub)
4
2
0
−2

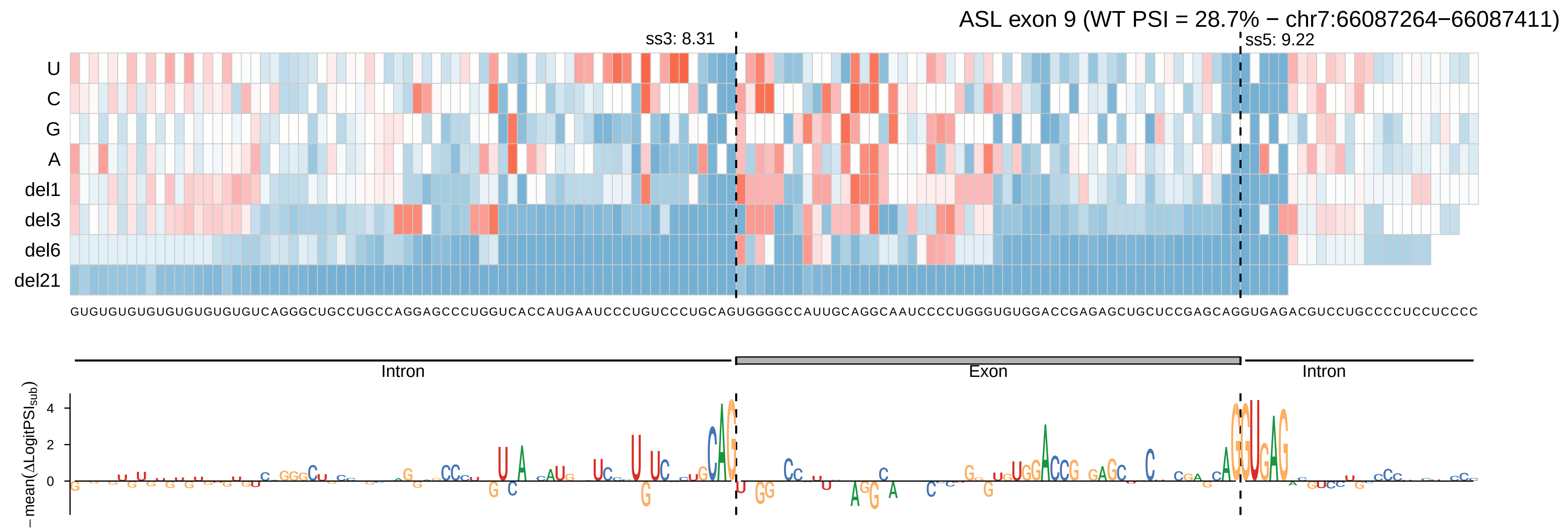

ASL exon 9 (WT PSI = 28.7% − chr7:66087264−66087411)
ss3: 8.31
ss5: 9.22
U
C
G
A
del1
del3
del6
del21
GUGUGUGUGUGUGUGUGUGUCAGGGCUGCCUGCCAGGAGCCCUGGUCACCAUGAAUCCCUGUCCCUGCAGUGGGGCCAUUGCAGGCAAUCCCCUGGGUGUGGACCGAGAGCUGCUCCGAGCAGGUGAGACGUCCUGCCCCUCCUCCCC
Intron
Exon
Intron
−mean(ΔLogitPSI_sub)
4
2
0

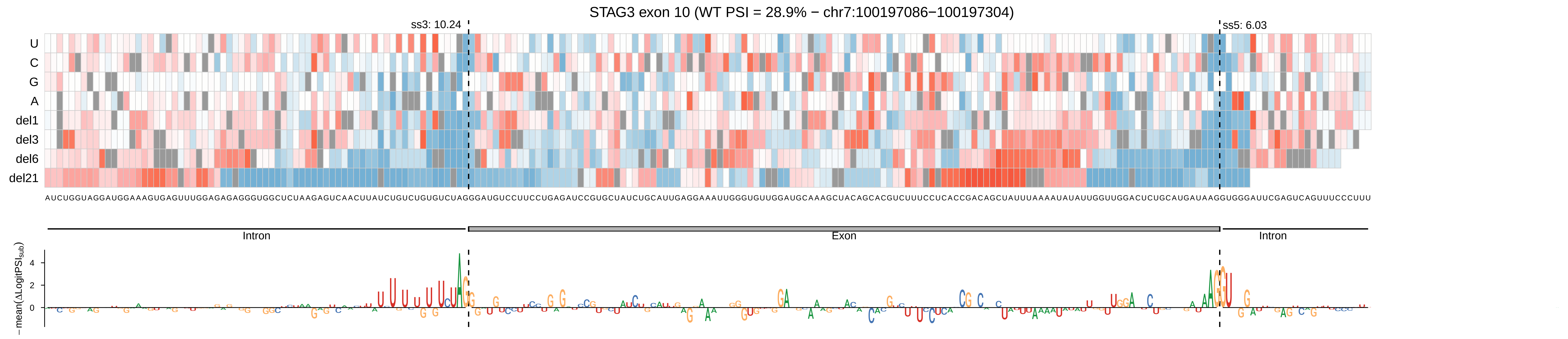
STAG3 exon 10 (WT PSI = 28.9% − chr7:100197086−100197304)
ss3: 10.24
ss5: 6.03
U
C
G
A
del1
del3
del6
del21
AUCUGGUAGGAUGGAAAGUGAGUUUGGAGAGAGGGUGGCUCUAAGAGUCAACUUAUCUGUCUGUGUCUAGGGAUGUCCUUCCUGAGAUCCGUGCUAUCUGCAUUGAGGAAAUUGGGUGUUGGAUGCAAAGCUACAGCACGUCUUUCCUCACCGACAGCUAUUUAAAAUAUAUUGGUUGGACUCUGCAUGAUAAGGUGGGAUUCGAGUCAGUUUCCCUUU
Intron
Exon
Intron
−mean(ΔLogitPSIsub)
4
2
0

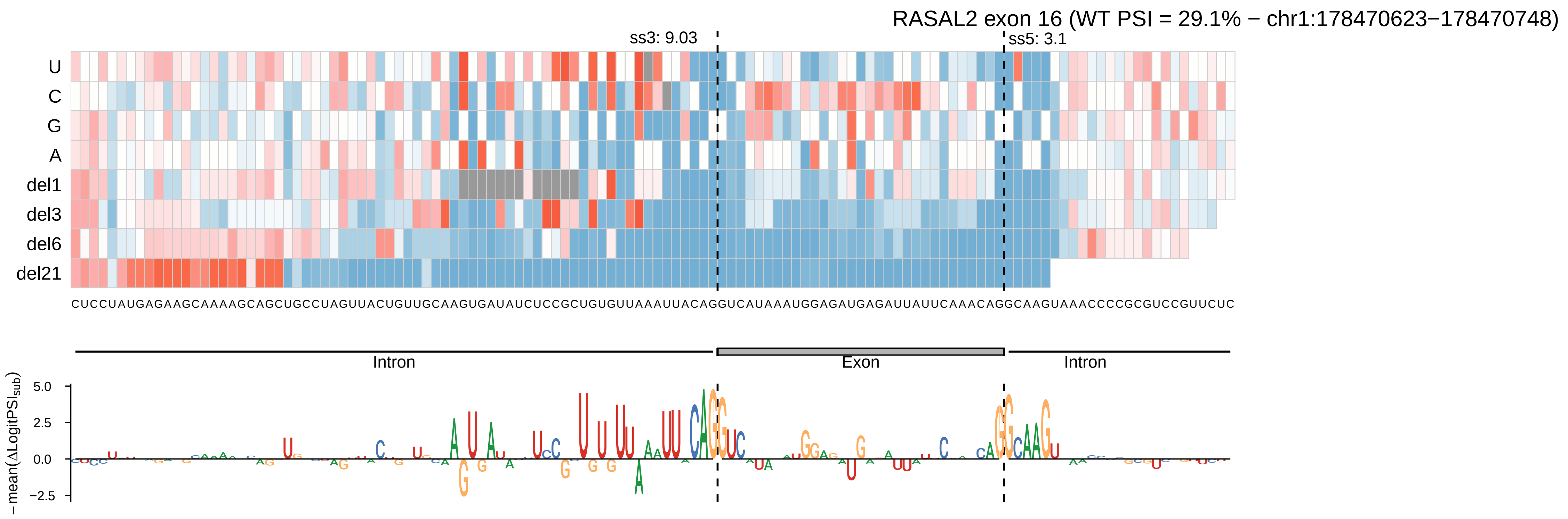

RASAL2 exon 16 (WT PSI = 29.1% − chr1:178470623−178470748)
ss3: 9.03
ss5: 3.1
U
C
G
A
del1
del3
del6
del21
CUCCUAUGAGAAGCAAAAGCAGCUGCCUAGUUACUGUUGCAAGUGAUAUCUCCGCUGUGUUAAAUUACAGGUCAUAAAUGGAGAUGAGAUUAUUCAAACAGGCAAGUAAACCCCGCGUCCGUUCUC
Intron
Exon
Intron
− mean(ΔLogitPSI$_{sub}$)
5.0
2.5
0.0
−2.5

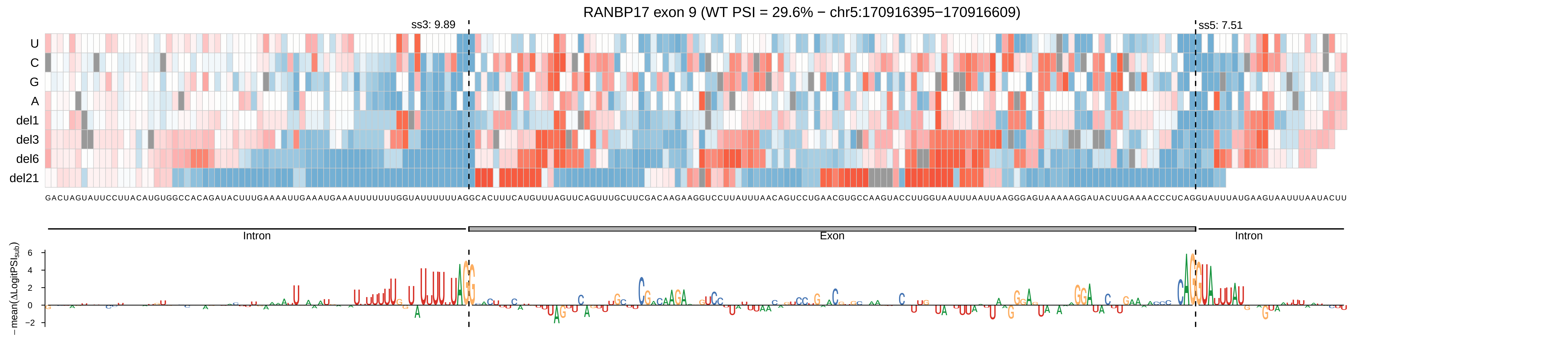

RANBP17 exon 9 (WT PSI = 29.6% − chr5:170916395−170916609)
ss3: 9.89
ss5: 7.51
U
C
G
A
del1
del3
del6
del21
Intron
Exon
Intron
− mean(ΔLogitPSIsub)
6
4
2
0
−2

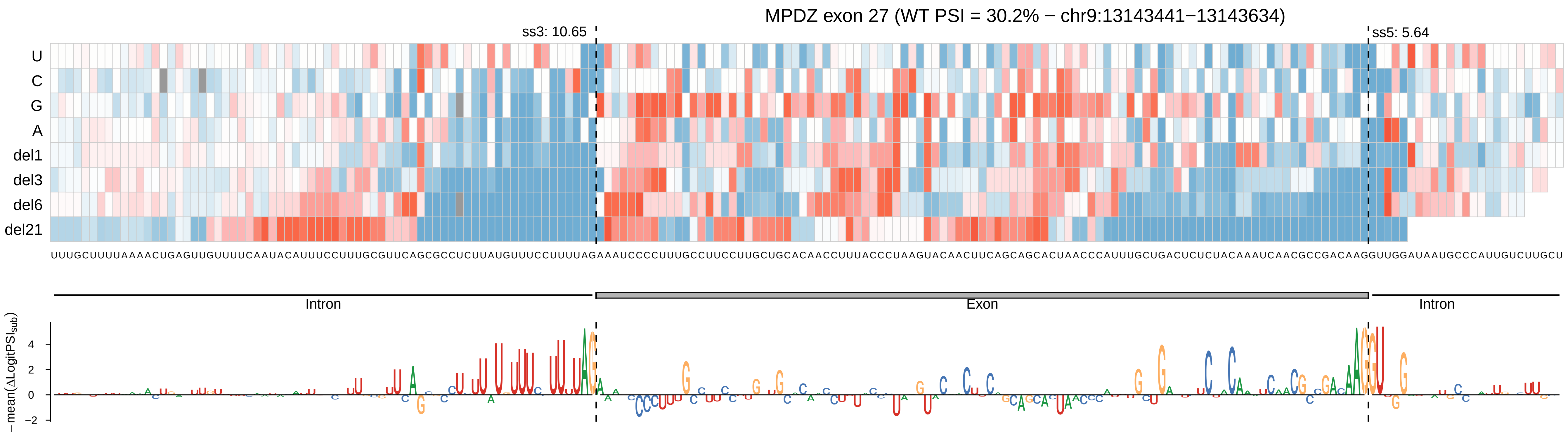

MPDZ exon 27 (WT PSI = 30.2% − chr9:13143441−13143634)
ss3: 10.65
ss5: 5.64
U
C
G
A
del1
del3
del6
del21
UUUGCUUUUAAAACUGAGUUGUUUUCAAUACAUUUCCUUUGCGUUCAGCGCCUCUUAUGUUUCCUUUUAGAAAUCCCCUUUGCCUUCCUUGCUGCACAACCUUUACCCUAAGUACAACUUCAGCAGCACUAACCCAUUUGCUGACUCUCUACAAAUCAACGCCGACAAGGUUGGAUAAUGCCCAUUGUCUUGCU
Intron
Exon
Intron
− mean(ΔLogitPSI_sub)
4
2
0
−2

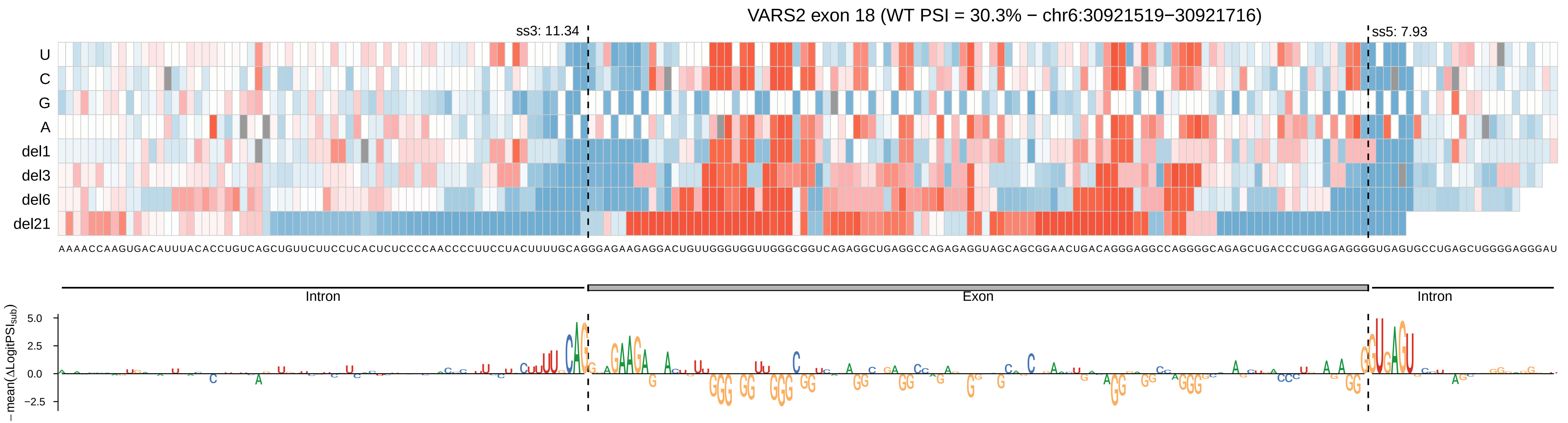

VARS2 exon 18 (WT PSI = 30.3% − chr6:30921519−30921716)
ss3: 11.34
ss5: 7.93
U
C
G
A
del1
del3
del6
del21
AAAACCAAGUGACAUUUACACCUGUCAGCUGUUCUUCCUCACUCUCCCCAACCCCUUCCUACUUUUGCAGGGAGAAGAGGACUGUUGGGUGGUUGGGCGGUCAGAGGCUGAGGCCAGAGAGGUAGCAGCGGAACUGACAGGGAGGCCAGGGGCAGAGCUGACCCUGGAGAGGGGUGAGUGCCUGAGCUGGGGAGGGAU
Intron
Exon
Intron
−mean(ΔLogitPSI_sub)
5.0
2.5
0.0
−2.5

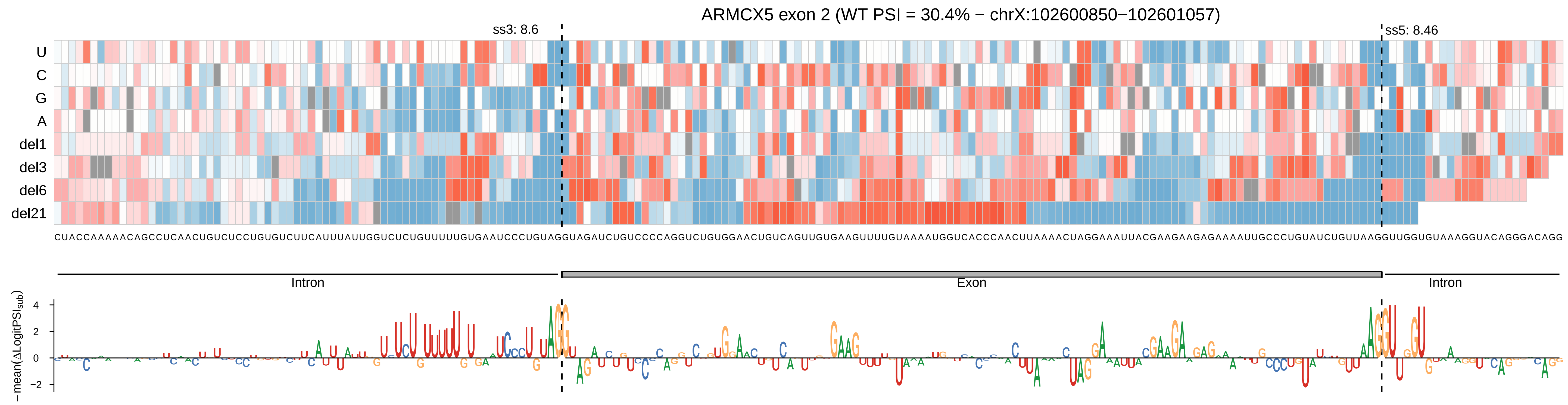
ARMCX5 exon 2 (WT PSI = 30.4% − chrX:102600850−102601057)
ss3: 8.6
ss5: 8.46
U
C
G
A
del1
del3
del6
del21
Intron
Exon
Intron
−mean(ΔLogitPSIsub)
4
2
0
−2

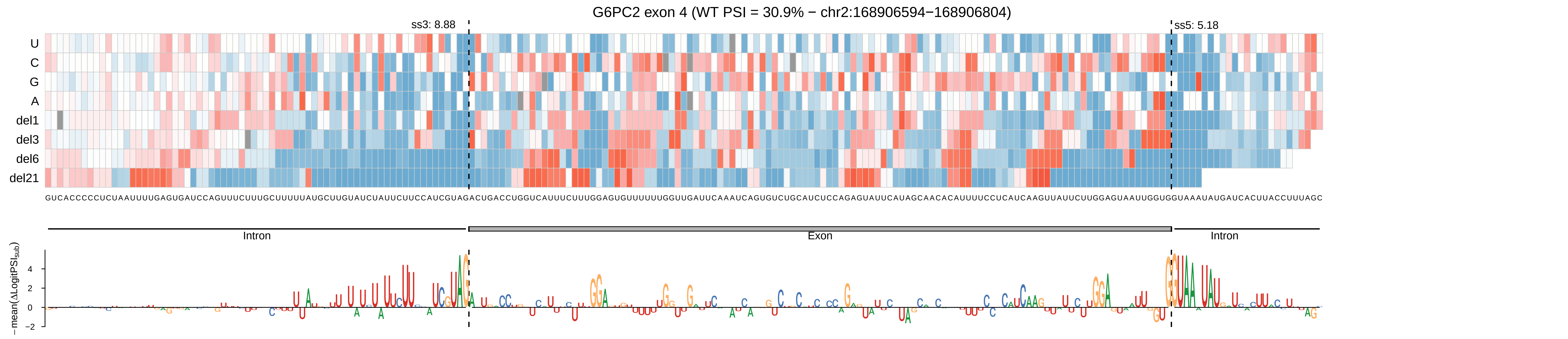

G6PC2 exon 4 (WT PSI = 30.9% − chr2:168906594−168906804)
ss3: 8.88
ss5: 5.18
U
C
G
A
del1
del3
del6
del21
GUCACCCCCUCUAAUUUUGAGUGAUCCAGUUUCUUUGCUUUUUAUGCUUGUAUCUAUUCUUCCAUCGUAGACUGACCUGGUCAUUUCUUUGGAGUGUUUUUUGGUUGAUUCAAAUCAGUGUCUGCAUCUCCAGAGUAUUCAUAGCAACACAUUUUCCUCAUCAAGUUAUUCUUGGAGUAAUUGGUGGUAAAUAUGAUCACUUACCUUUAGC
Intron
Exon
Intron
−mean(ΔLogitPSI_sub)
4
2
0
−2

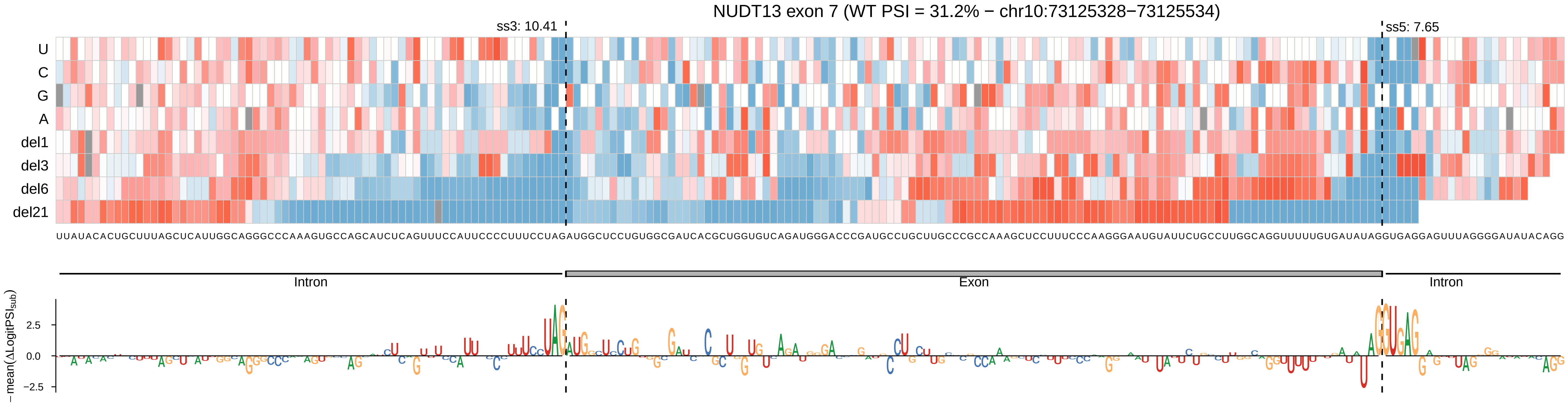

NUDT13 exon 7 (WT PSI = 31.2% − chr10:73125328−73125534)
ss3: 10.41
ss5: 7.65
U
C
G
A
del1
del3
del6
del21
UUAUACACUGCUUUAGCUCAUUGGCAGGGCCCAAAGUGCCAGCAUCUCAGUUUCCAUUCCCCUUUCCUAGAUGGCUCCUGUGGCGAUCACGCUGGUGUCAGAUGGGACCCGAUGCCUGCUUGCCCGCCAAAGCUCCUUUCCCAAGGGAAUGUAUUCUGCCUUGGCAGGUUUUUGUGAUAUAGGUGAGGAGUUUAGGGGAUAUACAGG
Intron
Exon
Intron
−mean(ΔLogitPSI_sub)
2.5
0.0
−2.5

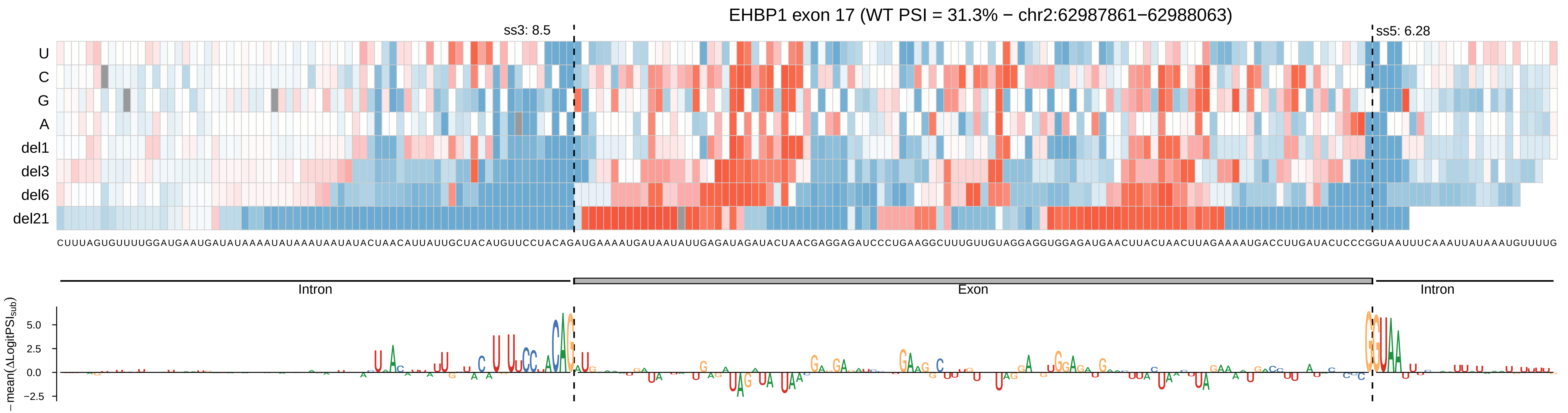
EHBP1 exon 17 (WT PSI = 31.3% − chr2:62987861−62988063)
ss3: 8.5
ss5: 6.28
U
C
G
A
del1
del3
del6
del21
CUUUAGUGUUUUGGAUGAAUGAUAUAAAAUAUAAAUAAUAUACUAACAUUAUUGCUACAUGUUCCUACAGAUGAAAAUGAUAAUAUUGAGAUAGAUACUAACGAGGAGAUCCCUGAAGGCUUUGUUGUAGGAGGUGGAGAUGAACUUACUAACUUAGAAAAUGACCUUGAUACUCCCGGUAAUUUCAAAUUAUAAAUGUUUUG
Intron
Exon
Intron
−mean(ΔLogitPSI_sub)
5.0
2.5
0.0
−2.5

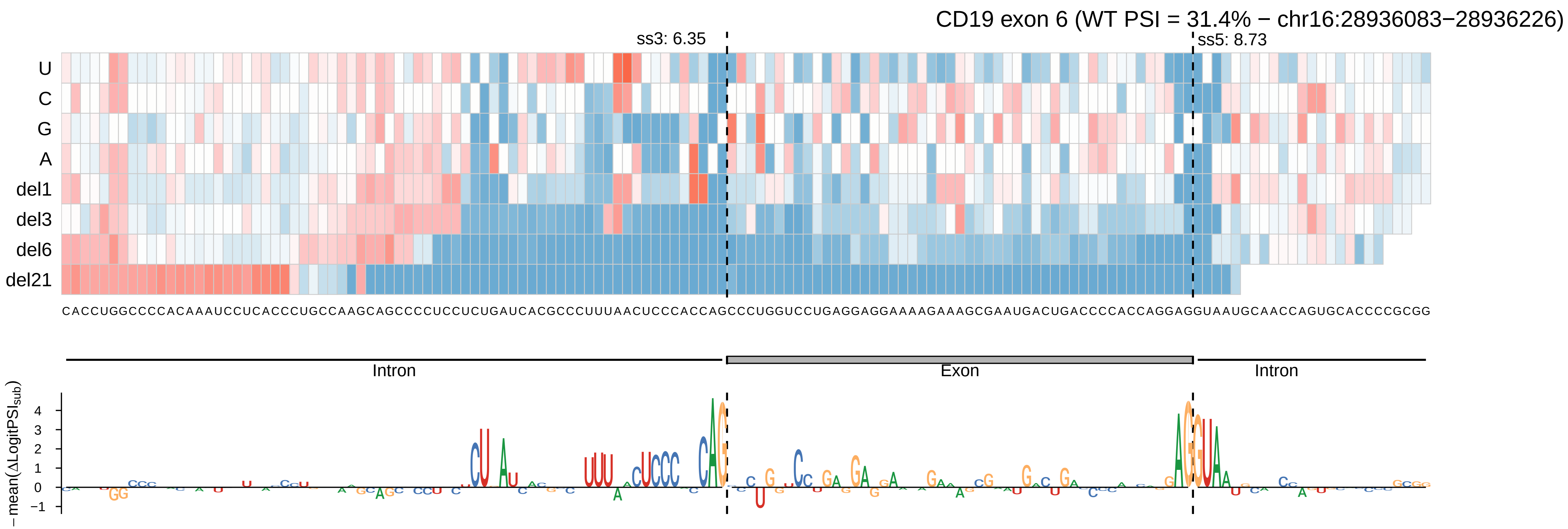

CD19 exon 6 (WT PSI = 31.4% − chr16:28936083−28936226)
ss3: 6.35
ss5: 8.73
U
C
G
A
del1
del3
del6
del21
CACCUGGCCCCACAAAUCCUCACCCUGCCAAGCAGCCCCUCCUCUGAUCACGCCCUUUAACUCCCACCAGCCCUGGUCCUGAGGAGGAAAAGAAAGCGAAUGACUGACCCCACCAGGAGGUAAUGCAACCAGUGCACCCCGCGG
Intron
Exon
Intron
−mean(ΔLogitPSI_sub)
4
3
2
1
0
−1

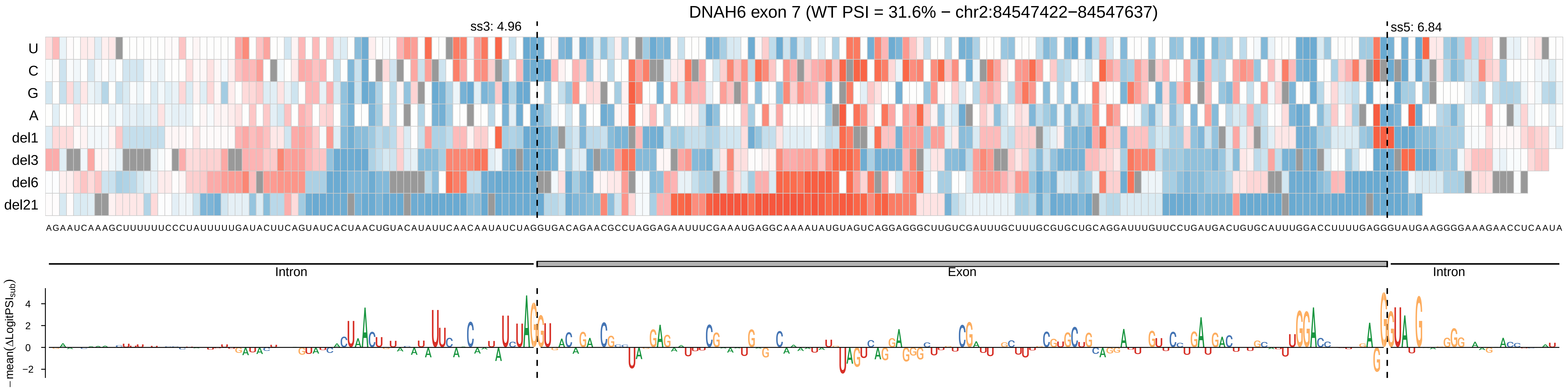
DNAH6 exon 7 (WT PSI = 31.6% − chr2:84547422−84547637)
ss3: 4.96
ss5: 6.84
U
C
G
A
del1
del3
del6
del21
AGAAUCAAAGCUUUUUUCCCUAUUUUUGAUACUUCAGUAUCACUAACUGUACAUAUUCAACAAUAUCUAGGUGACAGAACGCCUAGGAGAAUUUCGAAAUGAGGCAAAAUAUGUAGUCAGGAGGGCUUGUCGAUUUGCUUUGCGUGCUGCAGGAUUUGUUCCUGAUGACUGUGCAUUUGGACCUUUUGAGGGUAUGAAGGGGAAAGAACCUCAAUA
Intron
Exon
Intron
− mean(ΔLogitPSI_sub)
4
2
0
−2

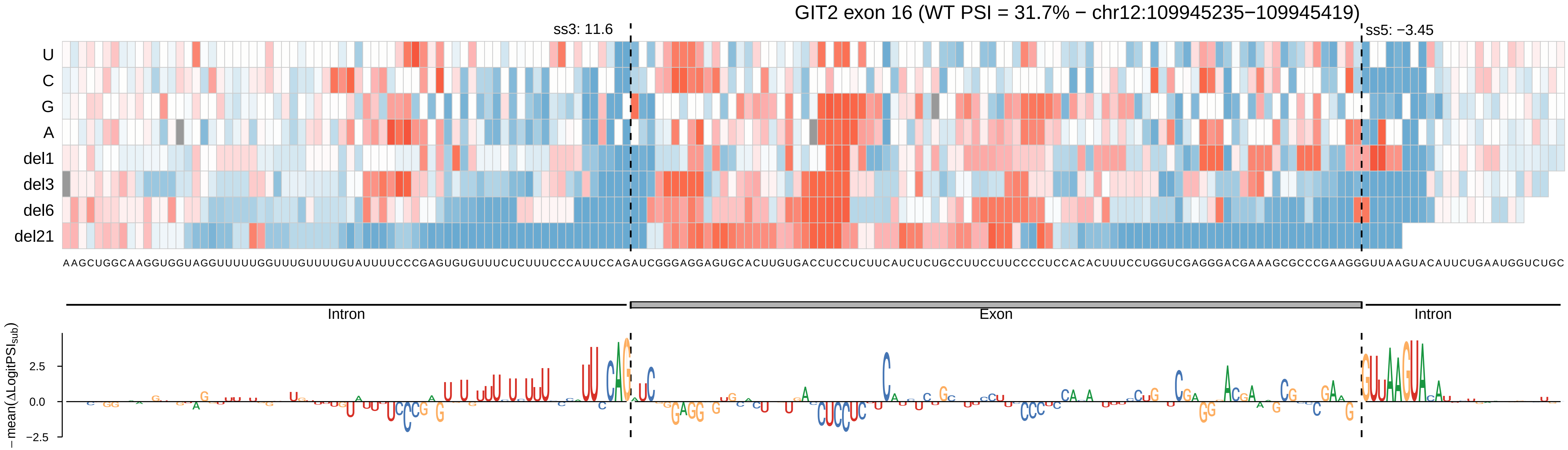
GIT2 exon 16 (WT PSI = 31.7% − chr12:109945235−109945419)
ss3: 11.6
ss5: −3.45
U
C
G
A
del1
del3
del6
del21
AAGCUGGCAAGGUGGUAGGUUUUUGGUUUGUUUUGUAUUUUCCCGAGUGUGUUUCUCUUUCCCAUUCCAGAUCGGGAGGAGUGCACUUGUGACCUCCUCUUCAUCUCUGCCUUCCUUCCCCUCCACACUUUCCUGGUCGAGGGACGAAAGCGCCCGAAGGGUUAAGUACAUUCUGAAUGGUCUGC
Intron
Exon
Intron
−mean(ΔLogitPSI_sub)
2.5
0.0
−2.5

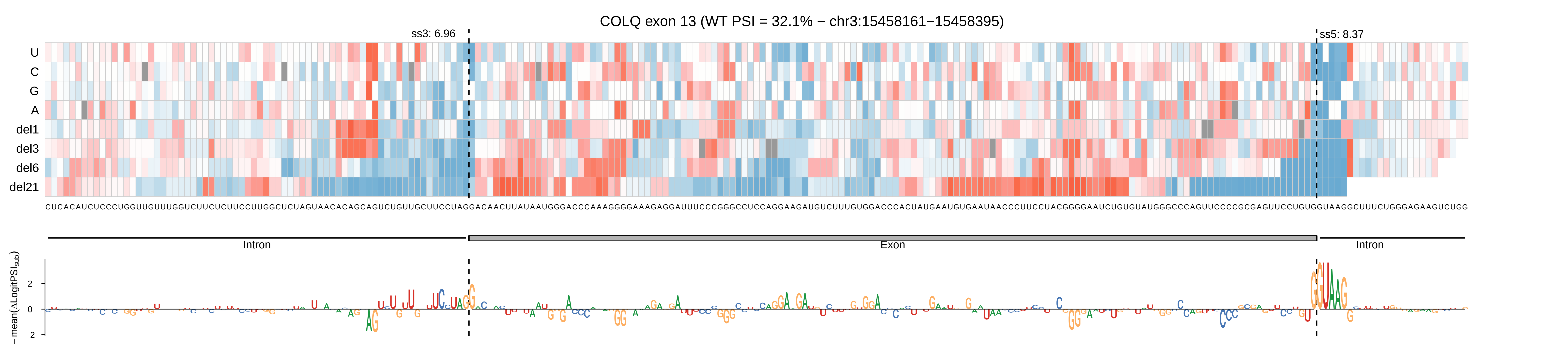

COLQ exon 13 (WT PSI = 32.1% − chr3:15458161−15458395)
ss3: 6.96
ss5: 8.37
U
C
G
A
del1
del3
del6
del21
CUCACAUCUCCCUGGUUGUUUGGUCUUCUCUUCCUUGGCUCUAGUAACACAGCAGUCUGUUGCUUCCUAGGACAACUUAUAAUGGGACCCAAAGGGGAAAGAGGAUUUCCCGGGCCUCCAGGAAGAUGUCUUUGUGGACCCACUAUGAAUGUGAAUAACCCUUCCUACGGGGAAUCUGUGUAUGGGCCCAGUUCCCCGCGAGUUCCUGUGGUAAGGCUUUCUGGGAGAAGUCUGG
Intron
Exon
Intron
− mean(ΔLogitPSI_sub)
2
0
−2

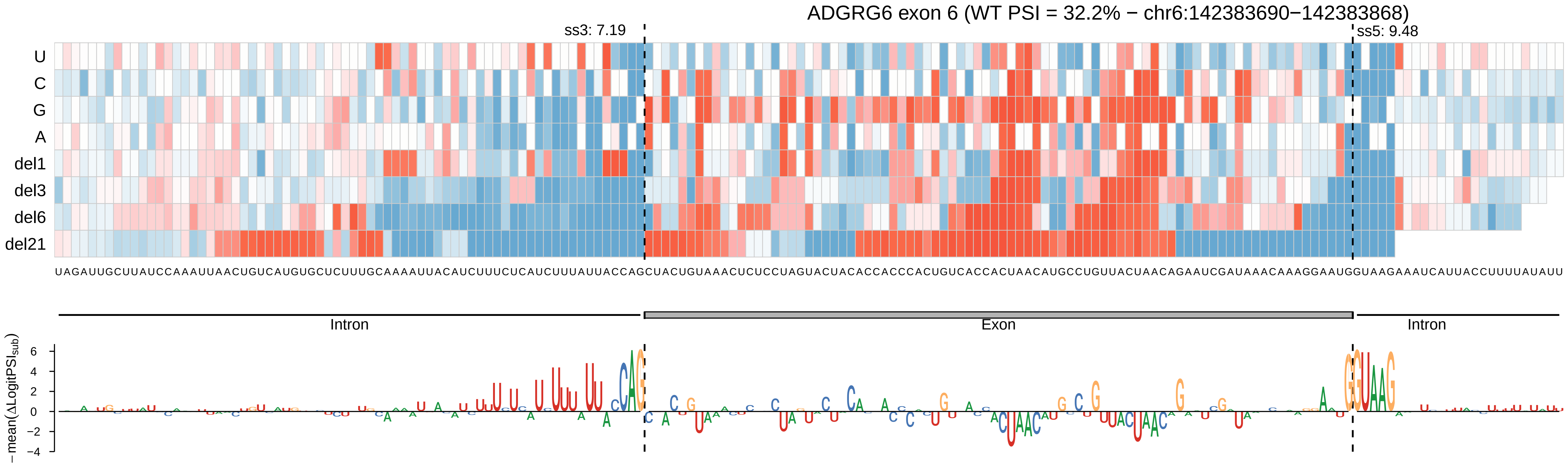
ADGRG6 exon 6 (WT PSI = 32.2% − chr6:142383690−142383868)
ss3: 7.19
ss5: 9.48
U
C
G
A
del1
del3
del6
del21
UAGAUUGCUUAUCCAAAUUAACUGUCAUGUGCUCUUUGCAAAAUUACAUCUUUCUCAUCUUUAUUACCAGCUACUGUAAACUCUCCUAGUACUACACCACCCACUGUCACCACUAACAUGCCUGUUACUAACAGAAUCGAUAAACAAAGGAAUGGUAAGAAAUCAUUACCUUUUAUAUU
Intron
Exon
Intron
−mean(ΔLogitPSI_sub)

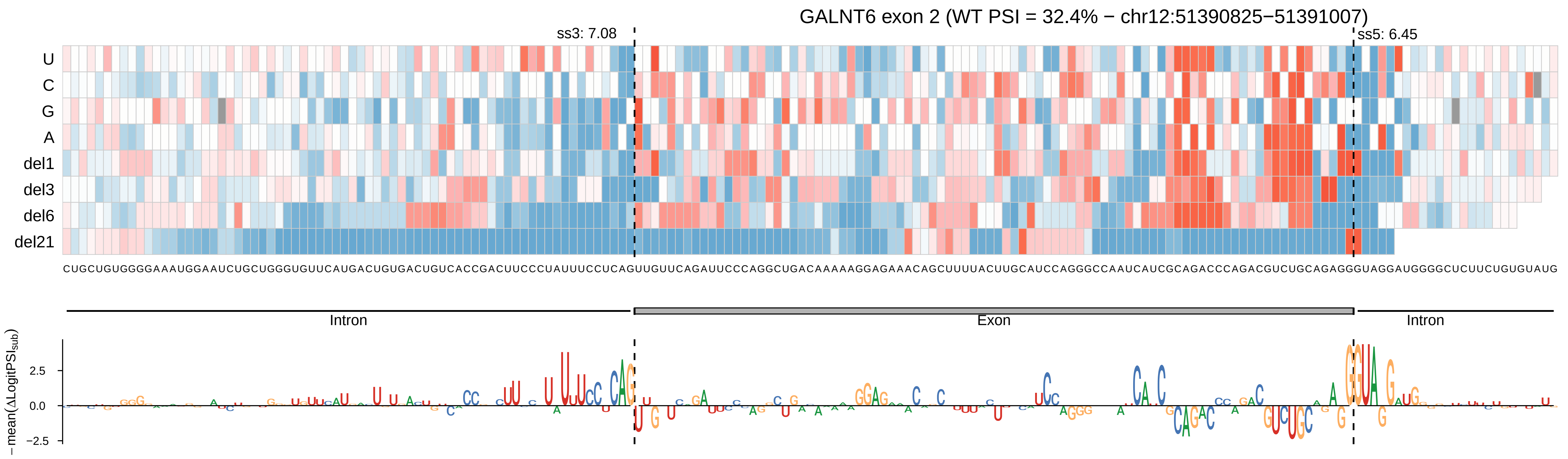
GALNT6 exon 2 (WT PSI = 32.4% − chr12:51390825−51391007)
ss3: 7.08
ss5: 6.45
U
C
G
A
del1
del3
del6
del21
CUGCUGUGGGGAAAUGGAAUCUGCUGGGUGUUCAUGACUGUGACUGUCACCGACUUCCCUAUUUCCUCAGUUGUUCAGAUUCCCAGGCUGACAAAAAGGAGAAACAGCUUUUACUUGCAUCCAGGGCCAAUCAUCGCAGACCCAGACGUCUGCAGAGGGUAGGAUGGGGCUCUUCUGUGUAUG
Intron
Exon
Intron
− mean(ΔLogitPSI_sub)
2.5
0.0
−2.5

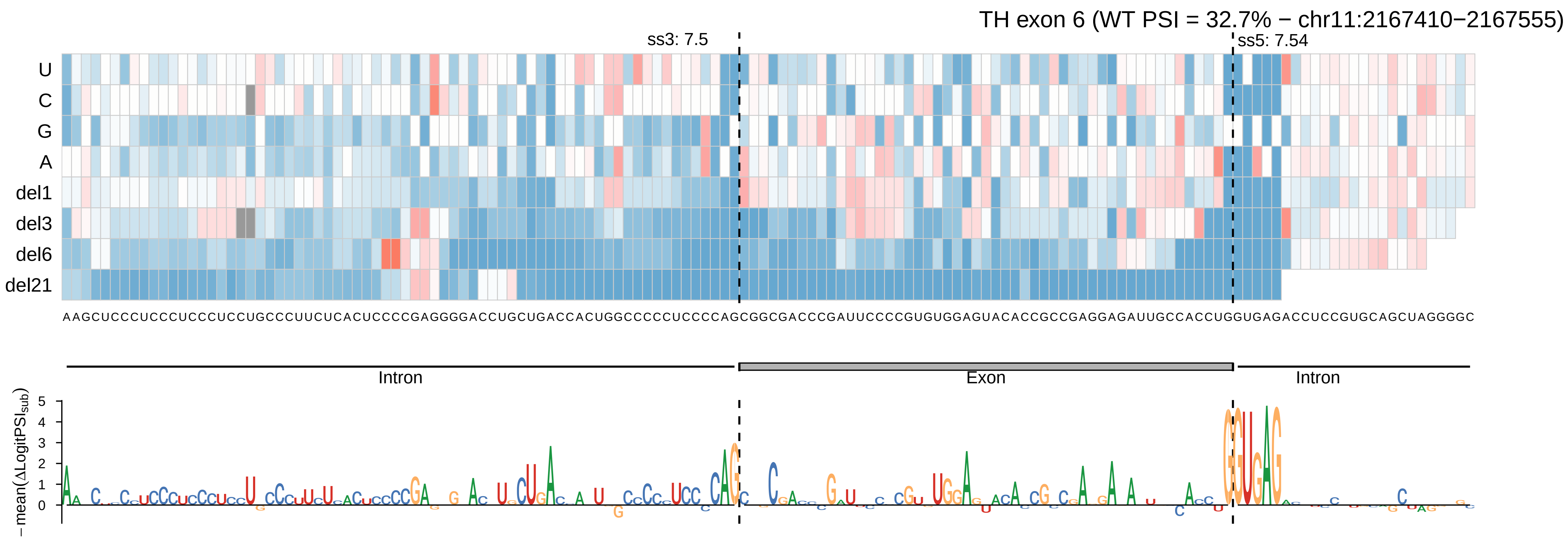

TH exon 6 (WT PSI = 32.7% − chr11:2167410−2167555)
ss3: 7.5
ss5: 7.54
U
C
G
A
del1
del3
del6
del21
AAGCUCCCUCCCUCCCUCCUGCCCUUCUCACUCCCCGAGGGGACCUGCUGACCACUGGCCCCCUCCCCAGCGGCGACCCGAUUCCCCGUGUGGAGUACACCGCCGAGGAGAUUGCCACCUGGUGAGACCUCCGUGCAGCUAGGGGC
Intron
Exon
Intron
−mean(ΔLogitPSI_sub)

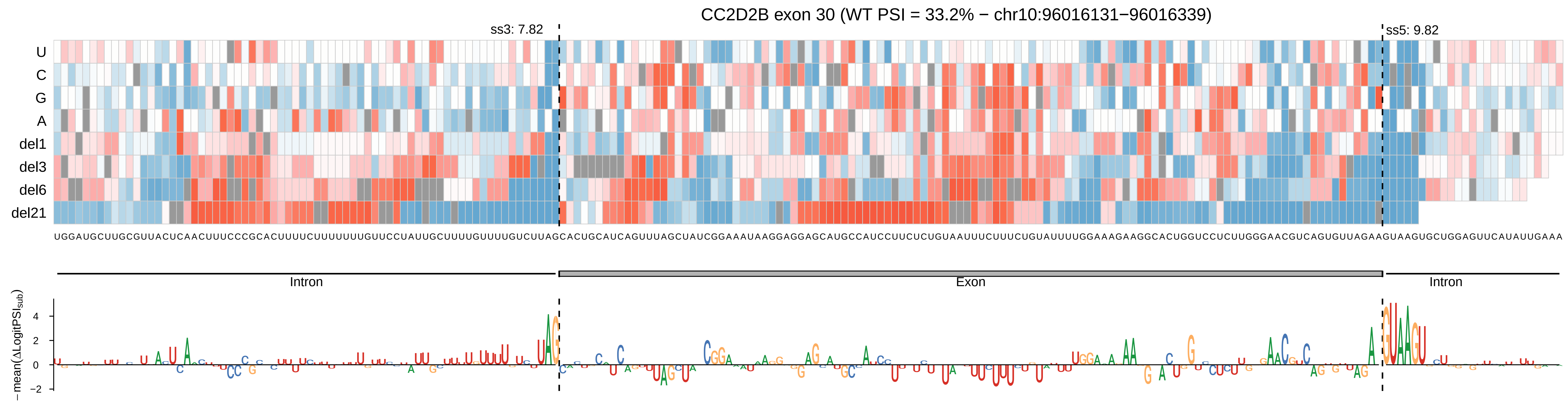

CC2D2B exon 30 (WT PSI = 33.2% − chr10:96016131−96016339)
ss3: 7.82
ss5: 9.82
U
C
G
A
del1
del3
del6
del21
UGGAUGCUUGCGUUACUCAACUUUCCCGCACUUUUCUUUUUUUGUUCCUAUUGCUUUUGUUUUGUCUUAGCACUGCAUCAGUUUAGCUAUCGGAAAUAAGGAGGAGCAUGCCAUCCUUCUCUGUAAUUUCUUUCUGUAUUUUGGAAAGAAGGCACUGGUCCUCUUGGGAACGUCAGUGUUAGAAGUAAGUGCUGGAGUUCAUAUUGAAA
Intron
Exon
Intron
−mean(ΔLogitPSI_sub)
4
2
0
−2

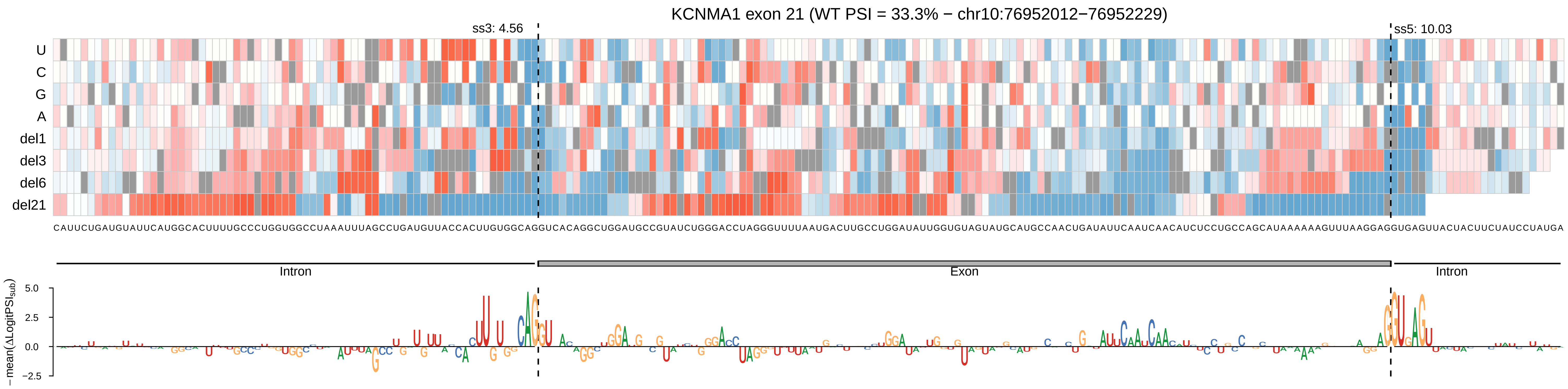

KCNMA1 exon 21 (WT PSI = 33.3% − chr10:76952012−76952229)
ss3: 4.56
ss5: 10.03
U
C
G
A
del1
del3
del6
del21
CAUUCUGAUGUAUUCAUGGCACUUUUGCCCUGGUGGCCUAAAUUUAGCCUGAUGUUACCACUUGUGGCAGGUCACAGGCUGGAUGCCGUAUCUGGGACCUAGGGUUUUAAUGACUUGCCUGGAUAUUGGUGUAGUAUGCAUGCCAACUGAUAUUCAAUCAACAUCUCCUGCCAGCAUAAAAAAGUUUAAGGAGGUGAGUUACUACUUCUAUCCUAUGA
Intron
Exon
Intron
5.0
2.5
0.0
−2.5
− mean(ΔLogitPSI_sub)

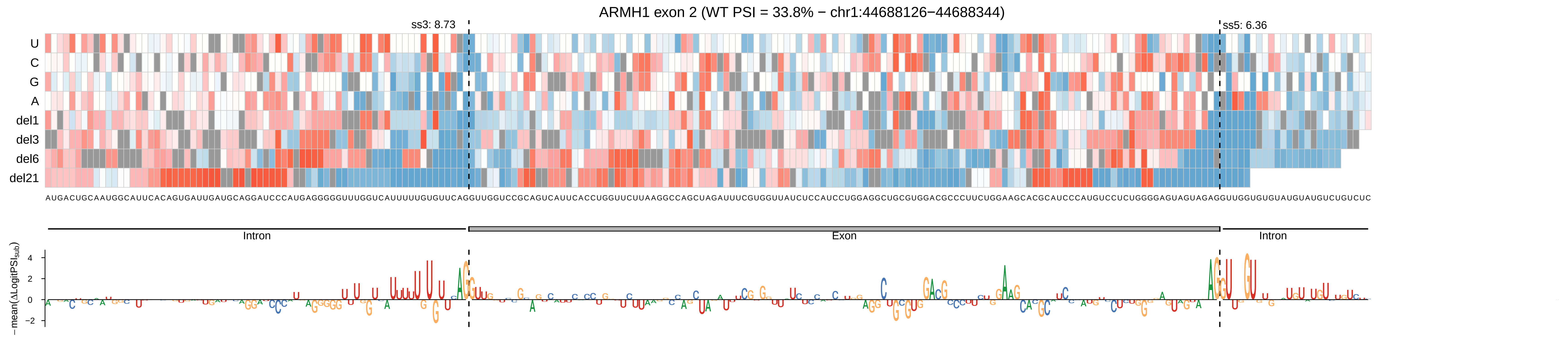

ARMH1 exon 2 (WT PSI = 33.8% − chr1:44688126−44688344)
ss3: 8.73
ss5: 6.36
U
C
G
A
del1
del3
del6
del21
Intron
Exon
Intron
−mean(ΔLogitPSI_sub)
4
2
0
−2

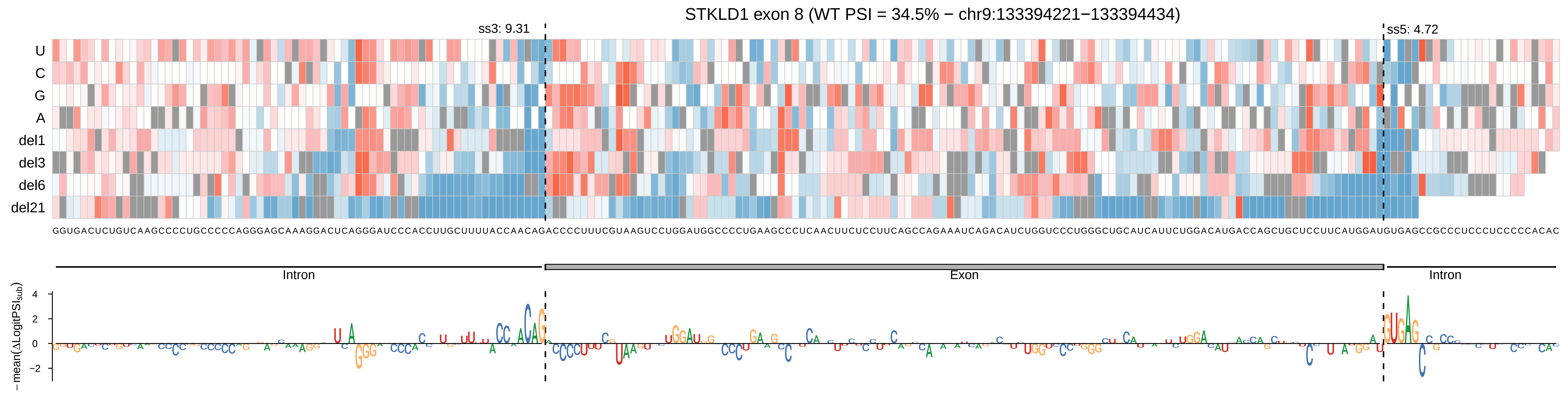
STKLD1 exon 8 (WT PSI = 34.5% − chr9:133394221−133394434)
ss3: 9.31
ss5: 4.72
U
C
G
A
del1
del3
del6
del21
Intron
Exon
Intron
−mean(ΔLogitPSI_sub)
4
2
0
−2

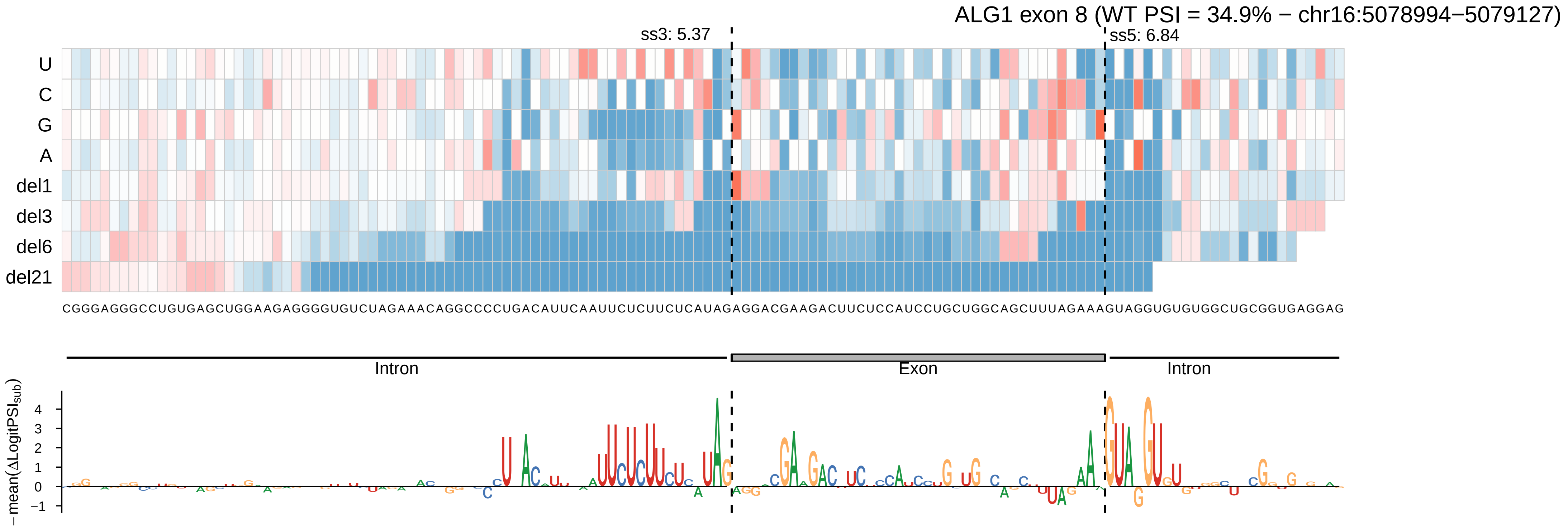
ALG1 exon 8 (WT PSI = 34.9% − chr16:5078994−5079127)
ss3: 5.37
ss5: 6.84
U
C
G
A
del1
del3
del6
del21
CGGGAGGGCCUGUGAGCUGGAAGAGGGGUGUCUAGAAACAGGCCCCUGACAUUCAAUUCUCUUCUCAUAGAGGACGAAGACUUCUCCAUCCUGCUGGCAGCUUUAGAAAGUAGGUGUGUGGCUGCGGUGAGGAG
Intron
Exon
Intron
− mean(ΔLogitPSI_sub)
4
3
2
1
0
−1

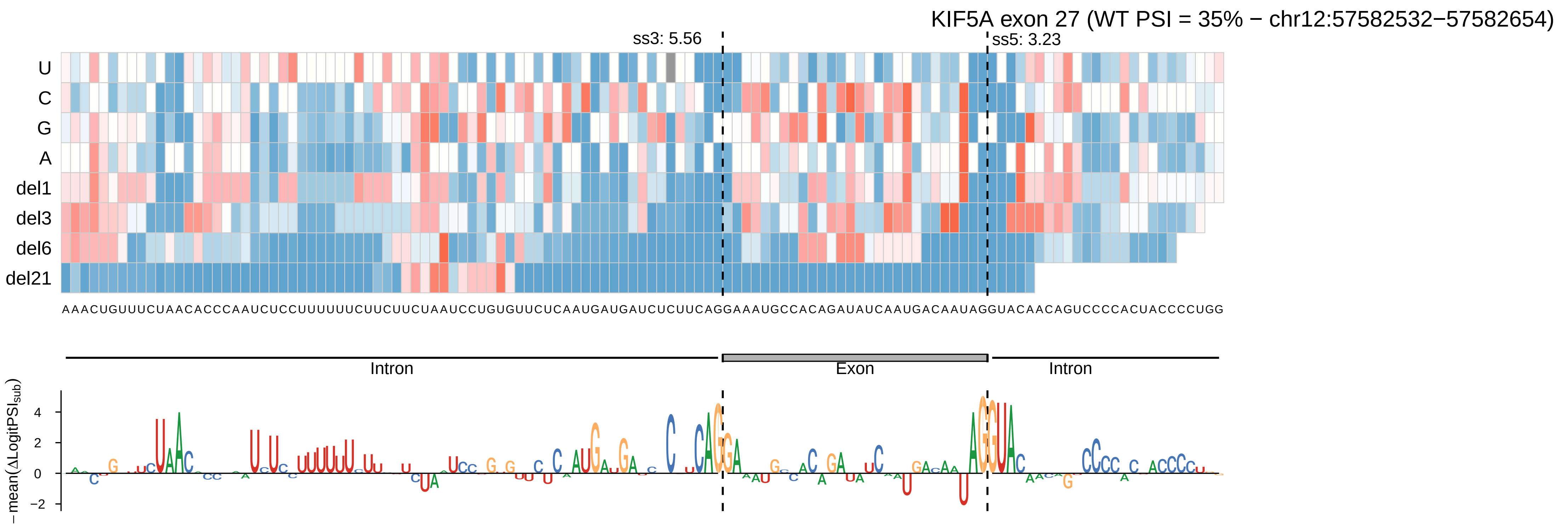

KIF5A exon 27 (WT PSI = 35% − chr12:57582532−57582654)
ss3: 5.56
ss5: 3.23
U
C
G
A
del1
del3
del6
del21
AAACUGUUUCUAACACCCAAUCUCCUUUUUUCUUCUUCUAAUCCUGUGUUCUCAAUGAUGAUCUCUUCAGGAAAUGCCACAGAUAUCAAUGACAAUAGGUACAACAGUCCCCACUACCCCUGG
Intron
Exon
Intron
− mean(ΔLogitPSI sub)
4
2
0
−2

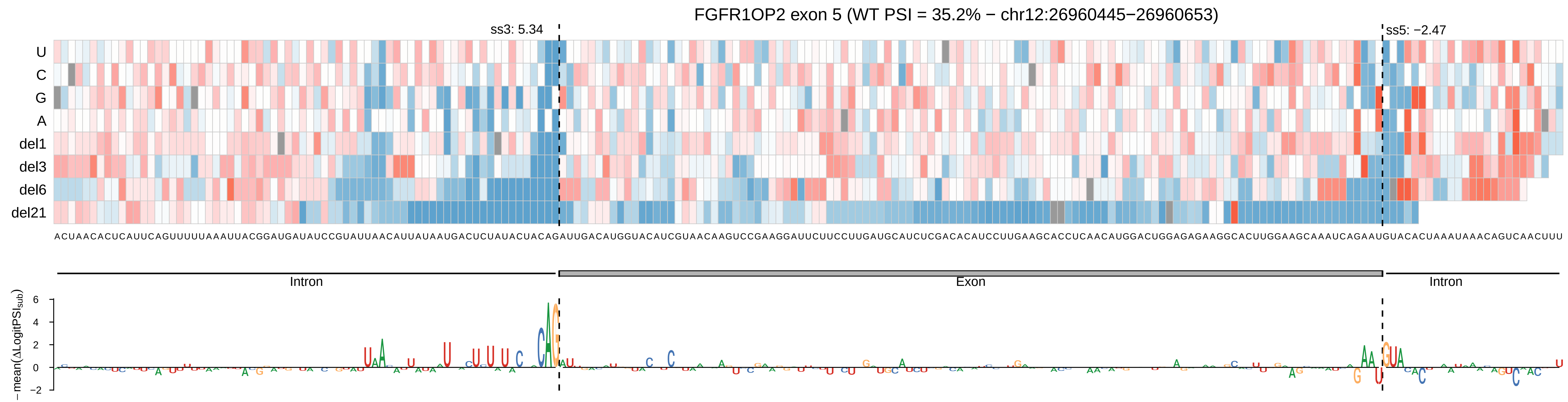

FGFR1OP2 exon 5 (WT PSI = 35.2% − chr12:26960445−26960653)
ss3: 5.34
ss5: −2.47
U
C
G
A
del1
del3
del6
del21
ACUAACACUCAUUCAGUUUUUAAAUUACGGAUGAUAUCCGUAUUAACAUUAUAAUGACUCUAUACUACAGAUUGACAUGGUACAUCGUAACAAGUCCGAAGGAUUCUUCCUUGAUGCAUCUCGACACAUCCUUGAAGCACCUCAACAUGGACUGGAGAGAAGGCACUUGGAAGCAAAUCAGAAUGUACACUAAAUAAACAGUCAACUUU
Intron
Exon
Intron
− mean(ΔLogitPSI_sub)
6
4
2
0
−2

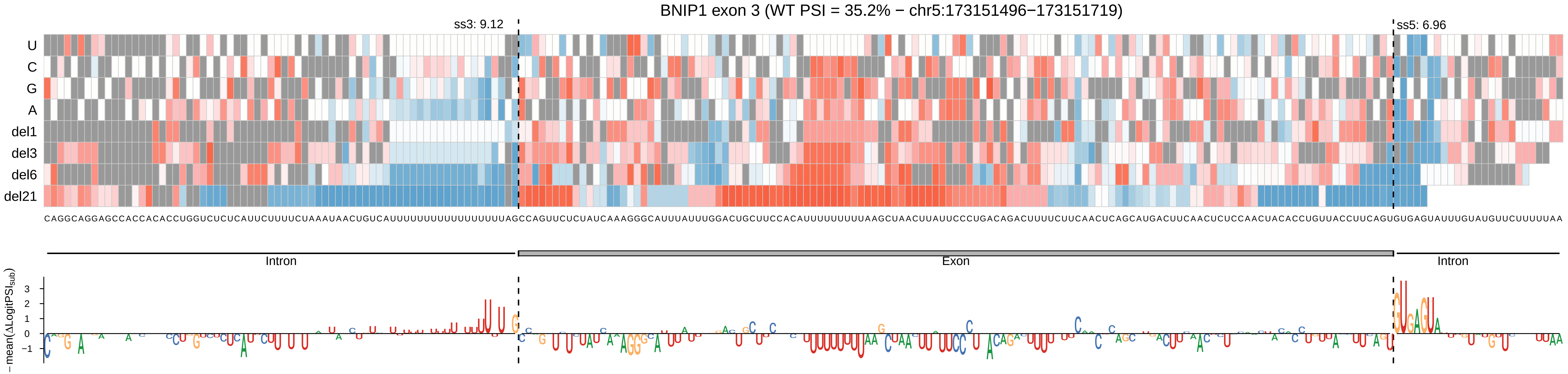
BNIP1 exon 3 (WT PSI = 35.2% − chr5:173151496−173151719)
ss3: 9.12
ss5: 6.96
U
C
G
A
del1
del3
del6
del21
Intron
Exon
Intron
−mean(ΔLogitPSI_sub)
3
2
1
0
−1

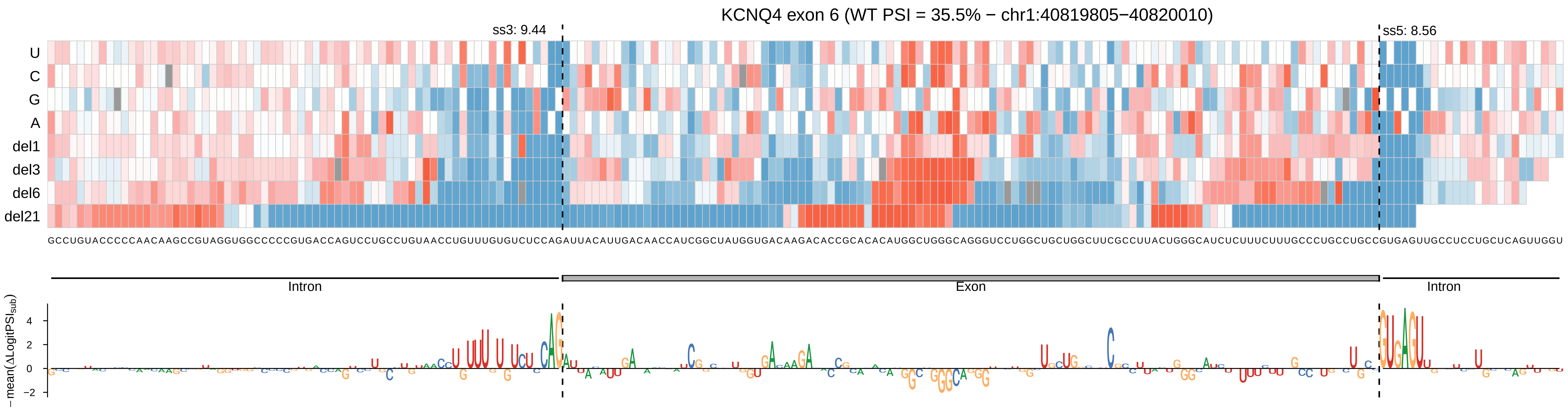

KCNQ4 exon 6 (WT PSI = 35.5% − chr1:40819805−40820010)
ss3: 9.44
ss5: 8.56
U
C
G
A
del1
del3
del6
del21
Intron
Exon
Intron
−mean(ΔLogitPSI_sub)
4
2
0
−2

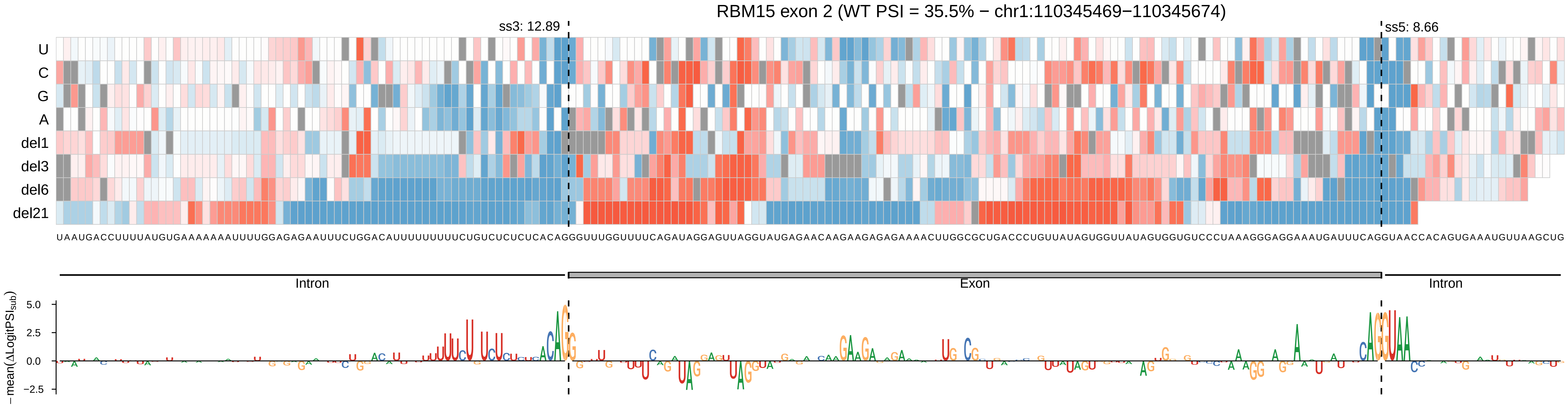
RBM15 exon 2 (WT PSI = 35.5% − chr1:110345469−110345674)
ss3: 12.89
ss5: 8.66
U
C
G
A
del1
del3
del6
del21
UAAUGACCUUUUAUGUGAAAAAAAUUUUGGAGAGAAUUUCUGGACAUUUUUUUUUCUGUCUCUCUCACAGGGUUUGGUUUUCAGAUAGGAGUUAGGUAUGAGAACAAGAAGAGAGAAAACUUGGCGCUGACCCUGUUAUAGUGGUUAUAGUGGUGUCCCUAAAGGGAGGAAAUGAUUUCAGGUAACCACAGUGAAAUGUUAAGCUG
Intron
Exon
Intron
−mean(ΔLogitPSI_sub)
5.0
2.5
0.0
−2.5

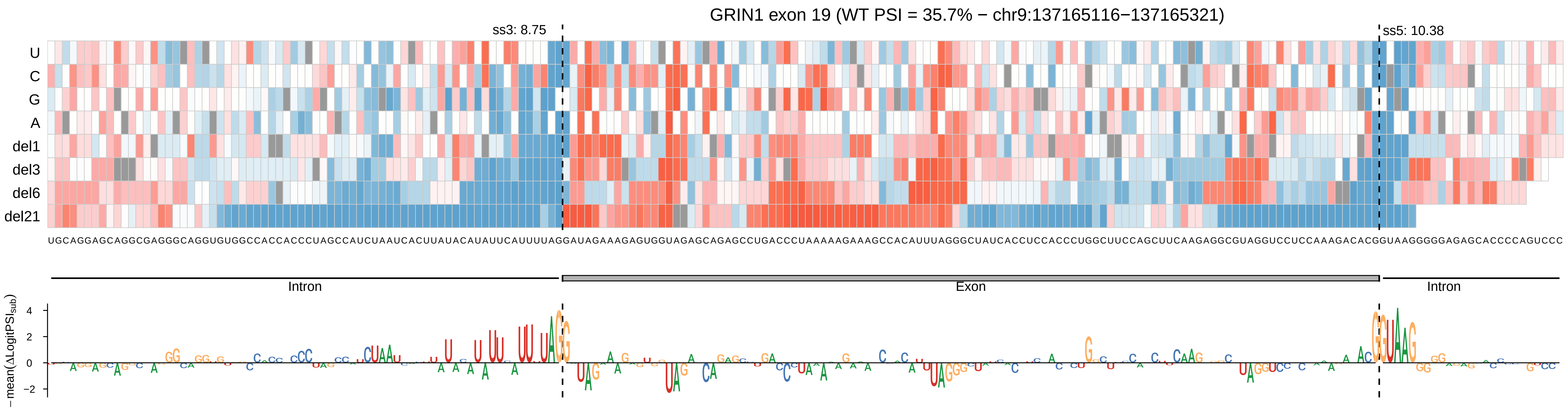

GRIN1 exon 19 (WT PSI = 35.7% − chr9:137165116−137165321)
ss3: 8.75
ss5: 10.38
U
C
G
A
del1
del3
del6
del21
UGCAGGAGCAGGCGAGGGCAGGUGUGGCCACCACCCUAGCCAUCUAAUCACUUAUACAUAUUCAUUUUAGGAUAGAAAGAGUGGUAGAGCAGAGCCUGACCCUAAAAAGAAAGCCACAUUUAGGGCUAUCACCUCCACCCUGGCUUCCAGCUUCAAGAGGCGUAGGUCCUCCAAAGACACGGUAAGGGGGAGAGCACCCCAGUCCC
Intron
Exon
Intron
−mean(ΔLogitPSI_sub)
4
2
0
−2

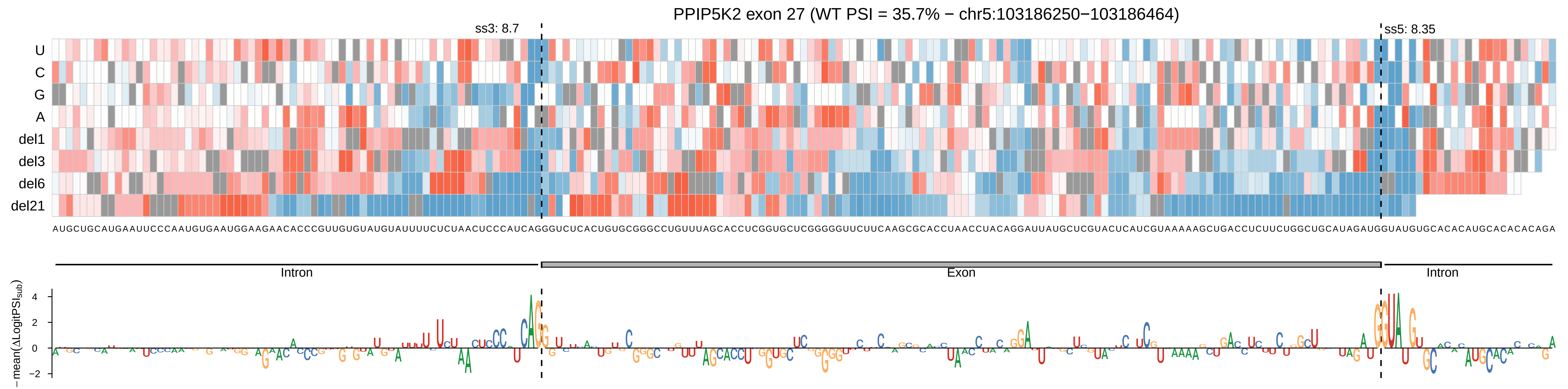
PPIP5K2 exon 27 (WT PSI = 35.7% − chr5:103186250−103186464)
ss3: 8.7
ss5: 8.35
U
C
G
A
del1
del3
del6
del21
AUGCUGCAUGAAUUCCCAAUGUGAAUGGAAGAACACCCGUUGUGUAUGUAUUUUCUCUAACUCCCAUCAGGGUCUCACUGUGCGGGCCUGUUUAGCACCUCGGUGCUCGGGGGUUCUUCAAGCGCACCUAACCUACAGGAUUAUGCUCGUACUCAUCGUAAAAAGCUGACCUCUUCUGGCUGCAUAGAUGGUAUGUGCACACAUGCACACACAGA
Intron
Exon
Intron
−mean(ΔLogitPSI_sub)
4
2
0
−2

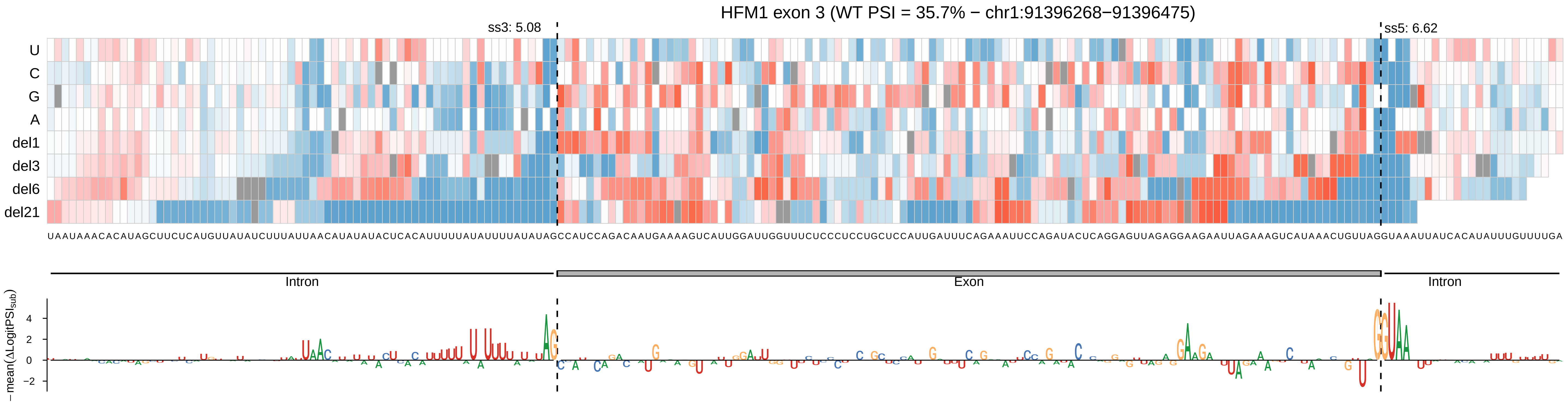

HFM1 exon 3 (WT PSI = 35.7% − chr1:91396268−91396475)
ss3: 5.08
ss5: 6.62
U
C
G
A
del1
del3
del6
del21
Intron
Exon
Intron
−mean(ΔLogitPSI_sub)
4
2
0
−2

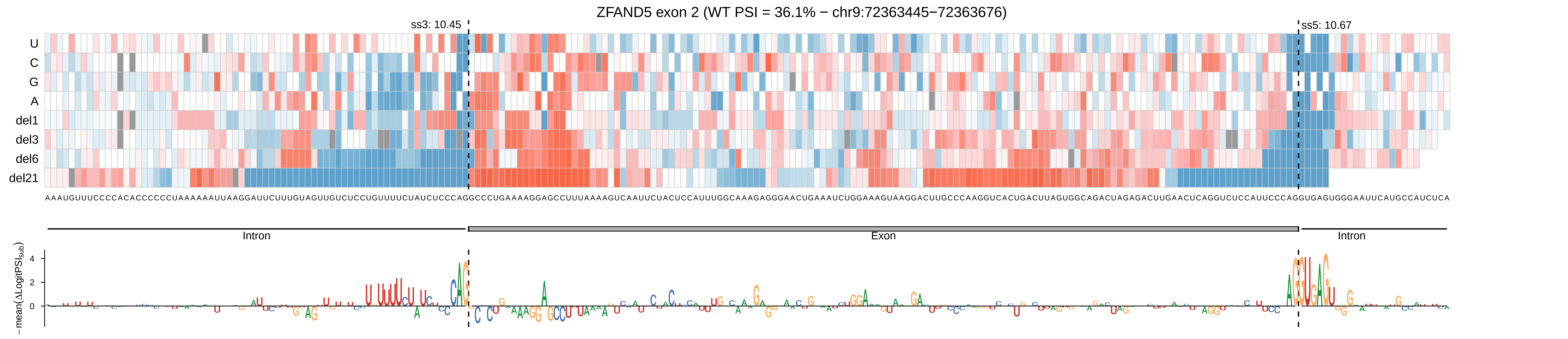

ZFAND5 exon 2 (WT PSI = 36.1% − chr9:72363445−72363676)
ss3: 10.45
ss5: 10.67
U
C
G
A
del1
del3
del6
del21
AAAUGUUUCCCCACACCCCCCUAAAAAAUUAAGGAUUCUUUGUAGUUGUCUCCUGUUUUCUAUCUCCCAGGCCCUGAAAAGGAGCCUUUAAAAGUCAAUUCUACUCCAUUUGGCAAAGAGGGAACUGAAAUCUGGAAAGUAAGGACUUGCCCAAGGUCACUGACUUAGUGGCAGACUAGAGACUUGAACUCAGGUCUCCAUUCCCAGGUGAGUGGGAAUUCAUGCCAUCUCA
Intron
Exon
Intron
−mean(ΔLogitPSI_sub)
4
2
0

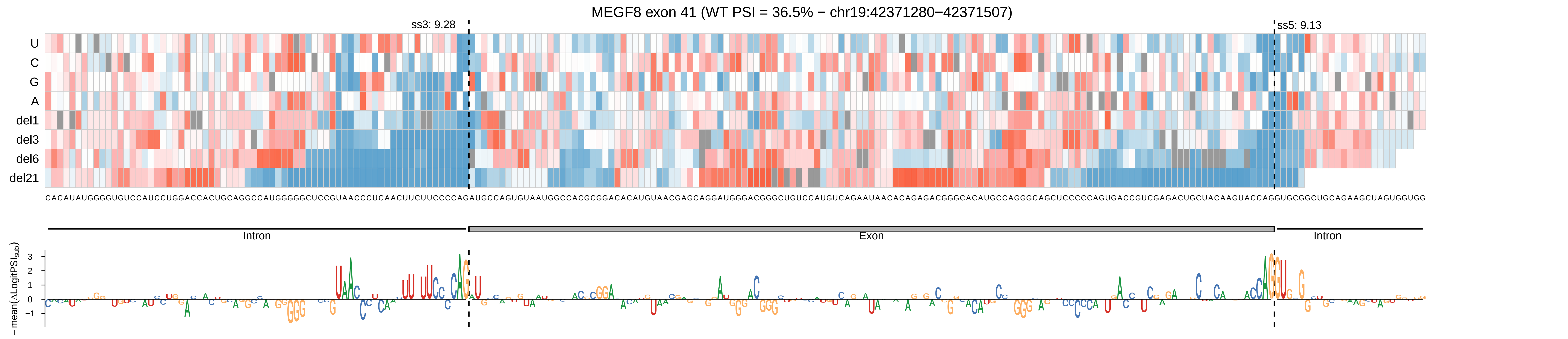

MEGF8 exon 41 (WT PSI = 36.5% − chr19:42371280−42371507)
ss3: 9.28
ss5: 9.13
U
C
G
A
del1
del3
del6
del21
Intron
Exon
Intron
−mean(ΔLogitPSI_sub)

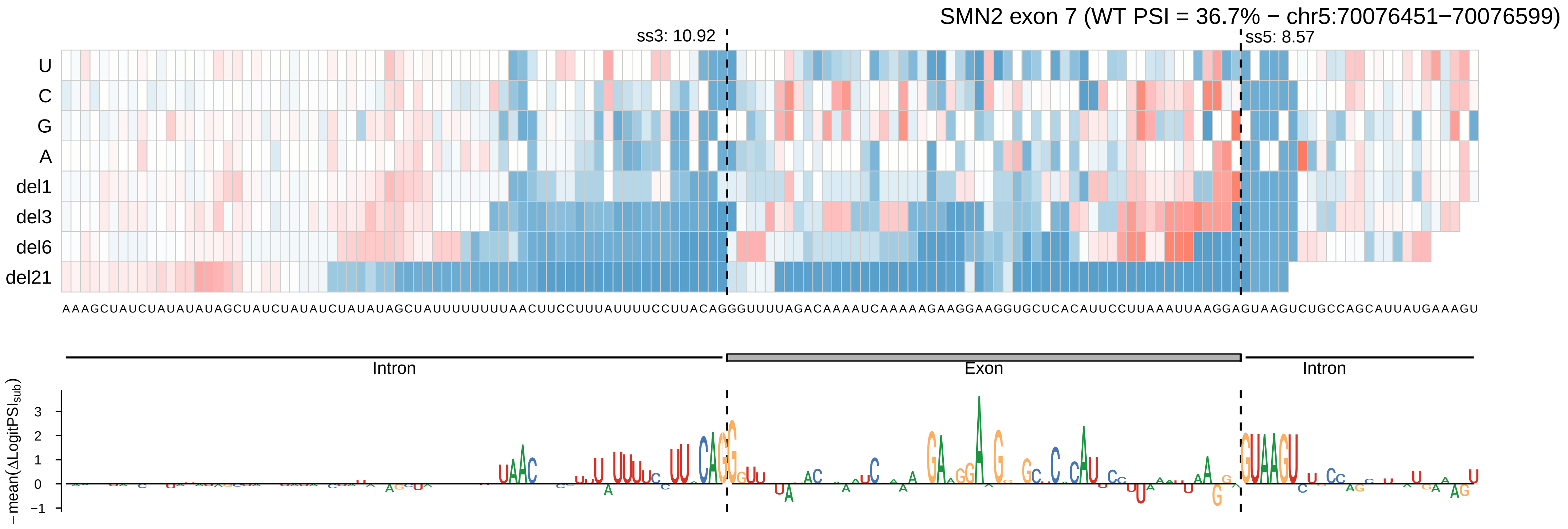
SMN2 exon 7 (WT PSI = 36.7% − chr5:70076451−70076599)
ss3: 10.92
ss5: 8.57
U
C
G
A
del1
del3
del6
del21
AAAGCUAUCUAUAUAUAGCUAUCUAUAUCUAUAUAGCUAUUUUUUUUAACUUCCUUUAUUUUCCUUACAGGGUUUUAGACAAAAUCAAAAAGAAGGAAGGUGCUCACAUUCCUUAAAUUAAGGAGUAAGUCUGCCAGCAUUAUGAAAGU
Intron
Exon
Intron
− mean(ΔLogitPSI_sub)
3
2
1
0
−1

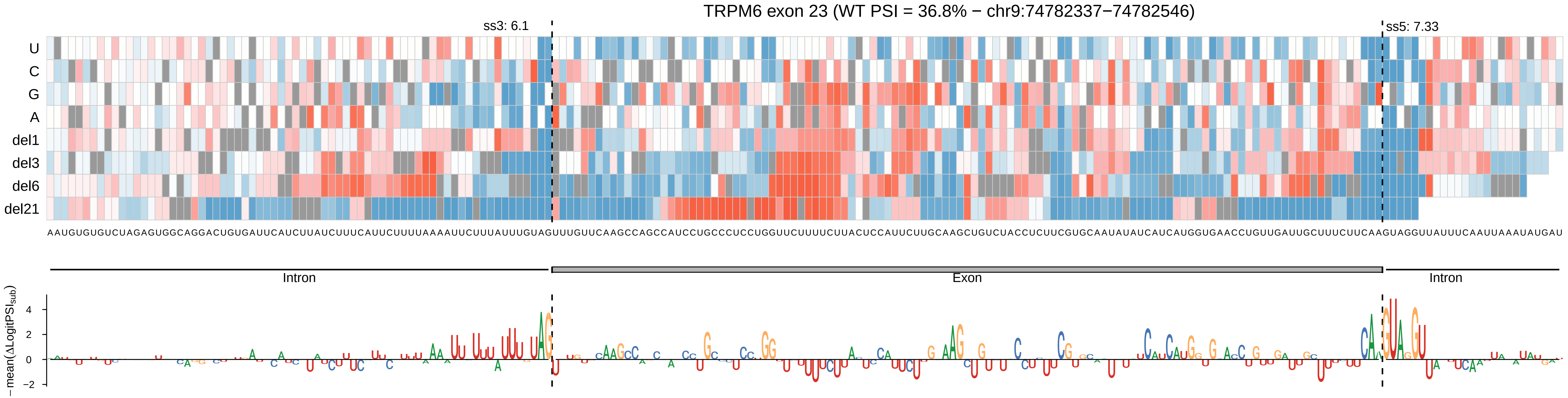
TRPM6 exon 23 (WT PSI = 36.8% − chr9:74782337−74782546)
ss3: 6.1
ss5: 7.33
U
C
G
A
del1
del3
del6
del21
Intron
Exon
Intron
−mean(ΔLogitPSIsub)
4
2
0
−2

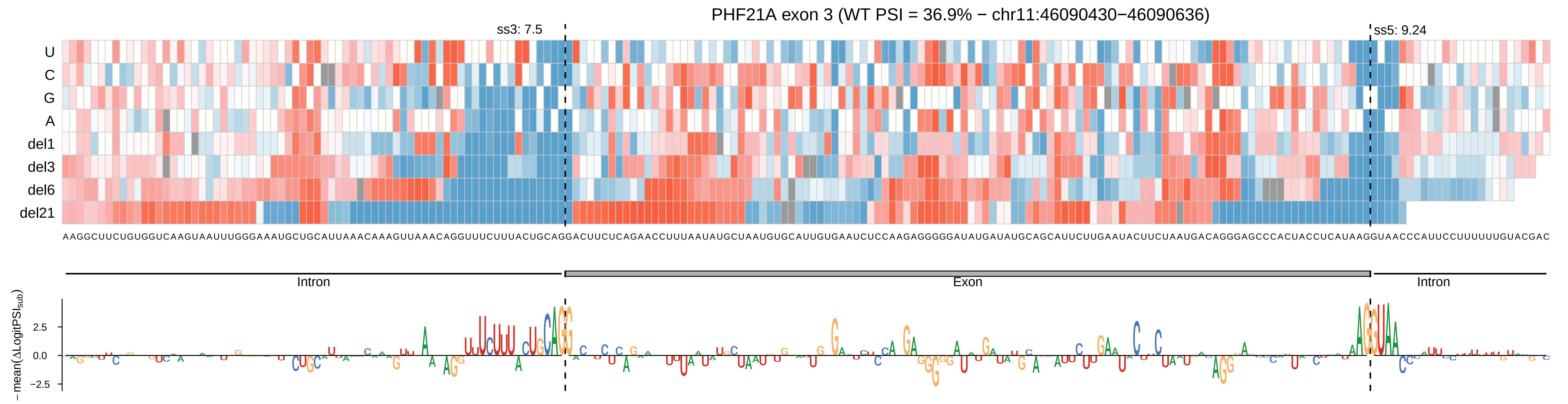

PHF21A exon 3 (WT PSI = 36.9% − chr11:46090430−46090636)
ss3: 7.5
ss5: 9.24
U
C
G
A
del1
del3
del6
del21
AAGGCUUCUGUGGUCAAGUAAUUUGGGAAAUGCUGCAUUAAACAAAGUUAAACAGGUUUCUUUACUGCAGGACUUCUCAGAACCUUUAAUAUGCUAAUGUGCAUUGUGAAUCUCCAAGAGGGGGAUAUGAUAUGCAGCAUUCUUGAAUACUUCUAAUGACAGGGAGCCCACUACCUCAUAAGGUAACCCAUUCCUUUUUUGUACGAC
Intron
Exon
Intron
−mean(ΔLogitPSI_sub)
2.5
0.0
−2.5

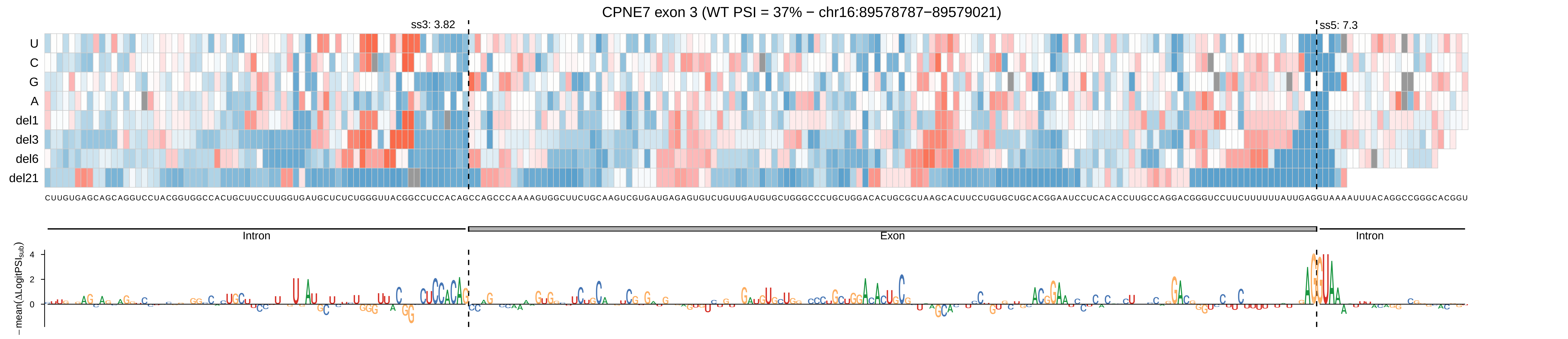
CPNE7 exon 3 (WT PSI = 37% − chr16:89578787−89579021)
ss3: 3.82
ss5: 7.3
U
C
G
A
del1
del3
del6
del21
CUUGUGAGCAGCAGGUCCUACGGUGGCCACUGCUUCCUUGGUGAUGCUCUCUGGGUUACGGCCUCCACAGCCAGCCCAAAAGUGGCUUCUGCAAGUCGUGAUGAGAGUGUCUGUUGAUGUGCUGGGCCCUGCUGGACACUGCGCUAAGCACUUCCUGUGCUGCACGGAAUCCUCACACCUUGCCAGGACGGGUCCUUCUUUUUUAUUGAGGUAAAAUUUACAGGCCGGGCACGGU
Intron
Exon
Intron
−mean(ΔLogitPSI_sub)
4
2
0

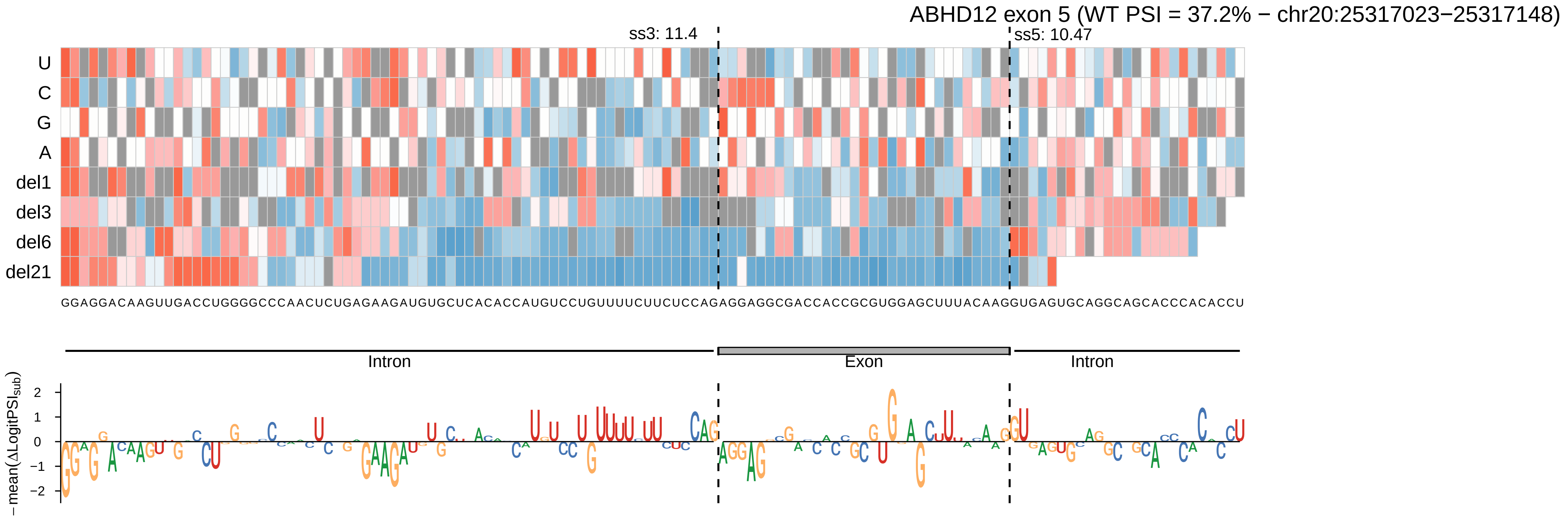
ABHD12 exon 5 (WT PSI = 37.2% − chr20:25317023−25317148)
ss3: 11.4
ss5: 10.47
U
C
G
A
del1
del3
del6
del21
GGAGGACAAGUUGACCUGGGGCCCAACUCUGAGAAGAUGUGCUCACACCAUGUCCUGUUUUCUUCUCCAGAGGAGGCGACCACCGCGUGGAGCUUUACAAGGUGAGUGCAGGCAGCACCCACACCU
Intron
Exon
Intron
−mean(ΔLogitPSI_sub)
2
1
0
−1
−2

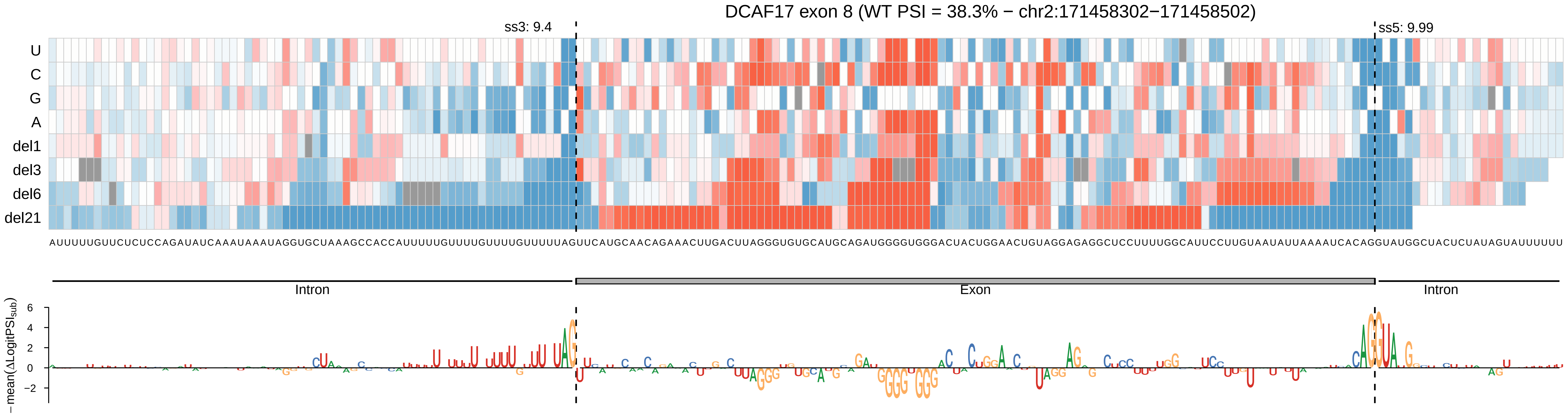
DCAF17 exon 8 (WT PSI = 38.3% − chr2:171458302−171458502)
ss3: 9.4
ss5: 9.99
U
C
G
A
del1
del3
del6
del21
Intron
Exon
Intron
−mean(ΔLogitPSI_sub)

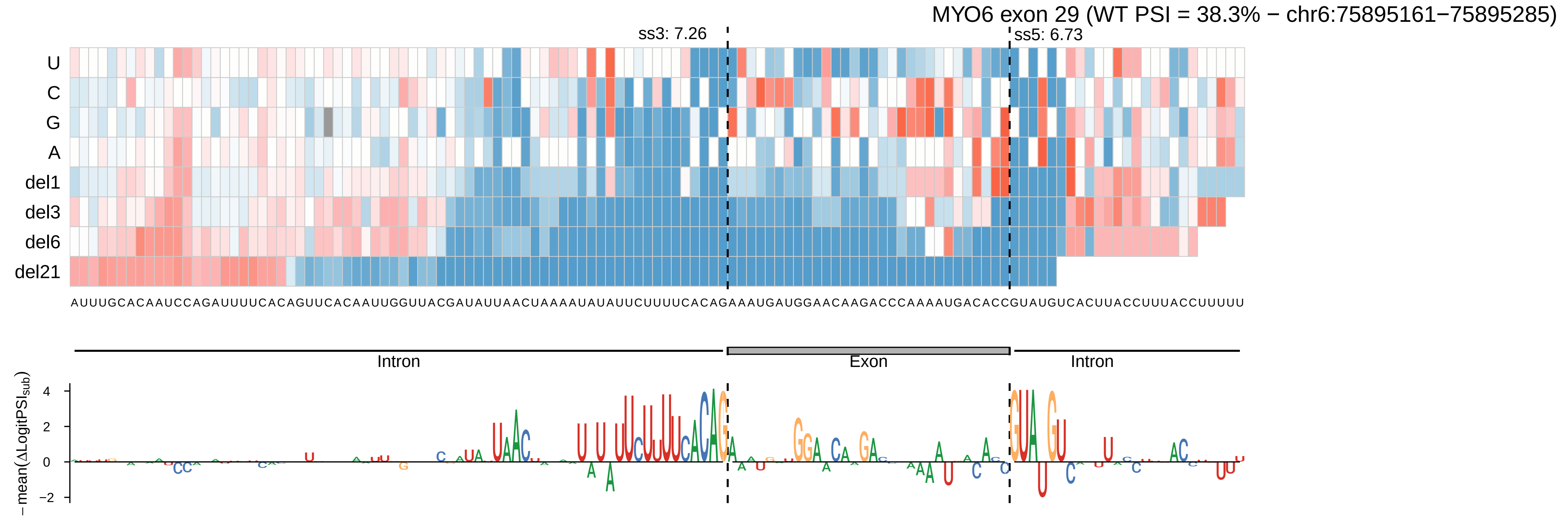
MYO6 exon 29 (WT PSI = 38.3% − chr6:75895161−75895285)
ss3: 7.26
ss5: 6.73
U
C
G
A
del1
del3
del6
del21
AUUUGCACAAUCCAGAUUUUCACAGUUCACAAUUGGUUACGAUAUUAACUAAAAUAUAUUCUUUUCACAGAAAUGAUGGAACAAGACCCAAAAUGACACCGUAUGUCACUUACCUUUACCUUUUU
Intron
Exon
Intron
− mean(ΔLogitPSI sub)
4
2
0
−2

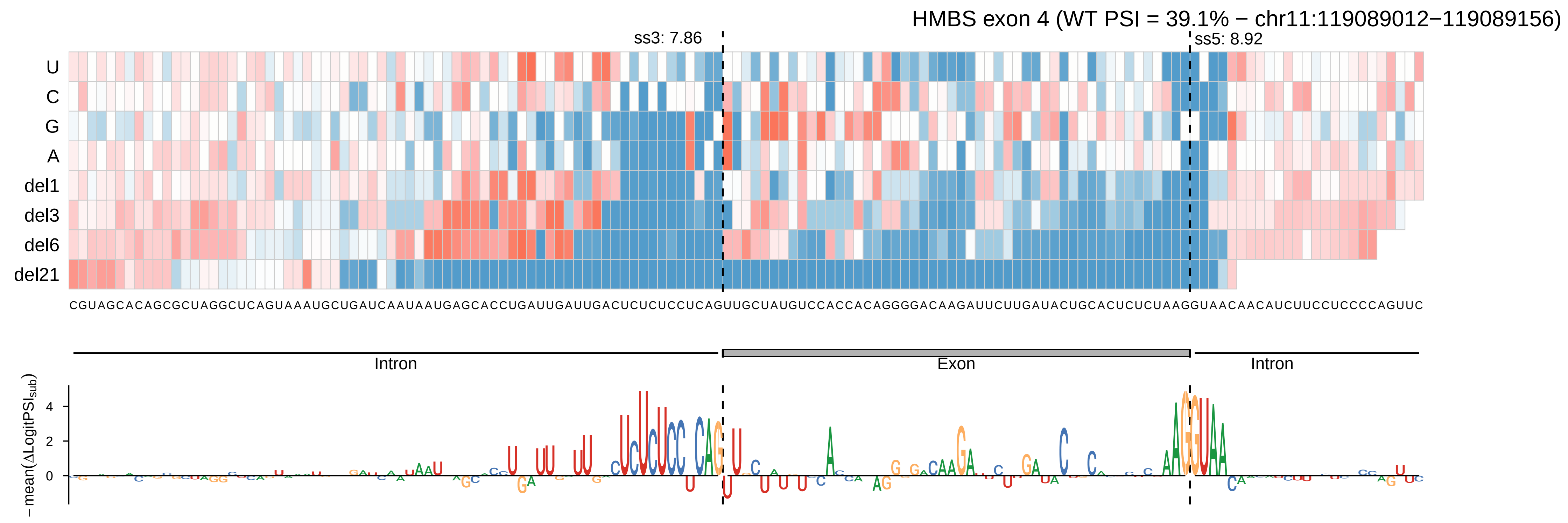
HMBS exon 4 (WT PSI = 39.1% − chr11:119089012−119089156)
ss3: 7.86
ss5: 8.92
U
C
G
A
del1
del3
del6
del21
CGUAGCACAGCGCUAGGCUCAGUAAAUGCUGAUCAAUAAUGAGCACCUGAUUGAUUGACUCUCUCCUCAGUUGCUAUGUCCACCACAGGGGACAAGAUUCUUGAUACUGCACUCUCUAAGGUAACAACAUCUUCCUCCCCAGUUC
Intron
Exon
Intron
−mean(ΔLogitPSI_sub)
4
2
0

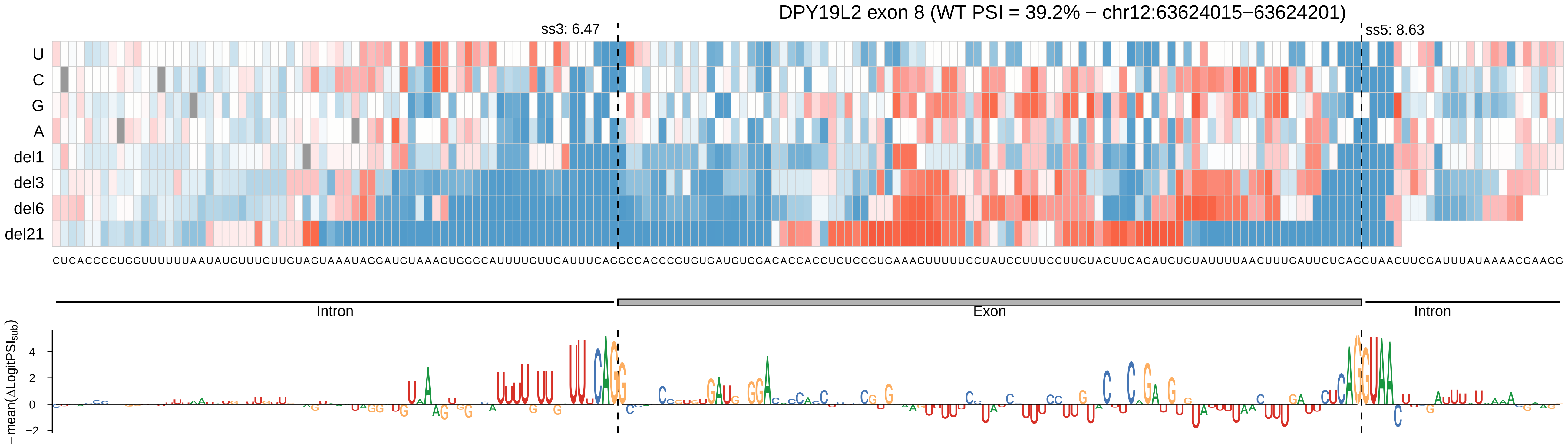
DPY19L2 exon 8 (WT PSI = 39.2% − chr12:63624015−63624201)
ss3: 6.47
ss5: 8.63
U
C
G
A
del1
del3
del6
del21
CUCACCCCUGGUUUUUUAAUAUGUUUGUUGUAGUAAAUAGGAUGUAAAGUGGGCAUUUUGUUGAUUUCAGGCCACCCGUGUGAUGUGGACACCACCUCUCCGUGAAAGUUUUUCCUAUCCUUUCCUUGUACUUCAGAUGUGUAUUUUAACUUUGAUUCUCAGGUAACUUCGAUUUAUAAAACGAAGG
Intron
Exon
Intron
−mean(ΔLogitPSI_sub)
4
2
0
−2

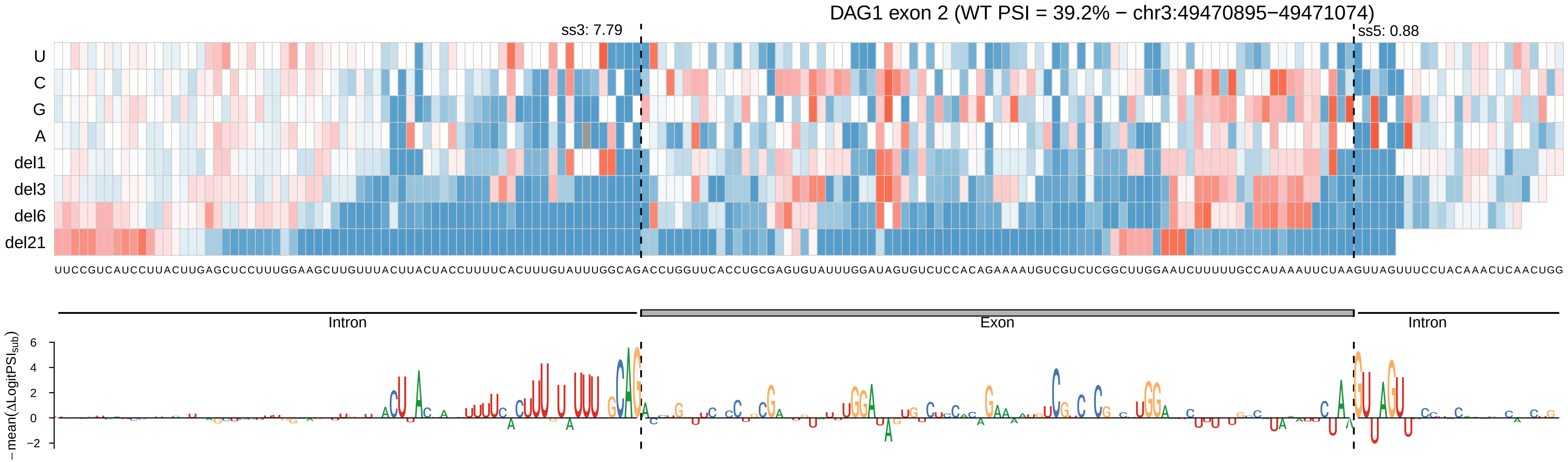
DAG1 exon 2 (WT PSI = 39.2% − chr3:49470895−49471074)
ss3: 7.79
ss5: 0.88
U
C
G
A
del1
del3
del6
del21
UUCCGUCAUCCUUACUUGAGCUCCUUUGGAAGCUUGUUUACUUACUACCUUUUCACUUUGUAUUUGGCAGACCUGGUUCACCUGCGAGUGUAUUUGGAUAGUGUCUCCACAGAAAAUGUCGUCUCGGCUUGGAAUCUUUUUGCCAUAAAUUCUAAGUUAGUUUCCUACAAACUCAACUGG
Intron
Exon
Intron
−mean(ΔLogitPSI_sub)
6
4
2
0
−2

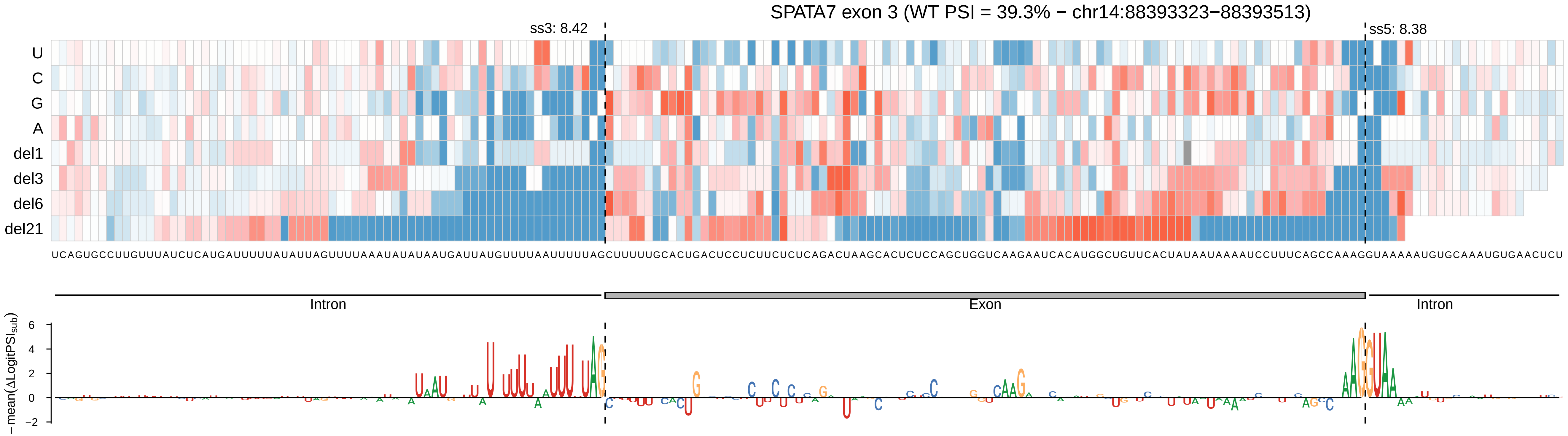

SPATA7 exon 3 (WT PSI = 39.3% − chr14:88393323−88393513)
ss3: 8.42
ss5: 8.38
U
C
G
A
del1
del3
del6
del21
UCAGUGCCUUGUUUAUCUCAUGAUUUUUAUAUUAGUUUUAAAUAUAUAAUGAUUAUGUUUUAAUUUUUAGCUUUUUGCACUGACUCCUCUUCUCUCAGACUAAGCACUCUCCAGCUGGUCAAGAAUCACAUGGCUGUUCACUAUAAUAAAAUCCUUUCAGCCAAAGGUAAAAAUGUGCAAAUGUGAACUCU
Intron
Exon
Intron
− mean(ΔLogitPSI_sub)
6
4
2
0
−2

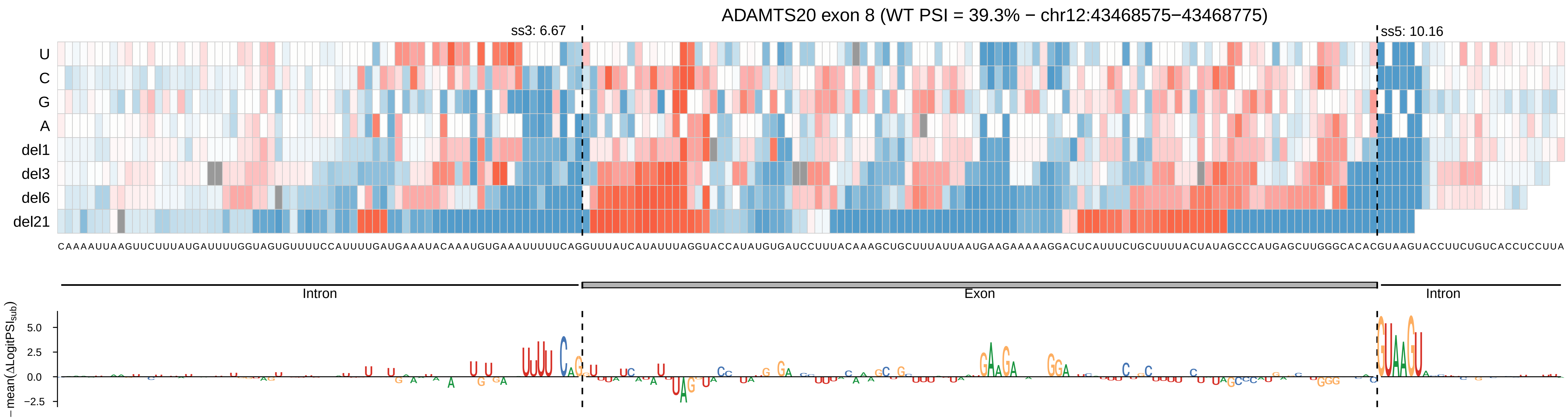

ADAMTS20 exon 8 (WT PSI = 39.3% − chr12:43468575−43468775)
ss3: 6.67
ss5: 10.16
U
C
G
A
del1
del3
del6
del21
CAAAAUUAAGUUCUUUAUGAUUUUGGUAGUGUUUUCCAUUUUGAUGAAAAUACAAAUGUGAAAUUUUUCAGGUUUAUCAUAUUUAGGUACCAUAUGUGAUCCUUUACAAAGCUGCUUUAUUAAUGAAGAAAAAGGACUCAUUUCUGCUUUUACUAUAGCCCAUGAGCUUGGGCACACGUAAGUACCUUCUGUCACCUCCUUA
Intron
Exon
Intron
−mean(ΔLogitPSI_sub)
5.0
2.5
0.0
−2.5

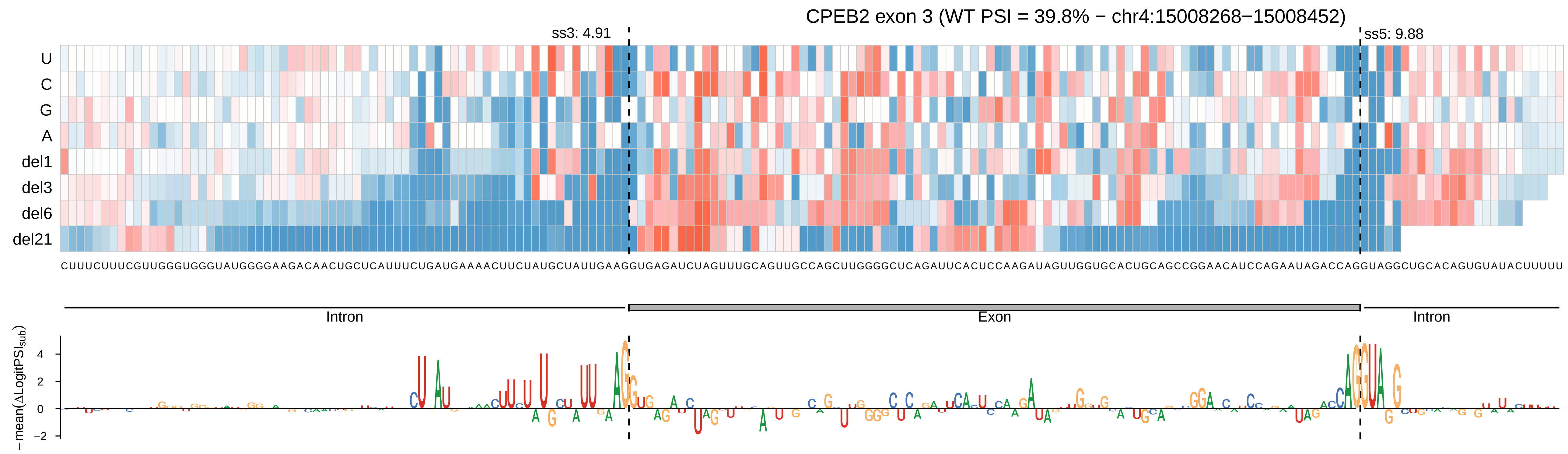
CPEB2 exon 3 (WT PSI = 39.8% − chr4:15008268−15008452)
ss3: 4.91
ss5: 9.88
U
C
G
A
del1
del3
del6
del21
Intron
Exon
Intron
−mean(ΔLogitPSI_sub)
4
2
0
−2

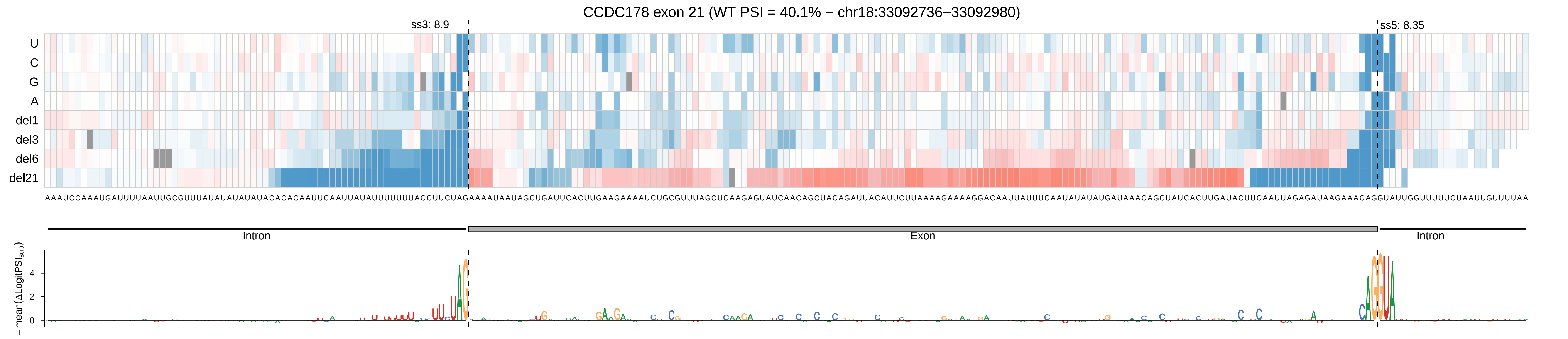
CCDC178 exon 21 (WT PSI = 40.1% − chr18:33092736−33092980)
ss3: 8.9
ss5: 8.35
U
C
G
A
del1
del3
del6
del21
Intron
Exon
Intron
−mean(ΔLogitPSIsub)
4
2
0

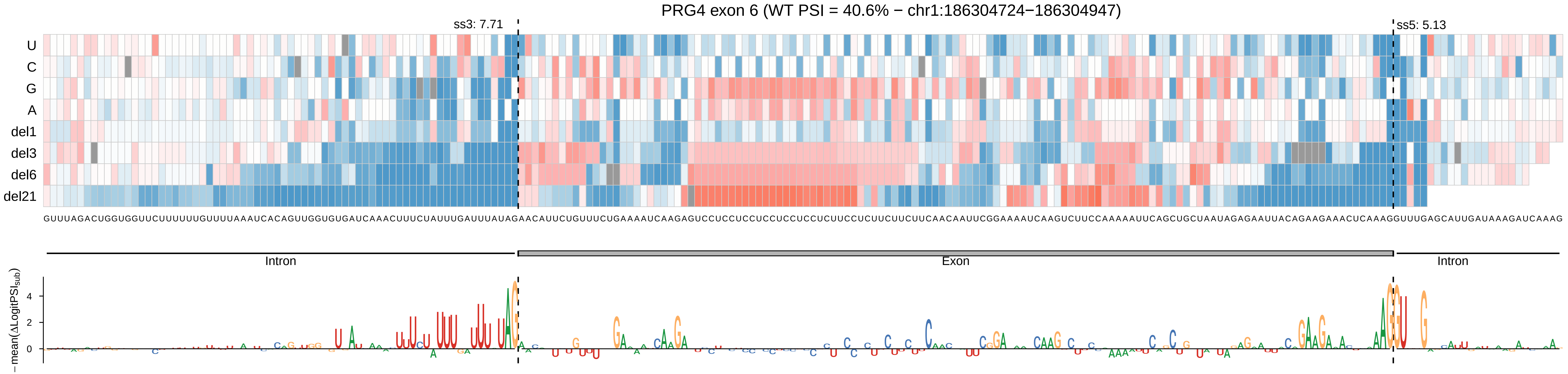

PRG4 exon 6 (WT PSI = 40.6% − chr1:186304724−186304947)
ss3: 7.71
ss5: 5.13
U
C
G
A
del1
del3
del6
del21
GUUUAGACUGGUGGUUCUUUUUUGUUUUAAAUCACAGUUGGUGUGAUCAAACUUUCUAUUUGAUUUAUAGAACAUUCUGUUUCUGAAAAUCAAGAGUCCUCCUCCUCCUCCUCCUCUUCCUCUUCUUCUUCAACAAUUCGGAAAAUCAAGUCUUCCAAAAAUUCAGCUGCUAAUAGAGAAUUACAGAAGAAACUCAAAGGUUUGAGCAUUGAUAAAGAUCAAAG
Intron
Exon
Intron
−mean(ΔLogitPSI_sub)
4
2
0

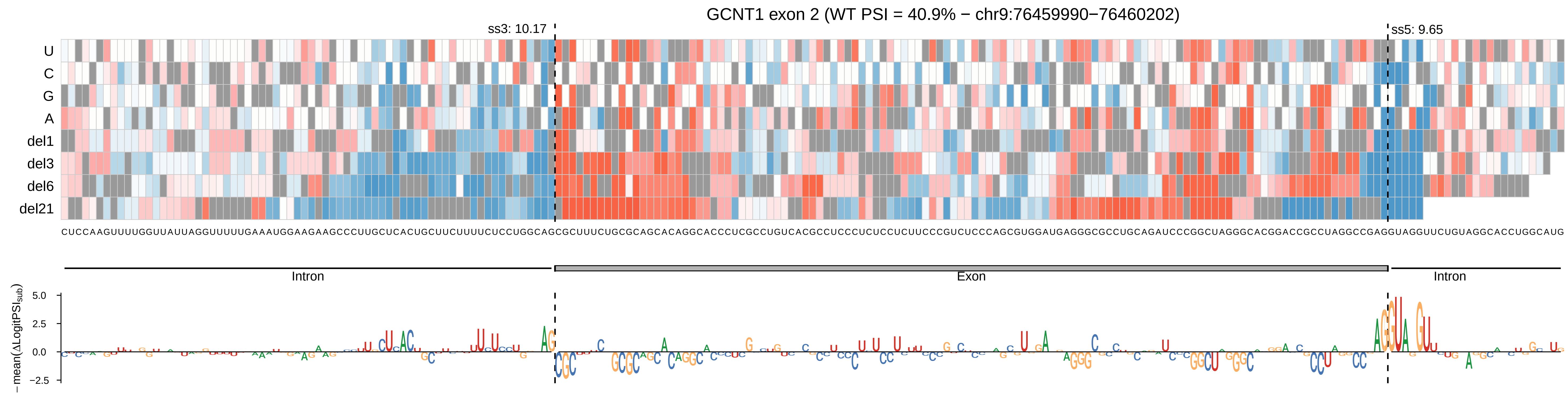
GCNT1 exon 2 (WT PSI = 40.9% − chr9:76459990−76460202)
ss3: 10.17
ss5: 9.65
U
C
G
A
del1
del3
del6
del21
CUCCAAGUUUUGGUUAUUAGGUUUUUGAAAUGGAAGAAGCCCUUGCUCACUGCUUCUUUUCUCCUGGCAGCGCUUUCUGCGCAGCACAGGCACCCUCGCCUGUCACGCCUCCCCUCUCCUCUUCCCGUCUCCCAGCGUGGAUGAGGGCGCCUGCAGAUCCCGGCUAGGGCACGGACCGCCUAGGCCGAGGUAGGUUCUGUAGGCACCUGGCAUG
Intron
Exon
Intron
5.0
2.5
0.0
−2.5
−mean(ΔLogitPSI_sub)

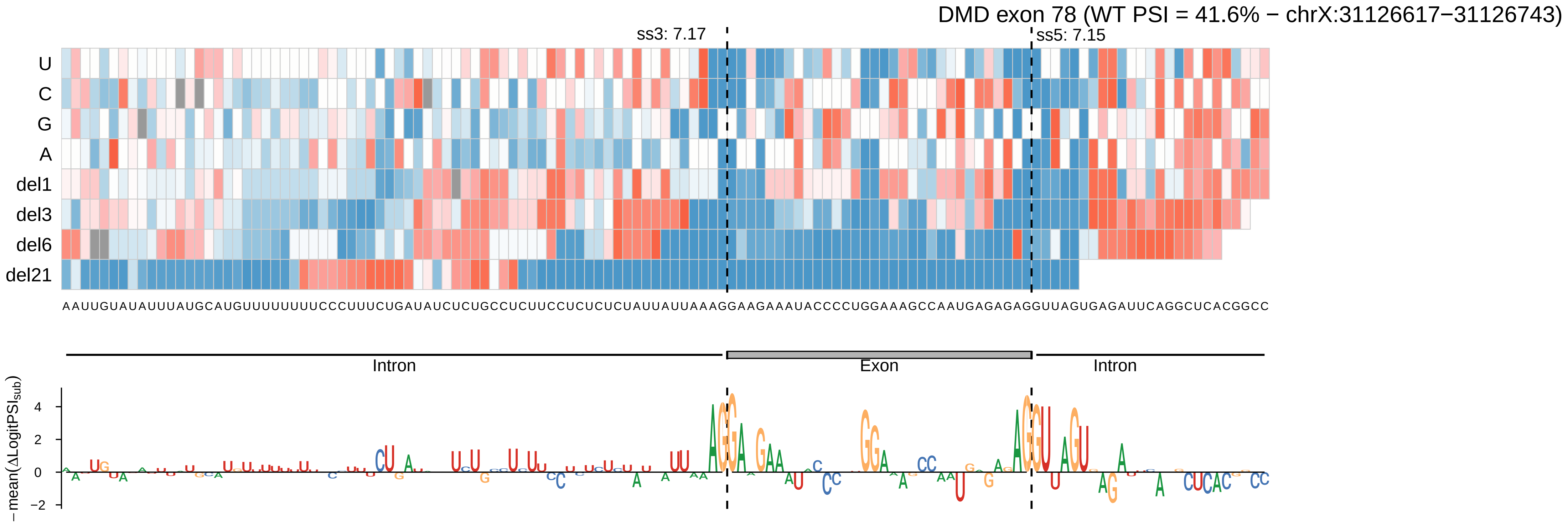
DMD exon 78 (WT PSI = 41.6% − chrX:31126617−31126743)
ss3: 7.17
ss5: 7.15
U
C
G
A
del1
del3
del6
del21
AAUUGUAUAUUUAUGCAUGUUUUUUUUCCCUUUCUGAUAUCUCUGCCUCUUCCUCUCUCUAUUAUUAAAGGAAGAAAUACCCCUGGAAAGCCAAUGAGAGAGGUUAGUGAGAUUCAGGCUCACGGCC
Intron
Exon
Intron
− mean(ΔLogitPSI_sub)
4
2
0
−2

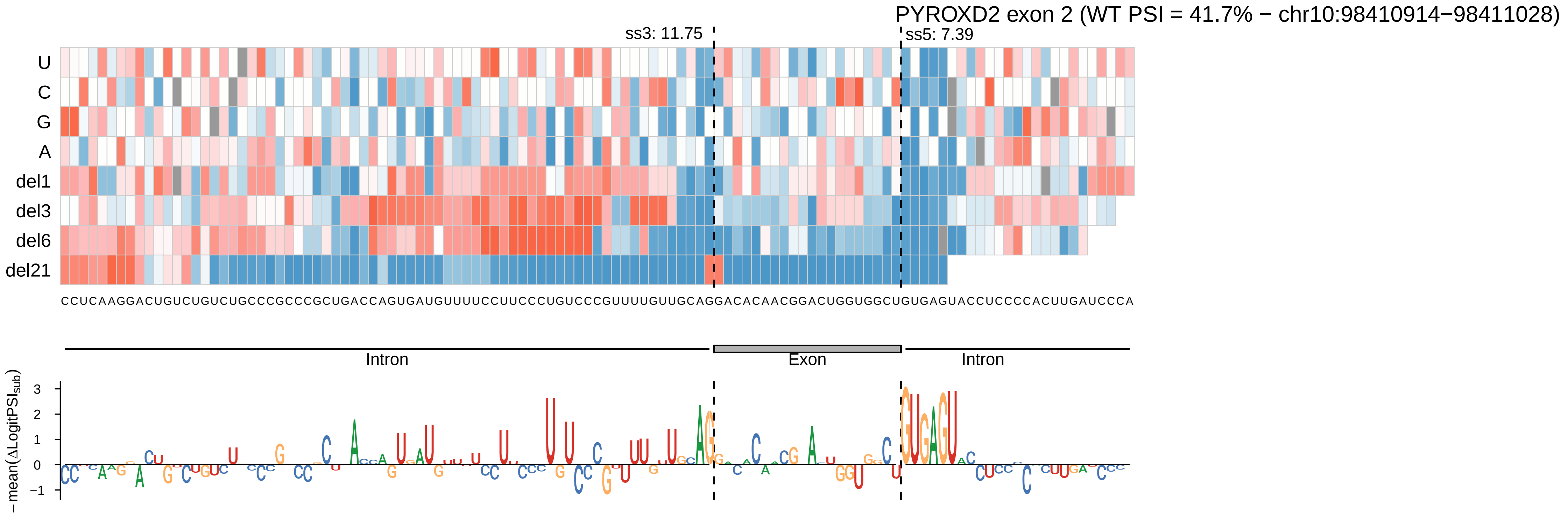
PYROXD2 exon 2 (WT PSI = 41.7% − chr10:98410914−98411028)
ss3: 11.75
ss5: 7.39
U
C
G
A
del1
del3
del6
del21
CCUCAAGGACUGUCUGUCUGCCCGCCCGCUGACCAGUGAUGUUUUCCUUCCCUGUCCCGUUUUGUUGCAGGACACAACGGACUGGUGGCUGUGAGUACCUCCCCACUUGAUCCCA
Intron
Exon
Intron
− mean(ΔLogitPSI sub)

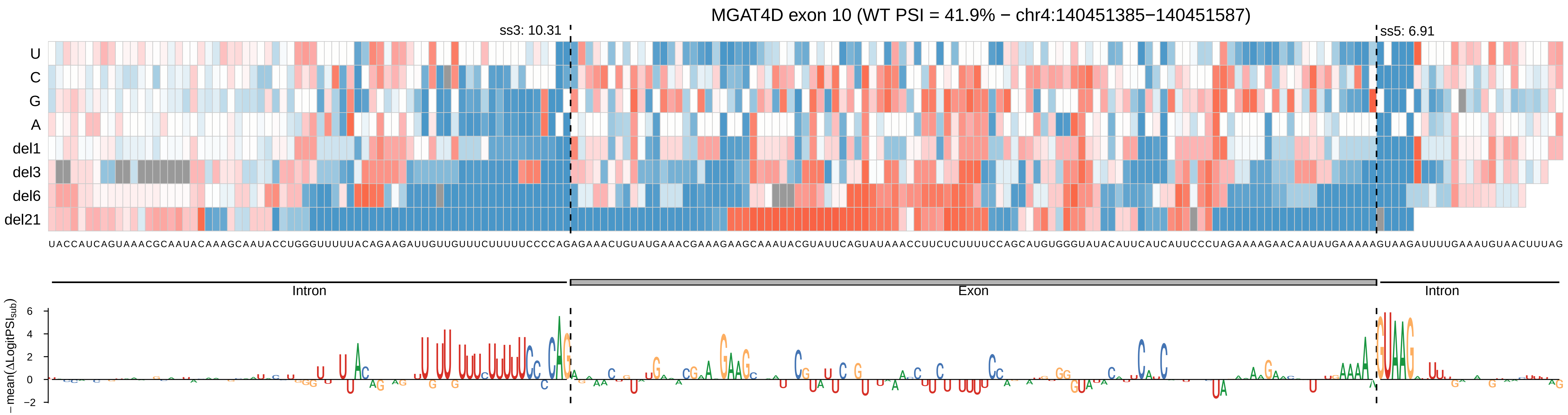
MGAT4D exon 10 (WT PSI = 41.9% − chr4:140451385−140451587)
ss3: 10.31
ss5: 6.91
U
C
G
A
del1
del3
del6
del21
UACCAUCAGUAAACGCAAUACAAAGCAAUACCUGGGUUUUUACAGAAGAUUGUUGUUUCUUUUUCCCCAGAGAAACUGUAUGAAACGAAAGAAGCAAAUACGUAUUCAGUAUAAACCUUCUCUUUUCCAGCAUGUGGGUAUACAUUCAUCAUUCCCUAGAAAAGAACAAUAUGAAAAAGUAAGAUUUUGAAAUGUAACUUUAG
Intron
Exon
Intron
− mean(ΔLogitPSI_sub)
6
4
2
0
−2

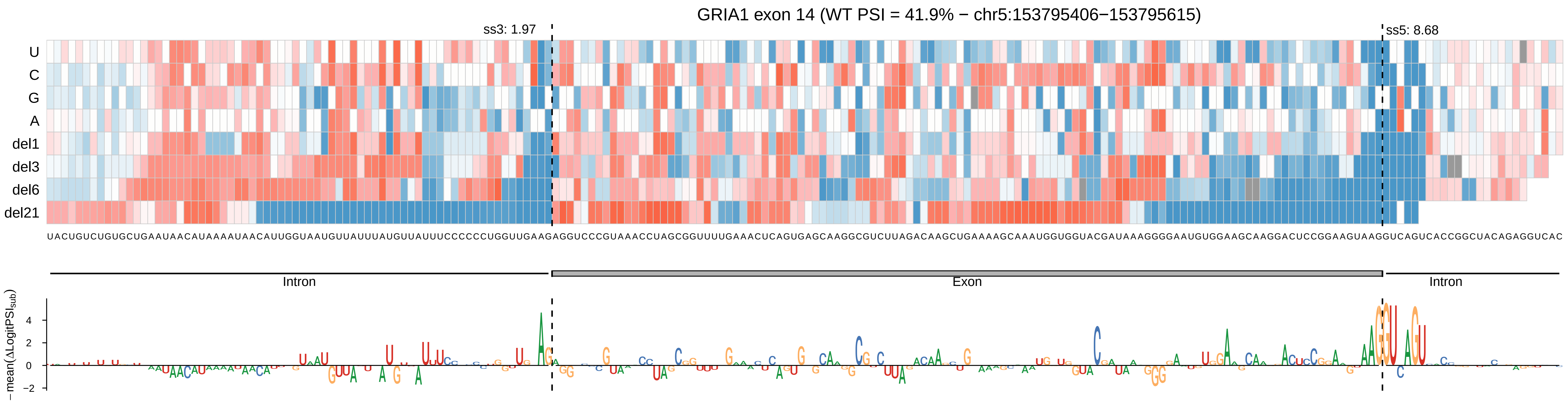
GRIA1 exon 14 (WT PSI = 41.9% − chr5:153795406−153795615)
ss3: 1.97
ss5: 8.68
U
C
G
A
del1
del3
del6
del21
UACUGUCUGUGCUGAAUAACAUAAAAUAACAUUGGUAAUGUUAUUUAUGUUAUUUCCCCCCUGGUUGAAGAGGUCCCGUAAACCUAGCGGUUUUGAAACUCAGUGAGCAAGGCGUCUUAGACAAGCUGAAAAGCAAAUGGUGGUACGAUAAAGGGGAAUGUGGAAGCAAGGACUCCGGAAGUAAGGUCAGUCACCGGCUACAGAGGUCAC
Intron
Exon
Intron
− mean(ΔLogitPSI_sub)
4
2
0
−2

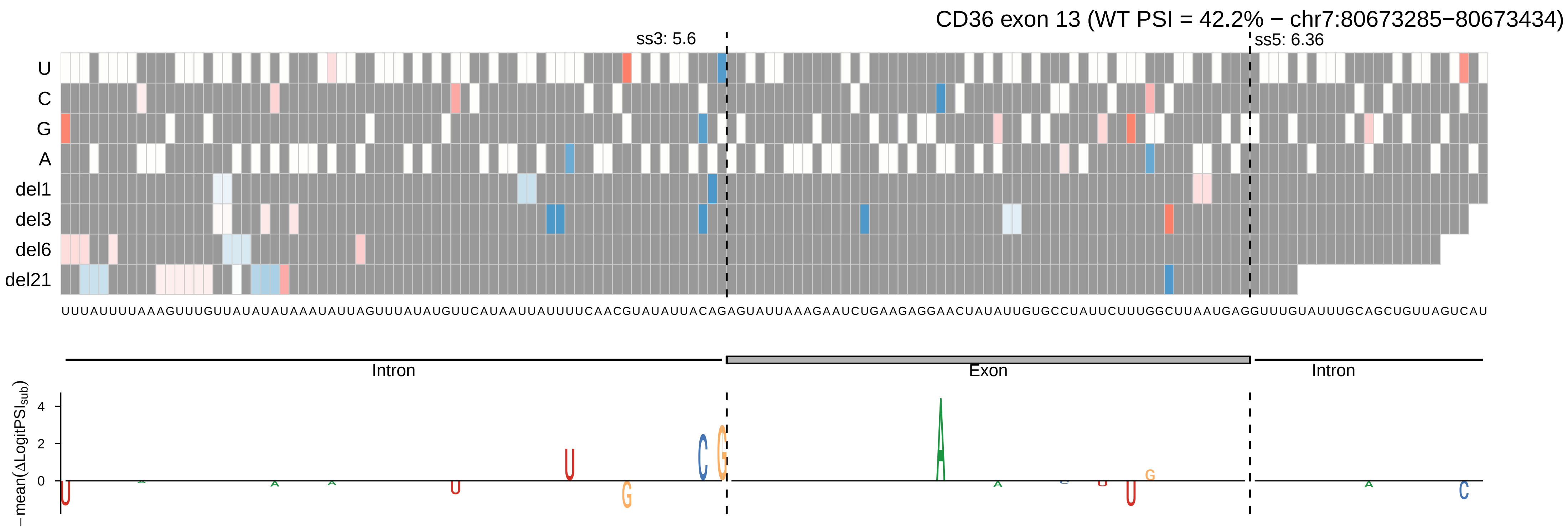
CD36 exon 13 (WT PSI = 42.2% − chr7:80673285−80673434)
ss3: 5.6
ss5: 6.36
U
C
G
A
del1
del3
del6
del21
UUUAUUUUAAAGUUUGUUAUAUAUAAAUAUUAGUUUAUAUGUUCAUAAUUAUUUUCAACGUAUAUUACAGAGUAUUAAAGAAUCUGAAGAGGAACUAUAUUGUGCCUAUUCUUUGGCUUAAUGAGGUUUGUAUUUGCAGCUGUUAGUCAU
Intron
Exon
Intron
−mean(ΔLogitPSI_sub)
4
2
0

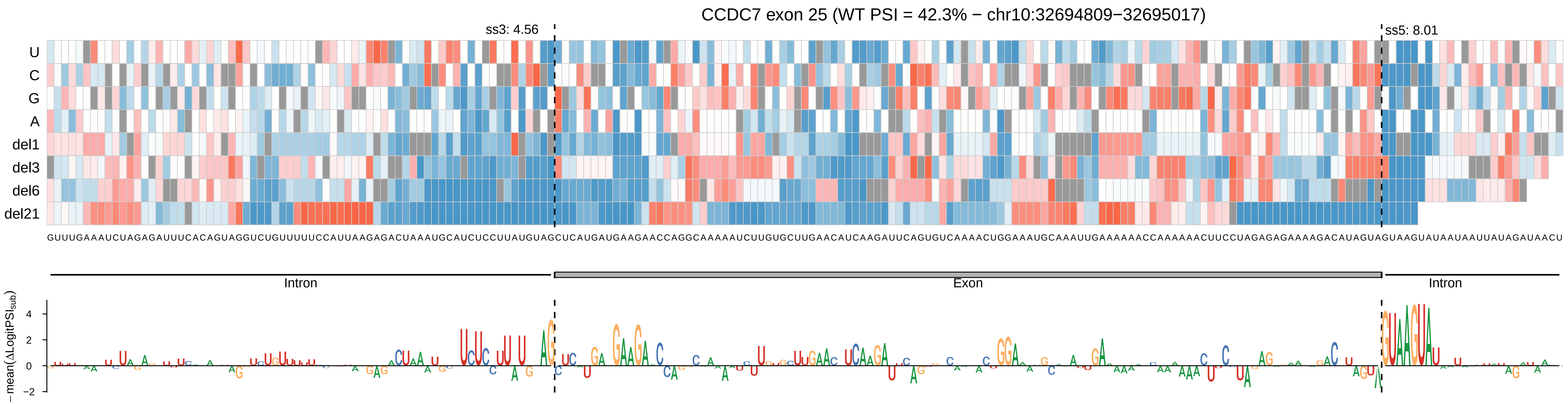
CCDC7 exon 25 (WT PSI = 42.3% − chr10:32694809−32695017)
ss3: 4.56
ss5: 8.01
U
C
G
A
del1
del3
del6
del21
GUUUGAAAUCUAGAGAUUUCACAGUAGGUCUGUUUUUCCAUUAAGAGACUAAAUGCAUCUCCUUAUGUAGCUCAUGAUGAAGAACCAGGCAAAAAUCUUGUGCUUGAACAUCAAGAUUCAGUGUCAAAACUGGAAAUGCAAAUUGAAAAAACCAAAAAACUUCCUAGAGAGAAAAGACAUAGUAGUAAGUAUAAUAAUUAUAGAUAACU
Intron
Exon
Intron
−mean(ΔLogitPSI_sub)
4
2
0
−2

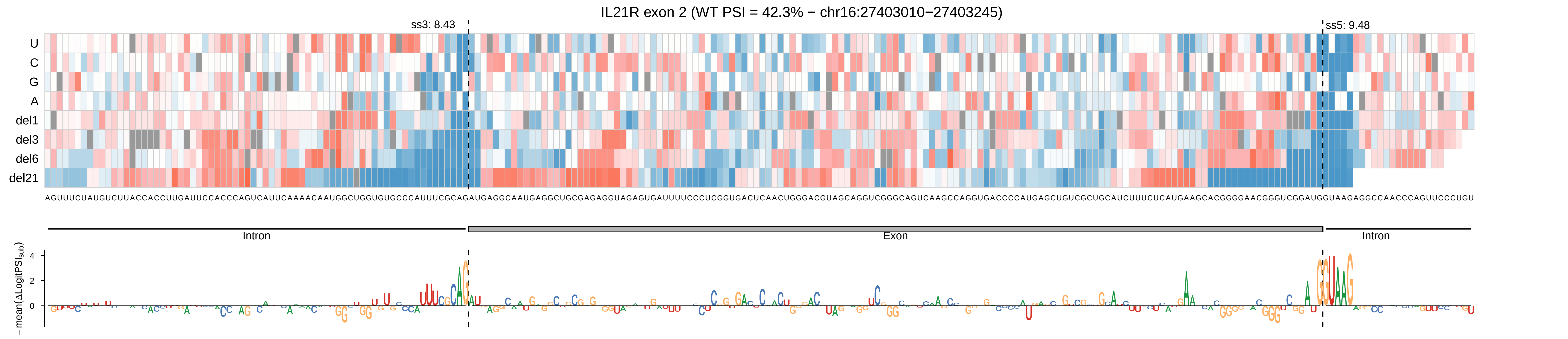

IL21R exon 2 (WT PSI = 42.3% − chr16:27403010−27403245)
ss3: 8.43
ss5: 9.48
U
C
G
A
del1
del3
del6
del21
Intron
Exon
Intron
−mean(ΔLogitPSI_sub)

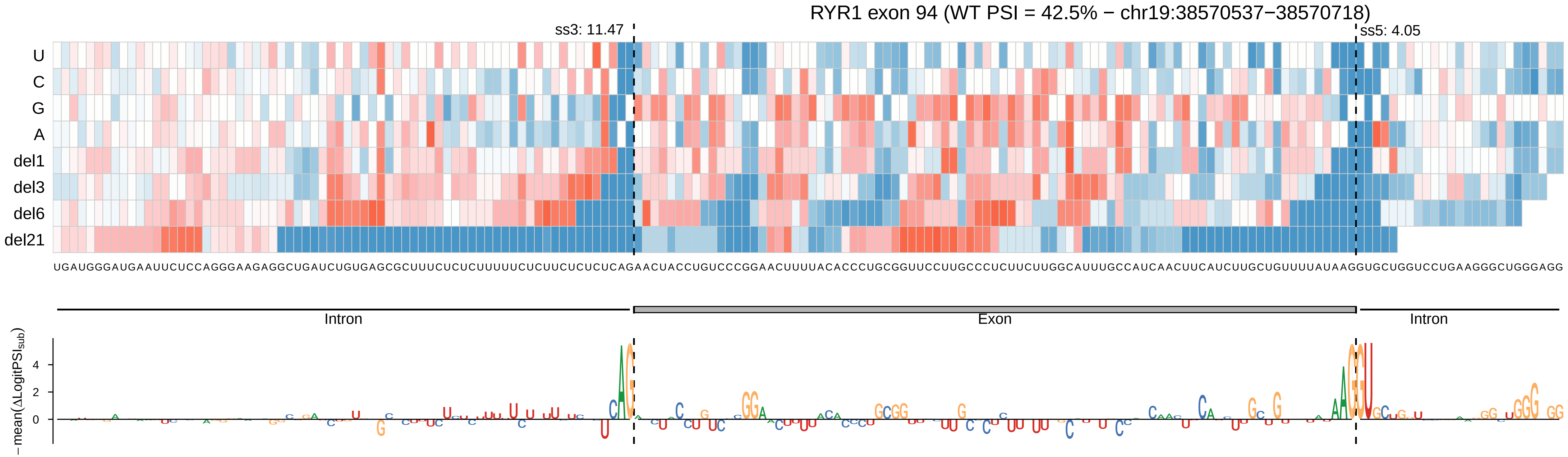

RYR1 exon 94 (WT PSI = 42.5% − chr19:38570537−38570718)
ss3: 11.47
ss5: 4.05
U
C
G
A
del1
del3
del6
del21
UGAUGGGAUGAAUUCUCCAGGGAAGAGGCUGAUCUGUGAGCGCUUUCUCUCUUUUUCUCUUCUCUCUCAGAACUACCUGUCCCGGAACUUUUACACCCUGCGGUUCCUUGCCCUCUUCUUGGCAUUUGCCAUCAACUUCAUCUUGCUGUUUUAUAAGGUGCUGGUCCUGAAGGGCUGGGAGG
Intron
Exon
Intron
−mean(ΔLogitPSI_sub)
4
2
0

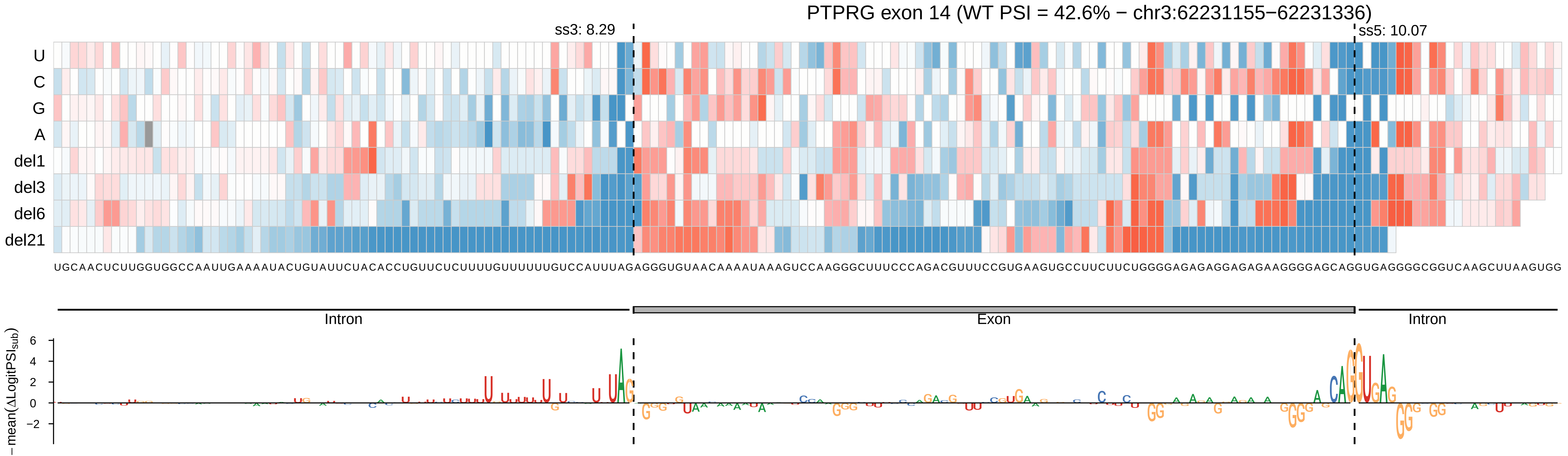

PTPRG exon 14 (WT PSI = 42.6% − chr3:62231155−62231336)
ss3: 8.29
ss5: 10.07
U
C
G
A
del1
del3
del6
del21
UGCAACUCUUGGUGGCCAAUUGAAAAUACUGUAUUCUACACCUGUUCUCUUUUGUUUUUUGUCCAUUUAGAGGGUGUAACAAAAUAAAGUCCAAGGGCUUUCCCAGACGUUUCCGUGAAGUGCCUUCUUCUGGGGAGAGAGGAGAGAAGGGGAGCAGGUGAGGGGCGGUCAAGCUUAAGUGG
Intron
Exon
Intron
−mean(ΔLogitPSI_sub)
6
4
2
0
−2

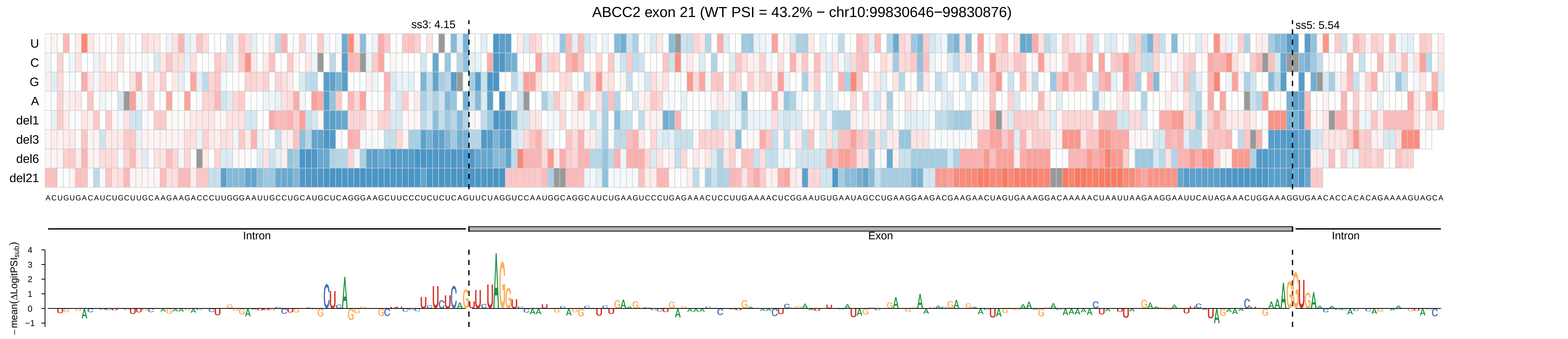
ABCC2 exon 21 (WT PSI = 43.2% − chr10:99830646−99830876)
ss3: 4.15
ss5: 5.54
U
C
G
A
del1
del3
del6
del21
Intron
Exon
Intron
−mean(ΔLogitPSI_sub)

NEK10 exon 11 (WT PSI = 43.5% − chr3:27143420−27143544)

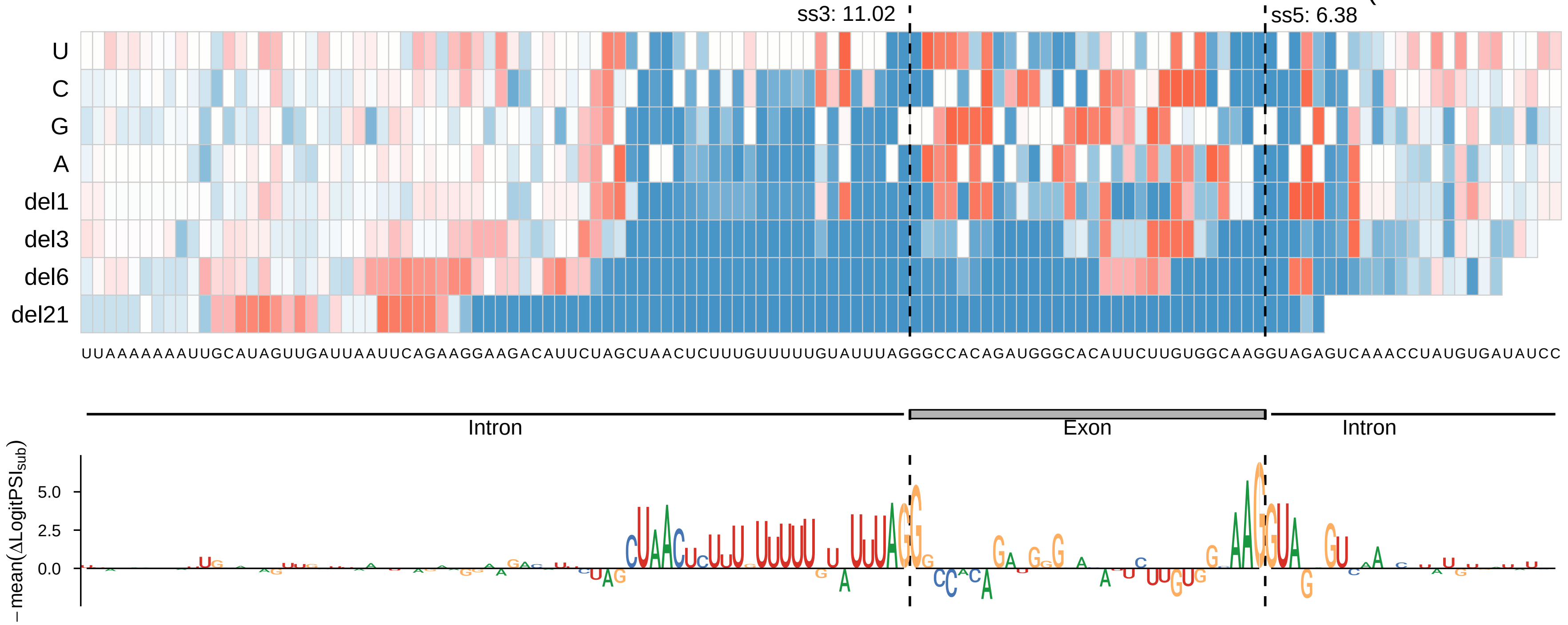

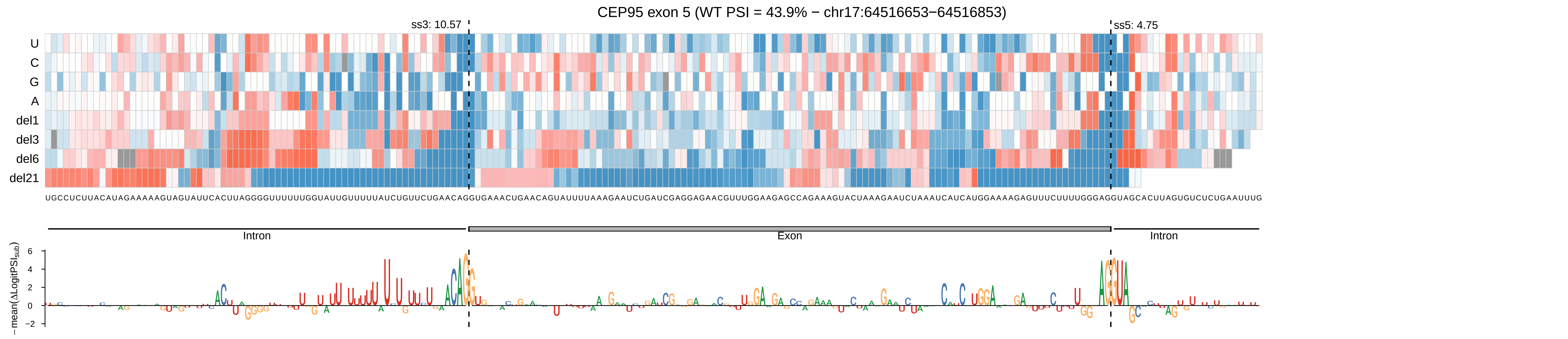

CEP95 exon 5 (WT PSI = 43.9% − chr17:64516653−64516853)
ss3: 10.57
ss5: 4.75
U
C
G
A
del1
del3
del6
del21
UGCCUCUUACAUAGAAAAAGUAGUAUUCACUUAGGGGUUUUUUGGUAUUGUUUUUAUCUGUUCUGAACAGGUGAAACUGAACAGUAUUUUAAAGAAUCUGAUCGAGGAGAACGUUUGGAAGAGCCAGAAAGUACUAAAGAAUCUAAAUCAUCAUGGAAAAGAGUUUCUUUUGGGAGGUAGCACUUAGUGUCUCUGAAUUUG
Intron
Exon
Intron
−mean(ΔLogitPSI_sub)
6
4
2
0
−2

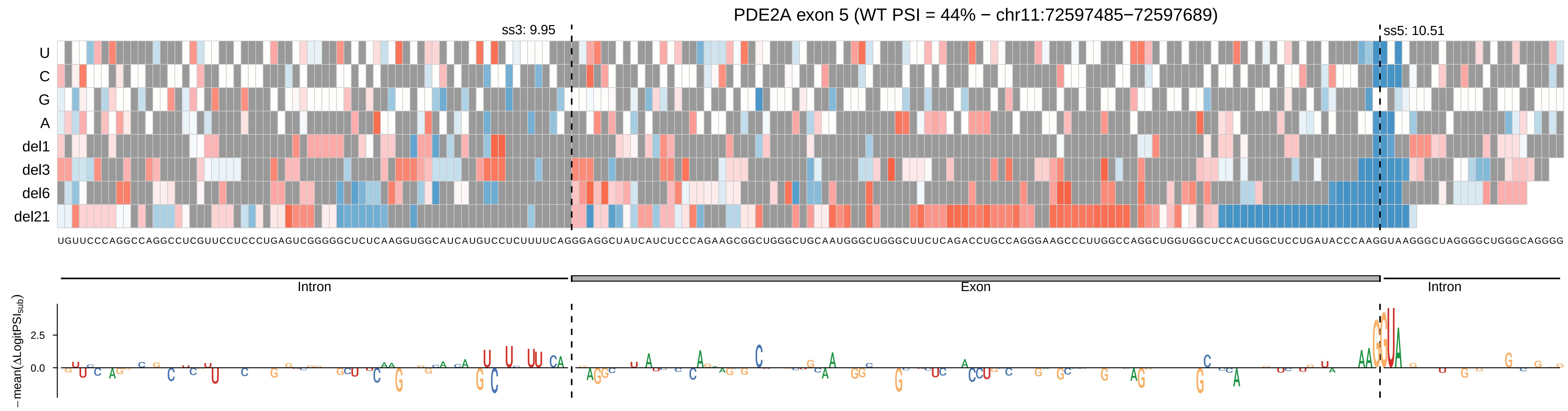

PDE2A exon 5 (WT PSI = 44% − chr11:72597485−72597689)
ss3: 9.95
ss5: 10.51
U
C
G
A
del1
del3
del6
del21
UGUUCCCAGGCCAGGCCUCGUUCCUCCCUGAGUCGGGGGCUCUCAAGGUGGCAUCAUGUCCUCUUUUCAGGGAGGCUAUCAUCUCCCAGAAGCGGCUGGGCUGCAAUGGGCUGGGCUUCUCAGACCUGCCAGGGAAGCCCUUGGCCAGGCUGGUGGCUCCACUGGCUCCUGAUACCCAAGGUAAGGGCUAGGGGCUGGGCAGGGG
Intron
Exon
Intron
− mean(ΔLogitPSI_sub)
2.5
0.0

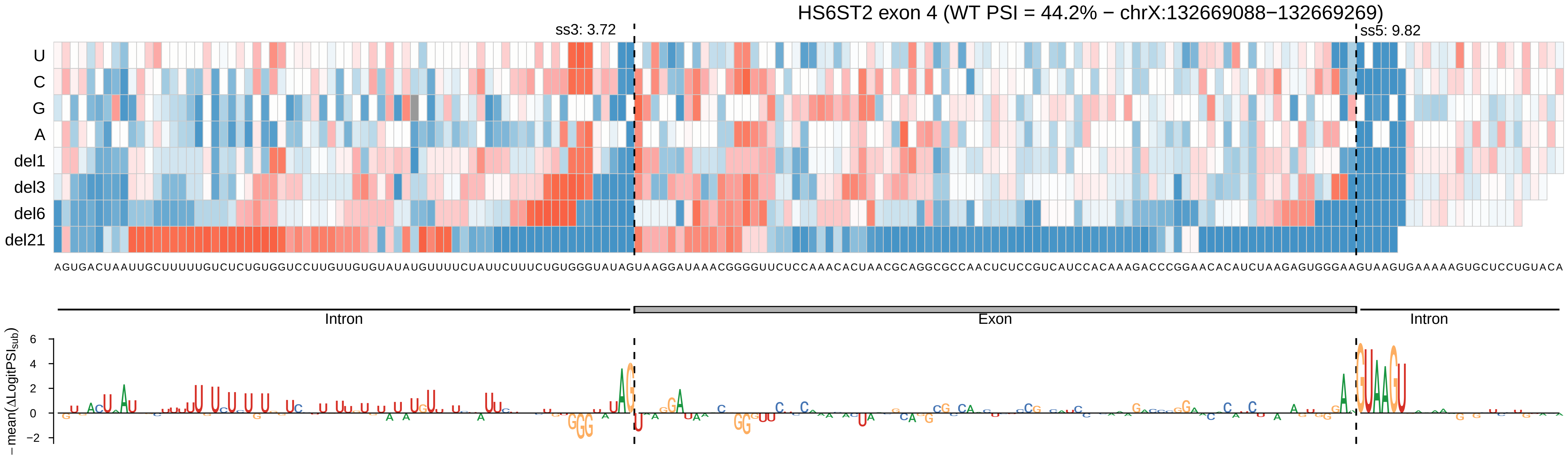
HS6ST2 exon 4 (WT PSI = 44.2% − chrX:132669088−132669269)
ss3: 3.72
ss5: 9.82
U
C
G
A
del1
del3
del6
del21
AGUGACUAAUUGCUUUUUGUCUCUGUGGUCCUUGUUGUGUAUAUGUUUUCUAUUCUUUCUGUGGGUAUAGUAAGGAUAAACGGGGUUCUCCAAACACUAACGCAGGCGCCAACUCUCCGUCAUCCACAAAGACCCGGAACACAUCUAAGAGUGGGAAGUAAGUGAAAAAGUGCUCCUGUACA
Intron
Exon
Intron
−mean(ΔLogitPSI_sub)
6
4
2
0
−2

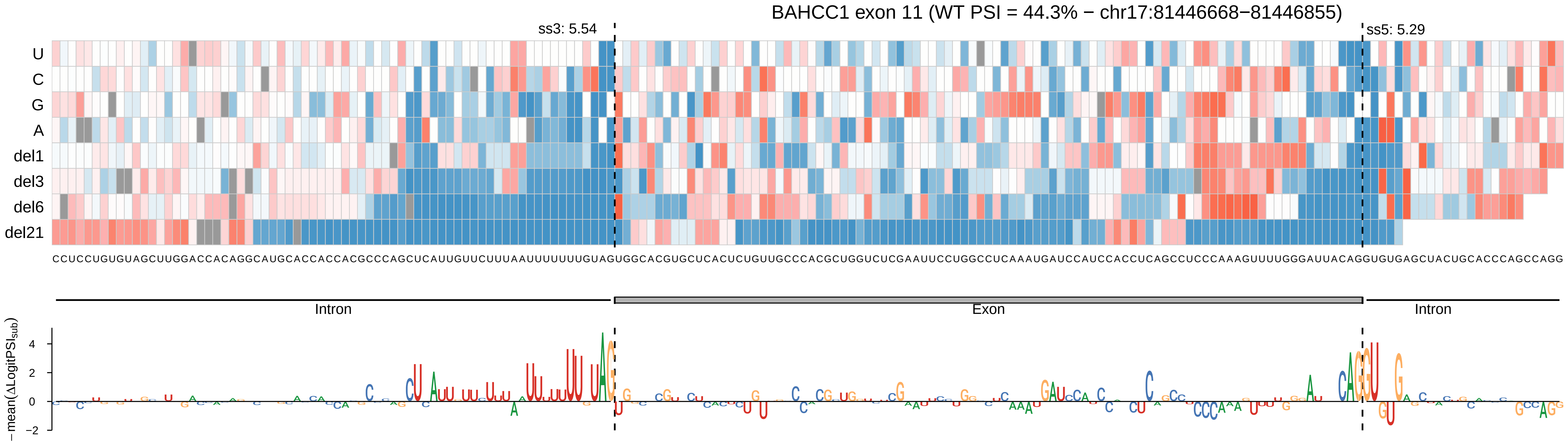

BAHCC1 exon 11 (WT PSI = 44.3% − chr17:81446668−81446855)
ss3: 5.54
ss5: 5.29
U
C
G
A
del1
del3
del6
del21
CCUCCUGUGUAGCUUGGACCACAGGCAUGCACCACCACGCCCAGCUCAUUGUUCUUUAAUUUUUUUGUAGUGGCACGUGCUCACUCUGUUGCCCACGCUGGUCUCGAAUUCCUGGCCUCAAAUGAUCCAUCCACCUCAGCCUCCCAAAGUUUUGGGAUUACAGGUGUGAGCUACUGCACCCAGCCAGG
Intron
Exon
Intron
− mean(ΔLogitPSI_sub)
4
2
0
−2

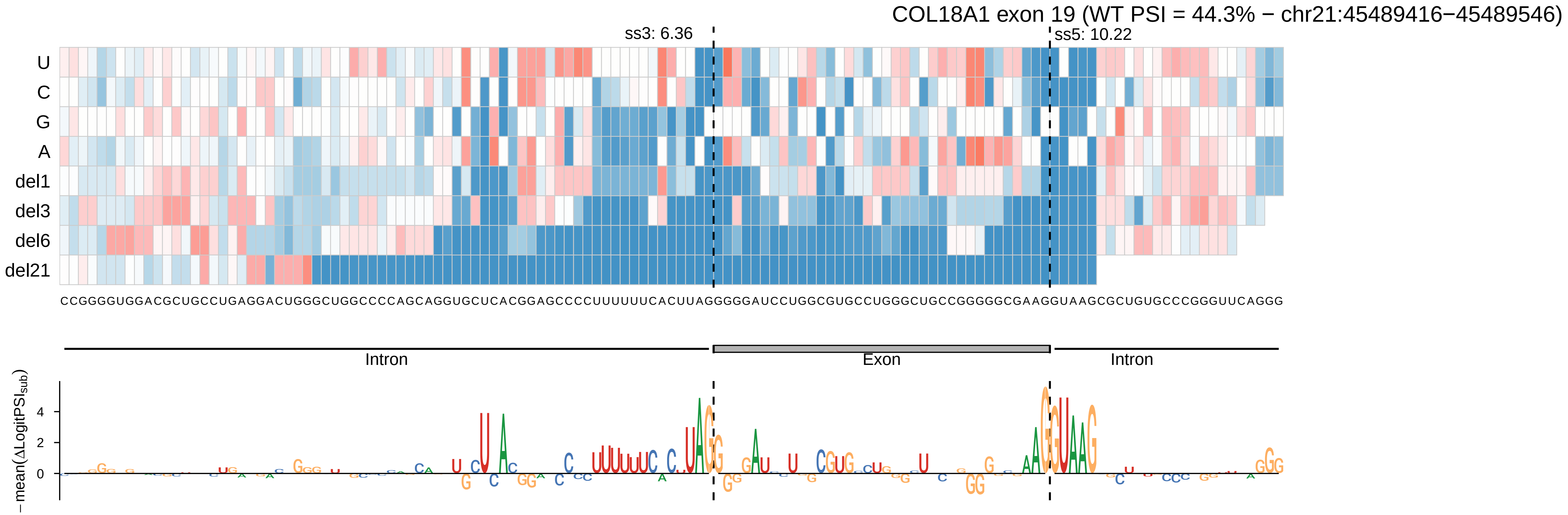

COL18A1 exon 19 (WT PSI = 44.3% − chr21:45489416−45489546)
ss3: 6.36
ss5: 10.22
U
C
G
A
del1
del3
del6
del21
CCGGGGUGGACGCUGCCUGAGGACUGGGCUGGCCCCAGCAGGUGCUCACGGAGCCCCUUUUUUCACUUAGGGGGAUCCUGGCGUGCCUGGGCUGCCGGGGGCGAAGGUAAGCGCUGUGCCCGGGUUCAGGG
Intron
Exon
Intron
− mean(ΔLogitPSI_sub)
4
2
0

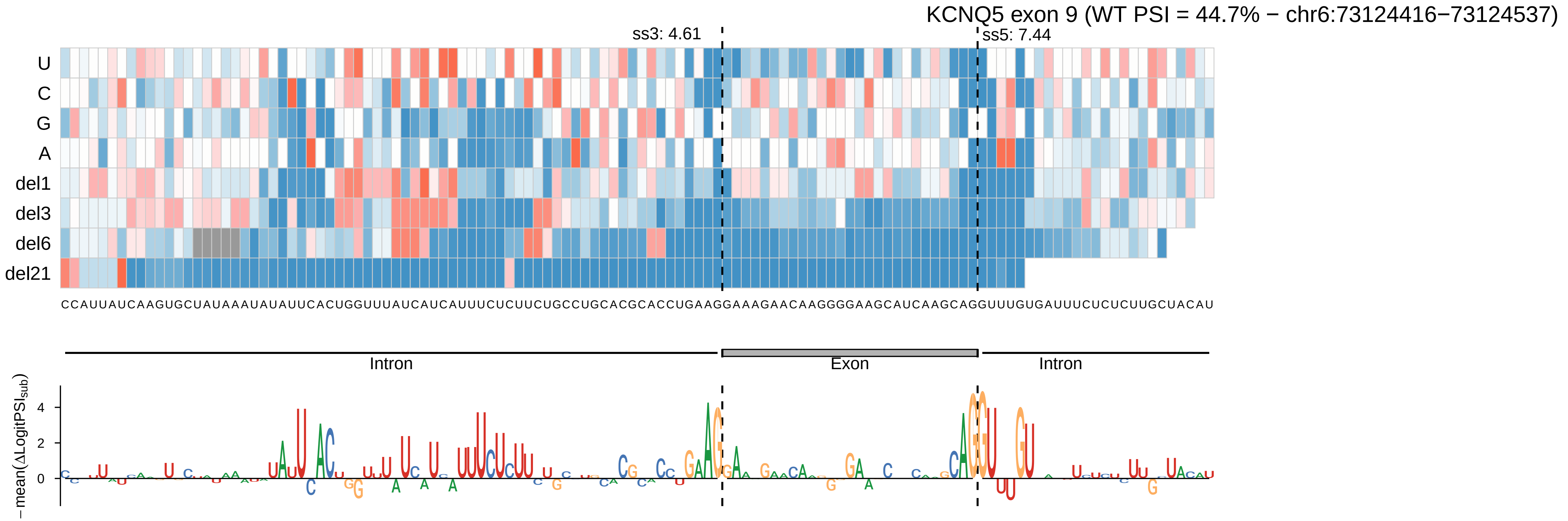

KCNQ5 exon 9 (WT PSI = 44.7% − chr6:73124416−73124537)
ss3: 4.61
ss5: 7.44
U
C
G
A
del1
del3
del6
del21
CCAUUAUCAAGUGCUAUAAAUAUAUUCACUGGUUUAUCAUCAUUUCUCUUCUGCCUGCACGCACCUGAAGGAAAGAACAAGGGGAAGCAUCAAGCAGGUUUGUGAUUUCUCUCUUGCUACAU
Intron
Exon
Intron
−mean(ΔLogitPSI_sub)
4
2
0

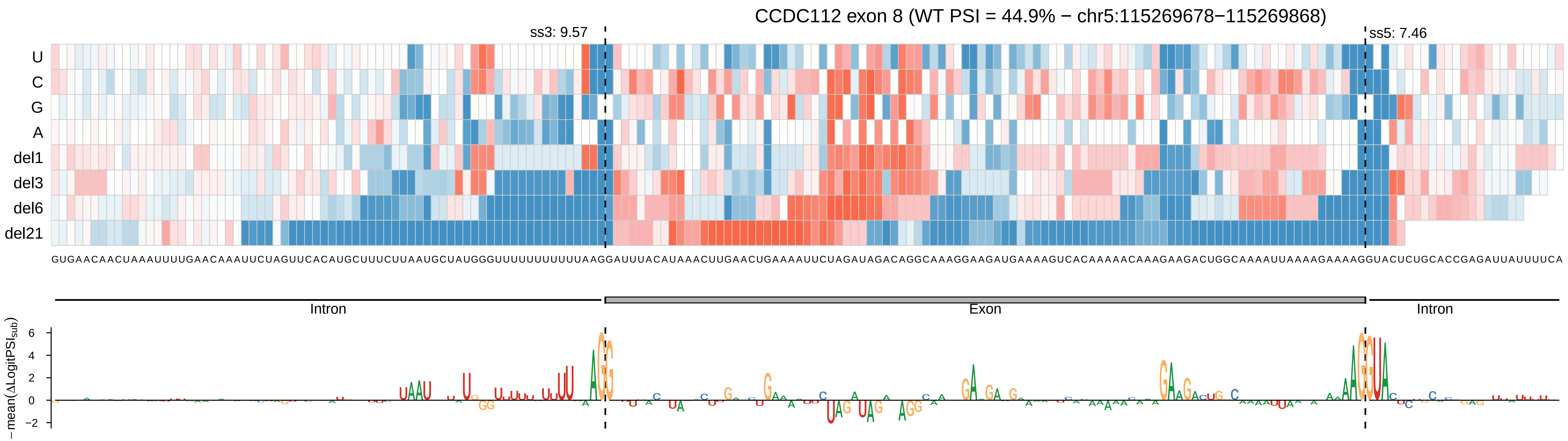
CCDC112 exon 8 (WT PSI = 44.9% − chr5:115269678−115269868)
ss3: 9.57
ss5: 7.46
U
C
G
A
del1
del3
del6
del21
Intron
Exon
Intron
−mean(ΔLogitPSI_sub)

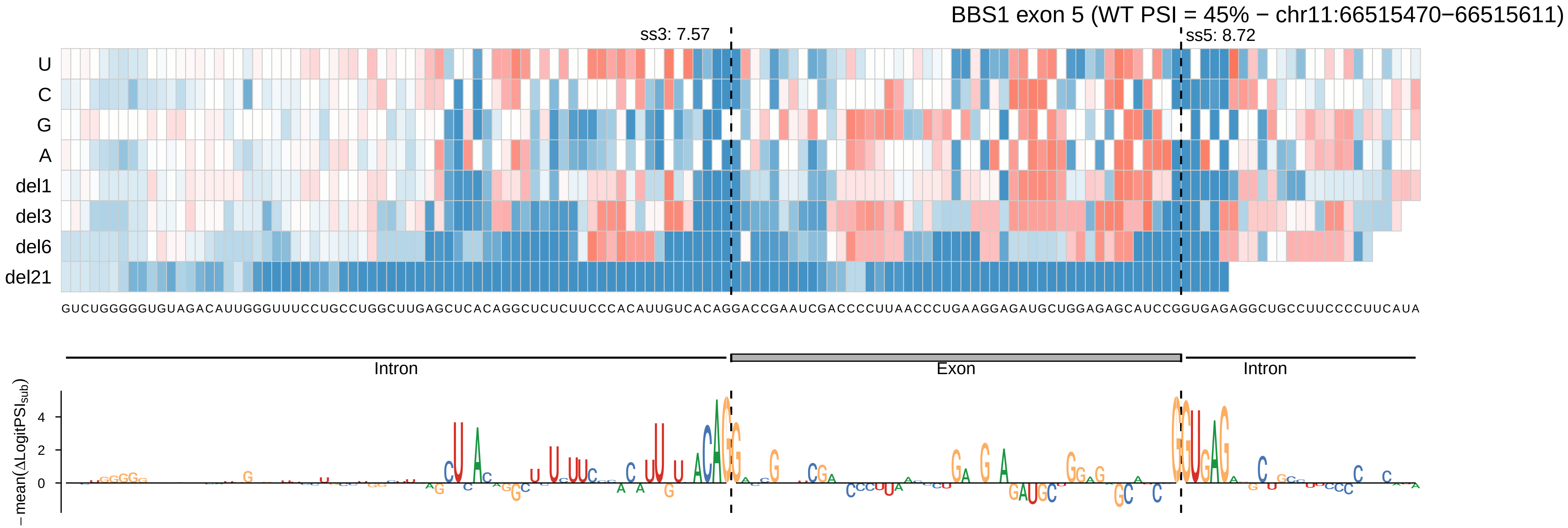

BBS1 exon 5 (WT PSI = 45% − chr11:66515470−66515611)
ss3: 7.57
ss5: 8.72
U
C
G
A
del1
del3
del6
del21
GUCUGGGGGUGUAGACAUUGGGUUUCCUGCCUGGCUUGAGCUCACAGGCUCUCUUCCCACAUUGUCACAGGACCGAAUCGACCCCUUAACCCUGAAGGAGAUGCUGGAGAGCAUCCGGUGAGAGGCUGCCUUCCCCUUCAUA
Intron
Exon
Intron
−mean(ΔLogitPSI_sub)
4
2
0

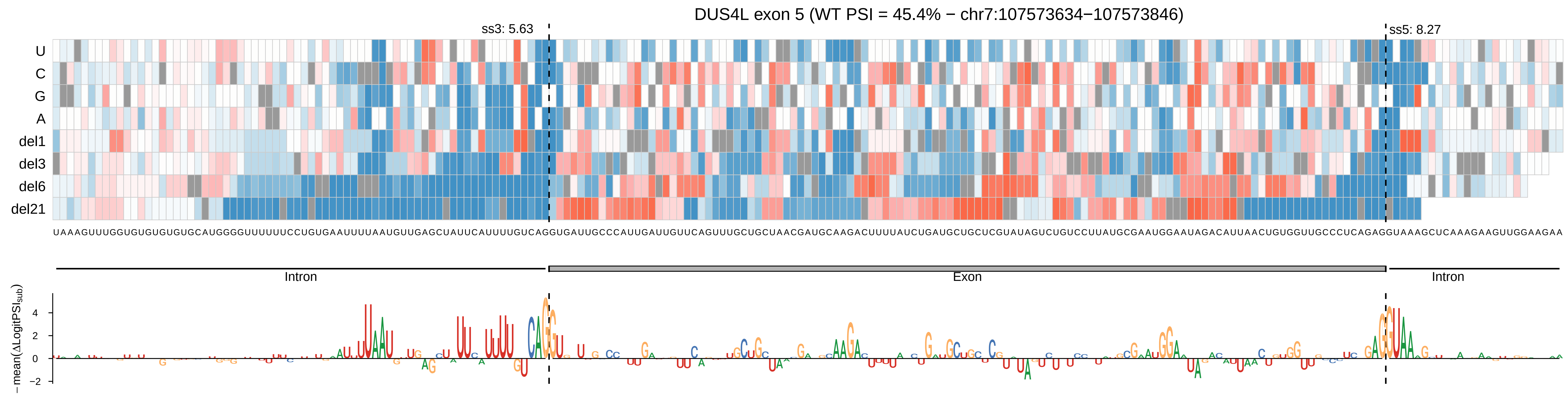

DUS4L exon 5 (WT PSI = 45.4% − chr7:107573634−107573846)
ss3: 5.63
ss5: 8.27
U
C
G
A
del1
del3
del6
del21
UAAAGUUUGGUGUGUGUGUGCAUGGGGUUUUUUCCUGUGAAUUUUAAUGUUGAGCUAUUCAUUUUGUCAGGUGAUUGCCCAUUGAUUGUUCAGUUUGCUGCUAACGAUGCAAGACUUUUAUCUGAUGCUGCUCGUAUAGUCUGUCCUUAUGCGAAUGGAAUAGACAUUAACUGUGGUUGCCCUCAGAGGUAAAGCUCAAAGAAGUUGGAAGAA
Intron
Exon
Intron
−mean(ΔLogitPSI_sub)
4
2
0
−2

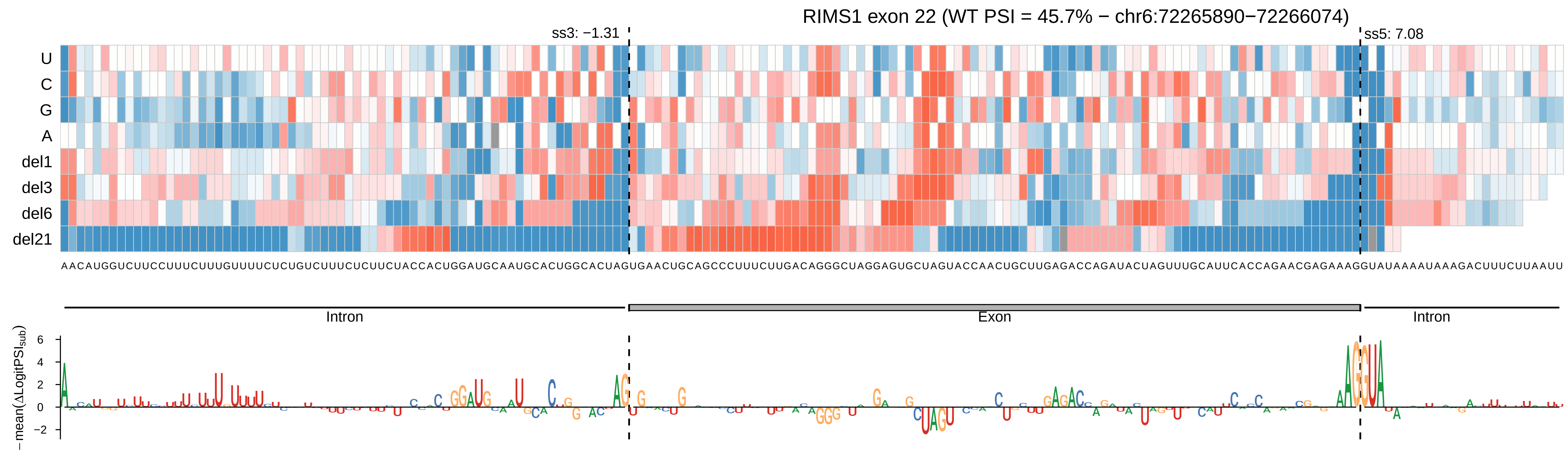

RIMS1 exon 22 (WT PSI = 45.7% − chr6:72265890−72266074)
ss3: −1.31
ss5: 7.08
U
C
G
A
del1
del3
del6
del21
AACAUGGUCUUCCUUUCUUUGUUUUCUCUGUCUUUCUCUUCUACCACUGGAUGCAAUGCACUGGCACUAGUGAACUGCAGCCCUUUCUUGACAGGGCUAGGAGUGCUAGUACCAACUGCUUGAGACCAGAUACUAGUUUGCAUUCACCAGAACGAGAAAGGUAUAAAAUAAAGACUUUCUUAAUU
Intron
Exon
Intron
−mean(ΔLogitPSI_sub)
6
4
2
0
−2

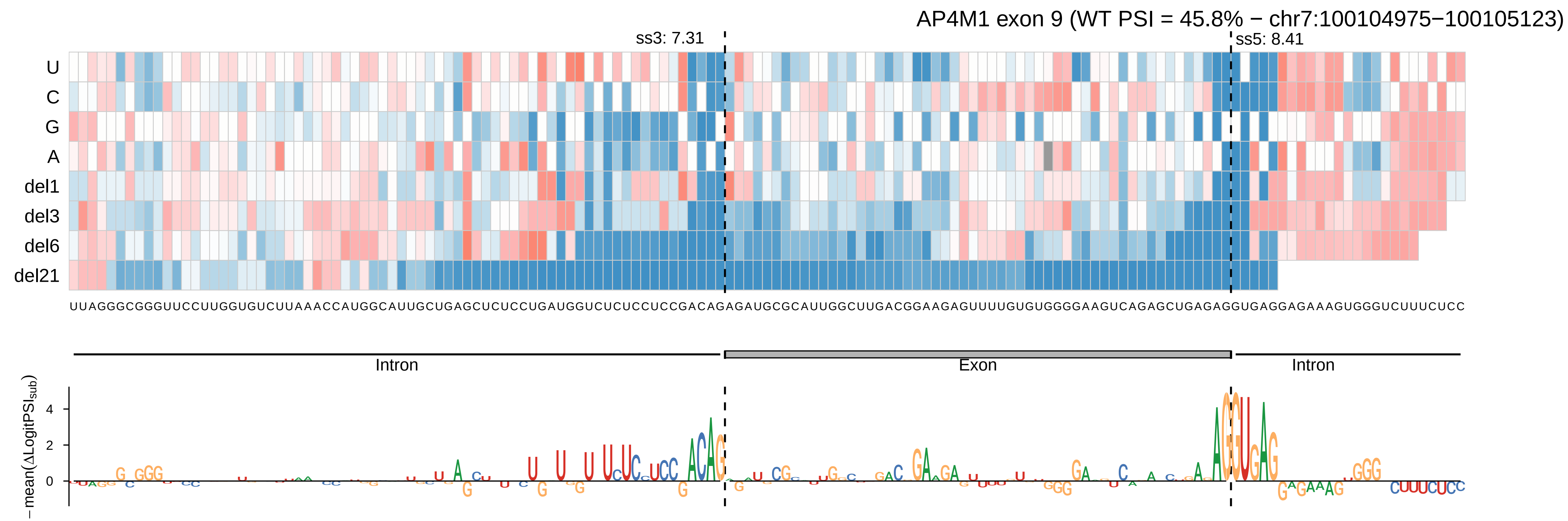
AP4M1 exon 9 (WT PSI = 45.8% − chr7:100104975−100105123)
ss3: 7.31
ss5: 8.41
U
C
G
A
del1
del3
del6
del21
UUAGGGCGGGUUCCUUGGUGUCUUAAACCAUGGCAUUGCUGAGCUCUCCUGAUGGUCUCUCCUCCGACAGAGAUGCGCAUUGGCUUGACGGAAGAGUUUUGUGUGGGGAAGUCAGAGCUGAGAGGUGAGGAGAAAGUGGGUCUUUCUCC
Intron
Exon
Intron
− mean(ΔLogitPSI_sub)
4
2
0

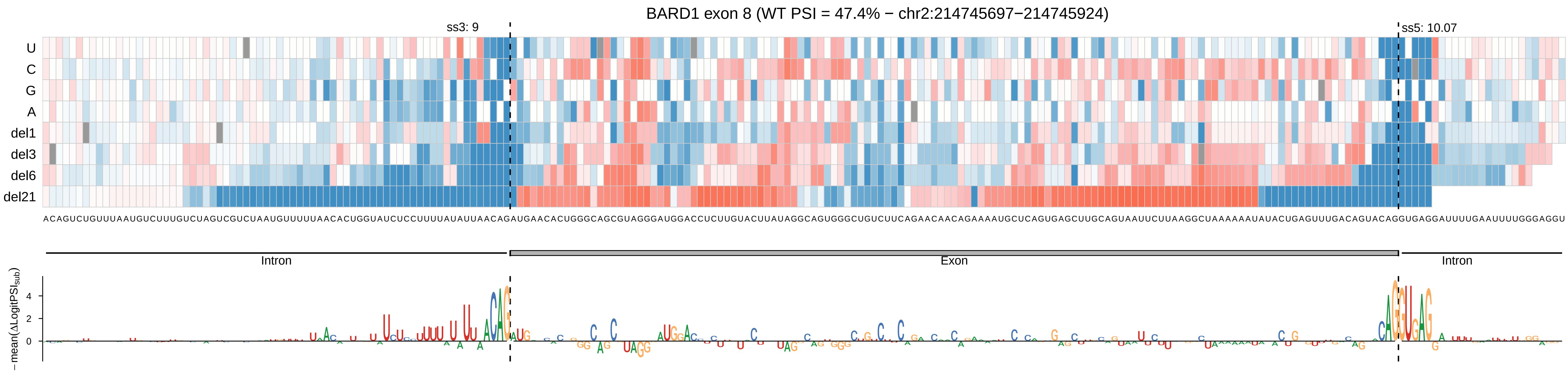

BARD1 exon 8 (WT PSI = 47.4% − chr2:214745697−214745924)
ss3: 9
ss5: 10.07
U
C
G
A
del1
del3
del6
del21
ACAGUCUGUUUAAUGUCUUUGUCUAGUCGUCUAAUGUUUUUAACACUGGUAUCUCCUUUUAUAUUAACAGAUGAACACUGGGCAGCGUAGGGAUGGACCUCUUGUACUUAUAGGCAGUGGGCUGUCUUCAGAACAACAGAAAAUGCUCAGUGAGCUUGCAGUAAUUCUUAAGGCUAAAAAAUAUACUGAGUUUGACAGUACAGGUGAGGAUUUUGAAUUUUGGGAGGU
Intron
Exon
Intron
−mean(ΔLogitPSIsub)
4
2
0

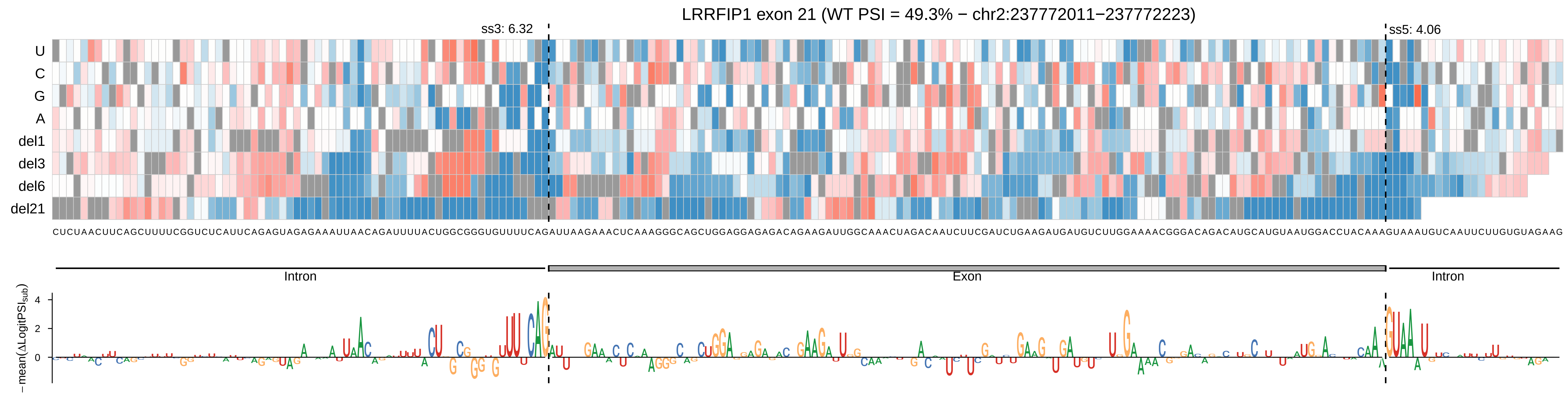
LRRFIP1 exon 21 (WT PSI = 49.3% − chr2:237772011−237772223)
ss3: 6.32
ss5: 4.06
U
C
G
A
del1
del3
del6
del21
Intron
Exon
Intron
−mean(ΔLogitPSIsub)
4
2
0

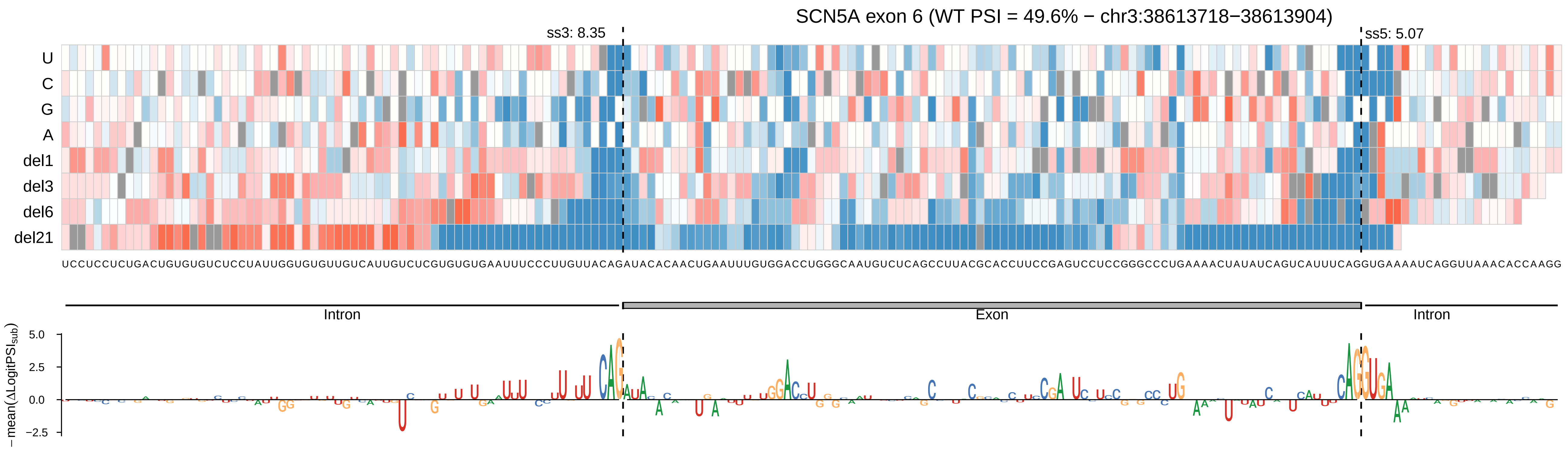
SCN5A exon 6 (WT PSI = 49.6% − chr3:38613718−38613904)
ss3: 8.35
ss5: 5.07
U
C
G
A
del1
del3
del6
del21
UCCUCCUCUGACUGUGUGUCUCCUAUUGGUGUGUUGUCAUUGUCUCGUGUGUGAAUUUCCCUUGUUACAGAUACACAACUGAAUUUGUGGACCUGGGCAAUGUCUCAGCCUUACGCACCUUCCGAGUCCUCCGGGCCCUGAAAACUAUAUCAGUCAUUUCAGGUGAAAAUCAGGUUAAACACCAAGG
Intron
Exon
Intron
− mean(ΔLogitPSI_sub)
5.0
2.5
0.0
−2.5

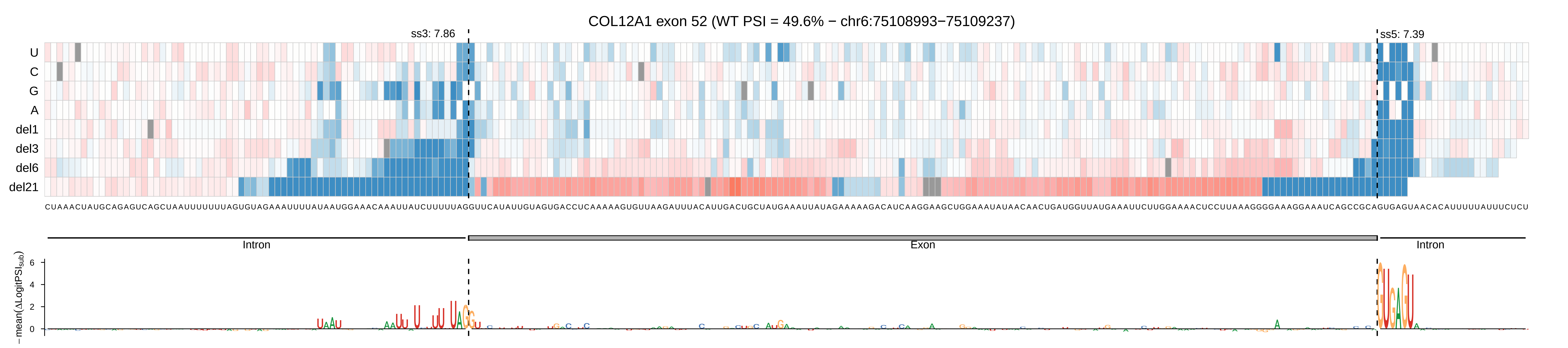
COL12A1 exon 52 (WT PSI = 49.6% − chr6:75108993−75109237)
ss3: 7.86
ss5: 7.39
U
C
G
A
del1
del3
del6
del21
Intron
Exon
Intron
−mean(ΔLogitPSI_sub)
6
4
2
0

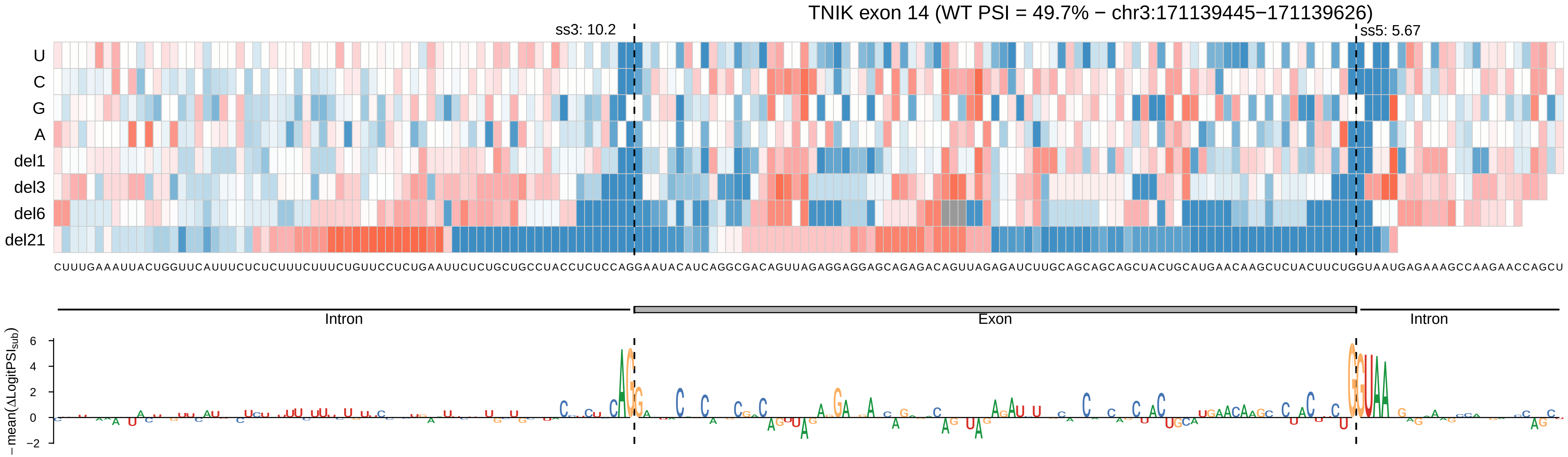
TNIK exon 14 (WT PSI = 49.7% − chr3:171139445−171139626)
ss3: 10.2
ss5: 5.67
U
C
G
A
del1
del3
del6
del21
CUUUGAAAUUACUGGUUCAUUUCUCUCUUUCUUUCUGUUCCUCUGAAUUCUCUGCUGCCUACCUCUCCAGGAAUACAUCAGGCGACAGUUAGAGGAGGAGCAGAGACAGUUAGAGAUCUUGCAGCAGCAGCUACUGCAUGAACAAGCUCUACUUCUGGUAAUGAGAAAGCCAAGAACCAGCU
Intron
Exon
Intron
− mean(ΔLogitPSI_sub)
6
4
2
0
−2

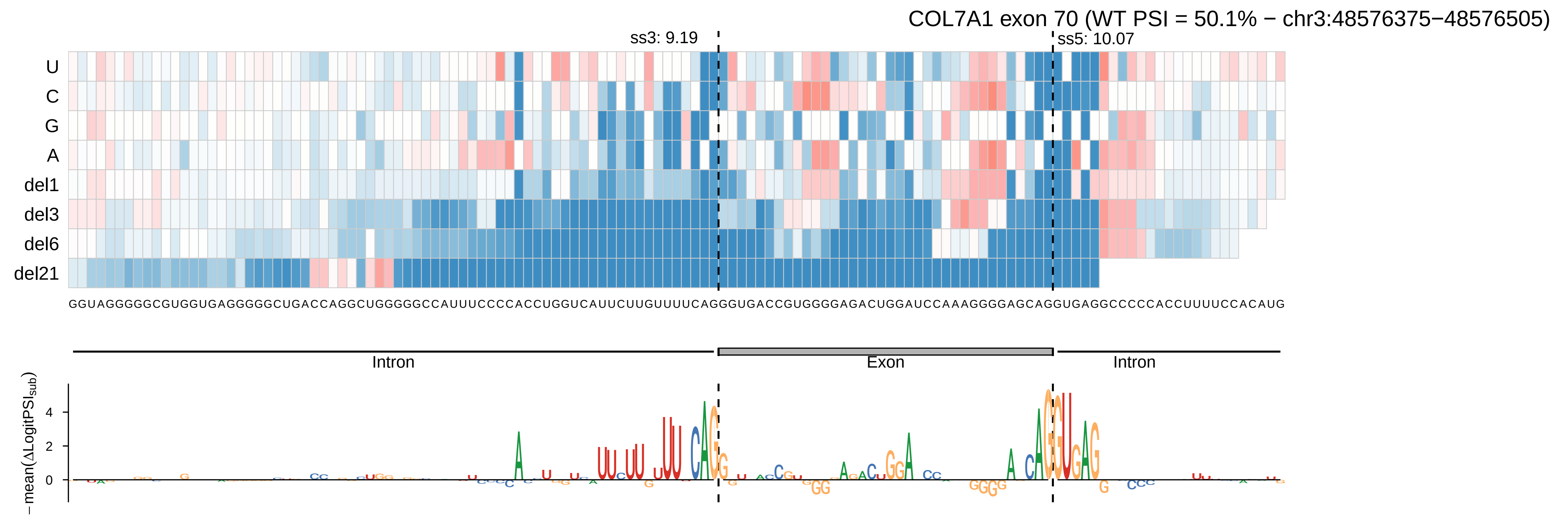
COL7A1 exon 70 (WT PSI = 50.1% − chr3:48576375−48576505)
ss3: 9.19
ss5: 10.07
U
C
G
A
del1
del3
del6
del21
GGUAGGGGGCGUGGUGAGGGGGCUGACCAGGCUGGGGGCCAUUUCCCCACCUGGUCAUUCUUGUUUUCAGGGUGACCGUGGGGAGACUGGAUCCAAAGGGGAGCAGGUGAGGCCCCCACCUUUUCCACAUG
Intron
Exon
Intron
− mean(ΔLogitPSI_sub)
4
2
0

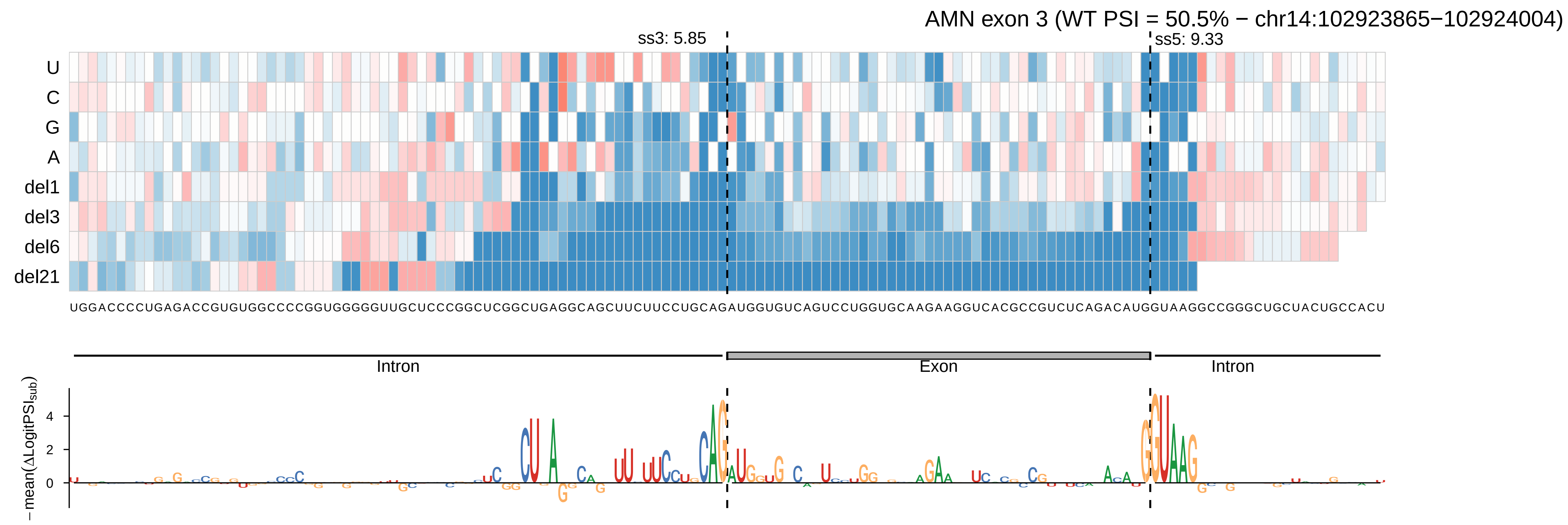
AMN exon 3 (WT PSI = 50.5% − chr14:102923865−102924004)
ss3: 5.85
ss5: 9.33
U
C
G
A
del1
del3
del6
del21
UGGACCCCUGAGACCGUGUGGCCCCGGUGGGGGUUGCUCCCGGCUCGGCUGAGGCAGCUUCUUCCUGCAGAUGGUGUCAGUCCUGGUGCAAGAAGGUCACGCCGUCUCAGACAUGGUAAGGCCGGGCUGCUACUGCCACU
Intron
Exon
Intron
−mean(ΔLogitPSI_sub)
4
2
0

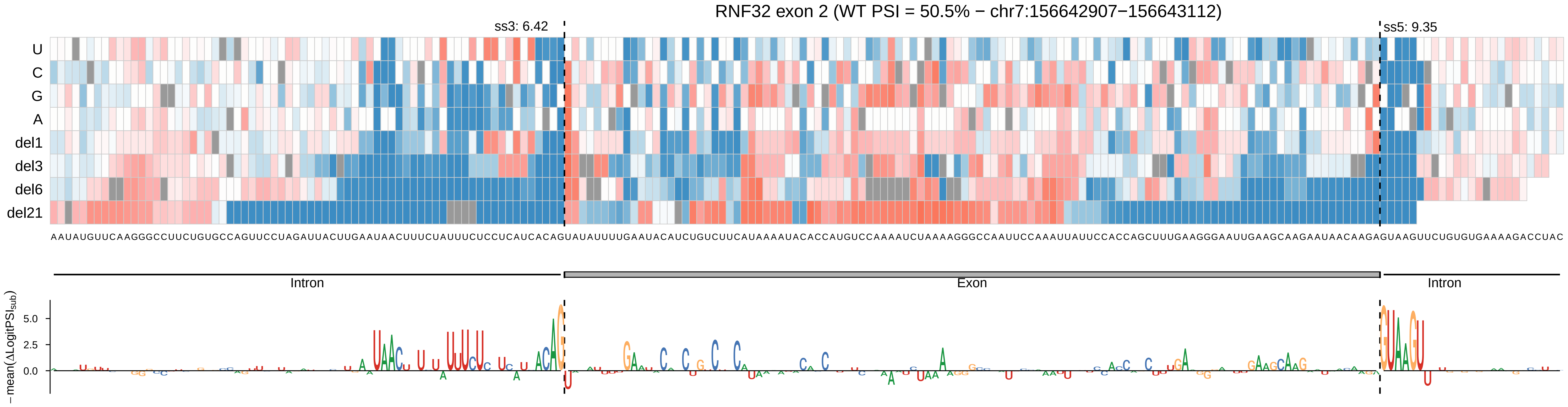
RNF32 exon 2 (WT PSI = 50.5% − chr7:156642907−156643112)
ss3: 6.42
ss5: 9.35
U
C
G
A
del1
del3
del6
del21
Intron
Exon
Intron
−mean(ΔLogitPSI_sub)
5.0
2.5
0.0

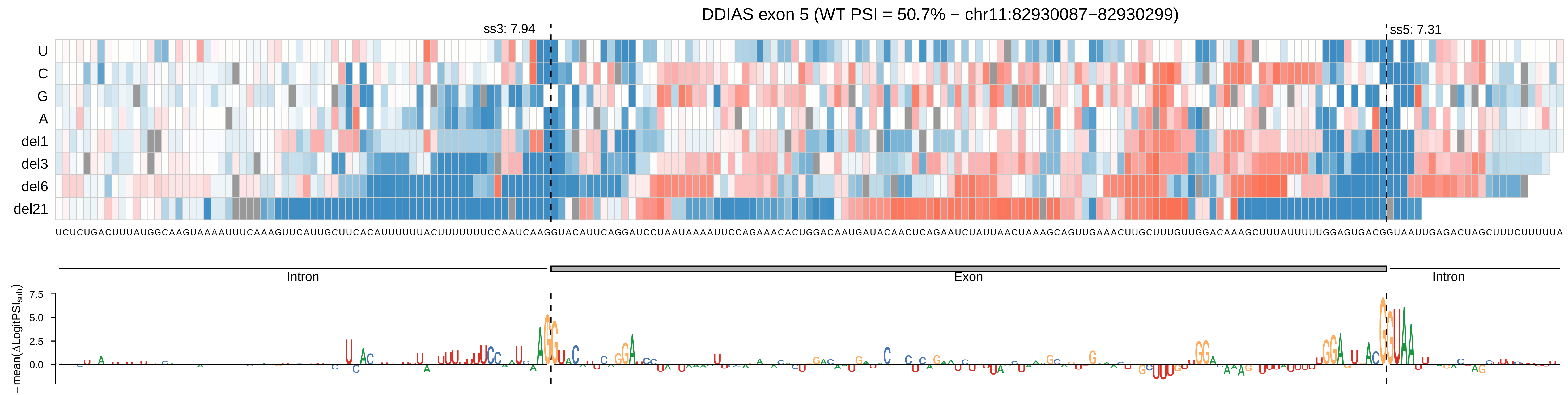
DDIAS exon 5 (WT PSI = 50.7% − chr11:82930087−82930299)
ss3: 7.94
ss5: 7.31
U
C
G
A
del1
del3
del6
del21
UCUCUGACUUUAUGGCAAGUAAAAUUUCAAAGUUCAUUGCUUCACAUUUUUUACUUUUUUUCCAAUCAAGGUACAUUCAGGAUCCUAAUAAAAUUCCAGAAACACUGGACAAUGAUACAACUCAGAAUCUAUUAACUAAAGCAGUUGAAACUUGCUUUGUUGGACAAAGCUUUAUUUUUGGAGUGACGGUAAUUGAGACUAGCUUUCUUUUUA
Intron
Exon
Intron
−mean(ΔLogitPSIsub)
7.5
5.0
2.5
0.0

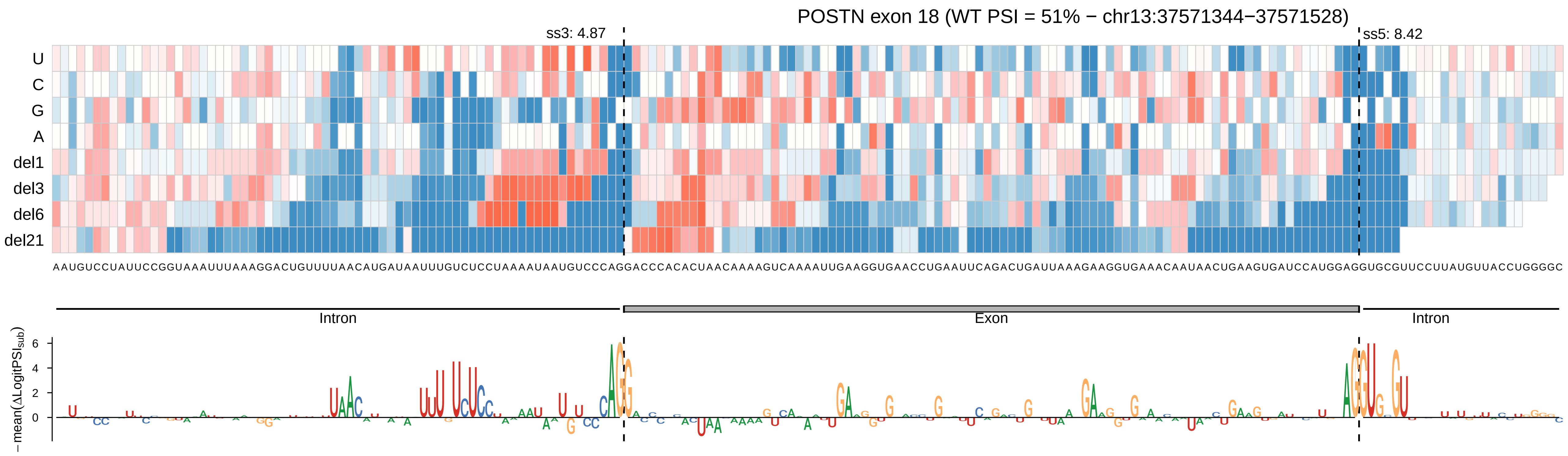
POSTN exon 18 (WT PSI = 51% − chr13:37571344−37571528)
ss3: 4.87
ss5: 8.42
U
C
G
A
del1
del3
del6
del21
AAUGUCCUAUUCCGGUAAAUUUAAAGGACUGUUUUAACAUGAUAAUUUGUCUCCUAAAAUAAUGUCCCAGGACCCACACUAACAAAAGUCAAAAUUGAAGGUGAACCUGAAUUCAGACUGAUUAAAGAAGGUGAAACAAUAACUGAAGUGAUCCAUGGAGGUGCGUUCCUUAUGUUACCUGGGGC
Intron
Exon
Intron
−mean(ΔLogitPSI_sub)
6
4
2
0

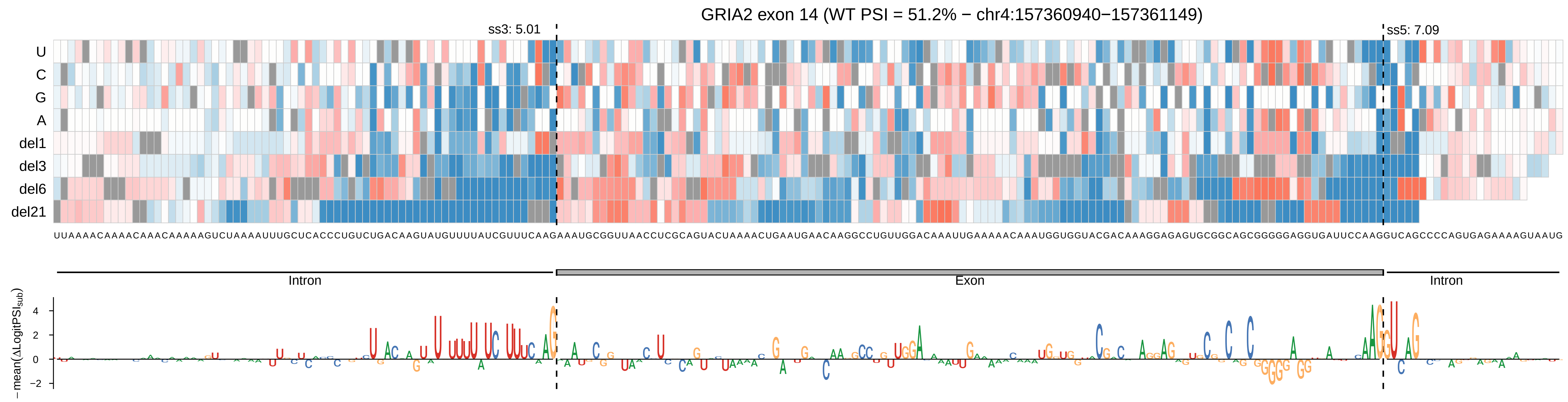
GRIA2 exon 14 (WT PSI = 51.2% − chr4:157360940−157361149)
ss3: 5.01
ss5: 7.09
U
C
G
A
del1
del3
del6
del21
Intron
Exon
Intron
−mean(ΔLogitPSI_sub)
4
2
0
−2

LIMCH1 exon 12 (WT PSI = 51.7% − chr4:41632907−41633110)

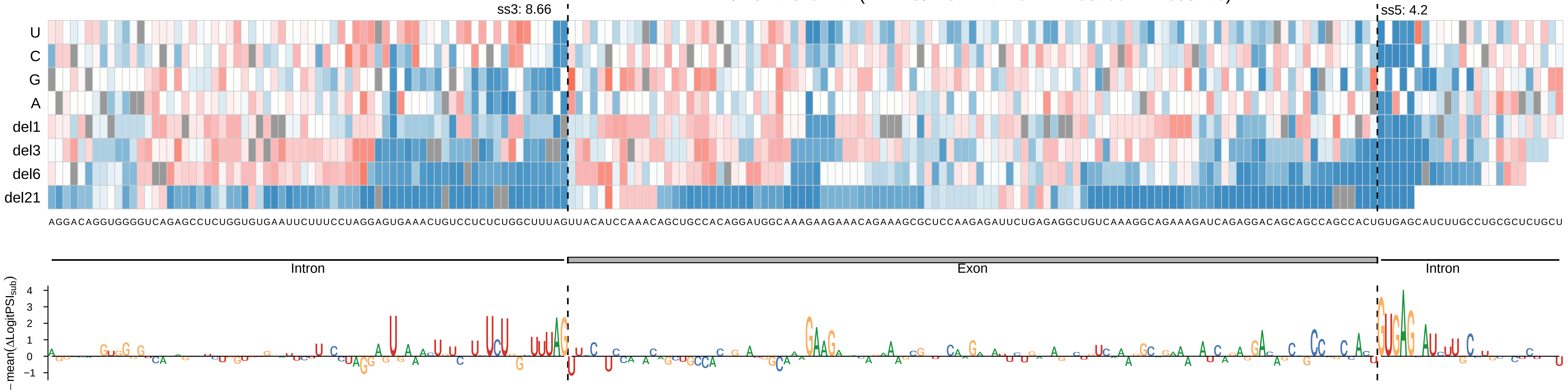

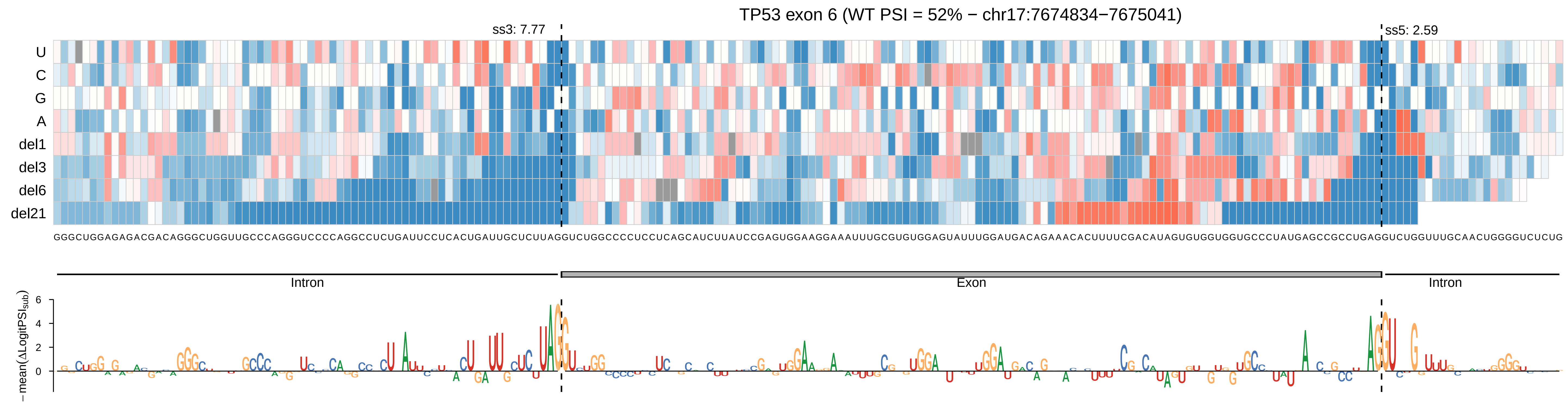

TP53 exon 6 (WT PSI = 52% − chr17:7674834−7675041)
ss3: 7.77
ss5: 2.59
U
C
G
A
del1
del3
del6
del21
Intron
Exon
Intron
−mean(ΔLogitPSI_sub)

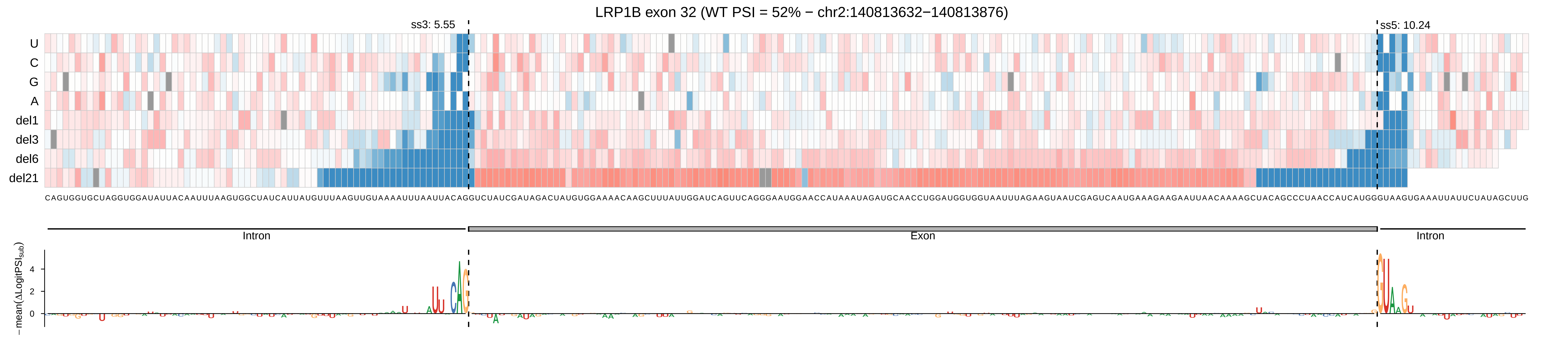
LRP1B exon 32 (WT PSI = 52% − chr2:140813632−140813876)
ss3: 5.55
ss5: 10.24
U
C
G
A
del1
del3
del6
del21
Intron
Exon
Intron
−mean(ΔLogitPSI_sub)
4
2
0

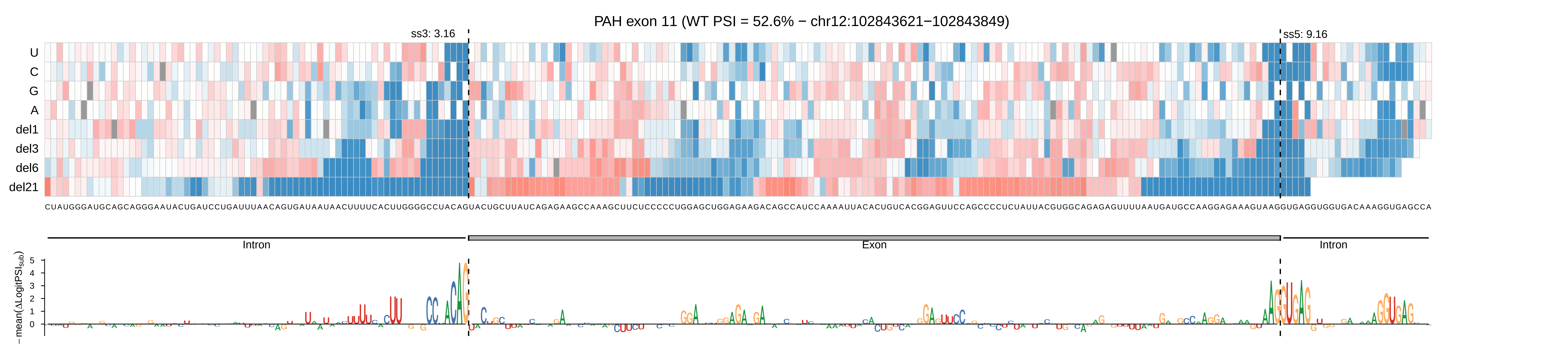
PAH exon 11 (WT PSI = 52.6% − chr12:102843621−102843849)
ss3: 3.16
ss5: 9.16
U
C
G
A
del1
del3
del6
del21
Intron
Exon
Intron
−mean(ΔLogitPSI_sub)
5
4
3
2
1
0

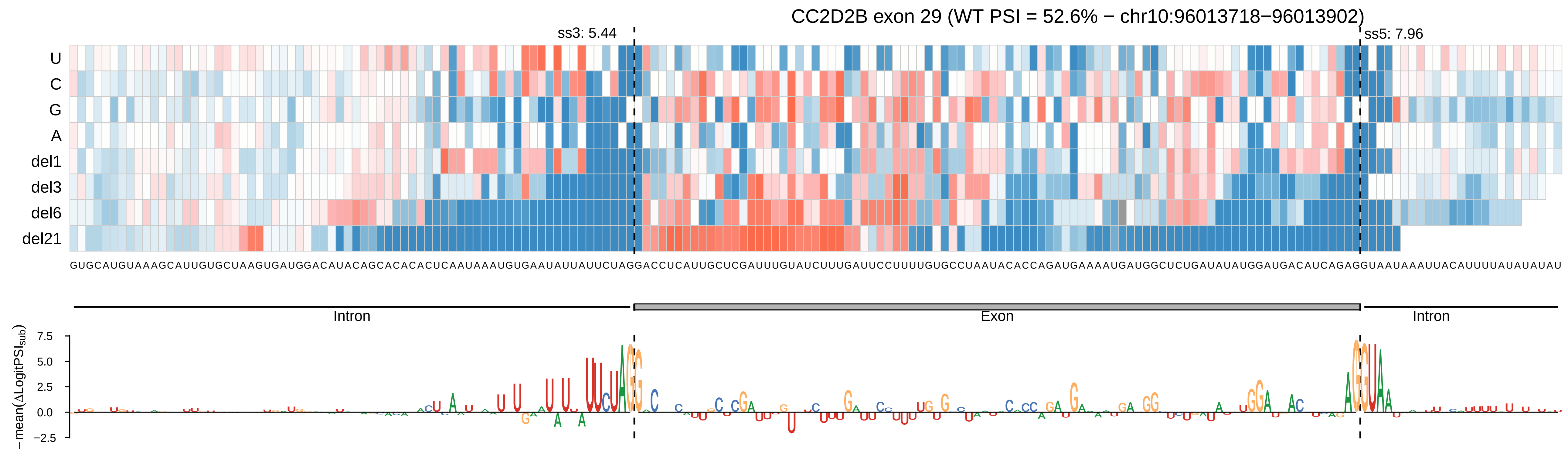
CC2D2B exon 29 (WT PSI = 52.6% − chr10:96013718−96013902)
ss3: 5.44
ss5: 7.96
U
C
G
A
del1
del3
del6
del21
GUGCAUGUAAAGCAUUGUGCUAAGUGAUGGACAUACAGCACACACUCAAUAAAUGUGAAUAUUAUUCUAGGACCUCAUUGCUCGAUUUGUAUCUUUGAUUCCUUUUGUGCCUAAUACACCAGAUGAAAAUGAUGGCUCUGAUAUAUGGAUGACAUCAGAGGUAAUAAAUUACAUUUUAUAUAUAU
Intron
Exon
Intron
−mean(ΔLogitPSI_sub)
7.5
5.0
2.5
0.0
−2.5

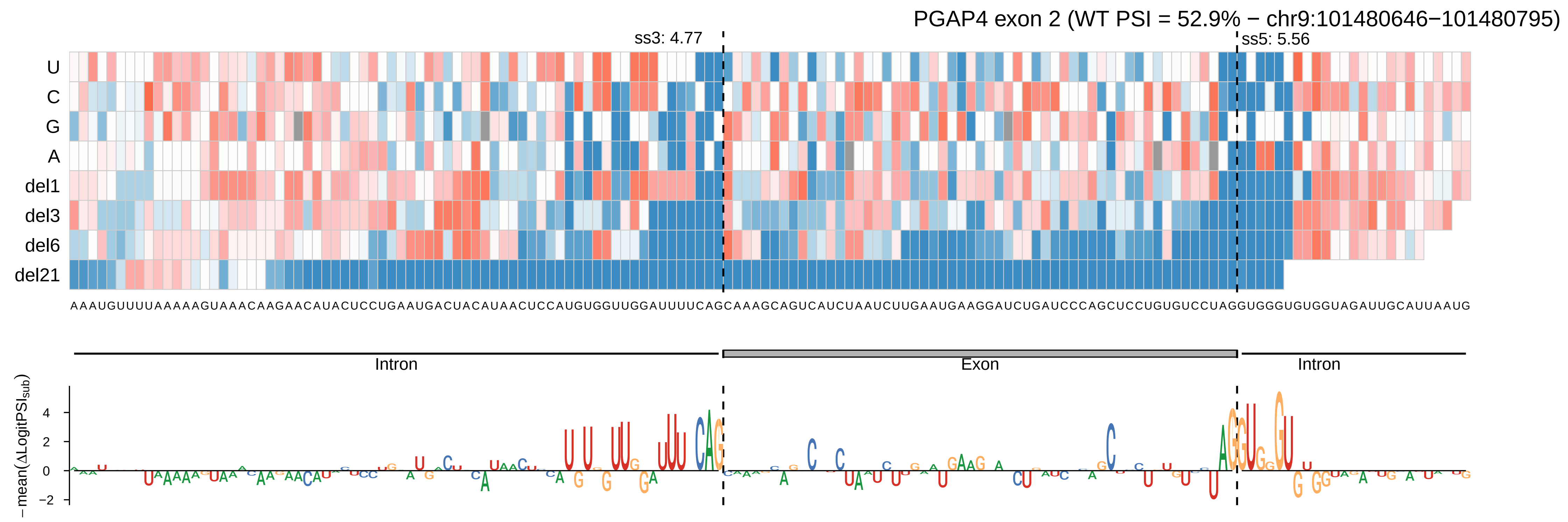
PGAP4 exon 2 (WT PSI = 52.9% − chr9:101480646−101480795)
ss3: 4.77
ss5: 5.56
U
C
G
A
del1
del3
del6
del21
AAAUGUUUUAAAAAGUAAACAAGAACAUACUCCUGAAUGACUACAUAACUCCAUGUGGUUGGAUUUUCAGCAAAGCAGUCAUCUAAUCUUGAAUGAAGGAUCUGAUCCCAGCUCCUGUGUCCUAGGUGGGUGUGGUAGAUUGCAUUAAUG
Intron
Exon
Intron
−mean(ΔLogitPSI$_{sub}$)
4
2
0
−2

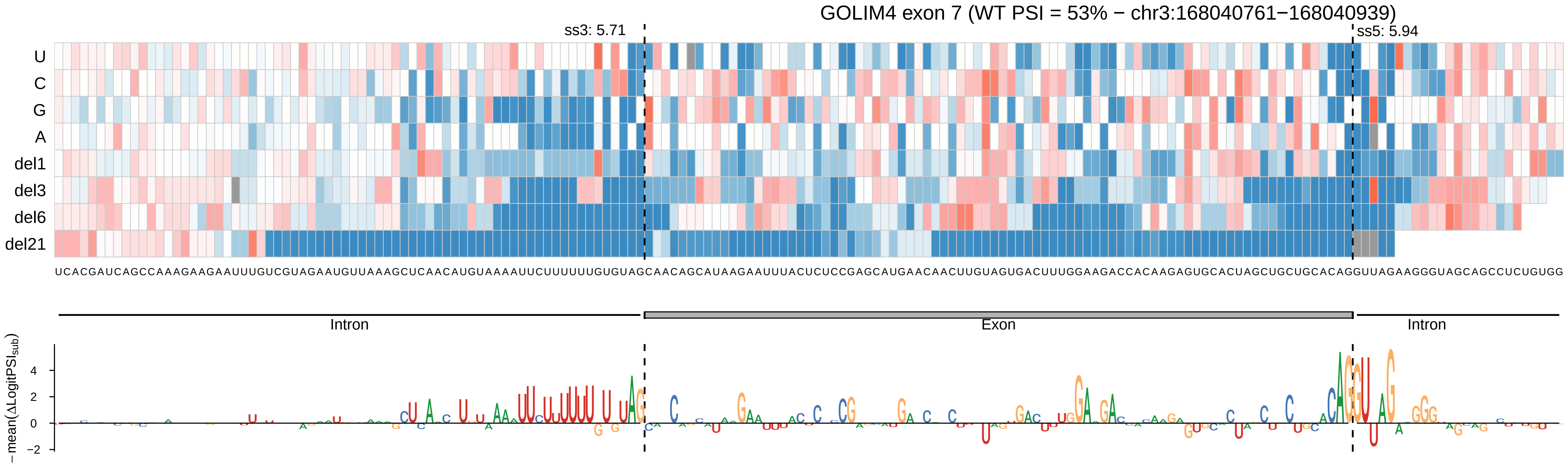

GOLIM4 exon 7 (WT PSI = 53% − chr3:168040761−168040939)
ss3: 5.71
ss5: 5.94
U
C
G
A
del1
del3
del6
del21
UCACGAUCAGCCAAAGAAGAAUUUGUCGUAGAAUGUUAAAGCUCAACAUGUAAAAUUCUUUUUUGUGUAGCAACAGCAUAAGAAUUUACUCUCCGAGCAUGAACAACUUGUAGUGACUUUGGAAGACCACAAGAGUGCACUAGCUGCUGCACAGGUUAGAAGGGUAGCAGCCUCUGUGG
Intron
Exon
Intron
−mean(ΔLogitPSI_sub)
4
2
0
−2

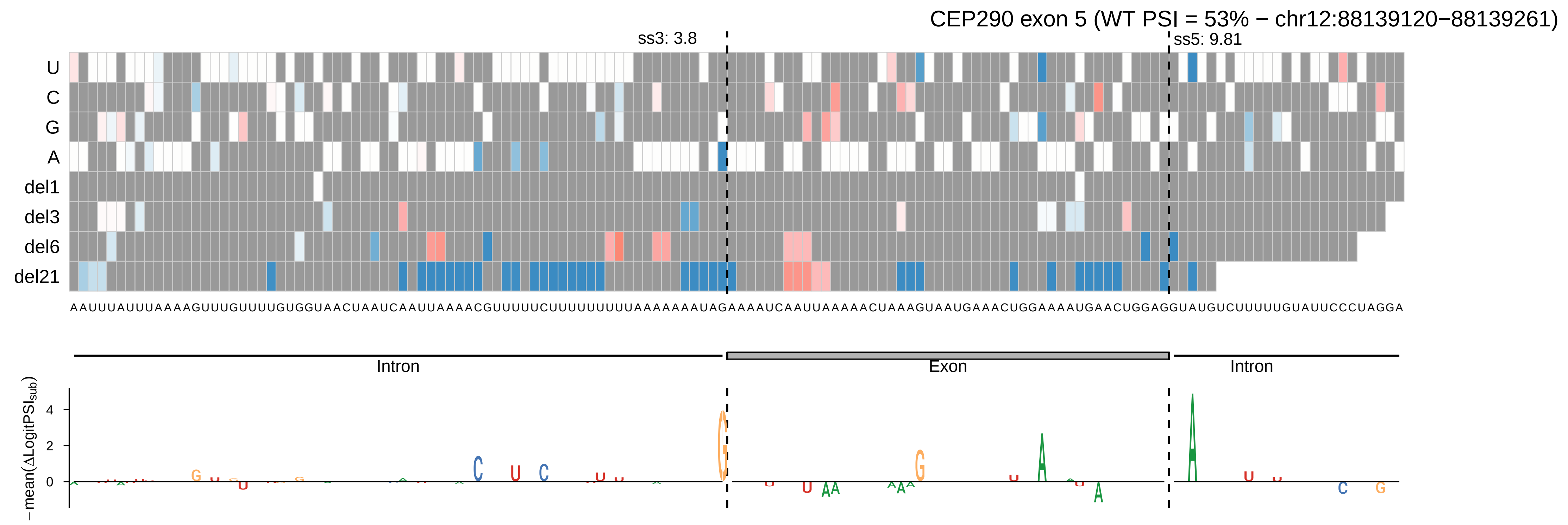

CEP290 exon 5 (WT PSI = 53% − chr12:88139120−88139261)
ss3: 3.8
ss5: 9.81
U
C
G
A
del1
del3
del6
del21
AAUUUAUUUAAAAGUUUGUUUUGUGGUAACUAAUCAAUUAAAACGUUUUUCUUUUUUUUUAAAAAAAAUAGAAAAUCAAUUAAAAACUAAAGUAAUGAAACUGGAAAAUGAACUGGAGGUAUGUCUUUUUGUAUUCCCUAGGA
Intron
Exon
Intron
− mean(ΔLogitPSI_sub)
4
2
0

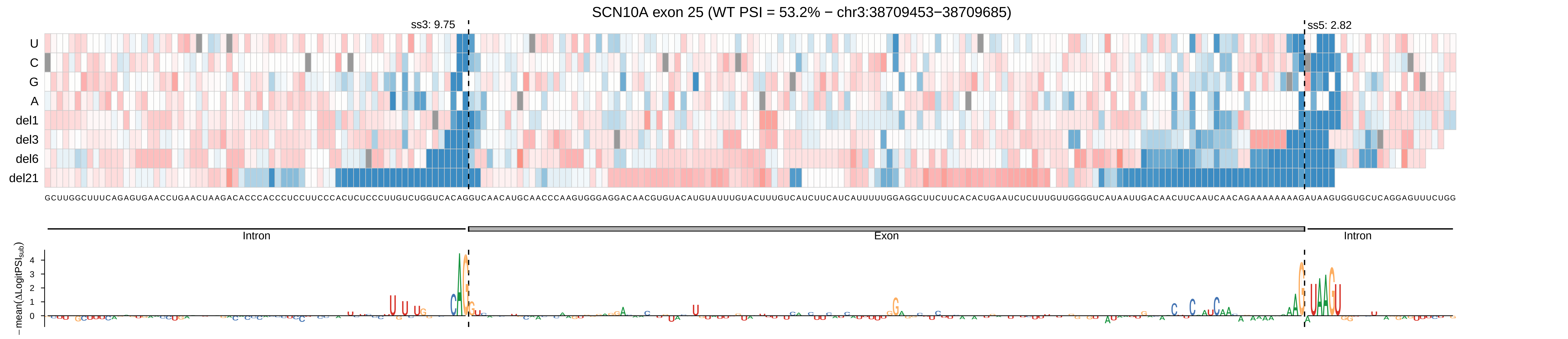

SCN10A exon 25 (WT PSI = 53.2% − chr3:38709453−38709685)
ss3: 9.75
ss5: 2.82
U
C
G
A
del1
del3
del6
del21
GCUUGGCUUUCAGAGUGAACCUGAACUAAGACACCCACCCUCCUUCCCACUCUCCCUUGUCUGGUCACAGGUCAACAUGCAACCCAAGUGGGAGGACAACGUGUACAUGUAUUUGUACUUUGUCAUCUUCAUCAUUUUUGGAGGCUUCUUCACACUGAAUCUCUUUGUUGGGGUCAUAAUUGACAACUUCAAUCAACAGAAAAAAAAGAUAAGUGGUGCUCAGGAGUUUCUGG
Intron
Exon
Intron
−mean(ΔLogitPSI_sub)
0
1
2
3
4

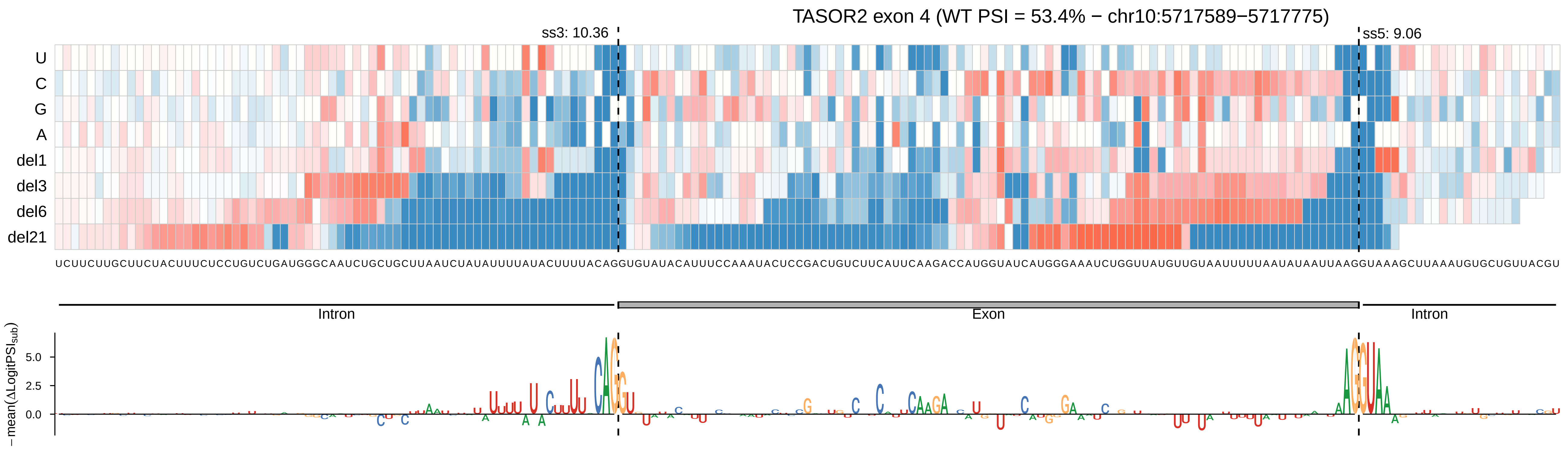
TASOR2 exon 4 (WT PSI = 53.4% − chr10:5717589−5717775)
ss3: 10.36
ss5: 9.06
U
C
G
A
del1
del3
del6
del21
Intron
Exon
Intron
−mean(ΔLogitPSI_sub)
5.0
2.5
0.0

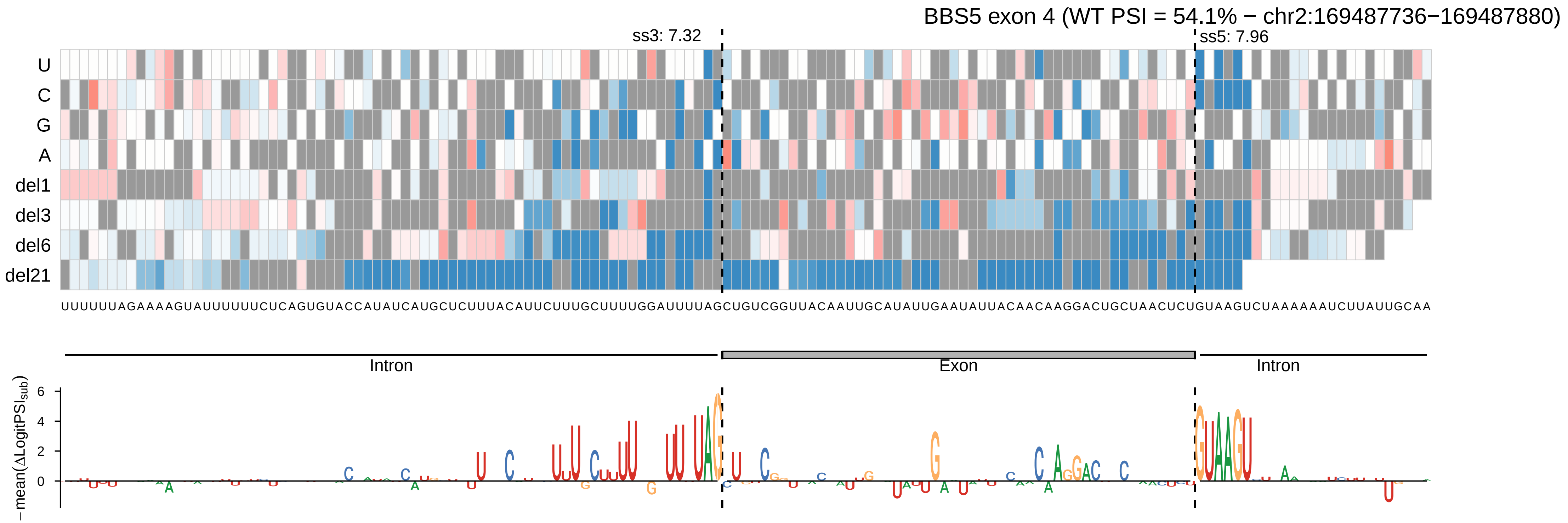

BBS5 exon 4 (WT PSI = 54.1% − chr2:169487736−169487880)
ss3: 7.32
ss5: 7.96
U
C
G
A
del1
del3
del6
del21
UUUUUUAGAAAAGUAUUUUUUCUCAGUGUACCAUAUCAUGCUCUUUACAUUCUUUGCUUUUGGAUUUUAGCUGUCGGUUACAAUUGCAUAUUGAAUAUUACAACAAGGACUGCUAACUCUGUAAGUCUAAAAAAUCUUAUUGCAA
Intron
Exon
Intron
−mean(ΔLogitPSI_sub)
6
4
2
0

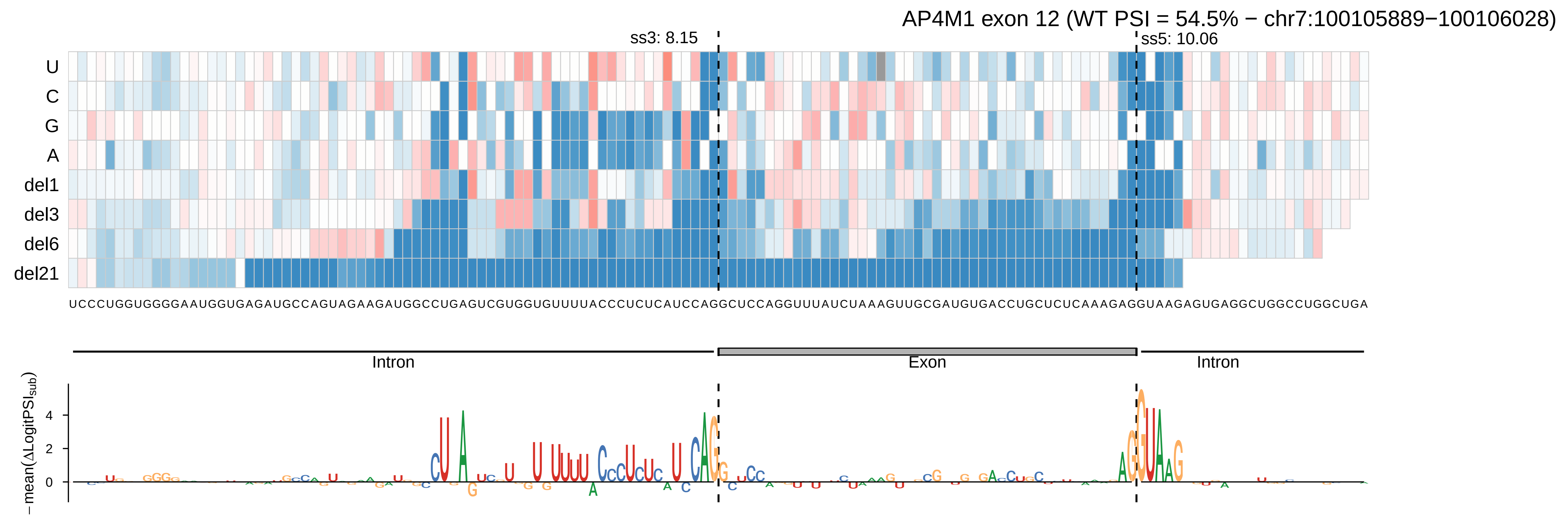

AP4M1 exon 12 (WT PSI = 54.5% − chr7:100105889−100106028)
ss3: 8.15
ss5: 10.06
U
C
G
A
del1
del3
del6
del21
UCCCUGGUGGGGAAUGGUGAGAUGCCAGUAGAAGAUGGCCUGAGUCGUGGUGUUUUACCCUCUCAUCCAGGCUCCAGGUUUAUCUAAAGUUGCGAUGUGACCUGCUCUCAAAGAGGUAAGAGUGAGGCUGGCCUGGCUGA
Intron
Exon
Intron
−mean(ΔLogitPSI_sub)
4
2
0

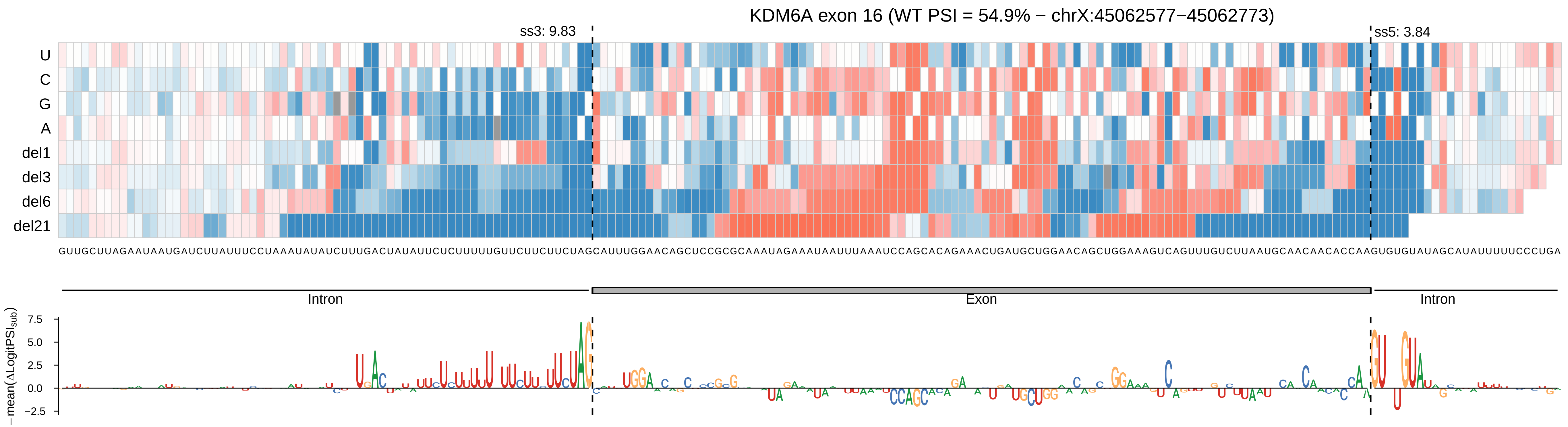

KDM6A exon 16 (WT PSI = 54.9% − chrX:45062577−45062773)
ss3: 9.83
ss5: 3.84
U
C
G
A
del1
del3
del6
del21
GUUGCUUAGAAUAAUGAUCUUAUUUCCUAAAUAUAUCUUUGACUAUAUUCUCUUUUUGUUCUUCUUCUAGCAUUUGGAACAGCUCCGCGCAAAUAGAAAUAAUUUAAAUCCAGCACAGAAACUGAUGCUGGAACAGCUGGAAAGUCAGUUUGUCUUAAUGCAACAACACCAAGUGUGUAUAGCAUAUUUUUCCCUGA
Intron
Exon
Intron
−mean(ΔLogitPSI_sub)
7.5
5.0
2.5
0.0
−2.5

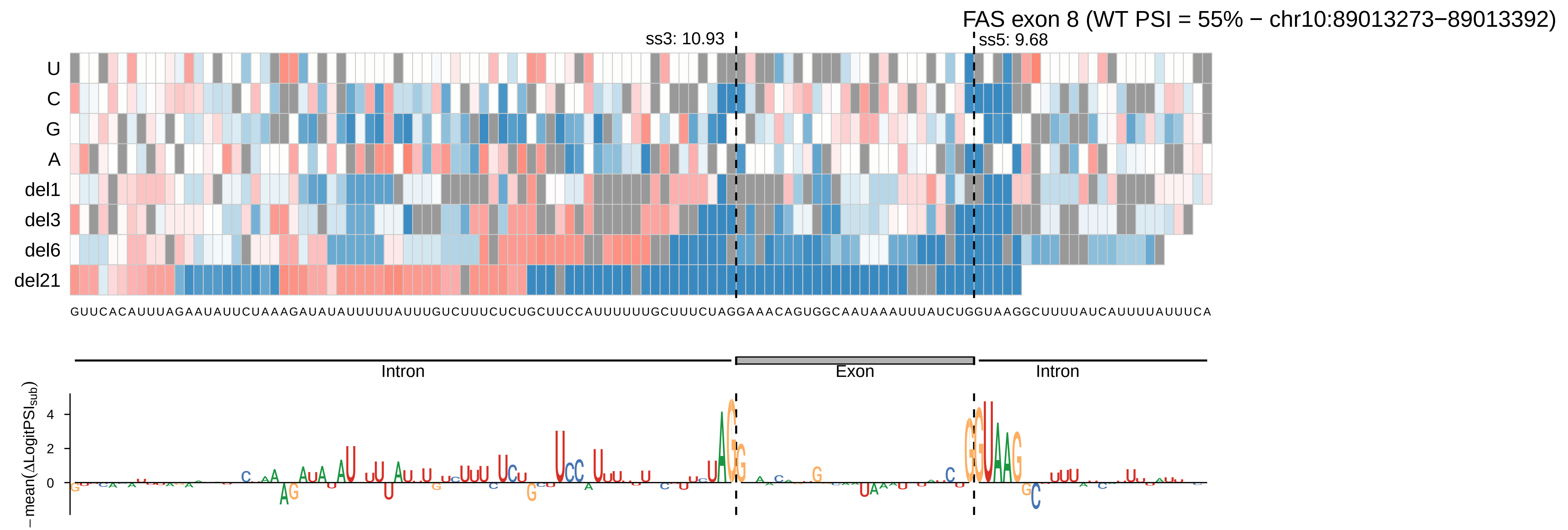
FAS exon 8 (WT PSI = 55% − chr10:89013273−89013392)
ss3: 10.93
ss5: 9.68
U
C
G
A
del1
del3
del6
del21
GUUCACAUUUAGAAUAUUCUAAAGAUAUAUUUUUAUUUGUCUUUCUCUGCUUCCAUUUUUUGCUUUCUAGGAAACAGUGGCAAUAAAUUUAUCUGGUAAGGCUUUUAUCAUUUUAUUUCA
Intron
Exon
Intron
−mean(ΔLogitPSI_sub)

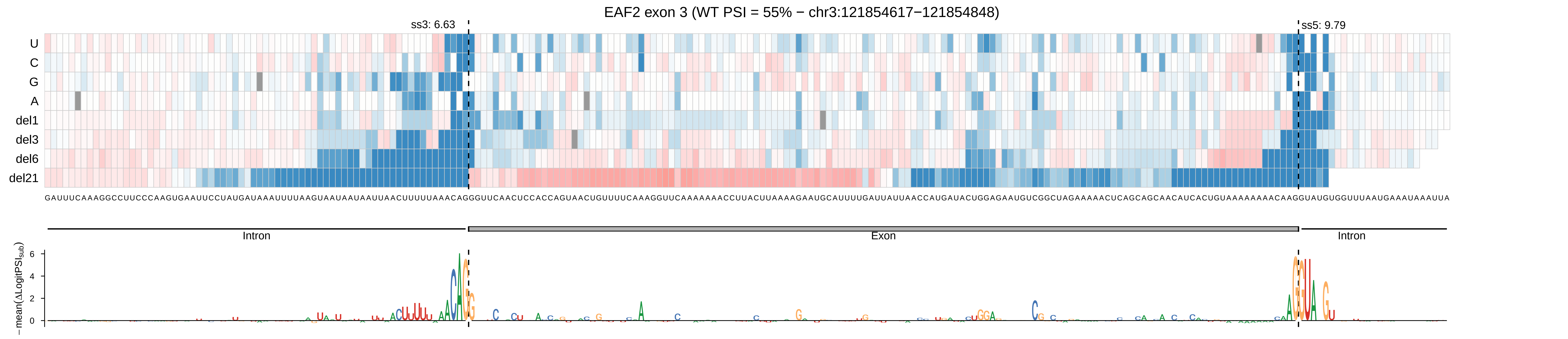

EAF2 exon 3 (WT PSI = 55% − chr3:121854617−121854848)
ss3: 6.63
ss5: 9.79
U
C
G
A
del1
del3
del6
del21
Intron
Exon
Intron
−mean(ΔLogitPSI_sub)
0
2
4
6

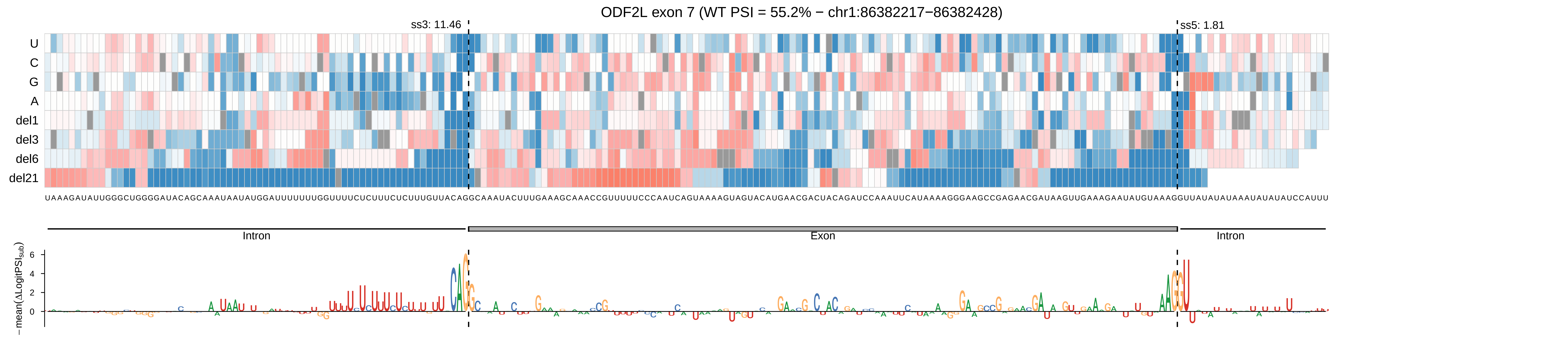

ODF2L exon 7 (WT PSI = 55.2% − chr1:86382217−86382428)
ss3: 11.46
ss5: 1.81
U
C
G
A
del1
del3
del6
del21
UAAAGAUAUUGGGCUGGGGAUACAGCAAAUAAUAUGGAUUUUUUUGGUUUUCUCUUUCUCUUUGUUACAGGCAAAUACUUUGAAAGCAAACCGUUUUUCCCAAUCAGUAAAAGUAGUACAUGAACGACUACAGAUCCAAAUUCAUAAAAGGGAAGCCGAGAACGAUAAGUUGAAAGAAUAUGUAAAGGUUAUAUAUAAAUAUAUAUCCAUUU
Intron
Exon
Intron
− mean(ΔLogitPSI_sub)
6
4
2
0

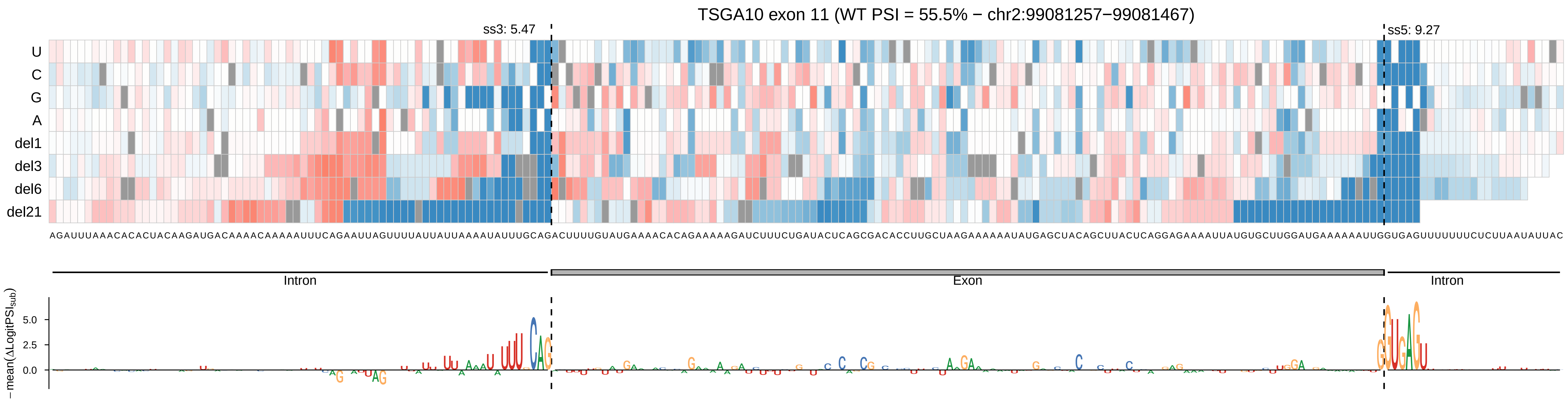

TSGA10 exon 11 (WT PSI = 55.5% − chr2:99081257−99081467)
ss3: 5.47
ss5: 9.27
U
C
G
A
del1
del3
del6
del21
Intron
Exon
Intron
−mean(ΔLogitPSI_sub)
5.0
2.5
0.0

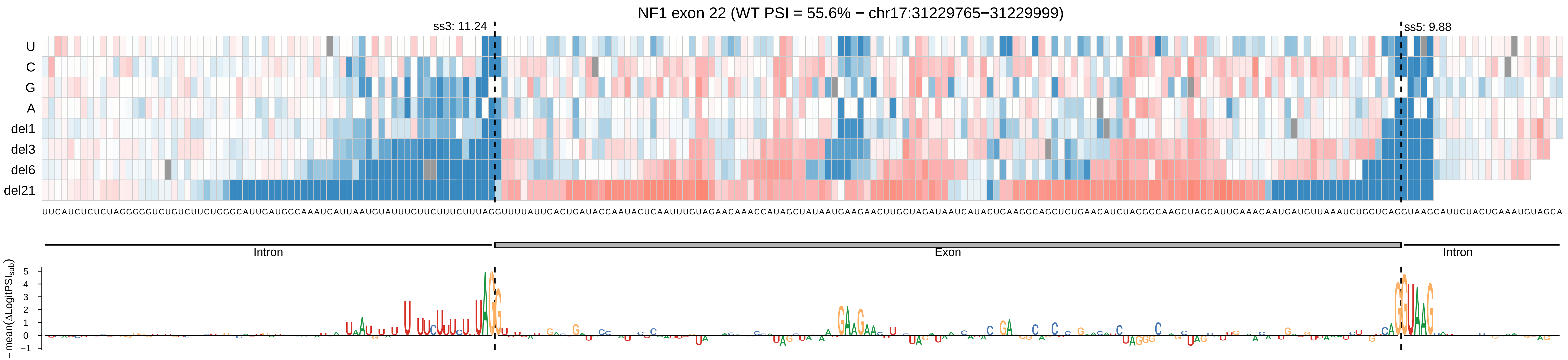
NF1 exon 22 (WT PSI = 55.6% − chr17:31229765−31229999)
ss3: 11.24
ss5: 9.88
U
C
G
A
del1
del3
del6
del21
Intron
Exon
Intron
− mean(ΔLogitPSI_sub)
5
4
3
2
1
0
−1

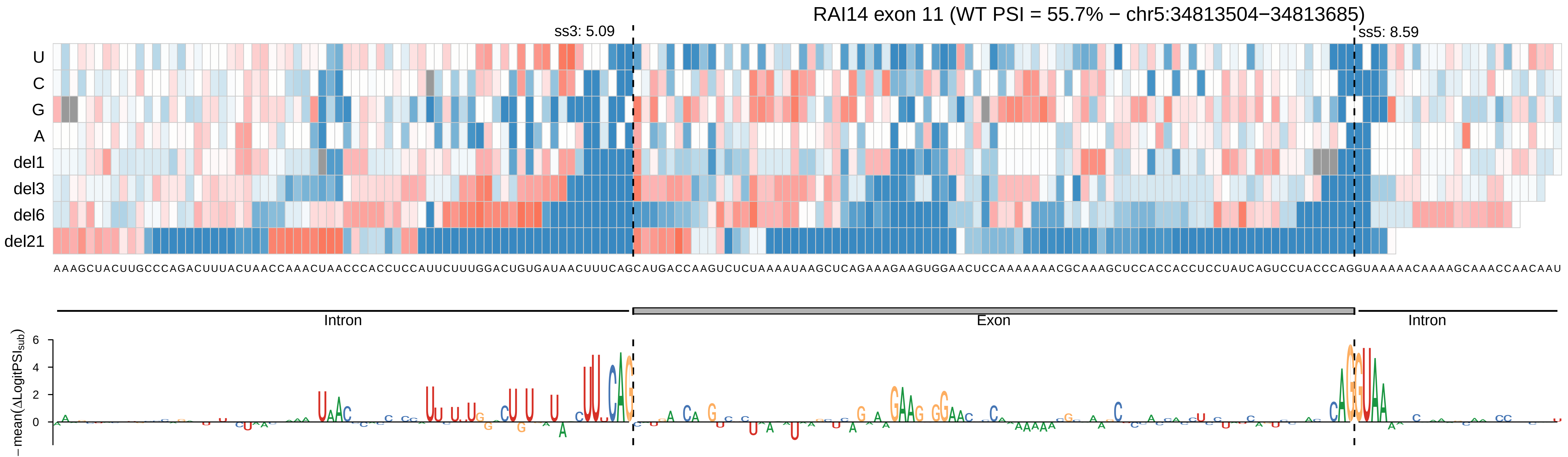
RAI14 exon 11 (WT PSI = 55.7% − chr5:34813504−34813685)
ss3: 5.09
ss5: 8.59
U
C
G
A
del1
del3
del6
del21
AAAGCUACUUGCCCAGACUUUACUAACCAAACUAACCCACCUCCAUUCUUUGGACUGUGAUAACUUUCAGCAUGACCAAGUCUCUAAAAUAAGCUCAGAAAGAAGUGGAACUCCAAAAAAACGCAAAGCUCCACCACCUCCUAUCAGUCCUACCCAGGUAAAAACAAAAGCAAACCAACAAU
Intron
Exon
Intron
− mean(ΔLogitPSI_sub)
6
4
2
0

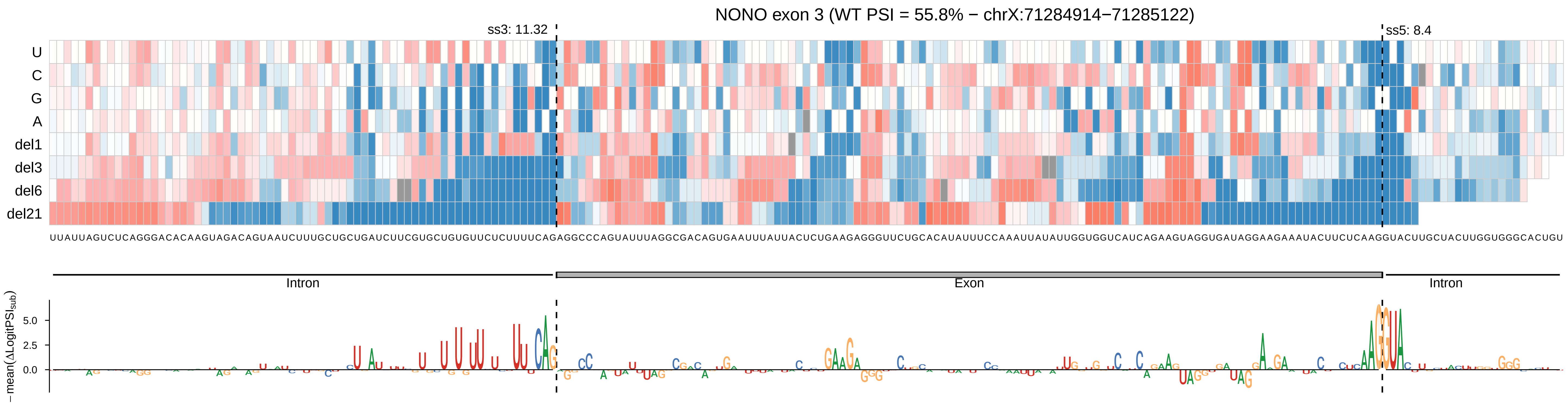

NONO exon 3 (WT PSI = 55.8% − chrX:71284914−71285122)
ss3: 11.32
ss5: 8.4
U
C
G
A
del1
del3
del6
del21
UUAUUAGUCUCAGGGACACAAGUAGACAGUAAUCUUUGCUGCUGAUCUUCGUGCUGUGUUCUCUUUUCAGAGGCCCAGUAUUUAGGCGACAGUGAAUUUAUUACUCUGAAGAGGGUUCUGCACAUAUUUCCAAAUUAUAUUGGUGGUCAUCAGAAGUAGGUGAUAGGAAGAAAUACUUCUCAAGGUACUUGCUACUUGGUGGGCACUGU
Intron
Exon
Intron
−mean(ΔLogitPSI_sub)
5.0
2.5
0.0

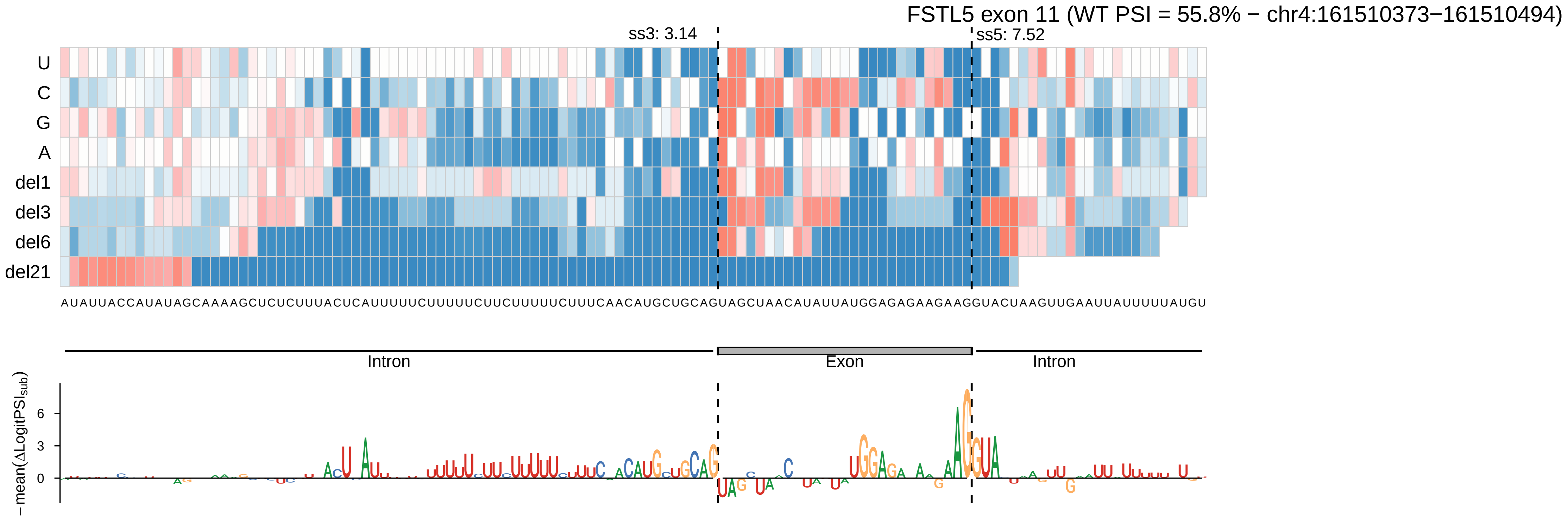

FSTL5 exon 11 (WT PSI = 55.8% − chr4:161510373−161510494)
ss3: 3.14
ss5: 7.52
U
C
G
A
del1
del3
del6
del21
AUAUUACCAUAUAGCAAAAGCUCUCUUUACUCAUUUUUCUUUUUCUUCUUUUUCUUUCAACAUGCUGCAGUAGCUAACAUAUUAUGGAGAGAAGAAGGUACUAAGUUGAAUUAUUUUUAUGU
Intron
Exon
Intron
−mean(ΔLogitPSI_sub)
6
3
0

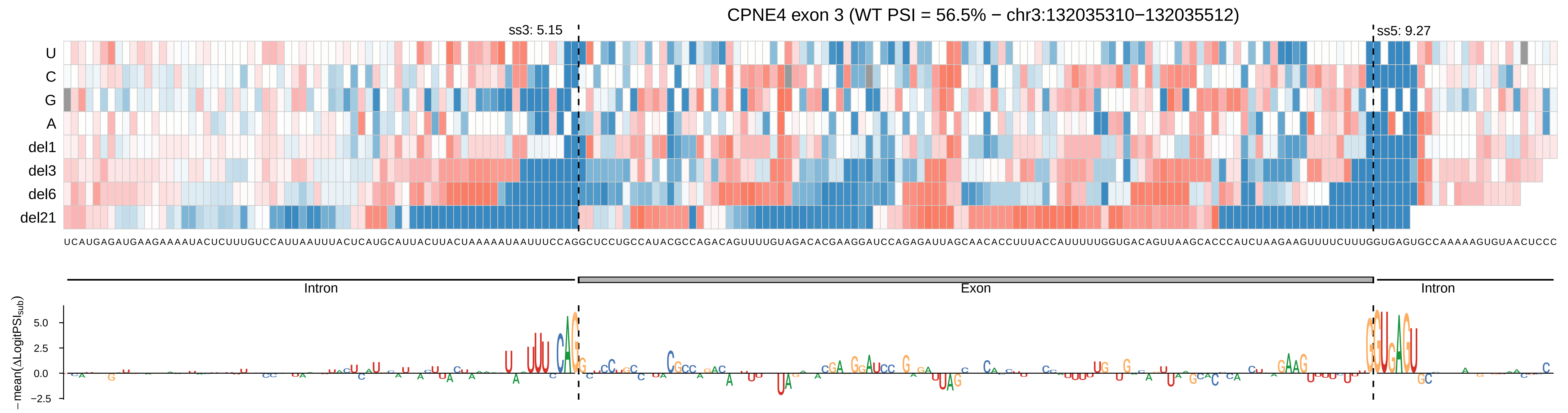

CPNE4 exon 3 (WT PSI = 56.5% − chr3:132035310−132035512)
ss3: 5.15
ss5: 9.27
U
C
G
A
del1
del3
del6
del21
UCAUGAGAUGAAGAAAAUACUCUUUGUCCAUUAAUUUACUCAUGCAUUACUUACUAAAAAUAAUUUCCAGGCUCCUGCCAUACGCCAGACAGUUUUGUAGACACGAAGGAUCCAGAGAUUAGCAACACCUUUACCAUUUUUGGUGACAGUUAAGCACCCAUCUAAGAAGUUUUCUUUGGUGAGUGCCAAAAAGUGUAACUCCC
Intron
Exon
Intron
− mean(ΔLogitPSI sub)
5.0
2.5
0.0
−2.5

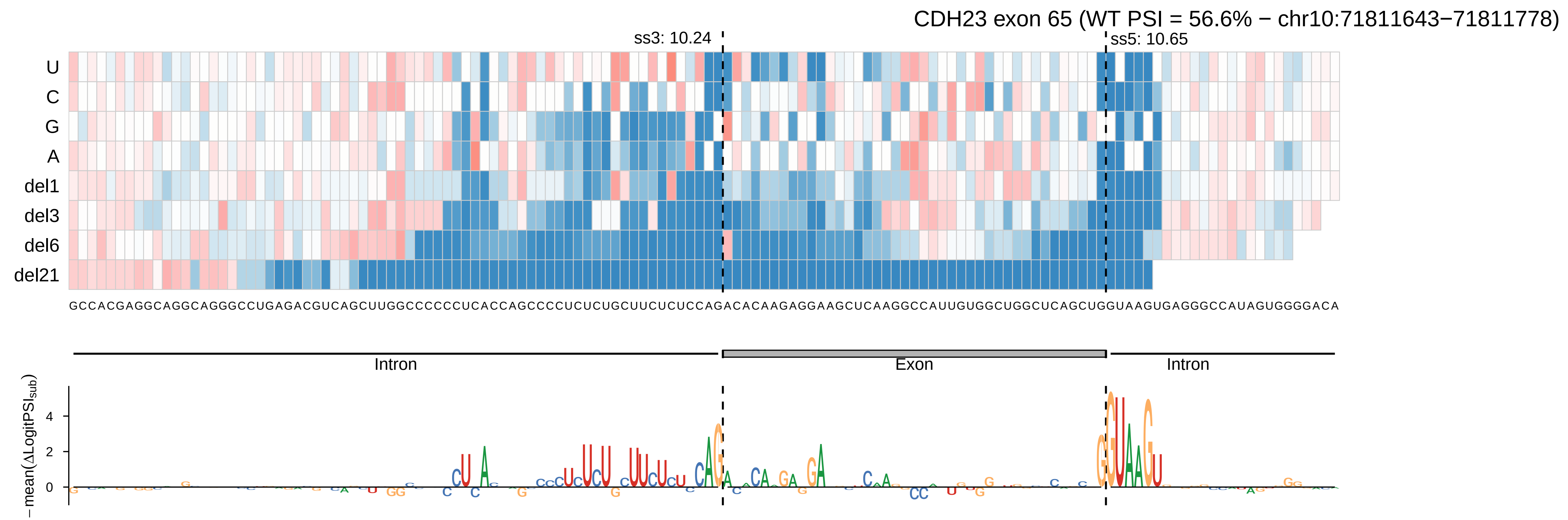
CDH23 exon 65 (WT PSI = 56.6% − chr10:71811643−71811778)
ss3: 10.24
ss5: 10.65
U
C
G
A
del1
del3
del6
del21
GCCACGAGGCAGGCAGGGCCUGAGACGUCAGCUUGGCCCCCCUCACCAGCCCCUCUCUGCUUCUCUCCAGACACAAGAGGAAGCUCAAGGCCAUUGUGGCUGGCUCAGCUGGUAAGUGAGGGCCAUAGUGGGGACA
Intron
Exon
Intron
−mean(ΔLogitPSI$_{sub}$)
4
2
0

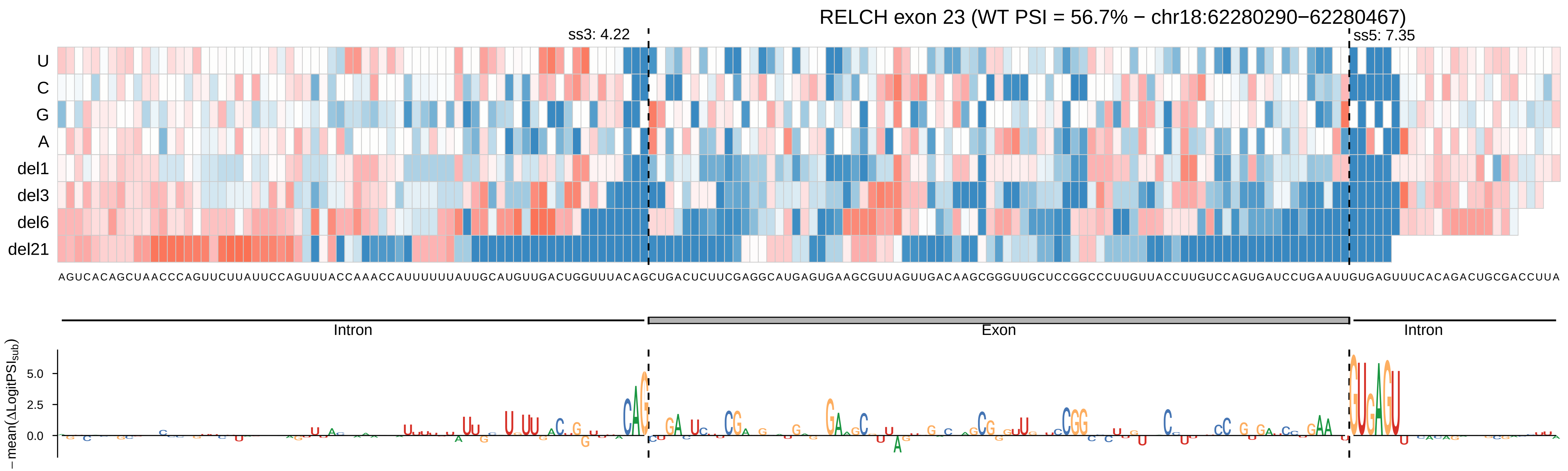

RELCH exon 23 (WT PSI = 56.7% − chr18:62280290−62280467)
ss3: 4.22
ss5: 7.35
U
C
G
A
del1
del3
del6
del21
AGUCACAGCUAACCCAGUUCUUAUUCCAGUUUACCAAACCAUUUUUUAUUGCAUGUUGACUGGUUUACAGCUGACUCUUCGAGGCAUGAGUGAAGCGUUAGUUGACAAGCGGGUUGCUCCGGCCCUUGUUACCUUGUCCAGUGAUCCUGAAUUGUGAGUUUCACAGACUGCGACCUUA
Intron
Exon
Intron
−mean(ΔLogitPSI_sub)
5.0
2.5
0.0

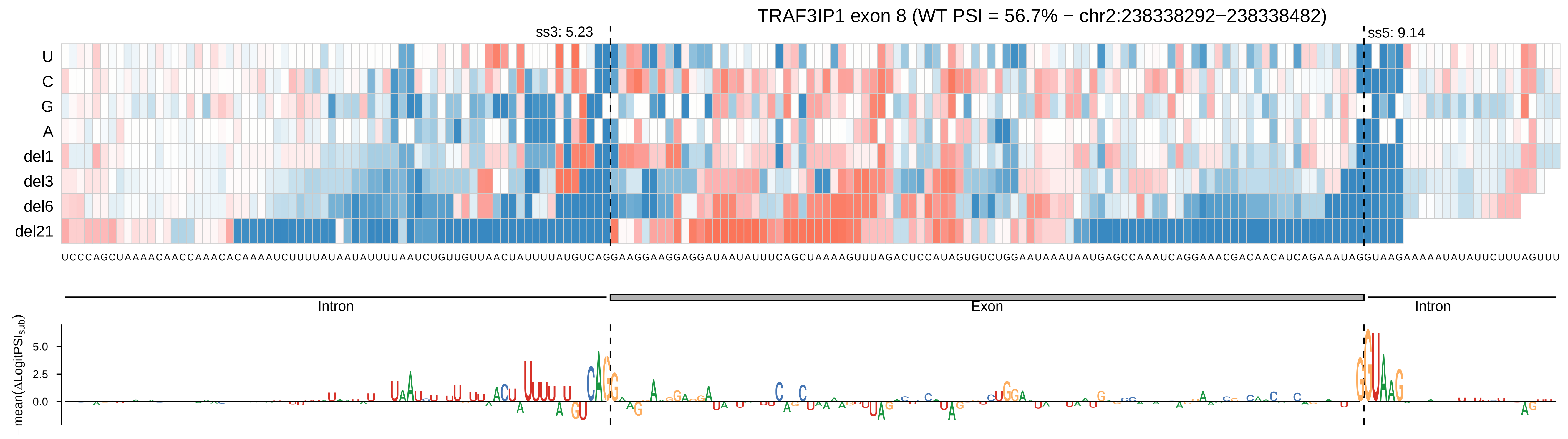

TRAF3IP1 exon 8 (WT PSI = 56.7% − chr2:238338292−238338482)
ss3: 5.23
ss5: 9.14
U
C
G
A
del1
del3
del6
del21
UCCCAGCUAAAACAACCAAACACAAAAUCUUUUAUAUAUUUUAAUCUGUUGUUAACUAUUUUAUGUCAGGAAGGAAGGAGGAUAAUAUUUCAGCUAAAAGUUUAGACUCCAUAGUGUCUGGAAUAAAUAAUGAGCCAAAUCAGGAAACGACAACAUCAGAAAUAGGUAAGAAAAAUAUAUUCUUUAGUUU
Intron
Exon
Intron
−mean(ΔLogitPSI_sub)
5.0
2.5
0.0

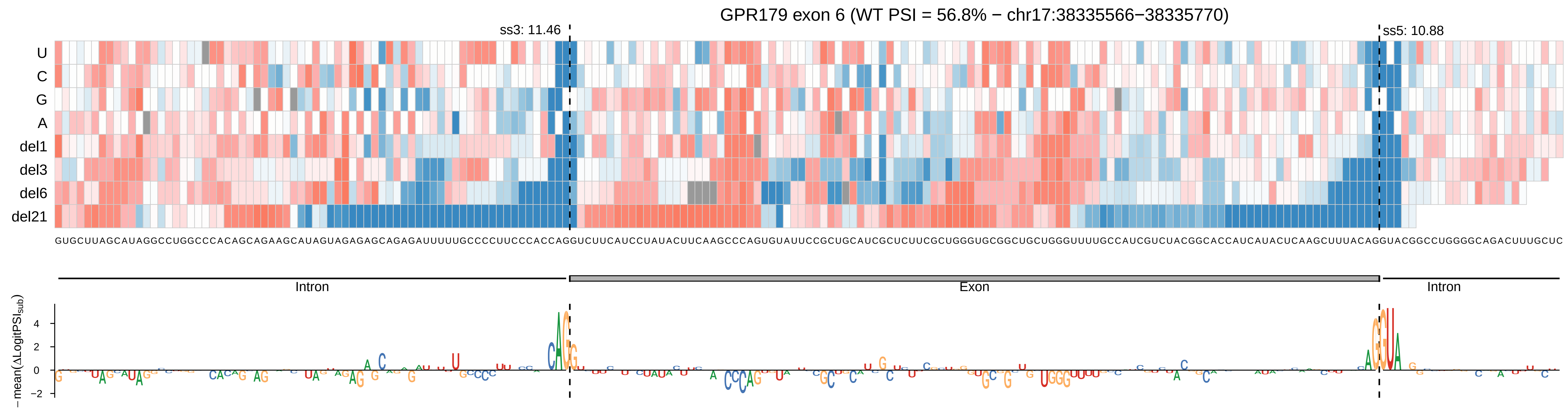

GPR179 exon 6 (WT PSI = 56.8% − chr17:38335566−38335770)
ss3: 11.46
ss5: 10.88
U
C
G
A
del1
del3
del6
del21
Intron
Exon
Intron
− mean(ΔLogitPSI_sub)
4
2
0
−2

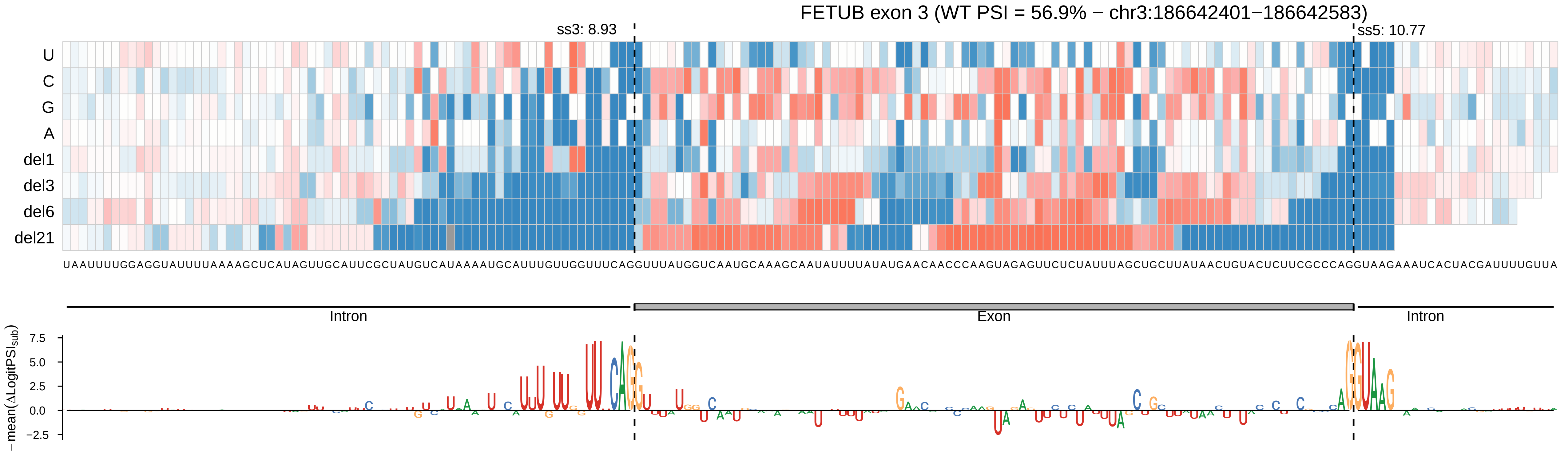

FETUB exon 3 (WT PSI = 56.9% − chr3:186642401−186642583)
ss3: 8.93
ss5: 10.77
U
C
G
A
del1
del3
del6
del21
UAAUUUUGGAGGUAUUUUAAAAGCUCAUAGUUGCAUUCGCUAUGUCAUAAAAUGCAUUUGUUGGUUUCAGGUUUAUGGUCAAUGCAAAGCAAUAUUUUAUAUGAACAACCCAAGUAGAGUUCUCUAUUUAGCUGCUUAUAACUGUACUCUUCGCCCAGGUAAGAAAUCACUACGAUUUUGUUA
Intron
Exon
Intron
−mean(ΔLogitPSI$_{sub}$)
7.5
5.0
2.5
0.0
−2.5

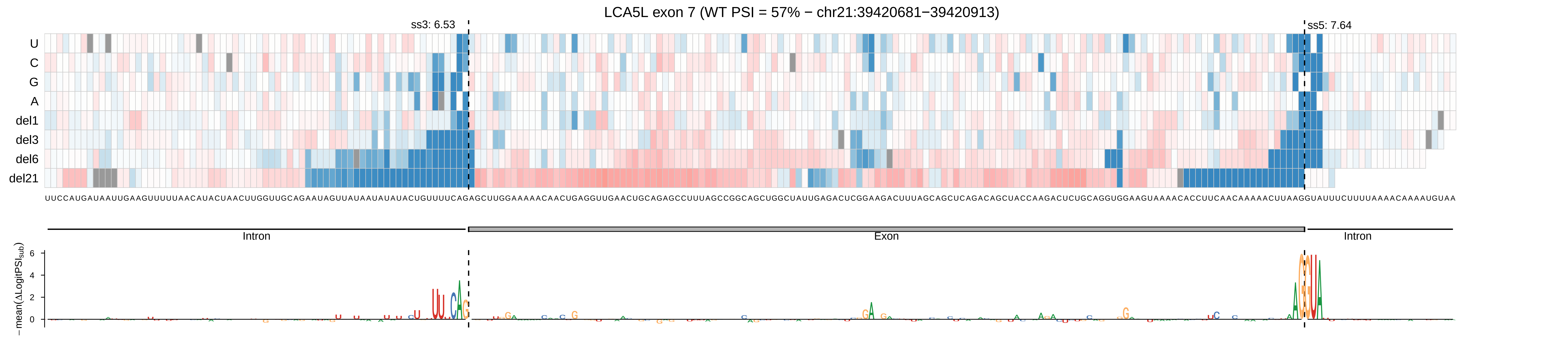

LCA5L exon 7 (WT PSI = 57% − chr21:39420681−39420913)
ss3: 6.53
ss5: 7.64
U
C
G
A
del1
del3
del6
del21
Intron
Exon
Intron
−mean(ΔLogitPSI_sub)

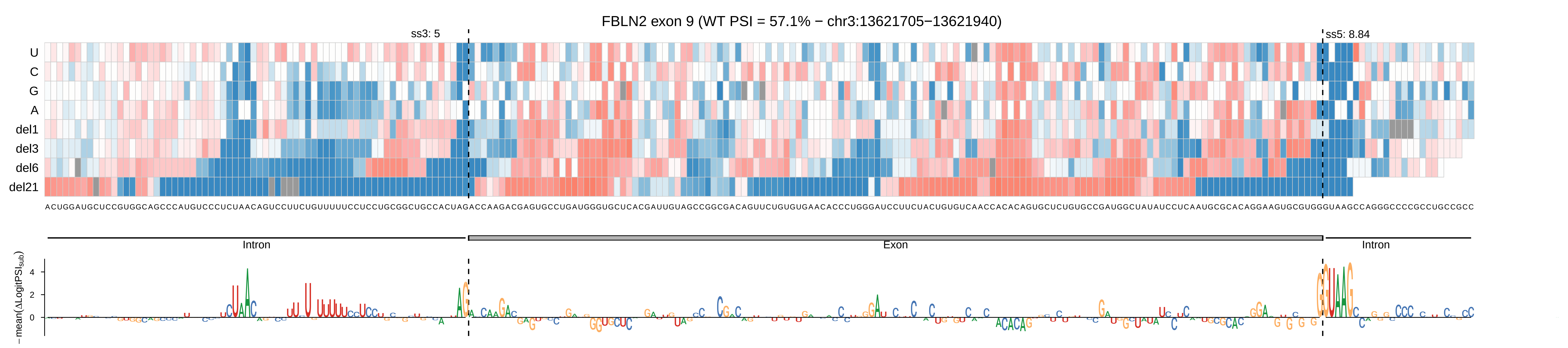
FBLN2 exon 9 (WT PSI = 57.1% − chr3:13621705−13621940)
ss3: 5
ss5: 8.84
U
C
G
A
del1
del3
del6
del21
ACUGGAUGCUCCGUGGCAGCCCAUGUCCCUCUAACAGUCCUUCUGUUUUUCCUCCUGCGGCUGCCACUAGACCAAGACGAGUGCCUGAUGGGUGCUCACGAUUGUAGCCGGCGACAGUUCUGUGUGAACACCCUGGGAUCCUUCUACUGUGUCAACCACACAGUGCUCUGUGCCGAUGGCUAUAUCCUCAAUGCGCACAGGAAGUGCGUGGGUAAGCCAGGCCCCGCCUGCCGCC
Intron
Exon
Intron
−mean(ΔLogitPSI_sub)
4
2
0

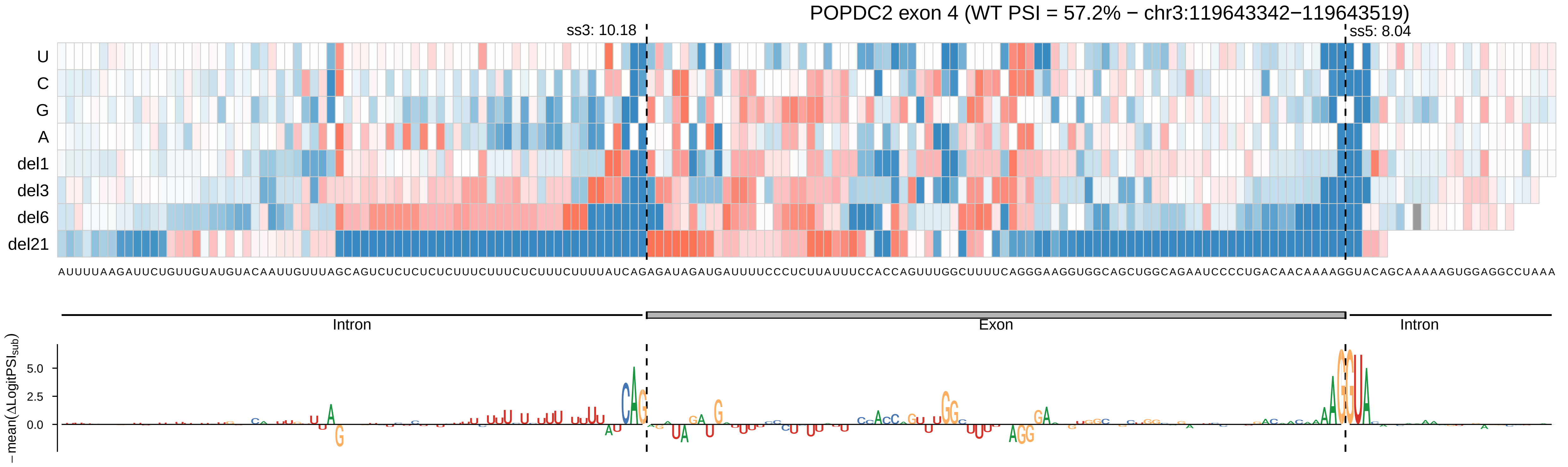
POPDC2 exon 4 (WT PSI = 57.2% − chr3:119643342−119643519)
ss3: 10.18
ss5: 8.04
U
C
G
A
del1
del3
del6
del21
AUUUUAAGAUUCUGUUGUAUGUACAAUUGUUUAGCAGUCUCUCUCUUUCUUUCUCUUUCUUUUAUCAGAGAUAGAUGAUUUUCCCUCUUAUUUCCACCAGUUUGGCUUUUCAGGGAAGGUGGCAGCUGGCAGAAUCCCCUGACAACAAAAGGUACAGCAAAAAGUGGAGGCCUAAA
Intron
Exon
Intron
−mean(ΔLogitPSI_sub)
5.0
2.5
0.0

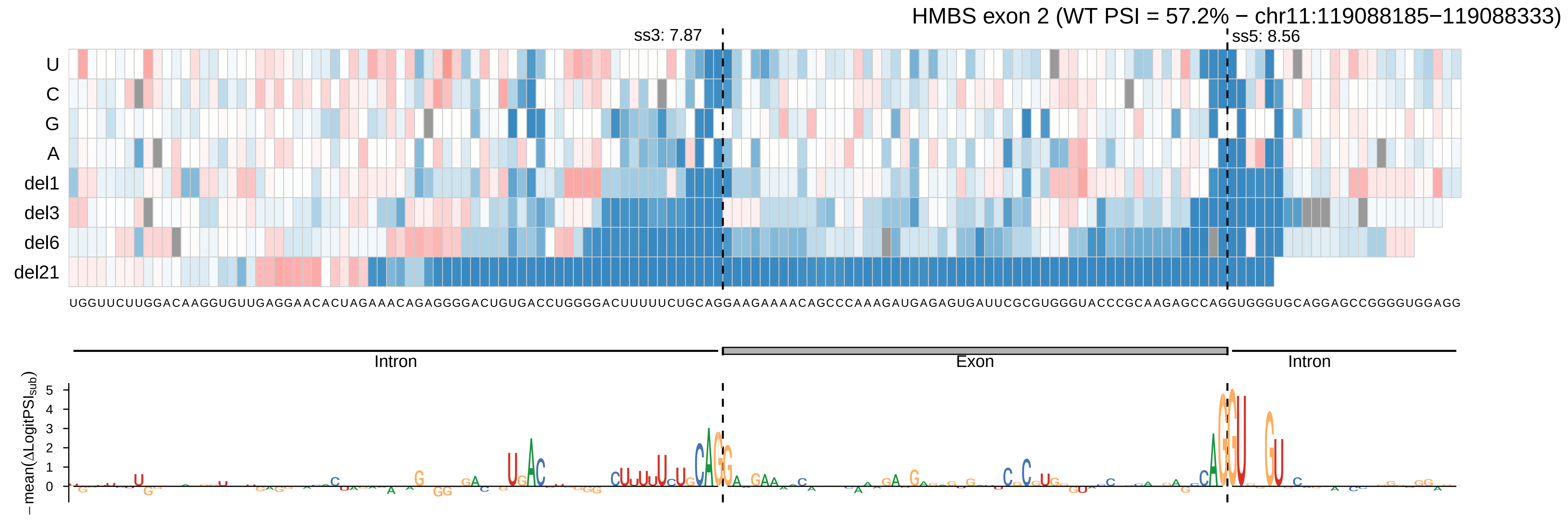
HMBS exon 2 (WT PSI = 57.2% − chr11:119088185−119088333)
ss3: 7.87
ss5: 8.56
U
C
G
A
del1
del3
del6
del21
UGGUUCUUGGACAAGGUGUUGAGGAACACUAGAAACAGAGGGGACUGUGACCUGGGGACUUUUUCUGCAGGAAGAAAACAGCCCAAAGAUGAGAGUGAUUCGCGUGGGUACCCGCAAGAGCCAGGUGGGUGCAGGAGCCGGGGUGGAGG
Intron
Exon
Intron
−mean(ΔLogitPSI_sub)
5
4
3
2
1
0

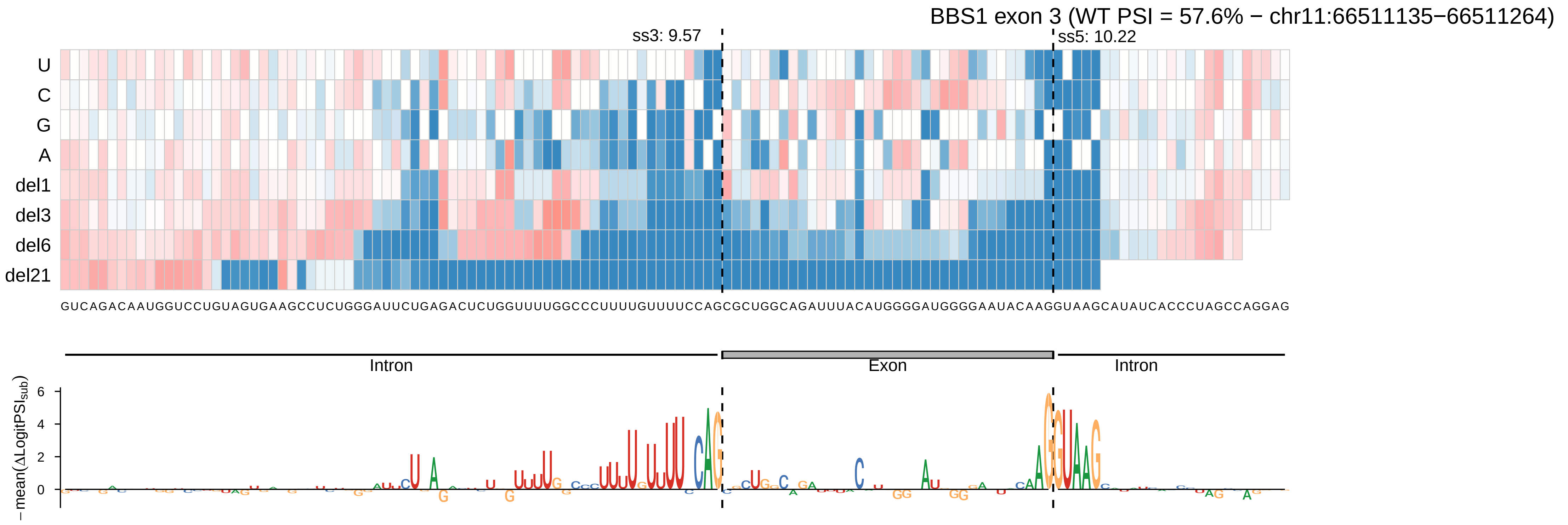

BBS1 exon 3 (WT PSI = 57.6% − chr11:66511135−66511264)
ss3: 9.57
ss5: 10.22
U
C
G
A
del1
del3
del6
del21
GUCAGACAAUGGUCCUGUAGUGAAGCCUCUGGGAUUCUGAGACUCUGGUUUUGGCCCUUUUGUUUUCCAGCGCUGGCAGAUUUACAUGGGGAUGGGGAAUACAAGGUAAGCAUAUCACCCUAGCCAGGAG
Intron
Exon
Intron
−mean(ΔLogitPSI_sub)
6
4
2
0

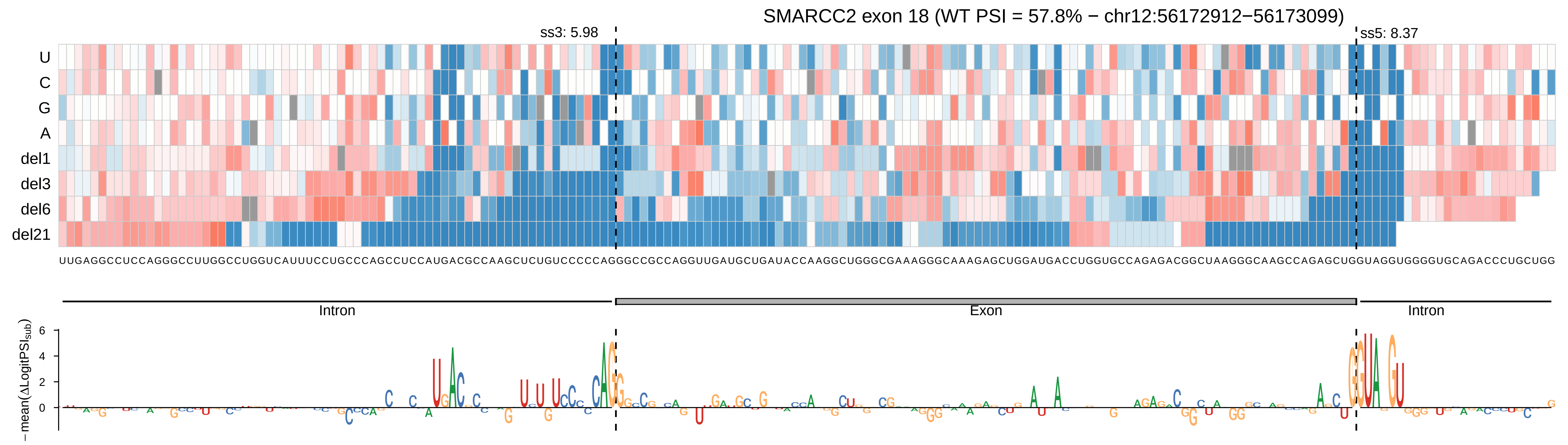
SMARCC2 exon 18 (WT PSI = 57.8% − chr12:56172912−56173099)
ss3: 5.98
ss5: 8.37
U
C
G
A
del1
del3
del6
del21
UUGAGGCCUCCAGGGCCUUGGCCUGGUCAUUUCCUGCCCAGCCUCCAUGACGCCAAGCUCUGUCCCCCAGGGCCGCCAGGUUGAUGCUGAUACCAAGGCUGGGCGAAAGGGCAAAGAGCUGGAUGACCUGGUGCCAGAGACGGCUAAGGGCAAGCCAGAGCUGGUAGGUGGGGUGCAGACCCUGCUGG
Intron
Exon
Intron
−mean(ΔLogitPSI_sub)
6
4
2
0

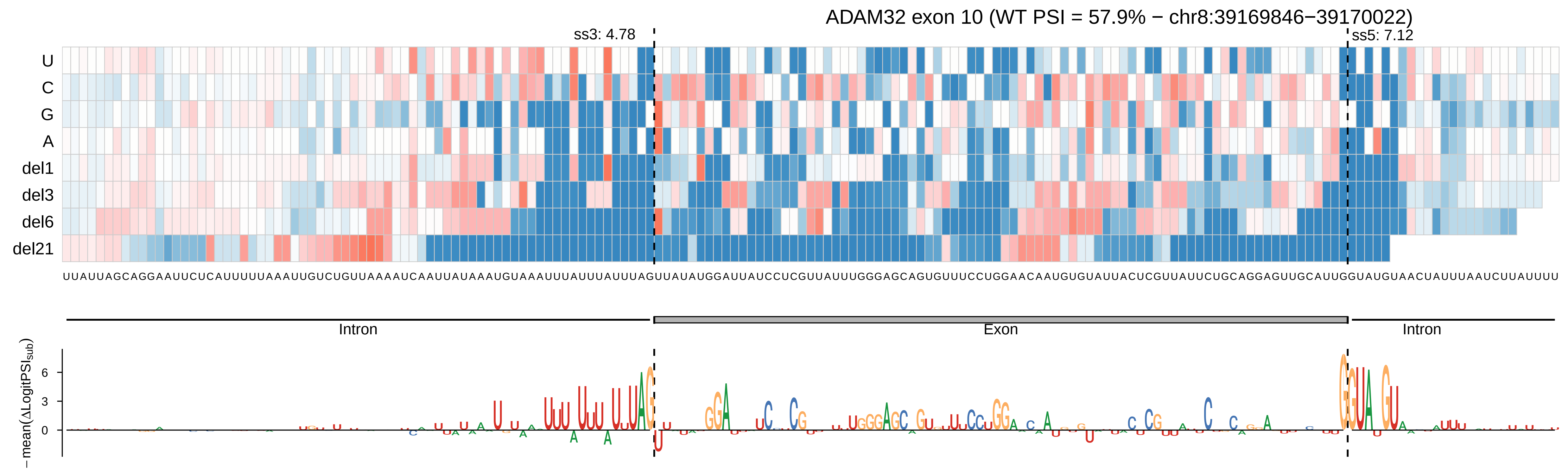

ADAM32 exon 10 (WT PSI = 57.9% − chr8:39169846−39170022)
ss3: 4.78
ss5: 7.12
U
C
G
A
del1
del3
del6
del21
Intron
Exon
Intron
−mean(ΔLogitPSI_sub)
6
3
0

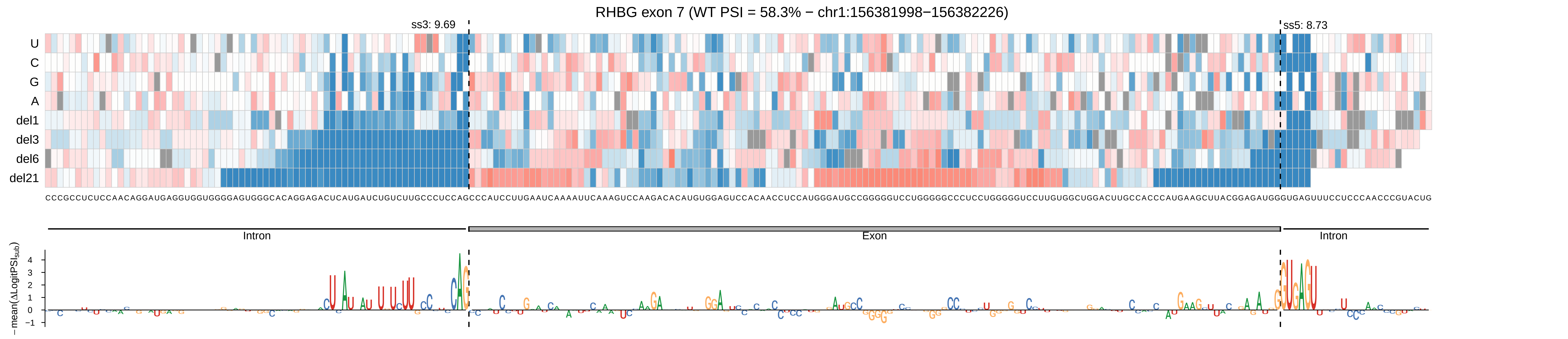

RHBG exon 7 (WT PSI = 58.3% − chr1:156381998−156382226)
ss3: 9.69
ss5: 8.73
U
C
G
A
del1
del3
del6
del21
Intron
Exon
Intron
−mean(ΔLogitPSI_sub)
4
3
2
1
0
−1

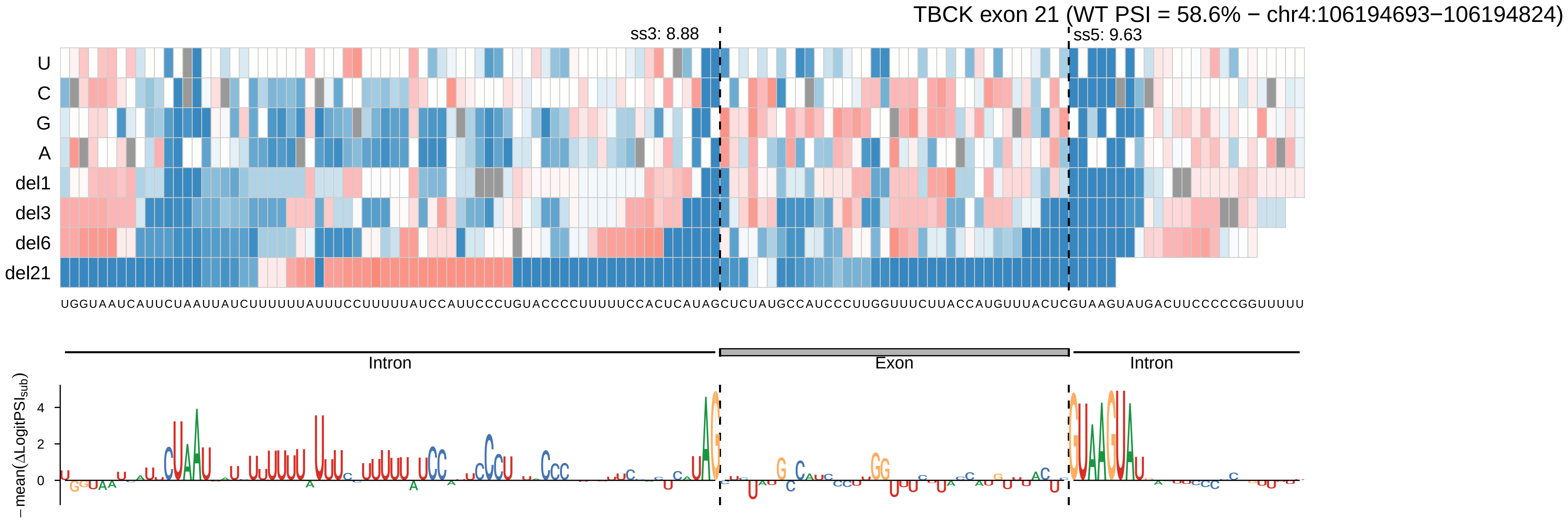
TBCK exon 21 (WT PSI = 58.6% − chr4:106194693−106194824)
ss3: 8.88
ss5: 9.63
U
C
G
A
del1
del3
del6
del21
UGGUAAUCAUUCUAAUUAUCUUUUUUAUUUCCUUUUUAUCCAUUCCCUGUACCCCUUUUUCCACUCAUAGCUCUAUGCCAUCCCUUGGUUUCUUACCAUGUUUACUCGUAAGUAUGACUUCCCCCGGUUUUU
Intron
Exon
Intron
−mean(ΔLogitPSI$_{sub}$)
4
2
0

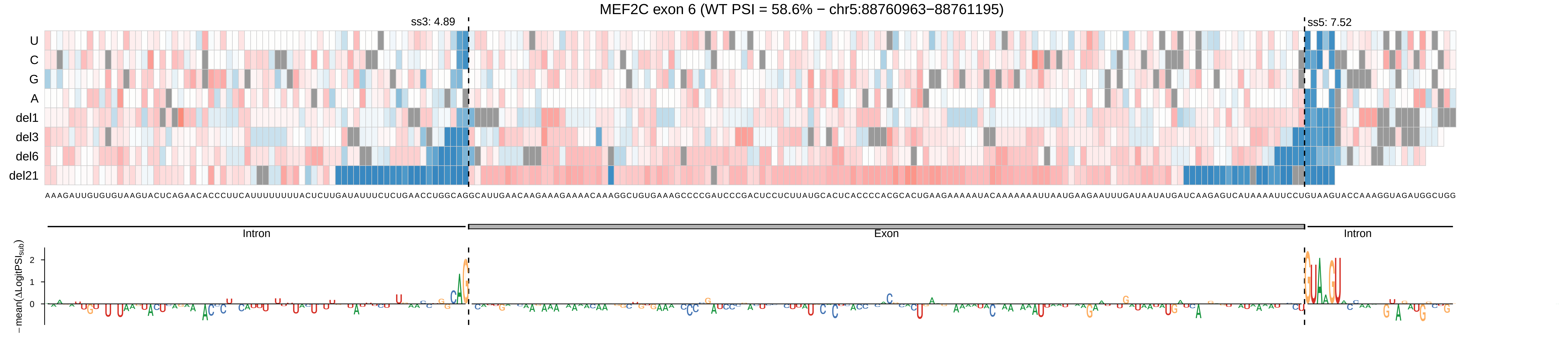
MEF2C exon 6 (WT PSI = 58.6% − chr5:88760963−88761195)
ss3: 4.89
ss5: 7.52
U
C
G
A
del1
del3
del6
del21
AAAGAUUGUGUGUAAGUACUCAGAACACCCUUCAUUUUUUUUACUCUUGAUAUUUCUCUGAACCUGGCAGGCAUUGAACAAGAAAGAAAACAAAGGCUGUGAAAGCCCCGAUCCCGACUCCUCUUAUGCACUCACCCCACGCACUGAAGAAAAAUACAAAAAAAUUAAUGAAGAAUUUGAUAAUAUGAUCAAGAGUCAUAAAAUUCCUGUAAGUACCAAAGGUAGAUGGCUGG
Intron
Exon
Intron
−mean(ΔLogitPSI_sub)
2
1
0

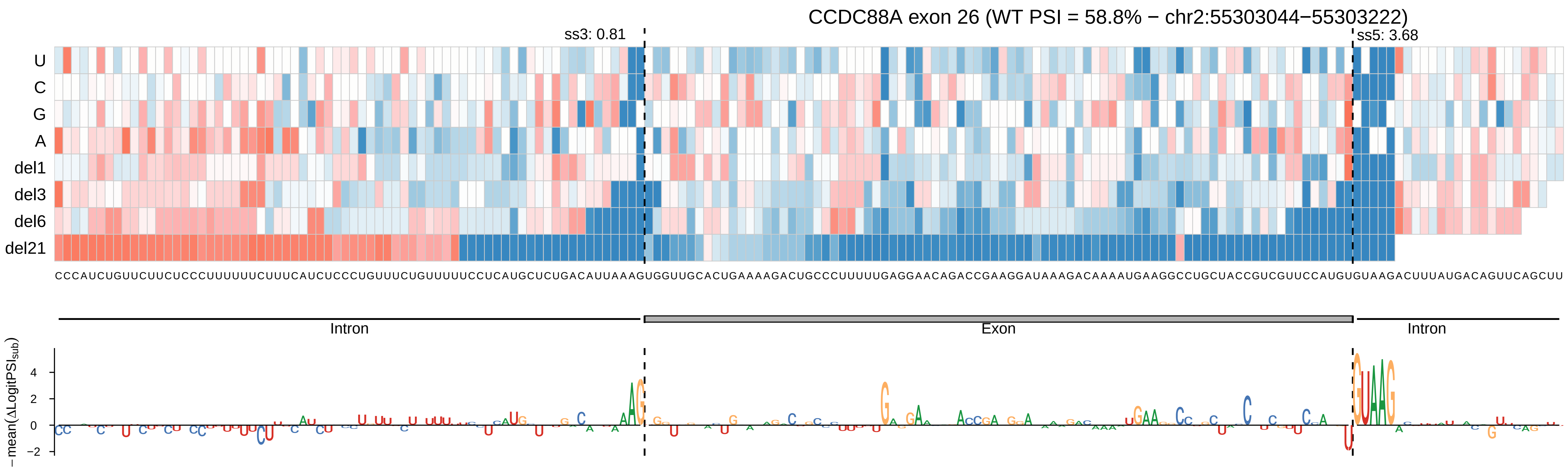

CCDC88A exon 26 (WT PSI = 58.8% − chr2:55303044−55303222)
ss3: 0.81
ss5: 3.68
U
C
G
A
del1
del3
del6
del21
Intron
Exon
Intron
−mean(ΔLogitPSI_sub)
4
2
0
−2

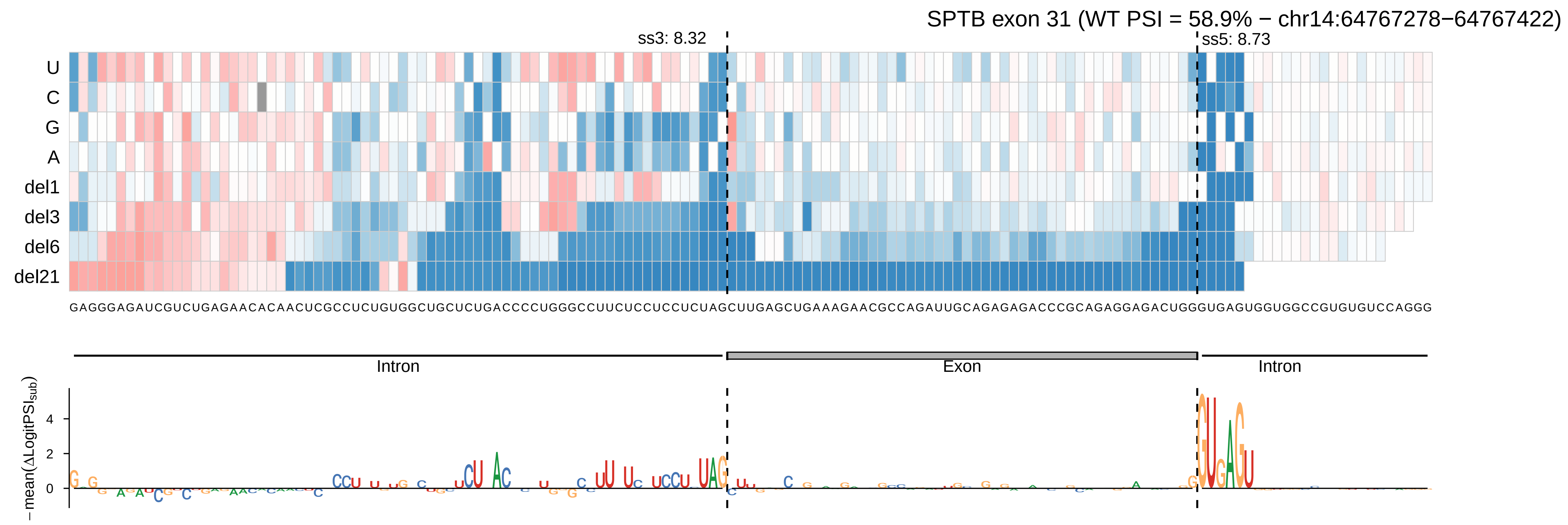

SPTB exon 31 (WT PSI = 58.9% − chr14:64767278−64767422)
ss3: 8.32
ss5: 8.73
U
C
G
A
del1
del3
del6
del21
GAGGGAGAUCGUCUGAGAACACAACUCGCCUCUGUGGCUGCUCUGACCCCUGGGCCUUCUCCUCCUCUAGCUUGAGCUGAAAGAACGCCAGAUUGCAGAGAGACCCGCAGAGGAGACUGGGUGAGUGGUGGCCGUGUGUCCAGGG
Intron
Exon
Intron
−mean(ΔLogitPSI_sub)
4
2
0

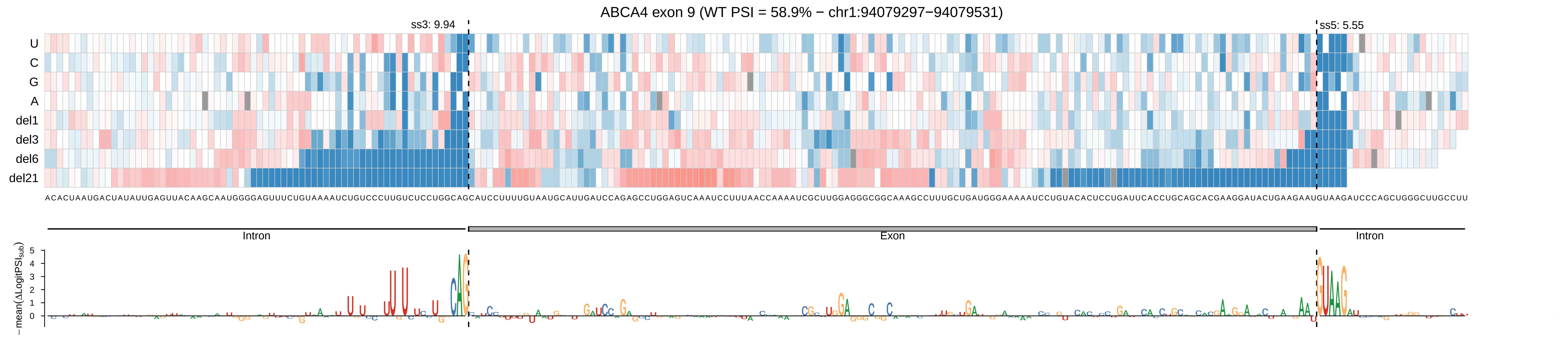

ABCA4 exon 9 (WT PSI = 58.9% − chr1:94079297−94079531)
ss3: 9.94
ss5: 5.55
U
C
G
A
del1
del3
del6
del21
ACACUAAUGACUAUAUUGAGUUACAAGCAAUGGGGAGUUUCUGUAAAAUCUGUCCCUUGUCUCCUGGCAGCAUCCUUUUGUAAUGCAUUGAUCCAGAGCCUGGAGUCAAAUCCUUUAACCAAAAUCGCUUGGAGGGCGGCAAAGCCUUUGCUGAUGGGAAAAAUCCUGUACACUCCUGAUUCACCUGCAGCACGAAGGAUACUGAAGAAUGUAAGAUCCCAGCUGGGCUUGCCUU
Intron
Exon
Intron
− mean(ΔLogitPSI$_{sub}$)
5
4
3
2
1
0

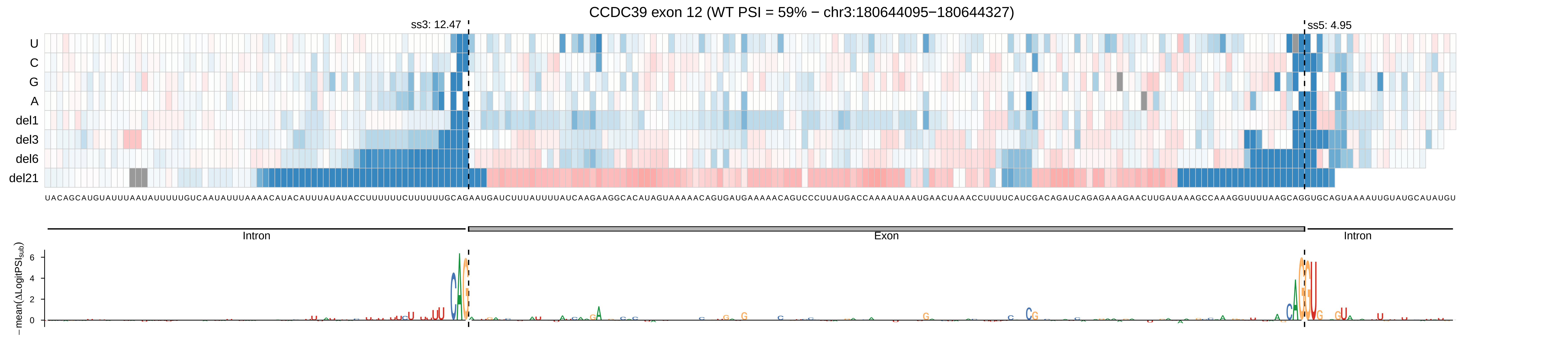

CCDC39 exon 12 (WT PSI = 59% − chr3:180644095−180644327)
ss3: 12.47
ss5: 4.95
U
C
G
A
del1
del3
del6
del21
Intron
Exon
Intron
−mean(ΔLogitPSI_sub)
6
4
2
0

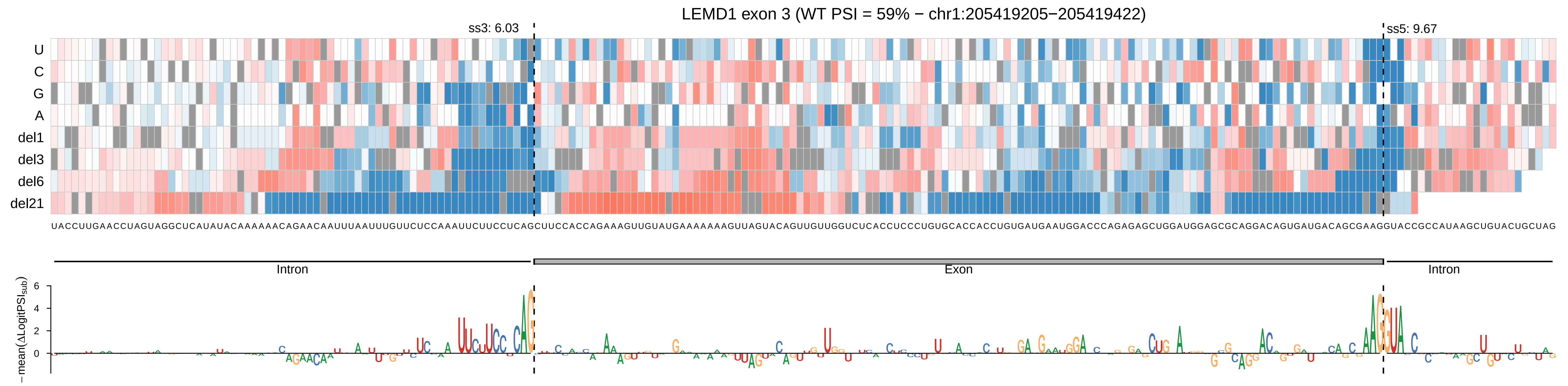
LEMD1 exon 3 (WT PSI = 59% − chr1:205419205−205419422)
ss3: 6.03
ss5: 9.67
U
C
G
A
del1
del3
del6
del21
UACCUUGAACCUAGUAGGCUCAUAUACAAAAAACAGAACAAUUUAAUUUGUUCUCCAAAUUCUUCCUCAGCUUCCACCAGAAAGUUGUAUGAAAAAAAGUUAGUACAGUUGUUGGUCUCACCUCCCUGUGCACCACCUGUGAUGAAUGGACCCAGAGAGCUGGAUGGAGCGCAGGACAGUGAUGACAGCGAAGGUACCGCCAUAAGCUGUACUGCUAG
Intron
Exon
Intron
−mean(ΔLogitPSI$_{sub}$)
6
4
2
0

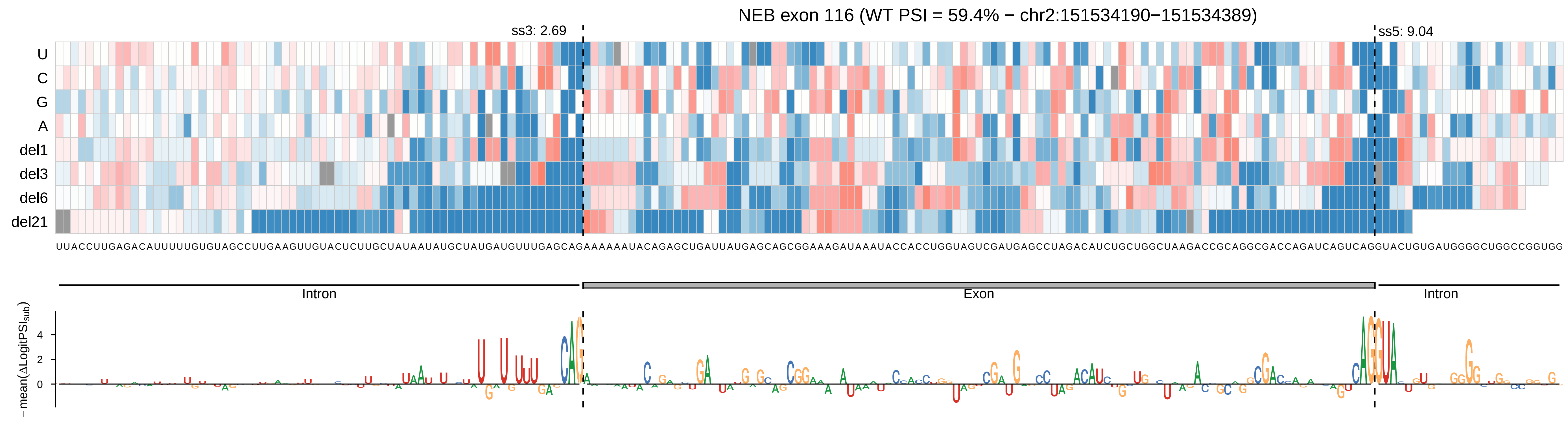
NEB exon 116 (WT PSI = 59.4% − chr2:151534190−151534389)
ss3: 2.69
ss5: 9.04
U
C
G
A
del1
del3
del6
del21
UUACCUUGAGACAUUUUUGUGUAGCCUUGAAGUUGUACUCUUGCUAUAAUAUGCUAUGAUGUUUGAGCAGAAAAAAUACAGAGCUGAUUAUGAGCAGCGGAAAGAUAAAUACCACCUGGUAGUCGAUGAGCCUAGACAUCUGCUGGCUAAGACCGCAGGCGACCAGAUCAGUCAGGUACUGUGAUGGGGCUGGCCGGUGG
Intron
Exon
Intron
−mean(ΔLogitPSI_sub)
4
2
0

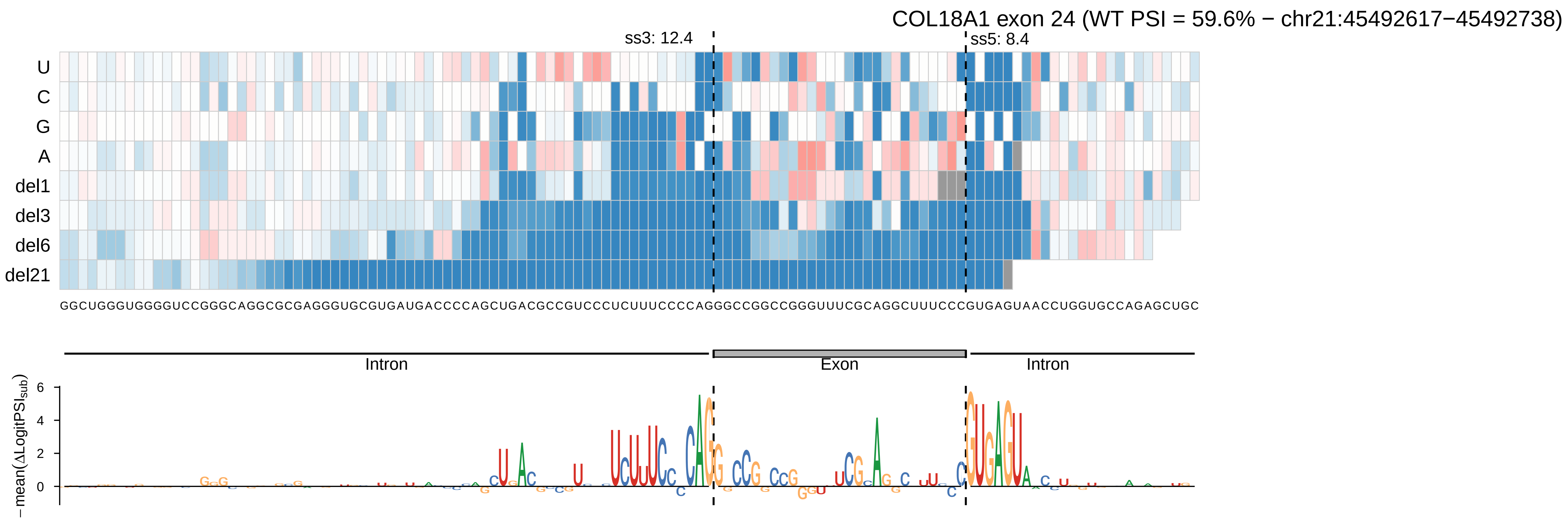

COL18A1 exon 24 (WT PSI = 59.6% − chr21:45492617−45492738)
ss3: 12.4
ss5: 8.4
U
C
G
A
del1
del3
del6
del21
GGCUGGGUGGGUCCGGGCAGGCGCGAGGGUGCGUGAUGACCCCAGCUGACGCCGUCCCUCUUUCCCCAGGGCCGGCCGGGUUUCGCAGGCUUUCCCGUGAGUAACCUGGUGCCAGAGCUGC
Intron
Exon
Intron
−mean(ΔLogitPSI_sub)
6
4
2
0

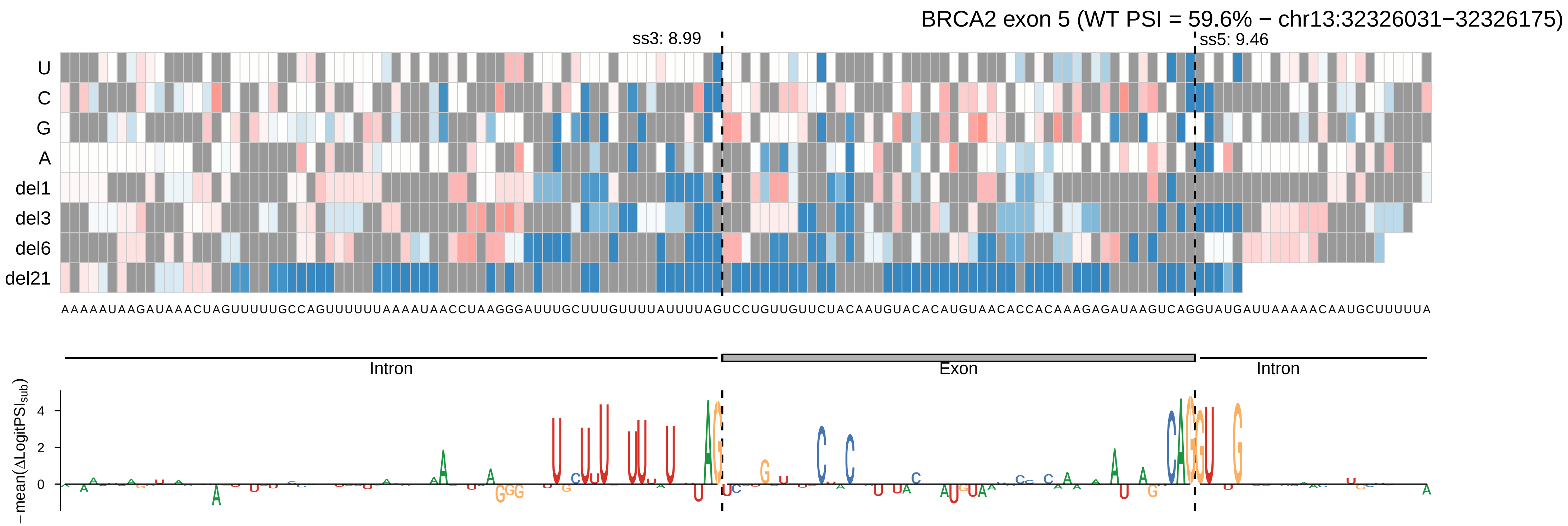
BRCA2 exon 5 (WT PSI = 59.6% − chr13:32326031−32326175)
ss3: 8.99
ss5: 9.46
U
C
G
A
del1
del3
del6
del21
Intron
Exon
Intron
−mean(ΔLogitPSI_sub)

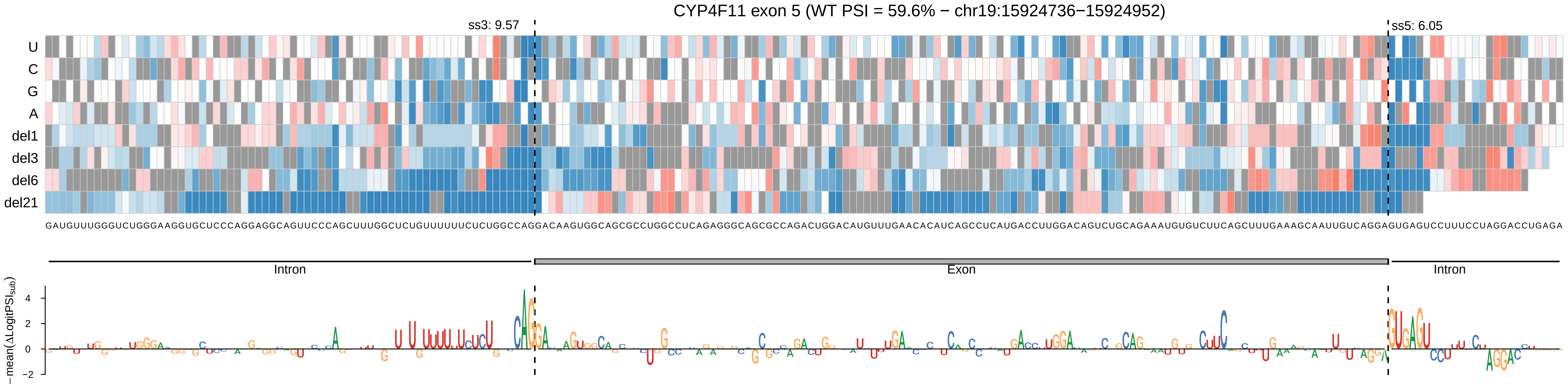

CYP4F11 exon 5 (WT PSI = 59.6% − chr19:15924736−15924952)
ss3: 9.57
ss5: 6.05
U
C
G
A
del1
del3
del6
del21
GAUGUUUGGGUCUGGGAAGGUGCUCCCAGGAGGCAGUUCCCAGCUUUGGCUCUGUUUUUUCUCUGGCCAGGACAAGUGGCAGCGCCUGGCCUCAGAGGGCAGCGCCAGACUGGACAUGUUUGAACACAUCAGCCUCAUGACCUUGGACAGUCUGCAGAAAUGUGUCUUCAGCUUUGAAAGCAAUUGUCAGGAGUGAGUCCUUUCCUAGGACCUGAGA
Intron
Exon
Intron
−mean(ΔLogitPSI_sub)

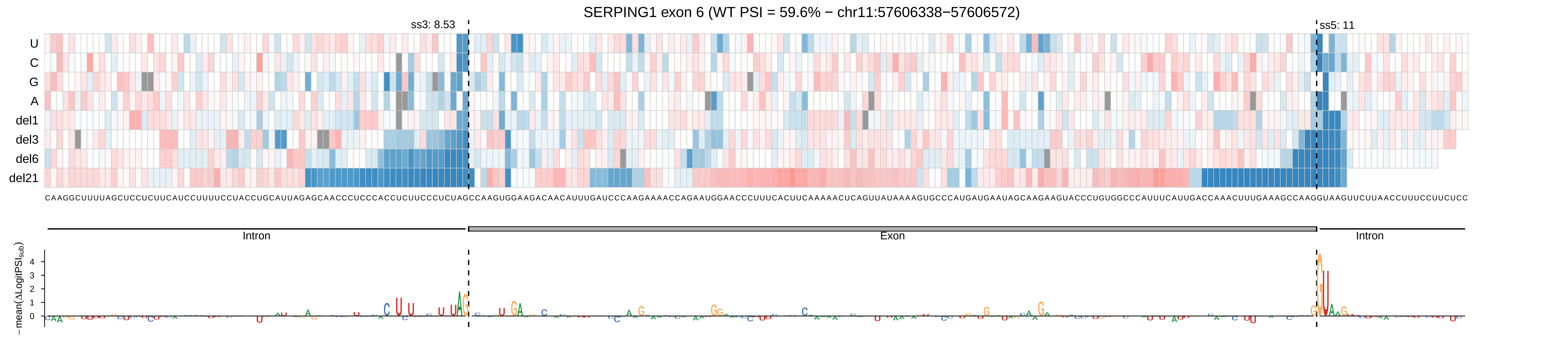
SERPING1 exon 6 (WT PSI = 59.6% − chr11:57606338−57606572)
ss3: 8.53
ss5: 11
U
C
G
A
del1
del3
del6
del21
CAAGGCUUUUAGCUCCUCUUCAUCCUUUUCCUACCUGCAUUAGAGCAACCCUCCCACCUCUUCCCUCUAGCCAAGUGGAAGACAACAUUUGAUCCCAAGAAAACCAGAAUGGAACCCUUUCACUUCAAAAACUCAGUUAUAAAAGUGCCCAUGAUGAAUAGCAAGAAGUACCCUGUGGCCCAUUUCAUUGACCAAACUUUGAAAGCCAAGGUAAGUUCUUAACCUUUCCUUCUCC
Intron
Exon
Intron
− mean(ΔLogitPSI_sub)
4
3
2
1
0

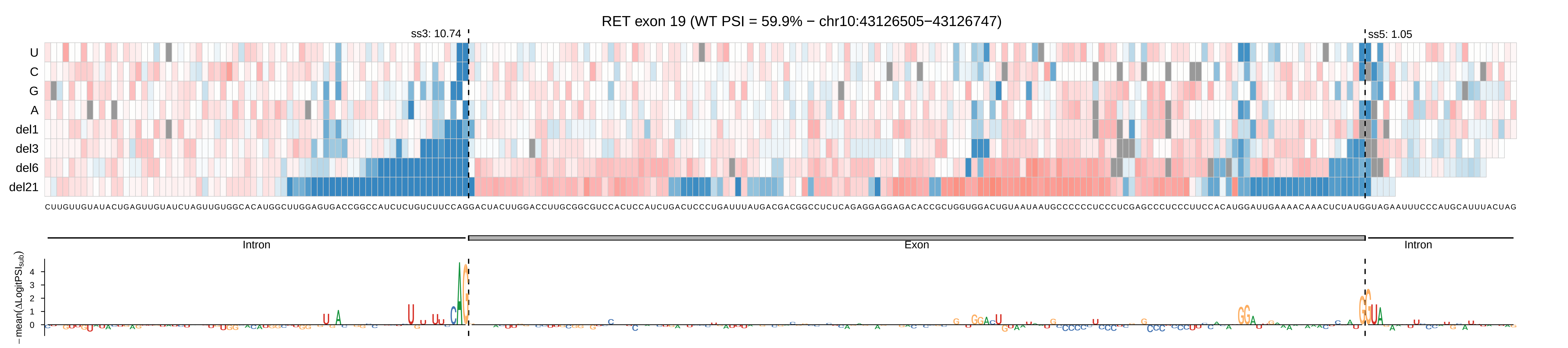
RET exon 19 (WT PSI = 59.9% − chr10:43126505−43126747)
ss3: 10.74
ss5: 1.05
U
C
G
A
del1
del3
del6
del21
Intron
Exon
Intron
− mean(ΔLogitPSI_sub)
4
3
2
1
0

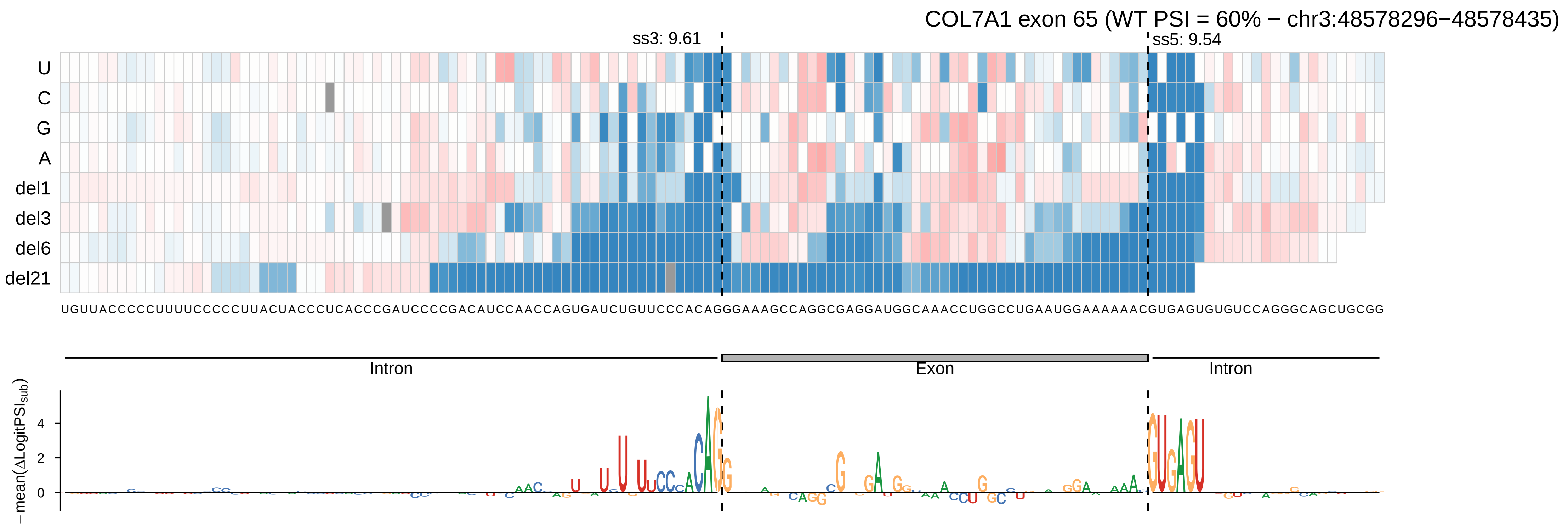
COL7A1 exon 65 (WT PSI = 60% − chr3:48578296−48578435)
ss3: 9.61
ss5: 9.54
U
C
G
A
del1
del3
del6
del21
UGUUACCCCCUUUUCCCCCUUACUACCCUCACCCGAUCCCCGACAUCCAACCAGUGAUCUGUUCCCACAGGGAAAGCCAGGCGAGGAUGGCAAACCUGGCCUGAAUGGAAAAAACGUGAGUGUGUCCAGGGCAGCUGCGG
Intron
Exon
Intron
−mean(ΔLogitPSIsub)
4
2
0

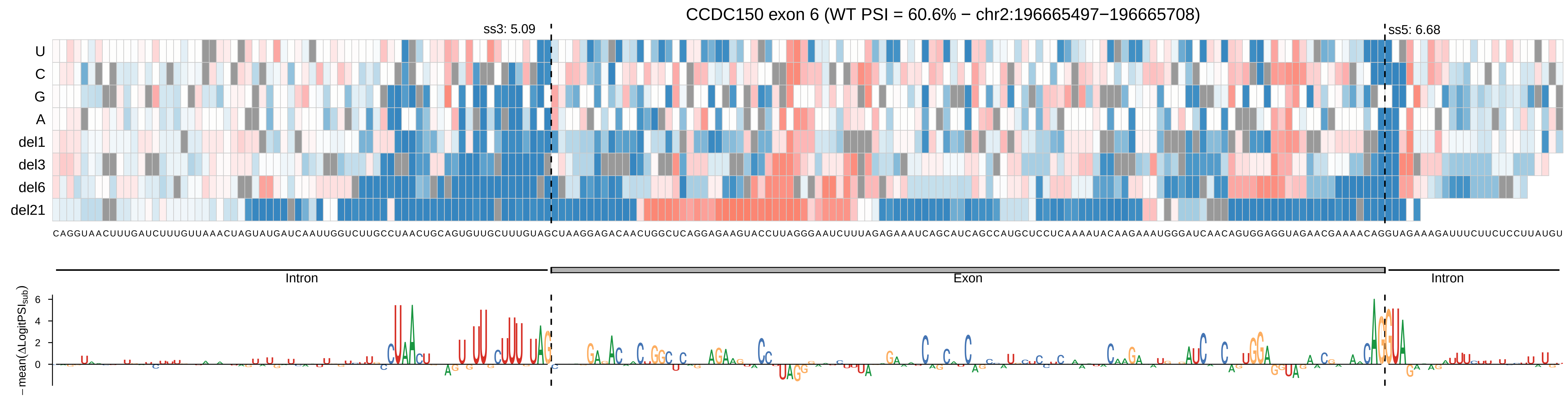

CCDC150 exon 6 (WT PSI = 60.6% − chr2:196665497−196665708)
ss3: 5.09
ss5: 6.68
U
C
G
A
del1
del3
del6
del21
CAGGUAACUUUGAUCUUUGUUAAACUAGUAUGAUCAAUUGGUCUUGCCUAACUGCAGUGUUGCUUUGUAGCUAAGGAGACAACUGGCUCAGGAGAAGUACCUUAGGGAAUCUUUAGAGAAAUCAGCAUCAGCCAUGCUCCUCAAAAUACAAGAAAUGGGAUCAACAGUGGAGGUAGAACGAAAACAGGUAGAAAGAUUUCUUCUCCUUAUGU
Intron
Exon
Intron
−mean(ΔLogitPSI$_{sub}$)

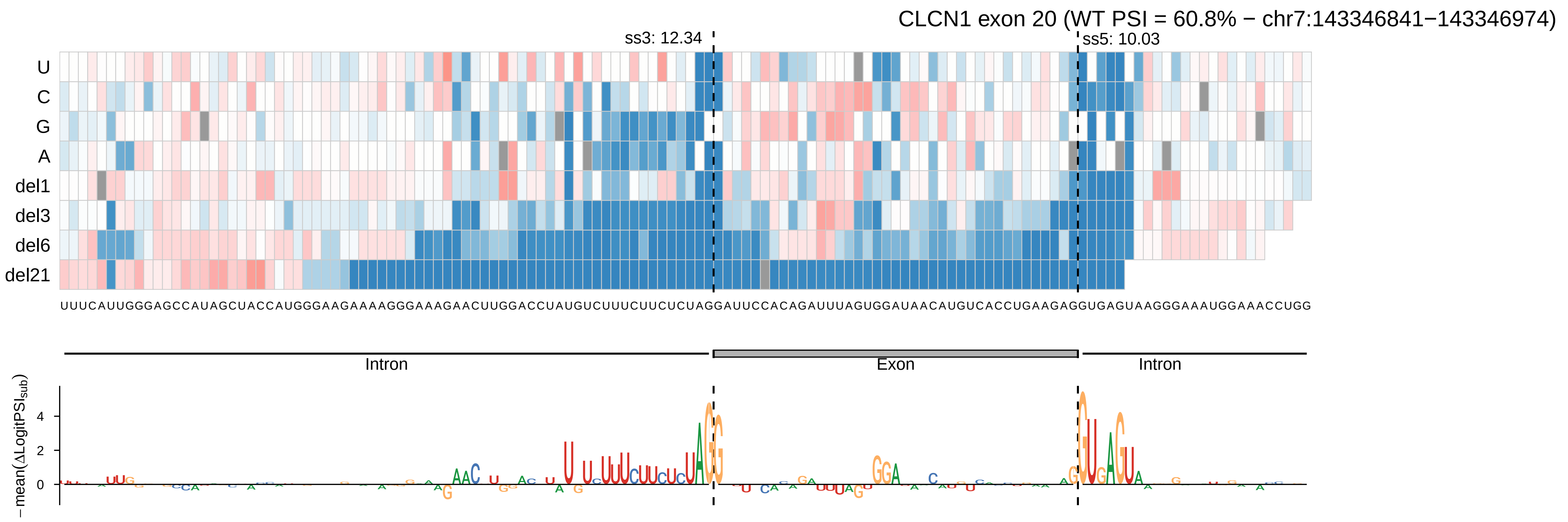

CLCN1 exon 20 (WT PSI = 60.8% − chr7:143346841−143346974)
ss3: 12.34
ss5: 10.03
U
C
G
A
del1
del3
del6
del21
UUUCAUUGGGAGCCAUAGCUACCAUGGGAAGAAAAGGGAAAGAACUUGGACCUAUGUCUUUCUUCUCUAGGAUUCCACAGAUUUAGUGGAUAACAUGUCACCUGAAGAGGUGAGUAAGGGAAAUGGAAACCUGG
Intron
Exon
Intron
−mean(ΔLogitPSI$_{sub}$)
4
2
0

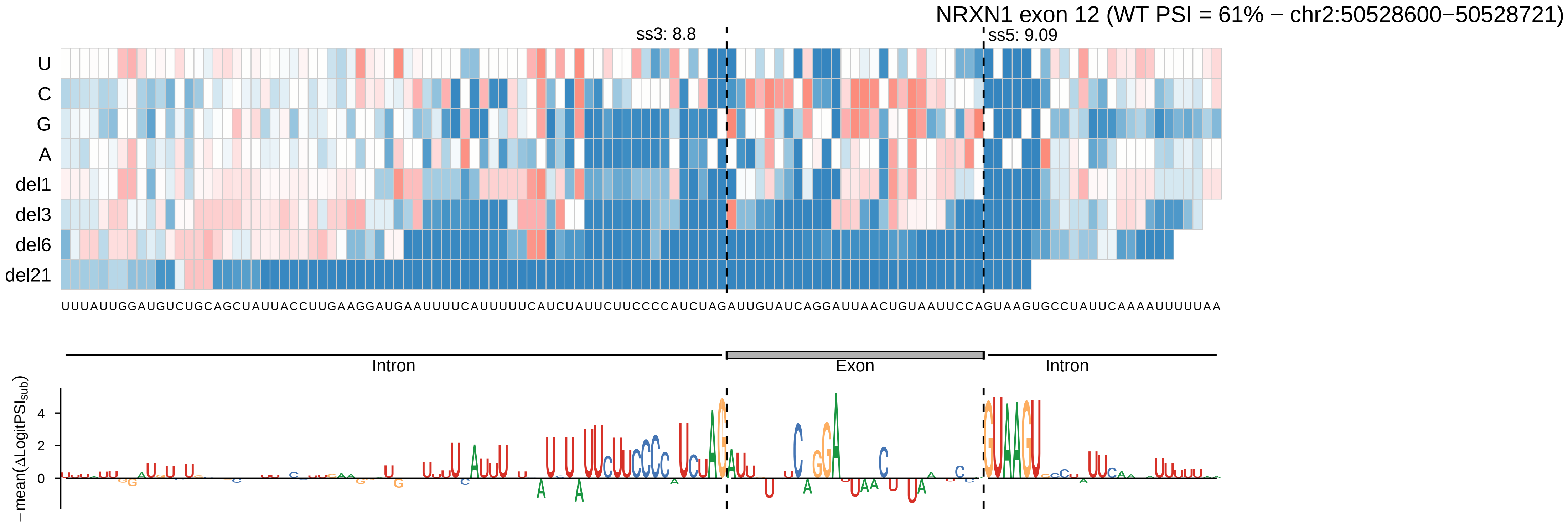
NRXN1 exon 12 (WT PSI = 61% − chr2:50528600−50528721)
ss3: 8.8
ss5: 9.09
U
C
G
A
del1
del3
del6
del21
UUUAUUGGAUGUCUGCAGCUAUUACCUUGAAGGAUGAAUUUUCAUUUUUCAUCUAUUCUUCCCCAUCUAGAUUGUAUCAGGAUUAACUGUAAUUCCAGUAAGUGCCUAUUCAAAAUUUUUAA
Intron
Exon
Intron
−mean(ΔLogitPSI_sub)
4
2
0

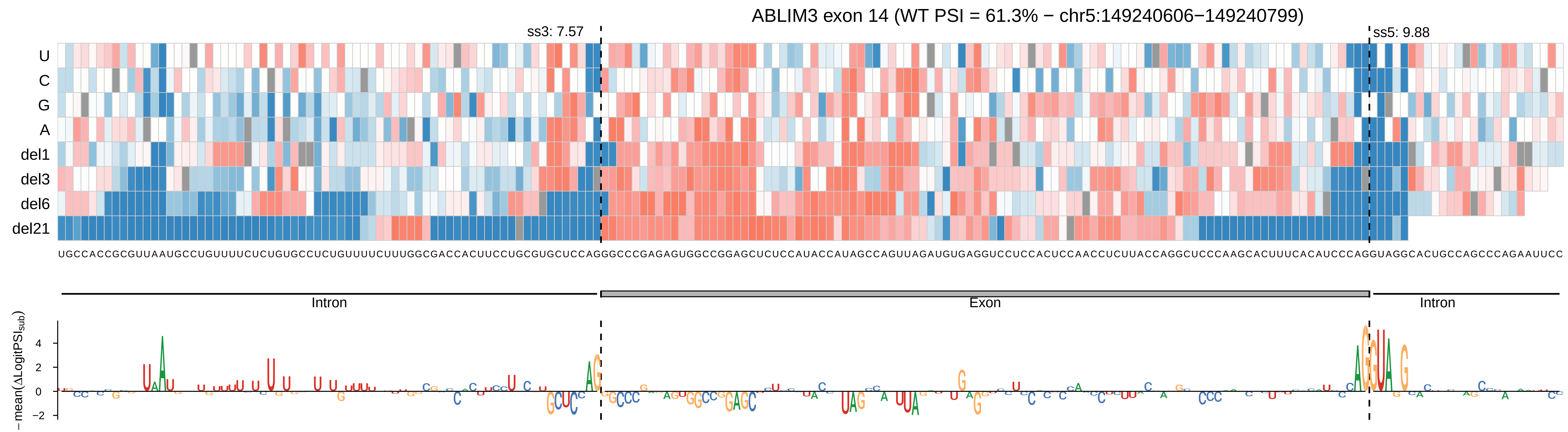

ABLIM3 exon 14 (WT PSI = 61.3% − chr5:149240606−149240799)
ss3: 7.57
ss5: 9.88
U
C
G
A
del1
del3
del6
del21
UGCCACCGCGUUAAUGCCUGUUUUCUCUGUGCCUCUGUUUUCUUUGGCGACCACUUCCUGCGUGCUCCAGGGCCCGAGAGUGGCCGGAGCUCUCCAUACCAUAGCCAGUUAGAUGUGAGGUCCUCCACUCCAACCUCUUACCAGGCUCCCAAGCACUUUCACAUCCCAGGUAGGCACUGCCAGCCCAGAAUUCC
Intron
Exon
Intron
− mean(ΔLogitPSI_sub)
4
2
0
−2

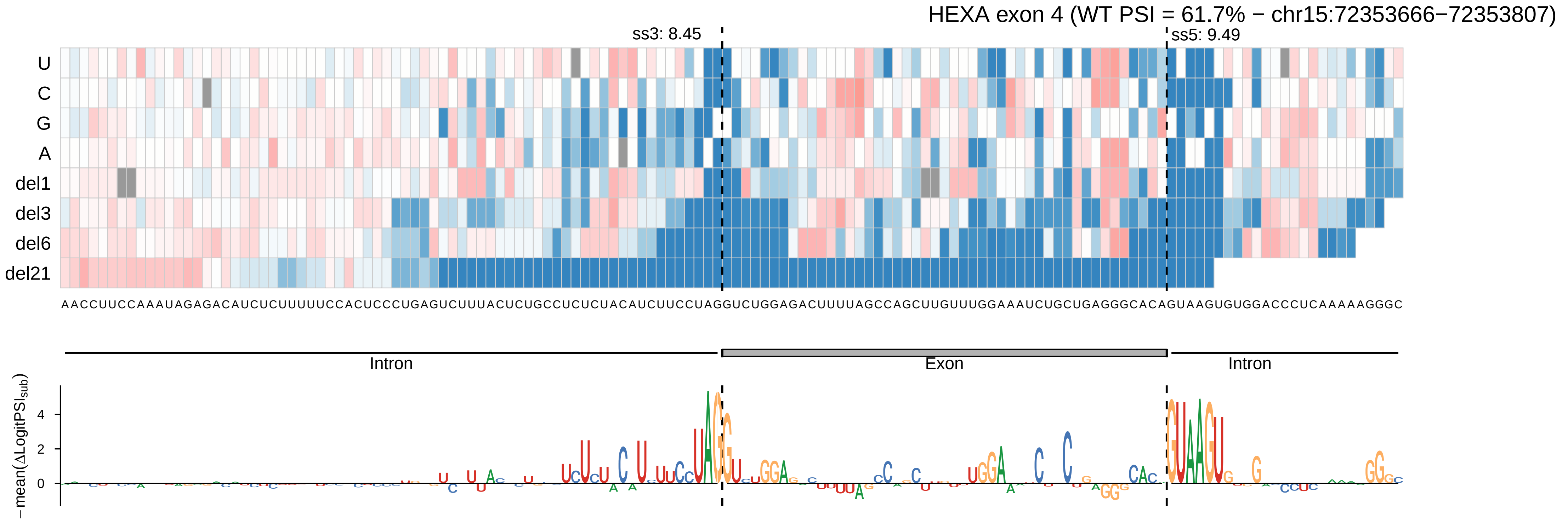

HEXA exon 4 (WT PSI = 61.7% − chr15:72353666−72353807)
ss3: 8.45
ss5: 9.49
U
C
G
A
del1
del3
del6
del21
AACCUUCCAAAUAGAGACAUCUCUUUUUCCACUCCCUGAGUCUUUACUCUGCCUCUCUACAUCUUCCUAGGUCUGGAGACUUUUAGCCAGCUUGUUUGGAAAUCUGCUGAGGGCACAGUAAGUGUGGACCCUCAAAAAGGGC
Intron
Exon
Intron
−mean(ΔLogitPSI_sub)
4
2
0

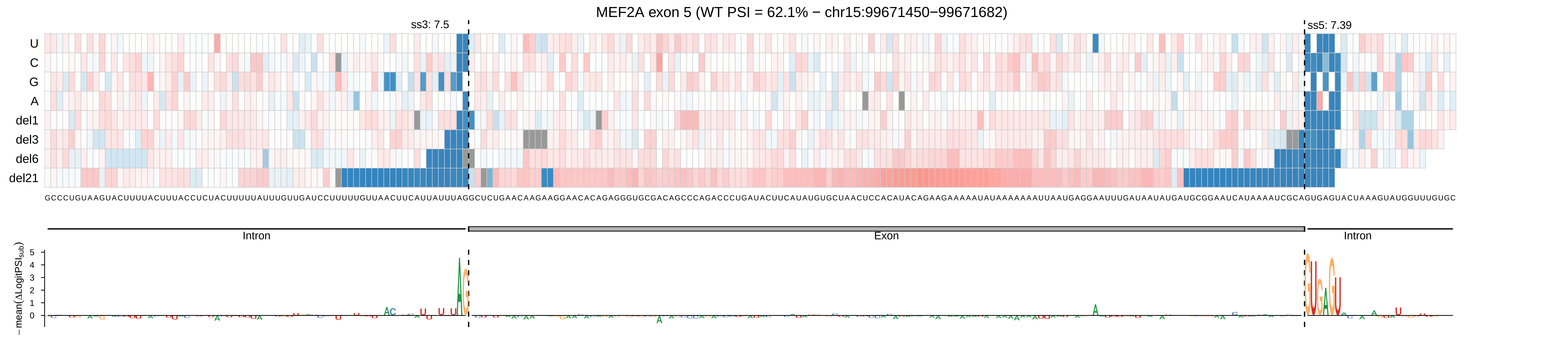

MEF2A exon 5 (WT PSI = 62.1% − chr15:99671450−99671682)
ss3: 7.5
ss5: 7.39
U
C
G
A
del1
del3
del6
del21
GCCCUGUAAGUACUUUUACUUUACCUCUACUUUUUAUUUGUUGAUCCUUUUUGUUAACUUCAUUAUUUAGGCUCUGAACAAGAAGGAACACAGAGGGUGCGACAGCCCAGACCCUGAUACUUCAUAUGUGCUAACUCCACAUACAGAAGAAAAAUAUAAAAAAAUUAAUGAGGAAUUUGAUAAUAUGAUGCGGAAUCAUAAAAUCGCAGUGAGUACUAAAGUAUGGUUUGUGC
Intron
Exon
Intron
− mean(ΔLogitPSI_sub)

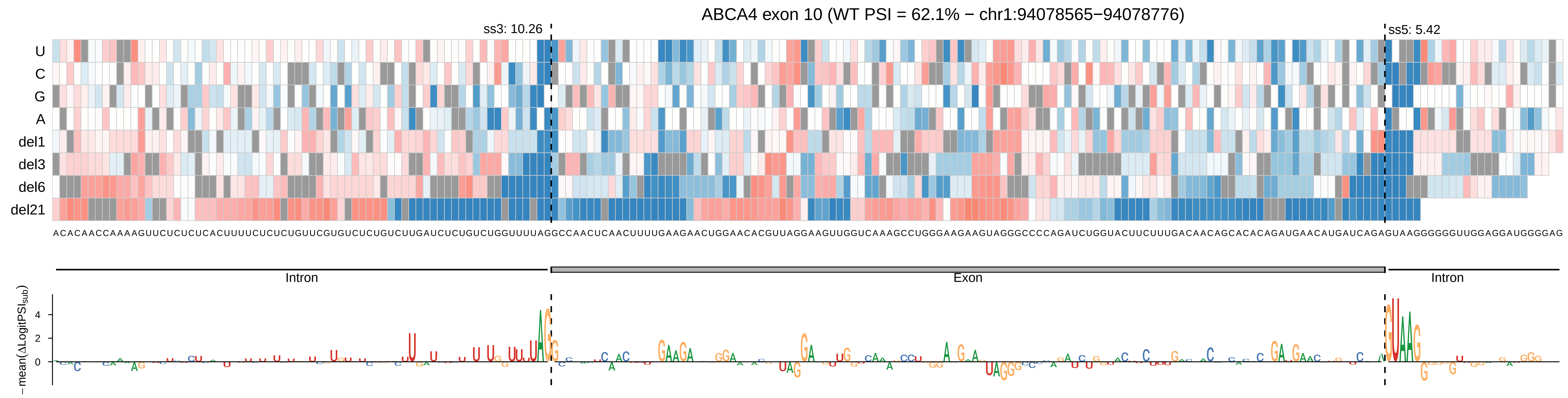

ABCA4 exon 10 (WT PSI = 62.1% − chr1:94078565−94078776)
ss3: 10.26
ss5: 5.42
U
C
G
A
del1
del3
del6
del21
ACACAACCAAAAGUUCUCUCUCACUUUUCUCUCUGUUCGUGUCUCUGUCUUGAUCUCUGUCUGGUUUUAGGCCAACUCAACUUUUGAAGAACUGGAACACGUUAGGAAGUUGGUCAAAGCCUGGGAAGAAGUAGGGCCCCAGAUCUGGUACUUCUUUGACAACAGCACACAGAUGAACAUGAUCAGAGUAAGGGGGGUUGGAGGAUGGGGAG
Intron
Exon
Intron
−mean(ΔLogitPSIsub)
4
2
0

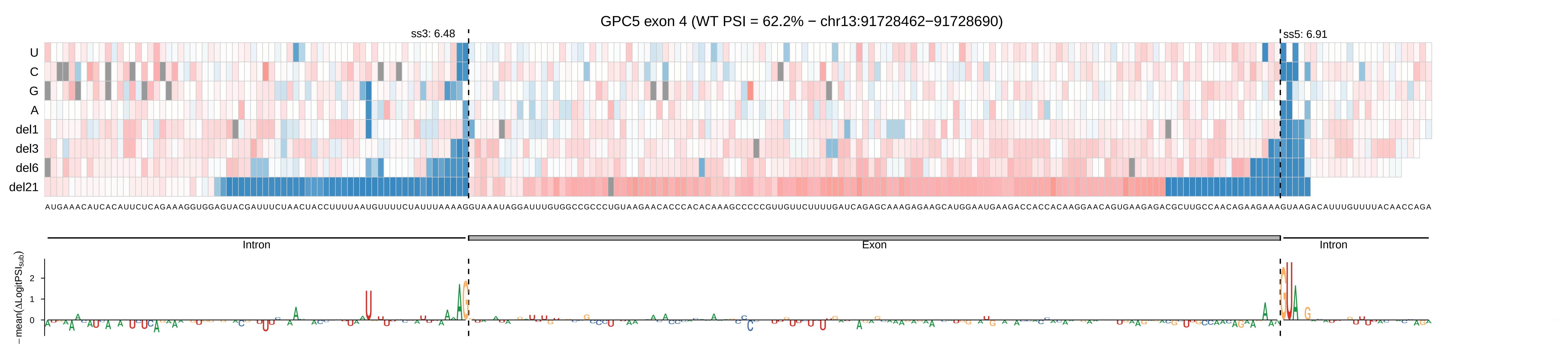
GPC5 exon 4 (WT PSI = 62.2% − chr13:91728462−91728690)
ss3: 6.48
ss5: 6.91
U
C
G
A
del1
del3
del6
del21
Intron
Exon
Intron
−mean(ΔLogitPSI_sub)
2
1
0

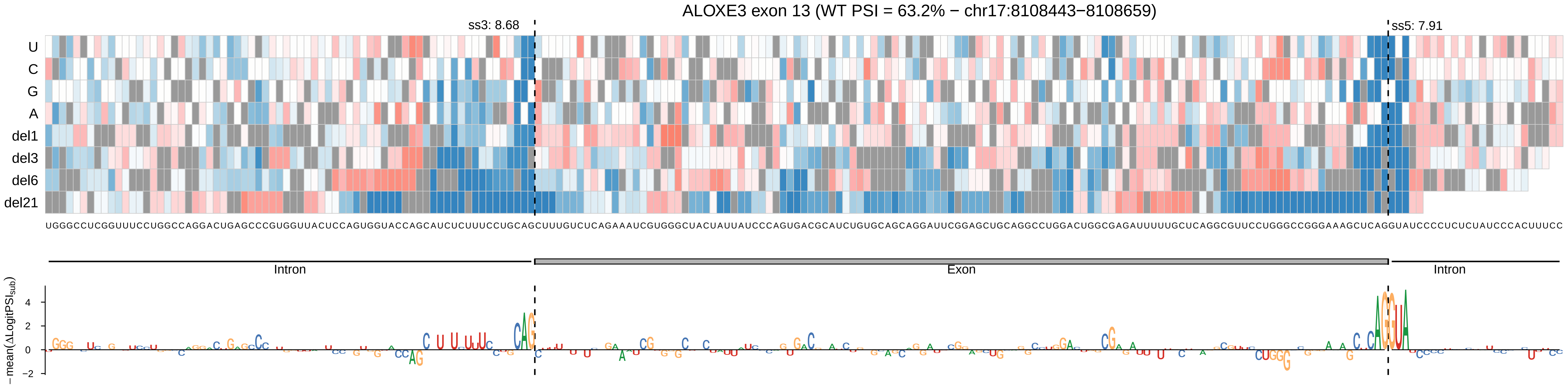
ALOXE3 exon 13 (WT PSI = 63.2% − chr17:8108443−8108659)
ss3: 8.68
ss5: 7.91
U
C
G
A
del1
del3
del6
del21
UGGGCCUCGGUUUCCUGGCCAGGACUGAGCCCGUGGUUACUCCAGUGGUACCAGCAUCUCUUUCCUGCAGCUUUGUCUCAGAAAUCGUGGGCUACUAUUAUCCCAGUGACGCAUCUGUGCAGCAGGAUUCGGAGCUGCAGGCCUGGACUGGCGAGAUUUUUGCUCAGGCGUUCCUGGGCCGGGAAAGCUCAGGUAUCCCCUCUCUAUCCCACUUUCC
Intron
Exon
Intron
− mean(ΔLogitPSI_sub)
4
2
0
−2

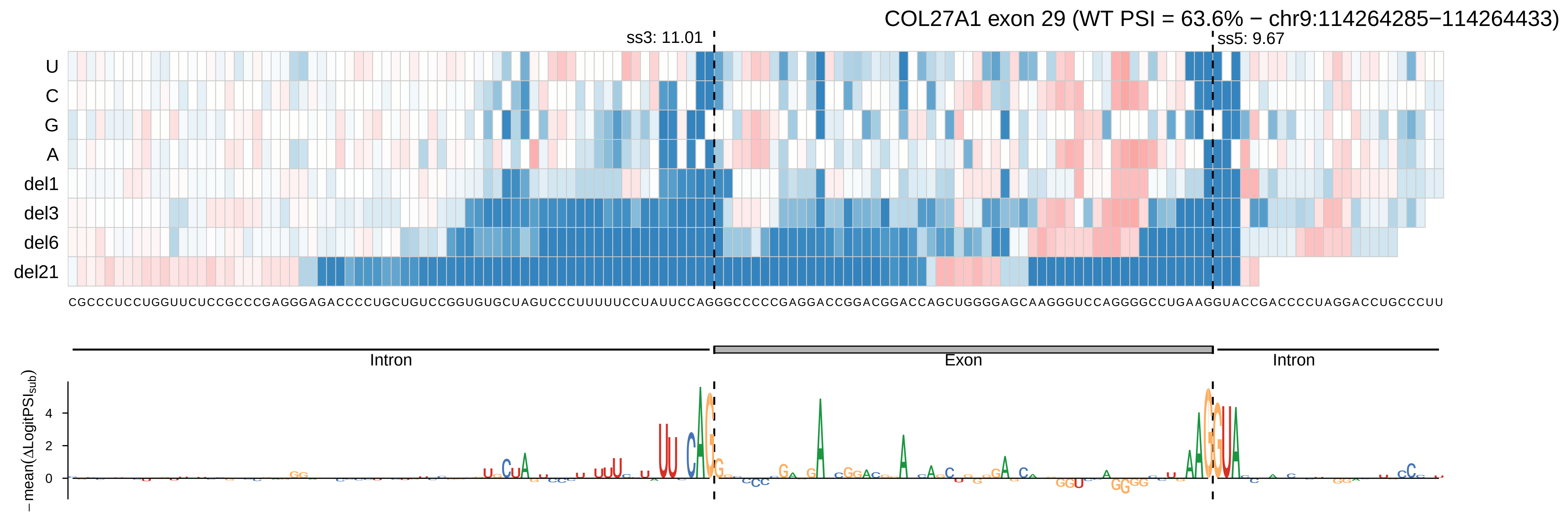

COL27A1 exon 29 (WT PSI = 63.6% − chr9:114264285−114264433)
ss3: 11.01
ss5: 9.67
U
C
G
A
del1
del3
del6
del21
CGCCCUCCUGGUUCUCCGCCCGAGGGAGACCCCUGCUGUCCGGUGUGCUAGUCCCUUUUUCCUAUUCCAGGGCCCCCGAGGACCGGACGGACCAGCUGGGGAGCAAGGGUCCAGGGGCCUGAAGGUACCGACCCCUAGGACCUGCCCUU
Intron
Exon
Intron
− mean(ΔLogitPSI_sub)
4
2
0

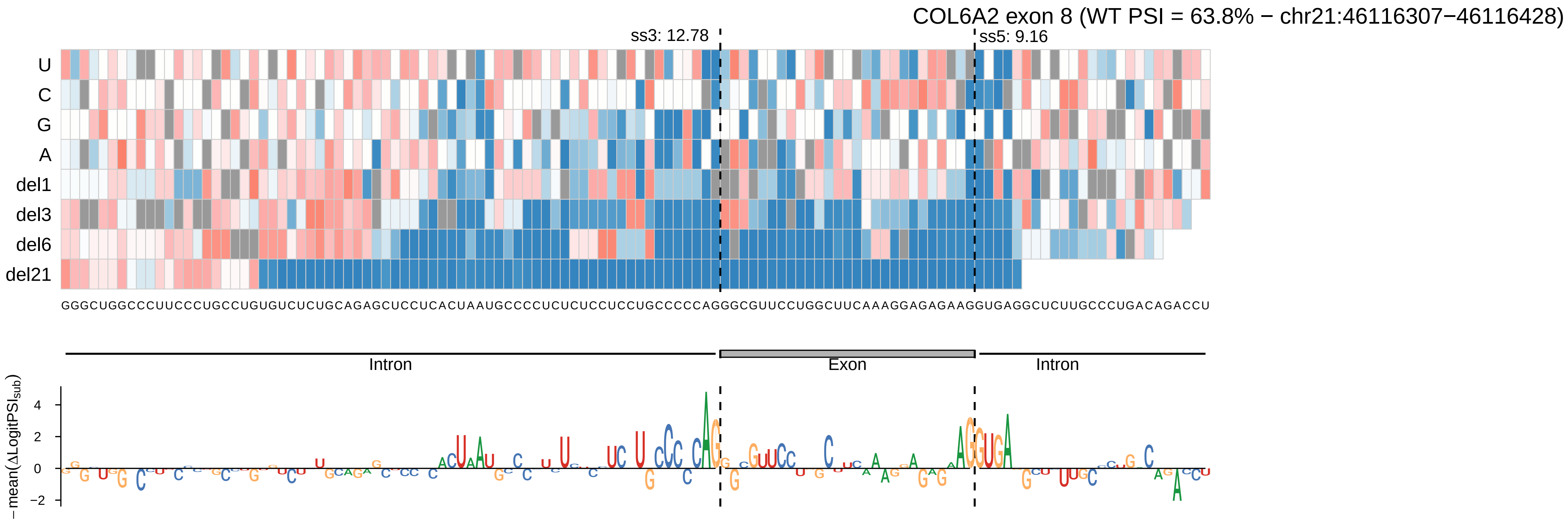

COL6A2 exon 8 (WT PSI = 63.8% − chr21:46116307−46116428)
ss3: 12.78
ss5: 9.16
U
C
G
A
del1
del3
del6
del21
GGGCUGGCCCUUCCCUGCCUGUGUCUCUGCAGAGCUCCUCACUAAUGCCCCUCUCUCCUCCUGCCCCCAGGGCGUUCCUGGCUUCAAAGGAGAGAAGGUGAGGCUCUUGCCCUGACAGACCU
Intron
Exon
Intron
−mean(ΔLogitPSI_sub)

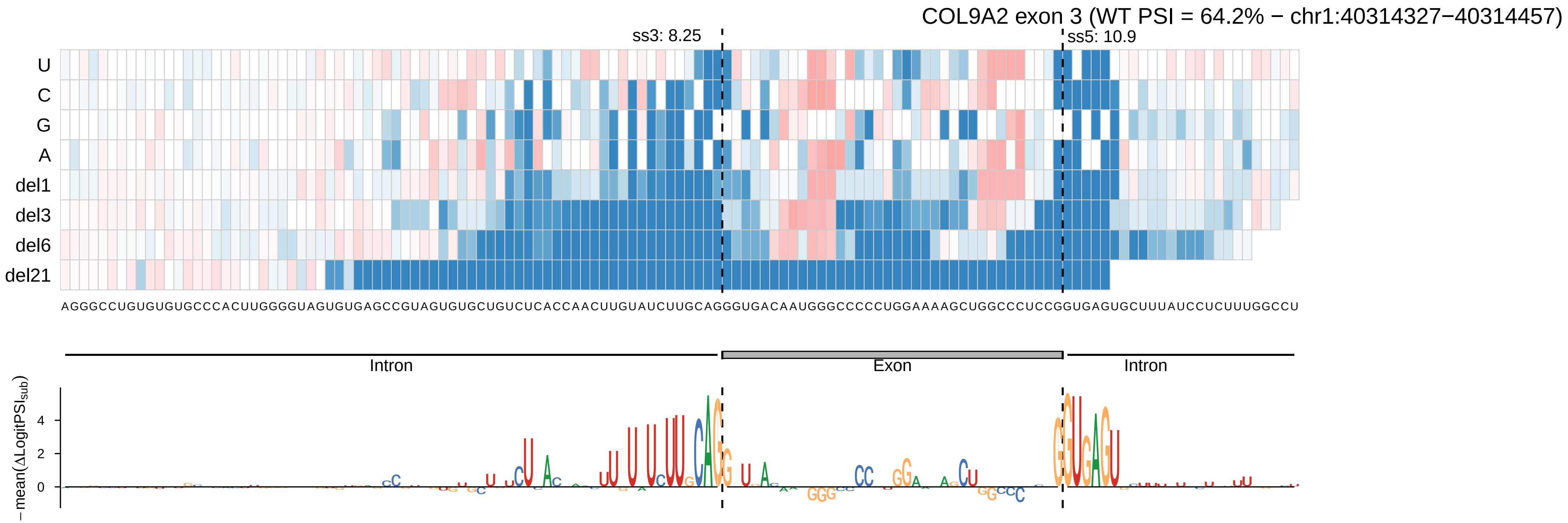
COL9A2 exon 3 (WT PSI = 64.2% − chr1:40314327−40314457)
ss3: 8.25
ss5: 10.9
U
C
G
A
del1
del3
del6
del21
AGGGCCUGUGUGUGCCCACUUGGGGUAGUGUGAGCCGUAGUGUGCUGUCUCACCAACUUGUAUCUUGCAGGGUGACAAUGGGCCCCCUGGAAAAGCUGGCCCUCCGGUGAGUGCUUUAUCCUCUUUGGCCU
Intron
Exon
Intron
−mean(ΔLogitPSI_sub)
4
2
0

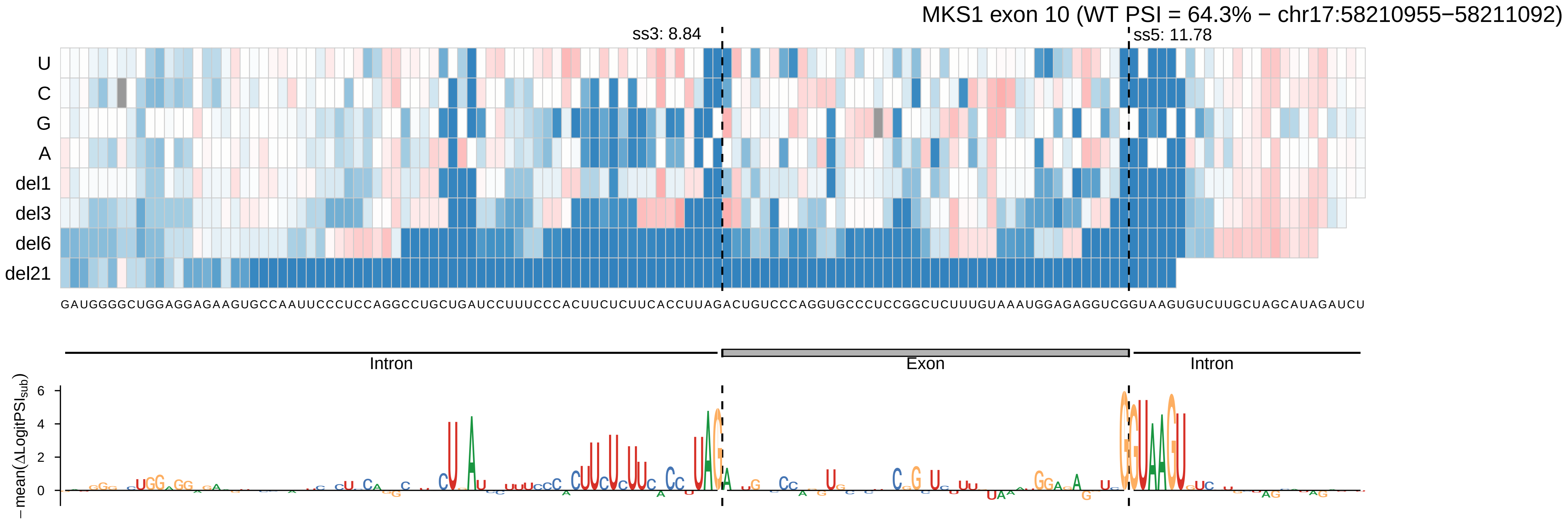
MKS1 exon 10 (WT PSI = 64.3% − chr17:58210955−58211092)
ss3: 8.84
ss5: 11.78
U
C
G
A
del1
del3
del6
del21
GAUGGGGCUGGAGGAGAAGUGCCAAUUCCCUCCAGGCCUGCUGAUCCUUUCCCACUUCUCUUCACCUUAGACUGUCCCAGGUGCCCUCCGGCUCUUUGUAAAUGGAGAGGUCGGUAAGUGUCUUGCUAGCAUAGAUCU
Intron
Exon
Intron
−mean(ΔLogitPSI_sub)
6
4
2
0

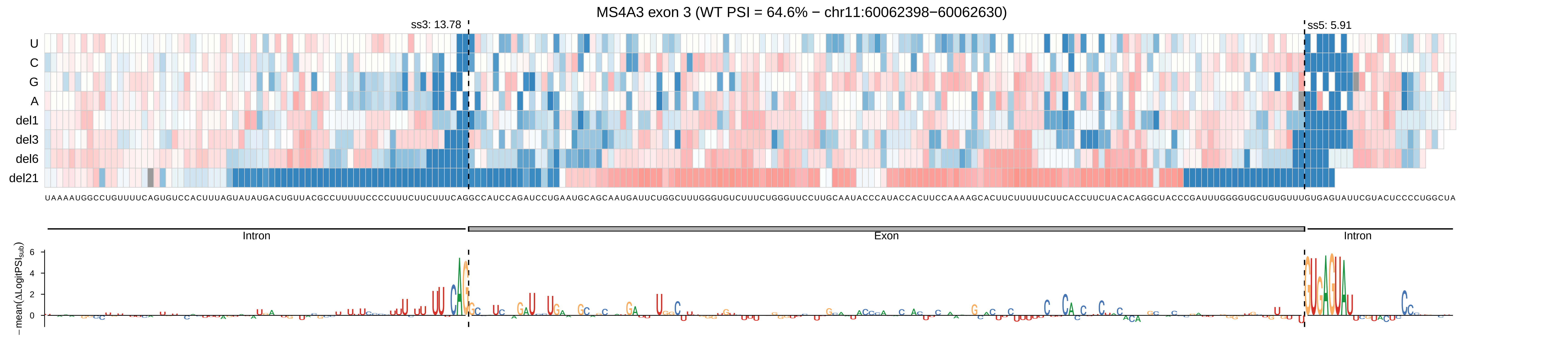
MS4A3 exon 3 (WT PSI = 64.6% − chr11:60062398−60062630)
ss3: 13.78
ss5: 5.91
U
C
G
A
del1
del3
del6
del21
UAAAAUGGCCUGUUUUCAGUGUCCACUUUAGUAUAUGACUGUUACGCCUUUUUCCCCUUUCUUCUUUCAGGCCAUCCAGAUCCUGAAUGCAGCAAUGAUUCUGGCUUUGGGUGUCUUUCUGGGUUCCUUGCAAUACCCAUACCACUUCCAAAAGCACUUCUUUUUCUUCACCUUCUACACAGGCUACCCGAUUUGGGGUGCUGUGUUUGUGAGUAUUCGUACUCCCCUGGCUA
Intron
Exon
Intron
−mean(ΔLogitPSI_sub)
6
4
2
0

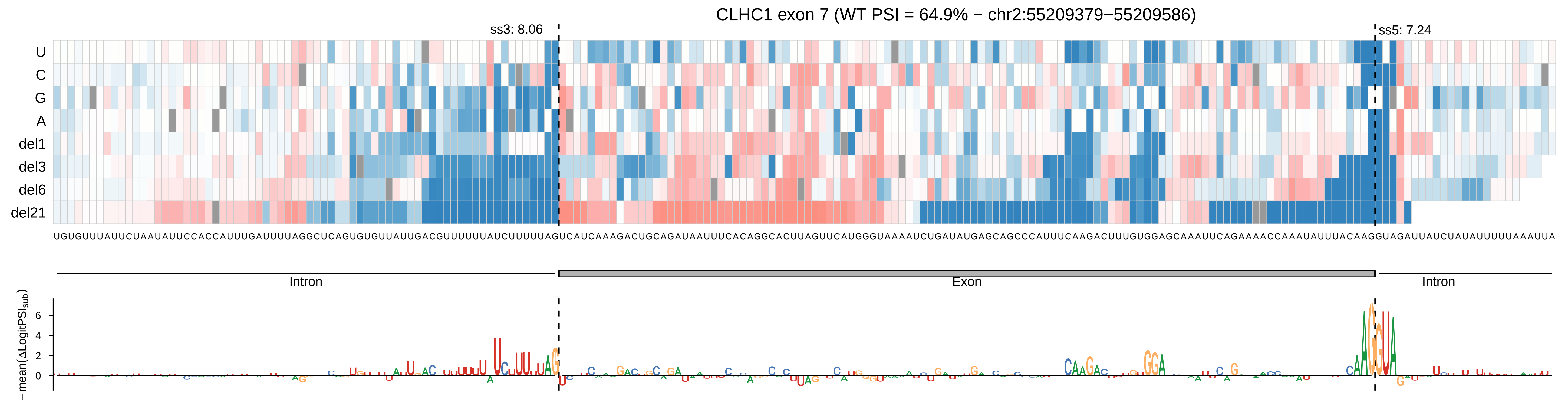

CLHC1 exon 7 (WT PSI = 64.9% − chr2:55209379−55209586)
ss3: 8.06
ss5: 7.24
U
C
G
A
del1
del3
del6
del21
UGUGUUUAUUCUAAUAUUCCACCAUUUGAUUUUAGGCUCAGUGUGUUAUUGACGUUUUUUAUCUUUUUAGUCAUCAAAGACUGCAGAUAAUUUCACAGGCACUUAGUUCAUGGGUAAAAUCUGAUAUGAGCAGCCCAUUUCAAGACUUUGUGGAGCAAAUUCAGAAAACCAAAUAUUUACAAGGUAGAUUAUCUAUAUUUUUAAAUUA
Intron
Exon
Intron
−mean(ΔLogitPSI_sub)
6
4
2
0

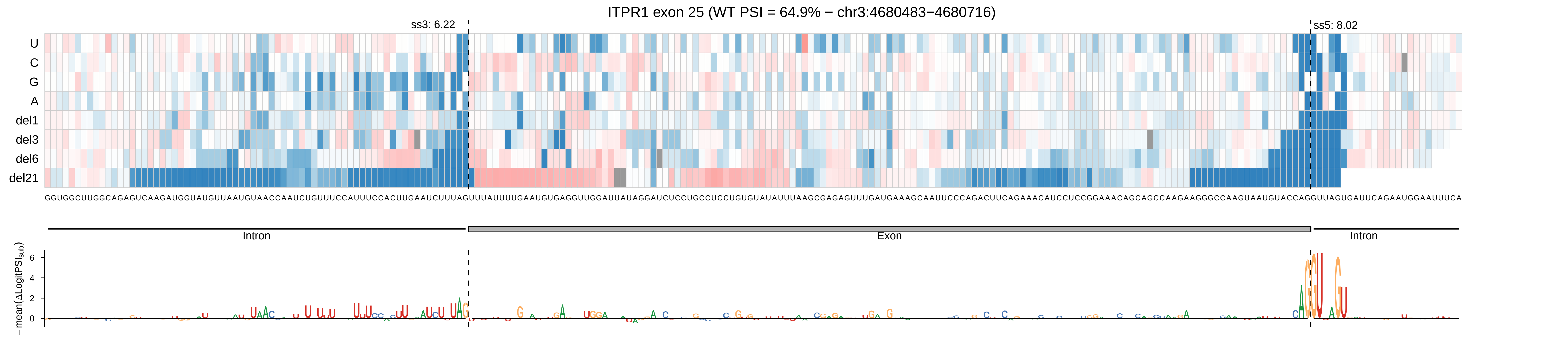
ITPR1 exon 25 (WT PSI = 64.9% − chr3:4680483−4680716)
ss3: 6.22
ss5: 8.02
U
C
G
A
del1
del3
del6
del21
GGUGGCUUGGCAGAGUCAAGAUGGUAUGUUAAUGUAACCAAUCUGUUUCCAUUUCCACUUGAAUCUUUAGUUUAUUUUGAAUGUGAGGUUGGAUUAUAGGAUCUCCUGCCUCCUGUGUAUAUUUAAGCGAGAGUUUGAUGAAAGCAAUUCCCAGACUUCAGAAACAUCCUCCGGAAACAGCAGCCAAGAAGGGCCAAGUAAUGUACCAGGUUAGUGAUUCAGAAUGGAAUUUCA
Intron
Exon
Intron
−mean(ΔLogitPSI_sub)
6
4
2
0

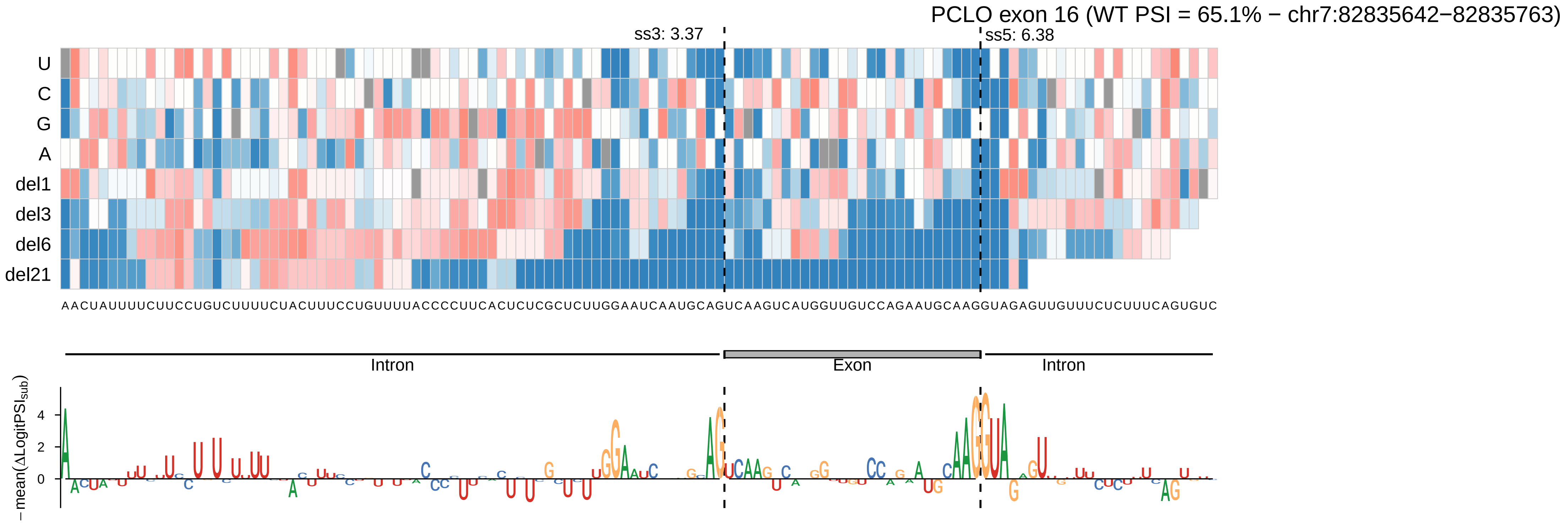
PCLO exon 16 (WT PSI = 65.1% − chr7:82835642−82835763)
ss3: 3.37
ss5: 6.38
U
C
G
A
del1
del3
del6
del21
AACUAUUUUCUUCCUGUCUUUUCUACUUUCCUGUUUUACCCCUUCACUCUCGCUCUUGGAAUCAAUGCAGUCAAGUCAUGGUUGUCCAGAAUGCAAGGUAGAGUUGUUUCUCUUUCAGUGUC
Intron
Exon
Intron
−mean(ΔLogitPSI_sub)
4
2
0

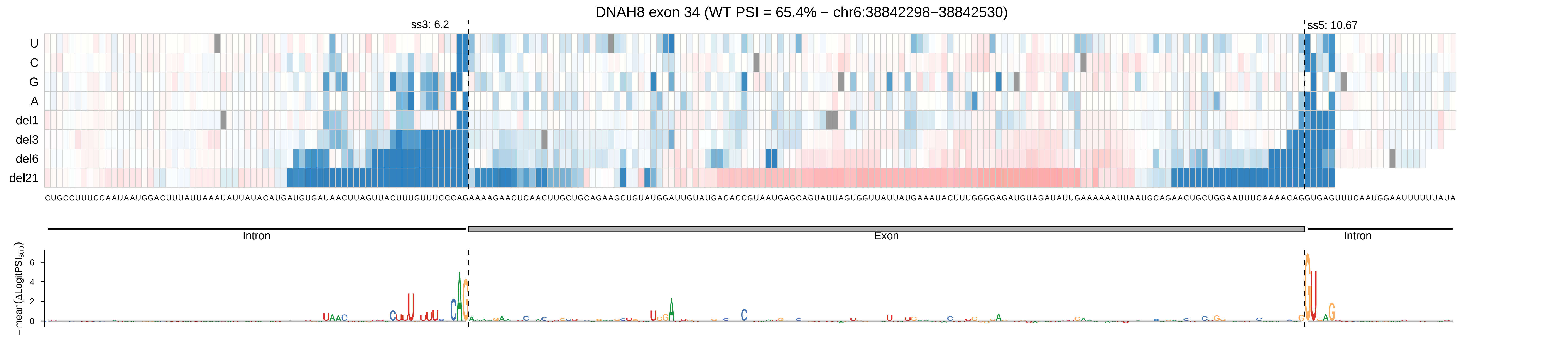

DNAH8 exon 34 (WT PSI = 65.4% − chr6:38842298−38842530)
ss3: 6.2
ss5: 10.67
U
C
G
A
del1
del3
del6
del21
CUGCCUUUCCAAUAAUGGACUUUAUUAAAUAUUAUACAUGAUGUGAUAACUUAGUUACUUUGUUUCCCAGAAAAGAACUCAACUUGCUGCAGAAGCUGUAUGGAUUGUAUGACACCGUAAUGAGCAGUAUUAGUGGUUAUUAUGAAAUACUUUGGGGAGAUGUAGAUAUUGAAAAAAUUAAUGCAGAACUGCUGGAAUUUCAAAACAGGUGAGUUUCAAUGGAAUUUUUUAUA
Intron
Exon
Intron
−mean(ΔLogitPSI_sub)

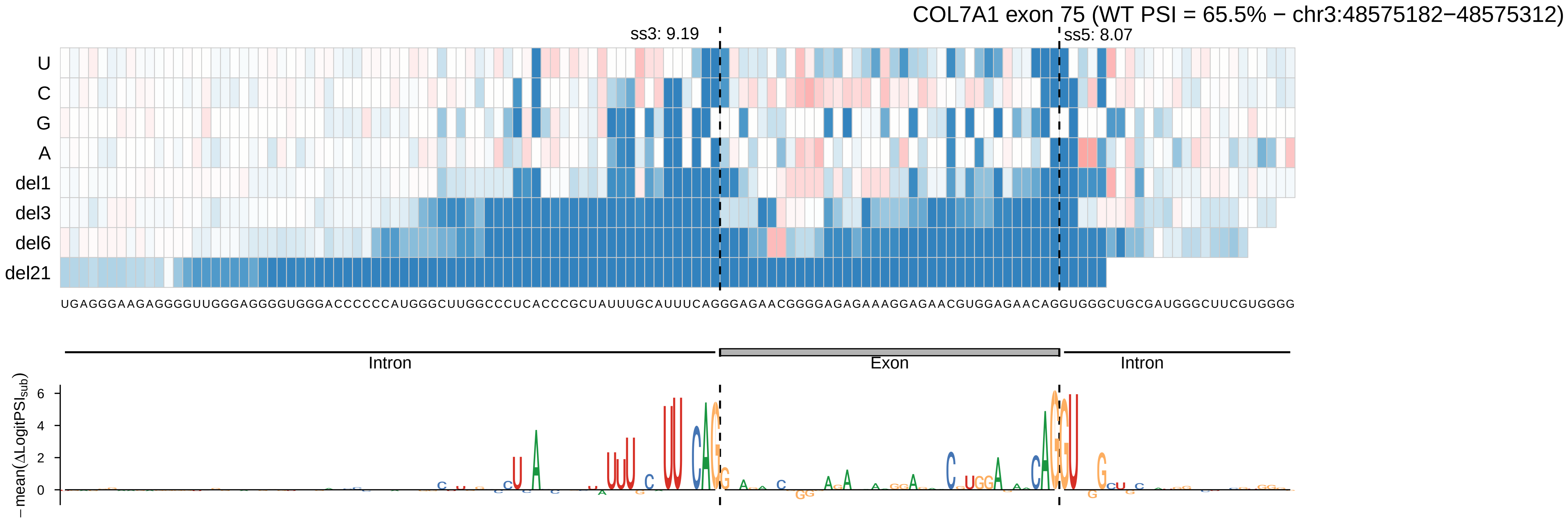
COL7A1 exon 75 (WT PSI = 65.5% − chr3:48575182−48575312)
ss3: 9.19
ss5: 8.07
U
C
G
A
del1
del3
del6
del21
UGAGGGAAGAGGGGUUGGGAGGGGUGGGACCCCCCAUGGGCUUGGCCCUCACCCGCUAUUUGCAUUUCAGGGAGAACGGGGAGAGAAAGGAGAACGUGGAGAACAGGUGGGCUGCGAUGGGCUUCGUGGGG
Intron
Exon
Intron
− mean(ΔLogitPSI_sub)
6
4
2
0

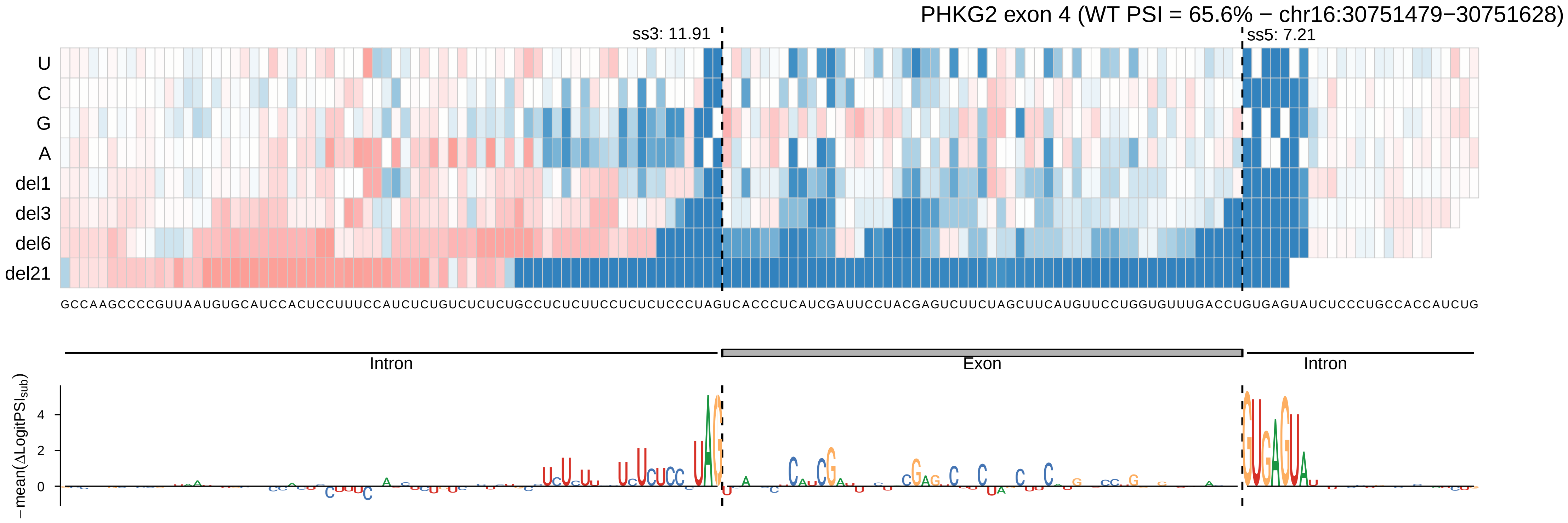
PHKG2 exon 4 (WT PSI = 65.6% − chr16:30751479−30751628)
ss3: 11.91
ss5: 7.21
U
C
G
A
del1
del3
del6
del21
GCCAAGCCCCGUUAAUGUGCAUCCACUCCUUUCCAUCUCUGUCUCUCUGCCUCUCUUCCUCUCUCCCUAGUCACCCUCAUCGAUUCCUACGAGUCUUCUAGCUUCAUGUUCCUGGUGUUUGACCUGUGAGUAUCUCCCUGCCACCAUCUG
Intron
Exon
Intron
−mean(ΔLogitPSI_sub)
4
2
0

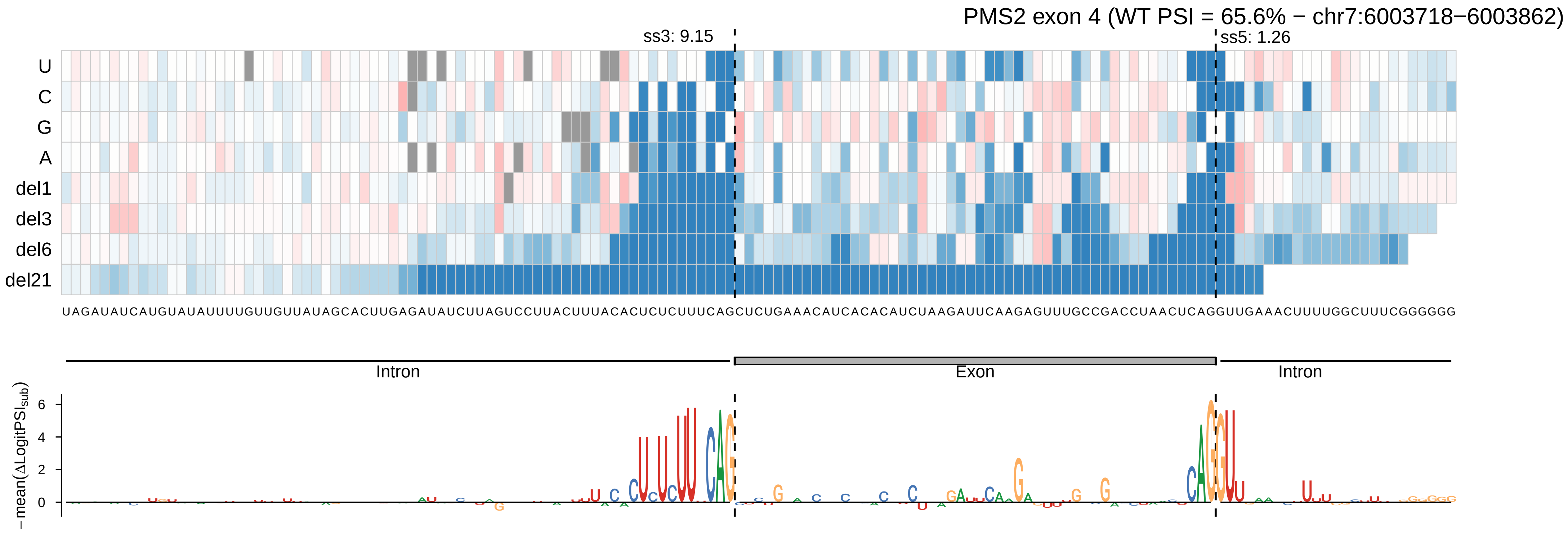
PMS2 exon 4 (WT PSI = 65.6% − chr7:6003718−6003862)
ss3: 9.15
ss5: 1.26
U
C
G
A
del1
del3
del6
del21
UAGAUAUCAUGUAUAUUUUGUUGUUAUAGCACUUGAGAUAUCUUAGUCCUUACUUUACACUCUCUUUCAGCUCUGAAACAUCACACAUCUAAGAUUCAAGAGUUUGCCGACCUAACUCAGGUUGAAACUUUUGGCUUUCGGGGGG
Intron
Exon
Intron
−mean(ΔLogitPSI_sub)
6
4
2
0

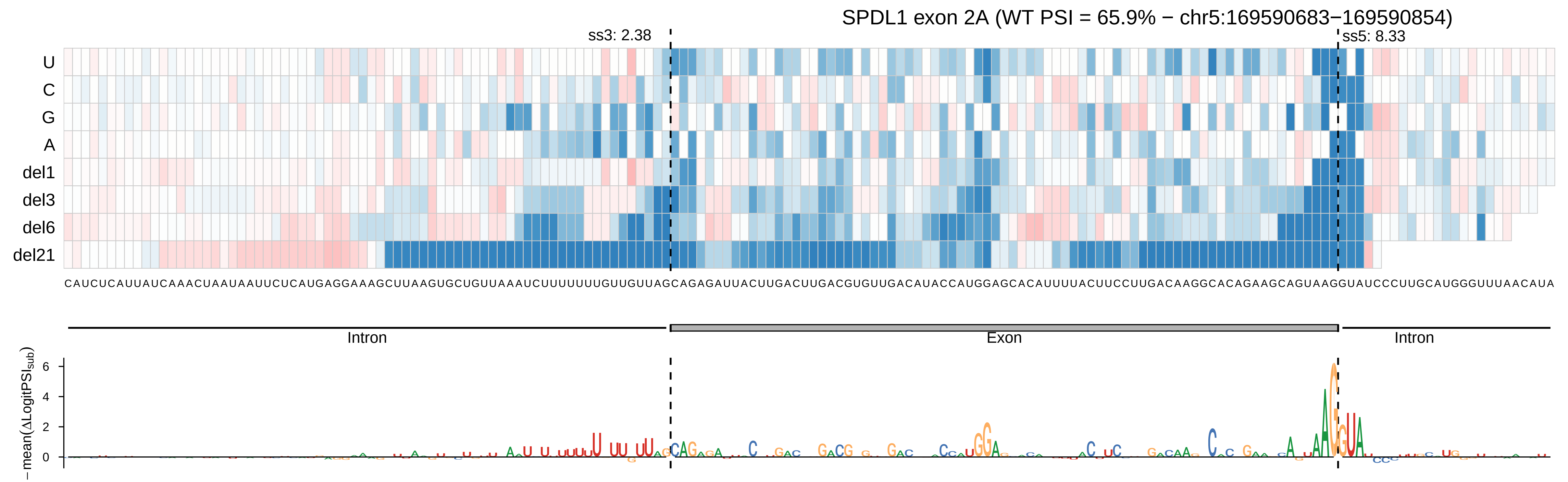

SPDL1 exon 2A (WT PSI = 65.9% − chr5:169590683−169590854)
ss3: 2.38
ss5: 8.33
U
C
G
A
del1
del3
del6
del21
CAUCUCAUUAUCAAACUAAUAAUUCUCAUGAGGAAAGCUUAAGUGCUGUUAAAUCUUUUUUUGUUGUUAGCAGAGAUUACUUGACUUGACGUGUUGACAUACCAUGGAGCACAUUUUACUUCCUUGACAAGGCACAGAAGCAGUAAGGUAUCCCUUGCAUGGGUUUAACAUA
Intron
Exon
Intron
− mean(ΔLogitPSI_sub)
6
4
2
0

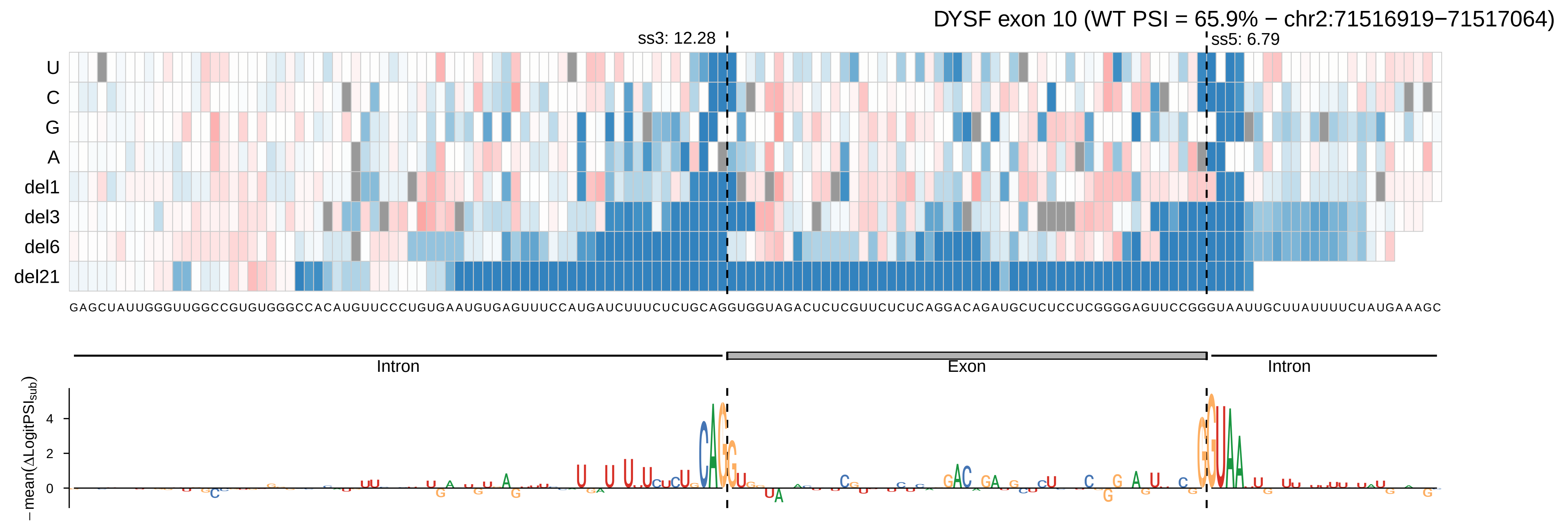
DYSF exon 10 (WT PSI = 65.9% − chr2:71516919−71517064)
ss3: 12.28
ss5: 6.79
U
C
G
A
del1
del3
del6
del21
GAGCUAUUGGGUUGGCCGUGUGGGCCACAUGUUCCCUGUGAAUGUGAGUUUCCAUGAUCUUUCUCUGCAGGUGGUAGACUCUCGUUCUCUCAGGACAGAUGCUCUCCUCGGGGAGUUCCGGGUAAUUGCUUAUUUUCUAUGAAAGC
Intron
Exon
Intron
−mean(ΔLogitPSI_sub)
4
2
0

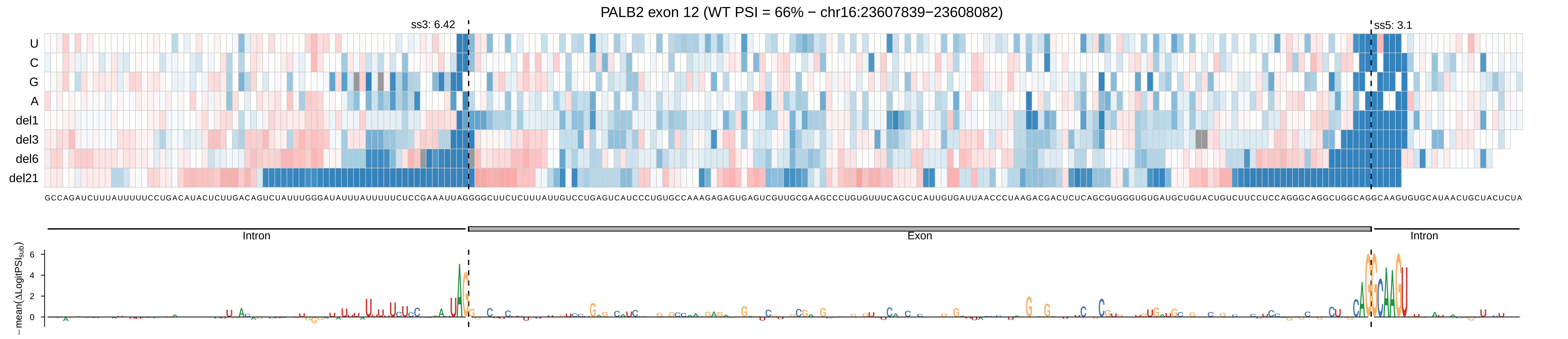

PALB2 exon 12 (WT PSI = 66% − chr16:23607839−23608082)
ss3: 6.42
ss5: 3.1
U
C
G
A
del1
del3
del6
del21
GCCAGAUCUUUAUUUUUCCUGACAUACUCUUGACAGUCUAUUUGGGAUAUUUAUUUUUCUCCGAAAUUAGGGGCUUCUCUUUAUUGUCCUGAGUCAUCCCUGUGCCAAAGAGAGUGAGUCGUUGCGAAGCCCUGUGUUUCAGCUCAUUGUGAUUAACCCUAAGACGACUCUCAGCGUGGGUGUGAUGCUGUACUGUCUUCCUCCAGGGCAGGCUGGCAGGCAAGUGUGCAUAACUGCUACUCUA
Intron
Exon
Intron
−mean(ΔLogitPSI_sub)
6
4
2
0

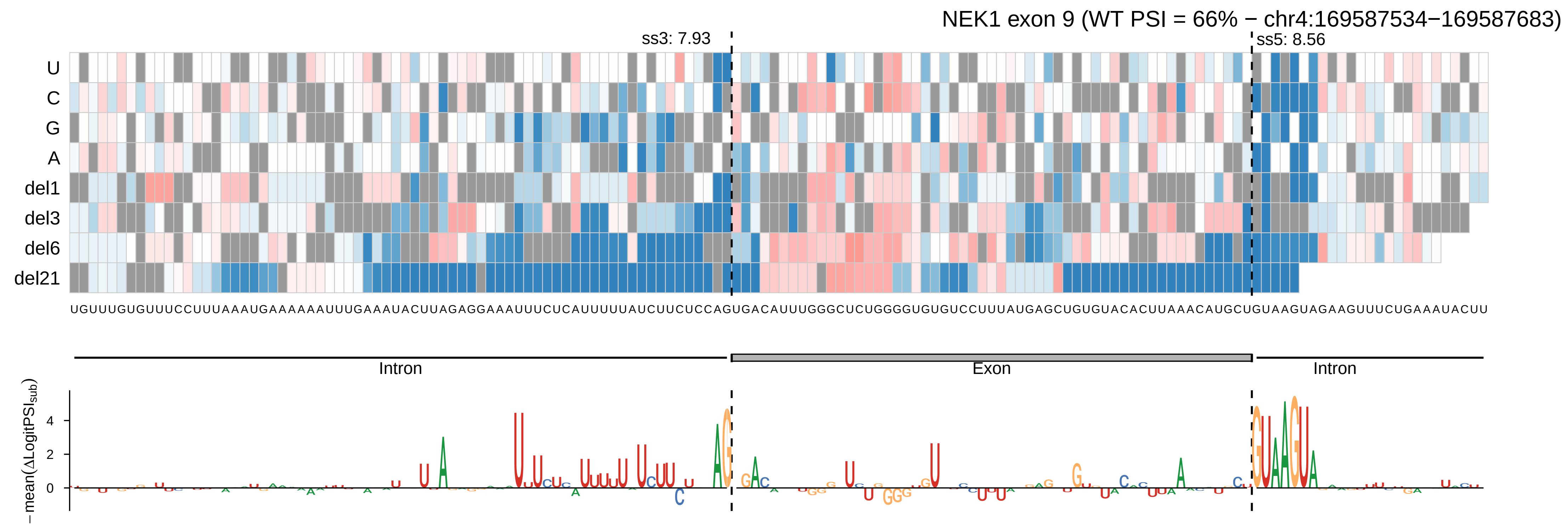

NEK1 exon 9 (WT PSI = 66% − chr4:169587534−169587683)
ss3: 7.93
ss5: 8.56
U
C
G
A
del1
del3
del6
del21
UGUUUGUGUUUCCUUUAAAUGAAAAAAUUUGAAAUACUUAGAGGAAAUUUCUCAUUUUUAUCUUCUCCAGUGACAUUUGGGCUCUGGGGUGUGUCCUUUAUGAGCUGUGUACACUUAAACAUGCUGUAAGUAGAAGUUUCUGAAAUACUU
Intron
Exon
Intron
−mean(ΔLogitPSI_sub)
4
2
0

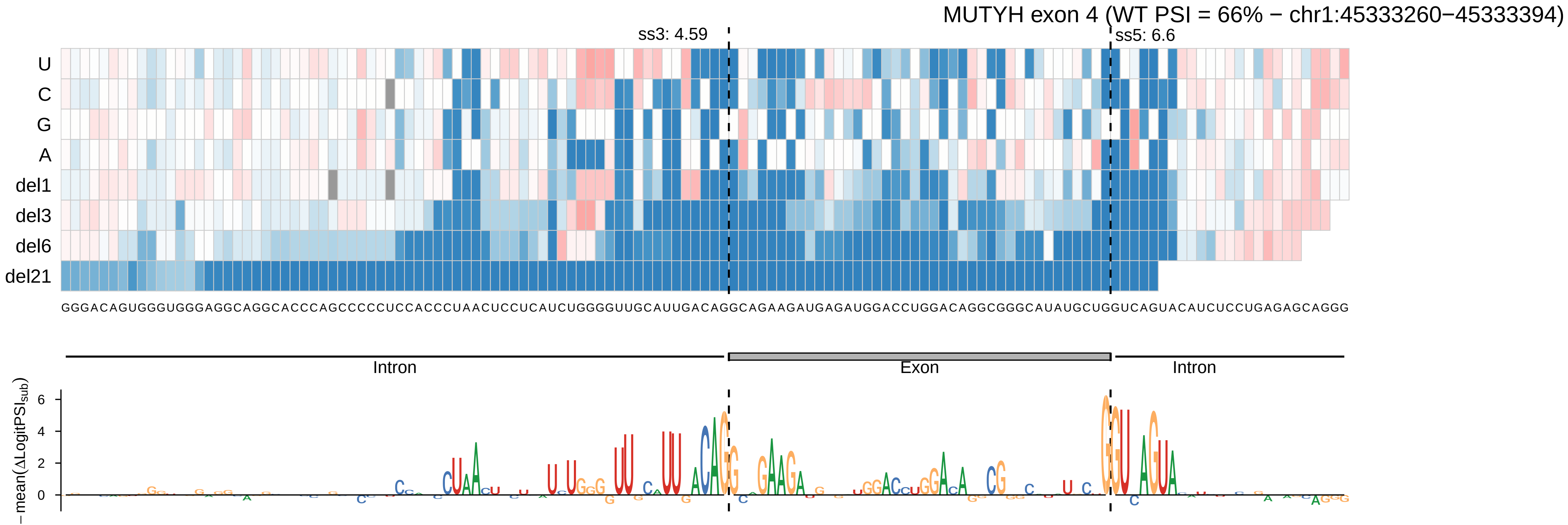
MUTYH exon 4 (WT PSI = 66% − chr1:45333260−45333394)
ss3: 4.59
ss5: 6.6
U
C
G
A
del1
del3
del6
del21
GGGACAGUGGGUGGGAGGCAGGCACCCAGCCCCCUCCACCCUAACUCCUCAUCUGGGGUUGCAUUGACAGGCAGAAGAUGAGAUGGACCUGGACAGGCGGGCAUAUGCUGGUCAGUACAUCUCCUGAGAGCAGGG
Intron
Exon
Intron
−mean(ΔLogitPSI_sub)
6
4
2
0

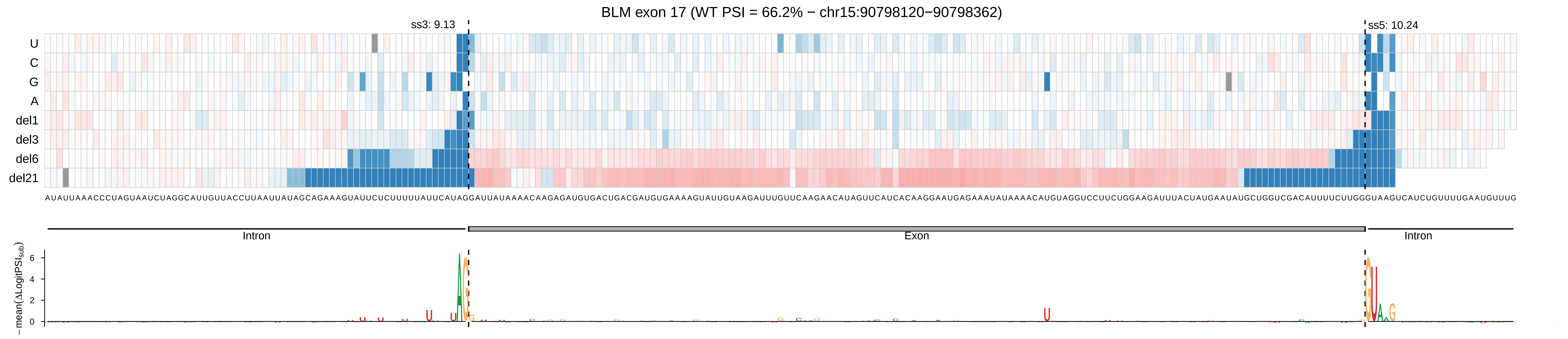
BLM exon 17 (WT PSI = 66.2% − chr15:90798120−90798362)
ss3: 9.13
ss5: 10.24
U
C
G
A
del1
del3
del6
del21
Intron
Exon
Intron
− mean(ΔLogitPSI_sub)
6
4
2
0

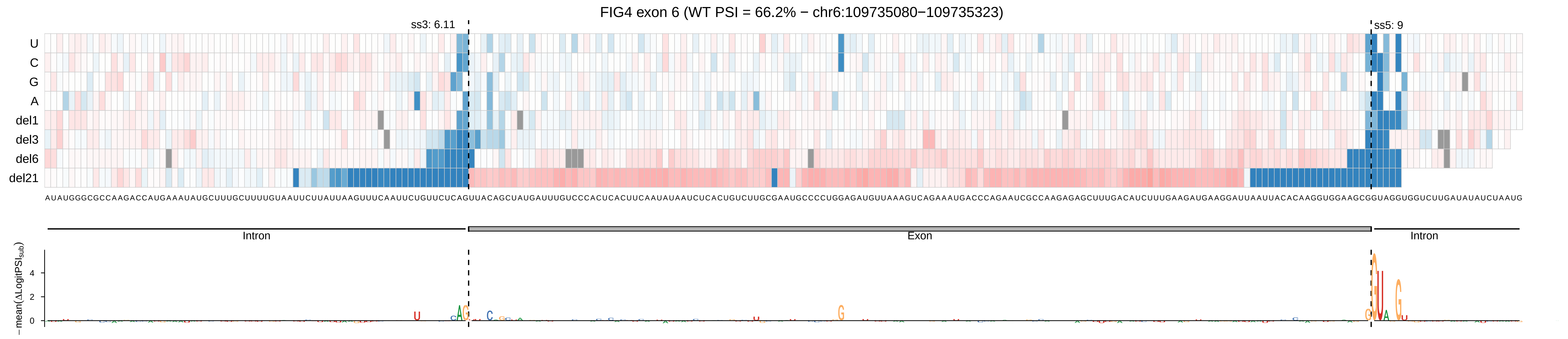
FIG4 exon 6 (WT PSI = 66.2% − chr6:109735080−109735323)
ss3: 6.11
ss5: 9
U
C
G
A
del1
del3
del6
del21
Intron
Exon
Intron
−mean(ΔLogitPSI_sub)
4
2
0

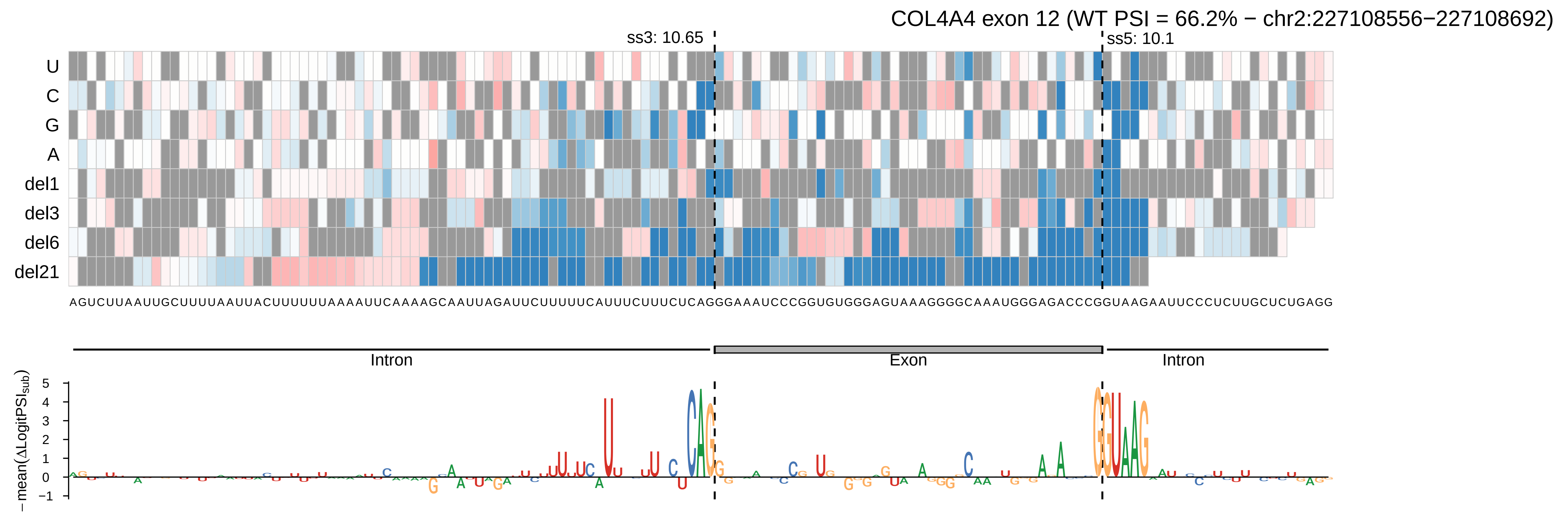
COL4A4 exon 12 (WT PSI = 66.2% − chr2:227108556−227108692)
ss3: 10.65
ss5: 10.1
U
C
G
A
del1
del3
del6
del21
AGUCUUAAUUGCUUUUAAUUACUUUUUUAAAAUUCAAAAGCAAUUAGAUUCUUUUUCAUUUCUUUCUCAGGGAAAUCCCGGUGUGGGAGUAAAGGGGCAAAUGGGAGACCCGGUAAGAAUUCCCUCUUGCUCUGAGG
Intron
Exon
Intron
− mean(ΔLogitPSI_sub)
5
4
3
2
1
0
−1

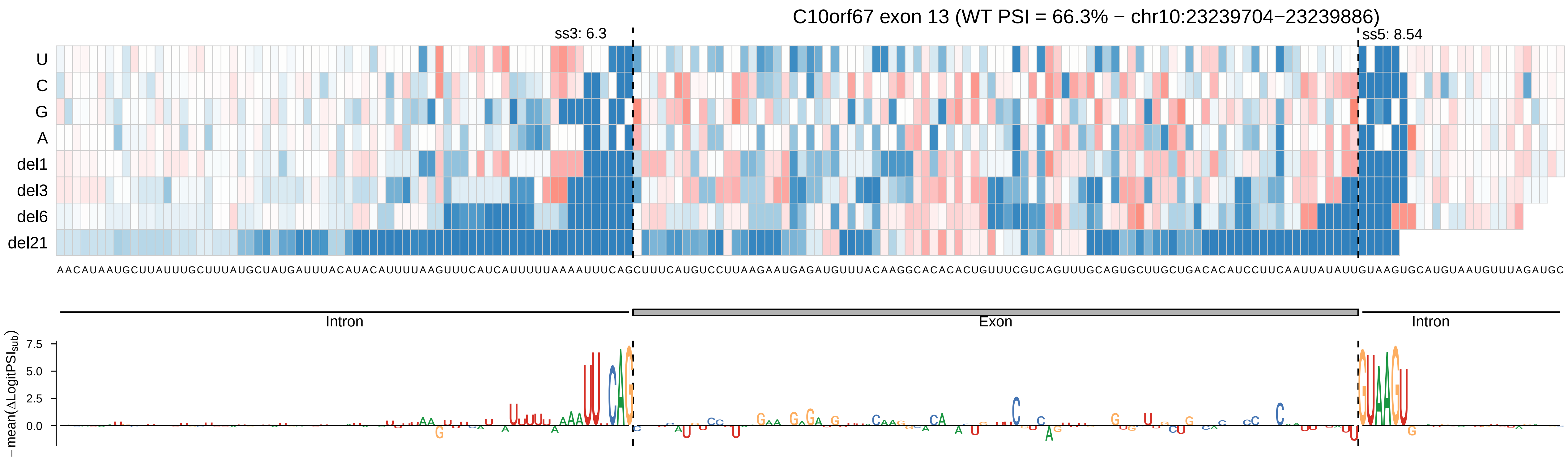
C10orf67 exon 13 (WT PSI = 66.3% − chr10:23239704−23239886)
ss3: 6.3
ss5: 8.54
U
C
G
A
del1
del3
del6
del21
Intron
Exon
Intron
−mean(ΔLogitPSI_sub)
7.5
5.0
2.5
0.0

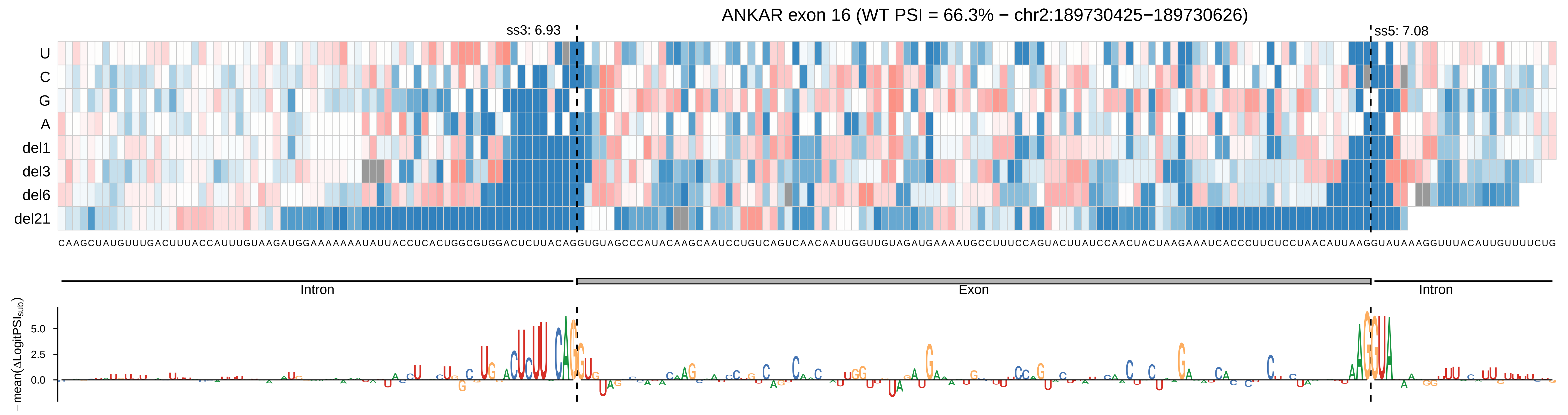

ANKAR exon 16 (WT PSI = 66.3% − chr2:189730425−189730626)
ss3: 6.93
ss5: 7.08
U
C
G
A
del1
del3
del6
del21
Intron
Exon
Intron
−mean(ΔLogitPSI_sub)
5.0
2.5
0.0

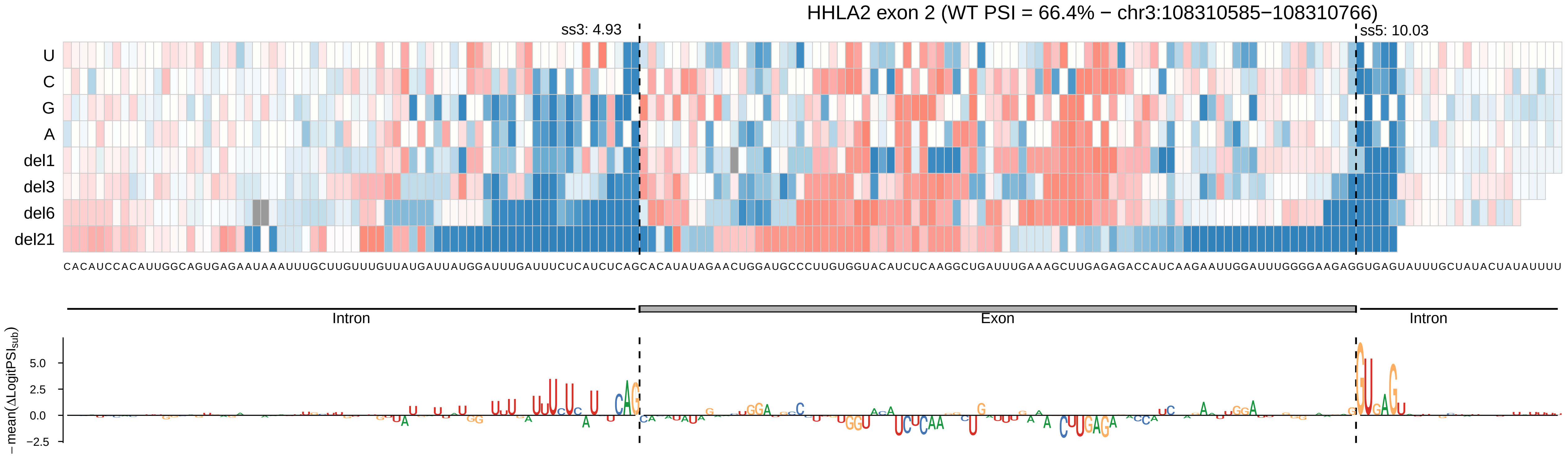

HHLA2 exon 2 (WT PSI = 66.4% − chr3:108310585−108310766)
ss3: 4.93
ss5: 10.03
U
C
G
A
del1
del3
del6
del21
CACAUCCACAUUGGCAGUGAGAAUAAAUUUGCUUGUUUGUUAUGAUUAUGGAUUUGAUUUCUCAUCUCAGCACAUAUAGAACUGGAUGCCCUUGUGGUACAUCUCAAGGCUGAUUUGAAAGCUUGAGAGACCAUCAAGAAUUGGAUUUGGGGAAGAGGUGAGUAUUUGCUAUACUAUAUUUU
Intron
Exon
Intron
− mean(ΔLogitPSI_sub)
5.0
2.5
0.0
−2.5

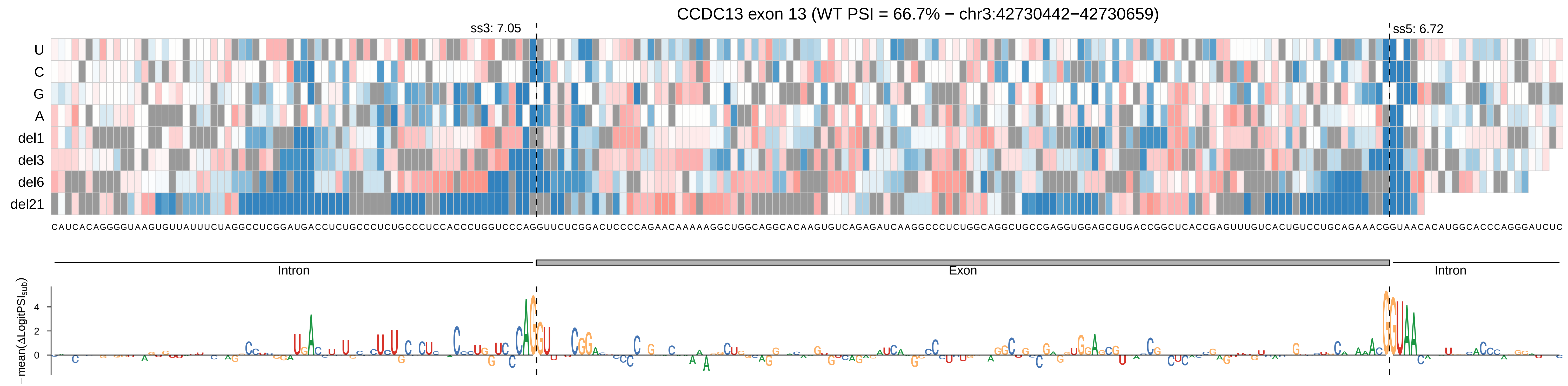

CCDC13 exon 13 (WT PSI = 66.7% − chr3:42730442−42730659)
ss3: 7.05
ss5: 6.72
U
C
G
A
del1
del3
del6
del21
CAUCACAGGGGUAAGUGUUAUUUCUAGGCCUCGGAUGACCUCUGCCCUCUGCCCUCCACCCUGGUCCCAGGUUCUCGGACUCCCCAGAACAAAAAGGCUGGCAGGCACAAGUGUCAGAGAUCAAGGCCCUCUGGCAGGCUGCCGAGGUGGAGCGUGACCGGCUCACCGAGUUUGUCACUGUCCUGCAGAAACGGUAACACAUGGCACCCAGGGAUCUC
Intron
Exon
Intron
− mean(ΔLogitPSI$_{sub}$)
4
2
0

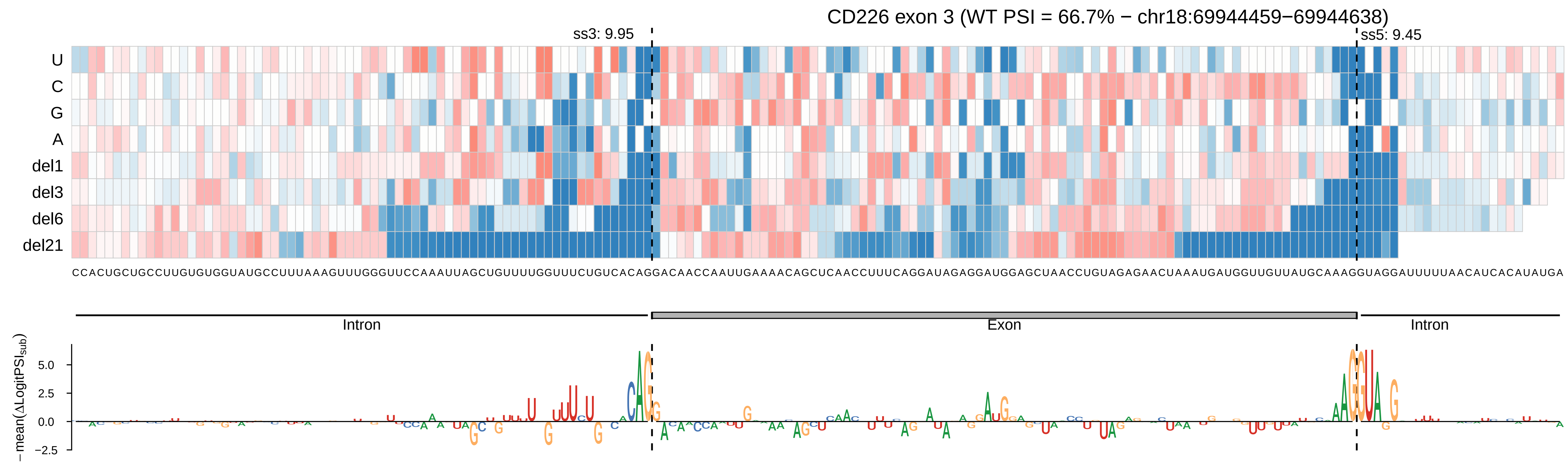
CD226 exon 3 (WT PSI = 66.7% − chr18:69944459−69944638)
ss3: 9.95
ss5: 9.45
U
C
G
A
del1
del3
del6
del21
CCACUGCUGCCUUGUGUGGUAUGCCUUUAAAGUUUGGGUUCCAAAUUAGCUGUUUUGGUUUCUGUCACAGGACAACCAAUUGAAAACAGCUCAACCUUUCAGGAUAGAGGAUGGAGCUAACCUGUAGAGAACUAAAUGAUGGUUGUUAUGCAAAGGUAGGAUUUUUAACAUCACAUAUGA
Intron
Exon
Intron
− mean(ΔLogitPSI_sub)
5.0
2.5
0.0
−2.5

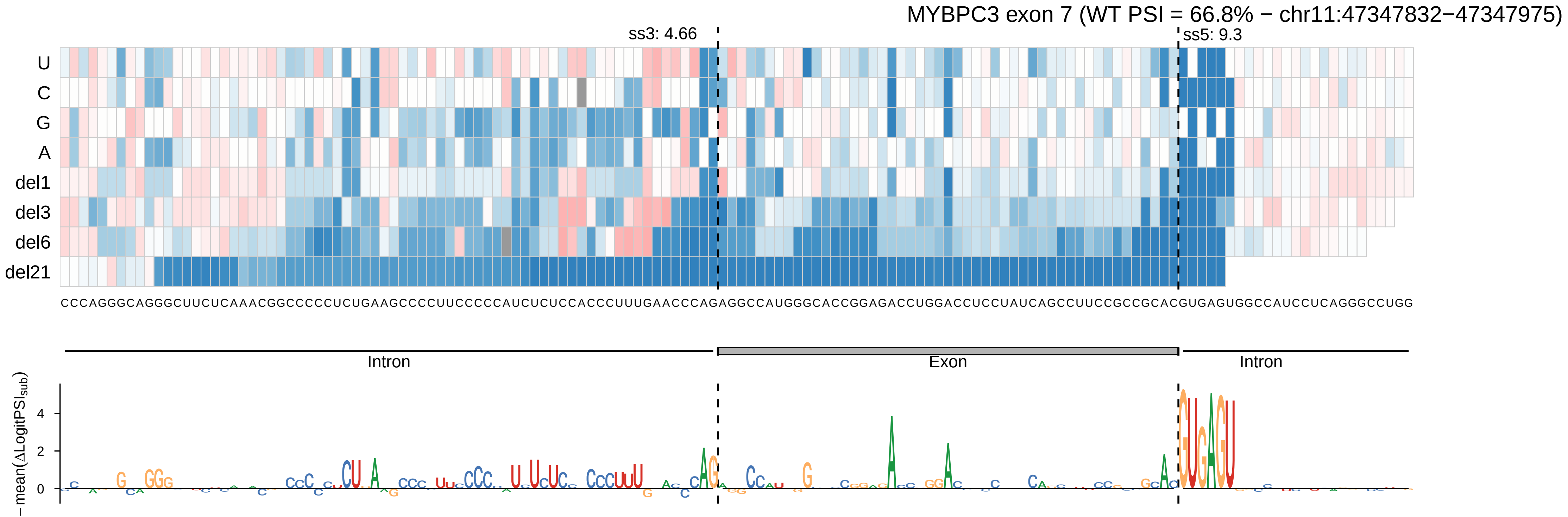
MYBPC3 exon 7 (WT PSI = 66.8% − chr11:47347832−47347975)
ss3: 4.66
ss5: 9.3
U
C
G
A
del1
del3
del6
del21
CCCAGGGCAGGGCUUCUCAAACGGCCCCCUCUGAAGCCCCUUCCCCCAUCUCUCCACCCUUUGAACCCAGAGGCCAUGGGCACCGGAGACCUGGACCUCCUAUCAGCCUUCCGCCGCACGUGAGUGGCCAUCCUCAGGGCCUGG
Intron
Exon
Intron
− mean(ΔLogitPSI_sub)
4
2
0

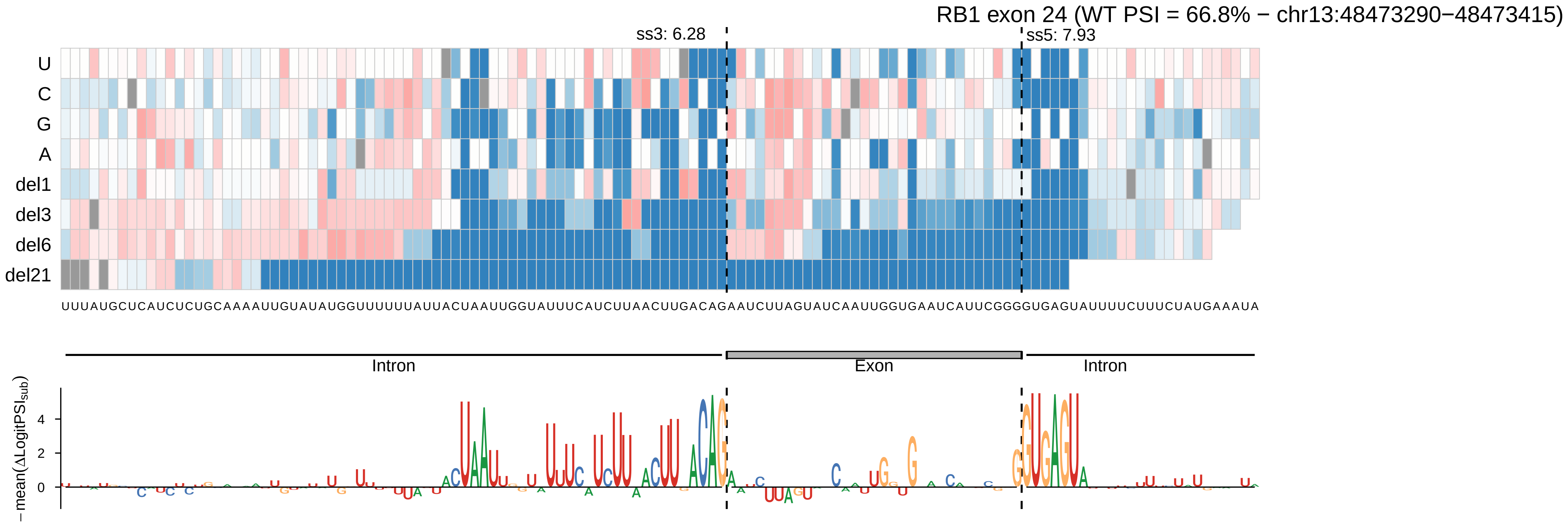
RB1 exon 24 (WT PSI = 66.8% − chr13:48473290−48473415)
ss3: 6.28
ss5: 7.93
U
C
G
A
del1
del3
del6
del21
UUUAUGCUCAUCUCUGCAAAAUUGUAUAUGGUUUUUUAUUACUAAUUGGUAUUUCAUCUUAACUUGACAGAAUCUUAGUAUCAAUUGGUGAAUCAUUCGGGGUGAGUAUUUUCUUUCUAUGAAAUA
Intron
Exon
Intron
−mean(ΔLogitPSI_sub)
4
2
0

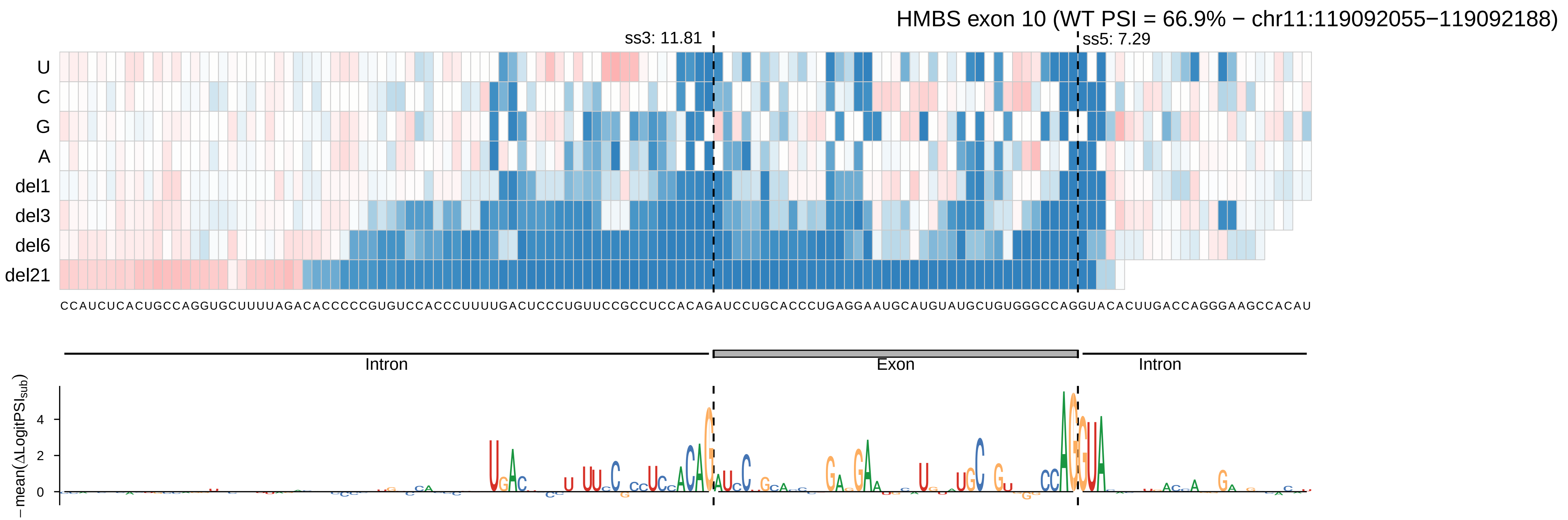
HMBS exon 10 (WT PSI = 66.9% − chr11:119092055−119092188)
ss3: 11.81
ss5: 7.29
U
C
G
A
del1
del3
del6
del21
CCAUCUCACUGCCAGGUGCUUUUAGACACCCCCGUGUCCACCCUUUUGACUCCCUGUUCCGCCUCCACAGAUCCUGCACCCUGAGGAAUGCAUGUAUGCUGUGGGCCAGGUACACUUGACCAGGGAAGCCACAU
Intron
Exon
Intron
−mean(ΔLogitPSI_sub)
4
2
0

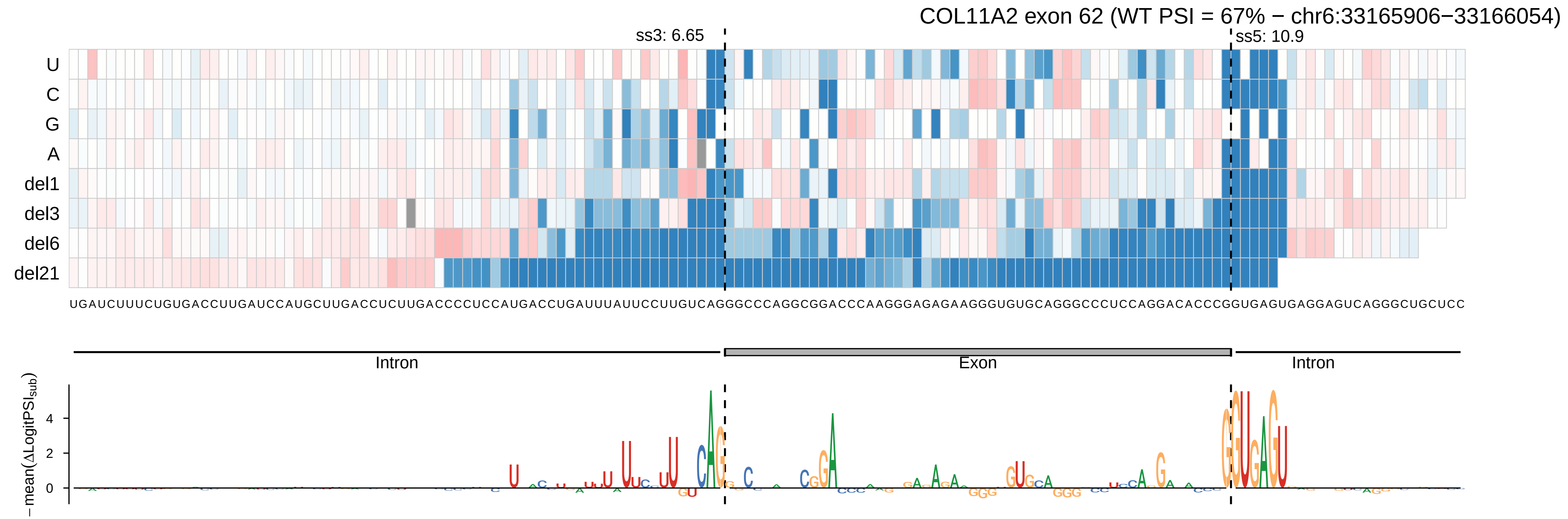
COL11A2 exon 62 (WT PSI = 67% − chr6:33165906−33166054)
ss3: 6.65
ss5: 10.9
U
C
G
A
del1
del3
del6
del21
UGAUCUUUCUGUGACCUUGAUCCAUGCUUGACCUCUUGACCCCUCCAUGACCUGAUUUAUUCCUUGUCAGGGCCCAGGCGGACCCAAGGGAGAGAAGGGUGUGCAGGGCCCUCCAGGACACCCGGUGAGUGAGGAGUCAGGGCUGCUCC
Intron
Exon
Intron
− mean(ΔLogitPSI_sub)
4
2
0

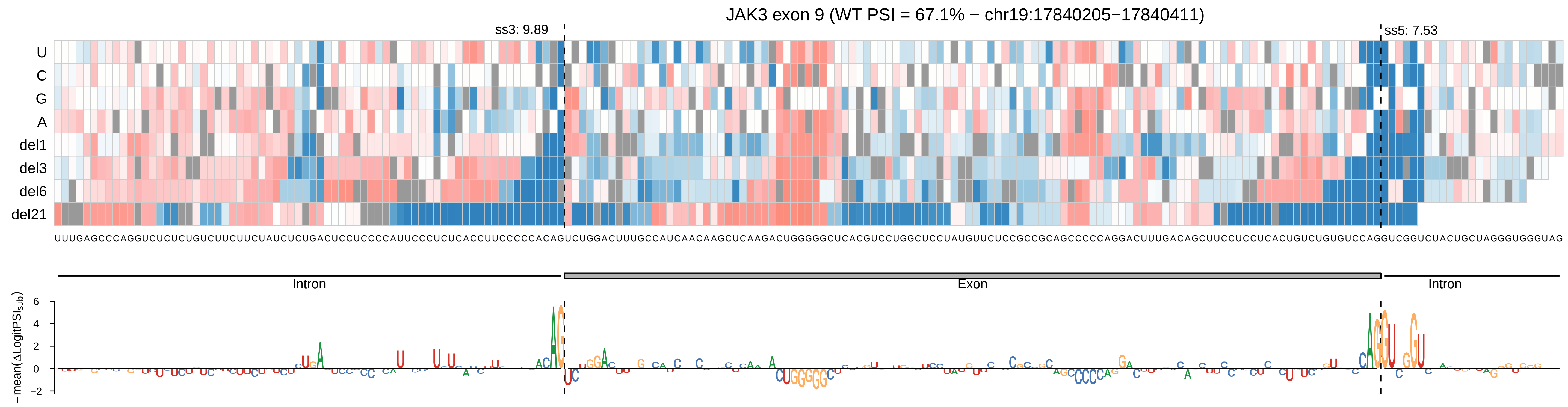

JAK3 exon 9 (WT PSI = 67.1% − chr19:17840205−17840411)
ss3: 9.89
ss5: 7.53
U
C
G
A
del1
del3
del6
del21
UUUGAGCCCAGGUCUCUCUGUCUUCUUCUAUCUCUGACUCCUCCCCAUUCCCUCUCACCUUCCCCCACAGUCUGGACUUUGCCAUCAACAAGCUCAAGACUGGGGGCUCACGUCCUGGCUCCUAUGUUCUCCGCCGCAGCCCCCAGGACUUUGACAGCUUCCUCCCUCACUGUCUGUGUCCAGGUCGGUCUACUGCUAGGGUGGGUAG
Intron
Exon
Intron
−mean(ΔLogitPSI$_{sub}$)

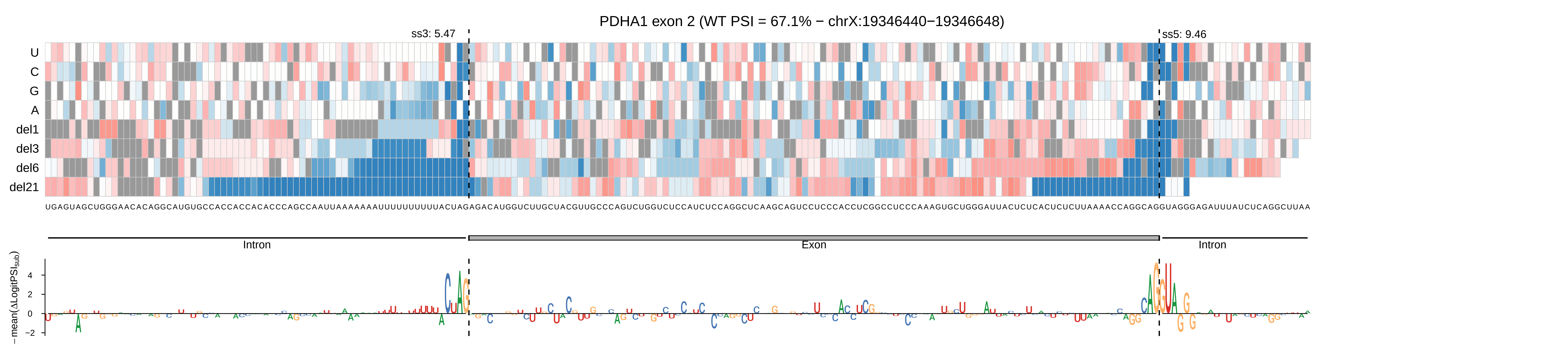
PDHA1 exon 2 (WT PSI = 67.1% − chrX:19346440−19346648)
ss3: 5.47
ss5: 9.46
U
C
G
A
del1
del3
del6
del21
UGAGUAGCUGGGAACACAGGCAUGUGCCACCACCACACCCAGCCAAUUAAAAAAAAUUUUUUUUUUACUAGAGACAUGGUCUUGCUACGUUGCCCAGUCUGGUCUCCAUCUCCAGGCUCAAGCAGUCCUCCCACCUCGGCCUCCCAAAGUGCUGGGAUUACUCUCACUCUCUUAAAACCAGGCAGGUAGGGAGAUUUAUCUCAGGCUUAA
Intron
Exon
Intron
−mean(ΔLogitPSI_sub)
4
2
0
−2

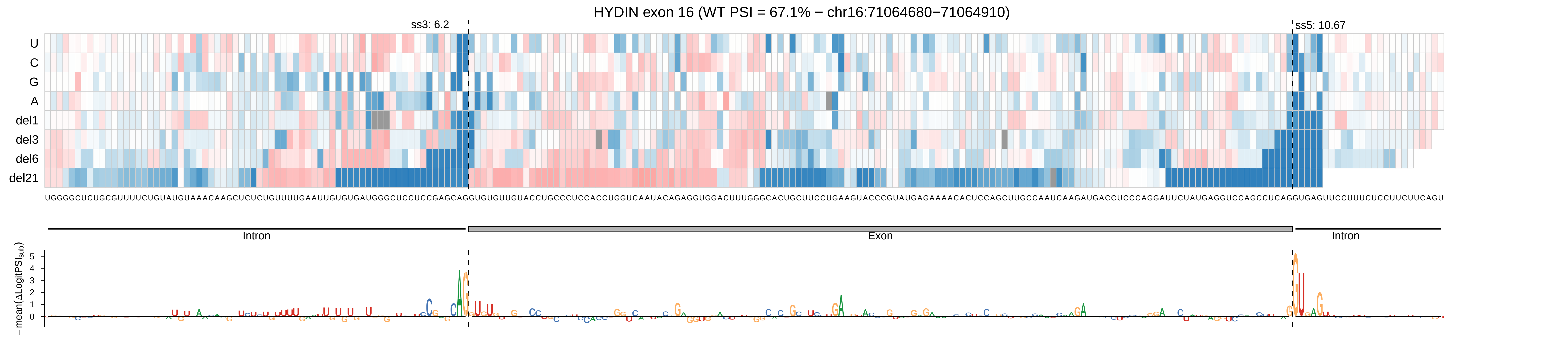
HYDIN exon 16 (WT PSI = 67.1% − chr16:71064680−71064910)
ss3: 6.2
ss5: 10.67
U
C
G
A
del1
del3
del6
del21
Intron
Exon
Intron
−mean(ΔLogitPSI_sub)
5
4
3
2
1
0

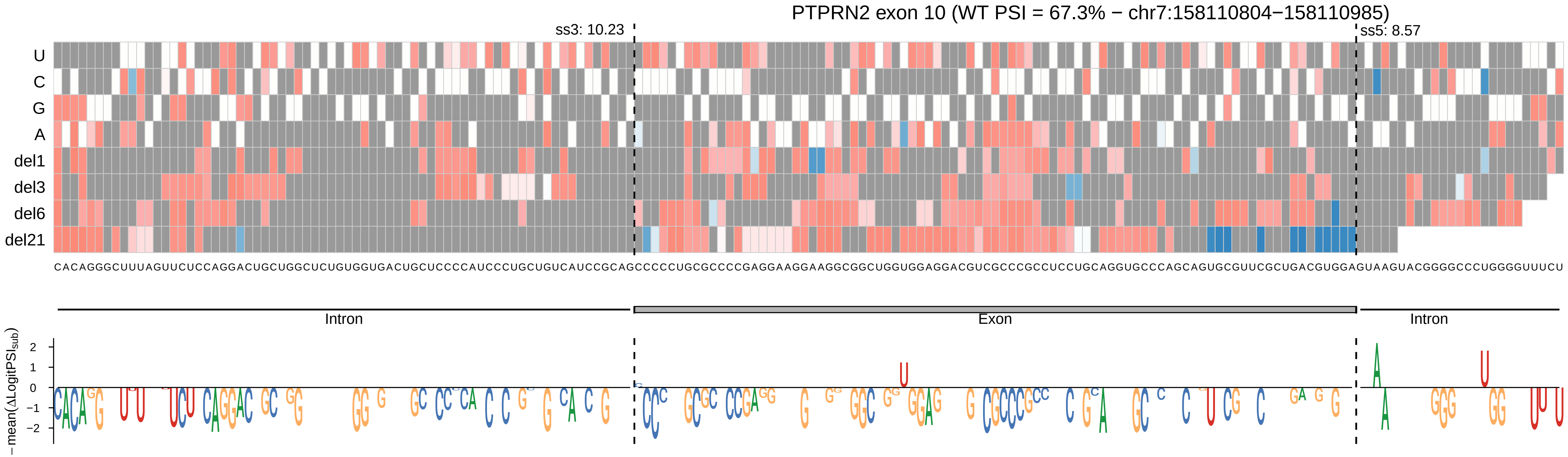
PTPRN2 exon 10 (WT PSI = 67.3% − chr7:158110804−158110985)
ss3: 10.23
ss5: 8.57
U
C
G
A
del1
del3
del6
del21
CACAGGGCUUUAGUUCUCCAGGACUGCUGGCUCUGUGGUGACUGCUCCCCAUCCCUGCUGUCAUCCGCAGCCCCCUGCGCCCCGAGGAAGGAAGGCGGCUGGUGGAGGACGUCGCCCGCCUCCUGCAGGUGCCCAGCAGUGCGUUCGCUGACGUGGAGUAAGUACGGGGCCCUGGGGUUUCU
Intron
Exon
Intron
−mean(ΔLogitPSI_sub)
2
1
0
−1
−2

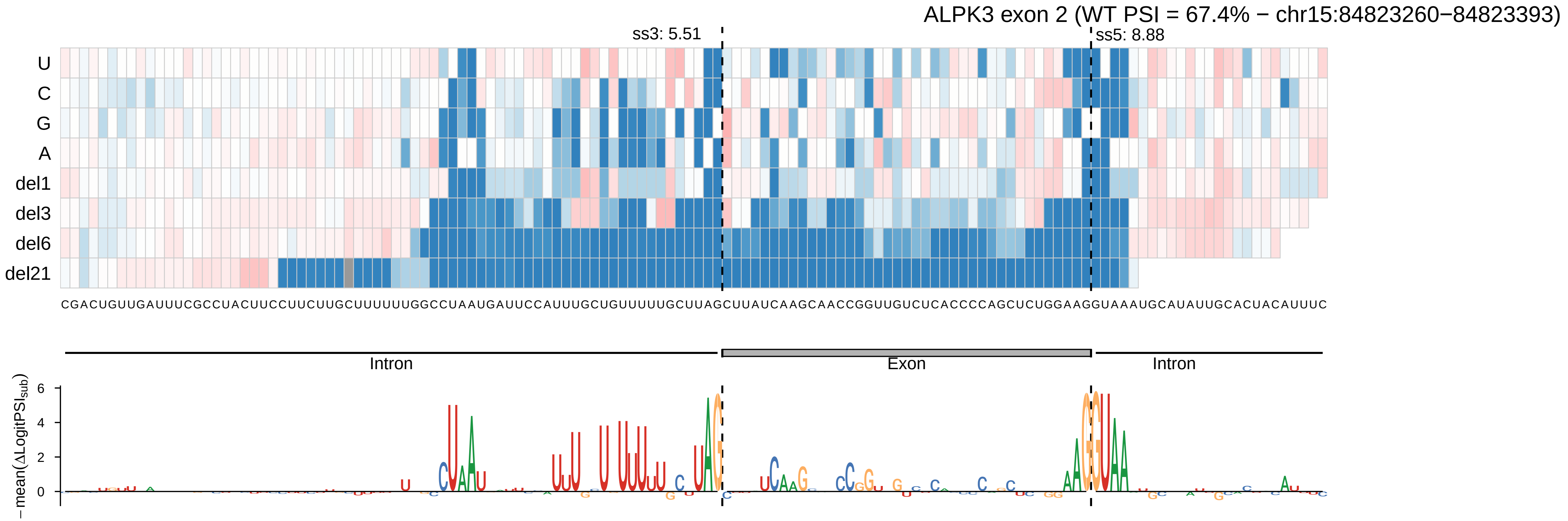
ALPK3 exon 2 (WT PSI = 67.4% − chr15:84823260−84823393)
ss3: 5.51
ss5: 8.88
U
C
G
A
del1
del3
del6
del21
CGACUGUUGAUUUCGCCUACUUCCUUCUUGCUUUUUUGGCCUAAUGAUUCCAUUUGCUGUUUUUGCUUAGCUUAUCAAGCAACCGGUUGUCUCACCCCAGCUCUGGAAGGUAAAUGCAUAUUGCACUACAUUUC
Intron
Exon
Intron
−mean(ΔLogitPSI_sub)
6
4
2
0

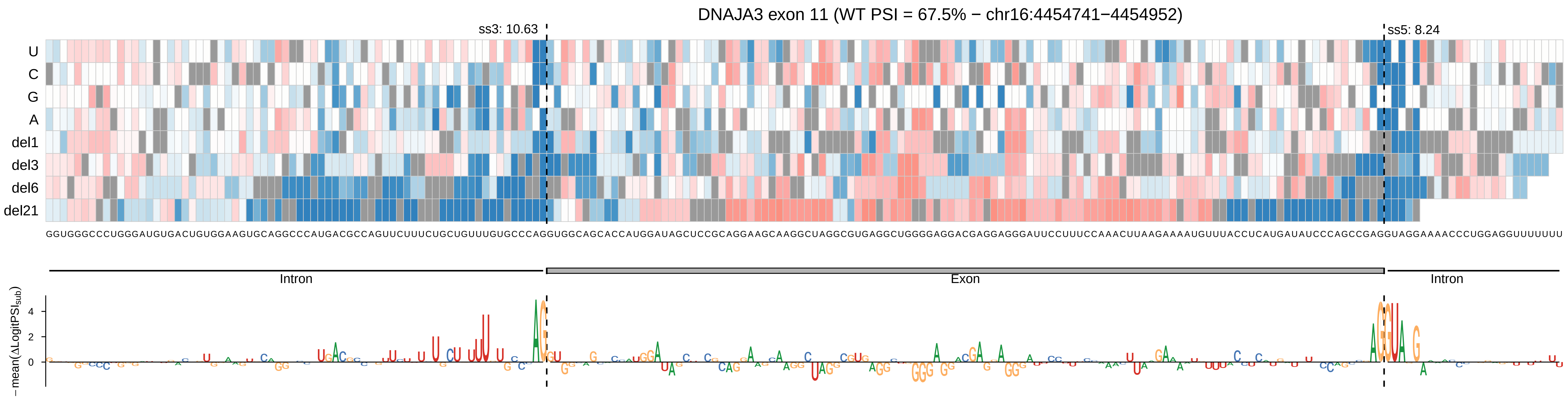
DNAJA3 exon 11 (WT PSI = 67.5% − chr16:4454741−4454952)
ss3: 10.63
ss5: 8.24
U
C
G
A
del1
del3
del6
del21
GGUGGGCCCUGGGAUGUGACUGUGGAAGUGCAGGCCCAUGACGCCAGUUCUUUCUGCUGUUUGUGCCCAGGUGGCAGCACCAUGGAUAGCUCCGCAGGAAGCAAGGCUAGGCGUGAGGCUGGGGAGGACGAGGAGGGAUUCCCUUUCCAAACUUAAGAAAAUGUUUACCUCAUGAUAUCCCAGCCGAGGUAGGAAAACCCUGGAGGUUUUUUU
Intron
Exon
Intron
4
2
0
−mean(ΔLogitPSI_sub)

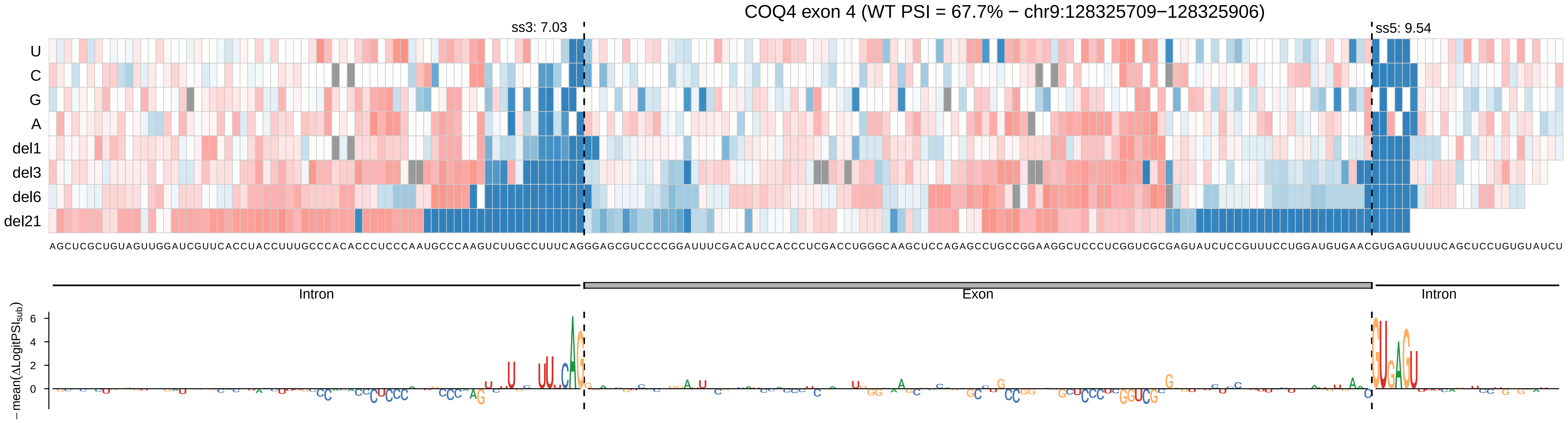

COQ4 exon 4 (WT PSI = 67.7% − chr9:128325709−128325906)
ss3: 7.03
ss5: 9.54
U
C
G
A
del1
del3
del6
del21
AGCUCGCUGUAGUUGGAUCGUUCACCUACCUUUGCCCACACCCUCCCAAUGCCCAAGUCUUGCCUUUCAGGGAGCGUCCCCGGAUUUCGACAUCCACCCUCGACCUGGGCAAGCUCCAGAGCCUGCCGGAAGGCUCCCUCGGUCGCGAGUAUCUCCGUUUCCUGGAUGUGAACGUGAGUUUUCAGCUCCUGUGUAUCU
Intron
Exon
Intron
−mean(ΔLogitPSIsub)

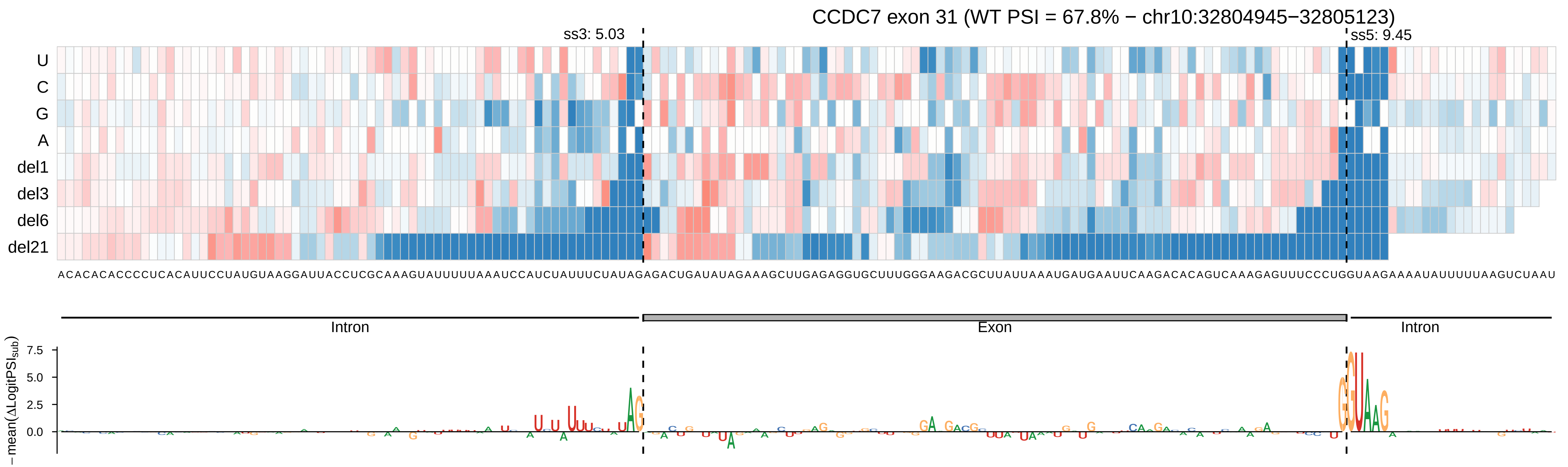

CCDC7 exon 31 (WT PSI = 67.8% − chr10:32804945−32805123)
ss3: 5.03
ss5: 9.45
U
C
G
A
del1
del3
del6
del21
ACACACACCCCUCACAUUCCUAUGUAAGGAUUACCUCGCAAAGUAUUUUUAAAUCCAUCUAUUUCUAUAGAGACUGAUAUAGAAAGCUUGAGAGGUGCUUUGGGAAGACGCUUAUUAAAUGAUGAAUUCAAGACACAGUCAAAGAGUUUCCCUGGUAAGAAAAUAUUUUUAAGUCUAAU
Intron
Exon
Intron
−mean(ΔLogitPSI_sub)
7.5
5.0
2.5
0.0

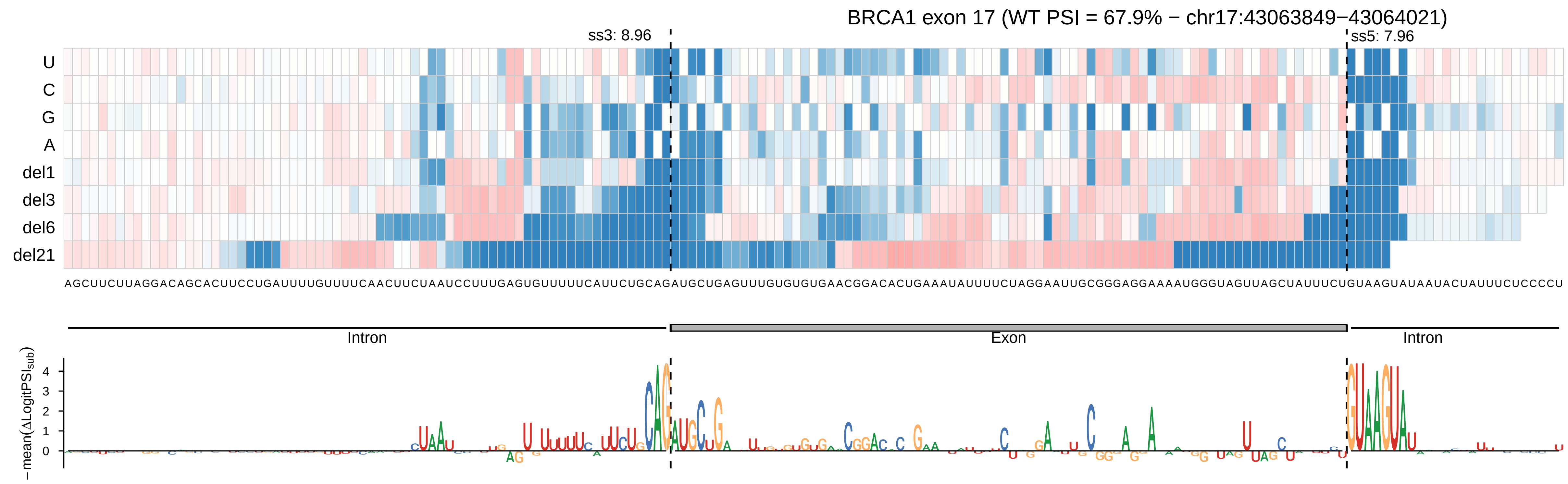

BRCA1 exon 17 (WT PSI = 67.9% − chr17:43063849−43064021)
ss3: 8.96
ss5: 7.96
U
C
G
A
del1
del3
del6
del21
AGCUUCUUAGGACAGCACUUCCUGAUUUUGUUUUCAACUUCUAAUCCUUUGAGUGUUUUUCAUUCUGCAGAUGCUGAGUUUGUGUGUGAACGGACACUGAAAUAUUUUCUAGGAAUUGCGGGAGGAAAAUGGGUAGUUAGCUAUUUCUGUAAGUAUAAUACUAUUUCUCCCCU
Intron
Exon
Intron
−mean(ΔLogitPSI_sub)
4
3
2
1
0

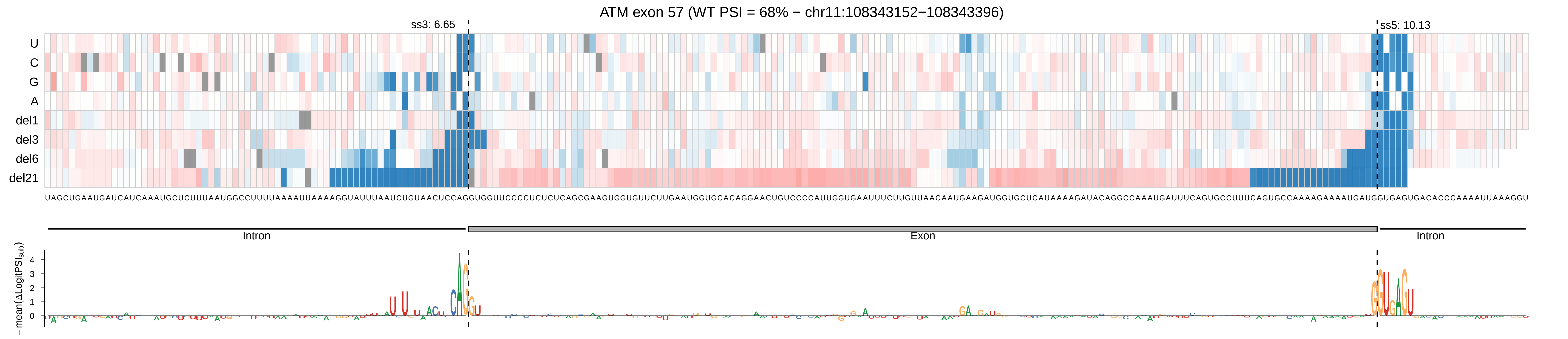
ATM exon 57 (WT PSI = 68% − chr11:108343152−108343396)
ss3: 6.65
ss5: 10.13
U
C
G
A
del1
del3
del6
del21
UAGCUGAAUGAUCAUCAAAUGCUCUUUAAUGGCCUUUUAAAAUUAAAAGGUAUUUAAUCUGUAACUCCAGGUGGUUCCCCUCUCUCAGCGAAGUGGUGUUCUUGAAUGGUGCACAGGAACUGUCCCCAUUGGUGAAUUUCUUGUUAACAAUGAAGAUGGUGCUCAUAAAAGAUACAGGCCAAAUGAUUUCAGUGCCUUUCAGUGCCAAAAGAAAAUGAUGGUGAGUGACACCCAAAAUUAAAGGU
Intron
Exon
Intron
−mean(ΔLogitPSI_sub)
4
3
2
1
0

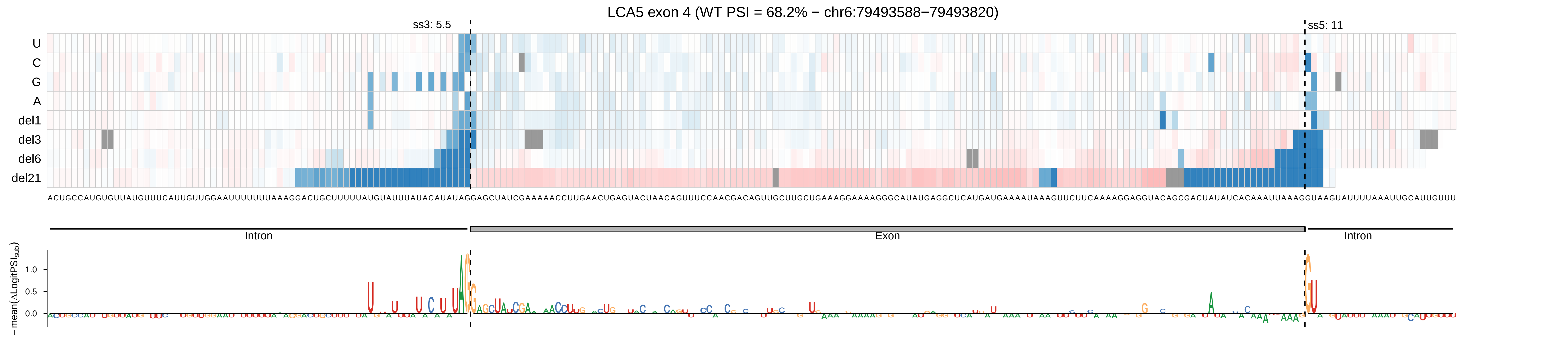
LCA5 exon 4 (WT PSI = 68.2% − chr6:79493588−79493820)
ss3: 5.5
ss5: 11
U
C
G
A
del1
del3
del6
del21
ACUGCCAUGUGUUAUGUUUCAUUGUUGGAAUUUUUUUAAAGGACUGCUUUUUAUGUAUUUAUACAUAUAGGAGCUAUCGAAAAACCUUGAACUGAGUACUAACAGUUUCCAACGACAGUUGCUUGCUGAAAGGAAAAGGGCAUAUGAGGCUCAUGAUGAAAAUAAAGUUCUUCAAAAGGAGGUACAGCGACUAUAUCACAAAUUAAAGGUAAGUAUUUUAAAUUGCAUUGUUU
Intron
Exon
Intron
−mean(ΔLogitPSI_sub)
1.0
0.5
0.0

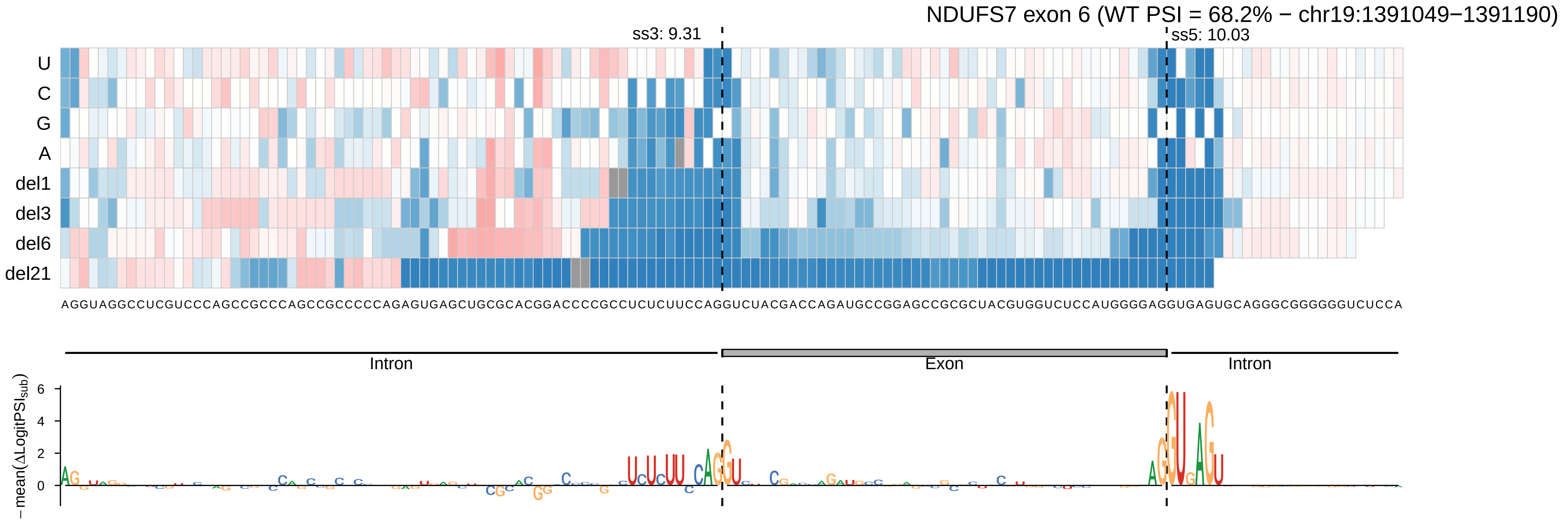
NDUFS7 exon 6 (WT PSI = 68.2% − chr19:1391049−1391190)
ss3: 9.31
ss5: 10.03
U
C
G
A
del1
del3
del6
del21
AGGUAGGCCUCGUCCCAGCCGCCCAGCCGCCCCCAGAGUGAGCUGCGCACGGACCCCGCCUCUCUUCCAGGUCUACGACCAGAUGCCGGAGCCGCGCUACGUGGUCUCCAUGGGGAGGUGAGUGCAGGGCGGGGGGUCUCCA
Intron
Exon
Intron
−mean(ΔLogitPSI_sub)
6
4
2
0

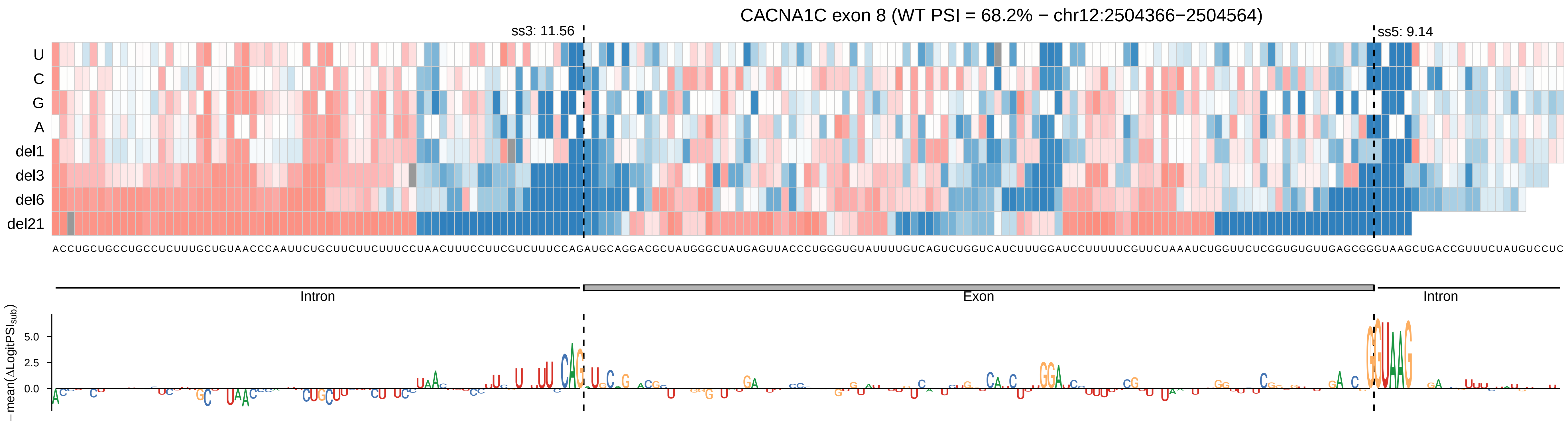

CACNA1C exon 8 (WT PSI = 68.2% − chr12:2504366−2504564)
ss3: 11.56
ss5: 9.14
U
C
G
A
del1
del3
del6
del21
Intron
Exon
Intron
−mean(ΔLogitPSI_sub)
5.0
2.5
0.0

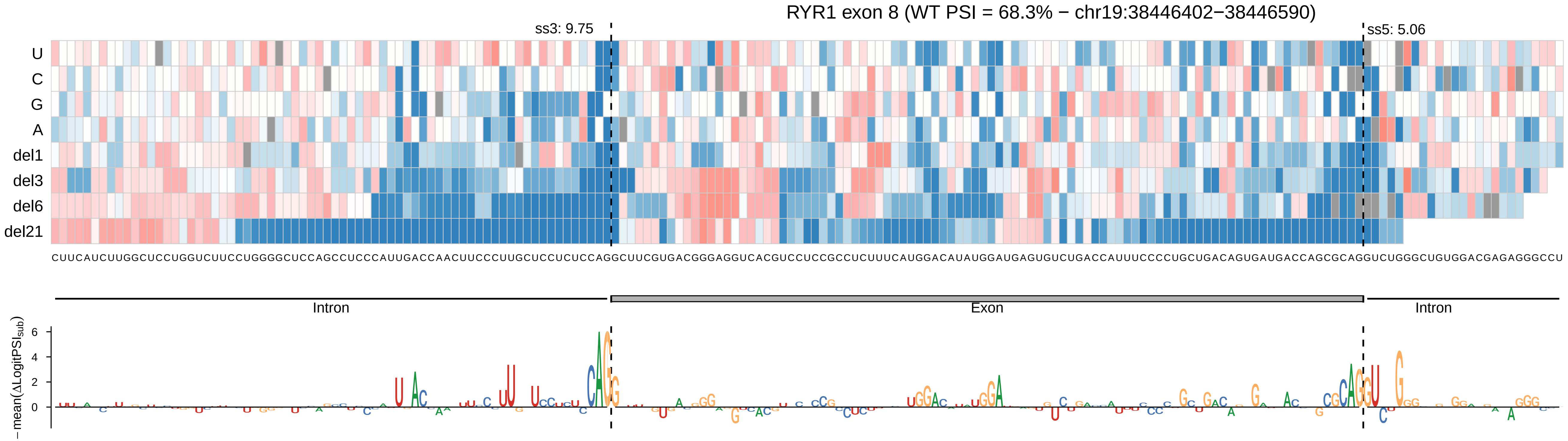
RYR1 exon 8 (WT PSI = 68.3% − chr19:38446402−38446590)
ss3: 9.75
ss5: 5.06
U
C
G
A
del1
del3
del6
del21
CUUCAUCUUGGCUCCUGGUCUUCCUGGGGCUCCAGCCUCCCAUUGACCAACUUCCCUUGCUCCUCUCCAGGCUUCGUGACGGGAGGUCACGUCCUCCGCCUCUUUCAUGGACAUAUGGAUGAGUGUCUGACCAUUUCCCCUGCUGACAGUGAUGACCAGCGCAGGUCUGGGCUGUGGACGAGAGGGCCU
Intron
Exon
Intron
−mean(ΔLogitPSI_sub)
6
4
2
0

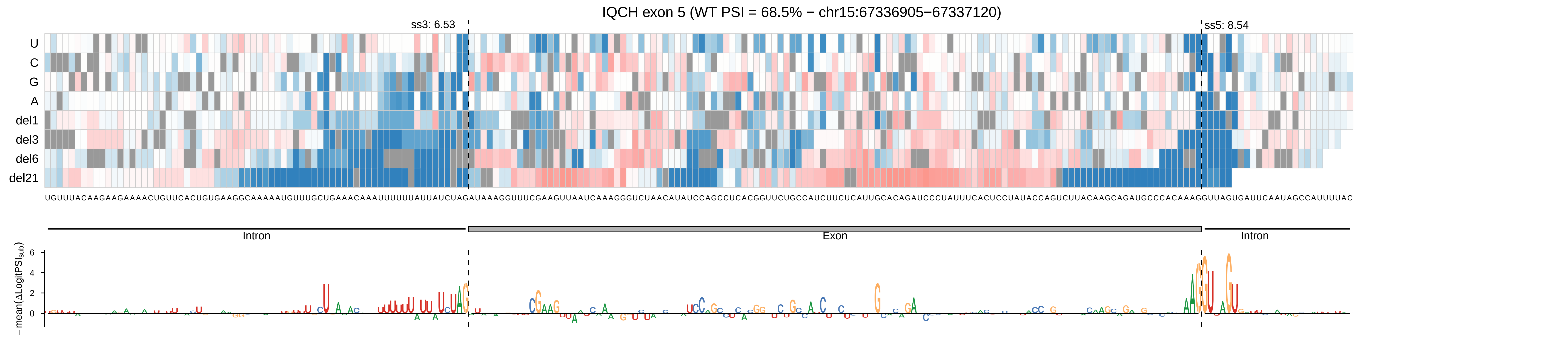

IQCH exon 5 (WT PSI = 68.5% − chr15:67336905−67337120)
ss3: 6.53
ss5: 8.54
U
C
G
A
del1
del3
del6
del21
UGUUUACAAGAAGAAAACUGUUCACUGUGAAGGCAAAAAUGUUUGCUGAAACAAAUUUUUUAUUAUCUAGAUAAAGGUUUCGAAGUUAAUCAAAGGGUCUAACAUAUCCAGCCUCACGGUUCUGCCAUCUUCUCAUUGCACAGAUCCCUAUUUCACUCCUAUACCAGUCUUACAAGCAGAUGCCCACAAAGGUUAGUGAUUCAAUAGCCAUUUUAC
Intron
Exon
Intron
−mean(ΔLogitPSI$_{sub}$)

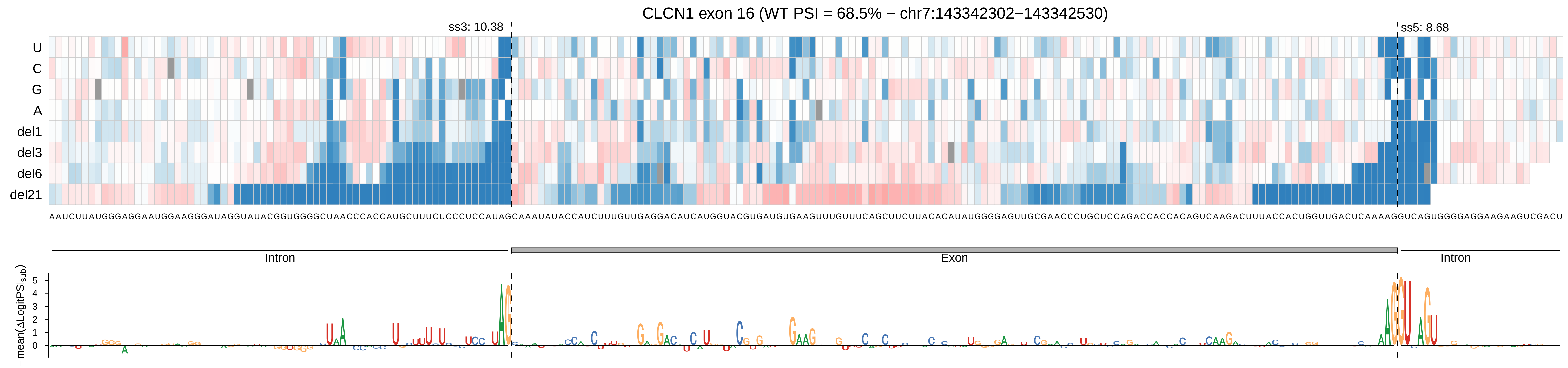
CLCN1 exon 16 (WT PSI = 68.5% − chr7:143342302−143342530)
ss3: 10.38
ss5: 8.68
U
C
G
A
del1
del3
del6
del21
AAUCUUAUGGGAGGAAUGGAAGGGAUAGGUAUACGGUGGGGCUAACCCACCAUGCUUUCUCCCUCCAUAGCAAAUAUACCAUCUUUGUUGAGGACAUCAUGGUACGUGAUGUGAAGUUUGUUUCAGCUUCUUACACAUAUGGGGAGUUGCGAACCCUGCUCCAGACCACCACAGUCAAGACUUUACCACUGGUUGACUCAAAAGGUCAGUGGGGAGGAAGAAGUCGACU
Intron
Exon
Intron
− mean(ΔLogitPSI$_{sub}$)
5
4
3
2
1
0

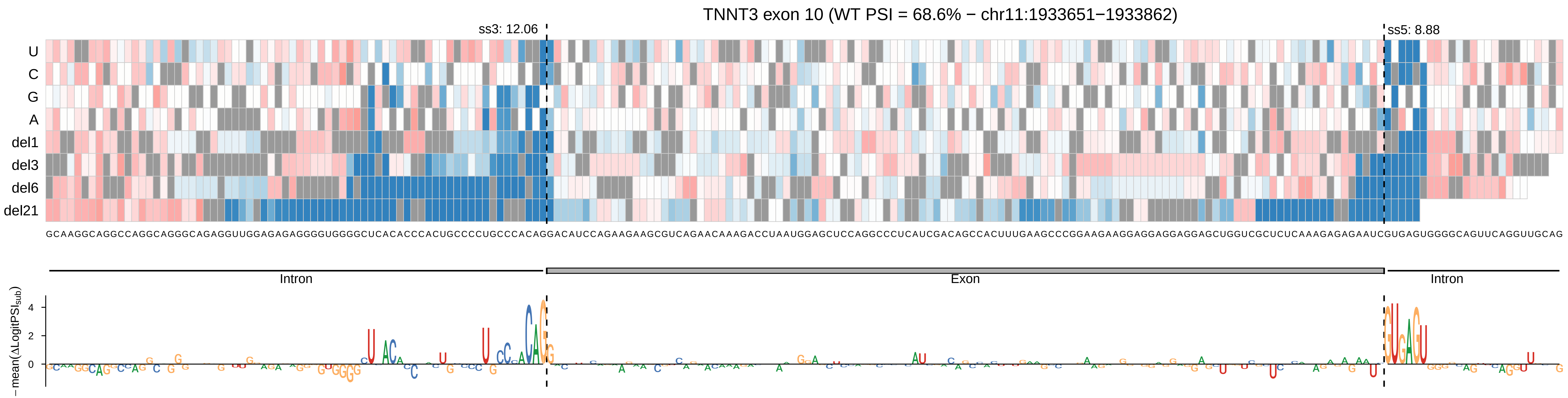
TNNT3 exon 10 (WT PSI = 68.6% − chr11:1933651−1933862)
ss3: 12.06
ss5: 8.88
U
C
G
A
del1
del3
del6
del21
Intron
Exon
Intron
−mean(ΔLogitPSI_sub)
4
2
0

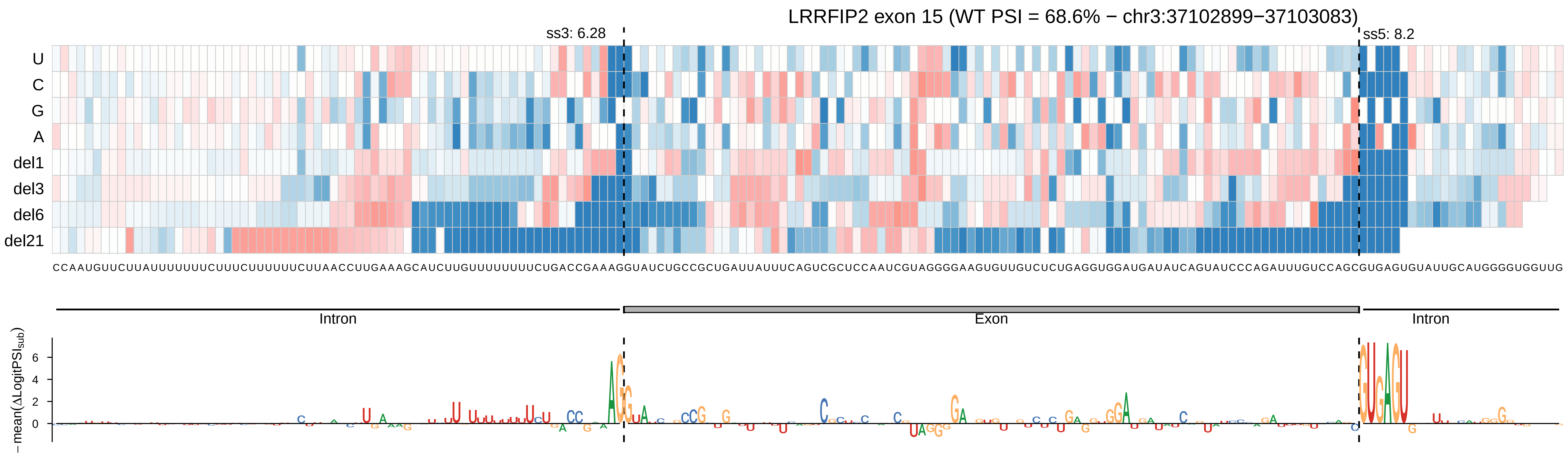
LRRFIP2 exon 15 (WT PSI = 68.6% − chr3:37102899−37103083)
ss3: 6.28
ss5: 8.2
U
C
G
A
del1
del3
del6
del21
CCAAUGUUCUUAUUUUUUUCUUUCUUUUUUCUUAACCUUGAAAGCAUCUUGUUUUUUUUCUGACCGAAAGGUAUCUGCCGCUGAUUAUUUCAGUCGCUCCAAUCGUAGGGGAAGUGUUGUCUCUGAGGUGGAUGAUAUCAGUAUCCCAGAUUUGUCCAGCGUGAGUGUAUUGCAUGGGGUGGUUG
Intron
Exon
Intron
−mean(ΔLogitPSIsub)
6
4
2
0

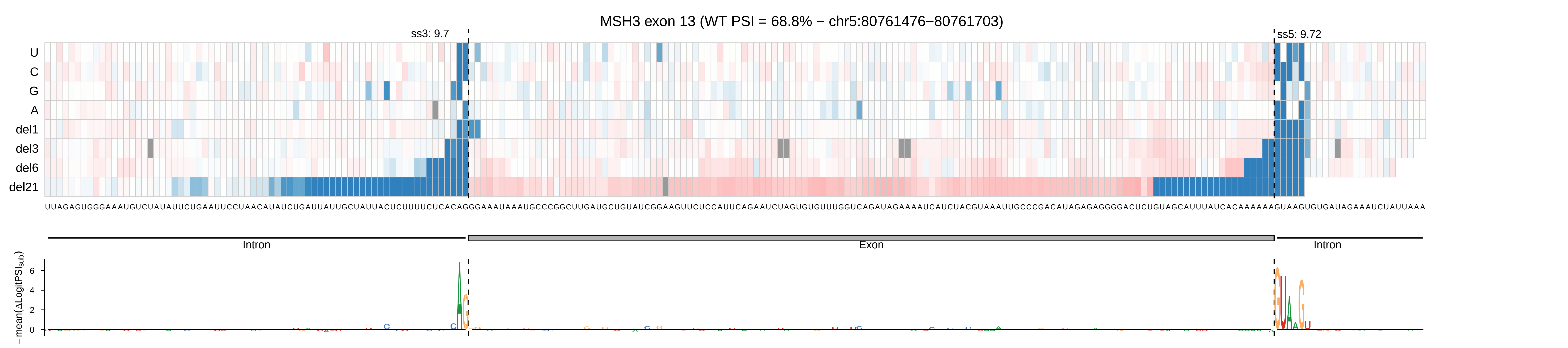
MSH3 exon 13 (WT PSI = 68.8% − chr5:80761476−80761703)
ss3: 9.7
ss5: 9.72
U
C
G
A
del1
del3
del6
del21
UUAGAGUGGGAAAUGUCUAUAUUCUGAAUUCCUAACAUAUCUGAUUAUUGCUAUUACUCUUUUCUCACAGGGAAAUAAAUGCCCGGCUUGAUGCUGUAUCGGAAGUUCUCCAUUCAGAAUCUAGUGUGUUUGGUCAGAUAGAAAAUCAUCUACGUAAAUUGCCCGACAUAGAGAGGGGACUCUGUAGCAUUUAUCACAAAAAAGUAAGUGUGAUAGAAAUCUAUUAAA
Intron
Exon
Intron
− mean(ΔLogitPSI_sub)
6
4
2
0

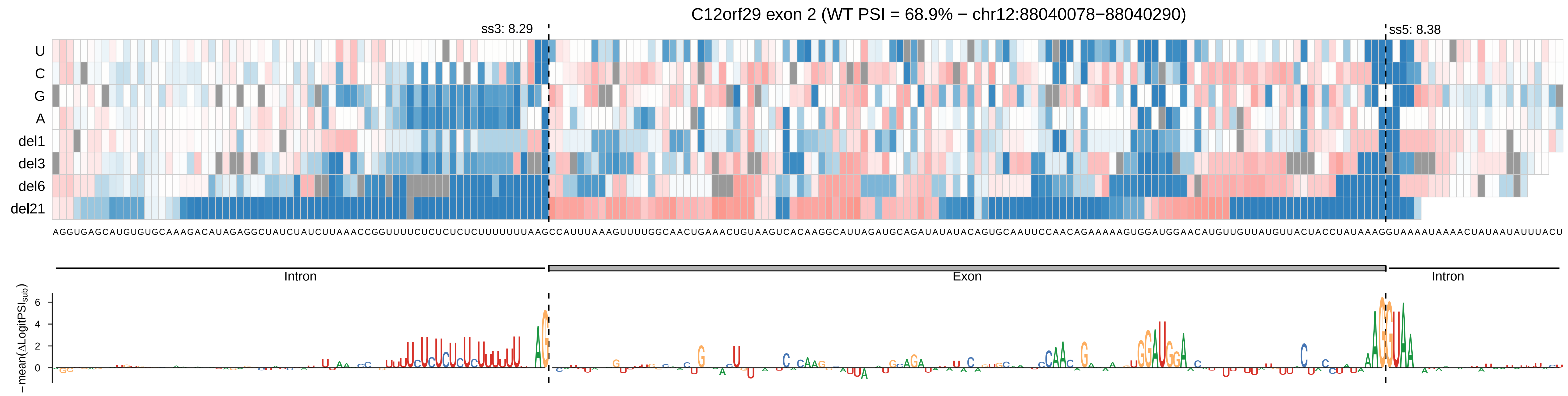
C12orf29 exon 2 (WT PSI = 68.9% − chr12:88040078−88040290)
ss3: 8.29
ss5: 8.38
U
C
G
A
del1
del3
del6
del21
AGGUGAGCAUGUGUGCAAAGACAUAGAGGCUAUCUAUCUUAAACCGGUUUUCUCUCUCUUUUUUUUAAGCCAUUUAAAGUUUUGGCAACUGAAACUGUAAGUCACAAGGCAUUAGAUGCAGAUAUAUACAGUGCAAUUCCAACAGAAAAAGUGGAUGGAACAUGUUGUUAUGUUACUACCUAUAAAGGUAAAAUAAAACUAUAAUAUUUACU
Intron
Exon
Intron
−mean(ΔLogitPSI_sub)
6
4
2
0

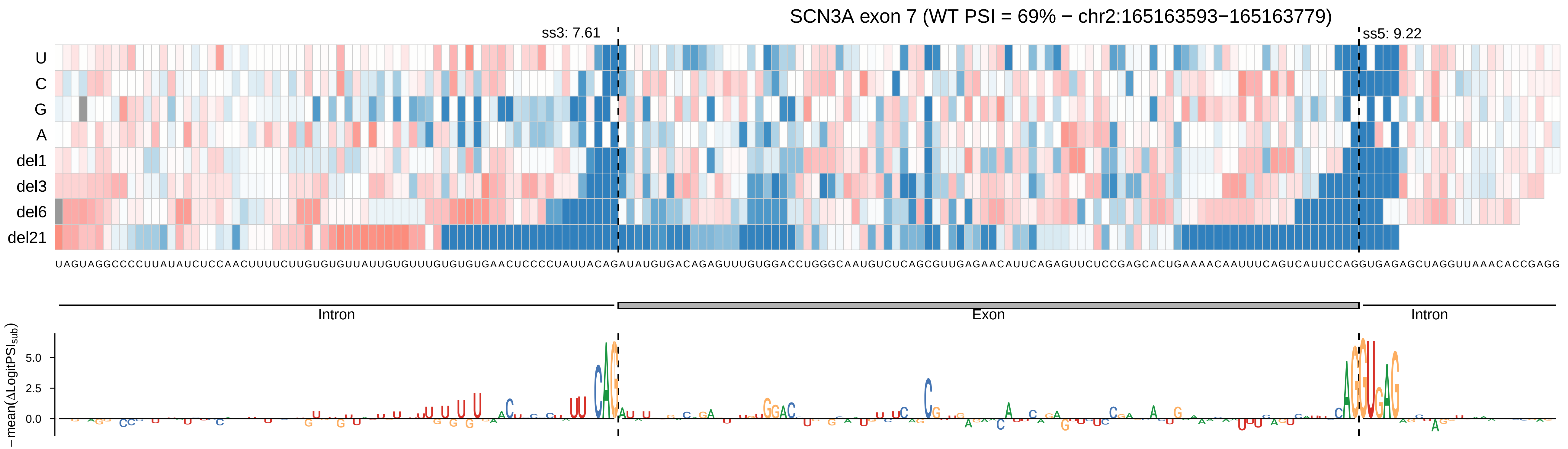

SCN3A exon 7 (WT PSI = 69% − chr2:165163593−165163779)
ss3: 7.61
ss5: 9.22
U
C
G
A
del1
del3
del6
del21
UAGUAGGCCCCUUAUAUCUCCAACUUUUCUUGUGUGUUAUUGUGUUUGUGUGUGAACUCCCCUAUUACAGAUAUGUGACAGAGUUUGUGGACCUGGGCAAUGUCUCAGCGUUGAGAACAUUCAGAGUUCUCCGAGCACUGAAAACAAUUUCAGUCAUUCCAGGUGAGAGCUAGGUUAAACACCGAGG
Intron
Exon
Intron
−mean(ΔLogitPSI_sub)
5.0
2.5
0.0

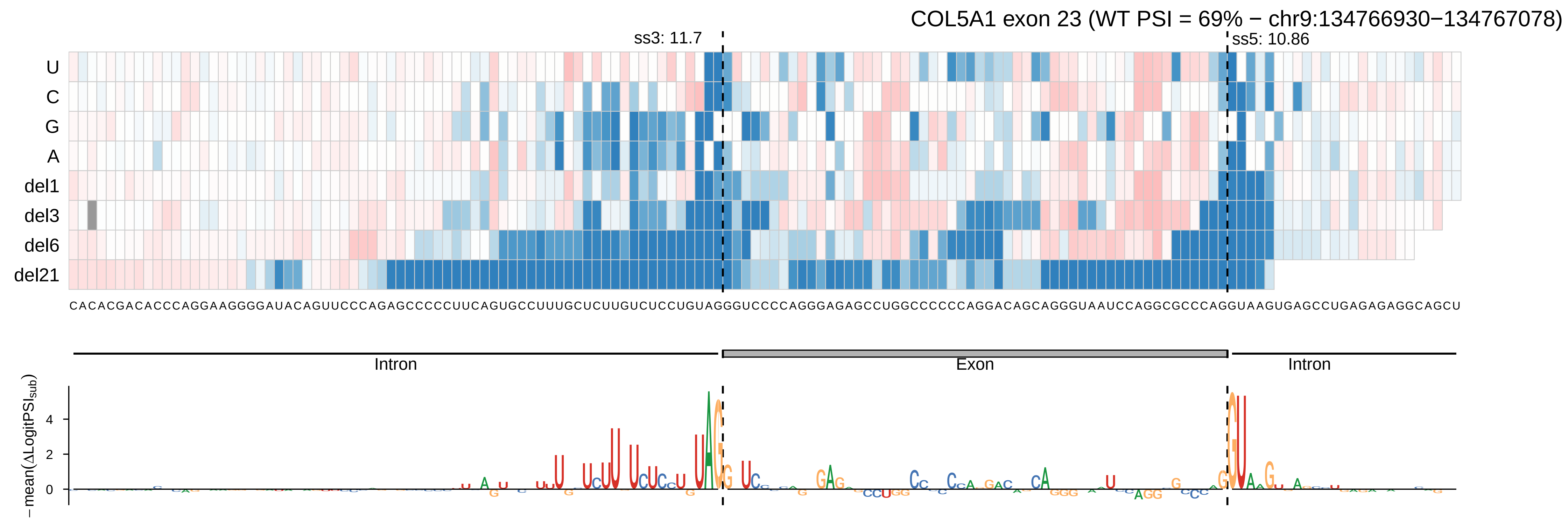
COL5A1 exon 23 (WT PSI = 69% − chr9:134766930−134767078)
ss3: 11.7
ss5: 10.86
U
C
G
A
del1
del3
del6
del21
CACACGACACCCAGGAAGGGGAUACAGUUCCCAGAGCCCCCUUCAGUGCCUUUGCUCUUGUCUCCUGUAGGGUCCCCAGGGAGAGCCUGGCCCCCCAGGACAGCAGGGUAAUCCAGGCGCCCAGGUAAGUGAGCCUGAGAGAGGCAGCU
Intron
Exon
Intron
−mean(ΔLogitPSI_sub)
4
2
0

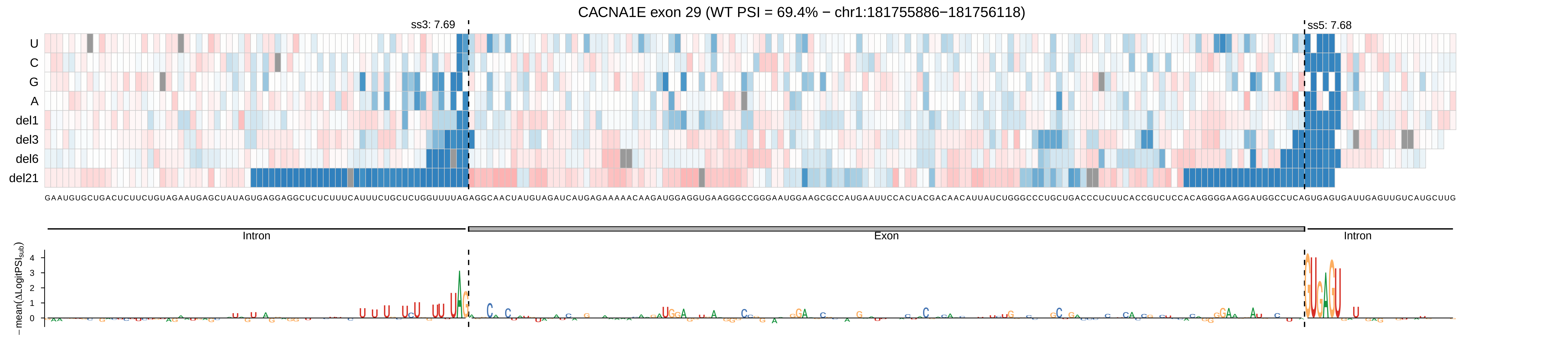

CACNA1E exon 29 (WT PSI = 69.4% − chr1:181755886−181756118)
ss3: 7.69
ss5: 7.68
U
C
G
A
del1
del3
del6
del21
Intron
Exon
Intron
−mean(ΔLogitPSI_sub)

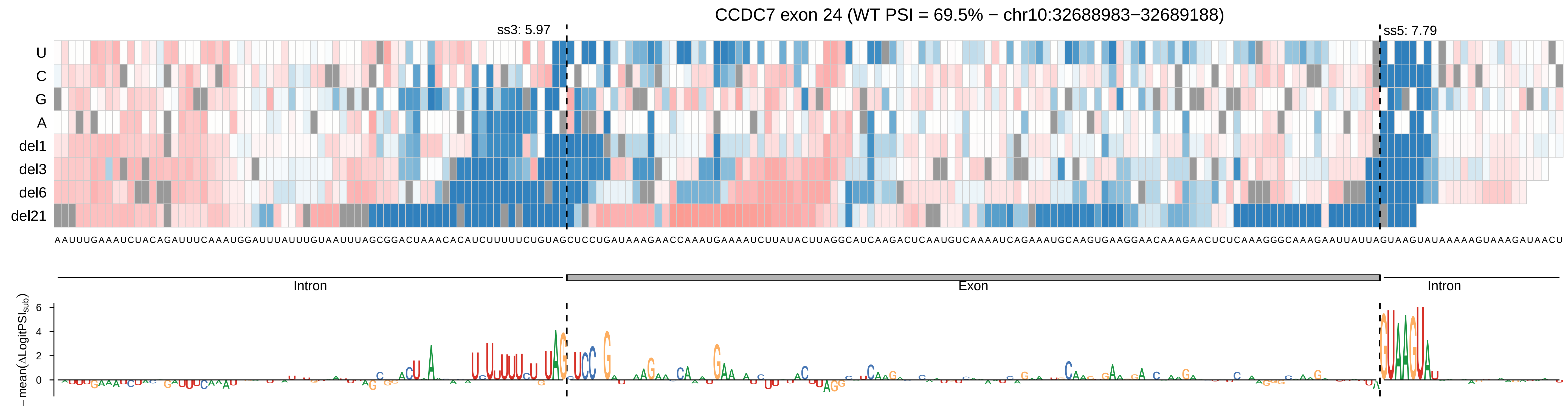

CCDC7 exon 24 (WT PSI = 69.5% − chr10:32688983−32689188)
ss3: 5.97
ss5: 7.79
U
C
G
A
del1
del3
del6
del21
AAUUUGAAAUCUACAGAUUUCAAAUGGAUUUAUUUGUAAUUUAGCGGACUAAACACAUCUUUUUCUGUAGCUCCUGAUAAAGAACCAAAUGAAAAUCUUAUACUUAGGCAUCAAGACUCAAUGUCAAAAUCAGAAAUGCAAGUGAAGGAACAAAGAACUCUCAAAGGGCAAAGAAUUAUUAGUAAGUAUAAAAAGUAAAGAUAACU
Intron
Exon
Intron
−mean(ΔLogitPSI_sub)
6
4
2
0

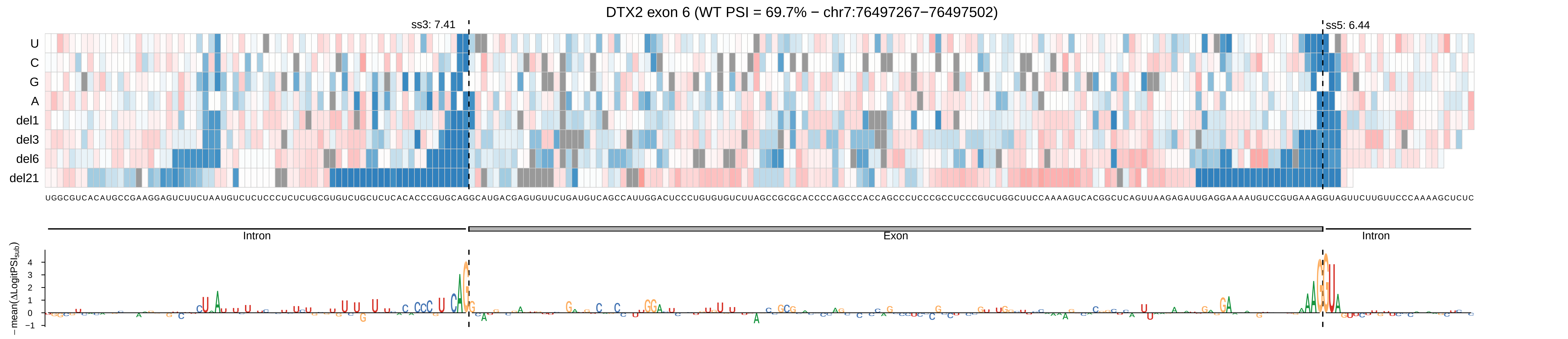
DTX2 exon 6 (WT PSI = 69.7% − chr7:76497267−76497502)
ss3: 7.41
ss5: 6.44
U
C
G
A
del1
del3
del6
del21
UGGCGUCACAUGCCGAAGGAGUCUUCUAAUGUCUCUCCCUCUCUGCGUGUCUGCUCUCACACCCGUGCAGGCAUGACGAGUGUUCUGAUGUCAGCCAUUGGACUCCCUGUGUGUCUUAGCCGCGCACCCCAGCCCACCAGCCCUCCCGCCUCCCGUCUGGCUUCCAAAAGUCACGGCUCAGUUAAGAGAUUGAGGAAAAUGUCCGUGAAAGGUAGUUCUUGUUCCCAAAAGCUCUC
Intron
Exon
Intron
− mean(ΔLogitPSI$_{sub}$)
4
3
2
1
0
−1

MYBPC3 exon 14 (WT PSI = 69.7% − chr11:47343235−47343332)

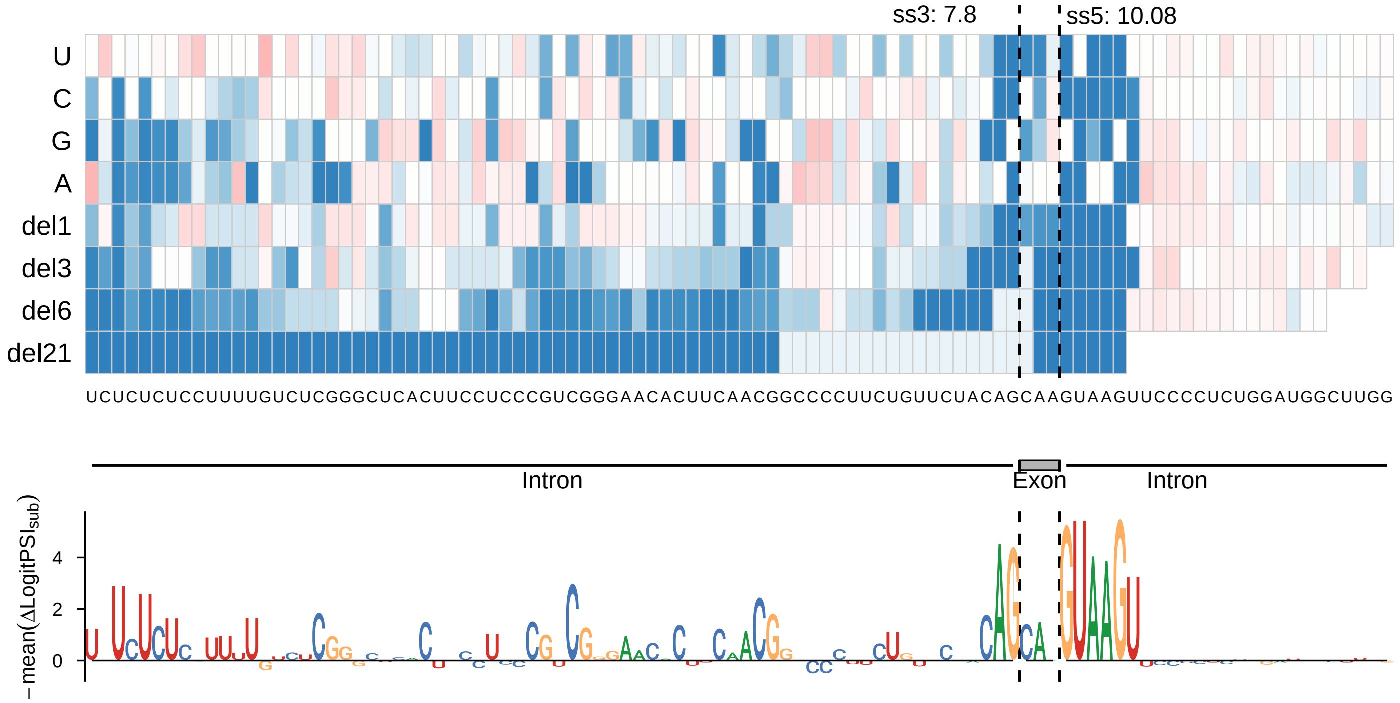

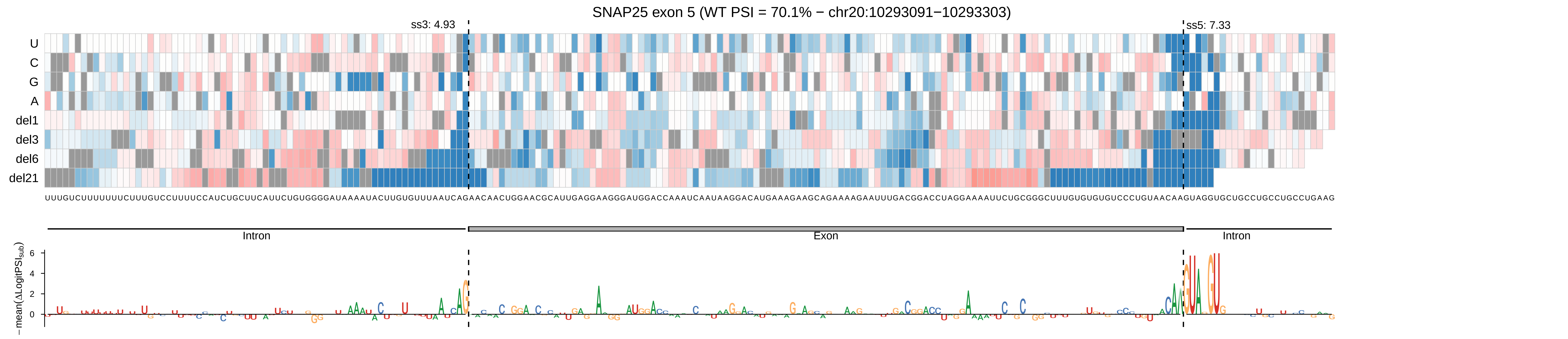
SNAP25 exon 5 (WT PSI = 70.1% − chr20:10293091−10293303)
ss3: 4.93
ss5: 7.33
U
C
G
A
del1
del3
del6
del21
Intron
Exon
Intron
−mean(ΔLogitPSI_sub)
6
4
2
0

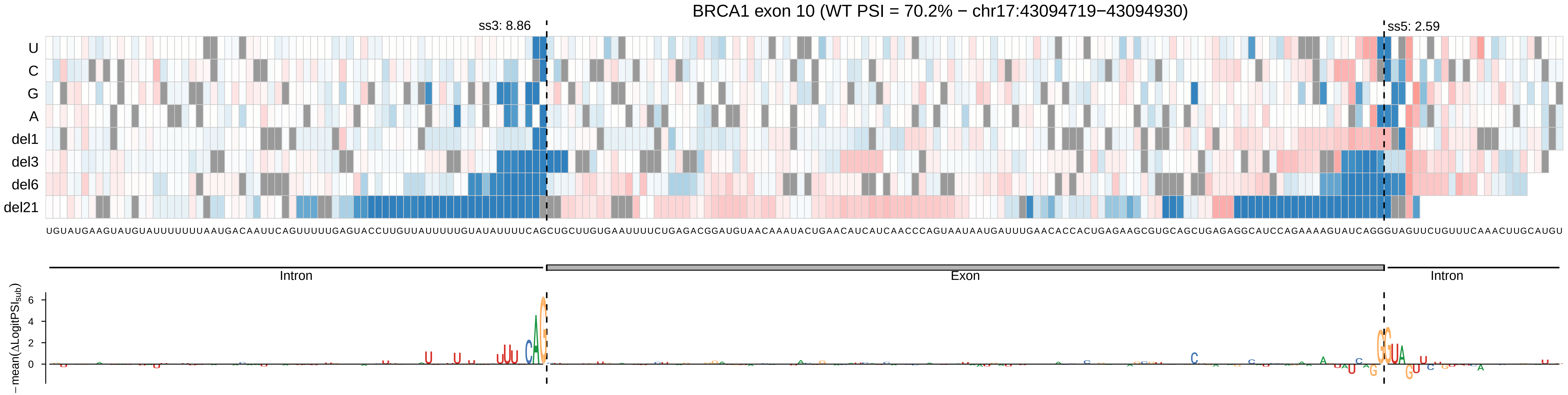

BRCA1 exon 10 (WT PSI = 70.2% − chr17:43094719−43094930)
ss3: 8.86
ss5: 2.59
U
C
G
A
del1
del3
del6
del21
UGUAUGAAGUAUGUAUUUUUUUAAUGACAAUUCAGUUUUUGAGUACCUUGUUAUUUUUGUAUAUUUUCAGCUGCUUGUGAAUUUUCUGAGACGGAUGUAACAAAUACUGAACAUCAUCAACCCAGUAAUAAUGAUUUGAACACCACUGAGAAGCGUGCAGCUGAGAGGCAUCCAGAAAAGUAUCAGGGUAGUUCUGUUUCAAACUUGCAUGU
Intron
Exon
Intron
−mean(ΔLogitPSI_sub)
6
4
2
0

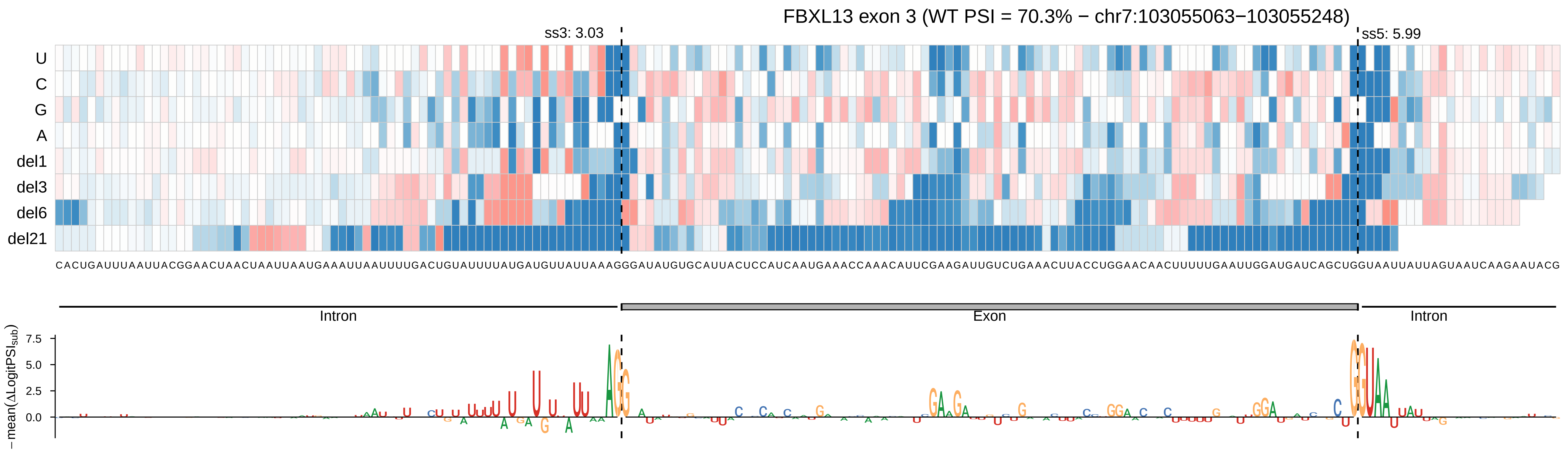

FBXL13 exon 3 (WT PSI = 70.3% − chr7:103055063−103055248)
ss3: 3.03
ss5: 5.99
U
C
G
A
del1
del3
del6
del21
Intron
Exon
Intron
7.5
5.0
2.5
0.0
− mean(ΔLogitPSI_sub)

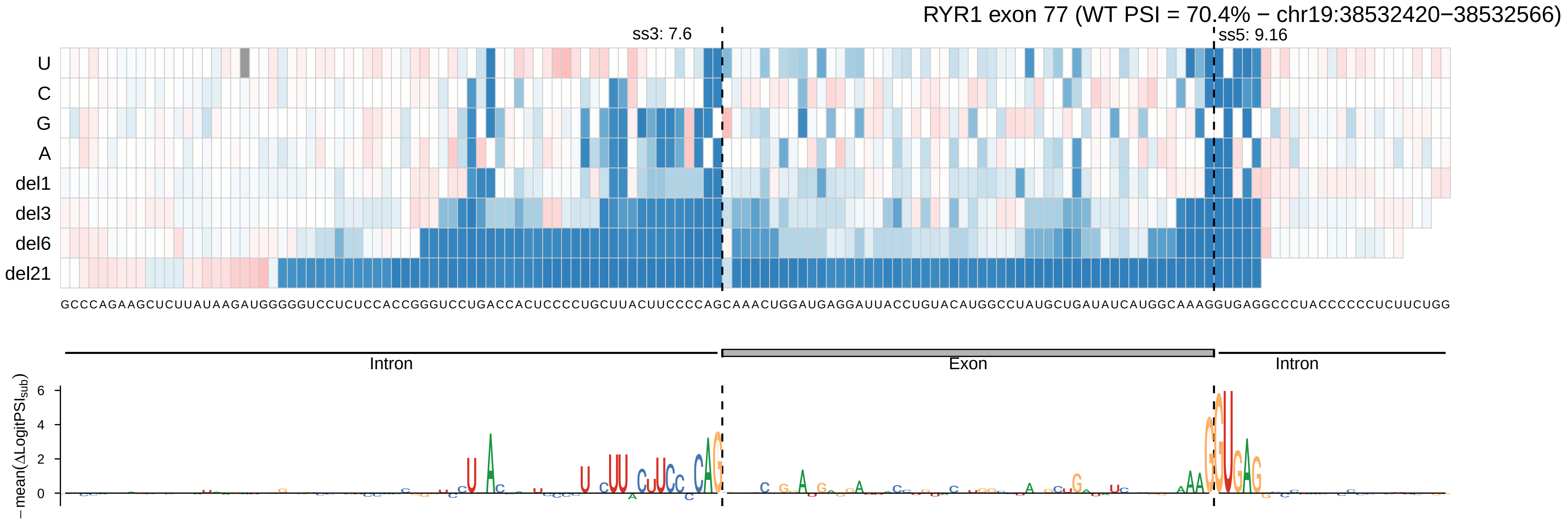

RYR1 exon 77 (WT PSI = 70.4% − chr19:38532420−38532566)
ss3: 7.6
ss5: 9.16
U
C
G
A
del1
del3
del6
del21
GCCCAGAAGCUCUUAUAAGAUGGGGUCCUCUCCACCGGGUCCUGACCACUCCCCUGCUUACUUCCCCAGCAAACUGGAUGAGGAUUACCUGUACAUGGCCUAUGCUGAUAUCAUGGCAAAGGUGAGGCCCUACCCCCCUCUUCUGG
Intron
Exon
Intron
− mean(ΔLogitPSI_sub)
6
4
2
0

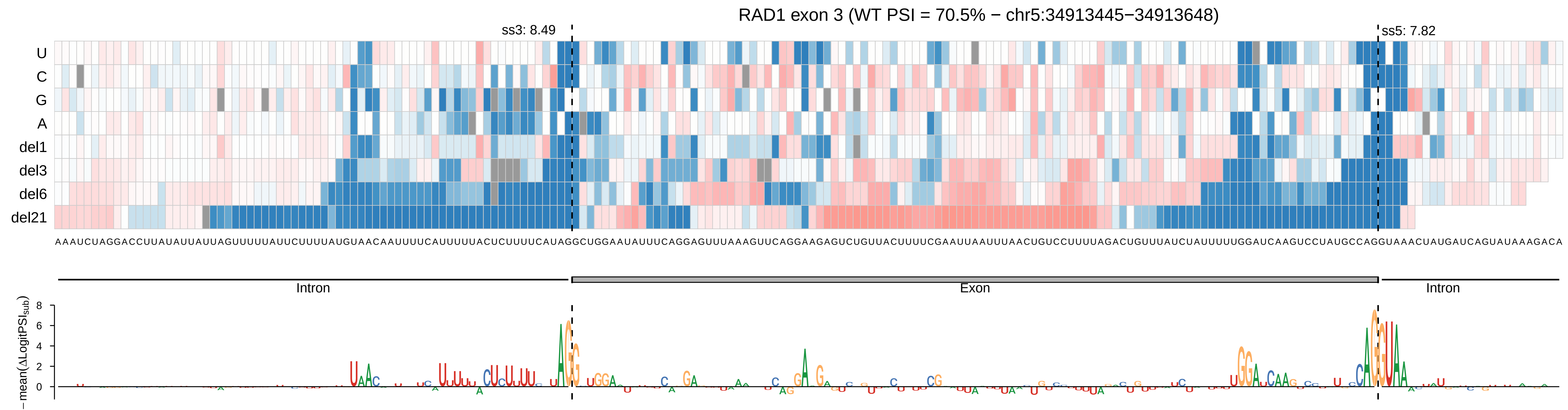

RAD1 exon 3 (WT PSI = 70.5% − chr5:34913445−34913648)
ss3: 8.49
ss5: 7.82
U
C
G
A
del1
del3
del6
del21
Intron
Exon
Intron
−mean(ΔLogitPSI_sub)

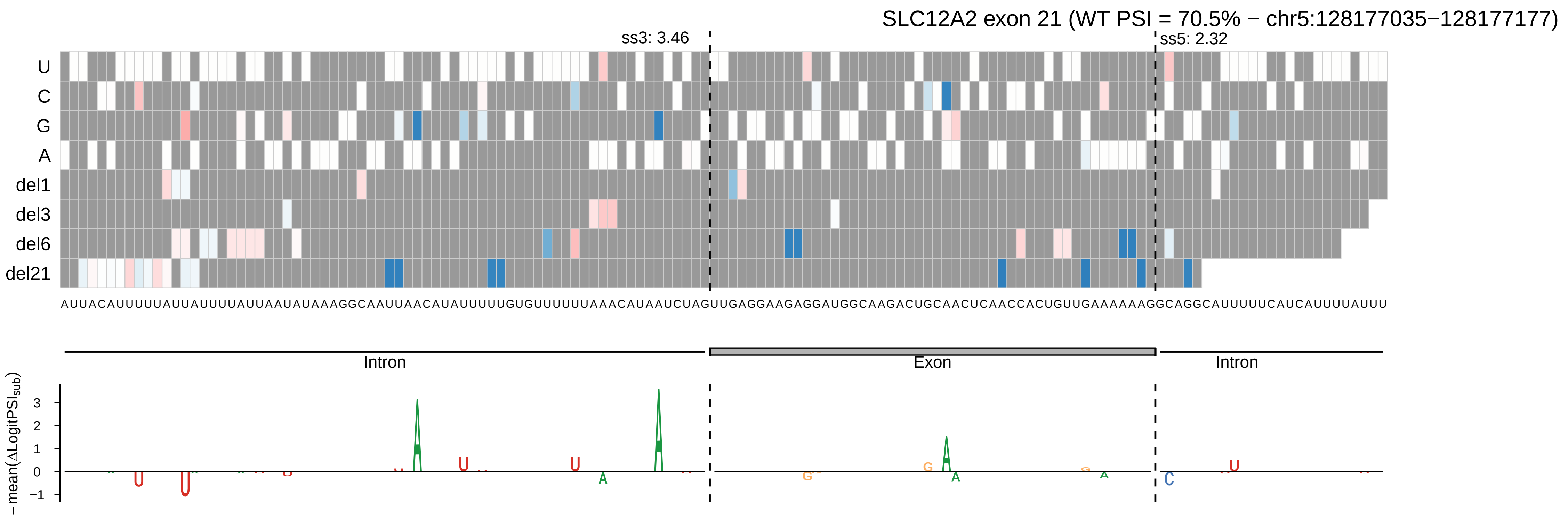
SLC12A2 exon 21 (WT PSI = 70.5% − chr5:128177035−128177177)
ss3: 3.46
ss5: 2.32
U
C
G
A
del1
del3
del6
del21
AUUACAUUUUUAUUAUUUUAUUAAUAUAAAGGCAAUUAACAUAUUUUUGUGUUUUUUAAACAUAAUCUAGUUGAGGAAGAGGAUGGCAAGACUGCAACUCAACCACUGUUGAAAAAAGGCAGGCAUUUUUCAUCAUUUUAUUU
Intron
Exon
Intron
−mean(ΔLogitPSI_sub)
3
2
1
0
−1

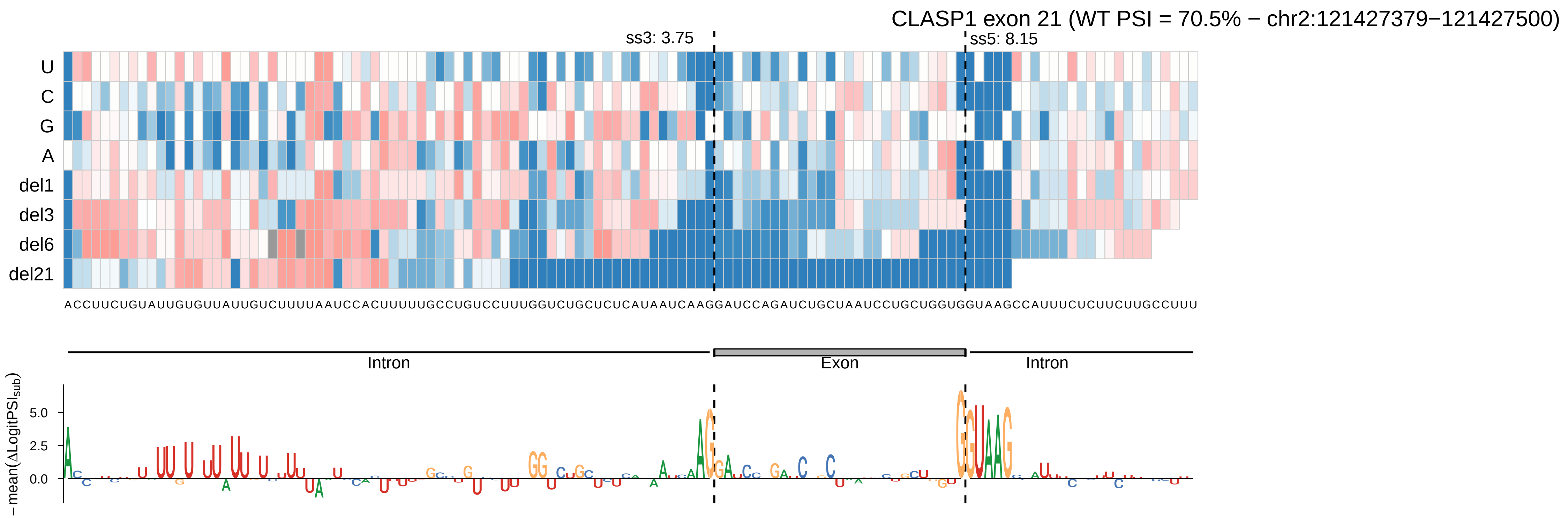
CLASP1 exon 21 (WT PSI = 70.5% − chr2:121427379−121427500)
ss3: 3.75
ss5: 8.15
U
C
G
A
del1
del3
del6
del21
ACCUUCUGUAUUGUGUUAUUGUCUUUUAAUCCACUUUUUGCCUGUCCUUUGGUCUGCUCUCAUAAUCAAGGAUCCAGAUCUGCUAAUCCUGCUGGUGGUAAGCCAUUUCUCUUCUUGCCUUU
Intron
Exon
Intron
− mean(ΔLogitPSI_sub)
5.0
2.5
0.0

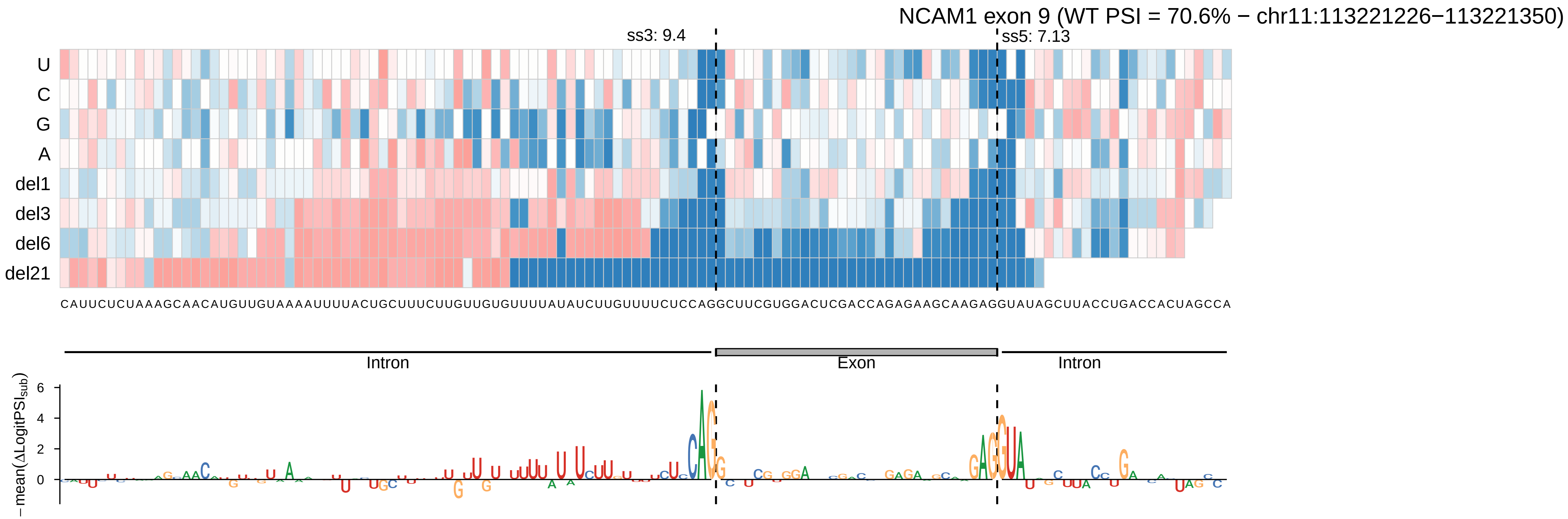

NCAM1 exon 9 (WT PSI = 70.6% − chr11:113221226−113221350)
ss3: 9.4
ss5: 7.13
U
C
G
A
del1
del3
del6
del21
CAUUCUCUAAAGCAACAUGUUGUAAAAUUUUACUGCUUUCUUGUUGUGUUUUAUAUCUUGUUUUCUCCAGGCUUCGUGGACUCGACCAGAGAAGCAAGAGGUAUAGCUUACCUGACCACUAGCCA
Intron
Exon
Intron
−mean(ΔLogitPSI_sub)
6
4
2
0

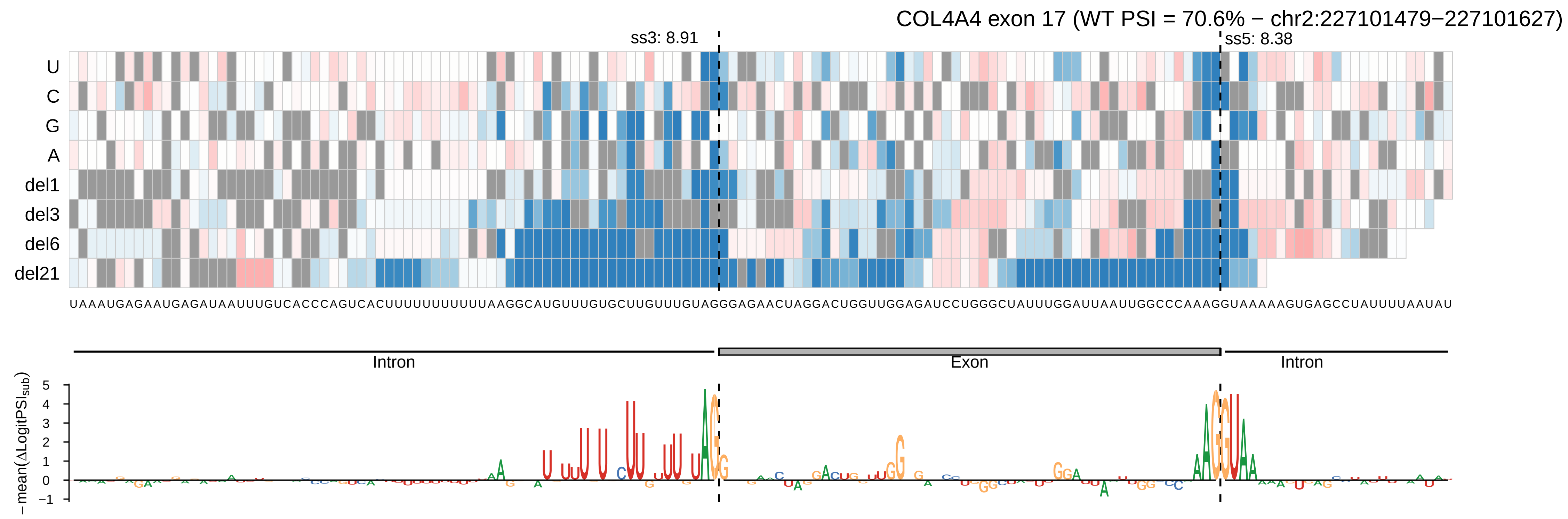
COL4A4 exon 17 (WT PSI = 70.6% − chr2:227101479−227101627)
ss3: 8.91
ss5: 8.38
U
C
G
A
del1
del3
del6
del21
Intron
Exon
Intron
− mean(ΔLogitPSI_sub)
5
4
3
2
1
0
−1

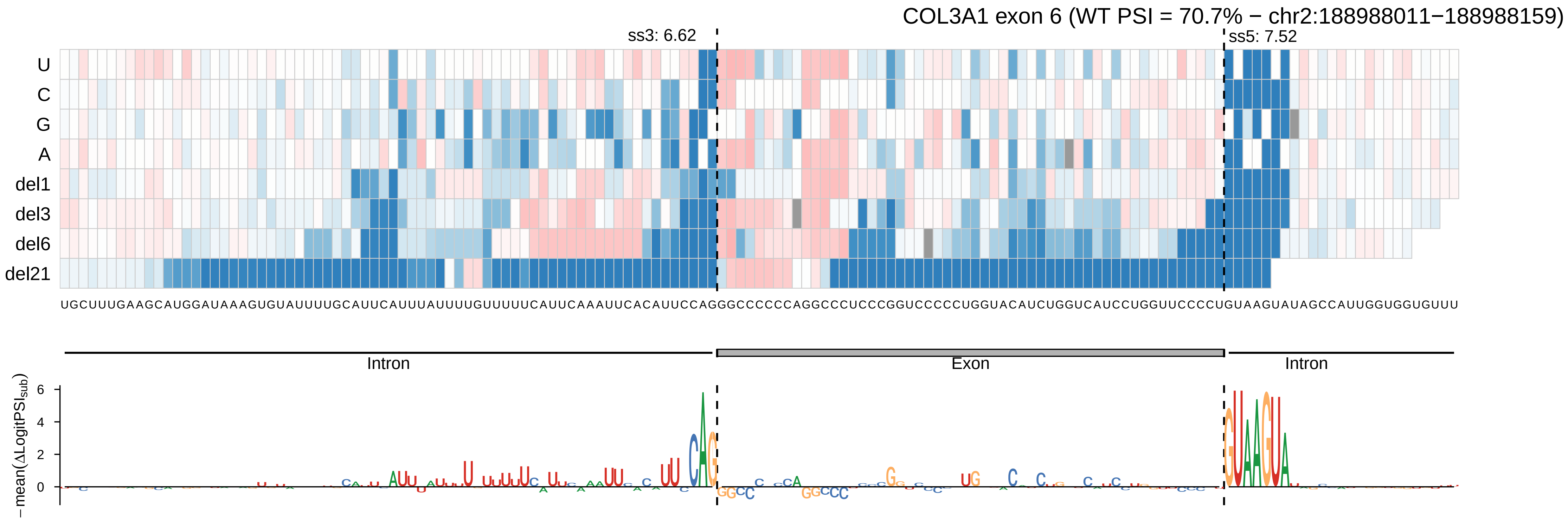
COL3A1 exon 6 (WT PSI = 70.7% − chr2:188988011−188988159)
ss3: 6.62
ss5: 7.52
U
C
G
A
del1
del3
del6
del21
UGCUUUGAAGCAUGGAUAAAGUGUAUUUUGCAUUCAUUUAUUUUGUUUUUCAUUCAAAUUCACAUUCCAGGGCCCCCCAGGCCCUCCCGGUCCCCCUGGUACAUCUGGUCAUCCUGGUUCCCCUGUAAGUAUAGCCAUUGGUGGUGUUU
Intron
Exon
Intron
−mean(ΔLogitPSI_sub)
6
4
2
0

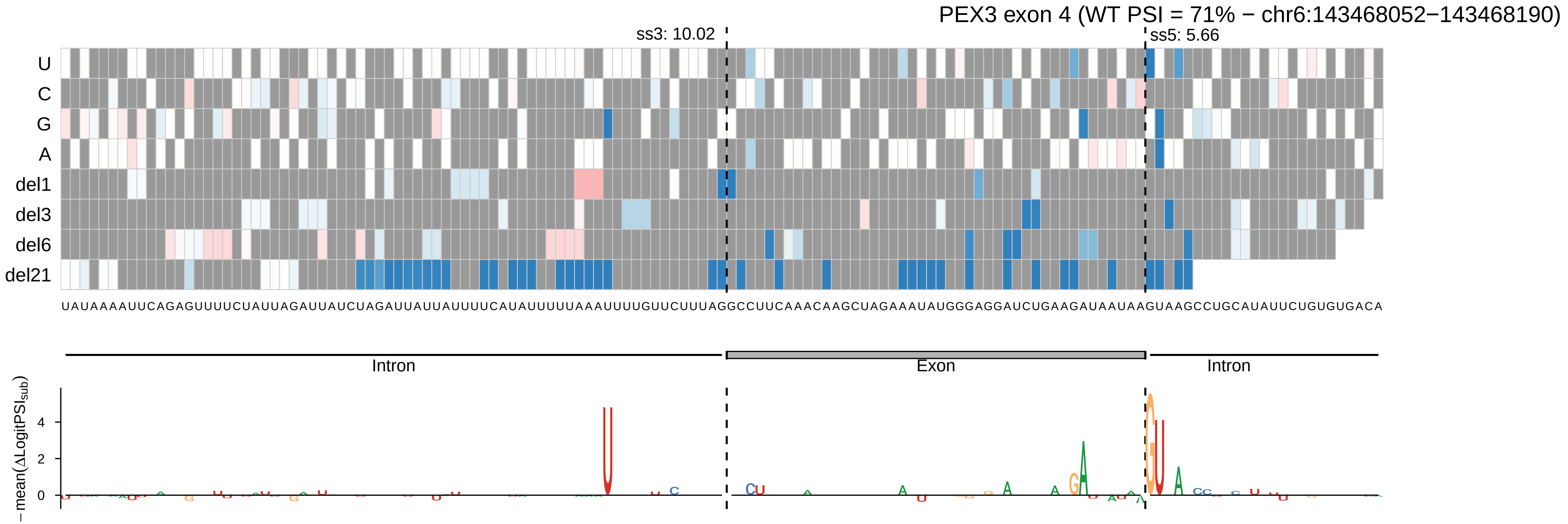

PEX3 exon 4 (WT PSI = 71% − chr6:143468052−143468190)
ss3: 10.02
ss5: 5.66
U
C
G
A
del1
del3
del6
del21
UAUAAAAUUCAGAGUUUUCUAUUAGAUUAUCUAGAUUAUUAUUUUCAUAUUUUUAAAUUUUGUUCUUUAGGCCUUCAAACAAGCUAGAAAUAUGGGAGGAUCUGAAGAUAAUAAGUAAGCCUGCAUAUUCUGUGUGACA
Intron
Exon
Intron
− mean(ΔLogitPSI_sub)
4
2
0

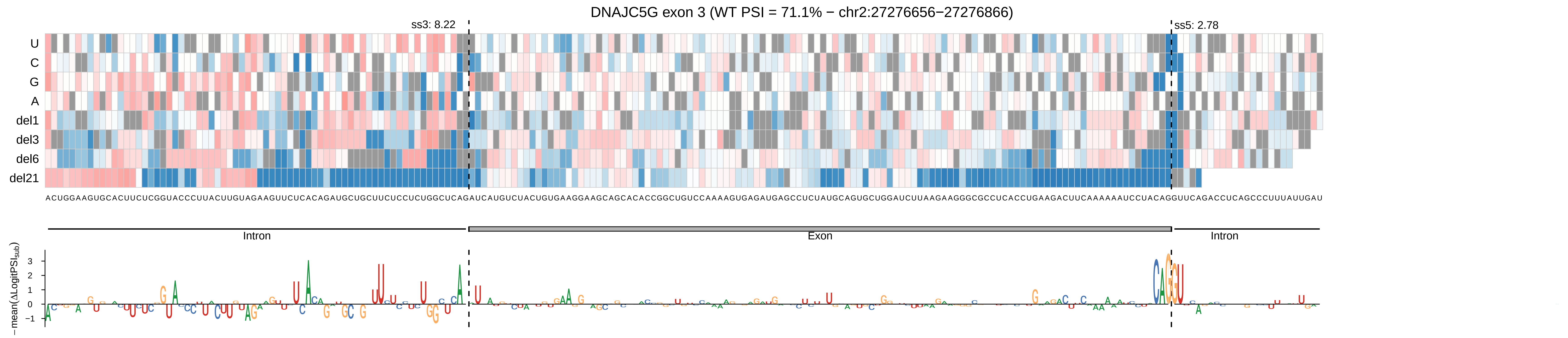
DNAJC5G exon 3 (WT PSI = 71.1% − chr2:27276656−27276866)
ss3: 8.22
ss5: 2.78
U
C
G
A
del1
del3
del6
del21
ACUGGAAGUGCACUUCUCGGUACCCUUACUUGUAGAAGUUCUCACAGAUGCUGCUUCUCCUCUGGCUCAGAUCAUGUCUACUGUGAAGGAAGCAGCACACCGGCUGUCCAAAAGUGAGAUGAGCCUCUAUGCAGUGCUGGAUCUUAAGAAGGGCGCCUCACCUGAAGACUUCAAAAAAUCCUACAGGUUCAGACCUCAGCCCUUUAUUGAU
Intron
Exon
Intron
− mean(ΔLogitPSI_sub)
3
2
1
0
−1

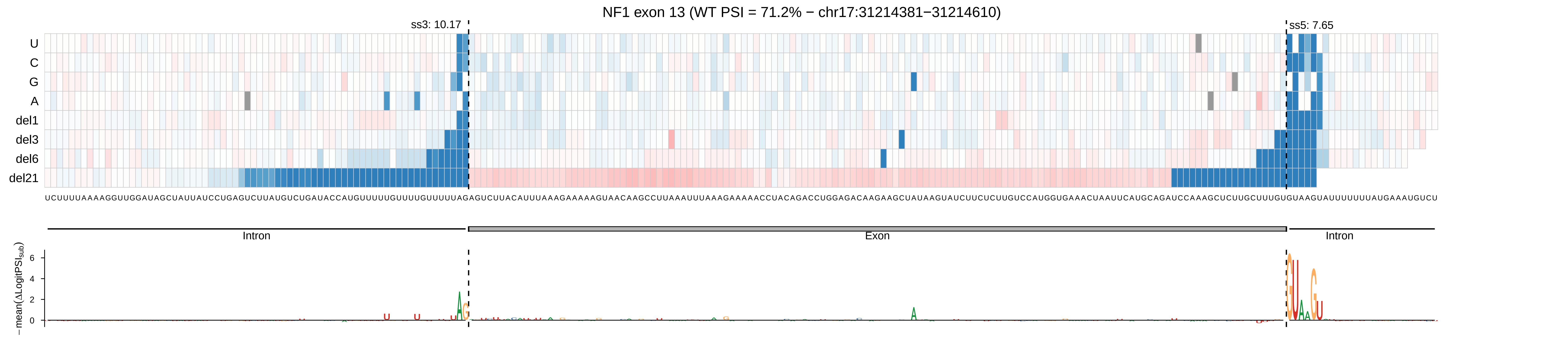

NF1 exon 13 (WT PSI = 71.2% − chr17:31214381−31214610)
ss3: 10.17
ss5: 7.65
U
C
G
A
del1
del3
del6
del21
Intron
Exon
Intron
− mean(ΔLogitPSI_sub)

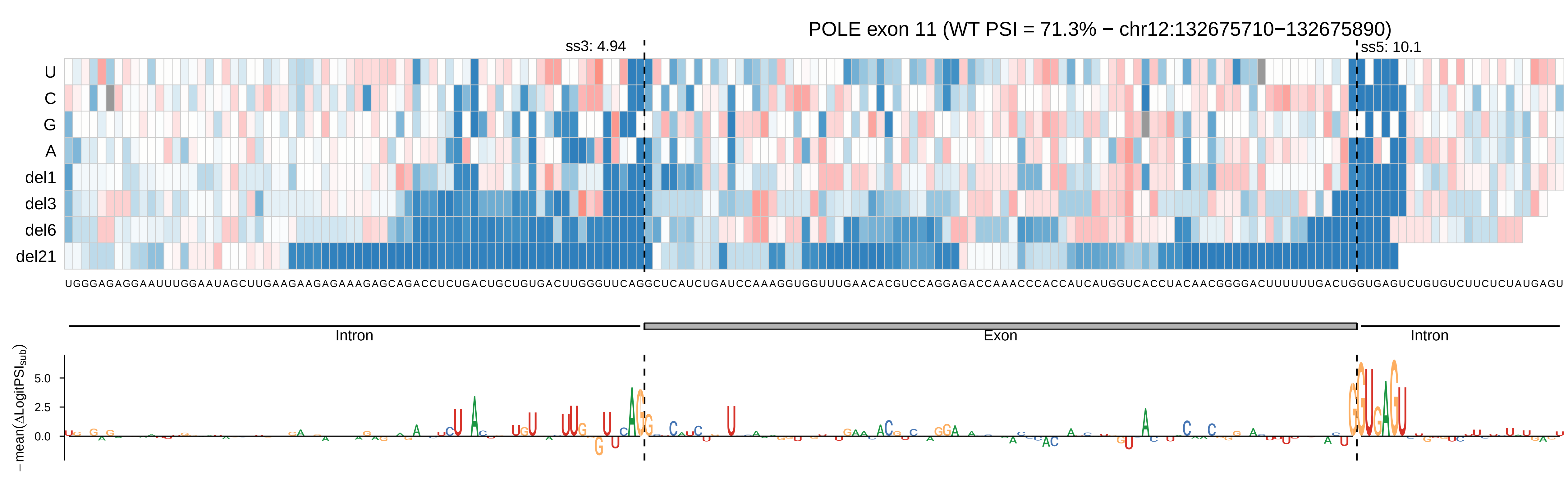

POLE exon 11 (WT PSI = 71.3% − chr12:132675710−132675890)
ss3: 4.94
ss5: 10.1
U
C
G
A
del1
del3
del6
del21
UGGGAGAGGAAUUUGGAAUAGCUUGAAGAAGAGAAAGAGCAGACCUCUGACUGCUGUGACUUGGGUUCAGGCUCAUCUGAUCCAAAGGUGGUUUGAACACGUCCAGGAGACCAAACCCACCAUCAUGGUCACCUACAACGGGGACUUUUUUGACUGGUGAGUCUGUGUCUUCUCUAUGAGU
Intron
Exon
Intron
−mean(ΔLogitPSIsub)
5.0
2.5
0.0

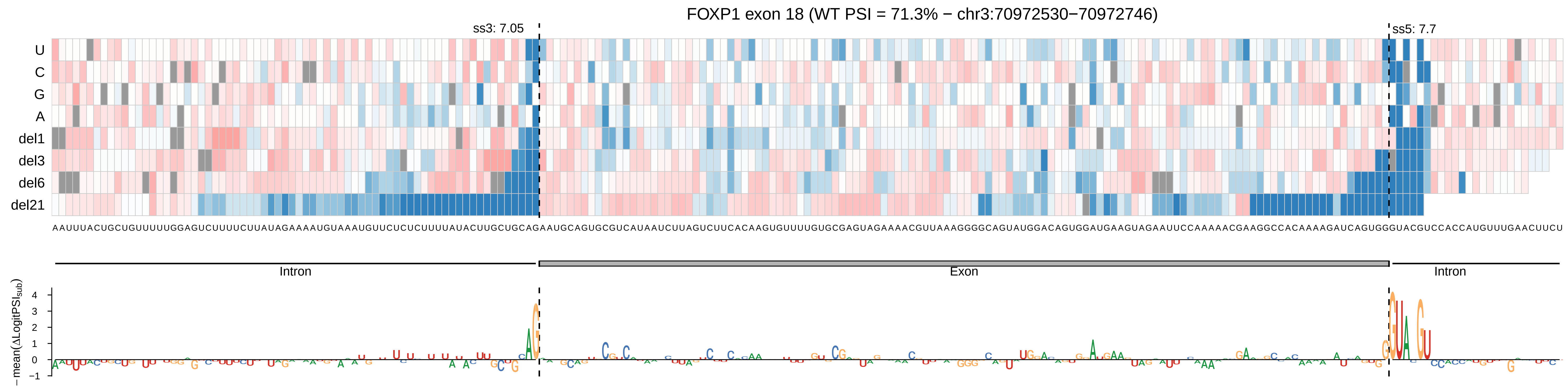
FOXP1 exon 18 (WT PSI = 71.3% − chr3:70972530−70972746)
ss3: 7.05
ss5: 7.7
U
C
G
A
del1
del3
del6
del21
AAUUUACUGCUGUUUUUGGAGUCUUUUCUUAUAGAAAAUGUAAAUGUUCUCUCUUUUAUACUUGCUGCAGAAUGCAGUGCGUCAUAAUCUUAGUCUUCACAAGUGUUUUGUGCGAGUAGAAAACGUUAAAGGGGCAGUAUGGACAGUGGAUGAAGUAGAAUUCCAAAAACGAAGGCCACAAAAGAUCAGUGGGUACGUCCACCAUGUUUGAACUUCU
Intron
Exon
Intron
− mean(ΔLogitPSI_sub)
4
3
2
1
0
−1

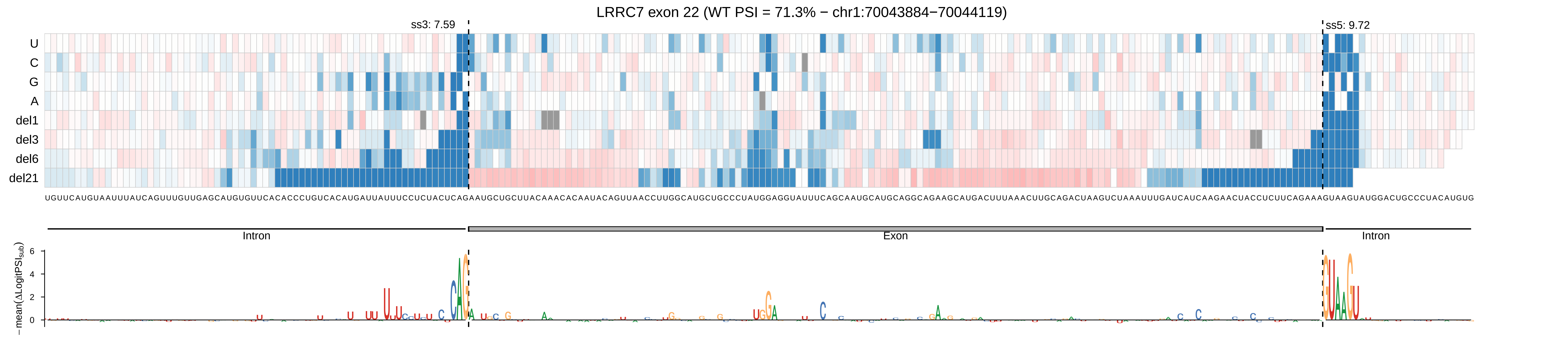
LRRC7 exon 22 (WT PSI = 71.3% − chr1:70043884−70044119)
ss3: 7.59
ss5: 9.72
U
C
G
A
del1
del3
del6
del21
UGUUCAUGUAAUUUAUCAGUUUGUUGAGCAUGUGUUCACACCCUGUCACAUGAUUAUUUCCUCUACUCAGAAUGCUGCUUACAAACACAAUACAGUUAACCUUGGCAUGCUGCCCUAUGGAGGUAUUUCAGCAAUGCAUGCAGGCAGAAGCAUGACUUUAAACUUGCAGACUAAGUCUAAAUUUGAUCAUCAAGAACUACCUCUUCAGAAAGUAAGUAUGGACUGCCCUACAUGUG
Intron
Exon
Intron
−mean(ΔLogitPSI_sub)
0
2
4
6

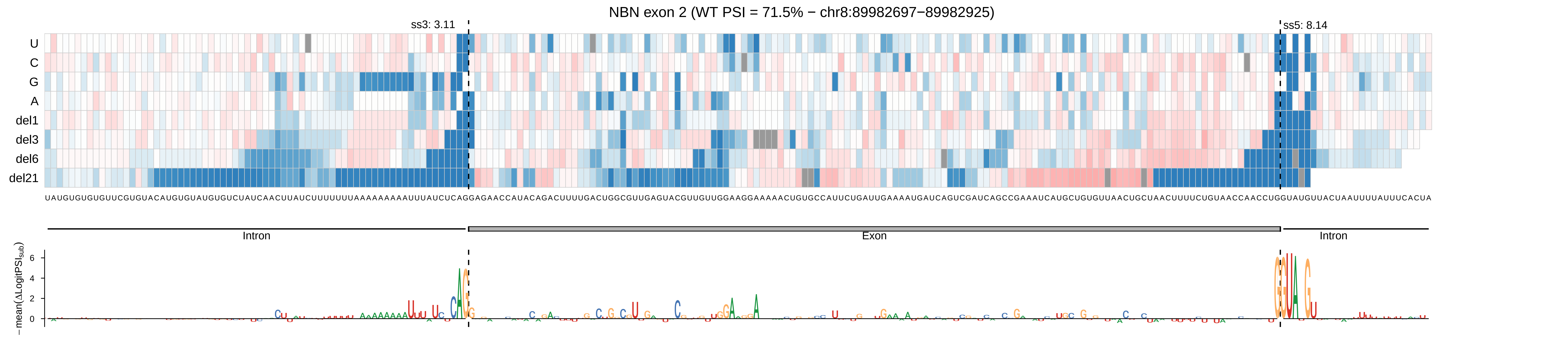

NBN exon 2 (WT PSI = 71.5% − chr8:89982697−89982925)
ss3: 3.11
ss5: 8.14
U
C
G
A
del1
del3
del6
del21
UAUGUGUGUGUUCGUGUACAUGUGUAUGUGUCUAUCAACUUAUCUUUUUUUUAAAAAAAAAUUUAUCUCAGGAGAACCAUACAGACUUUUGACUGGCGUUGAGUACGUUGUUGGAAGGAAAAACUGUGCCAUUCUGAUUGAAAAUGAUCAGUCGAUCAGCCGAAAUCAUGCUGUGUUAACUGCUAACUUUUCUGUAACCAACCUGGUAUGUUACUAAUUUUAUUUCACUA
Intron
Exon
Intron
−mean(ΔLogitPSI_sub)
6
4
2
0

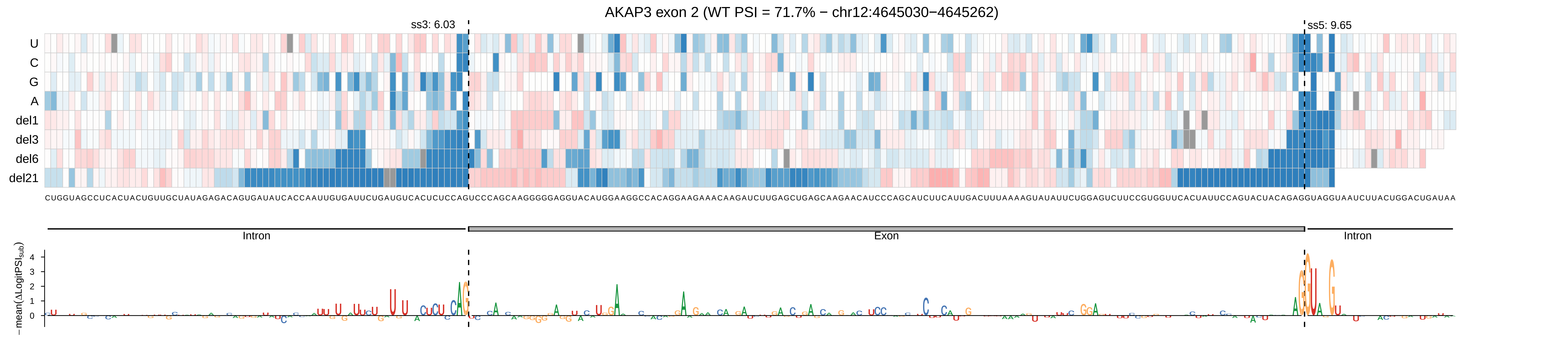

AKAP3 exon 2 (WT PSI = 71.7% − chr12:4645030−4645262)
ss3: 6.03
ss5: 9.65
U
C
G
A
del1
del3
del6
del21
CUGGUAGCCUCACUACUGUUGCUAUAGAGACAGUGAUAUCACCAAUUGUGAUUCUGAUGUCACUCUCCAGUCCCAGCAAGGGGGAGGUACAUGGAAGGCCACAGGAAGAAACAAGAUCUUGAGCUGAGCAAGAACAUCCCAGCAUCUUCAUUGACUUUAAAAGUAUAUUCUGGAGUCUUCCGUGGUUCACUAUUCCAGUACUACAGAGGUAGGUAAUCUUACUGGACUGAUAA
Intron
Exon
Intron
− mean(ΔLogitPSI_sub)
4
3
2
1
0

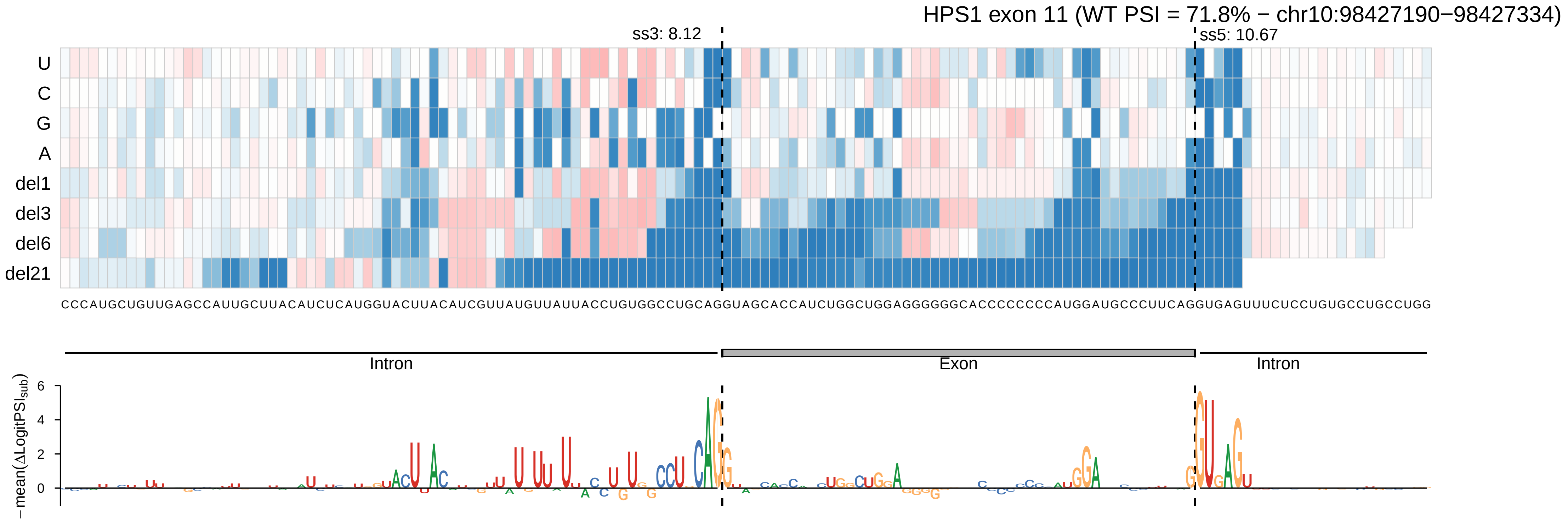

HPS1 exon 11 (WT PSI = 71.8% − chr10:98427190−98427334)
ss3: 8.12
ss5: 10.67
U
C
G
A
del1
del3
del6
del21
CCCAUGCUGUUGAGCCAUUGCUUACAUCUCAUGGUACUUACAUCGUUAUGUUAUUACCUGUGGCCUGCAGGUAGCACCAUCUGGCUGGAGGGGGGCACCCCCCCCAUGGAUGCCCUUCAGGUGAGUUUCUCCUGUGCCUGCCUGG
Intron
Exon
Intron
−mean(ΔLogitPSI_sub)
6
4
2
0

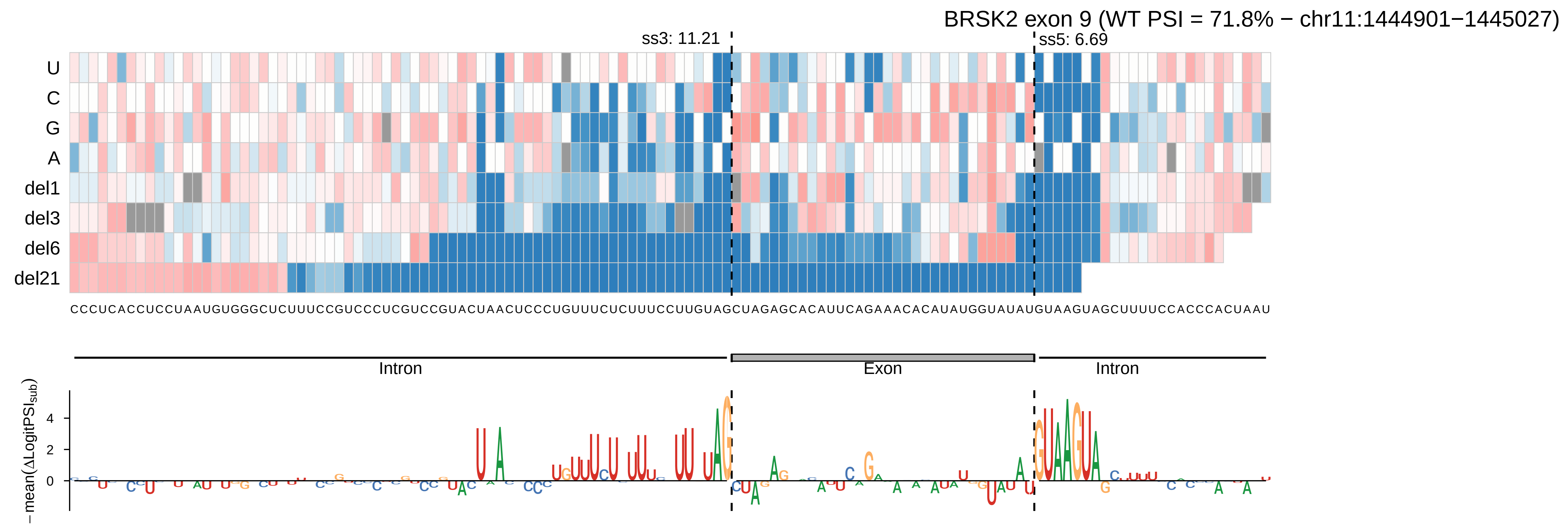
BRSK2 exon 9 (WT PSI = 71.8% − chr11:1444901−1445027)
ss3: 11.21
ss5: 6.69
U
C
G
A
del1
del3
del6
del21
Intron
Exon
Intron
−mean(ΔLogitPSI_sub)
4
2
0

BRIP1 exon 10 (WT PSI = 71.9% − chr17:61793572−61793799)

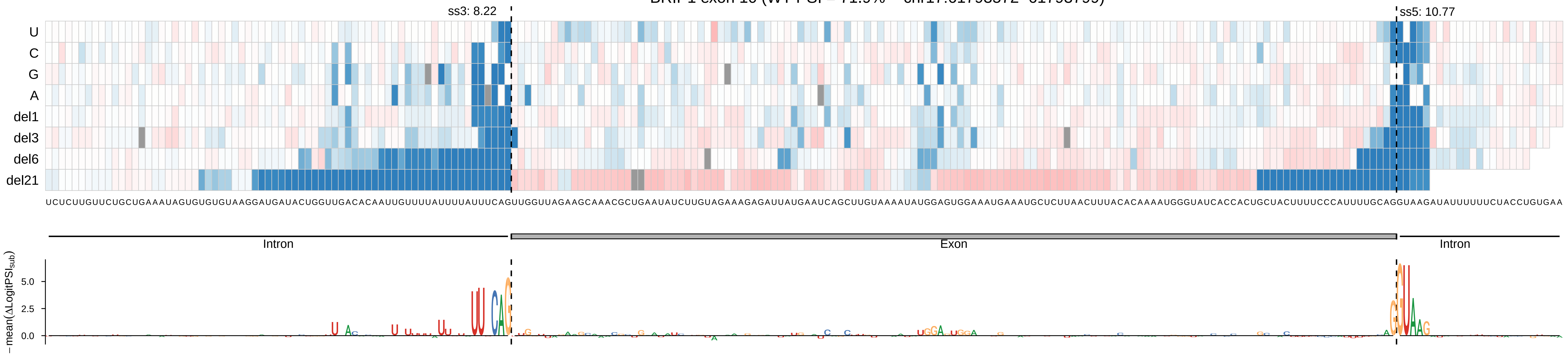

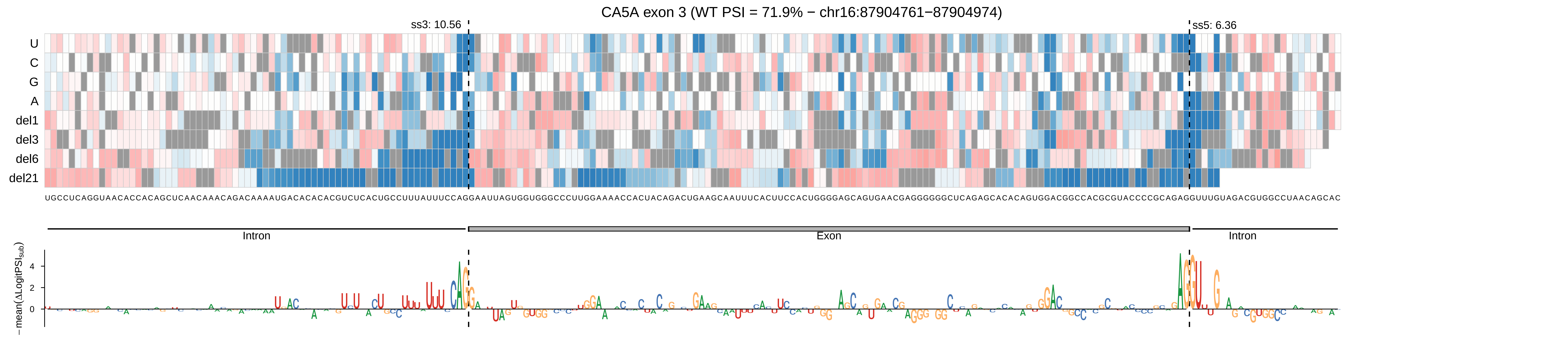

CA5A exon 3 (WT PSI = 71.9% − chr16:87904761−87904974)
ss3: 10.56
ss5: 6.36
U
C
G
A
del1
del3
del6
del21
UGCCUCAGGUAACACCACAGCUCAACAAACAGACAAAAUGACACACACGUCUCACUGCCUUUAUUUCCAGGAAUUAGUGGUGGGCCCUUGGAAAACCACUACAGACUGAAGCAAUUUCACUUCCACUGGGGAGCAGUGAACGAGGGGGGCUCAGAGCACACAGUGGACGGCCACGCGUACCCCGCAGAGGUUUGUAGACGUGGCCUAACAGCAC
Intron
Exon
Intron
−mean(ΔLogitPSI$_{sub}$)
4
2
0

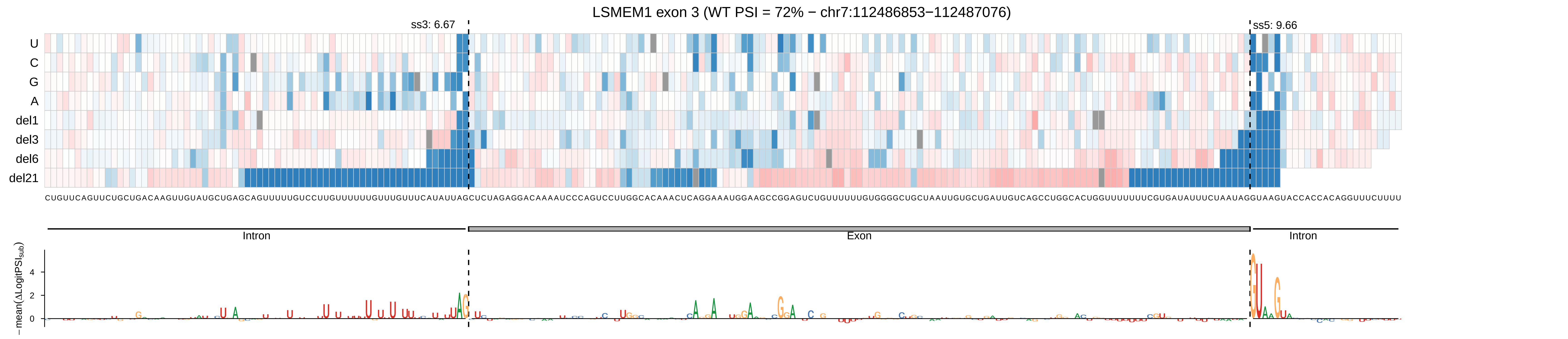
LSMEM1 exon 3 (WT PSI = 72% − chr7:112486853−112487076)
ss3: 6.67
ss5: 9.66
U
C
G
A
del1
del3
del6
del21
CUGUUCAGUUCUGCUGACAAGUUGUAUGCUGAGCAGUUUUUGUCCUUGUUUUUUGUUUGUUUCAUAUUAGCUCUAGAGGACAAAAUCCCAGUCCUUGGCACAAACUCAGGAAAUGGAAGCCGGAGUCUGUUUUUUGUGGGGCUGCUAAUUGUGCUGAUUGUCAGCCUGGCACUGGUUUUUUUCGUGAUAUUUCUAAUAGGUAAGUACCACCACAGGUUUCUUUU
Intron
Exon
Intron
− mean(ΔLogitPSI_sub)
4
2
0

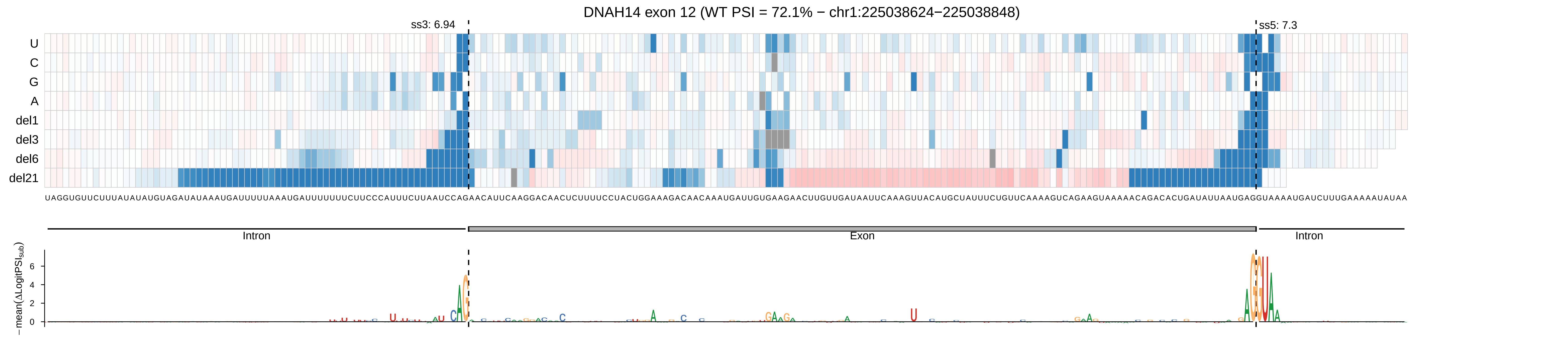
DNAH14 exon 12 (WT PSI = 72.1% − chr1:225038624−225038848)
ss3: 6.94
ss5: 7.3
U
C
G
A
del1
del3
del6
del21
UAGGUGUUCUUUAUAUAUGUAGAUAUAAAUGAUUUUUAAAUGAUUUUUUUCUUCCCAUUUCUUAAUCCAGAACAUUCAAGGACAACUCUUUUCCUACUGGAAAGACAACAAAUGAUUGUGAAGAACUUGUUGAUAAUUCAAAGUUACAUGCUAUUUCUGUUCAAAAGUCAGAAGUAAAAACAGACACUGAUAUUAAUGAGGUAAAAUGAUCUUUGAAAAAUAUAA
Intron
Exon
Intron
−mean(ΔLogitPSI$_{sub}$)
6
4
2
0

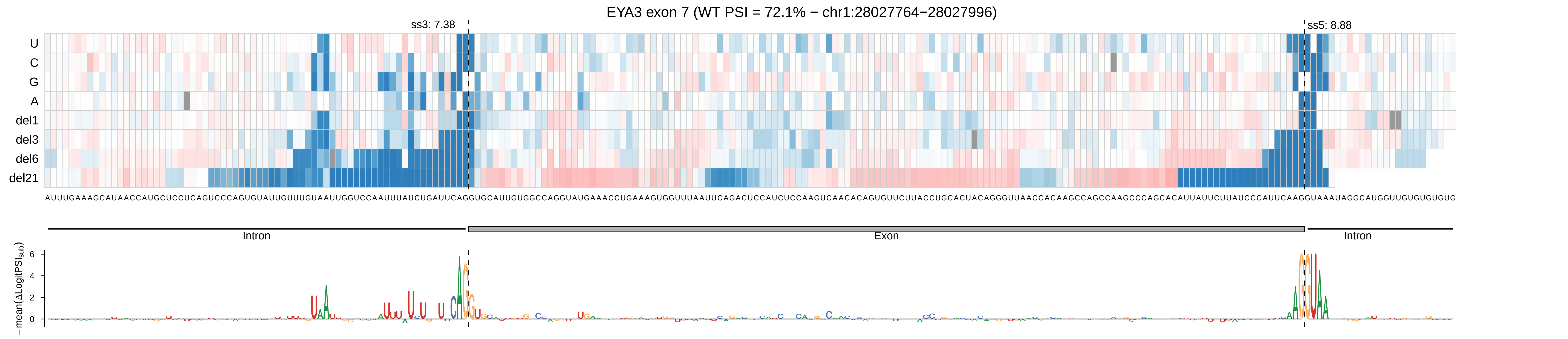
EYA3 exon 7 (WT PSI = 72.1% − chr1:28027764−28027996)
ss3: 7.38
ss5: 8.88
U
C
G
A
del1
del3
del6
del21
Intron
Exon
Intron
−mean(ΔLogitPSI_sub)
6
4
2
0

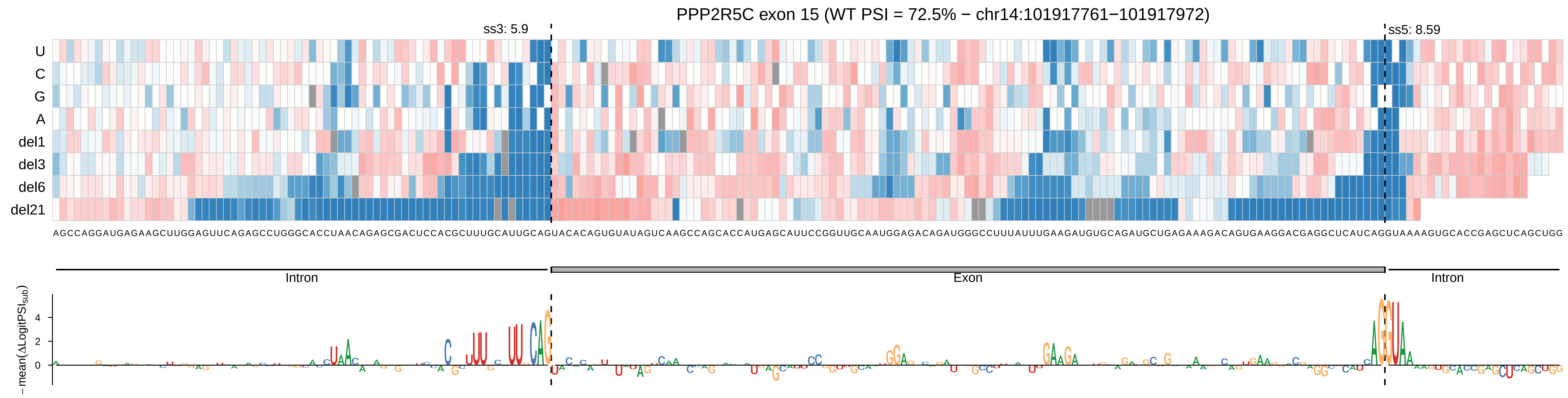

PPP2R5C exon 15 (WT PSI = 72.5% − chr14:101917761−101917972)
ss3: 5.9
ss5: 8.59
U
C
G
A
del1
del3
del6
del21
AGCCAGGAUGAGAAGCUUGGAGUUCAGAGCCUGGGCACCUAACAGAGCGACUCCACGCUUUGCAUUGCAGUACACAGUGUAUAGUCAAGCCAGCACCAUGAGCAUUCCGGUUGCAAUGGAGACAGAUGGGCCUUUAUUUGAAGAUGUGCAGAUGCUGAGAAAGACAGUGAAGGACGAGGCUCAUCAGGUAAAAGUGCACCGAGCUCAGCUGG
Intron
Exon
Intron
−mean(ΔLogitPSI_sub)
4
2
0

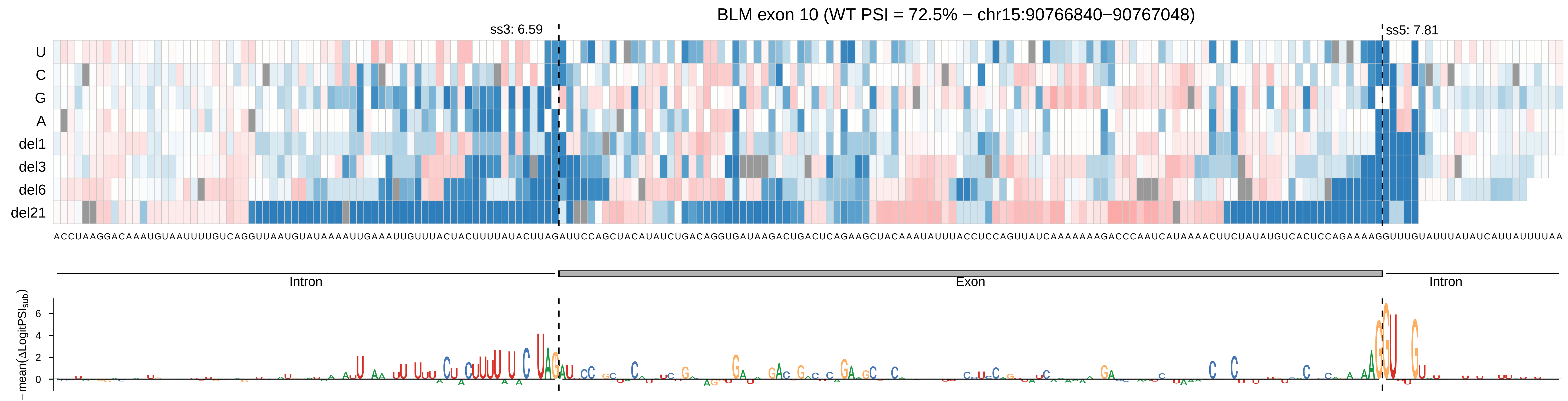
BLM exon 10 (WT PSI = 72.5% − chr15:90766840−90767048)
ss3: 6.59
ss5: 7.81
U
C
G
A
del1
del3
del6
del21
Intron
Exon
Intron
−mean(ΔLogitPSI_sub)
0
2
4
6

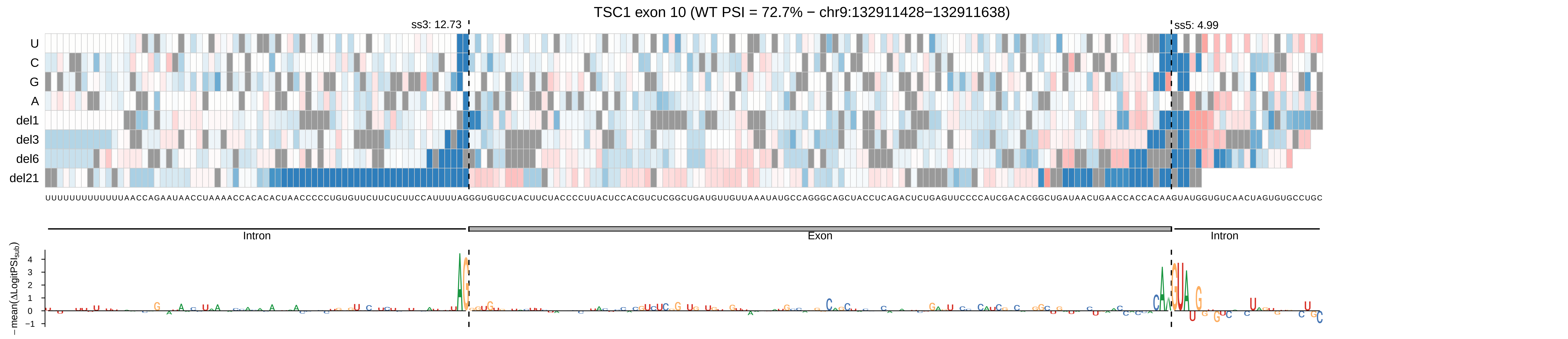
TSC1 exon 10 (WT PSI = 72.7% − chr9:132911428−132911638)
ss3: 12.73
ss5: 4.99
U
C
G
A
del1
del3
del6
del21
Intron
Exon
Intron
− mean(ΔLogitPSI_sub)
4
3
2
1
0
−1

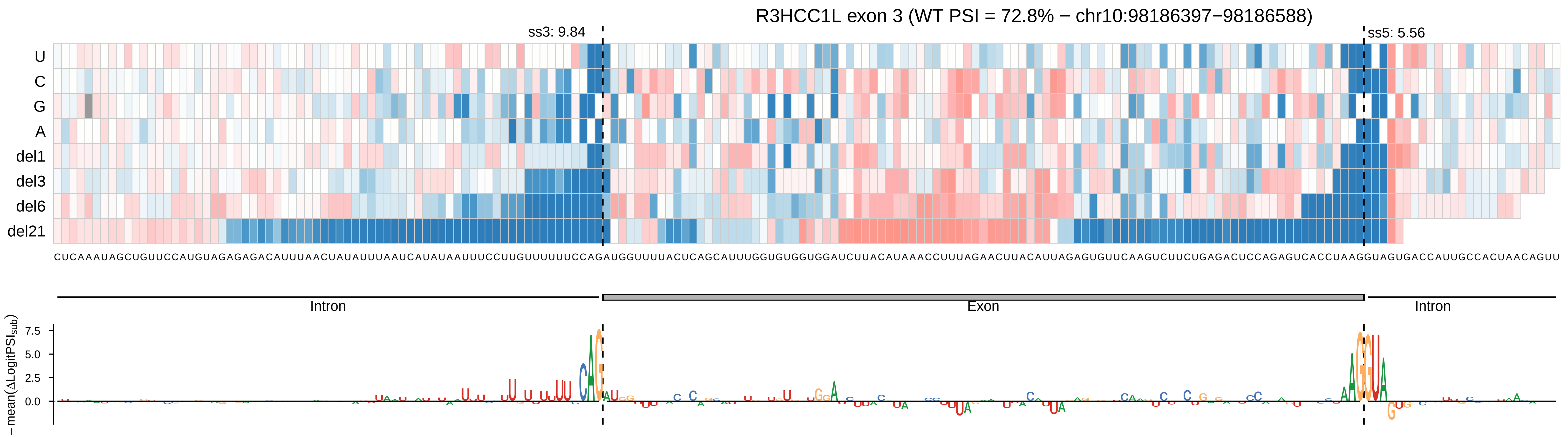

R3HCC1L exon 3 (WT PSI = 72.8% − chr10:98186397−98186588)
ss3: 9.84
ss5: 5.56
U
C
G
A
del1
del3
del6
del21
CUCAAAUAGCUGUUCCAUGUAGAGAGACAUUUAACUAUAUUUAAUCAUAUAAUUUCCUUGUUUUUUCCAGAUGGUUUUACUCAGCAUUUGGUGUGGUGGAUCUUACAUAAACCUUUAGAACUUACAUUAGAGUGUUCAAGUCUUCUGAGACUCCAGAGUCACCUAAGGUAGUGACCAUUGCCACUAACAGUU
Intron
Exon
Intron
−mean(ΔLogitPSI_sub)
7.5
5.0
2.5
0.0

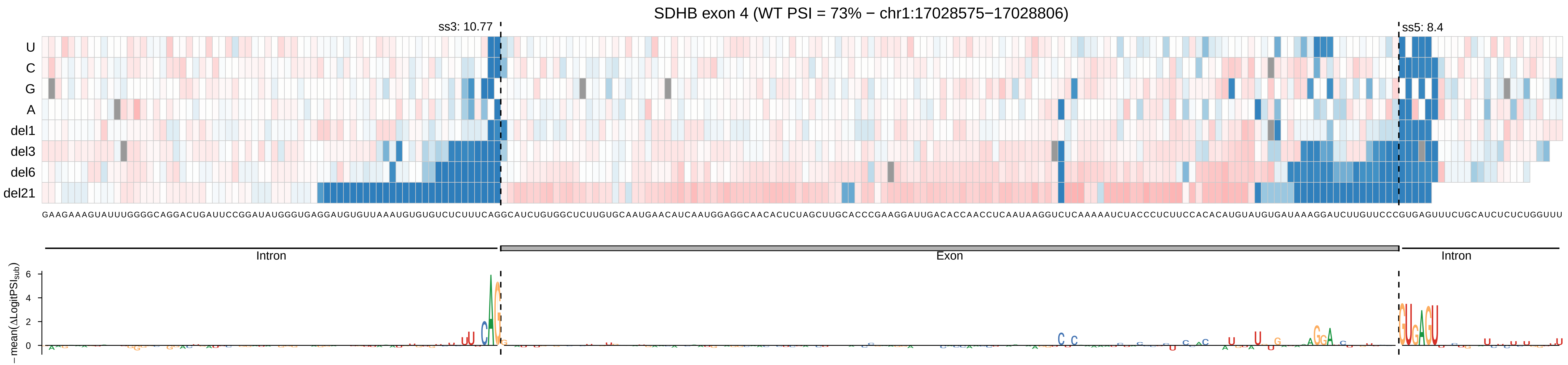
SDHB exon 4 (WT PSI = 73% − chr1:17028575−17028806)
ss3: 10.77
ss5: 8.4
U
C
G
A
del1
del3
del6
del21
GAAGAAAGUAUUUGGGGCAGGACUGAUUCCGGAUAUGGGUGAGGAUGUGUUAAAUGUGUGUCUCUUUCAGGCAUCUGUGGCUCUUGUGCAAUGAACAUCAAUGGAGGCAACACUCUAGCUUGCACCCGAAGGAUUGACACCAACCUCAAUAAGGUCUCAAAAAUCUACCCUCUUCCACACAUGUAUGUGAUAAAGGAUCUUGUUCCCGUGAGUUUCUGCAUCUCUCUGGUUU
Intron
Exon
Intron
−mean(ΔLogitPSI_sub)
6
4
2
0

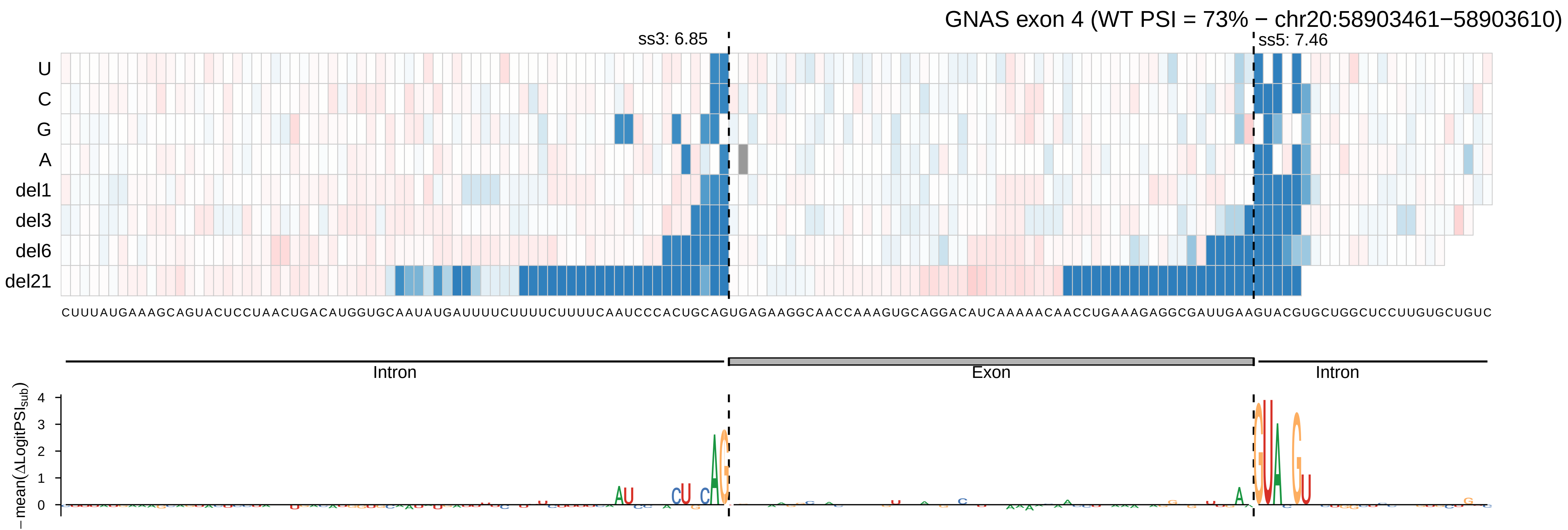
GNAS exon 4 (WT PSI = 73% − chr20:58903461−58903610)
ss3: 6.85
ss5: 7.46
U
C
G
A
del1
del3
del6
del21
CUUUAUGAAAGCAGUACUCCUAACUGACAUGGUGCAAUAUGAUUUUCUUUUCUUUUCAAUCCCACUGCAGUGAGAAGGCAACCAAAGUGCAGGACAUCAAAAACAACCUGAAAGAGGCGAUUGAAGUACGUGCUGGCUCCUUGUGCUGUC
Intron
Exon
Intron
−mean(ΔLogitPSI_sub)

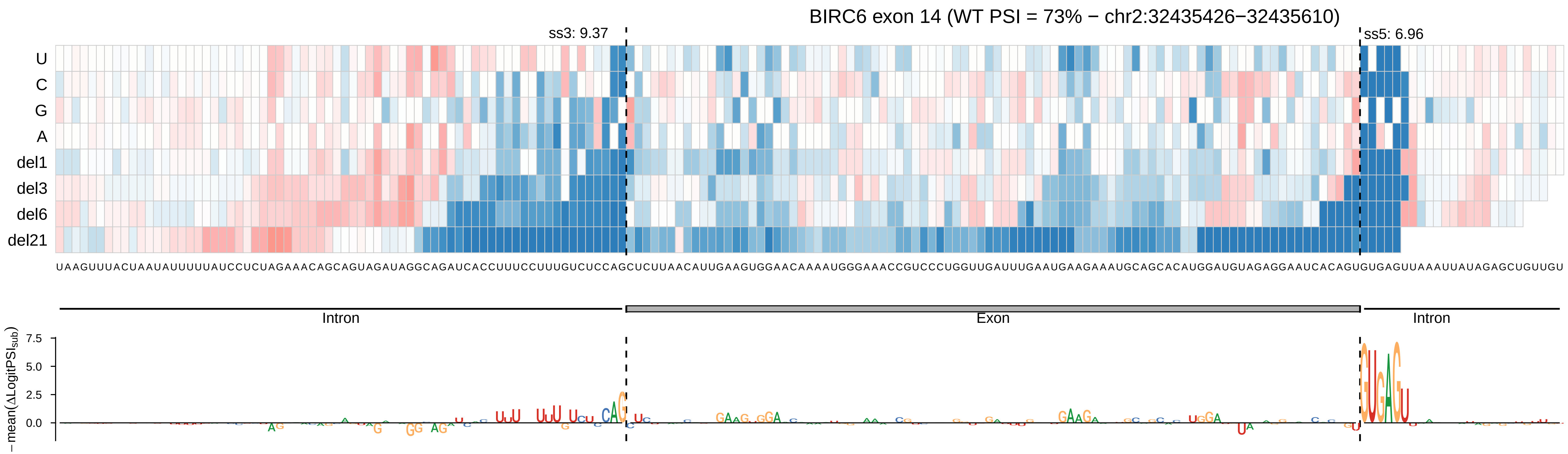

BIRC6 exon 14 (WT PSI = 73% − chr2:32435426−32435610)
ss3: 9.37
ss5: 6.96
U
C
G
A
del1
del3
del6
del21
UAAGUUUACUAAUAUUUUUAUCCUCUAGAAACAGCAGUAGAUAGGCAGAUCACCUUUCCUUUGUCUCCAGCUCUUAACAUUGAAGUGGAACAAAAUGGGAAACCGUCCCUGGUUGAUUUGAAUGAAGAAAUGCAGCACAUGGAUGUAGAGGAAUCACAGUGUGAGUUAAAUUAUAGAGCUGUUGU
Intron
Exon
Intron
−mean(ΔLogitPSI_sub)
7.5
5.0
2.5
0.0

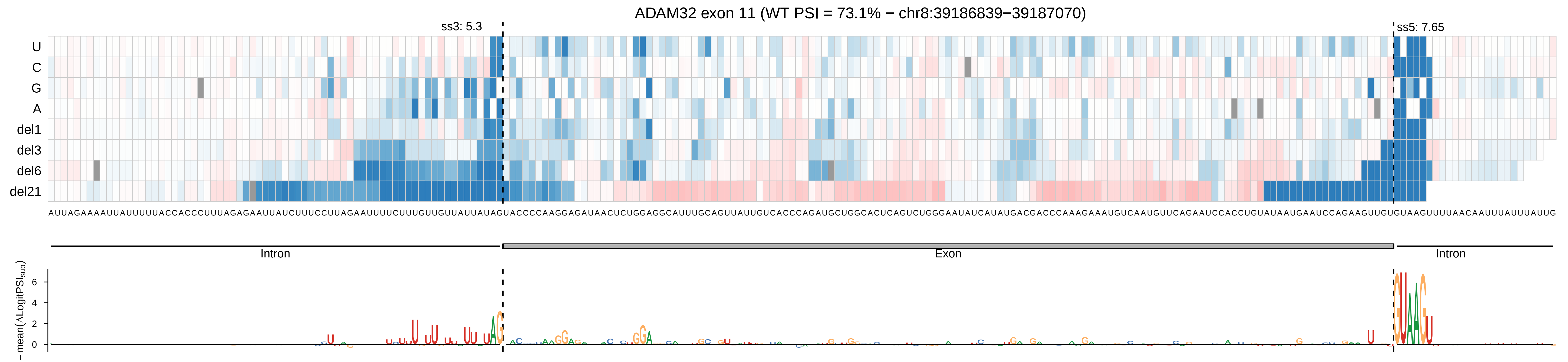

ADAM32 exon 11 (WT PSI = 73.1% − chr8:39186839−39187070)
ss3: 5.3
ss5: 7.65
U
C
G
A
del1
del3
del6
del21
AUUAGAAAAUUAUUUUUACCACCCUUUAGAGAAUUAUCUUUCCUUAGAAUUUUCUUUGUUGUUAUUAUAGUACCCCAAGGAGAUAACUCUGGAGGCAUUUGCAGUUAUUGUCACCCAGAUGCUGGCACUCAGUCUGGGAAUAUCAUAUGACGACCCAAAGAAAUGUCAAUGUUCAGAAUCCACCUGUAUAAUGAAUCCAGAAGUUGUGUAAGUUUUAACAAUUUAUUUAUUG
Intron
Exon
Intron
− mean(ΔLogitPSI_sub)
6
4
2
0

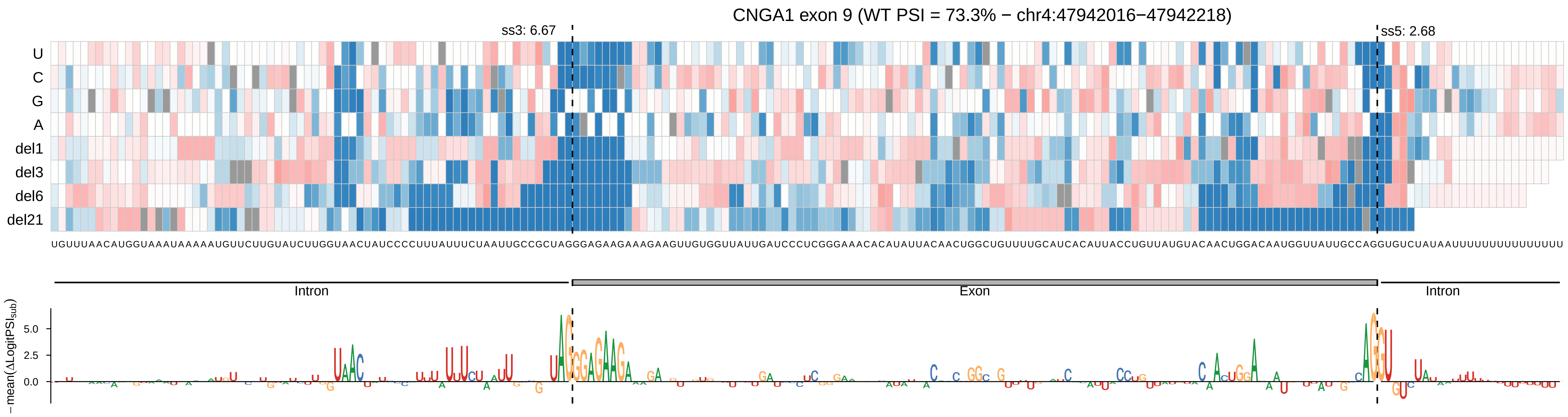

CNGA1 exon 9 (WT PSI = 73.3% − chr4:47942016−47942218)
ss3: 6.67
ss5: 2.68
U
C
G
A
del1
del3
del6
del21
UGUUUAACAUGGUAAAUAAAAAUGUUCUUGUAUCUUGGUAACUAUCCCCUUUAUUUCUAAUUGCCGCUAGGGAGAAGAAAGAAGUUGUGGUUAUUGAUCCCUCGGGAAACACAUAUUACAACUGGCUGUUUUGCAUCACAUUACCUGUUAUGUACAACUGGACAAUGGUUAUUGCCAGGUGUCUAUAAUUUUUUUUUUUUUU
Intron
Exon
Intron
−mean(ΔLogitPSI_sub)
5.0
2.5
0.0

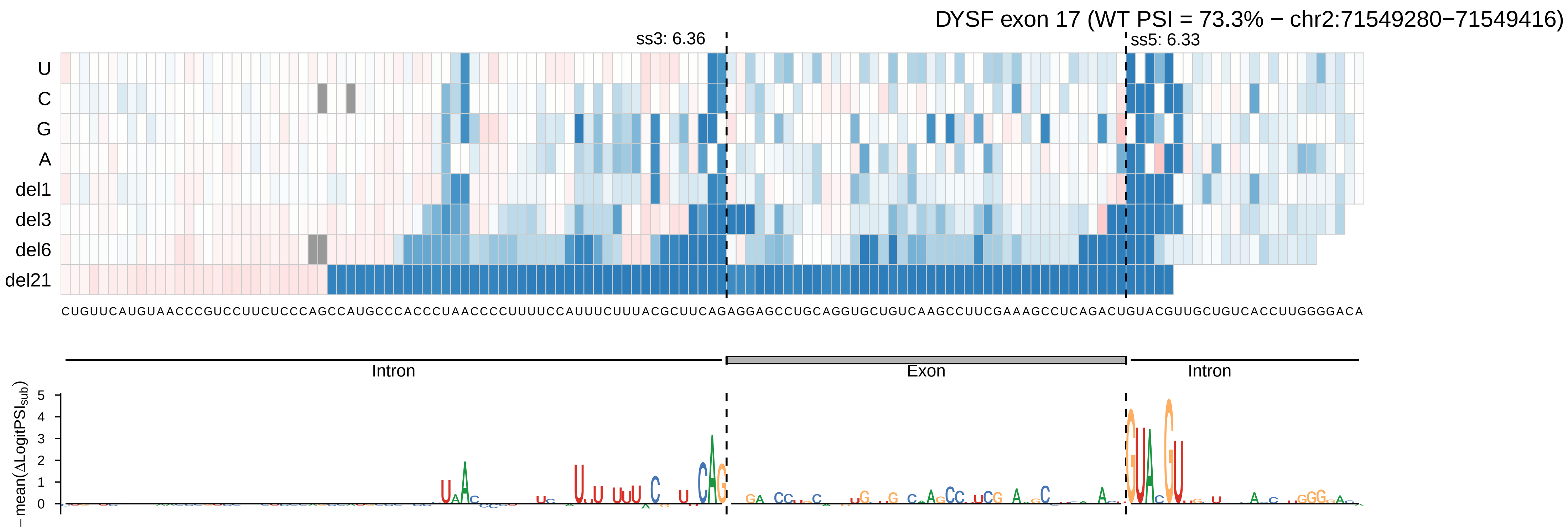

DYSF exon 17 (WT PSI = 73.3% − chr2:71549280−71549416)
ss3: 6.36
ss5: 6.33
U
C
G
A
del1
del3
del6
del21
CUGUUCAUGUAACCCGUCCUUCUCCCAGCCAUGCCCACCCUAACCCCUUUUCCAUUUCUUUACGCUUCAGAGGAGCCUGCAGGUGCUGUCAAGCCUUCGAAAGCCUCAGACUGUACGUUGCUGUCACCUUGGGGACA
Intron
Exon
Intron
− mean(ΔLogitPSI_sub)

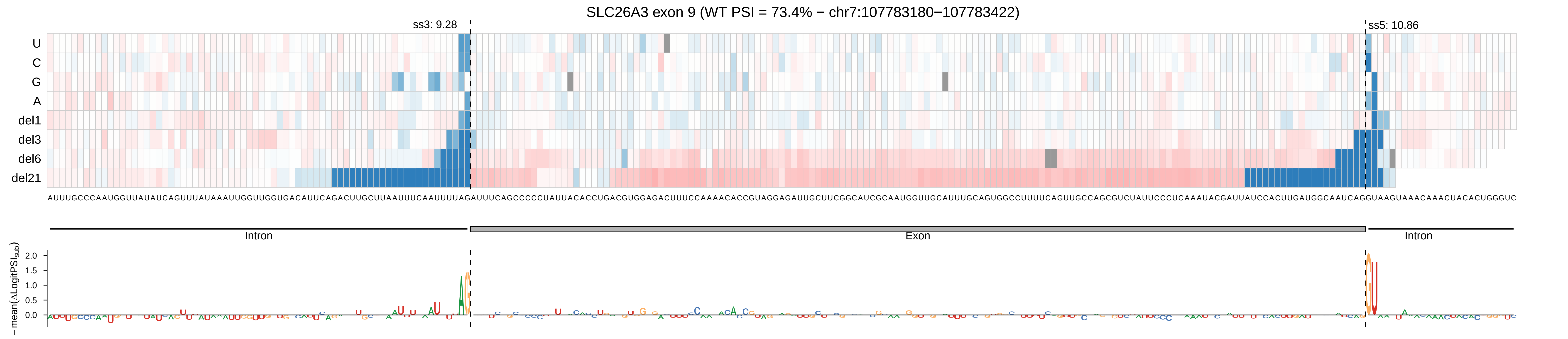

SLC26A3 exon 9 (WT PSI = 73.4% − chr7:107783180−107783422)
ss3: 9.28
ss5: 10.86
U
C
G
A
del1
del3
del6
del21
Intron
Exon
Intron
−mean(ΔLogitPSI_sub)
2.0
1.5
1.0
0.5
0.0

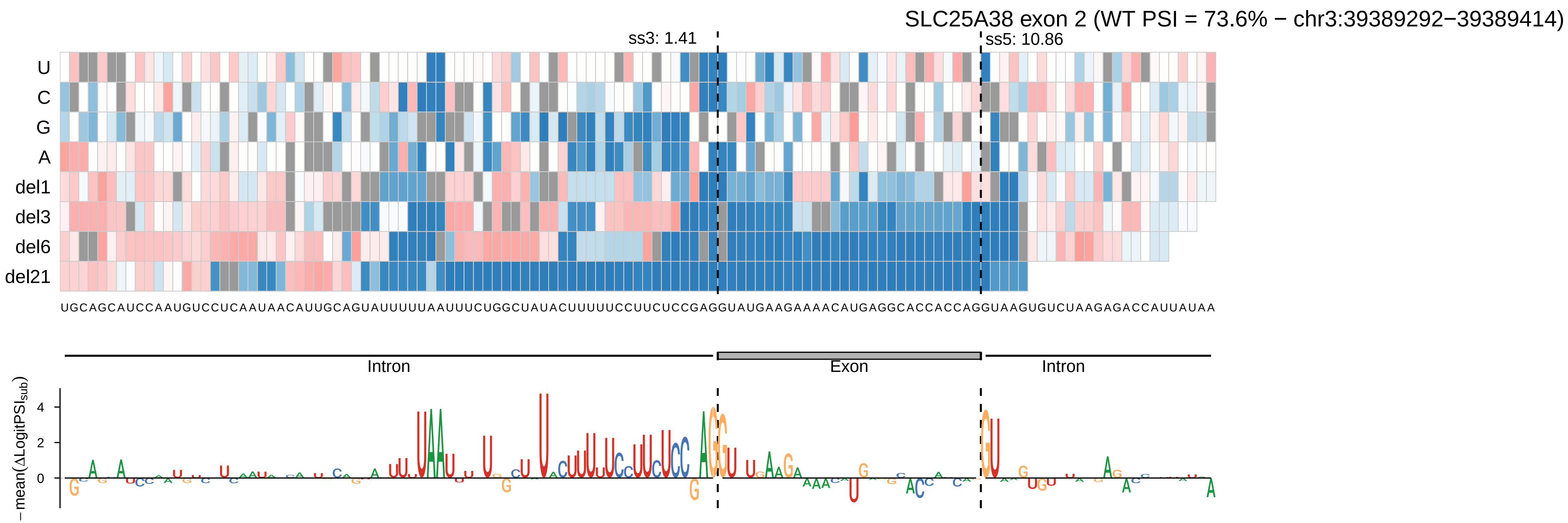

SLC25A38 exon 2 (WT PSI = 73.6% − chr3:39389292−39389414)
ss3: 1.41
ss5: 10.86
U
C
G
A
del1
del3
del6
del21
UGCAGCAUCCAAUGUCCUCAAUAACAUUGCAGUAUUUUUAAUUUCUGGCUAUACUUUUUCCUUCUCCGAGGUAUGAAGAAAACAUGAGGCACCACCAGGUAAGUGUCUAAGAGACCAUUAUAA
Intron
Exon
Intron
−mean(ΔLogitPSI_sub)
4
2
0

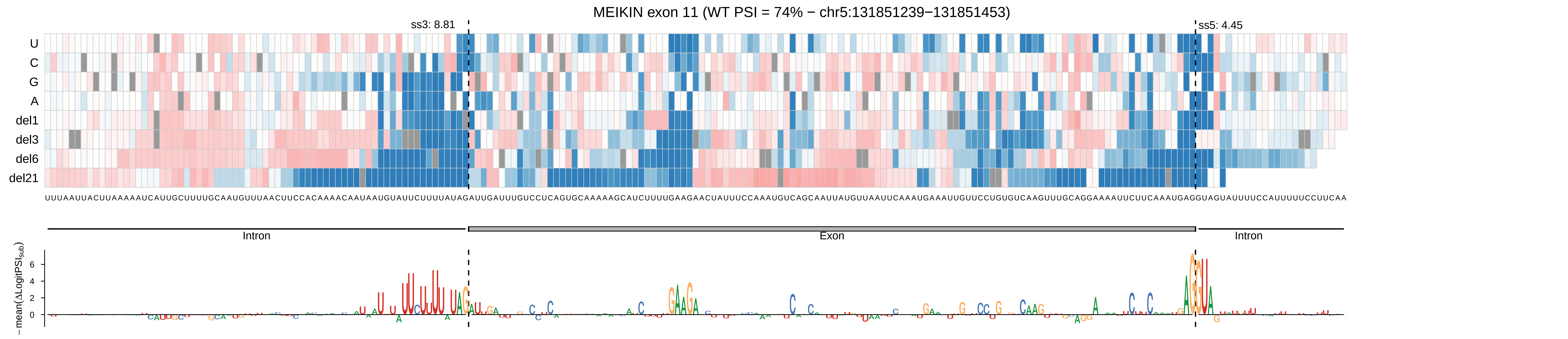

MEIKIN exon 11 (WT PSI = 74% − chr5:131851239−131851453)
ss3: 8.81
ss5: 4.45
U
C
G
A
del1
del3
del6
del21
UUUAAUUACUUAAAAAUCAUUGCUUUUGCAAUGUUUAACUUCCACAAAACAAUAAUGUAUUCUUUUAUAGAUUGAUUUGUCCUCAGUGCAAAAAGCAUCUUUUGAAGAACUAUUUCCAAAUGUCAGCAAUUAUGUUAAUUCAAAUGAAAUUGUUCCUGUGUCAAGUUUGCAGGAAAAUUCUUCAAAUGAGGUAGUAUUUUCCAUUUUUCCUUCAA
Intron
Exon
Intron
−mean(ΔLogitPSI_sub)
6
4
2
0

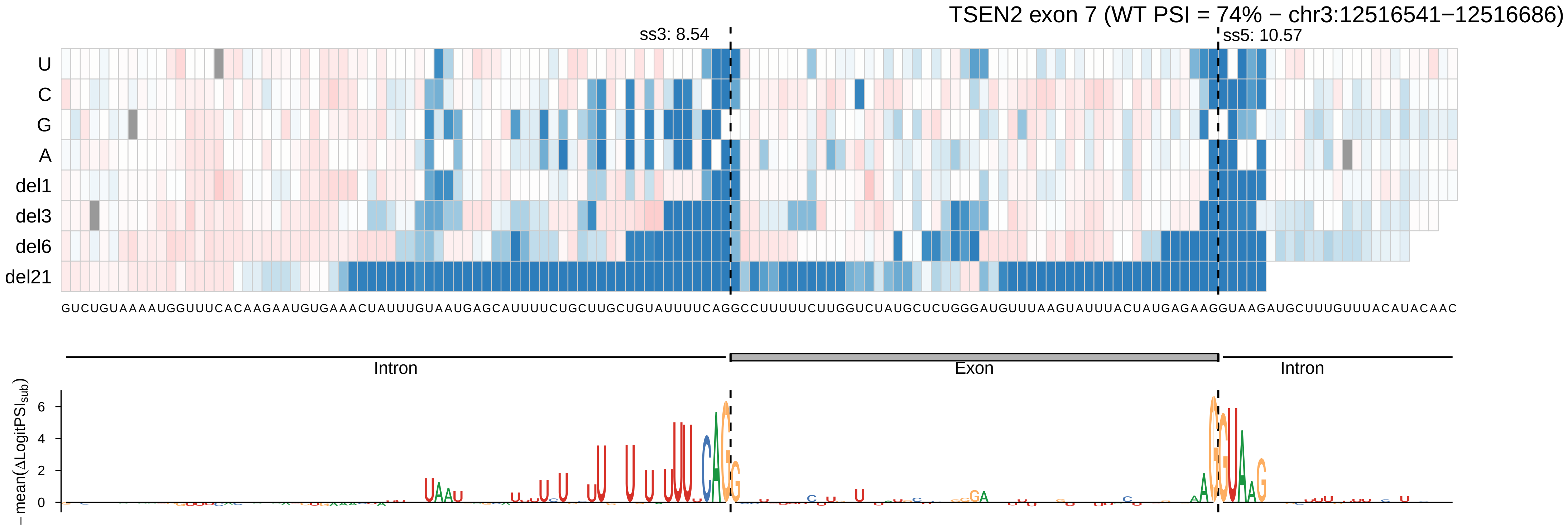

TSEN2 exon 7 (WT PSI = 74% − chr3:12516541−12516686)
ss3: 8.54
ss5: 10.57
U
C
G
A
del1
del3
del6
del21
GUCUGUAAAAUGGUUUCACAAGAAUGUGAAACUAUUUGUAAUGAGCAUUUUCUGCUUGCUGUAUUUUCAGGCCUUUUUCUUGGUCUAUGCUCUGGGAUGUUUAAGUAUUUACUAUGAGAAGGUAAGAUGCUUUGUUUACAUACAAC
Intron
Exon
Intron
6
4
2
0
−mean(ΔLogitPSI sub)

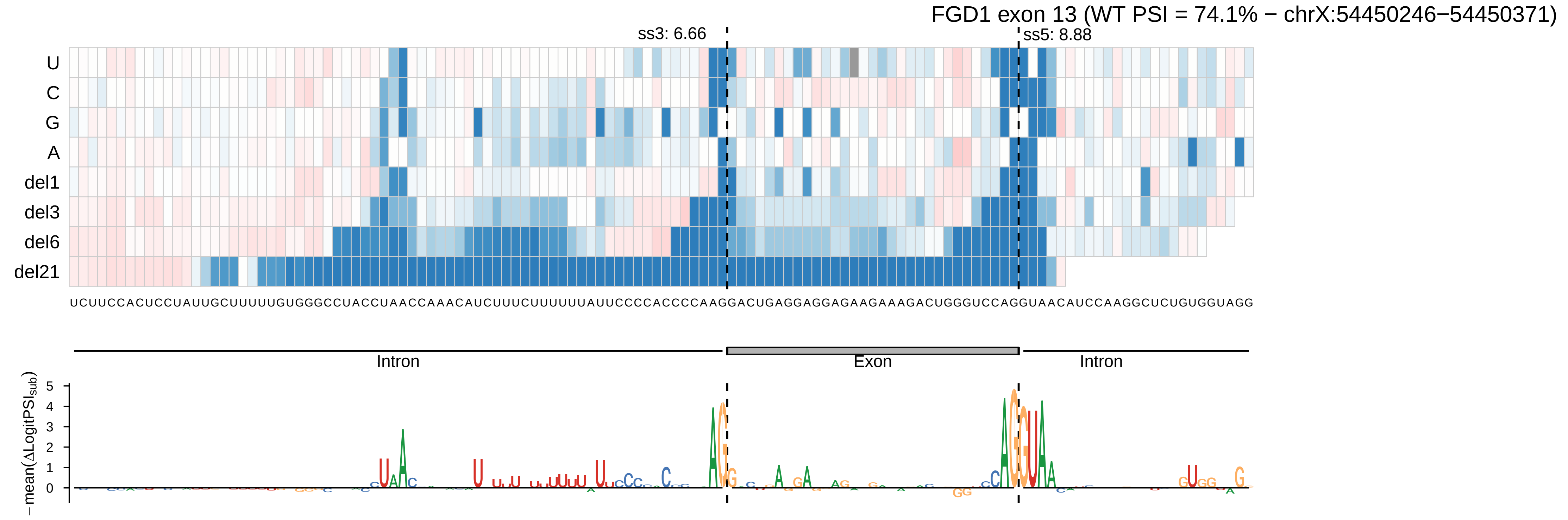
FGD1 exon 13 (WT PSI = 74.1% − chrX:54450246−54450371)
ss3: 6.66
ss5: 8.88
U
C
G
A
del1
del3
del6
del21
UCUUCCACUCCUAUUGCUUUUUGUGGGCCUACCUAACCAAACAUCUUUCUUUUUUAUUCCCCACCCCAAGGACUGAGGAGGAGAAGAAAGACUGGGUCCAGGUAACAUCCAAGGCUCUGUGGUAGG
Intron
Exon
Intron
−mean(ΔLogitPSI_sub)
5
4
3
2
1
0

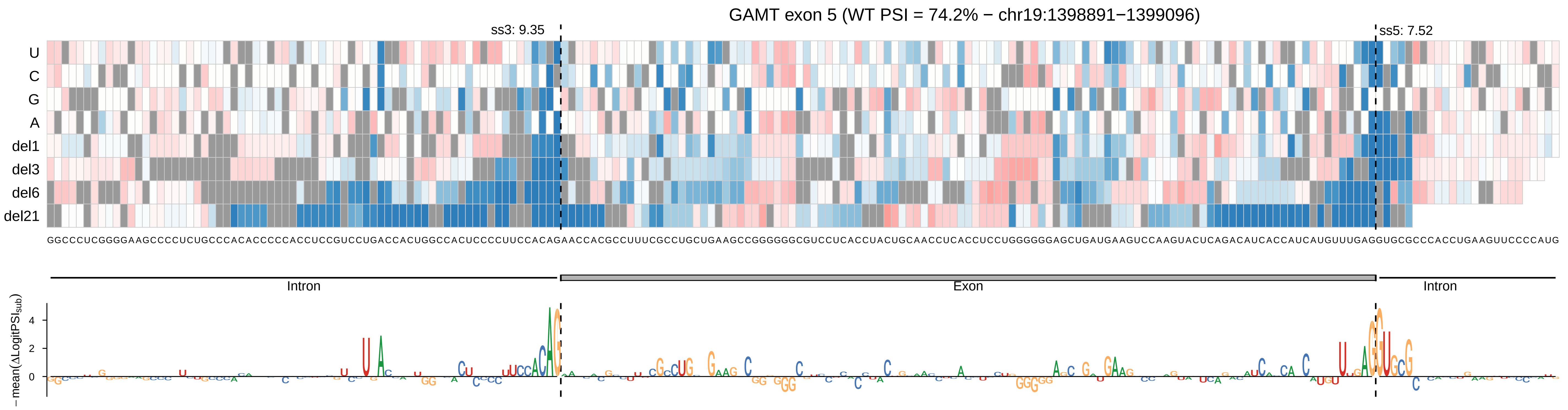
GAMT exon 5 (WT PSI = 74.2% − chr19:1398891−1399096)
ss3: 9.35
ss5: 7.52
U
C
G
A
del1
del3
del6
del21
GGCCCUCGGGGAAGCCCCUCUGCCCACACCCCCACCUCCGUCCUGACCACUGGCCACUCCCCUUCCACAGAACCACGCCUUUCGCCUGCUGAAGCCGGGGGGCGUCCUCACCUACUGCAACCUCACCUCCUGGGGGGAGCUGAUGAAGUCCAAGUACUCAGACAUCACCAUCAUGUUUGAGGUGCGCCCACCUGAAGUUCCCCAUG
Intron
Exon
Intron
−mean(ΔLogitPSIsub)
4
2
0

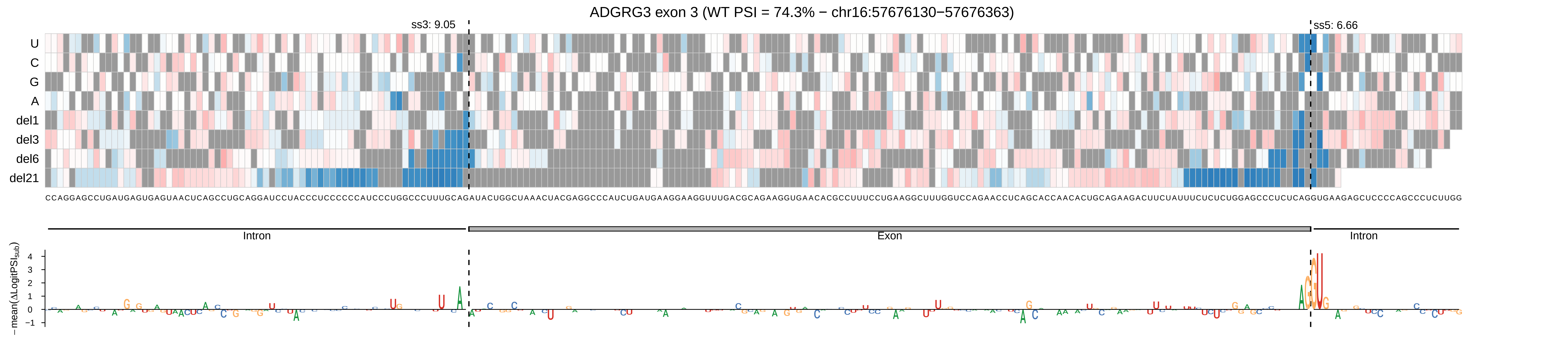

ADGRG3 exon 3 (WT PSI = 74.3% − chr16:57676130−57676363)
ss3: 9.05
ss5: 6.66
U
C
G
A
del1
del3
del6
del21
Intron
Exon
Intron
−mean(ΔLogitPSI$_{sub}$)
4
3
2
1
0
−1

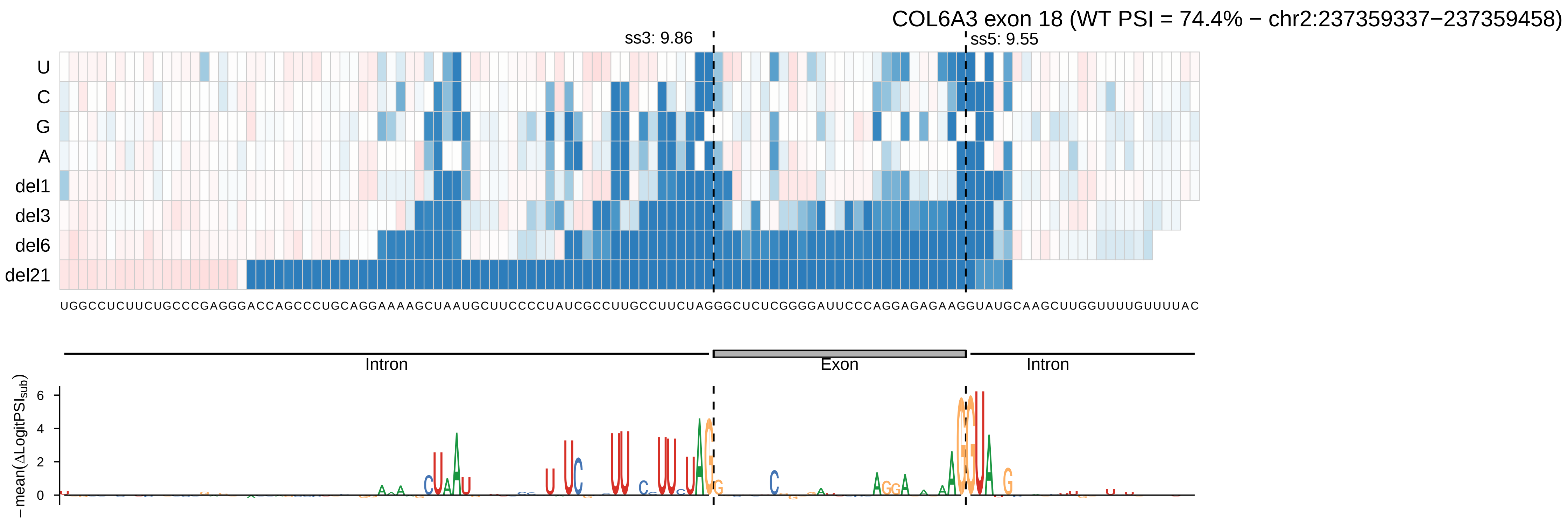
COL6A3 exon 18 (WT PSI = 74.4% − chr2:237359337−237359458)
ss3: 9.86
ss5: 9.55
U
C
G
A
del1
del3
del6
del21
UGGCCUCUUCUGCCCGAGGGACCAGCCCUGCAGGAAAAGCUAAUGCUUCCCCUAUCGCCUUGCCUUCUAGGGCUCUCGGGGAUUCCCAGGAGAGAAGGUAUGCAAGCUUGGUUUUGUUUUAC
Intron
Exon
Intron
−mean(ΔLogitPSIsub)
6
4
2
0

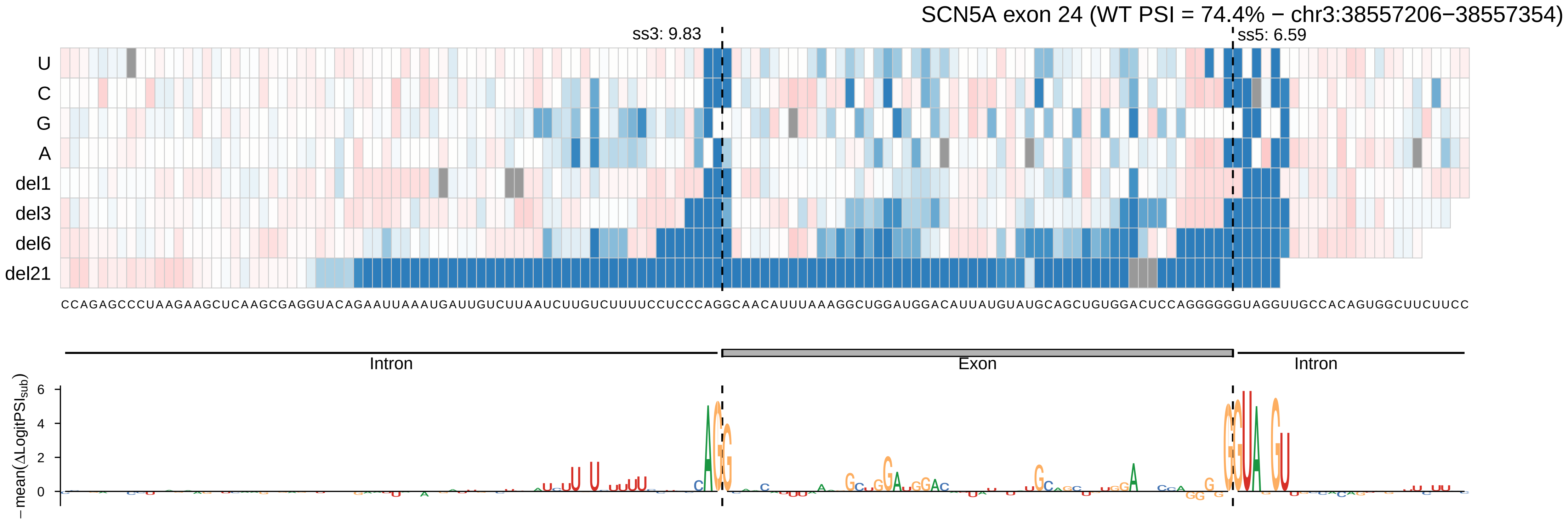

SCN5A exon 24 (WT PSI = 74.4% − chr3:38557206−38557354)
ss3: 9.83
ss5: 6.59
U
C
G
A
del1
del3
del6
del21
CCAGAGCCCUAAGAAGCUCAAGCGAGGUACAGAAUUAAAUGAUUGUCUUAAUCUUGUCUUUUCCUCCCAGGCAACAUUUAAAGGCUGGAUGGACAUUAUGUAUGCAGCUGUGGACUCCAGGGGGGUAGGUUGCCACAGUGGCUUCUUCC
Intron
Exon
Intron
− mean(ΔLogitPSI_sub)
6
4
2
0

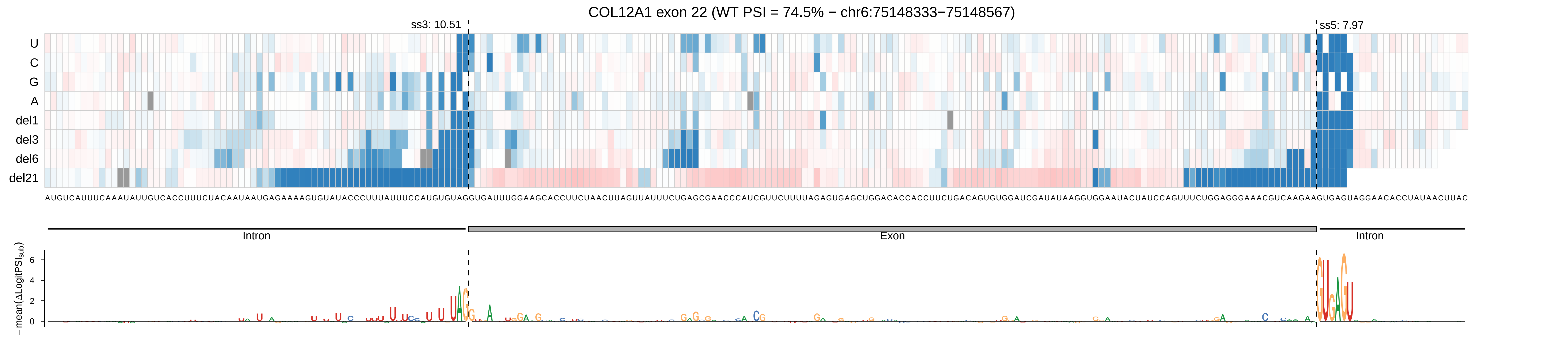
COL12A1 exon 22 (WT PSI = 74.5% − chr6:75148333−75148567)
ss3: 10.51
ss5: 7.97
U
C
G
A
del1
del3
del6
del21
Intron
Exon
Intron
− mean(ΔLogitPSI_sub)
6
4
2
0

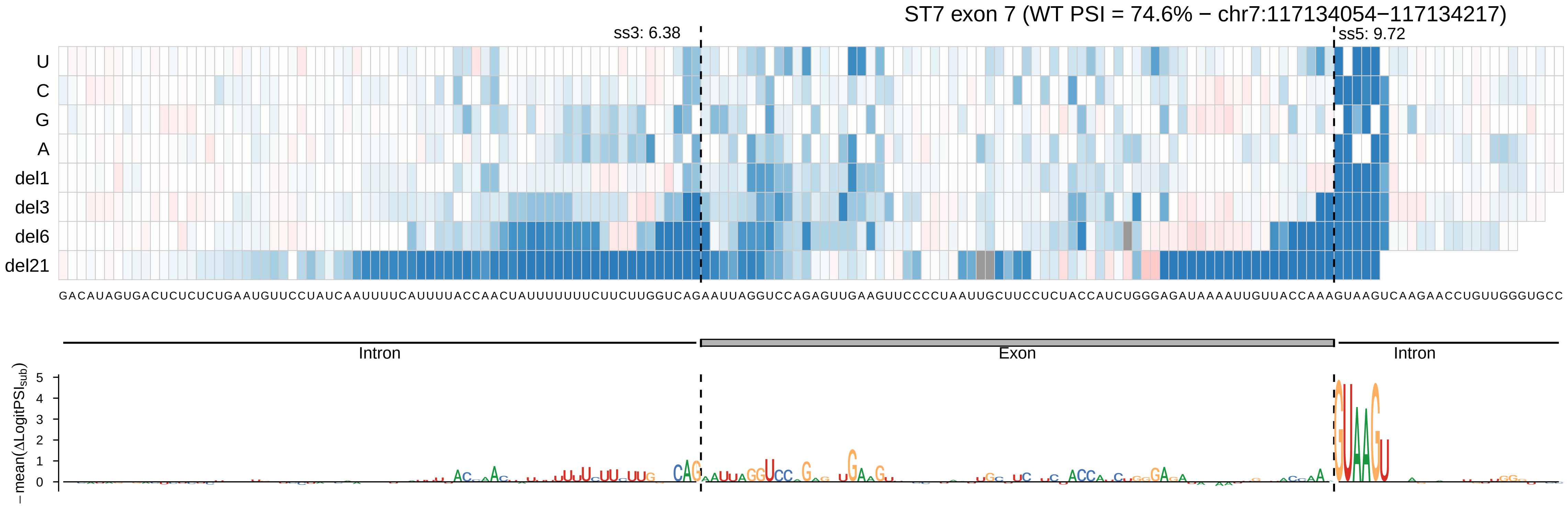

ST7 exon 7 (WT PSI = 74.6% − chr7:117134054−117134217)
ss3: 6.38
ss5: 9.72
U
C
G
A
del1
del3
del6
del21
GACAUAGUGACUCUCUCUGAAUGUUCCUAUCAAUUUUCAUUUUACCAACUAUUUUUUUCUUCUUGGUCAGAAUUAGGUCCAGAGUUGAAGUUCCCCUAAUUGCUUCCUCUACCAUCUGGGAGAUAAAAUUGUUACCAAAGUAAGUCAAGAACCUGUUGGGUGCC
Intron
Exon
Intron
−mean(ΔLogitPSI_sub)
0
1
2
3
4
5

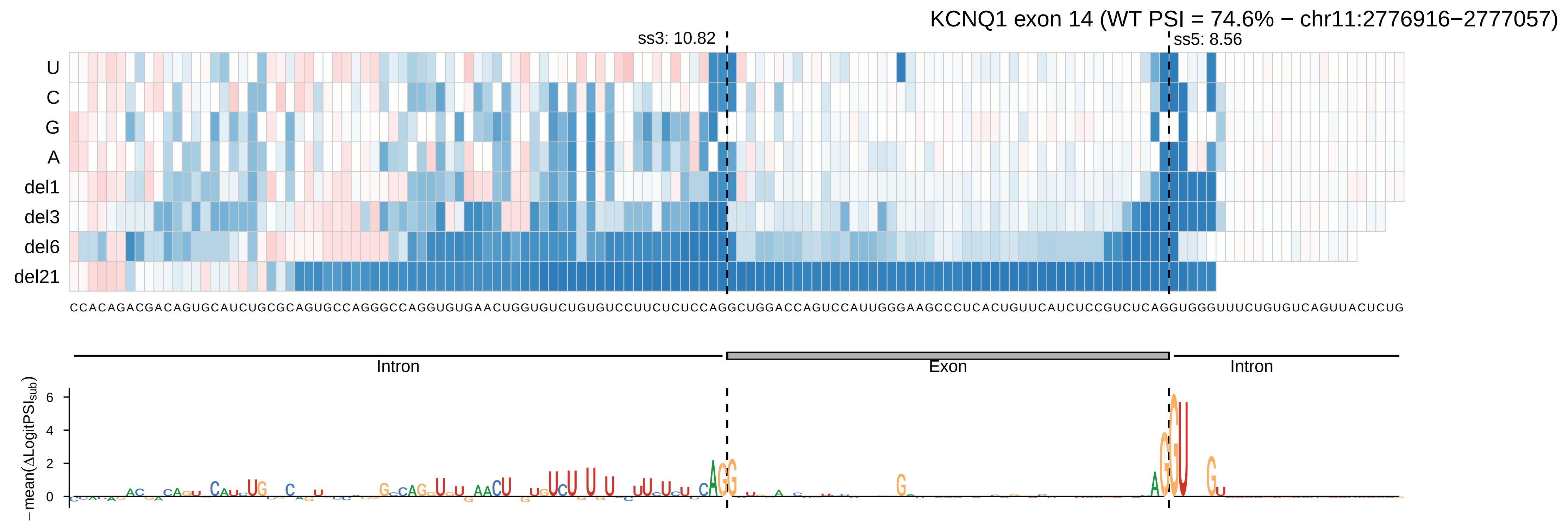

KCNQ1 exon 14 (WT PSI = 74.6% − chr11:2776916−2777057)
ss3: 10.82
ss5: 8.56
U
C
G
A
del1
del3
del6
del21
CCACAGACGACAGUGCAUCUGCGCAGUGCCAGGGCCAGGUGUGAACUGGUGUCUGUGUCCUUCUCUCCAGGCUGGACCAGUCCAUUGGGAAGCCCUCACUGUUCAUCUCCGUCUCAGGUGGGUUUCUGUGUCAGUUACUCUG
Intron
Exon
Intron
−mean(ΔLogitPSI_sub)
6
4
2
0

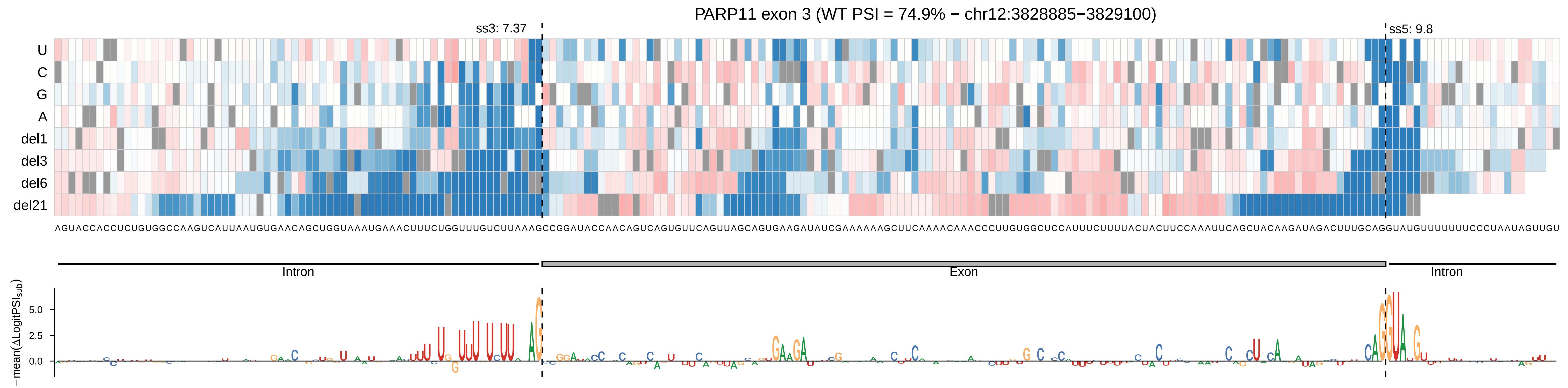
PARP11 exon 3 (WT PSI = 74.9% − chr12:3828885−3829100)
ss3: 7.37
ss5: 9.8
U
C
G
A
del1
del3
del6
del21
AGUACCACCUCUGUGGCCAAGUCAUUAAUGUGAACAGCUGGUAAAUGAAACUUUCUGGUUUGUCUUAAAGCCGGAUACCAACAGUCAGUGUUCAGUUAGCAGUGAAGAUAUCGAAAAAAGCUUCAAAACAAACCCUUGUGGCUCCAUUUCUUUUACUACUUCCAAAUUCAGCUACAAGAUAGACUUUGCAGGUAUGUUUUUUUCCCUAAUAGUUGU
Intron
Exon
Intron
− mean(ΔLogitPSI_sub)
5.0
2.5
0.0

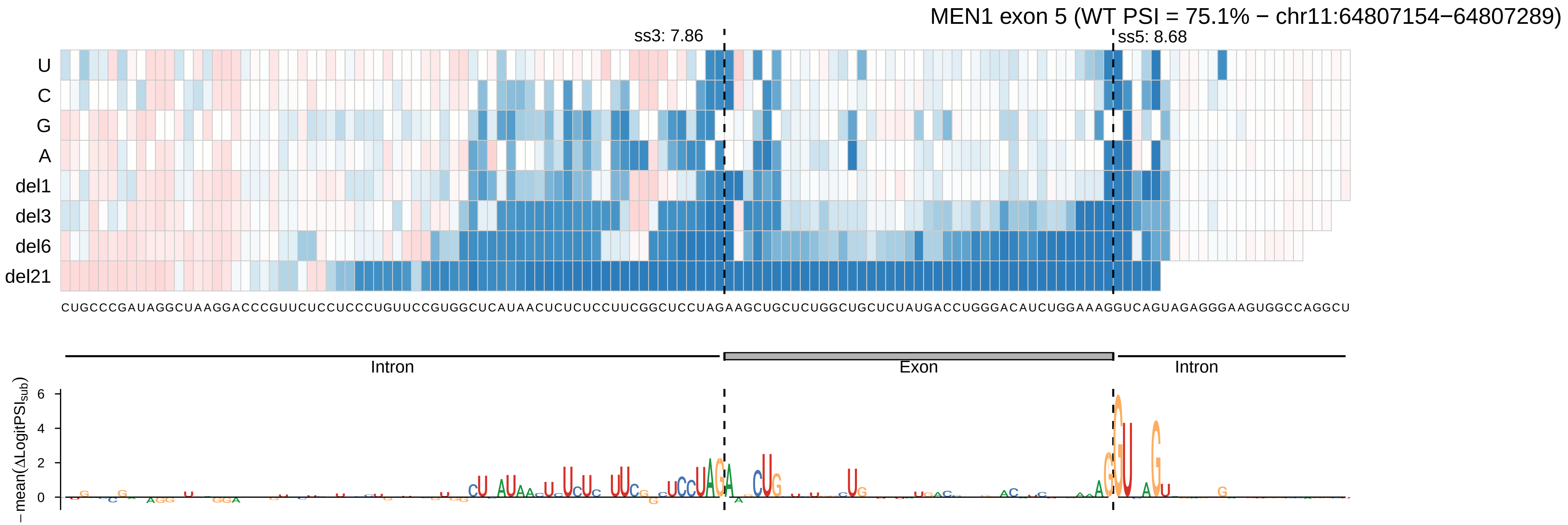

MEN1 exon 5 (WT PSI = 75.1% − chr11:64807154−64807289)
ss3: 7.86
ss5: 8.68
U
C
G
A
del1
del3
del6
del21
CUGCCCGAUAGGCUAAGGACCCGUUCUCCUCCCUGUUCCGUGGCUCAUAACUCUCUCCUUCGGCUCCUAGAAGCUGCUCUGGCUGCUCUAUGACCUGGGACAUCUGGAAAGGUCAGUAGAGGGAAGUGGCCAGGCU
Intron
Exon
Intron
− mean(ΔLogitPSI_sub)
6
4
2
0

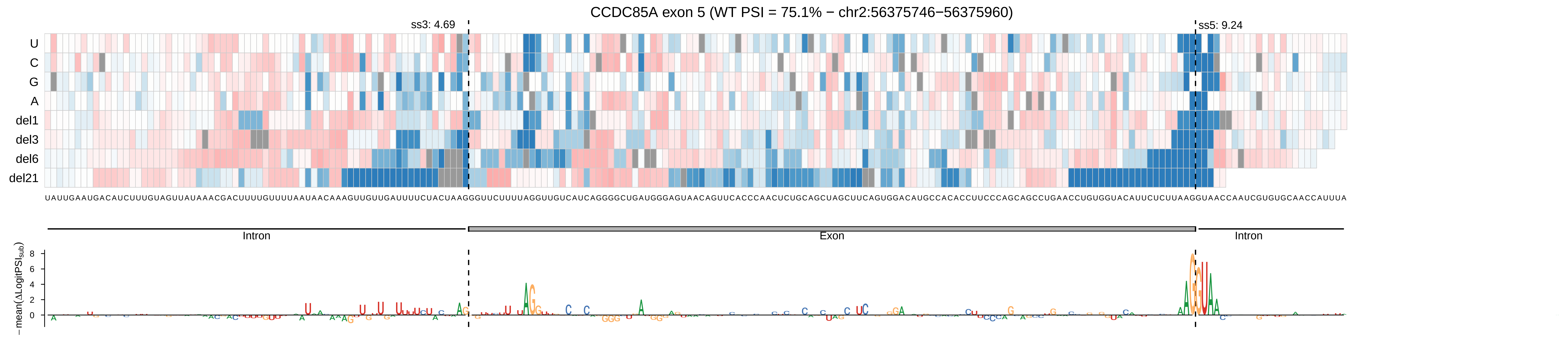

CCDC85A exon 5 (WT PSI = 75.1% − chr2:56375746−56375960)
ss3: 4.69
ss5: 9.24
U
C
G
A
del1
del3
del6
del21
Intron
Exon
Intron
−mean(ΔLogitPSI_sub)

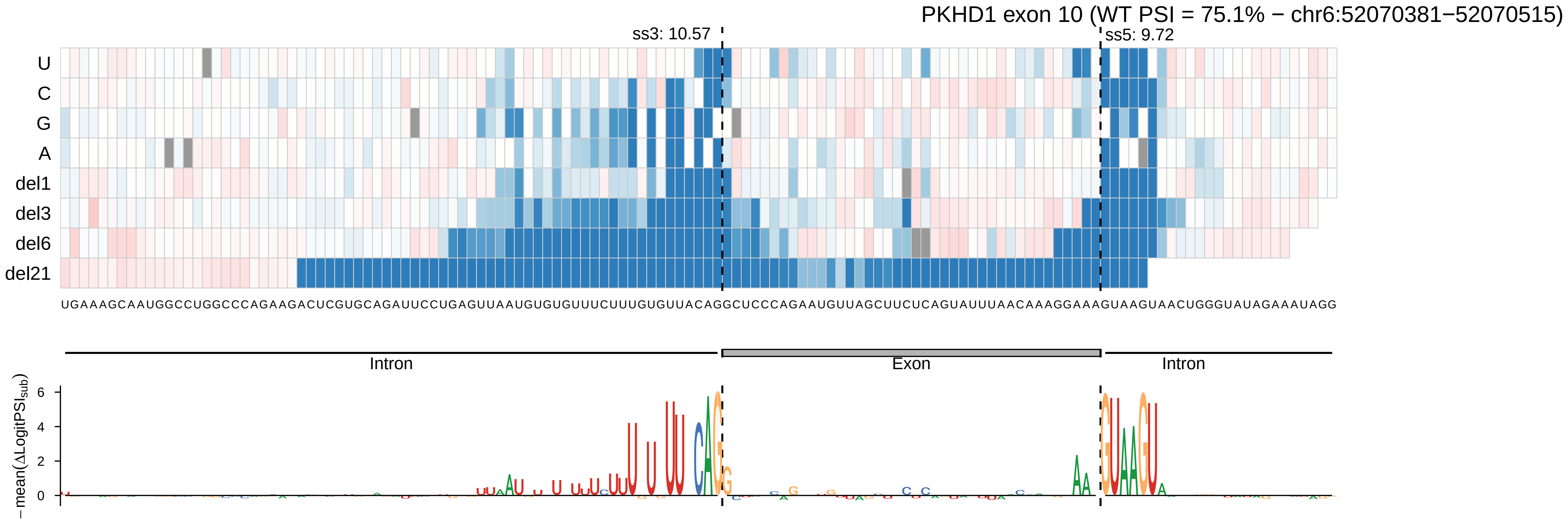
PKHD1 exon 10 (WT PSI = 75.1% − chr6:52070381−52070515)
ss3: 10.57
ss5: 9.72
U
C
G
A
del1
del3
del6
del21
UGAAAGCAAUGGCCUGGCCCAGAAGACUCGUGCAGAUUCCUGAGUUAAUGUGUGUUUCUUUGUGUUACAGGCUCCCAGAAUGUUAGCUUCUCAGUAUUUAACAAAGGAAAGUAAGUAACUGGGUAUAGAAAUAGG
Intron
Exon
Intron
−mean(ΔLogitPSI_sub)
6
4
2
0

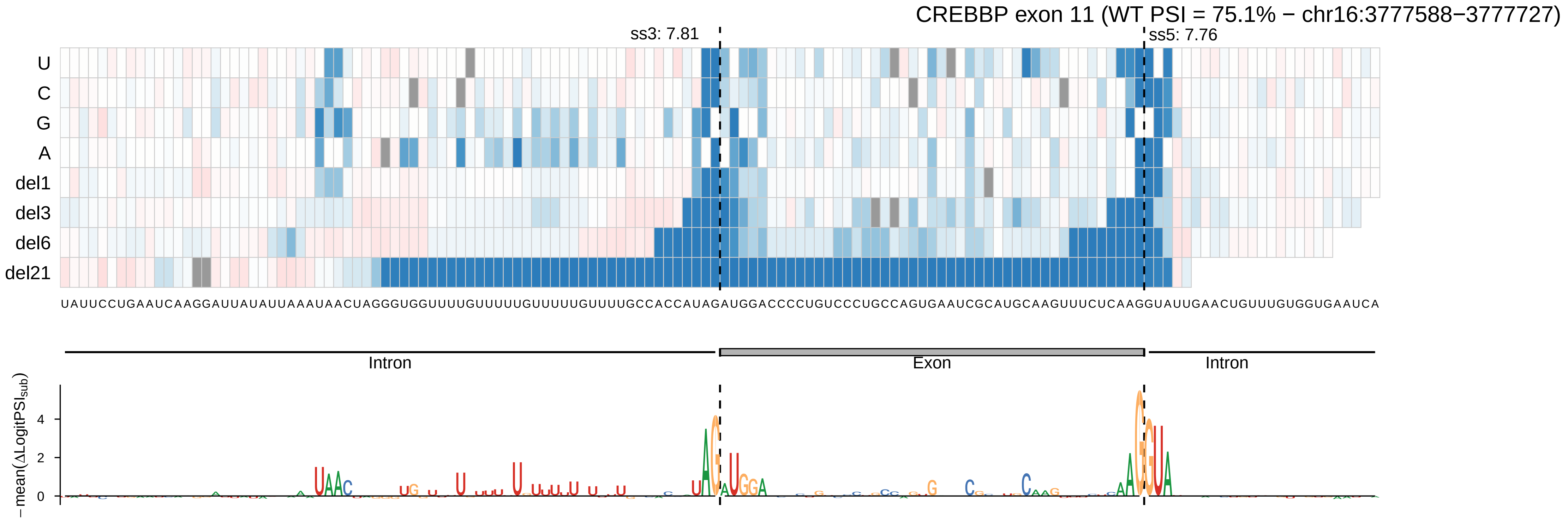

CREBBP exon 11 (WT PSI = 75.1% − chr16:3777588−3777727)
ss3: 7.81
ss5: 7.76
U
C
G
A
del1
del3
del6
del21
UAUUCCUGAAUCAAGGAUUAUAUUAAAUAACUAGGGUGGUUUUGUUUUUGUUUUUGUUUUGCCACCAUAGAUGGACCCCUGUCCCUGCCAGUGAAUCGCAUGCAAGUUUCUCAAGGUAUUGAACUGUUUGUGGUGAAUCA
Intron
Exon
Intron
− mean(ΔLogitPSI_sub)
4
2
0

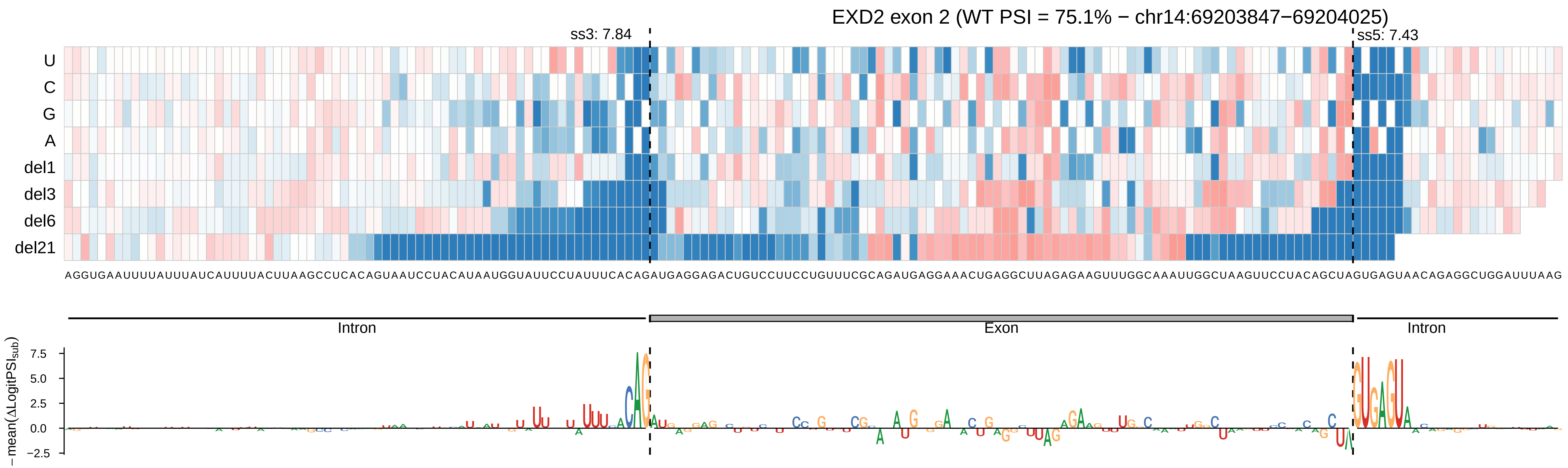
EXD2 exon 2 (WT PSI = 75.1% − chr14:69203847−69204025)
ss3: 7.84
ss5: 7.43
U
C
G
A
del1
del3
del6
del21
AGGUGAAUUUUAUUUAUCAUUUUACUUAAGCCUCACAGUAAUCCUACAUAAUGGUAUUCCUAUUUCACAGAUGAGGAGACUGUCCUUCCUGUUUCGCAGAUGAGGAAACUGAGGCUUAGAGAAGUUUGGCAAAUUGGCUAAGUUCCUACAGCUAGUGAGUAACAGAGGCUGGAUUUAAG
Intron
Exon
Intron
− mean(ΔLogitPSI_sub)
7.5
5.0
2.5
0.0
−2.5

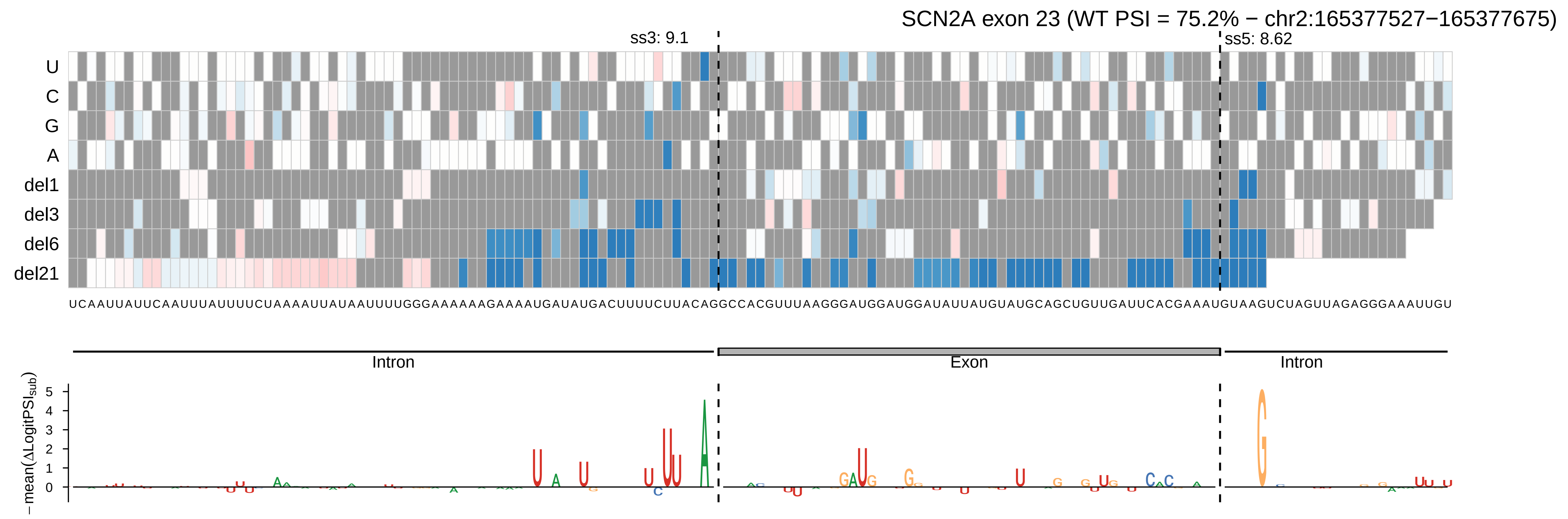

SCN2A exon 23 (WT PSI = 75.2% − chr2:165377527−165377675)
ss3: 9.1
ss5: 8.62
U
C
G
A
del1
del3
del6
del21
UCAAUUAUUCAAUUUAUUUUCUAAAAUUAUAAUUUGGGAAAAAAGAAAAUGAUAUGACUUUUCUUACAGGCCACGUUUAAGGGAUGGAUGGAUAUUAUGUAUGCAGCUGUUGAUUCACGAAAUGUAAGUCUAGUUAGAGGGAAAUUGU
Intron
Exon
Intron
−mean(ΔLogitPSI_sub)

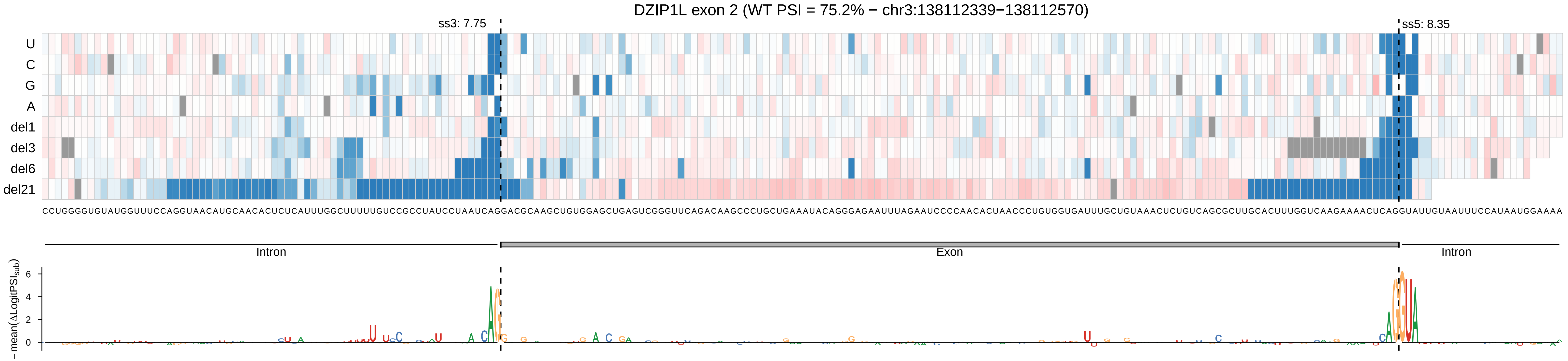

DZIP1L exon 2 (WT PSI = 75.2% − chr3:138112339−138112570)
ss3: 7.75
ss5: 8.35
U
C
G
A
del1
del3
del6
del21
CCUGGGGUGUAUGGUUUCCAGGUAACAUGCAACACUCUCAUUUGGCUUUUUGUCCGCCUAUCCUAAUCAGGACGCAAGCUGUGGAGCUGAGUCGGGUUCAGACAAGCCCUGCUGAAAUACAGGGAGAAUUUAGAAUCCCCAACACUAACCCUGUGGUGAUUUGCUGUAAACUCUGUCAGCGCUUGCACUUUGGUCAAGAAAACUCAGGUAUUGUAAUUUCCAUAAUGGAAAA
Intron
Exon
Intron
−mean(ΔLogitPSI_sub)
6
4
2
0

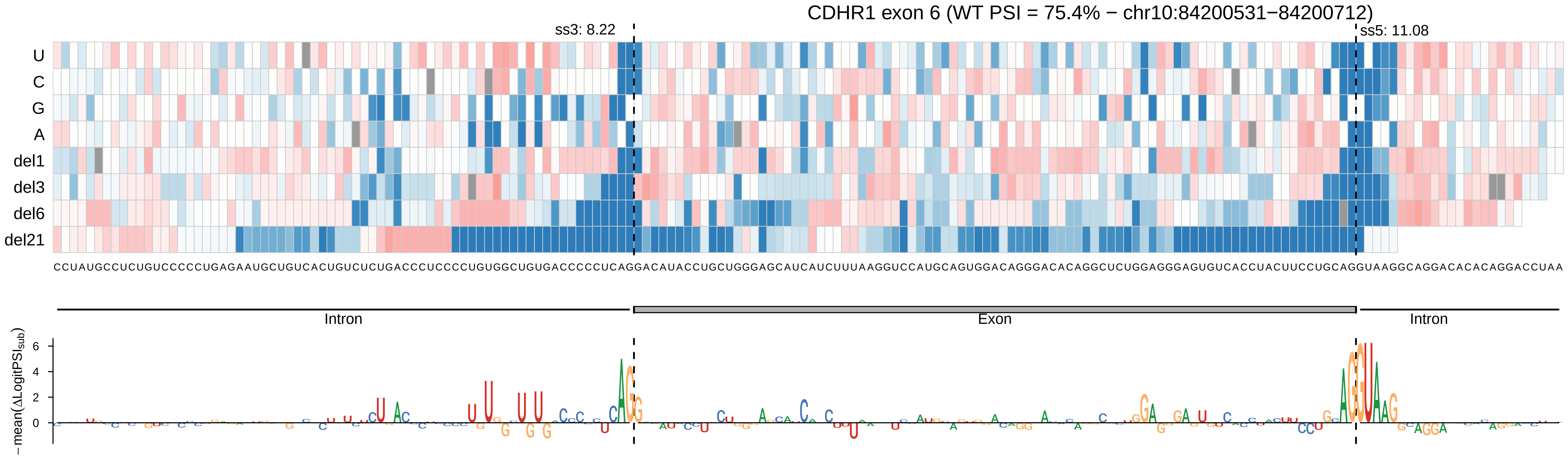

CDHR1 exon 6 (WT PSI = 75.4% − chr10:84200531−84200712)
ss3: 8.22
ss5: 11.08
U
C
G
A
del1
del3
del6
del21
CCUAUGCCUCUGUCCCCCUGAGAAUGCUGUCACUGUCUCUGACCCUCCCCUGUGGCUGUGACCCCCUCAGGACAUACCUGCUGGGAGCAUCAUCUUUAAGGUCCAUGCAGUGGACAGGGACACAGGCUCUGGAGGGAGUGUCACCUACUUCCUGCAGGUAAGGCAGGACACACAGGACCUAA
Intron
Exon
Intron
−mean(ΔLogitPSI_sub)
6
4
2
0

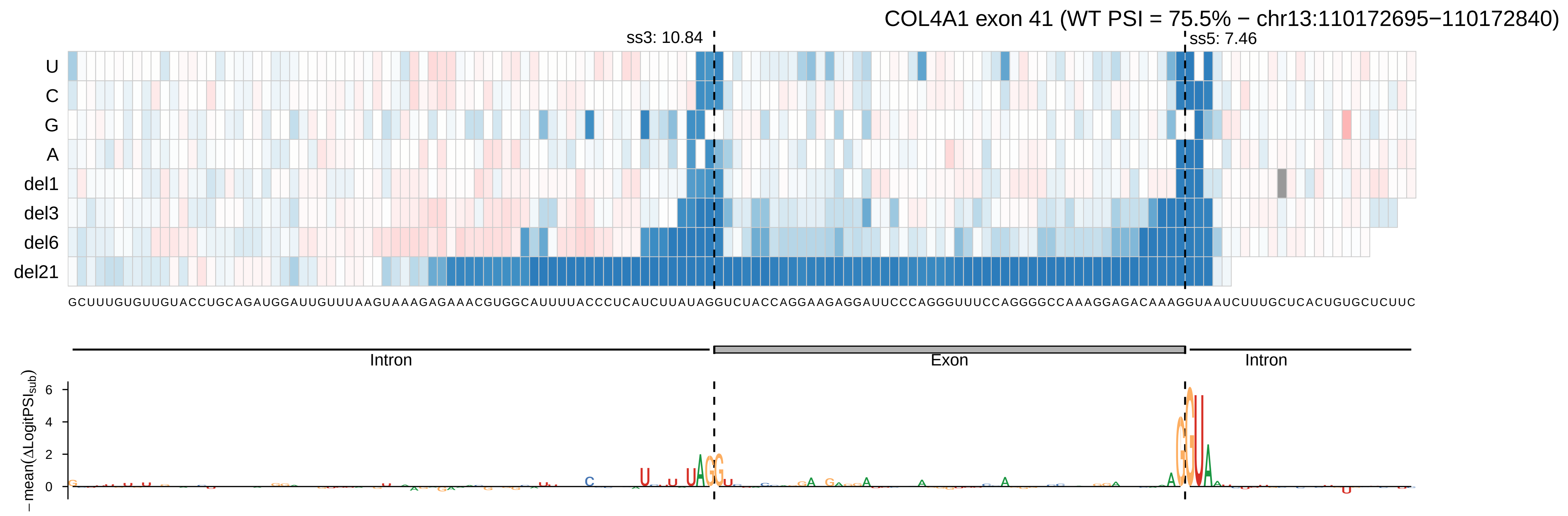

COL4A1 exon 41 (WT PSI = 75.5% − chr13:110172695−110172840)
ss3: 10.84
ss5: 7.46
U
C
G
A
del1
del3
del6
del21
GCUUUGUGUUGUACCUGCAGAUGGAUUGUUUAAGUAAAGAGAAACGUGGCAUUUUACCCUCAUCUUAUAGGUCUACCAGGAAGAGGAUUCCCAGGGUUUCCAGGGGCCAAAGGAGACAAAGGUAAUCUUUGCUCACUGUGCUCUUC
Intron
Exon
Intron
− mean(ΔLogitPSI_sub)
6
4
2
0

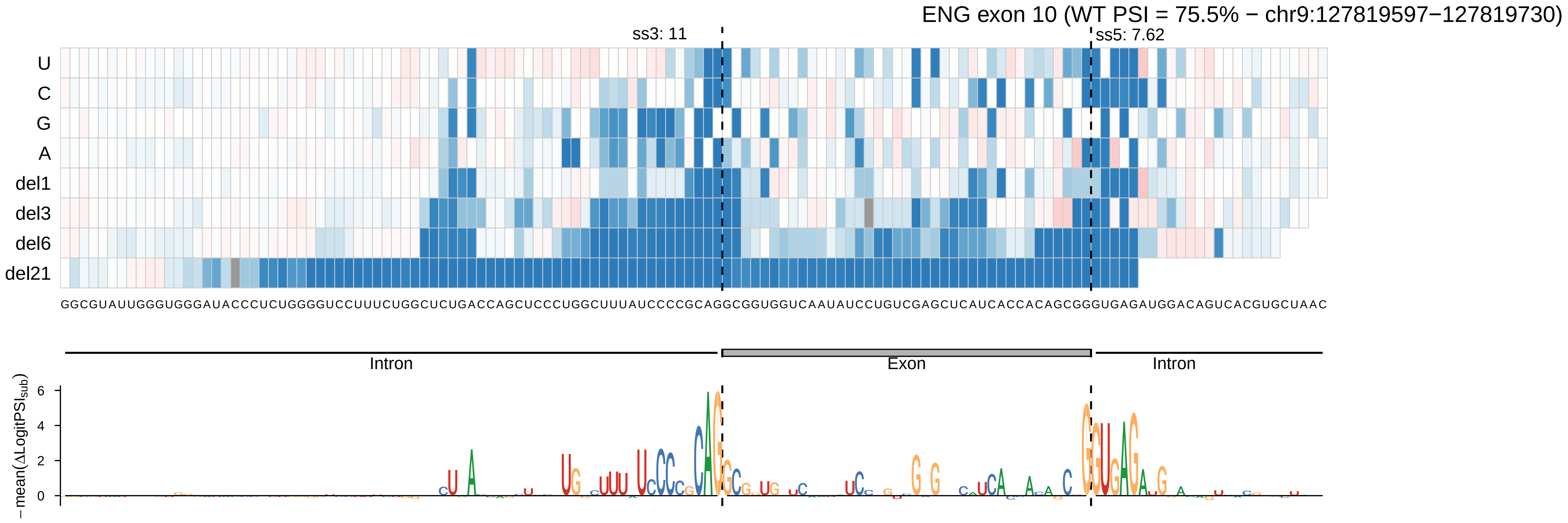
ENG exon 10 (WT PSI = 75.5% − chr9:127819597−127819730)
ss3: 11
ss5: 7.62
U
C
G
A
del1
del3
del6
del21
GGCGUAUUGGGUGGGAUACCCUCUGGGGUCCUUUCUGGCUCUGACCAGCUCCCUGGCUUUAUCCCCGCAGGCGGUGGUCAAUAUCCUGUCGAGCUCAUCACCACAGCGGGUGAGAUGGACAGUCACGUGCUAAC
Intron
Exon
Intron
−mean(ΔLogitPSI_sub)
6
4
2
0

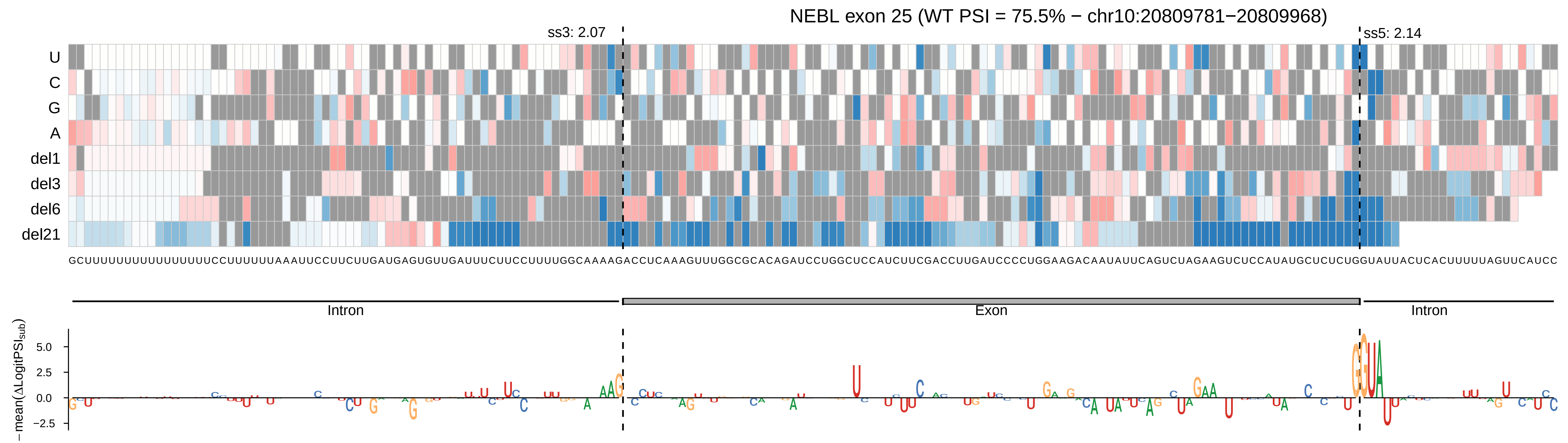

NEBL exon 25 (WT PSI = 75.5% − chr10:20809781−20809968)
ss3: 2.07
ss5: 2.14
U
C
G
A
del1
del3
del6
del21
Intron
Exon
Intron
−mean(ΔLogitPSI_sub)
5.0
2.5
0.0
−2.5

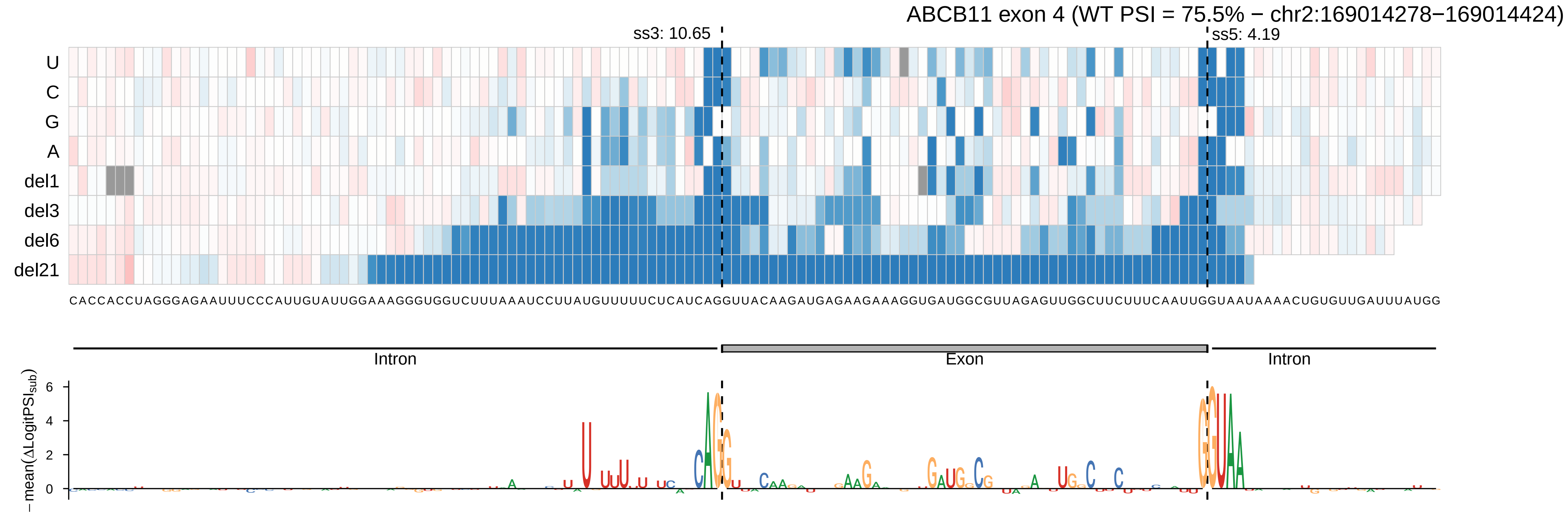
ABCB11 exon 4 (WT PSI = 75.5% − chr2:169014278−169014424)
ss3: 10.65
ss5: 4.19
U
C
G
A
del1
del3
del6
del21
CACCACCUAGGGAGAAUUUCCCAUUGUAUUGGAAAGGGUGGUCUUUAAAUCCUUAUGUUUUUCUCAUCAGGUUACAAGAUGAGAAGAAAGGUGAUGGCGUUAGAGUUGGCUUCUUUCAAUUGGUAAUAAAACUGUGUUGAUUUAUGG
Intron
Exon
Intron
−mean(ΔLogitPSI_sub)
6
4
2
0

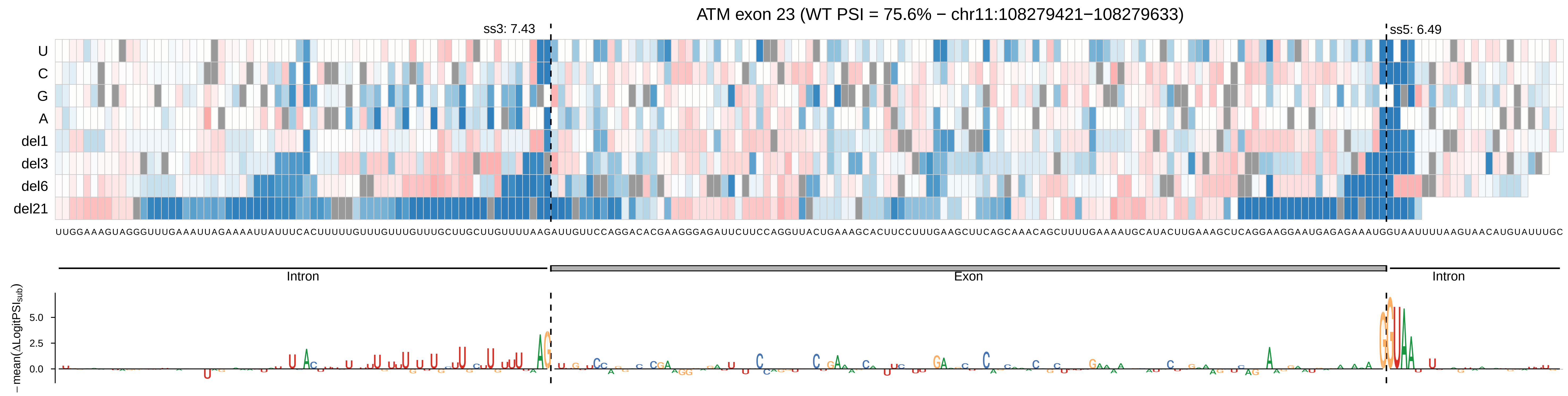
ATM exon 23 (WT PSI = 75.6% − chr11:108279421−108279633)
ss3: 7.43
ss5: 6.49
U
C
G
A
del1
del3
del6
del21
UUGGAAAGUAGGGUUUGAAAUUAGAAAAUUAUUUCACUUUUUGUUUGUUUGUUUGCUUGCUUGUUUUAAGAUUGUUCCAGGACACGAAGGGAGAUUCUUCCAGGUUACUGAAAGCACUUCCUUUGAAGCUUCAGCAAACAGCUUUUGAAAAUGCAUACUUGAAAGCUCAGGAAGGAAUGAGAGAAAUGGUAAUUUUAAGUAACAUGUAUUUGC
Intron
Exon
Intron
− mean(ΔLogitPSI_sub)
5.0
2.5
0.0

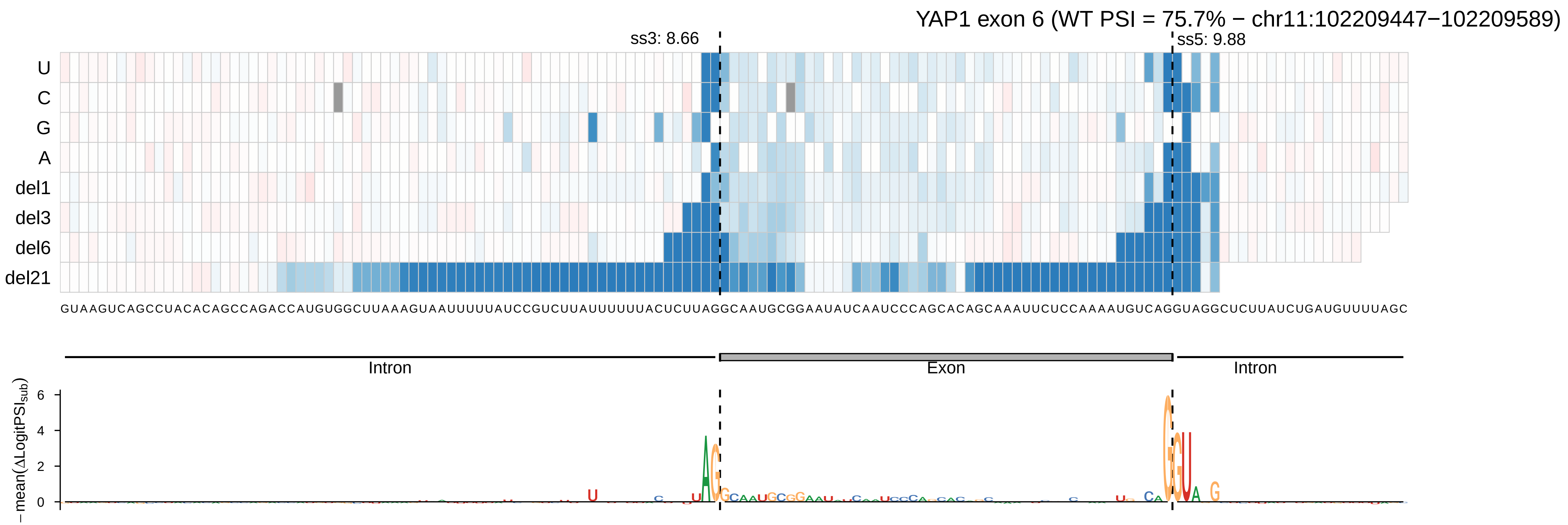
YAP1 exon 6 (WT PSI = 75.7% − chr11:102209447−102209589)
ss3: 8.66
ss5: 9.88
U
C
G
A
del1
del3
del6
del21
GUAAGUCAGCCUACACAGCCAGACCAUGUGGCUUAAAGUAAUUUUUAUCCGUCUUAUUUUUUACUCUUAGGCAAUGCGGAAUAUCAAUCCCAGCACAGCAAAUUCUCCAAAAUGUCAGGUAGGCUCUUAUCUGAUGUUUUAGC
Intron
Exon
Intron
−mean(ΔLogitPSI_sub)
6
4
2
0

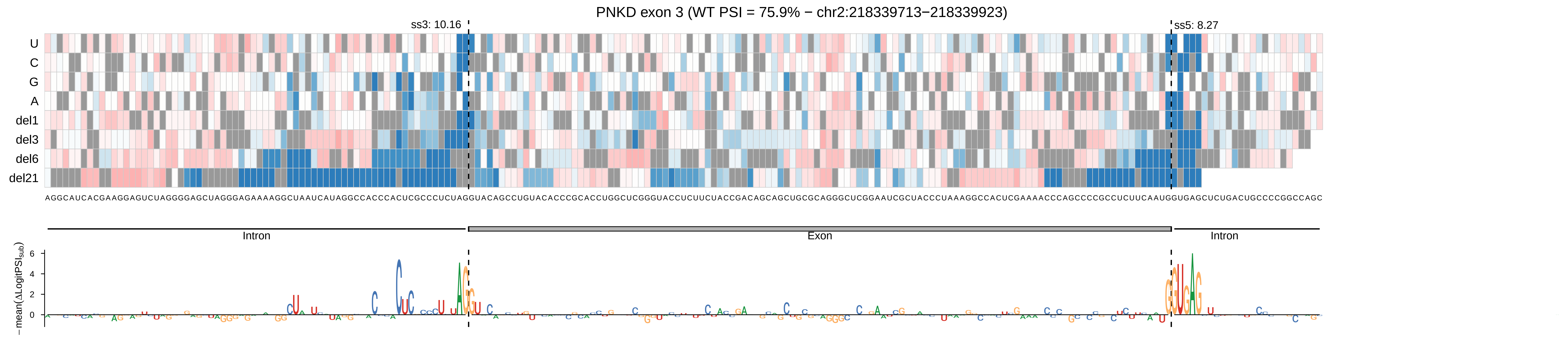
PNKD exon 3 (WT PSI = 75.9% − chr2:218339713−218339923)
ss3: 10.16
ss5: 8.27
U
C
G
A
del1
del3
del6
del21
AGGCAUCACGAAGGAGUCUAGGGGAGCUAGGGAGAAAAGGCUAAUCAUAGGCCACCCACUCGCCCUCUAGGUACAGCCUGUACACCCGCACCUGGCUCGGGUACCUCUUCUACCGACAGCAGCUGCGCAGGGCUCGGAAUCGCUACCCUAAAGGCCACUCGAAAACCCAGCCCCGCCUCUUCAAUGGUGAGCUCUGACUGCCCCGGCCAGC
Intron
Exon
Intron
−mean(ΔLogitPSIsub)
6
4
2
0

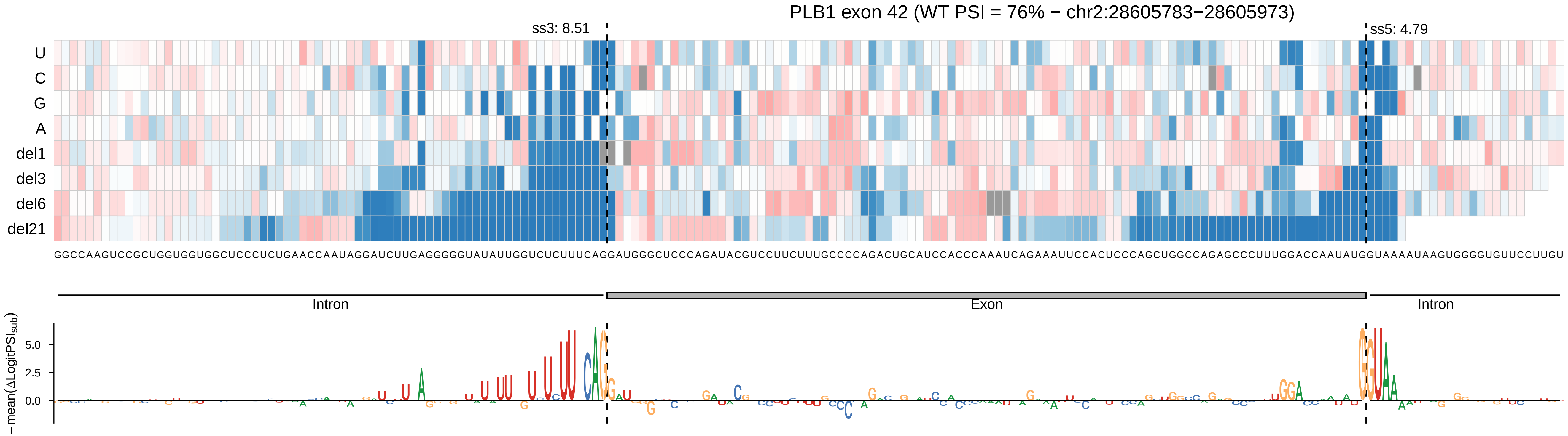

PLB1 exon 42 (WT PSI = 76% − chr2:28605783−28605973)
ss3: 8.51
ss5: 4.79
U
C
G
A
del1
del3
del6
del21
Intron
Exon
Intron
−mean(ΔLogitPSI_sub)
5.0
2.5
0.0

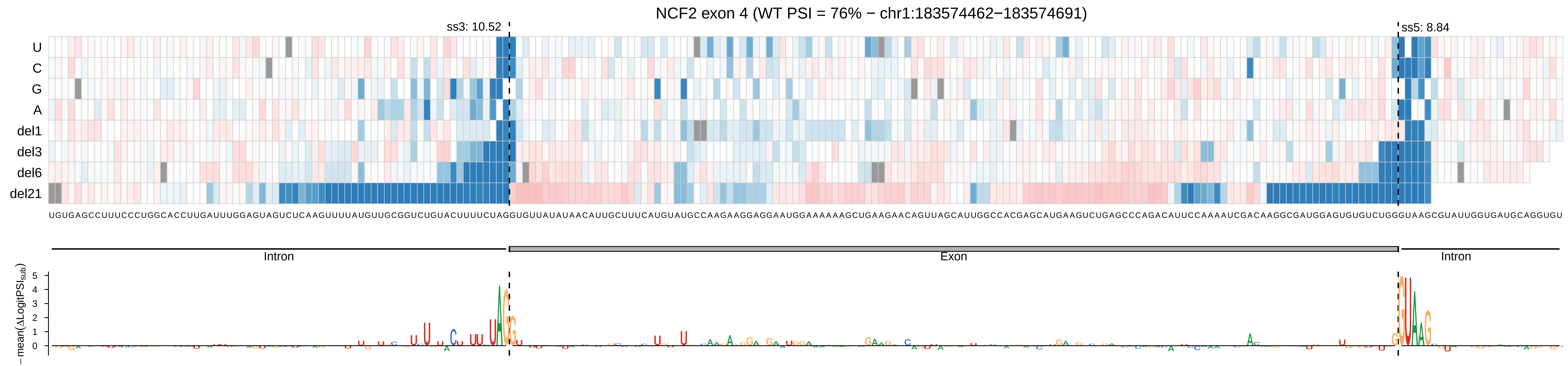

NCF2 exon 4 (WT PSI = 76% − chr1:183574462−183574691)
ss3: 10.52
ss5: 8.84
U
C
G
A
del1
del3
del6
del21
UGUGAGCCUUUCCCUGGCACCUUGAUUUGGAGUAGUCUCAAGUUUUAUGUUGCGGUCUGUACUUUUCUAGGUGUUAUAUAACAUUGCUUUCAUGUAUGCCAAGAAGGAGGAAUGGAAAAAAGCUGAAGAACAGUUAGCAUUGGCCACGAGCAUGAAGUCUGAGCCCAGACAUUCCAAAAUCGACAAGGCGAUGGAGUGUGUCUGGGUAAGCGUAUUGGUGAUGCAGGUGU
Intron
Exon
Intron
−mean(ΔLogitPSI_sub)
5
4
3
2
1
0

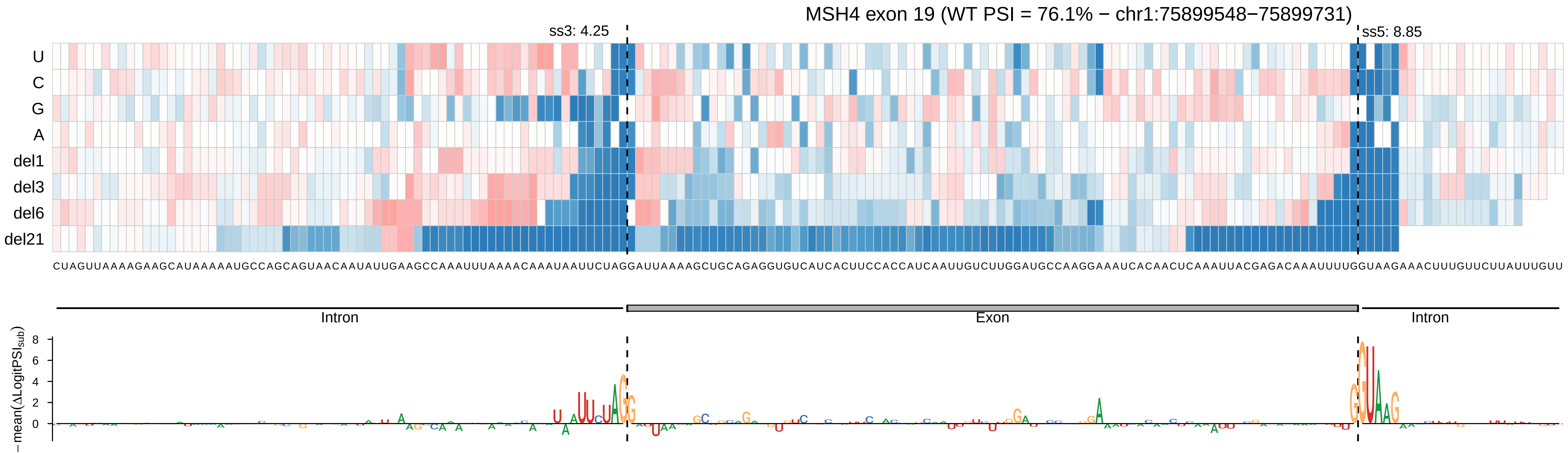

MSH4 exon 19 (WT PSI = 76.1% − chr1:75899548−75899731)
ss3: 4.25
ss5: 8.85
U
C
G
A
del1
del3
del6
del21
CUAGUUAAAAGAAGCAUAAAAAUGCCAGCAGUAACAAUAUUGAAGCCAAAUUUAAAACAAAUAAUUCUAGGAUUAAAAGCUGCAGAGGUGUCAUCACUUCCACCAUCAAUUGUCUUGGAUGCCAAGGAAAUCACAACUCAAAUUACGAGACAAAUUUUGGUAAGAAACUUUGUUCUUAUUUGUU
Intron
Exon
Intron
−mean(ΔLogitPSI$_{sub}$)
8
6
4
2
0

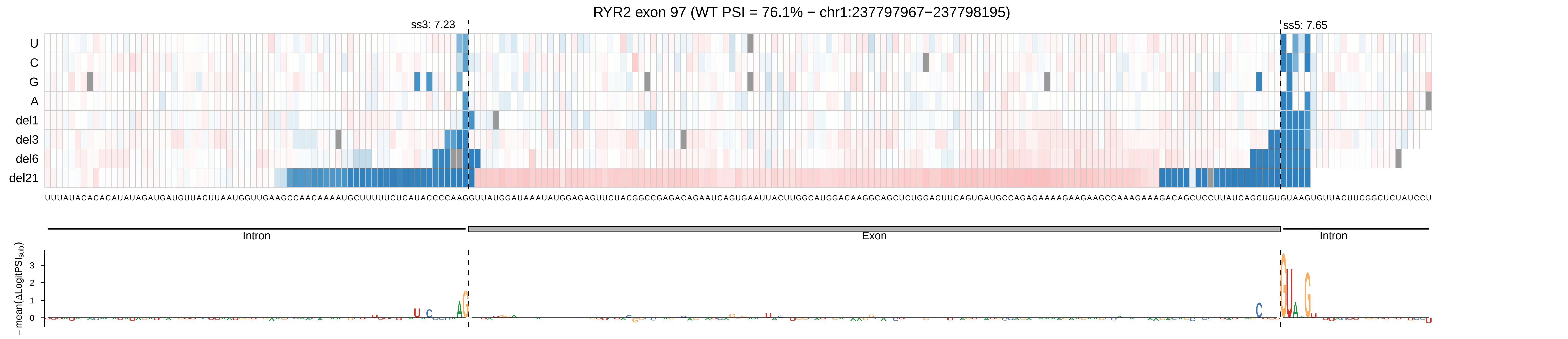
RYR2 exon 97 (WT PSI = 76.1% − chr1:237797967−237798195)
ss3: 7.23
ss5: 7.65
U
C
G
A
del1
del3
del6
del21
UUUAUACACACAUAUAGAUGAUGUUACUUAAUGGUUGAAGCCAACAAAAUGCUUUUUCUCAUACCCCAAGGUUAUGGAUAAAUAUGGAGAGUUCUACGGCCGAGACAGAAUCAGUGAAUUACUUGGCAUGGACAAGGCAGCUCUGGACUUCAGUGAUGCCAGAGAAAAGAAGAAGCCAAAGAAAGACAGCUCCUUAUCAGCUGUGUAAGUGUUACUUCGGCUCUAUCCU
Intron
Exon
Intron
− mean(ΔLogitPSI_sub)
3
2
1
0

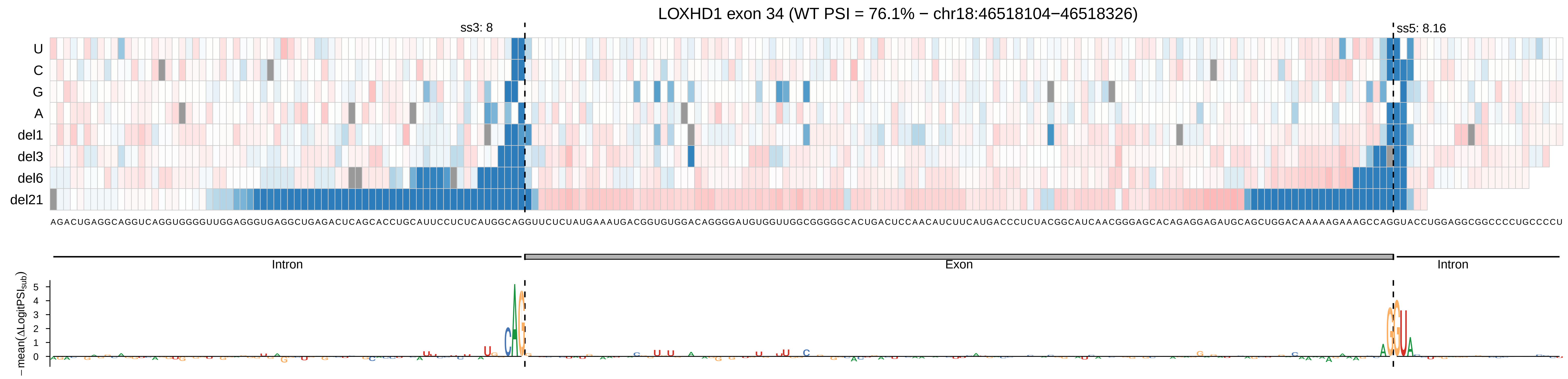

LOXHD1 exon 34 (WT PSI = 76.1% − chr18:46518104−46518326)
ss3: 8
ss5: 8.16
U
C
G
A
del1
del3
del6
del21
AGACUGAGGCAGGUCAGGUGGGGUUGGAGGGUGAGGCUGAGACUCAGCACCUGCAUUCCUCUCAUGGCAGGUUCUCUAUGAAAUGACGGUGUGGACAGGGGAUGUGGUUGGCGGGGGCACUGACUCCAACAUCUUCAUGACCCUCUACGGCAUCAACGGGAGCACAGAGGAGAUGCAGCUGGACAAAAAGAAAGCCAGGUACCUGGAGGCGGCCCCUGCCCCU
Intron
Exon
Intron
− mean(ΔLogitPSI_sub)
0
1
2
3
4
5

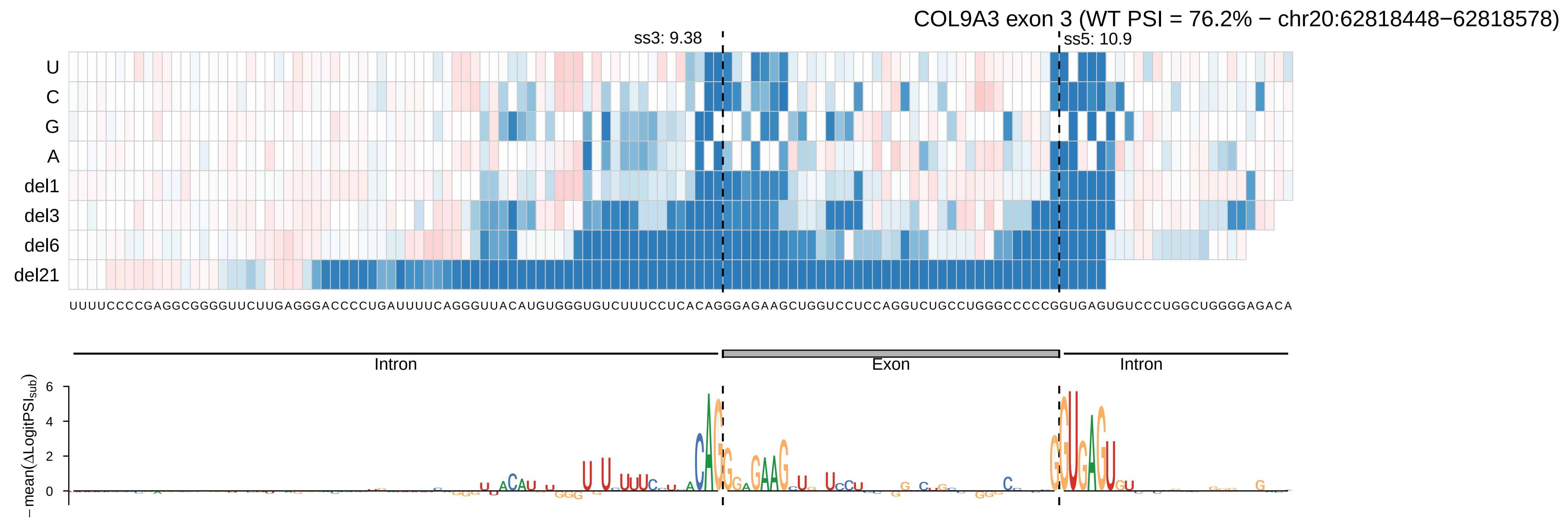

COL9A3 exon 3 (WT PSI = 76.2% − chr20:62818448−62818578)
ss3: 9.38
ss5: 10.9
U
C
G
A
del1
del3
del6
del21
UUUUCCCCGAGGCGGGGUUCUUGAGGGACCCCUGAUUUUCAGGGUUACAUGUGGGUGUCUUUCCUCACAGGGAGAAGCUGGUCCUCCAGGUCUGCCUGGGCCCCCGGUGAGUGUCCCUGGCUGGGGAGACA
Intron
Exon
Intron
−mean(ΔLogitPSI_sub)
6
4
2
0

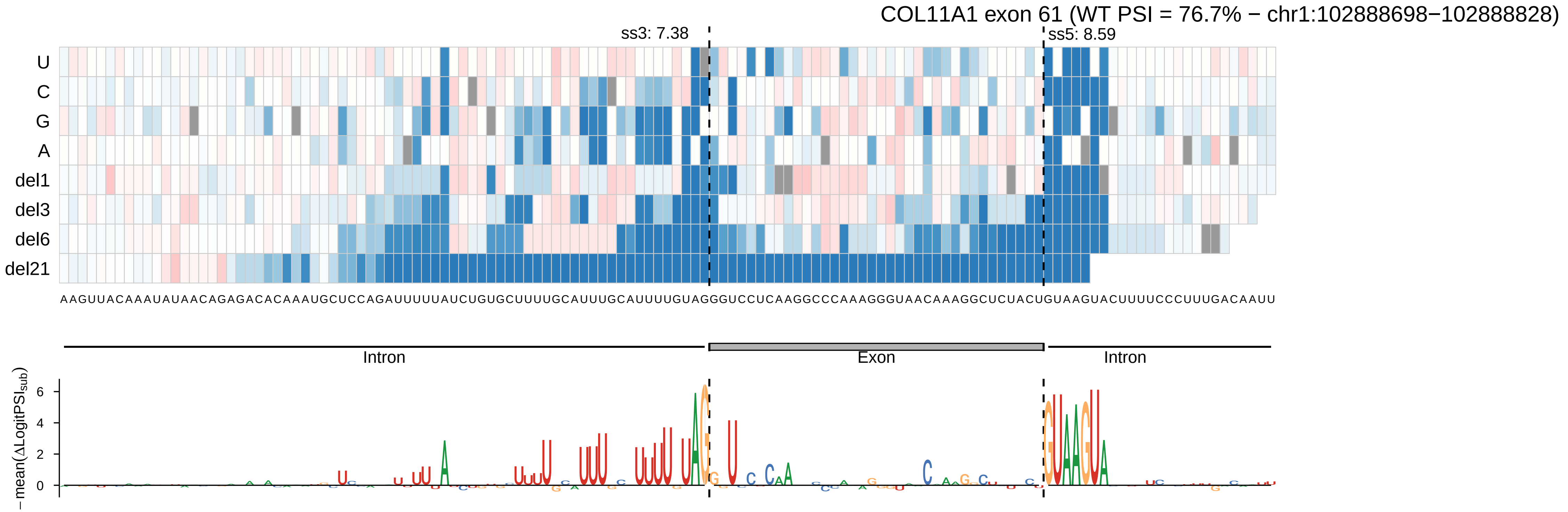

COL11A1 exon 61 (WT PSI = 76.7% − chr1:102888698−102888828)
ss3: 7.38
ss5: 8.59
U
C
G
A
del1
del3
del6
del21
AAGUUACAAAUAUAACAGAGACACAAAUGCUCCAGAUUUUUAUCUGUGCUUUUGCAUUUGCAUUUUGUAGGGUCCUCAAGGCCCAAAGGGUAACAAAGGCUCUACUGUAAGUACUUUUCCCUUUGACAAUU
Intron
Exon
Intron
−mean(ΔLogitPSI_sub)

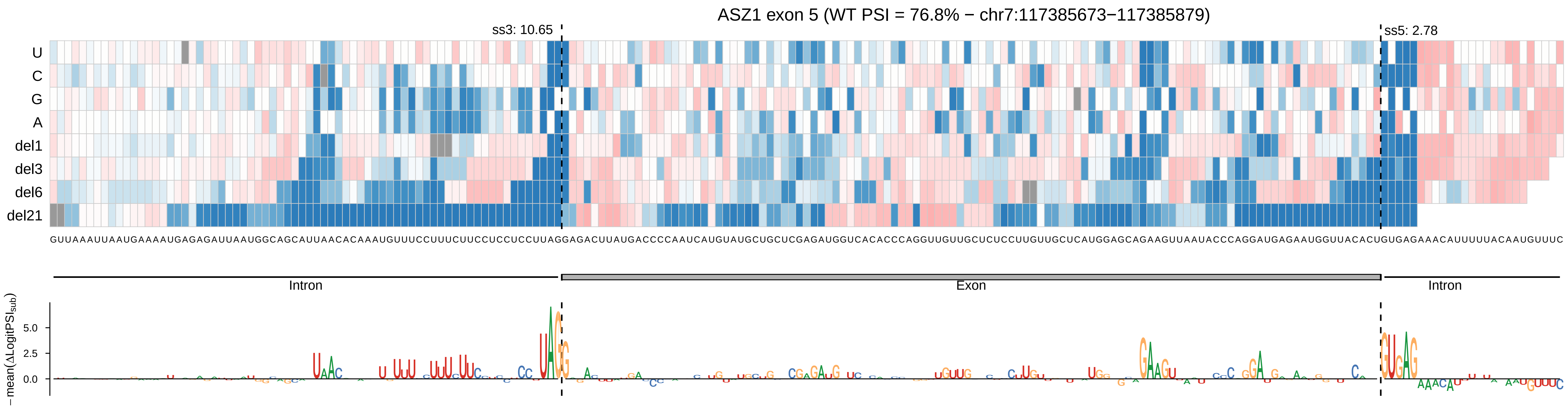

ASZ1 exon 5 (WT PSI = 76.8% − chr7:117385673−117385879)
ss3: 10.65
ss5: 2.78
U
C
G
A
del1
del3
del6
del21
GUUAAAUUAAUGAAAAUGAGAGAUUAAUGGCAGCAUUAACACAAAUGUUUCCUUUCUUCCUCCUCCUUAGGAGACUUAUGACCCCAAUCAUGUAUGCUGCUCGAGAUGGUCACACCCAGGUUGUUGCUCUCCUUGUUGCUCAUGGAGCAGAAGUUAAUACCCAGGAUGAGAAUGGUUACACUGUGAGAAACAUUUUUACAAUGUUUC
Intron
Exon
Intron
−mean(ΔLogitPSI_sub)
5.0
2.5
0.0

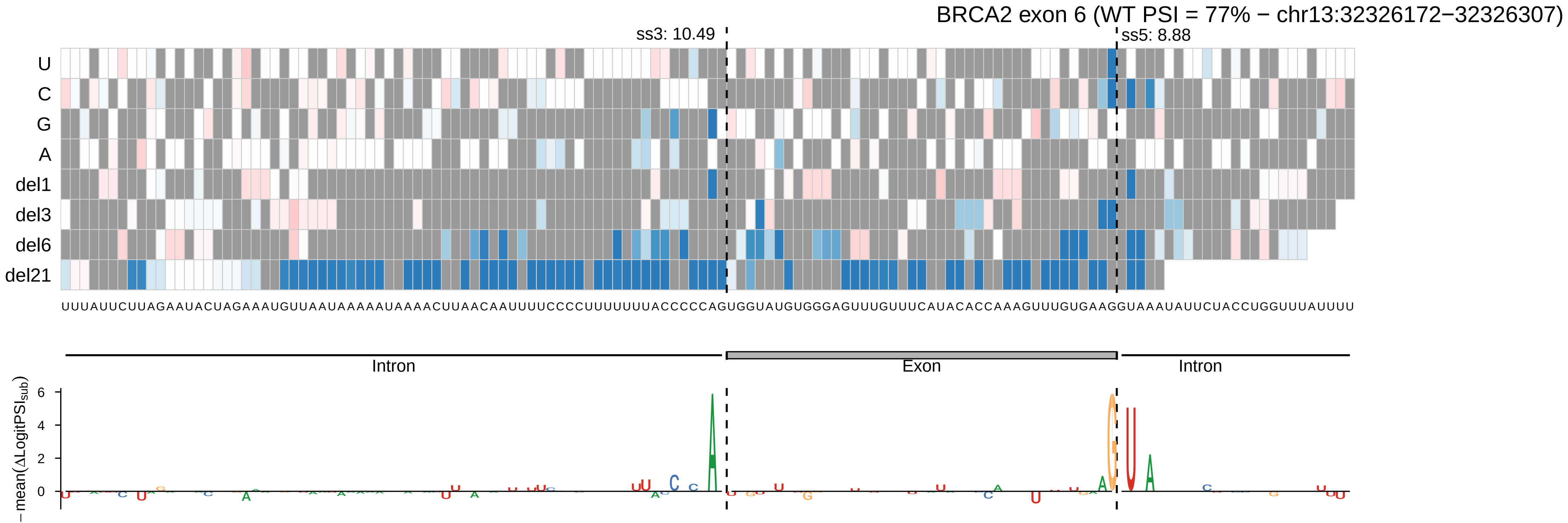
BRCA2 exon 6 (WT PSI = 77% − chr13:32326172−32326307)
ss3: 10.49
ss5: 8.88
U
C
G
A
del1
del3
del6
del21
UUUAUUCUUAGAAUACUAGAAAUGUUAAUAAAAAAUAAAACUUAACAAUUUUCCCCUUUUUUUUACCCCCAGUGGUAUGUGGGAGUUUGUUUCAUACACCAAAGUUUGUGAAGGUAAAAUAUUCUACCUGGUUUAUUUU
Intron
Exon
Intron
−mean(ΔLogitPSI_sub)
6
4
2
0

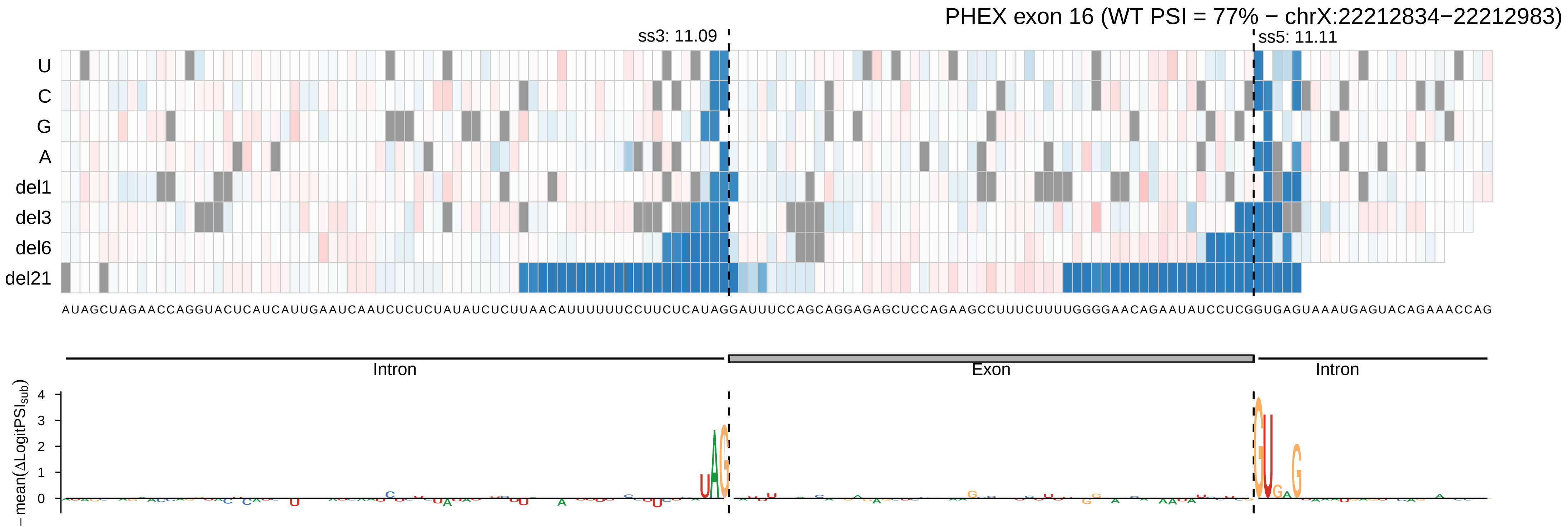

PHEX exon 16 (WT PSI = 77% − chrX:22212834−22212983)
ss3: 11.09
ss5: 11.11
U
C
G
A
del1
del3
del6
del21
AUAGCUAGAACCAGGUACUCAUCAUUGAAUCAAUCUCUCUAUAUCUCUUAACAUUUUUUCCUUCUCAUAGGAUUUCCAGCAGGAGAGCUCCAGAAGCCUUUCUUUUGGGGAACAGAAUAUCCUCGGUGAGUAAAUGAGUACAGAAACCAG
Intron
Exon
Intron
− mean(ΔLogitPSI_sub)
0
1
2
3
4

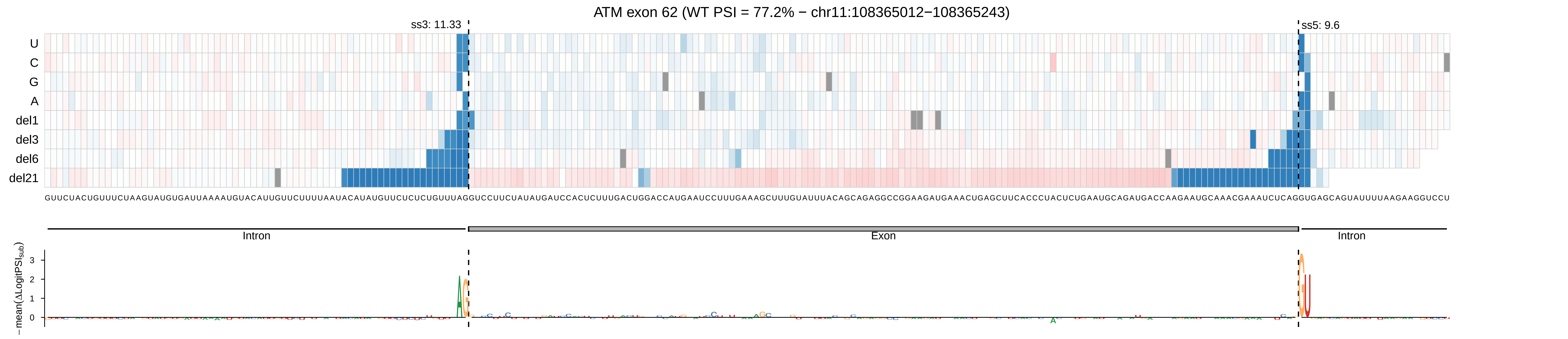

ATM exon 62 (WT PSI = 77.2% − chr11:108365012−108365243)
ss3: 11.33
ss5: 9.6
U
C
G
A
del1
del3
del6
del21
Intron
Exon
Intron
−mean(ΔLogitPSI_sub)
0
1
2
3

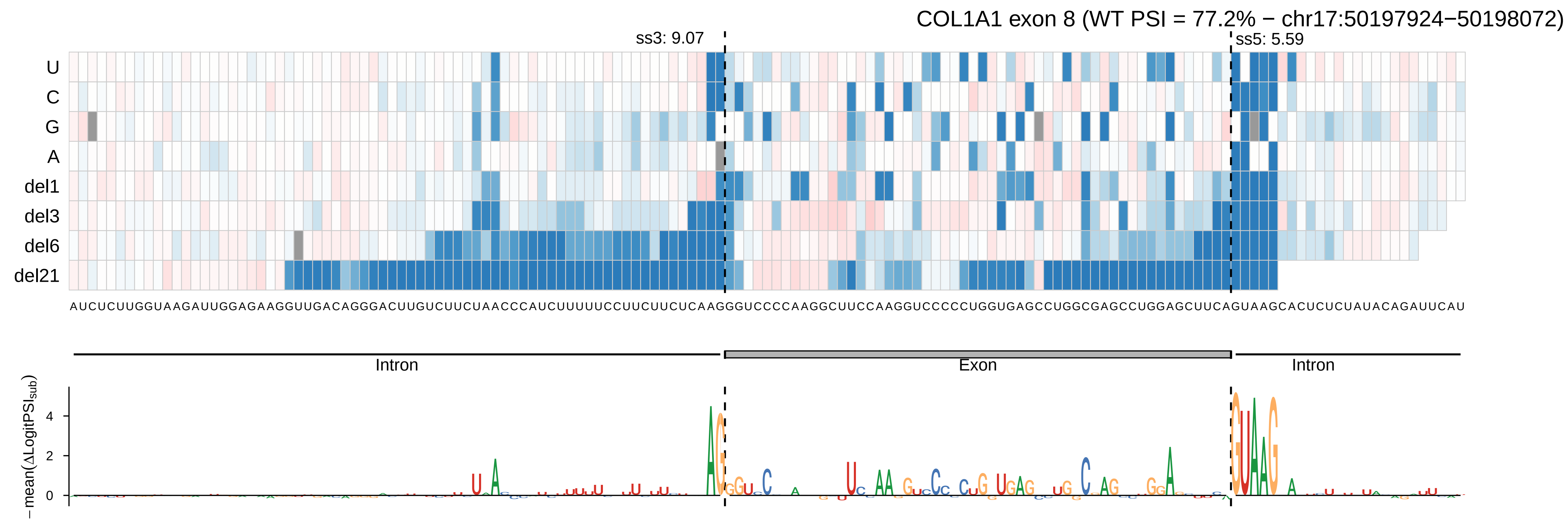
COL1A1 exon 8 (WT PSI = 77.2% − chr17:50197924−50198072)
ss3: 9.07
ss5: 5.59
U
C
G
A
del1
del3
del6
del21
AUCUCUUGGUAAGAUUGGAGAAGGUUGACAGGGACUUGUCUUCUAACCCAUCUUUUUCCUUCUUCUCAAGGGUCCCCAAGGCUUCCAAGGUCCCCCUGGUGAGCCUGGCGAGCCUGGAGCUUCAGUAAGCACUCUCUAUACAGAUUCAU
Intron
Exon
Intron
−mean(ΔLogitPSI_sub)
4
2
0

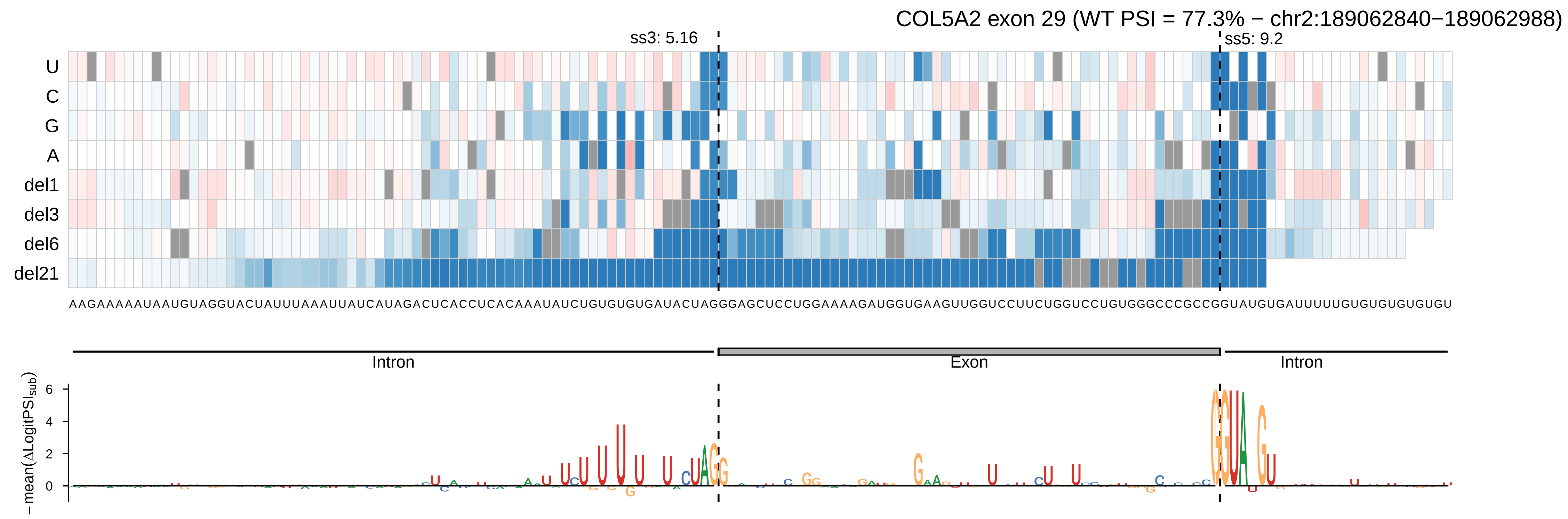
COL5A2 exon 29 (WT PSI = 77.3% − chr2:189062840−189062988)
ss3: 5.16
ss5: 9.2
U
C
G
A
del1
del3
del6
del21
AAGAAAAAUAAUGUAGGUACUAUUUAAAUUAUCAUAGACUCACCUCACAAAUAUCUGUGUGUGAUACUAGGGAGCUCCUGGAAAAGAUGGUGAAGUUGGUCCUUCUGGUCCUGUGGGCCCGCCGGUAUGUGAUUUUUGUGUGUGUGU
Intron
Exon
Intron
−mean(ΔLogitPSI_sub)
6
4
2
0

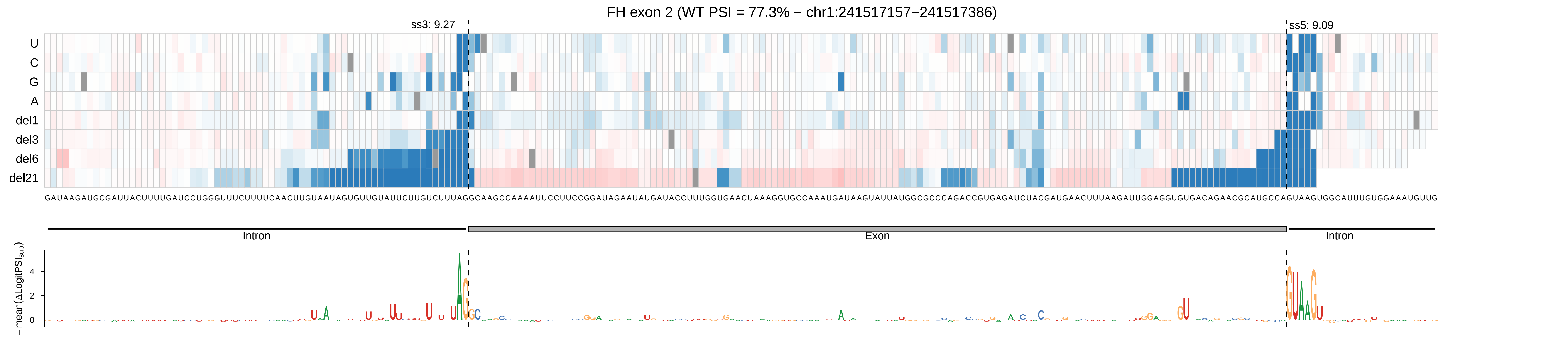
FH exon 2 (WT PSI = 77.3% − chr1:241517157−241517386)
ss3: 9.27
ss5: 9.09
U
C
G
A
del1
del3
del6
del21
GAUAAGAUGCGAUUACUUUUGAUCCUGGGUUUCUUUUCAACUUGUAAUAGUGUUGUAUUCUUGUCUUUAGGCAAGCCAAAAUUCCUUCCGGAUAGAAUAUGAUACCUUUGGUGAACUAAAGGUGCCAAAUGAUAAGUAUUAUGGCGCCCAGACCGUGAGAUCUACGAUGAACUUUAAGAUUGGAGGUGUGACAGAACGCAUGCCAGUAAGUGGCAUUUGUGGAAAUGUUG
Intron
Exon
Intron
− mean(ΔLogitPSI_sub)
4
2
0

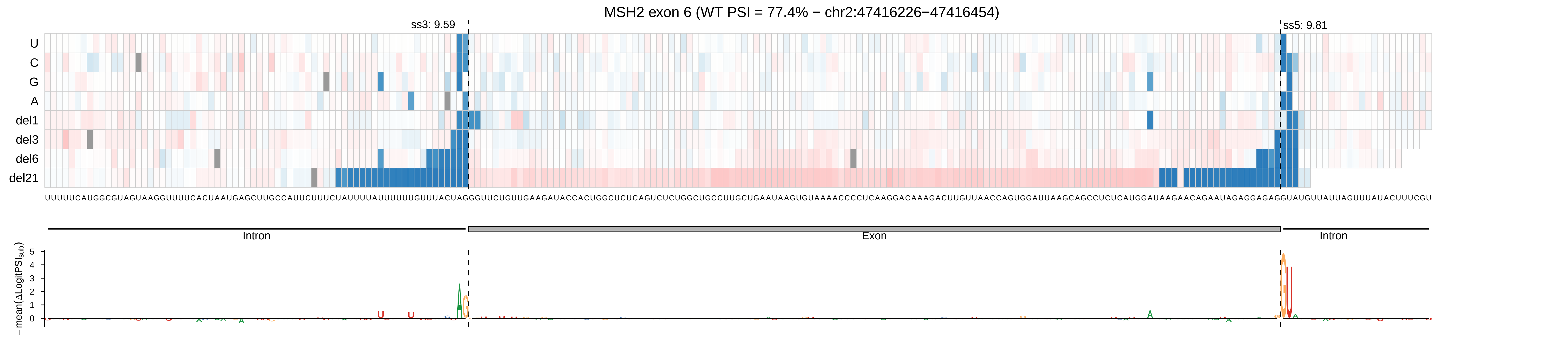

MSH2 exon 6 (WT PSI = 77.4% − chr2:47416226−47416454)
ss3: 9.59
ss5: 9.81
U
C
G
A
del1
del3
del6
del21
Intron
Exon
Intron
− mean(ΔLogitPSI_sub)
5
4
3
2
1
0

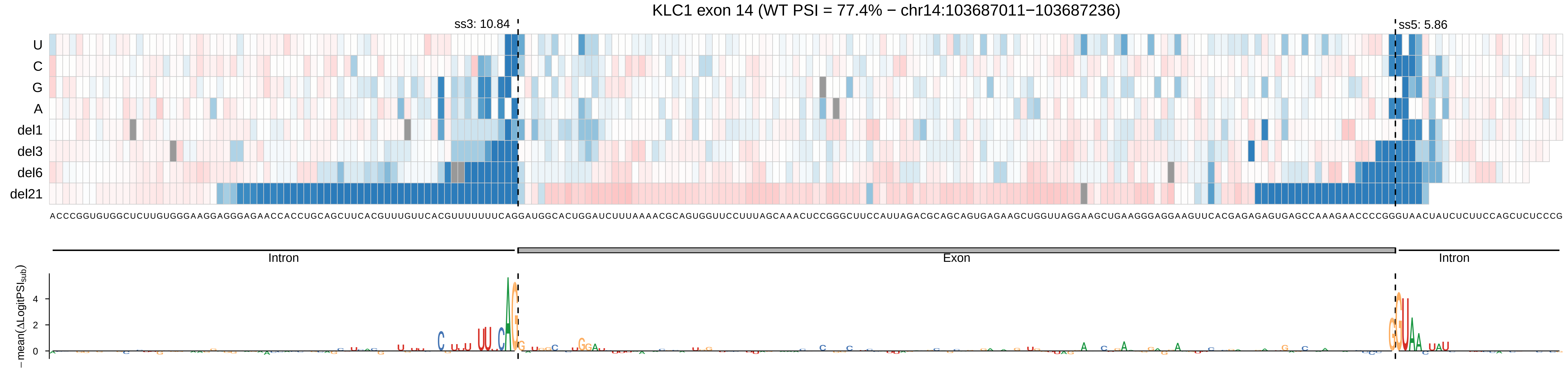
KLC1 exon 14 (WT PSI = 77.4% − chr14:103687011−103687236)
ss3: 10.84
ss5: 5.86
U
C
G
A
del1
del3
del6
del21
ACCCGGUGUGGCUCUUGUGGGAAGGAGGGAGAACCACCUGCAGCUUCACGUUUGUUCACGUUUUUUUCAGGAUGGCACUGGAUCUUUAAAACGCAGUGGUUCCUUUAGCAAACUCCGGGCUUCCAUUAGACGCAGCAGUGAGAAGCUGGUUAGGAAGCUGAAGGGAGGAAGUUCACGAGAGAGUGAGCCAAAGAACCCCGGGUAACUAUCUCUUCCAGCUCUCCCG
Intron
Exon
Intron
−mean(ΔLogitPSIsub)
4
2
0

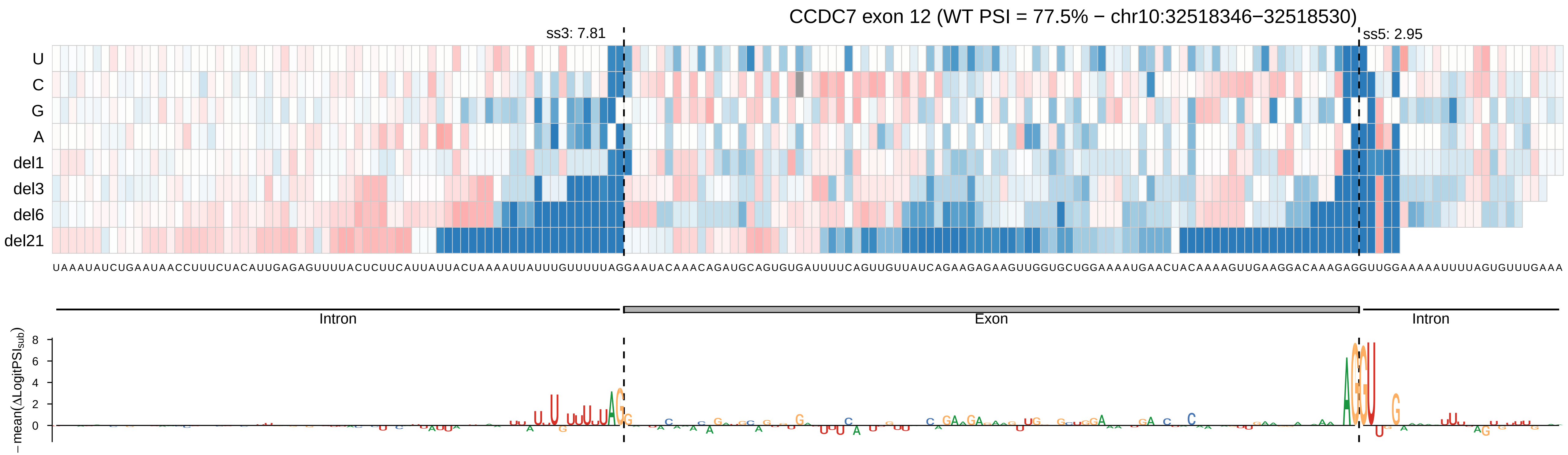
CCDC7 exon 12 (WT PSI = 77.5% − chr10:32518346−32518530)
ss3: 7.81
ss5: 2.95
U
C
G
A
del1
del3
del6
del21
UAAAUAUCUGAAUAACCUUUCUACAUUGAGAGUUUUACUCUUCAUUAUUACUAAAAUUAUUUGUUUUUAGGAAUACAAACAGAUGCAGUGUGAUUUUCAGUUGUUAUCAGAAGAGAAGUUGGUGCUGGAAAAUGAACUACAAAAGUUGAAGGACAAAGAGGUUGGAAAAAUUUUAGUGUUUGAAA
Intron
Exon
Intron
−mean(ΔLogitPSI_sub)
8
6
4
2
0

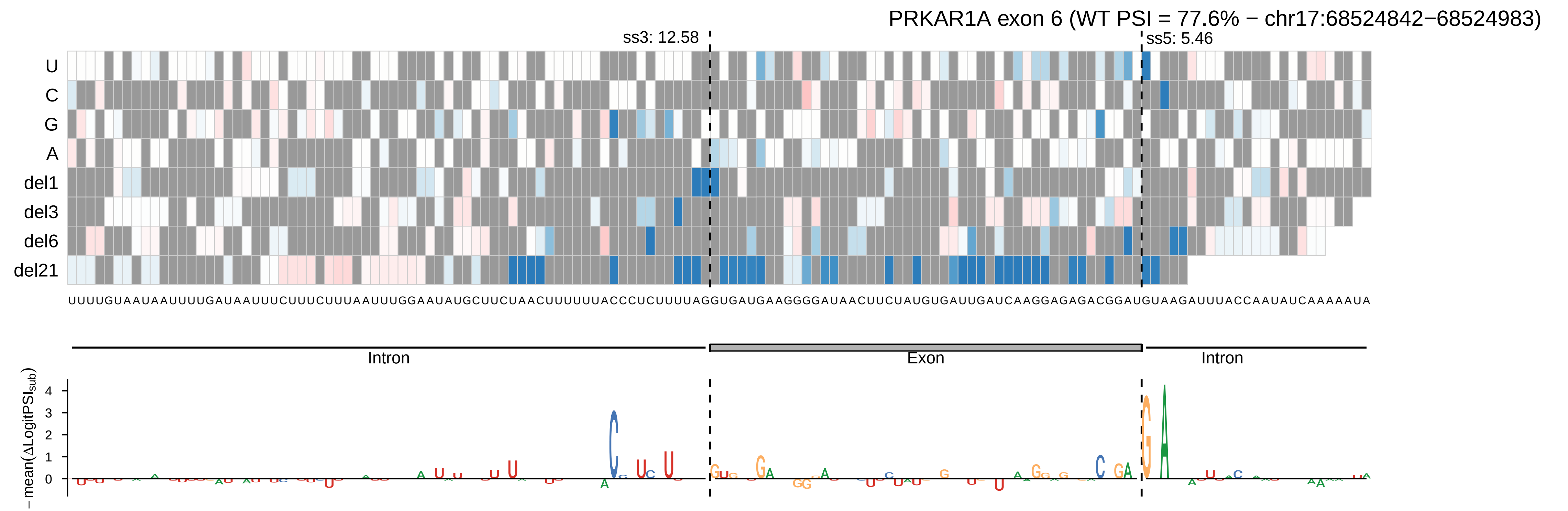

PRKAR1A exon 6 (WT PSI = 77.6% − chr17:68524842−68524983)
ss3: 12.58
ss5: 5.46
U
C
G
A
del1
del3
del6
del21
Intron
Exon
Intron
−mean(ΔLogitPSI_sub)
4
3
2
1
0

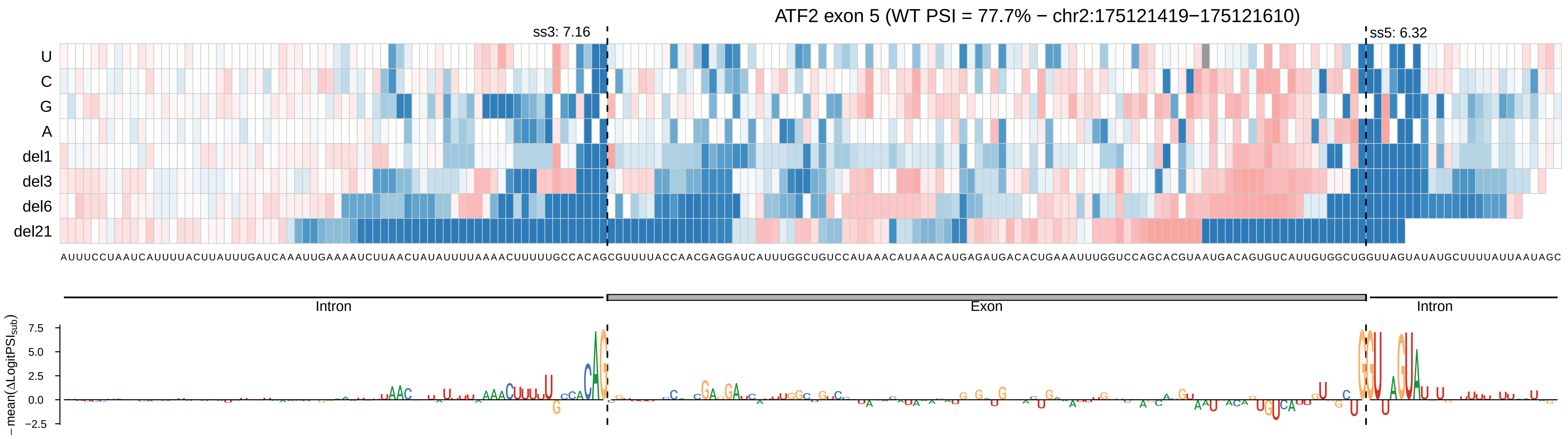
ATF2 exon 5 (WT PSI = 77.7% − chr2:175121419−175121610)
ss3: 7.16
ss5: 6.32
U
C
G
A
del1
del3
del6
del21
AUUUCCUAAUCAUUUUACUUAUUUGAUCAAAUUGAAAAUCUUAACUAUAUUUUAAAACUUUUUGCCACAGCGUUUUACCAACGAGGAUCAUUUGGCUGUCCAUAAACAUAAACAUGAGAUGACACUGAAAUUUGGUCCAGCACGUAAUGACAGUGUCAUUGUGGCUGGUUAGUAUAUGCUUUUAUUAAUAGC
Intron
Exon
Intron
− mean(ΔLogitPSIsub)
7.5
5.0
2.5
0.0
−2.5

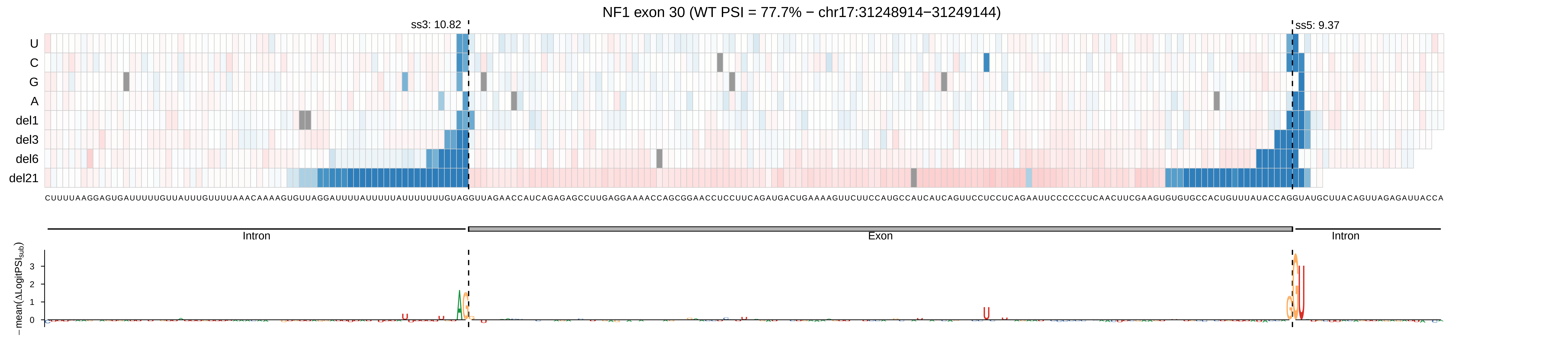

NF1 exon 30 (WT PSI = 77.7% − chr17:31248914−31249144)
ss3: 10.82
ss5: 9.37
U
C
G
A
del1
del3
del6
del21
Intron
Exon
Intron
−mean(ΔLogitPSI_sub)
0
1
2
3

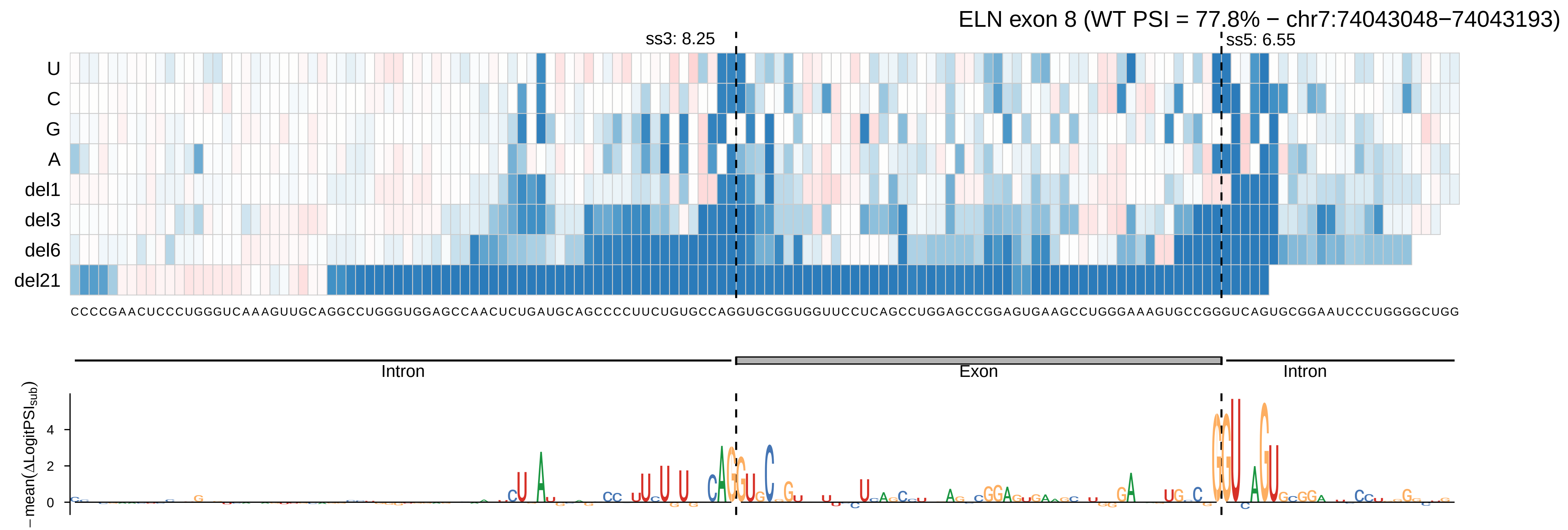

ELN exon 8 (WT PSI = 77.8% − chr7:74043048−74043193)
ss3: 8.25
ss5: 6.55
U
C
G
A
del1
del3
del6
del21
CCCCGAACUCCCUGGGUCAAAGUUGCAGGCCUGGGUGGAGCCAACUCUGAUGCAGCCCCUUCUGUGCCAGGUGCGGUGGUUCCUCAGCCUGGAGCCGGAGUGAAGCCUGGGAAAGUGCCGGGUCAGUGCGGAAUCCCUGGGGCUGG
Intron
Exon
Intron
−mean(ΔLogitPSI_sub)
4
2
0

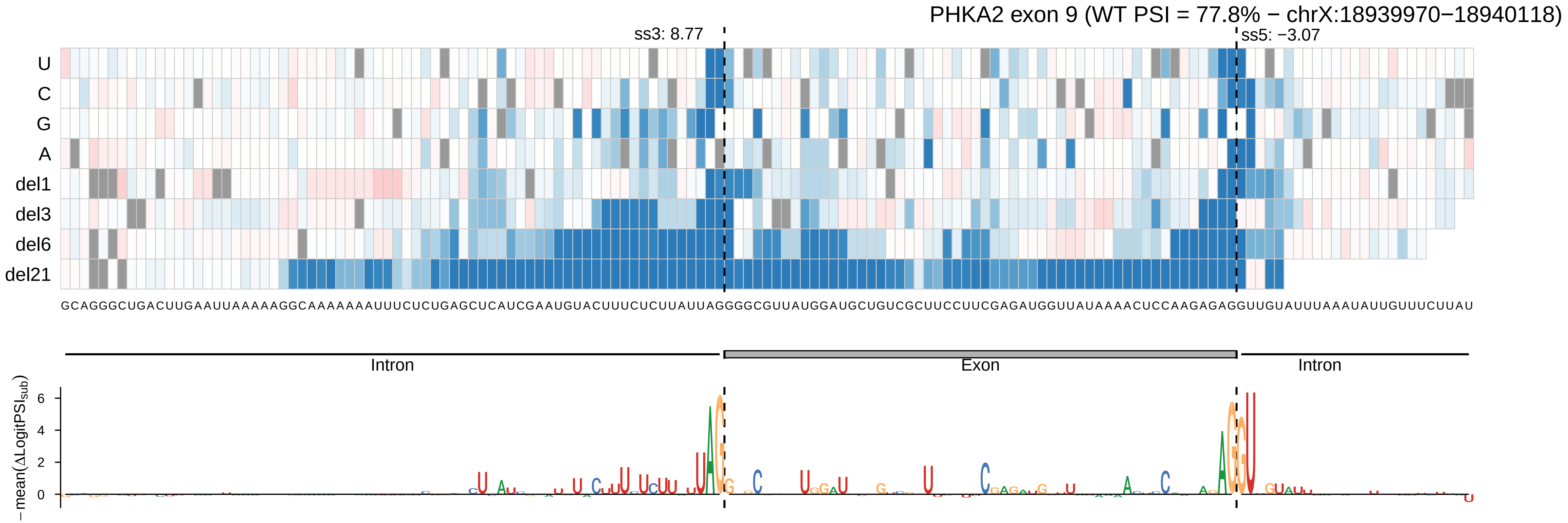
PHKA2 exon 9 (WT PSI = 77.8% − chrX:18939970−18940118)
ss3: 8.77
ss5: −3.07
U
C
G
A
del1
del3
del6
del21
Intron
Exon
Intron
−mean(ΔLogitPSI_sub)
6
4
2
0

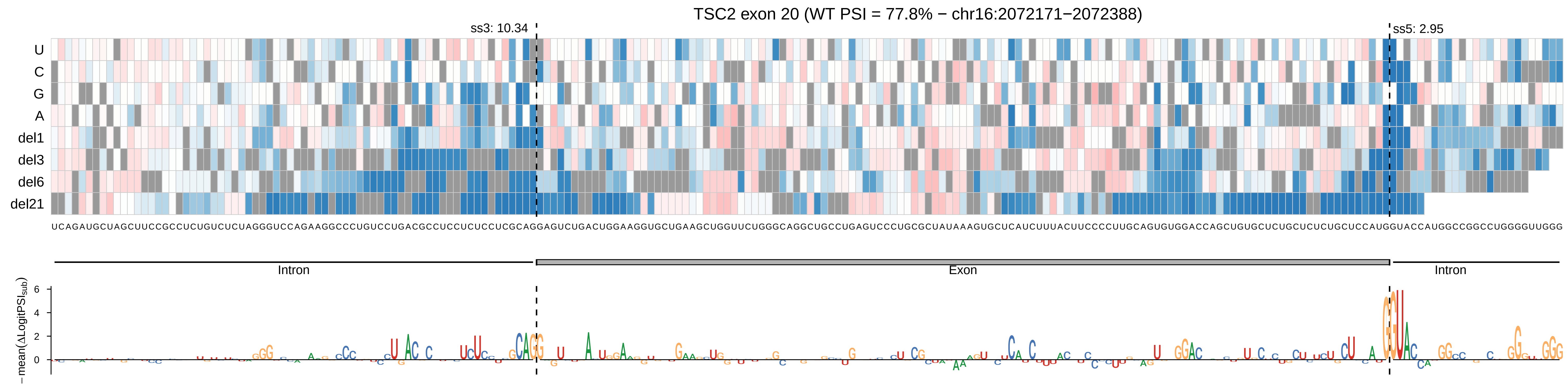

TSC2 exon 20 (WT PSI = 77.8% − chr16:2072171−2072388)
ss3: 10.34
ss5: 2.95
U
C
G
A
del1
del3
del6
del21
UCAGAUGCUAGCUUCCGCCUCUGUCUCUAGGGUCCAGAAGGCCCUGUCCUGACGCCUCCUCUCCUCGCAGGAGUCUGACUGGAAGGUGCUGAAGCUGGUUCUGGGCAGGCUGCCUGAGUCCCUGCGCUAUAAAGUGCUCAUCUUUACUUCCCCUUGCAGUGUGGACCAGCUGUGCUCUGCUCUCUGCUCCAUGGUACCAUGGCCGGCCUGGGGUUGGG
Intron
Exon
Intron
−mean(ΔLogitPSI_sub)
6
4
2
0

CACNA1D exon 43 (WT PSI = 77.9% − chr3:53802077−53802198)

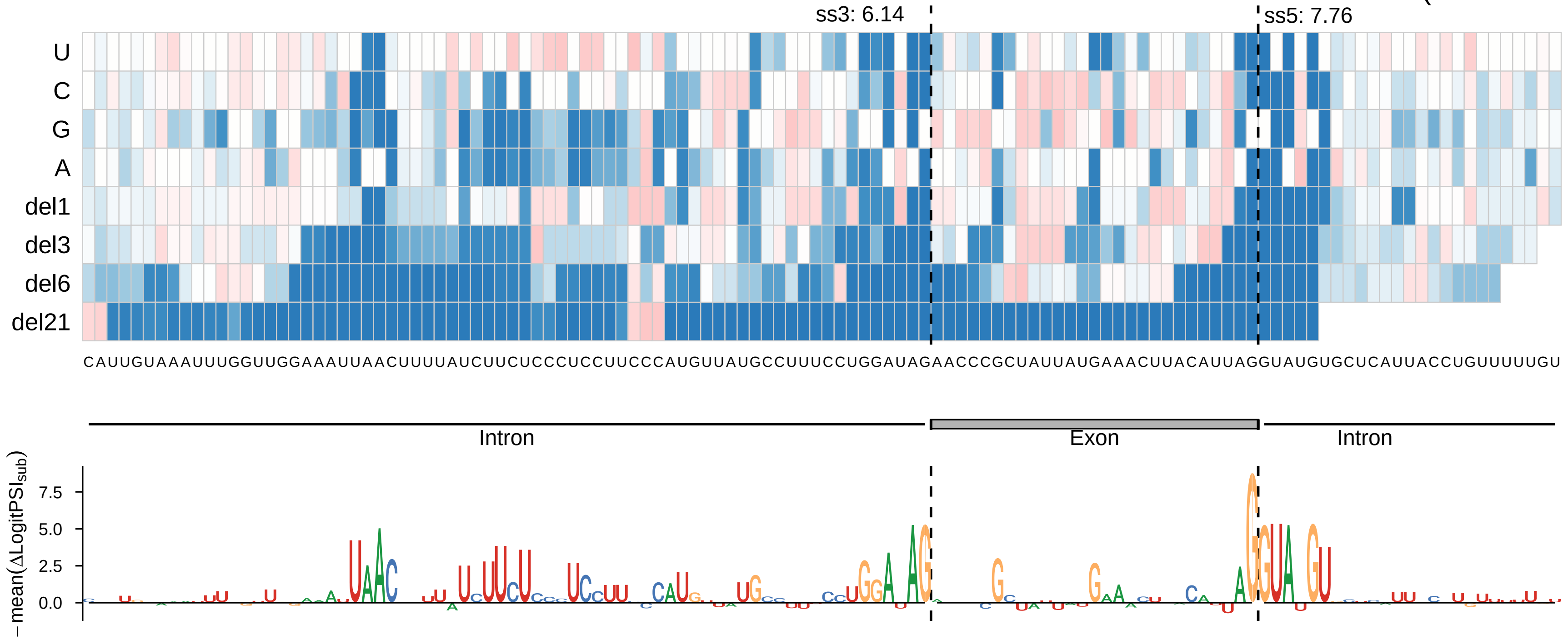

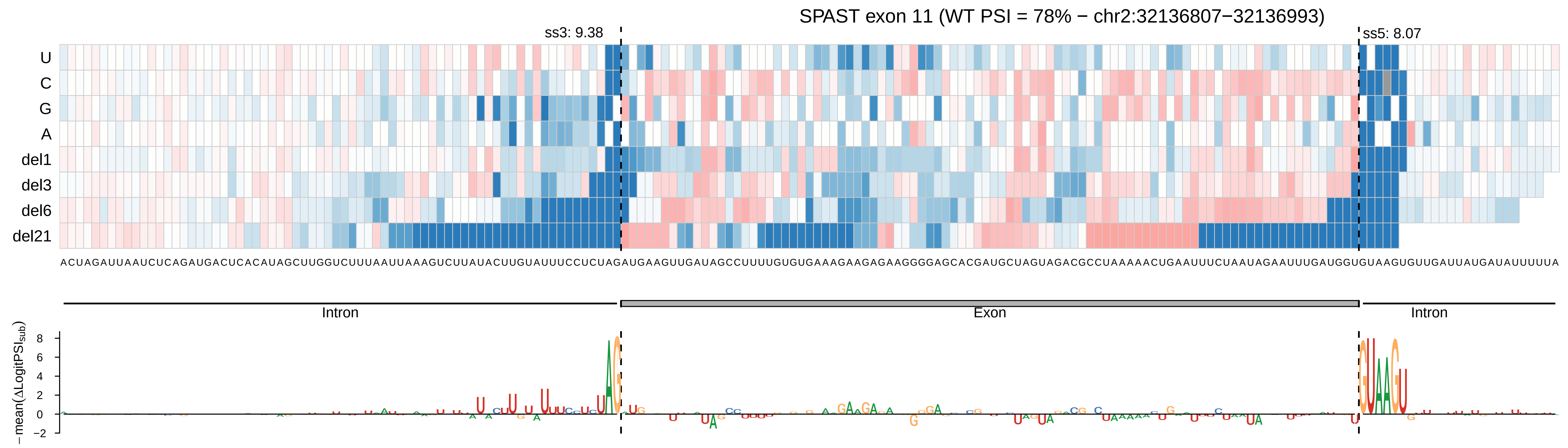

SPAST exon 11 (WT PSI = 78% − chr2:32136807−32136993)
ss3: 9.38
ss5: 8.07
U
C
G
A
del1
del3
del6
del21
ACUAGAUUAAUCUCAGAUGACUCACAUAGCUUGGUCUUUAAUUAAAGUCUUAUACUUGUAUUUCCUCUAGAUGAAGUUGAUAGCCUUUUGUGUGAAAGAAGAGAAGGGGAGCACGAUGCUAGUAGACGCCUAAAAACUGAAUUUCUAAUAGAAUUUGAUGGUGUAAGUGUUGAUUAUGAUAUUUUUA
Intron
Exon
Intron
− mean(ΔLogitPSI_sub)
8
6
4
2
0
−2

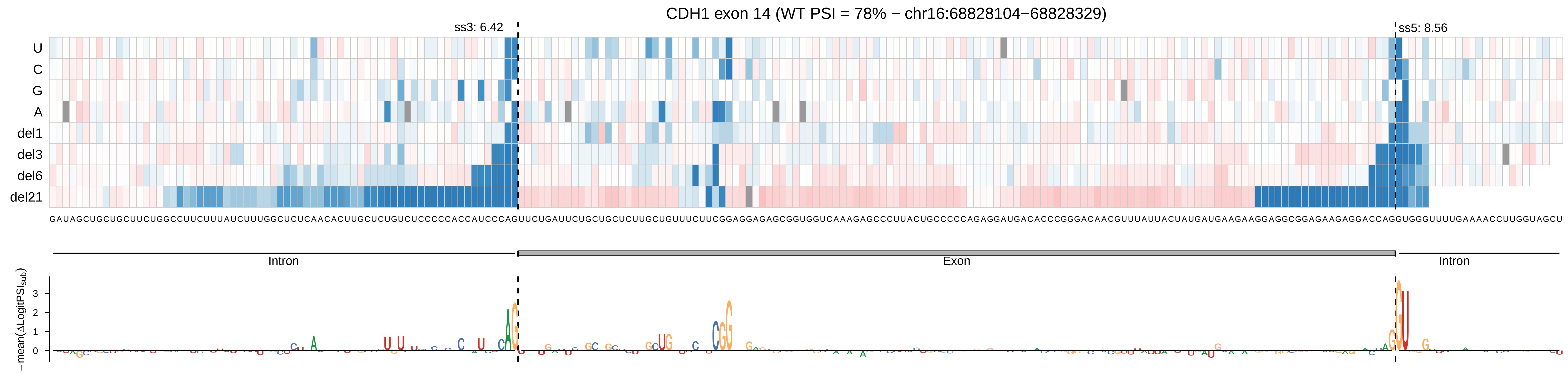
CDH1 exon 14 (WT PSI = 78% − chr16:68828104−68828329)
ss3: 6.42
ss5: 8.56
U
C
G
A
del1
del3
del6
del21
Intron
Exon
Intron
−mean(ΔLogitPSI_sub)
0
1
2
3

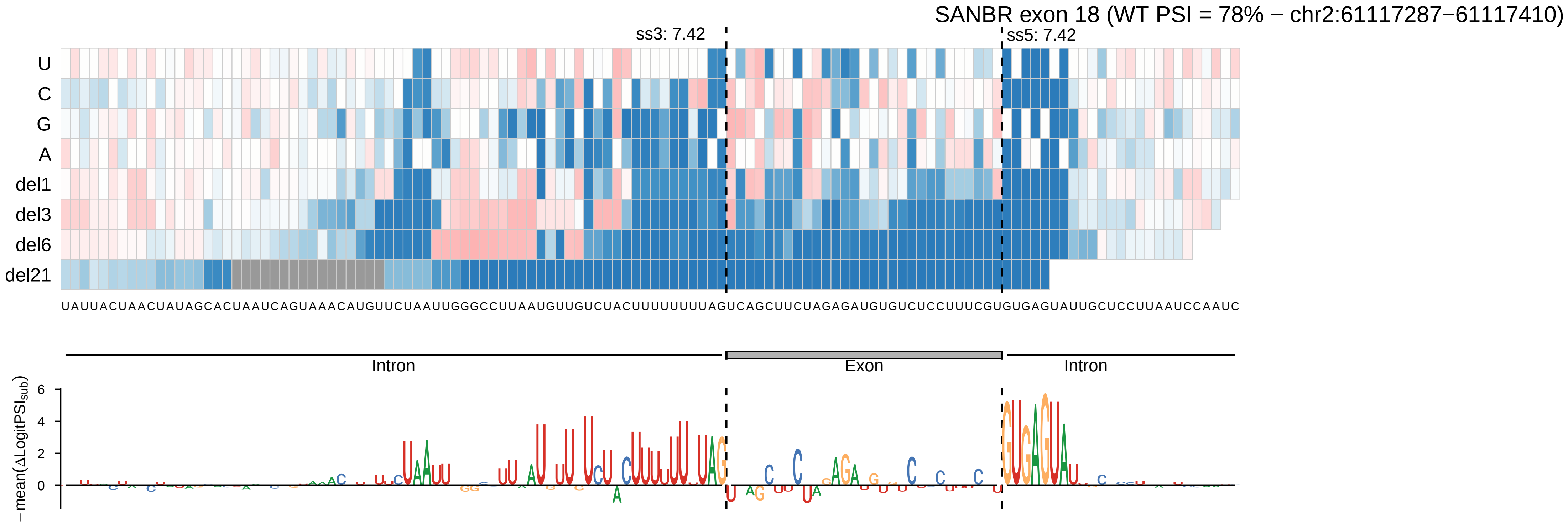

SANBR exon 18 (WT PSI = 78% − chr2:61117287−61117410)
ss3: 7.42
ss5: 7.42
U
C
G
A
del1
del3
del6
del21
UAUUACUAACUAUAGCACUAAUCAGUAAACAUGUUCUAAUUGGGCCUUAAUGUUGUCUACUUUUUUUUAGUCAGCUUCUAGAGAUGUGUCUCCUUUCGUGUGAGUAUUGCUCCUUAAUCCAAUC
Intron
Exon
Intron
− mean(ΔLogitPSI_sub)
6
4
2
0

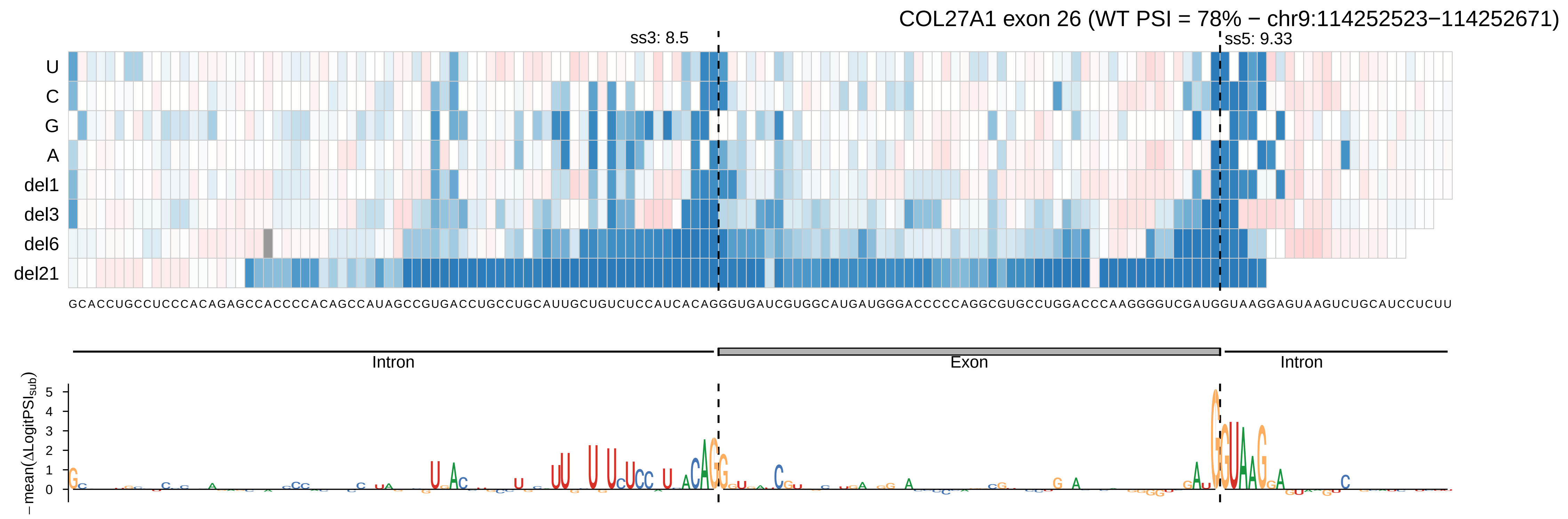
COL27A1 exon 26 (WT PSI = 78% − chr9:114252523−114252671)
ss3: 8.5
ss5: 9.33
U
C
G
A
del1
del3
del6
del21
GCACCUGCCUCCCACAGAGCCACCCCACAGCCAUAGCCGUGACCUGCCUGCAUUGCUGUCUCCAUCACAGGGUGAUCGUGGCAUGAUGGGACCCCCAGGCGUGCCUGGACCCAAGGGGUCGAUGGUAAGGAGUAAGUCUGCAUCCUCUU
Intron
Exon
Intron
−mean(ΔLogitPSI_sub)
5
4
3
2
1
0

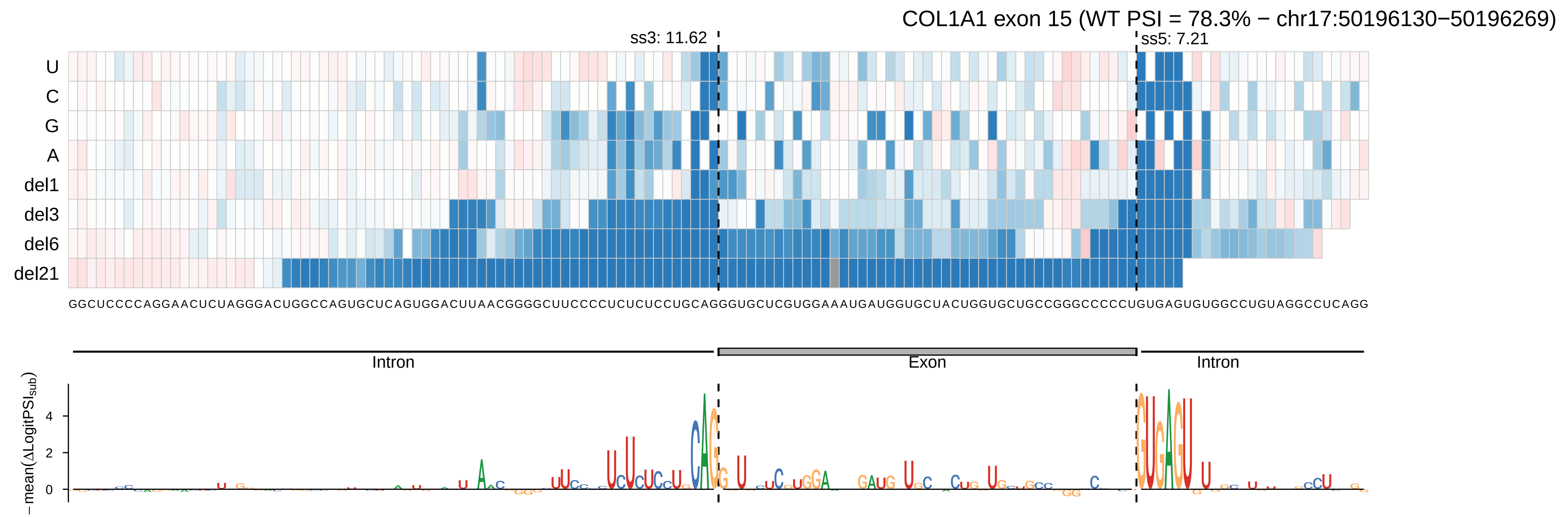
COL1A1 exon 15 (WT PSI = 78.3% − chr17:50196130−50196269)
ss3: 11.62
ss5: 7.21
U
C
G
A
del1
del3
del6
del21
GGCUCCCCAGGAACUCUAGGGACUGGCCAGUGCUCAGUGGACUUAACGGGGCUUCCCCUCUCUCCUGCAGGGUGCUCGUGGAAAUGAUGGUGCUACUGGUGCUGCCGGGCCCCCUGUGAGUGUGGCCUGUAGGCCUCAGG
Intron
Exon
Intron
− mean(ΔLogitPSI_sub)
0
2
4

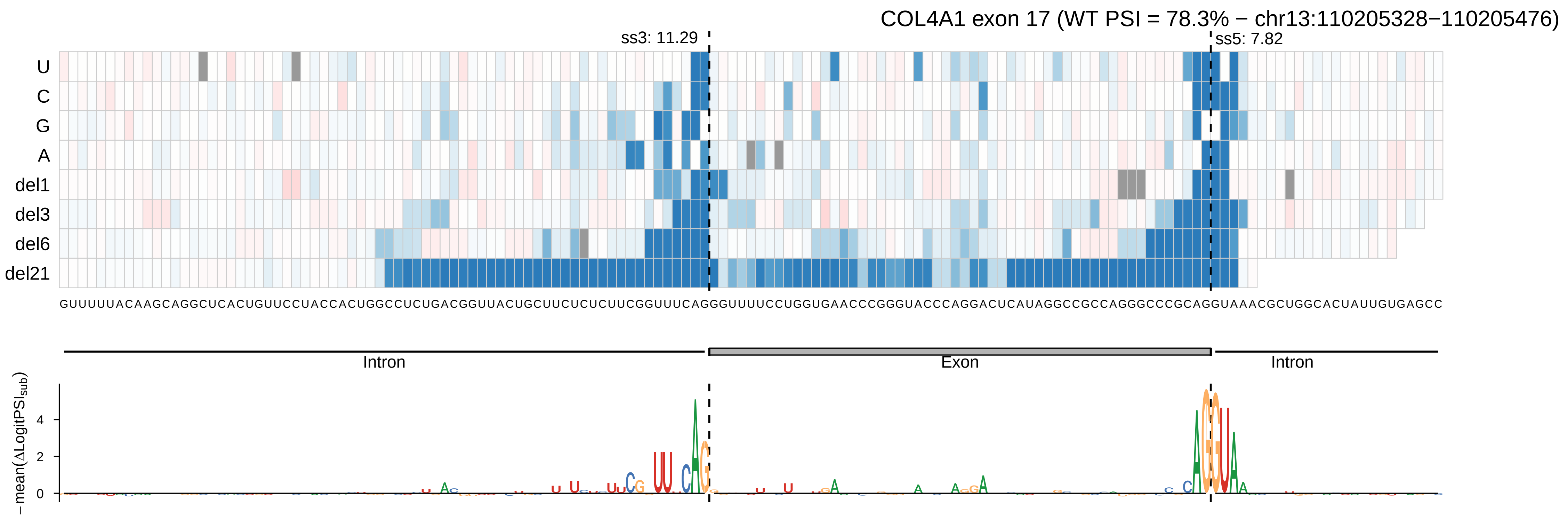
COL4A1 exon 17 (WT PSI = 78.3% − chr13:110205328−110205476)
ss3: 11.29
ss5: 7.82
U
C
G
A
del1
del3
del6
del21
GUUUUUACAAGCAGGCUCACUGUUCCUACCACUGGCCUCUGACGGUUACUGCUUCUCUCUUCGGUUUCAGGGUUUUCCUGGUGAACCCGGGUACCCAGGACUCAUAGGCCGCCAGGGCCCGCAGGUAAACGCUGGCACUAUUGUGAGCC
Intron
Exon
Intron
−mean(ΔLogitPSI_sub)
0
2
4

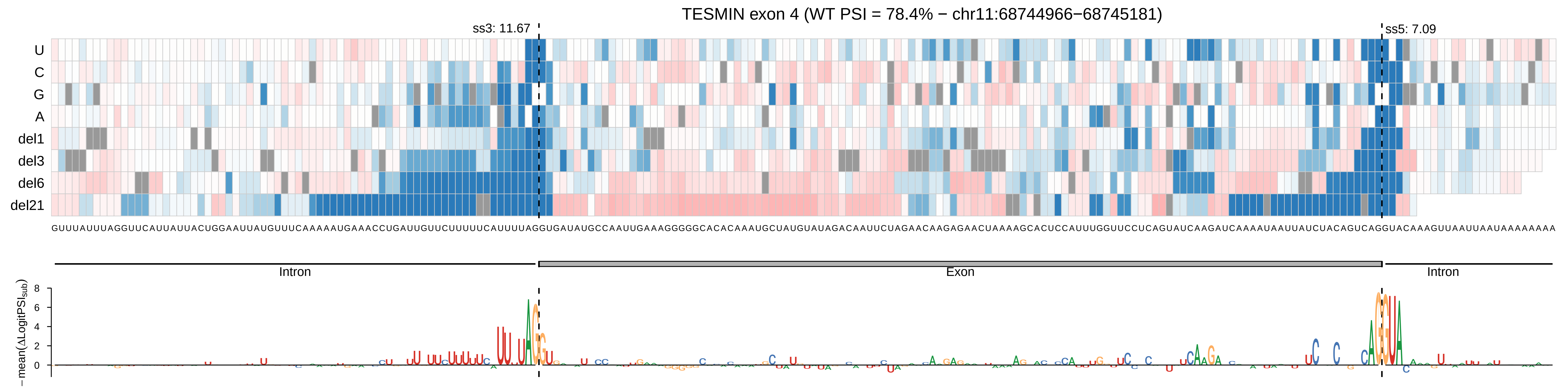

TESMIN exon 4 (WT PSI = 78.4% − chr11:68744966−68745181)
ss3: 11.67
ss5: 7.09
U
C
G
A
del1
del3
del6
del21
GUUUAUUUAGGUUCAUUAUUACUGGAAUUAUGUUUCAAAAAUGAAACCUGAUUGUUCUUUUUCAUUUUAGGUGAUAUGCCAAUUGAAAGGGGGCACACAAAUGCUAUGUAUAGACAAUUCUAGAACAAGAGAACUAAAAGCACUCCAUUUGGUUCCUCAGUAUCAAGAUCAAAAUAAUUAUCUACAGUCAGGUACAAAGUUAAUUAAUAAAAAAAA
Intron
Exon
Intron
−mean(ΔLogitPSI_sub)
8
6
4
2
0

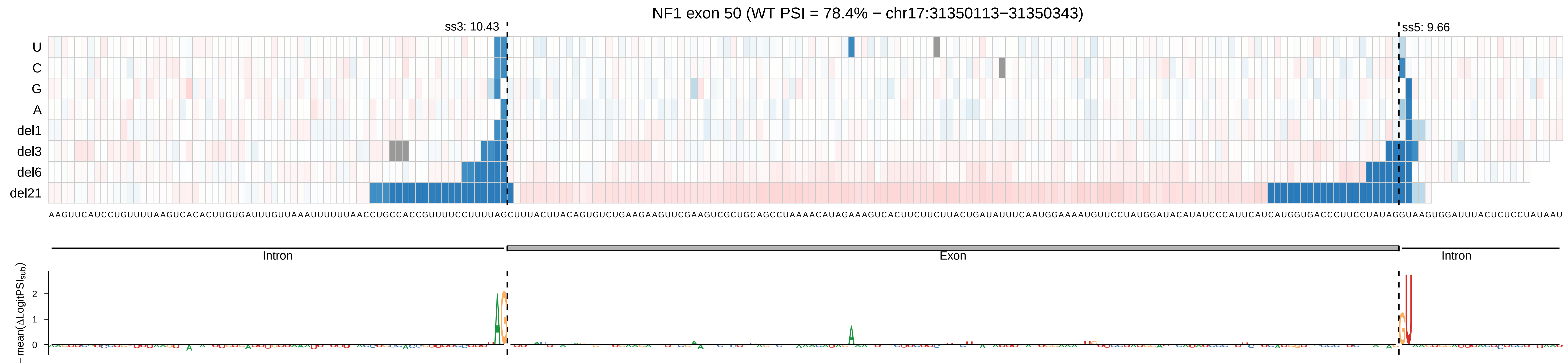

NF1 exon 50 (WT PSI = 78.4% − chr17:31350113−31350343)
ss3: 10.43
ss5: 9.66
U
C
G
A
del1
del3
del6
del21
AAGUUCAUCCUGUUUUAAGUCACACUUGUGAUUUGUUAAAUUUUUUAACCUGCCACCGUUUUCCUUUUAGCUUUACUUACAGUGUCUGAAGAAGUUCGAAGUCGCUGCAGCCUAAAACAUAGAAAGUCACUUCUUCUUACUGAUAUUUCAAUGGAAAAUGUUCCUAUGGAUACAUAUCCCAUUCAUCAUGGUGACCCUUCCUAUAGGUAAGUGGAUUUACUCUCCUAUAAU
Intron
Exon
Intron
− mean(ΔLogitPSI_sub)
2
1
0

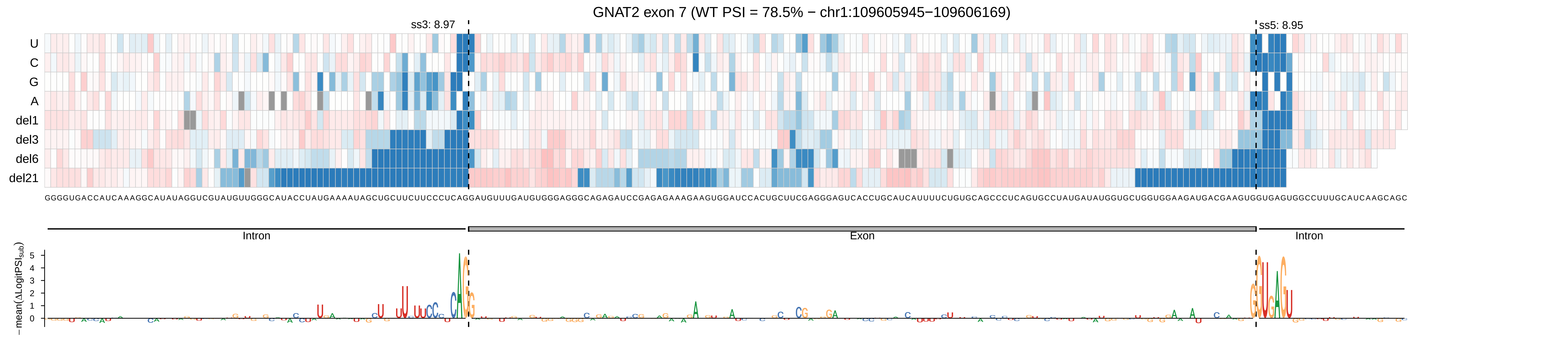
GNAT2 exon 7 (WT PSI = 78.5% − chr1:109605945−109606169)
ss3: 8.97
ss5: 8.95
U
C
G
A
del1
del3
del6
del21
Intron
Exon
Intron
− mean(ΔLogitPSI_sub)
5
4
3
2
1
0

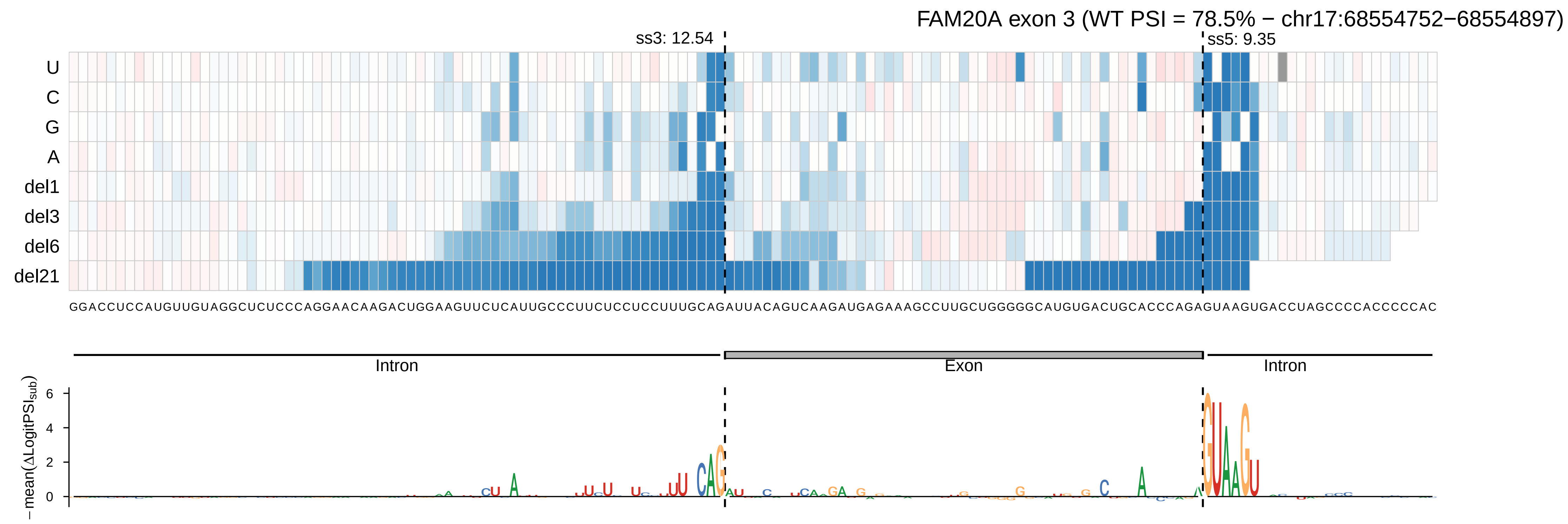

FAM20A exon 3 (WT PSI = 78.5% − chr17:68554752−68554897)
ss3: 12.54
ss5: 9.35
U
C
G
A
del1
del3
del6
del21
GGACCUCCAUGUUGUAGGCUCUCCCAGGAACAAGACUGGAAGUUCUCAUUGCCCUUCUCCUCCUUUGCAGAUUACAGUCAAGAUGAGAAAGCCUUGCUGGGGGCAUGUGACUGCACCCAGAGUAAGUGACCUAGCCCCACCCCCAC
Intron
Exon
Intron
−mean(ΔLogitPSIsub)
6
4
2
0

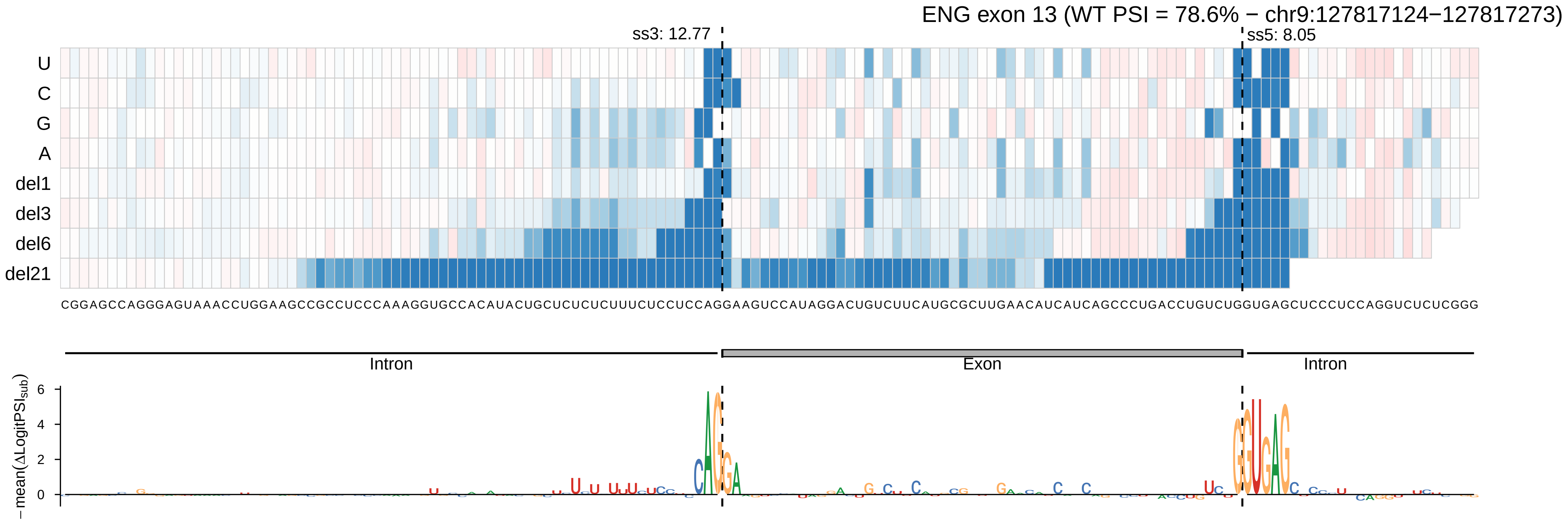

ENG exon 13 (WT PSI = 78.6% − chr9:127817124−127817273)
ss3: 12.77
ss5: 8.05
U
C
G
A
del1
del3
del6
del21
CGGAGCCAGGGAGUAAACCUGGAAGCCGCCUCCCAAAGGUGCCACAUACUGCUCUCUCUUUCUCCUCCAGGAAGUCCAUAGGACUGUCUUCAUGCGCUUGAACAUCAUCAGCCCUGACCUGUCUGGUGAGCUCCCUCCAGGUCUCUCGGG
Intron
Exon
Intron
− mean(ΔLogitPSIsub)
6
4
2
0

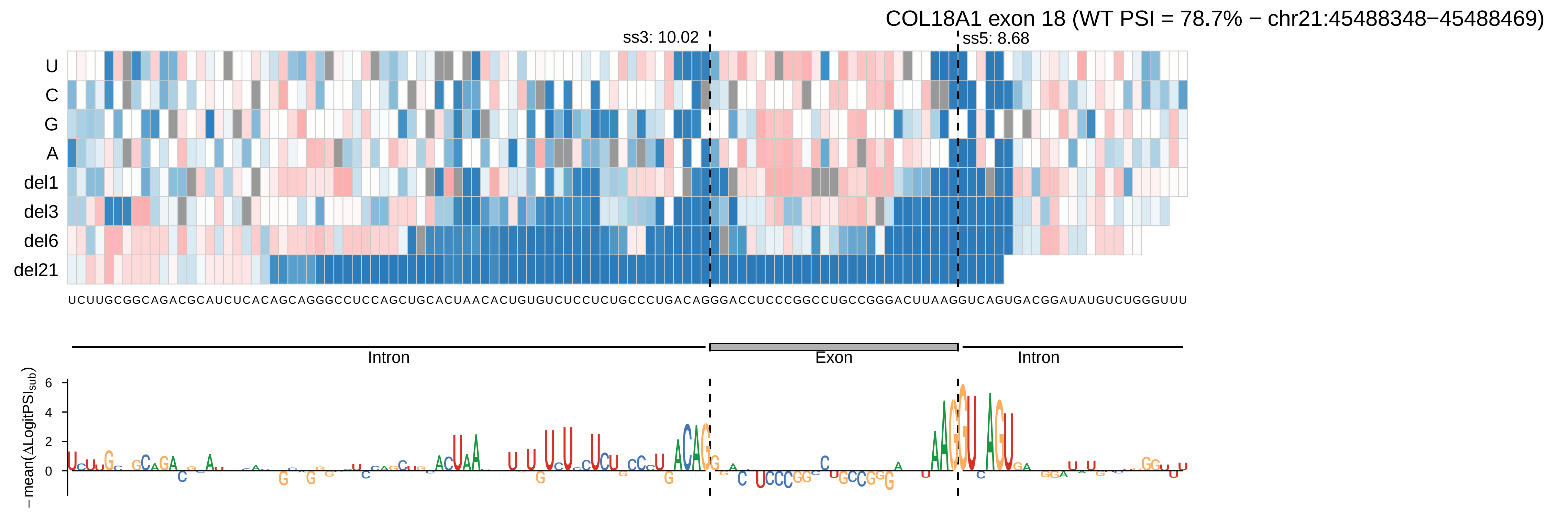

COL18A1 exon 18 (WT PSI = 78.7% − chr21:45488348−45488469)
ss3: 10.02
ss5: 8.68
U
C
G
A
del1
del3
del6
del21
Intron
Exon
Intron
−mean(ΔLogitPSI_sub)

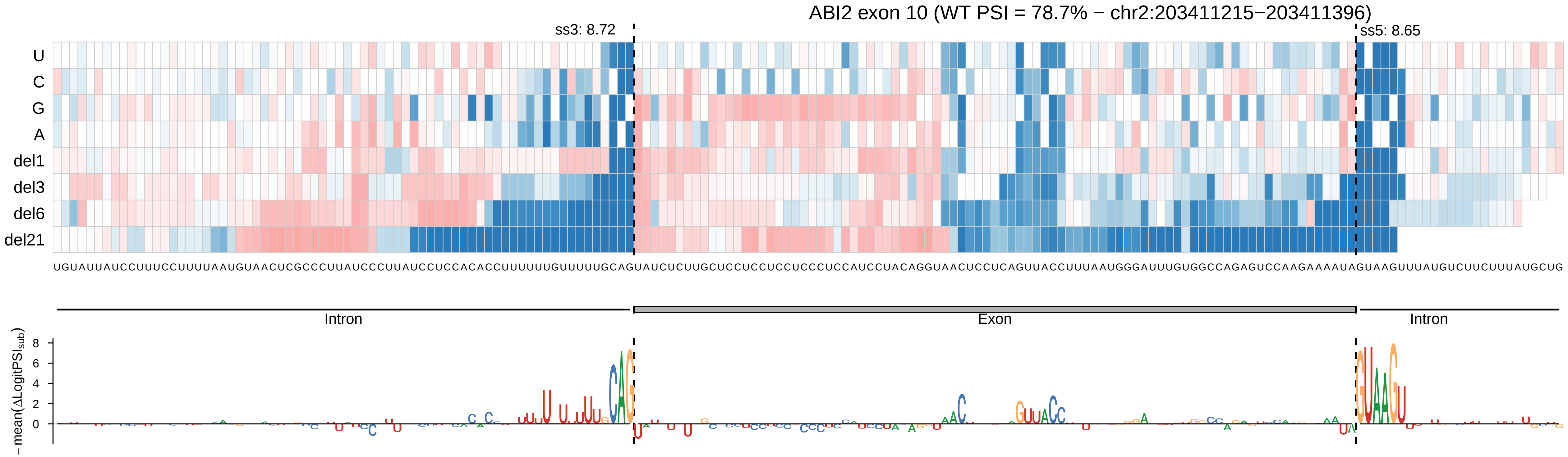
ABI2 exon 10 (WT PSI = 78.7% − chr2:203411215−203411396)
ss3: 8.72
ss5: 8.65
U
C
G
A
del1
del3
del6
del21
UGUAUUAUCCUUUCCUUUUAAUGUAACUCGCCCUUAUCCCUUAUCCUCCACACCUUUUUUGUUUUUGCAGUAUCUCUUGCUCCUCCUCCUCCCUCCAUCCUACAGGUAACUCCUCAGUUACCUUUAAUGGGAUUUGUGGCCAGAGUCCAAGAAAAUAGUAAGUUUAUGUCUUCUUUAUGCUG
Intron
Exon
Intron
− mean(ΔLogitPSI_sub)
8
6
4
2
0

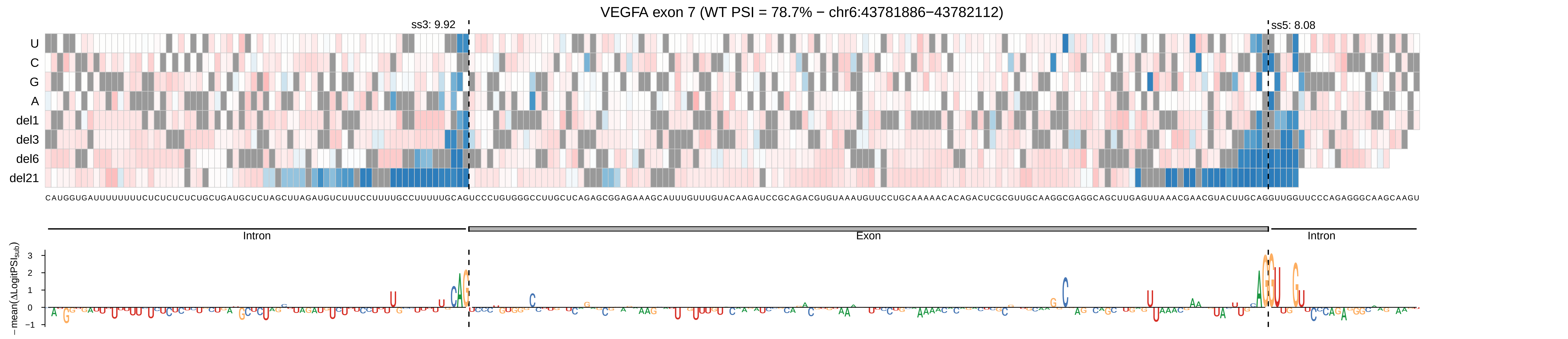

VEGFA exon 7 (WT PSI = 78.7% − chr6:43781886−43782112)
ss3: 9.92
ss5: 8.08
U
C
G
A
del1
del3
del6
del21
CAUGGUGAUUUUUUUUCUCUCUCUCUGCUGAUGCUCUAGCUUAGAUGUCUUUCCUUUUGCCUUUUUGCAGUCCCUGUGGGCCUUGCUCAGAGCGGAGAAAGCAUUUGUUUGUACAAGAUCCGCAGACGUGUAAAUGUUCCUGCAAAAACACAGACUCGCGUUGCAAGGCGAGGCAGCUUGAGUUAAACGAACGUACUUGCAGGUUGGUUCCCAGAGGGCAAGCAAGU
Intron
Exon
Intron
−mean(ΔLogitPSI_sub)
3
2
1
0
−1

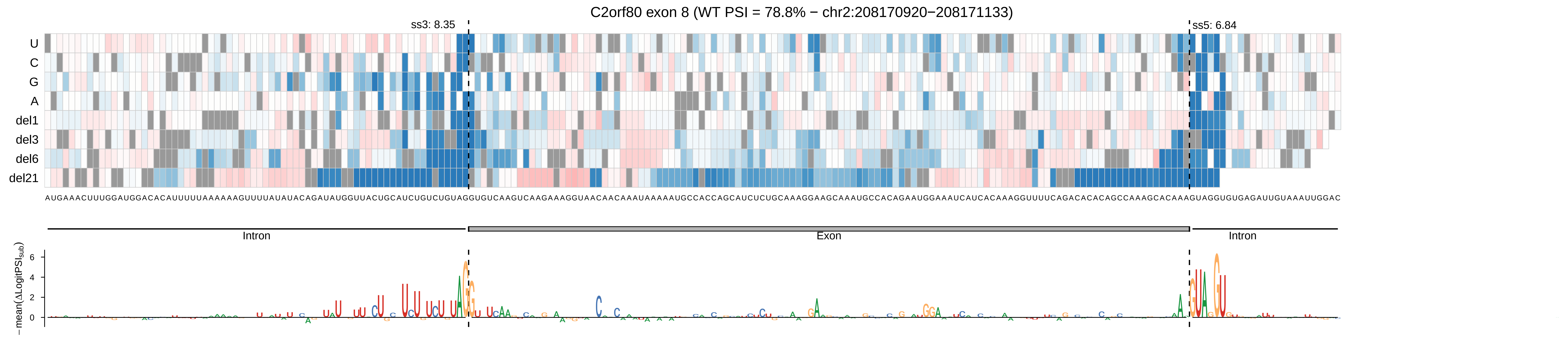
C2orf80 exon 8 (WT PSI = 78.8% − chr2:208170920−208171133)
ss3: 8.35
ss5: 6.84
U
C
G
A
del1
del3
del6
del21
AUGAAACUUUGGAUGGACACAUUUUUAAAAAAGUUUUAUAUACAGAUAUGGUUACUGCAUCUGUCUGUAGGUGUCAAGUCAAGAAAGGUAACAACAAAUAAAAAUGCCACCAGCAUCUCUGCAAAGGAAGCAAAUGCCACAGAAUGGAAAUCAUCACAAAGGUUUUCAGACACACAGCCAAAGCACAAAGUAGGUGUGAGAUUGUAAAUUGGAC
Intron
Exon
Intron
−mean(ΔLogitPSI_sub)
6
4
2
0

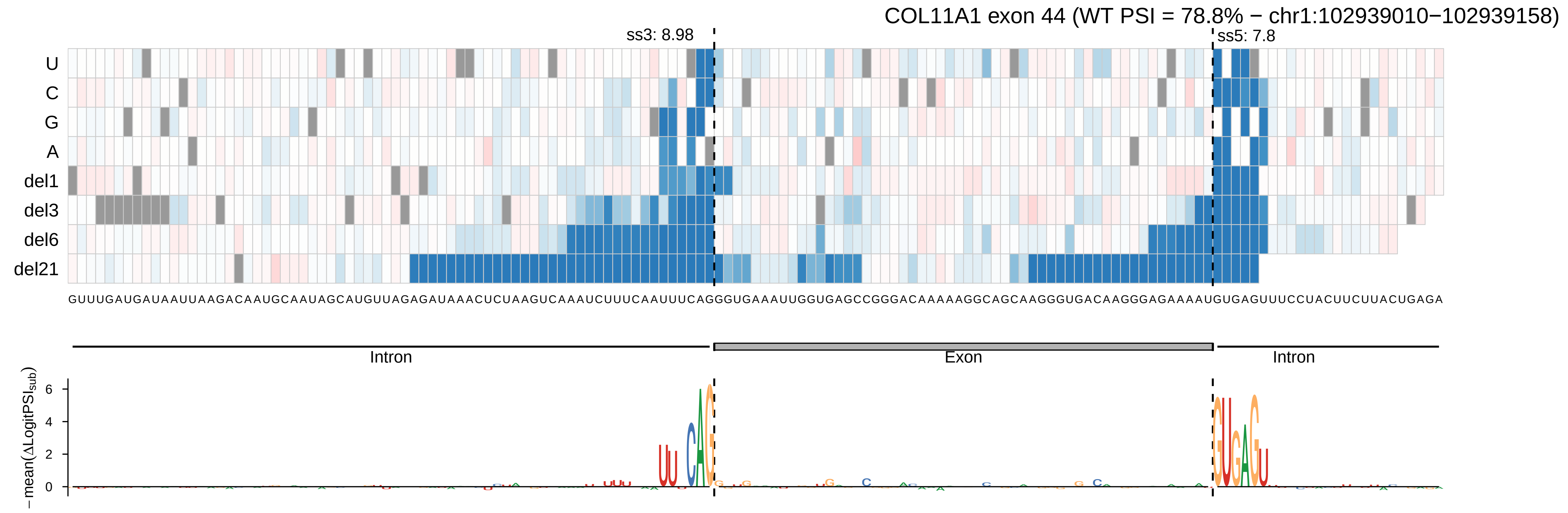

COL11A1 exon 44 (WT PSI = 78.8% − chr1:102939010−102939158)
ss3: 8.98
ss5: 7.8
U
C
G
A
del1
del3
del6
del21
GUUUGAUGAUAAUUAAGACAAUGCAAUAGCAUGUUAGAGAUAAACUCUAAGUCAAAUCUUUCAAUUUCAGGGUGAAAUUGGUGAGCCGGGACAAAAAGGCAGCAAGGGUGACAAGGGAGAAAAUGUGAGUUUCCUACUUCUUACUGAGA
Intron
Exon
Intron
− mean(ΔLogitPSI_sub)
6
4
2
0

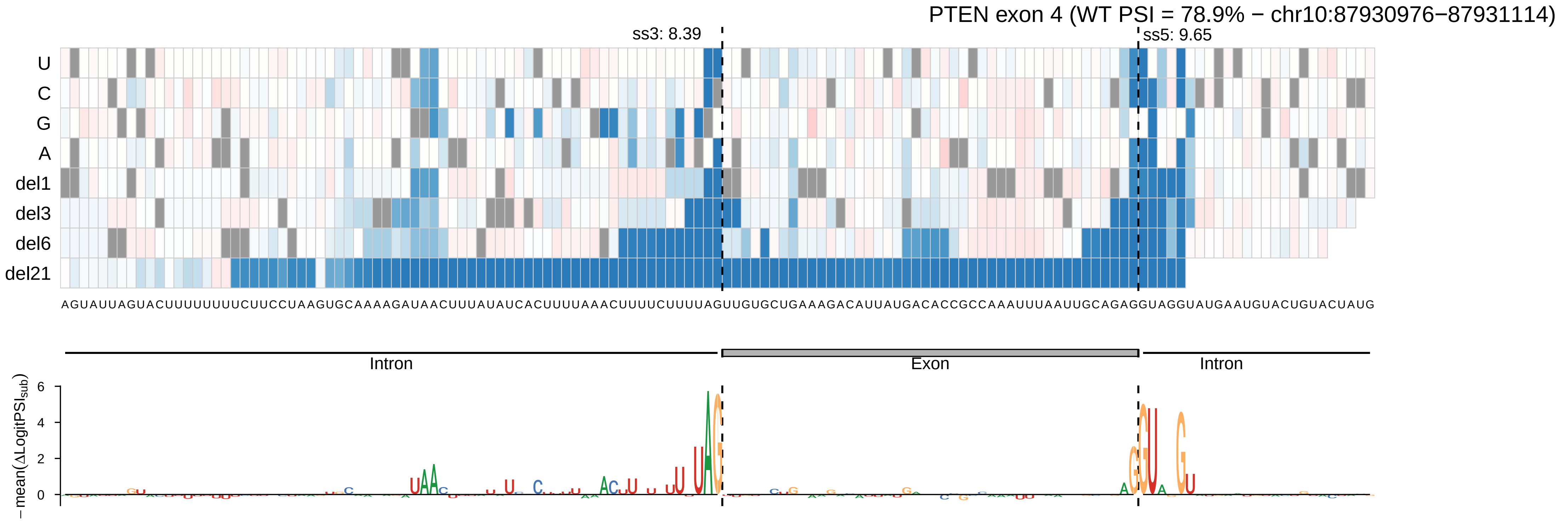
PTEN exon 4 (WT PSI = 78.9% − chr10:87930976−87931114)
ss3: 8.39
ss5: 9.65
U
C
G
A
del1
del3
del6
del21
AGUAUUAGUACUUUUUUUUCUUCCUAAGUGCAAAAGAUAACUUUAUAUCACUUUUAAACUUUUCUUUUAGUUGUGCUGAAAGACAUUAUGACACCGCCAAAUUUAAUUGCAGAGGUAGGUAUGAAUGUACUGUACUAUG
Intron
Exon
Intron
−mean(ΔLogitPSI_sub)
6
4
2
0

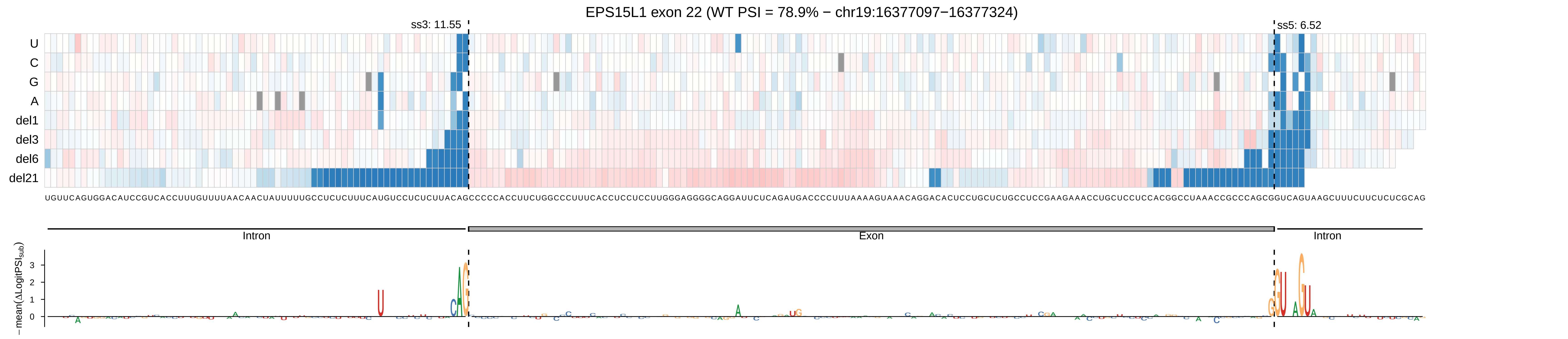

EPS15L1 exon 22 (WT PSI = 78.9% − chr19:16377097−16377324)
ss3: 11.55
ss5: 6.52
U
C
G
A
del1
del3
del6
del21
Intron
Exon
Intron
−mean(ΔLogitPSI_sub)
3
2
1
0

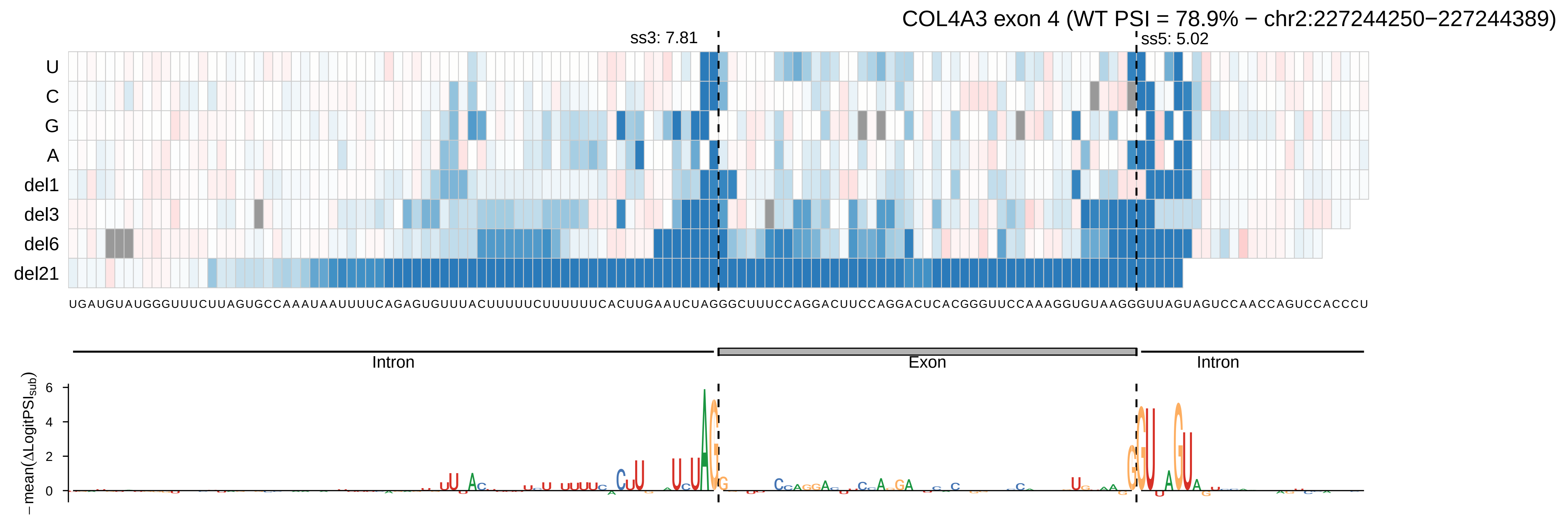

COL4A3 exon 4 (WT PSI = 78.9% − chr2:227244250−227244389)
ss3: 7.81
ss5: 5.02
U
C
G
A
del1
del3
del6
del21
UGAUGUAUGGGUUUCUUAGUGCCAAAUAAUUUUCAGAGUGUUUACUUUUUCUUUUUUCACUUGAAUCUAGGGCUUUCCAGGACUUCCAGGACUCACGGGUUCCAAAGGUGUAAGGGUUAGUAGUCCAACCAGUCCACCCU
Intron
Exon
Intron
−mean(ΔLogitPSI_sub)
0
2
4
6

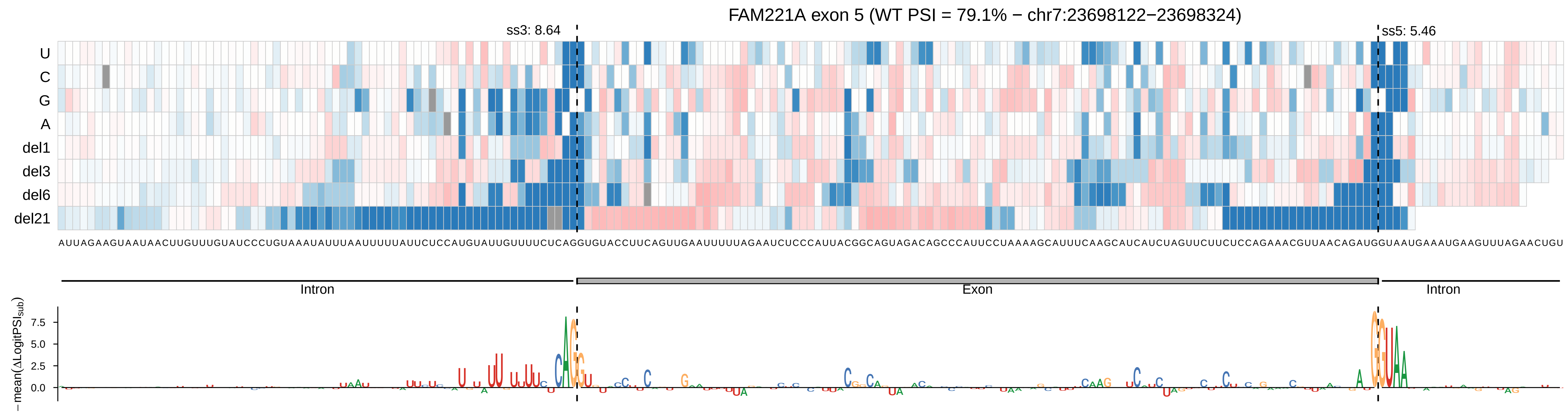

FAM221A exon 5 (WT PSI = 79.1% − chr7:23698122−23698324)
ss3: 8.64
ss5: 5.46
U
C
G
A
del1
del3
del6
del21
AUUAGAAGUAAUAACUUGUUUGUAUCCCUGUAAAUAUUUAAUUUUUAUUCUCCAUGUAUUGUUUUCUCAGGUGUACCUUCAGUUGAAUUUUUAGAAUCUCCCAUUACGGCAGUAGACAGCCCAUUCCUAAAAGCAUUUCAAGCAUCAUCUAGUUCUUCUCCAGAAACGUUAACAGAUGGUAAUGAAAUGAAGUUUAGAACUGU
Intron
Exon
Intron
−mean(ΔLogitPSI_sub)
7.5
5.0
2.5
0.0

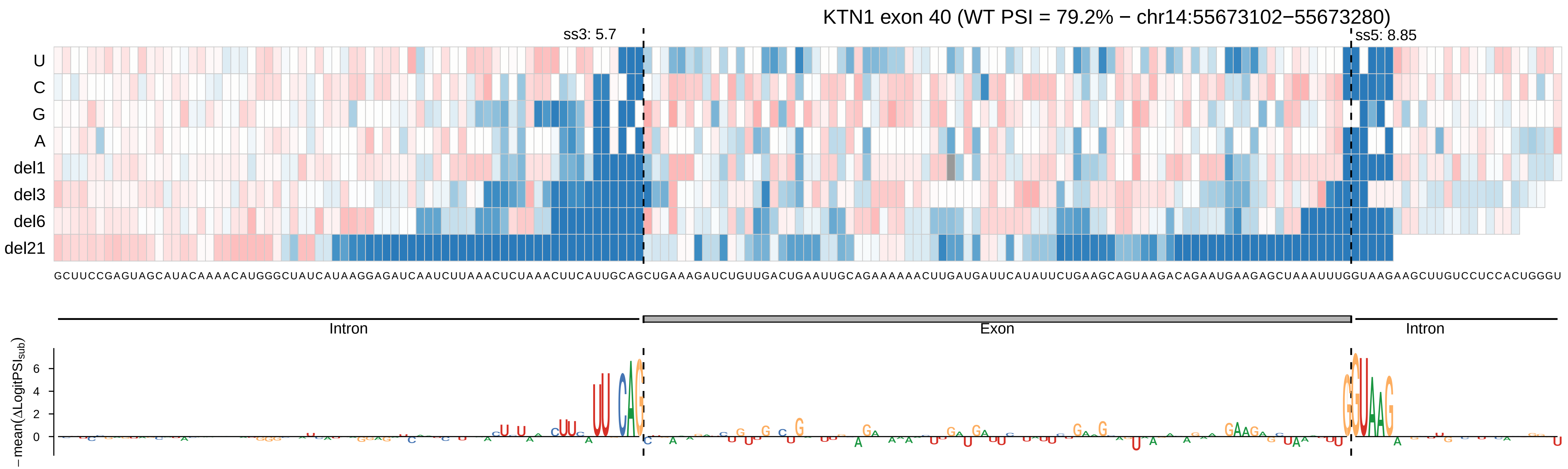

KTN1 exon 40 (WT PSI = 79.2% − chr14:55673102−55673280)
ss3: 5.7
ss5: 8.85
U
C
G
A
del1
del3
del6
del21
Intron
Exon
Intron
−mean(ΔLogitPSI_sub)
6
4
2
0

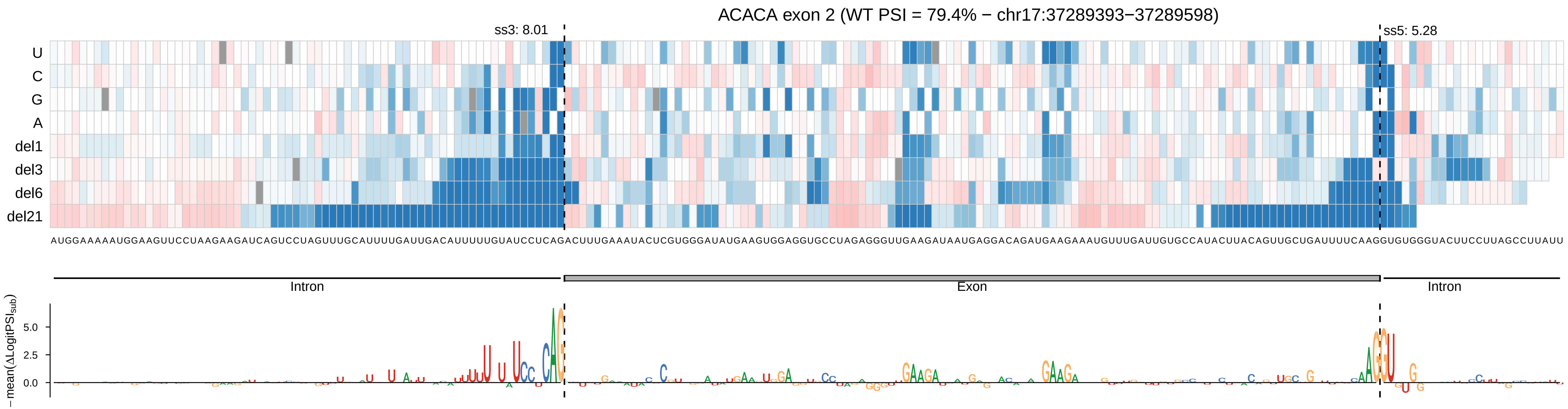
ACACA exon 2 (WT PSI = 79.4% − chr17:37289393−37289598)
ss3: 8.01
ss5: 5.28
U
C
G
A
del1
del3
del6
del21
AUGGAAAAAUGGAAGUUCCUAAGAAGAUCAGUCCUAGUUUGCAUUUUGAUUGACAUUUUUGUAUCCUCAGACUUUGAAAUACUCGUGGGAUAUGAAGUGGAGGUGCCUAGAGGGUUGAAGAUAAUGAGGACAGAUGAAGAAAUGUUUGAUUGUGCCAUACUUACAGUUGCUGAUUUUCAAGGUGUGGGUACUUCCUUAGCCUUAUU
Intron
Exon
Intron
−mean(ΔLogitPSI_sub)
5.0
2.5
0.0

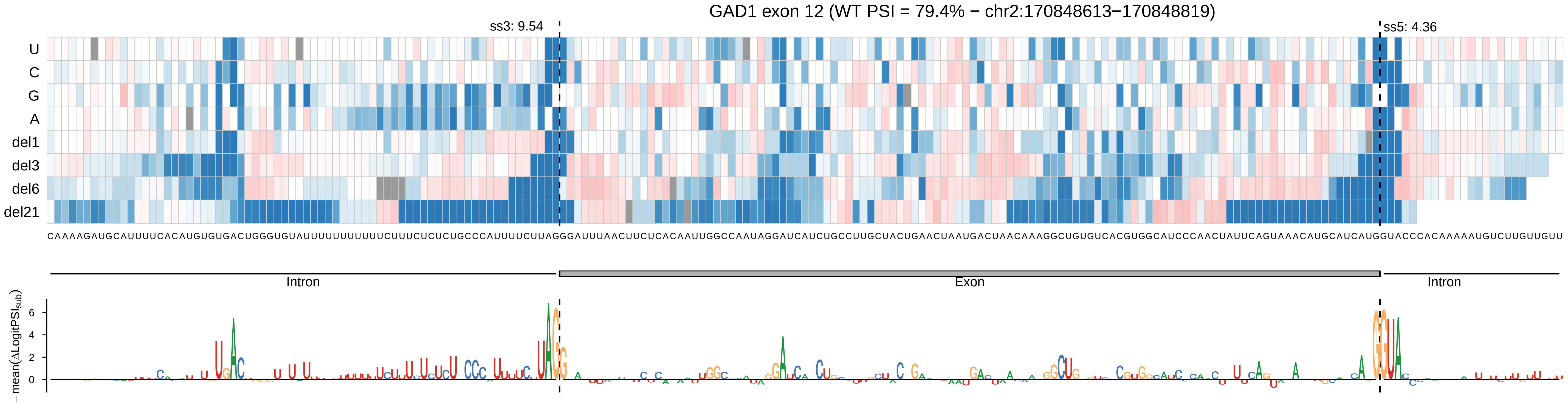

GAD1 exon 12 (WT PSI = 79.4% − chr2:170848613−170848819)
ss3: 9.54
ss5: 4.36
U
C
G
A
del1
del3
del6
del21
CAAAAGAUGCAUUUUCACAUGUGUGACUGGGUGUAUUUUUUUUUUUCUUUCUCUCUGCCCAUUUUCUUAGGGAUUUAACUUCUCACAAUUGGCCAAUAGGAUCAUCUGCCUUGCUACUGAACUAAUGACUAACAAAGGCUGUGUCACGUGGCAUCCCAACUAUUCAGUAAACAUGCAUCAUGGUACCCACAAAAAUGUCUUGUUGUU
Intron
Exon
Intron
−mean(ΔLogitPSI_sub)
6
4
2
0

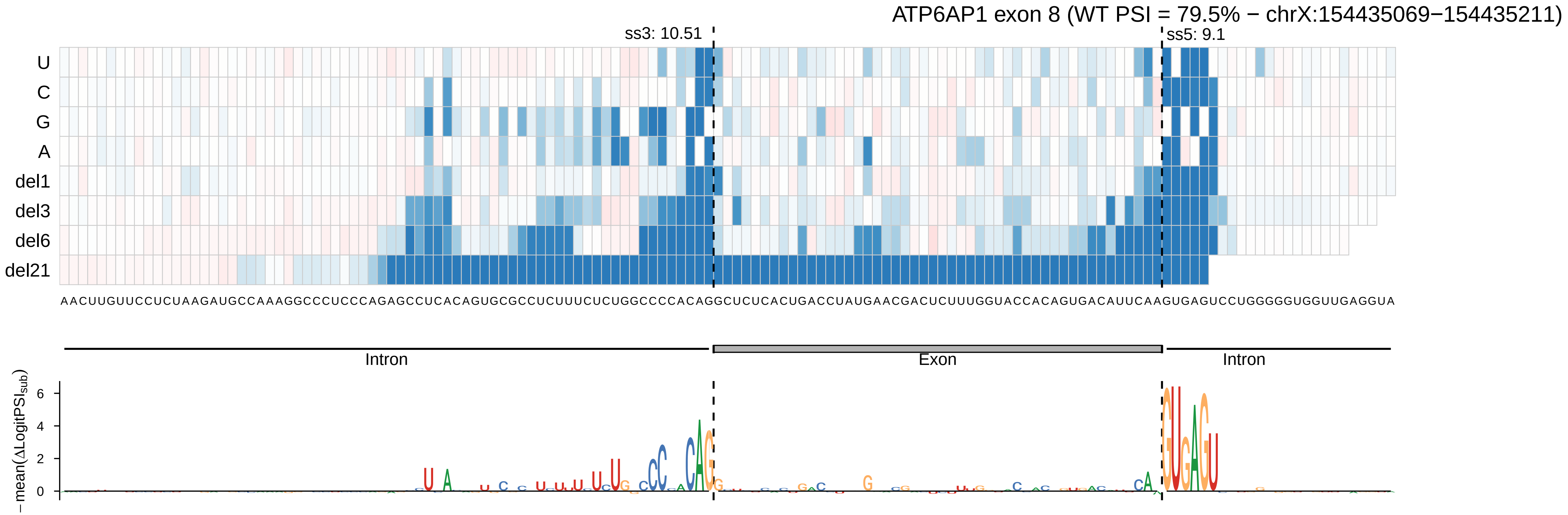

ATP6AP1 exon 8 (WT PSI = 79.5% − chrX:154435069−154435211)
ss3: 10.51
ss5: 9.1
U
C
G
A
del1
del3
del6
del21
AACUUGUUCCUCUAAGAUGCCAAAGGCCCUCCCAGAGCCUCACAGUGCGCCUCUUUCUCUGGCCCCACAGGCUCUCACUGACCUAUGAACGACUCUUUGGUACCACAGUGACAUUCAAGUGAGUCCUGGGGGUGGUUGAGGUA
Intron
Exon
Intron
−mean(ΔLogitPSI_sub)
6
4
2
0

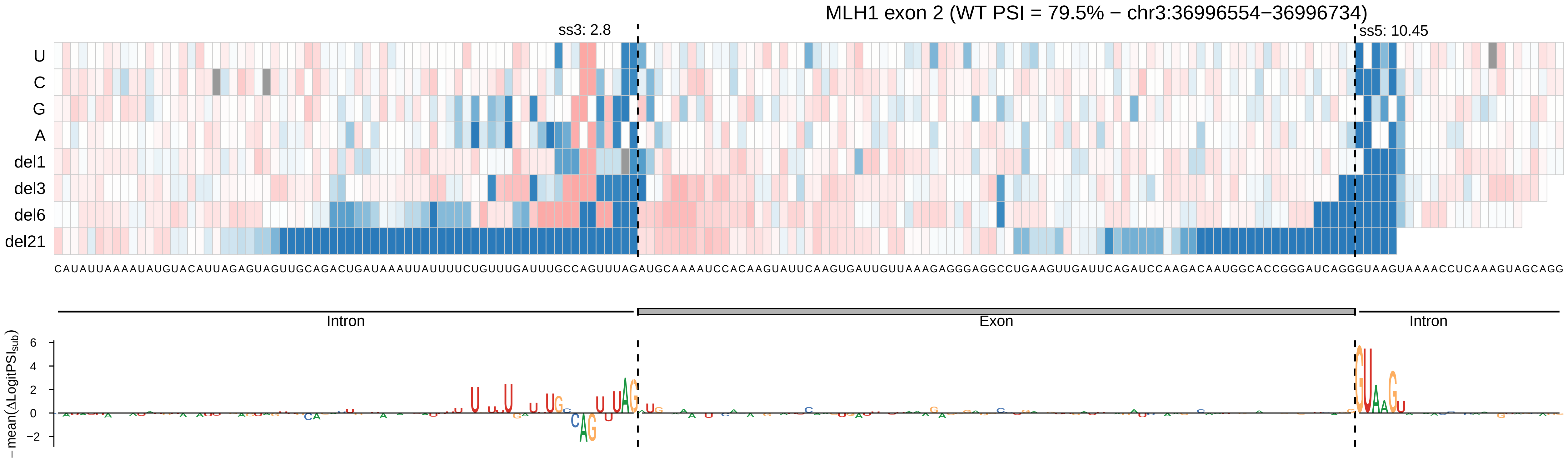

MLH1 exon 2 (WT PSI = 79.5% − chr3:36996554−36996734)
ss3: 2.8
ss5: 10.45
U
C
G
A
del1
del3
del6
del21
CAUAUUAAAAUAUGUACAUUAGAGUAGUUGCAGACUGAUAAAUUAUUUUCUGUUUGAUUUGCCAGUUUAGAUGCAAAAUCCACAAGUAUUCAAGUGAUUGUUAAAGAGGGAGGCCUGAAGUUGAUUCAGAUCCAAGACAAUGGCACCGGGAUCAGGGUAAGUAAAACCUCAAAGUAGCAGG
Intron
Exon
Intron
−mean(ΔLogitPSI_sub)
6
4
2
0
−2

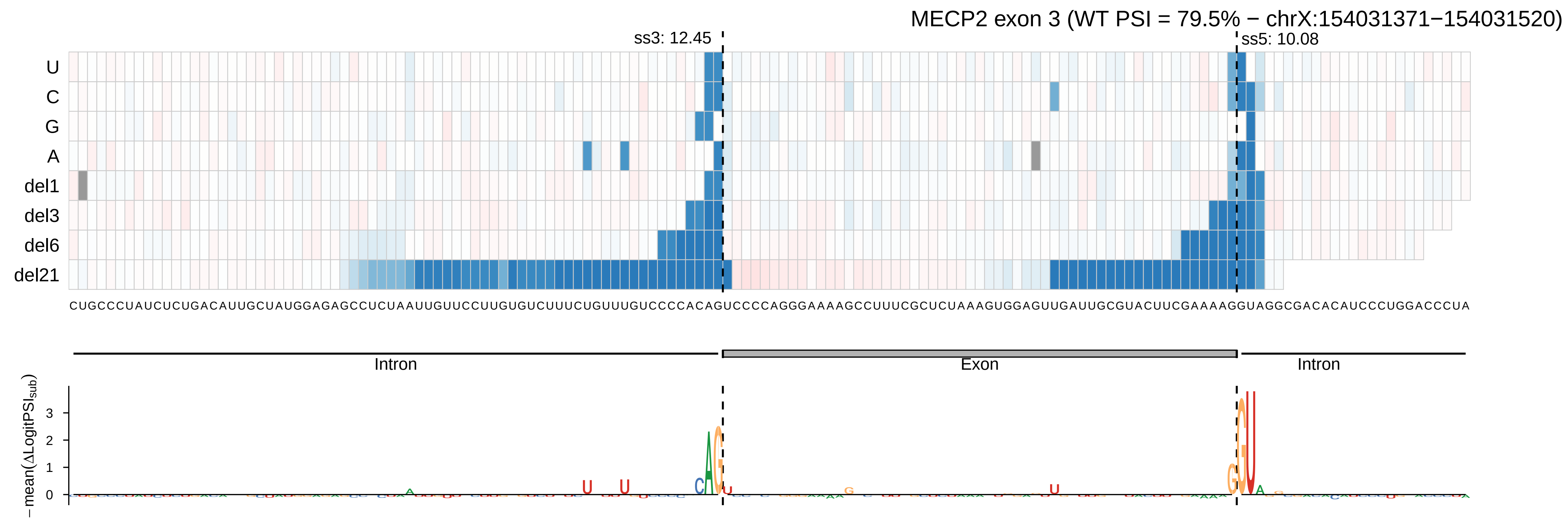

MECP2 exon 3 (WT PSI = 79.5% − chrX:154031371−154031520)
ss3: 12.45
ss5: 10.08
U
C
G
A
del1
del3
del6
del21
CUGCCCUAUCUCUGACAUUGCUAUGGAGAGCCUCUAAUUGUUCCUUGUGUCUUUCUGUUUGUCCCCACAGUCCCCAGGGAAAAGCCUUUCGCUCUAAAGUGGAGUUGAUUGCGUACUUCGAAAAGGUAGGCGACACAUCCCUGGACCCUA
Intron
Exon
Intron
−mean(ΔLogitPSI_sub)
0
1
2
3

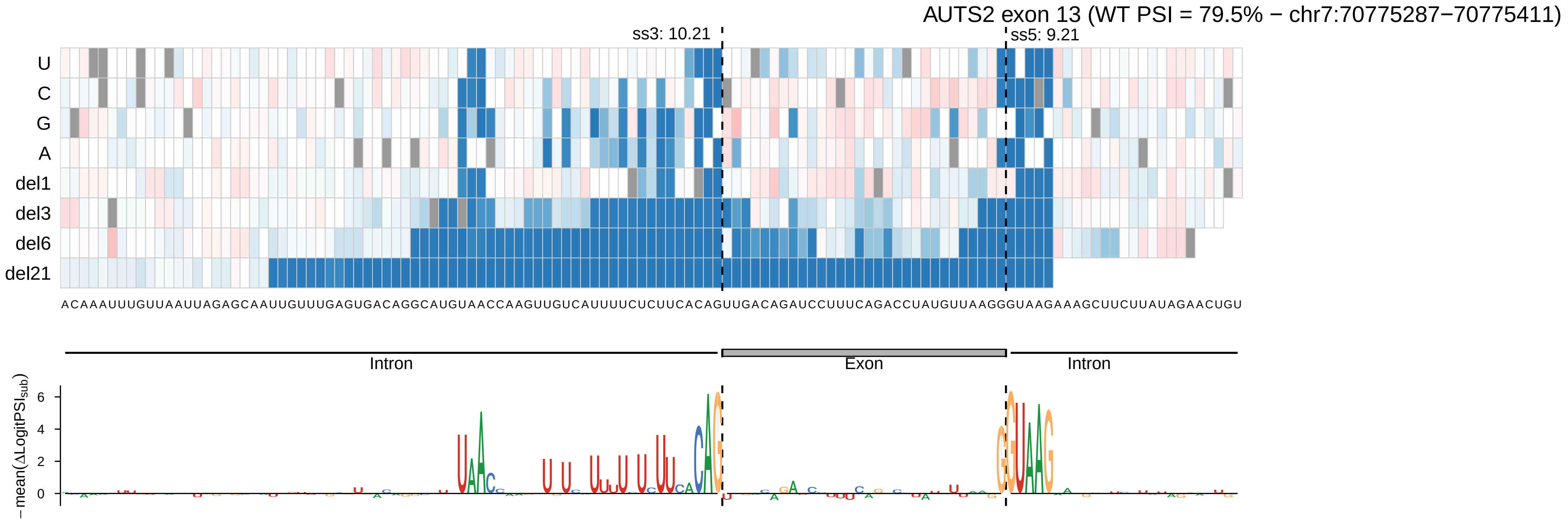

AUTS2 exon 13 (WT PSI = 79.5% − chr7:70775287−70775411)
ss3: 10.21
ss5: 9.21
U
C
G
A
del1
del3
del6
del21
ACAAAUUUGUUAAUUAGAGCAAUUGUUUGAGUGACAGGCAUGUAACCAAGUUGUCAUUUUCUCUUCACAGUUGACAGAUCCUUUCAGACCUAUGUUAAGGGUAAGAAAGCUUCUUAUAGAACUGU
Intron
Exon
Intron
−mean(ΔLogitPSI_sub)
0
2
4
6

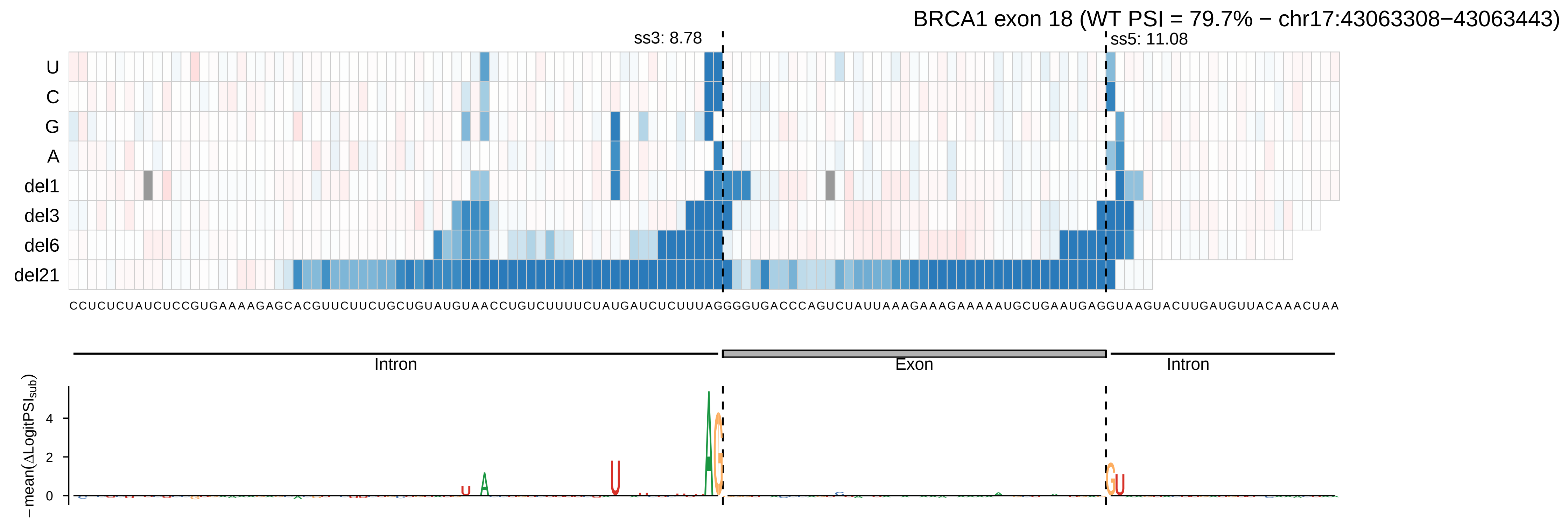
BRCA1 exon 18 (WT PSI = 79.7% − chr17:43063308−43063443)
ss3: 8.78
ss5: 11.08
U
C
G
A
del1
del3
del6
del21
CCUCUCUAUCUCCGUGAAAAGAGCACGUUCUUCUGCUGUAUGUAACCUGUCUUUUCUAUGAUCUCUUUAGGGGUGACCCAGUCUAUUAAAGAAAGAAAAAUGCUGAAUGAGGUAAGUACUUGAUGUUACAAACUAA
Intron
Exon
Intron
−mean(ΔLogitPSI_sub)
4
2
0

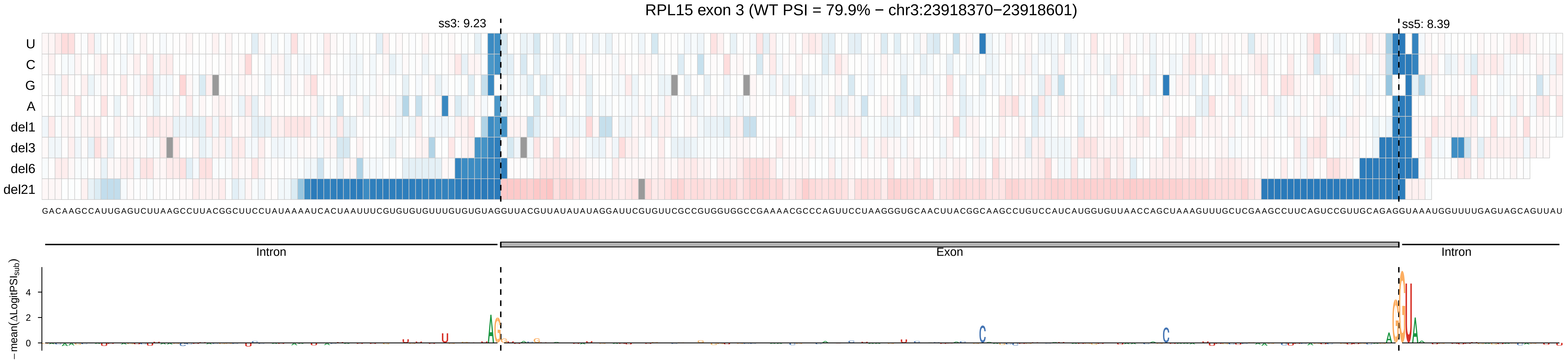

RPL15 exon 3 (WT PSI = 79.9% − chr3:23918370−23918601)
ss3: 9.23
ss5: 8.39
U
C
G
A
del1
del3
del6
del21
GACAAGCCAUUGAGUCUUAAGCCUUACGGCUUCCUAUAAAAUCACUAAUUUCGUGUGUGUUUGUGUGUAGGUUACGUUAUAUAUAGGAUUCGUGUUCGCCGUGGUGGCCGAAAACGCCCAGUUCCUAAGGGUGCAACUUACGGCAAGCCUGUCCAUCAUGGUGUUAACCAGCUAAAGUUUGCUCGAAGCCUUCAGUCCGUUGCAGAGGUAAAUGGUUUUGAGUAGCAGUUAU
Intron
Exon
Intron
−mean(ΔLogitPSI_sub)
4
2
0

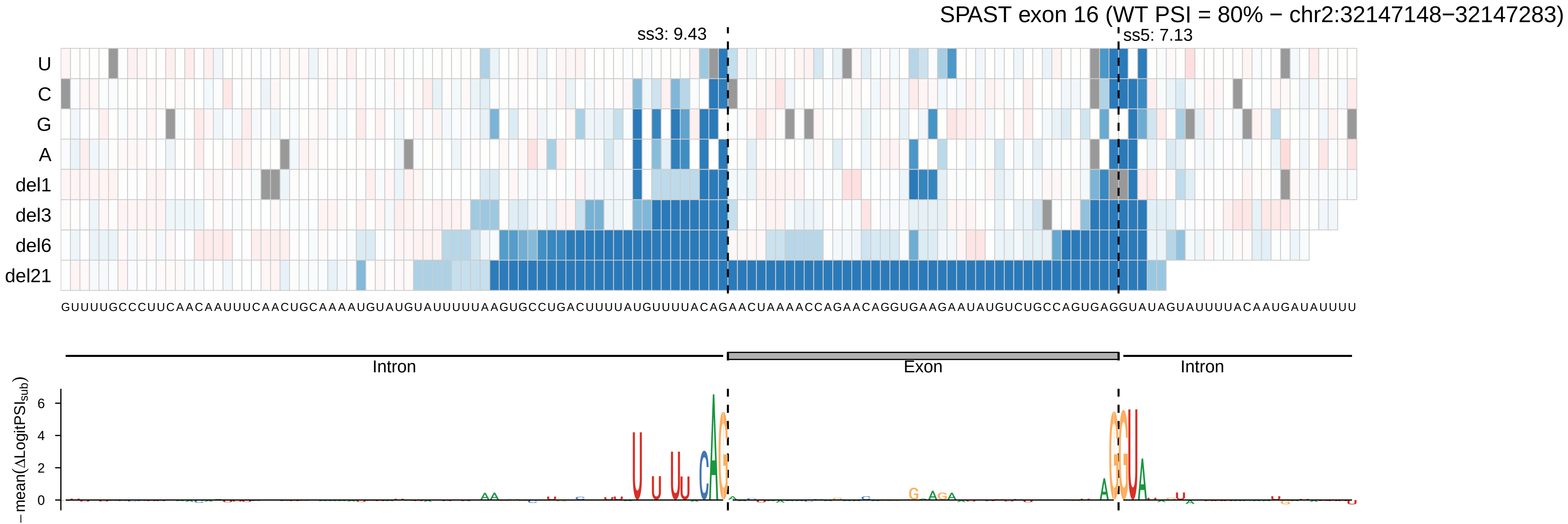
SPAST exon 16 (WT PSI = 80% − chr2:32147148−32147283)
ss3: 9.43
ss5: 7.13
U
C
G
A
del1
del3
del6
del21
GUUUUGCCCUUCAACAAUUUCAACUGCAAAAUGUAUGUAUUUUUAAGUGCCUGACUUUUAUGUUUUACAGAACUAAAACCAGAACAGGUGAAGAAUAUGUCUGCCAGUGAGGUAUAGUAUUUUACAAUGAUAUUUU
Intron
Exon
Intron
− mean(ΔLogitPSI_sub)
6
4
2
0

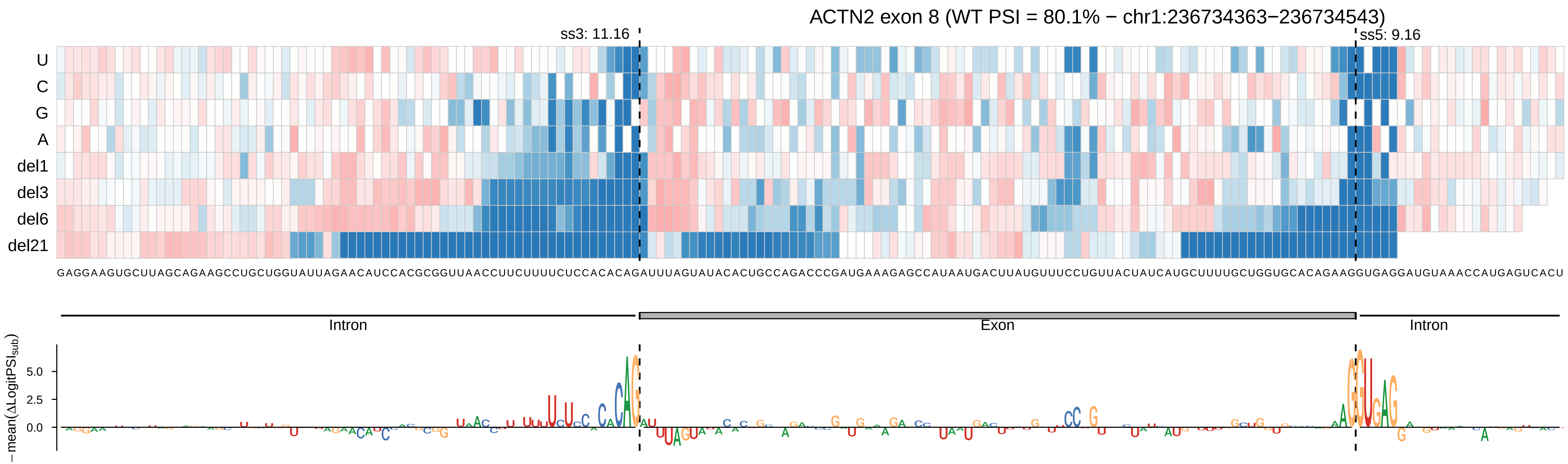

ACTN2 exon 8 (WT PSI = 80.1% − chr1:236734363−236734543)
ss3: 11.16
ss5: 9.16
U
C
G
A
del1
del3
del6
del21
Intron
Exon
Intron
−mean(ΔLogitPSI_sub)
5.0
2.5
0.0

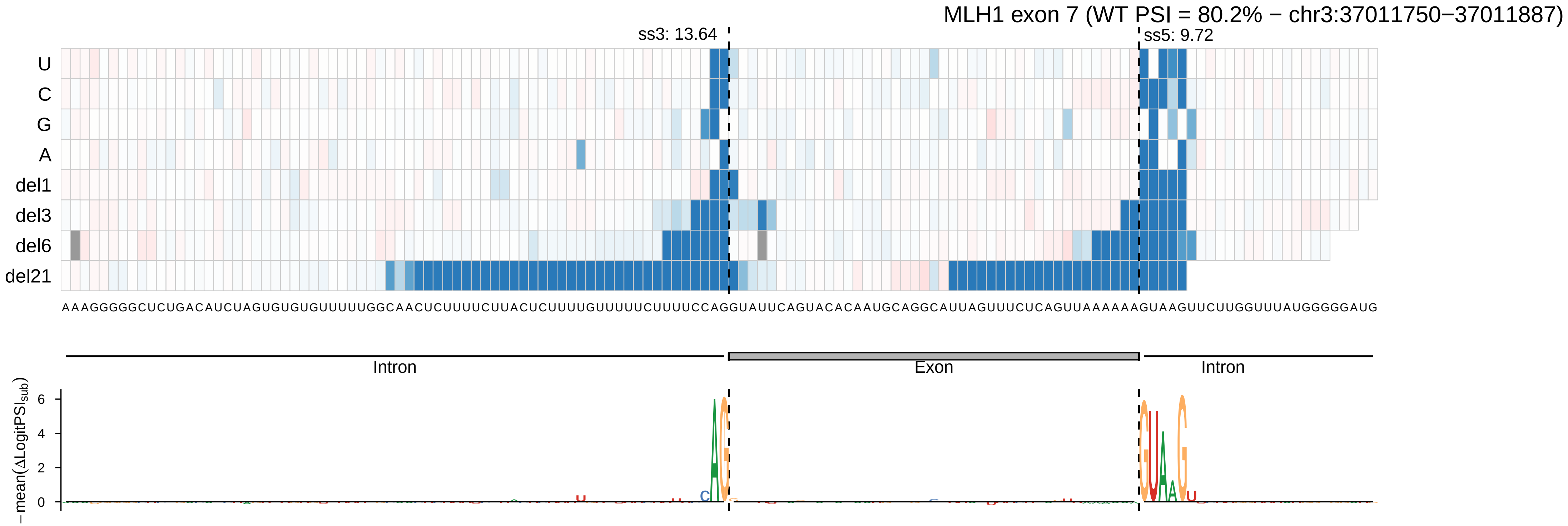

MLH1 exon 7 (WT PSI = 80.2% − chr3:37011750−37011887)
ss3: 13.64
ss5: 9.72
U
C
G
A
del1
del3
del6
del21
AAAGGGGGCUCUGACAUCUAGUGUGUUUUUGGCAACUCUUUUCUUACUCUUUUGUUUUUCUUUUCCAGGUAUUCAGUACACAAUGCAGGCAUUAGUUUCUCAGUUAAAAAAGUAAGUUCUUGGUUUAUGGGGAUG
Intron
Exon
Intron
−mean(ΔLogitPSI_sub)
6
4
2
0

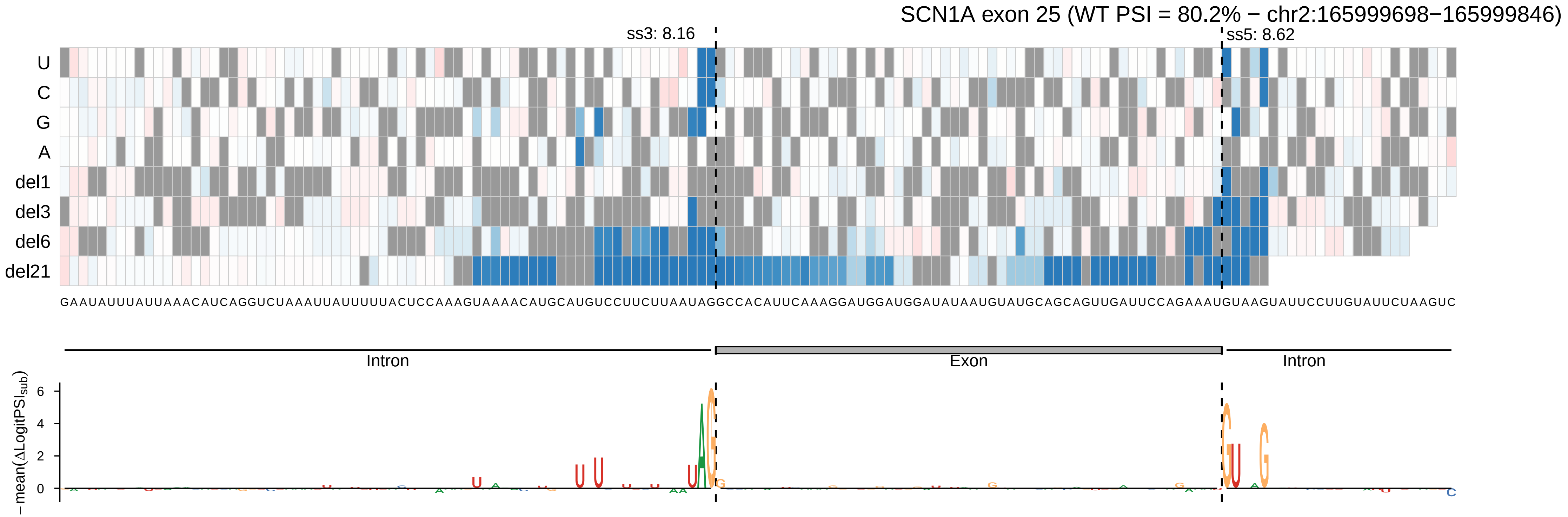

SCN1A exon 25 (WT PSI = 80.2% − chr2:165999698−165999846)
ss3: 8.16
ss5: 8.62
U
C
G
A
del1
del3
del6
del21
GAAUAUUUAUUAAACAUCAGGUCUAAAUUAUUUUUACUCCAAAGUAAAACAUGCAUGUCCUUCUUAAUAGGCCACAUUCAAAGGAUGGAUGGAUAUAAUGUAUGCAGCAGUUGAUUCCAGAAAUGUAAGUAUUCCUUGUAUUCUAAGUC
Intron
Exon
Intron
−mean(ΔLogitPSI_sub)
6
4
2
0

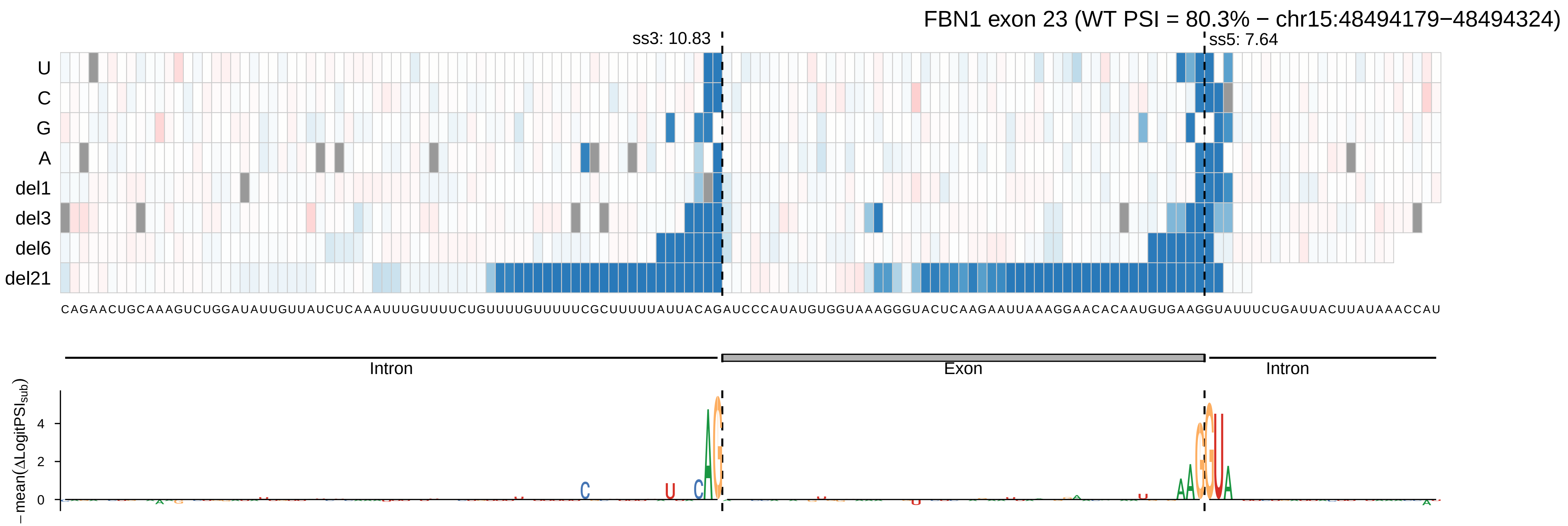

FBN1 exon 23 (WT PSI = 80.3% − chr15:48494179−48494324)
ss3: 10.83
ss5: 7.64
U
C
G
A
del1
del3
del6
del21
CAGAACUGCAAAGUCUGGAUAUUGUUAUCUCAAAUUUGUUUUCUGUUUUGUUUUUCGCUUUUUAUUACAGAUCCCAUAUGUGGUAAAGGGUACUCAAGAAUUAAAGGAACACAAUGUGAAGGUAUUUCUGAUUACUUAUAAACCAU
Intron
Exon
Intron
−mean(ΔLogitPSI_sub)
4
2
0

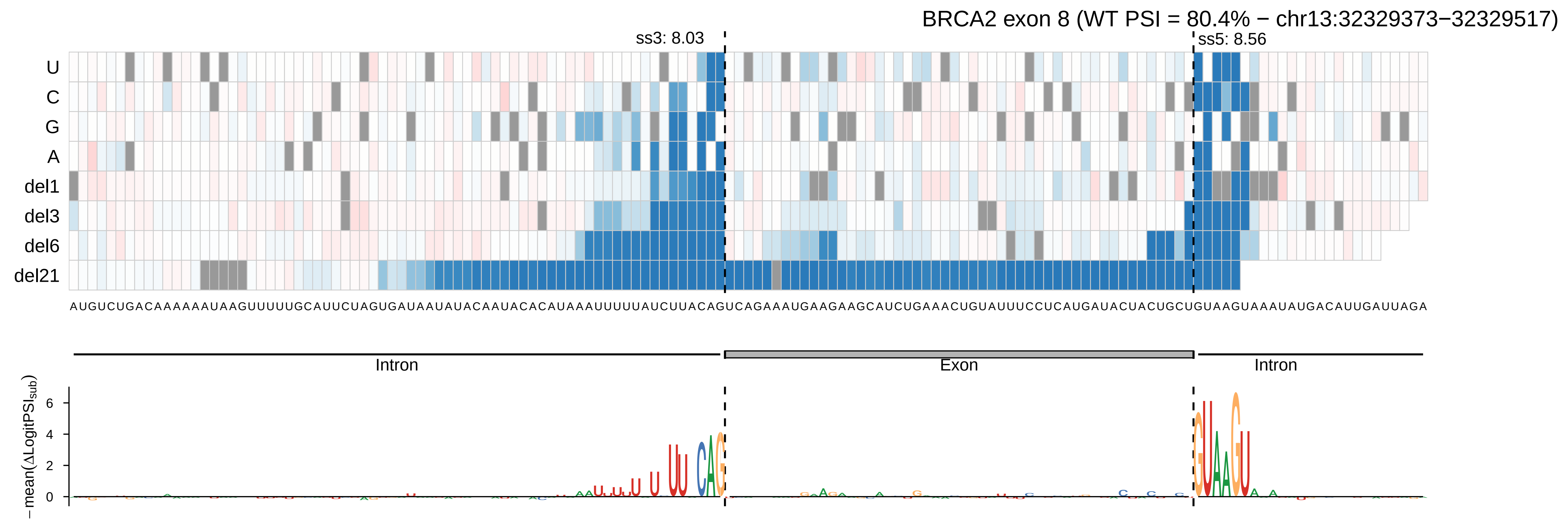
BRCA2 exon 8 (WT PSI = 80.4% − chr13:32329373−32329517)
ss3: 8.03
ss5: 8.56
U
C
G
A
del1
del3
del6
del21
AUGUCUGACAAAAAAUAAGUUUUUGCAUUCUAGUGAUAAUAUACAAUACACAUAAAUUUUUAUCUUACAGUCAGAAAUGAAGAAGCAUCUGAAACUGUAUUUCCUCAUGAUACUACUGCUGUAAGUAAAUAUGACAUUGAUUAGA
Intron
Exon
Intron
−mean(ΔLogitPSI_sub)
6
4
2
0

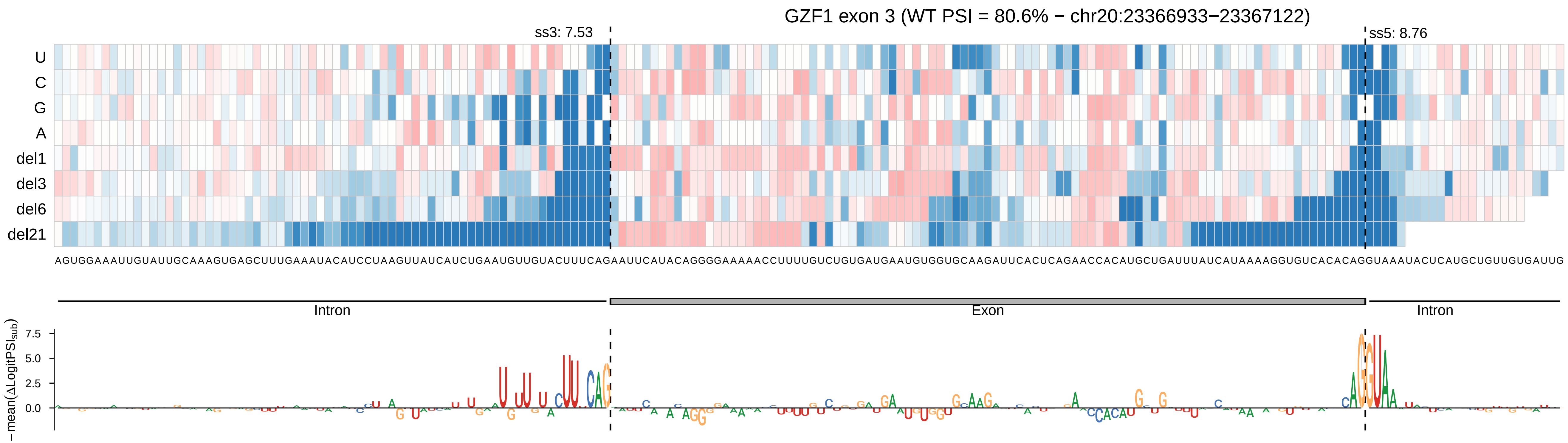

GZF1 exon 3 (WT PSI = 80.6% − chr20:23366933−23367122)
ss3: 7.53
ss5: 8.76
U
C
G
A
del1
del3
del6
del21
AGUGGAAAUUGUAUUGCAAAGUGAGCUUUGAAAUACAUCCUAAGUUAUCAUCUGAAUGUUGUACUUUCAGAAUUCAUACAGGGGAAAAACCUUUUGUCUGUGAUGAAUGUGGUGCAAGAUUCACUCAGAACCACAUGCUGAUUUAUCAUAAAAGGUGUCACACAGGUAAAUACUCAUGCUGUUGUGAUUG
Intron
Exon
Intron
7.5
5.0
2.5
0.0
−mean(ΔLogitPSI_sub)

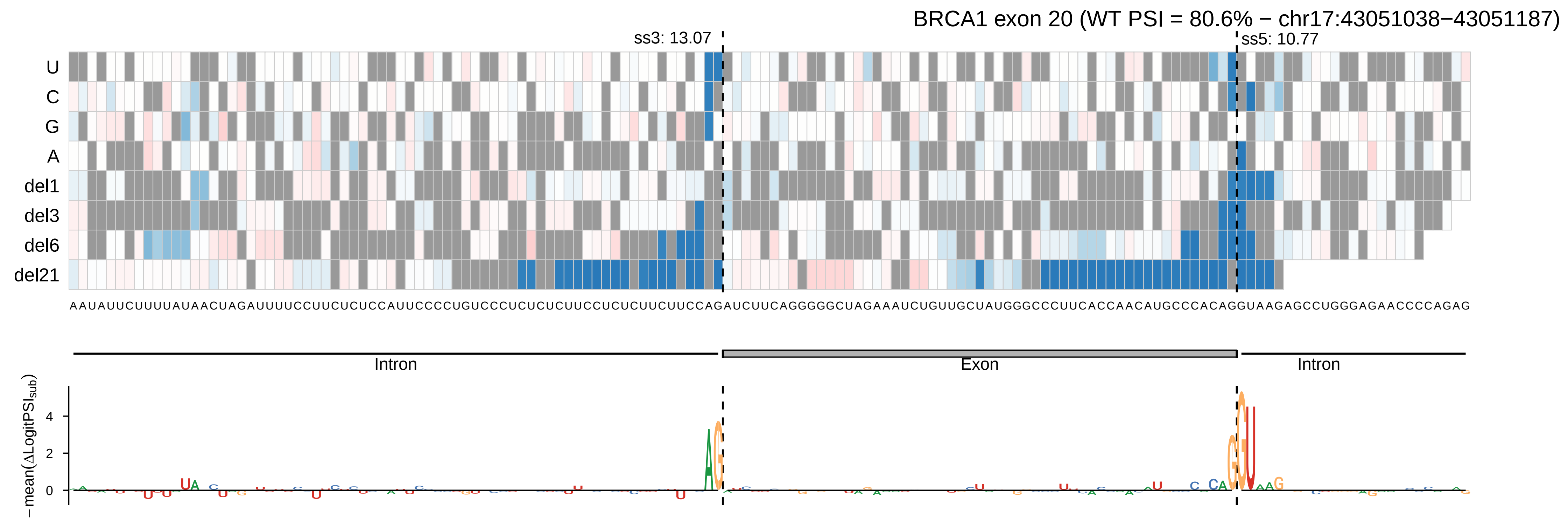

BRCA1 exon 20 (WT PSI = 80.6% − chr17:43051038−43051187)
ss3: 13.07
ss5: 10.77
U
C
G
A
del1
del3
del6
del21
AAUAUUCUUUUAUAACUAGAUUUUCCUUCUCUCCAUUCCCCUGUCCCUCUCUCUUCCUCUCUUCUUCCAGAUCUUCAGGGGGCUAGAAAUCUGUUGCUAUGGGCCCUUCACCAACAUGCCCACAGGUAAGAGCCUGGGAGAACCCCAGAG
Intron
Exon
Intron
−mean(ΔLogitPSI_sub)
4
2
0

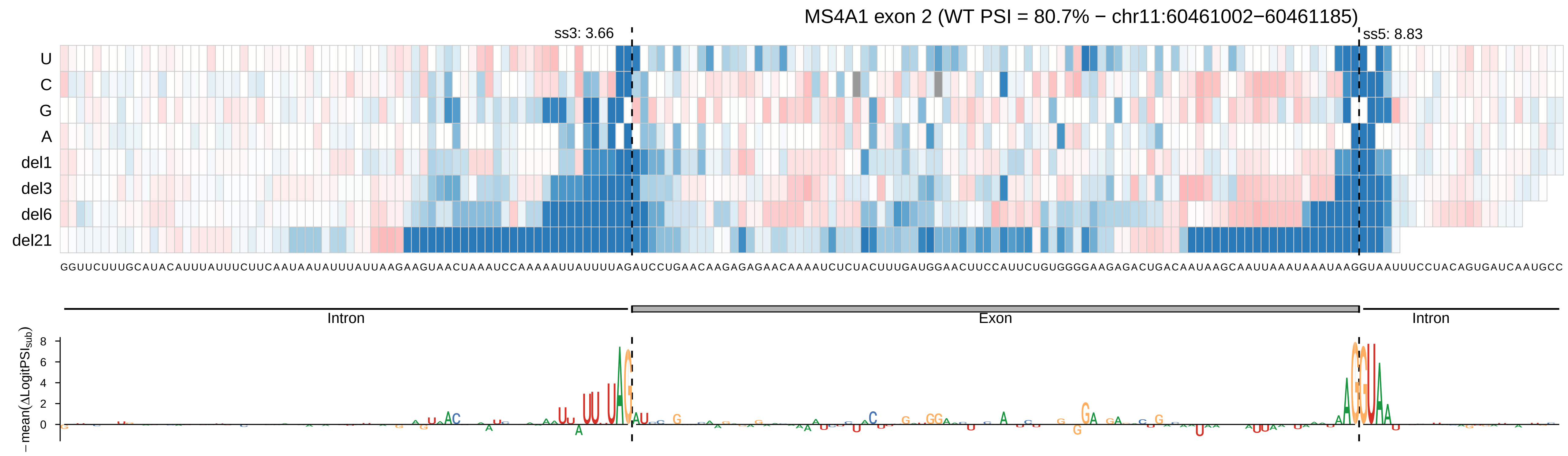
MS4A1 exon 2 (WT PSI = 80.7% − chr11:60461002−60461185)
ss3: 3.66
ss5: 8.83
U
C
G
A
del1
del3
del6
del21
GGUUCUUUGCAUACAUUUAUUUCUUCAAUAAUAUUUAUUAAGAAGUAACUAAAUCCAAAAAUUAUUUUAGAUCCUGAACAAGAGAGAACAAAAUCUCUACUUUGAUGGAACUUCCAUUCUGUGGGGAAGAGACUGACAAUAAGCAAUUAAAUAAAUAAGGUAAUUUCCUACAGUGAUCAAUGCC
Intron
Exon
Intron
− mean(ΔLogitPSI_sub)
8
6
4
2
0

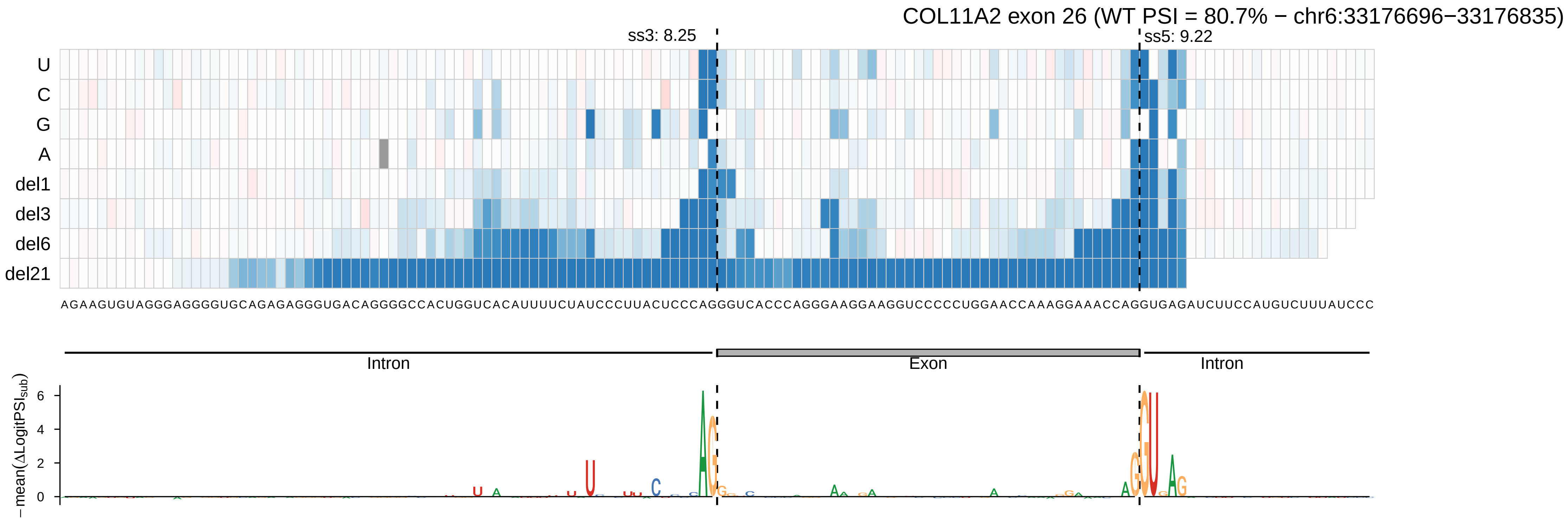
COL11A2 exon 26 (WT PSI = 80.7% − chr6:33176696−33176835)
ss3: 8.25
ss5: 9.22
U
C
G
A
del1
del3
del6
del21
AGAAGUGUAGGGAGGGGUGCAGAGAGGGUGACAGGGGCCACUGGUCACAUUUUCUAUCCCUUACUCCCAGGGUCACCCAGGGAAGGAAGGUCCCCCUGGAACCAAAGGAAACCAGGUGAGAUCUUCCAUGUCUUUAUCCC
Intron
Exon
Intron
− mean(ΔLogitPSI_sub)
6
4
2
0

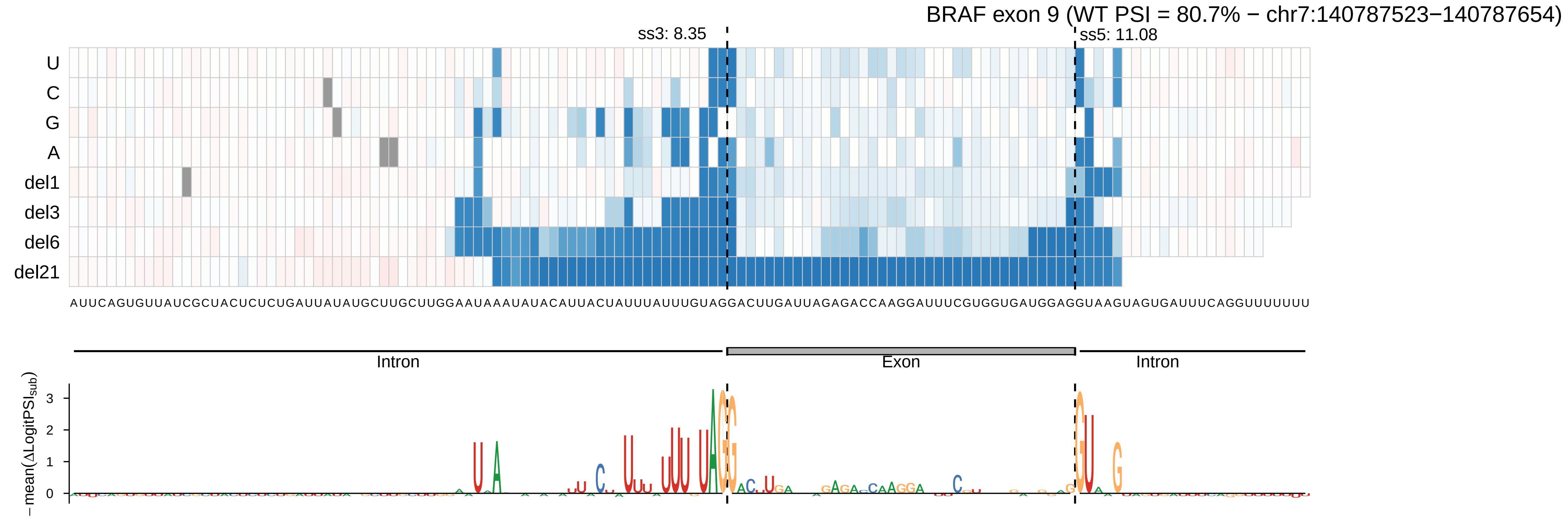
BRAF exon 9 (WT PSI = 80.7% − chr7:140787523−140787654)
ss3: 8.35
ss5: 11.08
U
C
G
A
del1
del3
del6
del21
AUUCAGUGUUAUCGCUACUCUCUGAUUAUAUGCUUGCUUGGAAUAAAUAUACAUUACUAUUUAUUUGUAGGACUUGAUUAGAGACCAAGGAUUUCGUGGUGAUGGAGGUAAGUAGUGAUUUCAGGUUUUUUU
Intron
Exon
Intron
−mean(ΔLogitPSIsub)
0
1
2
3

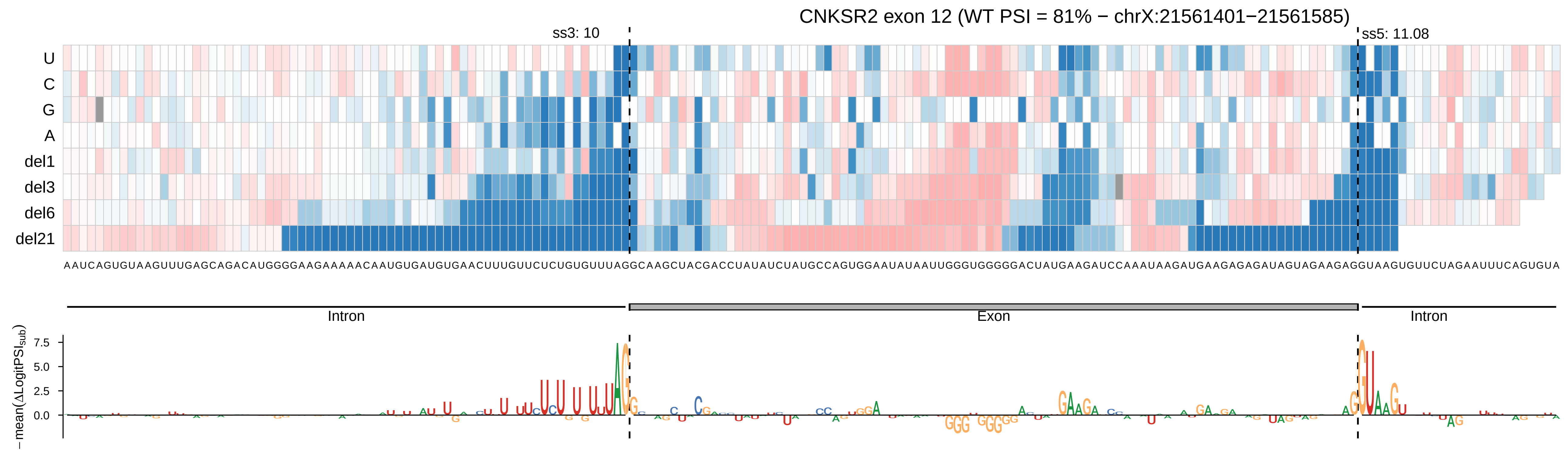
CNKSR2 exon 12 (WT PSI = 81% − chrX:21561401−21561585)
ss3: 10
ss5: 11.08
U
C
G
A
del1
del3
del6
del21
Intron
Exon
Intron
−mean(ΔLogitPSI_sub)
7.5
5.0
2.5
0.0

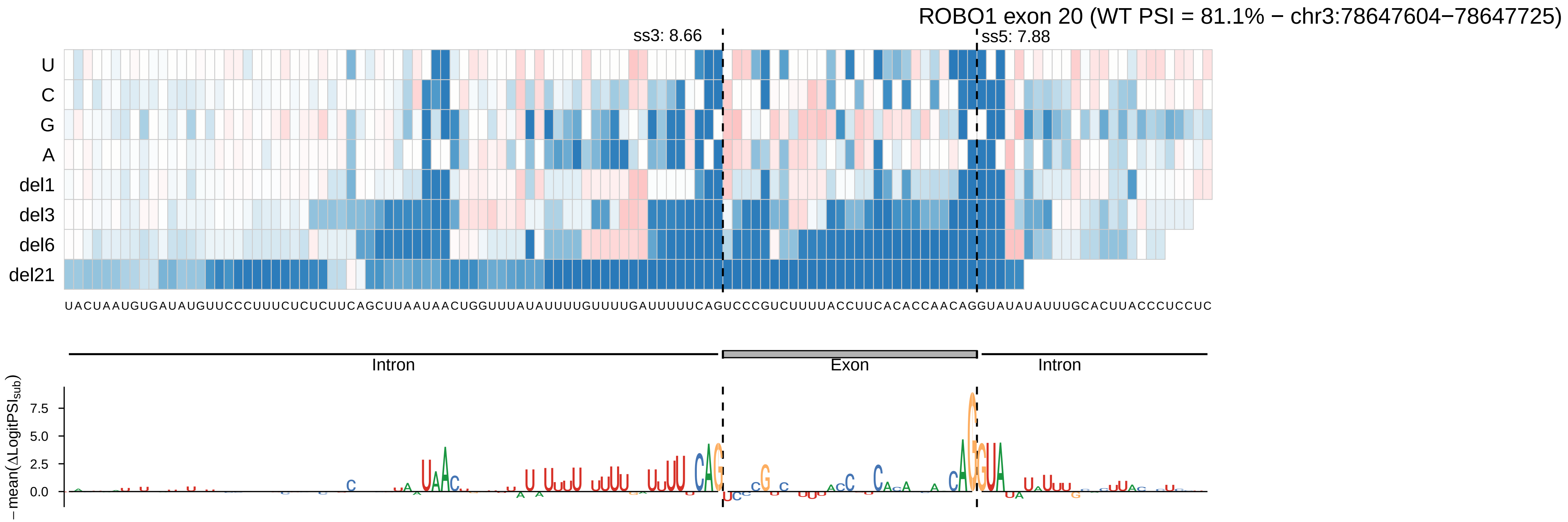

ROBO1 exon 20 (WT PSI = 81.1% − chr3:78647604−78647725)
ss3: 8.66
ss5: 7.88
U
C
G
A
del1
del3
del6
del21
UACUAAUGUGAUAUGUUCCCUUUCUCUCUUCAGCUUAAUAACUGGUUUAUAUUUUGUUUUGAUUUUUCAGUCCCGUCUUUUACCUUCACACCAACAGGUAUAUAUUUGCACUUACCCUCCUC
Intron
Exon
Intron
− mean(ΔLogitPSI_sub)
7.5
5.0
2.5
0.0

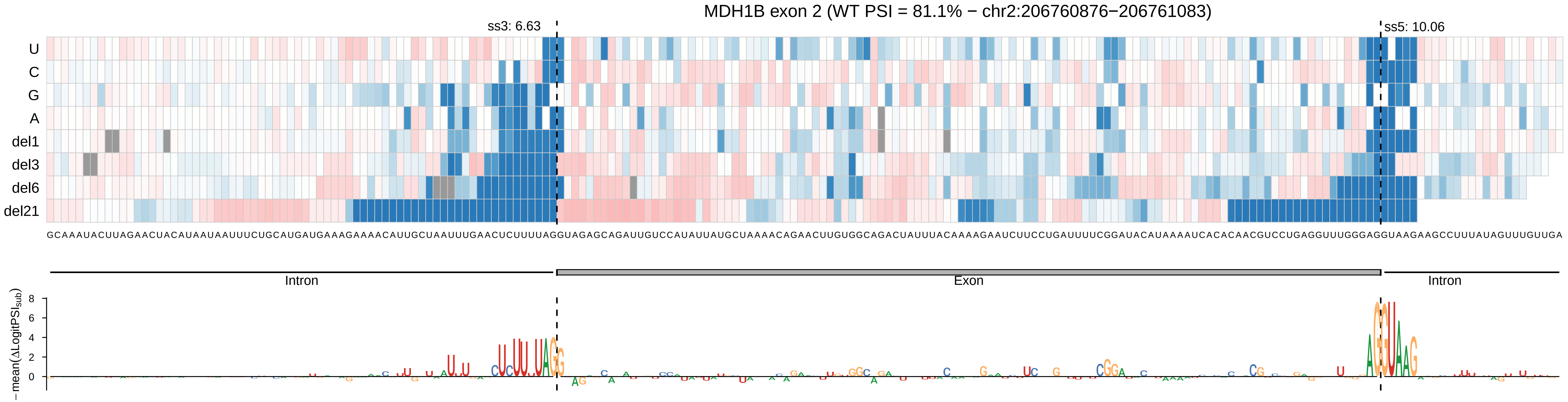

MDH1B exon 2 (WT PSI = 81.1% − chr2:206760876−206761083)
ss3: 6.63
ss5: 10.06
U
C
G
A
del1
del3
del6
del21
GCAAAUACUUAGAACUACAUAAUAAUUUCUGCAUGAUGAAAGAAAACAUUGCUAAUUUGAACUCUUUUAGGUAGAGCAGAUUGUCCAUAUUAUGCUAAAACAGAACUUGUGGCAGACUAUUUACAAAAGAAUCUUCCUGAUUUUCGGAUACAUAAAAUCACACAACGUCCUGAGGUUUGGGAGGUAAGAAGCCUUUAUAGUUUGUUGA
Intron
Exon
Intron
−mean(ΔLogitPSI_sub)
8
6
4
2
0

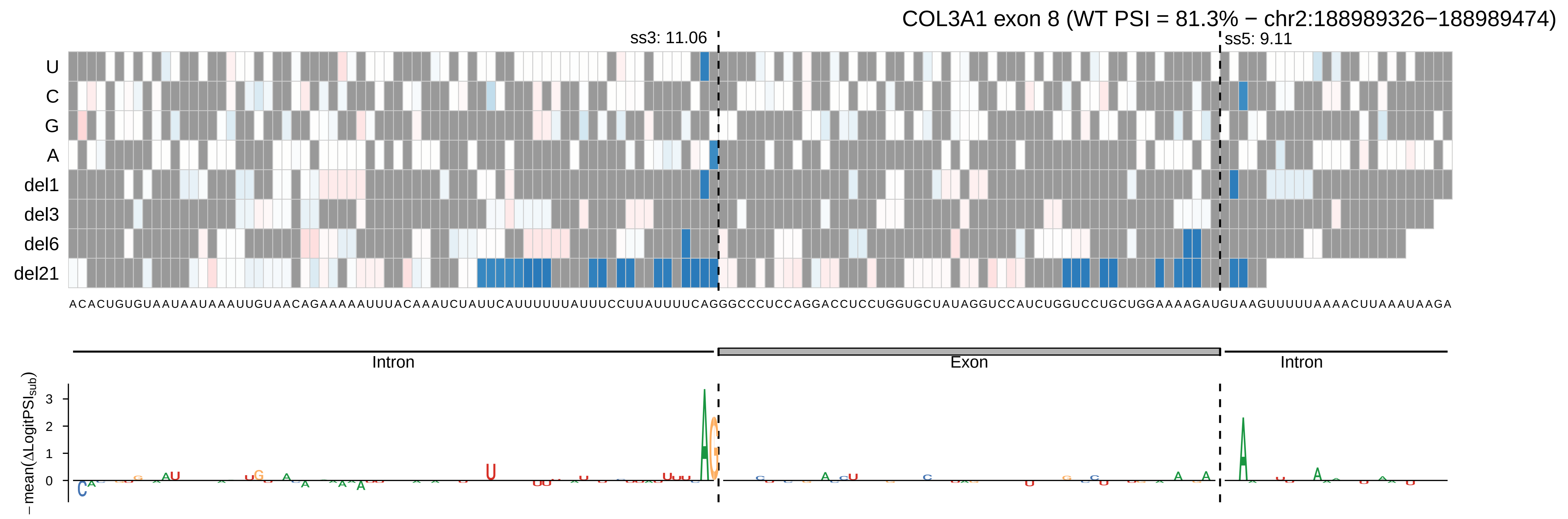
COL3A1 exon 8 (WT PSI = 81.3% − chr2:188989326−188989474)
ss3: 11.06
ss5: 9.11
U
C
G
A
del1
del3
del6
del21
ACACUGUGUAAUAAUAAAUUGUAACAGAAAAAUUUACAAAUCUAUUCAUUUUUUAUUUCCUUAUUUUCAGGGCCCCUCCAGGACCUCCUGGUGCUAUAGGUCCAUCUGGUCCUGCUGGAAAAGAUGUAAGUUUUUAAAACUUAAAUAAGA
Intron
Exon
Intron
−mean(ΔLogitPSI_sub)
3
2
1
0

COL6A2 exon 7 (WT PSI = 81.3% − chr21:46115939−46116078)

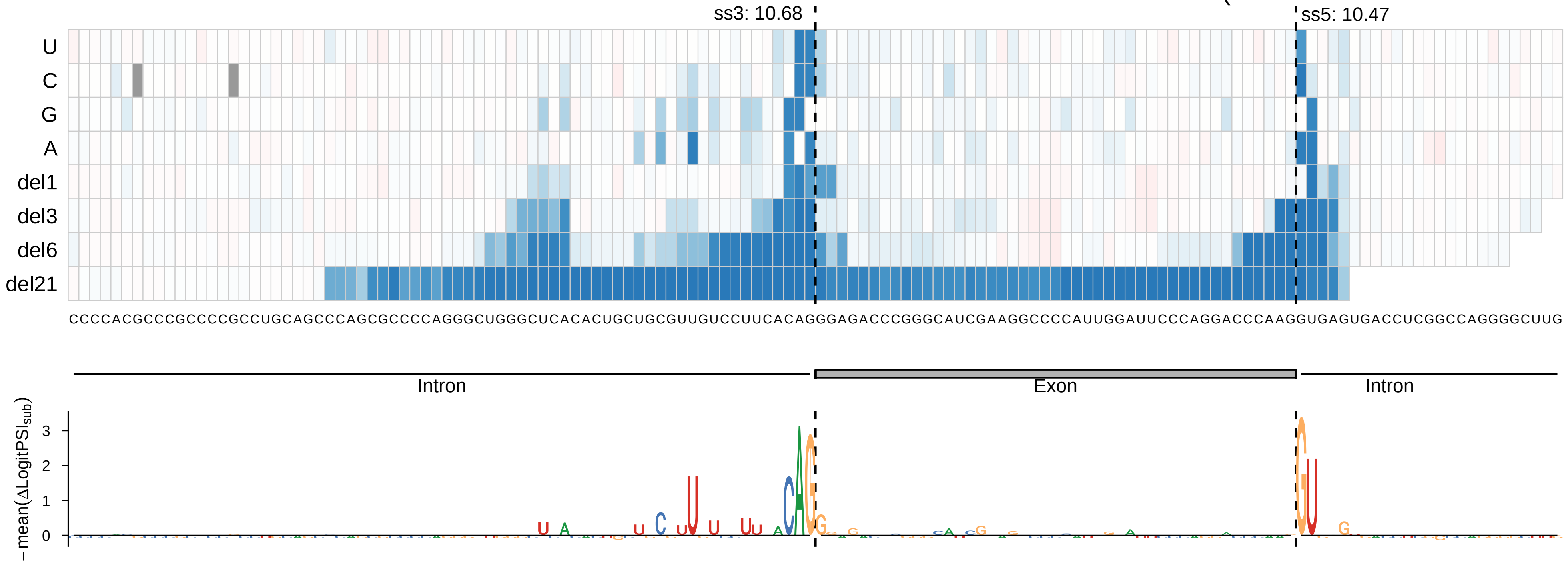

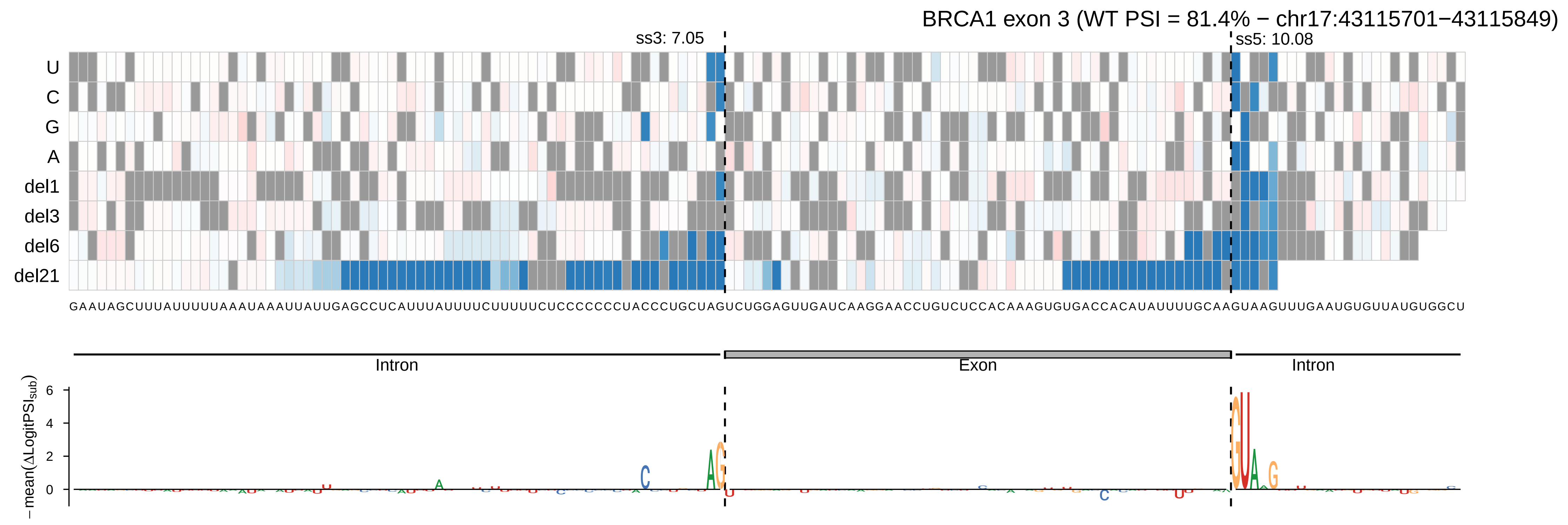
BRCA1 exon 3 (WT PSI = 81.4% − chr17:43115701−43115849)
ss3: 7.05
ss5: 10.08
U
C
G
A
del1
del3
del6
del21
GAAUAGCUUUAUUUUUAAAUAAAUUAUUGAGCCUCAUUUAUUUUCUUUUUCUCCCCCCCUACCCUGCUAGUCUGGAGUUGAUCAAGGAACCUGUCUCCACAAAGUGUGACCACAUAUUUUGCAAGUAAGUUUGAAUGUGUUAUGUGGCU
Intron
Exon
Intron
−mean(ΔLogitPSI_sub)
6
4
2
0

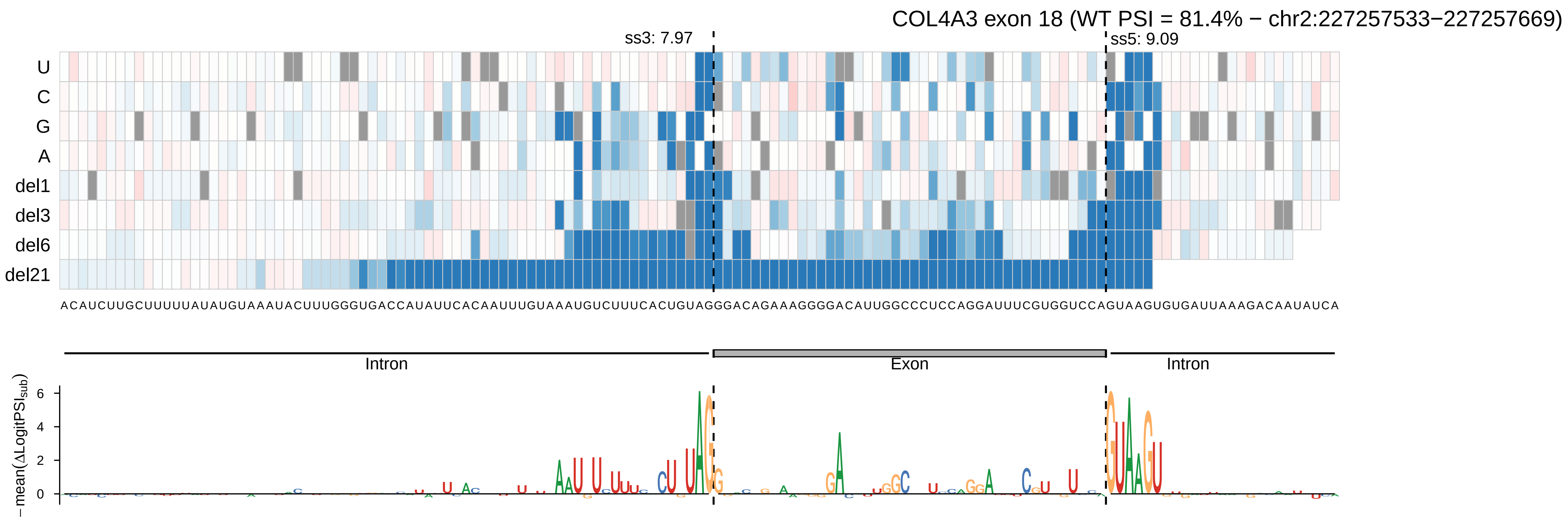
COL4A3 exon 18 (WT PSI = 81.4% − chr2:227257533−227257669)
ss3: 7.97
ss5: 9.09
U
C
G
A
del1
del3
del6
del21
ACAUCUUGCUUUUUAUAUGUAAAUACUUUGGGUGACCAUAUUCACAAUUUGUAAAUGUCUUUCACUGUAGGGACAGAAAGGGGACAUUGGCCCUCCAGGAUUUCGUGGUCCAGUAAGUGUGAUUAAAGACAAUAUCA
Intron
Exon
Intron
−mean(ΔLogitPSI_sub)
6
4
2
0

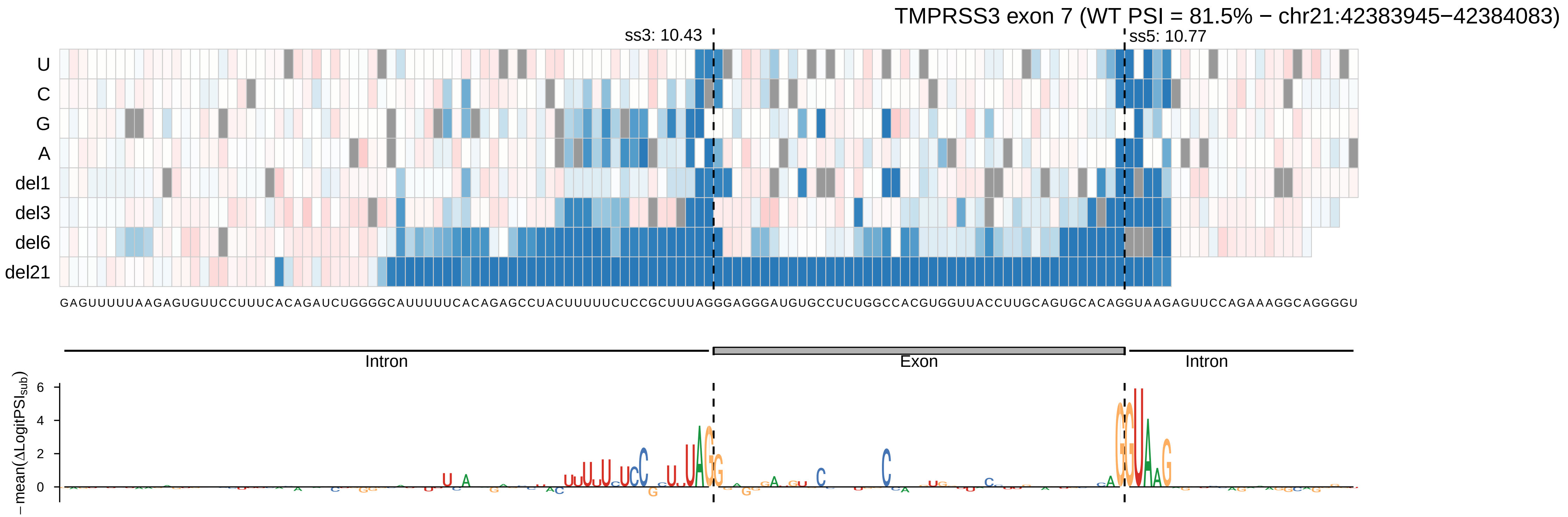

TMPRSS3 exon 7 (WT PSI = 81.5% − chr21:42383945−42384083)
ss3: 10.43
ss5: 10.77
U
C
G
A
del1
del3
del6
del21
GAGUUUUUAAGAGUGUUCCUUUCACAGAUCUGGGGCAUUUUUCACAGAGCCUACUUUUUCUCCGCUUUAGGGAGGGAUGUGCCUCUGGCCACGUGGUUACCUUGCAGUGCACAGGUAAGAGUUCCAGAAAGGCAGGGGU
Intron
Exon
Intron
− mean(ΔLogitPSI_sub)
6
4
2
0

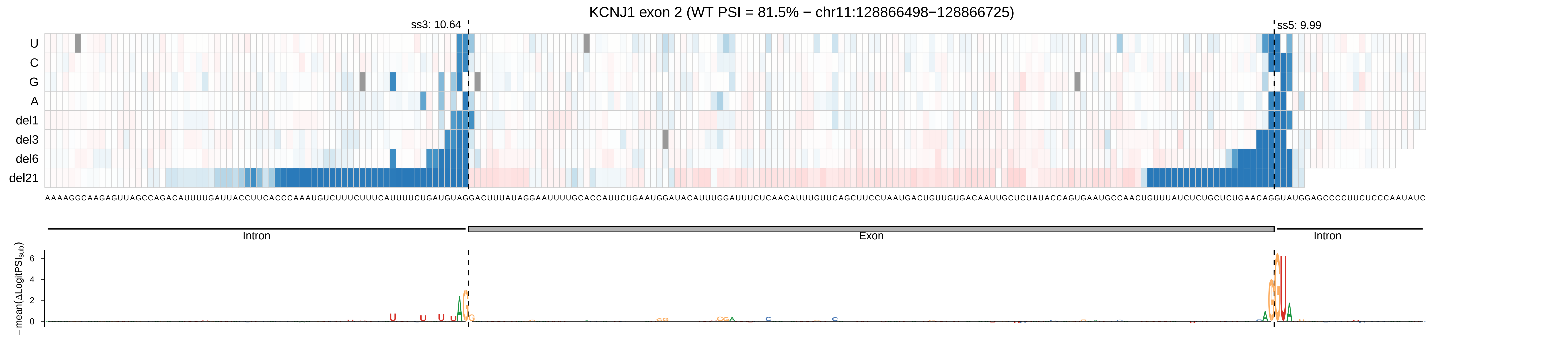

KCNJ1 exon 2 (WT PSI = 81.5% − chr11:128866498−128866725)
ss3: 10.64
ss5: 9.99
U
C
G
A
del1
del3
del6
del21
Intron
Exon
Intron
−mean(ΔLogitPSIsub)

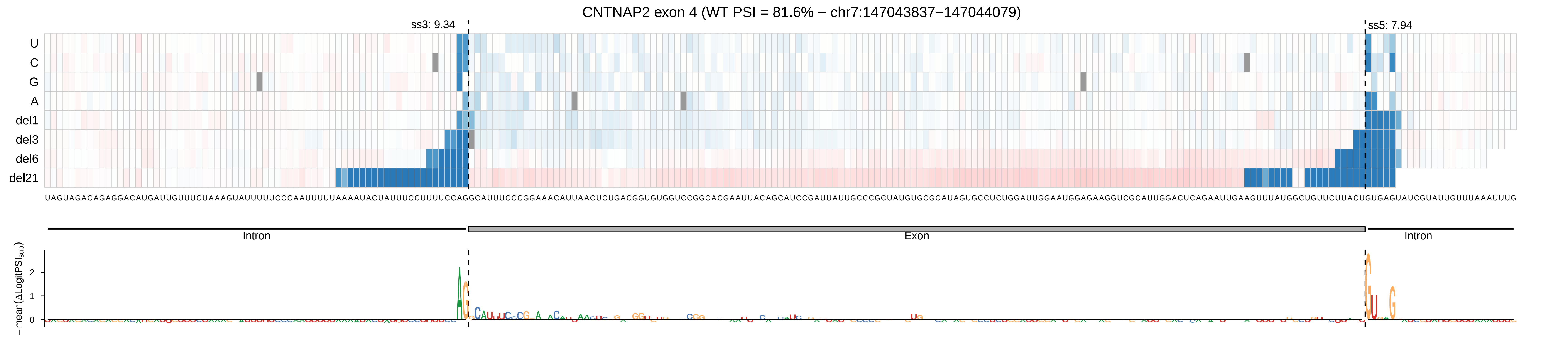

CNTNAP2 exon 4 (WT PSI = 81.6% − chr7:147043837−147044079)
ss3: 9.34
ss5: 7.94
U
C
G
A
del1
del3
del6
del21
Intron
Exon
Intron
− mean(ΔLogitPSI_sub)
2
1
0

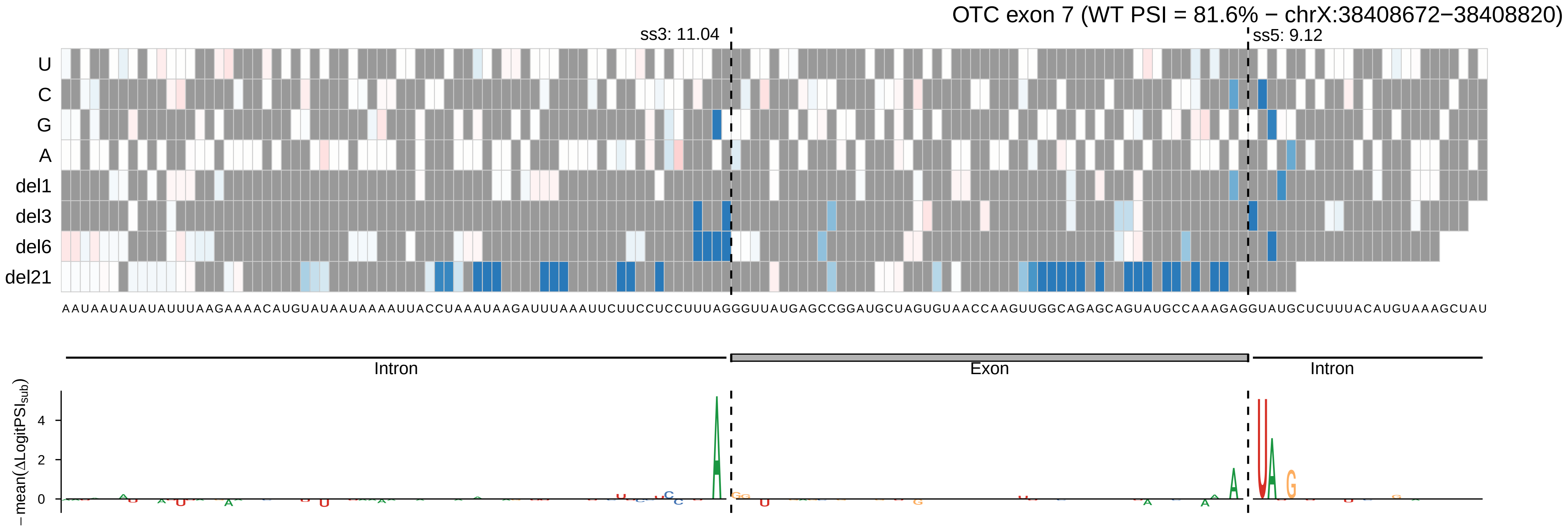
OTC exon 7 (WT PSI = 81.6% − chrX:38408672−38408820)
ss3: 11.04
ss5: 9.12
U
C
G
A
del1
del3
del6
del21
AAUAAUAUAUAUUUAAGAAAACAUGUAUAAUAAAAUUACCUAAAUAAGAUUUAAAUUCUUCCUCCUUUAGGGUUAUGAGCCGGAUGCUAGUGUAACCAAGUUGGCAGAGCAGUAUGCCAAAGAGGUAUGCUCUUUACAUGUAAAGCUAU
Intron
Exon
Intron
4
2
0

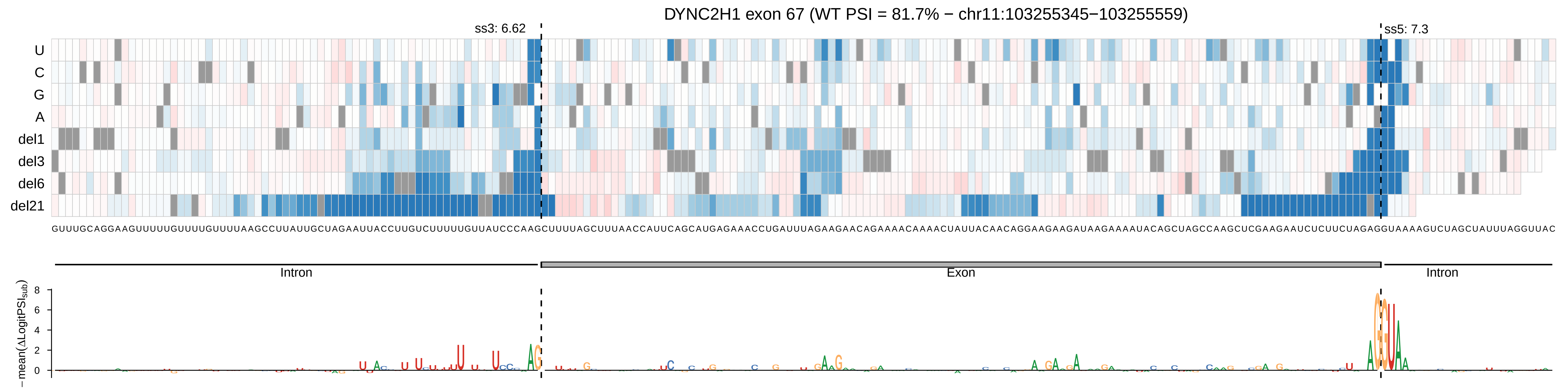

DYNC2H1 exon 67 (WT PSI = 81.7% − chr11:103255345−103255559)
ss3: 6.62
ss5: 7.3
U
C
G
A
del1
del3
del6
del21
GUUUGCAGGAAGUUUUUGUUUUGUUUUAAGCCUUAUUGCUAGAAUUACCUUGUCUUUUUGUUAUCCCAAGCUUUUAGCUUUAACCAUUCAGCAUGAGAAACCUGAUUUAGAAGAACAGAAAACAAAACUAUUACAACAGGAAGAAGAUAAGAAAAUACAGCUAGCCAAGCUCGAAGAAUCUCUUCUAGAGGUAAAAGUCUAGCUAUUUAGGUUAC
Intron
Exon
Intron
− mean(ΔLogitPSI_sub)
0
2
4
6
8

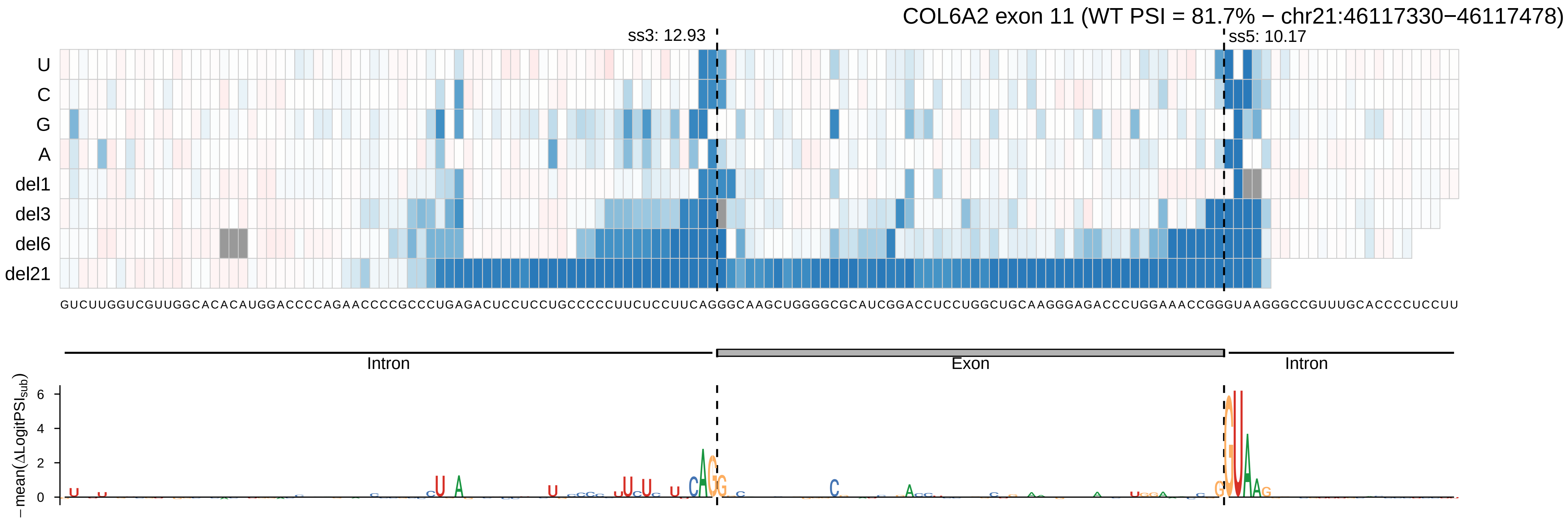
COL6A2 exon 11 (WT PSI = 81.7% − chr21:46117330−46117478)
ss3: 12.93
ss5: 10.17
U
C
G
A
del1
del3
del6
del21
GUCUUGGUCGUUGGCACACAUGGACCCCAGAACCCCGCCCUGAGACUCCUCCUGCCCCCUUCUCCUUCAGGGCAAGCUGGGGCGCAUCGGACCUCCUGGCUGCAAGGGAGACCCUGGAAACCGGGUAAGGGCCGUUUGCACCCCUCCUU
Intron
Exon
Intron
−mean(ΔLogitPSI_sub)
6
4
2
0

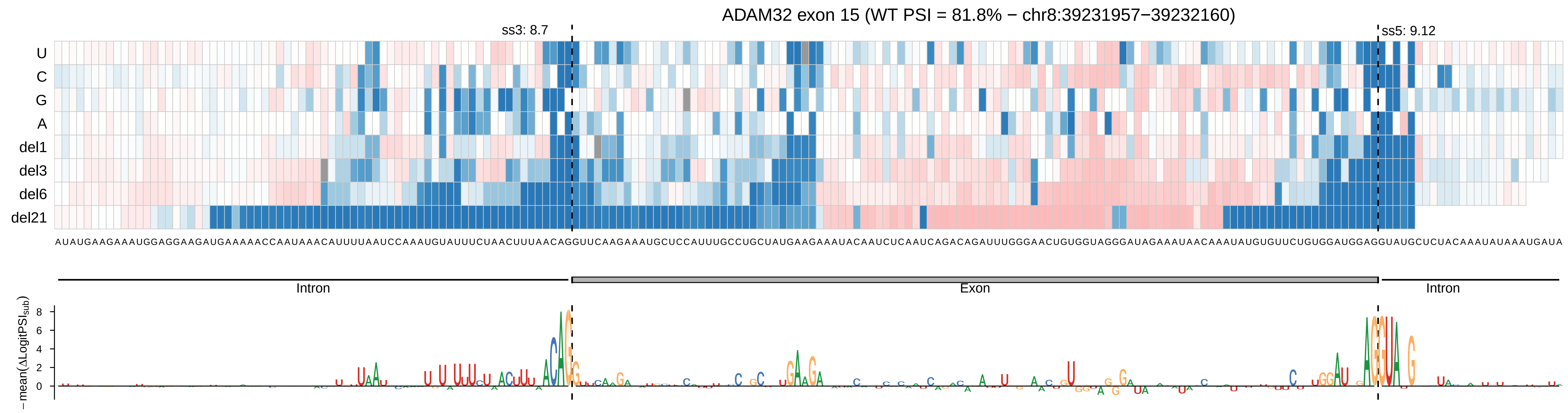

ADAM32 exon 15 (WT PSI = 81.8% − chr8:39231957−39232160)
ss3: 8.7
ss5: 9.12
U
C
G
A
del1
del3
del6
del21
Intron
Exon
Intron
−mean(ΔLogitPSI_sub)
8
6
4
2
0

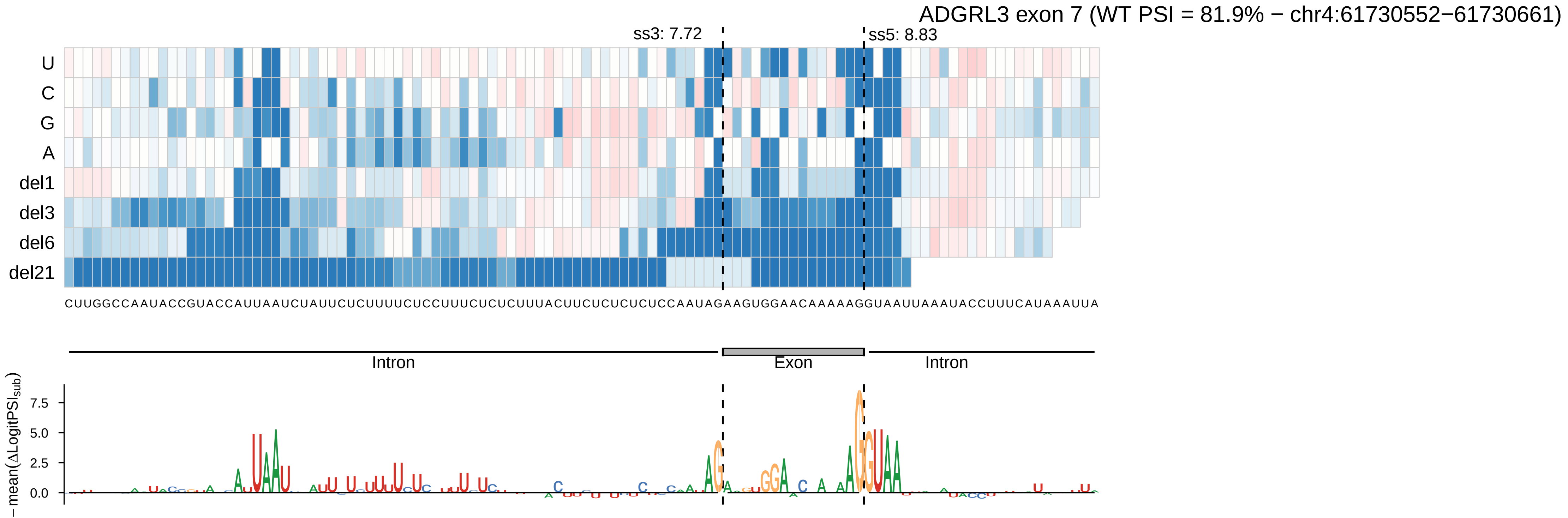
ADGRL3 exon 7 (WT PSI = 81.9% − chr4:61730552−61730661)
ss3: 7.72
ss5: 8.83
U
C
G
A
del1
del3
del6
del21
CUUGGCCAAUACCGUACCAUUAAUCUAUUCUCUUUUCUCCUUUCUCUCUUUACUUCUCUCUCUCCAAUAGAAGUGGAACAAAAAGGUAAUUAAAAUACCUUUCAUAAAUUA
Intron
Exon
Intron
−mean(ΔLogitPSI_sub)
7.5
5.0
2.5
0.0

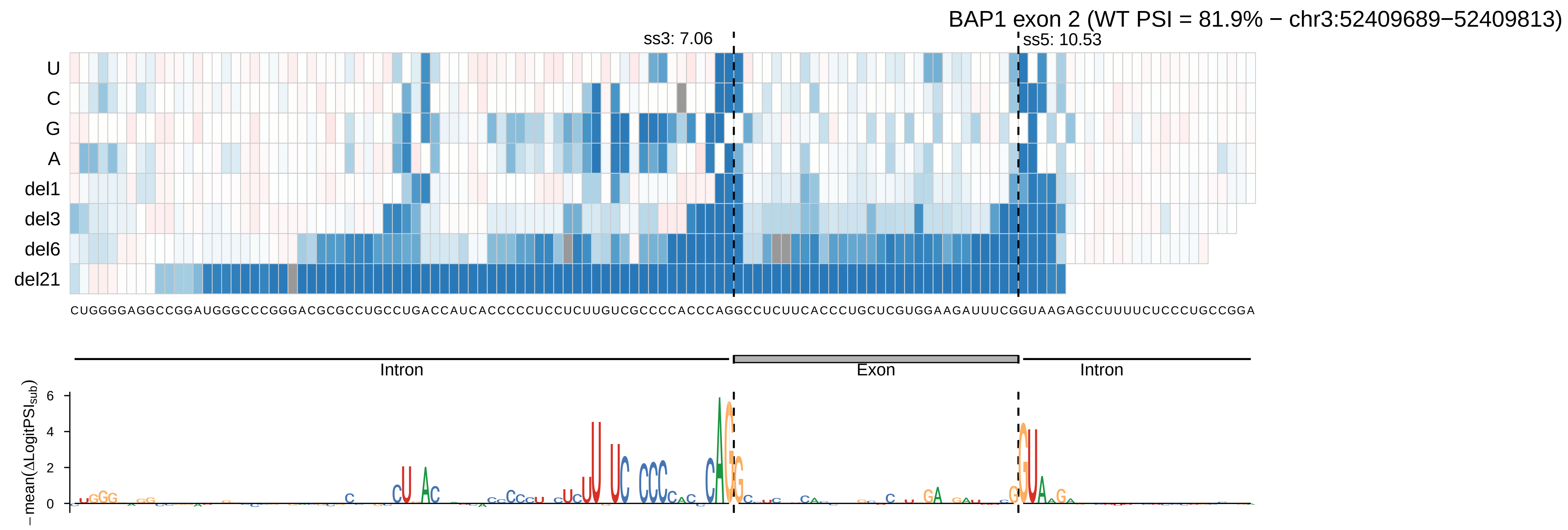

BAP1 exon 2 (WT PSI = 81.9% − chr3:52409689−52409813)
ss3: 7.06
ss5: 10.53
U
C
G
A
del1
del3
del6
del21
CUGGGGAGGCCGGAUGGGCCCGGGACGCGCCUGCCUGACCAUCACCCCCUCCUCUUGUCGCCCCACCCAGGCCUCUUCACCCUGCUCGUGGAAGAUUUCGGUAAGAGCCUUUUCUCCCUGCCGGA
Intron
Exon
Intron
−mean(ΔLogitPSI_sub)
0
2
4
6

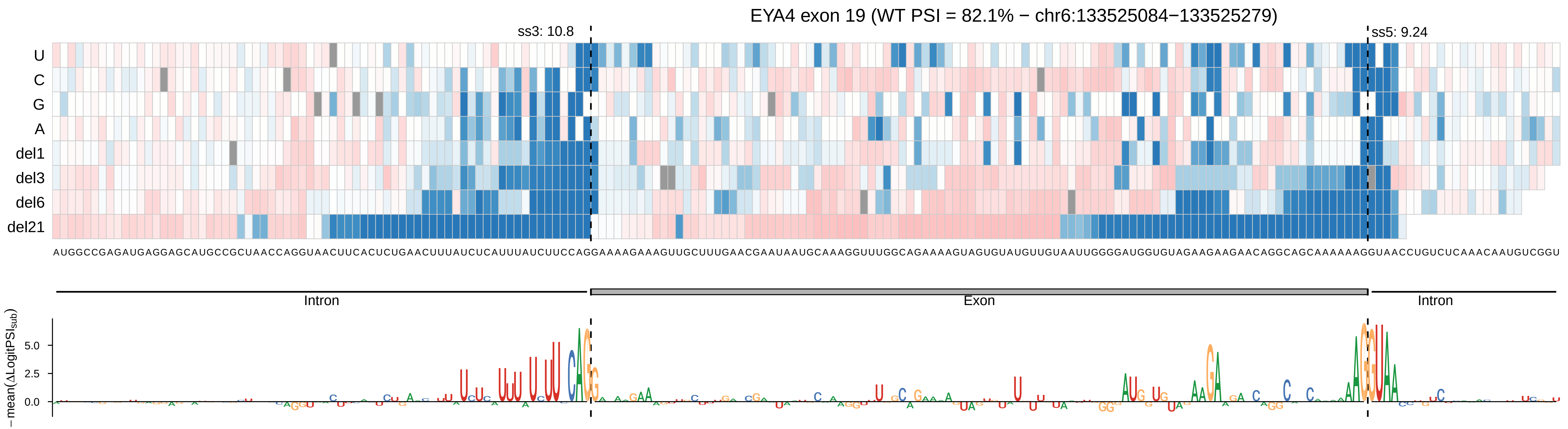

EYA4 exon 19 (WT PSI = 82.1% − chr6:133525084−133525279)
ss3: 10.8
ss5: 9.24
U
C
G
A
del1
del3
del6
del21
AUGGCCGAGAUGAGGAGCAUGCCGCUAACCAGGUAACUUCACUCUGAACUUUAUCUCAUUUAUCUUCCAGGAAAAGAAAGUUGCUUUGAACGAAUAAUGCAAAGGUUUGGCAGAAAAGUAGUGUAUGUUGUAAUUGGGGAUGGUGUAGAAGAAGAACAGGCAGCAAAAAAGGUAACCUGUCUCAAACAAUGUCGGU
Intron
Exon
Intron
−mean(ΔLogitPSI_sub)
5.0
2.5
0.0

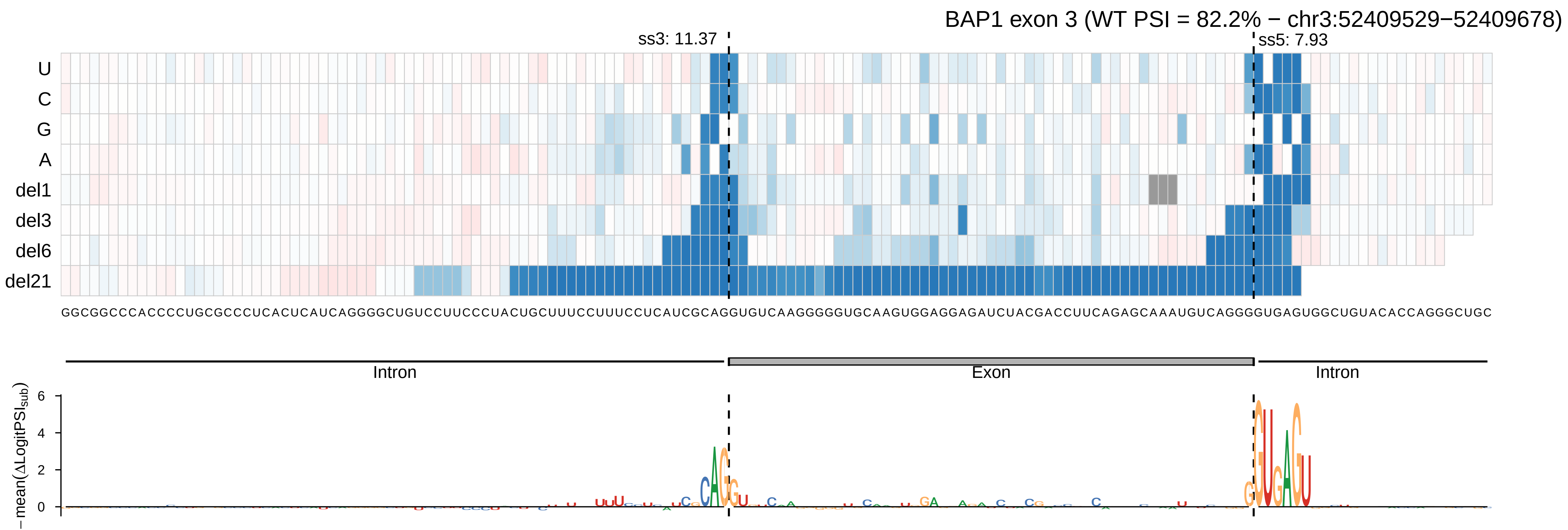

BAP1 exon 3 (WT PSI = 82.2% − chr3:52409529−52409678)
ss3: 11.37
ss5: 7.93
U
C
G
A
del1
del3
del6
del21
GGCGGCCCACCCCUGCGCCCUCACUCAUCAGGGGCUGUCCUUCCCUACUGCUUUCCUUUCCUCAUCGCAGGUGUCAAGGGGGUGCAAGUGGAGGAGAUCUACGACCUUCAGAGCAAAUGUCAGGGGUGAGUGGCUGUACACCAGGGCUGC
Intron
Exon
Intron
− mean(ΔLogitPSI_sub)
0
2
4
6

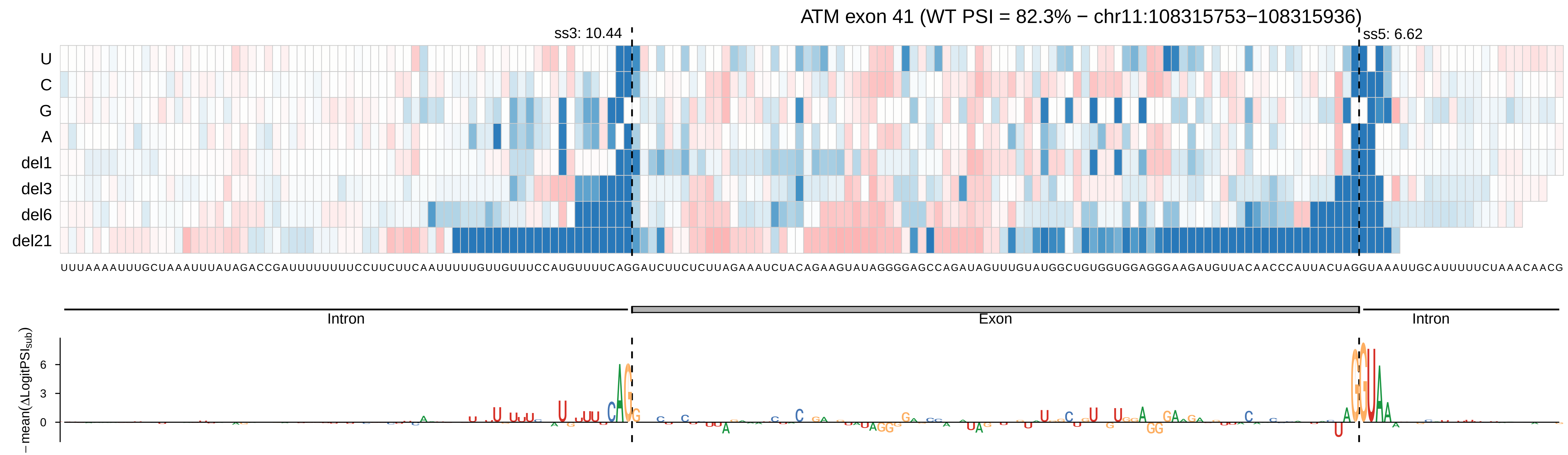
ATM exon 41 (WT PSI = 82.3% − chr11:108315753−108315936)
ss3: 10.44
ss5: 6.62
U
C
G
A
del1
del3
del6
del21
UUUAAAAUUUGCUAAAUUUAUAGACCGAUUUUUUUUCCUUCUUCAAUUUUUGUUGUUUCCAUGUUUUCAGGAUCUUCUCUUAGAAAUCUACAGAAGUAUAGGGGAGCCAGAUAGUUUGUAUGGCUGUGGUGGAGGGAAGAUGUUACAACCCAUUACUAGGUAAAUUGCAUUUUUCUAAACAACG
Intron
Exon
Intron
−mean(ΔLogitPSI_sub)
6
3
0

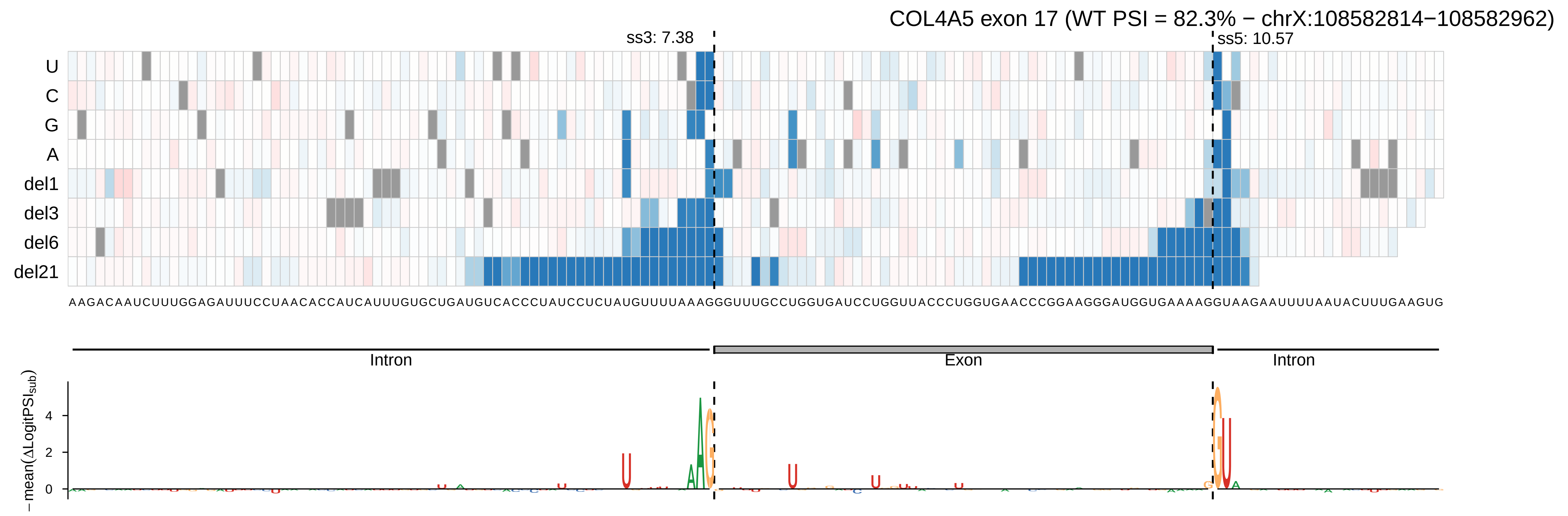

COL4A5 exon 17 (WT PSI = 82.3% − chrX:108582814−108582962)
ss3: 7.38
ss5: 10.57
U
C
G
A
del1
del3
del6
del21
AAGACAAUCUUUGGAGAUUUCCUAACACCAUCAUUUGUGCUGAUGUCACCCUAUCCUCUAUGUUUUAAAGGUUUGCCUGGUGAUCCUGGUUACCCUGGUGAACCCGGAAGGGAUGGUGAAAAGGUAAGAAUUUUAAUACUUUGAAGUG
Intron
Exon
Intron
−mean(ΔLogitPSIsub)
4
2
0

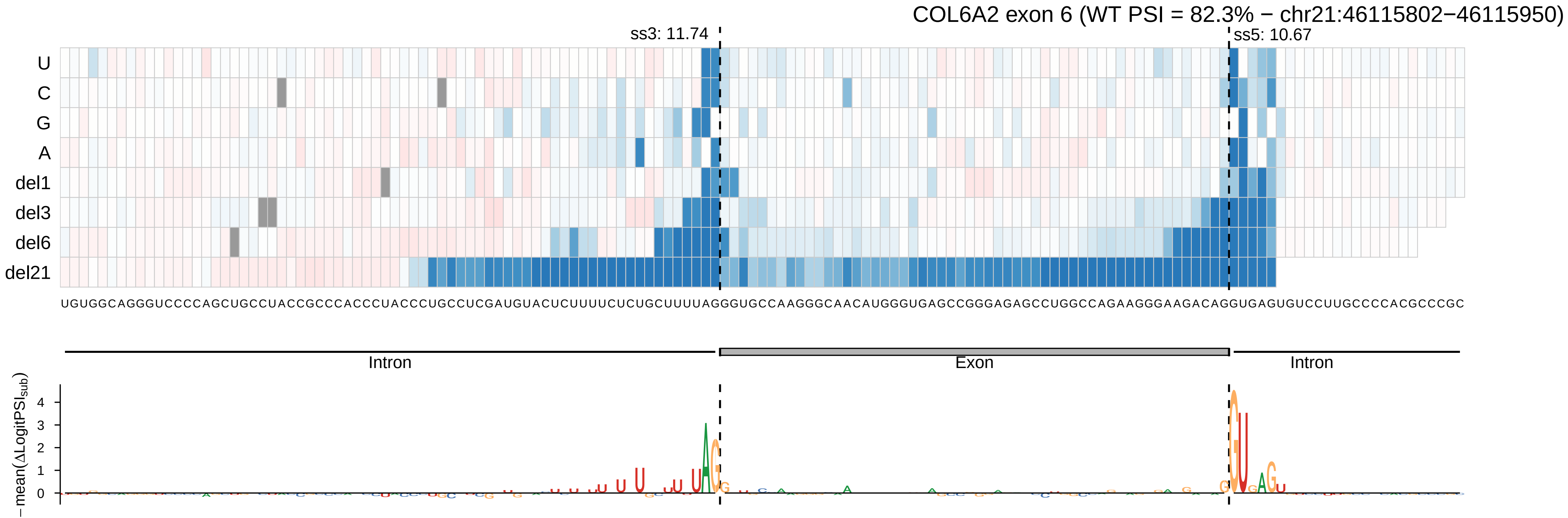

COL6A2 exon 6 (WT PSI = 82.3% − chr21:46115802−46115950)
ss3: 11.74
ss5: 10.67
U
C
G
A
del1
del3
del6
del21
UGUGGCAGGGUCCCCAGCUGCCUACCGCCCACCCUACCCUGCCUCGAUGUACUCUUUUCUCUGCUUUUAGGGUGCCAAGGGCAACAUGGGUGAGCCGGGAGAGCCUGGCCAGAAGGGAAGACAGGUGAGUGUCCUUGCCCCACGCCCGC
Intron
Exon
Intron
−mean(ΔLogitPSI$_{sub}$)
0
1
2
3
4

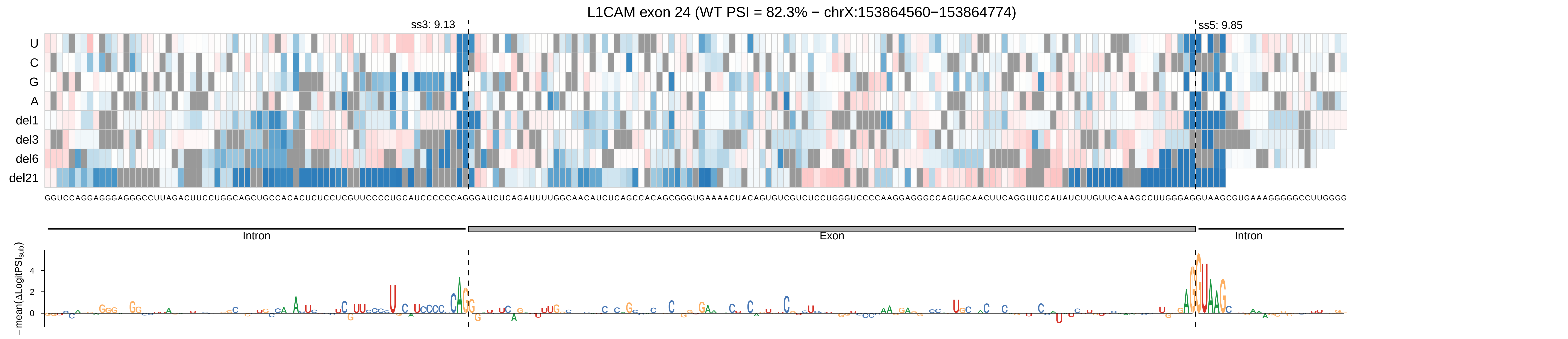

L1CAM exon 24 (WT PSI = 82.3% − chrX:153864560−153864774)
ss3: 9.13
ss5: 9.85
U
C
G
A
del1
del3
del6
del21
GGUCCAGGAGGGAGGGCCUUAGACUUCCUGGCAGCUGCCACACUCUCCUCGUUCCCCUGCAUCCCCCCAGGGAUCUCAGAUUUUGGCAACAUCUCAGCCACAGCGGGUGAAAACUACAGUGUCGUCUCCUGGGUCCCCAAGGAGGGCCAGUGCAACUUCAGGUUCCAUAUCUUGUUCAAAGCCUUGGGAGGUAAGCGUGAAAGGGGGCCUUGGGG
Intron
Exon
Intron
−mean(ΔLogitPSI_sub)
4
2
0

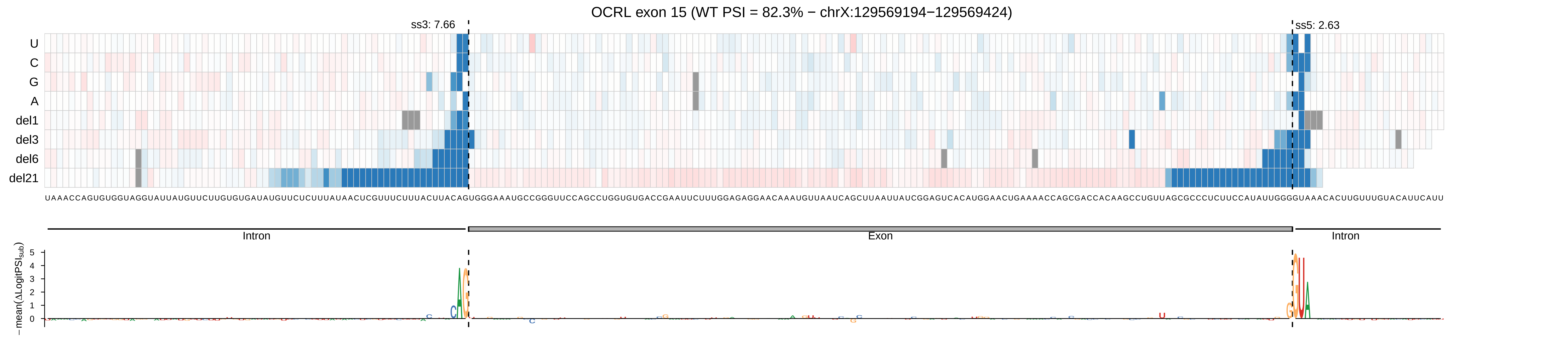
OCRL exon 15 (WT PSI = 82.3% − chrX:129569194−129569424)
ss3: 7.66
ss5: 2.63
U
C
G
A
del1
del3
del6
del21
Intron
Exon
Intron
−mean(ΔLogitPSI_sub)
5
4
3
2
1
0

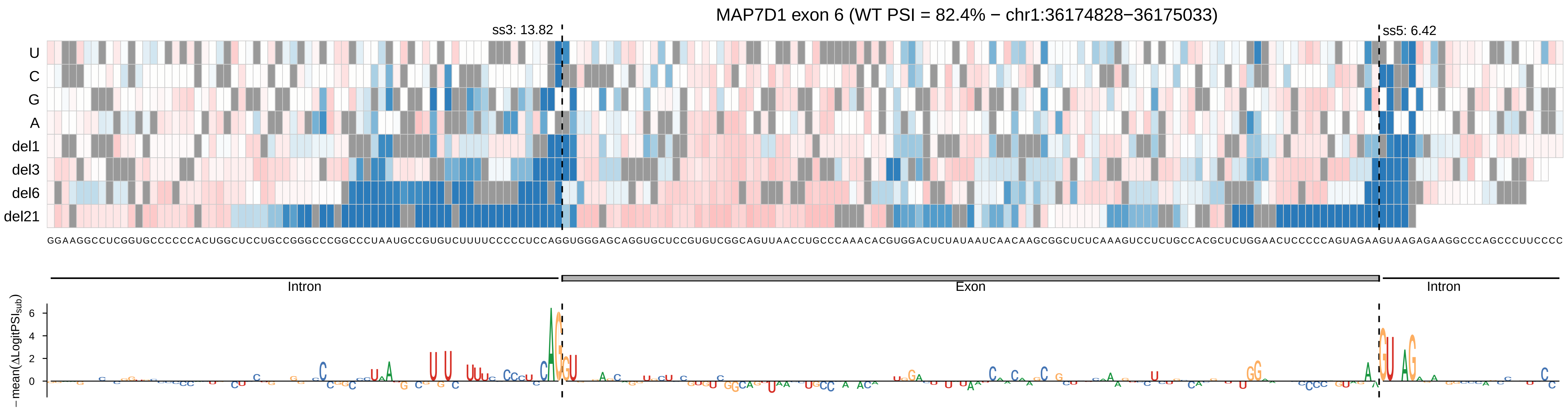

MAP7D1 exon 6 (WT PSI = 82.4% − chr1:36174828−36175033)
ss3: 13.82
ss5: 6.42
U
C
G
A
del1
del3
del6
del21
GGAAGGCCUCGGUGCCCCCCACUGGCUCCUGCCGGGCCCGGCCCUAAUGCCGUGUCUUUUCCCCCUCCAGGUGGGAGCAGGUGCUCCGUGUCGGCAGUUAACCUGCCCAAACACGUGGACUCUAUAAUCAACAAGCGGCUCUCAAAGUCCUCUGCCACGCUCUGGAACUCCCCCAGUAGAAGUAAGAGAAGGCCCAGCCCUUCCCC
Intron
Exon
Intron
−mean(ΔLogitPSI_sub)

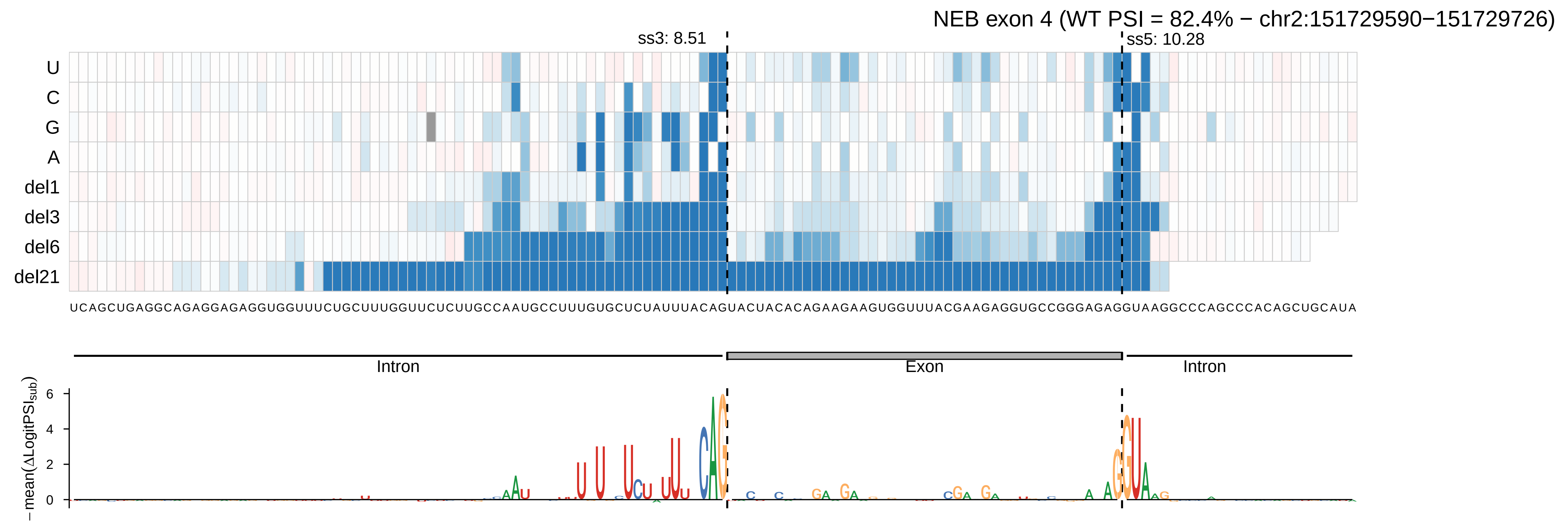

NEB exon 4 (WT PSI = 82.4% − chr2:151729590−151729726)
ss3: 8.51
ss5: 10.28
U
C
G
A
del1
del3
del6
del21
UCAGCUGAGGCAGAGGAGAGGUGGUUUCUGCUUUGGUUCUCUUGCCAAUGCCUUUGUGCUCUAUUUACAGUACUACACAGAAGAAGUGGUUUACGAAGAGGUGCCGGGAGAGGUAAGGCCCAGCCCACAGCUGCAUA
Intron
Exon
Intron
−mean(ΔLogitPSI_sub)
6
4
2
0

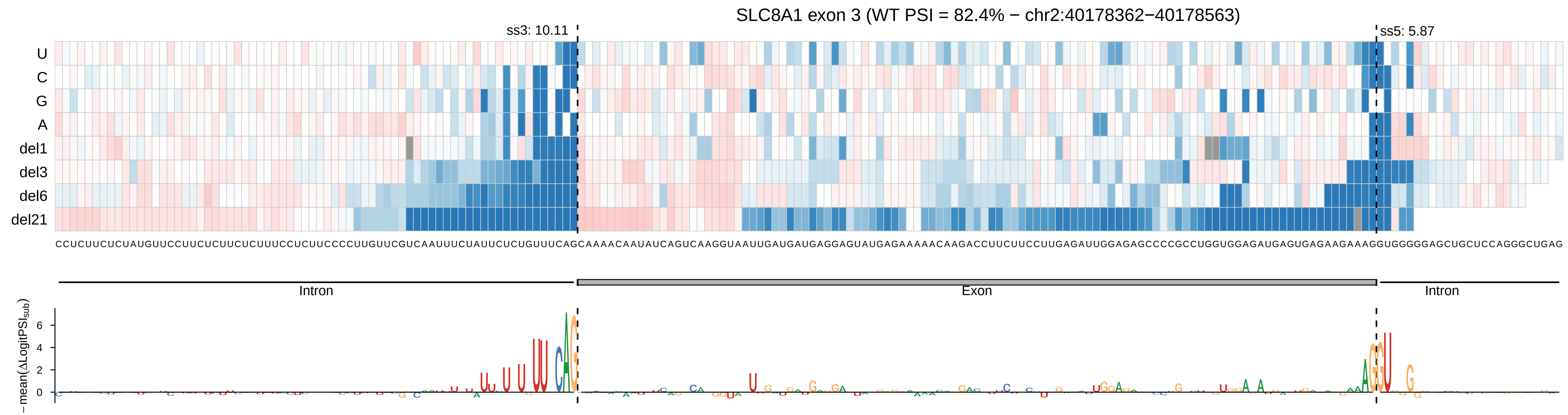

SLC8A1 exon 3 (WT PSI = 82.4% − chr2:40178362−40178563)
ss3: 10.11
ss5: 5.87
U
C
G
A
del1
del3
del6
del21
CCUCUUCUCUAUGUUCCUUCUCUUCUCUUUCCUCUUCCCCUUGUUCGUCAAUUUCUAUUCUCUGUUUCAGCAAAACAAUAUCAGUCAAGGUAAUUGAUGAUGAGGAGUAUGAGAAAAACAAGACCUUCUUCCUUGAGAUUGGAGAGCCCCGCCUGGUGGAGAUGAGUGAGAAGAAAGGUGGGGGAGCUGCUCCAGGGCUGAG
Intron
Exon
Intron
− mean(ΔLogitPSI_sub)
6
4
2
0

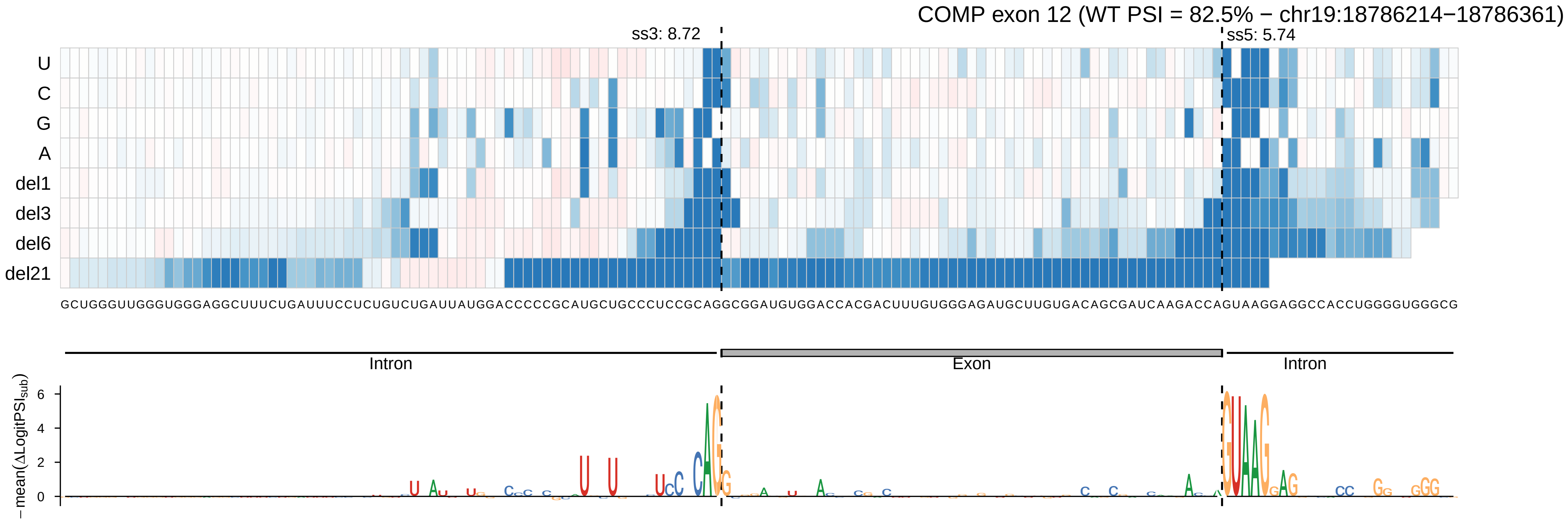
COMP exon 12 (WT PSI = 82.5% − chr19:18786214−18786361)
ss3: 8.72
ss5: 5.74
U
C
G
A
del1
del3
del6
del21
GCUGGGUUGGGUGGGAGGCUUUCUGAUUUCCUCUGUCUGAUUAUGGACCCCCGCAUGCUGCCCUCCGCAGGCGGAUGUGGACCACGACUUUGUGGGAGAUGCUUGUGACAGCGAUCAAGACCAGUAAGGAGGCCACCUGGGGUGGGCG
Intron
Exon
Intron
−mean(ΔLogitPSI_sub)
6
4
2
0

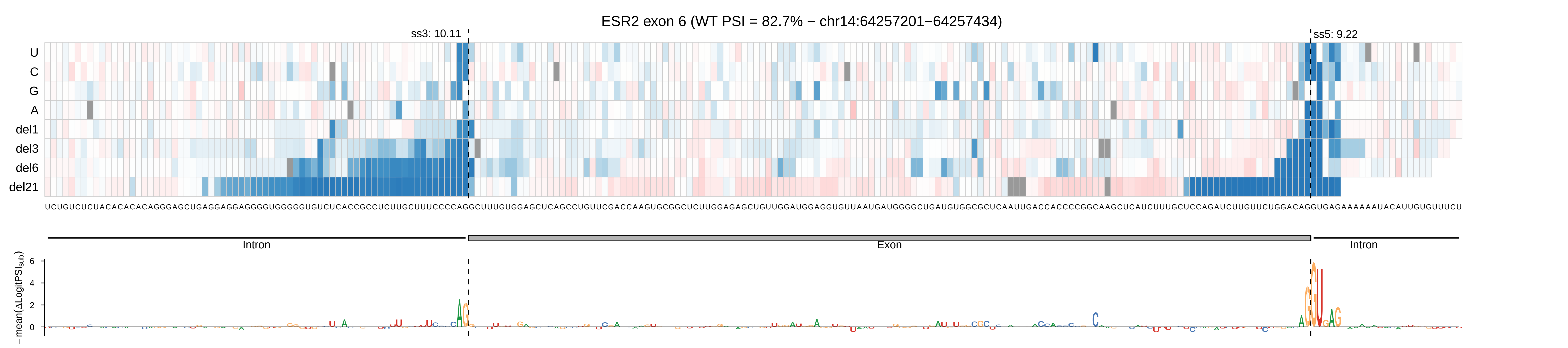

ESR2 exon 6 (WT PSI = 82.7% − chr14:64257201−64257434)
ss3: 10.11
ss5: 9.22
U
C
G
A
del1
del3
del6
del21
Intron
Exon
Intron
− mean(ΔLogitPSI_sub)
6
4
2
0

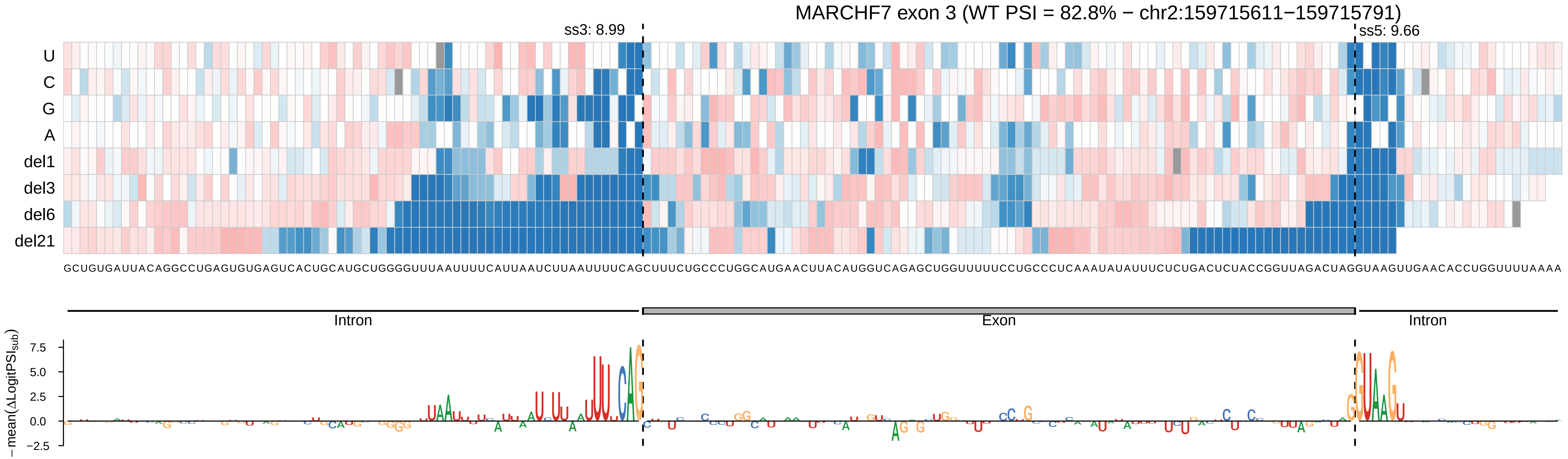
MARCHF7 exon 3 (WT PSI = 82.8% − chr2:159715611−159715791)
ss3: 8.99
ss5: 9.66
U
C
G
A
del1
del3
del6
del21
GCUGUGAUUACAGGCCUGAGUGUGAGUCACUGCAUGCUGGGGUUUAAUUUUCAUUAAUCUUAAUUUUCAGCUUUCUGCCCUGGCAUGAACUUACAUGGUCAGAGCUGGUUUUUCCUGCCCUCAAAUAUAUUUCUCUGACUCUACCGGUUAGACUAGGUAAGUUGAACACCUGGUUUUAAAA
Intron
Exon
Intron
− mean(ΔLogitPSI_sub)
7.5
5.0
2.5
0.0
−2.5

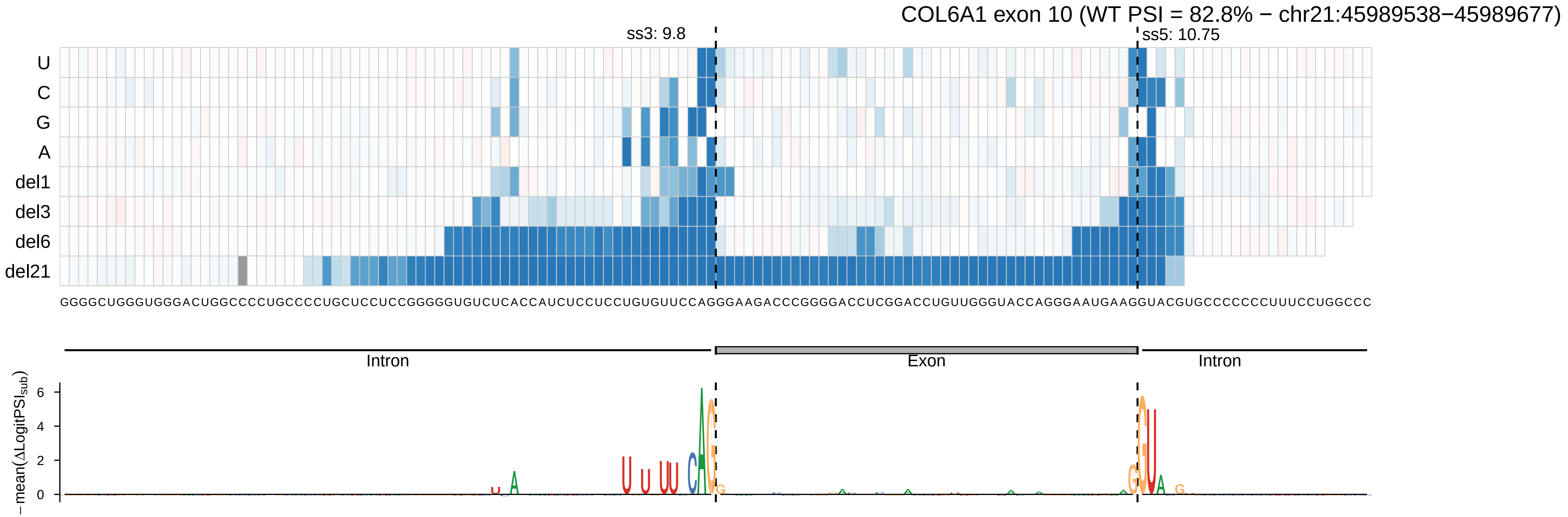

COL6A1 exon 10 (WT PSI = 82.8% − chr21:45989538−45989677)
ss3: 9.8
ss5: 10.75
U
C
G
A
del1
del3
del6
del21
GGGGCUGGGUGGGACUGGCCCCUGCCCCUGCUCCUCCGGGGGUGUCUCACCAUCUCCUCCUGUGUUCCAGGGAAGACCCGGGGACCUCGGACCUGUUGGGUACCAGGGAAUGAAGGUACGUGCCCCCCUUUCCUGGCCC
Intron
Exon
Intron
− mean(ΔLogitPSI_sub)
6
4
2
0

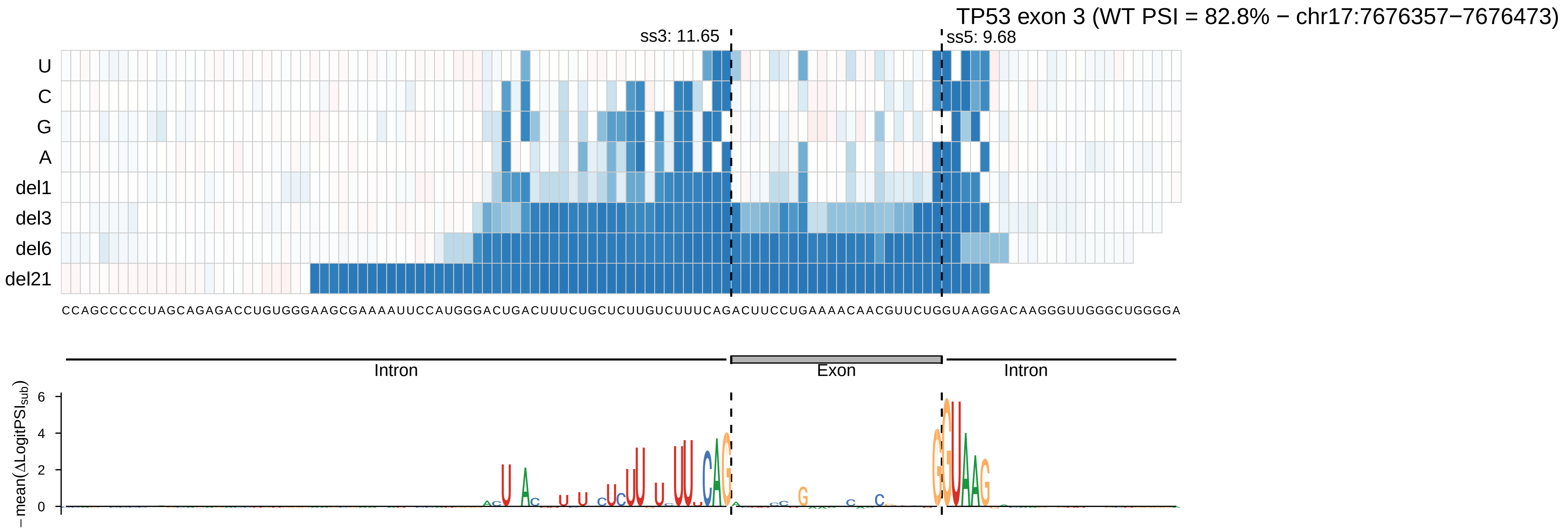
TP53 exon 3 (WT PSI = 82.8% − chr17:7676357−7676473)
ss3: 11.65
ss5: 9.68
U
C
G
A
del1
del3
del6
del21
CCAGCCCCCUAGCAGAGACCUGUGGGAAGCGAAAAUUCCAUGGGACUGACUUUCUGCUCUUGUCUUUCAGACUUCCUGAAAACAACGUUCUGGUAAGGACAAGGGUUGGGCUGGGGA
Intron
Exon
Intron
− mean(ΔLogitPSI_sub)
6
4
2
0

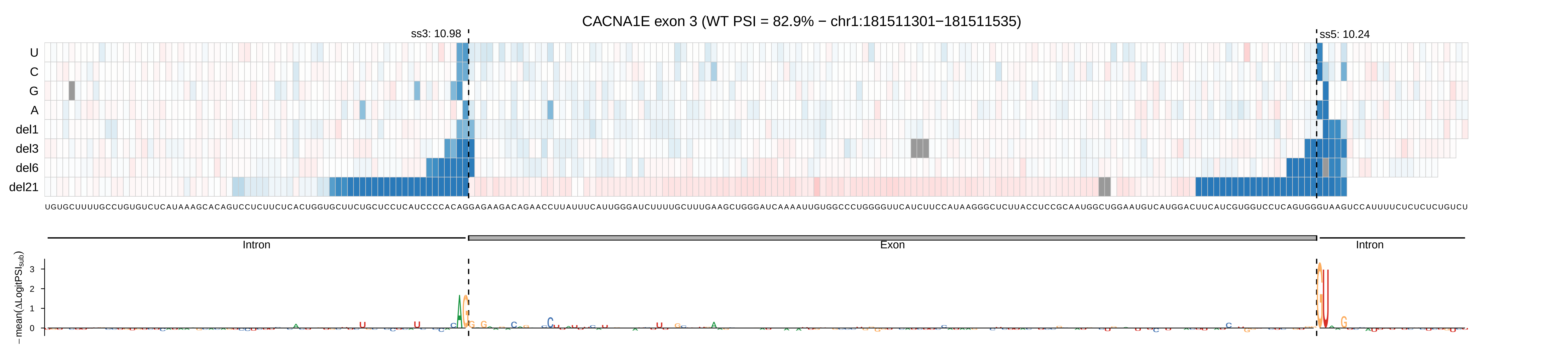
CACNA1E exon 3 (WT PSI = 82.9% − chr1:181511301−181511535)
ss3: 10.98
ss5: 10.24
U
C
G
A
del1
del3
del6
del21
Intron
Exon
Intron
−mean(ΔLogitPSI_sub)
3
2
1
0

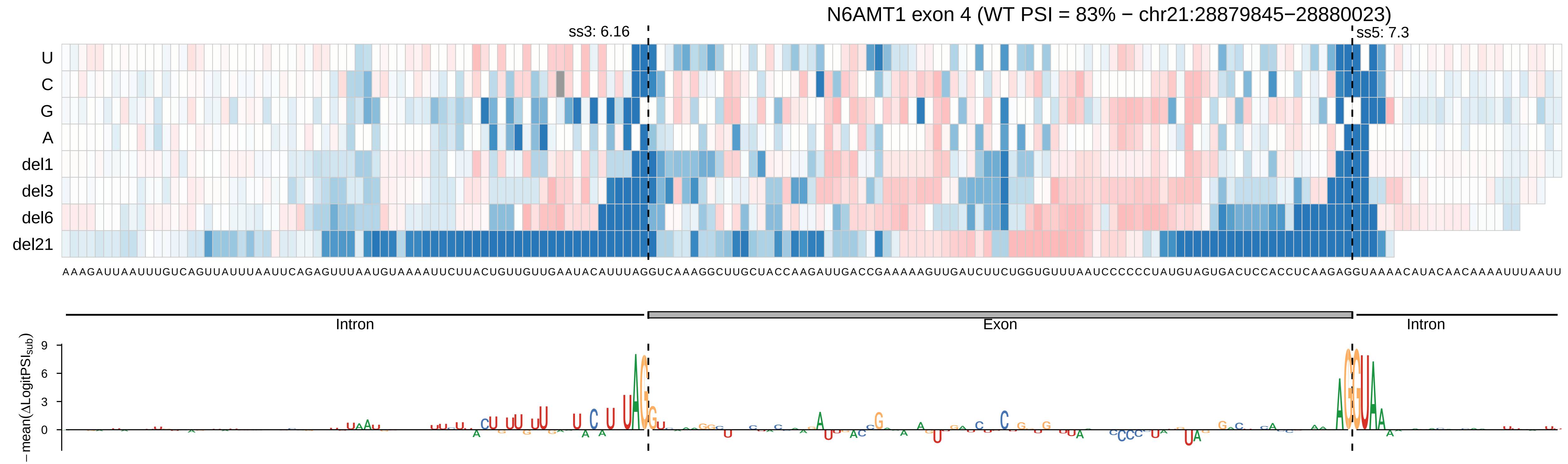
N6AMT1 exon 4 (WT PSI = 83% − chr21:28879845−28880023)
ss3: 6.16
ss5: 7.3
U
C
G
A
del1
del3
del6
del21
AAAGAUUAAUUUGUCAGUUAUUUAAUUCAGAGUUUAAUGUAAAAUUCUUACUGUUGUUGAAUACAUUUAGGUCAAAGGCUUGCUACCAAGAUUGACCGAAAAAGUUGAUCUUCUGGUGUUUAAUCCCCCCUAUGUAGUGACUCCACCUCAAGAGGUAAAACAUACAACAAAAUUUAAUU
Intron
Exon
Intron
−mean(ΔLogitPSIsub)
9
6
3
0

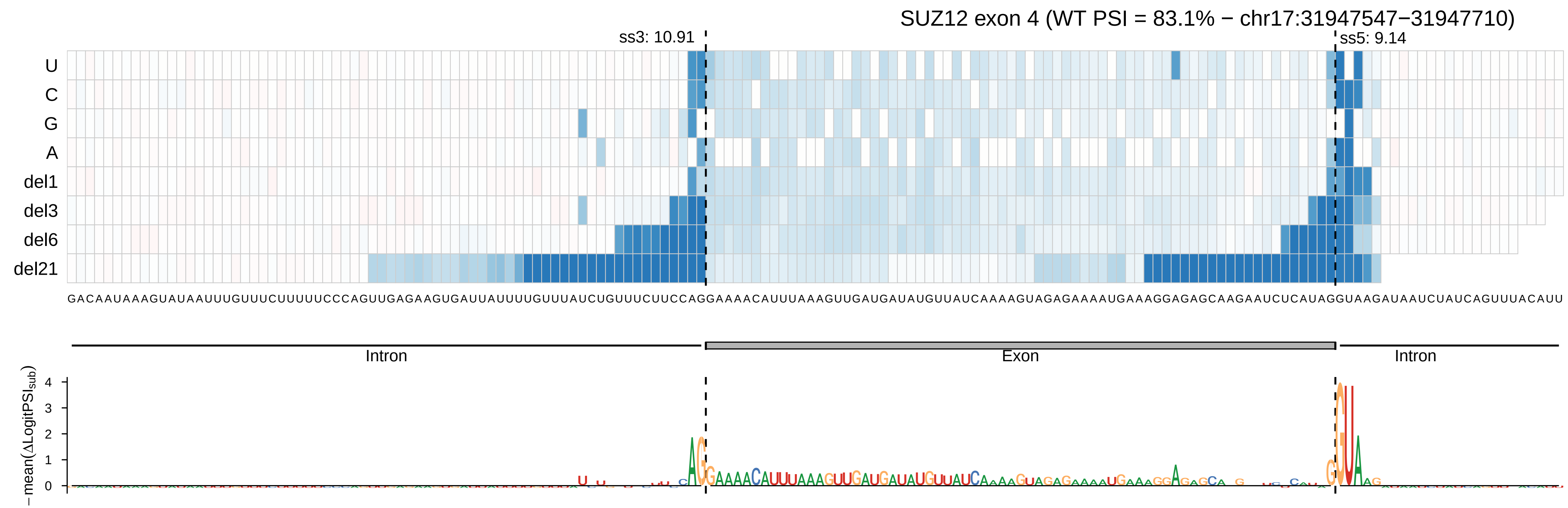

SUZ12 exon 4 (WT PSI = 83.1% − chr17:31947547−31947710)
ss3: 10.91
ss5: 9.14
U
C
G
A
del1
del3
del6
del21
GACAAUAAAGUAUAAUUUGUUUCUUUUUCCCAGUUGAGAAGUGAUUAUUUUGUUUAUCUGUUUCUUCCAGGAAAACAUUUAAAGUUGAUGAUAUGUUAUCAAAAGUAGAGAAAAUGAAAGGAGAGCAAGAAUCUCAUAGGUAAGAUAAUCUAUCAGUUUACAUU
Intron
Exon
Intron
− mean(ΔLogitPSI_sub)
0
1
2
3
4

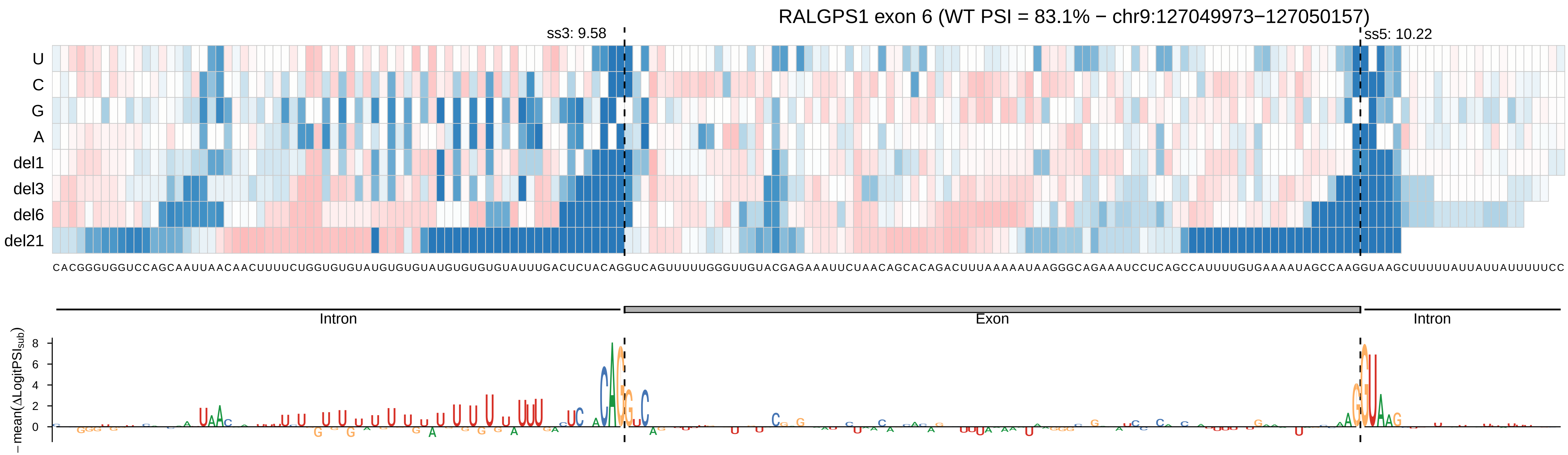
RALGPS1 exon 6 (WT PSI = 83.1% − chr9:127049973−127050157)
ss3: 9.58
ss5: 10.22
U
C
G
A
del1
del3
del6
del21
Intron
Exon
Intron
−mean(ΔLogitPSI_sub)
8
6
4
2
0

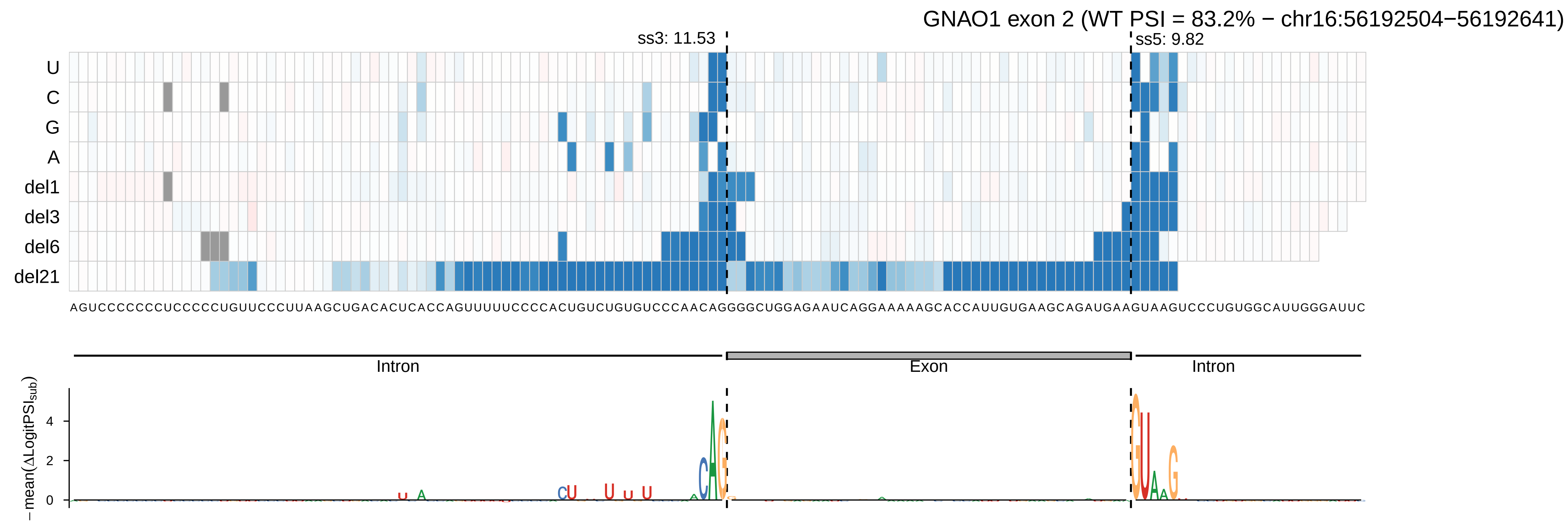

GNAO1 exon 2 (WT PSI = 83.2% − chr16:56192504−56192641)
ss3: 11.53
ss5: 9.82
U
C
G
A
del1
del3
del6
del21
AGUCCCCCCCUCCCCCUGUUCCCUUAAGCUGACACUCACCAGUUUUUCCCCACUGUCUGUGUCCCAACAGGGGCUGGAGAAUCAGGAAAAAGCACCAUUGUGAAGCAGAUGAAGUAAGUCCCUGUGGCAUUGGGAUUC
Intron
Exon
Intron
− mean(ΔLogitPSI$_{sub}$)
4
2
0

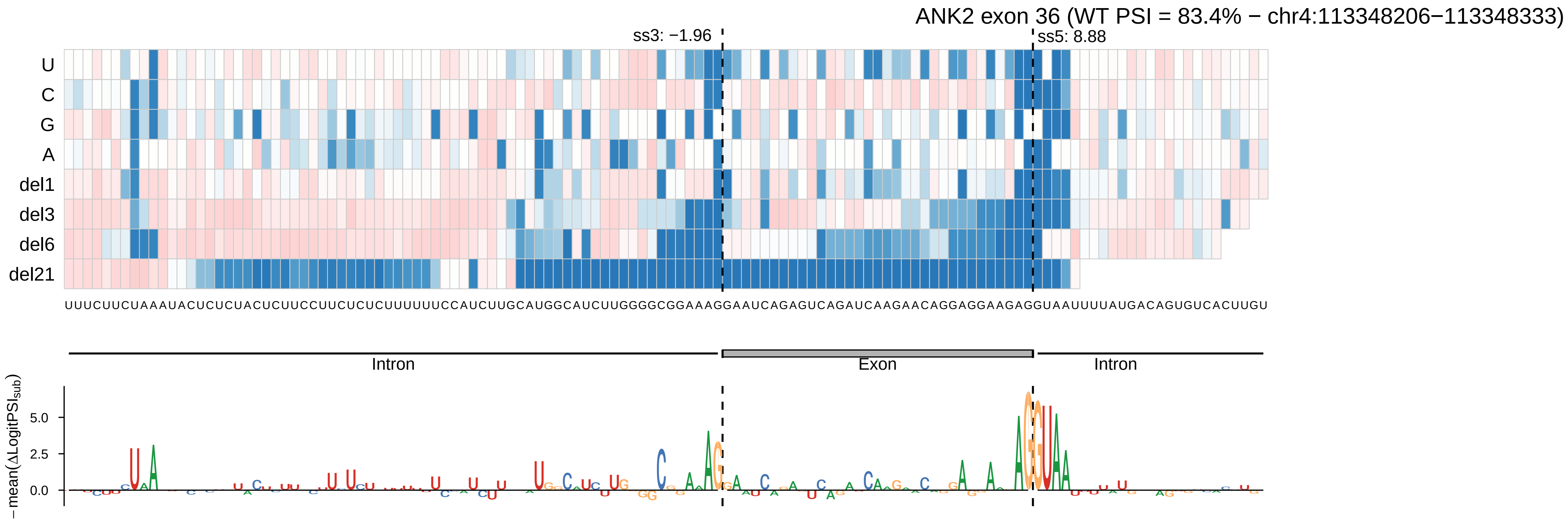

ANK2 exon 36 (WT PSI = 83.4% − chr4:113348206−113348333)
ss3: −1.96
ss5: 8.88
U
C
G
A
del1
del3
del6
del21
UUUCUUCUAAAUACUCUCUACUCUUCCUUCUCUCUUUUUUCCAUCUUGCAUGGCAUCUUGGGGCGGAAAGGAAUCAGAGUCAGAUCAAGAACAGGAGGAAGAGGUAAUUUUAUGACAGUGUCACUUGU
Intron
Exon
Intron
−mean(ΔLogitPSI_sub)
5.0
2.5
0.0

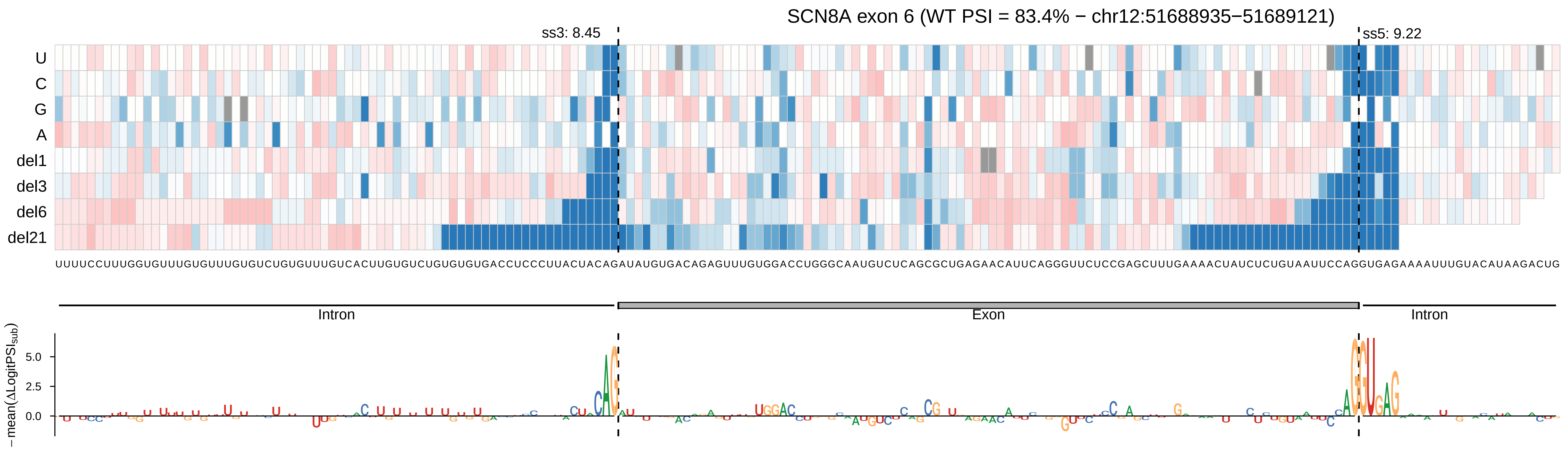

SCN8A exon 6 (WT PSI = 83.4% − chr12:51688935−51689121)
ss3: 8.45
ss5: 9.22
U
C
G
A
del1
del3
del6
del21
UUUUCCUUUGGUGUUUGUGUUUGUGUCUGUGUUUGUCACUUGUGUCUGUGUGUGACCUCCCUUACUACAGAUAUGUGACAGAGUUUGUGGACCUGGGCAAUGUCUCAGCGCUGAGAACAUUCAGGGUUCUCCGAGCUUUGAAAACUAUCUCUGUAAUUCCAGGUGAGAAAAUUUGUACAUAAGACUG
Intron
Exon
Intron
−mean(ΔLogitPSI$_{sub}$)
5.0
2.5
0.0

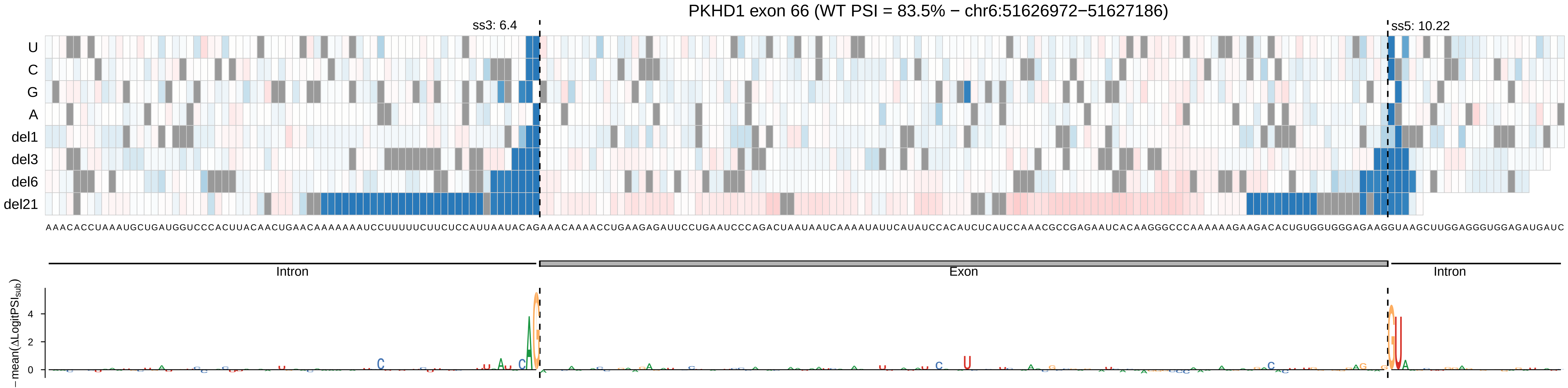

PKHD1 exon 66 (WT PSI = 83.5% − chr6:51626972−51627186)
ss3: 6.4
ss5: 10.22
U
C
G
A
del1
del3
del6
del21
AAACACCUAAAUGCUGAUGGUCCCACUUACAACUGAACAAAAAAAUCCUUUUUCUUCUCCAUUAAUACAGAAACAAAACCUGAAGAGAUUCCUGAAUCCCAGACUAAUAAUCAAAAUAUUCAUAUCCACAUCUCAUCCAAACGCCGAGAAUCACAAGGGCCCAAAAAAGAAGACACUGUGGUGGGAGAAGGUAAGCUUGGAGGGUGGAGAUGAUC
Intron
Exon
Intron
−mean(ΔLogitPSI_sub)

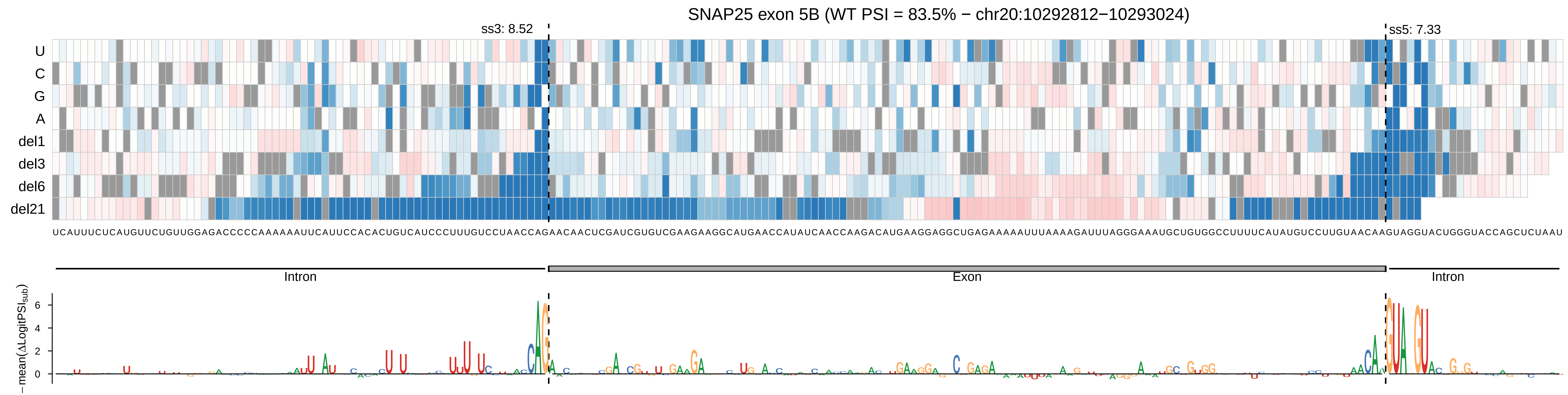
SNAP25 exon 5B (WT PSI = 83.5% − chr20:10292812−10293024)
ss3: 8.52
ss5: 7.33
U
C
G
A
del1
del3
del6
del21
Intron
Exon
Intron
−mean(ΔLogitPSI_sub)
6
4
2
0

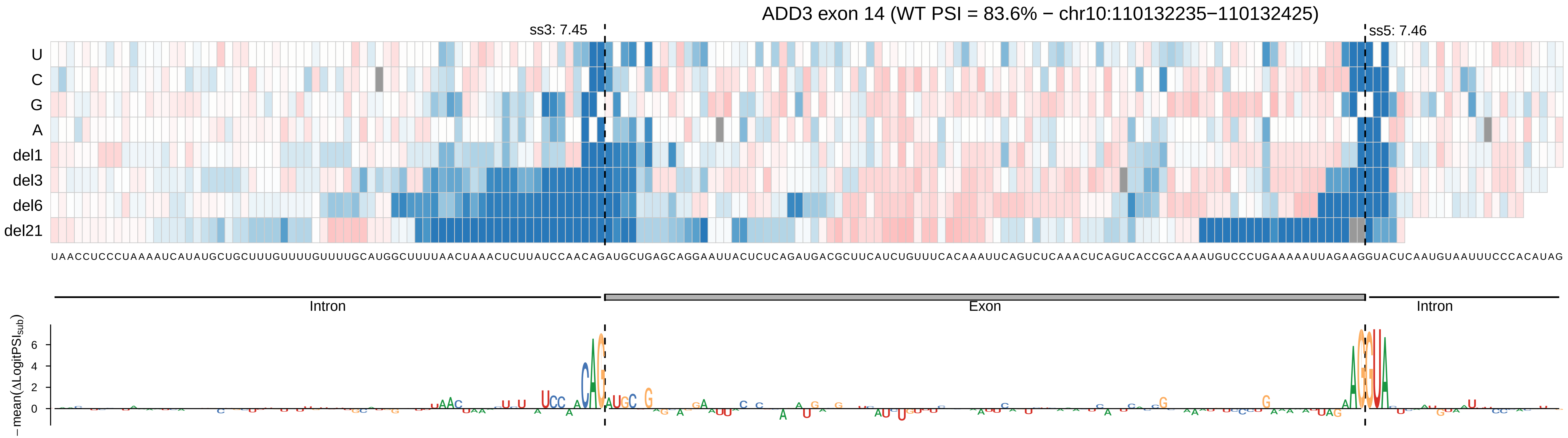
ADD3 exon 14 (WT PSI = 83.6% − chr10:110132235−110132425)
ss3: 7.45
ss5: 7.46
U
C
G
A
del1
del3
del6
del21
UAACCUCCCUAAAAUCAUAUGCUGCUUUGUUUUGUUUUGCAUGGCUUUUAACUAAACUCUUAUCCAACAGAUGCUGAGCAGGAAUUACUCUCAGAUGACGCUUCAUCUGUUUCACAAAUUCAGUCUCAAACUCAGUCACCGCAAAAUGUCCCUGAAAAAUUAGAAGGUACUCAAUGUAAUUUCCCACAUAG
Intron
Exon
Intron
−mean(ΔLogitPSI_sub)
6
4
2
0

SPAST exon 13 (WT PSI = 83.7% − chr2:32141834−32141971)

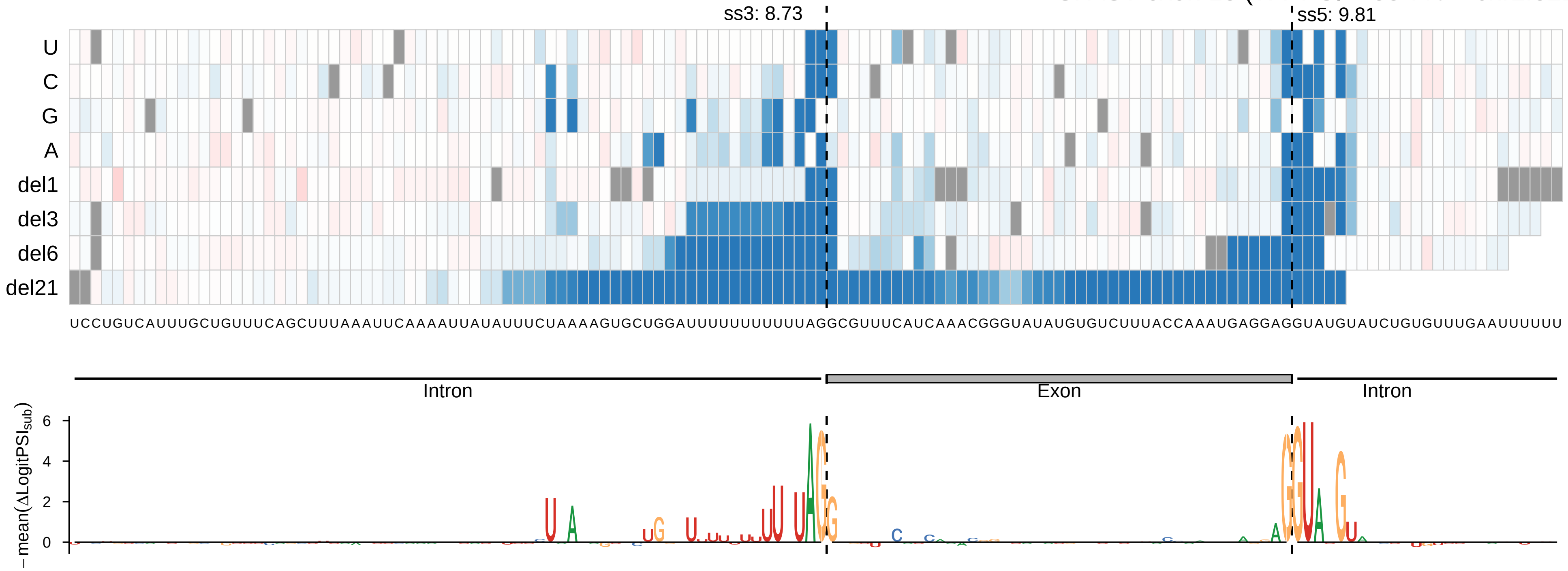

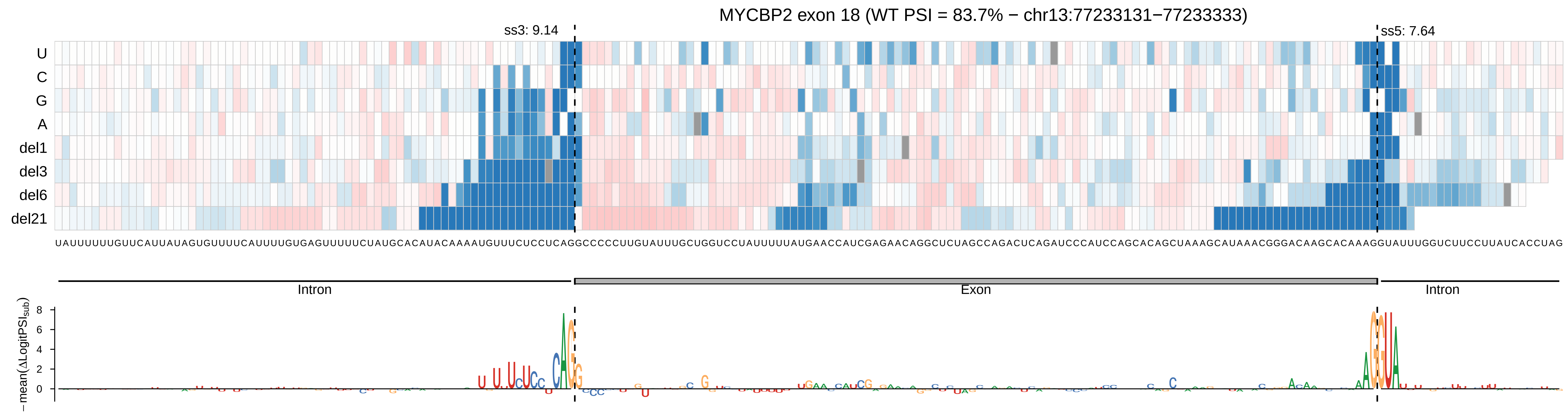

MYCBP2 exon 18 (WT PSI = 83.7% − chr13:77233131−77233333)
ss3: 9.14
ss5: 7.64
U
C
G
A
del1
del3
del6
del21
UAUUUUUUGUUCAUUAUAGUGUUUUCAUUUUGUGAGUUUUUCUAUGCACAUACAAAAUGUUUCUCCUCAGGCCCCCUUGUAUUUGCUGGUCCUAUUUUUAUGAACCAUCGAGAACAGGCUCUAGCCAGACUCAGAUCCCAUCCAGCACAGCUAAAGCAUAAACGGGACAAGCACAAAGGUAUUUGGUCUUCCUUAUCACCUAG
Intron
Exon
Intron
−mean(ΔLogitPSI_sub)

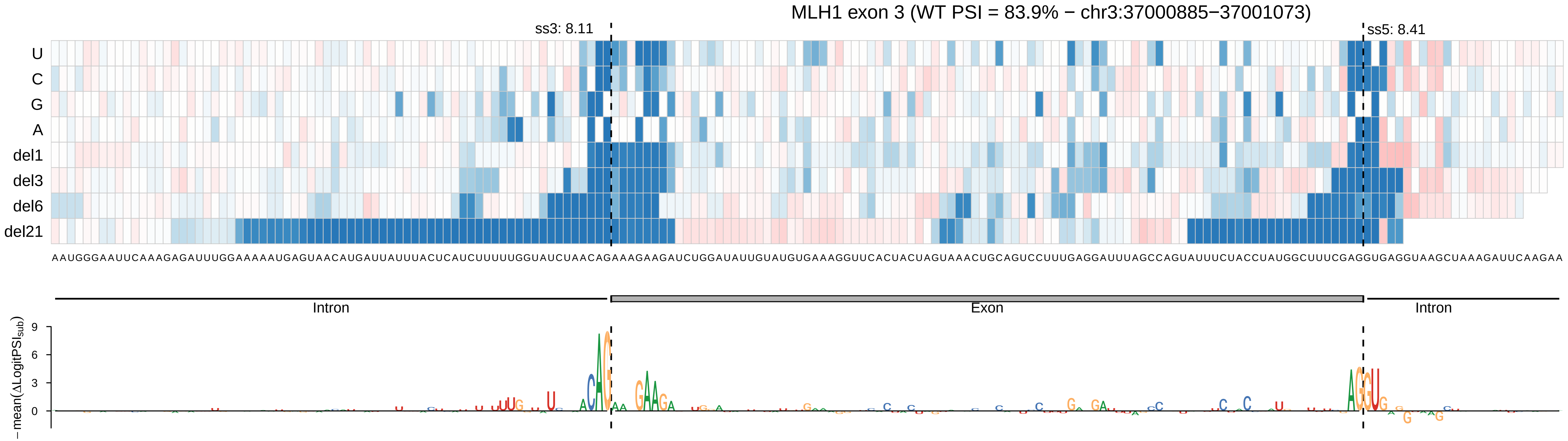

MLH1 exon 3 (WT PSI = 83.9% − chr3:37000885−37001073)
ss3: 8.11
ss5: 8.41
U
C
G
A
del1
del3
del6
del21
AAUGGGAAUUCAAAGAGAUUUGGAAAAAUGAGUAACAUGAUUAUUUACUCAUCUUUUUGGUAUCUAACAGAAAGAAGAUCUGGAUAUUGUAUGUGAAAGGUUCACUACUAGUAAACUGCAGUCCUUUGAGGAUUUAGCCAGUAUUUCUACCUAUGGCUUUCGAGGUGAGGUAAGCUAAAGAUUCAAGAA
Intron
Exon
Intron
−mean(ΔLogitPSI$_{sub}$)
9
6
3
0

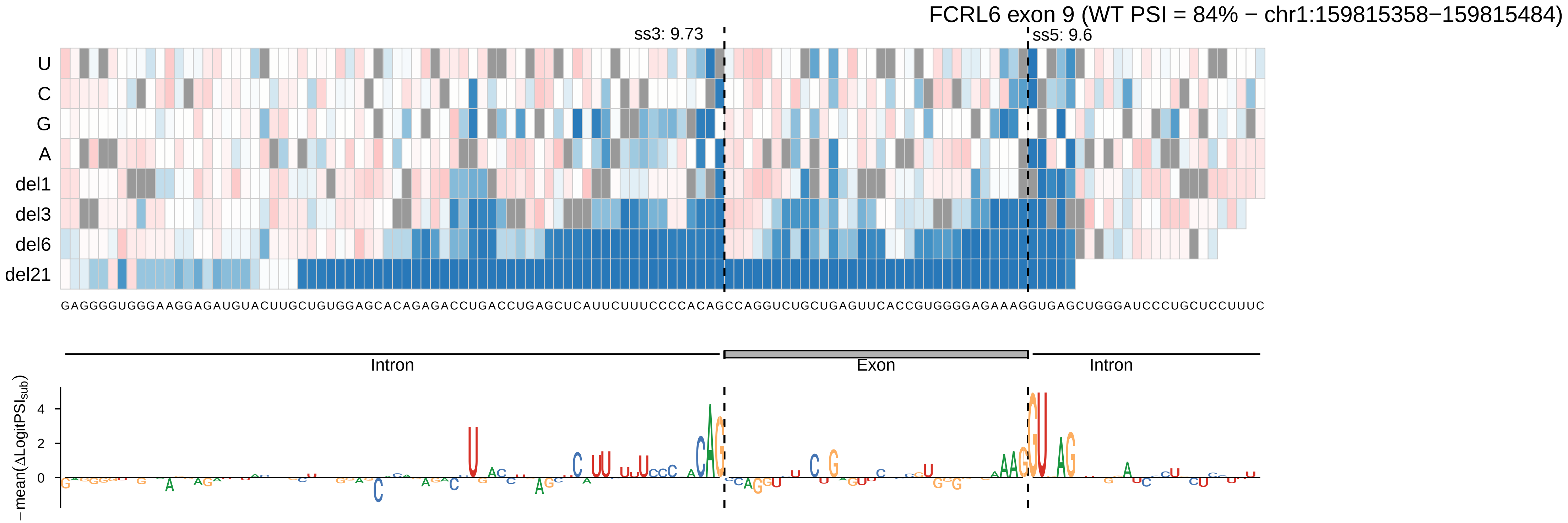
FCRL6 exon 9 (WT PSI = 84% − chr1:159815358−159815484)
ss3: 9.73
ss5: 9.6
U
C
G
A
del1
del3
del6
del21
GAGGGGUGGGAAGGAGAUGUACUUGCUGUGGAGCACAGAGACCUGACCUGAGCUCAUUCUUUCCCCACAGCCAGGUCUGCUGAGUUCACCGUGGGGAGAAAGGUGAGCUGGGAUCCCUGCUCCUUUC
Intron
Exon
Intron
−mean(ΔLogitPSI_sub)
4
2
0

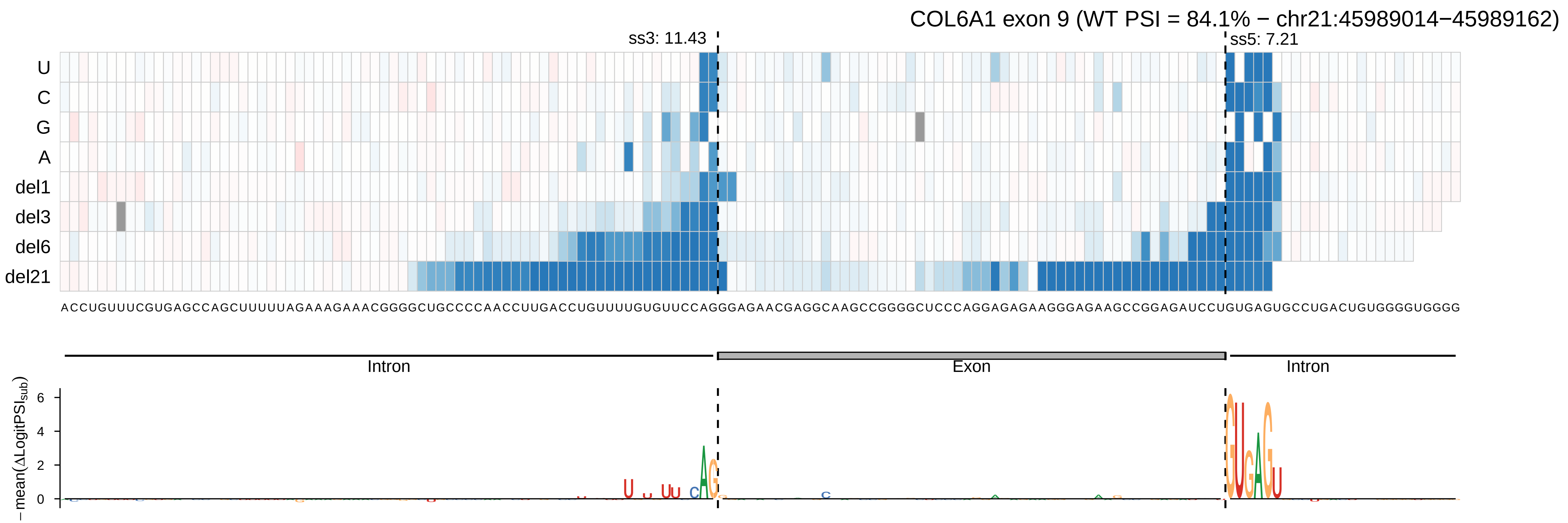
COL6A1 exon 9 (WT PSI = 84.1% − chr21:45989014−45989162)
ss3: 11.43
ss5: 7.21
U
C
G
A
del1
del3
del6
del21
ACCUGUUUCGUGAGCCAGCUUUUUAGAAAGAAACGGGGCUGCCCCAACCUUGACCUGUUUUGUGUUCCAGGGAGAACGAGGCAAGCCGGGGCUCCCAGGAGAGAAGGGAGAAGCCGGAGAUCCUGUGAGUGCCUGACUGUGGGGUGGGG
Intron
Exon
Intron
− mean(ΔLogitPSI_sub)
6
4
2
0

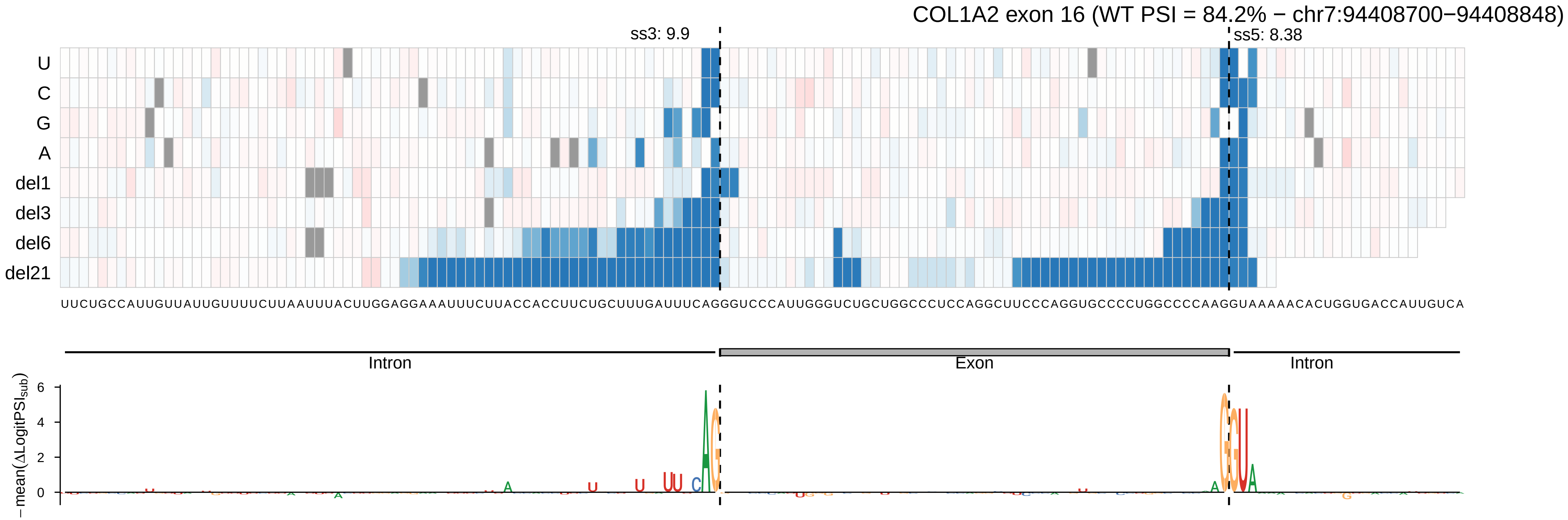

COL1A2 exon 16 (WT PSI = 84.2% − chr7:94408700−94408848)
ss3: 9.9
ss5: 8.38
U
C
G
A
del1
del3
del6
del21
UUCUGCCAUUGUUAUUGUUUCUUAAUUUACUUGGAGGAAAUUUCUUACCACCUUCUGCUUUGAUUUCAGGGUCCCAUUGGGUCUGCUGGCCCUCCAGGCUUCCCAGGUGCCCCUGGCCCCAAGGUAAAAACACUGGUGACCAUUGUCA
Intron
Exon
Intron
−mean(ΔLogitPSI_sub)
6
4
2
0

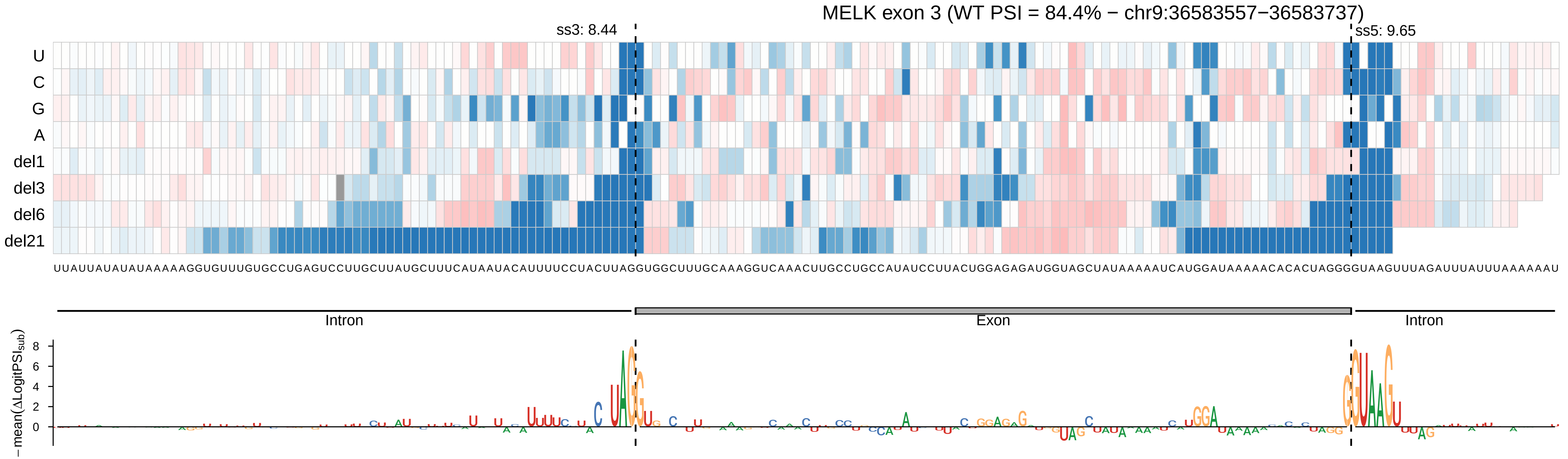

MELK exon 3 (WT PSI = 84.4% − chr9:36583557−36583737)
ss3: 8.44
ss5: 9.65
U
C
G
A
del1
del3
del6
del21
UUAUUAUAUAUAAAAAGGUGUUUGUGCCUGAGUCCUUGCUUAUGCUUUCAUAAUACAUUUUCCUACUUAGGUGGCUUUGCAAAGGUCAAACUUGCCUGCCAUAUCCUUACUGGAGAGAUGGUAGCUAUAAAAAUCAUGGAUAAAAACACACUAGGGGUAAGUUUAGAUUUAUUUAAAAAAU
Intron
Exon
Intron
−mean(ΔLogitPSI_sub)
8
6
4
2
0

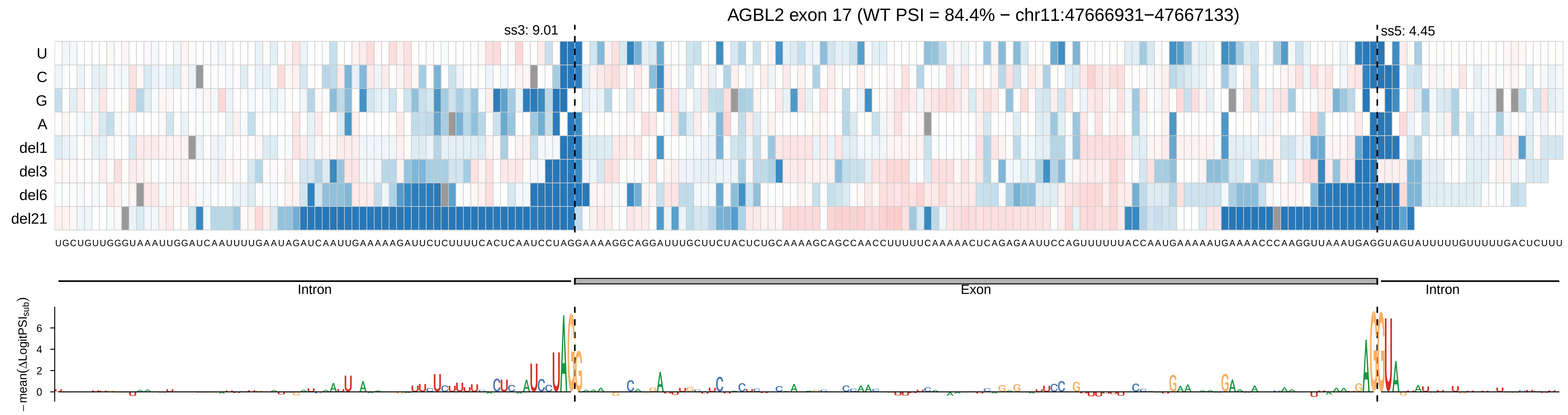
AGBL2 exon 17 (WT PSI = 84.4% − chr11:47666931−47667133)
ss3: 9.01
ss5: 4.45
U
C
G
A
del1
del3
del6
del21
Intron
Exon
Intron
−mean(ΔLogitPSI_sub)
6
4
2
0

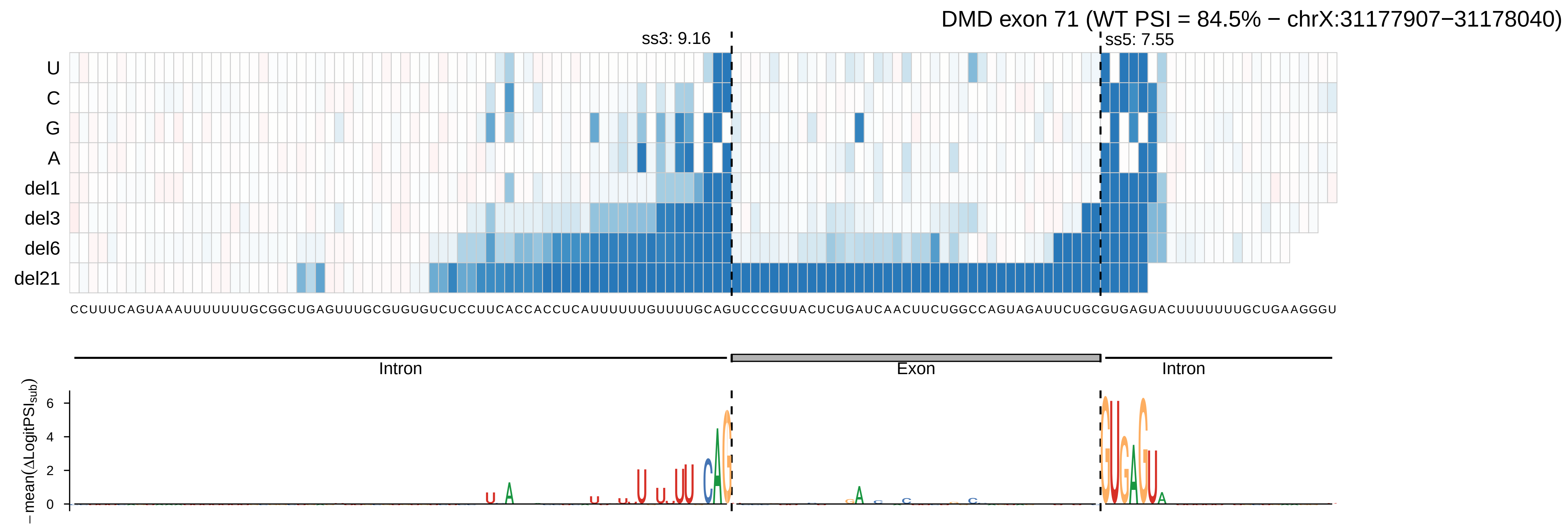

DMD exon 71 (WT PSI = 84.5% − chrX:31177907−31178040)
ss3: 9.16
ss5: 7.55
U
C
G
A
del1
del3
del6
del21
CCUUUCAGUAAAUUUUUUUGCGGCUGAGUUUGCGUGUGUCUCCUUCACCACCUCAUUUUUUGUUUUGCAGUCCCGUUACUCUGAUCAACUUCUGGCCAGUAGAUUCUGCGUGAGUACUUUUUUUGCUGAAGGGU
Intron
Exon
Intron
− mean(ΔLogitPSI_sub)
6
4
2
0

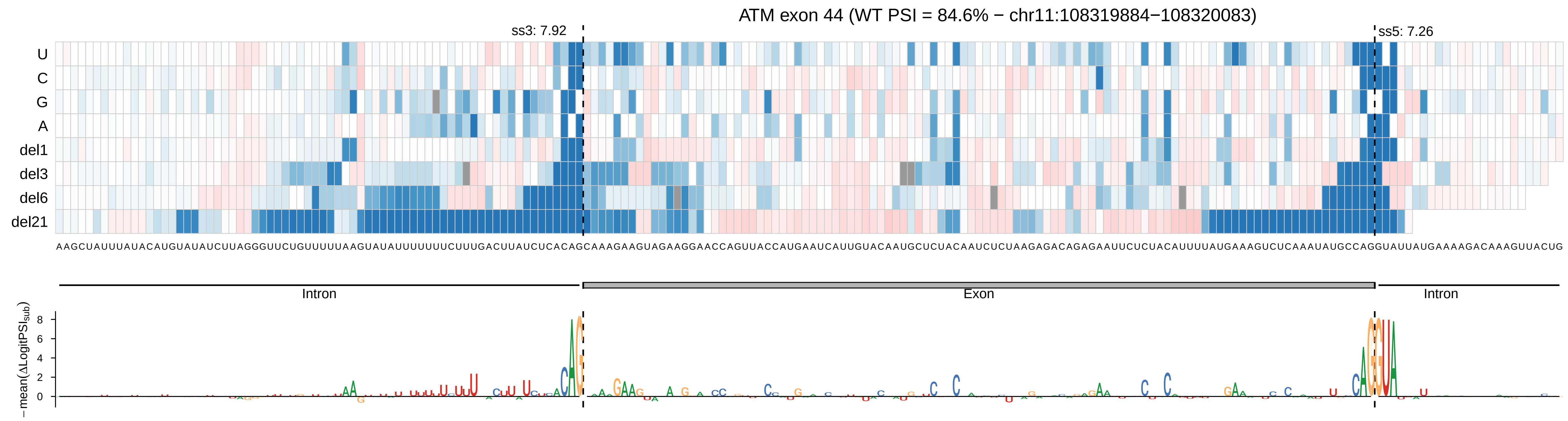
ATM exon 44 (WT PSI = 84.6% − chr11:108319884−108320083)
ss3: 7.92
ss5: 7.26
U
C
G
A
del1
del3
del6
del21
AAGCUAUUUAUACAUGUAUAUCUUAGGGUUCUGUUUUUAAGUAUAUUUUUUUCUUUGACUUAUCUCACAGCAAAGAAGUAGAAGGAACCAGUUACCAUGAAUCAUUGUACAAUGCUCUACAAUCUCUAAGAGACAGAGAAUUCUCUACAUUUUAUGAAAGUCUCAAAUAUGCCAGGUAUUAUGAAAAGACAAAGUUACUG
Intron
Exon
Intron
− mean(ΔLogitPSI_sub)
8
6
4
2
0

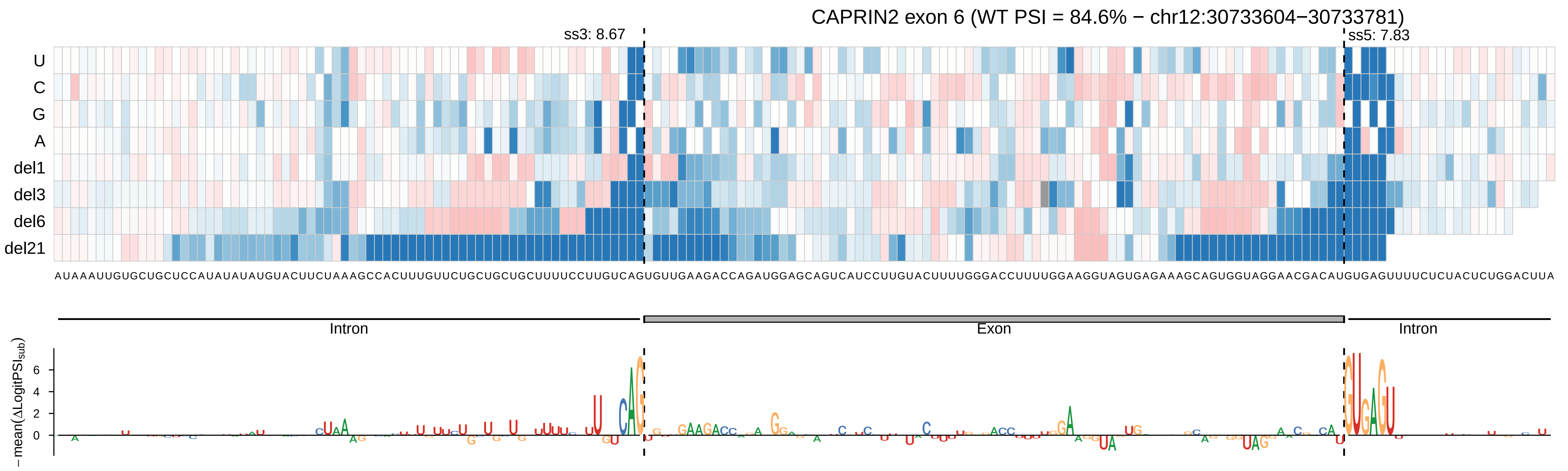

CAPRIN2 exon 6 (WT PSI = 84.6% − chr12:30733604−30733781)
ss3: 8.67
ss5: 7.83
U
C
G
A
del1
del3
del6
del21
AUAAAUUGUGCUGCUCCAUAUAUAUGUACUUCUAAAGCCACUUUGUUCUGCUGCUGCUUUUCCUUGUCAGUGUUGAAGACCAGAUGGAGCAGUCAUCCUUGUACUUUUGGGACCUUUUGGAAGGUAGUGAGAAAGCAGUGGUAGGAACGACAUGUGAGUUUUCUCUACUCUGGACUUA
Intron
Exon
Intron
− mean(ΔLogitPSI_sub)
6
4
2
0

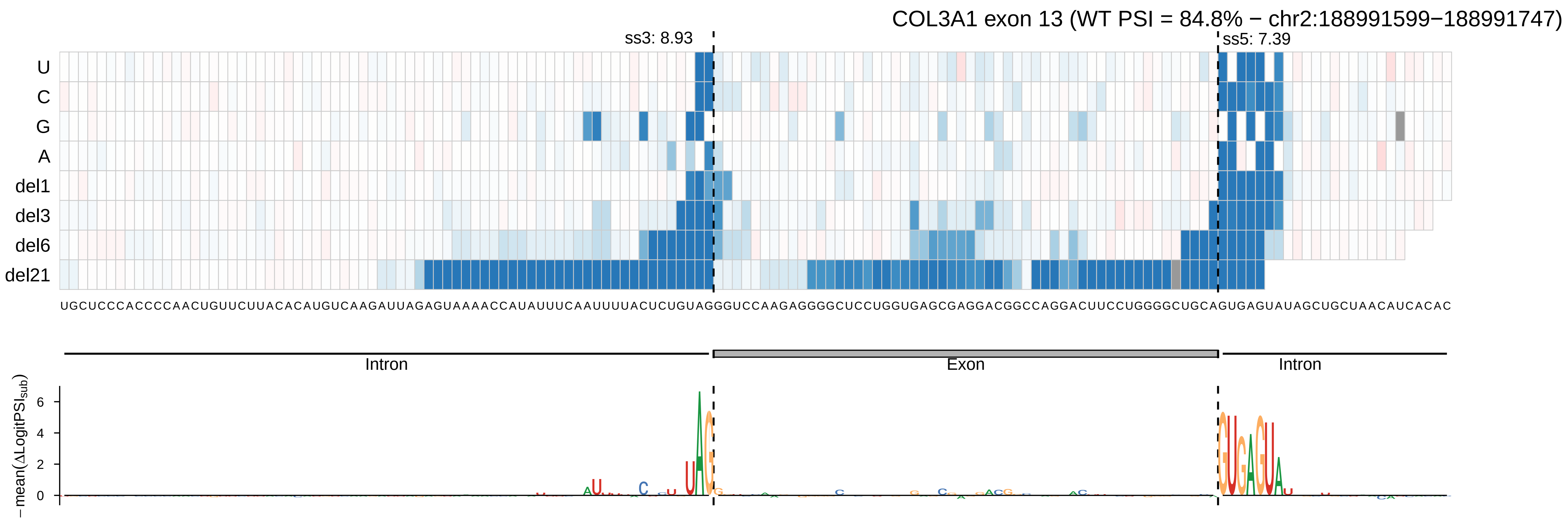

COL3A1 exon 13 (WT PSI = 84.8% − chr2:188991599−188991747)
ss3: 8.93
ss5: 7.39
U
C
G
A
del1
del3
del6
del21
UGCUCCCACCCCAACUGUUCUUACACAUGUCAAGAUUAGAGUAAAACCAUAUUUCAAUUUUACUCUGUAGGGUCCAAGAGGGGCUCCUGGUGAGCGAGGACGGCCAGGACUUCCUGGGGCUGCAGUGAGUAUAGCUGCUAACAUCACAC
Intron
Exon
Intron
−mean(ΔLogitPSI_sub)
0
2
4
6

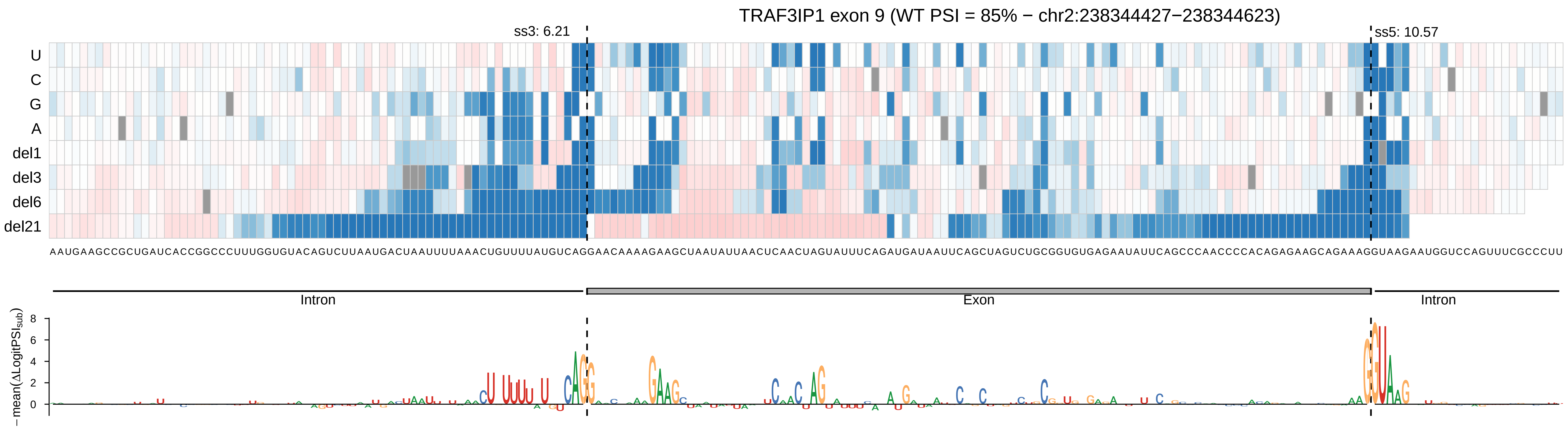

TRAF3IP1 exon 9 (WT PSI = 85% − chr2:238344427−238344623)
ss3: 6.21
ss5: 10.57
U
C
G
A
del1
del3
del6
del21
Intron
Exon
Intron
−mean(ΔLogitPSI_sub)

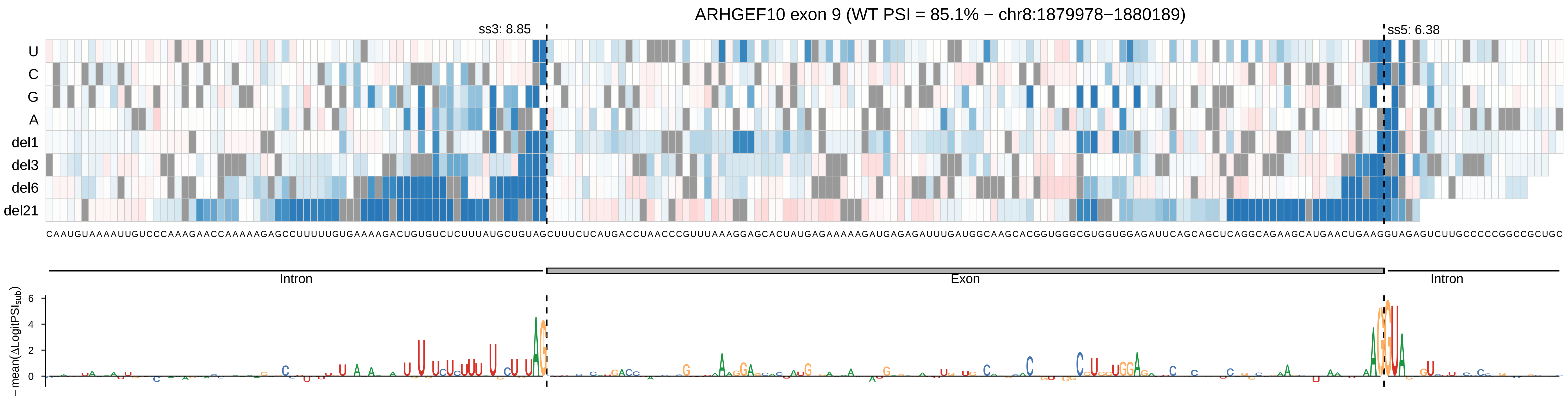

ARHGEF10 exon 9 (WT PSI = 85.1% − chr8:1879978−1880189)
ss3: 8.85
ss5: 6.38
U
C
G
A
del1
del3
del6
del21
Intron
Exon
Intron
−mean(ΔLogitPSI_sub)
6
4
2
0

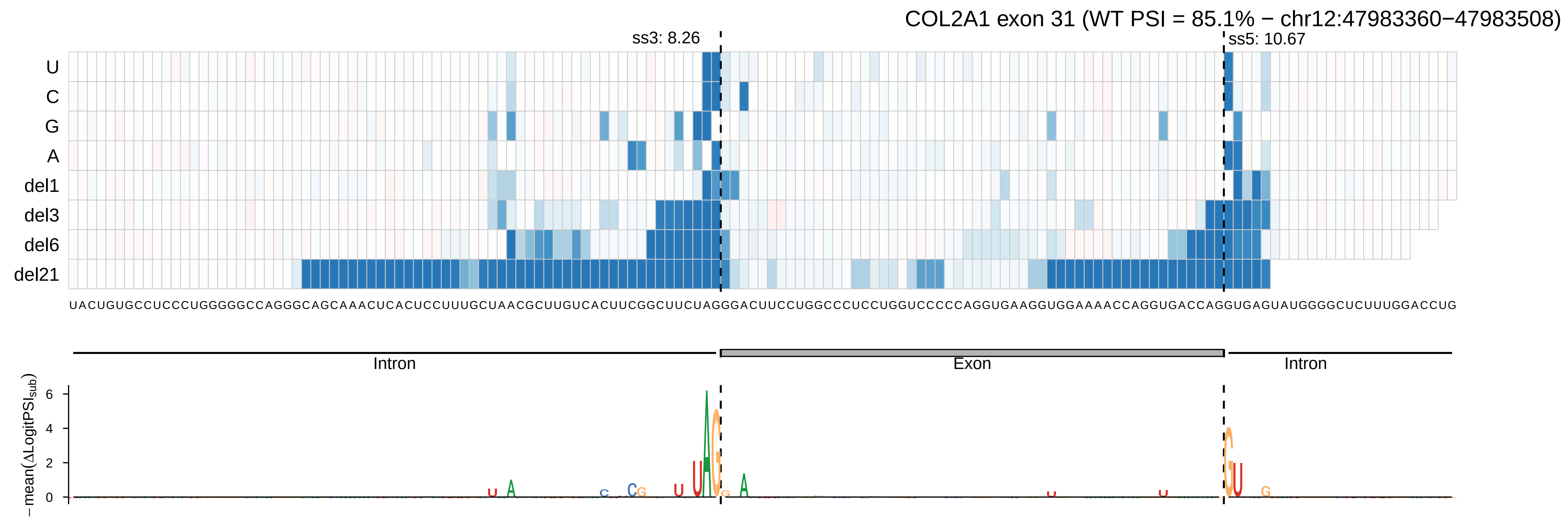

COL2A1 exon 31 (WT PSI = 85.1% − chr12:47983360−47983508)
ss3: 8.26
ss5: 10.67
U
C
G
A
del1
del3
del6
del21
UACUGUGCCUCCCUGGGGGCCAGGGCAGCAAACUCACUCCUUUGCUAACGCUUGUCACUUCGGCUUCUAGGGACUUCCUGGCCCUCCUGGUCCCCCAGGUGAAGGUGGAAAACCAGGUGACCAGGUGAGUAUGGGGCUCUUUGGACCUG
Intron
Exon
Intron
− mean(ΔLogitPSIsub)
6
4
2
0

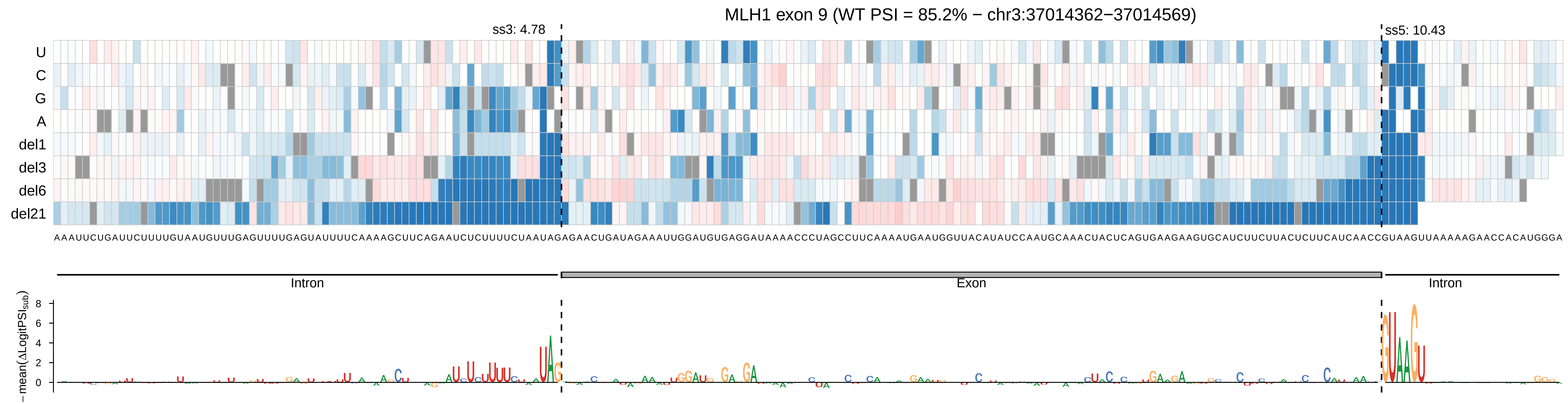

MLH1 exon 9 (WT PSI = 85.2% − chr3:37014362−37014569)
ss3: 4.78
ss5: 10.43
U
C
G
A
del1
del3
del6
del21
AAAUUCUGAUUCUUUUGUAAUGUUUGAGUUUUGAGUAUUUUCAAAAGCUUCAGAAUCUCUUUUCUAAUAGAGAACUGAUAGAAAUUGGAUGUGAGGAUAAAACCCUAGCCUUCAAAAUGAAUGGUUACAUAUCCAAUGCAAACUACUCAGUGAAGAAGUGCAUCUUCUUACUCUUCAUCAACCGUAAGUUAAAAAGAACCACAUGGGA
Intron
Exon
Intron
− mean(ΔLogitPSI_sub)
8
6
4
2
0

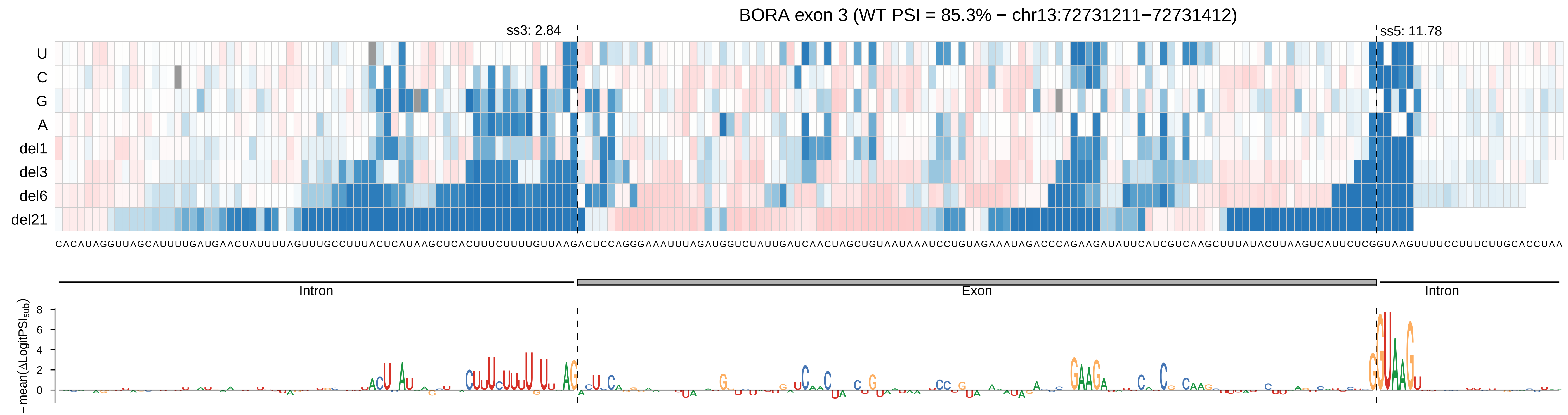

BORA exon 3 (WT PSI = 85.3% − chr13:72731211−72731412)
ss3: 2.84
ss5: 11.78
U
C
G
A
del1
del3
del6
del21
CACAUAGGUUAGCAUUUUGAUGAACUAUUUUAGUUUGCCUUUACUCAUAAGCUCACUUUCUUUUGUUAAGACUCCAGGGAAAUUUAGAUGGUCUAUUGAUCAACUAGCUGUAAUAAAUCCUGUAGAAAUAGACCCAGAAGAUAUUCAUCGUCAAGCUUUAUACUUAAGUCAUUCUCGGUAAGUUUUCCUUUCUUGCACCUAA
Intron
Exon
Intron
−mean(ΔLogitPSI_sub)
8
6
4
2
0

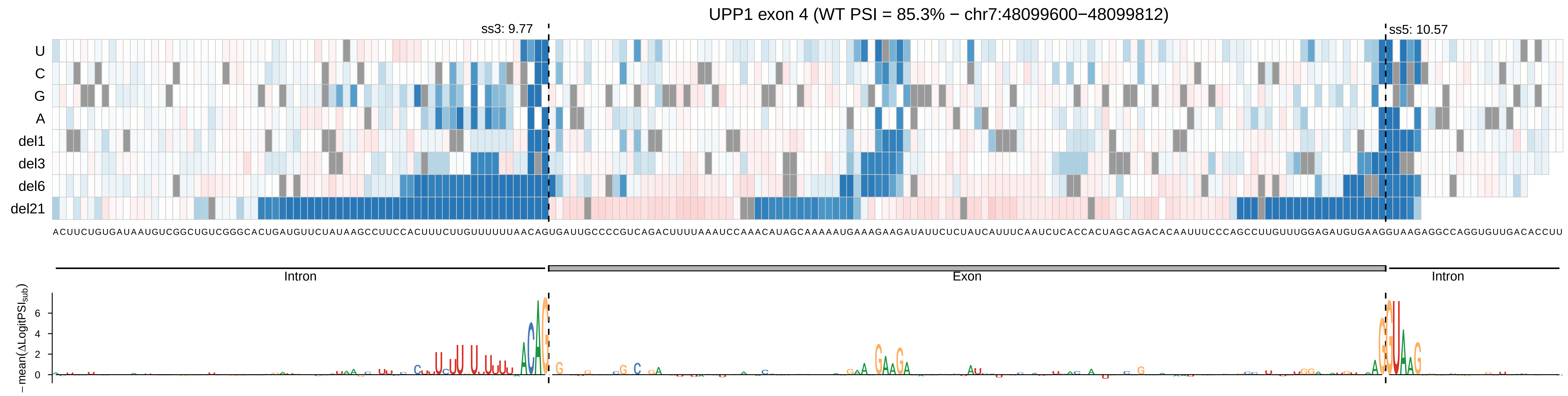

UPP1 exon 4 (WT PSI = 85.3% − chr7:48099600−48099812)
ss3: 9.77
ss5: 10.57
U
C
G
A
del1
del3
del6
del21
Intron
Exon
Intron
−mean(ΔLogitPSI_sub)

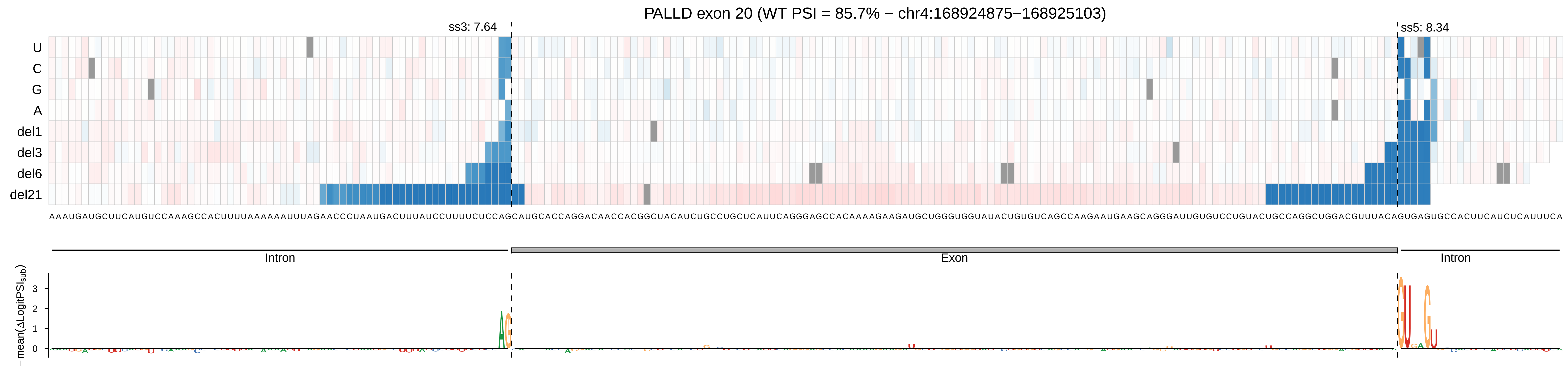
PALLD exon 20 (WT PSI = 85.7% − chr4:168924875−168925103)
ss3: 7.64
ss5: 8.34
U
C
G
A
del1
del3
del6
del21
AAAUGAUGCUUCAUGUCCAAAGCCACUUUUAAAAAAUUUAGAACCCUAAUGACUUUAUCCUUUUCUCCAGCAUGCACCAGGACAACCACGGCUACAUCUGCCUGCUCAUUCAGGGAGCCACAAAAGAAGAUGCUGGGUGGUAUACUGUGUCAGCCAAGAAUGAAGCAGGGAUUGUGUCCUGUACUGCCAGGCUGGACGUUUACAGUGAGUGCCACUUCAUCUCAUUUCA
Intron
Exon
Intron
−mean(ΔLogitPSIsub)
3
2
1
0

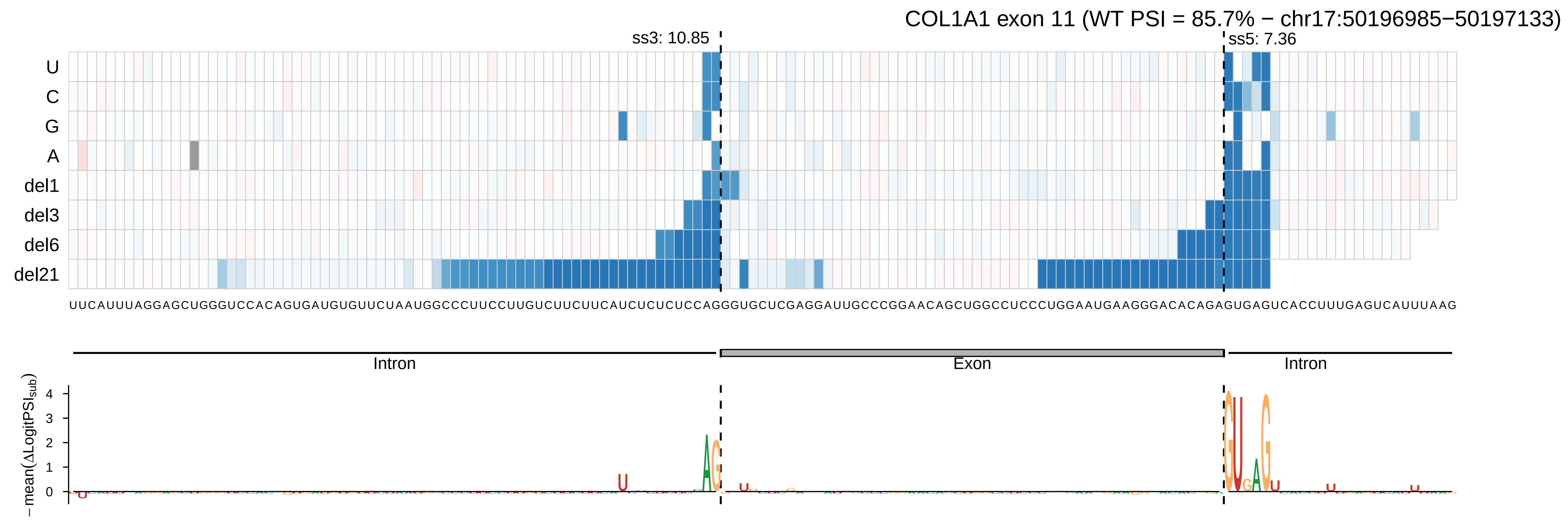
COL1A1 exon 11 (WT PSI = 85.7% − chr17:50196985−50197133)
ss3: 10.85
ss5: 7.36
U
C
G
A
del1
del3
del6
del21
Intron
Exon
Intron
−mean(ΔLogitPSI_sub)
4
3
2
1
0

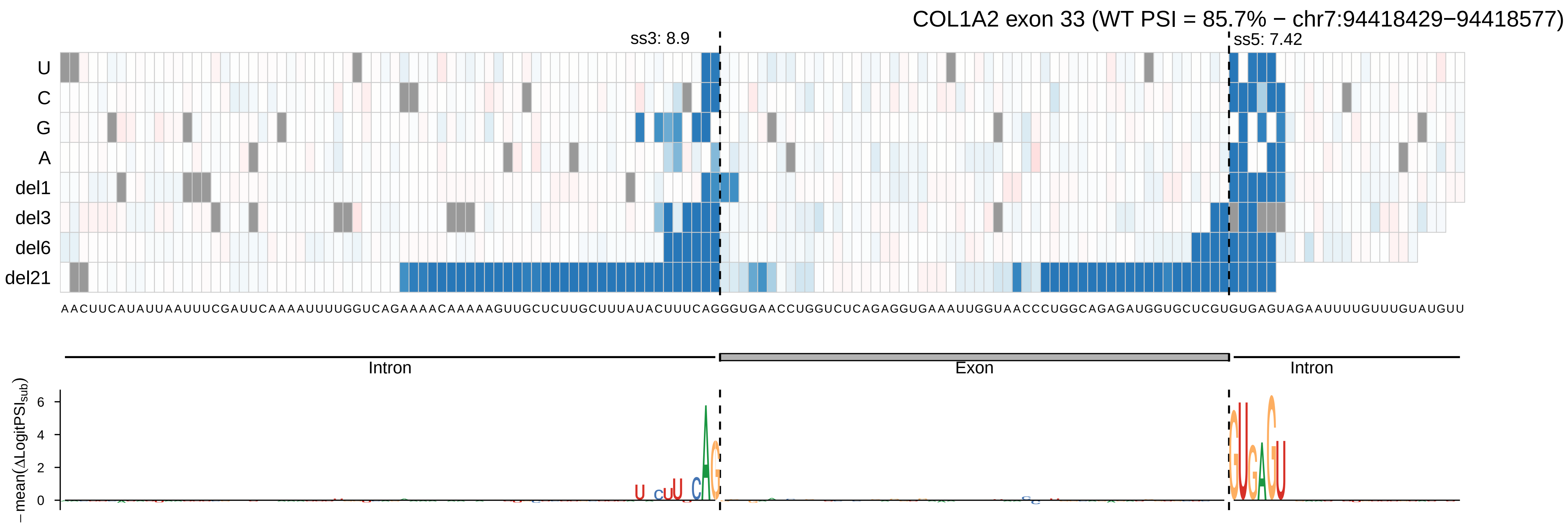

COL1A2 exon 33 (WT PSI = 85.7% − chr7:94418429−94418577)
ss3: 8.9
ss5: 7.42
U
C
G
A
del1
del3
del6
del21
AACUUCAUAUUAAUUUCGAUUCAAAAUUUUGGUCAGAAAACAAAAAGUUGCUCUUGCUUUAUACUUUCAGGGUGAACCUGGUCUCAGAGGUGAAAUUGGUAACCCUGGCAGAGAUGGUGCUCGUGUGAGUAGAAUUUUGUUUGUAUGUU
Intron
Exon
Intron
−mean(ΔLogitPSI_sub)
6
4
2
0

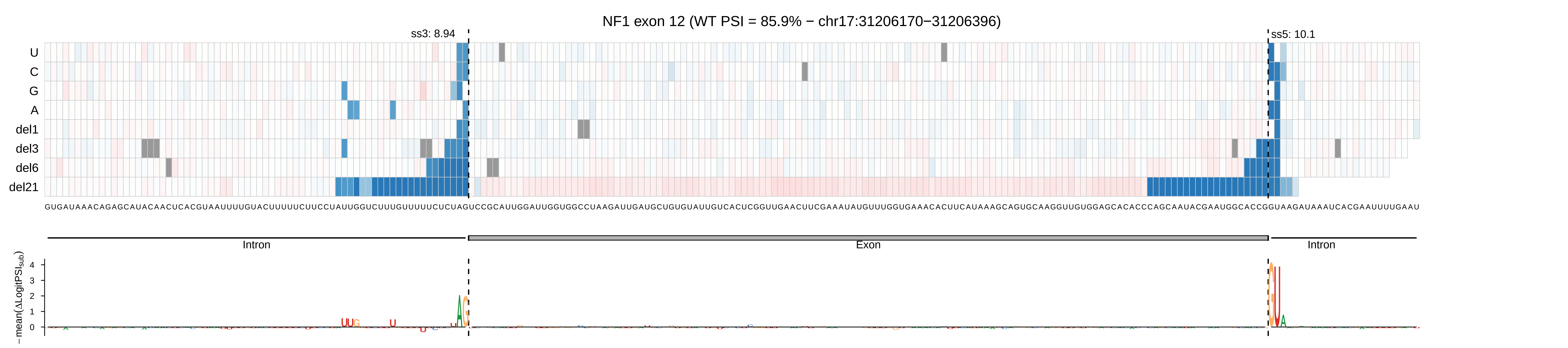

NF1 exon 12 (WT PSI = 85.9% − chr17:31206170−31206396)
ss3: 8.94
ss5: 10.1
U
C
G
A
del1
del3
del6
del21
GUGAUAAACAGAGCAUACAACUCACGUAAUUUUGUACUUUUUCUUCCUAUUGGUCUUUUGUUUUUCUCUAGUCCGCAUUGGAUUGGUGGCCUAAGAUUGAUGCUGUGUAUUGUCACUCGGUUGAACUUCGAAAUAUGUUUGGUGAAACACUUCAUAAAGCAGUGCAAGGUUGUGGAGCACACCCAGCAAUACGAAUGGCACCGGUAAGAUAAAUCACGAAUUUUGAAU
Intron
Exon
Intron
− mean(ΔLogitPSI_sub)
4
3
2
1
0

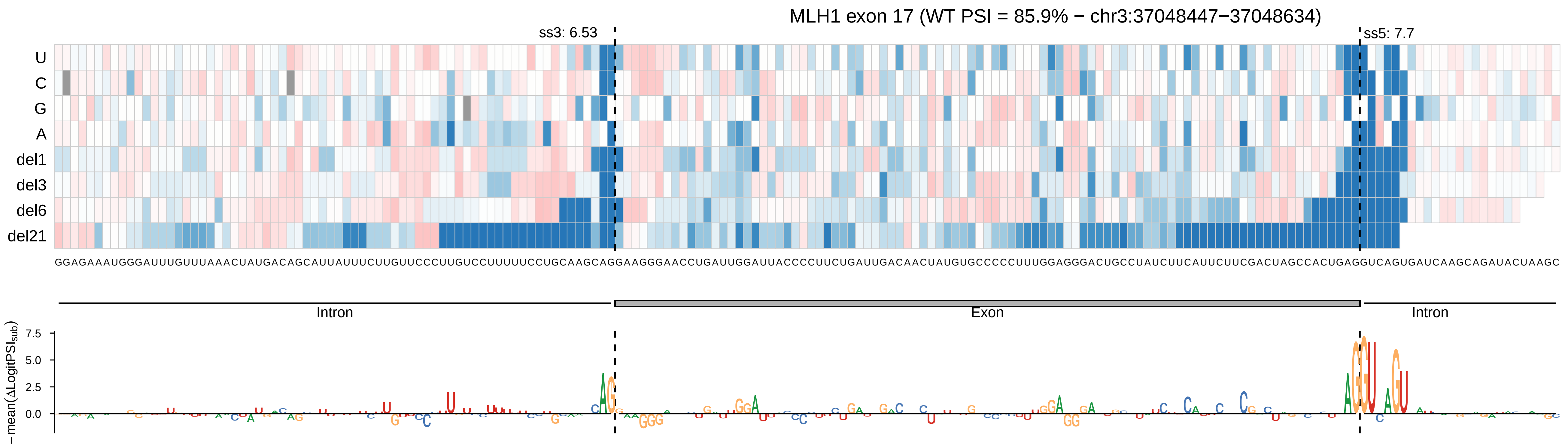

MLH1 exon 17 (WT PSI = 85.9% − chr3:37048447−37048634)
ss3: 6.53
ss5: 7.7
U
C
G
A
del1
del3
del6
del21
GGAGAAAUGGGAUUUGUUUAAACUAUGACAGCAUUAUUUCUUGUUCCCUUGUCCUUUUUCCUGCAAGCAGGAAGGGAACCUGAUUGGAUUACCCCUUCUGAUUGACAACUAUGUGCCCCCUUUGGAGGGACUGCCUAUCUUCAUUCUUCGACUAGCCACUGAGGUCAGUGAUCAAGCAGAUACUAAGC
Intron
Exon
Intron
−mean(ΔLogitPSI_sub)
7.5
5.0
2.5
0.0

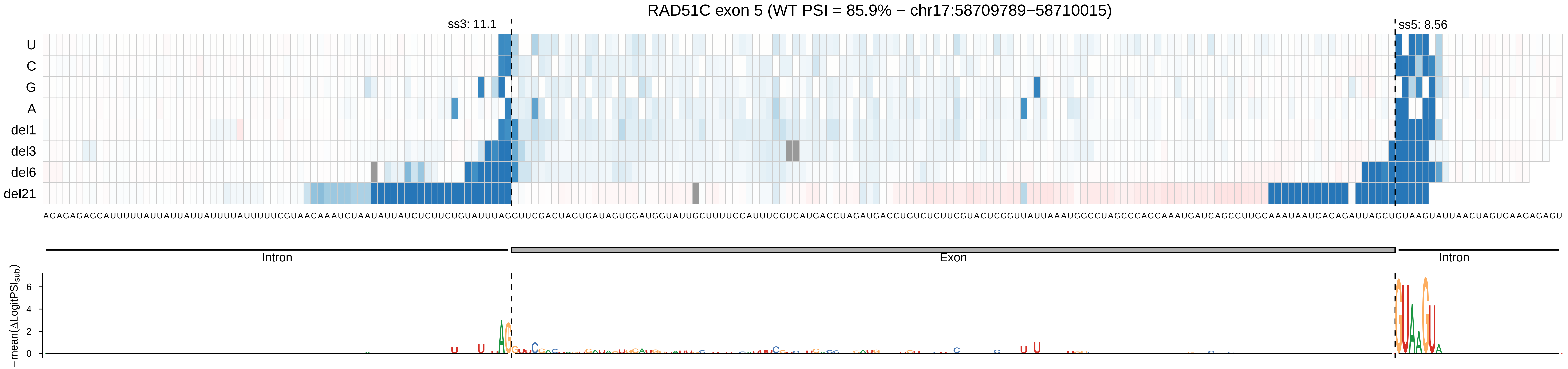

RAD51C exon 5 (WT PSI = 85.9% − chr17:58709789−58710015)
ss3: 11.1
ss5: 8.56
U
C
G
A
del1
del3
del6
del21
AGAGAGAGCAUUUUUAUUAUUAUUAUUUUAUUUUUCGUAACAAAUCUAAUAUUAUCUCUUCUGUAUUUAGGUUCGACUAGUGAUAGUGGAUGGUAUUGCUUUUCCAUUUCGUCAUGACCUAGAUGACCUGUCUCUUCGUACUCGGUUAUUAAAUGGCCUAGCCCAGCAAAUGAUCAGCCUUGCAAAUAAUCACAGAUUAGCUGUAAGUAUUAACUAGUGAAGAGAGU
Intron
Exon
Intron
− mean(ΔLogitPSI_sub)
0
2
4
6

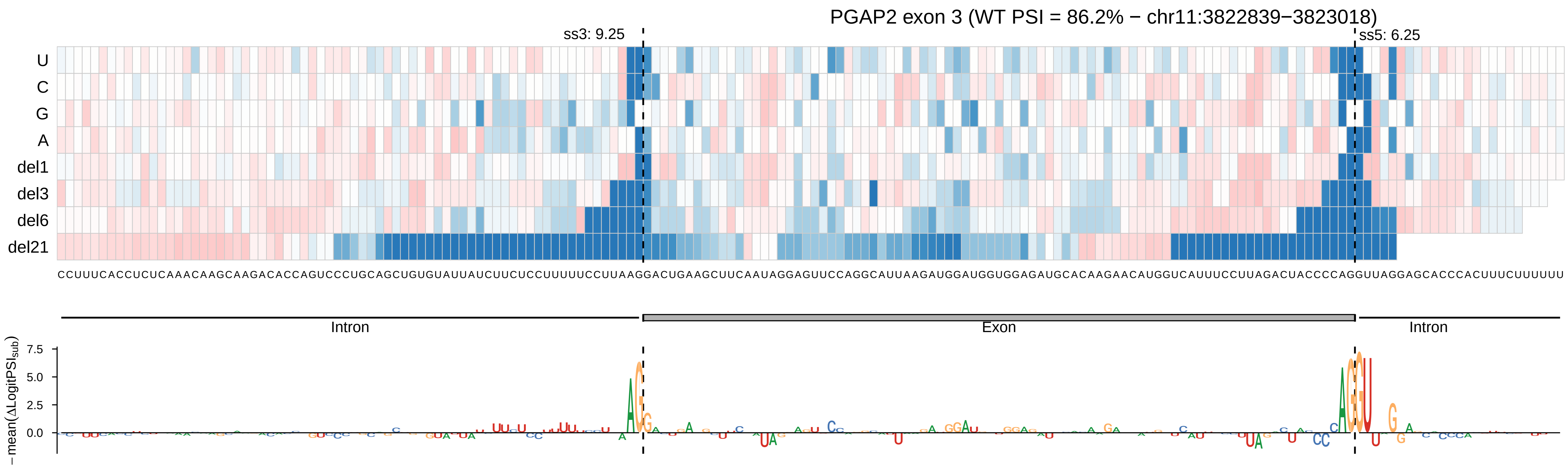

PGAP2 exon 3 (WT PSI = 86.2% − chr11:3822839−3823018)
ss3: 9.25
ss5: 6.25
U
C
G
A
del1
del3
del6
del21
CCUUUCACCUCUCAAACAAGCAAGACACCAGUCCCUGCAGCUGUGUAUUAUCUUCUCCUUUUUCCUUAAGGACUGAAGCUUCAAUAGGAGUUCCAGGCAUUAAGAUGGAUGGUGGAGAUGCACAAGAACAUGGUCAUUUCCUUAGACUACCCCAGGUUAGGAGCACCCACUUUCUUUUUU
Intron
Exon
Intron
7.5
5.0
2.5
0.0
−mean(ΔLogitPSI_sub)

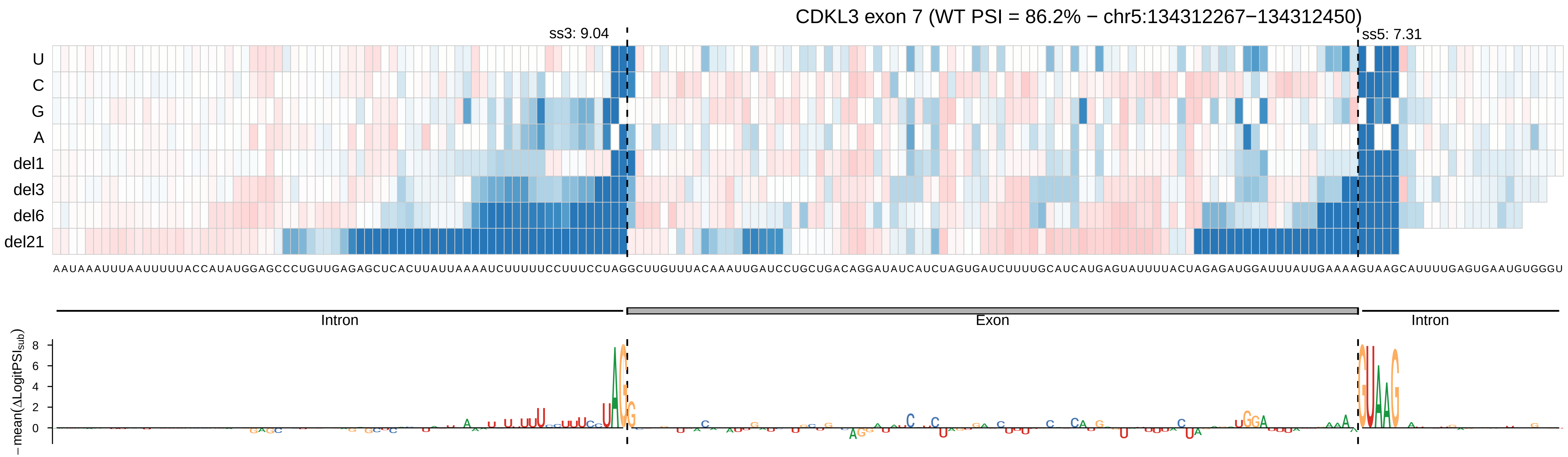

CDKL3 exon 7 (WT PSI = 86.2% − chr5:134312267−134312450)
ss3: 9.04
ss5: 7.31
U
C
G
A
del1
del3
del6
del21
AAUAAAUUUAAUUUUUACCAUAUGGAGCCCUGUUGAGAGCUCACUUAUUAAAAUCUUUUUCCUUUCCUAGGCUUGUUUACAAAUUGAUCCUGCUGACAGGAUAUCAUCUAGUGAUCUUUUGCAUCAUGAGUAUUUUACUAGAGAUGGAUUUAUUGAAAAGUAAGCAUUUUGAGUGAAUGUGGGU
Intron
Exon
Intron
− mean(ΔLogitPSI$_{sub}$)
8
6
4
2
0

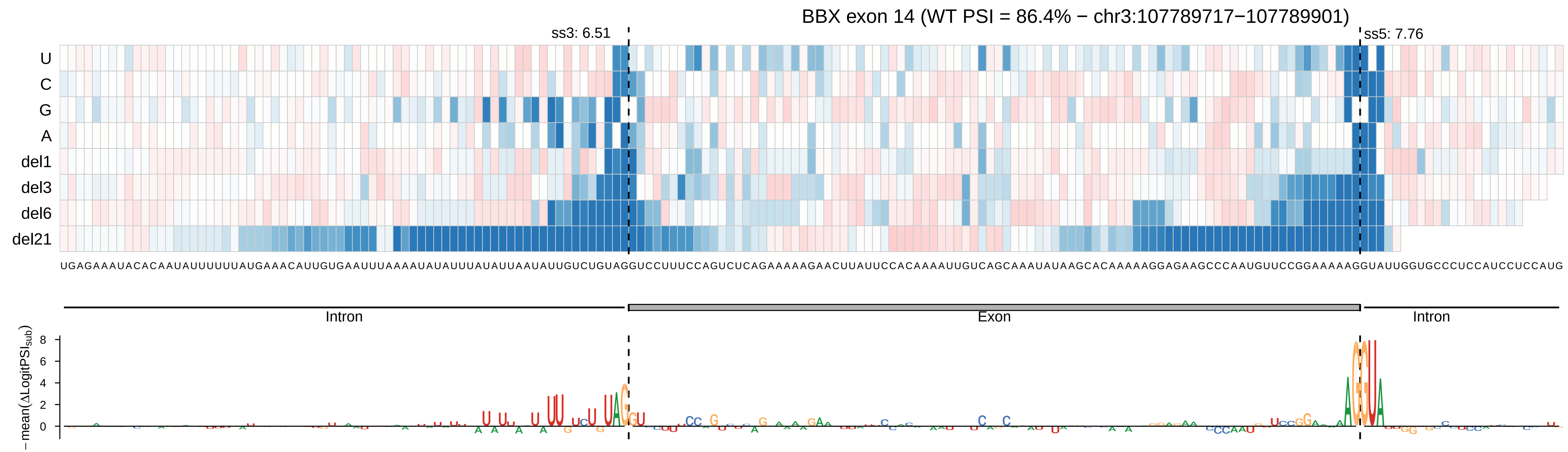
BBX exon 14 (WT PSI = 86.4% − chr3:107789717−107789901)
ss3: 6.51
ss5: 7.76
U
C
G
A
del1
del3
del6
del21
Intron
Exon
Intron
−mean(ΔLogitPSI_sub)

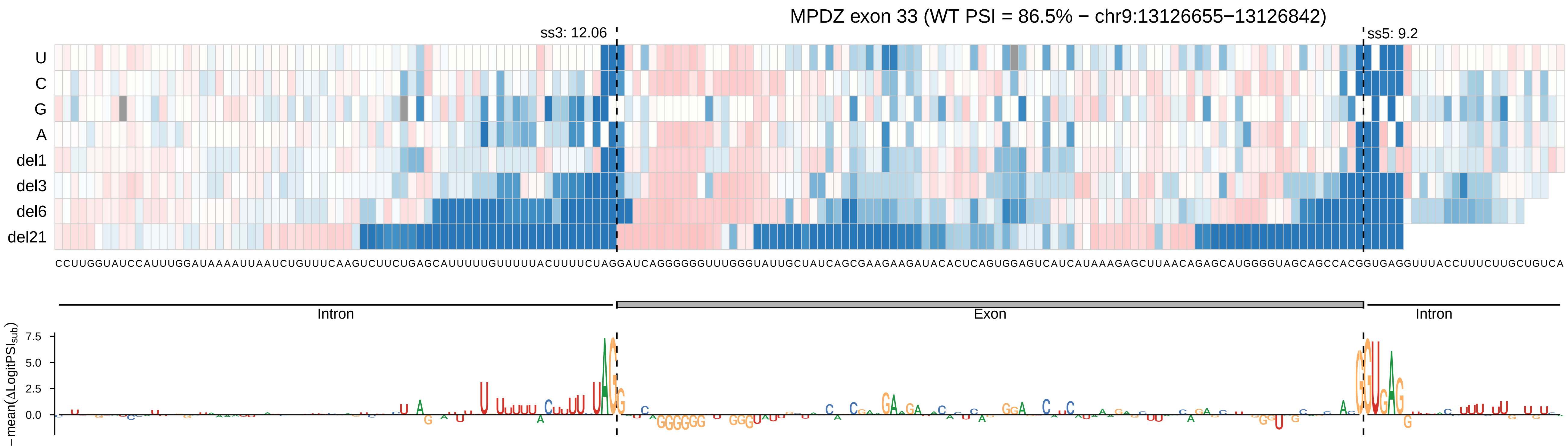

MPDZ exon 33 (WT PSI = 86.5% − chr9:13126655−13126842)
ss3: 12.06
ss5: 9.2
U
C
G
A
del1
del3
del6
del21
CCUUGGUAUCCAUUUGGAUAAAAUUAAUCUGUUUCAAGUCUUCUGAGCAUUUUUGUUUUUACUUUUCUAGGAUCAGGGGGGUUUGGGUAUUGCUAUCAGCGAAGAAGAUACACUCAGUGGAGUCAUCAUAAAGAGCUUAACAGAGCAUGGGGUAGCAGCCACGGUGAGGUUUACCUUUCUUGCUGUCA
Intron
Exon
Intron
−mean(ΔLogitPSI_sub)
7.5
5.0
2.5
0.0

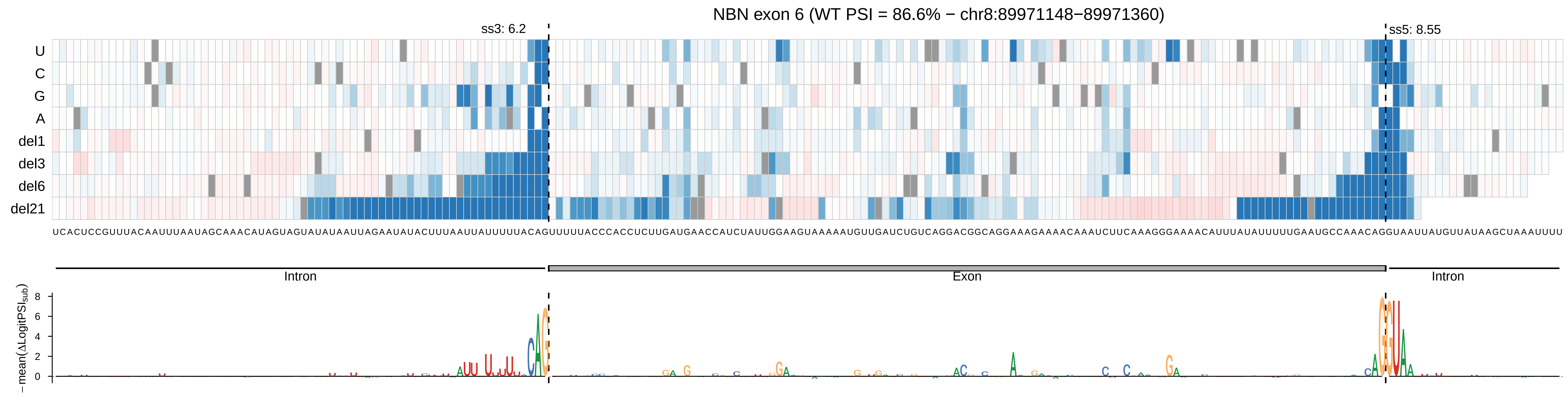
NBN exon 6 (WT PSI = 86.6% − chr8:89971148−89971360)
ss3: 6.2
ss5: 8.55
U
C
G
A
del1
del3
del6
del21
UCACUCCGUUUACAAUUUAAUAGCAAACAUAGUAGUAUAUAAUUAGAAUAUACUUUAAUUAUUUUUACAGUUUUACCCACCUCUUGAUGAACCAUCUAUUGGAAGUAAAAAUGUUGAUCUGUCAGGACGGCAGGAAAGAAAACAAAUCUUCAAAGGGAAAACAUUUAUAUUUUUGAAUGCCAAACAGGUAAUUAUGUUAUAAGCUAAAUUUU
Intron
Exon
Intron
− mean(ΔLogitPSI_sub)
8
6
4
2
0

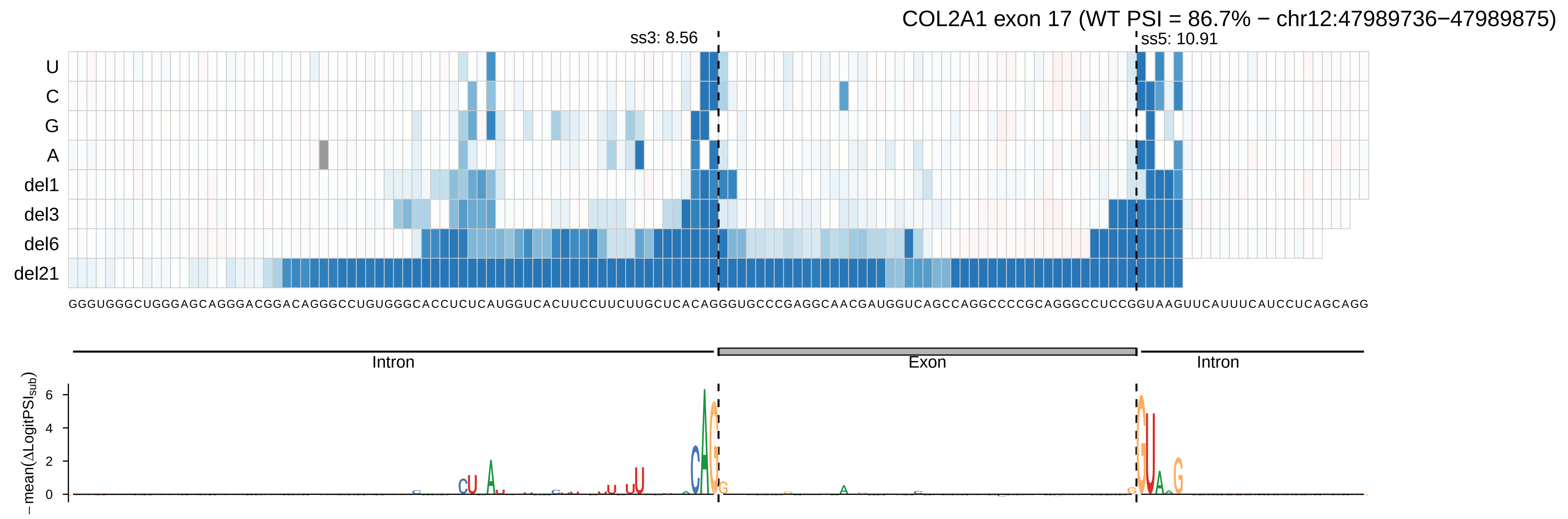
COL2A1 exon 17 (WT PSI = 86.7% − chr12:47989736−47989875)
ss3: 8.56
ss5: 10.91
U
C
G
A
del1
del3
del6
del21
GGGUGGGCUGGGAGCAGGGACGGACAGGGCCUGUGGGCACCUCUCAUGGUCACUUCCUUCUUGCUCACAGGGUGCCCGAGGCAACGAUGGUCAGCCAGGCCCCGCAGGGCCUCCGGUAAGUUCAUUUCAUCCUCAGCAGG
Intron
Exon
Intron
− mean(ΔLogitPSI_sub)
6
4
2
0

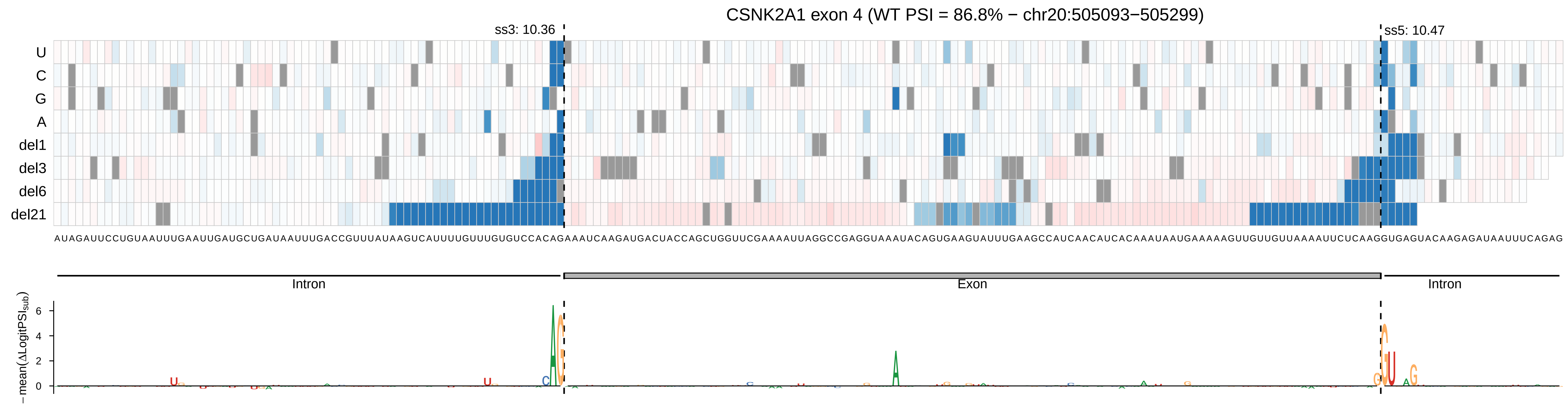

CSNK2A1 exon 4 (WT PSI = 86.8% − chr20:505093−505299)
ss3: 10.36
ss5: 10.47
U
C
G
A
del1
del3
del6
del21
AUAGAUUCCUGUAAUUUGAAUUGAUGCUGAUAAUUUGACCGUUUAUAAGUCAUUUUGUUUGUGUCCACAGAAAUCAAGAUGACUACCAGCUGGUUCGAAAAUUAGGCCGAGGUAAAAUACAGUGAAGUAUUUGAAGCCAUCAACAUCACAAAUAAUGAAAAAGUUGUUGUUAAAAUUCUCAAGGUGAGUACAAGAGAUAAUUUCAGAG
Intron
Exon
Intron
−mean(ΔLogitPSI_sub)
6
4
2
0

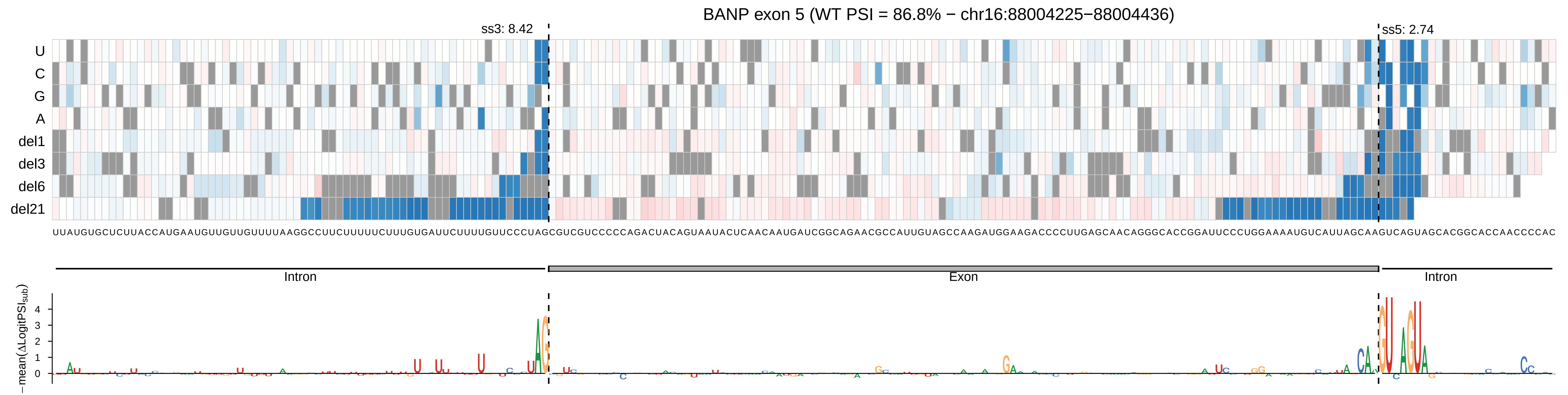

BANP exon 5 (WT PSI = 86.8% − chr16:88004225−88004436)
ss3: 8.42
ss5: 2.74
U
C
G
A
del1
del3
del6
del21
UUAUGUGCUCUUACCAUGAAUGUUGUUGUUUUAAGGCCUUCUUUUUCUUUGUGAUUCUUUUGUUCCCUAGCGUCGUCCCCCAGACUACAGUAAUACUCAACAAUGAUCGGCAGAACGCCAUUGUAGCCAAGAUGGAAGACCCCUUGAGCAACAGGGCACCGGAUUCCCUGGAAAAUGUCAUUAGCAAGUCAGUAGCACGGCACCAACCCCAC
Intron
Exon
Intron
− mean(ΔLogitPSI$_{sub}$)
0
1
2
3
4

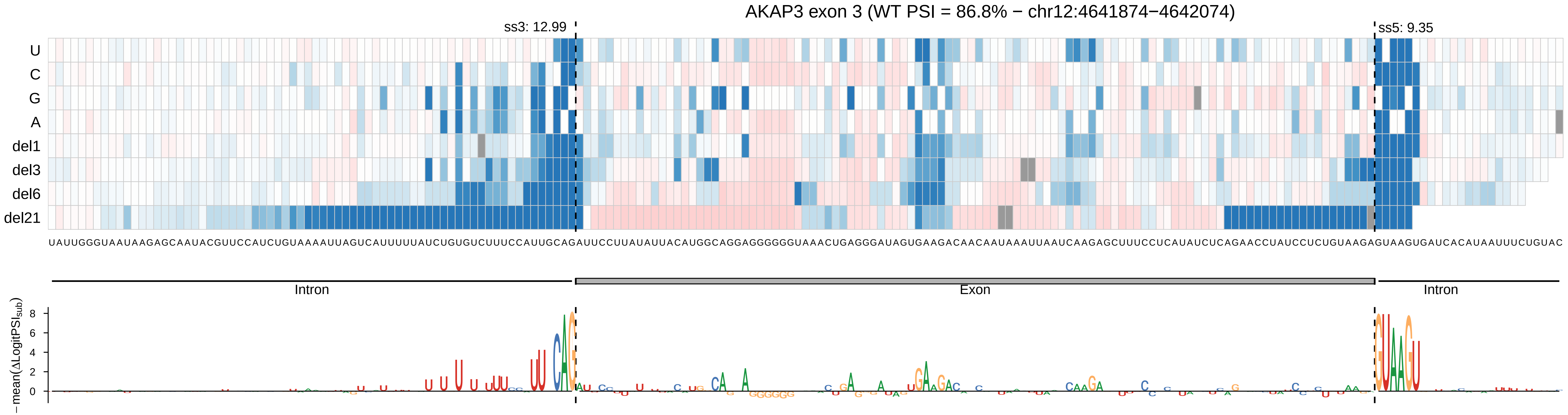

AKAP3 exon 3 (WT PSI = 86.8% − chr12:4641874−4642074)
ss3: 12.99
ss5: 9.35
U
C
G
A
del1
del3
del6
del21
UAUUGGGUAAUAAGAGCAAUACGUUCCAUCUGUAAAAUUAGUCAUUUUUAUCUGUGUCUUUCCAUUGCAGAUUCCUUAUAUUACAUGGCAGGAGGGGGGUAAACUGAGGGAUAGUGAAGACAACAAUAAAUUAAUCAAGAGCUUUCCUCAUAUCUCAGAACCUAUCCUCUGUAAGAGUAAGUGAUCACAUAAUUUCUGUAC
Intron
Exon
Intron
− mean(ΔLogitPSI_sub)
8
6
4
2
0

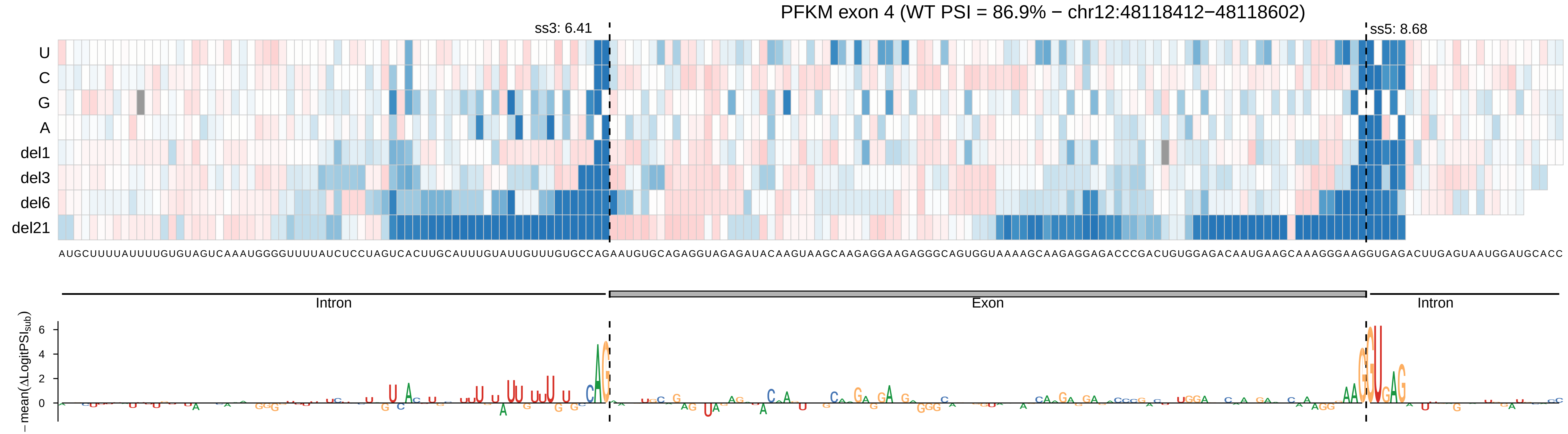

PFKM exon 4 (WT PSI = 86.9% − chr12:48118412−48118602)
ss3: 6.41
ss5: 8.68
U
C
G
A
del1
del3
del6
del21
AUGCUUUUAUUUUGUGUAGUCAAAUGGGGUUUUAUCUCCUAGUCACUUGCAUUUGUAUUGUUUGUGCCAGAAUGUGCAGAGGUAGAGAUACAAGUAAGCAAGAGGAAGAGGGCAGUGGUAAAAGCAAGAGGAGACCCGACUGUGGAGACAAUGAAGCAAAGGGAAGGUGAGACUUGAGUAAUGGAUGCACC
Intron
Exon
Intron
−mean(ΔLogitPSI_sub)
6
4
2
0

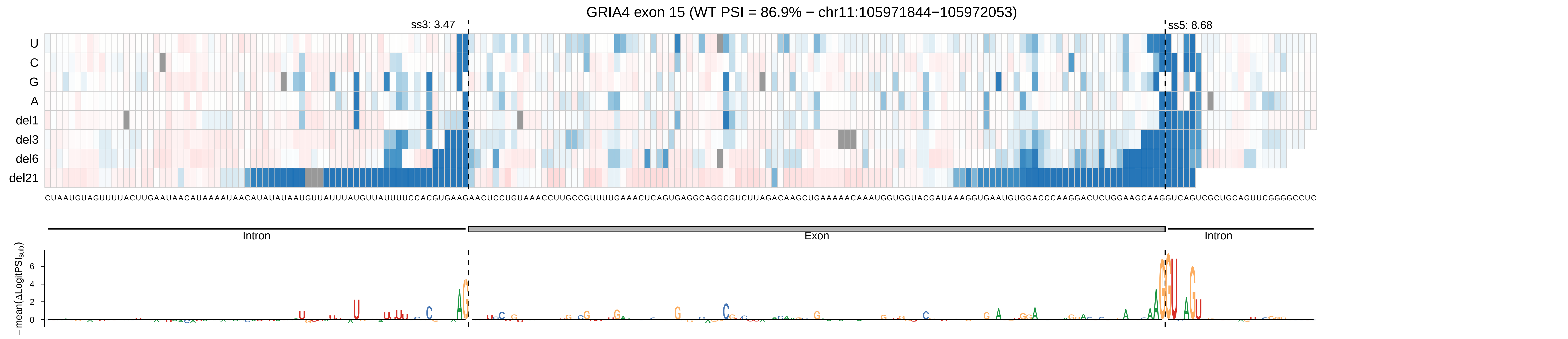

GRIA4 exon 15 (WT PSI = 86.9% − chr11:105971844−105972053)
ss3: 3.47
ss5: 8.68
U
C
G
A
del1
del3
del6
del21
CUAAUGUAGUUUUACUUGAAUAACAUAAAAUAACAUAUAUAAUGUUAUUUAUGUUAUUUUCCACGUGAAGAACUCCUGUAAACCUUGCCGUUUUGAAACUCAGUGAGGCAGGCGUCUUAGACAAGCUGAAAAACAAAUGGUGGUACGAUAAAGGUGAAUGUGGACCCAAGGACUCUGGAAGCAAGGUCAGUCGCUGCAGUUCGGGGCCUC
Intron
Exon
Intron
− mean(ΔLogitPSI_sub)
6
4
2
0

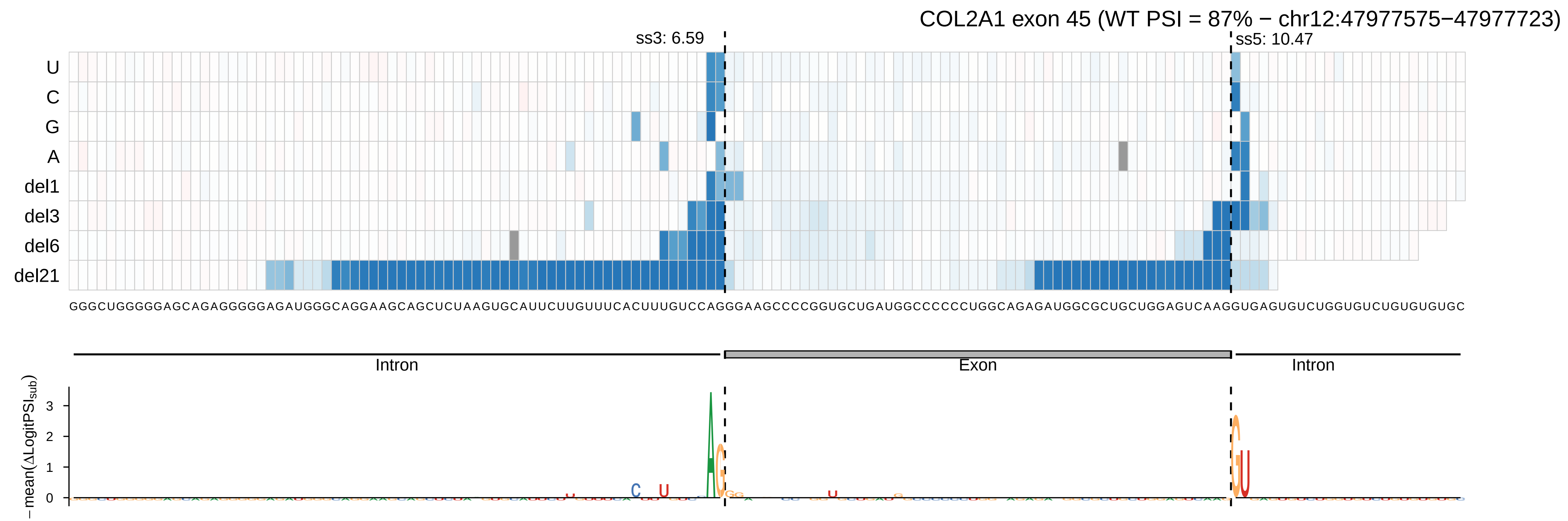
COL2A1 exon 45 (WT PSI = 87% − chr12:47977575−47977723)
ss3: 6.59
ss5: 10.47
U
C
G
A
del1
del3
del6
del21
Intron
Exon
Intron
− mean(ΔLogitPSI_sub)

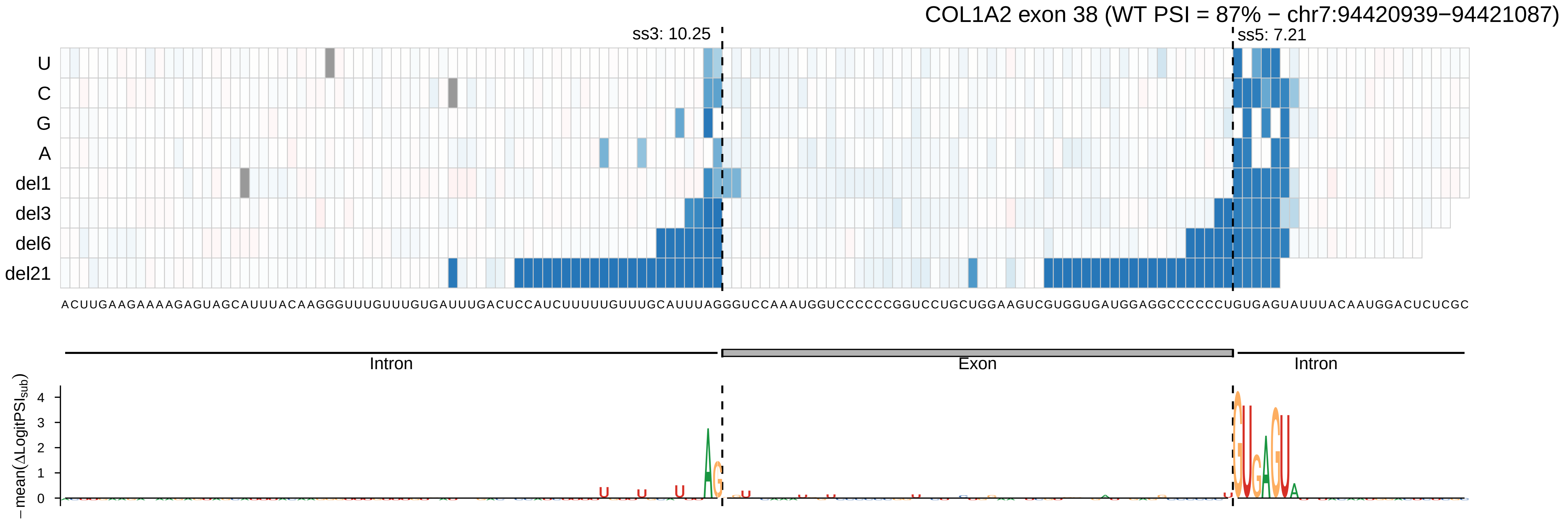
COL1A2 exon 38 (WT PSI = 87% − chr7:94420939−94421087)
ss3: 10.25
ss5: 7.21
U
C
G
A
del1
del3
del6
del21
ACUUGAAGAAAAGAGUAGCAUUUACAAGGGUUUGUUUGUGAUUUGACUCCAUCUUUUUGUUUGCAUUUAGGGUCCAAAUGGUCCCCCGGUCCUGCUGGAAGUCGUGGUGAUGGAGGCCCCCCUGUGAGUAUUUACAAUGGACUCUCGC
Intron
Exon
Intron
− mean(ΔLogitPSI_sub)
0
1
2
3
4

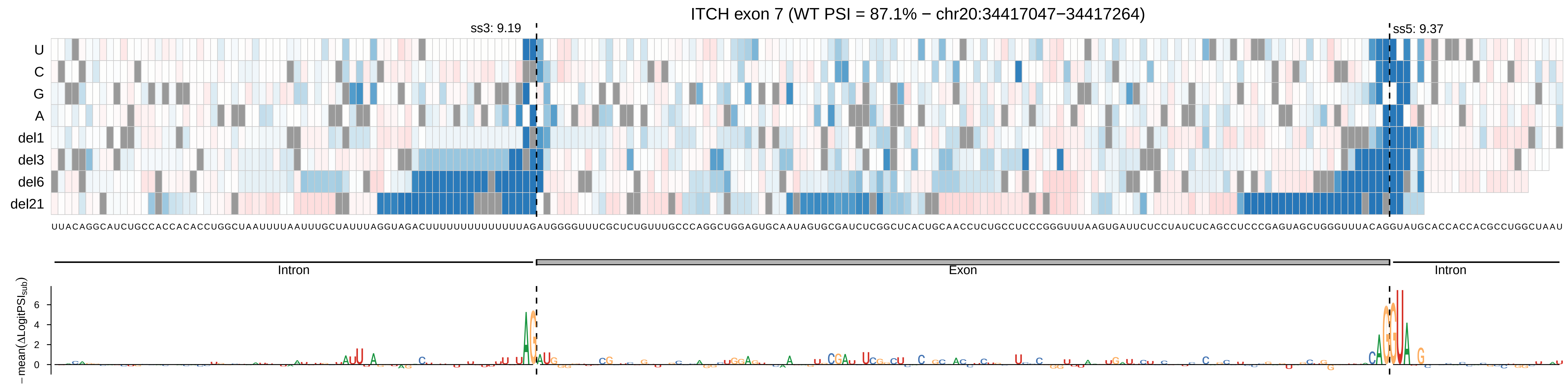

ITCH exon 7 (WT PSI = 87.1% − chr20:34417047−34417264)
ss3: 9.19
ss5: 9.37
U
C
G
A
del1
del3
del6
del21
Intron
Exon
Intron
−mean(ΔLogitPSI_sub)

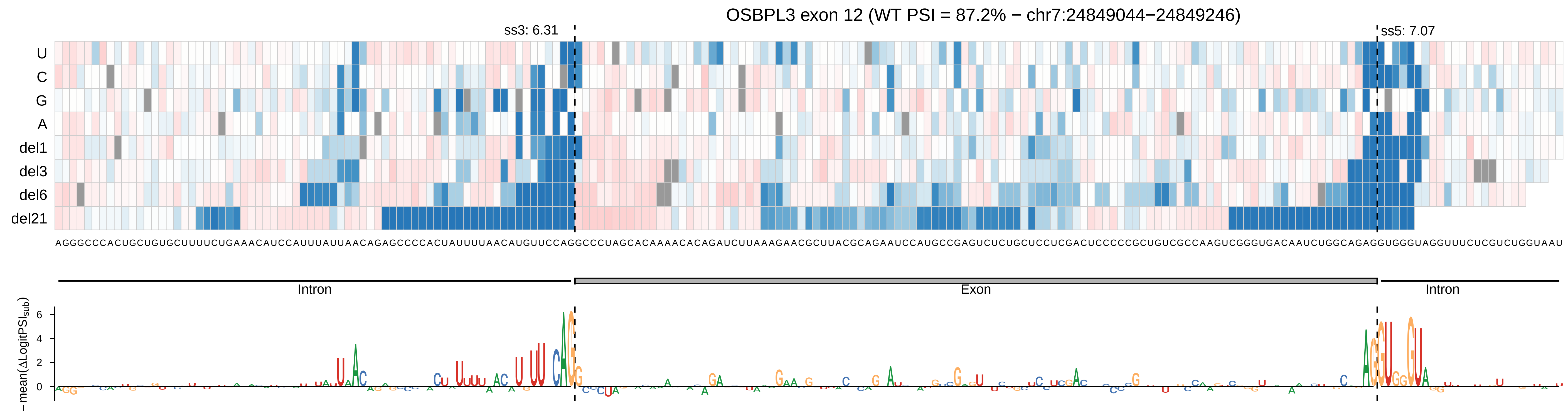
OSBPL3 exon 12 (WT PSI = 87.2% − chr7:24849044−24849246)
ss3: 6.31
ss5: 7.07
U
C
G
A
del1
del3
del6
del21
AGGGCCCACUGCUGUGCUUUUCUGAAACAUCCAUUUAUUAACAGAGCCCCACUAUUUUAACAUGUUCCAGGCCCUAGCACAAAACACAGAUCUUAAAGAACGCUUACGCAGAAUCCAUGCCGAGUCUCUGCUCCUCGACUCCCCCGCUGUCGCCAAGUCGGGUGACAAUCUGGCAGAGGUGGGUAGGUUUCUCGUCUGGUAAU
Intron
Exon
Intron
−mean(ΔLogitPSI_sub)
6
4
2
0

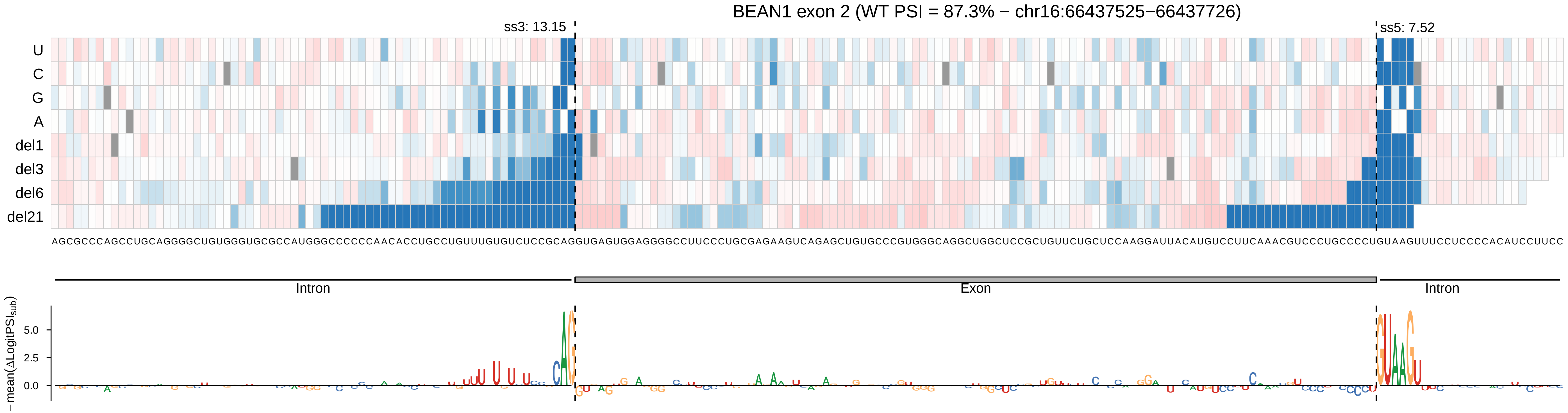

BEAN1 exon 2 (WT PSI = 87.3% − chr16:66437525−66437726)
ss3: 13.15
ss5: 7.52
U
C
G
A
del1
del3
del6
del21
AGCGCCCAGCCUGCAGGGGCUGUGGGUGCGCCAUGGGCCCCCCAACACCUGCCUGUUUGUGUCUCCGCAGGUGAGUGGAGGGGCCUUCCCUGCGAGAAGUCAGAGCUGUGCCCGUGGGCAGGCUGGCUCCGCUGUUCUGCUCCAAGGAUUACAUGUCCUUCAAACGUCCCUGCCCCUGUAAGUUUCCUCCCCACAUCCUUCC
Intron
Exon
Intron
−mean(ΔLogitPSI_sub)
5.0
2.5
0.0

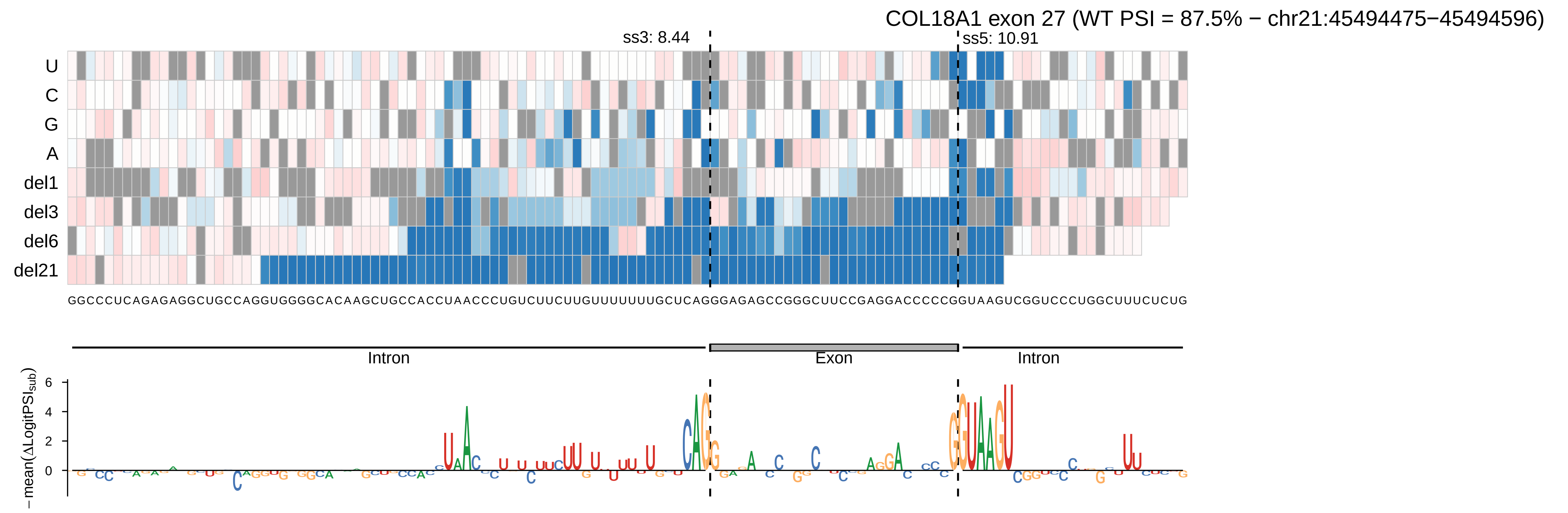
COL18A1 exon 27 (WT PSI = 87.5% − chr21:45494475−45494596)
ss3: 8.44
ss5: 10.91
U
C
G
A
del1
del3
del6
del21
GGCCCUCAGAGAGGCUGCCAGGUGGGGCACAAGCUGCCACCUAACCCUGUCUUCUUGUUUUUUUGCUCAGGGAGAGCCGGGCUUCCGAGGACCCCCGGUAAGUCGGUCCCUGGCUUUCUCUG
Intron
Exon
Intron
−mean(ΔLogitPSIsub)
6
4
2
0

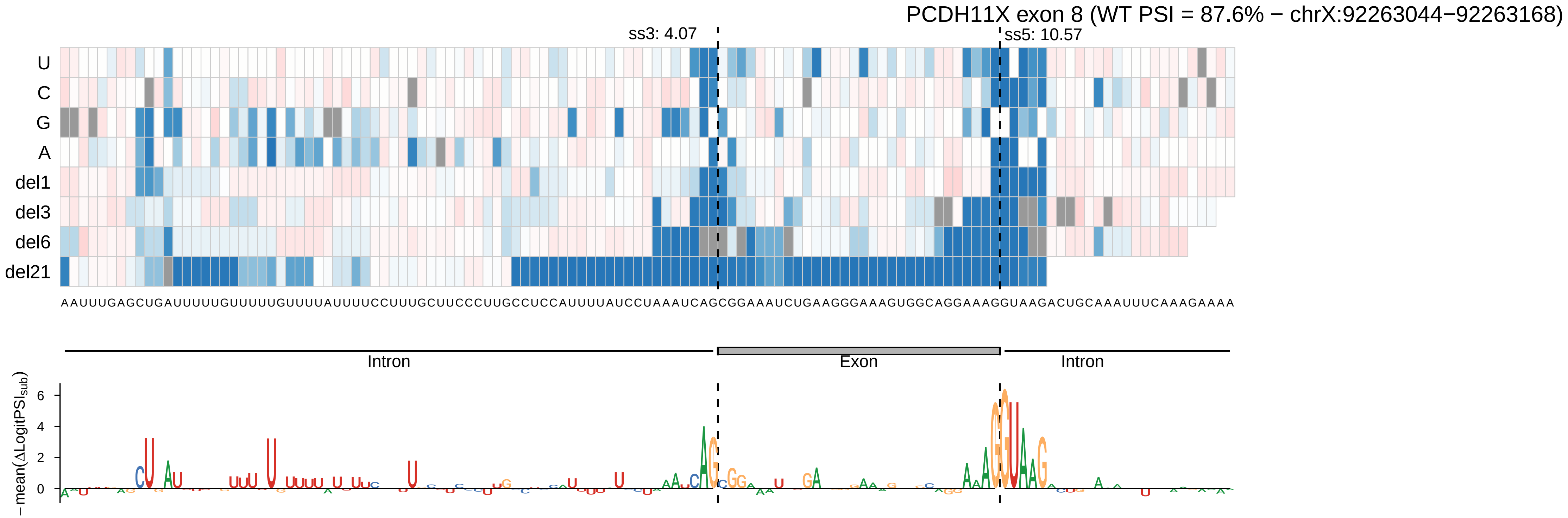
PCDH11X exon 8 (WT PSI = 87.6% − chrX:92263044−92263168)
ss3: 4.07
ss5: 10.57
U
C
G
A
del1
del3
del6
del21
AAUUUGAGCUGAUUUUUGUUUUUGUUUUAUUUUCCUUUGCUUCCCUUGCCUCCAUUUUAUCCUAAAUCAGCGGAAAUCUGAAGGGAAAGUGGCAGGAAAGGUAAGACUGCAAAUUUCAAAGAAAA
Intron
Exon
Intron
−mean(ΔLogitPSI$_{sub}$)
6
4
2
0

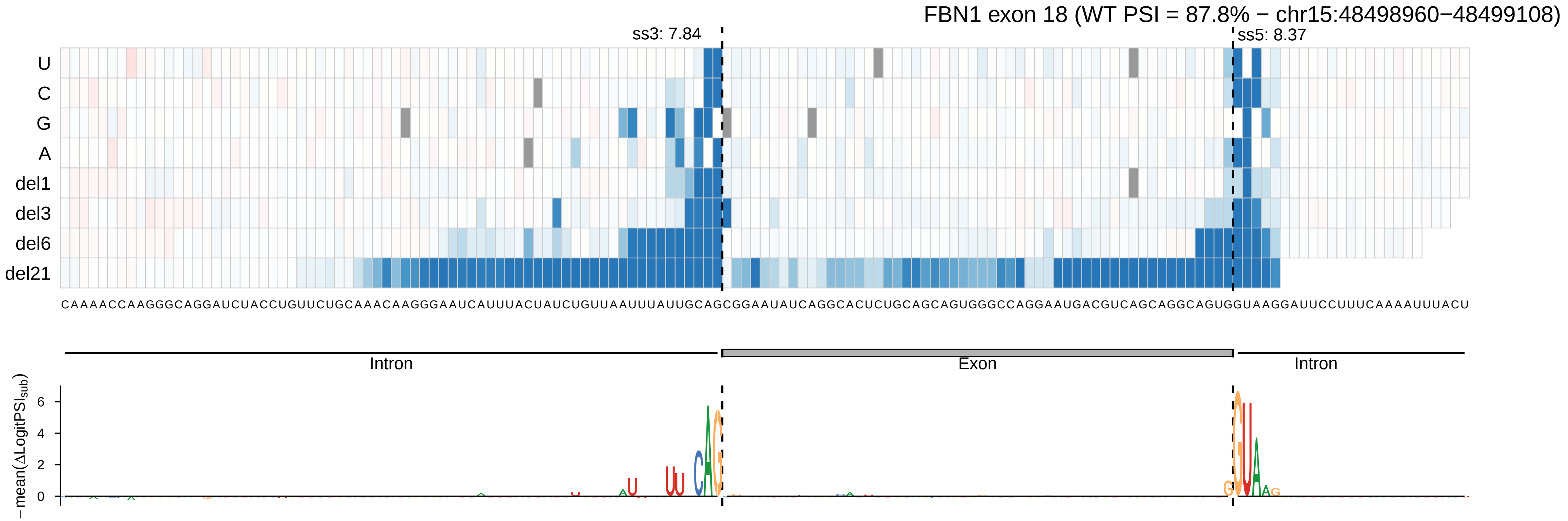
FBN1 exon 18 (WT PSI = 87.8% − chr15:48498960−48499108)
ss3: 7.84
ss5: 8.37
U
C
G
A
del1
del3
del6
del21
CAAAACCAAGGGCAGGAUCUACCUGUUCUGCAAACAAGGGAAUCAUUUACUAUCUGUUAAUUUAUUGCAGCGGAAUAUCAGGCACUCUGCAGCAGUGGGCCAGGAAUGACGUCAGCAGGCAGUGGUAAGGAUUCCUUUCAAAAUUUACU
Intron
Exon
Intron
− mean(ΔLogitPSI$_{sub}$)
6
4
2
0

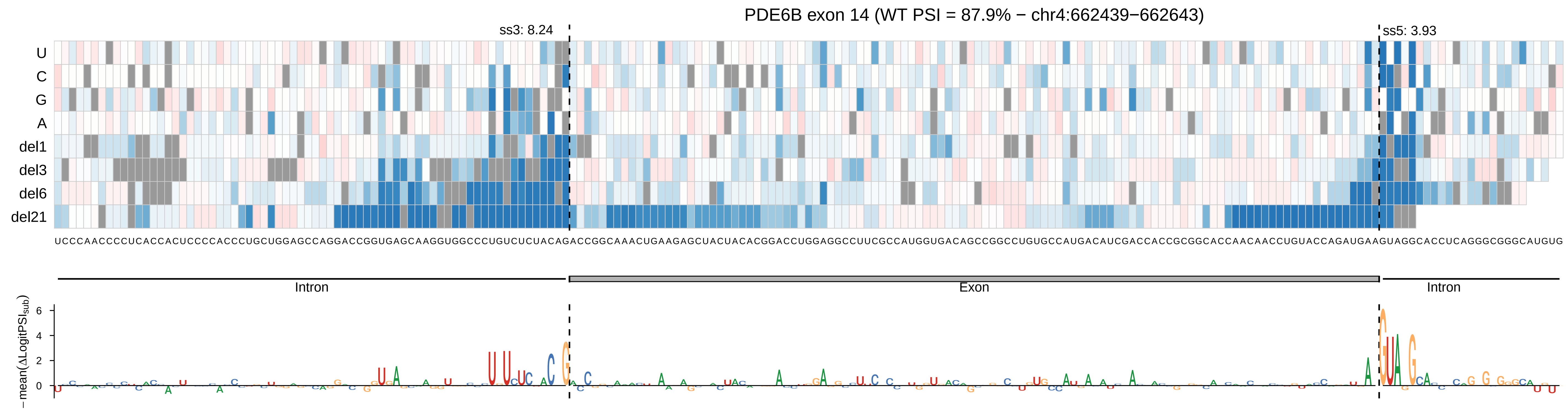
PDE6B exon 14 (WT PSI = 87.9% − chr4:662439−662643)
ss3: 8.24
ss5: 3.93
U
C
G
A
del1
del3
del6
del21
UCCCAACCCCUCACCACUCCCCACCCUGCUGGAGCCAGGACCGGUGAGCAAGGUGGCCCUGUCUCUACAGACCGGCAAACUGAAGAGCUACUACACGGACCUGGAGGCCUUCGCCAUGGUGACAGCCGGCCUGUGCCAUGACAUCGACCACCGCGGCACCAACAACCUGUACCAGAUGAAGUAGGCACCUCAGGGCGGGCAUGUG
Intron
Exon
Intron
−mean(ΔLogitPSI_sub)
6
4
2
0

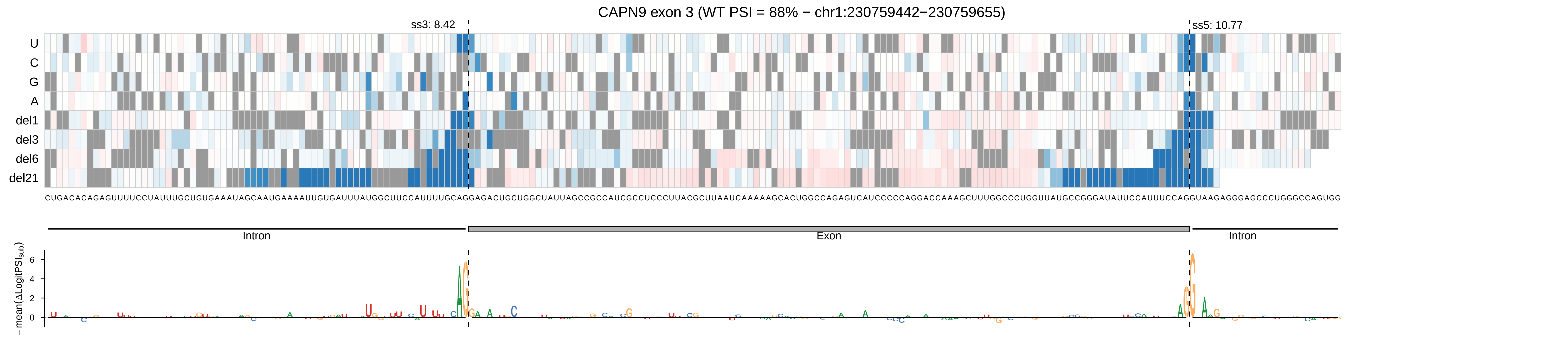

CAPN9 exon 3 (WT PSI = 88% − chr1:230759442−230759655)
ss3: 8.42
ss5: 10.77
U
C
G
A
del1
del3
del6
del21
CUGACACAGAGUUUUCCUAUUUGCUGUGAAAUAGCAAUGAAAAUUGUGAUUUAUGGCUUCCAUUUUGCAGGAGACUGCUGGCUAUUAGCCGCCAUCGCCUCCCUUACGCUUAAUCAAAAAGCACUGGCCAGAGUCAUCCCCCAGGACCAAAGCUUUGGCCCUGGUUAUGCCGGGAUAUUCCAUUUCCAGGUAAGAGGGAGCCCUGGGCCAGUGG
Intron
Exon
Intron
−mean(ΔLogitPSI_sub)
6
4
2
0

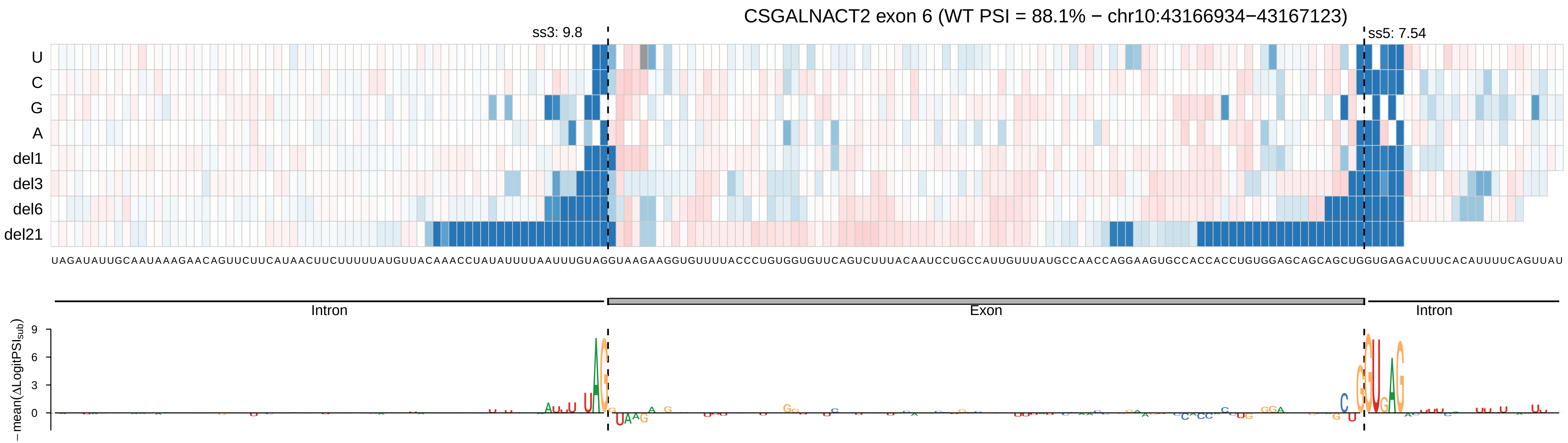

CSGALNACT2 exon 6 (WT PSI = 88.1% − chr10:43166934−43167123)
ss3: 9.8
ss5: 7.54
U
C
G
A
del1
del3
del6
del21
UAGAUAUUGCAAUAAAGAACAGUUCUUCAUAACUUCUUUUUAUGUUACAAACCUAUAUUUUAAUUUGUAGGUAAGAAGGUGUUUUACCCUGUGGGUGUUCAGUCUUUACAAUCCUGCCAUUGUUUAUGCCAACCAGGAAGUGCCACCACCUGUGGAGCAGCAGCUGGUGAGACUUUCACAUUUUCAGUUAU
Intron
Exon
Intron
−mean(ΔLogitPSI_sub)
9
6
3
0

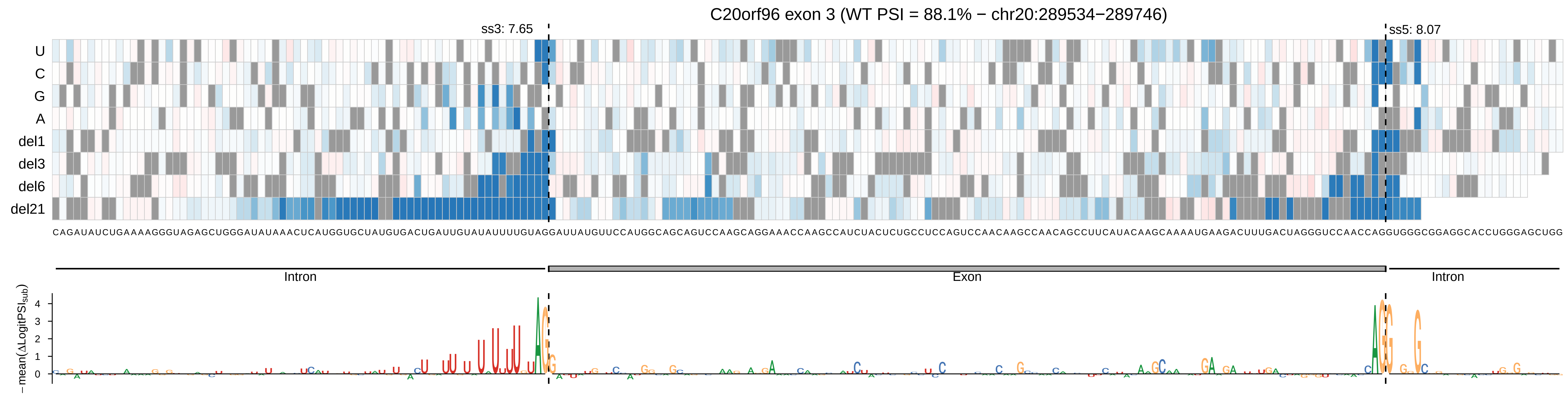
C20orf96 exon 3 (WT PSI = 88.1% − chr20:289534−289746)
ss3: 7.65
ss5: 8.07
U
C
G
A
del1
del3
del6
del21
CAGAUAUCUGAAAAGGGUAGAGCUGGGAUAUAAACUCAUGGUGCUAUGUGACUGAUUGUAUAUUUUGUAGGAUUAUGUUCCAUGGCAGCAGUCCAAGCAGGAAACCAAGCCAUCUACUCUGCCUCCAGUCCAACAAGCCAACAGCCUUCAUACAAGCAAAAUGAAGACUUUGACUAGGGUCCAACCAGGUGGGCGGAGGCACCUGGGAGCUGG
Intron
Exon
Intron
− mean(ΔLogitPSI_sub)
4
3
2
1
0

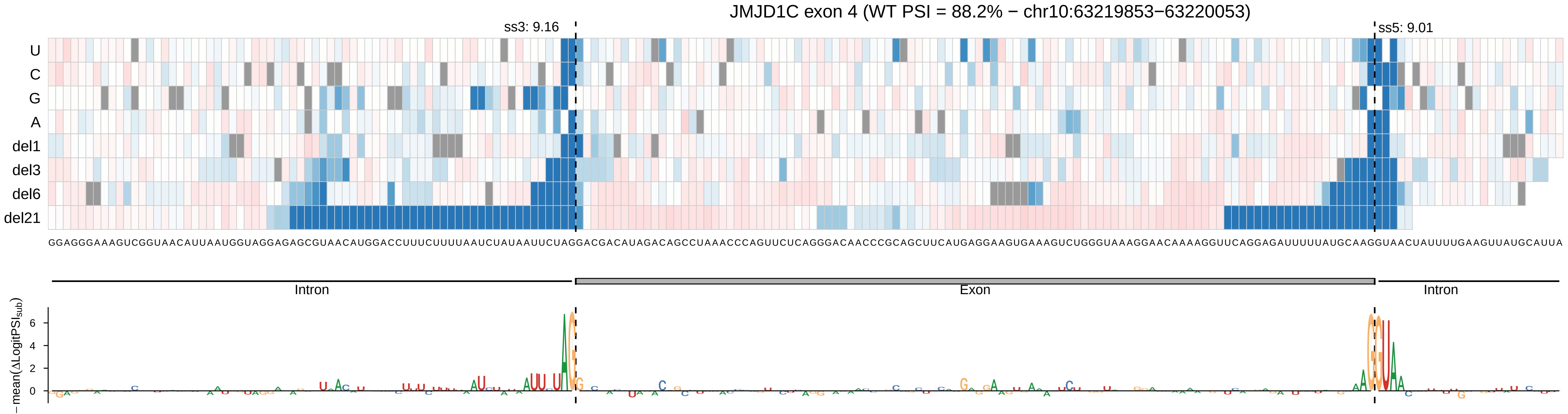
JMJD1C exon 4 (WT PSI = 88.2% − chr10:63219853−63220053)
ss3: 9.16
ss5: 9.01
U
C
G
A
del1
del3
del6
del21
Intron
Exon
Intron
−mean(ΔLogitPSIsub)
6
4
2
0

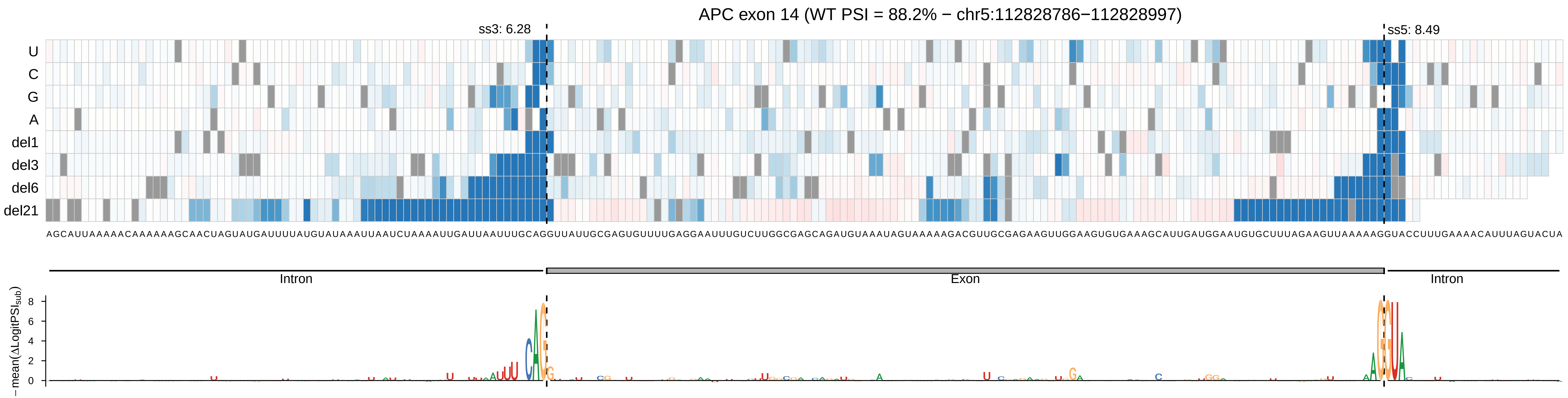
APC exon 14 (WT PSI = 88.2% − chr5:112828786−112828997)
ss3: 6.28
ss5: 8.49
U
C
G
A
del1
del3
del6
del21
Intron
Exon
Intron
− mean(ΔLogitPSI_sub)
8
6
4
2
0

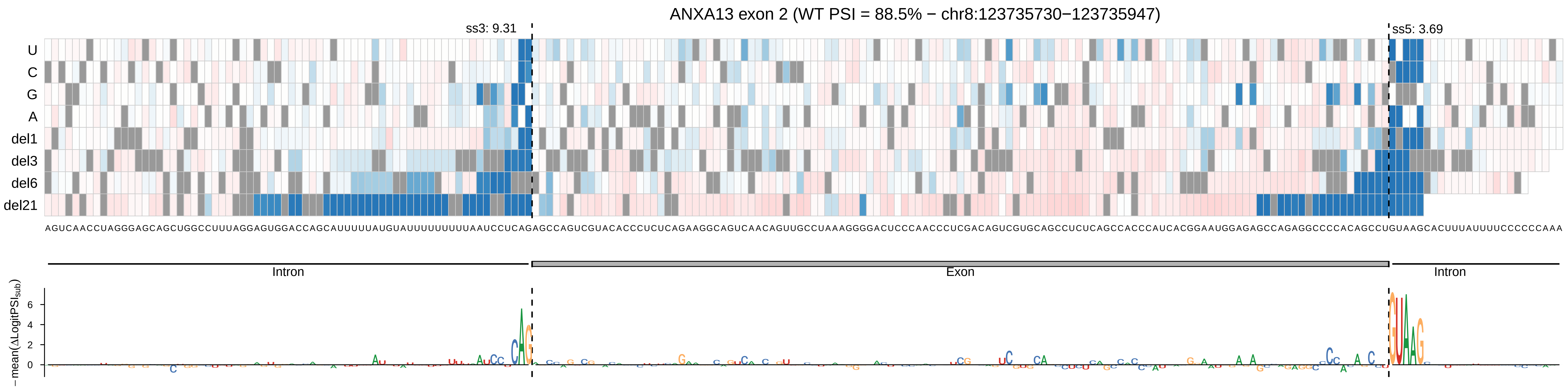
ANXA13 exon 2 (WT PSI = 88.5% − chr8:123735730−123735947)
ss3: 9.31
ss5: 3.69
U
C
G
A
del1
del3
del6
del21
Intron
Exon
Intron
−mean(ΔLogitPSI_sub)
0
2
4
6

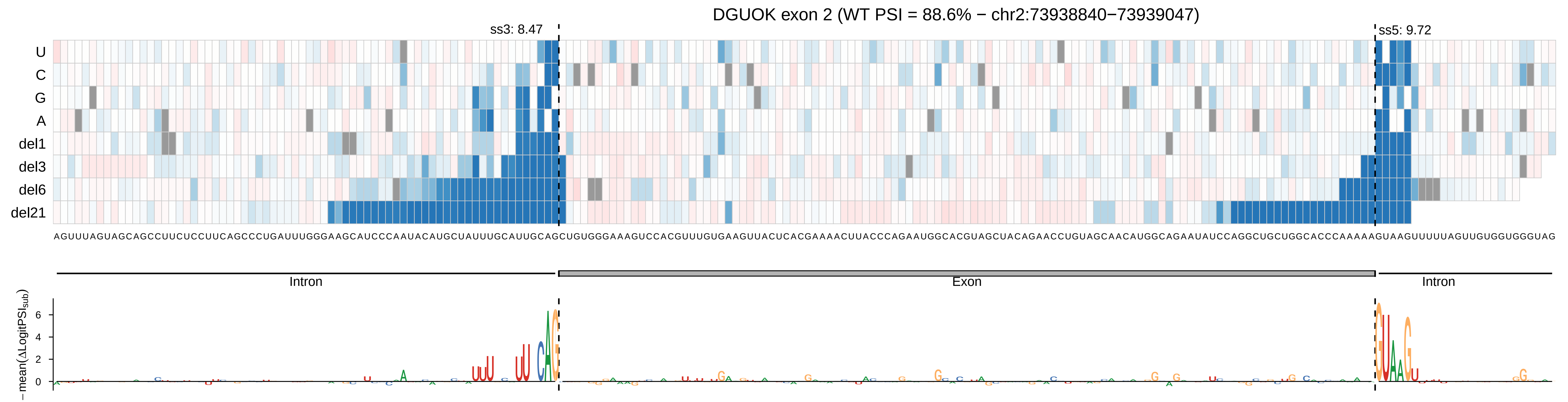

DGUOK exon 2 (WT PSI = 88.6% − chr2:73938840−73939047)
ss3: 8.47
ss5: 9.72
U
C
G
A
del1
del3
del6
del21
AGUUUAGUAGCAGCCUUCUCCUUCAGCCCUGAUUUGGGAAGCAUCCCAAUACAUGCUAUUUGCAUUGCAGCUGUGGGAAAGUCCACGUUUGUGAAGUUACUCACGAAAACUUACCCAGAAUGGCACGUAGCUACAGAACCUGUAGCAACAUGGCAGAAUAUCCAGGCUGCUGGCACCCAAAAAGUAAGUUUUUAGUUGUGGUGGGUAG
Intron
Exon
Intron
−mean(ΔLogitPSI_sub)
6
4
2
0

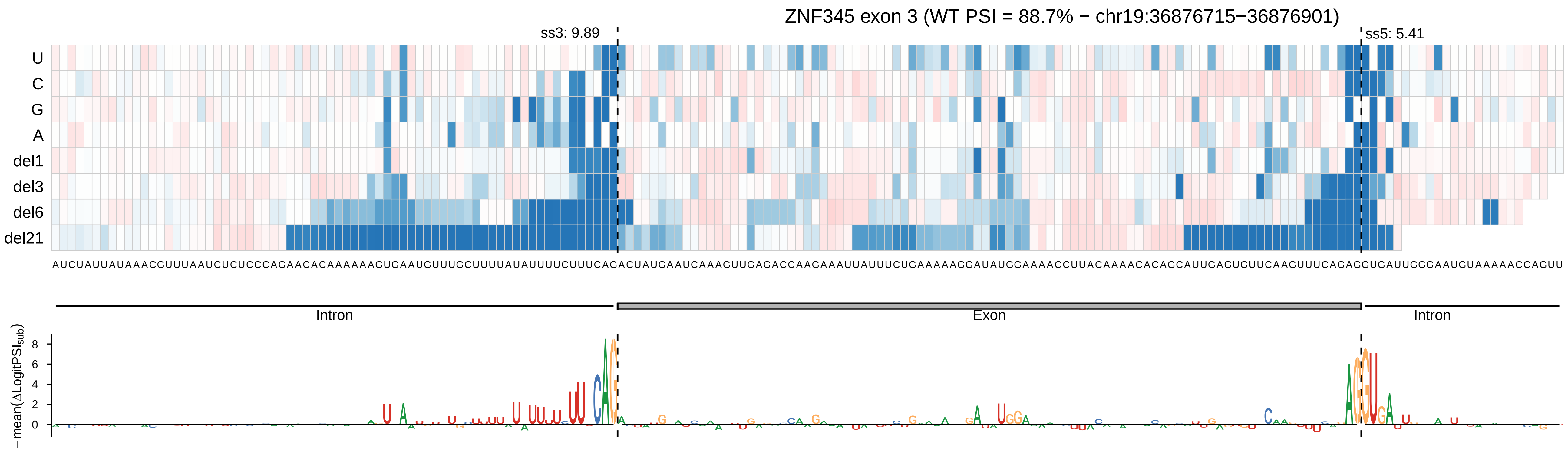
ZNF345 exon 3 (WT PSI = 88.7% − chr19:36876715−36876901)
ss3: 9.89
ss5: 5.41
U
C
G
A
del1
del3
del6
del21
AUCUAUUAUAAACGUUUAAUCUCUCCCAGAACACAAAAAAGUGAAUGUUUGCUUUUAUAUUUUCUUUCAGACUAUGAAUCAAAGUUGAGACCAAGAAAUUAUUUCUGAAAAAGGAUAUGGAAAACCUUACAAAACACAGCAUUGAGUGUUCAAGUUUCAGAGGUGAUUGGGAAUGUAAAAACCAGUU
Intron
Exon
Intron
− mean(ΔLogitPSI_sub)
8
6
4
2
0

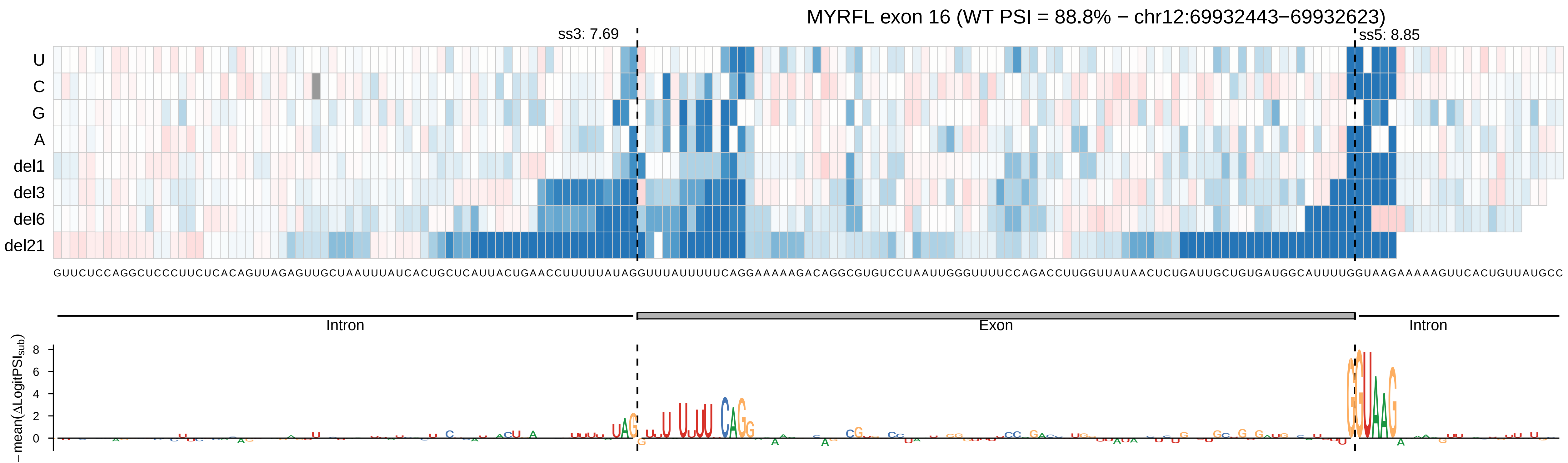

MYRFL exon 16 (WT PSI = 88.8% − chr12:69932443−69932623)
ss3: 7.69
ss5: 8.85
U
C
G
A
del1
del3
del6
del21
GUUCUCCAGGCUCCCUUCUCACAGUUAGAGUUGCUAAUUUAUCACUGCUCAUUACUGAACCUUUUUAUAGGUUUAUUUUUCAGGAAAAAGACAGGCGUGUCCUAAUUGGGUUUUCCAGACCUUGGUUAUAACUCUGAUUGCUGUGAUGGCAUUUUGGUAAGAAAAAGUUCACUGUUAUGCC
Intron
Exon
Intron
− mean(ΔLogitPSIsub)
8
6
4
2
0

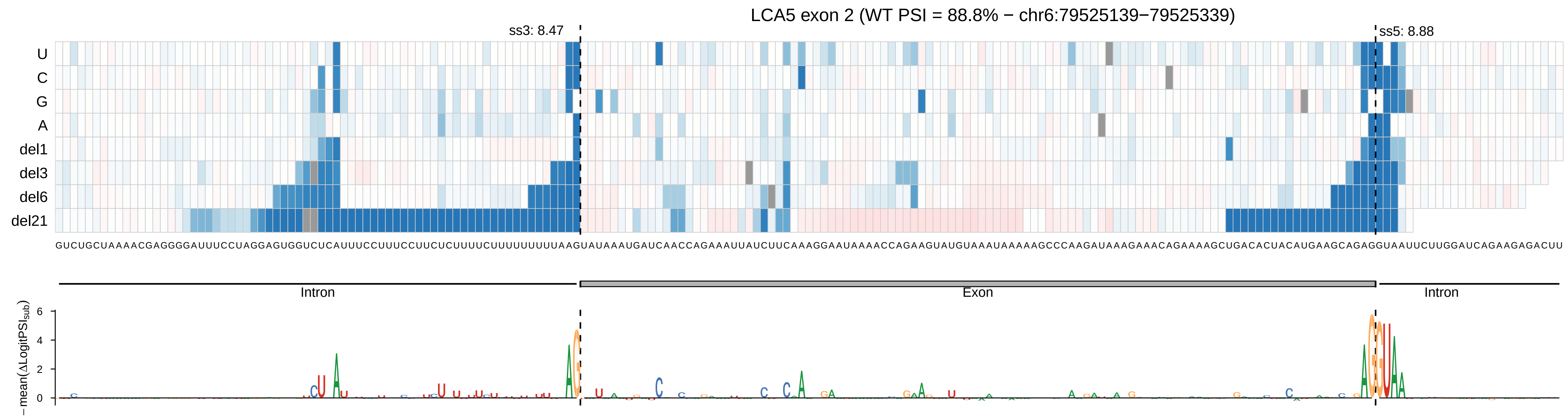
LCA5 exon 2 (WT PSI = 88.8% − chr6:79525139−79525339)
ss3: 8.47
ss5: 8.88
U
C
G
A
del1
del3
del6
del21
GUCUGCUAAAACGAGGGGAUUUCCUAGGAGUGGUCUCAUUUCCUUUCCUUCUCUUUUCUUUUUUUUUUAAGUAUAAAUGAUCAACCAGAAAUUAUCUUCAAAGGAAUAAAACCAGAAGUAUGUAAAUAAAAAGCCCAAGAUAAAGAAACAGAAAAGCUGACACUACAUGAAGCAGAGGUAAUUCUUGGAUCAGAAGAGACUU
Intron
Exon
Intron
− mean(ΔLogitPSI_sub)
6
4
2
0

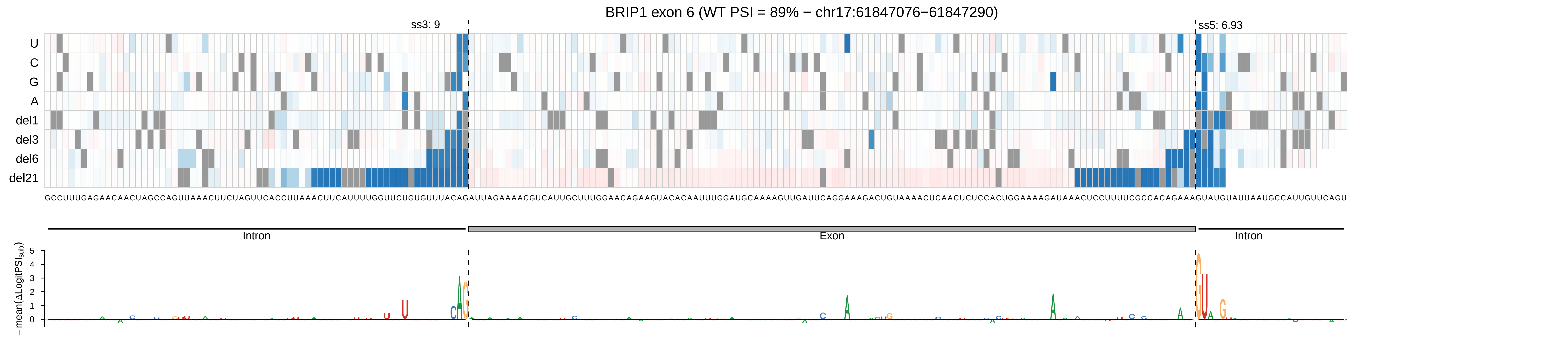
BRIP1 exon 6 (WT PSI = 89% − chr17:61847076−61847290)
ss3: 9
ss5: 6.93
U
C
G
A
del1
del3
del6
del21
GCCUUUGAGAACAACUAGCCAGUUAAACUUCUAGUUCACCUUAAACUUCAUUUUGGUUCUGUGUUUACAGAUUAGAAAACGUCAUUGCUUUGGAACAGAAGUACACAAUUUGGAUGCAAAAGUUGAUUCAGGAAAGACUGUAAAACUCAACUCUCCACUGGAAAAGAUAAACUCCUUUUCGCCACAGAAAGUAUGUAUUAAUGCCAUUGUUCAGU
Intron
Exon
Intron
−mean(ΔLogitPSI_sub)
5
4
3
2
1
0

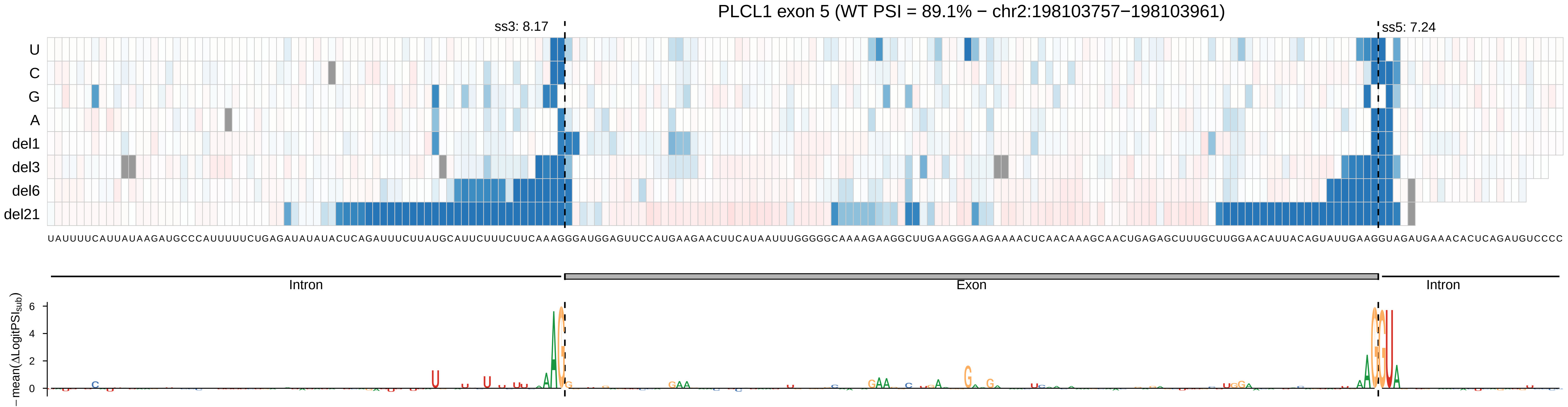

PLCL1 exon 5 (WT PSI = 89.1% − chr2:198103757−198103961)
ss3: 8.17
ss5: 7.24
U
C
G
A
del1
del3
del6
del21
UAUUUUCAUUAUAAGAUGCCCAUUUUUCUGAGAUAUAUACUCAGAUUUCUUAUGCAUUCUUUCUUCAAAGGGAUGGAGUUCCAUGAAGAACUUCAUAAUUUGGGGGCAAAAGAAGGCUUGAAGGGAAGAAAACUCAACAAAGCAACUGAGAGCUUUGCUUGGAACAUUACAGUAUUGAAGGUAGAUGAAACACUCAGAUGUCCCC
Intron
Exon
Intron
− mean(ΔLogitPSI_sub)
6
4
2
0

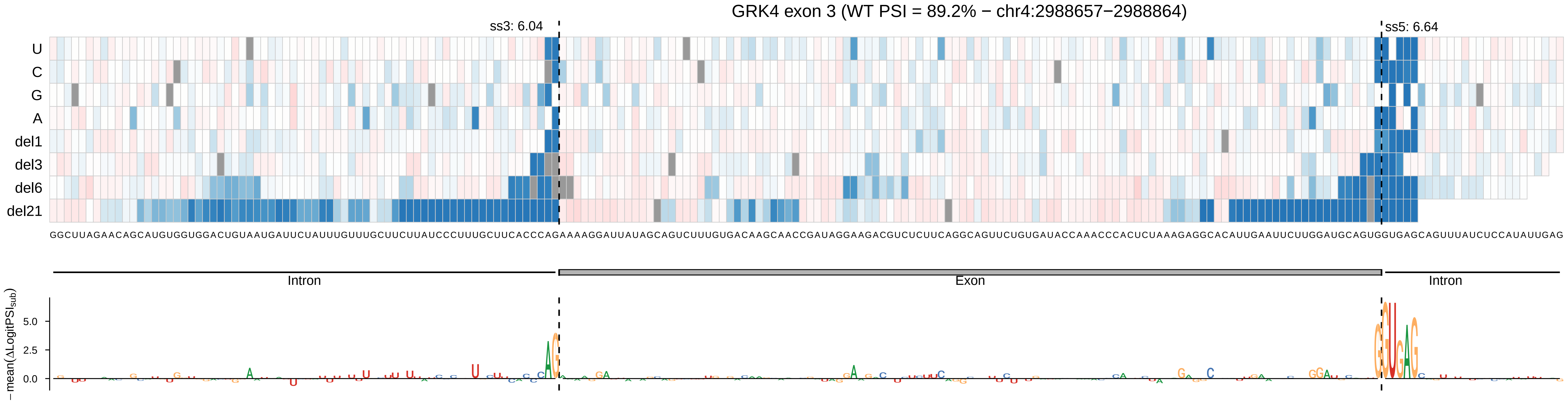
GRK4 exon 3 (WT PSI = 89.2% − chr4:2988657−2988864)
ss3: 6.04
ss5: 6.64
U
C
G
A
del1
del3
del6
del21
GGCUUAGAACAGCAUGUGGUGGACUGUAAUGAUUCUAUUUGUUUGCUUCUUAUCCCUUUGCUUCACCCAGAAAAGGAUUAUAGCAGUCUUUGUGACAAGCAACCGAUAGGAAGACGUCUCUUCAGGCAGUUCUGUGAUACCAAACCCACUCUAAAGAGGCACAUUGAAUUCUUGGAUGCAGUGGUGAGCAGUUUAUCUCCAUAUUGAG
Intron
Exon
Intron
− mean(ΔLogitPSI_sub)
5.0
2.5
0.0

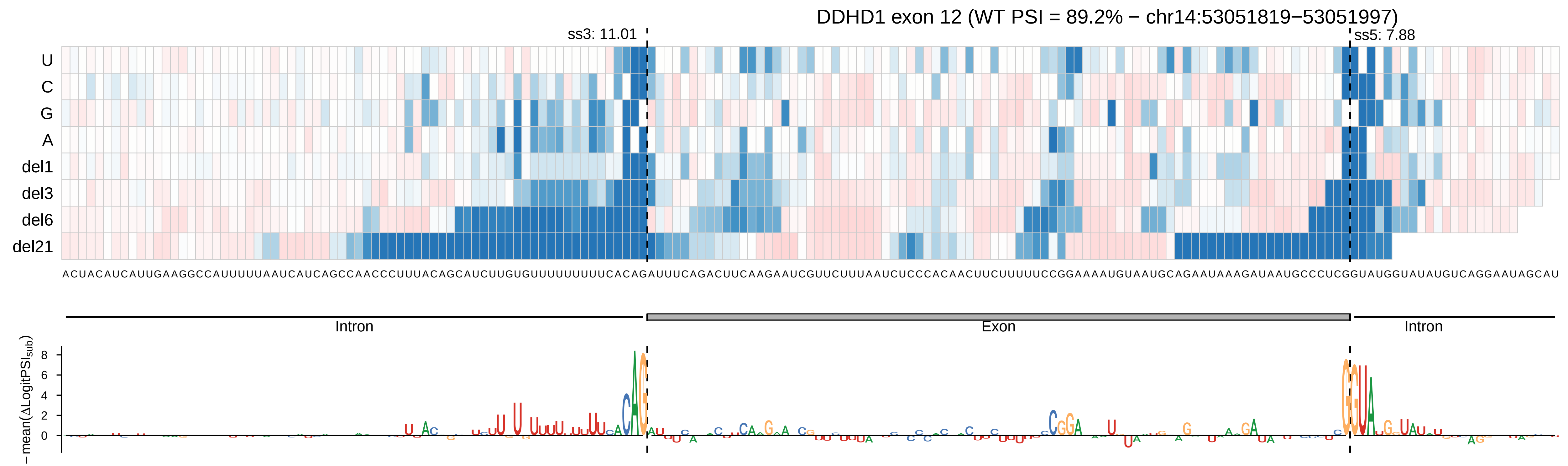
DDHD1 exon 12 (WT PSI = 89.2% − chr14:53051819−53051997)
ss3: 11.01
ss5: 7.88
U
C
G
A
del1
del3
del6
del21
ACUACAUCAUUGAAGGCCAUUUUUAAUCAUCAGCCAACCCUUUACAGCAUCUUGUGUUUUUUUUUCACAGAUUUCAGACUUCAAGAAUCGUUCUUUAAUCUCCCACAACUUCUUUUUCCGGAAAAUGUAAUGCAGAAUAAAGAUAAUGCCCUCGGUAUGGUAUAUGUCAGGAAUAGCAU
Intron
Exon
Intron
− mean(ΔLogitPSI_sub)
8
6
4
2
0

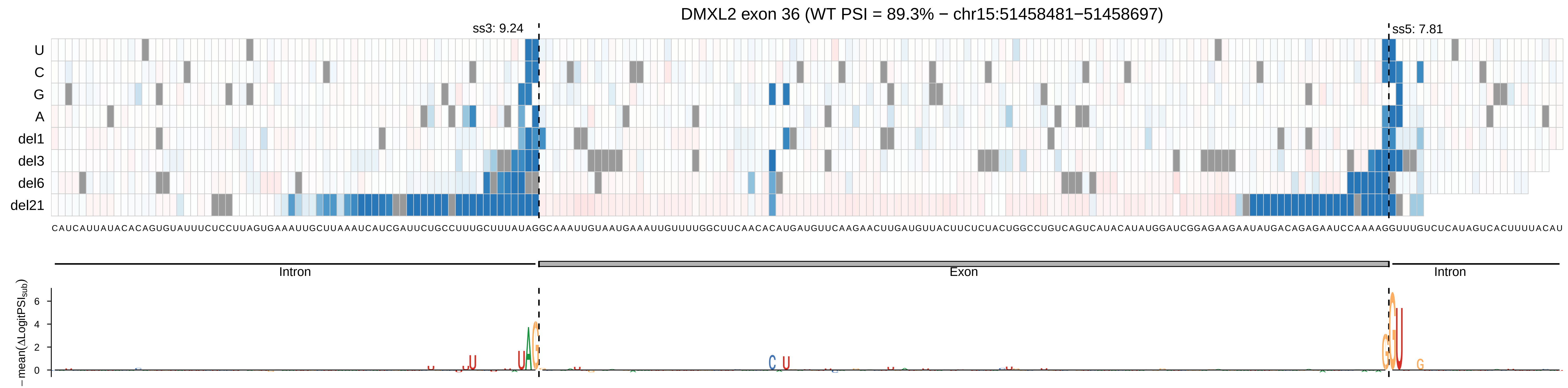
DMXL2 exon 36 (WT PSI = 89.3% − chr15:51458481−51458697)
ss3: 9.24
ss5: 7.81
U
C
G
A
del1
del3
del6
del21
CAUCAUUAUACACAGUGUAUUUCUCCUUAGUGAAAUUGCUUAAAUCAUCGAUUCUGCCUUUGCUUUAUAGGCAAAUUGUAAUGAAAUUGUUUUGGCUUCAACACAUGAUGUUCAAGAACUUGAUGUUACUUCUCUACUGGCCUGUCAGUCAUACAUAUGGAUCGGAGAAGAAUAUGACAGAGAAUCCAAAAGGUUUGUCUCAUAGUCACUUUUACAU
Intron
Exon
Intron
− mean(ΔLogitPSI_sub)
6
4
2
0

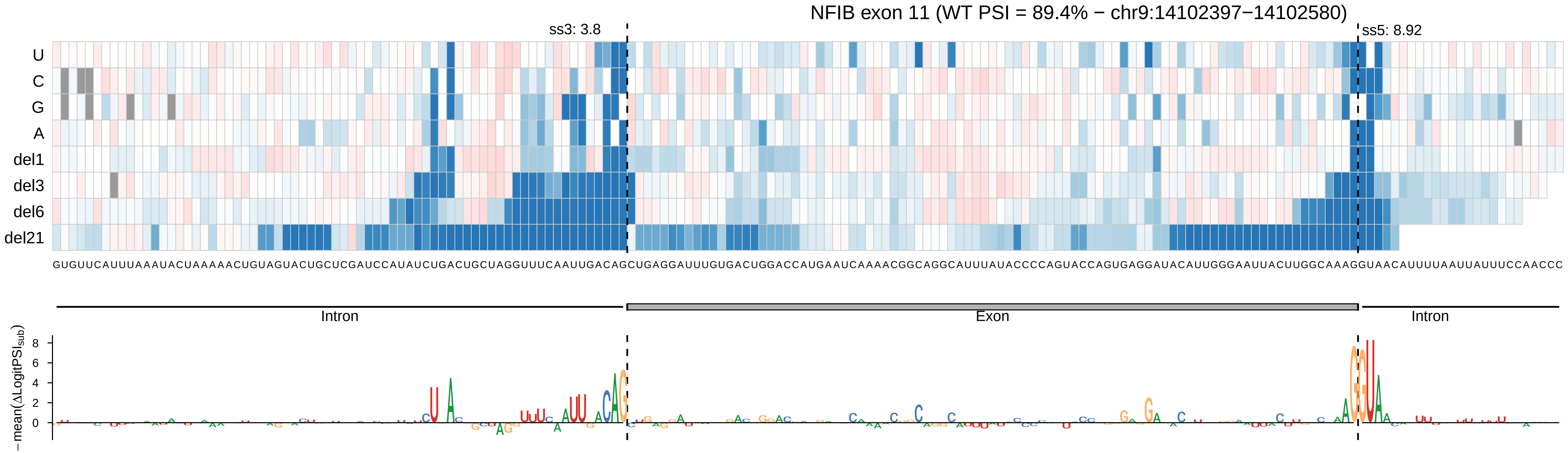
NFIB exon 11 (WT PSI = 89.4% − chr9:14102397−14102580)
ss3: 3.8
ss5: 8.92
U
C
G
A
del1
del3
del6
del21
GUGUUCAUUUAAAUACUAAAAACUGUAGUACUGCUCGAUCCAUAUCUGACUGCUAGGUUUCAAUUGACAGCUGAGGAUUUGUGACUGGACCAUGAAUCAAAACGGCAGGCAUUUAUACCCCAGUACCAGUGAGGAUACAUUGGGAAUUACUUGGCAAAGGUAACAUUUUAAUUAUUUCCAACCC
Intron
Exon
Intron
− mean(ΔLogitPSI$_{sub}$)
8
6
4
2
0

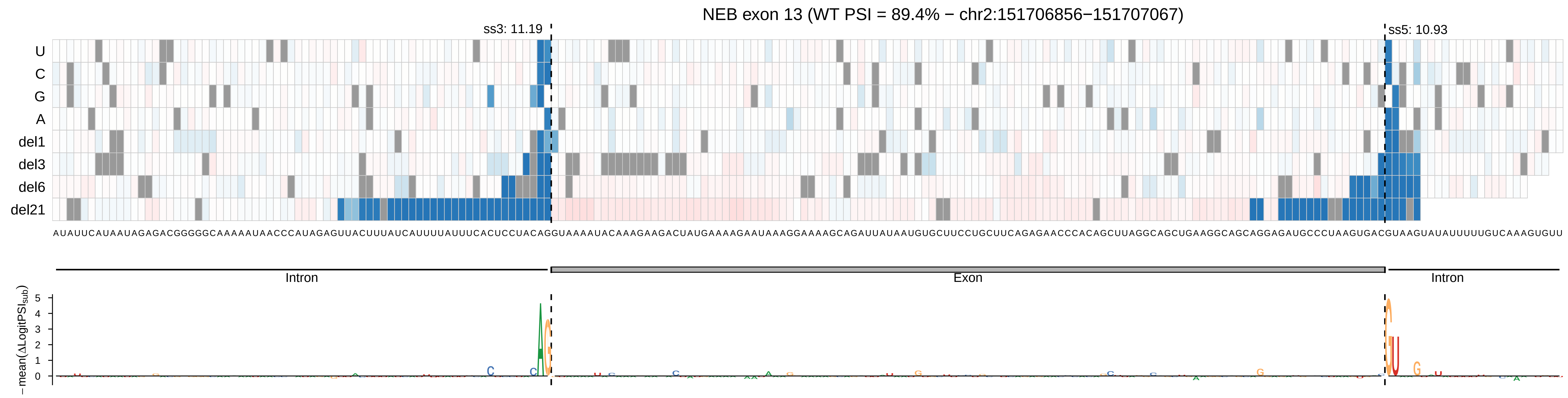
NEB exon 13 (WT PSI = 89.4% − chr2:151706856−151707067)
ss3: 11.19
ss5: 10.93
U
C
G
A
del1
del3
del6
del21
Intron
Exon
Intron
−mean(ΔLogitPSI_sub)
0
1
2
3
4
5

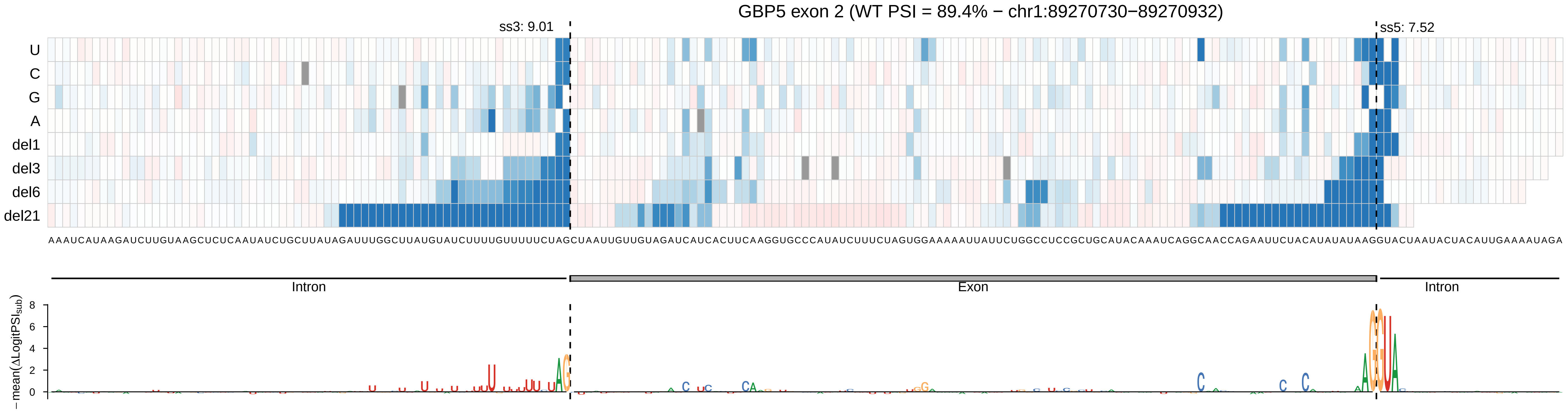
GBP5 exon 2 (WT PSI = 89.4% − chr1:89270730−89270932)
ss3: 9.01
ss5: 7.52
U
C
G
A
del1
del3
del6
del21
AAAUCAUAAGAUCUUGUAAGCUCUCAAUAUCUGCUUAUAGAUUUGGCUUAUGUAUCUUUUGUUUUUCUAGCUAAUUGUUGUAGAUCAUCACUUCAAGGUGCCCAUAUCUUUCUAGUGGAAAAAUUAUUCUGGCCUCCGCUGCAUACAAAUCAGGCAACCAGAAUUCUACAUAUAUAAGGUACUAAUACUACAUUGAAAAUAGA
Intron
Exon
Intron
− mean(ΔLogitPSIsub)
8
6
4
2
0

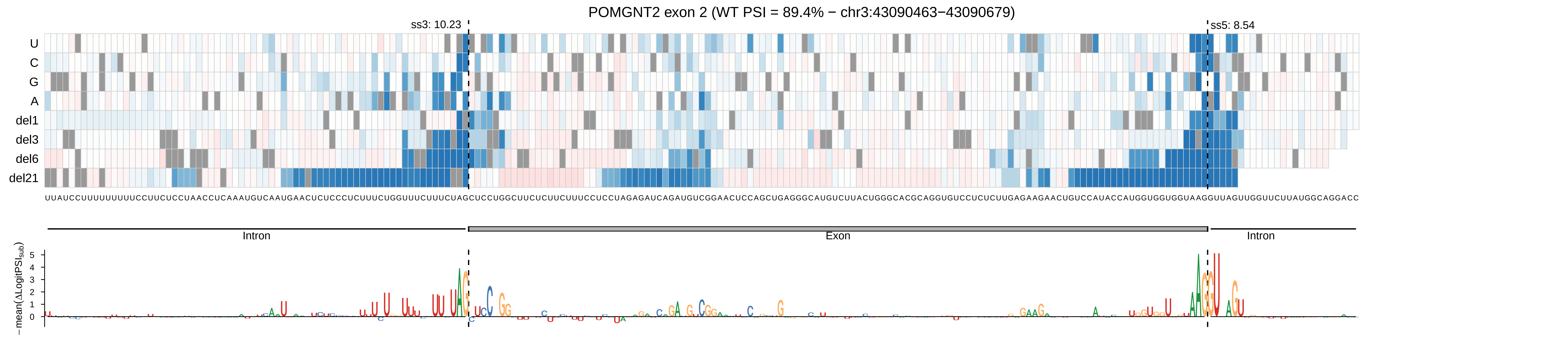

POMGNT2 exon 2 (WT PSI = 89.4% − chr3:43090463−43090679)
ss3: 10.23
ss5: 8.54
U
C
G
A
del1
del3
del6
del21
Intron
Exon
Intron
−mean(ΔLogitPSI$_{sub}$)

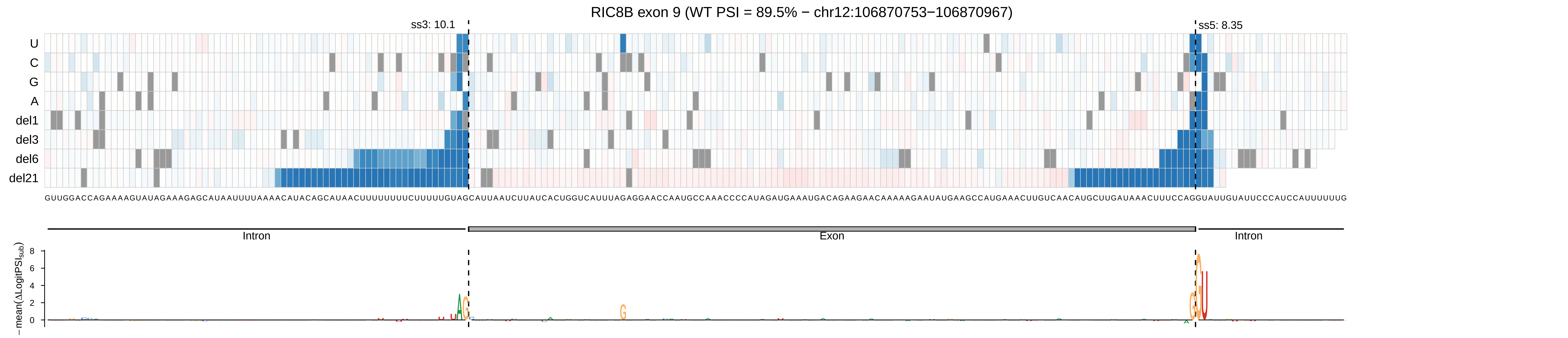

RIC8B exon 9 (WT PSI = 89.5% − chr12:106870753−106870967)
ss3: 10.1
ss5: 8.35
U
C
G
A
del1
del3
del6
del21
GUUGGACCAGAAAAGUAUAGAAAGAGCAUAAUUUUAAAACAUACAGCAUAACUUUUUUUUCUUUUUGUAGCAUUAAUCUUAUCACUGGUCAUUUAGAGGAACCAAUGCCAAACCCCAUAGAUGAAAUGACAGAAGAACAAAAAGAAUAUGAAGCCAUGAAACUUGUCAACAUGCUUGAUAAACUUUCCAGGUAUUGUAUUCCCAUCCAUUUUUUG
Intron
Exon
Intron
−mean(ΔLogitPSI_sub)
8
6
4
2
0

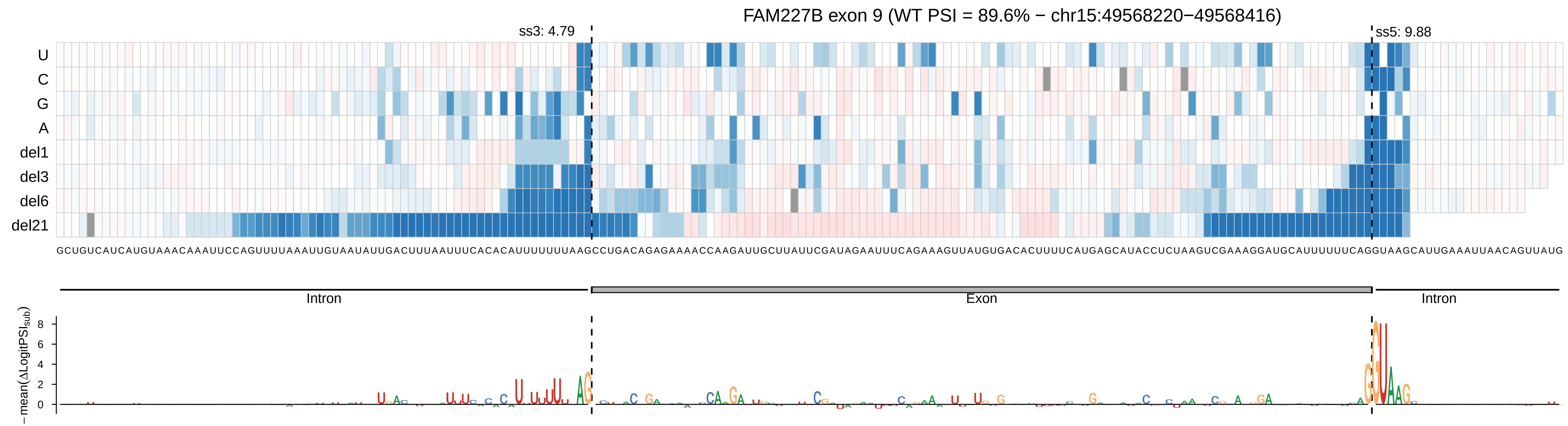
FAM227B exon 9 (WT PSI = 89.6% − chr15:49568220−49568416)
ss3: 4.79
ss5: 9.88
U
C
G
A
del1
del3
del6
del21
GCUGUCAUCAUGUAAACAAAUUCCAGUUUUAAAUUGUAAUAUUGACUUUAAUUUCACACAUUUUUUUAAGCCUGACAGAGAAAACCAAGAUUGCUUAUUCGAUAGAAUUUCAGAAAGUUAUGUGACACUUUUCAUGAGCAUACCUCUAAGUCGAAAGGAUGCAUUUUUUCAGGUAAGCAUUGAAAUUAACAGUUAUG
Intron
Exon
Intron
−mean(ΔLogitPSI_sub)
8
6
4
2
0

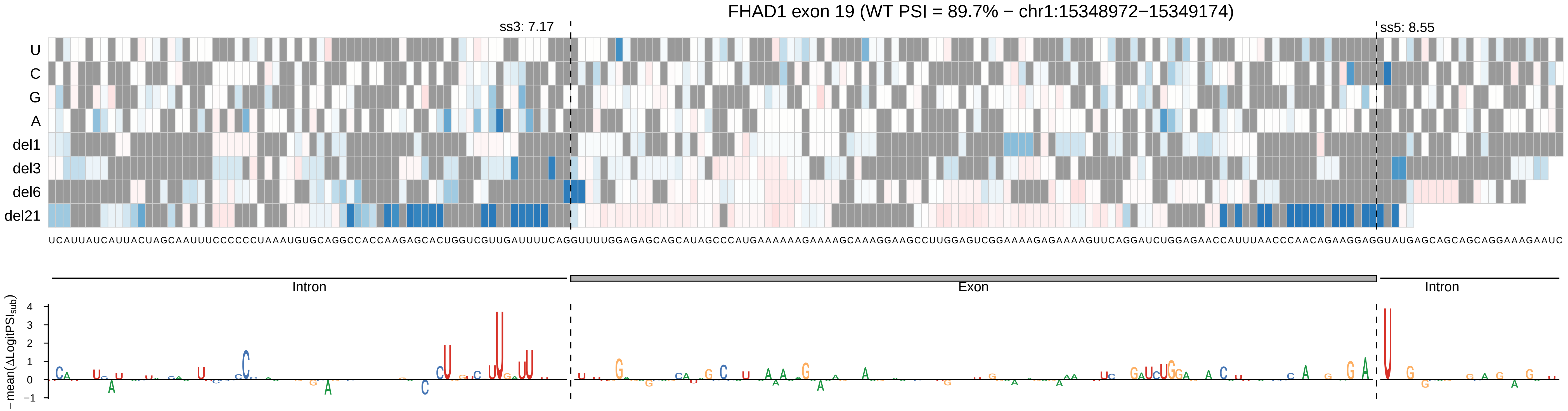

FHAD1 exon 19 (WT PSI = 89.7% − chr1:15348972−15349174)
ss3: 7.17
ss5: 8.55
U
C
G
A
del1
del3
del6
del21
UCAUUAUCAUUACUAGCAAUUUCCCCCCUAAAUGUGCAGGCCACCAAGAGCACUGGUCGUUGAUUUUCAGGUUUUGGAGAGCAGCAUAGCCCAUGAAAAAAGAAAAGCAAAGGAAGCCUUGGAGUCGGAAAAGAGAAAAGUUCAGGAUCUGGAGAACCAUUUAACCCAACAGAAGGAGGUAUGAGCAGCAGCAGGAAAGAAUC
Intron
Exon
Intron
−mean(ΔLogitPSI_sub)

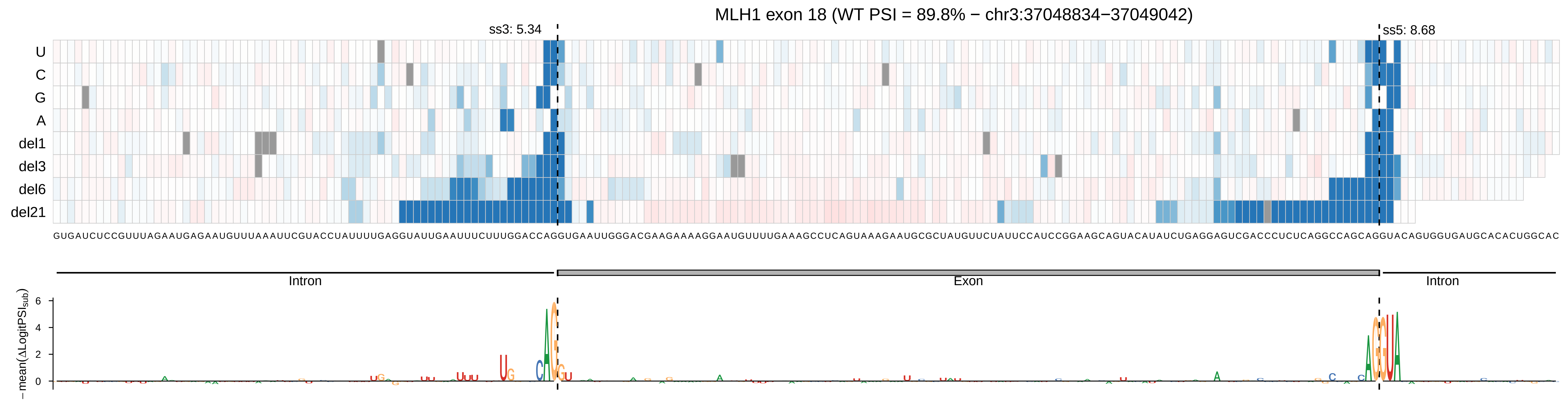

MLH1 exon 18 (WT PSI = 89.8% − chr3:37048834−37049042)
ss3: 5.34
ss5: 8.68
U
C
G
A
del1
del3
del6
del21
Intron
Exon
Intron
−mean(ΔLogitPSI_sub)
6
4
2
0

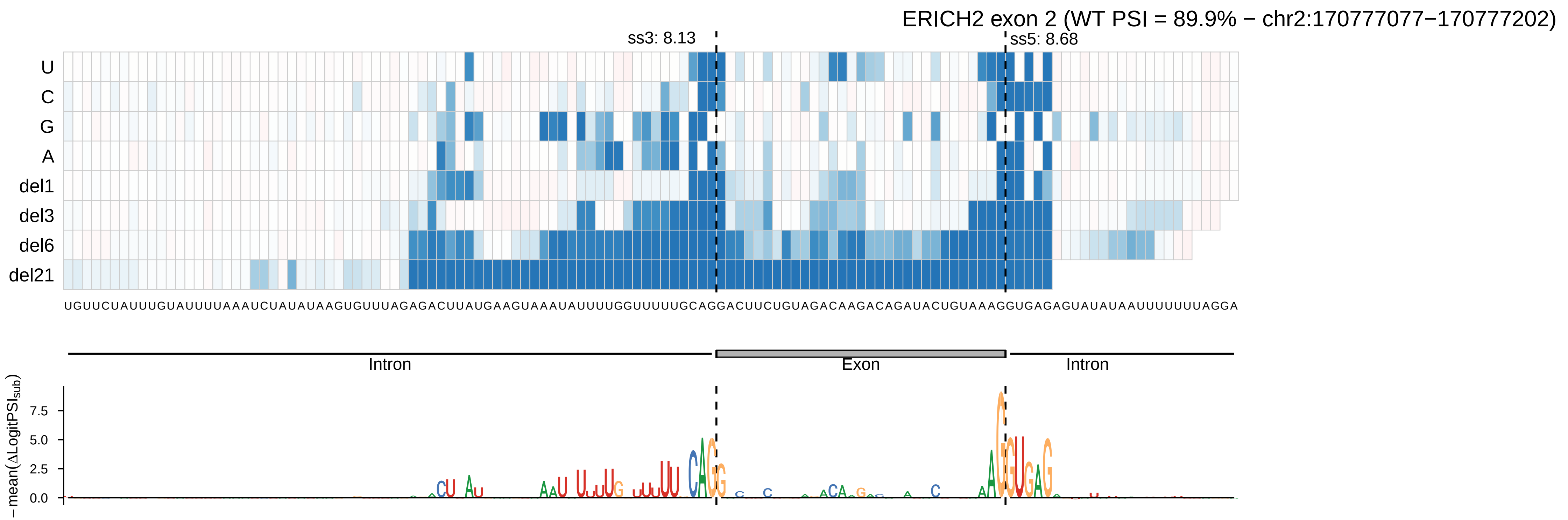
ERICH2 exon 2 (WT PSI = 89.9% − chr2:170777077−170777202)
ss3: 8.13
ss5: 8.68
U
C
G
A
del1
del3
del6
del21
UGUUCUAUUUGUAUUUUAAAUCUAUAUAAGUGUUUAGAGACUUAUGAAGUAAAUAUUUUGGUUUUUGCAGGACUUCUGUAGACAAGACAGAUACUGUAAAGGUGAGAGUAUAUAAUUUUUUUAGGA
Intron
Exon
Intron
− mean(ΔLogitPSI_sub)
7.5
5.0
2.5
0.0

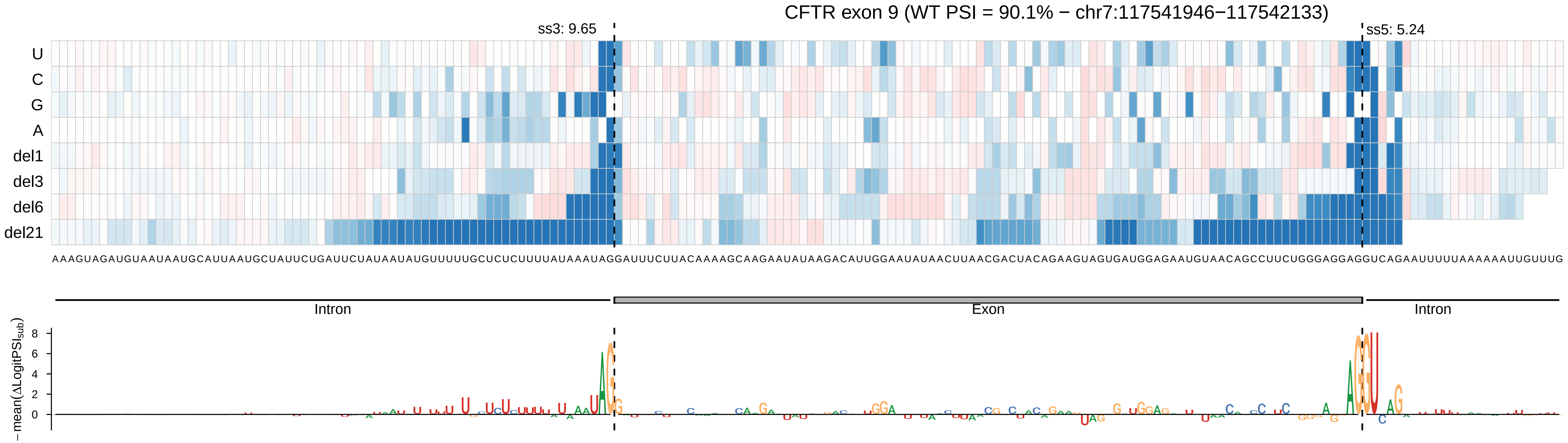

CFTR exon 9 (WT PSI = 90.1% − chr7:117541946−117542133)
ss3: 9.65
ss5: 5.24
U
C
G
A
del1
del3
del6
del21
AAAGUAGAUGUAAUAAUGCAUUAAUGCUAUUCUGAUUCUAUAAUAUGUUUUUGCUCUCUUUUAUAAAUAGGAUUUCUUACAAAAGCAAGAAUAUAAGACAUUGGAAUAUAACUUAACGACUACAGAAGUAGUGAUGGAGAAUGUAACAGCCUUCUGGGAGGAGGUCAGAAUUUUUAAAAAAUUGUUUG
Intron
Exon
Intron
−mean(ΔLogitPSI sub)
8
6
4
2
0

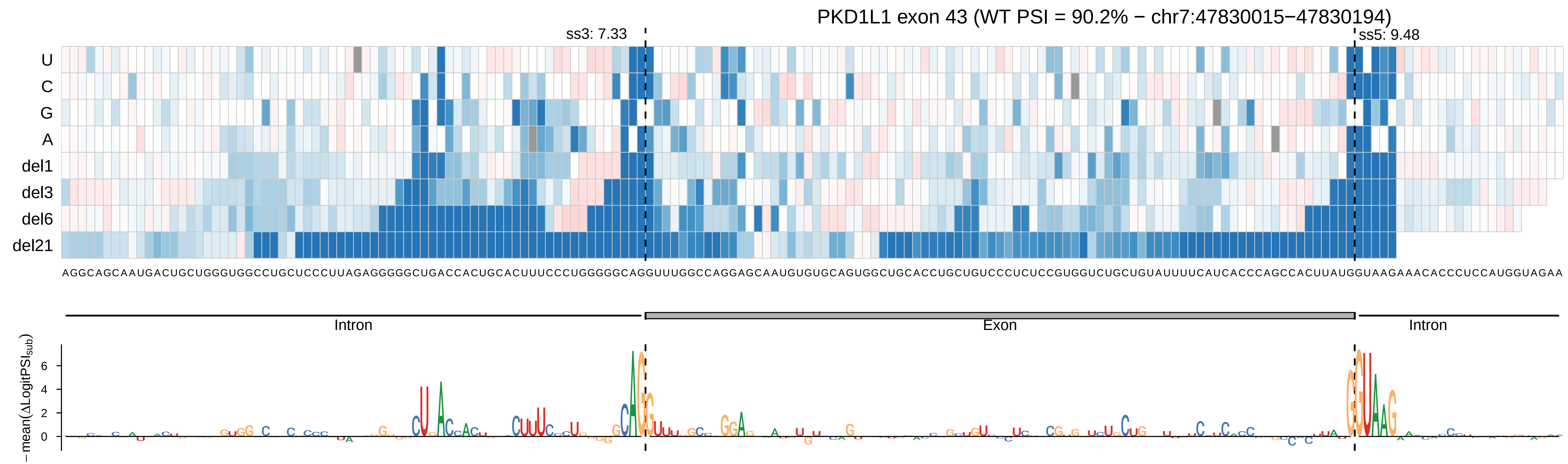

PKD1L1 exon 43 (WT PSI = 90.2% − chr7:47830015−47830194)
ss3: 7.33
ss5: 9.48
U
C
G
A
del1
del3
del6
del21
Intron
Exon
Intron
−mean(ΔLogitPSI_sub)
6
4
2
0

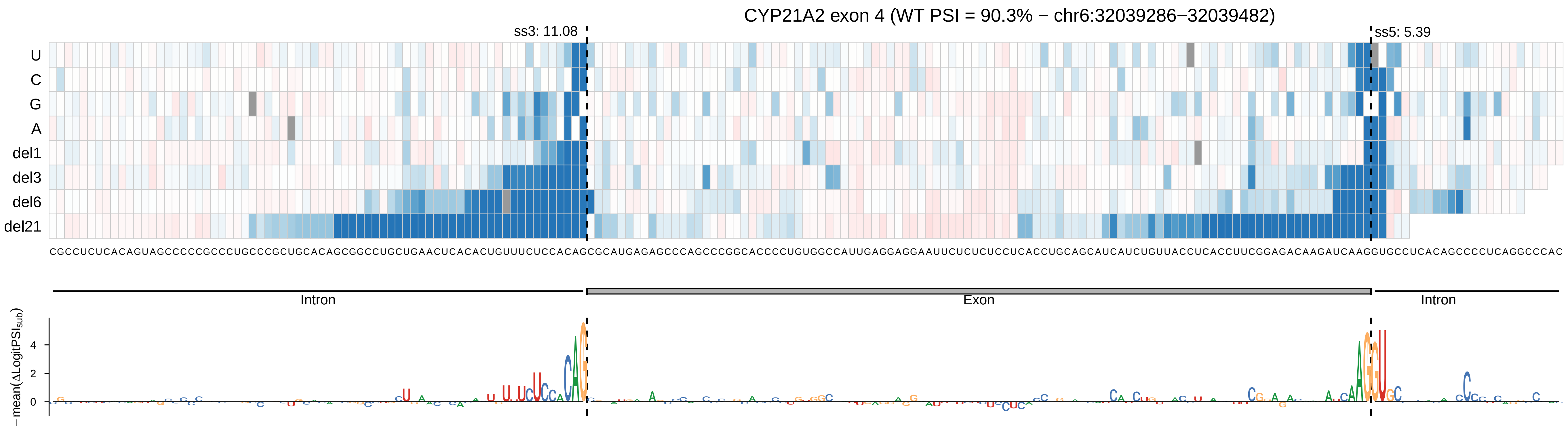

CYP21A2 exon 4 (WT PSI = 90.3% − chr6:32039286−32039482)
ss3: 11.08
ss5: 5.39
U
C
G
A
del1
del3
del6
del21
CGCCUCUCACAGUAGCCCCCGCCCUGCCCGCUGCACAGCGGCCUGCUGAACUCACACUGUUUCUCCACAGCGCAUGAGAGCCCAGCCCGGCACCCCUGUGGCCAUUGAGGAGGAAUUCUCUCUCCUCACCUGCAGCAUCAUCUGUUACCUCACCUUCGGAGACAAGAUCAAGGUGCCUCACAGCCCCUCAGGCCCAC
Intron
Exon
Intron
−mean(ΔLogitPSI$_{sub}$)
4
2
0

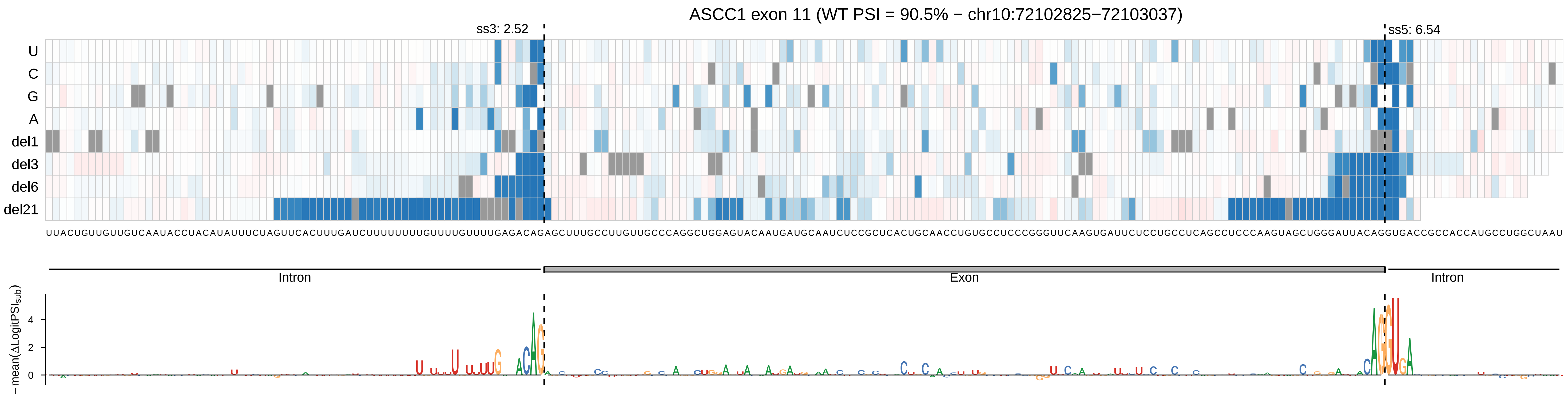
ASCC1 exon 11 (WT PSI = 90.5% − chr10:72102825−72103037)
ss3: 2.52
ss5: 6.54
U
C
G
A
del1
del3
del6
del21
UUACUGUUGUUGUCAAUACCUACAUAUUUCUAGUUCACUUUGAUCUUUUUUUUGUUUUGUUUUGAGACAGAGCUUUGCCUUGUUGCCCAGGCUGGAGUACAAUGAUGCAAUCUCCGCUCACUGCAACCUGUGCCUCCCGGGUUCAAGUGAUUCUCCUGCCUCAGCCUCCCAAGUAGCUGGGAUUACAGGUGACCGCCACCAUGCCUGGCUAAU
Intron
Exon
Intron
− mean(ΔLogitPSI_sub)
4
2
0

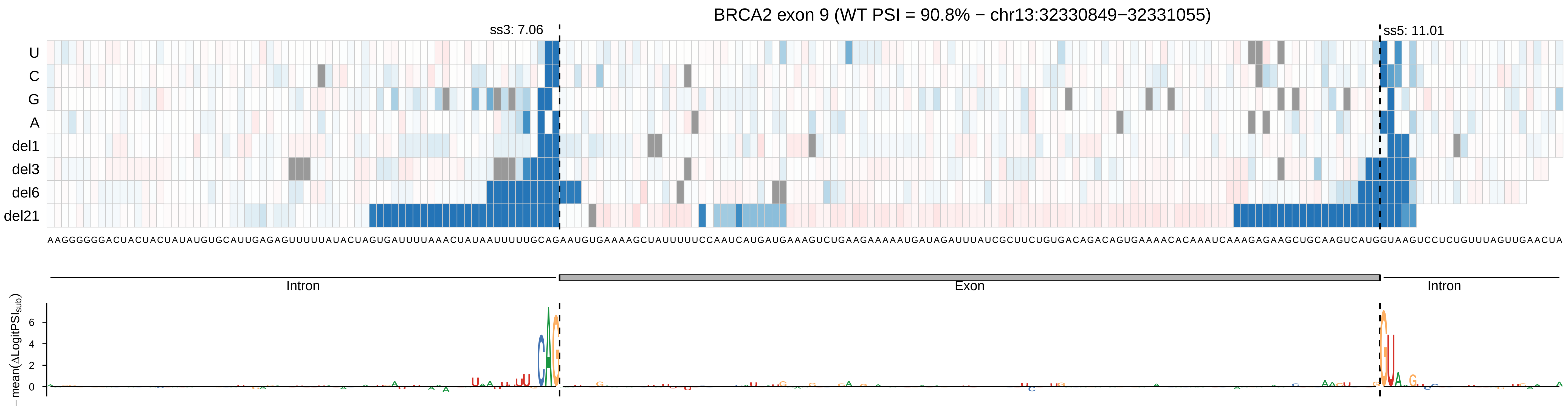
BRCA2 exon 9 (WT PSI = 90.8% − chr13:32330849−32331055)
ss3: 7.06
ss5: 11.01
U
C
G
A
del1
del3
del6
del21
AAGGGGGGACUACUACUAUAUGUGCAUUGAGAGUUUUUAUACUAGUGAUUUUAAACUAUAAUUUUUGCAGAAUGUGAAAAGCUAUUUUUCCAAUCAUGAUGAAAGUCUGAAGAAAAAUGAUAGAUUUAUCGCUUCUGUGACAGACAGUGAAAACACAAAUCAAAGAGAAGCUGCAAGUCAUGGUAAGUCCUCUGUUUAGUUGAACUA
Intron
Exon
Intron
− mean(ΔLogitPSI_sub)
6
4
2
0

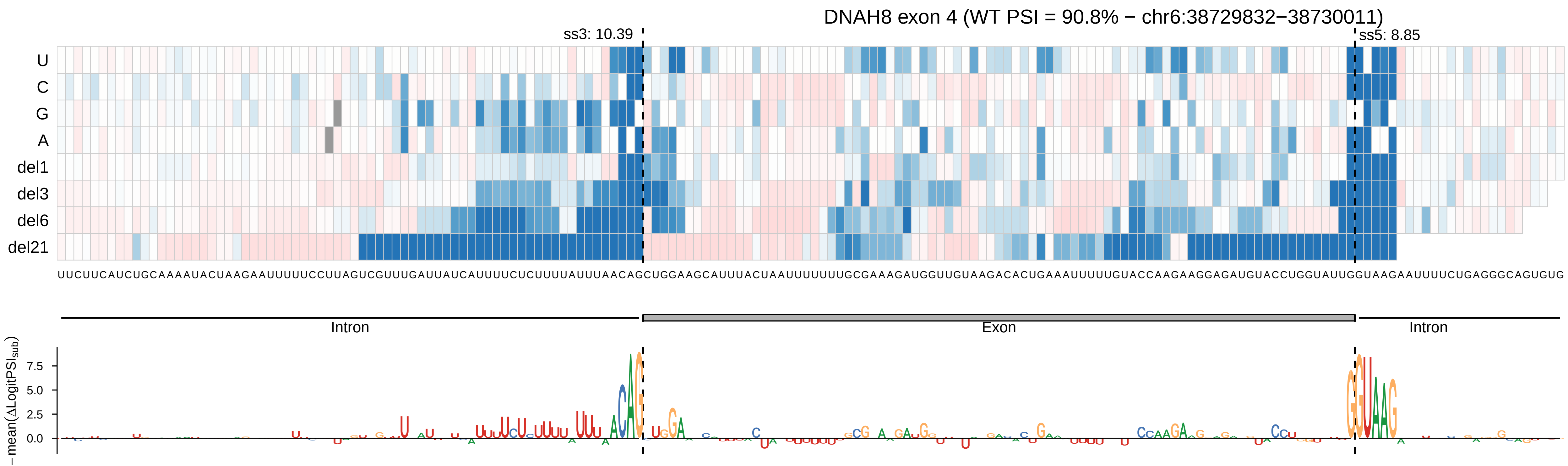

DNAH8 exon 4 (WT PSI = 90.8% − chr6:38729832−38730011)
ss3: 10.39
ss5: 8.85
U
C
G
A
del1
del3
del6
del21
UUCUUCAUCUGCAAAAUACUAAGAAUUUUUCCUUAGUCGUUUGAUUAUCAUUUUCUCUUUUAUUUAACAGCUGGAAGCAUUUACUAAUUUUUUUGCGAAAGAUGGUUGUAAGACACUGAAAUUUUUGUACCAAGAAGGAGAUGUACCUGGUAUUGGUAAGAAUUUUCUGAGGGCAGUGUG
Intron
Exon
Intron
−mean(ΔLogitPSIsub)
7.5
5.0
2.5
0.0

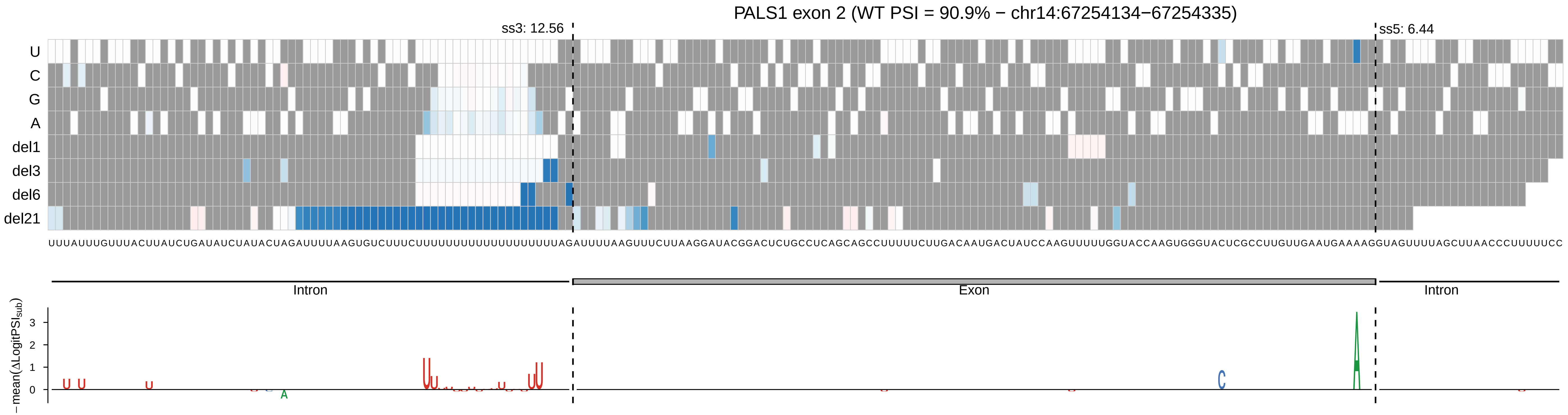

PALS1 exon 2 (WT PSI = 90.9% − chr14:67254134−67254335)
ss3: 12.56
ss5: 6.44
U
C
G
A
del1
del3
del6
del21
UUUAUUUGUUUACUUAUCUGAUAUCUAUACUAGAUUUUAAGUGUCUUUCUUUUUUUUUUUUUUUUUUAGAUUUUAAGUUUCUUAAGGAUACGGACUCUGCCUCAGCAGCCUUUUUCUUGACAAUGACUAUCCAAGUUUUUGGUACCAAGUGGGUACUCGCCUUGUUGAAUGAAAAGGUAGUUUUAGCUUAACCCUUUUUCC
Intron
Exon
Intron
−mean(ΔLogitPSI_sub)
0
1
2
3

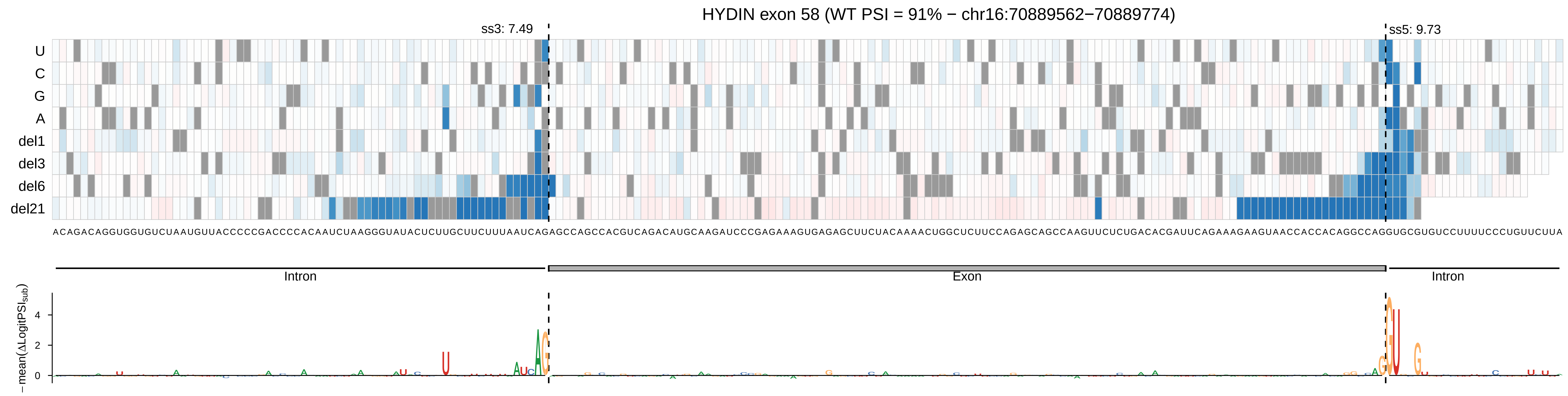

HYDIN exon 58 (WT PSI = 91% − chr16:70889562−70889774)
ss3: 7.49
ss5: 9.73
U
C
G
A
del1
del3
del6
del21
Intron
Exon
Intron
−mean(ΔLogitPSI_sub)

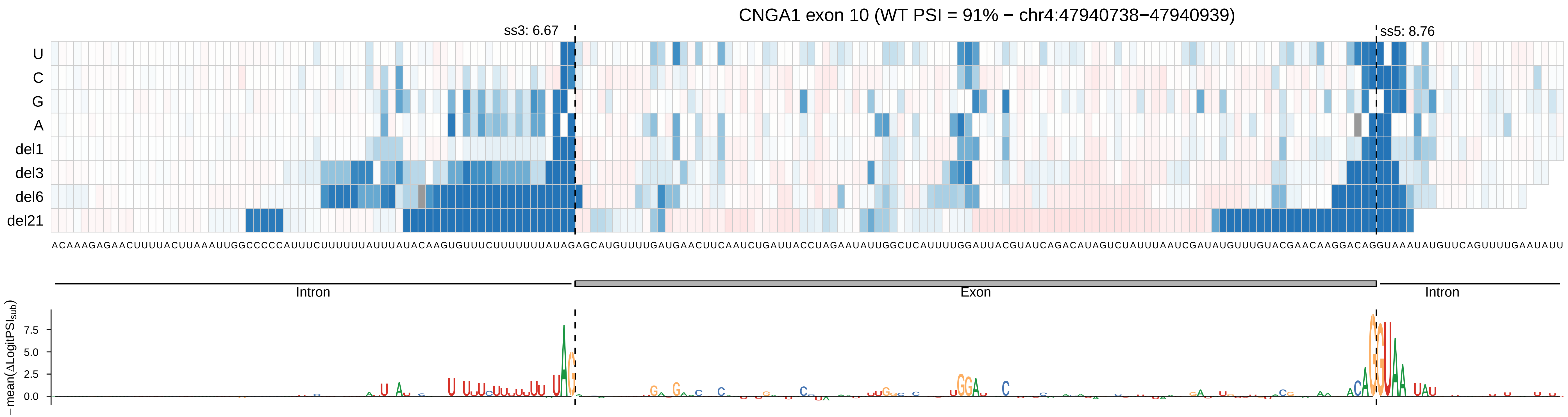

CNGA1 exon 10 (WT PSI = 91% − chr4:47940738−47940939)
ss3: 6.67
ss5: 8.76
U
C
G
A
del1
del3
del6
del21
ACAAAGAGAACUUUUACUUAAAUUGGCCCCCAUUUCUUUUUUAUUUAUACAAGUGUUUCUUUUUUUAUAGAGCAUGUUUUGAUGAACUUCAAUCUGAUUACCUAGAAUAUUGGCUCAUUUUGGAUUACGUAUCAGACAUAGUCUAUUUAAUCGAUAUGUUUGUACGAACAAGGACAGGUAAAUAUGUUCAGUUUUGAAUAUU
Intron
Exon
Intron
− mean(ΔLogitPSI_sub)
7.5
5.0
2.5
0.0

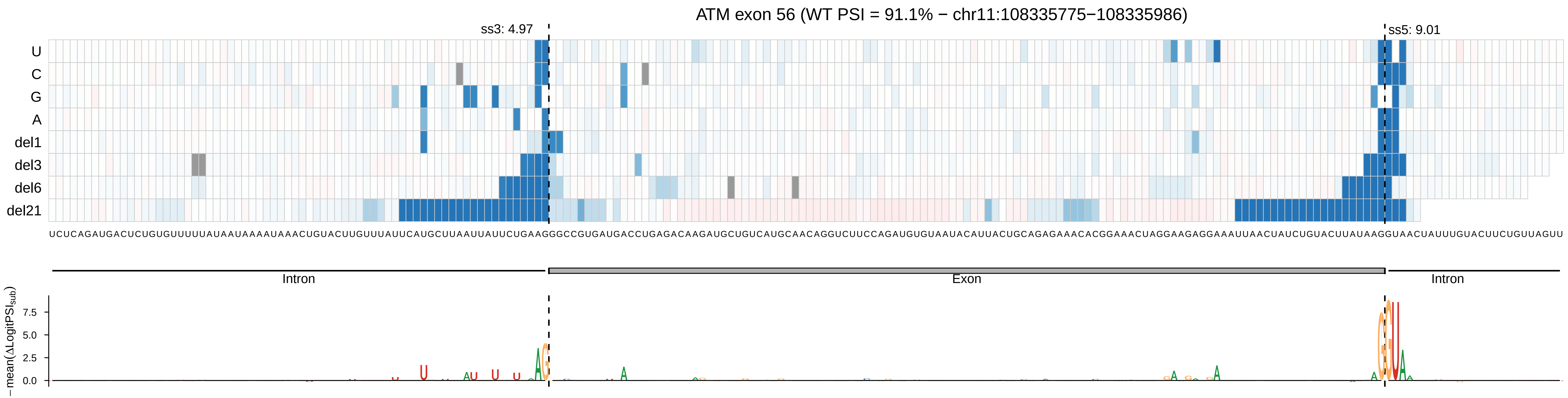
ATM exon 56 (WT PSI = 91.1% − chr11:108335775−108335986)
ss3: 4.97
ss5: 9.01
U
C
G
A
del1
del3
del6
del21
UCUCAGAUGACUCUGUGUUUUUAUAAUAAAAUAAACUGUACUUGUUUAUUCAUGCUUAAUUAUUCUGAAGGGCCGUGAUGACCUGAGACAAGAUGCUGUCAUGCAACAGGUCUUCCAGAUGUGUAAUACAUUACUGCAGAGAAACACGGAAACUAGGAAGAGGAAAUUAACUAUCUGUACUUAUAAGGUAACUAUUUGUACUUCUGUUAGUU
Intron
Exon
Intron
− mean(ΔLogitPSI_sub)
7.5
5.0
2.5
0.0

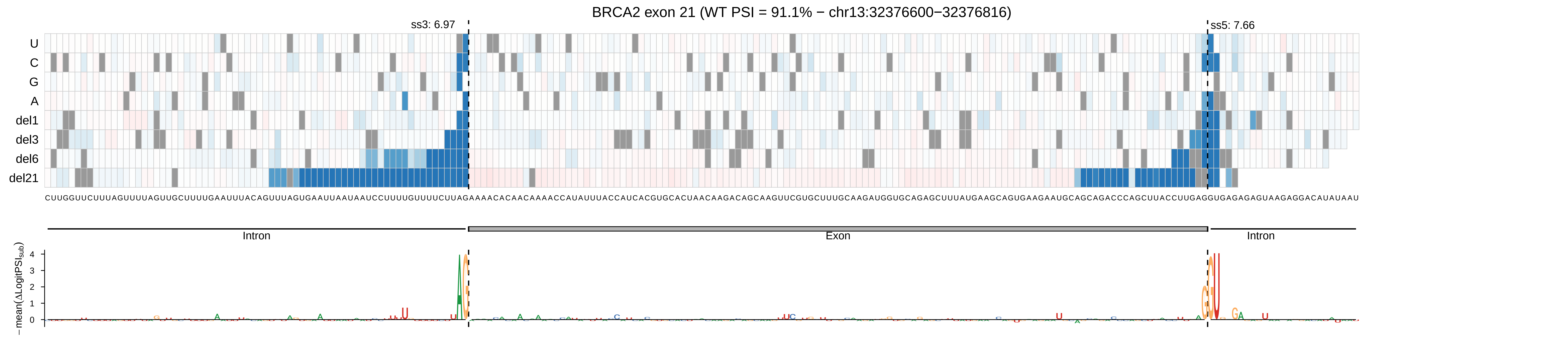

BRCA2 exon 21 (WT PSI = 91.1% − chr13:32376600−32376816)
ss3: 6.97
ss5: 7.66
U
C
G
A
del1
del3
del6
del21
Intron
Exon
Intron
−mean(ΔLogitPSI_sub)
0
1
2
3
4

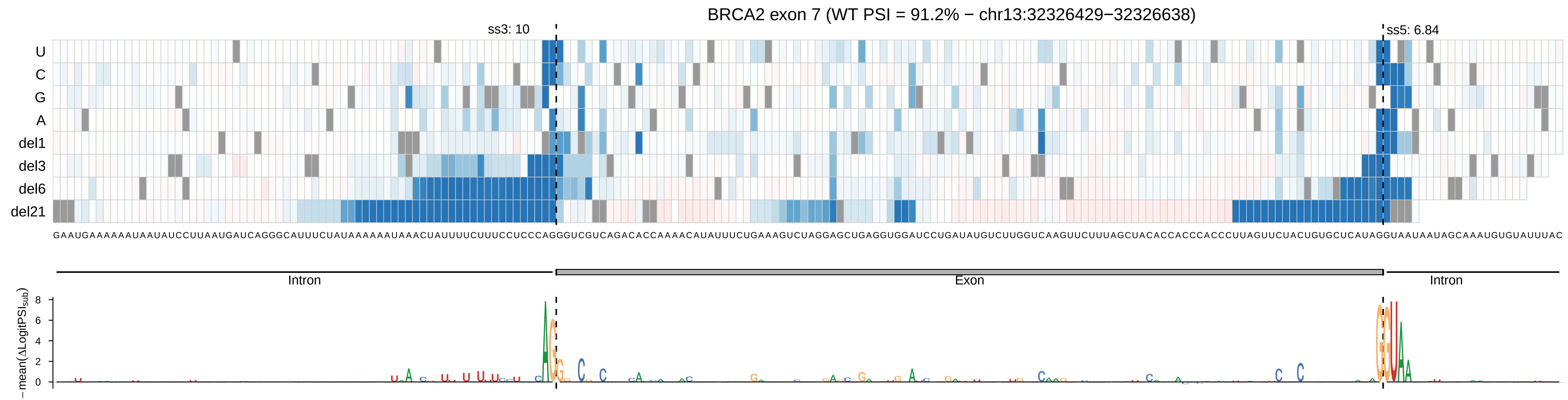

BRCA2 exon 7 (WT PSI = 91.2% − chr13:32326429−32326638)
ss3: 10
ss5: 6.84
U
C
G
A
del1
del3
del6
del21
GAAUGAAAAAAUAAUAUCCUUAAUGAUCAGGGCAUUUCUAUAAAAAAUAAACUAUUUUCUUUCCUCCCAGGGUCGUCAGACACCAAAACAUAUUUCUGAAAGUCUAGGAGCUGAGGUGGAUCCUGAUAUGUCUUGGUCAAGUUCUUUAGCUACACCACCCACCCUUAGUUCUACUGUGCUCAUAGGUAAUAAUAGCAAAUGUGUAUUUAC
Intron
Exon
Intron
− mean(ΔLogitPSIsub)
8
6
4
2
0

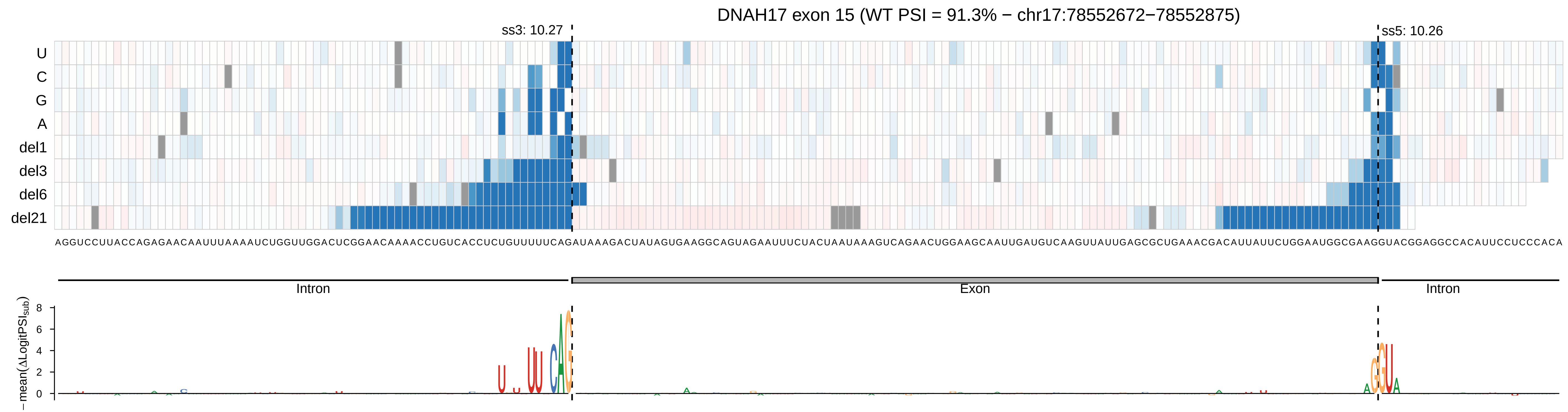

DNAH17 exon 15 (WT PSI = 91.3% − chr17:78552672−78552875)
ss3: 10.27
ss5: 10.26
U
C
G
A
del1
del3
del6
del21
AGGUCCUUACCAGAGAACAAUUUAAAAUCUGGUUGGACUCGGAACAAAACCUGUCACCUCUGUUUUUCAGAUAAAGACUAUAGUGAAGGCAGUAGAAUUUCUACUAAUAAAGUCAGAACUGGAAGCAAUUGAUGUCAAGUUAUUGAGCGCUGAAACGACAUUAUUCUGGAAUGGCGAAGGUACGGAGGCCACAUUCCUCCCACA
Intron
Exon
Intron
− mean(ΔLogitPSI_sub)
0
2
4
6
8

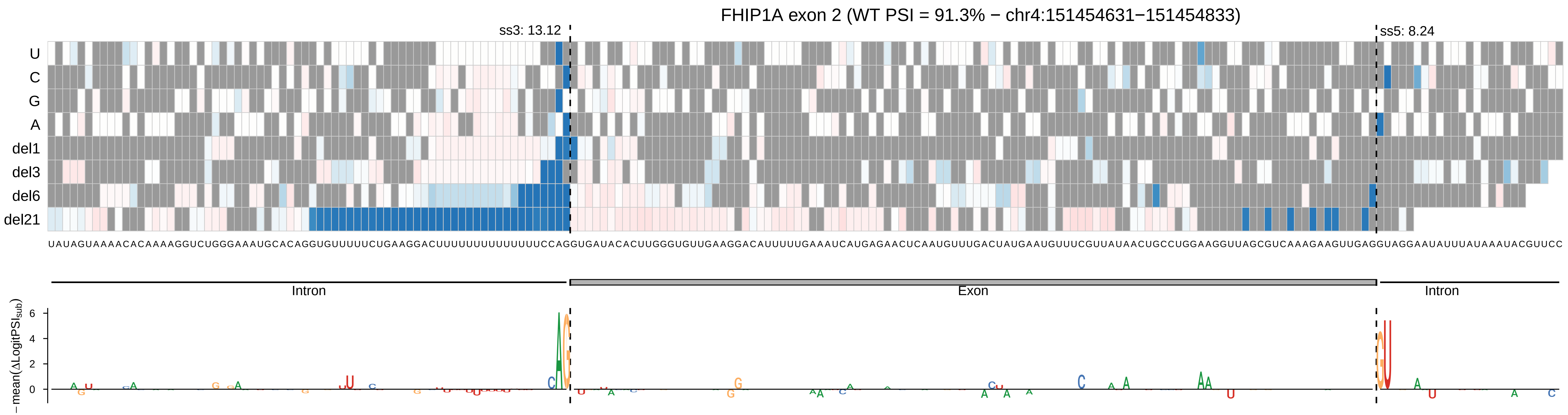

FHIP1A exon 2 (WT PSI = 91.3% − chr4:151454631−151454833)
ss3: 13.12
ss5: 8.24
U
C
G
A
del1
del3
del6
del21
UAUAGUAAAACACAAAAGGUCUGGGAAAUGCACAGGUGUUUUUCUGAAGGACUUUUUUUUUUUUUUCCAGGUGAUACACUUGGGUGUUGAAGGACAUUUUUGAAAUCAUGAGAACUCAAUGUUUGACUAUGAAUGUUUCGUUAUAACUGCCUGGAAGGUUAGCGUCAAAGAAGUUGAGGUAGGAAUAUUUAUAAAUACGUUCC
Intron
Exon
Intron
−mean(ΔLogitPSI_sub)
6
4
2
0

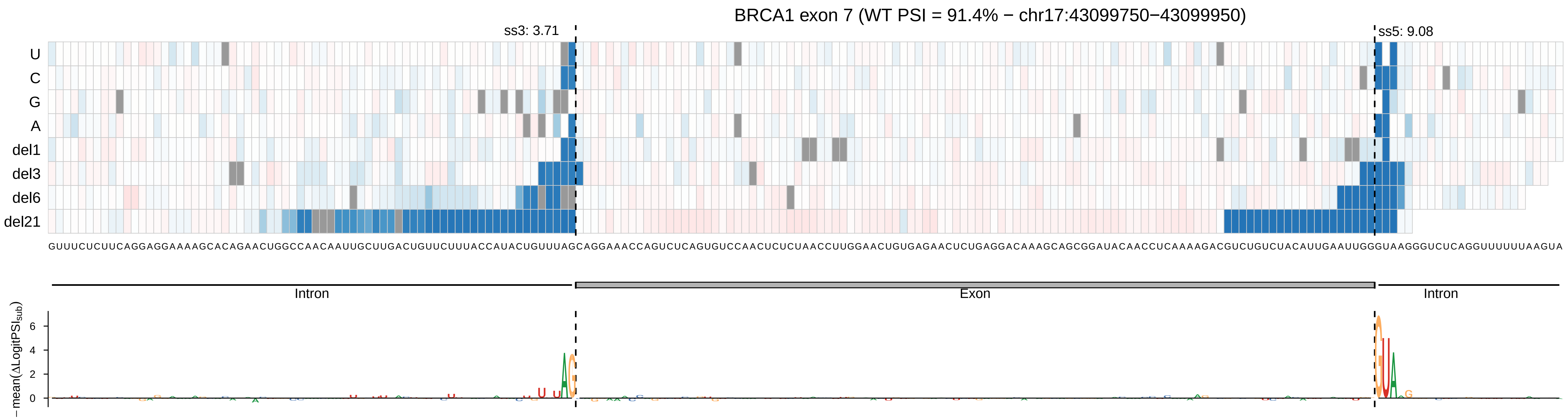
BRCA1 exon 7 (WT PSI = 91.4% − chr17:43099750−43099950)
ss3: 3.71
ss5: 9.08
U
C
G
A
del1
del3
del6
del21
GUUUCUCUUCAGGAGGAAAAGCACAGAACUGGCCAACAAUUGCUUGACUGUUCUUUACCAUACUGUUUAGCAGGAAACCAGUCUCAGUGUCCAACUCUCUAACCUUGGAACUGUGAGAACUCUGAGGACAAAGCAGCGGAUACAACCUCAAAAGACGUCUGUCUACAUUGAAUUGGGUAAGGGUCUCAGGUUUUUUAAGUA
Intron
Exon
Intron
− mean(ΔLogitPSI_sub)
6
4
2
0

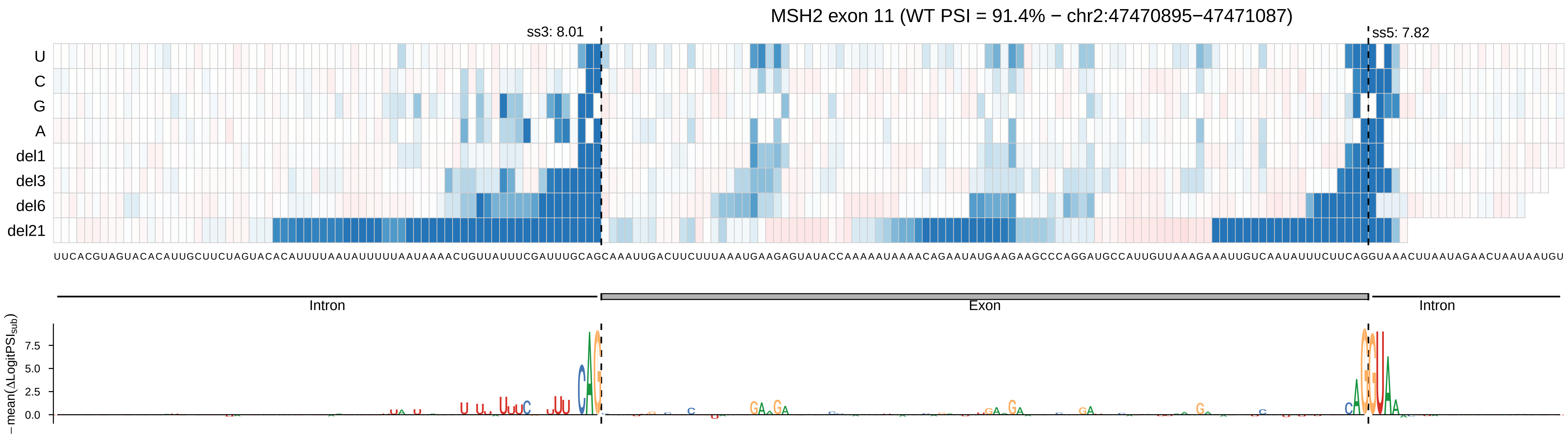

MSH2 exon 11 (WT PSI = 91.4% − chr2:47470895−47471087)
ss3: 8.01
ss5: 7.82
U
C
G
A
del1
del3
del6
del21
UUCACGUAGUACACAUUGCUUCUAGUACACAUUUUAAUAUUUUUAAUAAAACUGUUAUUUCGAUUUGCAGCAAAUUGACUUCUUUAAAUGAAGAGUAUACCAAAAAUAAAACAGAAUAUGAAGAAGCCCAGGAUGCCAUUGUUAAAGAAAUUGUCAAUAUUUCUUCAGGUAAACUUAAUAGAACUAAUAAUGU
Intron
Exon
Intron
−mean(ΔLogitPSI_sub)
7.5
5.0
2.5
0.0

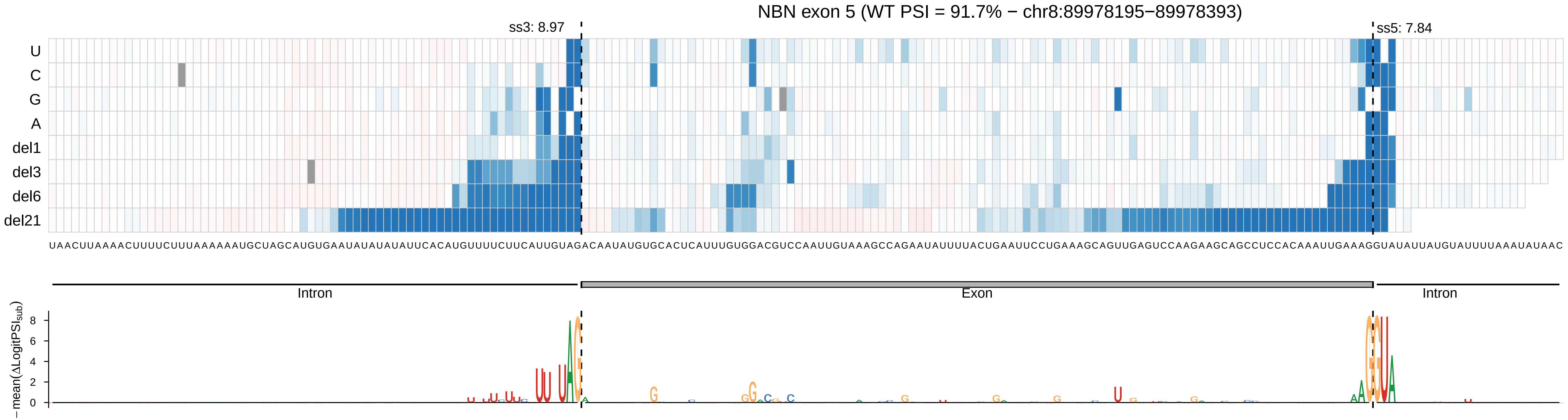
NBN exon 5 (WT PSI = 91.7% − chr8:89978195−89978393)
ss3: 8.97
ss5: 7.84
U
C
G
A
del1
del3
del6
del21
UAACUUAAAACUUUUCUUUAAAAAAUGCUAGCAUGUGAAUAUAUAUAUUCACAUGUUUUCUUCAUUGUAGACAAUAUGUGCACUCAUUUGUGGACGUCCAAUUGUAAAGCCAGAAUAUUUUACUGAAUUCCUGAAAGCAGUUGAGUCCAAGAAGCAGCCUCCACAAAUUGAAAGGUAUAUUAUGUAUUUUAAAUAUAAC
Intron
Exon
Intron
− mean(ΔLogitPSI_sub)
8
6
4
2
0

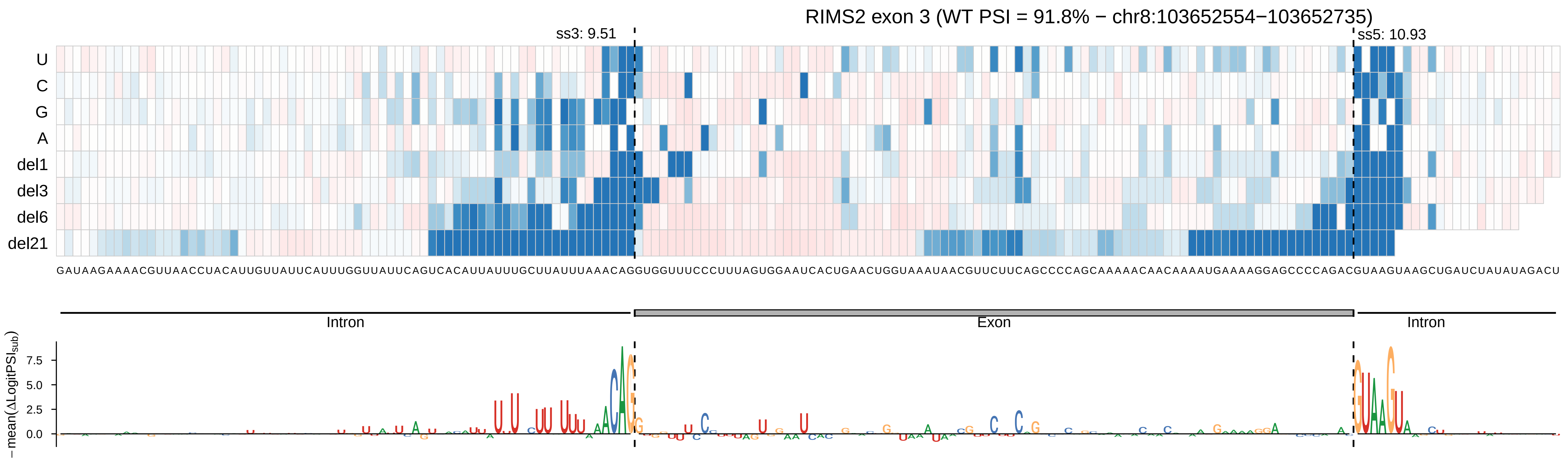
RIMS2 exon 3 (WT PSI = 91.8% − chr8:103652554−103652735)
ss3: 9.51
ss5: 10.93
U
C
G
A
del1
del3
del6
del21
Intron
Exon
Intron
− mean(ΔLogitPSI_sub)
7.5
5.0
2.5
0.0

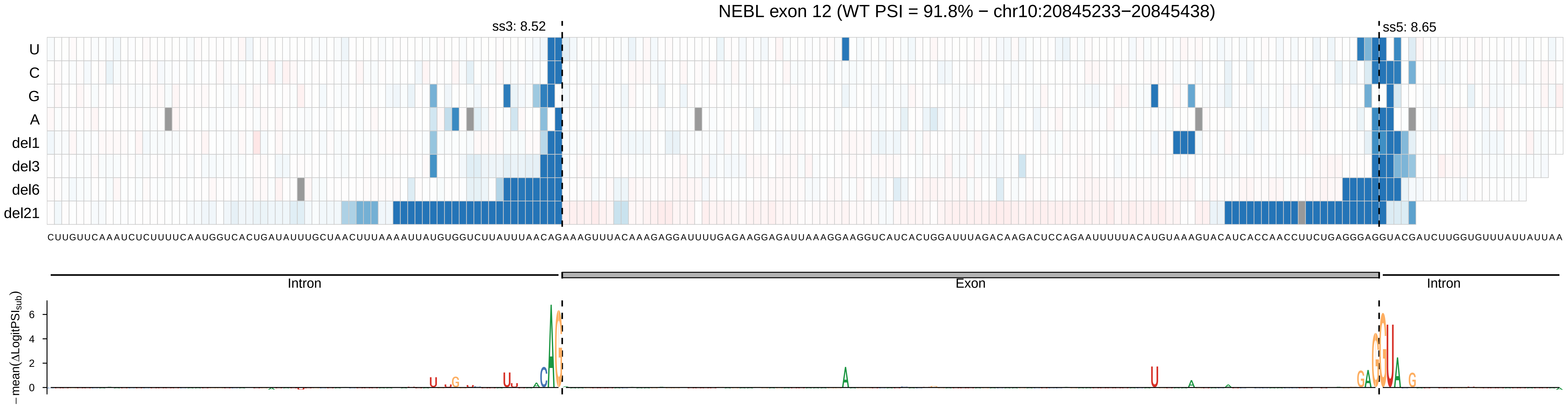

NEBL exon 12 (WT PSI = 91.8% − chr10:20845233−20845438)
ss3: 8.52
ss5: 8.65
U
C
G
A
del1
del3
del6
del21
CUUGUUCAAAUCUCUUUUCAAUGGUCACUGAUAUUUGCUAACUUUAAAAUUAUGUGGUCUUAUUUAACAGAAAGUUUACAAAGAGGAUUUUGAGAAGGAGAUUAAAGGAAGGUCAUCACUGGAUUUAGACAAGACUCCAGAAUUUUUACAUGUAAAGUACAUCACCAACCUUCUGAGGGAGGUACGAUCUUGGUGUUUAUUAUUAA
Intron
Exon
Intron
−mean(ΔLogitPSI_sub)
6
4
2
0

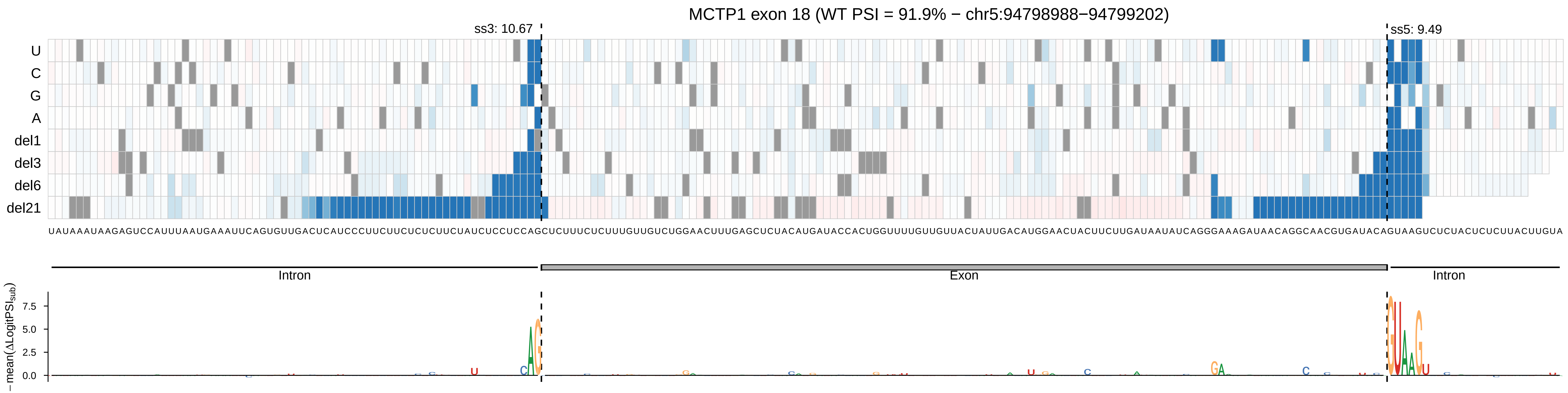
MCTP1 exon 18 (WT PSI = 91.9% − chr5:94798988−94799202)
ss3: 10.67
ss5: 9.49
U
C
G
A
del1
del3
del6
del21
UAUAAAUAAGAGUCCAUUUAAUGAAAUUCAGUGUUGACUCAUCCCUUCUUCUCUCUUCUAUCUCCUCCAGCUCUUUCUCUUUGUUGUCUGGAACUUUGAGCUCUACAUGAUACCACUGGUUUUGUUGUUACUAUUGACAUGGAACUACUUCUUGAUAAUAUCAGGGAAAGAUAACAGGCAACGUGAUACAGUAAGUCUCUACUCUCUUACUUGUA
Intron
Exon
Intron
− mean(ΔLogitPSI_sub)
7.5
5.0
2.5
0.0

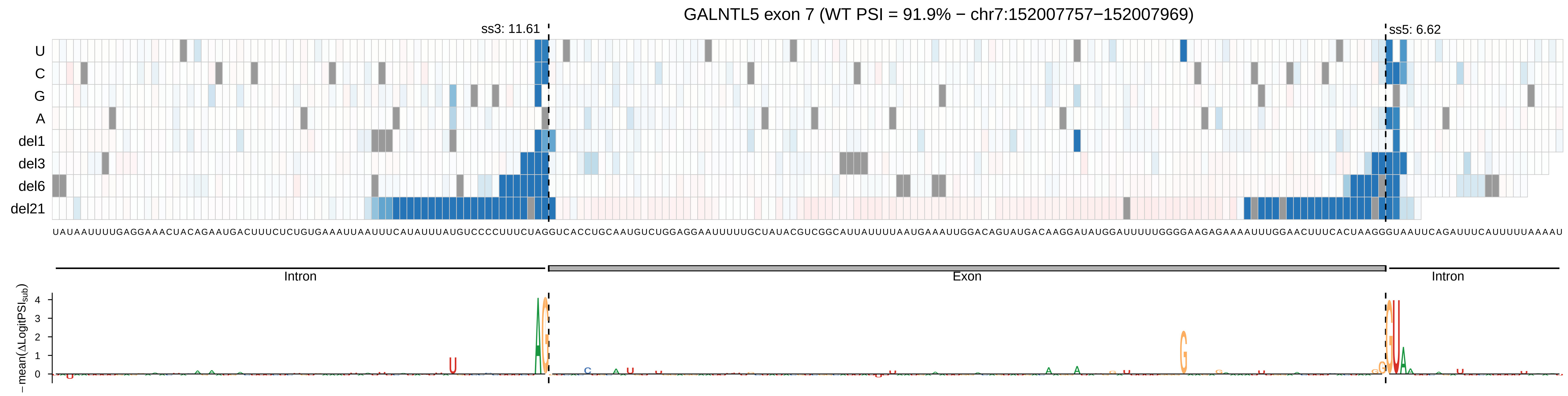
GALNTL5 exon 7 (WT PSI = 91.9% − chr7:152007757−152007969)
ss3: 11.61
ss5: 6.62
U
C
G
A
del1
del3
del6
del21
UAUAAUUUUGAGGAAACUACAGAAUGACUUUCUCUGUGAAAUUAAUUUCAUAUUUAUGUCCCCUUUCUAGGUCACCUGCAAUGUCUGGAGGAAUUUUUGCUAUACGUCGGCAUUAUUUUAAUGAAAUUGGACAGUAUGACAAGGAUAUGGAUUUUUGGGGAAGAGAAAAUUUGGAACUUUCACUAAGGGUAAUUCAGAUUUCAUUUUUAAAAU
Intron
Exon
Intron
− mean(ΔLogitPSI_sub)
4
3
2
1
0

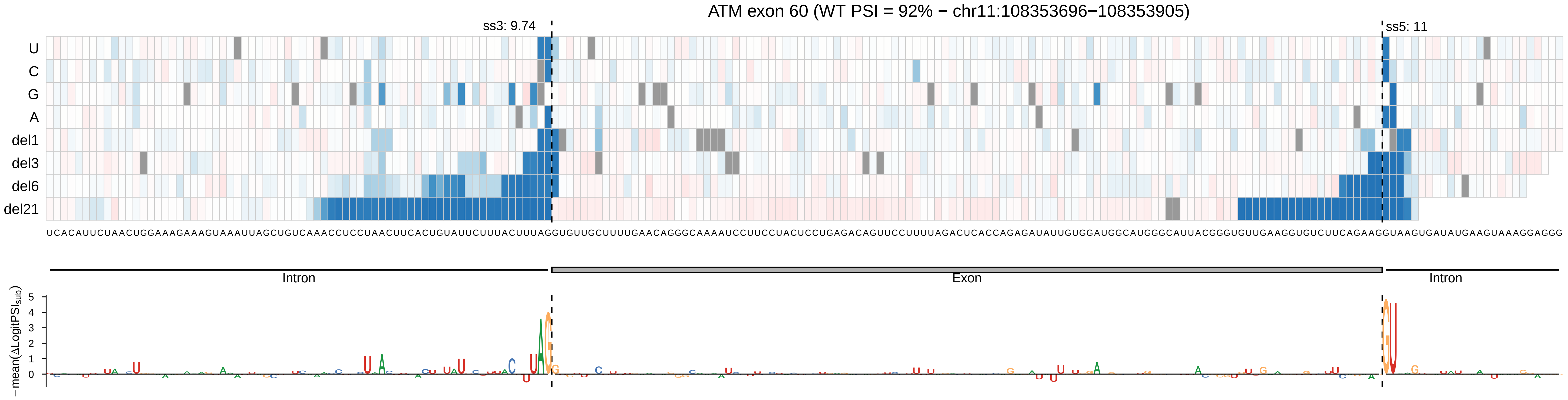

ATM exon 60 (WT PSI = 92% − chr11:108353696−108353905)
ss3: 9.74
ss5: 11
U
C
G
A
del1
del3
del6
del21
Intron
Exon
Intron
−mean(ΔLogitPSI_sub)

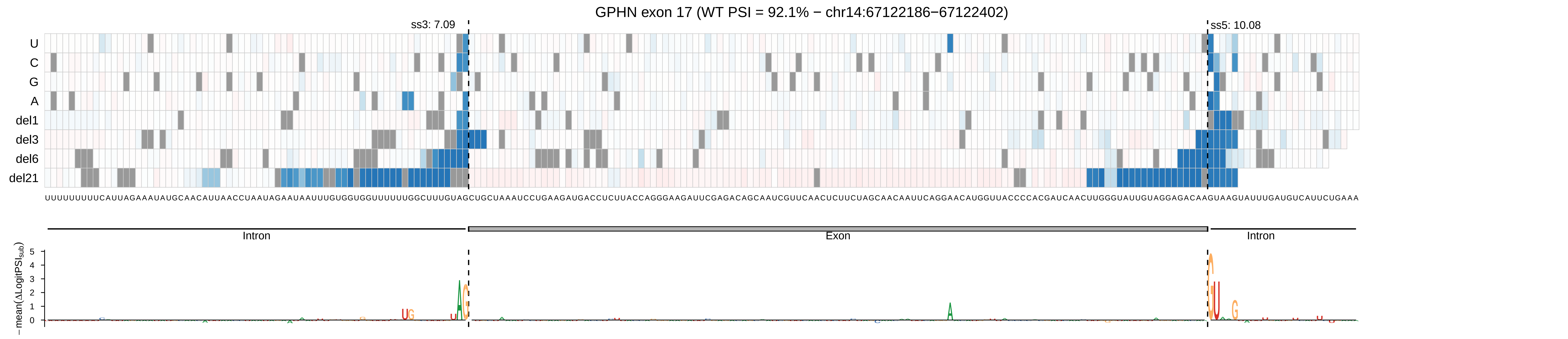
GPHN exon 17 (WT PSI = 92.1% − chr14:67122186−67122402)
ss3: 7.09
ss5: 10.08
U
C
G
A
del1
del3
del6
del21
Intron
Exon
Intron
−mean(ΔLogitPSI_sub)
5
4
3
2
1
0

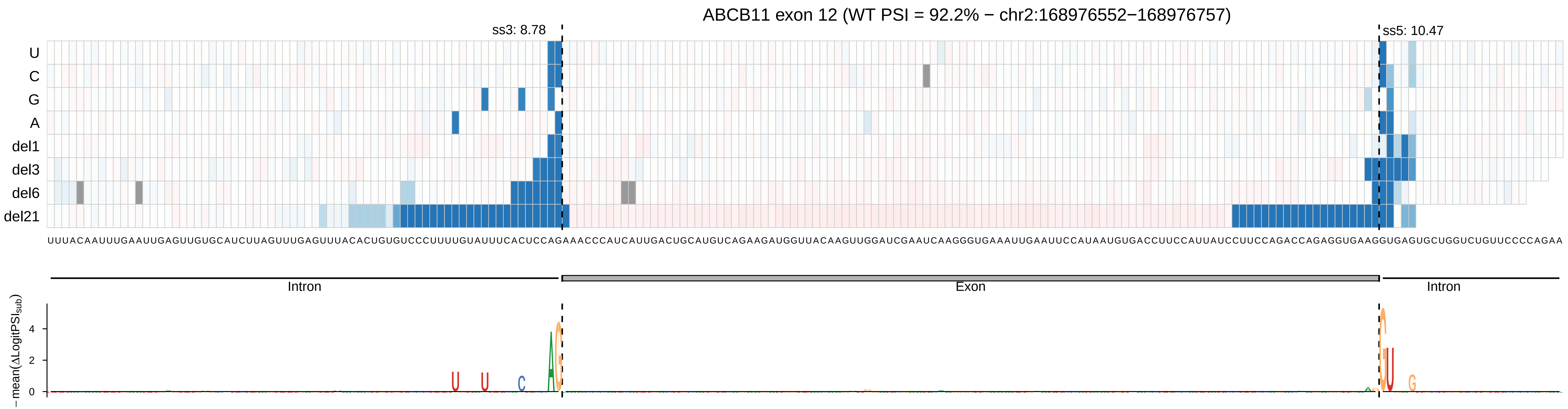

ABCB11 exon 12 (WT PSI = 92.2% − chr2:168976552−168976757)
ss3: 8.78
ss5: 10.47
U
C
G
A
del1
del3
del6
del21
UUUACAAUUUGAAUUGAGUUGUGCAUCUUAGUUUGAGUUUACACUGUGUCCCUUUUGUAUUUCACUCCAGAAACCCAUCAUUGACUGCAUGUCAGAAGAUGGUUACAAGUUGGAUCGAAUCAAGGGUGAAAUUGAAUUCCAUAAUGUGACCUUCCAUUAUCCUUCCAGACCAGAGGUGAAGGUGAGUGCUGGUCUGUUCCCCAGAA
Intron
Exon
Intron
−mean(ΔLogitPSI_sub)
4
2
0

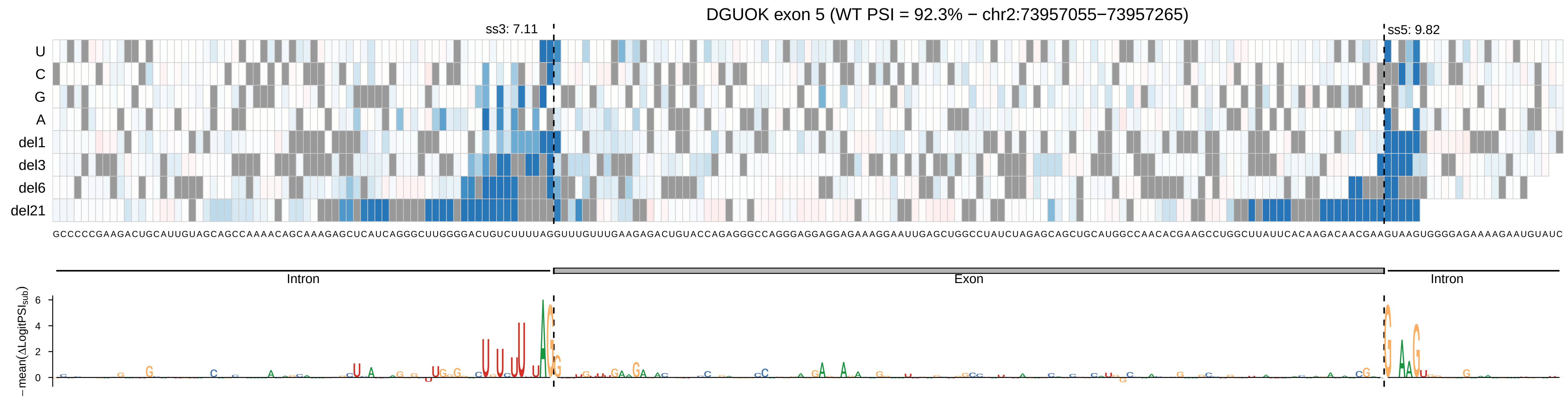

DGUOK exon 5 (WT PSI = 92.3% − chr2:73957055−73957265)
ss3: 7.11
ss5: 9.82
U
C
G
A
del1
del3
del6
del21
Intron
Exon
Intron
−mean(ΔLogitPSI_sub)

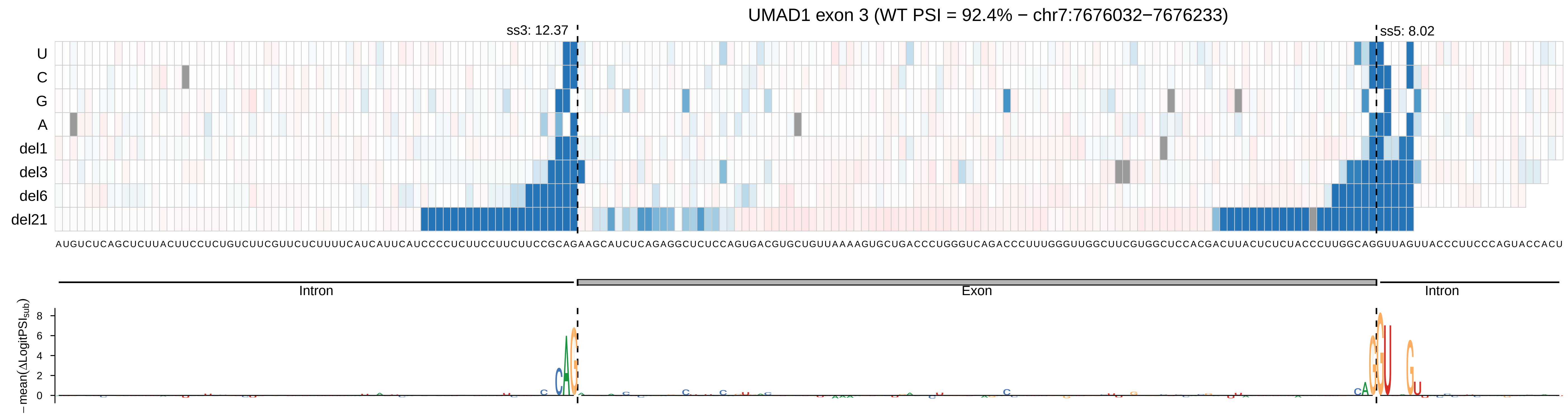

UMAD1 exon 3 (WT PSI = 92.4% − chr7:7676032−7676233)
ss3: 12.37
ss5: 8.02
U
C
G
A
del1
del3
del6
del21
Intron
Exon
Intron
− mean(ΔLogitPSI_sub)
8
6
4
2
0

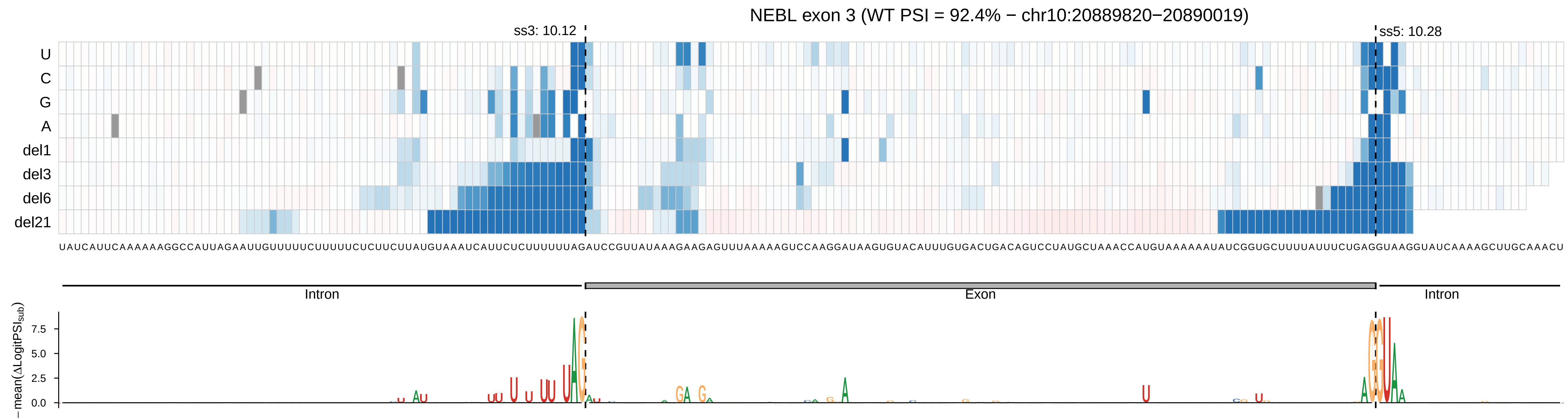
NEBL exon 3 (WT PSI = 92.4% − chr10:20889820−20890019)
ss3: 10.12
ss5: 10.28
U
C
G
A
del1
del3
del6
del21
Intron
Exon
Intron
− mean(ΔLogitPSI_sub)
7.5
5.0
2.5
0.0

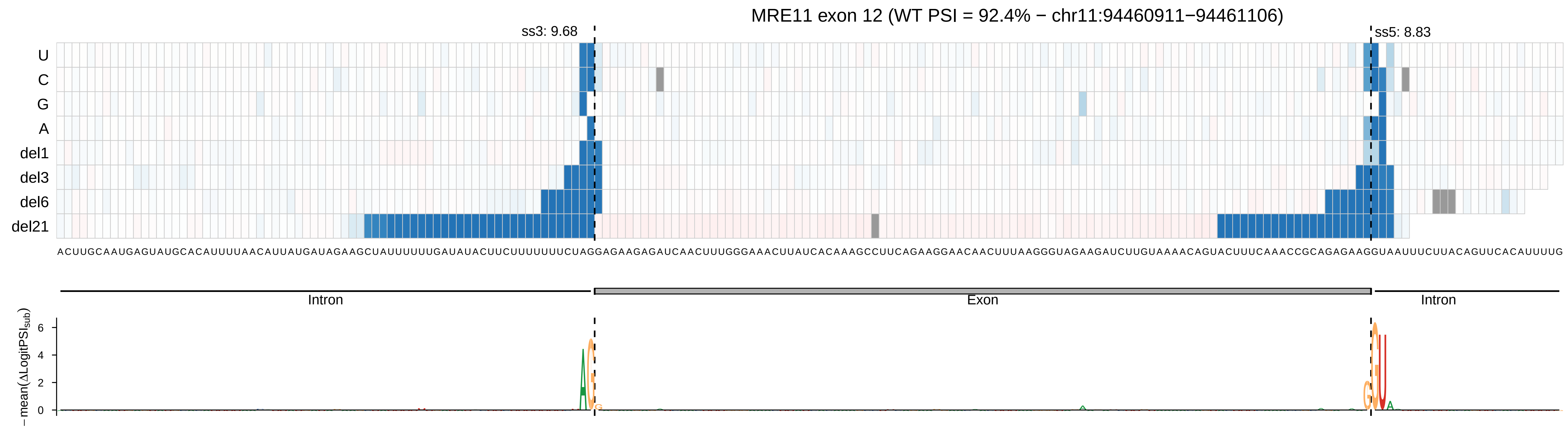
MRE11 exon 12 (WT PSI = 92.4% − chr11:94460911−94461106)
ss3: 9.68
ss5: 8.83
U
C
G
A
del1
del3
del6
del21
ACUUGCAAUGAGUAUGCACAUUUUAACAUUAUGAUAGAAGCUAUUUUUUGAUAUACUUCUUUUUUUCUAGGAGAAGAGAUCAACUUUGGGAAACUUAUCACAAAGCCUUCAGAAGGAACAACUUUAAGGGUAGAAGAUCUUGUAAAACAGUACUUUCAAACCGCAGAGAAGGUAAUUUCUUACAGUUCACAUUUUG
Intron
Exon
Intron
−mean(ΔLogitPSI_sub)
6
4
2
0

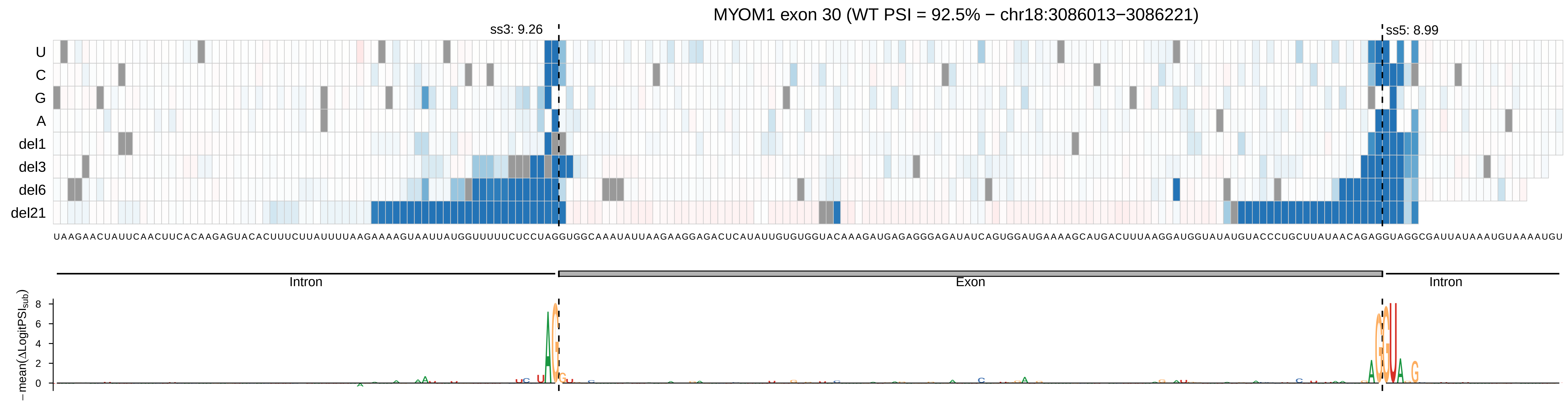
MYOM1 exon 30 (WT PSI = 92.5% − chr18:3086013−3086221)
ss3: 9.26
ss5: 8.99
U
C
G
A
del1
del3
del6
del21
UAAGAACUAUUCAACUUCACAAGAGUACACUUUCUUAUUUUAAGAAAAGUAAUUAUGGUUUUUCUCCUAGGUGGCAAAUAUUAAGAAGGAGACUCAUAUUGUGUGGUACAAAGAUGAGAGGGAGAUAUCAGUGGAUGAAAAGCAUGACUUUAAGGAUGGUAUAUGUACCCUGCUUAUAACAGAGGUAGGCGAUUAUAAAUGUAAAAUGU
Intron
Exon
Intron
− mean(ΔLogitPSI_sub)
8
6
4
2
0

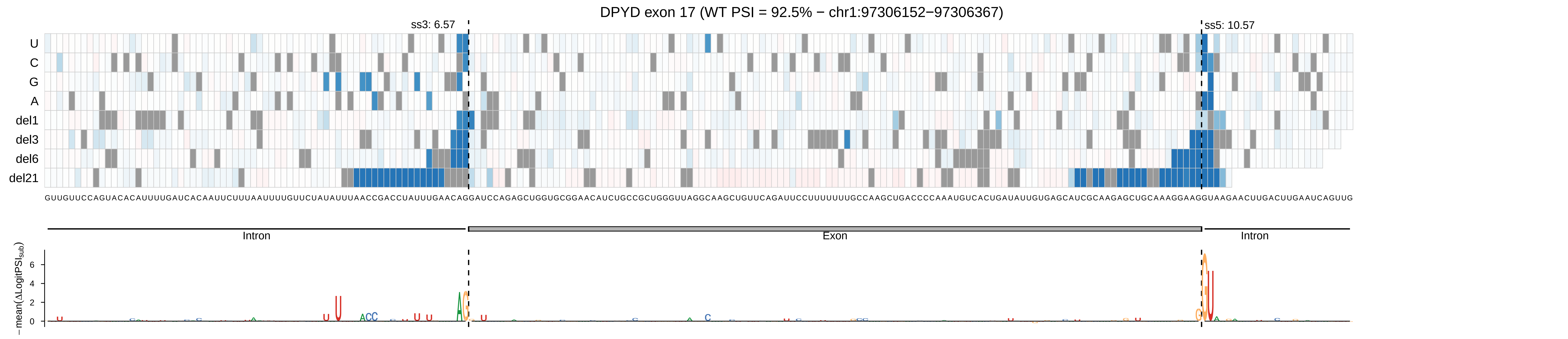
DPYD exon 17 (WT PSI = 92.5% − chr1:97306152−97306367)
ss3: 6.57
ss5: 10.57
U
C
G
A
del1
del3
del6
del21
GUUGUUCCAGUACACAUUUUGAUCACAAUUCUUUAAUUUUGUUCUAUAUUUAACCGACCUAUUUGAACAGGAUCCAGAGCUGGUGCGGAACAUCUGCCGCUGGGUUAGGCAAGCUGUUCAGAUUCCUUUUUUUGCCAAGCUGACCCCAAAUGUCACUGAUAUUGUGAGCAUCGCAAGAGCUGCAAAGGAAGGUAAGAACUUGACUUGAAUCAGUUG
Intron
Exon
Intron
−mean(ΔLogitPSIsub)
6
4
2
0

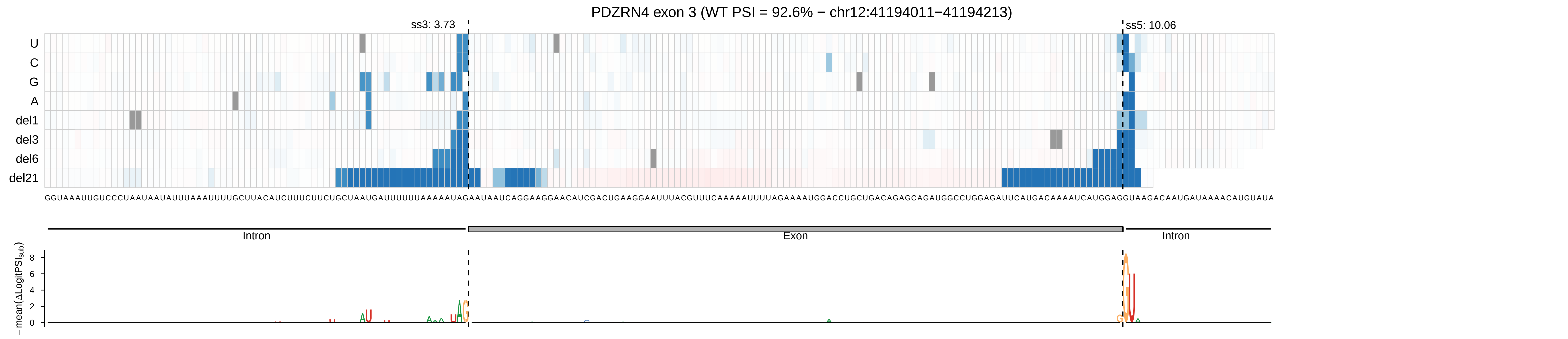

PDZRN4 exon 3 (WT PSI = 92.6% − chr12:41194011−41194213)
ss3: 3.73
ss5: 10.06
U
C
G
A
del1
del3
del6
del21
GGUAAAUUGUCCCUAAUAAUAUUUAAAUUUUGCUUACAUCUUUCUUCUGCUAAUGAUUUUUUAAAAAUAGAAUAAUCAGGAAGGAACAUCGACUGAAGGAAUUUACGUUUCAAAAAUUUUAGAAAAUGGACCUGCUGACAGAGCAGAUGGCCUGGAGAUUCAUGACAAAAUCAUGGAGGUAAGACAAUGAUAAAACAUGUAUA
Intron
Exon
Intron
− mean(ΔLogitPSI_sub)
8
6
4
2
0

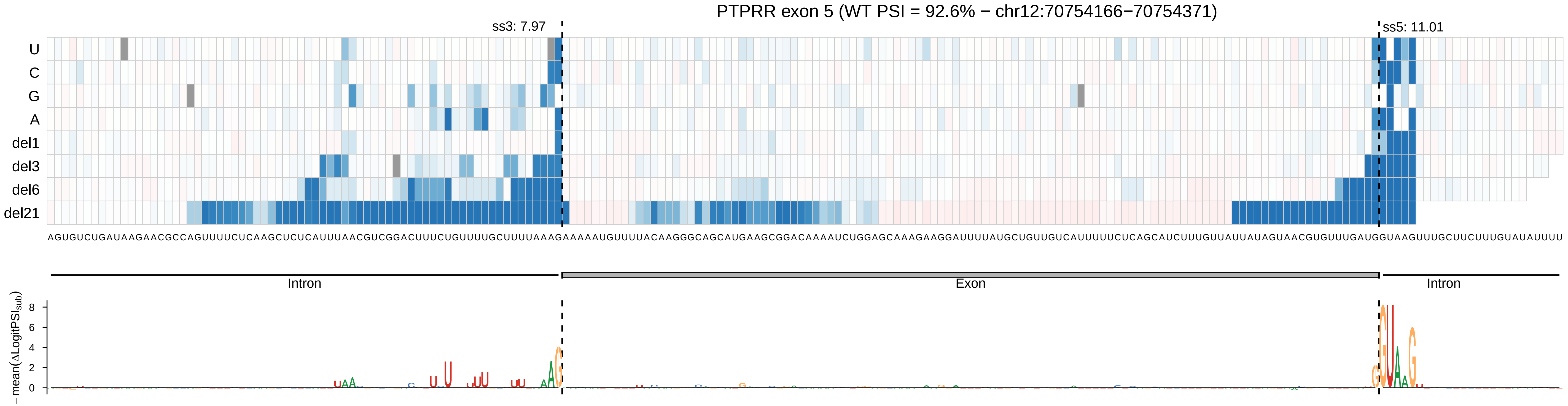
PTPRR exon 5 (WT PSI = 92.6% − chr12:70754166−70754371)
ss3: 7.97
ss5: 11.01
U
C
G
A
del1
del3
del6
del21
AGUGUCUGAUAAGAACGCCAGUUUUCUCAAGCUCUCAUUUAACGUCGGACUUUCUGUUUUGCUUUUAAAGAAAAAUGUUUUACAAGGGCAGCAUGAAGCGGACAAAAUCUGGAGCAAAGAAGGAUUUUAUGCUGUUGUCAUUUUUCUCAGCAUCUUUGUUAUUAUAGUAACGUGUUUGAUGGUAAGUUUGCUUCUUUGUAUAUUUU
Intron
Exon
Intron
−mean(ΔLogitPSI_sub)
8
6
4
2
0

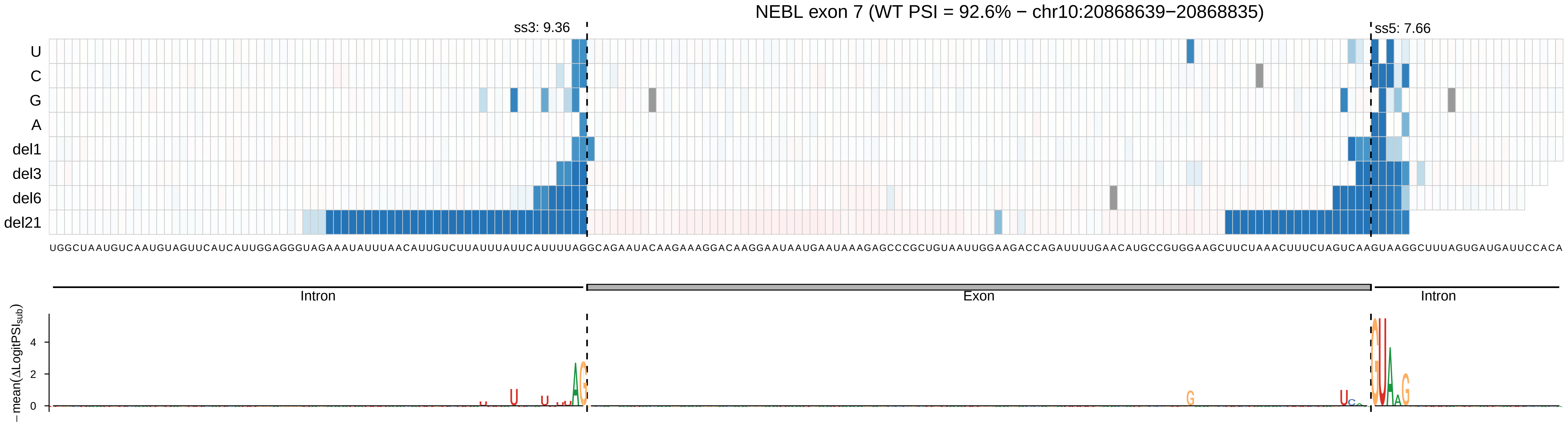
NEBL exon 7 (WT PSI = 92.6% − chr10:20868639−20868835)
ss3: 9.36
ss5: 7.66
U
C
G
A
del1
del3
del6
del21
UGGCUAAUGUCAAUGUAGUUCAUCAUUGGAGGGUAGAAAUAUUUAACAUUGUCUUAUUUAUUCAUUUUAGGCAGAAUACAAGAAAGGACAAGGAAUAAUGAAUAAAGAGCCCGCUGUAAUUGGAAGACCAGAUUUUGAACAUGCCGUGGAAGCUUCUAAACUUUCUAGUCAAGUAAGGCUUUAGUGAUGAUUCCACA
Intron
Exon
Intron
− mean(ΔLogitPSI_sub)
4
2
0

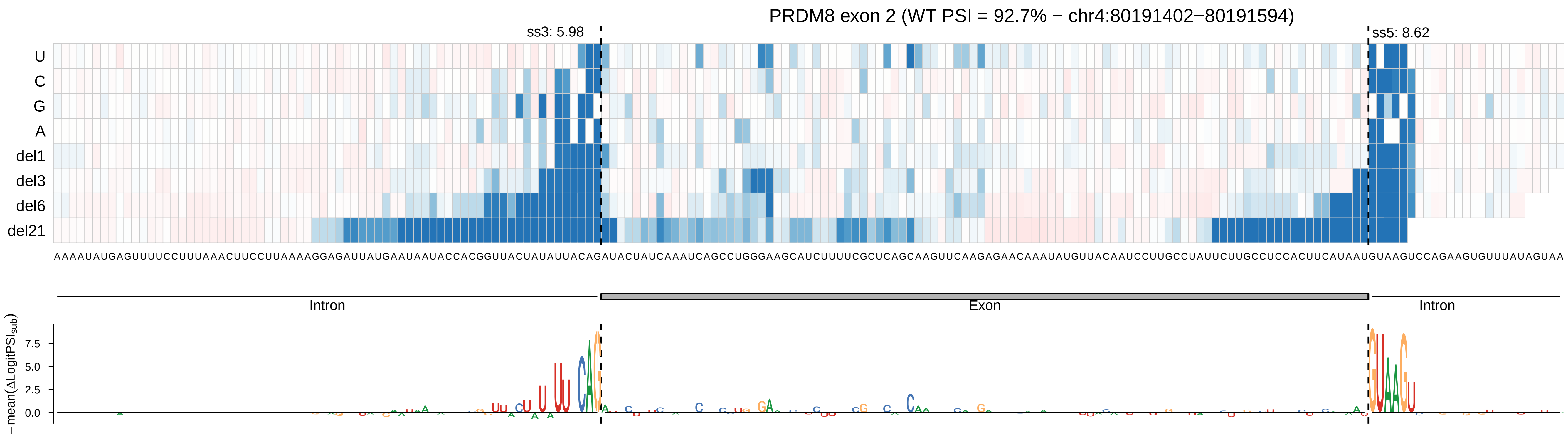

PRDM8 exon 2 (WT PSI = 92.7% − chr4:80191402−80191594)
ss3: 5.98
ss5: 8.62
U
C
G
A
del1
del3
del6
del21
AAAAUAUGAGUUUUCCUUUAAACUUCCUUAAAAGGAGAUUAUGAAUAAUACCACGGUUACUAUAUUACAGAUACUAUCAAAUCAGCCUGGGAAGCAUCUUUUCGCUCAGCAAGUUCAAGAGAACAAAUAUGUUACAAUCCUUGCCUAUUCUUGCCUCCACUUCAUAAUGUAAGUCCAGAAGUGUUUAUAGUAA
Intron
Exon
Intron
− mean(ΔLogitPSI_sub)
7.5
5.0
2.5
0.0

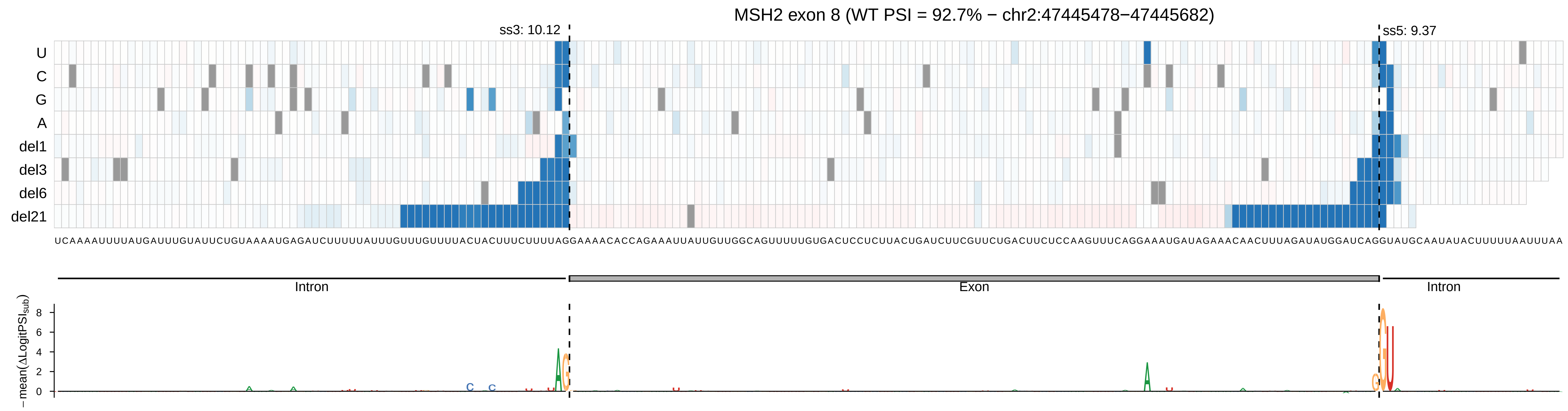

MSH2 exon 8 (WT PSI = 92.7% − chr2:47445478−47445682)
ss3: 10.12
ss5: 9.37
U
C
G
A
del1
del3
del6
del21
Intron
Exon
Intron
−mean(ΔLogitPSIsub)

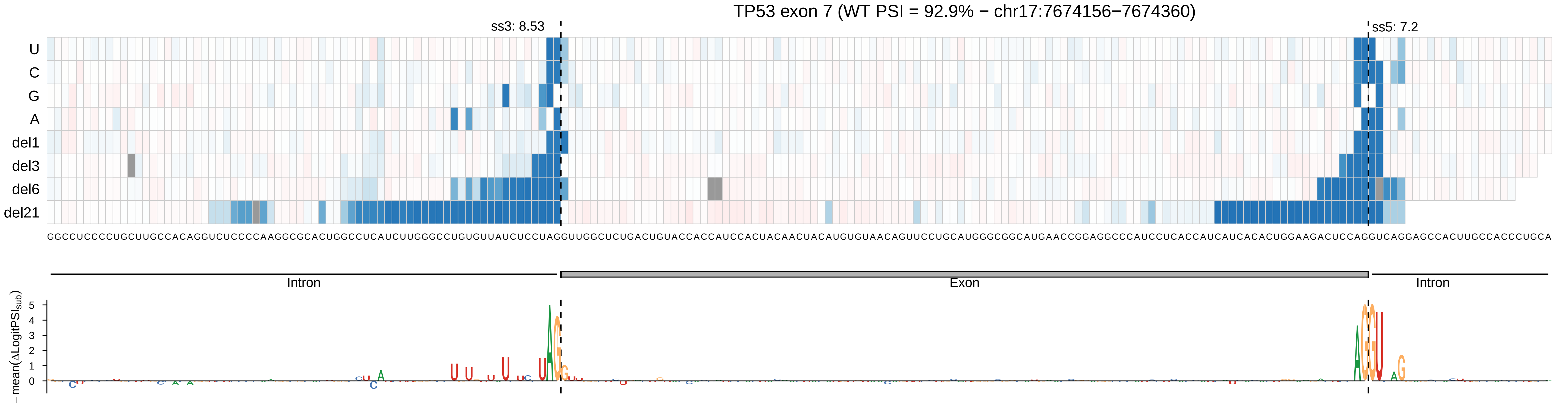

TP53 exon 7 (WT PSI = 92.9% − chr17:7674156−7674360)
ss3: 8.53
ss5: 7.2
U
C
G
A
del1
del3
del6
del21
GGCCUCCCCUGCUUGCCACAGGUCUCCCCAAGGCGCACUGGCCUCAUCUUGGGCCUGUGUUAUCUCCUAGGUUGGCUCUGACUGUACCACCAUCCACUACAACUACAUGUGUAACAGUUCCUGCAUGGGCGGCAUGAACCGGAGGCCCAUCCUCACCAUCAUCACACUGGAAGACUCCAGGUCAGGAGCCACUUGCCACCCUGCA
Intron
Exon
Intron
−mean(ΔLogitPSI_sub)
5
4
3
2
1
0

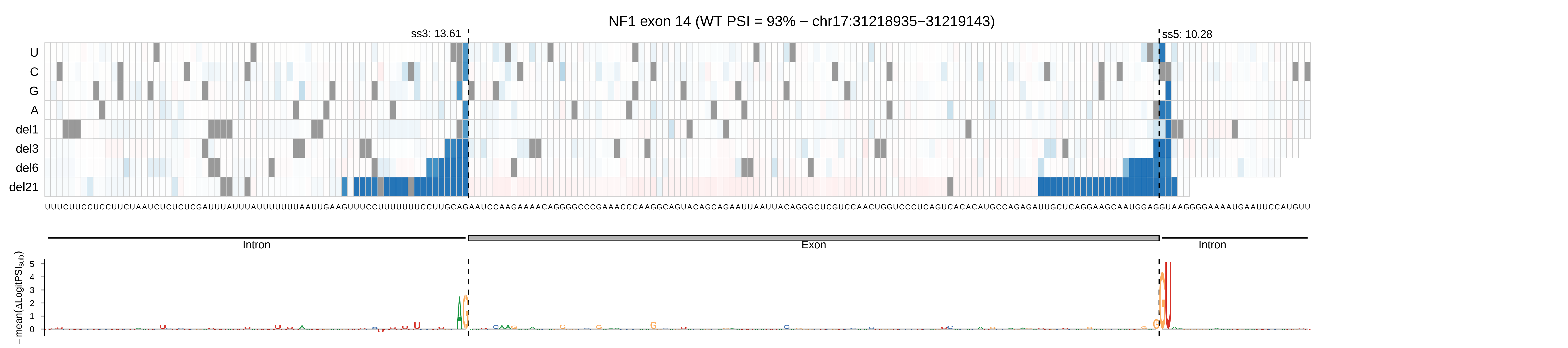
NF1 exon 14 (WT PSI = 93% − chr17:31218935−31219143)
ss3: 13.61
ss5: 10.28
U
C
G
A
del1
del3
del6
del21
Intron
Exon
Intron
−mean(ΔLogitPSI_sub)
0
1
2
3
4
5

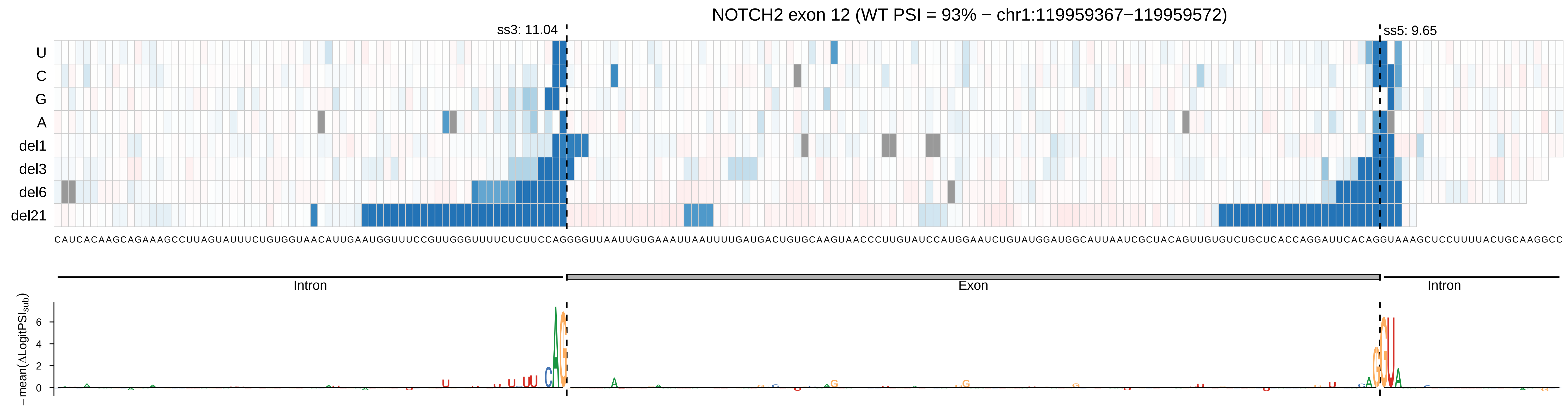

NOTCH2 exon 12 (WT PSI = 93% − chr1:119959367−119959572)
ss3: 11.04
ss5: 9.65
U
C
G
A
del1
del3
del6
del21
CAUCACAAGCAGAAAGCCUUAGUAUUUCUGUGGUAACAUUGAAUGGUUUCCGUUGGGUUUUCUCUUCCAGGGGUUAAUUGUGAAAUUAAUUUUGAUGACUGUGCAAGUAACCCUUGUAUCCAUGGAAUCUGUAUGGAUGGCAUUAAUCGCUACAGUUGUGUCUGCUCACCAGGAUUCACAGGUAAAGCUCCUUUUACUGCAAGGCC
Intron
Exon
Intron
−mean(ΔLogitPSI$_{sub}$)
0
2
4
6

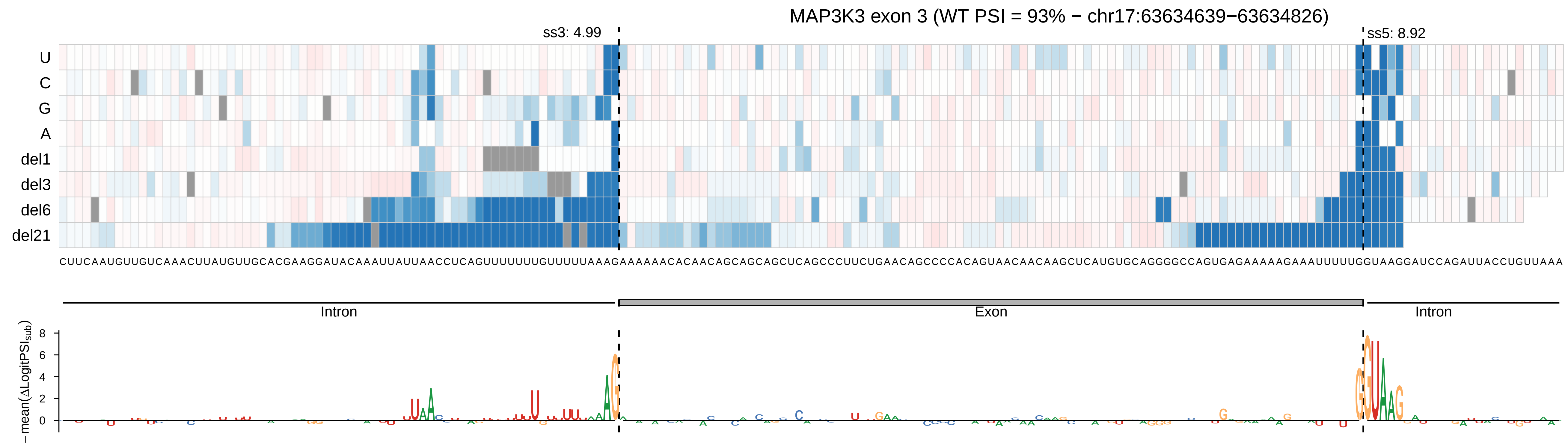

MAP3K3 exon 3 (WT PSI = 93% − chr17:63634639−63634826)
ss3: 4.99
ss5: 8.92
U
C
G
A
del1
del3
del6
del21
CUUCAAUGUUGUCAAACUUAUGUUGCACGAAGGAUACAAAUUAUUAACCUCAGUUUUUUUGUUUUUAAAGAAAAAACACAACAGCAGCAGCUCAGCCCUUCUGAACAGCCCCACAGUAACAACAAGCUCAUGUGCAGGGGCCAGUGAGAAAAAGAAAUUUUUGGUAAGGAUCCAGAUUACCUGUUAAA
Intron
Exon
Intron
−mean(ΔLogitPSIsub)

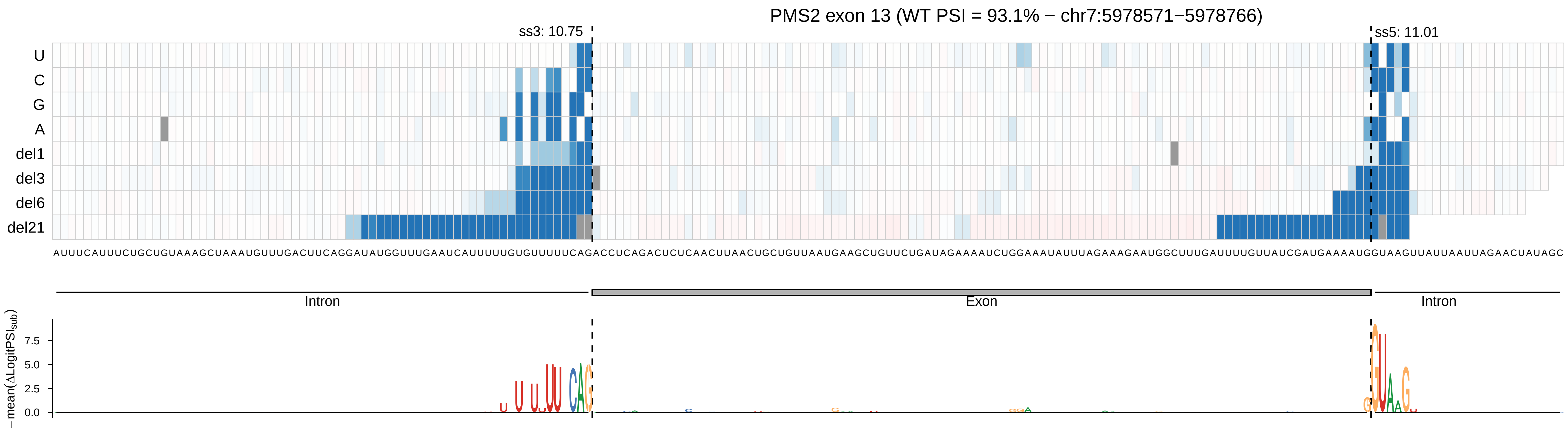
PMS2 exon 13 (WT PSI = 93.1% − chr7:5978571−5978766)
ss3: 10.75
ss5: 11.01
U
C
G
A
del1
del3
del6
del21
Intron
Exon
Intron
− mean(ΔLogitPSI_sub)
7.5
5.0
2.5
0.0

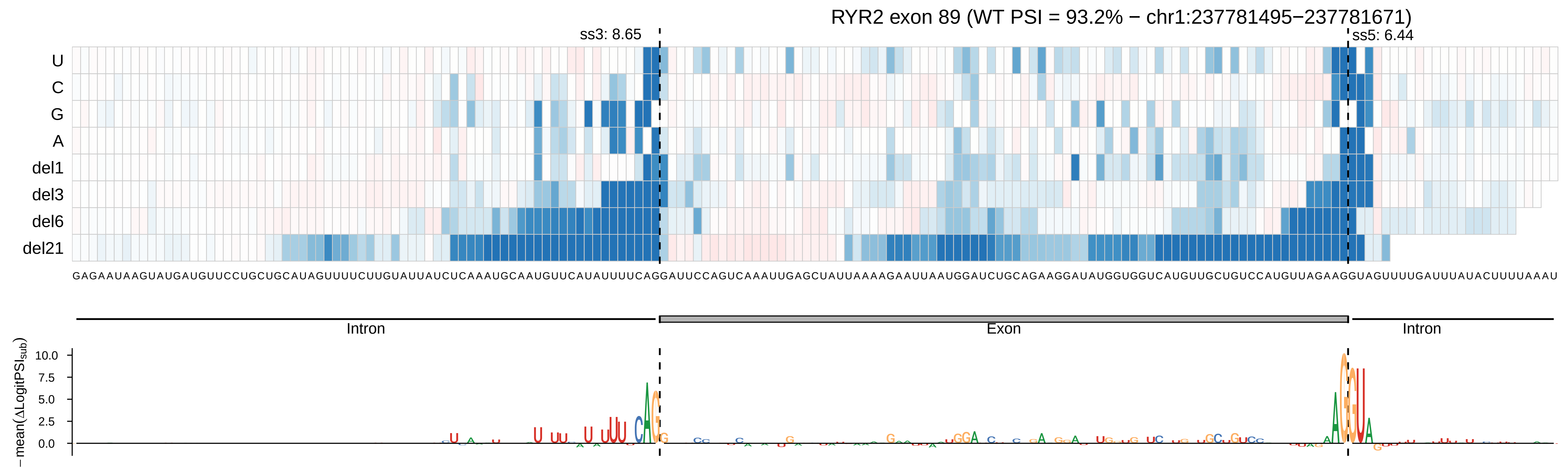
RYR2 exon 89 (WT PSI = 93.2% − chr1:237781495−237781671)
ss3: 8.65
ss5: 6.44
U
C
G
A
del1
del3
del6
del21
GAGAAUAAGUAUGAUGUUCCUGCUGCAUAGUUUUCUUGUAUUAUCUCAAAUGCAAUGUUCAUAUUUUCAGGAUUCCAGUCAAAUUGAGCUAUUAAAAGAAUUAAUGGAUCUGCAGAAGGAUAUGGUGGUCAUGUUGCUGUCCAUGUUAGAAGGUAGUUUUGAUUUAUACUUUUAAAU
Intron
Exon
Intron
− mean(ΔLogitPSI_sub)
10.0
7.5
5.0
2.5
0.0

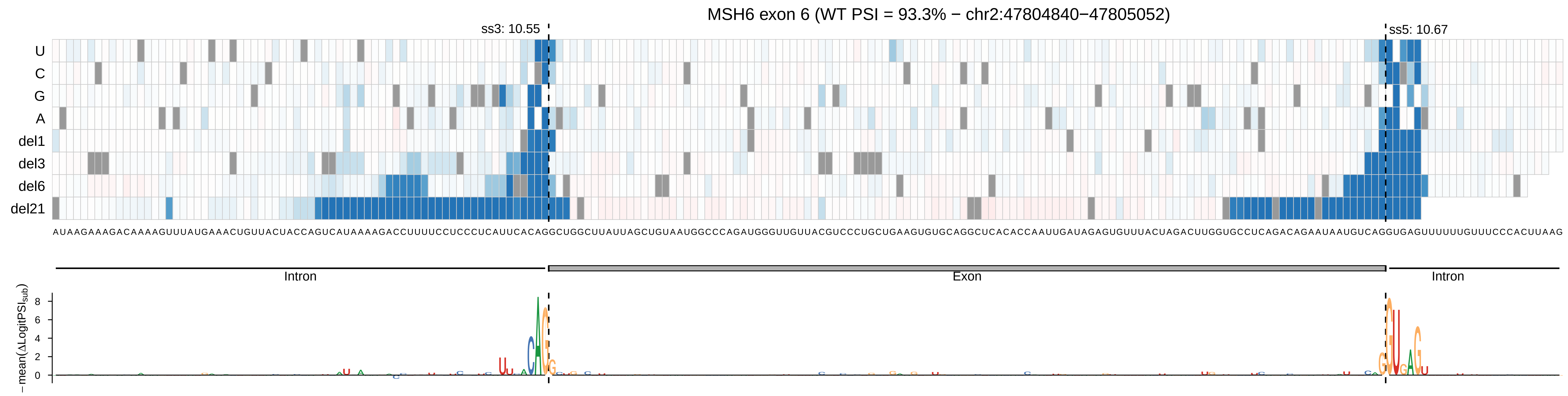
MSH6 exon 6 (WT PSI = 93.3% − chr2:47804840−47805052)
ss3: 10.55
ss5: 10.67
U
C
G
A
del1
del3
del6
del21
Intron
Exon
Intron
− mean(ΔLogitPSI_sub)
8
6
4
2
0

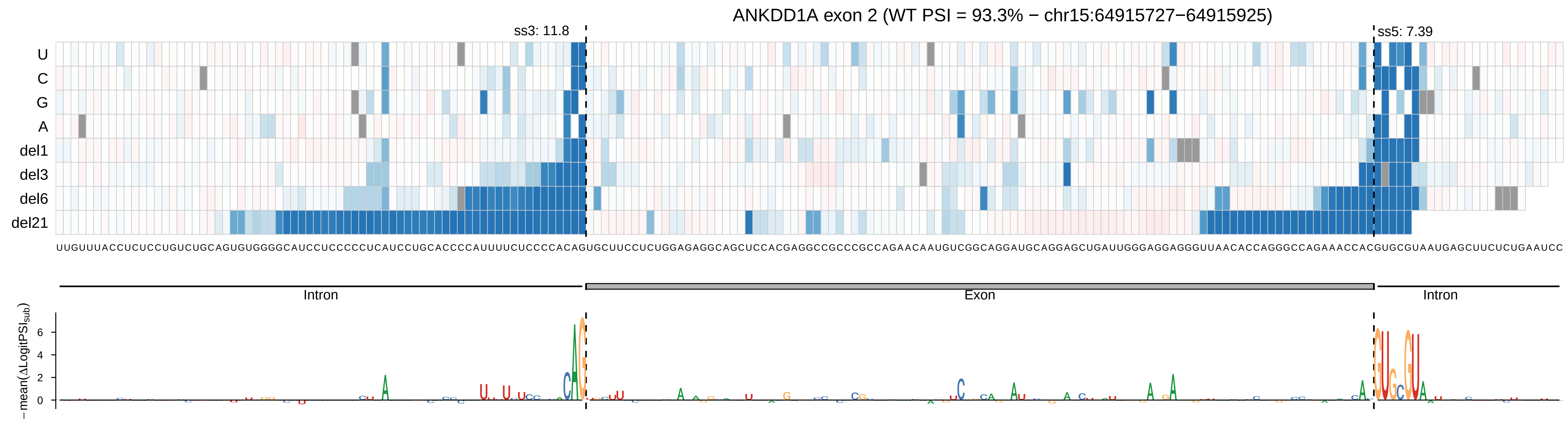

ANKDD1A exon 2 (WT PSI = 93.3% − chr15:64915727−64915925)
ss3: 11.8
ss5: 7.39
U
C
G
A
del1
del3
del6
del21
UUGUUUACCUCUCCUGUCUGCAGUGUGGGGCAUCCUCCCCCUCAUCCUGCACCCCAUUUUCUCCCCACAGUGCUUCCUCUGGAGAGGCAGCUCCACGAGGCCGCCCGCCAGAACAAUGUCGGCAGGAUGCAGGAGCUGAUUGGGAGGAGGGUUAACACCAGGGCCAGAAACCACGUGCGUAAUGAGCUUCUCUGAAUCC
Intron
Exon
Intron
− mean(ΔLogitPSI_sub)
6
4
2
0

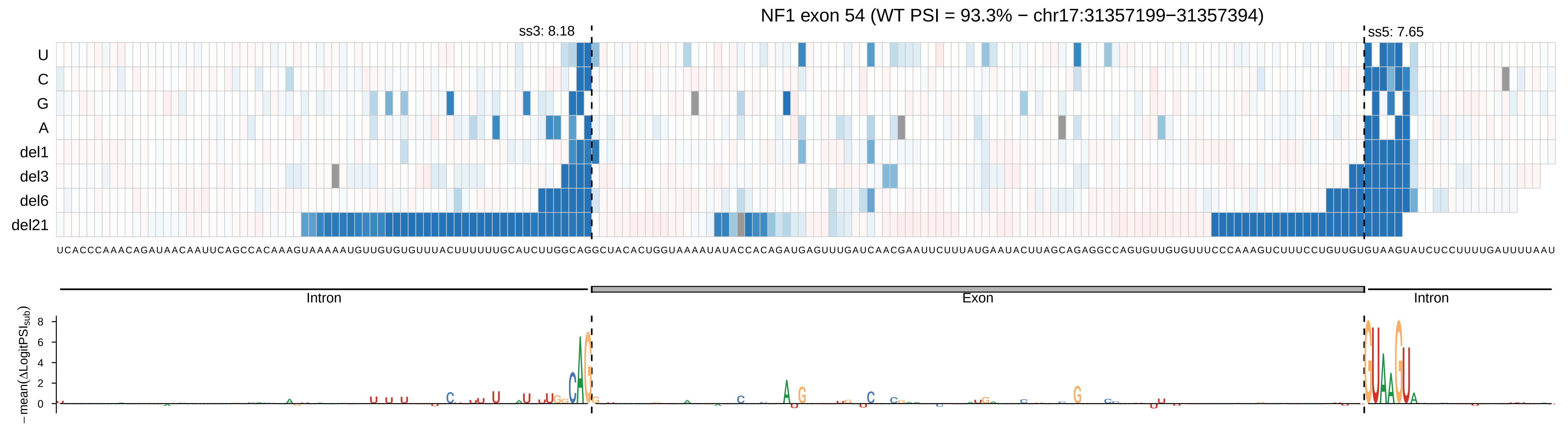
NF1 exon 54 (WT PSI = 93.3% − chr17:31357199−31357394)
ss3: 8.18
ss5: 7.65
U
C
G
A
del1
del3
del6
del21
UCACCCAAACAGAUAACAAUUCAGCCACAAAGUAAAAAUGUUGUGUGUUUACUUUUUUGCAUCUUGGCAGGCUACACUGGUAAAAUAUACCACAGAUGAGUUUGAUCAACGAAUUCUUUAUGAAUACUUAGCAGAGGCCAGUGUUGUGUUUCCCAAAGUCUUUCCUGUUGUGUAAGUAUCUCCUUUUGAUUUUAAU
Intron
Exon
Intron
− mean(ΔLogitPSI_sub)
8
6
4
2
0

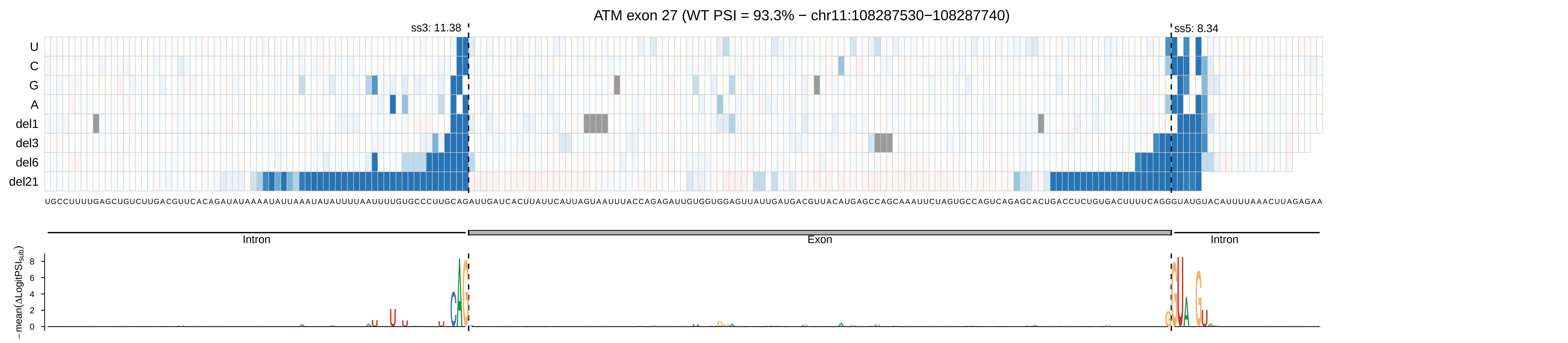
ATM exon 27 (WT PSI = 93.3% − chr11:108287530−108287740)
ss3: 11.38
ss5: 8.34
U
C
G
A
del1
del3
del6
del21
UGCCUUUUGAGCUGUCUUGACGUUCACAGAUAUAAAAUAUUAAAUAUAUUUUAAUUUUGUGCCCUUGCAGAUUGAUCACUUAUUCAUUAGUAAUUUACCAGAGAUUGUGGUGGAGUUAUUGAUGACGUUACAUGAGCCAGCAAAUUCUAGUGCCAGUCAGAGCACUGACCUCUGUGACUUUUCAGGGUAUGUACAUUUUAAACUUAGAGAA
Intron
Exon
Intron
− mean(ΔLogitPSIsub)
0
2
4
6
8

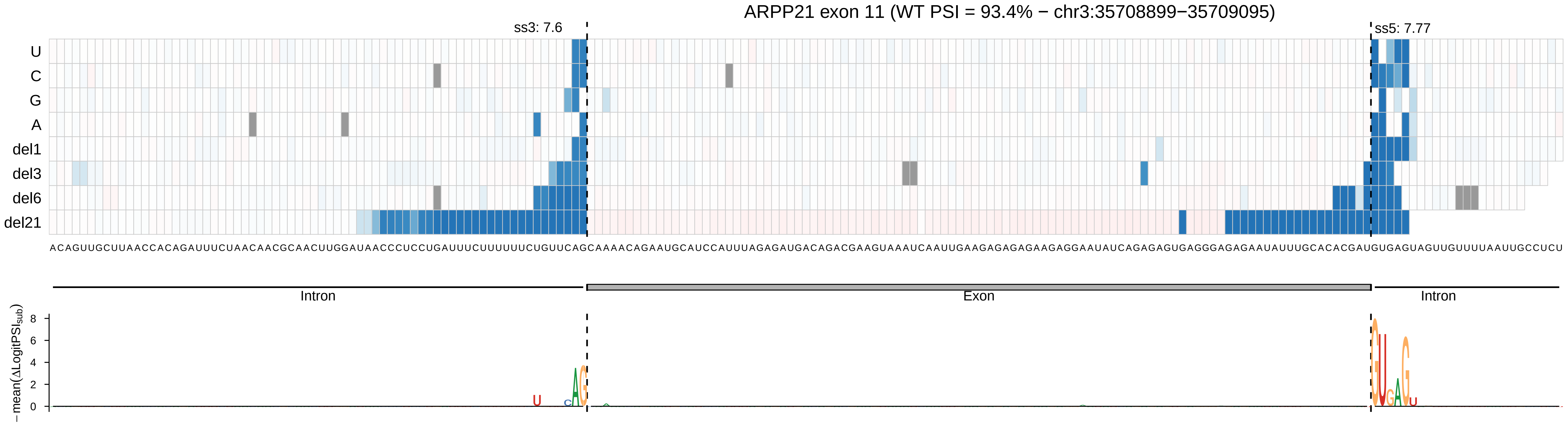

ARPP21 exon 11 (WT PSI = 93.4% − chr3:35708899−35709095)
ss3: 7.6
ss5: 7.77
U
C
G
A
del1
del3
del6
del21
Intron
Exon
Intron
−mean(ΔLogitPSI_sub)
8
6
4
2
0

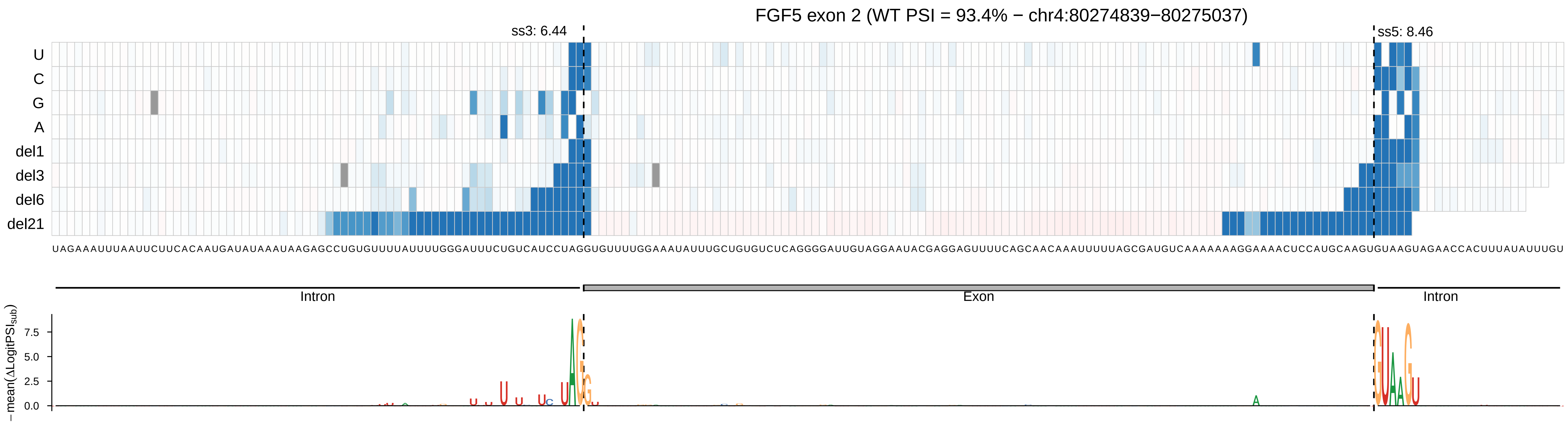
FGF5 exon 2 (WT PSI = 93.4% − chr4:80274839−80275037)
ss3: 6.44
ss5: 8.46
U
C
G
A
del1
del3
del6
del21
UAGAAAUUUAAUUCUUCACAAUGAUAUAAAUAAGAGCCUGUGUUUUAUUUUGGGAUUUCUGUCAUCCUAGGUGUUUUGGAAAUAUUUGCUGUGUCUCAGGGGAUUGUAGGAAUACGAGGAGUUUUCAGCAACAAAUUUUUAGCGAUGUCAAAAAAAGGAAAACUCCAUGCAAGUGUAAGUAGAACCACUUUAUAUUUGU
Intron
Exon
Intron
− mean(ΔLogitPSI_sub)
7.5
5.0
2.5
0.0

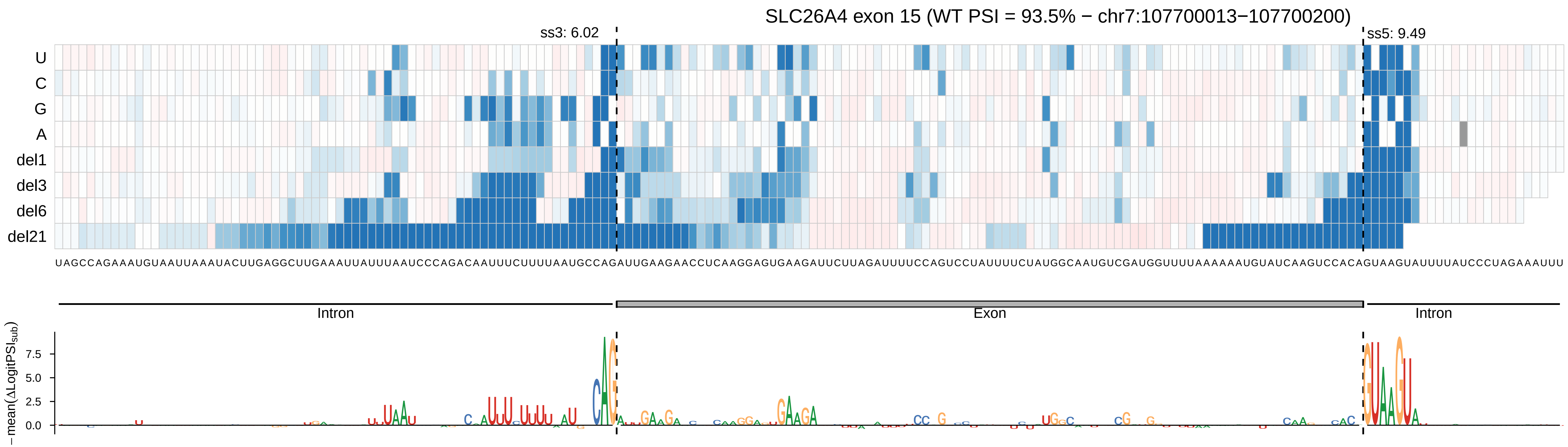
SLC26A4 exon 15 (WT PSI = 93.5% − chr7:107700013−107700200)
ss3: 6.02
ss5: 9.49
U
C
G
A
del1
del3
del6
del21
UAGCCAGAAAUGUAAUUAAAUACUUGAGGCUUGAAAUUAUUUAAUCCCAGACAAUUUCUUUUAAUGCCAGAUUGAAGAACCUCAAGGAGUGAAGAUUCUUAGAUUUUCCAGUCCUAUUUUCUAUGGCAAUGUCGAUGGUUUUAAAAAAUGUAUCAAGUCCACAGUAAGUAUUUUAUCCCUAGAAAUUU
Intron
Exon
Intron
− mean(ΔLogitPSI_sub)
7.5
5.0
2.5
0.0

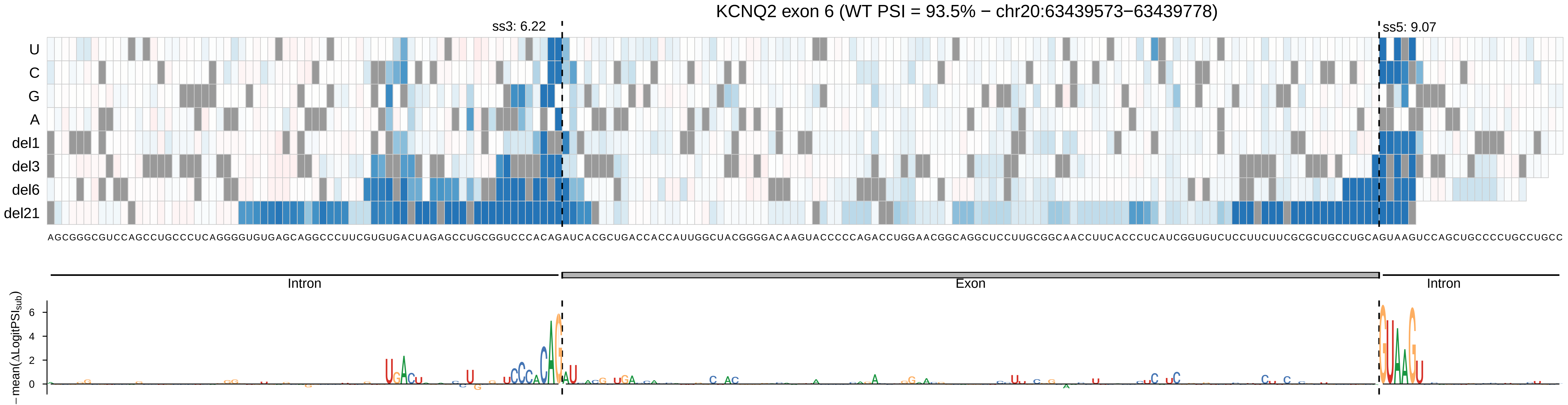
KCNQ2 exon 6 (WT PSI = 93.5% − chr20:63439573−63439778)
ss3: 6.22
ss5: 9.07
U
C
G
A
del1
del3
del6
del21
AGCGGGCGUCCAGCCUGCCCUCAGGGGUGUGAGCAGGCCCUUCGUGUGACUAGAGCCUGCGGUCCCACAGAUCACGCUGACCACCAUUGGCUACGGGGACAAGUACCCCCAGACCUGGAACGGCAGGCUCCUUGCGGCAACCUUCACCCUCAUCGGUGUCUCCUUCUUCGCGCUGCCUGCAGUAAGUCCAGCUGCCCCUGCCUGCC
Intron
Exon
Intron
−mean(ΔLogitPSI_sub)
6
4
2
0

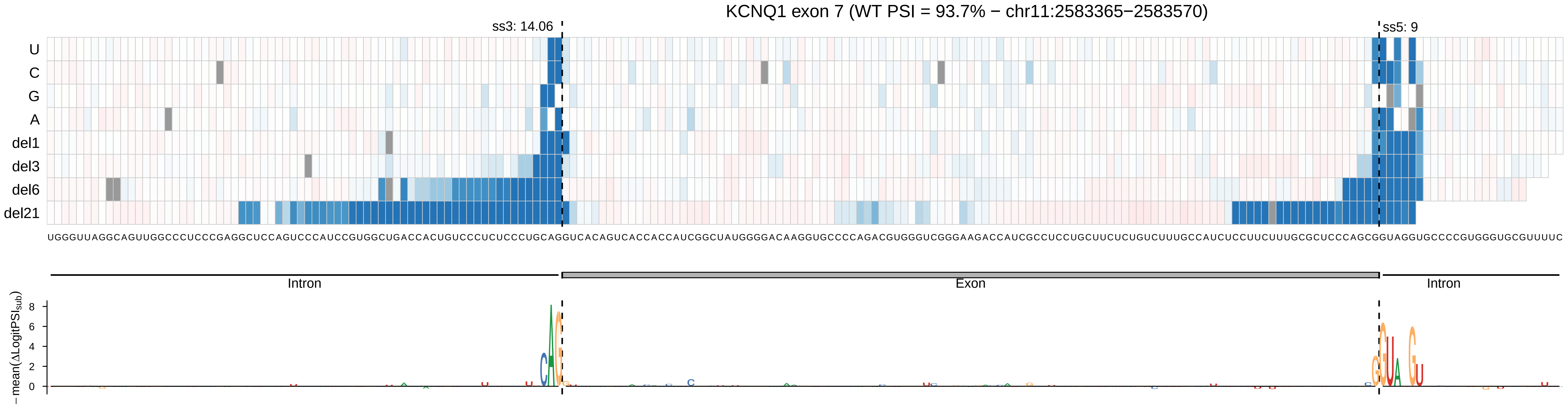

KCNQ1 exon 7 (WT PSI = 93.7% − chr11:2583365−2583570)
ss3: 14.06
ss5: 9
U
C
G
A
del1
del3
del6
del21
UGGGUUAGGCAGUUGGCCCUCCCGAGGCUCCAGUCCCAUCCGUGGCUGACCACUGUCCCUCUCCCUGCAGGUCACAGUCACCACCAUCGGCUAUGGGGACAAGGUGCCCCAGACGUGGGUCGGGAAGACCAUCGCCUCCUGCUUCUCUGUCUUUGCCAUCUCCUUCUUUGCGCUCCCAGCGGUAGGUGCCCCGUGGGUGCGUUUUC
Intron
Exon
Intron
− mean(ΔLogitPSI_sub)
8
6
4
2
0

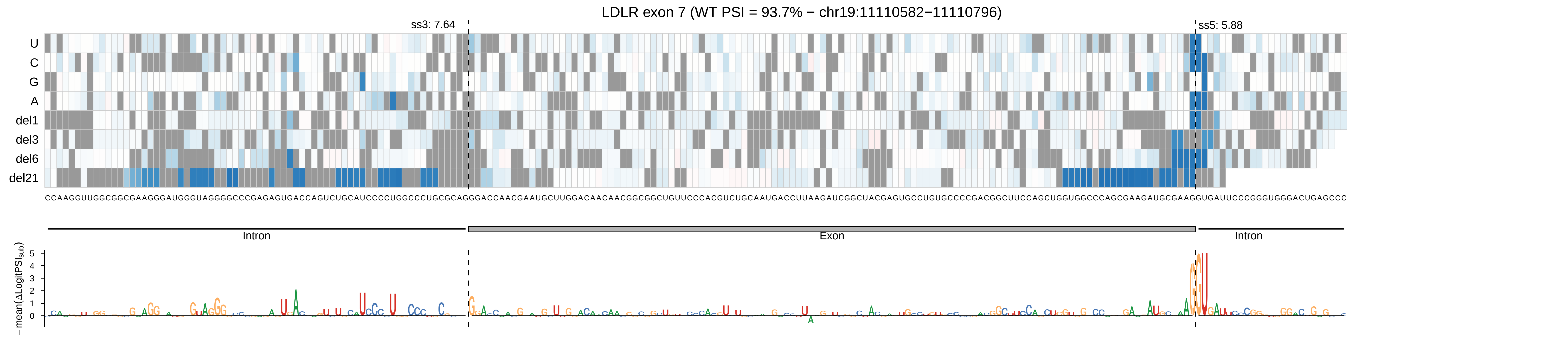

LDLR exon 7 (WT PSI = 93.7% − chr19:11110582−11110796)
ss3: 7.64
ss5: 5.88
U
C
G
A
del1
del3
del6
del21
CCAAGGUUGGCGGCGAAGGGAUGGGUAGGGGCCCGAGAGUGACCAGUCUGCAUCCCCUGGCCCUGCGCAGGGACCAACGAAUGCUUGGACAACAACGGCGGCUGUUCCCACGUCUGCAAUGACCUUAAGAUCGGCUACGAGUGCCUGUGCCCCGACGGCUUCCAGCUGGUGGCCCAGCGAAGAUGCGAAGGUGAUUCCCGGGUGGGACUGAGCCC
Intron
Exon
Intron
−mean(ΔLogitPSI_sub)

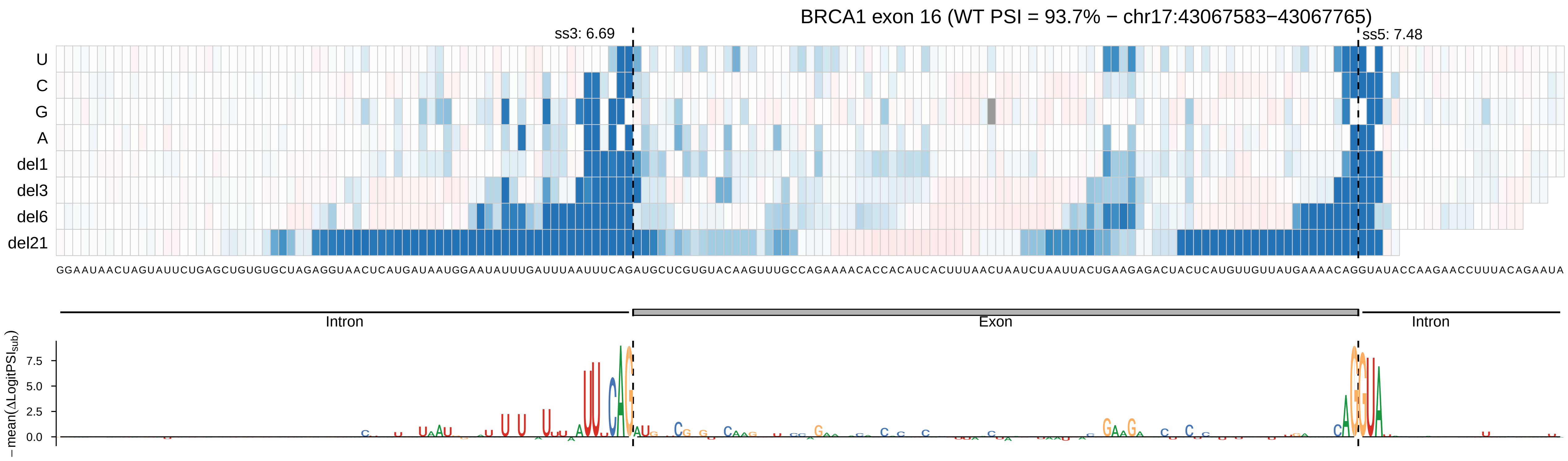
BRCA1 exon 16 (WT PSI = 93.7% − chr17:43067583−43067765)
ss3: 6.69
ss5: 7.48
U
C
G
A
del1
del3
del6
del21
GGAAUAACUAGUAUUCUGAGCUGUGUGCUAGAGGUAACUCAUGAUAAUGGAAUAUUUGAUUUAAUUUCAGAUGCUCGUGUACAAGUUUGCCAGAAAACACCACAUCACUUUAACUAAUCUAAUUACUGAAGAGACUACUCAUGUUGUUAUGAAAACAGGUAUACCAAGAACCUUUACAGAAUA
Intron
Exon
Intron
− mean(ΔLogitPSI_sub)
7.5
5.0
2.5
0.0

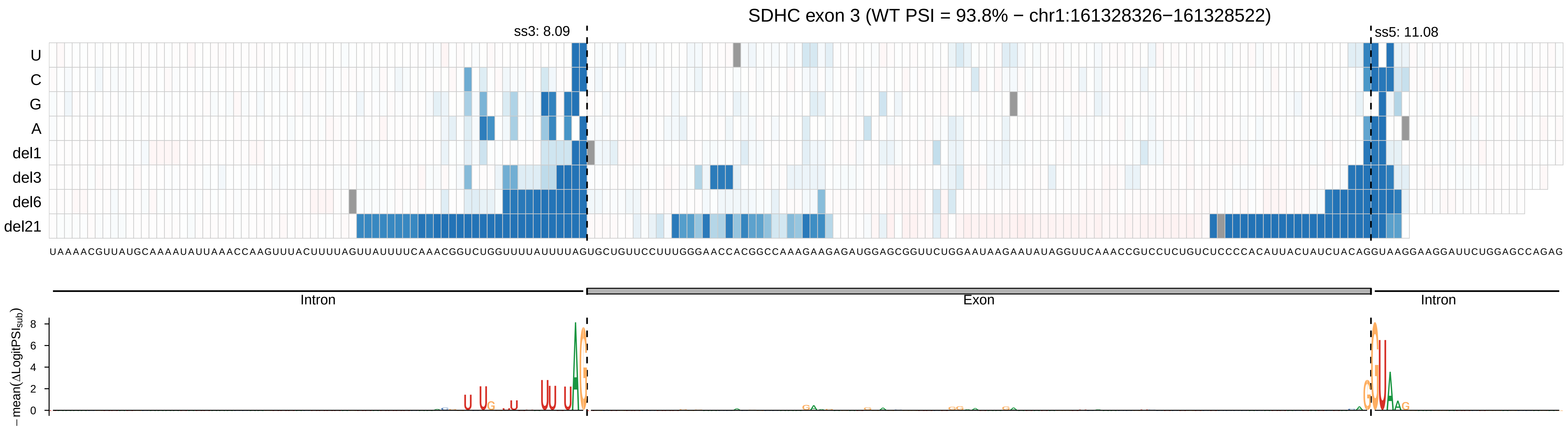

SDHC exon 3 (WT PSI = 93.8% − chr1:161328326−161328522)
ss3: 8.09
ss5: 11.08
U
C
G
A
del1
del3
del6
del21
UAAAACGUUAUGCAAAAUAUUAAACCAAGUUUACUUUUAGUUAUUUUCAAACGGUCUGGUUUUAUUUUAGUGCUGUUCCUUUGGGAACCACGGCCAAAGAAGAGAUGGAGCGGUUCUGGAAUAAGAAUAUAGGUUCAAACCGUCCUCUGUCUCCCCACAUUACUAUCUACAGGUAAGGAAGGAUUCUGGAGCCAGAG
Intron
Exon
Intron
− mean(ΔLogitPSI_sub)
8
6
4
2
0

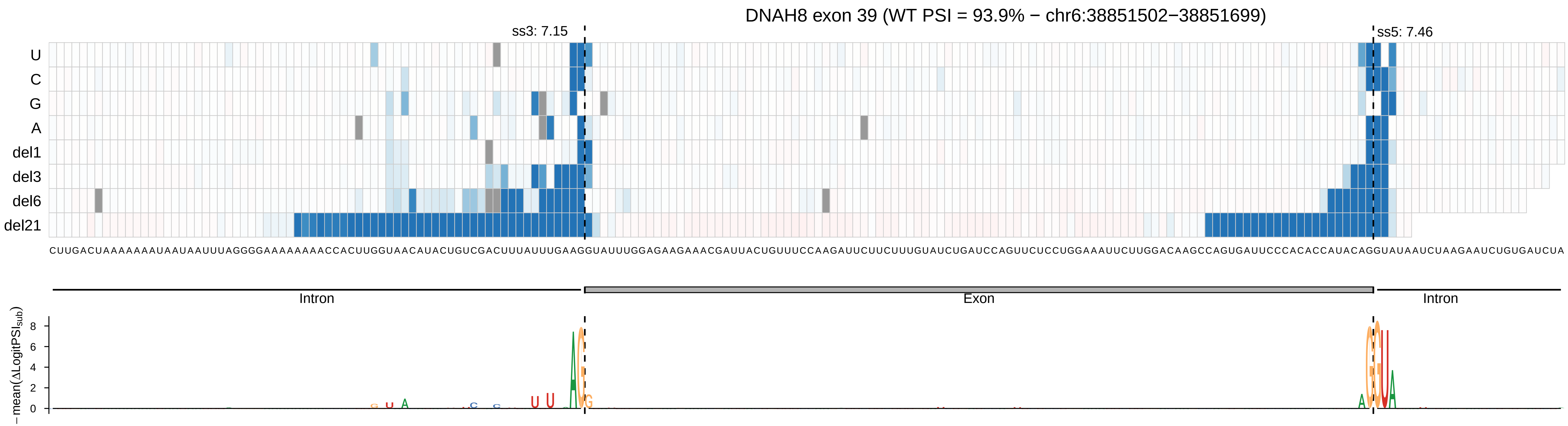

DNAH8 exon 39 (WT PSI = 93.9% − chr6:38851502−38851699)
ss3: 7.15
ss5: 7.46
U
C
G
A
del1
del3
del6
del21
CUUGACUAAAAAAAUAAUAAUUUAGGGGAAAAAAAACCACUUGGUAACAUACUGUCGACUUUAUUUGAAGGUAUUUGGAGAAGAAACGAUUACUGUUUCCAAGAUUCUUCUUUGUAUCUGAUCCAGUUCUCCUGGAAAUUCUUGGACAAGCCAGUGAUUCCCACACCAUACAGGUAUAAUCUAAGAAUCUGUGAUCUA
Intron
Exon
Intron
− mean(ΔLogitPSI_sub)
0
2
4
6
8

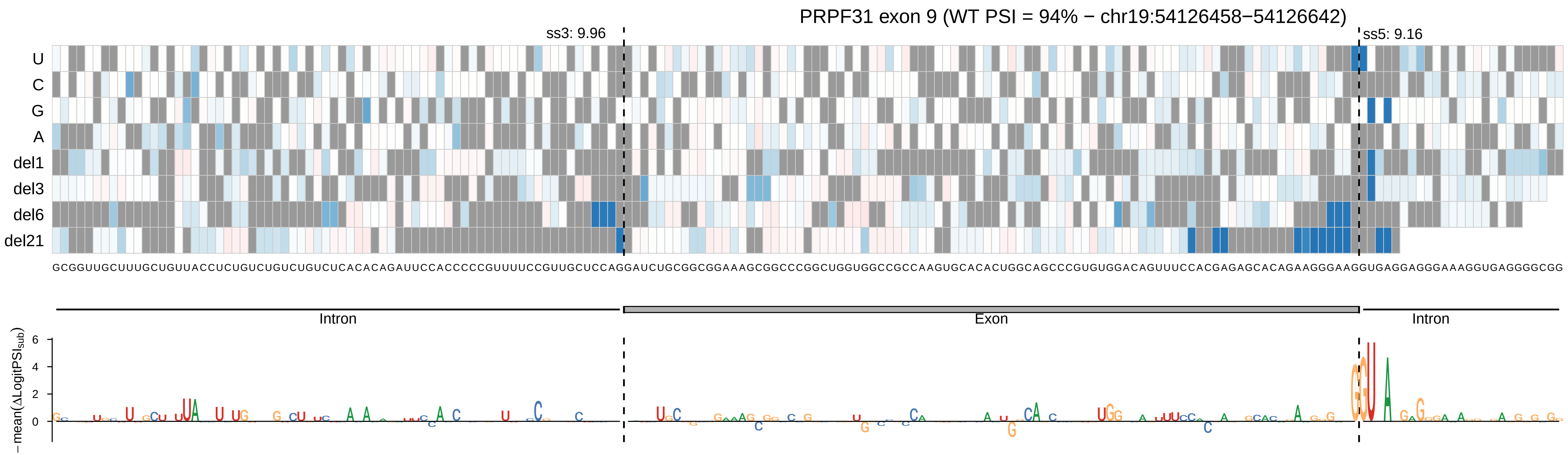
PRPF31 exon 9 (WT PSI = 94% − chr19:54126458−54126642)
ss3: 9.96
ss5: 9.16
U
C
G
A
del1
del3
del6
del21
GCGGUUGCUUUGCUGUUACCUCUGUCUGUCUGUCUCACACAGAUUCCACCCCCGUUUUCCGUUGCUCCAGGAUCUGCGGCGGAAAGCGGCCCGGCUGGUGGCCGCCAAGUGCACACUGGCAGCCCGUGUGGACAGUUUCCACGAGAGCACAGAAGGGAAGGUGAGGAGGGAAAGGUGAGGGGCGG
Intron
Exon
Intron
−mean(ΔLogitPSIsub)
6
4
2
0

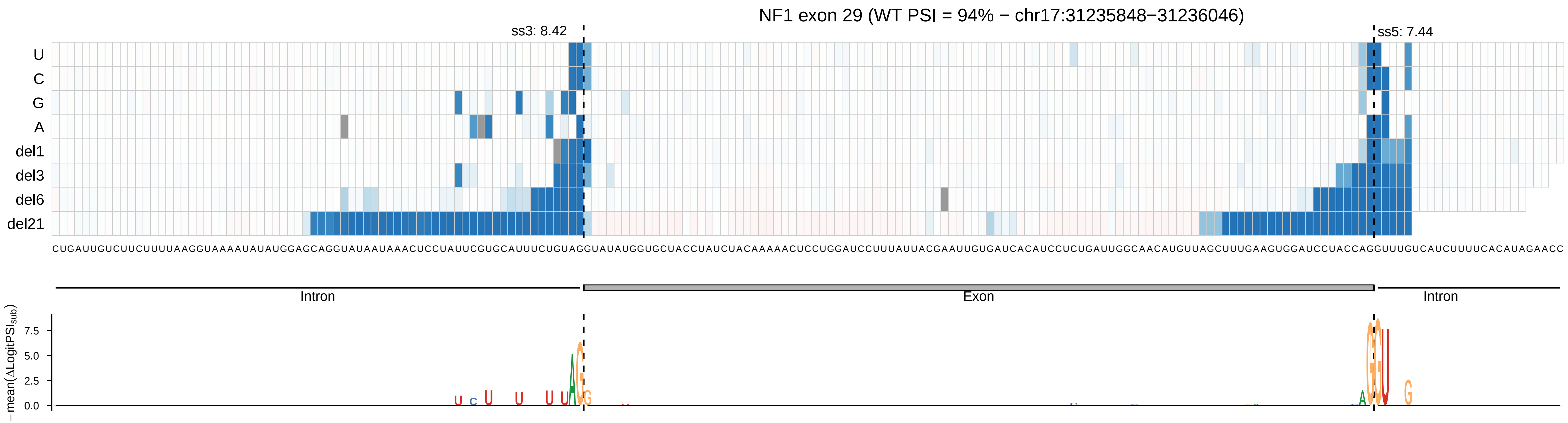
NF1 exon 29 (WT PSI = 94% − chr17:31235848−31236046)
ss3: 8.42
ss5: 7.44
U
C
G
A
del1
del3
del6
del21
CUGAUUGUCUUCUUUUAAGGUAAAAUAUAUGGAGCAGGUAUAAUAAACUCCUAUUCGUGCAUUUCUGUAGGUAUAUGGUGCUACCUAUCUACAAAAACUCCUGGAUCCUUUAUUACGAAUUGUGAUCACAUCCUCUGAUUGGCAACAUGUUAGCUUUGAAGUGGAUCCUACCAGGUUUGUCAUCUUUUCACAUAGAACC
Intron
Exon
Intron
− mean(ΔLogitPSI_sub)
7.5
5.0
2.5
0.0

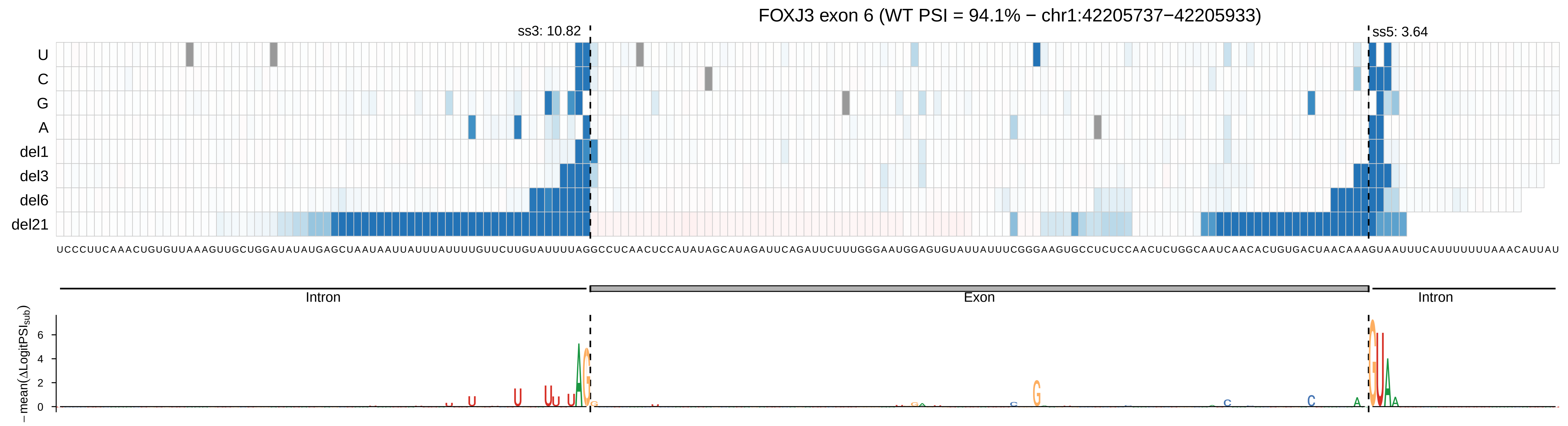
FOXJ3 exon 6 (WT PSI = 94.1% − chr1:42205737−42205933)
ss3: 10.82
ss5: 3.64
U
C
G
A
del1
del3
del6
del21
UCCCUUCAAACUGUGUUAAAGUUGCUGGAUAUAUGAGCUAAUAAUUAUUUAUUUUGUUCUUGUAUUUUAGGCCUCAACUCCAUAUAGCAUAGAUUCAGAUUCUUUGGGAAUGGAGUGUAUUAUUUCGGGAAGUGCCUCUCCAACUCUGGCAAUCAACACUGUGACUAACAAAGUAAUUUCAUUUUUUUAAACAUUAU
Intron
Exon
Intron
− mean(ΔLogitPSI_sub)
6
4
2
0

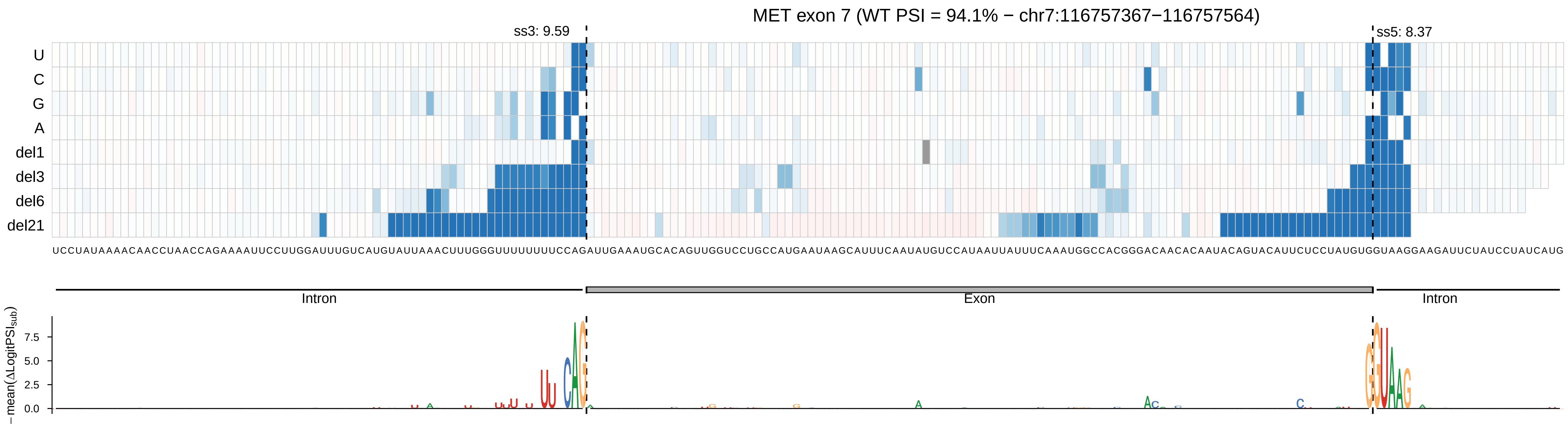
MET exon 7 (WT PSI = 94.1% − chr7:116757367−116757564)
ss3: 9.59
ss5: 8.37
U
C
G
A
del1
del3
del6
del21
UCCUAUAAAACAACCUAACCAGAAAAUUCCUUGGAUUUGUCAUGUAUUAAACUUUGGGUUUUUUUUCCAGAUUGAAAUGCACAGUUGGUCCUGCCAUGAAUAAGCAUUUCAAUAUGUCCAUAAUUAUUUCAAAUGGCCACGGGACAACACAAUACAGUACAUUCUCCUAUGUGGUAAGGAAGAUUCUAUCCUAUCAUG
Intron
Exon
Intron
− mean(ΔLogitPSI_sub)
7.5
5.0
2.5
0.0

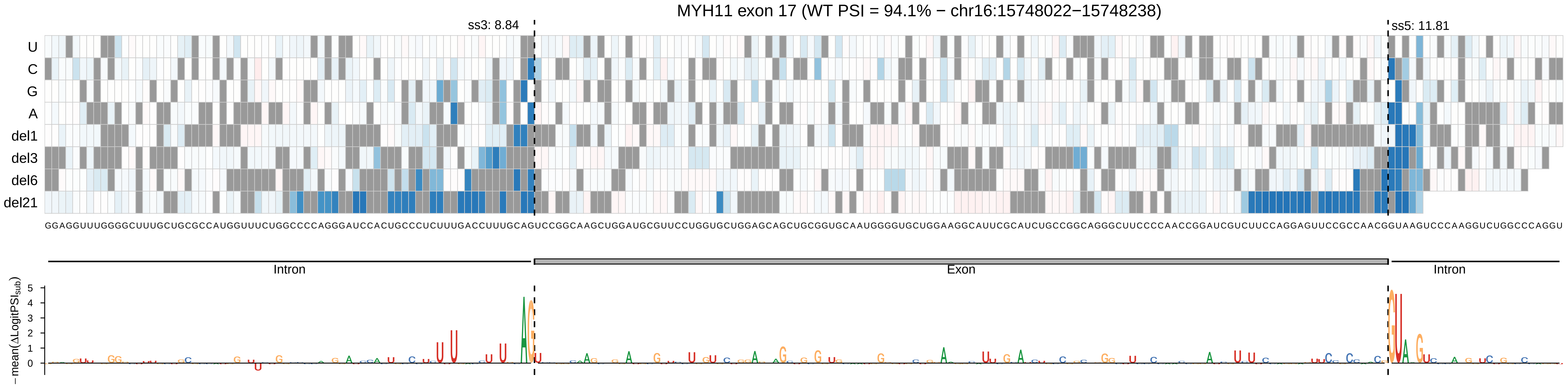

MYH11 exon 17 (WT PSI = 94.1% − chr16:15748022−15748238)
ss3: 8.84
ss5: 11.81
U
C
G
A
del1
del3
del6
del21
GGAGGUUUGGGGCUUUGCUGCGCCAUGGUUUCUGGCCCCAGGGAUCCACUGCCCUCUUUGACCUUUGCAGUCCGGCAAGCUGGAUGCGUUCCUGGUGCUGGAGCAGCUGCGGUGCAAUGGGGUGCUGGAAGGCAUUCGCAUCUGCCGGCAGGGCUUCCCCAACCGGAUCGUCUUCCAGGAGUUCCGCCAACGGUAAGUCCCAAGGUCUGGCCCAGGU
Intron
Exon
Intron
−mean(ΔLogitPSI_sub)
5
4
3
2
1
0

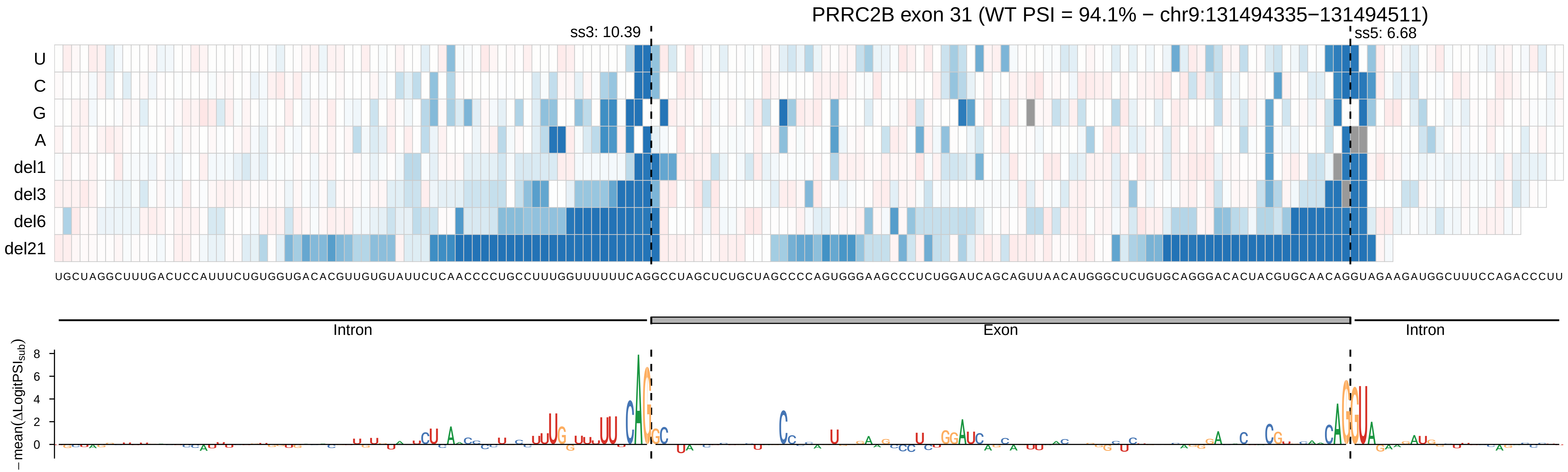
PRRC2B exon 31 (WT PSI = 94.1% − chr9:131494335−131494511)
ss3: 10.39
ss5: 6.68
U
C
G
A
del1
del3
del6
del21
UGCUAGGCUUUGACUCCAUUUCUGUGGUGACACGUUGUGUAUUCUCAACCCCUGCCUUUGGUUUUUUCAGGCCUAGCUCUGCUAGCCCCAGUGGGAAGCCCUCUGGAUCAGCAGUUAACAUGGGCUCUGUGCAGGGACACUACGUGCAACAGGUAGAAGAUGGCUUUCCAGACCCUU
Intron
Exon
Intron
− mean(ΔLogitPSI_sub)
8
6
4
2
0

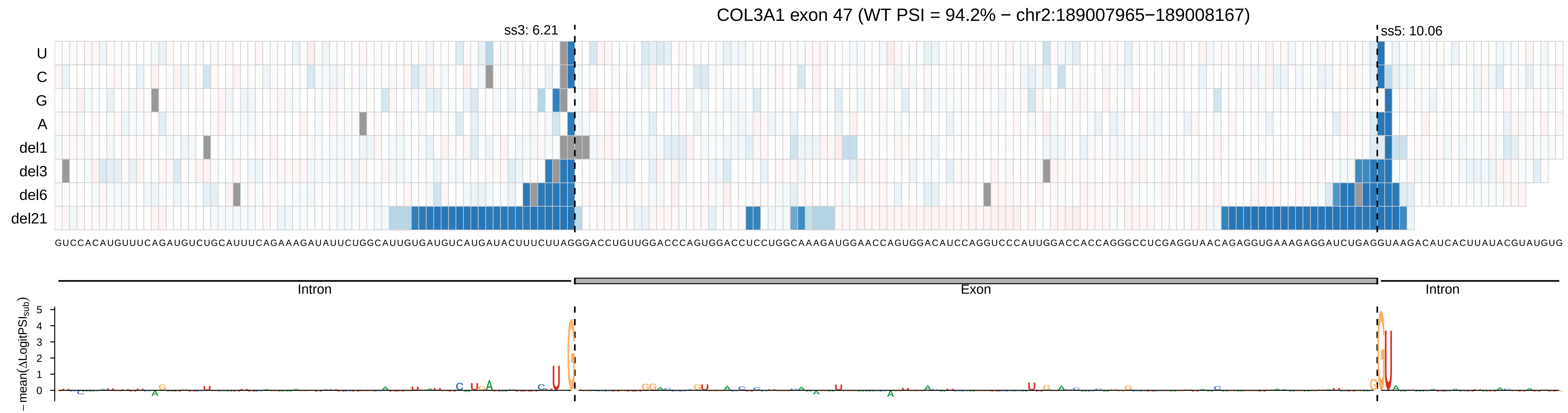

COL3A1 exon 47 (WT PSI = 94.2% − chr2:189007965−189008167)
ss3: 6.21
ss5: 10.06
U
C
G
A
del1
del3
del6
del21
GUCCACAUGUUUCAGAUGUCUGCAUUUCAGAAAGAUAUUCUGGCAUUGUGAUGUCAUGAUACUUUCUUAGGGACCUGUUGGACCCAGUGGACCUCCUGGCAAAGAUGGAACCAGUGGACAUCCAGGUCCCAUUGGACCACCAGGGCCUCGAGGUAACAGAGGUGAAAGAGGAUCUGAGGUAAGACAUCACUUAUACGUAUGUG
Intron
Exon
Intron
− mean(ΔLogitPSI_sub)
5
4
3
2
1
0

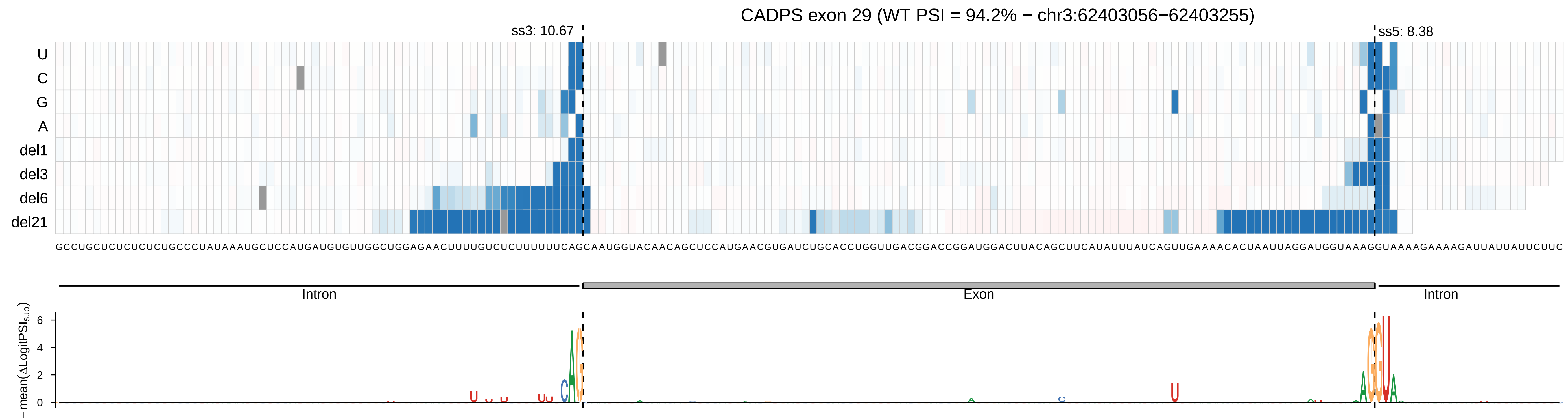
CADPS exon 29 (WT PSI = 94.2% − chr3:62403056−62403255)
ss3: 10.67
ss5: 8.38
U
C
G
A
del1
del3
del6
del21
Intron
Exon
Intron
−mean(ΔLogitPSI_sub)
6
4
2
0

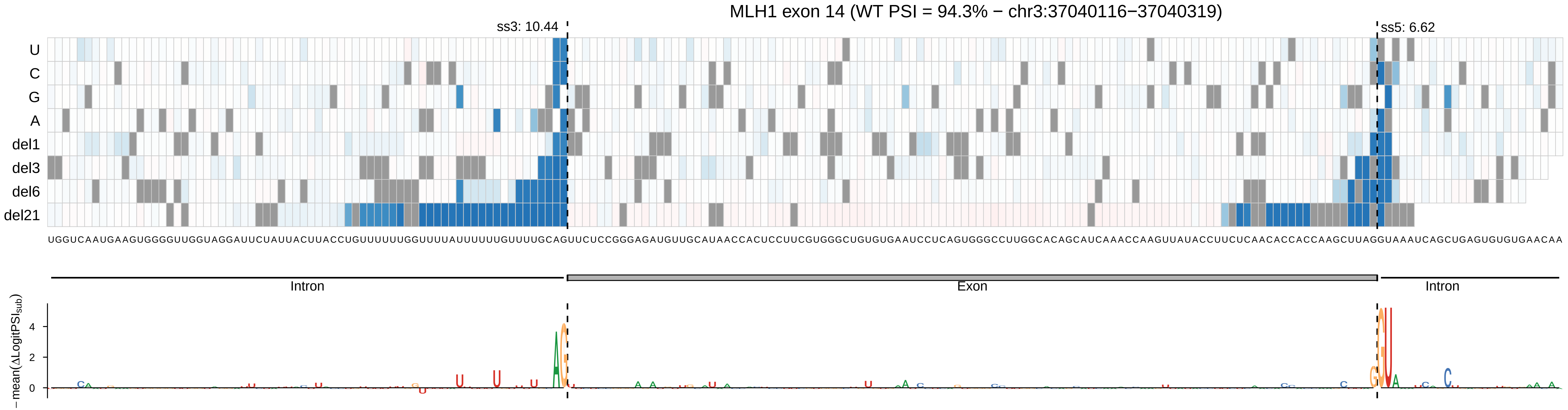

MLH1 exon 14 (WT PSI = 94.3% − chr3:37040116−37040319)
ss3: 10.44
ss5: 6.62
U
C
G
A
del1
del3
del6
del21
UGGUCAAUGAAGUGGGGUUGGUAGGAUUCUAUUACUUACCUGUUUUUUGGUUUUAUUUUUUGUUUUGCAGUUCUCCGGGAGAUGUUGCAUAACCACUCCUUCGUGGGCUGUGUGAAUCCUCAGUGGGCCUUGGCACAGCAUCAAACCAAGUUAUACCUUCUCAACACCACCAAGCUUAGGUAAAUCAGCUGAGUGUGUGAACAA
Intron
Exon
Intron
−mean(ΔLogitPSI_sub)
4
2
0

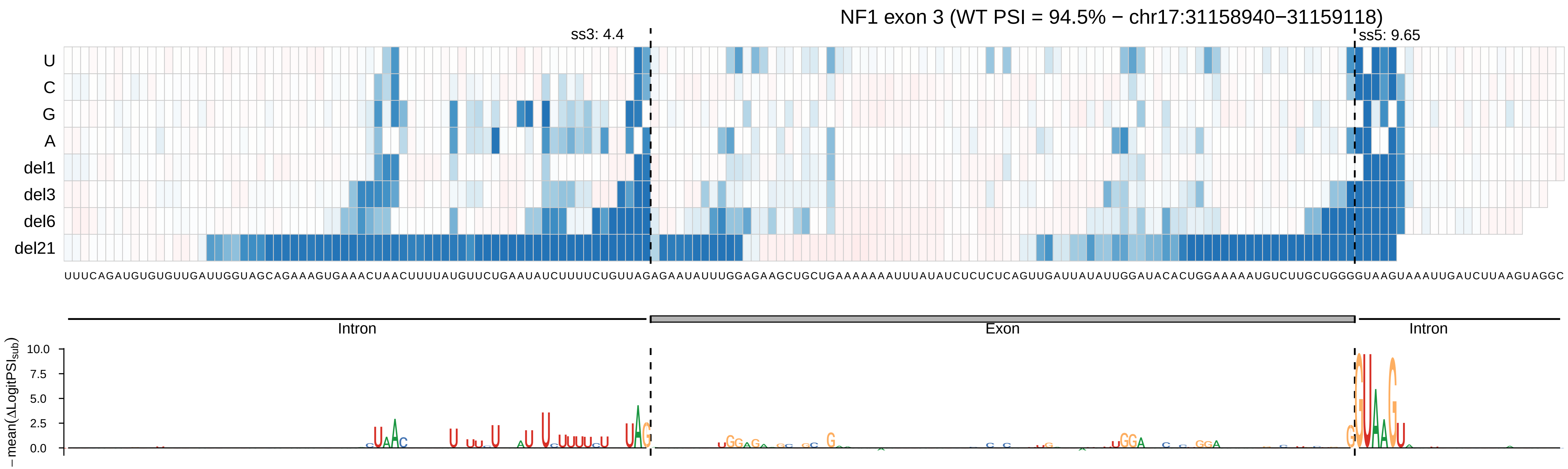

NF1 exon 3 (WT PSI = 94.5% − chr17:31158940−31159118)
ss3: 4.4
ss5: 9.65
U
C
G
A
del1
del3
del6
del21
UUUCAGAUGUGUGUUGAUUGGUAGCAGAAAGUGAAACUAACUUUUAUGUUCUGAAUAUCUUUUCUGUUAGAGAAUAUUUGGAGAAGCUGCUGAAAAAAAUUUAUAUCUCUCUCAGUUGAUUAUAUUGGAUACACUGGAAAAAUGUCUUGCUGGGGUAAGUAAAUUGAUCUUAAGUAGGC
Intron
Exon
Intron
− mean(ΔLogitPSI_sub)
10.0
7.5
5.0
2.5
0.0

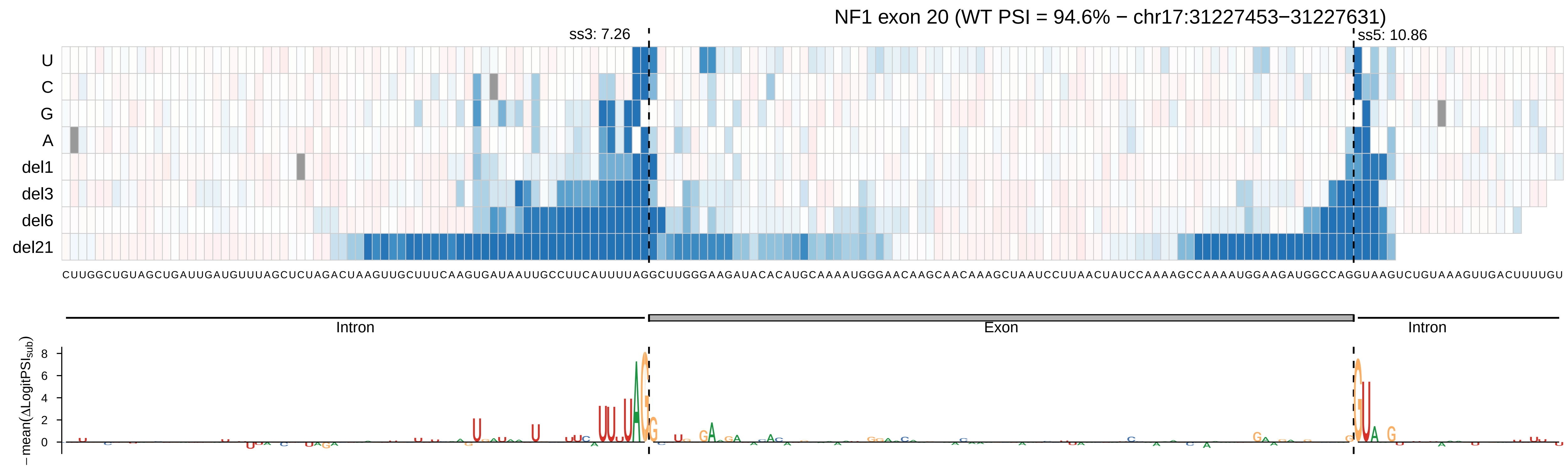

NF1 exon 20 (WT PSI = 94.6% − chr17:31227453−31227631)
ss3: 7.26
ss5: 10.86
U
C
G
A
del1
del3
del6
del21
CUUGGCUGUAGCUGAUUGAUGUUUAGCUCUAGACUAAGUUGCUUUCAAGUGAUAAUUGCCUUCAUUUUAGGCUUGGGAAGAUACACAUGCAAAAUGGGAACAAGCAACAAAGCUAAUCCUUAACUAUCCAAAAGCCAAAAUGGAAGAUGGCCAGGUAAGUCUGUAAAGUUGACUUUUGU
Intron
Exon
Intron
−mean(ΔLogitPSI_sub)
8
6
4
2
0

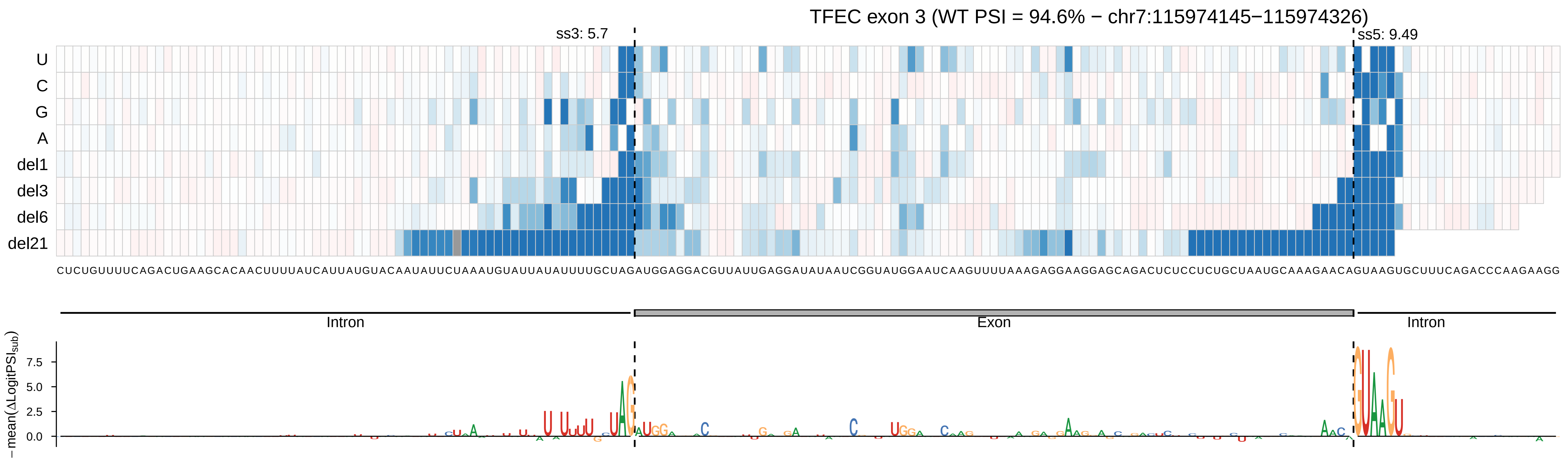
TFEC exon 3 (WT PSI = 94.6% − chr7:115974145−115974326)
ss3: 5.7
ss5: 9.49
U
C
G
A
del1
del3
del6
del21
CUCUGUUUUCAGACUGAAGCACAACUUUUAUCAUUAUGUACAAUAUUCUAAAUGUAUUAUAUUUUGCUAGAUGGAGGACGUUAUUGAGGAUAUAAUCGGUAUGGAAUCAAGUUUUAAAGAGGAAGGAGCAGACUCUCCUCUGCUAAUGCAAAGAACAGUAAGUGCUUUCAGACCCAAGAAGG
Intron
Exon
Intron
− mean(ΔLogitPSI_sub)
7.5
5.0
2.5
0.0

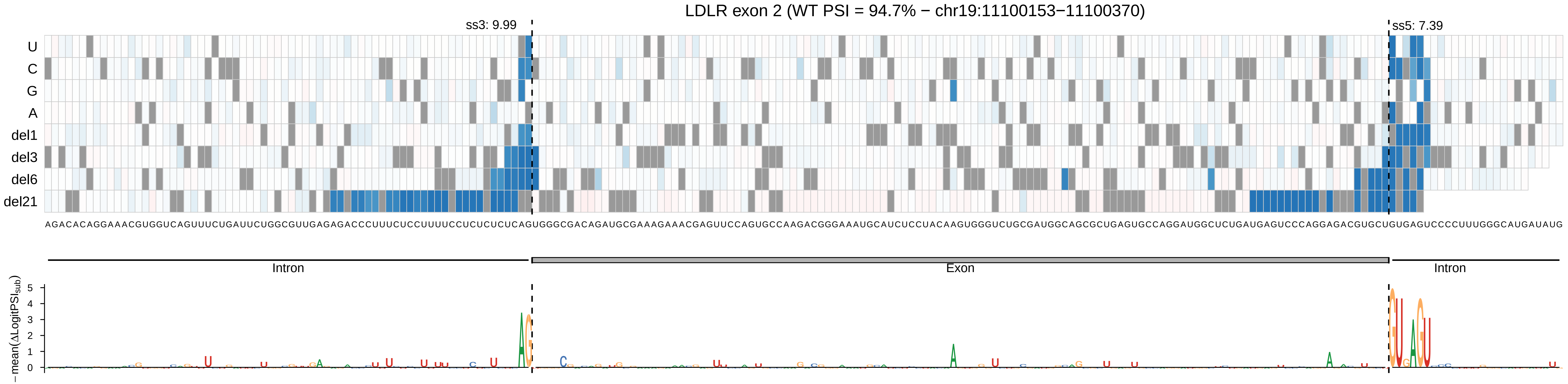

LDLR exon 2 (WT PSI = 94.7% − chr19:11100153−11100370)
ss3: 9.99
ss5: 7.39
U
C
G
A
del1
del3
del6
del21
AGACACAGGAAACGUGGUCAGUUUCUGAUUCUGGCGUUGAGAGACCCUUUCUCCUUUUCCUCUCUCUCAGUGGGCGACAGAUGCGAAAGAAACGAGUUCCAGUGCCAAGACGGGAAAUGCAUCUCCUACAAGUGGGUCUGCGAUGGCAGCGCUGAGUGCCAGGAUGGCUCUGAUGAGUCCCAGGAGACGUGCUGUGAGUCCCCUUUGGGCAUGAUAUG
Intron
Exon
Intron
−mean(ΔLogitPSI_sub)

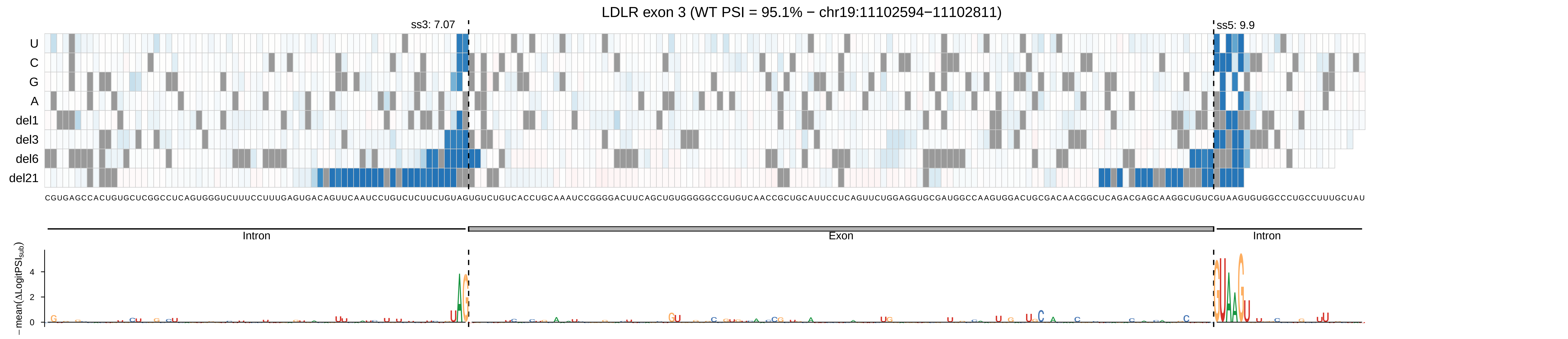
LDLR exon 3 (WT PSI = 95.1% − chr19:11102594−11102811)
ss3: 7.07
ss5: 9.9
U
C
G
A
del1
del3
del6
del21
CGUGAGCCACUGUGCUCGGCCUCAGUGGGUCUUUCCUUUGAGUGACAGUUCAAUCCUGUCUCUUCUGUAGUGUCUGUCACCUGCAAAUCCGGGGACUUCAGCUGUGGGGGCCGUGUCAACCGCUGCAUUCCUCAGUUCUGGAGGUGCGAUGGCCAAGUGGACUGCGACAACGGCUCAGACGAGCAAGGCUGUCGUAAGUGUGGCCCUGCCUUUGCUAU
Intron
Exon
Intron
− mean(ΔLogitPSI_sub)
4
2
0

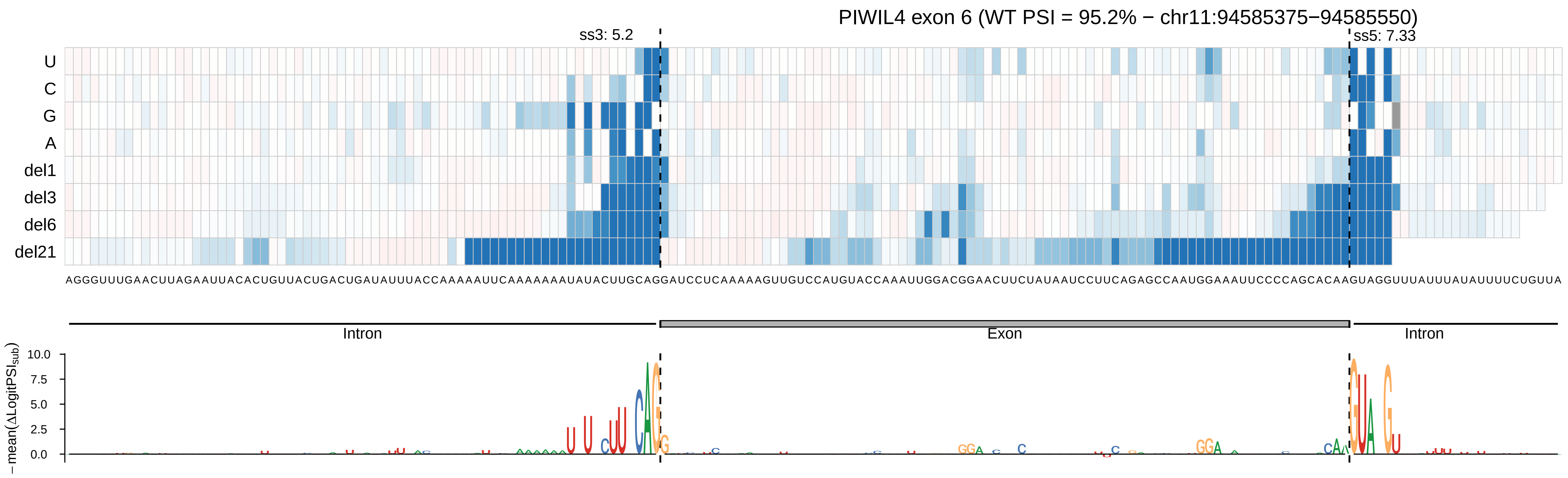

PIWIL4 exon 6 (WT PSI = 95.2% − chr11:94585375−94585550)
ss3: 5.2
ss5: 7.33
U
C
G
A
del1
del3
del6
del21
AGGGUUUGAACUUAGAAUUACACUGUUACUGACUGAUAUUUACCAAAAAUUCAAAAAAAAUAUACUUGCAGGAUCCUCAAAAAGUUGUCCAUGUACCAAAUUGGACGGAACUUCUAUAAUCCUUCAGAGCCAAUGGAAAUUCCCCAGCACAAGUAGGUUUAUUUAUAUUUUCUGUUA
Intron
Exon
Intron
− mean(ΔLogitPSI_sub)
10.0
7.5
5.0
2.5
0.0

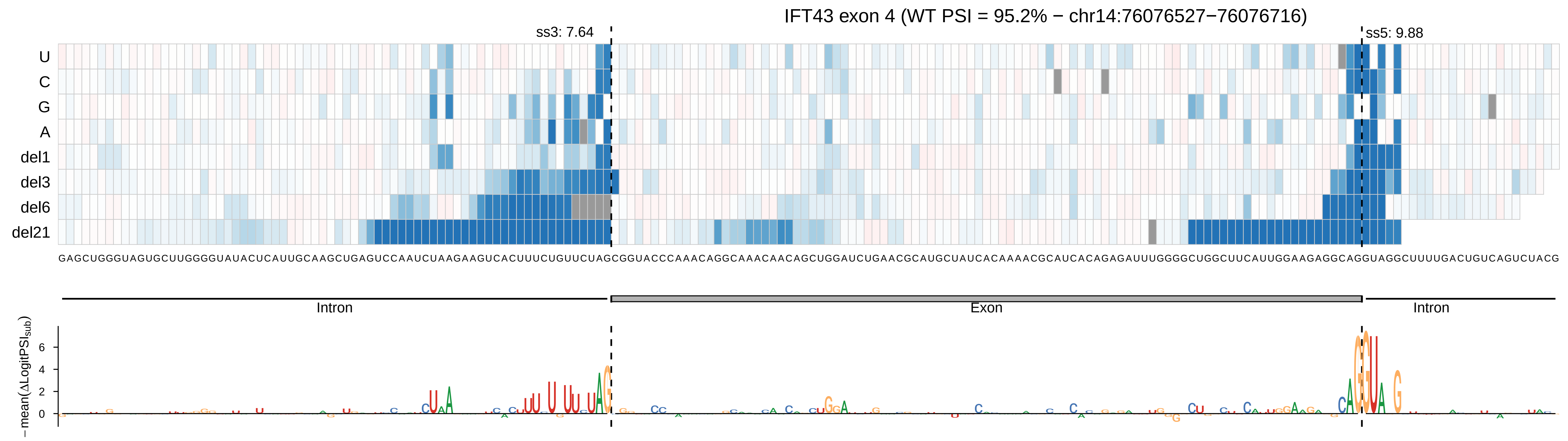
IFT43 exon 4 (WT PSI = 95.2% − chr14:76076527−76076716)
ss3: 7.64
ss5: 9.88
U
C
G
A
del1
del3
del6
del21
GAGCUGGGUAGUGCUUGGGGUAUACUCAUUGCAAGCUGAGUCCAAUCUAAGAAGUCACUUUCUGUUCUAGCGGUACCCAAACAGGCAAACAACAGCUGGAUCUGAACGCAUGCUAUCACAAAACGCAUCACAGAGAUUUGGGGCUGGCUUCAUUGGAAGAGGCAGGUAGGCUUUUGACUGUCAGUCUACG
Intron
Exon
Intron
−mean(ΔLogitPSIsub)
6
4
2
0

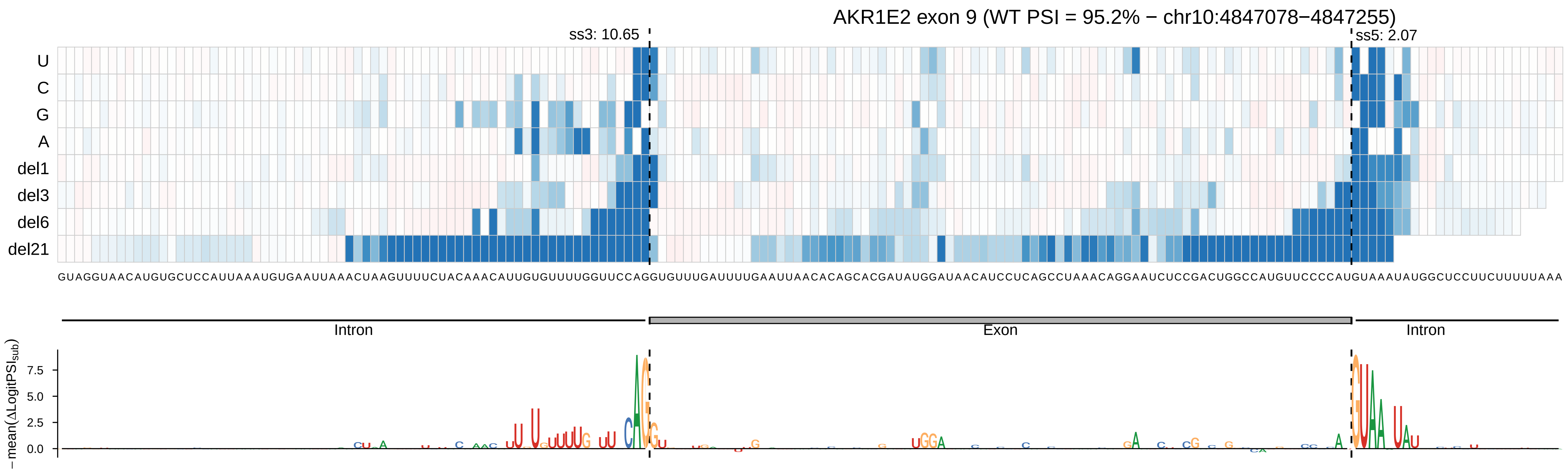
AKR1E2 exon 9 (WT PSI = 95.2% − chr10:4847078−4847255)
ss3: 10.65
ss5: 2.07
U
C
G
A
del1
del3
del6
del21
GUAGGUAACAUGUGCUCCAUUAAAUGUGAAUUAAACUAAGUUUUCUACAAACAUUGUGUUUUGGUUCCAGGUGUUUGAUUUUGAAUUAACACAGCACGAUAUGGAUAACAUCCUCAGCCUAAACAGGAAUCUCCGACUGGCCAUGUUCCCCAUGUAAAUAUGGCUCCUUCUUUUUAAA
Intron
Exon
Intron
− mean(ΔLogitPSI_sub)
7.5
5.0
2.5
0.0

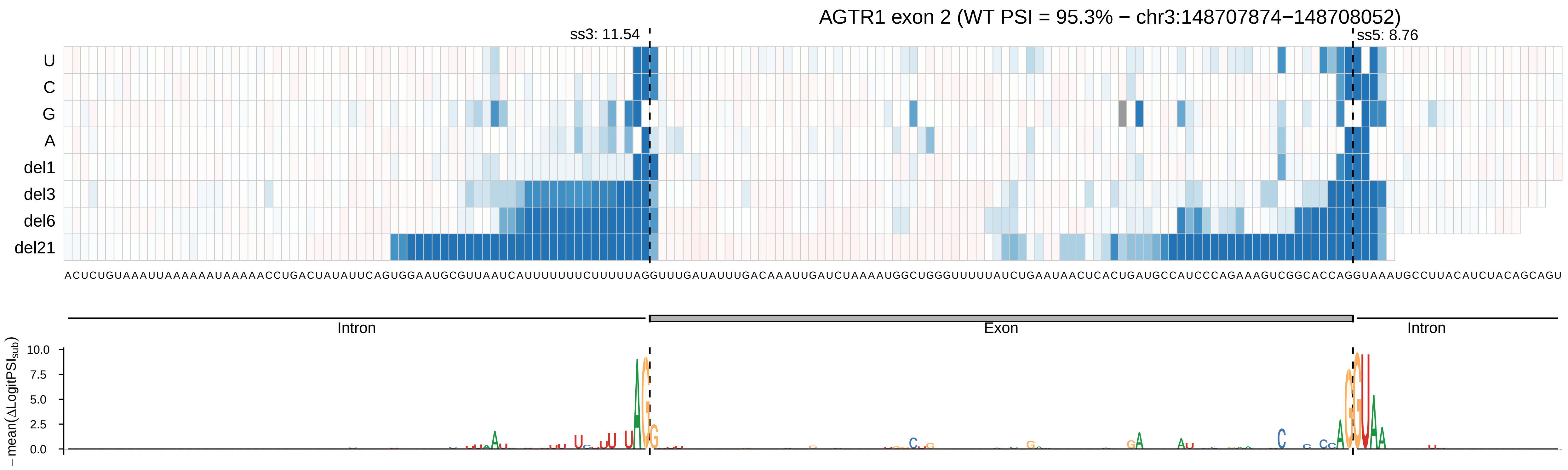

AGTR1 exon 2 (WT PSI = 95.3% − chr3:148707874−148708052)
ss3: 11.54
ss5: 8.76
U
C
G
A
del1
del3
del6
del21
ACUCUGUAAAUUAAAAAAUAAAAACCUGACUAUAUUCAGUGGAAUGCGUUAAUCAUUUUUUUCUUUUUAGGUUUGAUAUUUGACAAAUUGAUCUAAAAUGGCUGGGUUUUUAUCUGAAUAACUCACUGAUGCCAUCCCAGAAAGUCGGCACCAGGUAAAUGCCUUACAUCUACAGCAGU
Intron
Exon
Intron
−mean(ΔLogitPSI_sub)
10.0
7.5
5.0
2.5
0.0

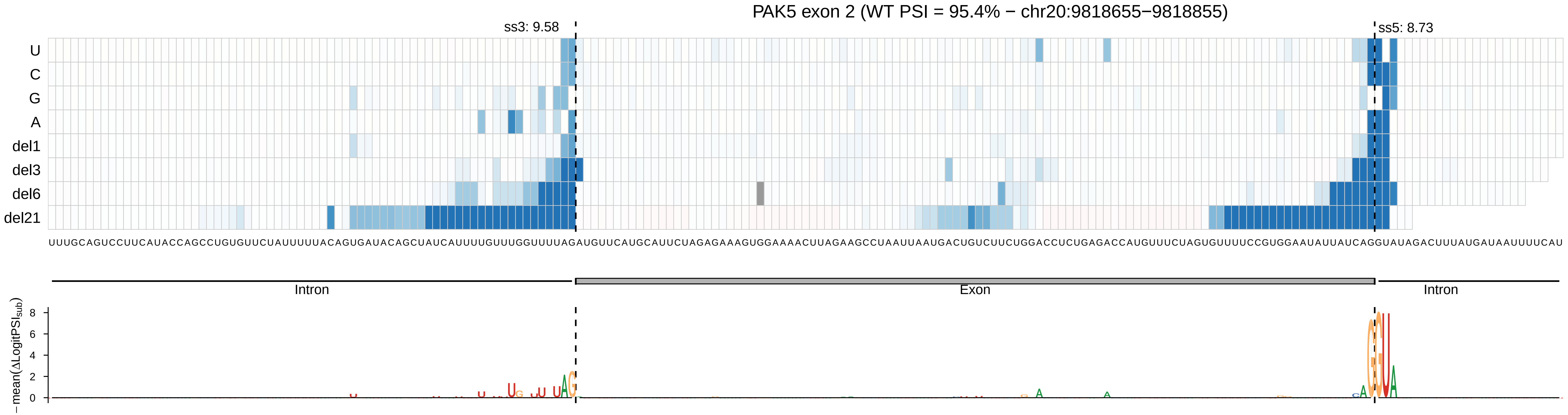
PAK5 exon 2 (WT PSI = 95.4% − chr20:9818655−9818855)
ss3: 9.58
ss5: 8.73
U
C
G
A
del1
del3
del6
del21
UUUGCAGUCCUUCAUACCAGCCUGUGUUCUAUUUUUACAGUGAUACAGCUAUCAUUUUGUUUGGUUUUAGAUGUUCAUGCAUUCUAGAGAAAGUGGAAAACUUAGAAGCCUAAUUAAUGACUGUCUUCUGGACCUCUGAGACCAUGUUUCUAGUGUUUUCCGUGGAAUAUUAUCAGGUAUAGACUUUAUGAUAAUUUUCAU
Intron
Exon
Intron
− mean(ΔLogitPSI_sub)
8
6
4
2
0

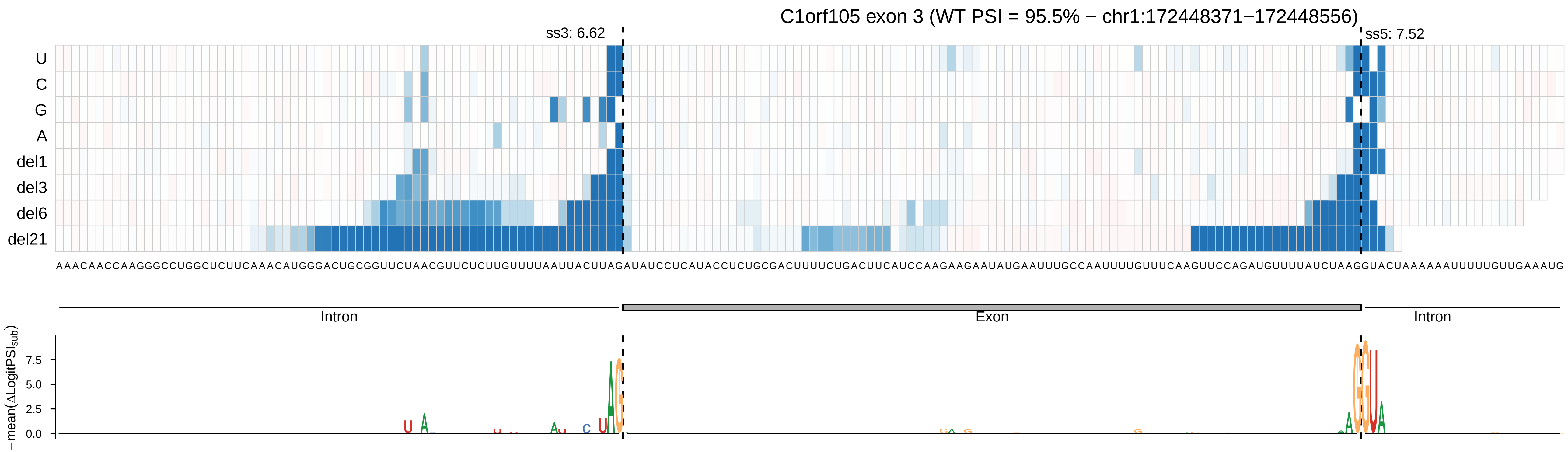
C1orf105 exon 3 (WT PSI = 95.5% − chr1:172448371−172448556)
ss3: 6.62
ss5: 7.52
U
C
G
A
del1
del3
del6
del21
AAACAACCAAGGGCCUGGCUCUUCAAACAUGGGACUGCGGUUCUAACGUUCUCUUGUUUUAAUUACUUAGAUAUCCUCAUACCUCUGCGACUUUUCUGACUUCAUCCAAGAAGAAUAUGAAUUUGCCAAUUUUGUUUCAAGUUCCAGAUGUUUUAUCUAAGGUACUAAAAAAUUUUUGUUGAAAUG
Intron
Exon
Intron
− mean(ΔLogitPSI_sub)
7.5
5.0
2.5
0.0

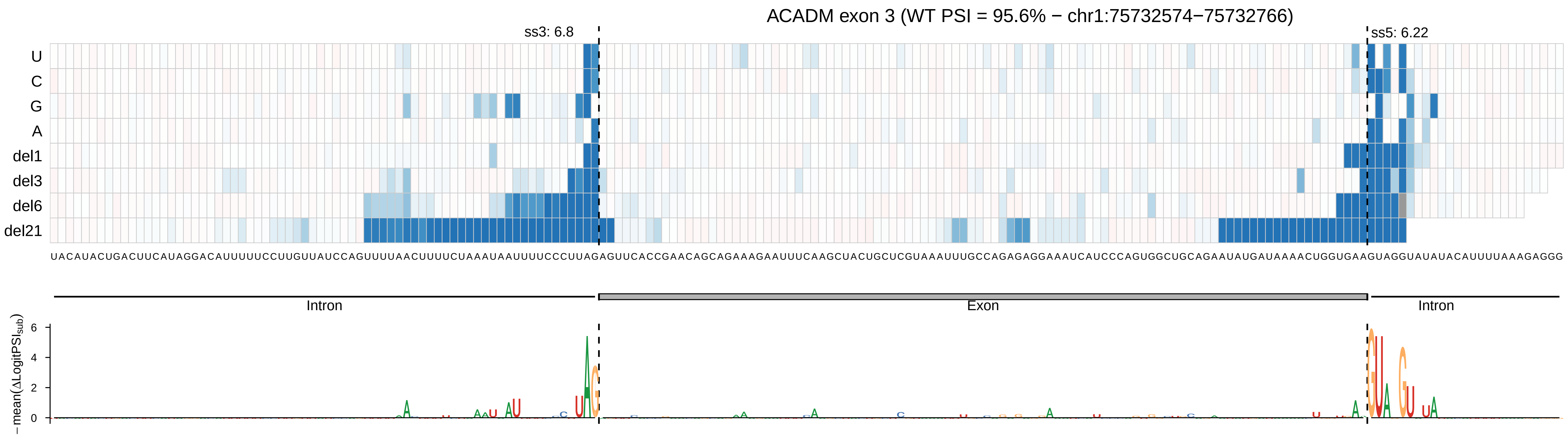
ACADM exon 3 (WT PSI = 95.6% − chr1:75732574−75732766)
ss3: 6.8
ss5: 6.22
U
C
G
A
del1
del3
del6
del21
UACAUACUGACUUCAUAGGACAUUUUUCCUUGUUAUCCAGUUUUAACUUUUCUAAAUAAUUUUCCCUUAGAGUUCACCGAACAGCAGAAAGAAUUUCAAGCUACUGCUCGUAAAUUUGCCAGAGAGGAAAUCAUCCCAGUGGCUGCAGAAUAUGAUAAAACUGGUGAAGUAGGUAUAUACAUUUUAAAGAGGG
Intron
Exon
Intron
− mean(ΔLogitPSI_sub)
6
4
2
0

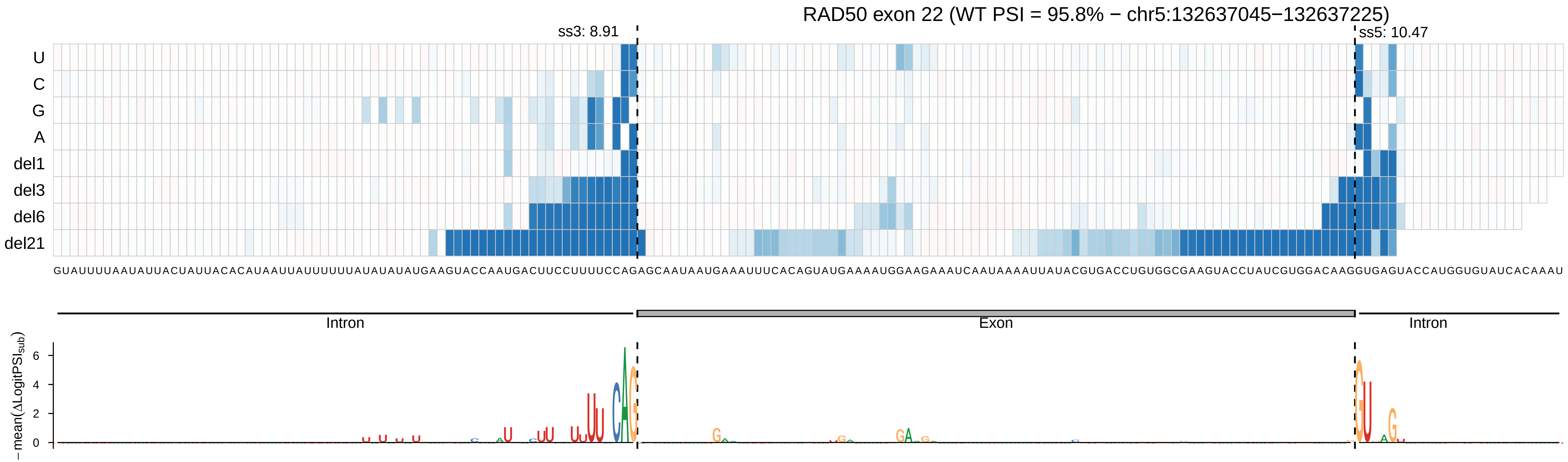

RAD50 exon 22 (WT PSI = 95.8% − chr5:132637045−132637225)
ss3: 8.91
ss5: 10.47
U
C
G
A
del1
del3
del6
del21
GUAUUUUAAUAUUACUAUUACACAUAAUUAUUUUUUAUAUAUAUGAAGUACCAAUGACUUCCUUUUCCAGAGCAAUAAUGAAAUUUCACAGUAUGAAAAUGGAAGAAAUCAAUAAAAUUAUACGUGACCUGUGGCGAAGUACCUAUCGUGGACAAGGUGAGUACCAUGGUGUAUCACAAAU
Intron
Exon
Intron
− mean(ΔLogitPSI_sub)
6
4
2
0

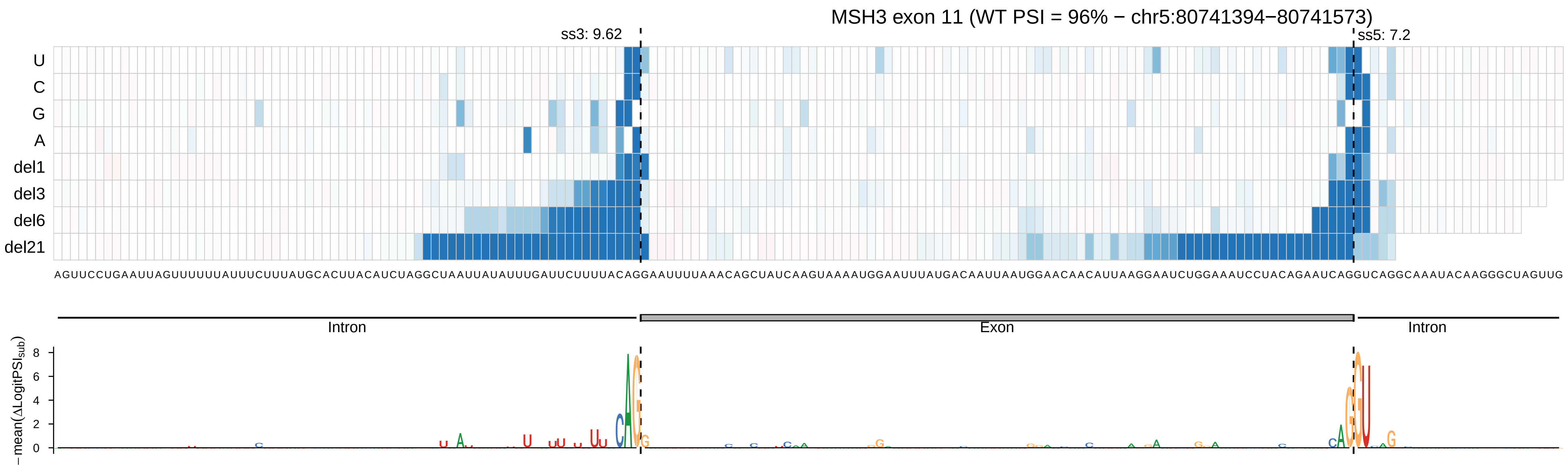
MSH3 exon 11 (WT PSI = 96% − chr5:80741394−80741573)
ss3: 9.62
ss5: 7.2
U
C
G
A
del1
del3
del6
del21
Intron
Exon
Intron
−mean(ΔLogitPSI_sub)
8
6
4
2
0

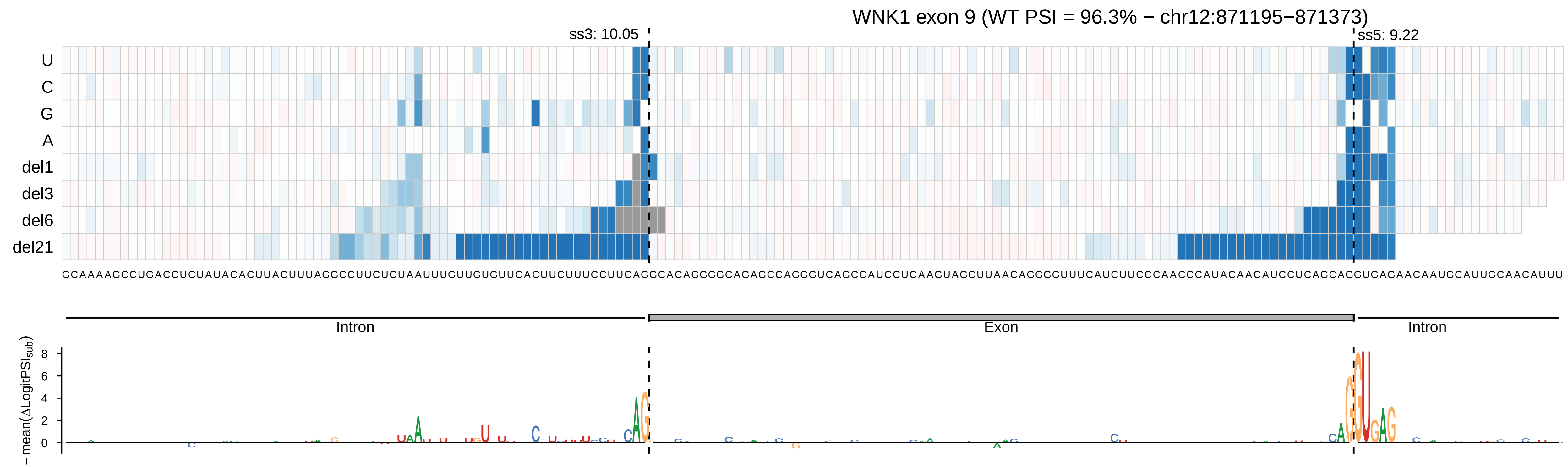

WNK1 exon 9 (WT PSI = 96.3% − chr12:871195−871373)
ss3: 10.05
ss5: 9.22
U
C
G
A
del1
del3
del6
del21
GCAAAAGCCUGACCUCUAUACACUUACUUUAGGCCUUCUCUAAUUUGUUGUGUUCACUUCUUUCCUUCAGGCACAGGGGCAGAGCCAGGGUCAGCCAUCCUCAAGUAGCUUAACAGGGGUUUCAUCUUCCCAACCCAUACAACAUCCUCAGCAGGUGAGAACAAUGCAUUGCAACAUUU
Intron
Exon
Intron
− mean(ΔLogitPSI_sub)
8
6
4
2
0

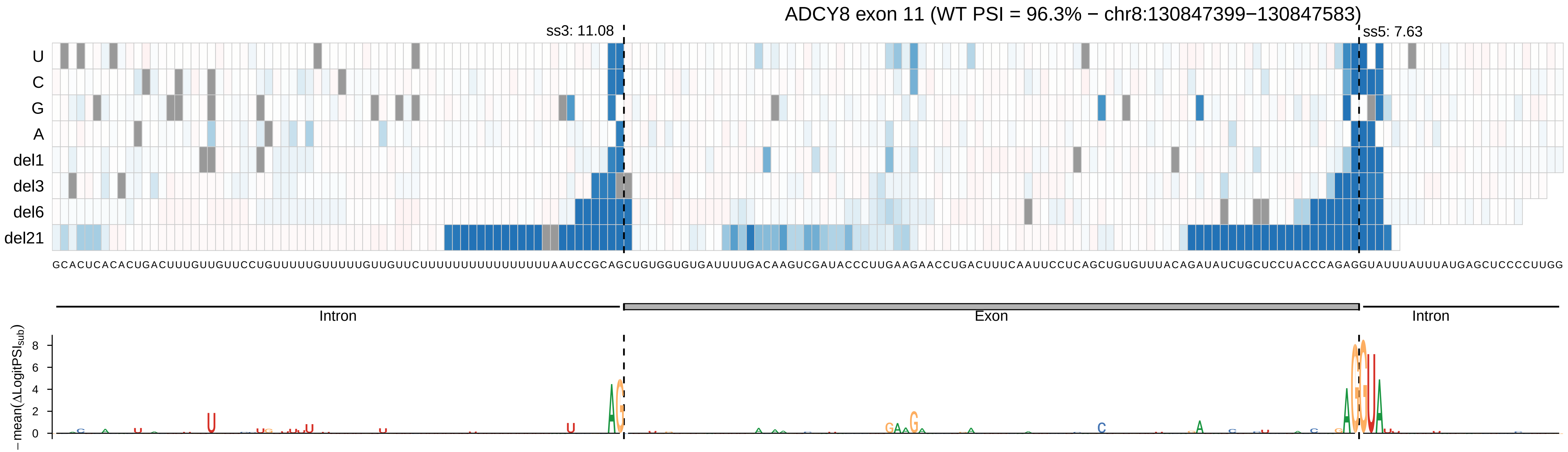

ADCY8 exon 11 (WT PSI = 96.3% − chr8:130847399−130847583)
ss3: 11.08
ss5: 7.63
U
C
G
A
del1
del3
del6
del21
GCACUCACACUGACUUUGUUGUUCCUGUUUUUGUUUUUGUUGUUCUUUUUUUUUUUUUUUUAAUCCGCAGCUGUGGUGUGAUUUUGACAAGUCGAUACCCUUGAAGAACCUGACUUUCAAUUCCUCAGCUGUGUUUACAGAUAUCUGCUCCUACCCAGAGGUAUUUAUUUAUGAGCUCCCCUUGG
Intron
Exon
Intron
− mean(ΔLogitPSI_sub)
8
6
4
2
0

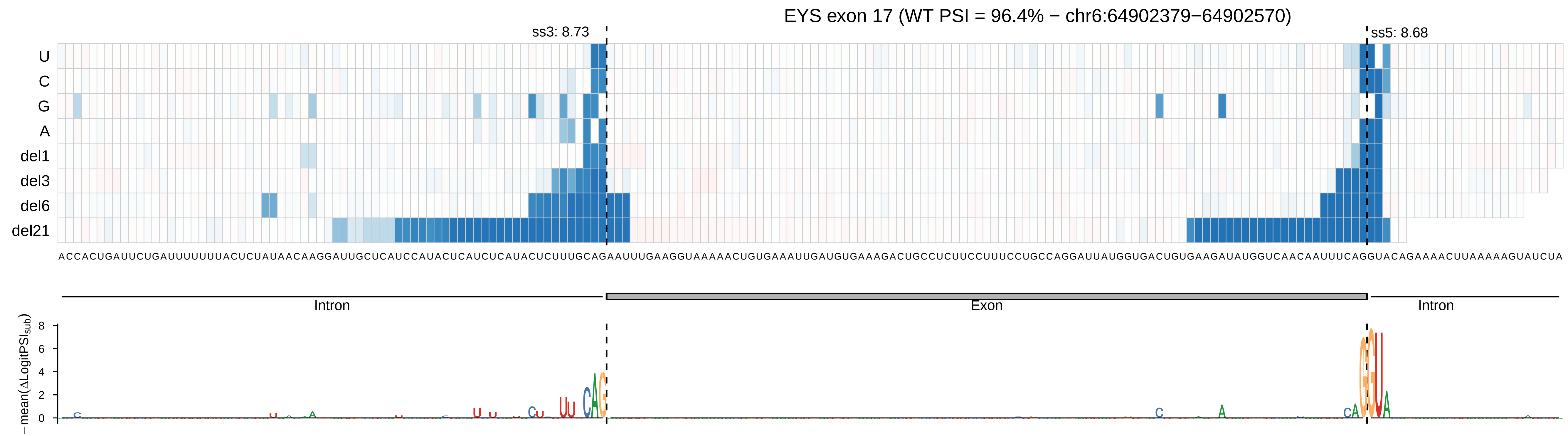
EYS exon 17 (WT PSI = 96.4% − chr6:64902379−64902570)
ss3: 8.73
ss5: 8.68
U
C
G
A
del1
del3
del6
del21
Intron
Exon
Intron
−mean(ΔLogitPSI_sub)
8
6
4
2
0

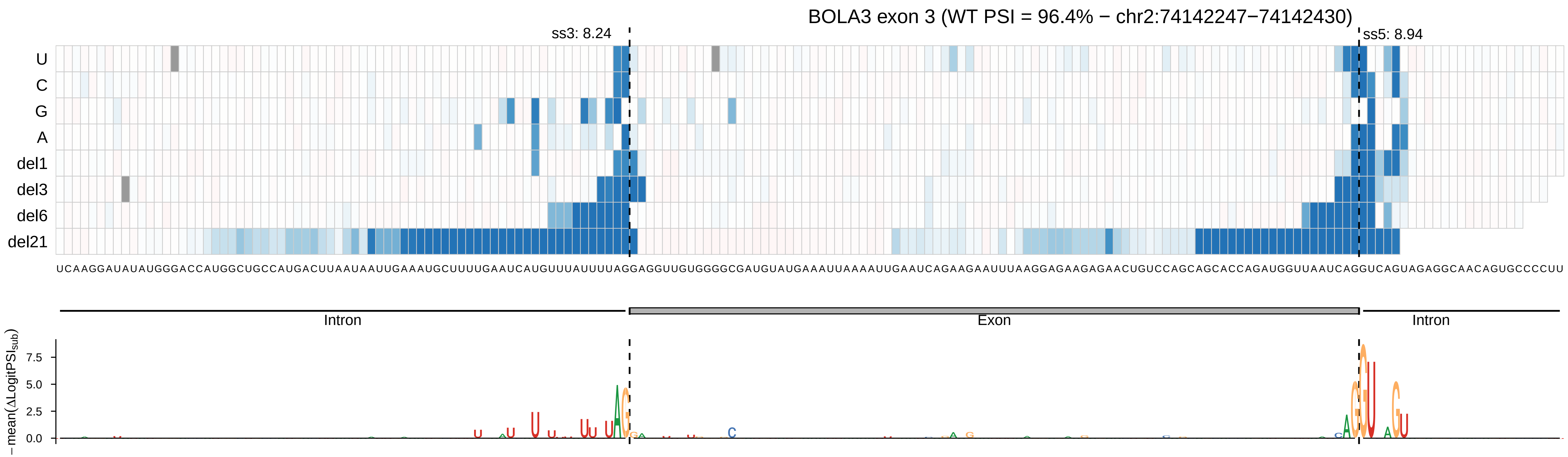

BOLA3 exon 3 (WT PSI = 96.4% − chr2:74142247−74142430)
ss3: 8.24
ss5: 8.94
U
C
G
A
del1
del3
del6
del21
UCAAGGAUAUAUGGGACCAUGGCUGCCAUGACUUAAUAAUUGAAAUGCUUUUGAAUCAUGUUUAUUUUAGGAGGUUGUGGGGCGAUGUAUGAAAUUAAAAUUGAAUCAGAAGAAUUUAAGGAGAAGAGAACUGUCCAGCAGCACCAGAUGGUUAAUCAGGUCAGUAGAGGCAACAGUGCCCCUU
Intron
Exon
Intron
− mean(ΔLogitPSI_sub)
7.5
5.0
2.5
0.0

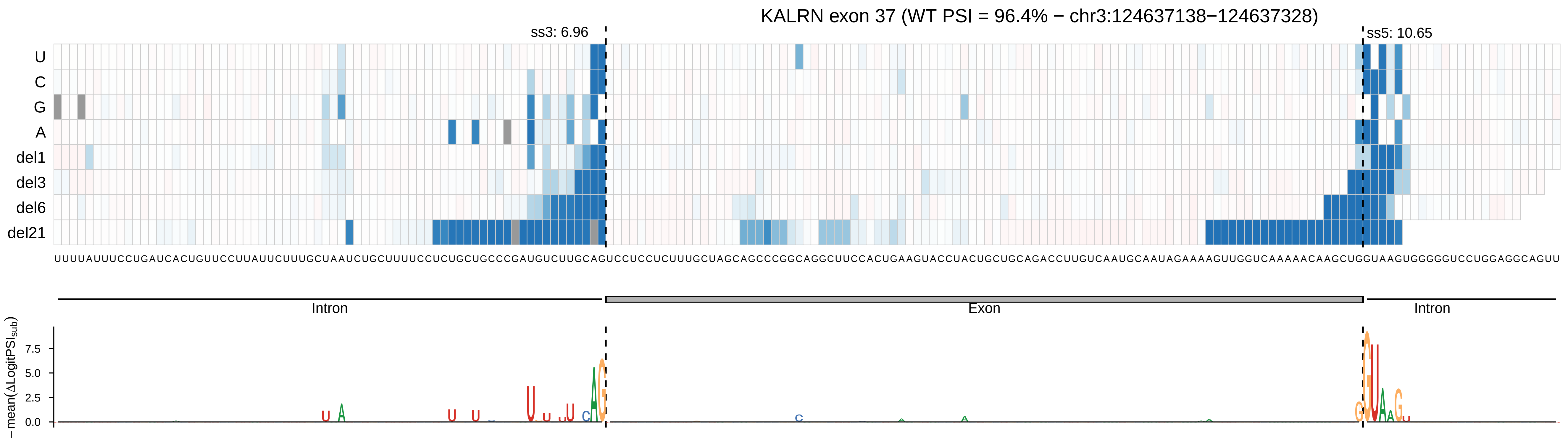

KALRN exon 37 (WT PSI = 96.4% − chr3:124637138−124637328)
ss3: 6.96
ss5: 10.65
U
C
G
A
del1
del3
del6
del21
UUUUAUUUCCUGAUCACUGUUCCUUAUUCUUUGCUAAUCUGCUUUUCCUCUGCUGCCCGAUGUCUUGCAGUCCUCCUCUUUGCUAGCAGCCCGGCAGGCUUCCACUGAAGUACCUACUGCUGCAGACCUUGUCAAUGCAAUAGAAAAGUUGGUCAAAAACAAGCUGGUAAGUGGGGGUCCUGGAGGCAGUU
Intron
Exon
Intron
− mean(ΔLogitPSI_sub)
7.5
5.0
2.5
0.0

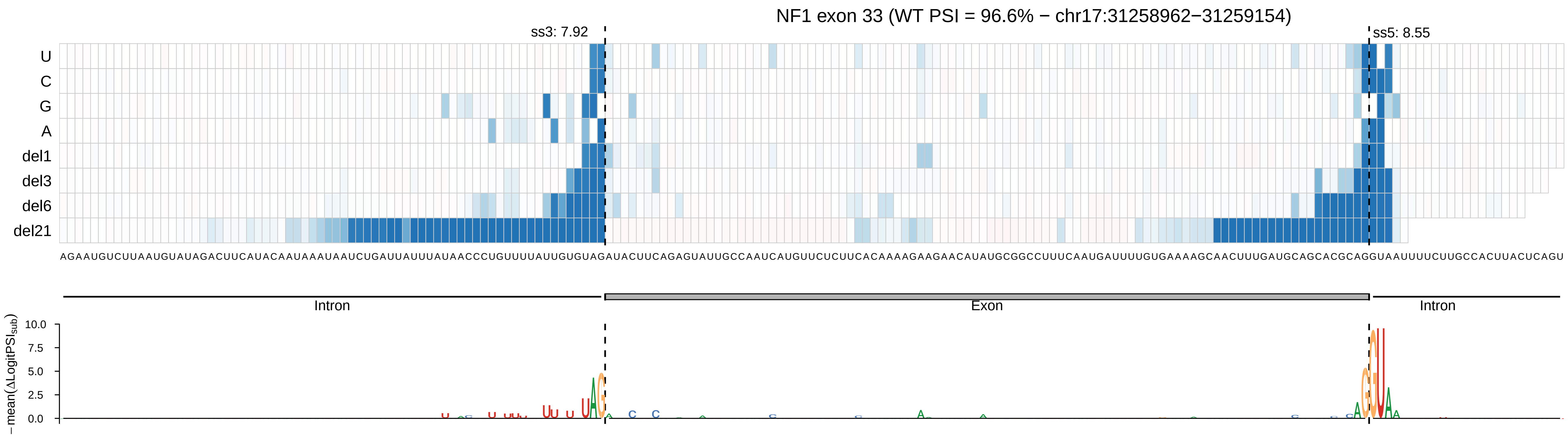
NF1 exon 33 (WT PSI = 96.6% − chr17:31258962−31259154)
ss3: 7.92
ss5: 8.55
U
C
G
A
del1
del3
del6
del21
Intron
Exon
Intron
−mean(ΔLogitPSI_sub)
10.0
7.5
5.0
2.5
0.0

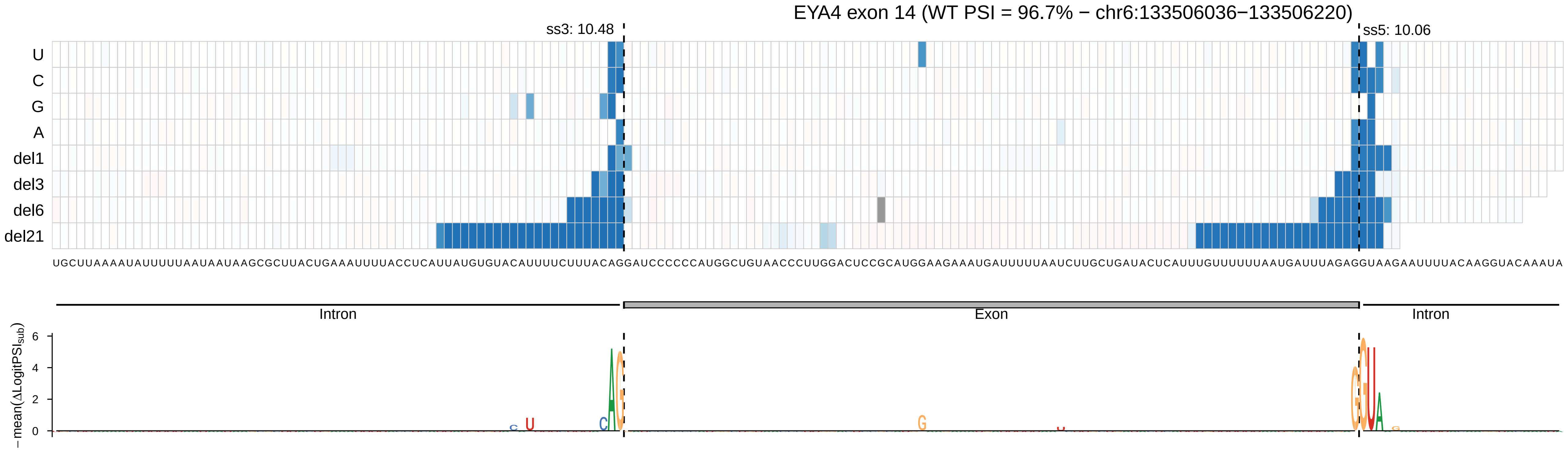
EYA4 exon 14 (WT PSI = 96.7% − chr6:133506036−133506220)
ss3: 10.48
ss5: 10.06
U
C
G
A
del1
del3
del6
del21
Intron
Exon
Intron
−mean(ΔLogitPSI_sub)
0
2
4
6

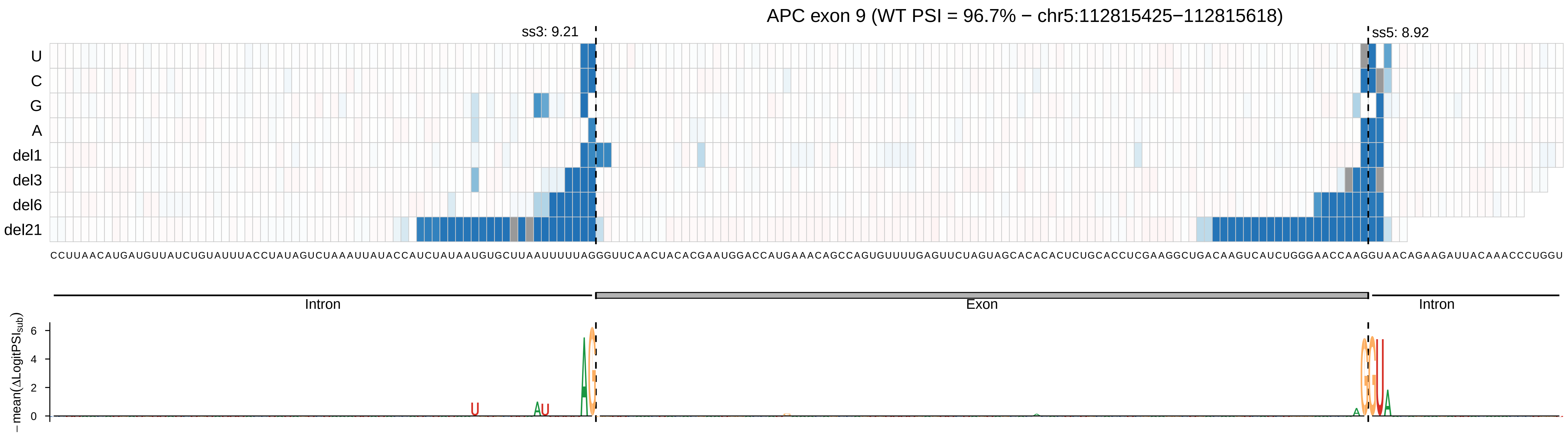
APC exon 9 (WT PSI = 96.7% − chr5:112815425−112815618)
ss3: 9.21
ss5: 8.92
U
C
G
A
del1
del3
del6
del21
Intron
Exon
Intron
− mean(ΔLogitPSI_sub)
6
4
2
0

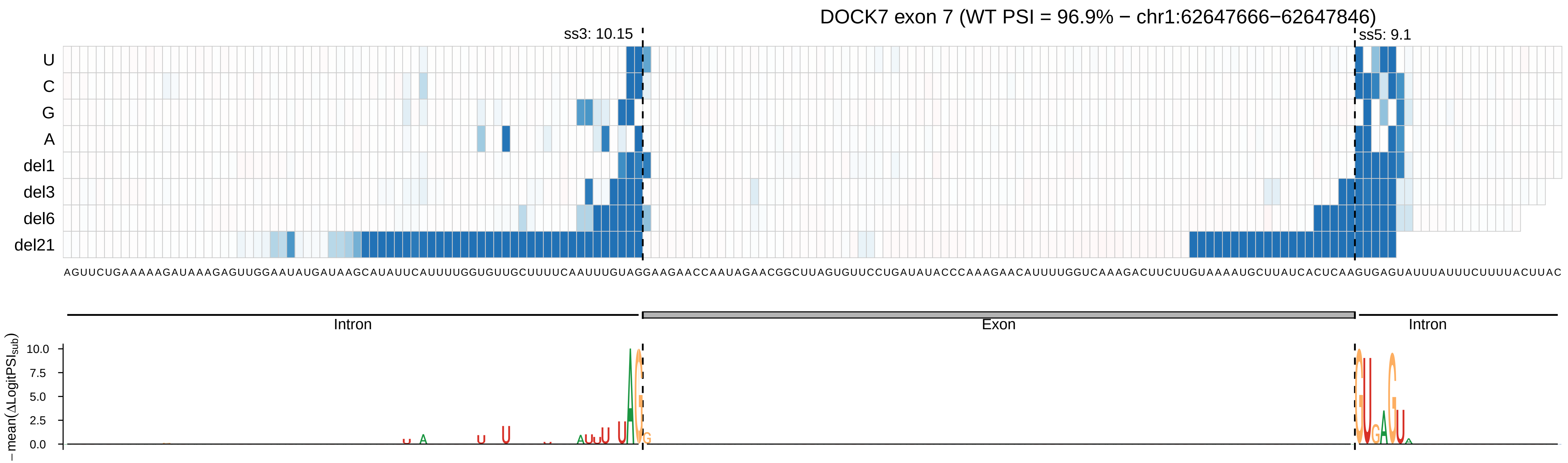

DOCK7 exon 7 (WT PSI = 96.9% − chr1:62647666−62647846)
ss3: 10.15
ss5: 9.1
U
C
G
A
del1
del3
del6
del21
AGUUCUGAAAAAGAUAAAGAGUUGGAAUAUGAUAAGCAUAUUCAUUUUGGUGUUGCUUUUCAAUUUGUAGGAAGAACCAAUAGAACGGCUUAGUGUUCCUGAUAUACCCAAAGAACAUUUUGGUCAAAGACUUCUUGUAAAAUGCUUAUCACUCAAGUGAGUAUUUAUUUCUUUUACUUAC
Intron
Exon
Intron
− mean(ΔLogitPSI_sub)
10.0
7.5
5.0
2.5
0.0

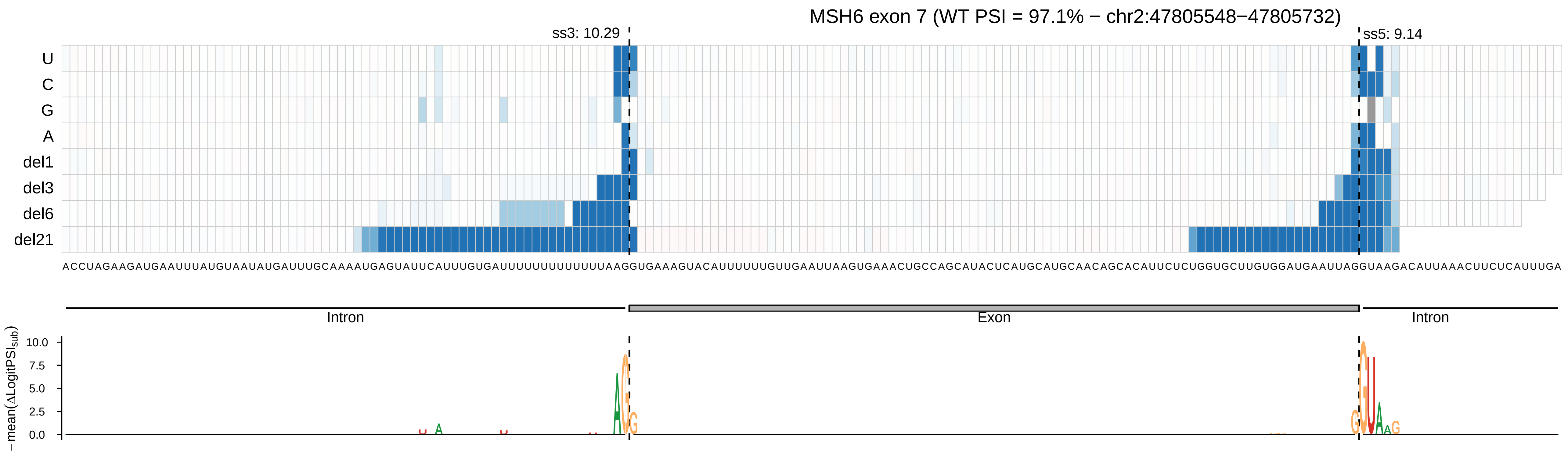

MSH6 exon 7 (WT PSI = 97.1% − chr2:47805548−47805732)
ss3: 10.29
ss5: 9.14
U
C
G
A
del1
del3
del6
del21
ACCUAGAAGAUGAAUUUAUGUAAUAUGAUUUGCAAAAUGAGUAUUCAUUUGUGAUUUUUUUUUUUUUUAAGGUGAAAGUACAUUUUUUGUUGAAUUAAGUGAAACUGCCAGCAUACUCAUGCAUGCAACAGCACAUUCUCUGGUGCUUGUGGAUGAAUUAGGUAAGACAUUAAACUUCUCAUUUGA
Intron
Exon
Intron
− mean(ΔLogitPSI_sub)
10.0
7.5
5.0
2.5
0.0

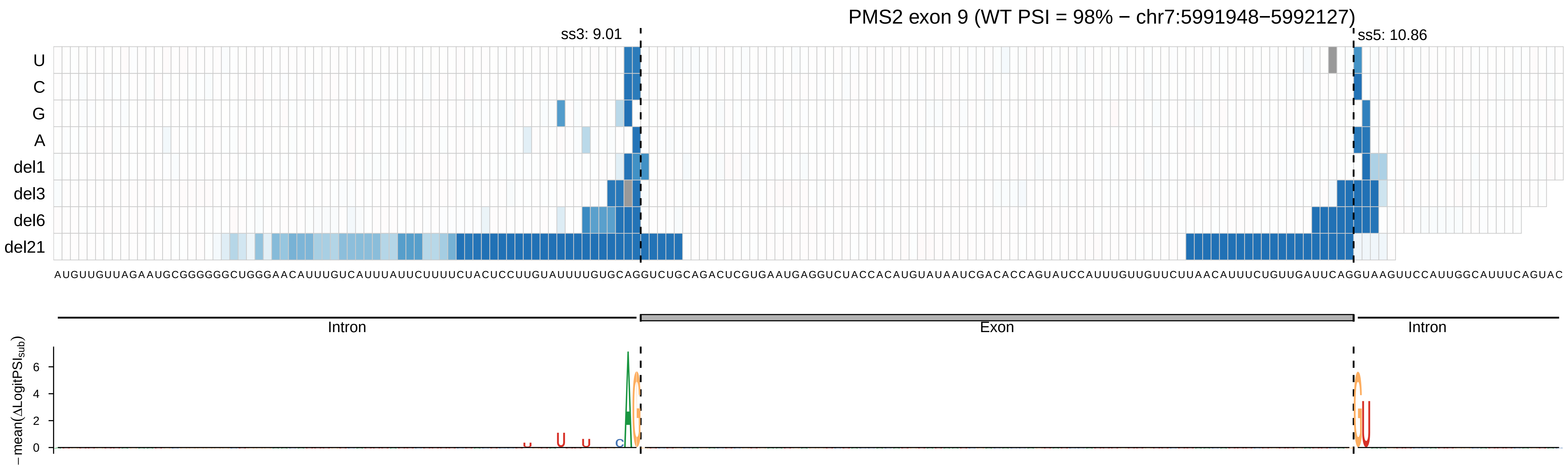
PMS2 exon 9 (WT PSI = 98% − chr7:5991948−5992127)
ss3: 9.01
ss5: 10.86
U
C
G
A
del1
del3
del6
del21
AUGUUGUUAGAAUGCGGGGGGCUGGGAACAUUUGUCAUUUAUUCUUUUCUACUCCUUGUAUUUUGUGCAGGUCUGCAGACUCGUGAAUGAGGUCUACCACAUGUAUAAUCGACACCAGUAUCCAUUUGUUGUUCUUAACAUUUCUGUUGAUUCAGGUAAGUUCCAUUGGCAUUUCAGUAC
Intron
Exon
Intron
− mean(ΔLogitPSI_sub)
0
2
4
6
