## Supplementary figures and images for "OpenSplice: the impact of half a million mutations on the alternative splicing of 600 human exons"

### Supplementary Figure 2

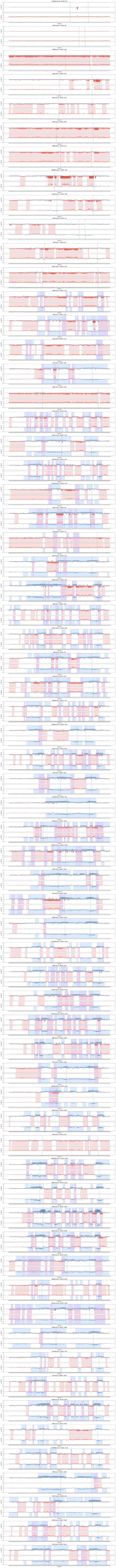

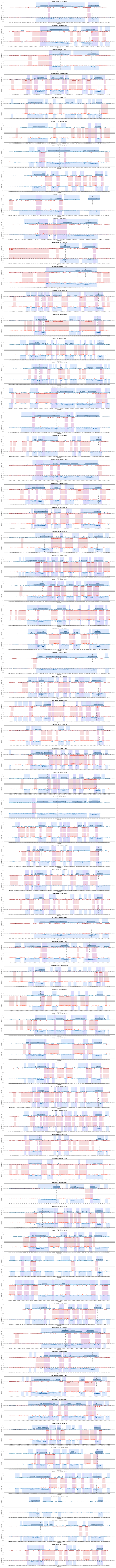

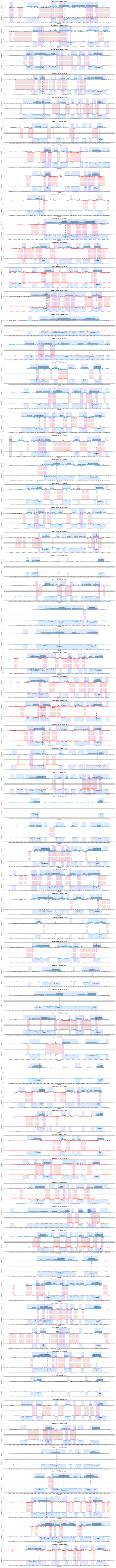

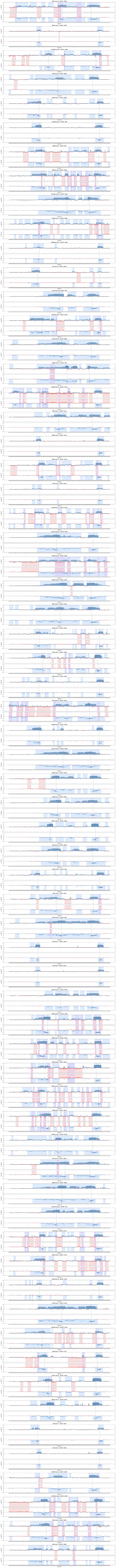

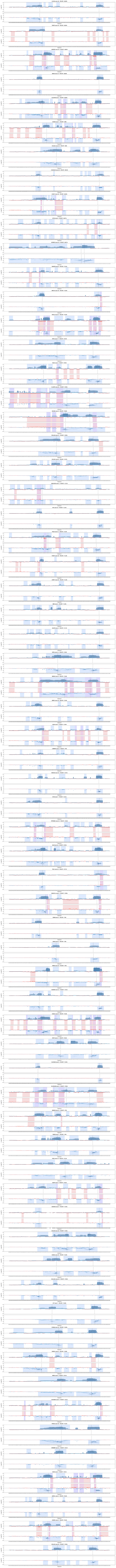

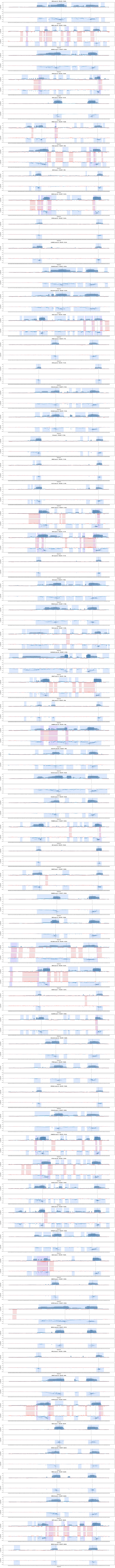

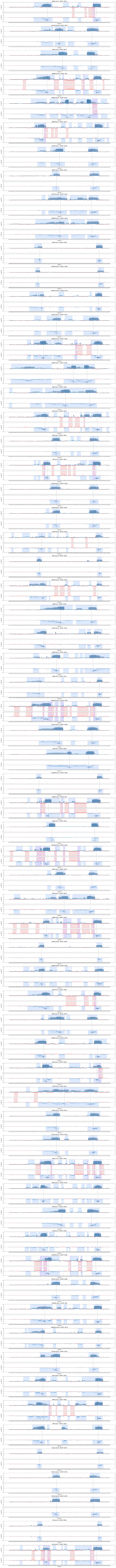

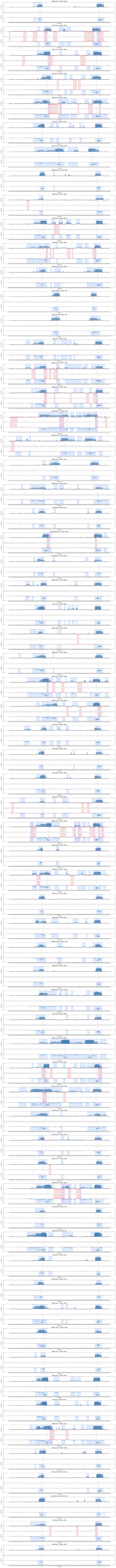

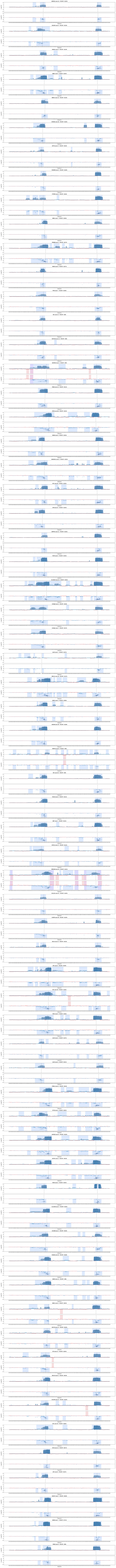
